# The genome of C57BL/6J “Eve”, the mother of the laboratory mouse genome reference strain

**DOI:** 10.1101/517466

**Authors:** Vishal Kumar Sarsani, Narayanan Raghupathy, Ian T. Fiddes, Joel Armstrong, Francoise Thibaud-Nissen, Oraya Zinder, Mohan Bolisetty, Kerstin Howe, Doug Hinerfeld, Xiaoan Ruan, Lucy Rowe, Mary Barter, Guruprasad Ananda, Benedict Paten, George M. Weinstock, Gary A. Churchill, Michael V. Wiles, Valerie A. Schneider, Anuj Srivastava, Laura G. Reinholdt

## Abstract

Isogenic laboratory mouse strains are used to enhance reproducibility as individuals within a strain are essentially genetically identical. For the most widely used isogenic strain, C57BL/6, there is also a wealth of genetic, phenotypic, and genomic data, including one of the highest quality reference genomes (GRCm38.p6). However, laboratory mouse strains are living reagents and hence genetic drift occurs and is an unavoidable source of accumulating genetic variability that can have an impact on reproducibility over time. Nearly 20 years after the first release of the mouse reference genome, individuals from the strain it represents (C57BL/6J) are at least 26 inbreeding generations removed from the individuals used to generate the mouse reference genome. Moreover, C57BL/6J is now maintained through the periodic reintroduction of mice from cryopreserved embryo stocks that are derived from a single breeder pair, aptly named C57BL/6J Adam and Eve. To more accurately represent the genome of today’s C57BL/6J mice, we have generated a *de novo* assembly of the C57BL/6J Eve genome (B6Eve) using high coverage, long-read sequencing, optical mapping, and short-read data. Using these data, we addressed recurring variants observed in previous mouse studies. We have also identified structural variations that impact coding sequences, closed gaps in the mouse reference assembly, some of which are in genes, and we have identified previously unannotated coding sequences through long read sequencing of cDNAs. This B6Eve assembly explains discrepant observations that have been associated with GRCm38-based analyses, and has provided data towards a reference genome that is more representative of the C57BL/6J mice that are in use today.

## Introduction

The inbred mouse strain C57BL/6 (B6) is the most commonly cited and well-characterized laboratory strain in biomedical research and comparative genomics. For that reason, this strain was selected by the Mouse Genome Sequencing Consortium (MGSC) to represent the laboratory mouse for reference genome sequencing^1,2^ in 1999 and it is the background strain on which the Knockout Mouse Project^3^ is creating and phenotyping null alleles for all protein-coding genes. The original whole genome shotgun (WGS) draft assembly of the C57BL/6 genome (MGSCv3) was later updated to a finished, clone-based assembly^4^. The finished assembly is comprised predominantly of Sanger sequencing of clones from two bacterial artificial clone (BAC) libraries, RPCI-23 and RPCI-24, derived from the DNA from pooled tissues of 3 females (kidney and brain) and one male (spleen and brain) mouse, respectively, representing inbreeding generation F204-F207 from production colonies at The Jackson Laboratory, hence the sub-strain designation C57BL/6J^4^. Since 2010, the Genome Reference Consortium GRC has actively maintained the mouse reference genome and produced updated assemblies, beginning with GRCm38 (GCA_000001635.2) in 2012 and its six subsequent patch releases.

Despite being one of the best-assembled mammalian reference genomes, GRCm38 still contains 523 gaps within chromosome sequences, and there are nearly 300 unresolved issues that have been reported to the GRC (https://genomereference.org). In addition to gaps, these issues include reports of localized sequence mis-assembly, missing genic and non-genic sequences, sequencing errors and suspect variation. These types of assembly issues inflate false positive rates in reference-based variant calling. For example, we reported an analysis of systemic exome variants called across a wide variety of mouse strains (including C57BL/6J) and showed that a significant fraction of these overlap with regions with annotated reference assembly issues and/or gaps^5^. The remaining fraction of recurrent false positive variant calls may represent unreported issues, reference-specific or private variation found only in earlier inbreeding generations, or regions where paralogous gene copies are not fully represented in the reference genome.

In an effort to close gaps and resolve other issues in the current mouse reference genome, to minimize variant calls associated with GRCm38-private variation (i.e. to bring the mouse reference genome sequence closer to the C57BL/6J mice that are currently in use), to provide a *de novo* assembly representing a single individual, and to identify additional data to support unannotated genes, we used high coverage, long-read sequencing, optical mapping and short-read data to generate a *de novo* genome assembly from C57BL/6J Eve.

## Results

The Jackson Laboratory manages the rate of genetic drift through periodic replenishment of foundation breeding colonies from pedigreed, cryopreserved embryo stock^6^ that are three generations removed from a single brother-sister breeder pair, “Adam” and “Eve” **(Figure 1)**. This process introduces a controlled bottleneck that minimizes the accumulation of genetic change. These two individual mice capture an evolutionary snapshot in time at inbreeding generation F223 **(Figure 1)**. Therefore, any C57BL/6J individual obtained from the production colonies at The Jackson Laboratory today is limited to a maximum of 24 inbreeding generations removed from the mice whose DNA was used to generate the C57BL/6J reference assembly, GRCm38 **(Figure 1).** Under the highly selective breeding paradigms employed for inbred laboratory strains, this genetic distance is sufficient for rapid fixation of 98.7% of variants, such that today’s C57BL/6J mice are by definition a sub-strain of the animals from which GRCm38 is derived ^7^. Therefore, we chose DNA isolated from one of these individuals as the material for our *de novo* assembly with the goal of providing a genome sequence from a single individual that is not more than eight generations removed from C57BL/6J mice sourced from The Jackson Laboratory, and that might also be used to improve the current C57BL/6J reference assembly (GRCm38). We chose C57BL/6J Eve (B6Eve) to get balanced representation of the X chromosome and the autosomes. Future efforts are focusing on a de novo assembly of Adam, where *de novo* assembly the Y chromosome will require more specialize approaches.

**Figure 1.**
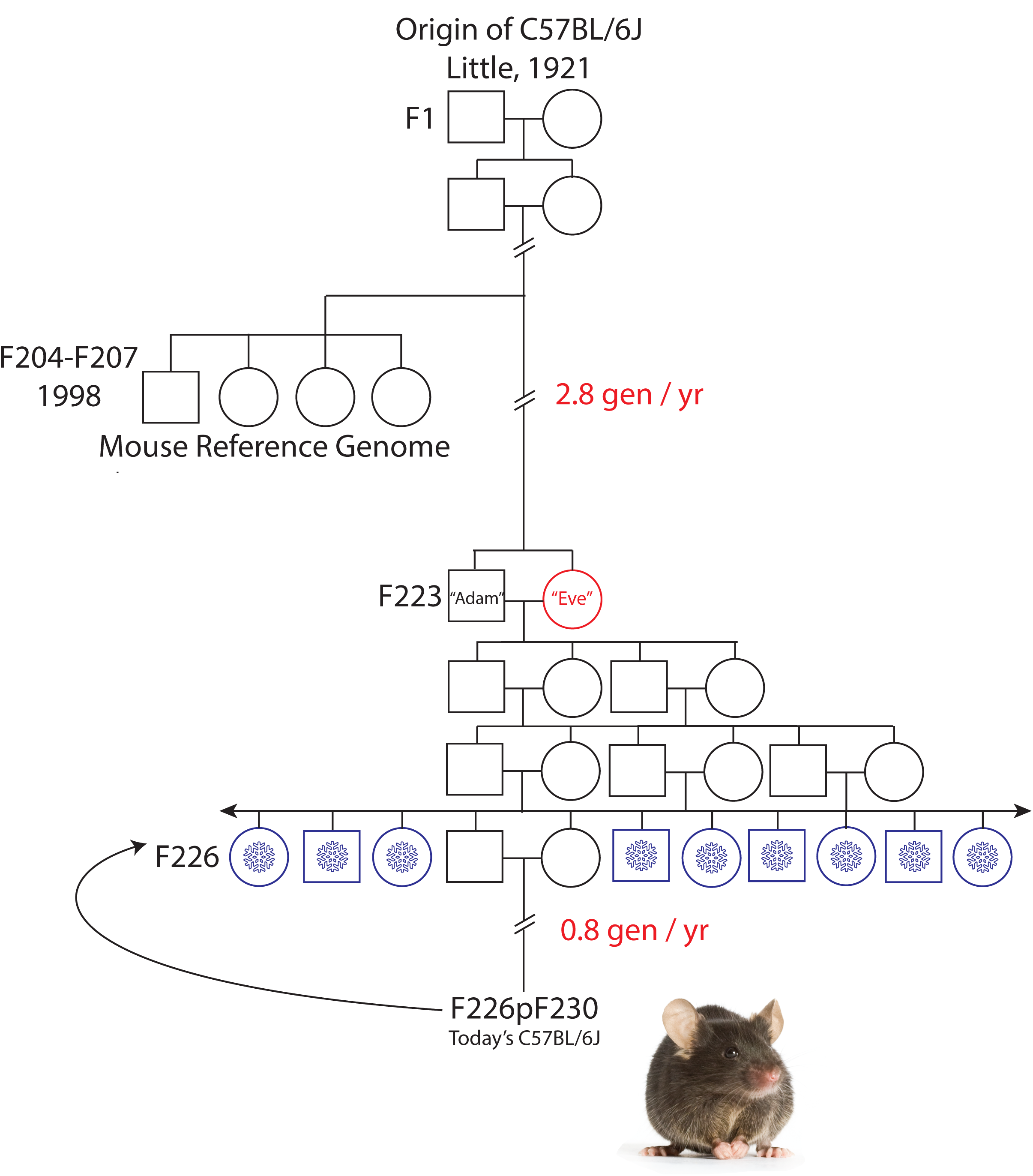
Origin of the inbred strain C57BL/6J. Inbred laboratory mouse strains are maintained by brother x sister mating. Filial (F) generations from which mice contributing to the reference assembly clone libraries and from which the B6Eve mouse were derived are shown. Cryopreserved embryo stock is represented by blue snowflakes at F226, 3 generations from Adam and Eve at F223. Generations subsequent to the cryopreservation event are F226p###, e.g. F226p230, which means embryos cryopreserved at F226 were recovered and there were an additional 4 generations of subsequent inbreeding.

### Sequence assembly and evaluation

To generate data for our *de novo* assembly of the B6Eve genome, we used a range of technologies, including Pacific BioSciences (PacBio) long read technology at 66X whole genome coverage **(Supplementary Table 1),** Illumina short read at 32X whole genome coverage, and Bionano Genomics (BNG) optical maps. The overall assembly procedure involved 1) correction of PacBio reads, 2) creation of contigs from PacBio, 3) extension of contigs to scaffolds using optical maps, 4) polishing of the assembly, and 5) further correction of the assembly using Illumina data **(Figure 2)**.

**Table 1:** Number of sequences, N50 size and assembly length for Bionano optical map, PacBio de novo assembly and scaffolded assemblies. Final hybrid assembly was submitted to the GenBank (**LXEJ02000000**)

|  | **Bionano**<br>**Genomics**<br>**optical**<br>**map** | **PacBio *de***<br>***novo***<br>**assembly** | **PacBio**<br>**only**<br>**Hybrid** | **Bionano**<br>**optical**<br>**only**<br>**Hybrid** | **Final Assembly**<br>**(LXEJ02000000)** | **Improvement**<br>**relative to**<br>**PacBio**<br>**assembly** |
| --- | --- | --- | --- | --- | --- | --- |
| **Number of sequences** | 3,016 | 14,551 | 3,732 | 1,652 | 12,690 |  |
| **N50 (in MB)** | 1.18 | 0.40 | 0.58 | 1.97 | 1.29 | 3.2 |
| **Assembly size (in MB)** | 2,482.74 | 2,535.01 | 1,820.29 | 2,470.31 | 2,789.93 | 1.3 |

**Figure 2:**
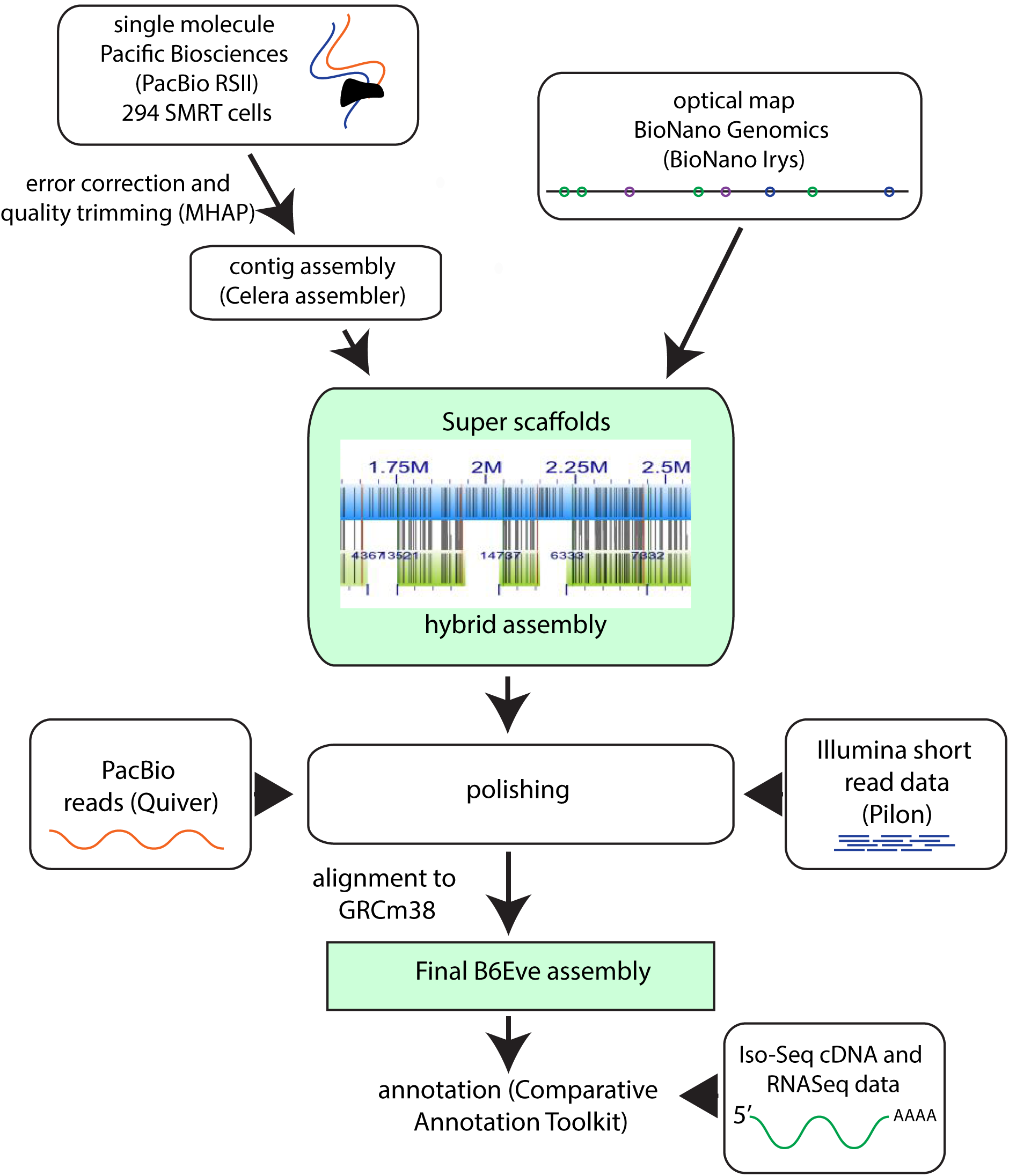
Schematic overview of the *de novo* assembly procedure for B6Eve. Details are described in Methods.

To assess base-pair level improvements in assembly quality afforded by each of these steps, we mapped the Illumina reads of B6Eve and called variants at each step using the GATK HaplotypeCaller ^8^ (**Supplementary Table 2**). The scaffolded assembly yielded 1,664,599 variants, the majority of which were insertions (~70%), followed by SNPs and small deletions.

**Table 2:**
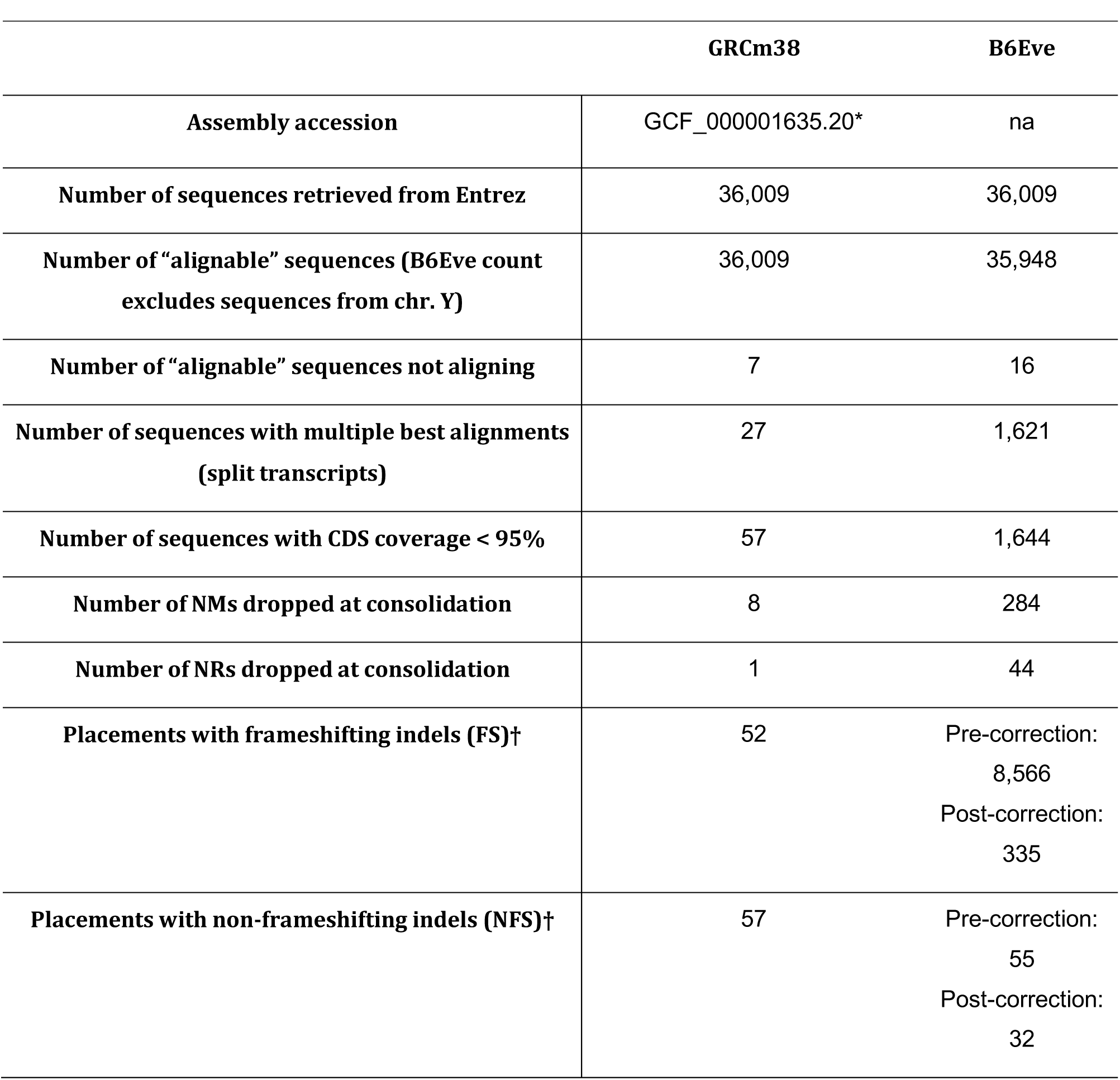
RefSeq Transcripts Alignment Table From NCBI

|  | **GRCm38** | **B6Eve** |
| --- | --- | --- |
| **Assembly accession** | GCF_000001635.20* | na |
| **Number of sequences retrieved from Entrez** | 36,009 | 36,009 |
| **Number of “alignable” sequences (B6Eve count excludes sequences from chr. Y)** | 36,009 | 35,948 |
| **Number of “alignable” sequences not aligning** | 7 | 16 |
| **Number of sequences with multiple best alignments (split transcripts)** | 27 | 1,621 |
| **Number of sequences with CDS coverage < 95%** | 57 | 1,644 |
| **Number of NMs dropped at consolidation** | 8 | 284 |
| **Number of NRs dropped at consolidation** | 1 | 44 |
| **Placements with frameshifting indels (FS)†** | 52 | Pre-correction:<br>8,566<br>Post-correction:<br>335 |
| **Placements with non-frameshifting indels (NFS)†** | 57 | Pre-correction:<br>55<br>Post-correction:<br>32 |

\* Transcripts were aligned to the GRCm38 full assembly (GCF\_000001635.20), which includes alternate loci scaffolds from a variety of mouse strains. Counts shown in Table 2 reflect only transcript alignments to the GRCm38 primary assembly unit (GCF\_000000055.19), which is comprised only of C57BL/6J sequences, unless noted.
† Frameshift counts are shown for alignments to the GRCm38 full assembly, including alternate loci scaffolds. Pre-correction: assembly prior to Quiver polishing and Pilon correction.

This pattern is reminiscent of the error profile generated from PacBio technology^9^. To improve base pair accuracy, we used Quiver software^10^ to polish the assembly and reduced the total number of variants to 505,782. We found that compared to the unpolished assembly, Quiver reduced the number of insertions to just 22%. Finally, we used B6Eve Illumina data itself to correct the polished assembly with Pilon^11^, a tool that improves the quality of the draft assemblies using read alignment analysis. After Pilon correction, the number of variants was narrowed down to 310,205; of which 227,523 were considered high quality (PASS) by GATK HaplotypeCaller **(Supplementary File1)**.

We used an automatic assembly quality evaluation tool, QUAST ^12^, to assess the overall quality of the Pilon corrected assembly. We found that during assembly-assembly alignment, 96.8% of the B6Eve assembly aligned to 97% of the GRCm38 (excluding chrY and alternate sequences) reference genome chromosomal sequences and 54% of unplaced & unlocalized sequences (**Supplementary Figure 1**). Only 166 and 2,456 B6Eve components comprising of 3.7 Mb and 7.2 Mb of sequences remained wholly or partially unaligned to the reference genome, respectively. We also found that the K-mer based completeness of B6Eve was very high at 97.4%, suggesting high coverage and per-base quality. The detailed QUAST report for complete and broken-down (breaks the assembly by continuous fragments of N's of length ≥ 10) versions of the assembly is found in **Supplementary File2.** Taken together, our resulting PacBio-only *de novo* B6Eve genome assembly was 2.53 Gb consisting of 14,551 contigs (longest contig = 4,574,471 and 2.3% of total contigs exceeding 1Mb) with an N50 size of 401,294 bp. Our complete PacBio-Bionano hybrid assembly yielded an N50 of 1,290,032 bp with a total assembly size of 2.79 Gb, which was a 3X improvement (in N50) over the PacBio-only assembly **(Table1)**.

### Gene Content Analysis

We also evaluated the gene content of the B6Eve assembly as another measure of assembly quality. Akin to previous analyses^13^ ^14^, we aligned 36,009 RefSeq transcripts to the GRCm38 primary assembly (also excluding unplaced sequences and alternate loci scaffolds from other mouse strains that are a part of the full assembly) and to the “Piloned” B6Eve assembly versions **(Table 2 and Supplemental Table 3).** We observed that B6Eve provides a comparable representation for total gene content, as compared to GRCm38, with only 19 non-chromosome and chrY associated sequences having no alignment. Consistent with the more fragmented nature of the B6Eve assembly, however, a greater number of aligned RefSeq transcripts exhibit partial alignments or alignments split over multiple scaffolds than in GRCm38. We also examined the co-placement of transcripts representing different genes as a proxy for measuring the collapse of segmental duplications. Although B6Eve shows a greater number of co-placed transcripts than GRCm38, these numbers are consistent with those seen in other high-quality long-read derived WGS assemblies, demonstrating the utility of this mouse assembly. To gauge the impact of the Illumina-read correction step on the quality of protein representation in the B6Eve assembly, we looked at the incidence of frameshifting indels in aligned RefSeq transcripts prior to and after this step^15^ **(Supplemental Table 3)**. Although the Pilon corrected assembly still exhibits more frameshifts than GRCm38, we find that this step resulted in a substantial improvement in functional representation (protein coding sequence).

**Table 3:** Counts of various structural variation classes detected in the comparison of B6Eve Sequences to GRCm38 using PacBio and Illumina data

| Technology | Duplication | Deletion | Inversion | Insertion | Trans |
| --- | --- | --- | --- | --- | --- |
| PacBio | 229 | 418 | 36 | 3,394 | 71 |
| Illumina | 289 | 221 | 111 | - | - |
| Common | 44 | 12 | 4 | - | - |

### Reference Assembly Gap Filling

The GRCm38 chromosome assemblies contain 440 gaps (excluding centromere, short arm, and telomere gaps). We assessed whether sequences in the B6Eve assembly could resolve these gaps. Based on our gap-filling methodology (**see Methods**), the B6Eve assembly spans 23 gaps in the GRCm38 chromosomes (**Supplementary Table 4 and Supplementary Figure 2**). In several instances, we observed discrepancies between the gap length reported in GRCm38 and the amount of sequence provided by the B6Eve assembly. For example, the B6Eve assembly spans a 1,760 bp intra-scaffold gap located at chr2:172,624,657-172,626,416 bp in GRCm38, with 1,620 bp (a 140 bp relative deletion) (**Supplementary Figure 2a**). In other cases, B6Eve spanning sequences are longer than assembly gaps. For example, a B6Eve assembly scaffold spans the 100 bp intra-scaffold gap at chr1:183,334,907-183,335,006 bp in GRCm38, with 595 bp sequence (**Supplementary Figure 2b**). These discrepancies are not unexpected, however, as the methods used to estimate reference assembly gap sizes do not always offer base-pair level resolution, and also because the GRC assigns default gap lengths when no sizing estimates are available (https://www.ncbi.nlm.nih.gov/projects/genome/assembly/grc/TPF_Specification_v1.8_20131106.docx). Consistent with prior reports that remaining reference assembly gaps are in complex genomic regions^16^, we observe that our gap spanning sequences have a repeat content of 45.9% (vs. 42.5% of GRCm38 total sequence) repeats, with simple repeats accounting for 18.1% (vs 2.6% in GRCm38 total sequence).

**Table 4:**
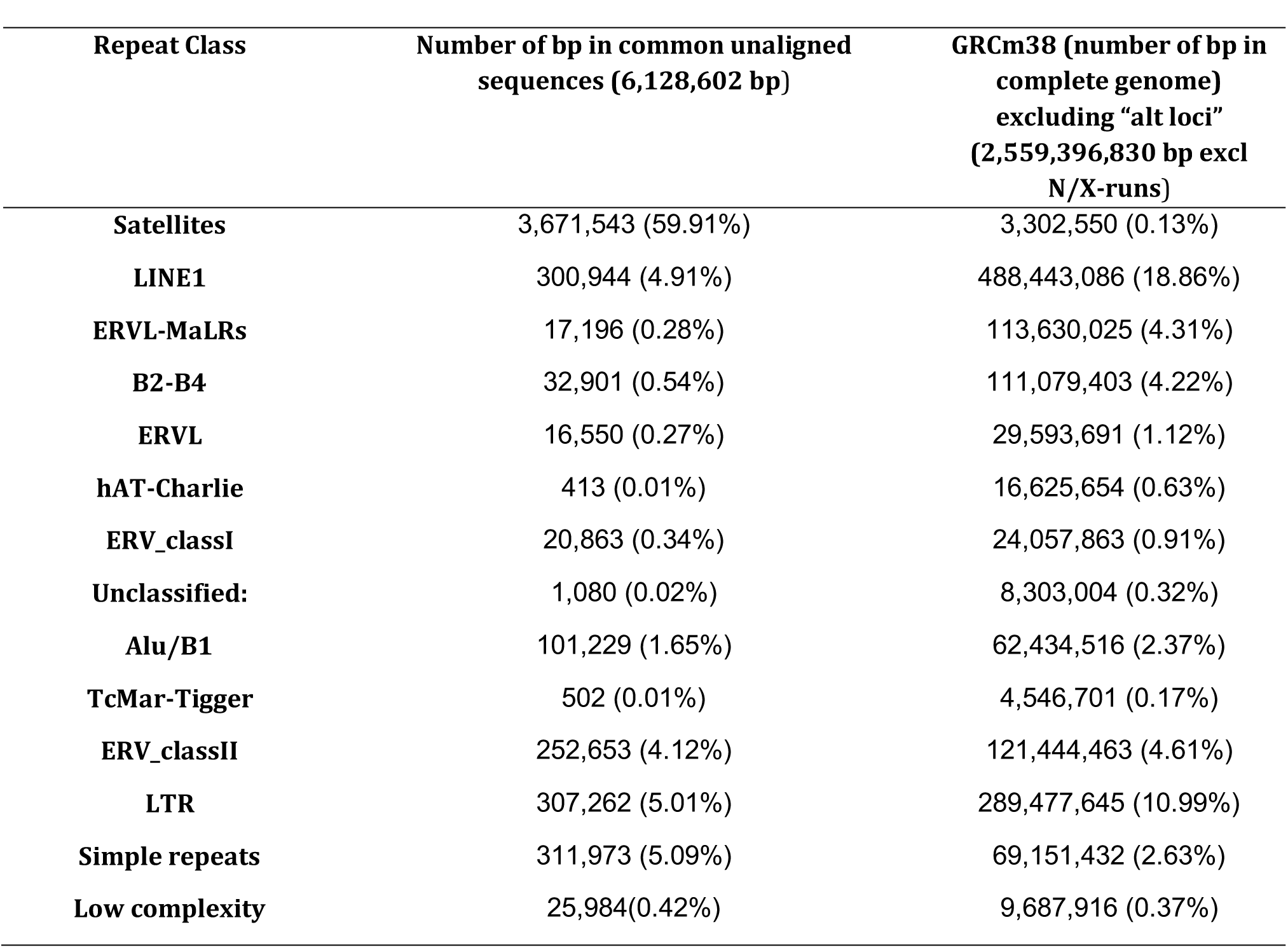
Comparison of various repeat class in common unaligned sequences with GRCm38

### Variant analysis

Previously, we reported whole-exome sequences from of a collection of nearly 200 unique strains of spontaneous mutant mice maintained at The Jackson Laboratory ^5^. In our analysis of these exomes, we found that there were 855 coding variants (SNPs) common across 75% or more of the samples, which we attributed to errors with the reference genome itself due to their significant inclusion within the component-mapped boundaries of GRC incident features. We investigated the subset of these exome variants (n=126) that were homozygous across all strains (100% allele frequency). We find that 10 (7.9%) matched the B6Eve assembly allele rather than the GRCm38 allele, supporting the assertion that high frequency alleles are putative indicators of reference assembly error **(Figure 3, Supplementary Table 5**).

**Figure 3.**
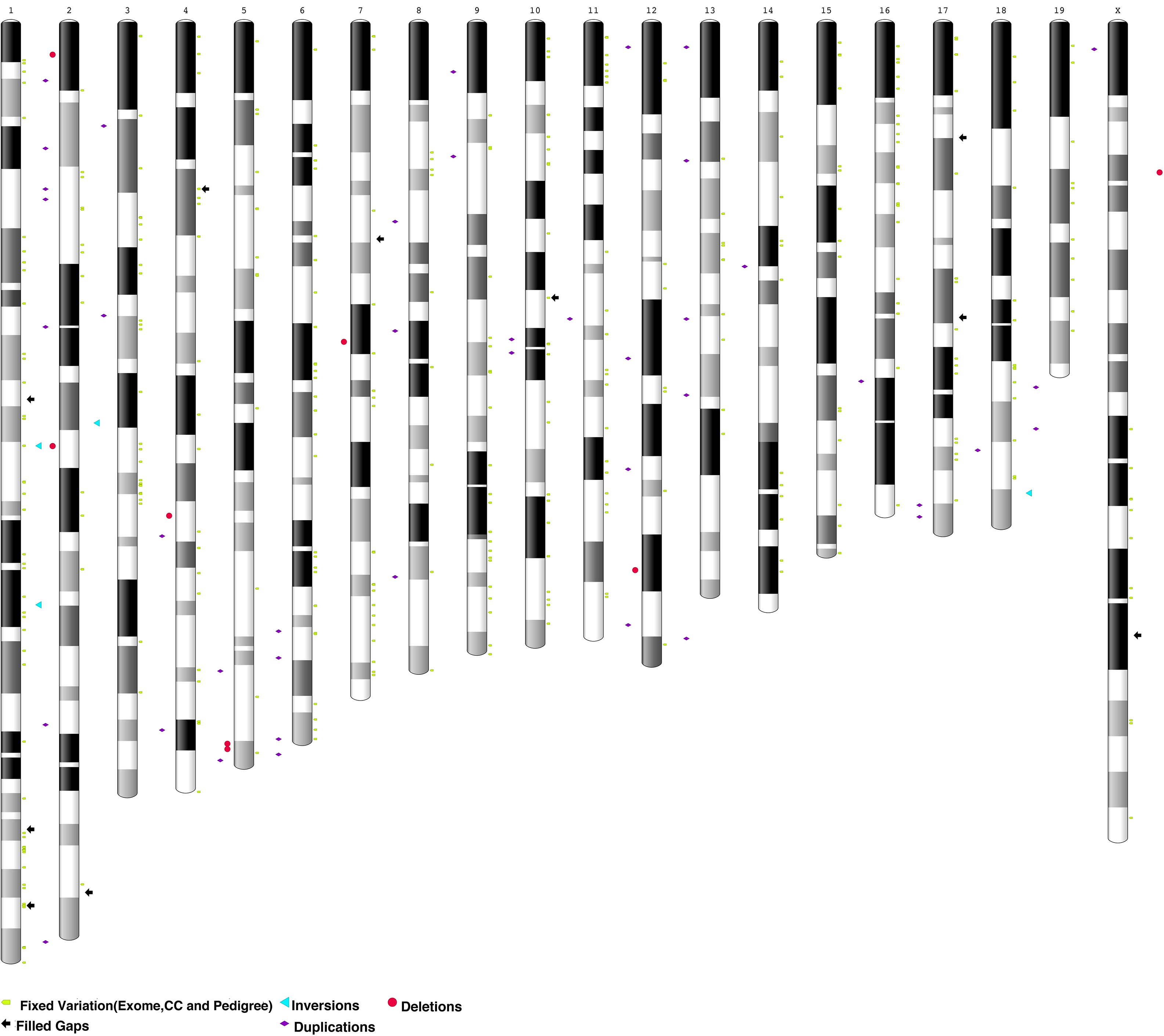
Ideogram of GRCm38 assembly annotated to highlight resolved gaps (vs. current reference), structural variants, and fixed variation using B6Eve data.

To extend this analysis to the whole genome, we performed a similar analysis using variant calls from sixty-nine multi-parent, recombinant inbred strains (Collaborative Cross (CC) strains, a panel derived from eight founder laboratory strains) ^17^. We found 14,757 variants (SNPs) shared across all strains, using C57BL6/J as a reference genome. Out of 14,757 variants, 2,407 are homozygous across all strains. Consistent with the results of the exome variant analysis, 307 of these variants (12.8%) (**Figure 3, Supplementary Table 5)** match the B6Eve assembly allele rather than GRCm38.

Finally, we analyzed variant calls from whole genome short read sequencing of 24 recent descendants of B6Eve, representing multiple inbred lineages. We found 3,203 homozygous variants (SNPs) common across these samples; of these, 2,194 (68.5%) met minimum alignment criteria for remapping to B6Eve. Of these 2,194, we found 393 cases (12.6% of the total variation and 17.9% of net variation) **(Figure 3, Supplementary Table 5)** where the reported alternate alleles matched the B6Eve assembly.

Taken together, our analyses identify 503 single nucleotide positions in GRCm38 (excluding variants common in three datasets) that are not representative of today’s C57BL/6J mice. Two of these are nonsynonymous SNPs in *Akap9 and Sfi1*. *Akap9* (A kinase anchoring protein 9) is a protein that is responsible for cytoskeletal organization and is required for formation and maintenance of the blood-testis barrier, and male fertility ^18,19^. Using allele specific PCR, we confirmed the presence of the alternate *Akap9* allele in B6Eve and in randomly selected descendants of Eve, and not in an ancestor of B6Eve. *Sfi1* is predicted to be a spindle assembly associated protein on the basis of homology with a yeast cytoskeletal protein of known function, however no phenotypic alleles have been reported in mice^20^. When we attempted to validate this variant, however, we discovered that the alternate allele is indeed represented in the mouse reference genome, though it amplifies from an unplaced scaffold (JH584304.1: 14,038-14,339). Flanking sequence variation between this scaffold and Chr11 allowed us to design allele specific primers, with which we confirmed that both alleles are present in DNA samples from B6Eve and from randomly selected descendants of B6Eve, which supports the idea that the unplaced scaffold indeed represents C57BL/6J sequence. Previously published SV data for C57BL/6J showed that the mouse genome potentially harbors 20-30 copies of this gene^21^. Therefore, the recurrent “variation” observed in this gene is likely not allelic, but due to mis-mapping of reads from paralogous gene copies to the *Sfi* locus that is currently represented on GRCm38 chromosome 11. Paralogous gene variation may be a previously underappreciated source of variation, since we observed a relative enrichment of variants within certain genes (e.g *Tulp4*, Supplemental Table 5). Previous studies have shown that GRCm38 is missing paralogous copies of many genes^4,16^, some of which may be represented on unplaced scaffolds as we found for *Sfi1*.

### Structural Variation

We aligned raw B6Eve PacBio reads to GRCm38 using NGMLR ^22^ and called structural variants (SVs) with Sniffles ^22^. We also aligned Illumina WGS data from B6Eve and called SVs with Delly 23 **(Table 3).** The median size of detected duplication, deletion and inversion events from Delly were 901, 2,610 and 12,362 bp, respectively. Similarly, from PacBio data, the median size of duplication, deletion, inversion and insertion events were 432, 77, 1352 and 92 bp, respectively.

We found 12 deletion, 43 duplication and 4 inversion calls that were common in both Illumina and PacBio data (**Supplementary Table 6)**. Of these common SVs, 8 deletions, 30 duplications, and 4 inversions overlapped genes (**Supplementary Table 6)**, though mostly within noncoding intronic regions. We used DGVa ^24^ to further investigate the SVs overlapping genes. Each of these were associated with multiple (21-124) DGVa entries representing germ line SV across genetically diverse inbred strains from multiple strain surveys of SV ^25-27^. Some of these regions contain genes that have been previously been shown to be subject to positive selection of copy number variants in inbred laboratory mouse strains. Our data show that even within a strain, we can detect SV in these regions, which suggests that these regions are by their very nature susceptible to rearrangements, i.e. through suppressed recombination. Alternatively, recurrent SV calls could reflect either private SVs in the reference assembly, or mis-assembly of these regions.

### Repeat analysis

Repetitive sequences present challenges to assembly, as highly identical repeat sequences from different genomic regions are often incorrectly assembled together. This is particularly a concern for data generated from short-read technologies, which are too short to span longer repeats. One advantage of long read sequencing reads are their ability to span a greater range of genomic repeats into unique sequence, enabling resolution of repetitive regions that cannot be resolved in unlinked short read assemblies. We used RepeatMasker ^28^ to compare repetitive sequence representation between B6Eve, GRCm38, and two Illumina WGS assemblies (GCA_000185125.1, GCA_000185105.2) ^29,30^. This analysis revealed that the B6Eve assembly consists of 42.0% repetitive sequence (1,065,403,997 out of 2,537,631,632 bp, excluding N’s). This fraction is very similar to GRCm38 (excluding the Y chromosome and alternate loci sequences), 42.5% of which is repetitive (1,088,395,156 out of 2,559,396,830 bp, excluding N’s and X’s). Consistent with the challenges of assembling repetitive sequence with shorter reads, the Illumina based assembly GCA_000185125.1 had 32.5% (833,318,654 out of 2,257,461,872 bp excluding N/X-runs) and GCA_000185105.2 had 33.9% (849,891,926 out of 2,279,058,378 bp excluding N/X-runs) annotated as repetitive sequence (**Supplementary Table 7a-d**).

We also used RepeatMasker analysis to assess repeat content in scaffolds from our B6Eve assembly that failed to align to GRCm38, as we surmised these scaffolds might contain repetitive sequences that could not be resolved with genomic clones. To do this we focused on sequences which are unaligned by all of three of the following methods a) Cactus based alignment using UCSC Comparative Annotation Toolkit b) NCBI assembly-assembly alignments and c) QUAST evaluation. The common unaligned sequences (total 6.12 MB) (**Supplementary File3)** between CACTUS, NCBI, and QUAST had significant enrichment for repeats relative to aligned sequences. The repeat content accounted to 77.6% (4,754,330 out of 6,128,602 bp) with the microsatellite repeat class showed significantly enrichment when compared with GRCm38 (59.9% vs 0.1%, chi-square test: X^2^=80,332,000, p-value < 2.2e-1 **(Table 4).** While more work is needed to determine the underlying cause of failed alignment, the enrichment of microsatellite repeats in these scaffolds is compelling. Microsatellite repeats are prone to slippage during DNA replication, and as a result their copy number is highly polymorphic in eukaryotic genomes; a phenomenon known as microsatellite instability (MSI). In inbred laboratory mouse strains and in the human population, mutations that change copy number occur at rates that are up to 10,000 times higher than single nucleotide mutation rates (1-3 × 10^−^4 9,22,23 per repeat per generation for microsatellite sequences vs. 2-4 × 10^−9^ ^30^ per nucleotide per generation for SNV in C57BL/6J). Similarly, in the human population, CNV are estimated to occur at rates that are 100-10,000 times higher than the point mutation rate ^31^. Taken together, CNV are a major source of intrastrain variation and divergence from isogenicity ^24,29^. Therefore, failed alignment of these microsatellite containing scaffolds could be due to repeat polymorphisms that have arisen over the intervening years in C57BL/6J. Alternatively, failed alignment could be due to assembly issues in either genome.

### Gene prediction

Long read sequencing of cDNAs (IsoSeq) provides full-length transcript sequences and highly accurate representations of splice junctions and isoforms. To determine if long read sequencing data of B6Eve cDNAs could support more accurate gene prediction for the mouse reference genome, we generated IsoSeq data from RNA extracted from archived B6Eve brain. We used the Comparative Annotation Toolkit (CAT)^32^ to identify 107,192 transcripts (82,187 protein-coding) representing 41,669 gene loci (20,182 protein-coding). 2,426 transcript predictions had splice junctions that were novel relative to GENCODE VM11, and we found additional support for these junctions in RNA-Seq data generated from the brain of a female C57BL/6J descendent of Eve. Analysis of the transcript predictions produced by AugustusPB and AugustusCGP revealed 206 exons with splice site shifts relative to GRCm38, nine putatively novel exons and ten putatively novel loci (**Supplementary Table 8)**. Three of the novel exons detected in the IsoSeq data reveal deletions in GRCm38: (1) 640 bp in *Mia3* (**Figure 4),** (2) *Traf5*, and *Slc26a6* (**Supplementary Figure 3 and 4).** In support of these data, there are GRC incident reports describing deletions at each of these loci in GRCm38. Contiguity analysis in the B6Eve assembly showed that 616 genes mapped across two or more scaffolds and four genes had projections split on the same scaffold. A total of 258 protein-coding genes exhibited signs of gene family collapse, with 156 pairs of genes being resolved to the same locus.

**Figure 4.**
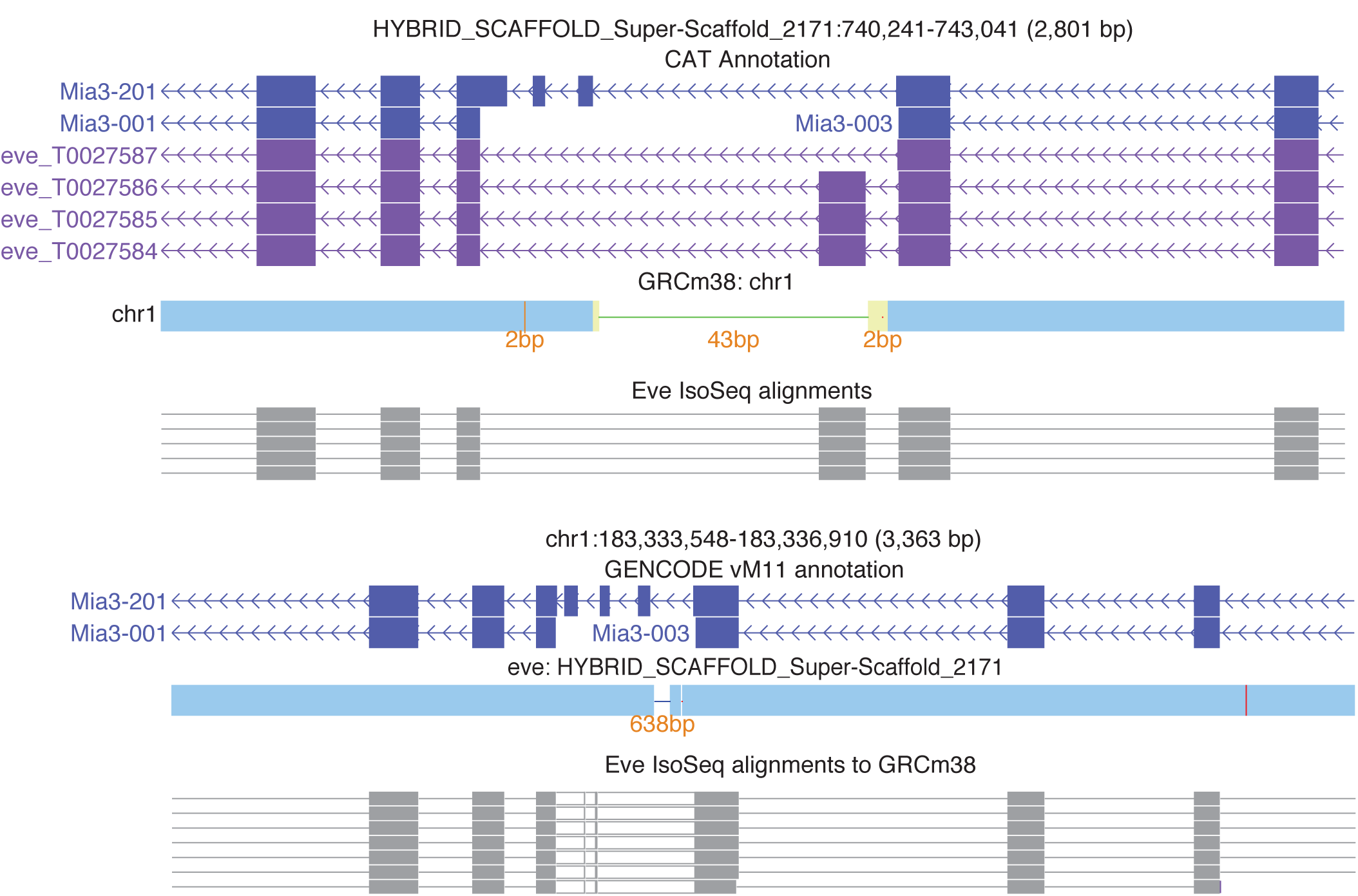
The *Mia3* locus from the perspective of both the B6Eve assembly (top) and the GRCm38 mouse reference (bottom). CAT annotation of B6Eve identified three isoforms with an IsoSeq supported exon not found in the reference. The cactus alignments (blue bars) show that there are 43 bp of reference sequence that does not align to B6Eve, and that there are 638 bp of B6Eve not seen in the reference. These 638 bp contain the extra exon. This result is confirmed in the B6Eve IsoSeq GRCm38 alignment, which shows an insertion (white blocks between grey exon alignments).

## Discussion

The value of isogenic mouse strain backgrounds in biomedical research was recognized by geneticists in the early 20^th^ century leading to the creation and description of the over 450 unique inbred mouse strains to date^33,34^. Twenty generations of sibling intercrosses are required for the generation of a new inbred strain; a breeding method that creates genomes in which more than 98% of loci are homozygous. Therefore, individuals within a generation, within the same vivarium are essentially, genetically identical. The remarkable genetic architecture of inbred laboratory mouse strains is shaped by the frequent bottlenecks required for the on-going maintenance of these strains. This accelerates genetic drift and is a major source of the often unexpected, genetic variation that can be observed across generations and/or between vivaria. For example, a reference-based alignment of the inbred laboratory strain C57BL/6J yields approximately 900 raw variant calls (SNPs/Indels), despite it being the same inbred strain as the mouse reference genome^5^. While some of these variant calls are certainly due to private reference alleles or intra-strain variation, we previously found that a significant percentage of these variants are located in regions of the reference genome where there are reported assembly issues^5^ and regions that contain missing paralogs, which are a known source of false positive variation due to mis-mapping of reads^16,27,31^. A major goal of this *de novo* assembly was to generate long read sequencing data that could potentially be used resolve these regions and to provide an updated representation of the variation that is present in the most recent inbreeding generations of C57BL/6J.

Using variant data from our B6Eve assembly, as well as data from several other large sequencing efforts^17^, we provide a “truth” call for over 500 high quality, recurring variants (SNP/Indels) that can be used to update the mouse reference genome (GRCm38.p6). We also found evidence for over 40 structural variants (inversions, deletions, and duplications) involving protein-coding genes in our B6Eve assembly compared to the reference genome. The majority of these SV calls were found in DGVa ^24^ across a variety of strains, suggesting that they are likely recurrent SV calls that, similar to recurrent variants, are due to mis-assembly of paralogous sequences or reference specific SVs. Further, our data fill 23 gaps of varying length in the mouse reference genome which will be used to inform the upcoming release of GRCm39.

Our IsoSeq data provided improved/more accurate gene models with previously unrecognized splice junctions for over 2,000 genes. This is likely an underrepresentation since our analysis is limited to only those genes expressed in brain. We also found evidence for novel exons, as well as evidence for novel loci (expressed regions that lack gene annotation). This demonstrates that even in a well-curated reference genome assembly, gene annotation remains subject to change as new technologies provide improved representation of transcribed sequences and access to more highly specialized cell types.

Whole genome sequencing data are now available for hundreds of standardized laboratory inbred mouse strains^17,27,35^. These data reveal the remarkable architecture of inbred genomes, and provide a stark reminder that isogenic mouse strains are subject to genetic drift; a feature that directly conflicts with the idea of a ‘reagent-grade’ laboratory mouse. Careful breeding practices, cryoarchiving, and routine sequencing are key steps towards maximizing reproducibility of studies that rely on these living reagents. Ultimately, *de novo* assembly captures the full spectrum of genetic variation resident in inbred strains, some of which harbor significantly more variation than distantly related human populations. Recently, genome graphs have been used to represent “population reference genomes” as a means to improve read mapping and to minimize false positive variant calls^36,37^. As applied to mouse genomes, this approach would ideally provide a framework for future representation of the laboratory mouse reference genome as a graph of many inbred strains upon which emergent variation can be more accurately discovered and used to guide experimental research involving laboratory mouse strains.

## Material & Methods

### Sample preparation and sequencing - PacBio

Genomic DNA samples were extracted using both kidney and brain samples from the C57BL/6J Eve female (MouseID 03-03685 (The Jackson Laboratory), Strain ID 00664, born 8/27/2003, generation F223) by phenol-chloroform extraction of a nuclei-enriched pellet. DNA samples were resuspended in TE buffer to a final concentration of 300-400 ng/ul, 260/280 1.8-1.9^14^. The PacBio data were generated from three libraries prepared using the Pacific Biosciences SMRTbell Template Prep Kit 1.0 (Pacific Biosciences, Menlo Park, CA, USA) using the “20-kb Template Preparation Using BluePippin Size-Selection System (15-kb Size cutoff)” protocol obtained from PacBio SampleNet. The BluePippin (Sage Science, Inc, Beverly, MA, USA) was set to collect from 7-50 kb. After sequence length QC, the resulting sized libraries were repaired using the “Procedure & Checklist-10 kb Template Preparation and Sequencing” protocol. All libraries were sequenced using 294 SMRT cells on Pacific Biosciences RS II platform (P6C4 chemistry). One of these libraries was generated and sequenced (10X) by Pacific Biosciences using the same protocols and chemistries. The other two libraries and the remaining coverage were generated and sequenced at The Jackson Laboratory.

For IsoSeq cDNA sequencing, RNA was extracted from archived whole brain samples from C57BL/6J Eve. 1 microgram of input RNA was used to generate cDNA (Clontech SMARTER cDNA synthesis kit), cDNA was size selected (3-6 kb) by BluePippin, and 400 ng of SMRT-Bell library was prepared as above. The library was sequenced on the Pacific Biosciences RSII platform (P6v2 chemistry), 532,941 reads were generated with a mean insert length of 3,004 bp. Quiver^10^ was used to predict consensus isoforms and for polishing. There were 31,076 high-quality isoforms, and 11,661 low-quality isoforms with average consensus read length of 3,042 bp.

### Sample preparation and sequencing – Illumina short read

Genomic DNA was fragmented and Illumina whole genome libraries were constructed using the methods described in Hodges et al., 2009 ^38^. Steps 1-28 were followed to produce a whole genome library that was then sequenced rather than used in the enrichment portion of the protocol. The library was quantified by QPCR and sequenced on six lanes of an Illumina HiSeq GAIIX (Illumina, San Diego, CA, USA) using a 100 base paired end sequencing protocol.

### Sample preparation and sequencing – Bionano optical mapping

#### DNA Isolation

High-molecular weight DNA was extracted from mouse spleen. 70 mg of mouse frozen spleen tissue was place on a Petri dish over ice and chopped with a razor blade into approximately 2 mm chunks. The tissue was then transferred into a 15 mL conical. 1 mL of fixing solution (2% (v/v) formaldehyde, 10 mM Tris, 10 mM EDTA, 100 mM NaCl, pH 7.5) and left on ice for 30 minutes. Fixing solution was pipetted out and discarded. Tissue was washed 3 times by adding 2 MB Buffer (10 mM Tris, 10 mM EDTA, 100 mM NaCl, pH 9.4), swirling tube, and pipetting off MB Buffer. 2 mL of MB Buffer was added after the third wash. The tissue was blended using a fixed rotor-stator homogenizer (TissueRuptor, Qiagen #9001271) on high speed for 10 seconds. The homogenate was transferred to a 2 mL microfuge tube and spun down at 2000 rcf for 5 min. at 4°C. Supernatant was removed, pellet was resuspended in 1.5 mL of MB Buffer, and spin was repeated. Supernatant was removed and final pellet was resuspended in MB Buffer.

Resuspended cells were embedded into low-melting point agarose gel plugs, using the CHEF Mammalian Genomic DNA Plug Kit (BioRad #170-3591), following the manufacturer’s instructions. Plugs were made with 10 mg, 15, mg, and 20 mg equivalents of the original starting material (mass equivalents calculated based on volume of final resuspended pellet). The plugs were incubated with Lysis Buffer (Bionano #20270) and Puregene Proteinase K (Qiagen #1588920) overnight at 50°C, then again the following morning for 2 hrs. (using new buffer and Proteinase K). The plug was washed, melted, and solubilized with GELase (Epicentre #G09200). The purified DNA was subjected to 4 hrs. of drop dialysis (Millipore, #VCWP04700) and quantified using the Quant-iT PicoGreen dsDNA Assay Kit (Invitrogen/Molecular Probes #P11496). The plug made with 10 mg equivalents of starting material had a concentration of 286 ng/µL and was clear and viscous, so it was selected for further processing.

#### DNA Fluorescent Labeling

DNA was labeled according to commercial protocols using the NLRS kit (Bionano Genomics, #80001). Briefly, 300 ng of purified genomic DNA was nicked with 7 U nicking endonuclease Nt.BspQI (New England BioLabs (NEB), #R0644) at 37°C for two hrs. in NEBuffer3. The nicked DNA was labeled with a fluorescent-dUTP nucleotide analog using Taq DNA Polymerase (NEB, #M0267) for one hr. at 72°C. After labeling, the nicks were ligated with Taq DNA Ligase (NEB, #M0208) in the presence of dNTPs. The backbone of fluorescently labeled DNA was counterstained using the DNA Stain from the NLRS DNA Labeling Kit.

#### Data Collection

The labeled DNA was loaded onto Irys chips (Bionano, #20247) and inserted into the Irys instrument. The instrument automated the electrophoresis of the DNA into nanochannels, thereby linearizing them with uniform stretch throughout the molecule. The stationary molecules were then imaged, and the automated process of electrophoresis followed by imaging was repeated for multiple cycles until the desired amount of data was collected. The stained DNA molecule backbones and locations of fluorescent labels along each molecule were automatically detected using the in-house software package, IrysView. A total of 244 Gbp (~80X coverage depth) of data were generated, using 3 Irys chips.

### Processing raw data - PacBio

#### Quality trimming of sequenced reads

Raw data from 294 SMRT cells were imported into the SMRT portal (http://files.pacb.com/Training/IntroductiontoSMRTPortal/story_content/external_files/Introduction <u>%20to%20SMRT%C2%AE%20Portal%20.pdf)</u> and subreads were extracted from the raw h5 files within PacBio SMRT portal. Subreads with polymerase read were further filtered according to the following criterion: quality < 75, read length < 50 and polymerase read length < 50.

Finally, after filtering, 32,210,376 subreads (mean subread length of 5,753) were extracted from 20,081,751 raw reads, providing theoretical coverage of 66X for *de novo* assembly.

#### Error correction

Error correction of reads was accomplished with the MinHash Alignment Process (MHAP)^39^ within PBcR. Continuous benchmarking of correction parameters (k-mer size, hash size, min-mer size, error rate) was done to obtain the best possible set of corrected subreads. Our analysis indicated that usage of more sensitive parameters (MhapSensitivity=high

OvlErrorRate=0.05) significantly increased run time but overall improved the quality of corrected reads.

### Sequence assembly - PacBio

Corrected reads from MHAP were assembled using the Celera assembler (CA 8.3) (default parameters)^40^, which requires ~60-70 gigabases of corrected sequence and consists of overlapper, unitigger, scaffolder and consensus steps to reconstruct genomes from corrected long reads.

### Hybrid scaffolding

Single molecule high-resolution maps of the B6Eve genome were obtained using the Bionano Irys System ^41^. Label positions captured in images and molecule map lengths were stored in CMAP format files (consensus map). The hybrid scaffold tool from Bionano genomics was used to further extend the scaffold size by combining the PacBio *de novo* assembly and genome map data of B6Eve. The hybrid scaffold pipeline created an alignment between the datasets and constructed super scaffolds excluding the conflicting alignments^33^.

### Polishing and assembly evaluation

The hybrid scaffolded assembly was polished using Quiver ^10^ to improve consensus accuracies in the range of Q60 and to reduce the high indel errors that are expected in the PacBio sequencing data ^9^. pbalign (https://github.com/PacificBiosciences/pbalign) was used to create a bam file from all h5 files of SMRT cells and Quiver trained on P6-C4 chemistries were used to obtain the consensus corrected assembly. This assembly was further improved by using Pilon^11^ with default parameters, that corrects bases, fixes mis-assemblies and fills gaps provided a draft assembly and paired-end Illumina sequencing data. Nearly 32X of Illumina data from B6Eve was used as an input to Pilon to fix the bases in B6Eve assembly. The GATK variant calling pipeline (following best practices) was used to call variant using Illumina data on the B6Eve assembly to judge the improvement in overall quality at each step of polishing. Finally, the QUAST tool ^12^ was used (--split-scaffolds) to compare the final assembly with GRCm38. The split-scaffolds option breaks the assembly and performs reconstruction of "contigs" which were used to build the scaffolds to compare the effectiveness of scaffolding.

### RefSeq Transcript Alignments

Murine “known” RefSeq transcripts (those with NM and NR prefixes) were queried from NCBI Entrez on September 11, 2017 and aligned to the Pilon-corrected B6Eve assembly and GRCm38 full assembly (GCF_000001635.20). From these analyses, the counts of transcripts with low quality alignments, split alignments or no alignment to the GRCm38 primary assembly unit and B6Eve were determined, as were the counts of transcripts dropped for co-location, as described on p.52 of the Supplementary Methods of Shi et al., 2016 ^14^. From the same set of RefSeq transcripts, we additionally identified alignments to GRCm38 and to the Pilon-corrected B6Eve containing frameshifting and non-frameshifting indels in CDS ^13^. The frameshift analysis of the pre-polished/corrected assembly used a set of known RefSeq transcripts queried on February 28, 2016.

### LiftOver construction

LiftOver was performed between the Eve assembly and GRCm38 reference genome using the samespecies lift over construction procedure^9^ outlined by University of California Santa Cruz Genome Bioinformatics Group. Same species lift over construction contained two steps a) BLAT alignments and b) chaining and netting to obtain the lift over file. Genome loci of the Eve assembly were further converted into GRCm38 coordinates using the LiftOver^8^ tool from UCSC utilities.

### Repeat content assessment

The repeat elements in the GRCm38.p6 (excluding chrY and alternate loci scaffolds) and B6Eve assembly were determined by RepeatMasker^3^ trained on the mouse model by excluding RNA elements (-norna). A chi-square test was performed to identify the repeat classes that are enriched in one genome over the other.

### Repeat analysis of unaligned B6Eve sequences

There were unaligned B6Eve sequences from three different alignment methods a) Cactus based alignment using UCSC Comparative Annotation Toolkit which was used for the B6Eve annotation, b) NCBI BLAST-based alignment of B6Eve to GRCm38, and c) QUAST (minimap2 aligner). A consensus set (common among all three set), was constructed using BEDTools ^42^. RepeatMasker was used to identify the repeat content in the common unaligned region. A chi-square test was used to test the differences in repeat content for each of the repeat classes of common unaligned sequence against GRCm38.

### Resolving recurring variants

Recurring variants present in ≥ 75% of strains, detected in previous whole exome sequencing efforts ^10^ were extracted. Fixed (homozygous) variants from the mouse Collaborative Cross genome^11^ project, as well as fixed variants from 24 C57BL/6J pedigrees descendent from Eve^12^ were also obtained (unpublished data). We used LiftOver to remap the genomic coordinates, and a recurring variant was said to be resolved if the ALT allele of a recurring variant in exome data matches the REF allele in the B6Eve assembly. The analysis was restricted to only homozygous variant calls.

### B6Eve annotation

B6Eve was annotated using the Comparative Annotation Toolkit ^32^ (https://github.com/ComparativeGenomicsToolkit/Comparative-Annotation-Toolkitcommitc7852b4). As input, CAT was given a progressive Cactus alignment generated with rat rn6 (GCA_000001895.4) and human GRCh38 as outgroups as well as the GENCODE VM11 annotation on mouse GRCm38. CAT was provided with extrinsic transcript information from RNA-seq as well as IsoSeq. For mouse GRCm38, the same RNA-seq used in the CAT publication was used. RNA-seq data were generated from whole brain of a female C57BL/6J Eve descendent and aligned to the B6Eve assembly. To guide AugustusPB in detecting novel isoforms, a total of 26,188 IsoSeq full length cDNA reads were aligned to the B6Eve assembly.

### Novel isoform detection

To detect novel isoforms, homGeneMapping ^43^ was used to map GENCODE VM11 annotation coordinates onto B6Eve, and these splice junction coordinates compared to AugustusPB and AugustusCGP transcript predictions filtered for IsoSeq support. Transcripts with annotation support were filtered out. The remaining candidate novel isoforms then were checked to see if they overlapped a comparatively annotated locus and if they contained either a fully novel exon or a splice site shift based on bedtools^21^ intersections.

### SV detection & gap filling

#### PacBio read alignment

Raw PacBio reads were aligned to GRCm38 using the long-read aligner NGMLR version 0.2.6. CoNvex Gap-cost alignments for Long Reads (NGMLR) ^22^ is a long-read aligner designed to align PacBio reads with the focus on identifying structural variations. Stringent alignment requirements were used for identifying SVs: -i 0.85 argument to disregard alignments with identity with less than 85% and -R 0.5 option to ignore alignments containing less than 50% of the read length.

#### Structural variant calling

Sniffles ^22^ (default parameters) was used to call SVs from the alignments produced by NGMLR. It identified 229 duplications, 418 deletions and 36 inversions, 3,394 insertions, and 71 translocations. GATK was used to process Illumina WGS data from B6Eve. Best practices were used to generate a BAM file and SVS were called using Delly v0.7.7. SURVIVOR-1.0.3^44^ package was used to perform the integration of PacBio and Illumina calls.

#### Gap filling

We extracted the coordinates of the gaps from the GRCm38 chromosomes and further extended this to include the 50 Kb flanking both sides of the gap. We aligned B6Eve scaffolds to these padded regions using minimap2. We filtered the candidate alignments according to the following criterion: a) Must be the reciprocal best hit, b) Total alignment length >= 80KB, and c) Align to one unique location to the reference (extracted this information from assembly-assembly alignments). The retained alignments were further visualized in Integrated Genomic Viewer (IGV) to inspect insertion/deletion patterns around the gap region. To confirm and extract the gap spanning a B6Eve scaffold, we performed the reciprocal alignments, aligning the padded gap regions to B6Eve scaffolds, using minimap2. We filtered out candidate alignments not satisfying criteria mentioned above and visualized the retained alignments in IGV to inspect whether we observe the opposite of previously found insertion/deletion pattern. The sequence and locus of confirmed gap spanning B6Eve scaffolds were extracted and subjected to GRC internal curation.

### Data deposition

This Whole Genome Shotgun project has been deposited at DDBJ/ENA/GenBank under the accession LXEJ00000000. The version described in this paper is version LXEJ02000000. The raw PacBio, Illumina and Bionano data used were deposited at NCBI BioProject under accession PRJNA318985. The B6Eve assembly along with annotation and an assembly hub are available at ftp link (ftp://ftp.jax.org/b6eve). Visualization of the assembly can be found at https://genome.ucsc.edu → MyData →Track Hubs → My Hubs with the following URL: ftp://ftp.jax.org/b6eve/assemblyhub/hub.txt. A file (LXEJ02_contigs.tsv) with mapping of B6Eve scaffold names to GenBank accession is also available at ftp.

## Author contributions

Conceptualization, A.S., G.C., M.V.W., D.H., G.W., L.G.R. Methodology, A.S., V.A.S., I.T.F. Validation, L.G.R., O.Z., V.K.S., N.R. Formal Analysis, V.K.S., A.S., N.R., I.T.F., F.T-N., M.By., K.H. Investigation, X.R., L.R., M.B., M.By.,A.H. Resources, L.R., M.G., L.G.R., G.A. Data Curation, V.K.S., A.S. Writing, original draft, V.K.S., A.S., V.A.G, I. T. F., L.G.R. Writing, editing, V.K.S., A.S., V.A.G., L.G.R., M.V.W., L.R., K.H., G.C., G.W. Visualization, I.T.F., V.A.S., L.G.R. Supervision, A.S., L.G.R., B.P., V.A.G. Project Administration, A.S., L.G.R. Funding Acquisition, A.S., G.C., M.V.W., D.H., G.W., L.G.R.

## Acknowledgements

LGR, VKS, AS, NR, and OZ were supported in part by NIH R24 OD02135 awarded to LGR and The Jackson Laboratory. The work of VAS and F. T-N was supported by the intramural research program of the National Library of Medicine, National Institutes of Health. We are grateful to the services provided by The Jackson Laboratory Genome Technologies and Computational Sciences Core, which are supported by a grant to The Jackson Laboratory Cancer Center, NCI P30 CA034196.

## Conflict of Interests

The authors declare no competing interests

## Supplementary Figures

**Supplementary Figure 1**: Non-N B6Eve coverage of GRCm38 chromosome assemblies showing that all the GRCm38 chromosomes (excluding chrY as we have female mouse) have 90% coverage in B6Eve assembly

**Supplementary Figure 2: a)** Figure showing IGV snapshot of B6Eve scaffold (#8912) partially filling a gap sized at 1,760 bp in chr2 (nt 172,624,657-172,626,416) with a deletion of 136 bp at the beginning of the gap **b)** Figure showing IGV snapshot of B6Eve scaffold (#2171) completely filling a gap of size 99bp in chr1 (183334907-183335006) by inserting 595bp at the beginning of the gap.

**Supplementary Figure 3:** The gene *Traf5* has a 50kb assembly gap between the last three exons and the remainder of the transcript in GRCm38 (bottom panel). In the B6Eve assembly, this gap is closed to 2,430 bp. Additionally, IsoSeq shows that this gap overlaps an exon that was included as a new isoform in the CAT annotation. BLAT alignment of the isoform back to the reference shows the inserted sequence.

**Supplementary Figure 4:** The gene *Slc26a6* has a 50kb assembly gap between the last exon and the remainder of the transcript in GRCm38 (bottom panel). In the B6Eve assembly, this gap is closed to 1,810bp (top panel). Additionally, IsoSeq shows that this gap overlaps an exon and removes an additional exon. Using the IsoSeq information CAT generated a new isoform that matches the IsoSeq alignment.

## Supplementary Tables

**Supplementary Table 1:** Summary PacBio and Illumina sequencing data.

**Supplementary Table 2:** Details for RefSeq transcript alignments to B6Eve and GRCm38

**Supplementary Table 3:** Progress of Assembly at each step

**Supplementary Table 4:** Coordinates of filled gaps with sequence in GRCm38 with B6Eve assembly

**Supplementary Table 5:** Locus of resolved variants in Exome, CC and Pedigree data, respectively

**Supplementary Table 6:** Coordinates of SVs common between Illumina and PacBio based analysis

**Supplementary Table 7a:** The complete distribution of classification of repeats in GRCm38

**Supplementary Table 7b**: The complete distribution of classification of repeats of B6Eve assembly

**Supplementary Table 7c:** The complete distribution of classification of repeats of Illumina GCA_000185125

**Supplementary Table 7d:** The complete distribution of classification of repeats of Illumina GCA_000185105

**Supplementary Table 8:** Coordinates of B6Eve splice site shifts relative to GRCm38, potential novel exons and loci

## Supplementary Files

**Supplementary File 1:** Variant calls made using the Illumina data (VCF file) of B6Eve to Pilon corrected assembly

**Supplementary File 2:** A detailed QUAST evaluation report (HTML format) reflecting the quality of the polished assembly

**Supplementary File 3:** Regions of the B6Eve assembly (BED file) remained unaligned to GRCm38 by a) Cactus based alignment using UCSC Comparative Annotation Toolkit b) NCBI assembly-assembly alignments and c) QUAST (minimap2 aligner)

