## Supplementary figures and images for "The genome of C57BL/6J “Eve”, the mother of the laboratory mouse genome reference strain"

### Supplementary Figure 1

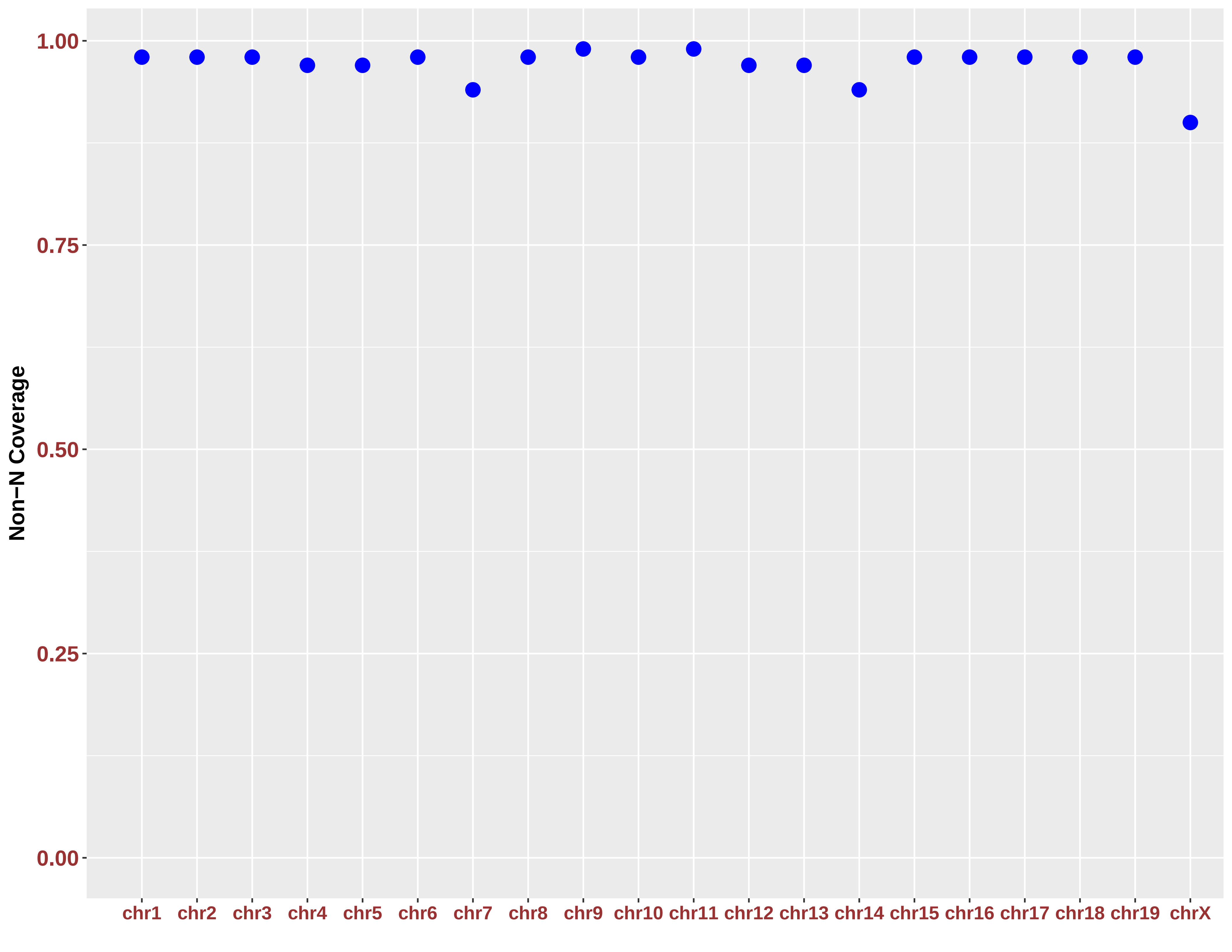

### Supplementary Figure 2

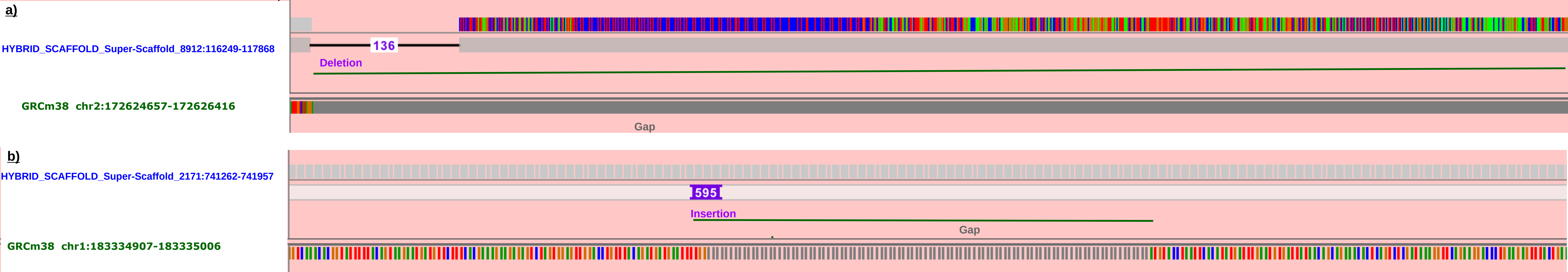

### Supplementary Figure 3

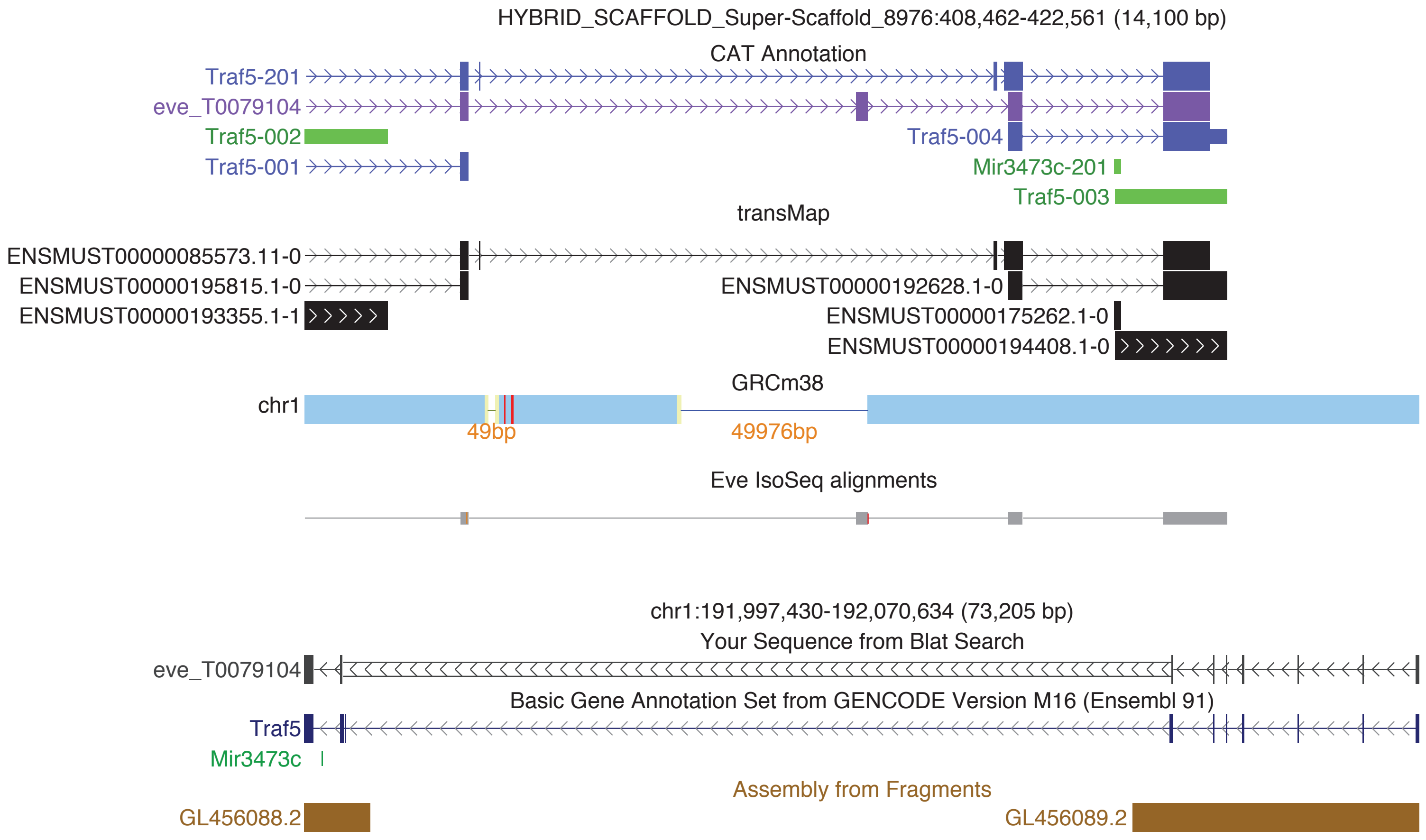

### Supplementary Figure 4

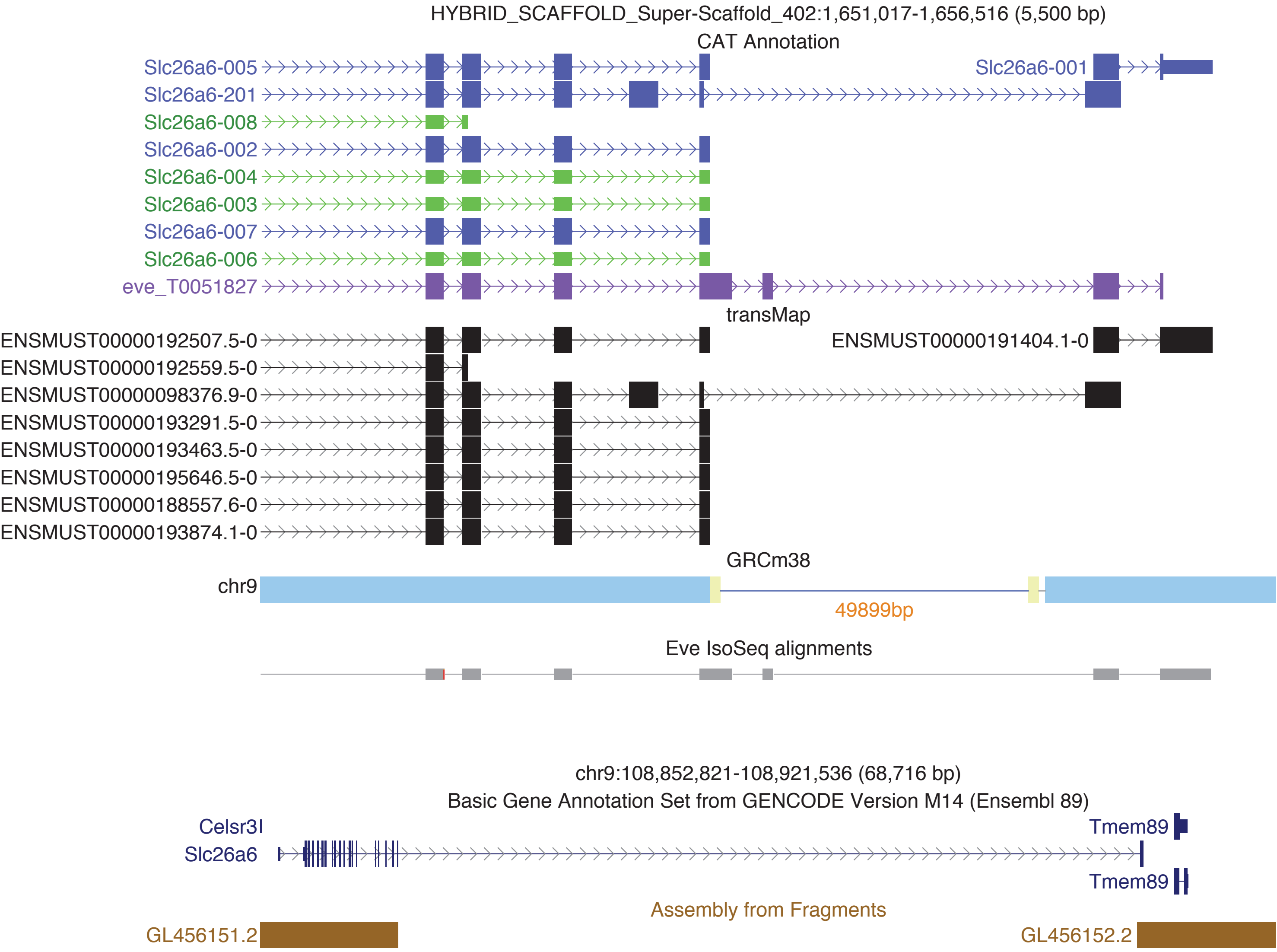
