## Supplementary File 2 for "The genome of C57BL/6J “Eve”, the mother of the laboratory mouse genome reference strain"

### QUAST

**Quality Assessment Tool for Genome Assemblies**
by CAB

Loading...

Aligned to
""

Combined reference
Estimated reference size:
 bp
|
 bp
|
 references
|
 fragments
|
 % G+C
|
 chromosomes
  
 reads
|
% mapped
|
% properly paired
  
 genes
|
 operons

Unfortunately, JavaScript in your
browser is disabled or is not supported.
We need JavaScript to build report and plots.

Worst
Median
Best

Show heatmap

|  |
| --- |
| Contigs are ordered from largest (contig #1) to smallest.  FRCurve: Y is the total number of aligned bases divided by the reference length, in the contigs having the total number of  at most X.  Contigs are broken into nonoverlapping 100 bp windows. Plot shows number of windows for each GC percentage.  Plot shows number of contigs with GC percentage in a certain range. |

{"subreports":[],"report":[["Genome statistics",[{"values":["96.848","96.841"],"quality":"More is better","isMain":true,"metricName":"Genome fraction (%)"},{"values":["1.123","1.017"],"quality":"Less is better","isMain":true,"metricName":"Duplication ratio"},{"values":["22994 + 2144 part","22953 + 2182 part"],"quality":"More is better","isMain":true,"metricName":"# genomic features"},{"values":[5570225,4577372],"quality":"More is better","isMain":true,"metricName":"Largest alignment"},{"values":[2526660255,2525333977],"quality":"More is better","isMain":true,"metricName":"Total aligned length"},{"values":[1390099,371965],"quality":"More is better","isMain":false,"metricName":"NG50"},{"values":[439613,151056],"quality":"More is better","isMain":false,"metricName":"NG75"},{"values":[422265,384295],"quality":"More is better","isMain":false,"metricName":"NA50"},{"values":[115689,168024],"quality":"More is better","isMain":false,"metricName":"NA75"},{"values":[476164,365098],"quality":"More is better","isMain":true,"metricName":"NGA50"},{"values":[146696,146364],"quality":"More is better","isMain":false,"metricName":"NGA75"},{"values":[524,1757],"quality":"Less is better","isMain":false,"metricName":"LG50"},{"values":[1281,4525],"quality":"Less is better","isMain":false,"metricName":"LG75"},{"values":[1516,1657],"quality":"Less is better","isMain":false,"metricName":"LA50"},{"values":[4821,4140],"quality":"Less is better","isMain":false,"metricName":"LA75"},{"values":[1336,1793],"quality":"Less is better","isMain":true,"metricName":"LGA50"},{"values":[3885,4626],"quality":"Less is better","isMain":false,"metricName":"LGA75"}]],["Reads mapping",[]],["Misassemblies",[{"values":[2590,1306],"quality":"Less is better","isMain":true,"metricName":"# misassemblies"},{"values":[1735,557],"quality":"Less is better","isMain":false,"metricName":" # relocations"},{"values":[827,724],"quality":"Less is better","isMain":false,"metricName":" # translocations"},{"values":[28,25],"quality":"Less is better","isMain":false,"metricName":" # inversions"},{"values":[1714,1166],"quality":"Less is better","isMain":false,"metricName":"# misassembled contigs"},{"values":[1509562627,220209949],"quality":"Less is better","isMain":true,"metricName":"Misassembled contigs length"},{"values":[3529,2962],"quality":"Less is better","isMain":false,"metricName":"# local misassemblies"},{"values":[88,null],"quality":"Less is better","isMain":false,"metricName":"# scaffold gap size mis."},{"values":[18,16],"quality":"Equal","isMain":false,"metricName":"# possible MGEs"},{"values":[69,51],"quality":"Less is better","isMain":false,"metricName":"# unaligned mis. contigs"}]],["Unaligned",[{"values":[166,21642],"quality":"Less is better","isMain":false,"metricName":"# fully unaligned contigs"},{"values":[3719105,3869437],"quality":"Less is better","isMain":false,"metricName":"Fully unaligned length"},{"values":[2456,1793],"quality":"Less is better","isMain":false,"metricName":"# partially unaligned contigs"},{"values":[7251439,6820765],"quality":"Less is better","isMain":false,"metricName":"Partially unaligned length"}]],["Mismatches",[{"values":[321280,320246],"quality":"Less is better","isMain":false,"metricName":"# mismatches"},{"values":[494061,492931],"quality":"Less is better","isMain":false,"metricName":"# indels"},{"values":[3818564,2606030],"quality":"Less is better","isMain":false,"metricName":"Indels length"},{"values":["12.93","12.89"],"quality":"Less is better","isMain":true,"metricName":"# mismatches per 100 kbp"},{"values":["19.89","19.85"],"quality":"Less is better","isMain":true,"metricName":"# indels per 100 kbp"},{"values":[465708,464749],"quality":"Less is better","isMain":false,"metricName":" # indels (<= 5 bp)"},{"values":[28353,28182],"quality":"Less is better","isMain":false,"metricName":" # indels (> 5 bp)"},{"values":[263669715,422],"quality":"Less is better","isMain":false,"metricName":"# N's"},{"values":["9412.42","0.02"],"quality":"Less is better","isMain":true,"metricName":"# N's per 100 kbp"}]],["Statistics without reference",[{"values":[12658,36182],"quality":"Equal","isMain":true,"metricName":"# contigs"},{"values":[12661,null],"quality":"Equal","isMain":false,"metricName":"# contigs (>= 0 bp)"},{"values":[12661,null],"quality":"Equal","isMain":false,"metricName":"# contigs (>= 1000 bp)"},{"values":[12656,14704],"quality":"Equal","isMain":false,"metricName":"# contigs (>= 5000 bp)"},{"values":[12645,14693],"quality":"Equal","isMain":false,"metricName":"# contigs (>= 10000 bp)"},{"values":[9722,11770],"quality":"Equal","isMain":false,"metricName":"# contigs (>= 25000 bp)"},{"values":[6787,8835],"quality":"Equal","isMain":false,"metricName":"# contigs (>= 50000 bp)"},{"values":[9177662,4577433],"quality":"More is better","isMain":true,"metricName":"Largest contig"},{"values":[2801295686,2537626393],"quality":"More is better","isMain":true,"metricName":"Total length"},{"values":[2801301292,null],"quality":"More is better","isMain":false,"metricName":"Total length (>= 0 bp)"},{"values":[2801301292,null],"quality":"More is better","isMain":true,"metricName":"Total length (>= 1000 bp)"},{"values":[2801286582,2537466957],"quality":"More is better","isMain":false,"metricName":"Total length (>= 5000 bp)"},{"values":[2801206759,2537387134],"quality":"More is better","isMain":true,"metricName":"Total length (>= 10000 bp)"},{"values":[2741388976,2477569351],"quality":"More is better","isMain":false,"metricName":"Total length (>= 25000 bp)"},{"values":[2638542761,2374723136],"quality":"More is better","isMain":true,"metricName":"Total length (>= 50000 bp)"},{"values":[1291016,391273],"quality":"More is better","isMain":false,"metricName":"N50"},{"values":[233769,172412],"quality":"More is better","isMain":false,"metricName":"N75"},{"values":[585,1624],"quality":"Less is better","isMain":false,"metricName":"L50"},{"values":[1674,4053],"quality":"Less is better","isMain":false,"metricName":"L75"},{"values":["41.80","41.80"],"quality":"Equal","isMain":false,"metricName":"GC (%)"}]],["K-mer-based statistics",[{"values":["97.38","97.38"],"quality":"More is better","isMain":true,"metricName":"K-mer-based completeness"},{"values":["74.76","87.29"],"quality":"More is better","isMain":false,"metricName":"K-mer-based cor. length (%)"},{"values":["24.45","11.86"],"quality":"Less is better","isMain":false,"metricName":"K-mer-based mis. length (%)"},{"values":["0.78","0.85"],"quality":"Less is better","isMain":false,"metricName":"K-mer-based undef. length (%)"},{"values":[1568,1493],"quality":"Less is better","isMain":true,"metricName":"# k-mer-based misjoins"},{"values":[1091,1024],"quality":"Equal","isMain":false,"metricName":" # k-mer-based translocations"},{"values":[477,469],"quality":"Equal","isMain":false,"metricName":" # k-mer-based 100kbp relocations"}]],["Predicted genes",[]],["Similarity statistics",[{"values":[9448,9448],"quality":"Equal","isMain":false,"metricName":"# similar correct contigs"},{"values":[3598,3598],"quality":"Equal","isMain":false,"metricName":"# similar misassembled blocks"}]],["Reference statistics",[{"values":[2639127076,2639127076],"quality":"Equal","isMain":false,"metricName":"Reference length"},{"values":[65,65],"quality":"Equal","isMain":false,"metricName":"Reference fragments"},{"values":["41.76","41.76"],"quality":"Equal","isMain":false,"metricName":"Reference GC (%)"},{"values":[25759,25759],"quality":"Equal","isMain":false,"metricName":"Reference genomic features"}]]],"assembliesWithNs":null,"referenceName":"GRCm38\_p5\_no\_chrY","date":"24 January 2018, Wednesday, 23:03:33","subreferences":[],"minContig":3000,"order":[0,1],"assembliesNames":["B6Eve","B6Eve\_broken"]}

{{ qualities }}

{{ mainMetrics }}

{"lists\_of\_lengths":[[148208263,101229800,90479788,82953928,77123851,71229642,66687435,62554156,58713099,55640244,53580815,50726652,48927206,46764613,44510324,42326681,41031336,39814437,37832458,36361922,35184222,33761653,32538799,31765118,30903947,30367014,29526310,28624730,27960719,27080580,26179928,25230518,24370512,23359866,22485558,21801690,21063218,20410129,19808094,19172013,18527921,17664925,16739414,16048388,15325602,14722504,14146201,13654079,13196272,12633396,12080739,11568845,10932462,10388818,9869157,9364570,8892506,8439980,8065517,7669859,7407238,7142018,6875525,6670350,6487358,6266908,6111826,5942714,5791175,5637698,5501227,5354436,5191301,5040032,4921950,4799885,4689509,4603659,4527010,4455649,4389210,4320236,4274611,4213866,4166189,4120765,4088350,4047331,4012184,3976970,3945151,3916094,3890053,3863546,3841313,3812068,3785739,3758242,3729120,3706735,3686140,3661036,3637663,3616677,3594438,3572145,3547704,3524607,3502874,3478206,3461294,3438628,3414794,3391384,3370432,3352126,3337045,3314165,3297837,3277329,3249540,3226011,3206920,3189652,3169129,3148721,3128728,3106664,3083379,3066876,3050378,3029860,3006610,2990948,2971629,2952970,2931869,2912583,2892851,2878828,2866724,2856605,2842068,2825171,2812199,2794571,2779764,2764537,2750740,2734557,2717547,2703928,2689861,2675730,2662475,2646534,2632172,2617909,2601490,2588606,2573412,2555250,2537127,2520943,2507193,2493064,2481011,2469266,2457432,2441137,2426434,2409785,2392321,2371566,2354780,2338222,2321695,2304941,2289381,2274873,2264341,2248863,2232734,2220620,2206459,2196013,2185320,2172964,2160714,2145206,2132237,2119694,2106845,2097311,2087348,2074160,2059686,2046578,2029374,2014293,2003439,1991493,1981038,1970992,1958305,1948613,1940478,1927303,1910868,1899935,1889283,1876151,1863950,1852654,1842385,1831612,1820844,1808165,1796559,1784406,1773466,1763404,1749349,1736491,1725388,1715869,1706914,1694573,1680772,1671107,1662109,1652399,1643791,1634229,1621455,1611107,1601004,1592522,1585943,1578197,1571067,1562795,1552286,1542962,1533908,1525355,1515501,1504412,1493233,1483346,1471454,1461651,1455043,1447655,1441683,1434036,1425507,1413630,1406877,1398938,1391866,1382906,1373715,1363590,1354744,1346125,1336908,1328179,1319674,1313047,1304294,1297730,1289118,1279173,1270092,1259448,1250297,1242761,1236173,1229229,1219512,1210072,1201905,1195744,1189186,1179748,1171792,1163289,1156264,1149775,1141237,1134134,1126412,1119935,1115002,1107906,1102418,1096079,1089381,1082258,1075166,1066948,1059904,1051839,1044076,1037661,1029092,1022369,1015674,1009360,1004408,997842,991028,983760,976599,970136,963550,958863,951750,943962,934486,926503,919884,916051,911356,905847,900508,893742,887400,881350,875279,870837,864906,860710,856939,851198,843728,838062,833897,830370,825349,820858,815616,810102,804273,798874,794393,788777,782687,778035,774085,770765,767315,763454,759762,756760,753775,750478,746710,742926,739297,735461,731186,726420,722305,717607,713858,709717,705234,701453,698095,694634,691542,688153,684773,681791,678790,676467,674138,670853,668183,665031,661730,658109,654610,651711,648830,645815,642827,640068,637972,635220,632497,629783,626876,623842,621115,618356,615513,613169,611028,608479,606019,604126,601124,598894,595983,593956,591615,589617,587392,585165,582690,580879,578976,576984,574780,571814,569210,566611,564211,561801,560049,558173,556082,554197,551731,549516,547701,545714,543889,541915,540212,538105,536510,535282,533590,531425,529753,528038,526341,524460,522535,520413,518518,517133,514952,513442,511903,510106,508184,506709,504911,503207,501956,500724,499616,497909,496122,494802,493495,492375,490969,489180,487336,486039,484552,482894,481736,480267,478724,477224,475925,474358,472803,471465,470186,469193,467443,465957,464536,462864,461824,460460,459043,457835,456471,455490,454627,453359,452365,451196,449989,448387,446729,445619,444335,443201,441852,440592,439204,438163,436920,435609,433983,432928,431718,430663,429753,428634,427672,426754,425460,424251,423002,421568,420432,419562,418478,416917,415652,414630,413653,412546,411651,410220,363445,63945],[71575825,54596436,47521487,42204996,38013644,35481179,32909482,31353391,30153069,28620183,27425155,26320517,25482364,24202212,23083929,22260960,21578921,20985233,20489736,19985110,19354599,18884520,18380252,17989713,17460443,17111288,16822691,16539818,16200553,15904797,15556681,15297927,15050211,14792817,14526714,14218012,13975183,13754261,13524121,13358614,13129382,12936695,12700218,12517265,12390682,12217268,12054390,11878339,11746568,11588416,11422889,11265031,11100637,10992337,10872716,10728069,10613107,10495205,10343557,10240875,10150375,10055948,9922369,9810637,9702990,9596319,9499243,9409194,9312023,9215527,9145152,9064710,8991950,8898246,8817689,8750234,8666690,8564749,8478632,8397712,8329925,8258054,8193423,8128226,8040943,7975898,7914771,7841422,7758850,7685097,7601690,7522209,7454388,7394747,7351425,7289461,7213198,7156248,7099991,7040557,6992320,6937040,6879274,6809340,6760071,6714120,6671612,6622946,6568019,6502884,6429169,6356996,6296302,6245098,6194242,6150107,6110066,6060602,6004273,5966948,5926056,5879839,5828329,5782124,5732100,5682623,5639296,5592174,5554533,5521295,5476538,5439685,5396990,5353640,5304235,5248425,5201268,5158927,5114138,5076696,5035058,4996518,4968409,4937379,4898520,4865613,4829621,4796970,4765022,4730676,4691951,4660993,4635394,4603005,4570435,4537439,4504661,4473695,4439859,4417389,4390097,4355704,4319062,4293120,4269854,4241627,4202708,4176593,4150179,4115427,4092342,4061496,4037845,4008237,3978959,3953199,3924897,3896790,3871728,3848099,3822289,3794752,3766515,3738863,3713556,3694066,3668582,3645152,3624430,3601082,3580206,3556320,3532036,3510024,3486048,3466689,3446140,3423552,3398454,3377447,3357895,3342869,3321163,3302893,3286210,3257544,3234093,3212441,3195325,3176503,3155374,3135695,3114541,3090152,3072417,3056539,3036614,3014001,2995877,2978334,2959754,2938181,2919506,2898597,2883190,2870583,2860113,2847650,2830349,2816401,2800576,2784976,2769231,2756520,2738767,2723545,2708352,2694607,2680201,2666936,2651846,2636754,2622614,2607235,2592768,2578612,2562215,2542510,2526022,2512021,2497432,2485354,2472438,2462333,2446385,2431415,2416314,2397027,2379053,2360193,2343158,2328102,2309708,2294596,2279192,2267776,2254799,2237875,2224694,2210981,2198675,2189781,2176974,2165077,2150339,2136393,2123926,2110711,2100299,2091037,2078997,2064011,2051168,2035480,2018645,2007427,1994898,1984504,1974515,1962848,1951512,1943038,1932201,1916471,1903153,1893190,1880877,1867526,1856080,1846403,1834869,1824378,1812797,1800014,1788686,1777057,1766520,1754798,1740347,1728695,1718947,1709775,1699472,1685044,1674097,1665154,1655485,1646599,1637458,1626214,1614217,1604451,1594783,1588391,1580655,1573670,1565474,1556144,1545507,1537237,1528133,1518970,1508330,1496607,1487102,1475030,1464729,1457117,1450122,1443761,1436654,1428674,1417324,1409021,1401658,1394190,1386008,1376892,1366948,1357626,1349329,1339467,1331328,1322119,1315568,1307250,1299549,1292352,1282253,1273584,1262896,1253270,1244750,1238421,1232081,1223076,1212592,1204599,1197583,1191909,1183067,1173999,1166054,1158593,1151994,1144352,1136128,1129301,1121548,1116884,1110331,1104194,1098132,1092013,1084144,1077924,1069607,1062103,1054871,1046165,1039953,1032029,1024541,1017864,1011269,1006024,1000353,993226,986134,979393,971927,965607,960367,954675,946593,937549,929017,921729,917107,913284,907604,902191,896299,889438,883408,877193,872326,866842,861829,858308,853628,845724,839995,835204,831513,827282,822222,817408,811915,806564,800307,796012,790909,784539,779270,775400,771699,768642,764878,760806,757669,754816,751602,748137,744017,740612,736735,732649,727960,723678,719124,715188,711133,706595,702802,699013,695894,692575,689372,685771,682786,679694,677274,675063,671854,669076,666033,662863,659391,655749,652612,649773,646816,643877,640860,638788,636048,633449,630747,627803,624946,621947,619218,616608,613757,611769,609382,606750,604856,602109,599656,596934,594638,592344,590237,588161,586020,583347,581523,579683,577583,575662,572806,570020,567475,564970,562618,560542,558910,556648,554942,552583,550122,548367,546346,544503,542587,540732,538914,536899,535696,534167,532222,530216,528687,526881,525127,523165,521190,518993,517736,515627,513860,512484,510658,508851,507185,505636,503605,502387,501114,499987,498643,496554,495244,493916,492782,491436,489862,487901,486439,485087,483389,482149,480790,479200,477696,476395,474890,473257,471966,470564,469600,468077,466320,465091,463395,462143,460948,459513,458215,456909,455794,454905,453813,452719,451582,450402,448972,447212,445950,444816,443588,442274,441024,439685,438466,437338,436111,434488,433271,432121,430998,430019,429077,427944,427057,425901,424690,423415,422092,420724,419870,418863,417444,416085,414894,414053,412838,411998,410767,408757,155214,266,168,168,168,168,168,168,168,168,168,168,168,168,168,168,168,168,168,168,168,168,168,168,168,168,168,168,168,168,168,168,168,168,168,168,168,168,168,168,168,168,168,168,168,168,168,168,168,168,168,168,168,168,168,168,168,168,168,168,168,168,168,168,168,168,168,168,168,168,168,168,168,168,168,168,168,168,168,168,168,168,168,168,168,168,168,168,168,168,168,168,168,168,168,168,168,168,168,168,168,168,168,168,168,168,168,168,168,168,168,168,168,168,168,168,168,168,168,168,168,168,168,168,168,168,168,168,168,168,168,168,168,168,168,168,168,168,168,168,168,168,168,168,168,168,168,168,168,168,168,168,168,168,168,168,168,168,168,168,168,168,168,168,168,168,168,168,168,168,168,168,168,168,168,168,168,168,168,168,168,168,168,168,168,168,168,168,168,168,168,168,168,168,168,168,168,168,168,168,168,168,168,168,168,168,168,168,168,168,168,168,168,168,168,168,168,168,168,168,168,168,168,168,168,168,168,168,168,168,168,168,168,168,168,168,168,168,168,168,168,168,168,168,168,168,168,168,168,168,168,168,168,168,168,168,168,168,168,168,168,168,168,168,168,168,168,168,168,168,168,168,168,168,168,168,168,168,168,168,168,168,168,168,168,168,168,168,168,168,168,168,168,168,168,168,168,168,168,168,168,168,168,168,168,168,168,168,168,168,168,168,168,168,168,168,168,168,168,168,168,168,168,168,168,168,168,168,168,168,168,168,168,168,168,168,168,168,168,168,168,168,168,168,168,168,168,168,168,168,168,168,168,168,168,168,168,168,168,168,168,168,168,168,168,168,168,168,168,168,168,168,168,168,168,168,168,168,168,168,168,168,168,168,168,168,168,168,168,168,168,168,168,168,168,168,168,168,168,168,168,168,168,168,168,168,168,168,168,168,168,168,168,168,168,168,168,168,168,168,168,168,168,168,168,168,168,168,168,168,168,168,168,168,168,168,168,168,168,168,168,168,168,168,168,168,168,168,168,168,168,168,168,168,168,168,168,168,168,168,168,168,168,168,168,168,168,168,168,168,168,168,168,168,168,168,168,168,168,168,168,168,168,168,168,168,168,168,168,168,168,168,168,168,168,168,168,168,168,168,168,168,168,168,168,168,168,168,168,168,168,168,168,168,168,168,168,168,168,168,168,168,168,168,168,168,168,168,168,168,168,168,168,168,168,168,168,168,168,168,168,168,168,168,168,168,168,168,168,168,168,168,168,168,168,168,168,168,168,168,168,168,168,168,168,168,168,168,168,168,168,168,168,168,168,168,168,168,168,168,168,168,168,168,168,168,168,168,168,168,168,168,168,168,168,168,168,168,168,168,168,168,168,168,168,168,168,168,168,168,168,168,168,168,168,168,168,168,168,168,168,168,168,168,168,168,168,168,168,168,168,168,168,168,168,168,168,168,168,168,168,168,168,168,168,168,168,168,168,168,168,168,168,168,168,168,168,168,168,168,168,168,168,168,168,168,168,168,168,168,168,168,168,168,168,168,168,168,168,168,168,168,168,168,168,168,168,168,168,168,168,168,168,168,168,168,168,168,168,168,168,168,168,168,168,168,168,168,168,168,168,168,168,168,168,168,168,168,168,168,168,168,168,168,168,168,168,168,168,168,168,168,168,168,168,168,168,168,168,168,168,168,168,168,168,168,168,168,168,168,168,168,168,168,168,168,168,168,168,168,168,168,168,168,168,168,168,168,168,168,168,168,168,168,168,168,168,168,168,168,168,168,168,168,168,168,168,168,168,168,168,168,168,168,168,168,168,168,168,168,168,168,168,168,168,168,168,168,168,168,168,168,168,168,168,168,168,168,168,168,168,168,168,168,168,168,168,168,168,168,168,168,168,168,168,168,168,168,168,168,168,168,168,168,168,168,168,168,168,168,168,168,168,168,168,168,168,168,168,168,168,168,168,168,168,168,168,168,168,168,168,168,168,168,168,168,168,168,168,168,168,168,168,168,168,168,168,168,168,168,168,168,168,168,168]],"filenames":["B6Eve","B6Eve\_broken"]}

{"assemblies\_lengths":[2801295686,2537626393],"filenames":["B6Eve","B6Eve\_broken"]}

{"reflen":[2635568227,3167367,391482,0,0,0,0,0,0,0,0,0,0,0,0,0,0,0,0,0,0,0,0,0,0,0,0,0,0,0,0,0,0,0,0,0,0,0,0,0,0,0,0,0,0,0,0,0,0,0,0,0,0,0,0,0,0,0,0,0,0,0,0,0,0,0,0,0,0,0,0,0,0,0,0,0,0,0,0,0,0,0,0,0,0,0,0,0,0,0,0,0,0,0,0,0,0,0,0,0,0,0,0,0,0,0,0,0,0,0,0,0,0,0,0,0,0,0,0,0,0,0,0,0,0,0,0,0,0,0,0,0,0,0,0,0,0,0,0,0,0,0,0,0,0,0,0,0,0,0,0,0,0,0,0,0,0,0,0,0,0,0,0,0,0,0,0,0,0,0,0,0,0,0,0,0,0,0,0,0,0,0,0,0,0,0,0,0,0,0,0,0,0,0,0,0,0,0,0,0,0,0,0,0,0,0,0,0,0,0,0,0,0,0,0,0,0,0,0,0,0,0,0,0,0,0,0,0,0,0,0,0,0,0,0,0,0,0,0,0,0,0,0,0,0,0,0,0,0,0,0,0,0,0,0,0,0,0,0,0,0,0,0,0,0,0,0,0,0,0,0,0,0,0,0,0,0,0,0,0,0,0,0,0,0,0,0,0,0,0,0,0,0,0,0,0,0,0,0,0,0,0,0,0,0,0,0,0,0,0,0,0,0,0,0,0,0,0,0,0,0,0,0,0,0,0,0,0,0,0,0,0,0,0,0,0,0,0,0,0,0,0,0,0,0,0,0,0,0,0,0,0,0,0,0,0,0,0,0,0,0,0,0,0,0,0,0,0,0,0,0,0,0,0,0,0,0,0,0,0,0,0,0,0,0,0,0,0,0,0,0,0,0,0,0,0,0,0,0,0,0,0,0,0,0,0,0,0,0,0,0,0,0,0,0,0,0,0,0,0,0,0,0,0,0,0,0,0,0,0,0,0,0,0,0,0,0,0,0,0,0,0,0,0,0,0,0,0,0,0,0,0,0,0,0,0,0,0,0,0,0,0,0,0,0,0,0,0,0,0,0,0,0,0,0,0,0,0,0,0,0,0,0,0,0,0,0,0,0,0,0,0,0,0,0,0,0,0,0,0,0,0,0,0,0,0,0,0,0,0,0,0,0,0,0,0,0,0,0,0,0,0,0,0,0,0,0,0,0,0,0,0,0,0,0,0,0,0,0,0,0,0,0,0,0,0,0,0,0,0,0,0,0,0,0,0,0,0,0,0,0,0,0,0,0,0,0,0,0,0,0,0,0,0,0,0,0,0,0,0,0,0,0,0,0,0,0,0,0,0,0,0,0,0,0,0,0,0,0,0,0,0,0,0,0,0,0,0,0,0,0,0,0,0,0,0,0,0,0,0,0,0,0,0,0,0,0,0,0,0,0,0,0,0,0,0,0,0,0,0,0,0,0,0,0,0,0,0,0,0,0,0,0,0,0,0,0,0,0,0,0,0,0,0,0,0,0,0,0,0,0,0,0,0,0,0,0,0,0,0,0,0,0,0,0,0,0,0,0,0,0,0,0,0,0,0,0,0,0,0,0,0,0,0,0,0,0,0,0,0,0,0,0,0,0,0,0,0,0,0,0,0,0,0,0,0,0,0,0,0,0,0,0,0,0,0,0,0,0,0,0,0,0,0,0,0,0,0,0,0,0,0,0,0,0,0,0,0,0,0,0,0,0,0,0,0,0,0,0,0,0,0,0,0,0,0,0,0,0,0,0,0,0,0,0,0,0,0,0,0,0,0,0,0,0,0,0,0,0,0,0,0,0,0,0,0,0,0,0,0,0,0,0,0,0,0,0,0,0,0,0,0,0,0,0,0,0,0,0,0,0,0,0,0,0,0,0,0,0,0,0,0,0,0,0,0,0,0,0,0,0,0,0,0,0,0,0,0,0,0,0,0,0,0,0,0,0,0,0,0,0,0,0,0,0,0,0,0,0,0,0,0,0,0,0,0,0,0,0,0,0,0,0,0,0,0,0,0,0,0,0,0,0,0,0,0,0,0,0,0,0,0,0,0,0,0,0,0,0,0,0,0,0,0,0,0,0,0,0,0,0,0,0,0,0,0,0,0,0,0,0,0,0,0,0,0,0,0,0,0,0,0,0,0,0,0,0,0,0,0,0,0,0,0,0,0,0,0,0,0,0,0,0,0,0,0,0,0,0,0,0,0,0,0,0,0,0,0,0,0,0,0,0,0,0,0,0,0,0,0,0,0,0,0,0,0,0,0,0,0,0,0,0,0,0,0,0,0,0,0,0,0,0,0,0,0,0,0,0,0,0,0,0,0,0,0,0,0,0,0,0,0,0,0,0,0,0,0,0,0,0,0,0,0,0,0,0,0,0,0,0,0,0,0,0,0,0,0,0,0,0,0,0,0,0,0,0,0,0,0,0,0,0,0,0,0,0,0,0,0,0,0,0,0,0,0,0,0,0,0,0,0,0,0,0,0,0,0,0,0,0,0,0,0,0,0,0,0,0,0,0,0,0,0,0,0,0,0,0,0,0,0,0,0,0,0,0,0,0,0,0,0,0,0,0,0,0,0,0,0,0,0,0,0,0,0,0,0,0,0,0,0,0,0,0,0,0,0,0,0,0,0,0,0,0,0,0,0,0,0,0,0,0,0,0,0,0,0,0,0,0,0,0,0,0,0,0,0,0,0,0,0,0,0,0,0,0,0,0,0,0,0,0,0,0,0,0,0,0,0,0,0,0,0,0,0,0,0,0,0,0,0,0,0,0,0,0,0,0,0,0,0,0,0,0,0,0,0,0,0,0,0,0,0,0,0,0,0,0,0,0,0,0,0,0,0,0,0,0,0,0,0,0,0,0,0,0,0,0,0,0,0,0,0,0,0,0,0,0,0,0,0,0,0,0,0,0,0,0,0,0,0,0,0,0,0,0,0,0,0,0,0,0,0,0,0,0,0,0,0,0,0,0,0,0,0,0,0,0,0,0,0,0,0,0,0,0,0,0,0,0,0,0,0,0,0,0,0,0,0,0,0,0,0,0,0,0,0,0,0,0,0,0,0,0,0,0,0,0,0,0,0,0,0,0,0,0,0,0,0,0,0,0,0,0,0,0,0,0,0,0,0,0,0,0,0,0,0,0,0,0,0,0,0,0,0,0,0,0,0,0,0,0,0,0,0,0,0,0,0,0,0,0,0,0,0,0,0,0,0,0,0,0,0,0,0,0,0,0,0,0,0,0,0,0,0,0,0,0,0,0,0,0,0,0,0,0,0,0,0,0,0,0,0,0,0,0,0,0,0,0,0,0,0,0,0,0,0,0,0,0,0,0,0,0,0,0,0,0,0,0,0,0,0,0,0,0,0,0,0,0,0,0,0,0,0,0]}

{"tickX":24}

{"coord\_y":[[9177662,9177662,9177662,8545921,8545921,8105515,8105515,7678524,7678524,7576305,7576305,7378928,7378928,7044041,7044041,7031885,7031885,6717911,6717911,6568745,6568745,6065545,6065545,5726933,5726933,5492634,5492634,5484239,5484239,5331103,5331103,5239597,5239597,5153703,5153703,5032164,5032164,4995302,4995302,4930397,4930397,4894719,4894719,4737590,4737590,4687971,4687971,4610929,4610929,4599233,4599233,4577433,4577433,4487000,4487000,4460343,4460343,4339348,4339348,4322873,4322873,4322607,4322607,4282844,4282844,4268630,4268630,4254679,4254679,4243097,4243097,4212005,4212005,4197143,4197143,4190444,4190444,4132050,4132050,4121404,4121404,4120018,4120018,4093307,4093307,4059383,4059383,4052957,4052957,4014129,4014129,3991828,3991828,3955295,3955295,3931750,3931750,3917426,3917426,3902112,3902112,3892790,3892790,3838091,3838091,3830475,3830475,3817316,3817316,3815258,3815258,3806206,3806206,3803369,3803369,3799092,3799092,3779752,3779752,3775352,3775352,3769984,3769984,3765926,3765926,3749294,3749294,3728928,3728928,3708854,3708854,3708510,3708510,3697295,3697295,3694626,3694626,3689986,3689986,3686968,3686968,3663345,3663345,3638833,3638833,3606267,3606267,3598289,3598289,3559489,3559489,3556693,3556693,3521952,3521952,3517958,3517958,3514244,3514244,3504872,3504872,3502077,3502077,3475268,3475268,3473192,3473192,3433515,3433515,3432172,3432172,3430570,3430570,3418776,3418776,3416165,3416165,3412672,3412672,3407297,3407297,3406991,3406991,3404775,3404775,3351511,3351511,3349211,3349211,3337798,3337798,3322174,3322174,3321597,3321597,3317426,3317426,3312148,3312148,3311016,3311016,3307065,3307065,3276704,3276704,3274586,3274586,3271197,3271197,3267823,3267823,3243538,3243538,3232772,3232772,3212681,3212681,3210577,3210577,3201269,3201269,3192631,3192631,3190374,3190374,3132265,3132265,3132065,3132065,3130918,3130918,3128809,3128809,3126809,3126809,3126350,3126350,3124726,3124726,3078505,3078505,3076147,3076147,3073465,3073465,3025583,3025583,3014745,3014745,3012150,3012150,3009817,3009817,3005418,3005418,3000898,3000898,2997064,2997064,2994220,2994220,2993266,2993266,2989348,2989348,2980124,2980124,2980015,2980015,2977842,2977842,2941234,2941234,2924104,2924104,2918614,2918614,2910562,2910562,2895199,2895199,2887686,2887686,2882559,2882559,2873313,2873313,2866269,2866269,2851370,2851370,2842036,2842036,2841561,2841561,2839786,2839786,2834966,2834966,2827103,2827103,2802283,2802283,2795603,2795603,2795403,2795403,2794252,2794252,2791386,2791386,2780916,2780916,2772159,2772159,2771104,2771104,2760613,2760613,2756147,2756147,2756120,2756120,2745829,2745829,2742641,2742641,2728229,2728229,2723223,2723223,2716440,2716440,2711999,2711999,2706266,2706266,2689651,2689651,2675702,2675702,2668282,2668282,2667768,2667768,2658370,2658370,2651871,2651871,2650157,2650157,2643903,2643903,2642853,2642853,2624267,2624267,2624248,2624248,2617590,2617590,2603784,2603784,2593181,2593181,2583820,2583820,2583408,2583408,2576963,2576963,2573282,2573282,2560632,2560632,2544737,2544737,2544111,2544111,2530857,2530857,2523272,2523272,2521447,2521447,2511634,2511634,2508805,2508805,2500778,2500778,2500275,2500275,2494664,2494664,2489798,2489798,2474762,2474762,2471936,2471936,2461260,2461260,2455247,2455247,2452784,2452784,2449995,2449995,2443779,2443779,2443407,2443407,2440898,2440898,2420981,2420981,2416063,2416063,2414770,2414770,2412943,2412943,2402324,2402324,2397047,2397047,2385965,2385965,2383102,2383102,2379882,2379882,2372340,2372340,2362326,2362326,2358386,2358386,2350464,2350464,2348559,2348559,2346215,2346215,2338545,2338545,2331880,2331880,2328218,2328218,2327577,2327577,2317381,2317381,2316702,2316702,2309639,2309639,2306952,2306952,2301170,2301170,2296669,2296669,2296294,2296294,2295112,2295112,2294499,2294499,2293484,2293484,2288634,2288634,2286899,2286899,2286284,2286284,2286015,2286015,2285735,2285735,2281879,2281879,2272901,2272901,2271386,2271386,2270123,2270123,2268893,2268893,2265226,2265226,2264978,2264978,2256195,2256195,2252146,2252146,2249058,2249058,2239984,2239984,2231669,2231669,2227605,2227605,2220894,2220894,2218949,2218949,2210495,2210495,2202575,2202575,2202519,2202519,2189003,2189003,2187457,2187457,2174888,2174888,2172107,2172107,2164150,2164150,2154790,2154790,2152358,2152358,2146480,2146480,2144492,2144492,2144338,2144338,2140171,2140171,2134174,2134174,2127527,2127527,2123602,2123602,2123094,2123094,2122261,2122261,2118515,2118515,2116945,2116945,2113871,2113871,2113099,2113099,2105097,2105097,2104306,2104306,2097739,2097739,2081089,2081089,2077920,2077920,2071952,2071952,2071923,2071923,2071400,2071400,2069509,2069509,2067256,2067256,2066989,2066989,2061832,2061832,2060156,2060156,2060098,2060098,2059482,2059482,2057946,2057946,2054864,2054864,2054812,2054812,2052044,2052044,2047492,2047492,2045570,2045570,2043046,2043046,2040752,2040752,2035964,2035964,2026746,2026746,2024728,2024728,2019778,2019778,2016279,2016279,2016141,2016141,2015861,2015861,2002346,2002346,2001001,2001001,1996023,1996023,1994860,1994860,1991308,1991308,1988578,1988578,1988510,1988510,1986541,1986541,1975709,1975709,1966462,1966462,1965509,1965509,1962210,1962210,1956557,1956557,1951019,1951019,1943830,1943830,1941283,1941283,1940234,1940234,1939680,1939680,1929652,1929652,1929047,1929047,1922563,1922563,1922033,1922033,1918314,1918314,1916270,1916270,1911814,1911814,1911731,1911731,1910899,1910899,1906360,1906360,1905676,1905676,1904056,1904056,1898073,1898073,1886477,1886477,1883902,1883902,1874736,1874736,1872784,1872784,1863127,1863127,1861138,1861138,1855142,1855142,1851959,1851959,1844968,1844968,1843082,1843082,1842844,1842844,1842549,1842549,1842200,1842200,1841691,1841691,1823858,1823858,1817481,1817481,1816196,1816196,1814273,1814273,1810922,1810922,1806830,1806830,1804357,1804357,1802899,1802899,1798623,1798623,1791305,1791305,1785297,1785297,1781834,1781834,1774795,1774795,1774084,1774084,1768111,1768111,1765749,1765749,1762213,1762213,1761786,1761786,1755144,1755144,1754975,1754975,1752047,1752047,1750167,1750167,1747078,1747078,1746514,1746514,1745345,1745345,1744263,1744263,1740988,1740988,1740466,1740466,1740277,1740277,1738364,1738364,1735414,1735414,1731481,1731481,1726284,1726284,1723102,1723102,1722951,1722951,1721867,1721867,1721040,1721040,1718233,1718233,1718060,1718060,1713434,1713434,1713349,1713349,1711128,1711128,1708918,1708918,1705511,1705511,1705237,1705237,1701917,1701917,1699969,1699969,1699075,1699075,1688141,1688141,1687570,1687570,1686065,1686065,1685643,1685643,1685095,1685095,1683730,1683730,1682013,1682013,1679784,1679784,1677607,1677607,1672525,1672525,1672200,1672200,1670635,1670635,1665921,1665921,1664440,1664440,1662798,1662798,1662703,1662703,1662019,1662019,1659368,1659368,1659110,1659110,1658184,1658184,1656392,1656392,1654374,1654374,1649774,1649774,1641521,1641521,1639876,1639876,1638175,1638175,1637215,1637215,1635474,1635474,1628599,1628599,1624655,1624655,1620797,1620797,1608723,1608723,1605987,1605987,1597597,1597597,1597152,1597152,1597058,1597058,1596447,1596447,1591866,1591866,1580807,1580807,1580488,1580488,1578664,1578664,1573842,1573842,1570495,1570495,1561813,1561813,1553720,1553720,1552537,1552537,1551827,1551827,1548852,1548852,1548666,1548666,1547340,1547340,1545223,1545223,1544196,1544196,1544180,1544180,1542725,1542725,1541621,1541621,1537847,1537847,1533216,1533216,1533059,1533059,1527337,1527337,1527068,1527068,1518981,1518981,1517873,1517873,1515099,1515099,1515067,1515067,1512256,1512256,1509615,1509615,1506673,1506673,1504694,1504694,1503217,1503217,1502011,1502011,1500402,1500402,1499001,1499001,1495695,1495695,1495575,1495575,1491376,1491376,1487334,1487334,1485427,1485427,1485388,1485388,1481575,1481575,1480948,1480948,1480325,1480325,1480278,1480278,1479297,1479297,1476014,1476014,1472333,1472333,1471469,1471469,1471204,1471204,1470551,1470551,1470360,1470360,1467273,1467273,1466774,1466774,1466642,1466642,1465412,1465412,1462511,1462511,1453002,1453002,1449242,1449242,1444715,1444715,1438204,1438204,1433571,1433571,1431707,1431707,1430578,1430578,1421471,1421471,1421449,1421449,1419391,1419391,1417510,1417510,1416945,1416945,1416180,1416180,1415807,1415807,1414324,1414324,1414261,1414261,1413116,1413116,1411414,1411414,1410598,1410598,1409769,1409769,1406926,1406926,1406704,1406704,1404122,1404122,1399307,1399307,1391461,1391461,1390099,1390099,1387096,1387096,1385631,1385631,1380258,1380258,1377236,1377236,1375582,1375582,1374901,1374901,1369807,1369807,1368232,1368232,1363105,1363105,1359163,1359163,1357890,1357890,1357445,1357445,1357052,1357052,1356598,1356598,1353923,1353923,1353048,1353048,1352934,1352934,1352121,1352121,1351940,1351940,1351529,1351529,1350615,1350615,1350352,1350352,1350163,1350163,1348701,1348701,1347240,1347240,1346467,1346467,1345607,1345607,1344384,1344384,1342535,1342535,1341402,1341402,1338940,1338940,1337112,1337112,1336607,1336607,1335995,1335995,1334556,1334556,1332753,1332753,1332375,1332375,1331669,1331669,1330274,1330274,1321765,1321765,1320175,1320175,1318842,1318842,1316375,1316375,1316233,1316233,1316073,1316073,1314580,1314580,1312790,1312790,1312554,1312554,1312025,1312025,1307281,1307281,1301385,1301385,1300822,1300822,1298261,1298261,1296626,1296626,1295715,1295715,1294502,1294502,1294242,1294242,1293986,1293986,1293304,1293304,1291461,1291461,1291016,1291016,1290461,1290461,1290161,1290161,1288327,1288327,1287838,1287838,1286271,1286271,1285220,1285220,1284834,1284834,1284780,1284780,1282305,1282305,1280671,1280671,1279363,1279363,1279127,1279127,1278898,1278898,1278787,1278787,1277791,1277791,1277765,1277765,1276382,1276382,1275837,1275837,1273293,1273293,1272028,1272028,1271730,1271730,1271464,1271464,1271346,1271346,1270729,1270729,1270170,1270170,1269444,1269444,1262549,1262549,1262037,1262037,1260998,1260998,1260926,1260926,1260056,1260056,1257979,1257979,1257774,1257774,1257206,1257206,1255178,1255178,1252132,1252132,1252068,1252068,1251675,1251675,1249245,1249245,1246712,1246712,1242196,1242196,1241286,1241286,1241175,1241175,1240076,1240076,1238333,1238333,1237891,1237891,1236961,1236961,1236184,1236184,1228171,1228171,1226500,1226500,1222956,1222956,1222381,1222381,1222089,1222089,1220958,1220958,1220046,1220046,1218810,1218810,1218138,1218138,1215373,1215373,1213861,1213861,1208803,1208803,1206953,1206953,1206906,1206906,1202220,1202220,1202183,1202183,1201365,1201365,1199827,1199827,1198140,1198140,1197376,1197376,1196171,1196171,1195858,1195858,1194811,1194811,1190010,1190010,1189851,1189851,1189599,1189599,1188079,1188079,1187703,1187703,1187554,1187554,1187257,1187257,1185330,1185330,1185309,1185309,1184512,1184512,1183781,1183781,1182785,1182785,1181150,1181150,1181048,1181048,1180986,1180986,1180180,1180180,1179944,1179944,1176652,1176652,1175600,1175600,1172738,1172738,1172265,1172265,1168633,1168633,1167229,1167229,1166723,1166723,1163245,1163245,1162688,1162688,1162613,1162613,1161834,1161834,1160842,1160842,1158648,1158648,1156236,1156236,1155986,1155986,1155060,1155060,1153750,1153750,1150817,1150817,1149709,1149709,1147293,1147293,1146797,1146797,1145001,1145001,1143663,1143663,1140915,1140915,1140368,1140368,1139470,1139470,1139021,1139021,1138081,1138081,1135645,1135645,1134074,1134074,1133507,1133507,1127378,1127378,1126712,1126712,1124836,1124836,1124753,1124753,1123857,1123857,1123754,1123754,1119028,1119028,1115116,1115116,1114433,1114433,1112280,1112280,1112176,1112176,1110820,1110820,1108895,1108895,1107978,1107978,1107636,1107636,1107325,1107325,1106254,1106254,1103190,1103190,1102824,1102824,1100026,1100026,1099047,1099047,1098353,1098353,1095919,1095919,1093330,1093330,1092934,1092934,1089734,1089734,1088594,1088594,1087160,1087160,1085851,1085851,1085504,1085504,1082835,1082835,1080417,1080417,1078239,1078239,1073572,1073572,1073356,1073356,1072548,1072548,1067302,1067302,1063401,1063401,1062006,1062006,1061725,1061725,1060522,1060522,1058970,1058970,1057958,1057958,1057682,1057682,1056119,1056119,1053863,1053863,1053813,1053813,1051294,1051294,1050984,1050984,1050937,1050937,1050743,1050743,1049608,1049608,1049048,1049048,1048849,1048849,1046683,1046683,1046410,1046410,1044720,1044720,1041112,1041112,1040497,1040497,1038051,1038051,1035523,1035523,1034114,1034114,1032347,1032347,1031286,1031286,1030862,1030862,1028854,1028854,1028438,1028438,1023362,1023362,1021228,1021228,1019935,1019935,1016571,1016571,1016001,1016001,1014672,1014672,1014171,1014171,1014132,1014132,1009918,1009918,1007924,1007924,1007906,1007906,1003876,1003876,1001487,1001487,999106,999106,995035,995035,994554,994554,993874,993874,993740,993740,990609,990609,990084,990084,987278,987278,985330,985330,983247,983247,981713,981713,980783,980783,980208,980208,978741,978741,978202,978202,973650,973650,972864,972864,972090,972090,969591,969591,962607,962607,961611,961611,961010,961010,960462,960462,959767,959767,959745,959745,959618,959618,958807,958807,958109,958109,953933,953933,950767,950767,949563,949563,949453,949453,945019,945019,940948,940948,940858,940858,940841,940841,939436,939436,938129,938129,938001,938001,937590,937590,936808,936808,933972,933972,933147,933147,932468,932468,932377,932377,932313,932313,932027,932027,929064,929064,927538,927538,926120,926120,923829,923829,921357,921357,921193,921193,920932,920932,918190,918190,917248,917248,916816,916816,916327,916327,914827,914827,913186,913186,912523,912523,909830,909830,908957,908957,907795,907795,906887,906887,906336,906336,904818,904818,903339,903339,902851,902851,902303,902303,902152,902152,901692,901692,901554,901554,899121,899121,897340,897340,895473,895473,894309,894309,894158,894158,889257,889257,887240,887240,887080,887080,886585,886585,885925,885925,885209,885209,883473,883473,883306,883306,882624,882624,878057,878057,877353,877353,876897,876897,873403,873403,871145,871145,869660,869660,867503,867503,866910,866910,865411,865411,864687,864687,864593,864593,864503,864503,863930,863930,861960,861960,861699,861699,860545,860545,860309,860309,858861,858861,857616,857616,855626,855626,854521,854521,853763,853763,851374,851374,850743,850743,847453,847453,847126,847126,847070,847070,845480,845480,845043,845043,843840,843840,843274,843274,841747,841747,840899,840899,840735,840735,839697,839697,839051,839051,838483,838483,837762,837762,836717,836717,836153,836153,835799,835799,831589,831589,831572,831572,826935,826935,826726,826726,825963,825963,825565,825565,824922,824922,824885,824885,823419,823419,823110,823110,822694,822694,822157,822157,819863,819863,819467,819467,816431,816431,816304,816304,814048,814048,813907,813907,813623,813623,812734,812734,812578,812578,810915,810915,810779,810779,809341,809341,808166,808166,807155,807155,805955,805955,804859,804859,803320,803320,801299,801299,799090,799090,797600,797600,794621,794621,793532,793532,792006,792006,790201,790201,790122,790122,789177,789177,788506,788506,788336,788336,787821,787821,787015,787015,786885,786885,785525,785525,784140,784140,783859,783859,779907,779907,779122,779122,778789,778789,777494,777494,774835,774835,773453,773453,772750,772750,772395,772395,771857,771857,771518,771518,771352,771352,770630,770630,769956,769956,769665,769665,769481,769481,767553,767553,762721,762721,761852,761852,760707,760707,760017,760017,758343,758343,755180,755180,755025,755025,754688,754688,752306,752306,750124,750124,748735,748735,748716,748716,748389,748389,746754,746754,744485,744485,744100,744100,736471,736471,735496,735496,733100,733100,730878,730878,726979,726979,724444,724444,723735,723735,723286,723286,720866,720866,719094,719094,716995,716995,715277,715277,709802,709802,709581,709581,709034,709034,708416,708416,707418,707418,707274,707274,705869,705869,705252,705252,703521,703521,703088,703088,701794,701794,700054,700054,698769,698769,696077,696077,693287,693287,692805,692805,692788,692788,692632,692632,691453,691453,690607,690607,688708,688708,687301,687301,685847,685847,684032,684032,683807,683807,683770,683770,682512,682512,680406,680406,679362,679362,677205,677205,675822,675822,675322,675322,672315,672315,671734,671734,671596,671596,666337,666337,666083,666083,665798,665798,665721,665721,664270,664270,663651,663651,663520,663520,663248,663248,663229,663229,661857,661857,660769,660769,659208,659208,657764,657764,656889,656889,654803,654803,654402,654402,652801,652801,652078,652078,649304,649304,647220,647220,645032,645032,644246,644246,643620,643620,642404,642404,640592,640592,640464,640464,638736,638736,635131,635131,631542,631542,630293,630293,628261,628261,628196,628196,626161,626161,625812,625812,625453,625453,625205,625205,625112,625112,624917,624917,623692,623692,623375,623375,623176,623176,622697,622697,621517,621517,619883,619883,619413,619413,618003,618003,617869,617869,615646,615646,615506,615506,611748,611748,610005,610005,608470,608470,608308,608308,608208,608208,605825,605825,605532,605532,604853,604853,603879,603879,602467,602467,601994,601994,601009,601009,596024,596024,595612,595612,595577,595577,594954,594954,594595,594595,594058,594058,593886,593886,593076,593076,590371,590371,589091,589091,588915,588915,588750,588750,588226,588226,586960,586960,586206,586206,585634,585634,585193,585193,584956,584956,583043,583043,582206,582206,580706,580706,580388,580388,579844,579844,578281,578281,577594,577594,576465,576465,575044,575044,574781,574781,574416,574416,572249,572249,571318,571318,569574,569574,569058,569058,566816,566816,566780,566780,565760,565760,562343,562343,562081,562081,561460,561460,559686,559686,559629,559629,559125,559125,558861,558861,558400,558400,557007,557007,555852,555852,555678,555678,555360,555360,555012,555012,554691,554691,554610,554610,554333,554333,553973,553973,553224,553224,552029,552029,551267,551267,549479,549479,548613,548613,547079,547079,546837,546837,546641,546641,545738,545738,544658,544658,541303,541303,540677,540677,537764,537764,536047,536047,535984,535984,533681,533681,533551,533551,533467,533467,532914,532914,532755,532755,532511,532511,531996,531996,530531,530531,528671,528671,528255,528255,528075,528075,527007,527007,526965,526965,526607,526607,525619,525619,523796,523796,522152,522152,519173,519173,518244,518244,516707,516707,515562,515562,514874,514874,514299,514299,511817,511817,510313,510313,509488,509488,508504,508504,508110,508110,507173,507173,505410,505410,505074,505074,504478,504478,504233,504233,503730,503730,503591,503591,503273,503273,502411,502411,500869,500869,500094,500094,499940,499940,499739,499739,498503,498503,497231,497231,496692,496692,495143,495143,494622,494622,493755,493755,492848,492848,491581,491581,490838,490838,490790,490790,490756,490756,488337,488337,487924,487924,487176,487176,483260,483260,482796,482796,482266,482266,482048,482048,481174,481174,479816,479816,478033,478033,477749,477749,475561,475561,474894,474894,473442,473442,472681,472681,469507,469507,468180,468180,467688,467688,464828,464828,464164,464164,464116,464116,462625,462625,462091,462091,461170,461170,460862,460862,460638,460638,459458,459458,457092,457092,456719,456719,455494,455494,455417,455417,453241,453241,452790,452790,450993,450993,448882,448882,448204,448204,446281,446281,446243,446243,445741,445741,443960,443960,443765,443765,443293,443293,443187,443187,443078,443078,442817,442817,442436,442436,441699,441699,440546,440546,439650,439650,439613,439613,436189,436189,435018,435018,431637,431637,430831,430831,430737,430737,430141,430141,426746,426746,426097,426097,425275,425275,425228,425228,425058,425058,423903,423903,423752,423752,420972,420972,420915,420915,419965,419965,418601,418601,416973,416973,416603,416603,416034,416034,415663,415663,413589,413589,413253,413253,412418,412418,411502,411502,411463,411463,411302,411302,411135,411135,411091,411091,410201,410201,409501,409501,409357,409357,408975,408975,408827,408827,404376,404376,404056,404056,402873,402873,402574,402574,401127,401127,400069,400069,398767,398767,397832,397832,395935,395935,395093,395093,394237,394237,393910,393910,393371,393371,392865,392865,392499,392499,392093,392093,390237,390237,389274,389274,388025,388025,386482,386482,383331,383331,382944,382944,382493,382493,381995,381995,381727,381727,381052,381052,379942,379942,379832,379832,378073,378073,377751,377751,376721,376721,376107,376107,375990,375990,375839,375839,374859,374859,374824,374824,374462,374462,371991,371991,371610,371610,370487,370487,370145,370145,368820,368820,367447,367447,367236,367236,366727,366727,364970,364970,364563,364563,363274,363274,361106,361106,360670,360670,359002,359002,358894,358894,358618,358618,357828,357828,356759,356759,356740,356740,356273,356273,356170,356170,355113,355113,354495,354495,354458,354458,354354,354354,353473,353473,352434,352434,352153,352153,349351,349351,349060,349060,348063,348063,347806,347806,346800,346800,345135,345135,344350,344350,344221,344221,343910,343910,343522,343522,343460,343460,343402,343402,342852,342852,341843,341843,341678,341678,341110,341110,340375,340375,340350,340350,339872,339872,338320,338320,335515,335515,334205,334205,333774,333774,333381,333381,332991,332991,332730,332730,332696,332696,332229,332229,331734,331734,331326,331326,330309,330309,329558,329558,329125,329125,328190,328190,326141,326141,326050,326050,323608,323608,323266,323266,322836,322836,322781,322781,321830,321830,320987,320987,320951,320951,320897,320897,320115,320115,319643,319643,319308,319308,318348,318348,316969,316969,316904,316904,316209,316209,316038,316038,315313,315313,313596,313596,312075,312075,311769,311769,311622,311622,311259,311259,311118,311118,310851,310851,310812,310812,310658,310658,310507,310507,309987,309987,309639,309639,309124,309124,308809,308809,308581,308581,308541,308541,308274,308274,308122,308122,307918,307918,307747,307747,307163,307163,307081,307081,306954,306954,306491,306491,306022,306022,305593,305593,304365,304365,303663,303663,303579,303579,302677,302677,301766,301766,300298,300298,299730,299730,299559,299559,299478,299478,298723,298723,298452,298452,297608,297608,297585,297585,296715,296715,296591,296591,296079,296079,295701,295701,295046,295046,294547,294547,293481,293481,292266,292266,292063,292063,291472,291472,291367,291367,291063,291063,290825,290825,290592,290592,289990,289990,289577,289577,288593,288593,287922,287922,287143,287143,286884,286884,286825,286825,286673,286673,286537,286537,285811,285811,285621,285621,285596,285596,284820,284820,284147,284147,283872,283872,283157,283157,282700,282700,282536,282536,282310,282310,281918,281918,280767,280767,280245,280245,280095,280095,279862,279862,279831,279831,278727,278727,278534,278534,278380,278380,278313,278313,278190,278190,277629,277629,277289,277289,277196,277196,276871,276871,276812,276812,276714,276714,276161,276161,275578,275578,275487,275487,275093,275093,275001,275001,274934,274934,274690,274690,274387,274387,273953,273953,273500,273500,273382,273382,273334,273334,272908,272908,272722,272722,270942,270942,270832,270832,270618,270618,270322,270322,269753,269753,269590,269590,269195,269195,269024,269024,268549,268549,268043,268043,265568,265568,265414,265414,265268,265268,265222,265222,265096,265096,264108,264108,263911,263911,263071,263071,262583,262583,262509,262509,262436,262436,262083,262083,262007,262007,261765,261765,261383,261383,261310,261310,261287,261287,261247,261247,260710,260710,260416,260416,260119,260119,259644,259644,259062,259062,258878,258878,257544,257544,257477,257477,256979,256979,256880,256880,256807,256807,256763,256763,256531,256531,256333,256333,256036,256036,256019,256019,255847,255847,255617,255617,255189,255189,255121,255121,255072,255072,254750,254750,254507,254507,254336,254336,254005,254005,253819,253819,253458,253458,252565,252565,252289,252289,251184,251184,250936,250936,250897,250897,250212,250212,249770,249770,249729,249729,249521,249521,249411,249411,249272,249272,248965,248965,248873,248873,248760,248760,248739,248739,248606,248606,247973,247973,247549,247549,247083,247083,246941,246941,246664,246664,246532,246532,246360,246360,246274,246274,246169,246169,244942,244942,244725,244725,244706,244706,244689,244689,244556,244556,243532,243532,243456,243456,243125,243125,243051,243051,242397,242397,242199,242199,242001,242001,241920,241920,241707,241707,241299,241299,241045,241045,240910,240910,240646,240646,239971,239971,239848,239848,239806,239806,238883,238883,238760,238760,238106,238106,237887,237887,237851,237851,237829,237829,237723,237723,237176,237176,236828,236828,236618,236618,236293,236293,236277,236277,235973,235973,235667,235667,235348,235348,235176,235176,235030,235030,234943,234943,234773,234773,234739,234739,233962,233962,233769,233769,233540,233540,233184,233184,232107,232107,231885,231885,231786,231786,231734,231734,231545,231545,231392,231392,231254,231254,231229,231229,231137,231137,230991,230991,230852,230852,229840,229840,229776,229776,229742,229742,229376,229376,229228,229228,228583,228583,228508,228508,228444,228444,228387,228387,227753,227753,227673,227673,227605,227605,227046,227046,226940,226940,226423,226423,226392,226392,225672,225672,225408,225408,225385,225385,225348,225348,225255,225255,225188,225188,224375,224375,224259,224259,223776,223776,223715,223715,223198,223198,222911,222911,222480,222480,222095,222095,222005,222005,221526,221526,220670,220670,220356,220356,220050,220050,219946,219946,219722,219722,219362,219362,219136,219136,218703,218703,218314,218314,218187,218187,217949,217949,217915,217915,217535,217535,217140,217140,216838,216838,216426,216426,216298,216298,216147,216147,216076,216076,215836,215836,215730,215730,215163,215163,215138,215138,214891,214891,214792,214792,214435,214435,213836,213836,213320,213320,213183,213183,212769,212769,212665,212665,212600,212600,212556,212556,211365,211365,211177,211177,210920,210920,210768,210768,210656,210656,210282,210282,209884,209884,209770,209770,209545,209545,209100,209100,208907,208907,208589,208589,208284,208284,208143,208143,207953,207953,207909,207909,207864,207864,207580,207580,207431,207431,207295,207295,206737,206737,206578,206578,206408,206408,206273,206273,206252,206252,206207,206207,206076,206076,205876,205876,205224,205224,205060,205060,204956,204956,204881,204881,204617,204617,204071,204071,203878,203878,203641,203641,203258,203258,202838,202838,202797,202797,202508,202508,202236,202236,202143,202143,201893,201893,201823,201823,201594,201594,201303,201303,200967,200967,200884,200884,200240,200240,200173,200173,199900,199900,199727,199727,199686,199686,199667,199667,199335,199335,199219,199219,199201,199201,198844,198844,198786,198786,198767,198767,198452,198452,198377,198377,198348,198348,198320,198320,197595,197595,197555,197555,197451,197451,197045,197045,196730,196730,196459,196459,196085,196085,195817,195817,195687,195687,195555,195555,195422,195422,195373,195373,195315,195315,195186,195186,195015,195015,194714,194714,194600,194600,193977,193977,193872,193872,193734,193734,193668,193668,193637,193637,193467,193467,193374,193374,193308,193308,193159,193159,193034,193034,192920,192920,192708,192708,192375,192375,192345,192345,192323,192323,192255,192255,192124,192124,192071,192071,191828,191828,191196,191196,191157,191157,191061,191061,190986,190986,190891,190891,190871,190871,190817,190817,190750,190750,190246,190246,190200,190200,190140,190140,189966,189966,189933,189933,189886,189886,189667,189667,189630,189630,189224,189224,189180,189180,188987,188987,188770,188770,188739,188739,188583,188583,187924,187924,187871,187871,187844,187844,187753,187753,187565,187565,187534,187534,187489,187489,187154,187154,187127,187127,187111,187111,187087,187087,187049,187049,186824,186824,186749,186749,186290,186290,185915,185915,185795,185795,185758,185758,185702,185702,185510,185510,185197,185197,185002,185002,184904,184904,184837,184837,184747,184747,184675,184675,184639,184639,184623,184623,184581,184581,184543,184543,184329,184329,184220,184220,184157,184157,184017,184017,183808,183808,183696,183696,183597,183597,183579,183579,183348,183348,183172,183172,183020,183020,182740,182740,182705,182705,182689,182689,182475,182475,182415,182415,182370,182370,182128,182128,182045,182045,182023,182023,181592,181592,181396,181396,181334,181334,181311,181311,181233,181233,181107,181107,181034,181034,180596,180596,180496,180496,180480,180480,180458,180458,180360,180360,180303,180303,180026,180026,179796,179796,179748,179748,179696,179696,179570,179570,179466,179466,179426,179426,179395,179395,179368,179368,179267,179267,179140,179140,179018,179018,178970,178970,178837,178837,178643,178643,178601,178601,178561,178561,178472,178472,178357,178357,178199,178199,178007,178007,177935,177935,177881,177881,177828,177828,177781,177781,177758,177758,177737,177737,177411,177411,177351,177351,177167,177167,177051,177051,176997,176997,176955,176955,176846,176846,176784,176784,176456,176456,176386,176386,176258,176258,176031,176031,175957,175957,175847,175847,175496,175496,175312,175312,175195,175195,175073,175073,174824,174824,174755,174755,174681,174681,174637,174637,174592,174592,174571,174571,174550,174550,174529,174529,174469,174469,174374,174374,174301,174301,174120,174120,173922,173922,173855,173855,173773,173773,173663,173663,173595,173595,173513,173513,173323,173323,173251,173251,173204,173204,173148,173148,173056,173056,173002,173002,172925,172925,172853,172853,172825,172825,172591,172591,172532,172532,172494,172494,172412,172412,172303,172303,172177,172177,172103,172103,172047,172047,171813,171813,171759,171759,171736,171736,171705,171705,171687,171687,171661,171661,171548,171548,171523,171523,171395,171395,171288,171288,171269,171269,171244,171244,171142,171142,171005,171005,170956,170956,170863,170863,170845,170845,170827,170827,170803,170803,170741,170741,170639,170639,170617,170617,170586,170586,170505,170505,170354,170354,170255,170255,170121,170121,170052,170052,170003,170003,169831,169831,169720,169720,169550,169550,169384,169384,169301,169301,169214,169214,169156,169156,169086,169086,169011,169011,168993,168993,168912,168912,168869,168869,168840,168840,168792,168792,168691,168691,168656,168656,168598,168598,168577,168577,168412,168412,168377,168377,168305,168305,168204,168204,168060,168060,167924,167924,167705,167705,167676,167676,167648,167648,167542,167542,167495,167495,167456,167456,167321,167321,167300,167300,167240,167240,167094,167094,167047,167047,166882,166882,166843,166843,166688,166688,166658,166658,166614,166614,166538,166538,166472,166472,166343,166343,166198,166198,166179,166179,166145,166145,166063,166063,165989,165989,165955,165955,165837,165837,165815,165815,165603,165603,165574,165574,165527,165527,165382,165382,165347,165347,165329,165329,165295,165295,165259,165259,165136,165136,165098,165098,164965,164965,164926,164926,164838,164838,164812,164812,164778,164778,164690,164690,164648,164648,164580,164580,164525,164525,164428,164428,164383,164383,164287,164287,164250,164250,164226,164226,164159,164159,164120,164120,163987,163987,163965,163965,163922,163922,163889,163889,163867,163867,163752,163752,163660,163660,163638,163638,163616,163616,163589,163589,163496,163496,163336,163336,163314,163314,163053,163053,163005,163005,162859,162859,162837,162837,162790,162790,162727,162727,162693,162693,162649,162649,162568,162568,162532,162532,162477,162477,162411,162411,162308,162308,162288,162288,162253,162253,162227,162227,162195,162195,162125,162125,161995,161995,161974,161974,161909,161909,161886,161886,161819,161819,161655,161655,161566,161566,161522,161522,161467,161467,161446,161446,161351,161351,161327,161327,161275,161275,161197,161197,161166,161166,161130,161130,161018,161018,160999,160999,160912,160912,160805,160805,160777,160777,160617,160617,160536,160536,160500,160500,160436,160436,160415,160415,160355,160355,160301,160301,160256,160256,160229,160229,160207,160207,160187,160187,160163,160163,160089,160089,160031,160031,159893,159893,159844,159844,159766,159766,159728,159728,159703,159703,159674,159674,159624,159624,159567,159567,159406,159406,159380,159380,159355,159355,159262,159262,159206,159206,159131,159131,159035,159035,158814,158814,158798,158798,158762,158762,158745,158745,158681,158681,158622,158622,158482,158482,158463,158463,158418,158418,158387,158387,158371,158371,158267,158267,158235,158235,158211,158211,158129,158129,158084,158084,157979,157979,157927,157927,157873,157873,157848,157848,157814,157814,157775,157775,157615,157615,157517,157517,157482,157482,157333,157333,157265,157265,157237,157237,157117,157117,157078,157078,157012,157012,156949,156949,156931,156931,156854,156854,156773,156773,156746,156746,156558,156558,156538,156538,156512,156512,156433,156433,156392,156392,156371,156371,156320,156320,156279,156279,156236,156236,156134,156134,156038,156038,156017,156017,155885,155885,155867,155867,155770,155770,155692,155692,155585,155585,155551,155551,155486,155486,155435,155435,155361,155361,155336,155336,155288,155288,155188,155188,155165,155165,155069,155069,154989,154989,154969,154969,154899,154899,154865,154865,154810,154810,154762,154762,154698,154698,154665,154665,154638,154638,154609,154609,154556,154556,154534,154534,154507,154507,154460,154460,154422,154422,154382,154382,154366,154366,154302,154302,154234,154234,154202,154202,154170,154170,154136,154136,154107,154107,154065,154065,153932,153932,153815,153815,153740,153740,153708,153708,153687,153687,153617,153617,153573,153573,153466,153466,153400,153400,153325,153325,153272,153272,153228,153228,153191,153191,153169,153169,153097,153097,152944,152944,152926,152926,152904,152904,152748,152748,152708,152708,152653,152653,152610,152610,152579,152579,152537,152537,152510,152510,152453,152453,152370,152370,152341,152341,152323,152323,152258,152258,152216,152216,152194,152194,152061,152061,152002,152002,151953,151953,151870,151870,151834,151834,151754,151754,151735,151735,151660,151660,151632,151632,151476,151476,151438,151438,151395,151395,151368,151368,151347,151347,151296,151296,151275,151275,151254,151254,151129,151129,151091,151091,151056,151056,151021,151021,150979,150979,150893,150893,150838,150838,150808,150808,150751,150751,150687,150687,150568,150568,150490,150490,150446,150446,150396,150396,150356,150356,150326,150326,150303,150303,150274,150274,150253,150253,150209,150209,150151,150151,149990,149990,149853,149853,149823,149823,149803,149803,149782,149782,149738,149738,149691,149691,149665,149665,149572,149572,149524,149524,149429,149429,149344,149344,149291,149291,149170,149170,149153,149153,149084,149084,149035,149035,148942,148942,148868,148868,148730,148730,148708,148708,148658,148658,148592,148592,148498,148498,148482,148482,148457,148457,148421,148421,148382,148382,148348,148348,148313,148313,148293,148293,148276,148276,148136,148136,148003,148003,147924,147924,147906,147906,147833,147833,147707,147707,147645,147645,147596,147596,147576,147576,147486,147486,147439,147439,147361,147361,147325,147325,147292,147292,147270,147270,147254,147254,147237,147237,147122,147122,147084,147084,147027,147027,146990,146990,146941,146941,146811,146811,146771,146771,146718,146718,146700,146700,146658,146658,146557,146557,146540,146540,146449,146449,146422,146422,146386,146386,146364,146364,146345,146345,146300,146300,146280,146280,146169,146169,146153,146153,146122,146122,146104,146104,145983,145983,145893,145893,145870,145870,145792,145792,145759,145759,145705,145705,145685,145685,145660,145660,145622,145622,145493,145493,145365,145365,145266,145266,145249,145249,145198,145198,145139,145139,145076,145076,145052,145052,145016,145016,144997,144997,144863,144863,144764,144764,144722,144722,144688,144688,144619,144619,144529,144529,144506,144506,144479,144479,144400,144400,144373,144373,144312,144312,144261,144261,144178,144178,144142,144142,144101,144101,144069,144069,143881,143881,143797,143797,143705,143705,143680,143680,143617,143617,143583,143583,143513,143513,143475,143475,143317,143317,143282,143282,143253,143253,143226,143226,143210,143210,143142,143142,143091,143091,143063,143063,142921,142921,142840,142840,142781,142781,142761,142761,142738,142738,142685,142685,142669,142669,142605,142605,142482,142482,142458,142458,142408,142408,142365,142365,142334,142334,142313,142313,142189,142189,142152,142152,142100,142100,142030,142030,141960,141960,141917,141917,141868,141868,141811,141811,141744,141744,141596,141596,141537,141537,141456,141456,141412,141412,141348,141348,141301,141301,141270,141270,141243,141243,141184,141184,141141,141141,141116,141116,141100,141100,141073,141073,141015,141015,140973,140973,140848,140848,140807,140807,140748,140748,140720,140720,140692,140692,140640,140640,140614,140614,140583,140583,140565,140565,140481,140481,140458,140458,140401,140401,140355,140355,140324,140324,140302,140302,140245,140245,140166,140166,140145,140145,140128,140128,140096,140096,140023,140023,139889,139889,139855,139855,139781,139781,139751,139751,139720,139720,139669,139669,139638,139638,139608,139608,139555,139555,139493,139493,139431,139431,139363,139363,139347,139347,139303,139303,139280,139280,139258,139258,139223,139223,139125,139125,139073,139073,139045,139045,139016,139016,138977,138977,138782,138782,138718,138718,138655,138655,138616,138616,138516,138516,138396,138396,138371,138371,138322,138322,138281,138281,138258,138258,138220,138220,138159,138159,138120,138120,138061,138061,138022,138022,138005,138005,137981,137981,137949,137949,137930,137930,137874,137874,137853,137853,137758,137758,137709,137709,137675,137675,137643,137643,137601,137601,137580,137580,137542,137542,137514,137514,137494,137494,137458,137458,137426,137426,137409,137409,137345,137345,137323,137323,137296,137296,137220,137220,137191,137191,137158,137158,137137,137137,137025,137025,137006,137006,136969,136969,136909,136909,136854,136854,136785,136785,136744,136744,136712,136712,136694,136694,136644,136644,136392,136392,136331,136331,136262,136262,136183,136183,136114,136114,136043,136043,136017,136017,135916,135916,135899,135899,135880,135880,135830,135830,135661,135661,135593,135593,135532,135532,135497,135497,135457,135457,135424,135424,135329,135329,135308,135308,135269,135269,135218,135218,135170,135170,135038,135038,135022,135022,134916,134916,134884,134884,134861,134861,134785,134785,134716,134716,134620,134620,134583,134583,134547,134547,134491,134491,134461,134461,134432,134432,134370,134370,134315,134315,134273,134273,134165,134165,134102,134102,134079,134079,134047,134047,134031,134031,133946,133946,133827,133827,133805,133805,133768,133768,133689,133689,133667,133667,133651,133651,133606,133606,133577,133577,133534,133534,133507,133507,133436,133436,133402,133402,133359,133359,133343,133343,133298,133298,133259,133259,133142,133142,133118,133118,133060,133060,133032,133032,133015,133015,132991,132991,132962,132962,132935,132935,132909,132909,132859,132859,132839,132839,132785,132785,132764,132764,132742,132742,132706,132706,132660,132660,132535,132535,132471,132471,132397,132397,132372,132372,132337,132337,132306,132306,132264,132264,132226,132226,132129,132129,132055,132055,132029,132029,131937,131937,131912,131912,131850,131850,131811,131811,131785,131785,131701,131701,131655,131655,131607,131607,131544,131544,131443,131443,131376,131376,131333,131333,131310,131310,131281,131281,131246,131246,131218,131218,131201,131201,131183,131183,130997,130997,130935,130935,130916,130916,130880,130880,130849,130849,130789,130789,130773,130773,130750,130750,130688,130688,130598,130598,130545,130545,130482,130482,130429,130429,130321,130321,130276,130276,130215,130215,130172,130172,130153,130153,130132,130132,130112,130112,130095,130095,130071,130071,130047,130047,129992,129992,129942,129942,129891,129891,129816,129816,129795,129795,129728,129728,129699,129699,129683,129683,129653,129653,129604,129604,129532,129532,129444,129444,129333,129333,129239,129239,129201,129201,129174,129174,129119,129119,128964,128964,128925,128925,128899,128899,128764,128764,128687,128687,128666,128666,128586,128586,128511,128511,128487,128487,128459,128459,128442,128442,128402,128402,128308,128308,128224,128224,128159,128159,128101,128101,128077,128077,128025,128025,127998,127998,127960,127960,127906,127906,127886,127886,127816,127816,127794,127794,127712,127712,127683,127683,127631,127631,127598,127598,127581,127581,127494,127494,127453,127453,127408,127408,127375,127375,127314,127314,127282,127282,127246,127246,127225,127225,127177,127177,127158,127158,127121,127121,127017,127017,126933,126933,126910,126910,126848,126848,126817,126817,126753,126753,126673,126673,126648,126648,126610,126610,126575,126575,126552,126552,126500,126500,126444,126444,126421,126421,126329,126329,126308,126308,126291,126291,126229,126229,126211,126211,126047,126047,126014,126014,125987,125987,125924,125924,125887,125887,125848,125848,125819,125819,125774,125774,125682,125682,125650,125650,125619,125619,125515,125515,125497,125497,125384,125384,125352,125352,125249,125249,125231,125231,125180,125180,125154,125154,125104,125104,125001,125001,124950,124950,124922,124922,124904,124904,124875,124875,124843,124843,124815,124815,124746,124746,124709,124709,124638,124638,124611,124611,124566,124566,124495,124495,124462,124462,124420,124420,124395,124395,124347,124347,124293,124293,124277,124277,124061,124061,124016,124016,123991,123991,123962,123962,123906,123906,123862,123862,123822,123822,123788,123788,123720,123720,123643,123643,123622,123622,123487,123487,123449,123449,123433,123433,123398,123398,123347,123347,123312,123312,123267,123267,123189,123189,123172,123172,123109,123109,123074,123074,123049,123049,123014,123014,122973,122973,122908,122908,122833,122833,122815,122815,122764,122764,122638,122638,122602,122602,122546,122546,122523,122523,122477,122477,122456,122456,122437,122437,122375,122375,122347,122347,122298,122298,122281,122281,122253,122253,122208,122208,122173,122173,122110,122110,122067,122067,121988,121988,121967,121967,121925,121925,121881,121881,121847,121847,121825,121825,121809,121809,121696,121696,121628,121628,121611,121611,121510,121510,121492,121492,121452,121452,121426,121426,121398,121398,121330,121330,121303,121303,121262,121262,121209,121209,121144,121144,121095,121095,121072,121072,121049,121049,120949,120949,120835,120835,120809,120809,120770,120770,120734,120734,120683,120683,120651,120651,120615,120615,120574,120574,120556,120556,120506,120506,120378,120378,120237,120237,120191,120191,120171,120171,120152,120152,120124,120124,120033,120033,120004,120004,119986,119986,119924,119924,119908,119908,119869,119869,119852,119852,119782,119782,119757,119757,119714,119714,119690,119690,119629,119629,119592,119592,119560,119560,119544,119544,119510,119510,119435,119435,119402,119402,119382,119382,119359,119359,119317,119317,119283,119283,119266,119266,119235,119235,119211,119211,119184,119184,119147,119147,119115,119115,119099,119099,119061,119061,119014,119014,118993,118993,118953,118953,118927,118927,118891,118891,118834,118834,118785,118785,118718,118718,118662,118662,118642,118642,118621,118621,118579,118579,118520,118520,118495,118495,118471,118471,118453,118453,118363,118363,118313,118313,118285,118285,118246,118246,118196,118196,118147,118147,118117,118117,118091,118091,118072,118072,117972,117972,117955,117955,117909,117909,117834,117834,117808,117808,117790,117790,117754,117754,117656,117656,117623,117623,117607,117607,117555,117555,117507,117507,117484,117484,117450,117450,117432,117432,117414,117414,117395,117395,117301,117301,117252,117252,117227,117227,117176,117176,117126,117126,117105,117105,117085,117085,117060,117060,117040,117040,117016,117016,116959,116959,116886,116886,116807,116807,116753,116753,116726,116726,116678,116678,116546,116546,116515,116515,116479,116479,116461,116461,116417,116417,116336,116336,116268,116268,116237,116237,116215,116215,116153,116153,116119,116119,116067,116067,116050,116050,116030,116030,115970,115970,115925,115925,115864,115864,115832,115832,115780,115780,115757,115757,115732,115732,115691,115691,115533,115533,115454,115454,115435,115435,115413,115413,115394,115394,115350,115350,115332,115332,115301,115301,115277,115277,115248,115248,115222,115222,115198,115198,115180,115180,115146,115146,115087,115087,115023,115023,115005,115005,114985,114985,114961,114961,114937,114937,114880,114880,114863,114863,114822,114822,114803,114803,114774,114774,114718,114718,114683,114683,114648,114648,114623,114623,114551,114551,114357,114357,114339,114339,114303,114303,114278,114278,114231,114231,114207,114207,114156,114156,114123,114123,114087,114087,114065,114065,114039,114039,114005,114005,113952,113952,113931,113931,113897,113897,113880,113880,113844,113844,113816,113816,113788,113788,113765,113765,113734,113734,113700,113700,113651,113651,113607,113607,113569,113569,113493,113493,113465,113465,113414,113414,113345,113345,113324,113324,113241,113241,113225,113225,113125,113125,113108,113108,113092,113092,113070,113070,113053,113053,113025,113025,112985,112985,112938,112938,112921,112921,112877,112877,112852,112852,112806,112806,112774,112774,112746,112746,112719,112719,112701,112701,112683,112683,112642,112642,112565,112565,112510,112510,112475,112475,112402,112402,112384,112384,112358,112358,112305,112305,112288,112288,112254,112254,112224,112224,112180,112180,112124,112124,112107,112107,112089,112089,112000,112000,111984,111984,111926,111926,111898,111898,111842,111842,111819,111819,111782,111782,111764,111764,111724,111724,111707,111707,111613,111613,111563,111563,111544,111544,111424,111424,111356,111356,111339,111339,111322,111322,111284,111284,111260,111260,111218,111218,111158,111158,111127,111127,111105,111105,111043,111043,111017,111017,110996,110996,110943,110943,110879,110879,110854,110854,110783,110783,110749,110749,110683,110683,110648,110648,110621,110621,110494,110494,110475,110475,110447,110447,110328,110328,110299,110299,110275,110275,110235,110235,110200,110200,110104,110104,110082,110082,110025,110025,109995,109995,109972,109972,109954,109954,109908,109908,109880,109880,109861,109861,109811,109811,109741,109741,109702,109702,109664,109664,109639,109639,109622,109622,109579,109579,109498,109498,109423,109423,109407,109407,109378,109378,109344,109344,109304,109304,109238,109238,109222,109222,109194,109194,109175,109175,109124,109124,109090,109090,109045,109045,109028,109028,109000,109000,108975,108975,108954,108954,108931,108931,108882,108882,108790,108790,108753,108753,108717,108717,108701,108701,108596,108596,108555,108555,108425,108425,108396,108396,108364,108364,108345,108345,108324,108324,108282,108282,108252,108252,108189,108189,108172,108172,108153,108153,108073,108073,108053,108053,108037,108037,108017,108017,107999,107999,107965,107965,107943,107943,107927,107927,107899,107899,107858,107858,107825,107825,107734,107734,107715,107715,107678,107678,107615,107615,107584,107584,107530,107530,107497,107497,107471,107471,107423,107423,107401,107401,107380,107380,107348,107348,107295,107295,107233,107233,107161,107161,107140,107140,107115,107115,107086,107086,107058,107058,107037,107037,107016,107016,106999,106999,106934,106934,106760,106760,106697,106697,106674,106674,106614,106614,106589,106589,106494,106494,106454,106454,106409,106409,106340,106340,106267,106267,106225,106225,106124,106124,105997,105997,105955,105955,105925,105925,105859,105859,105816,105816,105785,105785,105740,105740,105686,105686,105655,105655,105616,105616,105593,105593,105566,105566,105538,105538,105438,105438,105408,105408,105382,105382,105300,105300,105258,105258,105174,105174,105147,105147,105075,105075,105030,105030,105009,105009,104972,104972,104934,104934,104896,104896,104877,104877,104859,104859,104830,104830,104806,104806,104699,104699,104667,104667,104587,104587,104560,104560,104514,104514,104487,104487,104431,104431,104408,104408,104377,104377,104351,104351,104319,104319,104256,104256,104197,104197,104146,104146,104130,104130,104097,104097,104035,104035,104010,104010,103991,103991,103971,103971,103948,103948,103900,103900,103868,103868,103829,103829,103794,103794,103764,103764,103735,103735,103718,103718,103700,103700,103681,103681,103628,103628,103606,103606,103575,103575,103553,103553,103521,103521,103489,103489,103467,103467,103433,103433,103409,103409,103376,103376,103299,103299,103257,103257,103218,103218,103193,103193,103166,103166,103143,103143,103114,103114,103090,103090,103034,103034,103012,103012,102953,102953,102925,102925,102863,102863,102834,102834,102812,102812,102794,102794,102767,102767,102718,102718,102636,102636,102619,102619,102573,102573,102547,102547,102529,102529,102511,102511,102475,102475,102448,102448,102424,102424,102388,102388,102323,102323,102169,102169,102148,102148,102131,102131,102099,102099,102082,102082,102065,102065,102048,102048,101931,101931,101885,101885,101852,101852,101826,101826,101800,101800,101693,101693,101667,101667,101648,101648,101593,101593,101556,101556,101523,101523,101501,101501,101474,101474,101438,101438,101413,101413,101374,101374,101337,101337,101271,101271,101244,101244,101209,101209,101189,101189,101164,101164,101132,101132,101030,101030,100982,100982,100955,100955,100928,100928,100893,100893,100864,100864,100839,100839,100815,100815,100778,100778,100726,100726,100686,100686,100644,100644,100606,100606,100562,100562,100450,100450,100426,100426,100302,100302,100255,100255,100236,100236,100158,100158,100122,100122,100050,100050,100016,100016,100000,100000,99963,99963,99923,99923,99876,99876,99843,99843,99781,99781,99761,99761,99745,99745,99726,99726,99702,99702,99660,99660,99630,99630,99587,99587,99521,99521,99457,99457,99438,99438,99397,99397,99293,99293,99271,99271,99198,99198,99180,99180,99052,99052,99036,99036,99010,99010,98976,98976,98844,98844,98793,98793,98769,98769,98716,98716,98655,98655,98615,98615,98565,98565,98545,98545,98511,98511,98490,98490,98446,98446,98402,98402,98378,98378,98356,98356,98325,98325,98285,98285,98255,98255,98229,98229,98185,98185,98147,98147,98045,98045,98000,98000,97941,97941,97828,97828,97805,97805,97776,97776,97744,97744,97696,97696,97679,97679,97617,97617,97598,97598,97551,97551,97514,97514,97489,97489,97472,97472,97456,97456,97381,97381,97354,97354,97301,97301,97272,97272,97210,97210,97183,97183,97149,97149,97126,97126,97103,97103,97077,97077,97059,97059,97013,97013,96955,96955,96917,96917,96849,96849,96744,96744,96725,96725,96688,96688,96621,96621,96569,96569,96506,96506,96478,96478,96451,96451,96401,96401,96336,96336,96315,96315,96285,96285,96252,96252,96218,96218,96160,96160,96110,96110,96042,96042,95989,95989,95973,95973,95942,95942,95849,95849,95829,95829,95750,95750,95728,95728,95707,95707,95689,95689,95576,95576,95478,95478,95432,95432,95404,95404,95382,95382,95363,95363,95294,95294,95278,95278,95240,95240,95196,95196,95170,95170,95106,95106,95078,95078,94987,94987,94944,94944,94894,94894,94862,94862,94843,94843,94824,94824,94808,94808,94761,94761,94730,94730,94706,94706,94678,94678,94618,94618,94599,94599,94573,94573,94519,94519,94485,94485,94450,94450,94362,94362,94325,94325,94306,94306,94288,94288,94267,94267,94241,94241,94218,94218,94138,94138,94090,94090,93954,93954,93936,93936,93898,93898,93861,93861,93831,93831,93717,93717,93694,93694,93664,93664,93625,93625,93580,93580,93518,93518,93460,93460,93444,93444,93424,93424,93396,93396,93380,93380,93361,93361,93234,93234,93191,93191,93148,93148,93048,93048,92992,92992,92971,92971,92954,92954,92866,92866,92838,92838,92776,92776,92739,92739,92712,92712,92695,92695,92674,92674,92635,92635,92613,92613,92588,92588,92529,92529,92502,92502,92421,92421,92386,92386,92323,92323,92296,92296,92276,92276,92220,92220,92182,92182,92096,92096,92080,92080,92030,92030,91985,91985,91954,91954,91934,91934,91917,91917,91879,91879,91839,91839,91799,91799,91783,91783,91766,91766,91743,91743,91713,91713,91672,91672,91636,91636,91620,91620,91578,91578,91560,91560,91521,91521,91499,91499,91478,91478,91442,91442,91424,91424,91397,91397,91366,91366,91348,91348,91330,91330,91309,91309,91279,91279,91222,91222,91136,91136,91052,91052,91025,91025,90999,90999,90957,90957,90933,90933,90910,90910,90862,90862,90831,90831,90814,90814,90774,90774,90731,90731,90663,90663,90610,90610,90557,90557,90521,90521,90502,90502,90412,90412,90376,90376,90333,90333,90311,90311,90246,90246,90218,90218,90180,90180,90142,90142,90109,90109,90089,90089,90061,90061,90036,90036,90004,90004,89980,89980,89913,89913,89851,89851,89832,89832,89792,89792,89762,89762,89725,89725,89703,89703,89685,89685,89654,89654,89626,89626,89543,89543,89515,89515,89464,89464,89429,89429,89348,89348,89331,89331,89295,89295,89243,89243,89162,89162,89133,89133,89105,89105,89079,89079,89059,89059,89036,89036,88995,88995,88929,88929,88909,88909,88893,88893,88856,88856,88810,88810,88785,88785,88714,88714,88688,88688,88661,88661,88642,88642,88610,88610,88584,88584,88549,88549,88514,88514,88476,88476,88460,88460,88425,88425,88403,88403,88387,88387,88360,88360,88320,88320,88299,88299,88280,88280,88235,88235,88184,88184,88149,88149,88129,88129,88087,88087,88035,88035,88014,88014,87969,87969,87929,87929,87910,87910,87893,87893,87877,87877,87792,87792,87752,87752,87730,87730,87713,87713,87681,87681,87635,87635,87618,87618,87594,87594,87561,87561,87545,87545,87519,87519,87480,87480,87452,87452,87431,87431,87415,87415,87341,87341,87307,87307,87288,87288,87265,87265,87218,87218,87202,87202,87140,87140,87109,87109,87056,87056,87023,87023,87004,87004,86983,86983,86938,86938,86922,86922,86903,86903,86884,86884,86831,86831,86806,86806,86750,86750,86696,86696,86665,86665,86581,86581,86564,86564,86529,86529,86472,86472,86439,86439,86415,86415,86381,86381,86308,86308,86289,86289,86222,86222,86200,86200,86147,86147,86077,86077,86022,86022,85980,85980,85952,85952,85882,85882,85832,85832,85778,85778,85745,85745,85715,85715,85694,85694,85675,85675,85655,85655,85629,85629,85555,85555,85512,85512,85494,85494,85461,85461,85440,85440,85418,85418,85384,85384,85275,85275,85258,85258,85219,85219,85203,85203,85156,85156,85121,85121,85102,85102,85057,85057,85018,85018,84927,84927,84875,84875,84840,84840,84743,84743,84701,84701,84678,84678,84656,84656,84590,84590,84568,84568,84515,84515,84465,84465,84384,84384,84322,84322,84288,84288,84192,84192,84148,84148,84129,84129,84099,84099,84079,84079,84050,84050,83984,83984,83928,83928,83910,83910,83856,83856,83815,83815,83795,83795,83775,83775,83758,83758,83679,83679,83659,83659,83620,83620,83590,83590,83545,83545,83512,83512,83479,83479,83448,83448,83360,83360,83333,83333,83295,83295,83268,83268,83153,83153,83120,83120,83081,83081,83063,83063,83029,83029,83008,83008,82975,82975,82959,82959,82922,82922,82891,82891,82869,82869,82799,82799,82771,82771,82742,82742,82679,82679,82615,82615,82594,82594,82576,82576,82528,82528,82486,82486,82455,82455,82435,82435,82413,82413,82385,82385,82326,82326,82287,82287,82261,82261,82238,82238,82164,82164,82148,82148,82124,82124,82084,82084,82068,82068,82039,82039,82015,82015,81978,81978,81932,81932,81907,81907,81848,81848,81830,81830,81788,81788,81752,81752,81736,81736,81714,81714,81695,81695,81673,81673,81616,81616,81600,81600,81547,81547,81501,81501,81478,81478,81457,81457,81404,81404,81381,81381,81343,81343,81321,81321,81291,81291,81266,81266,81246,81246,81228,81228,81208,81208,81139,81139,81121,81121,81100,81100,81073,81073,81043,81043,81002,81002,80985,80985,80952,80952,80917,80917,80881,80881,80853,80853,80784,80784,80768,80768,80736,80736,80709,80709,80685,80685,80667,80667,80572,80572,80545,80545,80471,80471,80454,80454,80403,80403,80385,80385,80338,80338,80282,80282,80228,80228,80198,80198,80173,80173,80139,80139,80106,80106,80029,80029,79977,79977,79950,79950,79805,79805,79674,79674,79657,79657,79570,79570,79538,79538,79509,79509,79474,79474,79438,79438,79401,79401,79381,79381,79356,79356,79321,79321,79269,79269,79253,79253,79193,79193,79176,79176,79154,79154,79133,79133,79110,79110,79053,79053,79027,79027,78912,78912,78887,78887,78847,78847,78825,78825,78804,78804,78782,78782,78744,78744,78703,78703,78659,78659,78556,78556,78531,78531,78505,78505,78486,78486,78470,78470,78431,78431,78329,78329,78307,78307,78284,78284,78253,78253,78230,78230,78214,78214,78195,78195,78096,78096,78044,78044,78012,78012,77953,77953,77920,77920,77898,77898,77878,77878,77845,77845,77735,77735,77710,77710,77691,77691,77636,77636,77578,77578,77531,77531,77490,77490,77460,77460,77384,77384,77338,77338,77285,77285,77267,77267,77239,77239,77172,77172,77127,77127,77107,77107,77083,77083,77065,77065,77038,77038,77013,77013,76950,76950,76891,76891,76867,76867,76824,76824,76776,76776,76712,76712,76648,76648,76607,76607,76583,76583,76548,76548,76531,76531,76505,76505,76437,76437,76398,76398,76379,76379,76347,76347,76310,76310,76282,76282,76242,76242,76193,76193,76165,76165,76138,76138,76121,76121,76073,76073,76047,76047,76004,76004,75981,75981,75935,75935,75915,75915,75895,75895,75867,75867,75846,75846,75830,75830,75814,75814,75782,75782,75761,75761,75701,75701,75637,75637,75619,75619,75552,75552,75514,75514,75493,75493,75475,75475,75449,75449,75390,75390,75374,75374,75294,75294,75262,75262,75191,75191,75150,75150,75113,75113,75079,75079,75035,75035,74994,74994,74975,74975,74940,74940,74908,74908,74853,74853,74816,74816,74793,74793,74769,74769,74668,74668,74634,74634,74591,74591,74521,74521,74491,74491,74469,74469,74449,74449,74412,74412,74388,74388,74361,74361,74338,74338,74289,74289,74264,74264,74184,74184,74144,74144,74121,74121,74097,74097,74079,74079,74039,74039,74021,74021,73997,73997,73977,73977,73916,73916,73889,73889,73831,73831,73802,73802,73760,73760,73676,73676,73658,73658,73622,73622,73606,73606,73582,73582,73514,73514,73489,73489,73466,73466,73419,73419,73403,73403,73381,73381,73271,73271,73161,73161,73091,73091,73049,73049,73011,73011,72973,72973,72845,72845,72807,72807,72778,72778,72681,72681,72649,72649,72631,72631,72609,72609,72580,72580,72547,72547,72522,72522,72483,72483,72457,72457,72418,72418,72386,72386,72322,72322,72281,72281,72251,72251,72223,72223,72177,72177,72135,72135,72101,72101,72076,72076,72037,72037,72017,72017,71970,71970,71926,71926,71910,71910,71877,71877,71843,71843,71811,71811,71782,71782,71709,71709,71662,71662,71630,71630,71604,71604,71559,71559,71542,71542,71496,71496,71471,71471,71453,71453,71415,71415,71370,71370,71354,71354,71299,71299,71283,71283,71245,71245,71226,71226,71197,71197,71178,71178,71128,71128,71112,71112,71092,71092,71076,71076,71059,71059,71042,71042,71025,71025,70963,70963,70938,70938,70881,70881,70841,70841,70804,70804,70743,70743,70674,70674,70649,70649,70572,70572,70548,70548,70523,70523,70507,70507,70482,70482,70444,70444,70414,70414,70365,70365,70347,70347,70294,70294,70272,70272,70213,70213,70181,70181,70129,70129,70073,70073,70051,70051,70034,70034,69993,69993,69943,69943,69913,69913,69897,69897,69877,69877,69849,69849,69828,69828,69781,69781,69763,69763,69746,69746,69695,69695,69653,69653,69612,69612,69583,69583,69563,69563,69542,69542,69526,69526,69503,69503,69457,69457,69427,69427,69398,69398,69366,69366,69350,69350,69334,69334,69306,69306,69289,69289,69271,69271,69197,69197,69151,69151,69121,69121,69098,69098,69074,69074,69032,69032,68971,68971,68952,68952,68923,68923,68858,68858,68827,68827,68801,68801,68778,68778,68745,68745,68729,68729,68690,68690,68656,68656,68602,68602,68579,68579,68562,68562,68514,68514,68497,68497,68465,68465,68449,68449,68427,68427,68375,68375,68335,68335,68316,68316,68287,68287,68244,68244,68205,68205,68188,68188,68155,68155,68105,68105,68085,68085,68026,68026,68001,68001,67934,67934,67890,67890,67867,67867,67820,67820,67773,67773,67711,67711,67648,67648,67632,67632,67605,67605,67587,67587,67558,67558,67518,67518,67481,67481,67459,67459,67412,67412,67381,67381,67349,67349,67332,67332,67315,67315,67295,67295,67275,67275,67238,67238,67180,67180,67149,67149,67123,67123,67102,67102,67075,67075,67057,67057,67039,67039,67022,67022,67005,67005,66988,66988,66948,66948,66903,66903,66879,66879,66829,66829,66744,66744,66706,66706,66688,66688,66660,66660,66605,66605,66585,66585,66550,66550,66524,66524,66488,66488,66459,66459,66440,66440,66413,66413,66394,66394,66377,66377,66350,66350,66331,66331,66313,66313,66281,66281,66252,66252,66211,66211,66156,66156,66129,66129,66107,66107,66055,66055,66023,66023,66005,66005,65956,65956,65928,65928,65871,65871,65855,65855,65823,65823,65807,65807,65776,65776,65744,65744,65724,65724,65706,65706,65685,65685,65664,65664,65646,65646,65622,65622,65603,65603,65586,65586,65569,65569,65542,65542,65513,65513,65483,65483,65445,65445,65427,65427,65396,65396,65369,65369,65352,65352,65319,65319,65295,65295,65241,65241,65223,65223,65204,65204,65142,65142,65107,65107,65082,65082,65065,65065,65046,65046,65027,65027,64975,64975,64937,64937,64909,64909,64880,64880,64863,64863,64832,64832,64814,64814,64779,64779,64729,64729,64669,64669,64629,64629,64613,64613,64597,64597,64561,64561,64526,64526,64485,64485,64466,64466,64442,64442,64422,64422,64396,64396,64363,64363,64327,64327,64305,64305,64277,64277,64259,64259,64242,64242,64224,64224,64185,64185,64156,64156,64120,64120,64029,64029,63991,63991,63972,63972,63922,63922,63906,63906,63859,63859,63832,63832,63814,63814,63784,63784,63756,63756,63740,63740,63703,63703,63652,63652,63611,63611,63583,63583,63560,63560,63523,63523,63505,63505,63485,63485,63444,63444,63375,63375,63325,63325,63283,63283,63266,63266,63231,63231,63211,63211,63186,63186,63161,63161,63140,63140,63114,63114,63060,63060,63037,63037,62998,62998,62941,62941,62925,62925,62898,62898,62875,62875,62772,62772,62755,62755,62717,62717,62677,62677,62643,62643,62612,62612,62581,62581,62557,62557,62532,62532,62513,62513,62495,62495,62472,62472,62424,62424,62352,62352,62311,62311,62256,62256,62217,62217,62179,62179,62146,62146,62119,62119,62076,62076,62027,62027,62010,62010,61956,61956,61922,61922,61905,61905,61887,61887,61834,61834,61798,61798,61762,61762,61734,61734,61713,61713,61688,61688,61620,61620,61600,61600,61559,61559,61543,61543,61512,61512,61489,61489,61395,61395,61317,61317,61289,61289,61273,61273,61249,61249,61217,61217,61180,61180,61151,61151,61134,61134,61058,61058,60987,60987,60968,60968,60941,60941,60922,60922,60890,60890,60847,60847,60819,60819,60802,60802,60780,60780,60754,60754,60730,60730,60707,60707,60670,60670,60651,60651,60611,60611,60584,60584,60562,60562,60545,60545,60526,60526,60508,60508,60472,60472,60416,60416,60378,60378,60342,60342,60307,60307,60278,60278,60239,60239,60222,60222,60175,60175,60154,60154,60136,60136,60095,60095,60074,60074,60042,60042,60004,60004,59987,59987,59968,59968,59948,59948,59932,59932,59878,59878,59855,59855,59827,59827,59805,59805,59772,59772,59749,59749,59732,59732,59715,59715,59695,59695,59669,59669,59652,59652,59630,59630,59612,59612,59588,59588,59571,59571,59500,59500,59471,59471,59410,59410,59390,59390,59345,59345,59302,59302,59282,59282,59244,59244,59180,59180,59155,59155,59113,59113,59072,59072,59037,59037,58990,58990,58966,58966,58944,58944,58908,58908,58886,58886,58868,58868,58845,58845,58824,58824,58801,58801,58761,58761,58742,58742,58713,58713,58675,58675,58640,58640,58613,58613,58577,58577,58553,58553,58534,58534,58499,58499,58443,58443,58426,58426,58394,58394,58378,58378,58329,58329,58306,58306,58289,58289,58229,58229,58213,58213,58193,58193,58154,58154,58132,58132,58107,58107,58089,58089,58070,58070,58052,58052,58019,58019,58003,58003,57969,57969,57949,57949,57908,57908,57873,57873,57835,57835,57814,57814,57784,57784,57758,57758,57721,57721,57699,57699,57664,57664,57591,57591,57558,57558,57541,57541,57517,57517,57496,57496,57479,57479,57457,57457,57400,57400,57370,57370,57331,57331,57283,57283,57260,57260,57235,57235,57195,57195,57170,57170,57125,57125,57060,57060,57041,57041,57006,57006,56964,56964,56939,56939,56914,56914,56891,56891,56862,56862,56841,56841,56817,56817,56781,56781,56763,56763,56734,56734,56703,56703,56685,56685,56666,56666,56650,56650,56618,56618,56592,56592,56558,56558,56525,56525,56472,56472,56437,56437,56397,56397,56375,56375,56356,56356,56324,56324,56304,56304,56283,56283,56260,56260,56235,56235,56185,56185,56161,56161,56139,56139,56096,56096,56079,56079,56006,56006,55989,55989,55924,55924,55891,55891,55860,55860,55837,55837,55810,55810,55785,55785,55726,55726,55708,55708,55680,55680,55651,55651,55632,55632,55596,55596,55576,55576,55549,55549,55518,55518,55487,55487,55458,55458,55410,55410,55350,55350,55333,55333,55284,55284,55258,55258,55233,55233,55184,55184,55155,55155,55137,55137,55085,55085,55067,55067,55050,55050,55022,55022,55004,55004,54979,54979,54953,54953,54925,54925,54886,54886,54868,54868,54830,54830,54810,54810,54783,54783,54766,54766,54741,54741,54700,54700,54684,54684,54657,54657,54638,54638,54612,54612,54585,54585,54538,54538,54519,54519,54503,54503,54478,54478,54415,54415,54389,54389,54368,54368,54305,54305,54263,54263,54247,54247,54218,54218,54191,54191,54169,54169,54147,54147,54130,54130,54108,54108,54071,54071,54048,54048,54031,54031,54006,54006,53990,53990,53963,53963,53928,53928,53908,53908,53859,53859,53837,53837,53749,53749,53716,53716,53679,53679,53655,53655,53630,53630,53611,53611,53560,53560,53524,53524,53502,53502,53440,53440,53369,53369,53349,53349,53329,53329,53296,53296,53246,53246,53214,53214,53197,53197,53149,53149,53129,53129,53101,53101,53034,53034,53003,53003,52974,52974,52926,52926,52880,52880,52862,52862,52833,52833,52794,52794,52764,52764,52745,52745,52710,52710,52692,52692,52673,52673,52641,52641,52618,52618,52580,52580,52549,52549,52519,52519,52482,52482,52453,52453,52432,52432,52389,52389,52353,52353,52322,52322,52295,52295,52276,52276,52245,52245,52228,52228,52212,52212,52185,52185,52148,52148,52119,52119,52087,52087,52059,52059,52007,52007,51983,51983,51957,51957,51940,51940,51915,51915,51892,51892,51868,51868,51839,51839,51821,51821,51804,51804,51788,51788,51761,51761,51744,51744,51726,51726,51701,51701,51656,51656,51611,51611,51587,51587,51560,51560,51525,51525,51509,51509,51469,51469,51441,51441,51422,51422,51398,51398,51378,51378,51350,51350,51304,51304,51280,51280,51243,51243,51160,51160,51135,51135,51119,51119,51069,51069,51027,51027,51003,51003,50985,50985,50950,50950,50907,50907,50870,50870,50816,50816,50790,50790,50750,50750,50720,50720,50682,50682,50632,50632,50608,50608,50587,50587,50559,50559,50542,50542,50523,50523,50501,50501,50453,50453,50418,50418,50397,50397,50346,50346,50274,50274,50254,50254,50229,50229,50207,50207,50167,50167,50141,50141,50115,50115,50075,50075,50049,50049,50027,50027,50008,50008,49988,49988,49972,49972,49945,49945,49914,49914,49898,49898,49858,49858,49829,49829,49790,49790,49754,49754,49726,49726,49687,49687,49655,49655,49636,49636,49608,49608,49590,49590,49554,49554,49504,49504,49471,49471,49451,49451,49427,49427,49411,49411,49389,49389,49371,49371,49348,49348,49324,49324,49305,49305,49288,49288,49257,49257,49235,49235,49219,49219,49199,49199,49122,49122,49083,49083,49060,49060,49044,49044,49020,49020,48988,48988,48970,48970,48924,48924,48889,48889,48865,48865,48846,48846,48823,48823,48799,48799,48769,48769,48749,48749,48727,48727,48709,48709,48693,48693,48675,48675,48658,48658,48626,48626,48570,48570,48551,48551,48511,48511,48474,48474,48425,48425,48396,48396,48376,48376,48321,48321,48298,48298,48265,48265,48243,48243,48221,48221,48204,48204,48162,48162,48145,48145,48112,48112,48083,48083,48063,48063,48044,48044,48028,48028,47994,47994,47972,47972,47945,47945,47909,47909,47890,47890,47870,47870,47844,47844,47811,47811,47765,47765,47748,47748,47732,47732,47712,47712,47691,47691,47666,47666,47646,47646,47624,47624,47557,47557,47530,47530,47500,47500,47467,47467,47423,47423,47403,47403,47386,47386,47360,47360,47326,47326,47302,47302,47279,47279,47243,47243,47225,47225,47165,47165,47137,47137,47119,47119,47087,47087,47068,47068,46991,46991,46962,46962,46933,46933,46886,46886,46856,46856,46831,46831,46815,46815,46790,46790,46774,46774,46753,46753,46734,46734,46717,46717,46692,46692,46666,46666,46632,46632,46615,46615,46571,46571,46542,46542,46497,46497,46477,46477,46461,46461,46430,46430,46401,46401,46372,46372,46345,46345,46303,46303,46261,46261,46232,46232,46211,46211,46139,46139,46117,46117,46094,46094,46077,46077,46061,46061,46035,46035,46019,46019,46002,46002,45985,45985,45965,45965,45938,45938,45911,45911,45873,45873,45835,45835,45814,45814,45779,45779,45749,45749,45730,45730,45703,45703,45682,45682,45653,45653,45628,45628,45606,45606,45590,45590,45565,45565,45547,45547,45526,45526,45507,45507,45488,45488,45467,45467,45425,45425,45368,45368,45338,45338,45308,45308,45275,45275,45192,45192,45168,45168,45150,45150,45127,45127,45111,45111,45076,45076,45051,45051,45031,45031,45007,45007,44984,44984,44959,44959,44917,44917,44892,44892,44825,44825,44809,44809,44760,44760,44736,44736,44716,44716,44699,44699,44681,44681,44658,44658,44642,44642,44621,44621,44589,44589,44552,44552,44519,44519,44459,44459,44433,44433,44400,44400,44382,44382,44360,44360,44343,44343,44327,44327,44311,44311,44292,44292,44258,44258,44232,44232,44216,44216,44192,44192,44162,44162,44141,44141,44117,44117,44089,44089,44032,44032,44012,44012,43975,43975,43952,43952,43936,43936,43892,43892,43874,43874,43848,43848,43827,43827,43777,43777,43750,43750,43713,43713,43691,43691,43665,43665,43645,43645,43610,43610,43582,43582,43555,43555,43532,43532,43516,43516,43499,43499,43472,43472,43450,43450,43434,43434,43412,43412,43391,43391,43370,43370,43335,43335,43289,43289,43271,43271,43255,43255,43230,43230,43205,43205,43180,43180,43155,43155,43136,43136,43115,43115,43078,43078,43040,43040,42989,42989,42965,42965,42944,42944,42928,42928,42906,42906,42880,42880,42861,42861,42843,42843,42827,42827,42803,42803,42773,42773,42749,42749,42733,42733,42710,42710,42691,42691,42669,42669,42619,42619,42579,42579,42560,42560,42534,42534,42513,42513,42495,42495,42465,42465,42440,42440,42412,42412,42378,42378,42361,42361,42342,42342,42323,42323,42303,42303,42286,42286,42270,42270,42247,42247,42229,42229,42183,42183,42163,42163,42131,42131,42109,42109,42087,42087,42066,42066,42048,42048,42031,42031,42014,42014,41997,41997,41977,41977,41961,41961,41945,41945,41923,41923,41891,41891,41870,41870,41854,41854,41834,41834,41815,41815,41797,41797,41781,41781,41703,41703,41683,41683,41651,41651,41630,41630,41611,41611,41592,41592,41540,41540,41498,41498,41465,41465,41445,41445,41408,41408,41389,41389,41364,41364,41342,41342,41289,41289,41265,41265,41248,41248,41227,41227,41195,41195,41154,41154,41138,41138,41113,41113,41086,41086,41068,41068,41011,41011,40973,40973,40938,40938,40910,40910,40886,40886,40863,40863,40804,40804,40788,40788,40767,40767,40734,40734,40718,40718,40688,40688,40666,40666,40628,40628,40595,40595,40554,40554,40535,40535,40519,40519,40498,40498,40478,40478,40455,40455,40434,40434,40405,40405,40379,40379,40348,40348,40308,40308,40291,40291,40272,40272,40250,40250,40229,40229,40212,40212,40175,40175,40157,40157,40135,40135,40115,40115,40085,40085,40065,40065,40039,40039,40021,40021,40005,40005,39987,39987,39962,39962,39945,39945,39922,39922,39880,39880,39862,39862,39835,39835,39819,39819,39798,39798,39773,39773,39682,39682,39658,39658,39639,39639,39618,39618,39599,39599,39573,39573,39557,39557,39533,39533,39503,39503,39482,39482,39461,39461,39444,39444,39422,39422,39390,39390,39347,39347,39331,39331,39303,39303,39250,39250,39223,39223,39191,39191,39171,39171,39151,39151,39097,39097,39078,39078,39055,39055,39027,39027,38990,38990,38954,38954,38931,38931,38911,38911,38873,38873,38845,38845,38829,38829,38781,38781,38755,38755,38731,38731,38709,38709,38686,38686,38665,38665,38621,38621,38605,38605,38544,38544,38517,38517,38484,38484,38453,38453,38436,38436,38420,38420,38400,38400,38383,38383,38361,38361,38344,38344,38326,38326,38309,38309,38276,38276,38253,38253,38236,38236,38218,38218,38192,38192,38171,38171,38152,38152,38134,38134,38116,38116,38098,38098,38065,38065,38041,38041,38024,38024,38007,38007,37985,37985,37958,37958,37936,37936,37909,37909,37886,37886,37867,37867,37851,37851,37830,37830,37808,37808,37788,37788,37763,37763,37741,37741,37724,37724,37704,37704,37671,37671,37648,37648,37625,37625,37608,37608,37569,37569,37543,37543,37516,37516,37494,37494,37467,37467,37445,37445,37426,37426,37388,37388,37370,37370,37336,37336,37320,37320,37302,37302,37285,37285,37255,37255,37221,37221,37182,37182,37147,37147,37127,37127,37107,37107,37077,37077,37057,37057,37039,37039,37013,37013,36993,36993,36962,36962,36925,36925,36907,36907,36889,36889,36870,36870,36854,36854,36829,36829,36801,36801,36780,36780,36737,36737,36719,36719,36703,36703,36680,36680,36659,36659,36621,36621,36601,36601,36541,36541,36513,36513,36490,36490,36471,36471,36453,36453,36429,36429,36409,36409,36383,36383,36367,36367,36346,36346,36323,36323,36305,36305,36286,36286,36268,36268,36251,36251,36222,36222,36194,36194,36158,36158,36136,36136,36116,36116,36094,36094,36063,36063,36045,36045,36015,36015,35970,35970,35954,35954,35928,35928,35907,35907,35891,35891,35870,35870,35853,35853,35831,35831,35810,35810,35794,35794,35776,35776,35723,35723,35704,35704,35685,35685,35667,35667,35637,35637,35621,35621,35605,35605,35576,35576,35546,35546,35525,35525,35500,35500,35484,35484,35419,35419,35400,35400,35384,35384,35342,35342,35292,35292,35274,35274,35254,35254,35231,35231,35198,35198,35182,35182,35166,35166,35148,35148,35131,35131,35115,35115,35090,35090,35041,35041,35017,35017,34994,34994,34968,34968,34950,34950,34915,34915,34895,34895,34877,34877,34856,34856,34837,34837,34817,34817,34793,34793,34776,34776,34758,34758,34742,34742,34725,34725,34706,34706,34690,34690,34671,34671,34654,34654,34637,34637,34615,34615,34586,34586,34566,34566,34548,34548,34518,34518,34474,34474,34411,34411,34394,34394,34376,34376,34349,34349,34322,34322,34303,34303,34285,34285,34269,34269,34250,34250,34231,34231,34205,34205,34189,34189,34167,34167,34129,34129,34105,34105,34085,34085,34064,34064,34048,34048,34031,34031,34000,34000,33980,33980,33961,33961,33942,33942,33896,33896,33878,33878,33833,33833,33800,33800,33775,33775,33758,33758,33716,33716,33699,33699,33681,33681,33642,33642,33620,33620,33597,33597,33557,33557,33500,33500,33482,33482,33422,33422,33405,33405,33388,33388,33371,33371,33355,33355,33338,33338,33316,33316,33300,33300,33281,33281,33258,33258,33242,33242,33216,33216,33197,33197,33179,33179,33156,33156,33137,33137,33094,33094,33078,33078,33062,33062,33028,33028,33011,33011,32995,32995,32972,32972,32948,32948,32913,32913,32863,32863,32830,32830,32811,32811,32767,32767,32751,32751,32715,32715,32699,32699,32671,32671,32654,32654,32630,32630,32607,32607,32590,32590,32549,32549,32517,32517,32500,32500,32484,32484,32464,32464,32441,32441,32425,32425,32402,32402,32366,32366,32350,32350,32313,32313,32295,32295,32278,32278,32247,32247,32231,32231,32200,32200,32175,32175,32158,32158,32142,32142,32125,32125,32105,32105,32089,32089,32066,32066,32047,32047,32029,32029,31999,31999,31980,31980,31957,31957,31939,31939,31923,31923,31905,31905,31885,31885,31863,31863,31842,31842,31820,31820,31788,31788,31755,31755,31728,31728,31712,31712,31692,31692,31662,31662,31645,31645,31623,31623,31607,31607,31591,31591,31571,31571,31551,31551,31520,31520,31503,31503,31483,31483,31460,31460,31444,31444,31425,31425,31407,31407,31389,31389,31370,31370,31344,31344,31323,31323,31300,31300,31284,31284,31260,31260,31244,31244,31216,31216,31177,31177,31159,31159,31140,31140,31118,31118,31094,31094,31070,31070,31050,31050,31034,31034,31015,31015,30982,30982,30965,30965,30938,30938,30905,30905,30884,30884,30864,30864,30831,30831,30806,30806,30787,30787,30769,30769,30730,30730,30707,30707,30689,30689,30672,30672,30649,30649,30632,30632,30616,30616,30600,30600,30584,30584,30568,30568,30546,30546,30522,30522,30502,30502,30483,30483,30460,30460,30439,30439,30422,30422,30391,30391,30368,30368,30348,30348,30311,30311,30292,30292,30263,30263,30239,30239,30211,30211,30184,30184,30159,30159,30118,30118,30097,30097,30059,30059,30039,30039,30023,30023,29977,29977,29952,29952,29933,29933,29916,29916,29898,29898,29872,29872,29848,29848,29827,29827,29802,29802,29779,29779,29757,29757,29730,29730,29705,29705,29688,29688,29664,29664,29635,29635,29618,29618,29601,29601,29580,29580,29564,29564,29547,29547,29522,29522,29475,29475,29449,29449,29432,29432,29411,29411,29394,29394,29370,29370,29341,29341,29317,29317,29298,29298,29282,29282,29266,29266,29250,29250,29226,29226,29205,29205,29188,29188,29168,29168,29151,29151,29107,29107,29086,29086,29069,29069,29048,29048,29027,29027,28983,28983,28964,28964,28948,28948,28931,28931,28914,28914,28897,28897,28869,28869,28847,28847,28829,28829,28806,28806,28784,28784,28765,28765,28743,28743,28723,28723,28686,28686,28666,28666,28640,28640,28621,28621,28605,28605,28589,28589,28572,28572,28549,28549,28531,28531,28510,28510,28492,28492,28471,28471,28451,28451,28431,28431,28415,28415,28399,28399,28371,28371,28355,28355,28339,28339,28322,28322,28303,28303,28286,28286,28266,28266,28249,28249,28226,28226,28201,28201,28181,28181,28165,28165,28146,28146,28130,28130,28113,28113,28096,28096,28080,28080,28060,28060,28044,28044,28027,28027,28004,28004,27987,27987,27971,27971,27951,27951,27933,27933,27914,27914,27894,27894,27877,27877,27851,27851,27828,27828,27812,27812,27795,27795,27779,27779,27759,27759,27739,27739,27717,27717,27701,27701,27684,27684,27662,27662,27634,27634,27616,27616,27592,27592,27570,27570,27550,27550,27525,27525,27505,27505,27483,27483,27460,27460,27441,27441,27424,27424,27403,27403,27387,27387,27371,27371,27355,27355,27338,27338,27322,27322,27305,27305,27284,27284,27267,27267,27251,27251,27229,27229,27206,27206,27189,27189,27172,27172,27156,27156,27140,27140,27118,27118,27096,27096,27077,27077,27061,27061,27045,27045,27028,27028,27011,27011,26993,26993,26964,26964,26947,26947,26931,26931,26915,26915,26891,26891,26875,26875,26844,26844,26825,26825,26807,26807,26790,26790,26761,26761,26744,26744,26717,26717,26698,26698,26679,26679,26663,26663,26646,26646,26629,26629,26610,26610,26594,26594,26576,26576,26541,26541,26518,26518,26499,26499,26480,26480,26460,26460,26444,26444,26427,26427,26410,26410,26391,26391,26375,26375,26354,26354,26334,26334,26313,26313,26291,26291,26267,26267,26247,26247,26228,26228,26201,26201,26183,26183,26150,26150,26129,26129,26113,26113,26088,26088,26064,26064,26040,26040,26022,26022,26004,26004,25980,25980,25964,25964,25946,25946,25930,25930,25911,25911,25893,25893,25861,25861,25840,25840,25821,25821,25805,25805,25786,25786,25770,25770,25753,25753,25727,25727,25711,25711,25695,25695,25674,25674,25658,25658,25641,25641,25621,25621,25600,25600,25584,25584,25565,25565,25547,25547,25523,25523,25507,25507,25491,25491,25474,25474,25456,25456,25438,25438,25421,25421,25402,25402,25386,25386,25370,25370,25349,25349,25333,25333,25316,25316,25296,25296,25274,25274,25258,25258,25240,25240,25224,25224,25200,25200,25181,25181,25162,25162,25140,25140,25108,25108,25090,25090,25073,25073,25056,25056,25031,25031,25014,25014,24996,24996,24980,24980,24961,24961,24940,24940,24921,24921,24896,24896,24878,24878,24861,24861,24845,24845,24828,24828,24811,24811,24794,24794,24778,24778,24755,24755,24739,24739,24716,24716,24696,24696,24669,24669,24651,24651,24634,24634,24617,24617,24595,24595,24571,24571,24555,24555,24539,24539,24513,24513,24492,24492,24475,24475,24455,24455,24437,24437,24419,24419,24403,24403,24385,24385,24366,24366,24347,24347,24317,24317,24301,24301,24284,24284,24262,24262,24246,24246,24227,24227,24204,24204,24188,24188,24172,24172,24150,24150,24129,24129,24108,24108,24091,24091,24074,24074,24058,24058,24041,24041,24025,24025,24005,24005,23989,23989,23962,23962,23945,23945,23922,23922,23906,23906,23889,23889,23872,23872,23851,23851,23823,23823,23798,23798,23777,23777,23761,23761,23739,23739,23716,23716,23697,23697,23671,23671,23651,23651,23634,23634,23616,23616,23598,23598,23581,23581,23557,23557,23535,23535,23516,23516,23496,23496,23478,23478,23458,23458,23433,23433,23417,23417,23400,23400,23384,23384,23368,23368,23350,23350,23334,23334,23315,23315,23297,23297,23281,23281,23261,23261,23244,23244,23215,23215,23198,23198,23179,23179,23163,23163,23142,23142,23124,23124,23108,23108,23090,23090,23066,23066,23047,23047,23028,23028,23012,23012,22995,22995,22978,22978,22955,22955,22938,22938,22910,22910,22892,22892,22872,22872,22852,22852,22831,22831,22809,22809,22775,22775,22756,22756,22740,22740,22719,22719,22702,22702,22685,22685,22669,22669,22652,22652,22636,22636,22620,22620,22600,22600,22582,22582,22564,22564,22543,22543,22526,22526,22510,22510,22494,22494,22477,22477,22461,22461,22433,22433,22416,22416,22400,22400,22383,22383,22365,22365,22348,22348,22330,22330,22313,22313,22292,22292,22269,22269,22253,22253,22236,22236,22219,22219,22201,22201,22184,22184,22164,22164,22147,22147,22130,22130,22112,22112,22096,22096,22079,22079,22063,22063,22047,22047,22030,22030,22014,22014,21998,21998,21982,21982,21964,21964,21947,21947,21928,21928,21912,21912,21890,21890,21873,21873,21855,21855,21839,21839,21819,21819,21802,21802,21780,21780,21756,21756,21740,21740,21721,21721,21704,21704,21685,21685,21658,21658,21633,21633,21615,21615,21598,21598,21582,21582,21564,21564,21547,21547,21530,21530,21514,21514,21496,21496,21478,21478,21453,21453,21436,21436,21420,21420,21400,21400,21382,21382,21364,21364,21340,21340,21323,21323,21305,21305,21281,21281,21265,21265,21249,21249,21232,21232,21215,21215,21199,21199,21180,21180,21164,21164,21140,21140,21120,21120,21104,21104,21082,21082,21065,21065,21046,21046,21009,21009,20993,20993,20975,20975,20959,20959,20942,20942,20926,20926,20908,20908,20891,20891,20875,20875,20859,20859,20843,20843,20826,20826,20810,20810,20793,20793,20776,20776,20754,20754,20734,20734,20718,20718,20702,20702,20684,20684,20664,20664,20648,20648,20632,20632,20614,20614,20596,20596,20580,20580,20561,20561,20545,20545,20528,20528,20506,20506,20485,20485,20465,20465,20445,20445,20428,20428,20402,20402,20385,20385,20368,20368,20348,20348,20331,20331,20312,20312,20294,20294,20273,20273,20257,20257,20240,20240,20221,20221,20203,20203,20187,20187,20171,20171,20151,20151,20134,20134,20116,20116,20099,20099,20080,20080,20064,20064,20045,20045,20029,20029,20011,20011,19993,19993,19976,19976,19959,19959,19943,19943,19923,19923,19906,19906,19890,19890,19874,19874,19854,19854,19833,19833,19817,19817,19794,19794,19777,19777,19761,19761,19745,19745,19723,19723,19705,19705,19689,19689,19666,19666,19649,19649,19632,19632,19612,19612,19596,19596,19580,19580,19562,19562,19541,19541,19525,19525,19504,19504,19486,19486,19461,19461,19443,19443,19426,19426,19410,19410,19394,19394,19375,19375,19358,19358,19334,19334,19317,19317,19300,19300,19278,19278,19262,19262,19245,19245,19228,19228,19212,19212,19191,19191,19175,19175,19155,19155,19136,19136,19119,19119,19103,19103,19082,19082,19064,19064,19046,19046,19027,19027,19011,19011,18995,18995,18978,18978,18962,18962,18946,18946,18924,18924,18902,18902,18885,18885,18866,18866,18850,18850,18834,18834,18818,18818,18802,18802,18783,18783,18766,18766,18750,18750,18734,18734,18716,18716,18700,18700,18682,18682,18666,18666,18649,18649,18632,18632,18608,18608,18589,18589,18573,18573,18557,18557,18538,18538,18522,18522,18504,18504,18487,18487,18471,18471,18454,18454,18436,18436,18415,18415,18398,18398,18380,18380,18364,18364,18348,18348,18332,18332,18314,18314,18297,18297,18281,18281,18264,18264,18248,18248,18231,18231,18214,18214,18198,18198,18182,18182,18166,18166,18150,18150,18134,18134,18112,18112,18095,18095,18077,18077,18061,18061,18043,18043,18026,18026,18004,18004,17985,17985,17968,17968,17951,17951,17934,17934,17918,17918,17900,17900,17884,17884,17857,17857,17840,17840,17824,17824,17806,17806,17789,17789,17772,17772,17756,17756,17739,17739,17723,17723,17705,17705,17689,17689,17673,17673,17657,17657,17640,17640,17624,17624,17605,17605,17588,17588,17568,17568,17547,17547,17531,17531,17515,17515,17499,17499,17483,17483,17467,17467,17450,17450,17434,17434,17413,17413,17397,17397,17381,17381,17365,17365,17349,17349,17332,17332,17315,17315,17298,17298,17280,17280,17261,17261,17243,17243,17227,17227,17207,17207,17190,17190,17169,17169,17153,17153,17136,17136,17119,17119,17098,17098,17082,17082,17064,17064,17042,17042,17026,17026,16947,16947,16927,16927,16829,16829,16094,16094,14701,14701,12916,12916,11586,11586,10948,10948,8779,8779,8350,8350,7853,7853,7581,7581,7463,7463,7150,7150,6888,6888,6566,6566,5885,5885,5461,5461,4958,4958,4146,4146,0.0],[4577433,4577433,4577433,3912653,3912653,3775352,3775352,3418776,3418776,3327411,3327411,3243538,3243538,3212681,3212681,3126350,3126350,3039863,3039863,3029116,3029116,2895199,2895199,2851370,2851370,2788223,2788223,2765367,2765367,2757797,2757797,2742641,2742641,2650157,2650157,2554584,2554584,2516329,2516329,2515412,2515412,2481436,2481436,2474528,2474528,2465855,2465855,2453754,2453754,2446852,2446852,2428360,2428360,2421180,2421180,2405699,2405699,2402388,2402388,2348559,2348559,2344765,2344765,2317563,2317563,2300588,2300588,2299039,2299039,2294499,2294499,2273327,2273327,2272836,2272836,2267708,2267708,2245083,2245083,2243378,2243378,2208784,2208784,2194807,2194807,2192524,2192524,2164150,2164150,2138822,2138822,2131479,2131479,2128015,2128015,2126031,2126031,2103137,2103137,2067256,2067256,2066791,2066791,2065382,2065382,2060156,2060156,2057412,2057412,2053123,2053123,2045520,2045520,2021092,2021092,2020066,2020066,2019173,2019173,1988578,1988578,1965509,1965509,1956557,1956557,1952220,1952220,1944956,1944956,1913379,1913379,1908056,1908056,1906360,1906360,1900371,1900371,1896178,1896178,1872657,1872657,1871414,1871414,1866144,1866144,1863127,1863127,1861138,1861138,1860847,1860847,1859713,1859713,1842844,1842844,1841691,1841691,1838978,1838978,1824588,1824588,1815262,1815262,1774795,1774795,1763159,1763159,1741055,1741055,1733219,1733219,1733163,1733163,1727339,1727339,1698442,1698442,1696305,1696305,1689079,1689079,1686065,1686065,1685748,1685748,1682013,1682013,1676188,1676188,1661209,1661209,1649029,1649029,1642118,1642118,1641521,1641521,1637141,1637141,1629762,1629762,1624855,1624855,1616000,1616000,1609820,1609820,1604721,1604721,1601072,1601072,1597597,1597597,1596447,1596447,1591298,1591298,1574705,1574705,1565570,1565570,1564686,1564686,1559185,1559185,1553275,1553275,1552923,1552923,1549903,1549903,1547340,1547340,1546689,1546689,1541185,1541185,1533216,1533216,1532615,1532615,1531331,1531331,1530391,1530391,1525101,1525101,1524542,1524542,1522307,1522307,1516628,1516628,1515067,1515067,1512256,1512256,1506474,1506474,1503339,1503339,1499001,1499001,1492251,1492251,1480500,1480500,1470360,1470360,1463062,1463062,1460012,1460012,1445718,1445718,1442194,1442194,1435479,1435479,1434600,1434600,1421127,1421127,1418583,1418583,1415807,1415807,1415049,1415049,1413116,1413116,1411189,1411189,1407303,1407303,1401894,1401894,1395507,1395507,1391461,1391461,1389911,1389911,1385631,1385631,1381189,1381189,1376660,1376660,1374219,1374219,1369807,1369807,1368927,1368927,1366540,1366540,1360873,1360873,1355310,1355310,1351940,1351940,1351529,1351529,1350615,1350615,1347654,1347654,1346467,1346467,1340074,1340074,1338940,1338940,1332726,1332726,1332375,1332375,1331669,1331669,1328654,1328654,1324698,1324698,1323914,1323914,1321765,1321765,1319077,1319077,1316233,1316233,1314580,1314580,1312025,1312025,1308677,1308677,1307210,1307210,1304976,1304976,1304286,1304286,1303546,1303546,1302029,1302029,1296626,1296626,1295772,1295772,1291716,1291716,1286271,1286271,1284780,1284780,1283390,1283390,1279995,1279995,1279127,1279127,1278898,1278898,1278787,1278787,1277791,1277791,1275925,1275925,1273407,1273407,1271730,1271730,1270170,1270170,1265832,1265832,1262549,1262549,1262037,1262037,1260560,1260560,1259797,1259797,1256657,1256657,1255499,1255499,1252068,1252068,1250488,1250488,1248547,1248547,1246327,1246327,1242196,1242196,1241175,1241175,1235034,1235034,1229795,1229795,1229629,1229629,1228171,1228171,1226794,1226794,1225973,1225973,1222381,1222381,1221070,1221070,1219091,1219091,1218810,1218810,1215373,1215373,1207777,1207777,1201642,1201642,1198249,1198249,1195858,1195858,1194456,1194456,1185309,1185309,1184741,1184741,1182112,1182112,1180180,1180180,1173566,1173566,1172801,1172801,1169614,1169614,1167411,1167411,1167229,1167229,1163552,1163552,1163245,1163245,1162949,1162949,1162381,1162381,1161300,1161300,1160842,1160842,1160347,1160347,1159284,1159284,1158648,1158648,1157978,1157978,1155951,1155951,1153750,1153750,1150533,1150533,1145209,1145209,1144538,1144538,1140539,1140539,1140436,1140436,1140368,1140368,1133476,1133476,1130233,1130233,1129387,1129387,1125641,1125641,1125178,1125178,1124753,1124753,1123857,1123857,1122426,1122426,1118100,1118100,1114433,1114433,1110820,1110820,1108569,1108569,1108360,1108360,1107897,1107897,1107325,1107325,1106246,1106246,1103062,1103062,1099513,1099513,1098900,1098900,1098745,1098745,1097816,1097816,1097224,1097224,1092938,1092938,1092426,1092426,1090983,1090983,1090346,1090346,1089734,1089734,1087233,1087233,1086056,1086056,1083492,1083492,1082977,1082977,1082835,1082835,1082587,1082587,1077622,1077622,1077088,1077088,1076790,1076790,1076120,1076120,1075769,1075769,1075397,1075397,1075292,1075292,1066804,1066804,1066346,1066346,1063588,1063588,1063427,1063427,1061074,1061074,1059444,1059444,1059334,1059334,1057905,1057905,1056263,1056263,1056119,1056119,1055754,1055754,1054445,1054445,1053401,1053401,1049608,1049608,1049048,1049048,1043801,1043801,1031925,1031925,1029907,1029907,1028438,1028438,1027192,1027192,1023390,1023390,1023362,1023362,1022292,1022292,1021854,1021854,1020691,1020691,1016001,1016001,1014672,1014672,1014313,1014313,1014171,1014171,1013649,1013649,1011971,1011971,1011290,1011290,994554,994554,992754,992754,992571,992571,990609,990609,987976,987976,987265,987265,985847,985847,984689,984689,982438,982438,979583,979583,979302,979302,978176,978176,975339,975339,972970,972970,969648,969648,967736,967736,967249,967249,967004,967004,966414,966414,965674,965674,961611,961611,960899,960899,956394,956394,956064,956064,953933,953933,952790,952790,952299,952299,949000,949000,945019,945019,943035,943035,941916,941916,939436,939436,939225,939225,936943,936943,933972,933972,933732,933732,933147,933147,932743,932743,932377,932377,932313,932313,932027,932027,931263,931263,928556,928556,927885,927885,927538,927538,927251,927251,926548,926548,923165,923165,920972,920972,919906,919906,919600,919600,918190,918190,917696,917696,917414,917414,917248,917248,914827,914827,912133,912133,911581,911581,909830,909830,908967,908967,906969,906969,904818,904818,903982,903982,903339,903339,900782,900782,897340,897340,897070,897070,896641,896641,896359,896359,895309,895309,892776,892776,890862,890862,890193,890193,889257,889257,888304,888304,886645,886645,886055,886055,885925,885925,885209,885209,884874,884874,883473,883473,882723,882723,882624,882624,882061,882061,881938,881938,877353,877353,876897,876897,876255,876255,873733,873733,873010,873010,872243,872243,871648,871648,871145,871145,870963,870963,870900,870900,869660,869660,869507,869507,866810,866810,866722,866722,865411,865411,865340,865340,864734,864734,861758,861758,861699,861699,861568,861568,861338,861338,858153,858153,857616,857616,857443,857443,856866,856866,856160,856160,855626,855626,854521,854521,853949,853949,853763,853763,853027,853027,851502,851502,850782,850782,848359,848359,848236,848236,846634,846634,846027,846027,845480,845480,844151,844151,843381,843381,842663,842663,842447,842447,840899,840899,840351,840351,840085,840085,839575,839575,838600,838600,838483,838483,836707,836707,835412,835412,831742,831742,831589,831589,830403,830403,829834,829834,829788,829788,827933,827933,826935,826935,824931,824931,824885,824885,823451,823451,823419,823419,821036,821036,820565,820565,819968,819968,819467,819467,816431,816431,813266,813266,812606,812606,812578,812578,811770,811770,810990,810990,810915,810915,810779,810779,807335,807335,805746,805746,803832,803832,803665,803665,802953,802953,801332,801332,801299,801299,800601,800601,800312,800312,799767,799767,799580,799580,799090,799090,795196,795196,795121,795121,793532,793532,792382,792382,792342,792342,792006,792006,791141,791141,790122,790122,789177,789177,788805,788805,788518,788518,787368,787368,787301,787301,787015,787015,786885,786885,786853,786853,786247,786247,785650,785650,784698,784698,783383,783383,782911,782911,781854,781854,779907,779907,779122,779122,778789,778789,777700,777700,776008,776008,773453,773453,772750,772750,772197,772197,771857,771857,771213,771213,770630,770630,769841,769841,769665,769665,763632,763632,763196,763196,762737,762737,762721,762721,761852,761852,761632,761632,761391,761391,761373,761373,760707,760707,760017,760017,759892,759892,758837,758837,758613,758613,758338,758338,755831,755831,755674,755674,755180,755180,752779,752779,752511,752511,752306,752306,751664,751664,751465,751465,751198,751198,749332,749332,748896,748896,748851,748851,748716,748716,748163,748163,747801,747801,747325,747325,747114,747114,745601,745601,745248,745248,743765,743765,739256,739256,736805,736805,734637,734637,734561,734561,733107,733107,731702,731702,731587,731587,730878,730878,730609,730609,728135,728135,727794,727794,727386,727386,726979,726979,724911,724911,724443,724443,724270,724270,724234,724234,723735,723735,723230,723230,722469,722469,722239,722239,721741,721741,721025,721025,720866,720866,719461,719461,719094,719094,717365,717365,717048,717048,716995,716995,716813,716813,716557,716557,716494,716494,716397,716397,715632,715632,714229,714229,714181,714181,711760,711760,710807,710807,709802,709802,709744,709744,709581,709581,709409,709409,708599,708599,708489,708489,708421,708421,708126,708126,707868,707868,707749,707749,707681,707681,707418,707418,706964,706964,706494,706494,705869,705869,705815,705815,705252,705252,704382,704382,703521,703521,701794,701794,700919,700919,700883,700883,700054,700054,699133,699133,698769,698769,697105,697105,696467,696467,696430,696430,694908,694908,694440,694440,694367,694367,693472,693472,692805,692805,692788,692788,692420,692420,692322,692322,692154,692154,692109,692109,691453,691453,691361,691361,690986,690986,690875,690875,690607,690607,689873,689873,689734,689734,689383,689383,688815,688815,688734,688734,688708,688708,688552,688552,688028,688028,687884,687884,685847,685847,685377,685377,684231,684231,683807,683807,683770,683770,683279,683279,682512,682512,682383,682383,682035,682035,680406,680406,679362,679362,678860,678860,677452,677452,677266,677266,677205,677205,675822,675822,675322,675322,674625,674625,673905,673905,673813,673813,672621,672621,672315,672315,671596,671596,670506,670506,668738,668738,668179,668179,667753,667753,667681,667681,666917,666917,666695,666695,666494,666494,666337,666337,666062,666062,665938,665938,664838,664838,664115,664115,663520,663520,663248,663248,663229,663229,663212,663212,662721,662721,662189,662189,661905,661905,661857,661857,660506,660506,660299,660299,659755,659755,659101,659101,658513,658513,658050,658050,657840,657840,657148,657148,654250,654250,653107,653107,652165,652165,652078,652078,652047,652047,651297,651297,651112,651112,650592,650592,650428,650428,648401,648401,648147,648147,647317,647317,646772,646772,644396,644396,644246,644246,644121,644121,644022,644022,643786,643786,643723,643723,642371,642371,642342,642342,641709,641709,641586,641586,641490,641490,641147,641147,640730,640730,640592,640592,640518,640518,640464,640464,640052,640052,639340,639340,638734,638734,638506,638506,637811,637811,637466,637466,637252,637252,635274,635274,635131,635131,634962,634962,634663,634663,634441,634441,634302,634302,633696,633696,633351,633351,633277,633277,633142,633142,632131,632131,631975,631975,631542,631542,631284,631284,631237,631237,629785,629785,629592,629592,629322,629322,629119,629119,628261,628261,628043,628043,626600,626600,626495,626495,625812,625812,625229,625229,625205,625205,625112,625112,624917,624917,623692,623692,623590,623590,623375,623375,622728,622728,622604,622604,622561,622561,621517,621517,621293,621293,620954,620954,619077,619077,618940,618940,618782,618782,618003,618003,617918,617918,617869,617869,617403,617403,616944,616944,615646,615646,615506,615506,615490,615490,615384,615384,615095,615095,614707,614707,614280,614280,613830,613830,613385,613385,613248,613248,613019,613019,612657,612657,611870,611870,610005,610005,609779,609779,609641,609641,609334,609334,609176,609176,608660,608660,608470,608470,608208,608208,606566,606566,606107,606107,605951,605951,605532,605532,604381,604381,604003,604003,603963,603963,603879,603879,603285,603285,603096,603096,602842,602842,601994,601994,601376,601376,600853,600853,600714,600714,598899,598899,598712,598712,598468,598468,598448,598448,598198,598198,598149,598149,596024,596024,595612,595612,594954,594954,594490,594490,593076,593076,592765,592765,592554,592554,592456,592456,592053,592053,591305,591305,590371,590371,590311,590311,589448,589448,589188,589188,586638,586638,586462,586462,586325,586325,586206,586206,585799,585799,585634,585634,585488,585488,585193,585193,585041,585041,584949,584949,584783,584783,584398,584398,583915,583915,583225,583225,583105,583105,583043,583043,582936,582936,582206,582206,581831,581831,581758,581758,580847,580847,580577,580577,580513,580513,580388,580388,580012,580012,579735,579735,578636,578636,578335,578335,578172,578172,577966,577966,577594,577594,577105,577105,576558,576558,576237,576237,575886,575886,575341,575341,574781,574781,574416,574416,574179,574179,573187,573187,572249,572249,571828,571828,571318,571318,570988,570988,570740,570740,570644,570644,569574,569574,569355,569355,569071,569071,569045,569045,568969,568969,567824,567824,567762,567762,567563,567563,567156,567156,566833,566833,566769,566769,565989,565989,565963,565963,565362,565362,564013,564013,563592,563592,563356,563356,563112,563112,562857,562857,562081,562081,561460,561460,561264,561264,561080,561080,560679,560679,560503,560503,560351,560351,559686,559686,559629,559629,559237,559237,559118,559118,559006,559006,558660,558660,558507,558507,558448,558448,558400,558400,558209,558209,558110,558110,557777,557777,557161,557161,557007,557007,556812,556812,556642,556642,555852,555852,555770,555770,555561,555561,555122,555122,555012,555012,554959,554959,554610,554610,553973,553973,553224,553224,552029,552029,551267,551267,550189,550189,549479,549479,549004,549004,548891,548891,548613,548613,547712,547712,547574,547574,547129,547129,547079,547079,546861,546861,546604,546604,546396,546396,546376,546376,546152,546152,545891,545891,545835,545835,545738,545738,545706,545706,545341,545341,545038,545038,544744,544744,544693,544693,544658,544658,544467,544467,544089,544089,543740,543740,543093,543093,541935,541935,541464,541464,541387,541387,541303,541303,539788,539788,539769,539769,539065,539065,538869,538869,537831,537831,536613,536613,535322,535322,534980,534980,534909,534909,534149,534149,533796,533796,533681,533681,533635,533635,533467,533467,532914,532914,532755,532755,532511,532511,532151,532151,531544,531544,530941,530941,530531,530531,530056,530056,530015,530015,529832,529832,528958,528958,528817,528817,528622,528622,528428,528428,528255,528255,528214,528214,528075,528075,527925,527925,527049,527049,527007,527007,526965,526965,526607,526607,526317,526317,525729,525729,525619,525619,525342,525342,523887,523887,523796,523796,523702,523702,523508,523508,522883,522883,522799,522799,522398,522398,521952,521952,521855,521855,521717,521717,521181,521181,520818,520818,520457,520457,520113,520113,519930,519930,519723,519723,519698,519698,519510,519510,519185,519185,519021,519021,518998,518998,518598,518598,518530,518530,518073,518073,517846,517846,517751,517751,517621,517621,517442,517442,517362,517362,517262,517262,516939,516939,516707,516707,516067,516067,515897,515897,515880,515880,515762,515762,515744,515744,515617,515617,515244,515244,514782,514782,514445,514445,514309,514309,514199,514199,513571,513571,513529,513529,512325,512325,511624,511624,510860,510860,510662,510662,510582,510582,510313,510313,509780,509780,509730,509730,509488,509488,509160,509160,508662,508662,508511,508511,507336,507336,507173,507173,506691,506691,506123,506123,505856,505856,505540,505540,505493,505493,505454,505454,505410,505410,505074,505074,504366,504366,504233,504233,503619,503619,503591,503591,503485,503485,503277,503277,503257,503257,503189,503189,503087,503087,501917,501917,501553,501553,501097,501097,500932,500932,500869,500869,500094,500094,500046,500046,499940,499940,499739,499739,499603,499603,499496,499496,499328,499328,498231,498231,497231,497231,496877,496877,496692,496692,496516,496516,496430,496430,496076,496076,496036,496036,495644,495644,495136,495136,494835,494835,494622,494622,494539,494539,494204,494204,493969,493969,493917,493917,493755,493755,493418,493418,492414,492414,492260,492260,492218,492218,492102,492102,491889,491889,491718,491718,491581,491581,491457,491457,491401,491401,491098,491098,490979,490979,490790,490790,490756,490756,490720,490720,490556,490556,490178,490178,490048,490048,489743,489743,488884,488884,488772,488772,488337,488337,488234,488234,488201,488201,487924,487924,487495,487495,487409,487409,487008,487008,486874,486874,486405,486405,486375,486375,486234,486234,486054,486054,485911,485911,485401,485401,484654,484654,484328,484328,483836,483836,483603,483603,483386,483386,483260,483260,482915,482915,482796,482796,482266,482266,482090,482090,482021,482021,481198,481198,481174,481174,480101,480101,479881,479881,479816,479816,479108,479108,478631,478631,478276,478276,478033,478033,477799,477799,477759,477759,477672,477672,477219,477219,477162,477162,476492,476492,476376,476376,476342,476342,476164,476164,475561,475561,474991,474991,474894,474894,474581,474581,474555,474555,474401,474401,473906,473906,473442,473442,473376,473376,472733,472733,472712,472712,472681,472681,472170,472170,471832,471832,471661,471661,471381,471381,470994,470994,469959,469959,469792,469792,469705,469705,469399,469399,469267,469267,469142,469142,468847,468847,468764,468764,468180,468180,468157,468157,467705,467705,467426,467426,467341,467341,465246,465246,464997,464997,464940,464940,464922,464922,464828,464828,464610,464610,464547,464547,464359,464359,464164,464164,464116,464116,464015,464015,463842,463842,463190,463190,462628,462628,462200,462200,462091,462091,461698,461698,461340,461340,461181,461181,460999,460999,460950,460950,460862,460862,460784,460784,460642,460642,460008,460008,459884,459884,459859,459859,459787,459787,459458,459458,459428,459428,458744,458744,458498,458498,458464,458464,458358,458358,458085,458085,457768,457768,457545,457545,457519,457519,457460,457460,457092,457092,456719,456719,456553,456553,456384,456384,456091,456091,455908,455908,455539,455539,455417,455417,455308,455308,455054,455054,454953,454953,454466,454466,454381,454381,454341,454341,454232,454232,453241,453241,453160,453160,452956,452956,452662,452662,452621,452621,452562,452562,452374,452374,452005,452005,451729,451729,451646,451646,451321,451321,451215,451215,451148,451148,450993,450993,450562,450562,449918,449918,449895,449895,449278,449278,449110,449110,448534,448534,448204,448204,448070,448070,448036,448036,448004,448004,447719,447719,447100,447100,447036,447036,446749,446749,446281,446281,446243,446243,446187,446187,445875,445875,445741,445741,444987,444987,444791,444791,444092,444092,444068,444068,443960,443960,443765,443765,443628,443628,443559,443559,443293,443293,442893,442893,442863,442863,442817,442817,442440,442440,442194,442194,442167,442167,441997,441997,441929,441929,441699,441699,441415,441415,441357,441357,441308,441308,440994,440994,440876,440876,440703,440703,440546,440546,439650,439650,439613,439613,439491,439491,439448,439448,439182,439182,438375,438375,438290,438290,437792,437792,437435,437435,437156,437156,436885,436885,436755,436755,436316,436316,436189,436189,435904,435904,435539,435539,435481,435481,434825,434825,434809,434809,434610,434610,433834,433834,432988,432988,432565,432565,432450,432450,432364,432364,432259,432259,432028,432028,431926,431926,431843,431843,430925,430925,430842,430842,430771,430771,430737,430737,430383,430383,430244,430244,430141,430141,429799,429799,429665,429665,429477,429477,429410,429410,429288,429288,429001,429001,428648,428648,428533,428533,428322,428322,428019,428019,427983,427983,427778,427778,427354,427354,427333,427333,427154,427154,427017,427017,426746,426746,426378,426378,426133,426133,426052,426052,425810,425810,425739,425739,425449,425449,425413,425413,425376,425376,425275,425275,425228,425228,425082,425082,425058,425058,425030,425030,424805,424805,424464,424464,424408,424408,424228,424228,424051,424051,423998,423998,423903,423903,423756,423756,423058,423058,422937,422937,422683,422683,422388,422388,422269,422269,422203,422203,422164,422164,422065,422065,421979,421979,421718,421718,421590,421590,421030,421030,420972,420972,420924,420924,420901,420901,420875,420875,420797,420797,420704,420704,420668,420668,420586,420586,420380,420380,419965,419965,419623,419623,418963,418963,418874,418874,418705,418705,418640,418640,418601,418601,418462,418462,418216,418216,417605,417605,417403,417403,416973,416973,416720,416720,416603,416603,416564,416564,416510,416510,415851,415851,415663,415663,415621,415621,414969,414969,414782,414782,414735,414735,414205,414205,414114,414114,413877,413877,413837,413837,413589,413589,413394,413394,413274,413274,413253,413253,412418,412418,412223,412223,411957,411957,411604,411604,411502,411502,411463,411463,411302,411302,411135,411135,411091,411091,410836,410836,410798,410798,410558,410558,410201,410201,410162,410162,409962,409962,409823,409823,409501,409501,409357,409357,409114,409114,408975,408975,408825,408825,408627,408627,408496,408496,408082,408082,408045,408045,407463,407463,407194,407194,407047,407047,406942,406942,406836,406836,406659,406659,406485,406485,406387,406387,405991,405991,405685,405685,405630,405630,405239,405239,405030,405030,404917,404917,404811,404811,404722,404722,404597,404597,404565,404565,404384,404384,404056,404056,403501,403501,403391,403391,403218,403218,403158,403158,403022,403022,402873,402873,402574,402574,402519,402519,401859,401859,401802,401802,401706,401706,401457,401457,401239,401239,401127,401127,400884,400884,400349,400349,400217,400217,400069,400069,399993,399993,399976,399976,399864,399864,399585,399585,399540,399540,399504,399504,399484,399484,399167,399167,399141,399141,398906,398906,398789,398789,398767,398767,398536,398536,398373,398373,397844,397844,397821,397821,397486,397486,396978,396978,396722,396722,396610,396610,396387,396387,396093,396093,395935,395935,395864,395864,395700,395700,395657,395657,395235,395235,395093,395093,395075,395075,395033,395033,395017,395017,394995,394995,394934,394934,394590,394590,394312,394312,394279,394279,394237,394237,394166,394166,393910,393910,393569,393569,393371,393371,393129,393129,392865,392865,392631,392631,392499,392499,392252,392252,392136,392136,392093,392093,391840,391840,391355,391355,391273,391273,391169,391169,391018,391018,390770,390770,390270,390270,390237,390237,390030,390030,389719,389719,389610,389610,389480,389480,389274,389274,389162,389162,389078,389078,388822,388822,388776,388776,388700,388700,388245,388245,388122,388122,388025,388025,387821,387821,387700,387700,387658,387658,387612,387612,387522,387522,387426,387426,387080,387080,386961,386961,386570,386570,386482,386482,386319,386319,385960,385960,385565,385565,385489,385489,385326,385326,385171,385171,385121,385121,384528,384528,384027,384027,383764,383764,383728,383728,383703,383703,383579,383579,383483,383483,383379,383379,383331,383331,383310,383310,383007,383007,382967,382967,382944,382944,382797,382797,382682,382682,382493,382493,382396,382396,381995,381995,381799,381799,381764,381764,381727,381727,381524,381524,381440,381440,381123,381123,381052,381052,381031,381031,380990,380990,380965,380965,380688,380688,380646,380646,380447,380447,380338,380338,380099,380099,379948,379948,379832,379832,379666,379666,379485,379485,379030,379030,378726,378726,378546,378546,378091,378091,378073,378073,377876,377876,377827,377827,377751,377751,377662,377662,377533,377533,377516,377516,377402,377402,377231,377231,376721,376721,376607,376607,376299,376299,376159,376159,376107,376107,376064,376064,375990,375990,375839,375839,375758,375758,375632,375632,375528,375528,375500,375500,375404,375404,375219,375219,374905,374905,374859,374859,374824,374824,374570,374570,374462,374462,374292,374292,374041,374041,374003,374003,373893,373893,373181,373181,373055,373055,372880,372880,372767,372767,372720,372720,372571,372571,372072,372072,371991,371991,371965,371965,371907,371907,371667,371667,371610,371610,371579,371579,371173,371173,370487,370487,370318,370318,370145,370145,370123,370123,370061,370061,369858,369858,369782,369782,369302,369302,369218,369218,369124,369124,368871,368871,368820,368820,368784,368784,368713,368713,368660,368660,368415,368415,367916,367916,367875,367875,367845,367845,367670,367670,367548,367548,367447,367447,367352,367352,367236,367236,367093,367093,366956,366956,366880,366880,366850,366850,366727,366727,366660,366660,366271,366271,366221,366221,365985,365985,365851,365851,365787,365787,365740,365740,365647,365647,365500,365500,365244,365244,365140,365140,365087,365087,365068,365068,364970,364970,364891,364891,364799,364799,364759,364759,364563,364563,364519,364519,364477,364477,364360,364360,364118,364118,364002,364002,363667,363667,363577,363577,363429,363429,363274,363274,363151,363151,362519,362519,362424,362424,362292,362292,362193,362193,362091,362091,362017,362017,361950,361950,361844,361844,361183,361183,361106,361106,361064,361064,360984,360984,360832,360832,360707,360707,360670,360670,360568,360568,360460,360460,359952,359952,359744,359744,359547,359547,359441,359441,359002,359002,358978,358978,358894,358894,358618,358618,357892,357892,357828,357828,357729,357729,357239,357239,356846,356846,356774,356774,356740,356740,356703,356703,356356,356356,356273,356273,356170,356170,356091,356091,356044,356044,355899,355899,355503,355503,355457,355457,355113,355113,355092,355092,354955,354955,354780,354780,354495,354495,354458,354458,354365,354365,354298,354298,353473,353473,353354,353354,353303,353303,353227,353227,353002,353002,352722,352722,352688,352688,352562,352562,352434,352434,352312,352312,352281,352281,352265,352265,352247,352247,352153,352153,352057,352057,351755,351755,351694,351694,351623,351623,351201,351201,351099,351099,351074,351074,350940,350940,350729,350729,350122,350122,350043,350043,350005,350005,349826,349826,349635,349635,349617,349617,349586,349586,349357,349357,349240,349240,349223,349223,349181,349181,349137,349137,349060,349060,348507,348507,348289,348289,348192,348192,348104,348104,348063,348063,347835,347835,347806,347806,347586,347586,347464,347464,347439,347439,347355,347355,347306,347306,347247,347247,347131,347131,347108,347108,347049,347049,346800,346800,346526,346526,346445,346445,346333,346333,346181,346181,346033,346033,346017,346017,345829,345829,345805,345805,345135,345135,345059,345059,344915,344915,344810,344810,344769,344769,344590,344590,344535,344535,344517,344517,344359,344359,344273,344273,344221,344221,344070,344070,343910,343910,343522,343522,343474,343474,343454,343454,343402,343402,343136,343136,342852,342852,342595,342595,342577,342577,342489,342489,342252,342252,342188,342188,341945,341945,341867,341867,341846,341846,341678,341678,341623,341623,341594,341594,341384,341384,341269,341269,341161,341161,341110,341110,341083,341083,340844,340844,340818,340818,340650,340650,340564,340564,340375,340375,340350,340350,340320,340320,340267,340267,340148,340148,339964,339964,339872,339872,339843,339843,339608,339608,339441,339441,339302,339302,339127,339127,339109,339109,339078,339078,338951,338951,338407,338407,338359,338359,338320,338320,338263,338263,337699,337699,337672,337672,337622,337622,337591,337591,337536,337536,337391,337391,337126,337126,337007,337007,336832,336832,336791,336791,336595,336595,336339,336339,336051,336051,336011,336011,335515,335515,335457,335457,335285,335285,335052,335052,334922,334922,334229,334229,334205,334205,334073,334073,333973,333973,333826,333826,333774,333774,333758,333758,333639,333639,333441,333441,333381,333381,333126,333126,332991,332991,332943,332943,332730,332730,332696,332696,332547,332547,332506,332506,332229,332229,332185,332185,331841,331841,331778,331778,331734,331734,331419,331419,331350,331350,331326,331326,331288,331288,331214,331214,331006,331006,330944,330944,330773,330773,330697,330697,330633,330633,330309,330309,330215,330215,330083,330083,330051,330051,330001,330001,329962,329962,329585,329585,329558,329558,329488,329488,329125,329125,329105,329105,329053,329053,328949,328949,328829,328829,328723,328723,328697,328697,328365,328365,328190,328190,328033,328033,327896,327896,327573,327573,327481,327481,327305,327305,327220,327220,327202,327202,327134,327134,326899,326899,326511,326511,326141,326141,326088,326088,326050,326050,326019,326019,325915,325915,325424,325424,325134,325134,324894,324894,324773,324773,324516,324516,324464,324464,324414,324414,324322,324322,324253,324253,324208,324208,324180,324180,323985,323985,323921,323921,323608,323608,323290,323290,323274,323274,322993,322993,322864,322864,322836,322836,322781,322781,322579,322579,322563,322563,322458,322458,322166,322166,322115,322115,321897,321897,321830,321830,321603,321603,321507,321507,321484,321484,321448,321448,321047,321047,320987,320987,320951,320951,320897,320897,320782,320782,320239,320239,320115,320115,320019,320019,319956,319956,319677,319677,319643,319643,319378,319378,319312,319312,318881,318881,318573,318573,318348,318348,318078,318078,318007,318007,317858,317858,317719,317719,317665,317665,317588,317588,317471,317471,317412,317412,317213,317213,317129,317129,316969,316969,316904,316904,316739,316739,316647,316647,316607,316607,316209,316209,316149,316149,316038,316038,315946,315946,315868,315868,315752,315752,315707,315707,315448,315448,315389,315389,315313,315313,314813,314813,314303,314303,314174,314174,314077,314077,313883,313883,313778,313778,313718,313718,313596,313596,313373,313373,313264,313264,313175,313175,313083,313083,312977,312977,312514,312514,312107,312107,312075,312075,311979,311979,311846,311846,311792,311792,311769,311769,311649,311649,311629,311629,311578,311578,311259,311259,311215,311215,311171,311171,311155,311155,311118,311118,311027,311027,310851,310851,310812,310812,310662,310662,310507,310507,310177,310177,309987,309987,309953,309953,309926,309926,309745,309745,309650,309650,309274,309274,309124,309124,308921,308921,308809,308809,308581,308581,308541,308541,308508,308508,308419,308419,308274,308274,308165,308165,308122,308122,307979,307979,307941,307941,307918,307918,307895,307895,307747,307747,307725,307725,307707,307707,307565,307565,307510,307510,307432,307432,307191,307191,307163,307163,307081,307081,306954,306954,306929,306929,306853,306853,306829,306829,306583,306583,306540,306540,306491,306491,306416,306416,306196,306196,306022,306022,305774,305774,305593,305593,305327,305327,305200,305200,305118,305118,305099,305099,304951,304951,304932,304932,304700,304700,304602,304602,304381,304381,304365,304365,304307,304307,303663,303663,303579,303579,303385,303385,303270,303270,303212,303212,303074,303074,302921,302921,302768,302768,302677,302677,302316,302316,302246,302246,301870,301870,301766,301766,301607,301607,301196,301196,301167,301167,301108,301108,301089,301089,301010,301010,300905,300905,300846,300846,300372,300372,300327,300327,300298,300298,300252,300252,300105,300105,299944,299944,299757,299757,299730,299730,299601,299601,299559,299559,299478,299478,299425,299425,299363,299363,299037,299037,298956,298956,298906,298906,298813,298813,298723,298723,298668,298668,298452,298452,298125,298125,298105,298105,298056,298056,297972,297972,297880,297880,297800,297800,297608,297608,297592,297592,297150,297150,296927,296927,296873,296873,296755,296755,296715,296715,296591,296591,296389,296389,296289,296289,296232,296232,296212,296212,296134,296134,296079,296079,296012,296012,295914,295914,295813,295813,295701,295701,295571,295571,295469,295469,295357,295357,295233,295233,295046,295046,294954,294954,294846,294846,294822,294822,294654,294654,294594,294594,294547,294547,294367,294367,294346,294346,294300,294300,294194,294194,293836,293836,293626,293626,293508,293508,293481,293481,293095,293095,293075,293075,293059,293059,292972,292972,292874,292874,292839,292839,292716,292716,292608,292608,292578,292578,292445,292445,292318,292318,292266,292266,292220,292220,292173,292173,292063,292063,291958,291958,291916,291916,291877,291877,291829,291829,291755,291755,291472,291472,291440,291440,291367,291367,291218,291218,291063,291063,290904,290904,290825,290825,290627,290627,290592,290592,290447,290447,290339,290339,290053,290053,290031,290031,289990,289990,289964,289964,289789,289789,289721,289721,289670,289670,289607,289607,289577,289577,289187,289187,289132,289132,289116,289116,288954,288954,288923,288923,288858,288858,288808,288808,288593,288593,288228,288228,287922,287922,287721,287721,287672,287672,287554,287554,287405,287405,287345,287345,287143,287143,287112,287112,286990,286990,286884,286884,286825,286825,286723,286723,286673,286673,286596,286596,286571,286571,286537,286537,286485,286485,286021,286021,285861,285861,285811,285811,285621,285621,285596,285596,285243,285243,285082,285082,285011,285011,284820,284820,284774,284774,284297,284297,284244,284244,284210,284210,284147,284147,284107,284107,283872,283872,283485,283485,283350,283350,283236,283236,283164,283164,282978,282978,282700,282700,282582,282582,282562,282562,282536,282536,282414,282414,282352,282352,282310,282310,282285,282285,282255,282255,282139,282139,282120,282120,282066,282066,281918,281918,281856,281856,281797,281797,281729,281729,281691,281691,281551,281551,281431,281431,281311,281311,281090,281090,281023,281023,281007,281007,280969,280969,280790,280790,280767,280767,280664,280664,280584,280584,280488,280488,280463,280463,280419,280419,280283,280283,280245,280245,280095,280095,280032,280032,279900,279900,279875,279875,279831,279831,279719,279719,279343,279343,279234,279234,279038,279038,278978,278978,278932,278932,278848,278848,278814,278814,278727,278727,278690,278690,278534,278534,278489,278489,278469,278469,278453,278453,278380,278380,278313,278313,278190,278190,278117,278117,278028,278028,277900,277900,277689,277689,277657,277657,277629,277629,277482,277482,277416,277416,277376,277376,277356,277356,277289,277289,277196,277196,277010,277010,276871,276871,276845,276845,276812,276812,276764,276764,276714,276714,276461,276461,276167,276167,276109,276109,276049,276049,275886,275886,275578,275578,275487,275487,275471,275471,275397,275397,275271,275271,275152,275152,275093,275093,275001,275001,274934,274934,274870,274870,274762,274762,274690,274690,274396,274396,274054,274054,273985,273985,273966,273966,273903,273903,273750,273750,273638,273638,273616,273616,273500,273500,273382,273382,273334,273334,273075,273075,273015,273015,272908,272908,272885,272885,272790,272790,272722,272722,272456,272456,272370,272370,272209,272209,271873,271873,271635,271635,271619,271619,271468,271468,271249,271249,271190,271190,271174,271174,270942,270942,270844,270844,270682,270682,270655,270655,270618,270618,270451,270451,270372,270372,270322,270322,270253,270253,269753,269753,269590,269590,269444,269444,269345,269345,269195,269195,269151,269151,269024,269024,268938,268938,268773,268773,268549,268549,268338,268338,268303,268303,268141,268141,268043,268043,267942,267942,267130,267130,267104,267104,267032,267032,266994,266994,266709,266709,266593,266593,266462,266462,266383,266383,266337,266337,266228,266228,266119,266119,265874,265874,265751,265751,265568,265568,265491,265491,265414,265414,265361,265361,265306,265306,265268,265268,265222,265222,265096,265096,264823,264823,264560,264560,264445,264445,264355,264355,264263,264263,264108,264108,264066,264066,264039,264039,263911,263911,263776,263776,263620,263620,263457,263457,263434,263434,263153,263153,263071,263071,262893,262893,262840,262840,262728,262728,262695,262695,262609,262609,262583,262583,262509,262509,262436,262436,262349,262349,262146,262146,262083,262083,262051,262051,262007,262007,261765,261765,261642,261642,261589,261589,261490,261490,261383,261383,261310,261310,261287,261287,261247,261247,261140,261140,261045,261045,260752,260752,260710,260710,260464,260464,260416,260416,260375,260375,260256,260256,260181,260181,260119,260119,259758,259758,259648,259648,259449,259449,259303,259303,259263,259263,259126,259126,259090,259090,259062,259062,259037,259037,259015,259015,258899,258899,258878,258878,258744,258744,258573,258573,258468,258468,258392,258392,258367,258367,258201,258201,258171,258171,258137,258137,257685,257685,257595,257595,257562,257562,257544,257544,257477,257477,257411,257411,257367,257367,257218,257218,257197,257197,257159,257159,257059,257059,256979,256979,256880,256880,256807,256807,256763,256763,256576,256576,256531,256531,256476,256476,256448,256448,256407,256407,256333,256333,256036,256036,256020,256020,255909,255909,255847,255847,255779,255779,255628,255628,255612,255612,255465,255465,255366,255366,255320,255320,255189,255189,255121,255121,255072,255072,255012,255012,254950,254950,254868,254868,254750,254750,254689,254689,254618,254618,254507,254507,254411,254411,254365,254365,254336,254336,254005,254005,253969,253969,253949,253949,253903,253903,253854,253854,253819,253819,253559,253559,253495,253495,253458,253458,253421,253421,253069,253069,253003,253003,252978,252978,252807,252807,252716,252716,252616,252616,252565,252565,252518,252518,252360,252360,252289,252289,252101,252101,252052,252052,251979,251979,251922,251922,251463,251463,251399,251399,251376,251376,251291,251291,251184,251184,251134,251134,251054,251054,250969,250969,250936,250936,250897,250897,250869,250869,250851,250851,250335,250335,250212,250212,250068,250068,250045,250045,249774,249774,249729,249729,249666,249666,249521,249521,249462,249462,249411,249411,249318,249318,249272,249272,249237,249237,249187,249187,249140,249140,248965,248965,248913,248913,248873,248873,248848,248848,248803,248803,248764,248764,248739,248739,248664,248664,248606,248606,248437,248437,248285,248285,248136,248136,248082,248082,248035,248035,248018,248018,247991,247991,247973,247973,247549,247549,247467,247467,247444,247444,247420,247420,247372,247372,247343,247343,247277,247277,247104,247104,247083,247083,247067,247067,246941,246941,246853,246853,246664,246664,246532,246532,246505,246505,246466,246466,246360,246360,246317,246317,246274,246274,246251,246251,246169,246169,246132,246132,245972,245972,245859,245859,245809,245809,245041,245041,244960,244960,244942,244942,244922,244922,244851,244851,244725,244725,244706,244706,244689,244689,244666,244666,244612,244612,244556,244556,244495,244495,244290,244290,244196,244196,244175,244175,244062,244062,244009,244009,243859,243859,243679,243679,243643,243643,243532,243532,243456,243456,243125,243125,243051,243051,242884,242884,242865,242865,242849,242849,242697,242697,242571,242571,242397,242397,242284,242284,242237,242237,242199,242199,242001,242001,241968,241968,241920,241920,241815,241815,241770,241770,241707,241707,241595,241595,241520,241520,241356,241356,241299,241299,241267,241267,241251,241251,241211,241211,241045,241045,240910,240910,240851,240851,240804,240804,240728,240728,240646,240646,240191,240191,240062,240062,239971,239971,239930,239930,239848,239848,239806,239806,239627,239627,239591,239591,239575,239575,239471,239471,239416,239416,239268,239268,239238,239238,239100,239100,238968,238968,238883,238883,238819,238819,238760,238760,238564,238564,238323,238323,238106,238106,237967,237967,237887,237887,237851,237851,237829,237829,237723,237723,237641,237641,237511,237511,237470,237470,237434,237434,237410,237410,237203,237203,237176,237176,237114,237114,237082,237082,236828,236828,236656,236656,236618,236618,236484,236484,236355,236355,236303,236303,236277,236277,236235,236235,235973,235973,235883,235883,235793,235793,235667,235667,235556,235556,235513,235513,235348,235348,235267,235267,235176,235176,235030,235030,234971,234971,234943,234943,234888,234888,234845,234845,234773,234773,234739,234739,234606,234606,234441,234441,234380,234380,234192,234192,234148,234148,234060,234060,233967,233967,233849,233849,233769,233769,233540,233540,233511,233511,233430,233430,233382,233382,233184,233184,233075,233075,233024,233024,232960,232960,232903,232903,232715,232715,232654,232654,232606,232606,232571,232571,232451,232451,232195,232195,232107,232107,232085,232085,231985,231985,231893,231893,231838,231838,231786,231786,231734,231734,231679,231679,231608,231608,231545,231545,231523,231523,231392,231392,231371,231371,231350,231350,231277,231277,231261,231261,231229,231229,231213,231213,231194,231194,231137,231137,231044,231044,230991,230991,230929,230929,230886,230886,230852,230852,230821,230821,230729,230729,230595,230595,230365,230365,230332,230332,230274,230274,230189,230189,230060,230060,229948,229948,229840,229840,229776,229776,229742,229742,229574,229574,229424,229424,229404,229404,229376,229376,229228,229228,228862,228862,228803,228803,228583,228583,228508,228508,228447,228447,228400,228400,228340,228340,228320,228320,228260,228260,228146,228146,228090,228090,227793,227793,227753,227753,227673,227673,227605,227605,227556,227556,227492,227492,227471,227471,227398,227398,227350,227350,227261,227261,227172,227172,227152,227152,227046,227046,226945,226945,226786,226786,226737,226737,226614,226614,226543,226543,226423,226423,226392,226392,226236,226236,226086,226086,226068,226068,225854,225854,225801,225801,225672,225672,225408,225408,225385,225385,225350,225350,225255,225255,225188,225188,225158,225158,225107,225107,225089,225089,224982,224982,224707,224707,224683,224683,224660,224660,224543,224543,224375,224375,224259,224259,224128,224128,224100,224100,224081,224081,224029,224029,223866,223866,223826,223826,223778,223778,223749,223749,223715,223715,223580,223580,223355,223355,223287,223287,223198,223198,223044,223044,223002,223002,222969,222969,222911,222911,222808,222808,222789,222789,222715,222715,222480,222480,222337,222337,222236,222236,222147,222147,222095,222095,222020,222020,221921,221921,221866,221866,221563,221563,221537,221537,221419,221419,221194,221194,221018,221018,220892,220892,220742,220742,220670,220670,220356,220356,220328,220328,220159,220159,220103,220103,220050,220050,220028,220028,219946,219946,219778,219778,219722,219722,219640,219640,219500,219500,219362,219362,219241,219241,219136,219136,219088,219088,218931,218931,218703,218703,218530,218530,218331,218331,218314,218314,218187,218187,218155,218155,218087,218087,218019,218019,217949,217949,217915,217915,217703,217703,217663,217663,217535,217535,217489,217489,217358,217358,217164,217164,217140,217140,217121,217121,217059,217059,216913,216913,216838,216838,216788,216788,216772,216772,216686,216686,216628,216628,216512,216512,216426,216426,216298,216298,216147,216147,216076,216076,216060,216060,216024,216024,215987,215987,215957,215957,215859,215859,215836,215836,215820,215820,215730,215730,215604,215604,215583,215583,215370,215370,215163,215163,215138,215138,214949,214949,214908,214908,214891,214891,214859,214859,214792,214792,214772,214772,214738,214738,214435,214435,214186,214186,214162,214162,214087,214087,213836,213836,213801,213801,213727,213727,213686,213686,213638,213638,213563,213563,213407,213407,213320,213320,213222,213222,213183,213183,213081,213081,212985,212985,212908,212908,212886,212886,212769,212769,212681,212681,212665,212665,212617,212617,212600,212600,212567,212567,212533,212533,212517,212517,212490,212490,212159,212159,212046,212046,211617,211617,211571,211571,211493,211493,211383,211383,211365,211365,211315,211315,211224,211224,211179,211179,211065,211065,211049,211049,210920,210920,210856,210856,210791,210791,210768,210768,210656,210656,210562,210562,210384,210384,210282,210282,210194,210194,210113,210113,209968,209968,209884,209884,209824,209824,209770,209770,209604,209604,209545,209545,209528,209528,209509,209509,209374,209374,209233,209233,209132,209132,209100,209100,208915,208915,208887,208887,208871,208871,208624,208624,208603,208603,208291,208291,208273,208273,208254,208254,208143,208143,208033,208033,207959,207959,207929,207929,207909,207909,207864,207864,207795,207795,207763,207763,207666,207666,207580,207580,207460,207460,207431,207431,207392,207392,207351,207351,207317,207317,207295,207295,207234,207234,207206,207206,207157,207157,207088,207088,206949,206949,206737,206737,206652,206652,206628,206628,206596,206596,206578,206578,206551,206551,206535,206535,206408,206408,206379,206379,206313,206313,206278,206278,206252,206252,206230,206230,206207,206207,206076,206076,205974,205974,205876,205876,205549,205549,205224,205224,205206,205206,205155,205155,205060,205060,205008,205008,204980,204980,204956,204956,204881,204881,204806,204806,204689,204689,204617,204617,204593,204593,204349,204349,204316,204316,204274,204274,204210,204210,204151,204151,204110,204110,204071,204071,204042,204042,203968,203968,203894,203894,203878,203878,203768,203768,203641,203641,203532,203532,203497,203497,203479,203479,203281,203281,203260,203260,203194,203194,203148,203148,203109,203109,202847,202847,202821,202821,202797,202797,202681,202681,202568,202568,202508,202508,202328,202328,202236,202236,202143,202143,202123,202123,202063,202063,202027,202027,201924,201924,201893,201893,201874,201874,201823,201823,201739,201739,201679,201679,201594,201594,201568,201568,201443,201443,201303,201303,201264,201264,201193,201193,201155,201155,200967,200967,200884,200884,200823,200823,200797,200797,200697,200697,200668,200668,200637,200637,200604,200604,200343,200343,200301,200301,200272,200272,200240,200240,200187,200187,200073,200073,199995,199995,199942,199942,199900,199900,199757,199757,199738,199738,199686,199686,199667,199667,199591,199591,199469,199469,199379,199379,199335,199335,199219,199219,199201,199201,199153,199153,199109,199109,199029,199029,198886,198886,198844,198844,198786,198786,198767,198767,198510,198510,198452,198452,198434,198434,198401,198401,198377,198377,198348,198348,198320,198320,198257,198257,198211,198211,198173,198173,198143,198143,198044,198044,198010,198010,197876,197876,197712,197712,197691,197691,197595,197595,197555,197555,197505,197505,197451,197451,197276,197276,197149,197149,197056,197056,196776,196776,196757,196757,196730,196730,196472,196472,196332,196332,196286,196286,196240,196240,196085,196085,196049,196049,195996,195996,195910,195910,195817,195817,195715,195715,195687,195687,195555,195555,195422,195422,195373,195373,195341,195341,195316,195316,195226,195226,195194,195194,195117,195117,195094,195094,195015,195015,194978,194978,194962,194962,194867,194867,194825,194825,194752,194752,194714,194714,194600,194600,194584,194584,194496,194496,194329,194329,194260,194260,194196,194196,194136,194136,194081,194081,194030,194030,193977,193977,193910,193910,193872,193872,193803,193803,193768,193768,193734,193734,193668,193668,193637,193637,193620,193620,193541,193541,193467,193467,193408,193408,193374,193374,193308,193308,193212,193212,193196,193196,193159,193159,193104,193104,193034,193034,193014,193014,192992,192992,192920,192920,192811,192811,192708,192708,192674,192674,192653,192653,192480,192480,192393,192393,192375,192375,192345,192345,192323,192323,192255,192255,192137,192137,192112,192112,192071,192071,191945,191945,191889,191889,191828,191828,191593,191593,191491,191491,191442,191442,191288,191288,191227,191227,191196,191196,191157,191157,191134,191134,191112,191112,191061,191061,191014,191014,190986,190986,190891,190891,190871,190871,190817,190817,190750,190750,190530,190530,190246,190246,190200,190200,190152,190152,190040,190040,190019,190019,189966,189966,189933,189933,189886,189886,189778,189778,189741,189741,189676,189676,189630,189630,189544,189544,189356,189356,189224,189224,189180,189180,189119,189119,189050,189050,188987,188987,188968,188968,188929,188929,188902,188902,188798,188798,188770,188770,188739,188739,188646,188646,188622,188622,188583,188583,188554,188554,188500,188500,188382,188382,188271,188271,188108,188108,187924,187924,187883,187883,187844,187844,187753,187753,187635,187635,187565,187565,187544,187544,187489,187489,187325,187325,187154,187154,187127,187127,187111,187111,187087,187087,187049,187049,186960,186960,186942,186942,186899,186899,186863,186863,186837,186837,186816,186816,186781,186781,186749,186749,186677,186677,186392,186392,186290,186290,186196,186196,186060,186060,186012,186012,185915,185915,185857,185857,185795,185795,185758,185758,185702,185702,185642,185642,185512,185512,185377,185377,185328,185328,185197,185197,185083,185083,185043,185043,185002,185002,184950,184950,184919,184919,184900,184900,184866,184866,184837,184837,184747,184747,184675,184675,184639,184639,184623,184623,184581,184581,184548,184548,184513,184513,184331,184331,184276,184276,184251,184251,184220,184220,184157,184157,184129,184129,184102,184102,184017,184017,183917,183917,183847,183847,183821,183821,183802,183802,183702,183702,183597,183597,183579,183579,183521,183521,183495,183495,183443,183443,183348,183348,183326,183326,183249,183249,183172,183172,183124,183124,183031,183031,183011,183011,182924,182924,182865,182865,182740,182740,182705,182705,182689,182689,182657,182657,182519,182519,182475,182475,182415,182415,182370,182370,182227,182227,182181,182181,182128,182128,182085,182085,182045,182045,182023,182023,181838,181838,181593,181593,181536,181536,181499,181499,181396,181396,181334,181334,181311,181311,181233,181233,181194,181194,181153,181153,181107,181107,181051,181051,181034,181034,180899,180899,180596,180596,180496,180496,180480,180480,180458,180458,180360,180360,180303,180303,180136,180136,180107,180107,180082,180082,180026,180026,179986,179986,179832,179832,179796,179796,179748,179748,179696,179696,179671,179671,179570,179570,179466,179466,179426,179426,179395,179395,179368,179368,179299,179299,179267,179267,179140,179140,179101,179101,179033,179033,178970,178970,178870,178870,178850,178850,178643,178643,178601,178601,178561,178561,178541,178541,178472,178472,178357,178357,178339,178339,178313,178313,178222,178222,178199,178199,178145,178145,178032,178032,178016,178016,177979,177979,177956,177956,177935,177935,177881,177881,177828,177828,177781,177781,177758,177758,177739,177739,177688,177688,177426,177426,177351,177351,177234,177234,177167,177167,177051,177051,177029,177029,176997,176997,176975,176975,176955,176955,176846,176846,176823,176823,176797,176797,176739,176739,176593,176593,176456,176456,176434,176434,176386,176386,176258,176258,176096,176096,176031,176031,175957,175957,175909,175909,175884,175884,175847,175847,175507,175507,175312,175312,175233,175233,175195,175195,175168,175168,175073,175073,174831,174831,174755,174755,174681,174681,174637,174637,174592,174592,174571,174571,174550,174550,174529,174529,174469,174469,174374,174374,174321,174321,174301,174301,174210,174210,174153,174153,174120,174120,174037,174037,173969,173969,173922,173922,173896,173896,173855,173855,173773,173773,173739,173739,173663,173663,173611,173611,173595,173595,173564,173564,173513,173513,173474,173474,173440,173440,173323,173323,173251,173251,173215,173215,173148,173148,173056,173056,173009,173009,172987,172987,172925,172925,172853,172853,172825,172825,172591,172591,172532,172532,172494,172494,172412,172412,172303,172303,172177,172177,172103,172103,172047,172047,171855,171855,171813,171813,171791,171791,171759,171759,171736,171736,171705,171705,171687,171687,171661,171661,171548,171548,171523,171523,171471,171471,171412,171412,171395,171395,171288,171288,171269,171269,171244,171244,171155,171155,171005,171005,170956,170956,170863,170863,170845,170845,170827,170827,170803,170803,170741,170741,170640,170640,170617,170617,170597,170597,170530,170530,170505,170505,170354,170354,170255,170255,170124,170124,170052,170052,170003,170003,169887,169887,169849,169849,169831,169831,169786,169786,169720,169720,169564,169564,169492,169492,169384,169384,169311,169311,169215,169215,169156,169156,169086,169086,169011,169011,168993,168993,168943,168943,168912,168912,168869,168869,168840,168840,168792,168792,168769,168769,168691,168691,168656,168656,168598,168598,168577,168577,168543,168543,168443,168443,168412,168412,168377,168377,168313,168313,168204,168204,168091,168091,168069,168069,167924,167924,167705,167705,167676,167676,167648,167648,167542,167542,167495,167495,167456,167456,167321,167321,167300,167300,167240,167240,167098,167098,167047,167047,167020,167020,166882,166882,166843,166843,166688,166688,166658,166658,166614,166614,166538,166538,166472,166472,166343,166343,166249,166249,166198,166198,166179,166179,166145,166145,166063,166063,165998,165998,165955,165955,165837,165837,165815,165815,165603,165603,165574,165574,165527,165527,165403,165403,165382,165382,165347,165347,165329,165329,165295,165295,165279,165279,165259,165259,165136,165136,165098,165098,164965,164965,164926,164926,164882,164882,164838,164838,164812,164812,164778,164778,164690,164690,164648,164648,164580,164580,164525,164525,164428,164428,164383,164383,164287,164287,164250,164250,164226,164226,164159,164159,164120,164120,163987,163987,163965,163965,163922,163922,163889,163889,163867,163867,163752,163752,163660,163660,163638,163638,163616,163616,163589,163589,163496,163496,163336,163336,163314,163314,163053,163053,163005,163005,162859,162859,162837,162837,162790,162790,162727,162727,162693,162693,162649,162649,162568,162568,162532,162532,162477,162477,162411,162411,162308,162308,162288,162288,162253,162253,162227,162227,162195,162195,162125,162125,161995,161995,161974,161974,161909,161909,161886,161886,161819,161819,161655,161655,161566,161566,161522,161522,161467,161467,161446,161446,161351,161351,161327,161327,161275,161275,161197,161197,161166,161166,161130,161130,161018,161018,160999,160999,160912,160912,160805,160805,160777,160777,160689,160689,160617,160617,160536,160536,160500,160500,160436,160436,160415,160415,160355,160355,160301,160301,160256,160256,160229,160229,160207,160207,160187,160187,160163,160163,160089,160089,160031,160031,159893,159893,159844,159844,159766,159766,159728,159728,159703,159703,159674,159674,159624,159624,159567,159567,159406,159406,159380,159380,159355,159355,159262,159262,159206,159206,159131,159131,159035,159035,158814,158814,158798,158798,158762,158762,158745,158745,158681,158681,158622,158622,158482,158482,158463,158463,158418,158418,158387,158387,158371,158371,158267,158267,158235,158235,158211,158211,158129,158129,158084,158084,157979,157979,157927,157927,157873,157873,157848,157848,157814,157814,157775,157775,157615,157615,157517,157517,157482,157482,157333,157333,157265,157265,157237,157237,157117,157117,157078,157078,157012,157012,156949,156949,156931,156931,156854,156854,156773,156773,156746,156746,156558,156558,156538,156538,156512,156512,156433,156433,156392,156392,156371,156371,156320,156320,156279,156279,156236,156236,156134,156134,156038,156038,156017,156017,155885,155885,155867,155867,155770,155770,155692,155692,155585,155585,155551,155551,155486,155486,155435,155435,155361,155361,155336,155336,155288,155288,155188,155188,155165,155165,155069,155069,154989,154989,154969,154969,154899,154899,154865,154865,154810,154810,154762,154762,154698,154698,154665,154665,154638,154638,154609,154609,154556,154556,154534,154534,154507,154507,154460,154460,154422,154422,154382,154382,154366,154366,154302,154302,154234,154234,154202,154202,154170,154170,154136,154136,154107,154107,154065,154065,153932,153932,153815,153815,153740,153740,153708,153708,153687,153687,153617,153617,153573,153573,153466,153466,153400,153400,153325,153325,153272,153272,153228,153228,153191,153191,153169,153169,153097,153097,152944,152944,152926,152926,152904,152904,152748,152748,152708,152708,152653,152653,152610,152610,152579,152579,152537,152537,152510,152510,152453,152453,152370,152370,152341,152341,152323,152323,152258,152258,152216,152216,152194,152194,152061,152061,152002,152002,151953,151953,151870,151870,151834,151834,151754,151754,151735,151735,151660,151660,151632,151632,151476,151476,151438,151438,151395,151395,151368,151368,151347,151347,151296,151296,151275,151275,151254,151254,151129,151129,151091,151091,151056,151056,151021,151021,150979,150979,150893,150893,150838,150838,150808,150808,150751,150751,150687,150687,150568,150568,150490,150490,150446,150446,150396,150396,150356,150356,150326,150326,150303,150303,150274,150274,150253,150253,150209,150209,150151,150151,149990,149990,149853,149853,149823,149823,149803,149803,149782,149782,149738,149738,149691,149691,149665,149665,149572,149572,149524,149524,149429,149429,149344,149344,149291,149291,149170,149170,149153,149153,149084,149084,149035,149035,148942,148942,148868,148868,148730,148730,148708,148708,148658,148658,148592,148592,148498,148498,148482,148482,148457,148457,148421,148421,148382,148382,148348,148348,148313,148313,148293,148293,148276,148276,148136,148136,148003,148003,147924,147924,147906,147906,147833,147833,147707,147707,147645,147645,147596,147596,147576,147576,147486,147486,147439,147439,147361,147361,147325,147325,147292,147292,147270,147270,147254,147254,147237,147237,147122,147122,147084,147084,147027,147027,146990,146990,146941,146941,146811,146811,146771,146771,146718,146718,146700,146700,146658,146658,146557,146557,146540,146540,146449,146449,146422,146422,146386,146386,146364,146364,146345,146345,146300,146300,146280,146280,146169,146169,146153,146153,146122,146122,146104,146104,145983,145983,145893,145893,145870,145870,145792,145792,145759,145759,145705,145705,145685,145685,145660,145660,145622,145622,145493,145493,145365,145365,145266,145266,145249,145249,145198,145198,145139,145139,145076,145076,145052,145052,145016,145016,144997,144997,144863,144863,144764,144764,144722,144722,144688,144688,144619,144619,144529,144529,144506,144506,144479,144479,144400,144400,144373,144373,144312,144312,144261,144261,144178,144178,144142,144142,144101,144101,144069,144069,143881,143881,143797,143797,143705,143705,143680,143680,143617,143617,143583,143583,143513,143513,143475,143475,143317,143317,143282,143282,143253,143253,143226,143226,143210,143210,143142,143142,143091,143091,143063,143063,142921,142921,142840,142840,142781,142781,142761,142761,142738,142738,142685,142685,142669,142669,142605,142605,142482,142482,142458,142458,142408,142408,142365,142365,142334,142334,142313,142313,142189,142189,142152,142152,142100,142100,142030,142030,141960,141960,141917,141917,141868,141868,141811,141811,141744,141744,141596,141596,141537,141537,141456,141456,141412,141412,141348,141348,141301,141301,141270,141270,141243,141243,141184,141184,141141,141141,141116,141116,141100,141100,141073,141073,141015,141015,140973,140973,140848,140848,140807,140807,140748,140748,140720,140720,140692,140692,140640,140640,140614,140614,140583,140583,140565,140565,140481,140481,140458,140458,140401,140401,140355,140355,140324,140324,140302,140302,140245,140245,140166,140166,140145,140145,140128,140128,140096,140096,140023,140023,139889,139889,139855,139855,139781,139781,139751,139751,139720,139720,139669,139669,139638,139638,139608,139608,139555,139555,139493,139493,139431,139431,139363,139363,139347,139347,139303,139303,139280,139280,139258,139258,139223,139223,139125,139125,139073,139073,139045,139045,139016,139016,138977,138977,138782,138782,138718,138718,138655,138655,138616,138616,138516,138516,138396,138396,138371,138371,138322,138322,138281,138281,138258,138258,138220,138220,138159,138159,138120,138120,138061,138061,138022,138022,138005,138005,137981,137981,137949,137949,137930,137930,137874,137874,137853,137853,137758,137758,137709,137709,137675,137675,137643,137643,137601,137601,137580,137580,137542,137542,137514,137514,137494,137494,137458,137458,137426,137426,137409,137409,137345,137345,137323,137323,137296,137296,137220,137220,137191,137191,137158,137158,137137,137137,137025,137025,137006,137006,136969,136969,136909,136909,136854,136854,136785,136785,136744,136744,136712,136712,136694,136694,136644,136644,136392,136392,136331,136331,136262,136262,136183,136183,136114,136114,136043,136043,136017,136017,135916,135916,135899,135899,135880,135880,135830,135830,135661,135661,135593,135593,135532,135532,135497,135497,135457,135457,135424,135424,135329,135329,135308,135308,135269,135269,135218,135218,135170,135170,135038,135038,135022,135022,134916,134916,134884,134884,134861,134861,134785,134785,134716,134716,134620,134620,134583,134583,134547,134547,134491,134491,134461,134461,134432,134432,134370,134370,134315,134315,134273,134273,134165,134165,134102,134102,134079,134079,134047,134047,134031,134031,133946,133946,133827,133827,133805,133805,133768,133768,133689,133689,133667,133667,133651,133651,133606,133606,133577,133577,133534,133534,133507,133507,133436,133436,133402,133402,133359,133359,133343,133343,133298,133298,133259,133259,133142,133142,133118,133118,133060,133060,133032,133032,133015,133015,132991,132991,132962,132962,132935,132935,132909,132909,132859,132859,132839,132839,132785,132785,132764,132764,132742,132742,132706,132706,132660,132660,132535,132535,132471,132471,132397,132397,132372,132372,132337,132337,132306,132306,132264,132264,132226,132226,132129,132129,132055,132055,132029,132029,131937,131937,131912,131912,131850,131850,131811,131811,131785,131785,131701,131701,131655,131655,131607,131607,131544,131544,131443,131443,131376,131376,131333,131333,131310,131310,131281,131281,131246,131246,131218,131218,131201,131201,131183,131183,130997,130997,130935,130935,130916,130916,130880,130880,130849,130849,130789,130789,130773,130773,130750,130750,130688,130688,130598,130598,130545,130545,130482,130482,130429,130429,130321,130321,130276,130276,130215,130215,130172,130172,130153,130153,130132,130132,130112,130112,130095,130095,130071,130071,130047,130047,129992,129992,129942,129942,129891,129891,129816,129816,129795,129795,129728,129728,129699,129699,129683,129683,129653,129653,129604,129604,129532,129532,129444,129444,129333,129333,129239,129239,129201,129201,129174,129174,129119,129119,128964,128964,128925,128925,128899,128899,128764,128764,128687,128687,128666,128666,128586,128586,128511,128511,128487,128487,128459,128459,128442,128442,128402,128402,128308,128308,128224,128224,128159,128159,128101,128101,128077,128077,128025,128025,127998,127998,127960,127960,127906,127906,127886,127886,127816,127816,127794,127794,127712,127712,127683,127683,127631,127631,127598,127598,127581,127581,127494,127494,127453,127453,127408,127408,127375,127375,127314,127314,127282,127282,127246,127246,127225,127225,127177,127177,127158,127158,127121,127121,127017,127017,126933,126933,126910,126910,126848,126848,126817,126817,126753,126753,126673,126673,126648,126648,126610,126610,126575,126575,126552,126552,126500,126500,126444,126444,126421,126421,126329,126329,126308,126308,126291,126291,126229,126229,126211,126211,126047,126047,126014,126014,125987,125987,125924,125924,125887,125887,125848,125848,125819,125819,125774,125774,125682,125682,125650,125650,125619,125619,125515,125515,125497,125497,125384,125384,125352,125352,125249,125249,125231,125231,125180,125180,125154,125154,125104,125104,125001,125001,124950,124950,124922,124922,124904,124904,124875,124875,124843,124843,124815,124815,124746,124746,124709,124709,124638,124638,124611,124611,124566,124566,124495,124495,124462,124462,124420,124420,124395,124395,124347,124347,124293,124293,124277,124277,124061,124061,124016,124016,123991,123991,123962,123962,123906,123906,123862,123862,123822,123822,123788,123788,123720,123720,123643,123643,123622,123622,123487,123487,123449,123449,123433,123433,123398,123398,123347,123347,123312,123312,123267,123267,123189,123189,123172,123172,123109,123109,123074,123074,123049,123049,123014,123014,122973,122973,122908,122908,122833,122833,122815,122815,122764,122764,122638,122638,122602,122602,122546,122546,122523,122523,122477,122477,122456,122456,122437,122437,122375,122375,122347,122347,122298,122298,122281,122281,122253,122253,122208,122208,122173,122173,122110,122110,122067,122067,121988,121988,121967,121967,121925,121925,121881,121881,121847,121847,121825,121825,121809,121809,121696,121696,121628,121628,121611,121611,121510,121510,121492,121492,121452,121452,121426,121426,121398,121398,121330,121330,121303,121303,121262,121262,121209,121209,121144,121144,121095,121095,121072,121072,121049,121049,120949,120949,120835,120835,120809,120809,120770,120770,120734,120734,120683,120683,120651,120651,120615,120615,120574,120574,120556,120556,120506,120506,120378,120378,120237,120237,120191,120191,120171,120171,120152,120152,120124,120124,120033,120033,120004,120004,119986,119986,119924,119924,119908,119908,119869,119869,119852,119852,119782,119782,119757,119757,119714,119714,119690,119690,119629,119629,119592,119592,119560,119560,119544,119544,119510,119510,119435,119435,119402,119402,119382,119382,119359,119359,119317,119317,119283,119283,119266,119266,119235,119235,119211,119211,119184,119184,119147,119147,119115,119115,119099,119099,119061,119061,119014,119014,118993,118993,118953,118953,118927,118927,118891,118891,118834,118834,118785,118785,118718,118718,118662,118662,118642,118642,118621,118621,118579,118579,118520,118520,118495,118495,118471,118471,118453,118453,118363,118363,118313,118313,118285,118285,118246,118246,118196,118196,118147,118147,118117,118117,118091,118091,118072,118072,117972,117972,117955,117955,117909,117909,117834,117834,117808,117808,117790,117790,117754,117754,117656,117656,117623,117623,117607,117607,117555,117555,117507,117507,117484,117484,117450,117450,117432,117432,117414,117414,117395,117395,117301,117301,117252,117252,117227,117227,117176,117176,117126,117126,117105,117105,117085,117085,117060,117060,117040,117040,117016,117016,116959,116959,116886,116886,116807,116807,116753,116753,116726,116726,116678,116678,116546,116546,116515,116515,116479,116479,116461,116461,116417,116417,116336,116336,116268,116268,116237,116237,116215,116215,116153,116153,116119,116119,116067,116067,116050,116050,116030,116030,115970,115970,115925,115925,115864,115864,115832,115832,115780,115780,115757,115757,115732,115732,115691,115691,115533,115533,115454,115454,115435,115435,115413,115413,115394,115394,115350,115350,115332,115332,115301,115301,115277,115277,115248,115248,115222,115222,115198,115198,115180,115180,115146,115146,115087,115087,115023,115023,115005,115005,114985,114985,114961,114961,114937,114937,114880,114880,114863,114863,114822,114822,114803,114803,114774,114774,114718,114718,114683,114683,114648,114648,114623,114623,114551,114551,114357,114357,114339,114339,114303,114303,114278,114278,114231,114231,114207,114207,114156,114156,114123,114123,114087,114087,114065,114065,114039,114039,114005,114005,113952,113952,113931,113931,113897,113897,113880,113880,113844,113844,113816,113816,113788,113788,113765,113765,113734,113734,113700,113700,113651,113651,113607,113607,113569,113569,113493,113493,113465,113465,113414,113414,113345,113345,113324,113324,113241,113241,113225,113225,113125,113125,113108,113108,113092,113092,113070,113070,113053,113053,113025,113025,112985,112985,112938,112938,112921,112921,112877,112877,112852,112852,112806,112806,112774,112774,112746,112746,112719,112719,112701,112701,112683,112683,112642,112642,112565,112565,112510,112510,112475,112475,112402,112402,112384,112384,112358,112358,112305,112305,112288,112288,112254,112254,112224,112224,112180,112180,112124,112124,112107,112107,112089,112089,112000,112000,111984,111984,111926,111926,111898,111898,111842,111842,111819,111819,111782,111782,111764,111764,111724,111724,111707,111707,111613,111613,111563,111563,111544,111544,111424,111424,111356,111356,111339,111339,111322,111322,111284,111284,111260,111260,111218,111218,111158,111158,111127,111127,111105,111105,111043,111043,111017,111017,110996,110996,110943,110943,110879,110879,110854,110854,110783,110783,110749,110749,110683,110683,110648,110648,110621,110621,110494,110494,110475,110475,110447,110447,110328,110328,110299,110299,110275,110275,110235,110235,110200,110200,110104,110104,110082,110082,110025,110025,109995,109995,109972,109972,109954,109954,109908,109908,109880,109880,109861,109861,109811,109811,109741,109741,109702,109702,109664,109664,109639,109639,109622,109622,109579,109579,109498,109498,109423,109423,109407,109407,109378,109378,109344,109344,109304,109304,109238,109238,109222,109222,109194,109194,109175,109175,109124,109124,109090,109090,109045,109045,109028,109028,109000,109000,108975,108975,108954,108954,108931,108931,108882,108882,108790,108790,108753,108753,108717,108717,108701,108701,108596,108596,108555,108555,108425,108425,108396,108396,108364,108364,108345,108345,108324,108324,108282,108282,108252,108252,108189,108189,108172,108172,108153,108153,108073,108073,108053,108053,108037,108037,108017,108017,107999,107999,107965,107965,107943,107943,107927,107927,107899,107899,107858,107858,107825,107825,107734,107734,107715,107715,107678,107678,107615,107615,107584,107584,107530,107530,107497,107497,107471,107471,107423,107423,107401,107401,107380,107380,107348,107348,107295,107295,107233,107233,107161,107161,107140,107140,107115,107115,107086,107086,107058,107058,107037,107037,107016,107016,106999,106999,106934,106934,106760,106760,106697,106697,106674,106674,106614,106614,106589,106589,106494,106494,106454,106454,106409,106409,106340,106340,106267,106267,106225,106225,106124,106124,105997,105997,105955,105955,105925,105925,105859,105859,105816,105816,105785,105785,105740,105740,105686,105686,105655,105655,105616,105616,105593,105593,105566,105566,105538,105538,105438,105438,105408,105408,105382,105382,105300,105300,105258,105258,105174,105174,105147,105147,105075,105075,105030,105030,105009,105009,104972,104972,104934,104934,104896,104896,104877,104877,104859,104859,104830,104830,104806,104806,104699,104699,104667,104667,104587,104587,104560,104560,104514,104514,104487,104487,104431,104431,104408,104408,104377,104377,104351,104351,104319,104319,104256,104256,104197,104197,104146,104146,104130,104130,104097,104097,104035,104035,104010,104010,103991,103991,103971,103971,103948,103948,103900,103900,103868,103868,103829,103829,103794,103794,103764,103764,103735,103735,103718,103718,103700,103700,103681,103681,103628,103628,103606,103606,103575,103575,103553,103553,103521,103521,103489,103489,103467,103467,103433,103433,103409,103409,103376,103376,103299,103299,103257,103257,103218,103218,103193,103193,103166,103166,103143,103143,103114,103114,103090,103090,103034,103034,103012,103012,102953,102953,102925,102925,102863,102863,102834,102834,102812,102812,102794,102794,102767,102767,102718,102718,102636,102636,102619,102619,102573,102573,102547,102547,102529,102529,102511,102511,102475,102475,102448,102448,102424,102424,102388,102388,102323,102323,102169,102169,102148,102148,102131,102131,102099,102099,102082,102082,102065,102065,102048,102048,101931,101931,101885,101885,101852,101852,101826,101826,101800,101800,101693,101693,101667,101667,101648,101648,101593,101593,101556,101556,101523,101523,101501,101501,101474,101474,101438,101438,101413,101413,101374,101374,101337,101337,101271,101271,101244,101244,101209,101209,101189,101189,101164,101164,101132,101132,101030,101030,100982,100982,100955,100955,100928,100928,100893,100893,100864,100864,100839,100839,100815,100815,100778,100778,100726,100726,100686,100686,100644,100644,100606,100606,100562,100562,100450,100450,100426,100426,100302,100302,100255,100255,100236,100236,100158,100158,100122,100122,100050,100050,100016,100016,100000,100000,99963,99963,99923,99923,99876,99876,99843,99843,99781,99781,99761,99761,99745,99745,99726,99726,99702,99702,99660,99660,99630,99630,99587,99587,99521,99521,99457,99457,99438,99438,99397,99397,99293,99293,99271,99271,99198,99198,99180,99180,99052,99052,99036,99036,99010,99010,98976,98976,98844,98844,98793,98793,98769,98769,98716,98716,98655,98655,98615,98615,98565,98565,98545,98545,98511,98511,98490,98490,98446,98446,98402,98402,98378,98378,98356,98356,98325,98325,98285,98285,98255,98255,98229,98229,98185,98185,98147,98147,98045,98045,98000,98000,97941,97941,97828,97828,97805,97805,97776,97776,97744,97744,97696,97696,97679,97679,97617,97617,97598,97598,97551,97551,97514,97514,97489,97489,97472,97472,97456,97456,97381,97381,97354,97354,97301,97301,97272,97272,97210,97210,97183,97183,97149,97149,97126,97126,97103,97103,97077,97077,97059,97059,97013,97013,96955,96955,96917,96917,96849,96849,96744,96744,96725,96725,96688,96688,96621,96621,96569,96569,96506,96506,96478,96478,96451,96451,96401,96401,96336,96336,96315,96315,96285,96285,96252,96252,96218,96218,96160,96160,96110,96110,96042,96042,95989,95989,95973,95973,95942,95942,95849,95849,95829,95829,95750,95750,95728,95728,95707,95707,95689,95689,95576,95576,95478,95478,95432,95432,95404,95404,95382,95382,95363,95363,95294,95294,95278,95278,95240,95240,95196,95196,95170,95170,95106,95106,95078,95078,94987,94987,94944,94944,94894,94894,94862,94862,94843,94843,94824,94824,94808,94808,94761,94761,94730,94730,94706,94706,94678,94678,94618,94618,94599,94599,94573,94573,94519,94519,94485,94485,94450,94450,94362,94362,94325,94325,94306,94306,94288,94288,94267,94267,94241,94241,94218,94218,94138,94138,94090,94090,93954,93954,93936,93936,93898,93898,93861,93861,93831,93831,93717,93717,93694,93694,93664,93664,93625,93625,93580,93580,93518,93518,93460,93460,93444,93444,93424,93424,93396,93396,93380,93380,93361,93361,93234,93234,93191,93191,93148,93148,93048,93048,92992,92992,92971,92971,92954,92954,92866,92866,92838,92838,92776,92776,92739,92739,92712,92712,92695,92695,92674,92674,92635,92635,92613,92613,92588,92588,92529,92529,92502,92502,92421,92421,92386,92386,92323,92323,92296,92296,92276,92276,92220,92220,92182,92182,92096,92096,92080,92080,92030,92030,91985,91985,91954,91954,91934,91934,91917,91917,91879,91879,91839,91839,91799,91799,91783,91783,91766,91766,91743,91743,91713,91713,91672,91672,91636,91636,91620,91620,91578,91578,91560,91560,91521,91521,91499,91499,91478,91478,91442,91442,91424,91424,91397,91397,91366,91366,91348,91348,91330,91330,91309,91309,91279,91279,91222,91222,91136,91136,91052,91052,91025,91025,90999,90999,90957,90957,90933,90933,90910,90910,90862,90862,90831,90831,90814,90814,90774,90774,90731,90731,90663,90663,90610,90610,90557,90557,90521,90521,90502,90502,90412,90412,90376,90376,90333,90333,90311,90311,90246,90246,90218,90218,90180,90180,90142,90142,90109,90109,90089,90089,90061,90061,90036,90036,90004,90004,89980,89980,89913,89913,89851,89851,89832,89832,89792,89792,89762,89762,89725,89725,89703,89703,89685,89685,89654,89654,89626,89626,89543,89543,89515,89515,89464,89464,89429,89429,89348,89348,89331,89331,89295,89295,89243,89243,89162,89162,89133,89133,89105,89105,89079,89079,89059,89059,89036,89036,88995,88995,88929,88929,88909,88909,88893,88893,88856,88856,88810,88810,88785,88785,88714,88714,88688,88688,88661,88661,88642,88642,88610,88610,88584,88584,88549,88549,88514,88514,88476,88476,88460,88460,88425,88425,88403,88403,88387,88387,88360,88360,88320,88320,88299,88299,88280,88280,88235,88235,88184,88184,88149,88149,88129,88129,88087,88087,88035,88035,88014,88014,87969,87969,87929,87929,87910,87910,87893,87893,87877,87877,87792,87792,87752,87752,87730,87730,87713,87713,87681,87681,87635,87635,87618,87618,87594,87594,87561,87561,87545,87545,87519,87519,87480,87480,87452,87452,87431,87431,87415,87415,87341,87341,87307,87307,87288,87288,87265,87265,87218,87218,87202,87202,87140,87140,87109,87109,87056,87056,87023,87023,87004,87004,86983,86983,86938,86938,86922,86922,86903,86903,86884,86884,86831,86831,86806,86806,86750,86750,86696,86696,86665,86665,86581,86581,86564,86564,86529,86529,86472,86472,86439,86439,86415,86415,86381,86381,86308,86308,86289,86289,86222,86222,86200,86200,86147,86147,86077,86077,86022,86022,85980,85980,85952,85952,85882,85882,85832,85832,85778,85778,85745,85745,85715,85715,85694,85694,85675,85675,85655,85655,85629,85629,85555,85555,85512,85512,85494,85494,85461,85461,85440,85440,85418,85418,85384,85384,85275,85275,85258,85258,85219,85219,85203,85203,85156,85156,85121,85121,85102,85102,85057,85057,85018,85018,84927,84927,84875,84875,84840,84840,84743,84743,84701,84701,84678,84678,84656,84656,84590,84590,84568,84568,84515,84515,84465,84465,84384,84384,84322,84322,84288,84288,84192,84192,84148,84148,84129,84129,84099,84099,84079,84079,84050,84050,83984,83984,83928,83928,83910,83910,83856,83856,83815,83815,83795,83795,83775,83775,83758,83758,83679,83679,83659,83659,83620,83620,83590,83590,83545,83545,83512,83512,83479,83479,83448,83448,83360,83360,83333,83333,83295,83295,83268,83268,83153,83153,83120,83120,83081,83081,83063,83063,83029,83029,83008,83008,82975,82975,82959,82959,82922,82922,82891,82891,82869,82869,82799,82799,82771,82771,82742,82742,82679,82679,82615,82615,82594,82594,82576,82576,82528,82528,82486,82486,82455,82455,82435,82435,82413,82413,82385,82385,82326,82326,82287,82287,82261,82261,82238,82238,82164,82164,82148,82148,82124,82124,82084,82084,82068,82068,82039,82039,82015,82015,81978,81978,81932,81932,81907,81907,81848,81848,81830,81830,81788,81788,81752,81752,81736,81736,81714,81714,81695,81695,81673,81673,81616,81616,81600,81600,81547,81547,81501,81501,81478,81478,81457,81457,81404,81404,81381,81381,81343,81343,81321,81321,81291,81291,81266,81266,81246,81246,81228,81228,81208,81208,81139,81139,81121,81121,81100,81100,81073,81073,81043,81043,81002,81002,80985,80985,80952,80952,80917,80917,80881,80881,80853,80853,80784,80784,80768,80768,80736,80736,80709,80709,80685,80685,80667,80667,80572,80572,80545,80545,80471,80471,80454,80454,80403,80403,80385,80385,80338,80338,80282,80282,80228,80228,80198,80198,80173,80173,80139,80139,80106,80106,80029,80029,79977,79977,79950,79950,79805,79805,79674,79674,79657,79657,79570,79570,79538,79538,79509,79509,79474,79474,79438,79438,79401,79401,79381,79381,79356,79356,79321,79321,79269,79269,79253,79253,79193,79193,79176,79176,79154,79154,79133,79133,79110,79110,79053,79053,79027,79027,78912,78912,78887,78887,78847,78847,78825,78825,78804,78804,78782,78782,78744,78744,78703,78703,78659,78659,78556,78556,78531,78531,78505,78505,78486,78486,78470,78470,78431,78431,78329,78329,78307,78307,78284,78284,78253,78253,78230,78230,78214,78214,78195,78195,78096,78096,78044,78044,78012,78012,77953,77953,77920,77920,77898,77898,77878,77878,77845,77845,77735,77735,77710,77710,77691,77691,77636,77636,77578,77578,77531,77531,77490,77490,77460,77460,77384,77384,77338,77338,77285,77285,77267,77267,77239,77239,77172,77172,77127,77127,77107,77107,77083,77083,77065,77065,77038,77038,77013,77013,76950,76950,76891,76891,76867,76867,76824,76824,76776,76776,76712,76712,76648,76648,76607,76607,76583,76583,76548,76548,76531,76531,76505,76505,76437,76437,76398,76398,76379,76379,76347,76347,76310,76310,76282,76282,76242,76242,76193,76193,76165,76165,76138,76138,76121,76121,76073,76073,76047,76047,76004,76004,75981,75981,75935,75935,75915,75915,75895,75895,75867,75867,75846,75846,75830,75830,75814,75814,75782,75782,75761,75761,75701,75701,75637,75637,75619,75619,75552,75552,75514,75514,75493,75493,75475,75475,75449,75449,75390,75390,75374,75374,75294,75294,75262,75262,75191,75191,75150,75150,75113,75113,75079,75079,75035,75035,74994,74994,74975,74975,74940,74940,74908,74908,74853,74853,74816,74816,74793,74793,74769,74769,74668,74668,74634,74634,74591,74591,74521,74521,74491,74491,74469,74469,74449,74449,74412,74412,74388,74388,74361,74361,74338,74338,74289,74289,74264,74264,74184,74184,74144,74144,74121,74121,74097,74097,74079,74079,74039,74039,74021,74021,73997,73997,73977,73977,73916,73916,73889,73889,73831,73831,73802,73802,73760,73760,73676,73676,73658,73658,73622,73622,73606,73606,73582,73582,73514,73514,73489,73489,73466,73466,73419,73419,73403,73403,73381,73381,73271,73271,73161,73161,73091,73091,73049,73049,73011,73011,72973,72973,72845,72845,72807,72807,72778,72778,72681,72681,72649,72649,72631,72631,72609,72609,72580,72580,72547,72547,72522,72522,72483,72483,72457,72457,72418,72418,72386,72386,72322,72322,72281,72281,72251,72251,72223,72223,72177,72177,72135,72135,72101,72101,72076,72076,72037,72037,72017,72017,71970,71970,71926,71926,71910,71910,71877,71877,71843,71843,71811,71811,71782,71782,71709,71709,71662,71662,71630,71630,71604,71604,71559,71559,71542,71542,71496,71496,71471,71471,71453,71453,71415,71415,71370,71370,71354,71354,71299,71299,71283,71283,71245,71245,71226,71226,71197,71197,71178,71178,71128,71128,71112,71112,71092,71092,71076,71076,71059,71059,71042,71042,71025,71025,70963,70963,70938,70938,70881,70881,70841,70841,70804,70804,70743,70743,70674,70674,70649,70649,70572,70572,70548,70548,70523,70523,70507,70507,70482,70482,70444,70444,70414,70414,70365,70365,70347,70347,70294,70294,70272,70272,70213,70213,70181,70181,70129,70129,70073,70073,70051,70051,70034,70034,69993,69993,69943,69943,69913,69913,69897,69897,69877,69877,69849,69849,69828,69828,69781,69781,69763,69763,69746,69746,69695,69695,69653,69653,69612,69612,69583,69583,69563,69563,69542,69542,69526,69526,69503,69503,69457,69457,69427,69427,69398,69398,69366,69366,69350,69350,69334,69334,69306,69306,69289,69289,69271,69271,69197,69197,69151,69151,69121,69121,69098,69098,69074,69074,69032,69032,68971,68971,68952,68952,68923,68923,68858,68858,68827,68827,68801,68801,68778,68778,68745,68745,68729,68729,68690,68690,68656,68656,68602,68602,68579,68579,68562,68562,68514,68514,68497,68497,68465,68465,68449,68449,68427,68427,68375,68375,68335,68335,68316,68316,68287,68287,68244,68244,68205,68205,68188,68188,68155,68155,68105,68105,68085,68085,68026,68026,68001,68001,67934,67934,67890,67890,67867,67867,67820,67820,67773,67773,67711,67711,67648,67648,67632,67632,67605,67605,67587,67587,67558,67558,67518,67518,67481,67481,67459,67459,67412,67412,67381,67381,67349,67349,67332,67332,67315,67315,67295,67295,67275,67275,67238,67238,67180,67180,67149,67149,67123,67123,67102,67102,67075,67075,67057,67057,67039,67039,67022,67022,67005,67005,66988,66988,66948,66948,66903,66903,66879,66879,66829,66829,66744,66744,66706,66706,66688,66688,66660,66660,66605,66605,66585,66585,66550,66550,66524,66524,66488,66488,66459,66459,66440,66440,66413,66413,66394,66394,66377,66377,66350,66350,66331,66331,66313,66313,66281,66281,66252,66252,66211,66211,66156,66156,66129,66129,66107,66107,66055,66055,66023,66023,66005,66005,65956,65956,65928,65928,65871,65871,65855,65855,65823,65823,65807,65807,65776,65776,65744,65744,65724,65724,65706,65706,65685,65685,65664,65664,65646,65646,65622,65622,65603,65603,65586,65586,65569,65569,65542,65542,65513,65513,65483,65483,65445,65445,65427,65427,65396,65396,65369,65369,65352,65352,65319,65319,65295,65295,65241,65241,65223,65223,65204,65204,65142,65142,65107,65107,65082,65082,65065,65065,65046,65046,65027,65027,64975,64975,64937,64937,64909,64909,64880,64880,64863,64863,64832,64832,64814,64814,64779,64779,64729,64729,64669,64669,64629,64629,64613,64613,64597,64597,64561,64561,64526,64526,64485,64485,64466,64466,64442,64442,64422,64422,64396,64396,64363,64363,64327,64327,64305,64305,64277,64277,64259,64259,64242,64242,64224,64224,64185,64185,64156,64156,64120,64120,64029,64029,63991,63991,63972,63972,63922,63922,63906,63906,63859,63859,63832,63832,63814,63814,63784,63784,63756,63756,63740,63740,63703,63703,63652,63652,63611,63611,63583,63583,63560,63560,63523,63523,63505,63505,63485,63485,63444,63444,63375,63375,63325,63325,63283,63283,63266,63266,63231,63231,63211,63211,63186,63186,63161,63161,63140,63140,63114,63114,63060,63060,63037,63037,62998,62998,62941,62941,62925,62925,62898,62898,62875,62875,62772,62772,62755,62755,62717,62717,62677,62677,62643,62643,62612,62612,62581,62581,62557,62557,62532,62532,62513,62513,62495,62495,62472,62472,62424,62424,62352,62352,62311,62311,62256,62256,62217,62217,62179,62179,62146,62146,62119,62119,62076,62076,62027,62027,62010,62010,61956,61956,61922,61922,61905,61905,61887,61887,61834,61834,61798,61798,61762,61762,61734,61734,61713,61713,61688,61688,61620,61620,61600,61600,61559,61559,61543,61543,61512,61512,61489,61489,61395,61395,61317,61317,61289,61289,61273,61273,61249,61249,61217,61217,61180,61180,61151,61151,61134,61134,61058,61058,60987,60987,60968,60968,60941,60941,60922,60922,60890,60890,60847,60847,60819,60819,60802,60802,60780,60780,60754,60754,60730,60730,60707,60707,60670,60670,60651,60651,60611,60611,60584,60584,60562,60562,60545,60545,60526,60526,60508,60508,60472,60472,60416,60416,60378,60378,60342,60342,60307,60307,60278,60278,60239,60239,60222,60222,60175,60175,60154,60154,60136,60136,60095,60095,60074,60074,60042,60042,60004,60004,59987,59987,59968,59968,59948,59948,59932,59932,59878,59878,59855,59855,59827,59827,59805,59805,59772,59772,59749,59749,59732,59732,59715,59715,59695,59695,59669,59669,59652,59652,59630,59630,59612,59612,59588,59588,59571,59571,59500,59500,59471,59471,59410,59410,59390,59390,59345,59345,59302,59302,59282,59282,59244,59244,59180,59180,59155,59155,59113,59113,59072,59072,59037,59037,58990,58990,58966,58966,58944,58944,58908,58908,58886,58886,58868,58868,58845,58845,58824,58824,58801,58801,58761,58761,58742,58742,58713,58713,58675,58675,58640,58640,58613,58613,58577,58577,58553,58553,58534,58534,58499,58499,58443,58443,58426,58426,58394,58394,58378,58378,58329,58329,58306,58306,58289,58289,58229,58229,58213,58213,58193,58193,58154,58154,58132,58132,58107,58107,58089,58089,58070,58070,58052,58052,58019,58019,58003,58003,57969,57969,57949,57949,57908,57908,57873,57873,57835,57835,57814,57814,57784,57784,57758,57758,57721,57721,57699,57699,57664,57664,57591,57591,57558,57558,57541,57541,57517,57517,57496,57496,57479,57479,57457,57457,57400,57400,57370,57370,57331,57331,57283,57283,57260,57260,57235,57235,57195,57195,57170,57170,57125,57125,57060,57060,57041,57041,57006,57006,56964,56964,56939,56939,56914,56914,56891,56891,56862,56862,56841,56841,56817,56817,56781,56781,56763,56763,56734,56734,56703,56703,56685,56685,56666,56666,56650,56650,56618,56618,56592,56592,56558,56558,56525,56525,56472,56472,56437,56437,56397,56397,56375,56375,56356,56356,56324,56324,56304,56304,56283,56283,56260,56260,56235,56235,56185,56185,56161,56161,56139,56139,56096,56096,56079,56079,56006,56006,55989,55989,55924,55924,55891,55891,55860,55860,55837,55837,55810,55810,55785,55785,55726,55726,55708,55708,55680,55680,55651,55651,55632,55632,55596,55596,55576,55576,55549,55549,55518,55518,55487,55487,55458,55458,55410,55410,55350,55350,55333,55333,55284,55284,55258,55258,55233,55233,55184,55184,55155,55155,55137,55137,55085,55085,55067,55067,55050,55050,55022,55022,55004,55004,54979,54979,54953,54953,54925,54925,54886,54886,54868,54868,54830,54830,54810,54810,54783,54783,54766,54766,54741,54741,54700,54700,54684,54684,54657,54657,54638,54638,54612,54612,54585,54585,54538,54538,54519,54519,54503,54503,54478,54478,54415,54415,54389,54389,54368,54368,54305,54305,54263,54263,54247,54247,54218,54218,54191,54191,54169,54169,54147,54147,54130,54130,54108,54108,54071,54071,54048,54048,54031,54031,54006,54006,53990,53990,53963,53963,53928,53928,53908,53908,53859,53859,53837,53837,53749,53749,53716,53716,53679,53679,53655,53655,53630,53630,53611,53611,53560,53560,53524,53524,53502,53502,53440,53440,53369,53369,53349,53349,53329,53329,53296,53296,53246,53246,53214,53214,53197,53197,53149,53149,53129,53129,53101,53101,53034,53034,53003,53003,52974,52974,52926,52926,52880,52880,52862,52862,52833,52833,52794,52794,52764,52764,52745,52745,52710,52710,52692,52692,52673,52673,52641,52641,52618,52618,52580,52580,52549,52549,52519,52519,52482,52482,52453,52453,52432,52432,52389,52389,52353,52353,52322,52322,52295,52295,52276,52276,52245,52245,52228,52228,52212,52212,52185,52185,52148,52148,52119,52119,52087,52087,52059,52059,52007,52007,51983,51983,51957,51957,51940,51940,51915,51915,51892,51892,51868,51868,51839,51839,51821,51821,51804,51804,51788,51788,51761,51761,51744,51744,51726,51726,51701,51701,51656,51656,51611,51611,51587,51587,51560,51560,51525,51525,51509,51509,51469,51469,51441,51441,51422,51422,51398,51398,51378,51378,51350,51350,51304,51304,51280,51280,51243,51243,51160,51160,51135,51135,51119,51119,51069,51069,51027,51027,51003,51003,50985,50985,50950,50950,50907,50907,50870,50870,50816,50816,50790,50790,50750,50750,50720,50720,50682,50682,50632,50632,50608,50608,50587,50587,50559,50559,50542,50542,50523,50523,50501,50501,50453,50453,50418,50418,50397,50397,50346,50346,50274,50274,50254,50254,50229,50229,50207,50207,50167,50167,50141,50141,50115,50115,50075,50075,50049,50049,50027,50027,50008,50008,49988,49988,49972,49972,49945,49945,49914,49914,49898,49898,49858,49858,49829,49829,49790,49790,49754,49754,49726,49726,49687,49687,49655,49655,49636,49636,49608,49608,49590,49590,49554,49554,49504,49504,49471,49471,49451,49451,49427,49427,49411,49411,49389,49389,49371,49371,49348,49348,49324,49324,49305,49305,49288,49288,49257,49257,49235,49235,49219,49219,49199,49199,49122,49122,49083,49083,49060,49060,49044,49044,49020,49020,48988,48988,48970,48970,48924,48924,48889,48889,48865,48865,48846,48846,48823,48823,48799,48799,48769,48769,48749,48749,48727,48727,48709,48709,48693,48693,48675,48675,48658,48658,48626,48626,48570,48570,48551,48551,48511,48511,48474,48474,48425,48425,48396,48396,48376,48376,48321,48321,48298,48298,48265,48265,48243,48243,48221,48221,48204,48204,48162,48162,48145,48145,48112,48112,48083,48083,48063,48063,48044,48044,48028,48028,47994,47994,47972,47972,47945,47945,47909,47909,47890,47890,47870,47870,47844,47844,47811,47811,47765,47765,47748,47748,47732,47732,47712,47712,47691,47691,47666,47666,47646,47646,47624,47624,47557,47557,47530,47530,47500,47500,47467,47467,47423,47423,47403,47403,47386,47386,47360,47360,47326,47326,47302,47302,47279,47279,47243,47243,47225,47225,47165,47165,47137,47137,47119,47119,47087,47087,47068,47068,46991,46991,46962,46962,46933,46933,46886,46886,46856,46856,46831,46831,46815,46815,46790,46790,46774,46774,46753,46753,46734,46734,46717,46717,46692,46692,46666,46666,46632,46632,46615,46615,46571,46571,46542,46542,46497,46497,46477,46477,46461,46461,46430,46430,46401,46401,46372,46372,46345,46345,46303,46303,46261,46261,46232,46232,46211,46211,46139,46139,46117,46117,46094,46094,46077,46077,46061,46061,46035,46035,46019,46019,46002,46002,45985,45985,45965,45965,45938,45938,45911,45911,45873,45873,45835,45835,45814,45814,45779,45779,45749,45749,45730,45730,45703,45703,45682,45682,45653,45653,45628,45628,45606,45606,45590,45590,45565,45565,45547,45547,45526,45526,45507,45507,45488,45488,45467,45467,45425,45425,45368,45368,45338,45338,45308,45308,45275,45275,45192,45192,45168,45168,45150,45150,45127,45127,45111,45111,45076,45076,45051,45051,45031,45031,45007,45007,44984,44984,44959,44959,44917,44917,44892,44892,44825,44825,44809,44809,44760,44760,44736,44736,44716,44716,44699,44699,44681,44681,44658,44658,44642,44642,44621,44621,44589,44589,44552,44552,44519,44519,44459,44459,44433,44433,44400,44400,44382,44382,44360,44360,44343,44343,44327,44327,44311,44311,44292,44292,44258,44258,44232,44232,44216,44216,44192,44192,44162,44162,44141,44141,44117,44117,44089,44089,44032,44032,44012,44012,43975,43975,43952,43952,43936,43936,43892,43892,43874,43874,43848,43848,43827,43827,43777,43777,43750,43750,43713,43713,43691,43691,43665,43665,43645,43645,43610,43610,43582,43582,43555,43555,43532,43532,43516,43516,43499,43499,43472,43472,43450,43450,43434,43434,43412,43412,43391,43391,43370,43370,43335,43335,43289,43289,43271,43271,43255,43255,43230,43230,43205,43205,43180,43180,43155,43155,43136,43136,43115,43115,43078,43078,43040,43040,42989,42989,42965,42965,42944,42944,42928,42928,42906,42906,42880,42880,42861,42861,42843,42843,42827,42827,42803,42803,42773,42773,42749,42749,42733,42733,42710,42710,42691,42691,42669,42669,42619,42619,42579,42579,42560,42560,42534,42534,42513,42513,42495,42495,42465,42465,42440,42440,42412,42412,42378,42378,42361,42361,42342,42342,42323,42323,42303,42303,42286,42286,42270,42270,42247,42247,42229,42229,42183,42183,42163,42163,42131,42131,42109,42109,42087,42087,42066,42066,42048,42048,42031,42031,42014,42014,41997,41997,41977,41977,41961,41961,41945,41945,41923,41923,41891,41891,41870,41870,41854,41854,41834,41834,41815,41815,41797,41797,41781,41781,41703,41703,41683,41683,41651,41651,41630,41630,41611,41611,41592,41592,41540,41540,41498,41498,41465,41465,41445,41445,41408,41408,41389,41389,41364,41364,41342,41342,41289,41289,41265,41265,41248,41248,41227,41227,41195,41195,41154,41154,41138,41138,41113,41113,41086,41086,41068,41068,41011,41011,40973,40973,40938,40938,40910,40910,40886,40886,40863,40863,40804,40804,40788,40788,40767,40767,40734,40734,40718,40718,40688,40688,40666,40666,40628,40628,40595,40595,40554,40554,40535,40535,40519,40519,40498,40498,40478,40478,40455,40455,40434,40434,40405,40405,40379,40379,40348,40348,40308,40308,40291,40291,40272,40272,40250,40250,40229,40229,40212,40212,40175,40175,40157,40157,40135,40135,40115,40115,40085,40085,40065,40065,40039,40039,40021,40021,40005,40005,39987,39987,39962,39962,39945,39945,39922,39922,39880,39880,39862,39862,39835,39835,39819,39819,39798,39798,39773,39773,39682,39682,39658,39658,39639,39639,39618,39618,39599,39599,39573,39573,39557,39557,39533,39533,39503,39503,39482,39482,39461,39461,39444,39444,39422,39422,39390,39390,39347,39347,39331,39331,39303,39303,39250,39250,39223,39223,39191,39191,39171,39171,39151,39151,39097,39097,39078,39078,39055,39055,39027,39027,38990,38990,38954,38954,38931,38931,38911,38911,38873,38873,38845,38845,38829,38829,38781,38781,38755,38755,38731,38731,38709,38709,38686,38686,38665,38665,38621,38621,38605,38605,38544,38544,38517,38517,38484,38484,38453,38453,38436,38436,38420,38420,38400,38400,38383,38383,38361,38361,38344,38344,38326,38326,38309,38309,38276,38276,38253,38253,38236,38236,38218,38218,38192,38192,38171,38171,38152,38152,38134,38134,38116,38116,38098,38098,38065,38065,38041,38041,38024,38024,38007,38007,37985,37985,37958,37958,37936,37936,37909,37909,37886,37886,37867,37867,37851,37851,37830,37830,37808,37808,37788,37788,37763,37763,37741,37741,37724,37724,37704,37704,37671,37671,37648,37648,37625,37625,37608,37608,37569,37569,37543,37543,37516,37516,37494,37494,37467,37467,37445,37445,37426,37426,37388,37388,37370,37370,37336,37336,37320,37320,37302,37302,37285,37285,37255,37255,37221,37221,37182,37182,37147,37147,37127,37127,37107,37107,37077,37077,37057,37057,37039,37039,37013,37013,36993,36993,36962,36962,36925,36925,36907,36907,36889,36889,36870,36870,36854,36854,36829,36829,36801,36801,36780,36780,36737,36737,36719,36719,36703,36703,36680,36680,36659,36659,36621,36621,36601,36601,36541,36541,36513,36513,36490,36490,36471,36471,36453,36453,36429,36429,36409,36409,36383,36383,36367,36367,36346,36346,36323,36323,36305,36305,36286,36286,36268,36268,36251,36251,36222,36222,36194,36194,36158,36158,36136,36136,36116,36116,36094,36094,36063,36063,36045,36045,36015,36015,35970,35970,35954,35954,35928,35928,35907,35907,35891,35891,35870,35870,35853,35853,35831,35831,35810,35810,35794,35794,35776,35776,35723,35723,35704,35704,35685,35685,35667,35667,35637,35637,35621,35621,35605,35605,35576,35576,35546,35546,35525,35525,35500,35500,35484,35484,35419,35419,35400,35400,35384,35384,35342,35342,35292,35292,35274,35274,35254,35254,35231,35231,35198,35198,35182,35182,35166,35166,35148,35148,35131,35131,35115,35115,35090,35090,35041,35041,35017,35017,34994,34994,34968,34968,34950,34950,34915,34915,34895,34895,34877,34877,34856,34856,34837,34837,34817,34817,34793,34793,34776,34776,34758,34758,34742,34742,34725,34725,34706,34706,34690,34690,34671,34671,34654,34654,34637,34637,34615,34615,34586,34586,34566,34566,34548,34548,34518,34518,34474,34474,34411,34411,34394,34394,34376,34376,34349,34349,34322,34322,34303,34303,34285,34285,34269,34269,34250,34250,34231,34231,34205,34205,34189,34189,34167,34167,34129,34129,34105,34105,34085,34085,34064,34064,34048,34048,34031,34031,34000,34000,33980,33980,33961,33961,33942,33942,33896,33896,33878,33878,33833,33833,33800,33800,33775,33775,33758,33758,33716,33716,33699,33699,33681,33681,33642,33642,33620,33620,33597,33597,33557,33557,33500,33500,33482,33482,33422,33422,33405,33405,33388,33388,33371,33371,33355,33355,33338,33338,33316,33316,33300,33300,33281,33281,33258,33258,33242,33242,33216,33216,33197,33197,33179,33179,33156,33156,33137,33137,33094,33094,33078,33078,33062,33062,33028,33028,33011,33011,32995,32995,32972,32972,32948,32948,32913,32913,32863,32863,32830,32830,32811,32811,32767,32767,32751,32751,32715,32715,32699,32699,32671,32671,32654,32654,32630,32630,32607,32607,32590,32590,32549,32549,32517,32517,32500,32500,32484,32484,32464,32464,32441,32441,32425,32425,32402,32402,32366,32366,32350,32350,32313,32313,32295,32295,32278,32278,32247,32247,32231,32231,32200,32200,32175,32175,32158,32158,32142,32142,32125,32125,32105,32105,32089,32089,32066,32066,32047,32047,32029,32029,31999,31999,31980,31980,31957,31957,31939,31939,31923,31923,31905,31905,31885,31885,31863,31863,31842,31842,31820,31820,31788,31788,31755,31755,31728,31728,31712,31712,31692,31692,31662,31662,31645,31645,31623,31623,31607,31607,31591,31591,31571,31571,31551,31551,31520,31520,31503,31503,31483,31483,31460,31460,31444,31444,31425,31425,31407,31407,31389,31389,31370,31370,31344,31344,31323,31323,31300,31300,31284,31284,31260,31260,31244,31244,31216,31216,31177,31177,31159,31159,31140,31140,31118,31118,31094,31094,31070,31070,31050,31050,31034,31034,31015,31015,30982,30982,30965,30965,30938,30938,30905,30905,30884,30884,30864,30864,30831,30831,30806,30806,30787,30787,30769,30769,30730,30730,30707,30707,30689,30689,30672,30672,30649,30649,30632,30632,30616,30616,30600,30600,30584,30584,30568,30568,30546,30546,30522,30522,30502,30502,30483,30483,30460,30460,30439,30439,30422,30422,30391,30391,30368,30368,30348,30348,30311,30311,30292,30292,30263,30263,30239,30239,30211,30211,30184,30184,30159,30159,30118,30118,30097,30097,30059,30059,30039,30039,30023,30023,29977,29977,29952,29952,29933,29933,29916,29916,29898,29898,29872,29872,29848,29848,29827,29827,29802,29802,29779,29779,29757,29757,29730,29730,29705,29705,29688,29688,29664,29664,29635,29635,29618,29618,29601,29601,29580,29580,29564,29564,29547,29547,29522,29522,29475,29475,29449,29449,29432,29432,29411,29411,29394,29394,29370,29370,29341,29341,29317,29317,29298,29298,29282,29282,29266,29266,29250,29250,29226,29226,29205,29205,29188,29188,29168,29168,29151,29151,29107,29107,29086,29086,29069,29069,29048,29048,29027,29027,28983,28983,28964,28964,28948,28948,28931,28931,28914,28914,28897,28897,28869,28869,28847,28847,28829,28829,28806,28806,28784,28784,28765,28765,28743,28743,28723,28723,28686,28686,28666,28666,28640,28640,28621,28621,28605,28605,28589,28589,28572,28572,28549,28549,28531,28531,28510,28510,28492,28492,28471,28471,28451,28451,28431,28431,28415,28415,28399,28399,28371,28371,28355,28355,28339,28339,28322,28322,28303,28303,28286,28286,28266,28266,28249,28249,28226,28226,28201,28201,28181,28181,28165,28165,28146,28146,28130,28130,28113,28113,28096,28096,28080,28080,28060,28060,28044,28044,28027,28027,28004,28004,27987,27987,27971,27971,27951,27951,27933,27933,27914,27914,27894,27894,27877,27877,27851,27851,27828,27828,27812,27812,27795,27795,27779,27779,27759,27759,27739,27739,27717,27717,27701,27701,27684,27684,27662,27662,27634,27634,27616,27616,27592,27592,27570,27570,27550,27550,27525,27525,27505,27505,27483,27483,27460,27460,27441,27441,27424,27424,27403,27403,27387,27387,27371,27371,27355,27355,27338,27338,27322,27322,27305,27305,27284,27284,27267,27267,27251,27251,27229,27229,27206,27206,27189,27189,27172,27172,27156,27156,27140,27140,27118,27118,27096,27096,27077,27077,27061,27061,27045,27045,27028,27028,27011,27011,26993,26993,26964,26964,26947,26947,26931,26931,26915,26915,26891,26891,26875,26875,26844,26844,26825,26825,26807,26807,26790,26790,26761,26761,26744,26744,26717,26717,26698,26698,26679,26679,26663,26663,26646,26646,26629,26629,26610,26610,26594,26594,26576,26576,26541,26541,26518,26518,26499,26499,26480,26480,26460,26460,26444,26444,26427,26427,26410,26410,26391,26391,26375,26375,26354,26354,26334,26334,26313,26313,26291,26291,26267,26267,26247,26247,26228,26228,26201,26201,26183,26183,26150,26150,26129,26129,26113,26113,26088,26088,26064,26064,26040,26040,26022,26022,26004,26004,25980,25980,25964,25964,25946,25946,25930,25930,25911,25911,25893,25893,25861,25861,25840,25840,25821,25821,25805,25805,25786,25786,25770,25770,25753,25753,25727,25727,25711,25711,25695,25695,25674,25674,25658,25658,25641,25641,25621,25621,25600,25600,25584,25584,25565,25565,25547,25547,25523,25523,25507,25507,25491,25491,25474,25474,25456,25456,25438,25438,25421,25421,25402,25402,25386,25386,25370,25370,25349,25349,25333,25333,25316,25316,25296,25296,25274,25274,25258,25258,25240,25240,25224,25224,25200,25200,25181,25181,25162,25162,25140,25140,25108,25108,25090,25090,25073,25073,25056,25056,25031,25031,25014,25014,24996,24996,24980,24980,24961,24961,24940,24940,24921,24921,24896,24896,24878,24878,24861,24861,24845,24845,24828,24828,24811,24811,24794,24794,24778,24778,24755,24755,24739,24739,24716,24716,24696,24696,24669,24669,24651,24651,24634,24634,24617,24617,24595,24595,24571,24571,24555,24555,24539,24539,24513,24513,24492,24492,24475,24475,24455,24455,24437,24437,24419,24419,24403,24403,24385,24385,24366,24366,24347,24347,24317,24317,24301,24301,24284,24284,24262,24262,24246,24246,24227,24227,24204,24204,24188,24188,24172,24172,24150,24150,24129,24129,24108,24108,24091,24091,24074,24074,24058,24058,24041,24041,24025,24025,24005,24005,23989,23989,23962,23962,23945,23945,23922,23922,23906,23906,23889,23889,23872,23872,23851,23851,23823,23823,23798,23798,23777,23777,23761,23761,23739,23739,23716,23716,23697,23697,23671,23671,23651,23651,23634,23634,23616,23616,23598,23598,23581,23581,23557,23557,23535,23535,23516,23516,23496,23496,23478,23478,23458,23458,23433,23433,23417,23417,23400,23400,23384,23384,23368,23368,23350,23350,23334,23334,23315,23315,23297,23297,23281,23281,23261,23261,23244,23244,23215,23215,23198,23198,23179,23179,23163,23163,23142,23142,23124,23124,23108,23108,23090,23090,23066,23066,23047,23047,23028,23028,23012,23012,22995,22995,22978,22978,22955,22955,22938,22938,22910,22910,22892,22892,22872,22872,22852,22852,22831,22831,22809,22809,22775,22775,22756,22756,22740,22740,22719,22719,22702,22702,22685,22685,22669,22669,22652,22652,22636,22636,22620,22620,22600,22600,22582,22582,22564,22564,22543,22543,22526,22526,22510,22510,22494,22494,22477,22477,22461,22461,22433,22433,22416,22416,22400,22400,22383,22383,22365,22365,22348,22348,22330,22330,22313,22313,22292,22292,22269,22269,22253,22253,22236,22236,22219,22219,22201,22201,22184,22184,22164,22164,22147,22147,22130,22130,22112,22112,22096,22096,22079,22079,22063,22063,22047,22047,22030,22030,22014,22014,21998,21998,21982,21982,21964,21964,21947,21947,21928,21928,21912,21912,21890,21890,21873,21873,21855,21855,21839,21839,21819,21819,21802,21802,21780,21780,21756,21756,21740,21740,21721,21721,21704,21704,21685,21685,21658,21658,21633,21633,21615,21615,21598,21598,21582,21582,21564,21564,21547,21547,21530,21530,21514,21514,21496,21496,21478,21478,21453,21453,21436,21436,21420,21420,21400,21400,21382,21382,21364,21364,21340,21340,21323,21323,21305,21305,21281,21281,21265,21265,21249,21249,21232,21232,21215,21215,21199,21199,21180,21180,21164,21164,21140,21140,21120,21120,21104,21104,21082,21082,21065,21065,21046,21046,21009,21009,20993,20993,20975,20975,20959,20959,20942,20942,20926,20926,20908,20908,20891,20891,20875,20875,20859,20859,20843,20843,20826,20826,20810,20810,20793,20793,20776,20776,20754,20754,20734,20734,20718,20718,20702,20702,20684,20684,20664,20664,20648,20648,20632,20632,20614,20614,20596,20596,20580,20580,20561,20561,20545,20545,20528,20528,20506,20506,20485,20485,20465,20465,20445,20445,20428,20428,20402,20402,20385,20385,20368,20368,20348,20348,20331,20331,20312,20312,20294,20294,20273,20273,20257,20257,20240,20240,20221,20221,20203,20203,20187,20187,20171,20171,20151,20151,20134,20134,20116,20116,20099,20099,20080,20080,20064,20064,20045,20045,20029,20029,20011,20011,19993,19993,19976,19976,19959,19959,19943,19943,19923,19923,19906,19906,19890,19890,19874,19874,19854,19854,19833,19833,19817,19817,19794,19794,19777,19777,19761,19761,19745,19745,19723,19723,19705,19705,19689,19689,19666,19666,19649,19649,19632,19632,19612,19612,19596,19596,19580,19580,19562,19562,19541,19541,19525,19525,19504,19504,19486,19486,19461,19461,19443,19443,19426,19426,19410,19410,19394,19394,19375,19375,19358,19358,19334,19334,19317,19317,19300,19300,19278,19278,19262,19262,19245,19245,19228,19228,19212,19212,19191,19191,19175,19175,19155,19155,19136,19136,19119,19119,19103,19103,19082,19082,19064,19064,19046,19046,19027,19027,19011,19011,18995,18995,18978,18978,18962,18962,18946,18946,18924,18924,18902,18902,18885,18885,18866,18866,18850,18850,18834,18834,18818,18818,18802,18802,18783,18783,18766,18766,18750,18750,18734,18734,18716,18716,18700,18700,18682,18682,18666,18666,18649,18649,18632,18632,18608,18608,18589,18589,18573,18573,18557,18557,18538,18538,18522,18522,18504,18504,18487,18487,18471,18471,18454,18454,18436,18436,18415,18415,18398,18398,18380,18380,18364,18364,18348,18348,18332,18332,18314,18314,18297,18297,18281,18281,18264,18264,18248,18248,18231,18231,18214,18214,18198,18198,18182,18182,18166,18166,18150,18150,18134,18134,18112,18112,18095,18095,18077,18077,18061,18061,18043,18043,18026,18026,18004,18004,17985,17985,17968,17968,17951,17951,17934,17934,17918,17918,17900,17900,17884,17884,17857,17857,17840,17840,17824,17824,17806,17806,17789,17789,17772,17772,17756,17756,17739,17739,17723,17723,17705,17705,17689,17689,17673,17673,17657,17657,17640,17640,17624,17624,17605,17605,17588,17588,17568,17568,17547,17547,17531,17531,17515,17515,17499,17499,17483,17483,17467,17467,17450,17450,17434,17434,17413,17413,17397,17397,17381,17381,17365,17365,17349,17349,17332,17332,17315,17315,17298,17298,17280,17280,17261,17261,17243,17243,17227,17227,17207,17207,17190,17190,17169,17169,17153,17153,17136,17136,17119,17119,17098,17098,17082,17082,17064,17064,17042,17042,17026,17026,16947,16947,16927,16927,16829,16829,16094,16094,14701,14701,12916,12916,11586,11586,10948,10948,8779,8779,8350,8350,7853,7853,7581,7581,7463,7463,7150,7150,6888,6888,6566,6566,5885,5885,5461,5461,4958,4958,4146,4146,7,7,0.0]],"coord\_x":[[0.0,1e-10,0.3276220373974474,0.3276220374974474,0.6326923319297183,0.6326923320297183,0.9220411157981557,0.9220411158981557,1.196147274543727,1.196147274643727,1.4666044432697563,1.4666044433697563,1.730015693887732,1.730015693987732,1.9814722264916949,1.9814722265916949,2.23249481704303,2.23249481714303,2.4723092369764212,2.4723092370764213,2.706798763834601,2.706798763934601,2.9233251744635713,2.9233251745635713,3.127763892897381,3.1277638929973812,3.323838660279149,3.323838660379149,3.5196137449090408,3.519613745009041,3.7099222163297187,3.7099222164297188,3.8969641279060605,3.8969641280060605,4.080939815505074,4.080939815605074,4.260576832230912,4.260576832330912,4.438897957878767,4.438897957978767,4.614902119975635,4.614902120075635,4.789632657150353,4.789632657250353,4.958754039933219,4.958754040033219,5.126104135227658,5.126104135327658,5.29070400317605,5.29070400327605,5.45488635004452,5.45488635014452,5.6182904855978135,5.6182904856978135,5.778466365010495,5.778466365110495,5.937690649054889,5.937690649154889,6.09259568181122,6.09259568191122,6.246912593860325,6.246912593960325,6.401220010303475,6.401220010403475,6.5541079764472965,6.5541079765472965,6.7064885345345155,6.7064885346345156,6.858371073077788,6.858371073177788,7.009840160086549,7.009840160186549,7.160199332131481,7.160199332231481,7.310027963966957,7.310027964066957,7.459617456462966,7.459617456562966,7.607122413567305,7.607122413667305,7.754247332246811,7.754247332346811,7.901322773821599,7.901322773921599,8.047444692348696,8.047444692448696,8.192355599836525,8.192355599936525,8.33703711347521,8.33703711357521,8.48033255422648,8.48033255432648,8.622831899081431,8.622831899181431,8.76402709742366,8.76402709752366,8.904381791847731,8.904381791947731,9.044225151460859,9.044225151560859,9.183521835473957,9.183521835573957,9.322485744905402,9.322485745005402,9.459497022193322,9.459497022293322,9.596236425289666,9.596236425389666,9.732506081473328,9.732506081573328,9.868702271653019,9.868702271753019,10.004575325648077,10.004575325748077,10.14034710507886,10.14034710517886,10.27596620516125,10.27596620526125,10.41089491043467,10.41089491053467,10.545666545534386,10.545666545634386,10.680246555022181,10.680246555122181,10.8146817029725,10.8146817030725,10.948523125666213,10.948523125766213,11.081637527642272,11.081637527742272,11.214035332648566,11.214035332748566,11.346420857623096,11.346420857723096,11.478406032143512,11.478406032243512,11.61029592932447,11.61029592942447,11.742020188867702,11.742020188967702,11.87363671255088,11.87363671265088,12.004409947890093,12.004409947990093,12.134308159570699,12.134308159670699,12.263043837779287,12.263043837879287,12.391494719204733,12.391494719304733,12.518560527280233,12.518560527380233,12.645526524399894,12.645526524499894,12.771252345404854,12.771252345504854,12.89683558952941,12.89683558962941,13.02228625214825,13.02228625224825,13.147402355297098,13.147402355397098,13.272418683187876,13.272418683287876,13.396477989649823,13.396477989749823,13.520463187547993,13.520463187647993,13.643032005154774,13.643032005254774,13.76555288066081,13.76555288076081,13.88801656834451,13.88801656844451,14.010059236567146,14.010059236667146,14.132008697920796,14.132008698020796,14.253833466975182,14.253833467075182,14.37546636053328,14.37546636063328,14.497088330574753,14.497088330674753,14.618631194365106,14.618631194465106,14.738272652307222,14.738272652407222,14.857832005385811,14.857832005485811,14.976983939852467,14.976983939952467,15.095578132411402,15.095578132511402,15.214151727358923,15.214151727458923,15.332576426921324,15.332576427021325,15.450812713670812,15.450812713770812,15.569008590548338,15.569008590648338,15.68706342554943,15.68706342564943,15.804034440654188,15.804034440754188,15.920929847888967,15.920929847988967,16.037704275392226,16.037704275492224,16.154358258630467,16.154358258730465,16.270145321603156,16.270145321703154,16.385548062418998,16.385548062518996,16.500233599403046,16.500233599503044,16.614844028285845,16.614844028385843,16.729122182355727,16.729122182455725,16.84309197911641,16.843091979216407,16.956981206017534,16.956981206117533,17.068796071390516,17.068796071490514,17.180603797210146,17.180603797310145,17.29237057769131,17.29237057779131,17.4040620715824,17.404062071682397,17.515682169939986,17.515682170039984,17.627285883022633,17.62728588312263,17.738831622932075,17.738831623032073,17.84872737636451,17.84872737646451,17.958538954462945,17.958538954562943,18.068254791150956,18.068254791250954,18.176261347371383,18.17626134747138,18.28388101119576,18.283881011295758,18.39140803931542,18.391408039415417,18.498851784545245,18.498851784645243,18.606138495299135,18.606138495399133,18.71326385214731,18.71326385224731,18.82025234375776,18.820252343857756,18.927139310919568,18.927139311019566,19.033992222411896,19.033992222511895,19.140705270054095,19.140705270154093,19.24708904149578,19.24708904159578,19.353468921880886,19.353468921980884,19.459771231018845,19.459771231118843,19.564766716311574,19.564766716411572,19.669150698859855,19.669150698959854,19.773338700668674,19.773338700768672,19.87723926405961,19.87723926415961,19.980591402659947,19.980591402759945,20.083675343938683,20.08367534403868,20.186576262767286,20.186576262867284,20.289147120044507,20.289147120144506,20.39146652225273,20.39146652235273,20.49325406343413,20.493254063534128,20.594708401660675,20.594708401760673,20.69614578344801,20.69614578354801,20.797519801699362,20.79751980179936,20.898721756714973,20.89872175681497,20.99964302019062,20.99964302029062,21.099678265095505,21.099678265195504,21.199475048918487,21.199475049018485,21.29926469318812,21.29926469328812,21.399013249328224,21.399013249428222,21.498659495668818,21.498659495768816,21.59793198639153,21.597931986491528,21.6968918717708,21.696891871870797,21.795814096006144,21.79581409610614,21.8943618149705,21.8943618150705,21.992750107708552,21.99275010780855,22.091137436606896,22.091137436706894,22.189157399787607,22.189157399887605,22.28706355848791,22.28706355858791,22.3844552409738,22.3844552410738,22.48166822043933,22.48166822053933,22.57863906195299,22.57863906205299,22.675451369684506,22.675451369784504,22.772059022119237,22.772059022219235,22.8680735561594,22.868073556259397,22.963590142051146,22.963590142151144,23.0588418505136,23.058841850613597,23.154075210323942,23.15407521042394,23.248973082522358,23.248973082622356,23.343638954934654,23.343638955034653,23.438243641374743,23.43824364147474,23.53262507398157,23.53262507408157,23.626969023933306,23.626969024033304,23.720649495192205,23.720649495292204,23.814329288193534,23.814329288293532,23.907771405463837,23.907771405563835,24.00072067936637,24.00072067946637,24.09329144984804,24.093291449948037,24.18552805353515,24.18552805363515,24.277749949742365,24.277749949842363,24.369741773842865,24.369741773942863,24.46160219446395,24.461602194563948,24.553011038335637,24.553011038435635,24.643852466204812,24.64385246630481,24.734671547272,24.734671547372,24.82501749013867,24.825017490238668,24.91509266544453,24.915092665544528,25.00510269232607,25.005102692426068,25.094762417022476,25.094762417122475,25.184321152736747,25.184321152836745,25.273593342477305,25.273593342577303,25.36284757624119,25.36284757634119,25.45190150983583,25.45190150993583,25.54078173809746,25.54078173819746,25.62912521473822,25.62912521483822,25.717367809490142,25.71736780959014,25.805229294884224,25.805229294984223,25.892876129606833,25.89287612970683,25.980435040729933,25.98043504082993,26.067894390781568,26.067894390881566,26.155131843515072,26.15513184361507,26.24235601667935,26.242356016779347,26.329490624146842,26.32949062424684,26.415914239193956,26.415914239293954,26.502162292624185,26.502162292724183,26.588364188842007,26.588364188942005,26.67450086523997,26.67450086533997,26.760258467052807,26.760258467152806,26.845827691750497,26.845827691850495,26.931001313797047,26.931001313897045,27.01607273313739,27.016072733237387,27.101029205668794,27.101029205768793,27.18571644564336,27.18571644574336,27.270046208181682,27.27004620828168,27.354235321519,27.354235321618997,27.438141637148117,27.438141637248116,27.521979948531573,27.52197994863157,27.605734584349765,27.605734584449763,27.689215418296975,27.689215418396973,27.772458326628787,27.772458326728785,27.855570509738758,27.855570509838756,27.938659810580237,27.938659810680235,28.021385136991924,28.021385137091922,28.104086224619987,28.104086224719985,28.18653517892149,28.186535179021487,28.268888213323727,28.268888213423725,28.351034843238608,28.351034843338606,28.433020797505343,28.43302079760534,28.514993365109547,28.514993365209545,28.596923737953453,28.59692373805345,28.678832228066337,28.678832228166335,28.760704484945972,28.76070448504597,28.84240360765686,28.842403607756857,28.924040794742435,28.924040794842433,29.005656027701463,29.00565602780146,29.08726165796123,29.087261658061227,29.168857292846308,29.168857292946306,29.250315277142793,29.25031527724279,29.331452766889385,29.331452766989383,29.41253617451935,29.412536174619348,29.493574495869908,29.493574495969906,29.57456890896736,29.57456890906736,29.655432418354142,29.65543241845414,29.736287074694765,29.736287074794763,29.81682819755001,29.816828197650008,29.89722478014768,29.89722478024768,29.977511128041627,29.977511128141625,30.05747355440007,30.057473554500067,30.137139153827977,30.137139153927976,30.216659677531805,30.216659677631803,30.295940633522967,30.295940633622966,30.3751521573578,30.375152157457798,30.454061892272517,30.454061892372515,30.532688900874565,30.532688900974563,30.611313910401673,30.61131391050167,30.689456428913374,30.689456429013372,30.767543758677675,30.767543758777673,30.845182403211684,30.845182403311682,30.922721772256356,30.922721772356354,30.999977094170987,30.999977094270985,31.07689828498883,31.076898285088827,31.153732658837928,31.153732658937926,31.230357201214066,31.230357201314064,31.306910776429905,31.306910776529904,31.383458854189662,31.38345885428966,31.459858179355365,31.459858179455363,31.53604342501386,31.53604342511386,31.611991387616765,31.611991387716763,31.68779923648517,31.687799236585168,31.763588950888064,31.763588950988062,31.839348929051255,31.839348929151253,31.914975183380196,31.914975183480195,31.990545392215335,31.990545392315333,32.06600586611548,32.066005866215484,32.1414387813397,32.1414387814397,32.216586043034376,32.21658604313438,32.29170506779555,32.29170506789555,32.366589665322465,32.36658966542247,32.44087989503297,32.44087989513297,32.51505699852065,32.51505699862065,32.58902105773635,32.589021057836355,32.66298408171682,32.66298408181682,32.73692843576528,32.73692843586528,32.810805285336805,32.81080528543681,32.88460170783985,32.88460170793985,32.95838859903917,32.95838859913917,33.03199139685535,33.03199139695535,33.10553436521445,33.105534365314455,33.17907526310309,33.17907526320309,33.25259417116741,33.25259417126741,33.326058247461994,33.326058247561996,33.39941230323945,33.39941230333945,33.47276450273304,33.472764502833044,33.54601789080826,33.54601789090826,33.61910878264923,33.61910878274923,33.692131063382504,33.692131063482506,33.7650632429525,33.7650632430525,33.837913531845565,33.83791353194557,33.91059289983142,33.91059289993142,33.98294320580337,33.982943205903375,34.05522147368202,34.05522147378202,34.12732303761524,34.12732303771524,34.19929969506261,34.19929969516261,34.27127142621816,34.271271426318165,34.343233161999024,34.343233162099025,34.41471244246224,34.41471244256224,34.48614370942918,34.48614370952918,34.55739727291323,34.55739727301323,34.62860931989455,34.62860931999455,34.69969456840837,34.69969456850837,34.77068236201896,34.77068236211896,34.841667728181406,34.84166772828141,34.912582805441126,34.91258280554113,34.983111204491394,34.983111204591395,35.053309506292514,35.053309506392516,35.12347378812191,35.123473788221915,35.19352030301881,35.19352030311881,35.263365018440254,35.263365018540256,35.33301203962944,35.33301203972944,35.40240242957344,35.402402429673444,35.47170189730553,35.47170189740553,35.5409639180803,35.5409639181803,35.610206162292286,35.61020616239229,35.67909042929929,35.679090429399295,35.747953099157414,35.747953099257415,35.81658430469592,35.816584304795924,35.885196590418055,35.885196590518056,35.95367611614564,35.95367611624564,36.022082675637975,36.02208267573798,36.090330165881674,36.090330165981676,36.15857469321073,36.15857469331073,36.22678951999785,36.22678952009785,36.29484231462169,36.29484231472169,36.36287069197307,36.36287069207307,36.430841238942314,36.430841239042316,36.498598206173085,36.49859820627309,36.565941222100605,36.56594122220061,36.633192316278745,36.63319231637875,36.70011620472685,36.70011620482685,36.766970411134245,36.76697041123425,36.833479884208124,36.833479884308126,36.89991835442394,36.89991835452394,36.96614278083031,36.96614278093031,37.032253581245115,37.03225358134512,37.09811481857257,37.09811481867257,37.16390872991192,37.16390873001192,37.22969414518279,37.22969414528279,37.29546902961246,37.295469029712464,37.36123145552154,37.36123145562154,37.426975711267346,37.42697571136735,37.492083368738676,37.49208336883868,37.556963381551434,37.556963381651435,37.621797522733914,37.621797522833916,37.686563017110934,37.686563017210936,37.75120888827157,37.75120888837157,37.81570868417066,37.81570868427066,37.88012019949257,37.880120199592575,37.94447966747056,37.94447966757056,38.008686491797924,38.008686491897926,38.0726320798682,38.0726320799682,38.136363195755834,38.136363195855836,38.19997069027721,38.199970690377214,38.26332690821843,38.26332690831843,38.326657745047484,38.326657745147486,38.38977535911573,38.38977535921573,38.45280865505892,38.45280865515892,38.51571572369887,38.51571572379887,38.578607549392416,38.57860754949242,38.6412622705192,38.6412622706192,38.703910958723405,38.70391095882341,38.766455123866564,38.766455123966566,38.82893217720823,38.82893217730823,38.8912989601484,38.8912989602484,38.953645609548126,38.95364560964813,39.015950528258514,39.015950528358516,39.07821682198528,39.07821682208528,39.140366205525936,39.14036620562594,39.20249695483235,39.20249695493235,39.264620957260846,39.26462095736085,39.32667666986155,39.32667666996155,39.38862707405033,39.388627074150335,39.450437078922484,39.450437079022485,39.51206156250083,39.512061562600834,39.573572455785374,39.573572455885376,39.63507795870714,39.635077958807145,39.69654476524975,39.69654476534975,39.819448320815354,39.819448320915356,39.88078540167359,39.88078540177359,39.94211630681817,39.94211630691817,40.00328207409377,40.00328207419377,40.06444480705919,40.06444480715919,40.12552825528465,40.12552825538465,40.186532811445595,40.1865328115456,40.24741574531522,40.247415745415225,40.30828889799675,40.308288898096755,40.369043534092675,40.36904353419268,40.42972863093896,40.42972863103896,40.49038181398178,40.49038181408178,40.55064467764293,40.55064467774293,40.61088715787927,40.61088715797927,40.67107591297665,40.67107591307665,40.73124960361646,40.73124960371646,40.79140373188009,40.79140373198009,40.85150913269211,40.85150913279211,40.91155324043861,40.91155324053861,40.97151777786303,40.97151777796303,41.03140460124922,41.03140460134922,41.09111000858479,41.091110008684794,41.150803814146165,41.15080381424617,41.21044175270257,41.21044175280257,41.26991141198652,41.26991141208652,41.3293282028779,41.329328202977905,41.38868637803628,41.388686378136285,41.44804116190682,41.448041162006824,41.5073715285049,41.5073715286049,41.56660726032332,41.566607260423325,41.62583378211792,41.625833782217924,41.68502724778051,41.68502724788051,41.74415674304508,41.74415674314508,41.80321420021635,41.803214200316354,41.86210744766056,41.862107447760565,41.920706081435775,41.920706081535776,41.97924599238468,41.979245992484685,42.03772518143235,42.03772518153235,42.09617010062393,42.09617010072393,42.1545528700036,42.1545528701036,42.212690217236855,42.212690217336856,42.270686772478044,42.270686772578046,42.3285456057351,42.3285456058351,42.38597342415641,42.38597342425641,42.44330357348789,42.44330357358789,42.50033421855632,42.50033421865632,42.55734897811855,42.55734897821855,42.6143603820907,42.6143603821907,42.67134997472737,42.67134997482737,42.728176035894556,42.72817603599456,42.78460731545923,42.784607315559235,42.841027207436326,42.84102720753633,42.89738198668686,42.89738198678686,42.953564631306115,42.95356463140612,43.00962779550006,43.00962779560006,43.06538103168307,43.06538103178307,43.120845365839756,43.12084536593976,43.23173130392634,43.23173130402634,43.28712806221057,43.28712806231057,43.342418619638714,43.342418619738716,43.39770253728224,43.397702537382244,43.45293911968706,43.45293911978706,43.50810012991967,43.50810013001967,43.56322447854582,43.56322447864582,43.6183482560077,43.6183482561077,43.673420093218965,43.67342009331897,43.72845252009573,43.72845252019573,43.78335022360078,43.78335022370078,43.838082610748,43.838082610848005,43.89280939334585,43.89280939344585,43.947331913322344,43.947331913422346,44.00184483059958,44.00184483069958,44.056069060037096,44.0560690601371,44.11025373634906,44.11025373644906,44.16433938705605,44.164339387156055,44.218423895434505,44.21842389553451,44.27240805739063,44.27240805749063,44.32629794154475,44.326297941644754,44.3800828028691,44.3800828029691,44.433797018313044,44.433797018413046,44.4874585081555,44.4874585082555,44.54107694649125,44.541076946591254,44.594637947120304,44.594637947220306,44.648148935178135,44.64814893527814,44.70154190641909,44.70154190651909,44.75493059392803,44.75493059402803,44.80816938651438,44.80816938661438,44.86126388872753,44.86126388882753,44.914290315299475,44.91429031539948,44.96731534965852,44.96731534975852,45.02020426843295,45.02020426853295,45.07307080470762,45.07307080480762,45.125915101273606,45.12591510137361,45.17875772004455,45.17875772014455,45.23156531930632,45.23156531940632,45.28425572279984,45.28425572289984,45.33681472281395,45.33681472291395,45.38934287995759,45.38934288005759,45.44186157719303,45.44186157729303,45.49435696378679,45.494356963886794,45.5468455321071,45.5468455322071,45.59922390142145,45.599223901521455,45.65158445755019,45.651584457650195,45.70394030157372,45.70394030167372,45.75625223734414,45.756252237444144,45.808460613893224,45.808460613993226,45.86032954037827,45.86032954047827,45.912064243260325,45.91206424336033,45.9636373423523,45.9636373424523,46.01497801328496,46.01497801338496,46.06615329646426,46.06615329656426,46.117262039006334,46.117262039106336,46.16833047876975,46.16833047886975,46.21907381897135,46.219073819071355,46.269816373822096,46.2698163739221,46.32048546266886,46.320485462768865,46.37108740401637,46.37108740411637,46.42166917612567,46.42166917622567,46.47222363944339,46.472223639543394,46.522764787494125,46.52276478759413,46.57325299575677,46.57325299585677,46.6237389550601,46.6237389551601,46.6741840404205,46.6741840405205,46.72456836818189,46.72456836828189,46.77492356656562,46.77492356666562,46.82524917150071,46.82524917160071,46.87547328768492,46.87547328778492,46.92568947896491,46.925689479064914,46.97581349861116,46.97581349871116,47.025765633510495,47.0257656336105,47.0754376837319,47.0754376838319,47.12506111359499,47.12506111369499,47.17457734306453,47.17457734316453,47.22404127530577,47.224041275405774,47.27331340344627,47.27331340354627,47.32247765293563,47.322477653035634,47.371582858318796,47.3715828584188,47.4206637535228,47.420663753622804,47.46956280430298,47.46956280440298,47.51840563110052,47.51840563120052,47.56706543544793,47.56706543554793,47.615584519198805,47.61558451929881,47.66405815969261,47.66405815979261,47.7125159146802,47.712515914780205,47.76095964044547,47.76095964054547,47.809387159424624,47.809387159524626,47.85771918687772,47.857719186977725,47.90601997878492,47.90601997888492,47.9543167011467,47.954316701246704,48.00258440122411,48.002584401324114,48.05084564000575,48.05084564010575,48.09909220700524,48.099092207105244,48.147306146245924,48.147306146345926,48.195510696973955,48.195510697073956,48.24370850082407,48.24370850092407,48.2918541145392,48.2918541146392,48.3399475738171,48.3399475739171,48.3880134387213,48.388013438821304,48.4360486035461,48.4360486036461,48.484040110002155,48.48404011010216,48.53196561128749,48.53196561138749,48.5798506670031,48.5798506671031,48.62764783481696,48.627647834916964,48.675379747113205,48.675379747213206,48.72309363203724,48.72309363213724,48.770785669928024,48.770785670028026,48.81842633873246,48.81842633883246,48.86600264446343,48.86600264456343,48.913565456438576,48.91356545653858,48.96110306579039,48.96110306589039,49.008590876757594,49.008590876857596,49.05577493542751,49.05577493552751,49.10290223464828,49.102902234748285,49.14998194874598,49.14998194884598,49.19697359645311,49.196973596553114,49.243960175077355,49.24396017517736,49.29094104205892,49.29094104215892,49.337868612274725,49.33786861237473,49.38473228348805,49.38473228358805,49.431587530028416,49.43158753012842,49.478423892450174,49.478423892550175,49.52509090466646,49.525090904766465,49.57154744284999,49.57154744294999,49.61798388319083,49.61798388329083,49.66432890155102,49.664328901651025,49.71061555406258,49.71061555416258,49.756869685908626,49.75686968600863,49.8030805163636,49.8030805164636,49.849282065399215,49.84928206549922,49.89547447580655,49.89547447590655,49.94164254033696,49.94164254043696,49.98774481388324,49.98774481398324,50.03383120192332,50.03383120202332,50.079897777702854,50.079897777802856,50.12595364415237,50.12595364425237,50.17194404089765,50.17194404099765,50.217916981434996,50.217916981535,50.26383398357184,50.263833983671844,50.309713467355834,50.309713467455836,50.35557917180186,50.35557917190186,50.40144294856848,50.40144294866848,50.44721837336239,50.44721837346239,50.49293546800543,50.49293546810543,50.53860586996956,50.53860587006956,50.58426784726073,50.584267847360735,50.629921649763325,50.62992164986333,50.675571489813805,50.67557148991381,50.721185774888596,50.7211857749886,50.76679913182146,50.76679913192146,50.8123631187429,50.8123631188429,50.85790765038147,50.85790765048147,50.903361366901414,50.903361367001416,50.948769925746426,50.94876992584643,50.99416784665694,50.994167846756945,51.0395562719615,51.0395562720615,51.08494048492959,51.08494048502959,51.130302672375585,51.13030267247559,51.22100680663384,51.22100680673384,51.266323122449556,51.26632312254956,51.31139330216353,51.31139330226353,51.35644520462093,51.356445204720934,51.40146001709867,51.40146001719867,51.44647225933721,51.446472259437215,51.491453444518676,51.49145344461868,51.53636048543859,51.536360485538594,51.58126020831633,51.58126020841633,51.626139654862556,51.62613965496256,51.71582601009296,51.71582601019296,51.76052432617069,51.76052432627069,51.80522035759134,51.80522035769134,51.84990235978966,51.84990235988966,51.894497616414775,51.894497616514776,51.93900245059671,51.93900245069671,51.983346073664,51.983346073764004,52.02765721176354,52.02765721186354,52.07196438741098,52.07196438751098,52.11623233121275,52.11623233131275,52.16043805380708,52.16043805390708,52.2046279979885,52.204627998088505,52.248784743246844,52.248784743346846,52.29291375134042,52.29291375144042,52.33675671322902,52.336756713329024,52.38054002414938,52.380540024249385,52.467979776127066,52.46797977622707,52.51161604794618,52.51161604804618,52.5552418960174,52.5552418961174,52.598827369914424,52.598827370014426,52.64238028744817,52.64238028754817,52.68588908254221,52.68588908264221,52.729373888737,52.729373888837,52.772759990606716,52.77275999070672,52.81609211745311,52.81609211755311,52.85924368499527,52.85924368509527,52.90232921166895,52.90232921176895,52.94541306054758,52.94541306064758,52.98832962969122,52.988329629791224,53.031244878017496,53.0312448781175,53.074130925570564,53.074130925670566,53.11696206995837,53.11696207005837,53.15973299221366,53.159732992313664,53.202476641375156,53.20247664147516,53.24517727472772,53.245177274827725,53.28786673467929,53.28786673477929,53.33051881906907,53.33051881916907,53.372999518480675,53.37299951858068,53.415474541947376,53.41547454204738,53.45794056957685,53.45794056967685,53.500352336600855,53.50035233670086,53.54275068126457,53.54275068136457,53.585143706961034,53.585143707061036,53.627526130420776,53.62752613052078,53.66983976428399,53.669839764383994,53.712152648494104,53.712152648594106,53.75443708158411,53.75443708168411,53.79669541960663,53.79669541970663,53.83891820265345,53.838918202753455,53.88108261985165,53.88108261995165,53.923243395877634,53.923243395977636,53.965401958642076,53.96540195874208,54.00753174900652,54.00753174910652,54.04965311469801,54.04965311479801,54.09165696334136,54.09165696344136,54.133623257934076,54.13362325803408,54.17548738551836,54.17548738561836,54.217334628058964,54.217334628158966,54.259052216310735,54.25905221641074,54.300719684897985,54.300719684997986,54.342369090415254,54.342369090515255,54.38389433909977,54.38389433919977,54.425399704128196,54.4253997042282,54.46690239182412,54.46690239192412,54.50837727095975,54.50837727105975,54.54981673791076,54.54981673801076,54.591177883961514,54.591177884061516,54.632452926998866,54.63245292709887,54.67371904559453,54.67371904569453,54.714952108058185,54.71495210815819,54.75613840644739,54.75613840654739,54.797220003286725,54.79722000338673,54.83826204700049,54.83826204710049,54.87921784490979,54.87921784500979,54.92015593672677,54.920155936826774,54.96102991535468,54.96102991545468,55.00185613037067,55.00185613047067,55.04258424792362,55.04258424802362,55.083292838798165,55.08329283889817,55.12396937307817,55.12396937317817,55.1646298790609,55.1646298791609,55.20525682914288,55.205256829242884,55.24579681946506,55.245796819565065,55.28628072859568,55.28628072869568,55.32674439709254,55.32674439719254,55.366989273976984,55.366989274076985,55.407210376148775,55.40721037624878,55.44736450931014,55.44736450941014,55.487515679556864,55.487515679656866,55.52763486460458,55.52763486470458,55.56775037278232,55.56775037288232,55.60769717331439,55.607697173414394,55.64750432418293,55.647504324282934,55.68728709347678,55.687287093576785,55.726993005478825,55.72699300557883,55.76669520491312,55.76669520501312,55.80634899817569,55.806348998275695,55.84593407323728,55.84593407333728,55.88548641344675,55.88548641354675,55.925026545019996,55.92502654512,55.96455557458778,55.96455557468778,56.00404637184737,56.004046371947375,56.04342779114964,56.04342779124964,56.08279614506928,56.08279614516928,56.122064616637545,56.122064616737546,56.161298140092164,56.161298140192166,56.200506889296655,56.20050688939666,56.23962875013687,56.239628750236875,56.27865818945897,56.278658189558975,56.317673492465445,56.31767349256545,56.356574562618306,56.35657456271831,56.39543493731708,56.39543493741708,56.43424412141832,56.43424412151832,56.47300657714289,56.47300657724289,56.511756645742395,56.5117566458424,56.55041143700244,56.55041143710244,56.58897991106248,56.58897991116248,56.627470635386544,56.627470635486546,56.665794758233176,56.66579475833318,56.70411117036218,56.704111170462184,56.74239873869566,56.742398738795664,56.78049903654476,56.78049903664476,56.81846007740577,56.81846007750577,56.85637131988215,56.85637131998215,56.894272531286084,56.894272531386086,56.93213079827661,56.93213079837661,56.96993366233314,56.96993366243314,57.00770040024972,57.00770040034972,57.04545728558267,57.04545728568267,57.08315837530619,57.08315837540619,57.12077893086792,57.12077893096792,57.15839770154131,57.15839770164131,57.19592654954026,57.19592654964026,57.23344433123152,57.23344433133152,57.27096043512773,57.270960435227735,57.3084696136572,57.308469613757204,57.34593827522141,57.34593827532141,57.383386946036225,57.38338694613623,57.42082851299547,57.42082851309547,57.45819275859193,57.45819275869193,57.49554725869806,57.49554725879806,57.53284142957838,57.532841429678385,57.57000680291627,57.57000680301627,57.60715022212761,57.60715022222761,57.64420632460146,57.644206324701464,57.68117218312098,57.68117218322098,57.71808774348714,57.71808774358714,57.75494022589945,57.754940225999455,57.79175483298124,57.79175483308124,57.82855430420992,57.828554304309925,57.90208145845836,57.902081458558364,57.938794398300445,57.93879439840045,57.9753261362785,57.9753261363785,58.011781695222304,58.011781695322306,58.048191096953694,58.048191097053696,58.08448041139774,58.08448041149774,58.12074937811474,58.12074937821474,58.15697090249972,58.156970902599724,58.19317454230357,58.19317454240357,58.2293767898945,58.229376789994504,58.265428607096354,58.265428607196355,58.3014092429513,58.3014092430513,58.337389236246445,58.337389236346446,58.373225367541586,58.37322536764159,58.40897621687195,58.40897621697195,58.444642069819686,58.44464206991969,58.48016259715898,58.48016259725898,58.515665953872464,58.515665953972466,58.55114503610455,58.55114503620455,58.5866193348359,58.586619334935904,58.62198186385955,58.621981863959554,58.65732565155566,58.65732565165566,58.692569271318256,58.69256927141826,58.72774335183123,58.72774335193123,58.76284307389606,58.76284307399606,58.79788803558669,58.79788803568669,58.83289979835424,58.83289979845424,58.86789103490591,58.86789103500591,58.90282990283376,58.90282990293376,58.937749529665325,58.93774952976533,58.97250666026264,58.97250666036264,59.00723573241529,59.00723573251529,59.041937174496475,59.04193717459648,59.07654940785854,59.076549407958545,59.110912328017626,59.11091232811763,59.14523969320103,59.14523969330103,59.17954560402661,59.17954560412661,59.213831952476006,59.21383195257601,59.24809349097752,59.24809349107752,59.28235424412816,59.28235424422816,59.31661046366242,59.31661046376242,59.35083773230785,59.350837732407854,59.38504008391208,59.38504008401208,59.41909336164237,59.41909336174237,59.45303362024311,59.45303362034311,59.486930898732695,59.486930898832696,59.52082425046793,59.520824250567934,59.554559318305394,59.554559318405396,59.58814906053441,59.588149060634414,59.62173558996442,59.621735590064425,59.655321512532396,59.6553215126324,59.68885727973809,59.68885727983809,59.72234638996263,59.72234639006263,59.75583093087304,59.75583093097304,59.78930080000131,59.78930080010131,59.82274275347597,59.82274275357597,59.85608346808413,59.85608346818413,59.889394732034724,59.889394732134726,59.922681757201694,59.922681757301696,59.95596553387188,59.955965533971884,59.98924702588501,59.98924702598501,60.022518308336835,60.02251830843684,60.05568381830578,60.05568381840578,60.08879485348267,60.08879485358267,60.1218552692263,60.1218552693263,60.154833901386304,60.154833901486306,60.1877242886669,60.1877242887669,60.22060882151374,60.220608821613745,60.253484037243474,60.253484037343476,60.28626136969677,60.28626136979677,60.31900507485378,60.31900507495378,60.35173335857556,60.35173335867556,60.3844441860894,60.3844441861894,60.417101466953106,60.41710146705311,60.44970016778157,60.44970016788157,60.48227520099069,60.482275201090694,60.51475410011394,60.51475410021394,60.547201835086824,60.547201835186826,60.57960808925474,60.57960808935474,60.611981929850444,60.611981929950446,60.644336100976666,60.64433610107667,60.67663608289296,60.676636082992964,60.70888326781223,60.708883267912235,60.741113032221335,60.74111303232134,60.77332323425425,60.77332323435425,60.80552804592439,60.80552804602439,60.83771643662182,60.83771643672182,60.869899901027445,60.86989990112745,60.90199651276656,60.90199651286656,60.93402954678309,60.93402954688309,60.965995933069095,60.9659959331691,60.9979207671546,60.997920767254605,61.02984021087733,61.02984021097733,61.06158469984521,61.061584699945215,61.09325718641756,61.09325718651756,61.12492396134722,61.12492396144722,61.15657306588234,61.15657306598234,61.18819860989141,61.18819860999141,61.21979859429948,61.219798594399485,61.251336607384474,61.251336607484475,61.28286865894242,61.28286865904242,61.314376364623435,61.31437636472344,61.345721038603706,61.34572103870371,61.377040581356184,61.377040581456185,61.408343845927014,61.408343846027016,61.439522382500826,61.43952238260083,61.47062031351731,61.47062031361731,61.50166523335017,61.50166523345017,61.53263315310014,61.532633153200145,61.56357990407443,61.56357990417443,61.59447314409636,61.594473144196364,61.62534053893517,61.62534053903517,61.656204578183896,61.6562045782839,61.68706540463362,61.68706540473362,61.717905776262995,61.717905776363,61.74867582329186,61.748675823391864,61.77943655320362,61.77943655330362,61.84091674640875,61.84091674650875,61.87162785642472,61.871627856524725,61.90228727607443,61.902287276174434,61.93290225200454,61.93290225210454,61.96344618937881,61.96344618947881,61.99395068072082,61.993950680820824,62.024428113155636,62.02442811325564,62.05482026362568,62.05482026372568,62.085189888804905,62.08518988890491,62.11544206833152,62.11544206843152,62.14568257468841,62.14568257478841,62.17592108197035,62.17592108207035,62.206102829803164,62.206102829903166,62.23626897771191,62.23626897781191,62.26639218120725,62.26639218130725,62.29649517976661,62.29649517986661,62.32654366783614,62.32654366793614,62.356561884199465,62.35656188429947,62.386574246129044,62.386574246229046,62.41654955377674,62.41654955387674,62.44650180066711,62.44650180076711,62.47643377122596,62.476433771325965,62.50634000369499,62.50634000379499,62.53620893199776,62.536208932097765,62.566057726760086,62.56605772686009,62.59589388451298,62.595893884612984,62.625579754667854,62.625579754767855,62.655265017960694,62.655265018060696,62.684784750709106,62.68478475080911,62.714297022624265,62.71429702272427,62.7437820571434,62.7437820572434,62.77325288395136,62.77325288405136,62.8027007570953,62.8027007571953,62.83214730942187,62.83214730952187,62.861541528822386,62.86154152892239,62.890924717612975,62.89092471771298,62.9202930561326,62.9202930562326,62.949642224951475,62.94964222505148,62.97890950309342,62.97890950319342,63.008162644919736,63.00816264501974,63.03730740832619,63.037307408426194,63.06644763811627,63.066447638216275,63.095507333744564,63.095507333844566,63.12456199598774,63.12456199608774,63.15360652006516,63.15360652016516,63.18261930882794,63.18261930892794,63.2116265287391,63.211626528839105,63.24057438326416,63.24057438336416,63.269517382892936,63.26951738299294,63.29840904913313,63.29840904923313,63.32725877049739,63.32725877059739,63.35607240141946,63.35607240151946,63.38484319502143,63.384843195121434,63.41357486387105,63.41357486397105,63.44225159385763,63.44225159395763,63.470856178657606,63.47085617875761,63.49938190709083,63.49938190719083,63.527854445851595,63.5278544459516,63.55622064096521,63.55622064106521,63.58454796121083,63.58454796131083,63.612820806664395,63.6128208067644,63.64102921764896,63.641029217748965,63.66923480850996,63.669234808609964,63.697406664981386,63.69740666508139,63.72555456825131,63.72555456835131,63.7536964029009,63.7536964030009,63.7818198532006,63.781819853300604,63.809914531100304,63.809914531200306,63.83800456829033,63.83800456839033,63.866046056517575,63.86604605661758,63.89403810333787,63.89403810343787,63.92202011908571,63.92202011918571,63.94986105725935,63.94986105735935,63.977673972686084,63.977673972786086,64.00547500075649,64.00547500085649,64.03322980021896,64.03322980031896,64.06088967931963,64.06088967941963,64.08850022410665,64.08850022420665,64.11608567336366,64.11608567346366,64.14365844991345,64.14365845001345,64.17121202106475,64.17121202116475,64.1987534906731,64.1987534907731,64.22628903445218,64.22628903455218,64.25379880444366,64.25379880454366,64.28128451414037,64.28128451424037,64.30875983578693,64.30875983588693,64.33622858904442,64.33622858914443,64.36362851700761,64.36362851710761,64.39085595336186,64.39085595346187,64.41805236835681,64.41805236845681,64.44520790940882,64.44520790950882,64.47233881900178,64.47233881910178,64.49940997053318,64.49940997063318,64.52636821002837,64.52636821012837,64.5533209163697,64.5533209164697,64.58026159256363,64.58026159266363,64.60711723667717,64.60711723677717,64.63389498826366,64.63389498836366,64.66062315565212,64.66062315575212,64.68735064478302,64.68735064488303,64.71406646074419,64.7140664608442,64.74072391085673,64.74072391095673,64.7673003627365,64.76730036283651,64.79386307097607,64.79386307107607,64.8201534409531,64.82015344105311,64.84640900560755,64.84640900570756,64.87257903841287,64.87257903851287,64.89866975078046,64.89866975088046,64.92462127755549,64.92462127765549,64.95048231049181,64.95048231059181,64.9763180337115,64.9763180338115,65.00213772863391,65.00213772873391,65.02787103496078,65.02787103506078,65.05354108484498,65.05354108494498,65.07913620511677,65.07913620521677,65.10466999662526,65.10466999672526,65.1300083428608,65.1300083429608,65.1553387998899,65.1553387999899,65.18064973024057,65.18064973034058,65.20593859937141,65.20593859947141,65.23119184213101,65.23119184223101,65.25643994441221,65.25643994451221,65.28163789133112,65.28163789143112,65.30681381272794,65.30681381282794,65.33192794129052,65.33192794139052,65.35702661272009,65.35702661282009,65.38207909123949,65.38207909133949,65.40706945564474,65.40706945574475,65.43201394841972,65.43201394851972,65.4568623428066,65.4568623429066,65.48161114042425,65.48161114052425,65.50634273171832,65.50634273181832,65.53107371615036,65.53107371625036,65.5557991317308,65.5557991318308,65.58048245964422,65.58048245974422,65.60513558724696,65.60513558734696,65.62972092479066,65.62972092489066,65.65425603557654,65.65425603567654,65.67873924180955,65.67873924190955,65.70315765659592,65.70315765669592,65.72756803938476,65.72756803948477,65.75197710135623,65.75197710145623,65.77634125553713,65.77634125563713,65.80063023022126,65.80063023032126,65.8248819364369,65.8248819365369,65.84905664256965,65.84905664266965,65.87318197869098,65.87318197879098,65.89728946592895,65.89728946602895,65.92128960998228,65.92128961008228,65.94526901363314,65.94526901373314,65.96924349099218,65.96924349109219,65.99303023379589,65.9930302338959,66.01680790936683,66.01680790946683,66.04057541107426,66.04057541117426,66.06434016405365,66.06434016415365,66.08805311957347,66.08805311967348,66.11174397817568,66.11174397827568,66.13543016037043,66.13543016047043,66.15910663277265,66.15910663287265,66.18278242691729,66.18278242701729,66.20640924372594,66.20640924382595,66.22999722136437,66.22999722146437,66.2535294747889,66.25352947488891,66.27701018063824,66.27701018073824,66.30045965094168,66.30045965104168,66.32383465570368,66.32383465580368,66.3471953456612,66.34719534576121,66.37049888349416,66.37049888359417,66.39377661184176,66.39377661194176,66.41695531458438,66.41695531468439,66.4400596231811,66.4400596232811,66.46308582506417,66.46308582516417,66.48608396850257,66.48608396860257,66.50905976513899,66.50905976523899,66.53199215329103,66.53199215339103,66.55485985708971,66.55485985718971,66.57772299157426,66.57772299167426,66.60052444031787,66.60052444041787,66.64599857596039,66.64599857606039,66.66854321496999,66.66854321506999,66.69104326746891,66.69104326756892,66.71347078210579,66.71347078220579,66.73589597638784,66.73589597648784,66.75824852571454,66.75824852581454,66.78058861652065,66.78058861662065,66.80291589182849,66.80291589192849,66.82523431409018,66.82523431419018,66.84754941645957,66.84754941655957,66.86985755776443,66.86985755786444,66.89212196930502,66.89212196940503,66.91437506465357,66.91437506475357,66.93662105614652,66.93662105624652,66.9588499484092,66.9588499485092,66.9810367173071,66.98103671740711,67.00316515605415,67.00316515615415,67.02527681685082,67.02527681695082,67.04733814379637,67.04733814389637,67.06939468724117,67.06939468734117,67.09137187455049,67.09137187465049,67.11334406417245,67.11334406427245,67.13518210158698,67.13518210168698,67.15695791779405,67.15695791789405,67.17867893792915,67.17867893802915,67.20039417502605,67.20039417512605,67.22210584234627,67.22210584244627,67.24373244188833,67.24373244198833,67.26534858198471,67.26534858208471,67.30855623071845,67.30855623081845,67.3301133624064,67.3301133625064,67.3516200888477,67.3516200889477,67.37310993024533,67.37310993034534,67.39456460934271,67.39456460944271,67.41584133507283,67.41584133517283,67.43710335332305,67.43710335342305,67.45836412215144,67.45836412225144,67.47960265127114,67.47960265137114,67.50082836489257,67.50082836499257,67.52203490881327,67.52203490891327,67.54323531271807,67.54323531281807,67.56440680143182,67.56440680153182,67.5854817276865,67.5854817277865,67.60651096079972,67.60651096089973,67.62753391110601,67.62753391120602,67.6485509712808,67.6485509713808,67.6695493258258,67.6695493259258,67.69050248699808,67.69050248709809,67.71142873205424,67.71142873215425,67.73233455798783,67.73233455808783,67.75322464120626,67.75322464130626,67.77410626405398,67.77410626415399,67.81580097003726,67.81580097013726,67.83658442402641,67.83658442412641,67.87809771395906,67.87809771405907,67.89881626940827,67.89881626950827,67.91951540527236,67.91951540537237,67.94015874552701,67.94015874562702,67.96077756141591,67.96077756151591,67.98135607452615,67.98135607462615,68.00188386110983,68.00188386120983,68.02240225918086,68.02240225928087,68.04290762756702,68.04290762766702,68.06333563889264,68.06333563899264,68.0837304155974,68.0837304156974,68.10406293539695,68.10406293549696,68.12437703514887,68.12437703524887,68.14461110050772,68.14461110060772,68.16484388074697,68.16484388084697,68.20527263682796,68.20527263692796,68.2455432875,68.2455432876,68.26560832393328,68.26560832403328,68.28565119205342,68.28565119215342,68.30563073233533,68.30563073243533,68.32560823784455,68.32560823794455,68.34556775167933,68.34556775177933,68.36551784130367,68.36551784140367,68.38545147425755,68.38545147435755,68.40533538022234,68.40533538032234,68.42517805526653,68.42517805536653,68.44501451889931,68.44501451899932,68.46483963064226,68.46483963074226,68.48465231956239,68.48465231966239,68.50445354949937,68.50445354959938,68.52425188791727,68.52425188801728,68.54404033805376,68.54404033815376,68.56381593699423,68.56381593709423,68.5835647983074,68.58356479840741,68.6032710007893,68.60327100088931,68.62295000157295,68.62295000167295,68.64256517474963,68.64256517484964,68.66214943366032,68.66214943376032,68.6816789321968,68.6816789322968,68.70119979187373,68.70119979197374,68.72071365478838,68.72071365488839,68.74019528261965,68.74019528271965,68.75963835686284,68.75963835696284,68.77896166509856,68.77896166519857,68.79826262653232,68.79826262663232,68.8174596003715,68.81745960047151,68.8365952811452,68.8365952812452,68.85572871295957,68.85572871305958,68.87477993281713,68.87477993291714,68.893826511965,68.893826512065,68.91287009250047,68.91287009260047,68.93189393217092,68.93189393227092,68.95091209589647,68.95091209599647,68.96992154936693,68.96992154946693,68.9889126184875,68.9889126185875,69.00785139037978,69.00785139047979,69.02672376442591,69.02672376452591,69.04558128820108,69.04558128830108,69.06443238637823,69.06443238647823,69.08324535934048,69.08324535944048,69.10205683299652,69.10205683309653,69.12085552685208,69.12085552695208,69.13961895131409,69.13961895141409,69.1583172987473,69.15831729884731,69.17695695905198,69.17695695915198,69.1954902757095,69.1954902758095,69.21399042914173,69.21399042924173,69.23243571510645,69.23243571520645,69.25084012712823,69.25084012722823,69.26921997908649,69.26921997918649,69.28757930482887,69.28757930492887,69.30585002871418,69.30585002881418,69.32406706315828,69.32406706325828,69.36047125301573,69.36047125311573,69.37862371019979,69.37862371029979,69.39676210246375,69.39676210256376,69.41486704592026,69.41486704602026,69.432909054214,69.432909054314,69.45093906805809,69.45093906815809,69.46894780603321,69.46894780613322,69.48694779805547,69.48694779815547,69.50492983410106,69.50492983420106,69.52290690815707,69.52290690825707,69.54087263032325,69.54087263042325,69.55880758101449,69.5588075811145,69.5766874857494,69.5766874858494,69.59453972471509,69.59453972481509,69.61238646622469,69.6123864663247,69.63022603248316,69.63022603258317,69.64802147630195,69.64802147640195,69.66577151256142,69.66577151266142,69.68350230772461,69.68350230782461,69.7011778070471,69.7011778071471,69.71883470783312,69.71883470793313,69.73646065865536,69.73646065875536,69.75405423160316,69.75405423170317,69.7716025754805,69.7716025755805,69.78912439591713,69.78912439601713,69.80664450286095,69.80664450296095,69.8241633960807,69.82416339618071,69.8415959364027,69.8415959365027,69.85901373354702,69.85901373364702,69.8764048287618,69.87640482886181,69.893656131522,69.893656131622,69.9108908705184,69.9108908706184,69.92810668969845,69.92810668979845,69.94531472676533,69.94531472686533,69.96249156398409,69.96249156408409,69.97961992363558,69.97961992373558,69.99668463416896,69.99668463426896,70.01373920653658,70.01373920663659,70.03071567219055,70.03071567229055,70.04766832743411,70.04766832753411,70.06456914952034,70.06456914962034,70.09834330641239,70.09834330651239,70.11510365778645,70.11510365788645,70.13181663822404,70.13181663832404,70.14851205536037,70.14851205546037,70.1651053768838,70.1651053769838,70.1816749950901,70.18167499519011,70.19824289980362,70.19824289990362,70.21475757914689,70.21475757924689,70.23125319588273,70.23125319598273,70.24771593497539,70.24771593507539,70.26416767915588,70.26416767925588,70.28061142703662,70.28061142713662,70.2970130515526,70.2970130516526,70.31333021515245,70.31333021525245,70.32963406348529,70.32963406358529,70.34589418205387,70.34589418215387,70.3621515518944,70.3621515519944,70.37833124339448,70.37833124349449,70.39449483520177,70.39449483530177,70.4105942781222,70.4105942782222,70.42661836305702,70.42661836315702,70.44261824490597,70.44261824500597,70.45854947994947,70.45854948004947,70.47447935847784,70.47447935857784,70.49039131672728,70.49039131682729,70.50623969725414,70.50623969735415,70.5220811167165,70.52208111681651,70.53790568683294,70.53790568693294,70.55372647298611,70.55372647308612,70.5695433680827,70.5695433681827,70.58535094606218,70.58535094616218,70.60114492319252,70.60114492329252,70.61691259106875,70.61691259116876,70.63263909941993,70.63263909951993,70.64833362257211,70.64833362267211,70.66402682490691,70.66402682500691,70.67959779808835,70.67959779818835,70.69512696918493,70.69512696928493,70.71053544613213,70.71053544623213,70.72591515067931,70.72591515077931,70.74129149963643,70.74129149973643,70.75664657272456,70.75664657282456,70.77188045189457,70.77188045199458,70.78709116321396,70.78709116331396,70.80227253096909,70.8022725310691,70.81745222092917,70.81745222102917,70.83262584226891,70.83262584236891,70.84775823268804,70.84775823278804,70.8628852327444,70.8628852328444,70.87791299300919,70.87791299310919,70.89293871850128,70.89293871860129,70.90793053111496,70.90793053121496,70.92287365197477,70.92287365207477,70.93775865687033,70.93775865697033,70.95263045359218,70.95263045369218,70.96748193828475,70.96748193838475,70.98232017910587,70.98232017920587,70.99708438275873,70.99708438285873,71.01183659196197,71.01183659206197,71.02655899352997,71.02655899362998,71.04124869594362,71.04124869604362,71.05593700614438,71.05593700624438,71.07061956900468,71.07061956910468,71.08529617033794,71.08529617043794,71.09997120096946,71.09997120106947,71.11461446058858,71.11461446068859,71.12923273177097,71.12923273187097,71.14384586247495,71.14384586257495,71.15844535663201,71.15844535673202,71.17303956751961,71.17303956761961,71.20206902713946,71.20206902723946,71.21649292398189,71.21649292408189,71.23087459036626,71.23087459046626,71.24524558311835,71.24524558321835,71.25956492120197,71.25956492130197,71.27384649104836,71.27384649114836,71.28808158240244,71.28808158250244,71.3022832963446,71.3022832964446,71.31641729162324,71.31641729172324,71.33052122938228,71.33052122948229,71.34459460985298,71.34459460995298,71.35865631715394,71.35865631725395,71.37269878335864,71.37269878345865,71.38672318649336,71.38672318659336,71.40073452424544,71.40073452434544,71.41473136870422,71.41473136880423,71.42866195810791,71.42866195820791,71.44255817056222,71.44255817066222,71.4703058661691,71.4703058662691,71.48410241045865,71.48410241055865,71.49778647108515,71.49778647118515,71.51145671667592,71.51145671677592,71.5251108625739,71.5251108626739,71.53874723098403,71.53874723108403,71.55237403239266,71.55237403249266,71.56597673780875,71.56597673790876,71.57953981870374,71.57953981880374,71.59309897284439,71.59309897294439,71.60659533461332,71.60659533471332,71.62008020170136,71.62008020180136,71.63352830008964,71.63352830018964,71.64695448004913,71.64695448014913,71.66037648336992,71.66037648346992,71.67379309632793,71.67379309642793,71.68717472547452,71.68717472557452,71.70055510519927,71.70055510529927,71.71392256233247,71.71392256243247,71.72720181028401,71.72720181038402,71.74046745738643,71.74046745748643,71.75369301589679,71.75369301599679,71.76690636577091,71.76690636587091,71.78007241610409,71.78007241620409,71.79318945340353,71.79318945350353,71.80629895847417,71.80629895857417,71.81939029338155,71.81939029348155,71.83241890731273,71.83241890741273,71.84543299225285,71.84543299235285,71.85840106277163,71.85840106287164,71.8712917405321,71.8712917406321,71.88416685406625,71.88416685416625,71.89698242372548,71.89698242382548,71.90979413802589,71.90979413812589,71.92259599974267,71.92259599984267,71.93536966022373,71.93536966032373,71.94810515979212,71.94810515989212,71.96083998110295,71.96083998120295,71.9735581315567,71.9735581316567,71.98627260514047,71.98627260524047,71.9989493461848,71.9989493462848,72.01160402600927,72.01160402610927,72.02425738501637,72.02425738511637,72.03690703145573,72.03690703155573,72.04952522816258,72.04952522826258,72.06210633488978,72.06210633498978,72.07467741054451,72.07467741064451,72.08714846105681,72.08714846115681,72.09960912351899,72.09960912361899,72.11203419530771,72.11203419540772,72.12445009277039,72.12445009287039,72.13683007827971,72.13683007837972,72.14915062700739,72.14915062710739,72.16144315298817,72.16144315308817,72.17373107395704,72.17373107405705,72.18600789292044,72.18600789302045,72.19827086115036,72.19827086125036,72.21053161611873,72.21053161621873,72.22279030061662,72.22279030071662,72.23502935134282,72.23502935144282,72.24723238302235,72.24723238312235,72.25942952457036,72.25942952467037,72.27160638978687,72.27160638988687,72.28375701714482,72.28375701724482,72.29590675205858,72.29590675215859,72.30803942343985,72.30803942353985,72.32011669188712,72.32011669198712,72.33209382809866,72.33209382819867,72.34402420023575,72.34402420033575,72.35593918663537,72.35593918673537,72.36784014381266,72.36784014391266,72.3797271788609,72.3797271789609,72.39160489679203,72.39160489689203,72.4034814009991,72.4034814010991,72.4272172387874,72.42721723888741,72.43905940174285,72.43905940184285,72.45088700000947,72.45088700010947,72.46267829364729,72.46267829374729,72.47444277826229,72.47444277836229,72.48619180574428,72.48619180584429,72.49790745581436,72.49790745591436,72.50954996116036,72.50954996126036,72.5211892180096,72.5211892181096,72.53274130091242,72.53274130101242,72.54428117517902,72.54428117527902,72.55580569940591,72.55580569950591,72.56732826025564,72.56732826035564,72.57881687252917,72.57881687262918,72.59027539158535,72.59027539168535,72.60173262552192,72.60173262562192,72.61318793177908,72.61318793187908,72.62461532238264,72.62461532248264,72.6360258636403,72.6360258637403,72.64742444614609,72.64742444624609,72.65878875879581,72.65878875889581,72.67010384422518,72.67010384432518,72.6814166092997,72.6814166093997,72.69270456442634,72.69270456452634,72.70398641523485,72.70398641533485,72.7265241288777,72.7265241289777,72.7377188057398,72.7377188058398,72.74885918629869,72.74885918639869,72.75998864334095,72.75998864344095,72.7711128528115,72.7711128529115,72.78222410399272,72.78222410409272,72.79333032178882,72.79333032188882,72.8044270082812,72.8044270083812,72.81552230256067,72.81552230266067,72.82661209938406,72.82661209948407,72.83769650584469,72.83769650594469,72.8487623494666,72.8487623495666,72.85981577026567,72.85981577036567,72.87085080671487,72.87085080681487,72.88187459836755,72.88187459846755,72.89289025092941,72.89289025102941,72.90390447558059,72.90390447568059,72.91490916892805,72.91490916902805,72.92590843621497,72.92590843631497,72.93690042115747,72.93690042125748,72.94788630178185,72.94788630188185,72.95885133491045,72.95885133501045,72.96981344082218,72.96981344092218,72.98077101311753,72.98077101321753,72.99171205734687,72.99171205744688,73.00263635932362,73.00263635942362,73.01354534695842,73.01354534705843,73.02441049773566,73.02441049783566,73.03525058868063,73.03525058878063,73.04608768101319,73.04608768111319,73.06772941640834,73.06772941650834,73.07850178868979,73.0785017887898,73.08922175664965,73.08922175674965,73.09992144827798,73.09992144837798,73.11061503558821,73.11061503568821,73.12130573137934,73.12130573147934,73.13196947535656,73.13196947545656,73.14262354523899,73.14262354533899,73.15324748620628,73.15324748630628,73.16387060612495,73.16387060622495,73.17446266898652,73.17446266908652,73.18505030532503,73.18505030542504,73.19561966440696,73.19561966450696,73.20617552973306,73.20617552983306,73.21670801302194,73.21670801312194,73.2272226831252,73.22722268322521,73.23769929940913,73.23769929950913,73.24813254290643,73.24813254300643,73.25855853975709,73.25855853985709,73.2689634392276,73.2689634393276,73.2793645904326,73.27936459053261,73.2897548895165,73.2897548896165,73.30013669253192,73.30013669263192,73.3105101779677,73.3105101780677,73.32086217334788,73.32086217344788,73.3311994255504,73.3311994256504,73.34150155115043,73.34150155125043,73.35177972354897,73.35177972364897,73.36203008738721,73.36203008748721,73.37227120550386,73.37227120560387,73.38251021745228,73.38251021755228,73.39274380334015,73.39274380344015,73.40297253433174,73.40297253443174,73.4234036156653,73.4234036157653,73.4438023191244,73.4438023192244,73.45399745851749,73.4539974586175,73.46416489644356,73.46416489654356,73.47430830977262,73.47430830987263,73.48444190621582,73.48444190631582,73.4945499787558,73.4945499788558,73.50464173741636,73.50464173751637,73.51472764164318,73.51472764174318,73.52480547817471,73.52480547827471,73.53486932118169,73.53486932128169,73.54489207605913,73.54489207615913,73.55489619670232,73.55489619680232,73.5648949626805,73.5648949627805,73.58488414135944,73.58488414145944,73.59487348312719,73.59487348322719,73.60482341456046,73.60482341466046,73.61476645632474,73.61476645642475,73.62470400063295,73.62470400073295,73.63463915319078,73.63463915329078,73.6445699149233,73.6445699150233,73.65448065020867,73.65448065030867,73.66437924825334,73.66437924835334,73.67427452640571,73.67427452650571,73.68415820278388,73.68415820288388,73.69403977299382,73.69403977309382,73.70391784482261,73.70391784492261,73.71377617578639,73.7137761758864,73.72361369495216,73.72361369505217,73.74328494932041,73.74328494942041,73.7531051550693,73.75310515516931,73.76292207662365,73.76292207672365,73.77273660642763,73.77273660652763,73.78254242597652,73.78254242607652,73.79233742910209,73.79233742920209,73.80211693939688,73.80211693949688,73.81188027860334,73.81188027870334,73.82163940547332,73.82163940557332,73.8313968188505,73.8313968189505,73.84113902497903,73.84113902507903,73.85087459132295,73.85087459142295,73.86054661564206,73.86054661574207,73.87021471320682,73.87021471330682,73.8798751714495,73.8798751715495,73.88952506315323,73.88952506325323,73.89915464282767,73.89915464292767,73.90877840376612,73.90877840386612,73.91838806408671,73.91838806418671,73.92799162008919,73.92799162018919,73.93757821965247,73.93757821975247,73.94714675614576,73.94714675624576,73.95662694066634,73.95662694076634,73.96610162773085,73.96610162783085,73.9755711029214,73.97557110302141,73.9850389360147,73.9850389361147,74.00396960451393,74.00396960461393,74.01339767029506,74.01339767039507,74.02281870361614,74.02281870371614,74.03220975081315,74.03220975091315,74.04158337749998,74.04158337759998,74.05095436255208,74.05095436265208,74.06032274166719,74.0603227417672,74.06967851947066,74.06967851957066,74.07903158424384,74.07903158434384,74.08837601015746,74.08837601025746,74.0977067995242,74.0977067996242,74.10703498295396,74.10703498305396,74.11636234533508,74.11636234543508,74.13501549939559,74.1350154994956,74.1443222641653,74.1443222642653,74.15361853379159,74.15361853389159,74.16290420118114,74.16290420128114,74.17217291213149,74.17217291223149,74.1814208469816,74.1814208470816,74.19066221344261,74.19066221354261,74.19985595908278,74.19985595918278,74.20904731297259,74.20904731307259,74.21822088937455,74.21822088947455,74.2273909316976,74.2273909317976,74.24572787494023,74.24572787504023,74.25489374062457,74.25489374072457,74.26405132442702,74.26405132452702,74.27320184007166,74.27320184017167,74.28234175347957,74.28234175357957,74.29148106002545,74.29148106012545,74.30061422655545,74.30061422665545,74.30973918259909,74.30973918269909,74.31884885999857,74.31884886009857,74.3279561099499,74.3279561100499,74.33706161071066,74.33706161081066,74.34615561679054,74.34615561689054,74.35524094831308,74.35524094841308,74.36432017551752,74.36432017561752,74.37338758676117,74.37338758686117,74.38244835822019,74.38244835832019,74.39149624278542,74.39149624288542,74.40051224924494,74.40051224934494,74.40951840312083,74.40951840322083,74.41848511096447,74.41848511106447,74.42744296576195,74.42744296586196,74.43639942834653,74.43639942844653,74.44533143796089,74.44533143806089,74.45424766916233,74.45424766926233,74.46316243675534,74.46316243685534,74.47206977921287,74.47206977931287,74.48097319491606,74.48097319501606,74.48987164862967,74.48987164872968,74.49875914312888,74.49875914322888,74.50764335343356,74.50764335353357,74.51652352989059,74.5165235299906,74.52540295669452,74.52540295679452,74.53427763569547,74.53427763579548,74.54312971801006,74.54312971811007,74.55196666447156,74.55196666457157,74.56078697577375,74.56078697587375,74.56960221799307,74.56960221809307,74.57840757193098,74.57840757203098,74.58720821376369,74.58720821386369,74.59600271558052,74.59600271568053,74.60479414738941,74.60479414748941,74.61358183093279,74.61358183103279,74.62232571331636,74.62232571341636,74.63980558887707,74.63980558897707,74.64854104658768,74.64854104668768,74.65727589743626,74.65727589753627,74.66600600048187,74.66600600058187,74.67469954901433,74.67469954911434,74.69208386165344,74.69208386175345,74.70076288119483,74.70076288129484,74.70943925910147,74.70943925920147,74.71809229066866,74.71809229076867,74.72673825407804,74.72673825417804,74.73537714932961,74.73537714942961,74.74401315306206,74.74401315316206,74.75264155317019,74.7526415532702,74.76125538858949,74.76125538868949,74.76986015677603,74.76986015687604,74.77846010576407,74.77846010586407,74.78705063054169,74.78705063064169,74.79561705932674,74.79561705942675,74.80417909728648,74.80417909738648,74.81273963594002,74.81273963604002,74.82126722555485,74.82126722565485,74.82979042434437,74.82979042444437,74.83829027679444,74.83829027689444,74.84678231143359,74.84678231153359,74.85527306095312,74.85527306105313,74.8637630251218,74.8637630252218,74.8722492053272,74.8722492054272,74.88071585885417,74.88071585895418,74.88917008955833,74.88917008965834,74.89761682373147,74.89761682383147,74.90605195613041,74.90605195623041,74.91448651736509,74.9144865174651,74.92291022647868,74.92291022657868,74.93132301207564,74.93132301217564,74.93972441008499,74.939724410185,74.95652070911018,74.95652070921018,74.96491075522971,74.96491075532971,74.97329769564354,74.97329769574354,74.98167856743703,74.98167856753703,74.99005822550644,74.99005822560645,74.99841014641109,74.99841014651109,75.00675517764675,75.00675517774675,75.01509203409383,75.01509203419383,75.02341618213595,75.02341618223595,75.03170188368327,75.03170188378327,75.03997966032637,75.03997966042637,75.04825390289056,75.04825390299057,75.05652628917089,75.05652628927089,75.0647919285733,75.0647919286733,75.07305210621739,75.07305210631739,75.08130735756968,75.08130735766969,75.08956171647779,75.08956171657779,75.09781279119137,75.09781279129137,75.106058654031,75.106058654131,75.11429955488104,75.11429955498105,75.12250432959115,75.12250432969115,75.1389112373766,75.1389112374766,75.14711251370557,75.14711251380557,75.15530072465188,75.15530072475188,75.16348365232874,75.16348365242874,75.17164355494603,75.17164355504603,75.1798007802308,75.17980078033081,75.18795572085853,75.18795572095853,75.19610862671352,75.19610862681353,75.20423890018442,75.20423890028442,75.21236631783397,75.21236631793397,75.22861872568492,75.22861872578493,75.23672376083472,75.23672376093472,75.24482501202124,75.24482501212124,75.25290780746235,75.25290780756235,75.26098949627269,75.26098949637269,75.26904548269097,75.26904548279097,75.27709204489882,75.27709204499882,75.28513778605804,75.28513778615805,75.2931822063999,75.2931822064999,75.3092675130047,75.3092675131047,75.31730622170386,75.31730622180386,75.32531590811867,75.32531590821867,75.33332145359253,75.33332145369253,75.34130975704505,75.34130975714505,75.34929588293379,75.34929588303379,75.35726355307713,75.35726355317713,75.36522097796141,75.36522097806142,75.37316301710823,75.37316301720823,75.38109131261483,75.38109131271483,75.38901639532243,75.38901639542243,75.39692437879977,75.39692437889977,75.40480180498875,75.40480180508875,75.41266802207898,75.41266802217898,75.42052331565259,75.42052331575259,75.42837489665844,75.42837489675844,75.43621848136455,75.43621848146455,75.44404921487464,75.44404921497464,75.45187188068942,75.45187188078943,75.45967908937122,75.45967908947122,75.46747241162174,75.46747241172174,75.48305448680864,75.48305448690864,75.49083477937431,75.49083477947431,75.49861385821589,75.49861385831589,75.50637937190612,75.50637937200612,75.51413078497848,75.51413078507848,75.52187141732527,75.52187141742527,75.52959734219218,75.52959734229218,75.53731869774492,75.53731869784492,75.5450346629349,75.5450346630349,75.55274809358343,75.55274809368343,75.56045295676795,75.56045295686795,75.56815403598918,75.56815403608918,75.57583487457669,75.57583487467669,75.58351482072,75.58351482082,75.59118594951494,75.59118594961494,75.59885354423096,75.59885354433096,75.60650839484425,75.60650839494426,75.61414186249527,75.61414186259528,75.62175691009864,75.62175691019864,75.62936706710796,75.62936706720797,75.64457213818021,75.64457213828021,75.65216380374635,75.65216380384635,75.65975314895766,75.65975314905766,75.66734092346722,75.66734092356722,75.6748861819366,75.6748861820366,75.68242472922581,75.68242472932582,75.68995410218898,75.68995410228898,75.6974780490916,75.6974780491916,75.70499799784434,75.70499799794435,75.71250459563232,75.71250459573233,75.71999698570913,75.71999698580913,75.72748530624054,75.72748530634054,75.73496559477442,75.73496559487442,75.74242999780209,75.7424299979021,75.7498875111608,75.7498875112608,75.75733367262966,75.75733367272966,75.76476894627967,75.76476894637968,75.77219918654457,75.77219918664457,75.77962264423378,75.77962264433378,75.78704453122126,75.78704453132126,75.79446481180923,75.79446481190924,75.80187495423145,75.80187495433145,75.81668977731756,75.81668977741757,75.82408974587626,75.82408974597627,75.8314697950811,75.8314697951811,75.83884416834104,75.83884416844104,75.84621247298062,75.84621247308063,75.85357595842169,75.85357595852169,75.86093869420966,75.86093869430967,75.86829982359814,75.86829982369814,75.87565627657915,75.87565627667915,75.88300559000682,75.88300559010682,75.89033162849057,75.89033162859057,75.90497742265113,75.90497742275113,75.91229389413338,75.91229389423339,75.91960768828314,75.91960768838314,75.92691205822247,75.92691205832247,75.93419693718116,75.93419693728116,75.94147492647086,75.94147492657086,75.95602237328401,75.95602237338402,75.96327822995833,75.96327823005834,75.97051909357062,75.97051909367062,75.97775849357446,75.97775849367446,75.98498757692371,75.98498757702372,75.99220695048041,75.99220695058041,75.9994230041448,75.9994230042448,76.00663013336751,76.00663013346751,76.01383476374653,76.01383476384653,76.02103121933698,76.02103121943698,76.0282172868773,76.0282172869773,76.03539135996799,76.03539136006799,76.04256247014403,76.04256247024404,76.0497105909583,76.0497105910583,76.05685632002219,76.0568563201222,76.06399230359575,76.06399230369576,76.07112211145568,76.07112211155568,76.07825045570716,76.07825045580717,76.08537812170107,76.08537812180107,76.09249393603642,76.09249393613642,76.09960560943084,76.09960560953084,76.10671664026545,76.10671664036545,76.11381492699732,76.11381492709732,76.12091114325872,76.12091114335873,76.12800668126256,76.12800668136256,76.13509097446988,76.13509097456988,76.14217259034469,76.14217259044469,76.14925317098425,76.14925317108425,76.15633275208636,76.15633275218636,76.16338645230755,76.16338645240755,76.1704387246181,76.1704387247181,76.17748728436088,76.17748728446088,76.18452135081037,76.18452135091037,76.19154417246334,76.19154417256334,76.19855732002152,76.19855732012152,76.20555711661493,76.20555711671493,76.21254734620685,76.21254734630685,76.21953293508909,76.2195329351891,76.22651381186613,76.22651381196613,76.24047063912838,76.24047063922838,76.24744501891186,76.24744501901186,76.25441732822487,76.25441732832488,76.26138503252598,76.26138503262598,76.26834663250897,76.26834663260897,76.27529748746416,76.27529748756416,76.28224427287395,76.28224427297395,76.28916881857505,76.28916881867505,76.29608961601063,76.29608961611063,76.30300548715442,76.30300548725442,76.30991900224559,76.3099190023456,76.316831410706,76.316831410806,76.32373775054606,76.32373775064606,76.3306407704938,76.3306407705938,76.33754143438894,76.33754143448894,76.34443677931684,76.34443677941684,76.35132766202389,76.3513276621239,76.35821447518553,76.35821447528554,76.36509372042063,76.36509372052063,76.37196107829939,76.3719610783994,76.37882736524516,76.37882736534516,76.38569286684005,76.38569286694005,76.3925559409868,76.3925559410868,76.39941433872612,76.39941433882612,76.40627084448379,76.40627084458379,76.41311867568413,76.41311867578413,76.4199439458959,76.4199439459959,76.42676782389476,76.42676782399477,76.43358827490802,76.43358827500802,76.44722607122881,76.44722607132881,76.45404045362172,76.45404045372172,76.46085412205929,76.46085412215929,76.46766586281746,76.46766586291746,76.48128673839682,76.48128673849682,76.48807809573016,76.48807809583016,76.49486781096624,76.49486781106624,76.50165538433632,76.50165538443632,76.50843674629498,76.50843674639498,76.51521693022734,76.51521693032734,76.52199543636466,76.52199543646466,76.52876612469106,76.52876612479106,76.5355354922001,76.5355354923001,76.54229036641583,76.54229036651583,76.54904366992982,76.54904367002982,76.55579008377482,76.55579008387483,76.56252875120445,76.56252875130446,76.56926631200331,76.56926631210331,76.58273565056281,76.58273565066281,76.58944411768177,76.58944411778177,76.59615069281908,76.59615069291908,76.6028563041167,76.6028563042167,76.60955866691754,76.60955866701754,76.61625431853822,76.61625431863823,76.62294886352815,76.62294886362815,76.62964180211857,76.62964180221857,76.63632278195712,76.63632278205712,76.64300279795597,76.64300279805597,76.64968224279056,76.64968224289056,76.65636083087874,76.65636083097874,76.66303806245179,76.66303806255179,76.67638417232047,76.67638417242047,76.68971960855689,76.68971960865689,76.70303576799925,76.70303576809926,76.7096725183048,76.7096725184048,76.71630498487835,76.71630498497835,76.72293613063451,76.72293613073451,76.72956527731574,76.72956527741574,76.73618757002576,76.73618757012576,76.74279868933479,76.74279868943479,76.74940284757929,76.74940284767929,76.75600350744266,76.75600350754266,76.76260177555565,76.76260177565565,76.76919683086965,76.76919683096965,76.77578931594442,76.77578931604442,76.7823805158996,76.7823805159996,76.79556205906356,76.79556205916356,76.80215118854825,76.80215118864825,76.80873896151782,76.80873896161782,76.81531909516531,76.81531909526531,76.82189533775622,76.82189533785622,76.82846933138782,76.82846933148782,76.83503832733207,76.83503832743207,76.84159986244308,76.84159986254308,76.84815739940422,76.84815739950422,76.85471140228644,76.85471140238644,76.86126476260885,76.86126476270886,76.86780987674716,76.86780987684716,76.87434870807851,76.87434870817852,76.88088211334932,76.88088211344932,76.88740552324543,76.88740552334544,76.89392768371971,76.89392768381971,76.90044927302972,76.90044927312972,76.90696322301766,76.90696322311766,76.91347503113957,76.91347503123957,76.91998523286199,76.91998523296199,76.93299646198078,76.93299646208078,76.939495061929,76.939495062029,76.94599287652635,76.94599287662635,76.95247530538624,76.95247530548625,76.95895073748385,76.95895073758385,76.96542395631991,76.96542395641991,76.97189635410733,76.97189635420733,76.97836596746896,76.97836596756896,76.98483108291197,76.98483108301197,76.991293592418,76.991293592518,76.9977404663022,76.9977404664022,77.00418377040974,77.00418377050974,77.010626503353,77.010626503453,77.01706845094539,77.01706845104539,77.02350690015663,77.02350690025663,77.02994331459517,77.02994331469517,77.03636984075233,77.03636984085233,77.04278815642313,77.04278815652313,77.04920475860112,77.04920475870112,77.05561950449525,77.05561950459526,77.06844417708498,77.06844417718499,77.07485071249276,77.07485071259276,77.08125581998986,77.08125582008986,77.0876598208562,77.0876598209562,77.09406285788283,77.09406285798283,77.10046228943501,77.10046228953502,77.11325619054982,77.11325619064982,77.11964673335808,77.11964673345808,77.12603556267355,77.12603556277355,77.13241964418604,77.13241964428605,77.13879680033178,77.13879680043178,77.15154918494385,77.15154918504385,77.16429903501447,77.16429903511447,77.17704420867766,77.17704420877766,77.1834111552621,77.18341115536211,77.19613887271734,77.19613887281734,77.20249332508342,77.20249332518343,77.2088452072103,77.2088452073103,77.22154697238912,77.22154697248912,77.22789503485495,77.22789503495495,77.23424141952574,77.23424141962575,77.2405869831479,77.24058698324791,77.24693179711697,77.24693179721697,77.25326497361407,77.25326497371407,77.25959600824517,77.25959600834517,77.26592047448717,77.26592047458718,77.27224079978825,77.27224079988825,77.2785591974099,77.2785591975099,77.28487609572537,77.28487609582537,77.29118910298425,77.29118910308425,77.2974998969816,77.29749989708161,77.30379898211145,77.30379898221145,77.31009556839763,77.31009556849763,77.31638758536967,77.31638758546967,77.32267149894865,77.32267149904865,77.3352364345875,77.3352364346875,77.3415137797774,77.3415137798774,77.34777859505118,77.34777859515118,77.35403684193587,77.35403684203587,77.36029091218184,77.36029091228184,77.36654062730028,77.36654062740028,77.37903102600215,77.37903102610215,77.38526938923077,77.38526938933077,77.39150511082464,77.39150511092464,77.39773926171677,77.39773926181677,77.4039718062094,77.4039718063094,77.41020360104892,77.41020360114892,77.4226664053771,77.4226664054771,77.43512731772364,77.43512731782364,77.44135547139096,77.44135547149097,77.44758023377044,77.44758023387044,77.46002708119688,77.46002708129689,77.4662427763436,77.46624277644361,77.47245140333251,77.47245140343252,77.47865763857104,77.47865763867104,77.4848609465927,77.4848609466927,77.49106032786001,77.49106032796001,77.49725728167918,77.49725728177918,77.5034513082815,77.5034513083815,77.50963855230812,77.50963855240812,77.51582322609552,77.51582322619552,77.5220062220879,77.5220062221879,77.52818721900535,77.52818721910535,77.53436493172823,77.53436493182824,77.54054071677173,77.54054071687173,77.55288900266417,77.55288900276418,77.55905946874043,77.55905946884043,77.56522893527921,77.56522893537921,77.57139004854056,77.57139004864057,77.57754905563368,77.57754905573368,77.58370670621166,77.58370670631166,77.58986142957278,77.58986142967278,77.5960122618773,77.59601226197731,77.60215859626322,77.60215859636322,77.60830228901442,77.60830228911442,77.62058767544184,77.62058767554184,77.62672101584067,77.62672101594067,77.63285242856008,77.63285242866009,77.63898302023088,77.63898302033088,77.64511250527089,77.64511250537089,77.65124134775111,77.65124134785111,77.65736926208939,77.65736926218939,77.66349314258002,77.66349314268003,77.6696161306265,77.6696161307265,77.67573454935881,77.67573454945881,77.68184914843009,77.68184914853009,77.6879630692438,77.68796306934381,77.69407609761335,77.69407609771335,77.70018548481069,77.70018548491069,77.70628998141397,77.70628998151398,77.7123927288267,77.7123927289267,77.71849215634711,77.71849215644711,77.72459094130772,77.72459094140773,77.73068908370853,77.73068908380853,77.74288411908832,77.74288411918832,77.7611734415101,77.7611734416101,77.76726487273075,77.76726487283075,77.77335551860054,77.77335551870054,77.79162581411265,77.79162581421265,77.80380192967606,77.80380192977606,77.8098831870332,77.8098831871332,77.81596091031142,77.81596091041142,77.8220338500889,77.8220338501889,77.82810432672048,77.82810432682048,77.83417305416148,77.83417305426148,77.8402356415866,77.8402356416866,77.84629426655962,77.84629426665963,77.8523468229123,77.8523468230123,77.85839345343568,77.85839345353568,77.86443712104442,77.86443712114442,77.87047768294747,77.87047768304747,77.87651617438003,77.87651617448003,77.88255216696892,77.88255216706892,77.8946209393477,77.8946209394477,77.9006536120443,77.9006536121443,77.90668339322177,77.90668339332177,77.91271163939528,77.91271163949528,77.91873885033355,77.91873885043356,77.92476434777903,77.92476434787903,77.93078623975006,77.93078623985006,77.93680688229925,77.93680688239925,77.94282545437797,77.94282545447797,77.9548616346957,77.9548616347957,77.9668908896467,77.96689088974671,77.97290157251896,77.97290157261897,77.98492008244216,77.98492008254216,77.99092458960078,77.99092458970078,77.996923956281,77.996923956381,78.00891779904737,78.00891779914737,78.02089861212887,78.02089861222888,78.02688427086665,78.02688427096665,78.03286893006695,78.03286893016696,78.03884980530398,78.03884980540398,78.05080969949418,78.05080969959418,78.05678750472327,78.05678750482328,78.06873779621421,78.06873779631421,78.07471003259168,78.07471003269168,78.08068012710315,78.08068012720315,78.0985846989949,78.0985846990949,78.10454790383739,78.10454790393739,78.11050521854835,78.11050521864836,78.11646114104643,78.11646114114643,78.12241153039065,78.12241153049065,78.12836084880188,78.12836084890188,78.13430859651136,78.13430859661136,78.1462011290157,78.1462011291157,78.1521438076423,78.1521438077423,78.158081881257,78.158081881357,78.16401477869552,78.16401477879552,78.16994699787647,78.16994699797647,78.1818097941411,78.1818097942411,78.1877378723811,78.18773787248111,78.19959128013308,78.19959128023308,78.211440868224,78.211440868324,78.21736087876873,78.21736087886873,78.22328010396258,78.22328010406258,78.22919176122988,78.22919176132989,78.23510238326195,78.23510238336195,78.24101132749897,78.24101132759897,78.24691509555981,78.24691509565982,78.25872084679318,78.25872084689318,78.26462272287239,78.26462272297239,78.27642476153801,78.27642476163801,78.28232413877355,78.28232413887355,78.28821912518377,78.28821912528377,78.29411275507887,78.29411275517887,78.305895124275,78.305895124375,78.31178261415421,78.31178261425421,78.31766696262994,78.31766696272994,78.32355038296375,78.32355038306375,78.32943258957349,78.32943258967349,78.33531165477974,78.33531165487975,78.3411892206798,78.3411892207798,78.3470643591317,78.34706435923171,78.35293753420645,78.35293753430645,78.35880724659782,78.35880724669782,78.36467535258969,78.36467535268969,78.37054003159594,78.37054003169594,78.37640338978483,78.37640338988483,78.3822658912273,78.38226589132731,78.38812600091941,78.38812600101942,78.39398471839863,78.39398471849863,78.39983868807485,78.39983868817485,78.40569187240023,78.40569187250023,78.41739617058047,78.41739617068048,78.42324664187556,78.42324664197557,78.42909632781978,78.42909632791978,78.44079141597621,78.44079141607621,78.45832444551161,78.45832444561161,78.46416595666724,78.46416595676725,78.470006682472,78.470006682572,78.47584644443707,78.47584644453707,78.48168288650983,78.48168288660983,78.49334955924392,78.49334955934393,78.50501016335767,78.50501016345767,78.51083079131976,78.51083079141976,78.51664970578904,78.51664970588904,78.52828185878269,78.52828185888269,78.53409477602715,78.53409477612715,78.53990601547659,78.53990601557659,78.54571500596671,78.54571500606671,78.55152278273276,78.55152278283276,78.55732898879708,78.55732898889708,78.56893833091777,78.56893833101778,78.58634656106061,78.58634656116061,78.59214662710905,78.59214662720905,78.59794433710486,78.59794433720486,78.60953561615558,78.60953561625558,78.6153289353261,78.6153289354261,78.62691382447657,78.62691382457658,78.63849664315657,78.63849664325657,78.64428664243479,78.6442866425348,78.65007414286933,78.65007414296933,78.65585700259419,78.65585700269419,78.66742168680868,78.66742168690868,78.6732014765256,78.67320147662561,78.68475969944431,78.68475969954432,78.69053627636222,78.69053627646223,78.6963069988464,78.6963069989464,78.70207454422932,78.70207454432932,78.71360785010683,78.71360785020683,78.71937186141085,78.71937186151085,78.73089859889929,78.73089859899929,78.73665846926235,78.73665846936235,78.7481770319622,78.7481770320622,78.75393418929501,78.75393418939501,78.759688562202,78.759688562302,78.76544182847822,78.76544182857822,78.7769468260267,78.7769468261267,78.78269480903344,78.78269480913345,78.79418956132288,78.79418956142288,78.79993376036634,78.79993376046635,78.80567413974877,78.80567413984878,78.81141351959373,78.81141351969373,78.81714718779601,78.81714718789601,78.82861159698369,78.8286115970837,78.84007150825278,78.84007150835278,78.8515278140474,78.8515278141474,78.8572542713008,78.8572542714008,78.87443110851954,78.87443110861955,78.88015349622754,78.88015349632754,78.89731866027655,78.89731866037656,78.90303847774533,78.90303847784533,78.90875750986324,78.90875750996324,78.91447582802581,78.91447582812582,78.920193289442,78.920193289542,78.9316252493597,78.9316252494597,78.93733799867066,78.93733799877066,78.94304582168981,78.94304582178981,78.9487518955184,78.9487518956184,78.95445518492117,78.95445518502117,78.9601571178088,78.9601571179088,78.97155969846447,78.97155969856448,78.97725970367271,78.97725970377272,78.9829579239926,78.9829579240926,78.9886541095398,78.9886541096398,78.99434454774654,78.99434454784654,79.00003405781135,79.00003405791135,79.01141193561243,79.01141193571243,79.0227857438681,79.0227857439681,79.0284690425215,79.0284690426215,79.03414966384237,79.03414966394237,79.03982685817766,79.03982685827766,79.04549616330648,79.04549616340648,79.05116489727104,79.05116489737104,79.056832346116,79.056832346216,79.06249918809891,79.06249918819891,79.07383055170992,79.07383055180992,79.07949300286754,79.07949300296754,79.09081240772667,79.09081240782668,79.09646918293937,79.09646918303937,79.10212435175256,79.10212435185257,79.11343315437497,79.11343315447498,79.11908664539314,79.11908664549314,79.12473642384354,79.12473642394355,79.13603433864711,79.13603433874711,79.14168211802259,79.14168211812259,79.14732697018118,79.14732697028118,79.15297021594029,79.15297021604029,79.16425274500637,79.16425274510637,79.16989038621608,79.16989038631608,79.1755260997464,79.17552609984641,79.1867965986651,79.1867965987651,79.19806499141555,79.19806499151555,79.20933067113573,79.20933067123573,79.21495717464222,79.21495717474222,79.22058017976757,79.22058017986757,79.22620193547108,79.22620193557108,79.23181837220736,79.23181837230736,79.24304846125408,79.24304846135408,79.24866147100474,79.24866147110474,79.25427019702339,79.25427019712339,79.27109476867984,79.27109476877985,79.28230690174989,79.28230690184989,79.28790963054372,79.28790963064372,79.29351171677776,79.29351171687776,79.29911105428376,79.29911105438376,79.30470750027064,79.30470750037064,79.31589939270695,79.31589939280695,79.32148816367399,79.32148816377399,79.32707622068568,79.32707622078568,79.33266334955545,79.33266334965545,79.33824765830164,79.33824765840164,79.3438305034394,79.34383050353941,79.36057786082637,79.36057786092637,79.37173976713859,79.3717397672386,79.37731857842871,79.37731857852872,79.38289585471485,79.38289585481485,79.3884694898288,79.3884694899288,79.39403969795711,79.39403969805711,79.39960915643233,79.39960915653234,79.40517390280235,79.40517390290235,79.41073800661256,79.41073800671256,79.4162986477394,79.4162986478394,79.42185650444043,79.42185650454043,79.42741054148041,79.42741054158041,79.43851711625418,79.43851711635418,79.44406761921526,79.44406761931526,79.44961630159024,79.44961630169024,79.45516234233047,79.45516234243047,79.47179910803604,79.47179910813604,79.4773425428393,79.4773425429393,79.48288240786589,79.48288240796589,79.48842145184383,79.48842145194384,79.49949600572083,79.49949600582083,79.51609935831672,79.51609935841672,79.52163140553239,79.52163140563239,79.52716095390439,79.52716095400439,79.5326892885523,79.5326892886523,79.53821565982307,79.53821565992307,79.54374031760102,79.54374031770102,79.54926269072189,79.54926269082189,79.55478388581648,79.55478388591648,79.56030411707134,79.56030411717134,79.56582331309099,79.56582331319099,79.57685974175308,79.57685974185308,79.58237626044021,79.58237626054022,79.58789181528765,79.58789181538765,79.59340569234004,79.59340569244004,79.5989182128773,79.5989182129773,79.60442930550388,79.60442930560389,79.6099398269662,79.6099398270662,79.61544806377144,79.61544806387144,79.62095387312856,79.62095387322856,79.63196381404772,79.63196381414772,79.65398080436697,79.65398080446697,79.65948311534294,79.65948311544294,79.67598876315122,79.67598876325123,79.68148853958574,79.68148853968574,79.6869835682173,79.6869835683173,79.69247442021013,79.69247442031013,79.69796259487046,79.69796259497046,79.70893765907152,79.70893765917153,79.72539754234283,79.72539754244283,79.73088132617787,79.73088132627787,79.73636353931114,79.73636353941114,79.74184193278339,79.74184193288339,79.74731797020303,79.74731797030303,79.75826736770979,79.7582673678098,79.76373883581527,79.76373883591528,79.76920873321903,79.76920873331903,79.77467730980541,79.77467730990541,79.78014510104093,79.78014510114093,79.78561032203724,79.78561032213725,79.79107008127524,79.79107008137524,79.80198852881823,79.80198852891823,79.80744686014556,79.80744686024556,79.8183576326687,79.8183576327687,79.82380896723375,79.82380896733375,79.82925833842161,79.82925833852161,79.85105307444506,79.85105307454506,79.8564998039982,79.8564998040982,79.86739165670497,79.86739165680497,79.87283592311219,79.8728359232122,79.87827815474671,79.87827815484671,79.88371742346659,79.8837174235666,79.88915565695123,79.88915565705123,79.89459324787609,79.89459324797609,79.90546589518434,79.90546589528434,79.91089966644813,79.91089966654813,79.91633265236106,79.91633265246107,79.92176089047102,79.92176089057102,79.92718702241274,79.92718702251274,79.938037322919,79.938037323019,79.94345874275551,79.94345874285551,79.94887887747241,79.94887887757241,79.95429615636797,79.95429615646798,79.96513000020377,79.96513000030377,79.97596006007629,79.97596006017629,79.98137298384431,79.98137298394431,79.98678033876071,79.98678033886071,79.99218633716198,79.99218633726198,80.00840194061541,80.00840194071542,80.02461404568771,80.02461404578771,80.0300167955922,80.0300167956922,80.0354177249106,80.0354177250106,80.04081790457589,80.04081790467589,80.05161733576453,80.05161733586453,80.06241155507924,80.06241155517924,80.07319970577358,80.07319970587358,80.07859206762795,80.07859206772795,80.08398318006049,80.08398318016049,80.08937279318681,80.08937279328681,80.0947593363052,80.0947593364052,80.10014391604642,80.10014391614642,80.10552742485464,80.10552742495464,80.11090889889016,80.11090889899016,80.11628808826859,80.11628808836859,80.12704207620017,80.12704207630017,80.13241423311855,80.13241423321855,80.14315658357816,80.14315658367816,80.15389586402983,80.15389586412984,80.15926323744748,80.15926323754748,80.16462953993212,80.16462954003212,80.16999502136812,80.16999502146813,80.180724627725,80.180724627825,80.18608832427267,80.18608832437268,80.19681386108414,80.19681386118414,80.2021739164596,80.2021739165596,80.20752822449461,80.20752822459461,80.21287764193558,80.21287764203558,80.21822598844355,80.21822598854355,80.22357362099618,80.22357362109618,80.2342677437729,80.2342677438729,80.23961305597071,80.23961305607071,80.26633658264949,80.26633658274949,80.28236609364485,80.28236609374486,80.28770548001337,80.28770548011337,80.29304315288907,80.29304315298907,80.29837743447695,80.29837743457695,80.30370868175463,80.30370868185463,80.30903803705068,80.30903803715069,80.31969189274652,80.31969189284652,80.33034121482598,80.33034121492598,80.34098714561759,80.34098714571759,80.35162897116602,80.35162897126602,80.35694586794148,80.35694586804148,80.37820981387111,80.37820981397111,80.38883339786074,80.38883339796074,80.39414194135877,80.39414194145877,80.39944869996847,80.39944870006848,80.40475310252558,80.40475310262558,80.41535801658318,80.41535801668319,80.42065849238594,80.42065849248594,80.42595807574453,80.42595807584453,80.4365556360622,80.4365556361622,80.44185254208827,80.44185254218827,80.44714823439028,80.44714823449029,80.45244267727045,80.45244267737046,80.46303067058662,80.46303067068662,80.46832379264943,80.46832379274943,80.47361191702488,80.47361191712488,80.47889529359736,80.47889529369736,80.48417585004627,80.48417585014627,80.48945576393537,80.48945576403537,80.49473307188751,80.49473307198751,80.50528279719772,80.50528279729772,80.51582577563003,80.51582577573004,80.52109462321144,80.52109462331144,80.52636275683751,80.52636275693752,80.5368956684996,80.5368956685996,80.54215891153163,80.54215891163163,80.55268261316988,80.55268261326988,80.5579417866565,80.5579417867565,80.56845874141685,80.56845874151685,80.5737159515263,80.5737159516263,80.57897259047148,80.57897259057148,80.59474140067626,80.59474140077626,80.59999332751623,80.59999332761623,80.60524389784106,80.60524389794107,80.61049243339319,80.61049243349319,80.61573964812796,80.61573964822796,80.62098511367215,80.62098511377215,80.62622593850666,80.62622593860667,80.63670562479851,80.63670562489851,80.64194312974071,80.64194312984071,80.65241728287872,80.65241728297872,80.65765264595493,80.65765264605493,80.66288440355667,80.66288440365668,80.66811555429639,80.6681155543964,80.67334345653934,80.67334345663934,80.6785703949426,80.6785703950426,80.68379604822624,80.68379604832624,80.69424621246499,80.69424621256499,80.6994704021402,80.6994704022402,80.70991671102013,80.70991671112013,80.7151385803405,80.7151385804405,80.72557810664476,80.72557810674476,80.73079544234875,80.73079544244875,80.73601167142196,80.73601167152196,80.74122725793538,80.74122725803538,80.75165382595031,80.75165382605032,80.75686188023495,80.75686188033495,80.76206911347094,80.76206911357094,80.77248043853947,80.77248043863948,80.77768370932336,80.77768370942336,80.78288505242784,80.78288505252785,80.78808568157699,80.78808568167699,80.79328541828197,80.79328541838197,80.80368314250137,80.80368314260137,80.8088769176793,80.8088769177793,80.81406612354309,80.81406612364309,80.81925179532797,80.81925179542797,80.82962228215133,80.82962228225134,80.84517504947173,80.84517504957174,80.85035618764023,80.85035618774023,80.85553507684943,80.85553507694944,80.86071310931223,80.86071310941223,80.86588985665543,80.86588985675543,80.87624231610657,80.87624231620657,80.88658924240389,80.88658924250389,80.89175699390985,80.89175699400985,80.90725810656177,80.90725810666177,80.91242314503747,80.91242314513747,80.92275022337645,80.92275022347646,80.9330717685616,80.9330717686616,80.93823031004375,80.93823031014375,80.95886244119964,80.95886244129964,80.96401719871852,80.96401719881852,80.97432546433444,80.97432546443444,80.98978388245732,80.98978388255732,81.00008507991541,81.00008508001541,81.01553118230876,81.01553118240876,81.02582316974303,81.02582316984304,81.03611272972917,81.03611272982917,81.04125567128725,81.04125567138725,81.04639190166517,81.04639190176518,81.05666082834213,81.05666082844213,81.06179077591311,81.06179077601311,81.07204927884217,81.07204927894217,81.07717608500968,81.07717608510968,81.08742823373626,81.08742823383626,81.09255132733603,81.09255132743603,81.09767306442066,81.09767306452066,81.10278916125814,81.10278916135815,81.1079040086738,81.1079040087738,81.11813216850112,81.11813216860112,81.13347192724731,81.13347192734732,81.14880940133644,81.14880940143644,81.15391925106474,81.15391925116474,81.16413709423747,81.16413709433748,81.1692441238422,81.1692441239422,81.17434608436405,81.17434608446405,81.17944515336679,81.17944515346679,81.1845421162013,81.1845421163013,81.1947349709373,81.1947349710373,81.19983039876783,81.19983039886783,81.20492393461673,81.20492393471673,81.21001689930137,81.21001689940137,81.21510757932892,81.21510757942892,81.22019386853117,81.22019386863117,81.22527930098701,81.22527930108701,81.23036294855451,81.23036294865452,81.23544506111806,81.23544506121806,81.24052606705082,81.24052606715082,81.25068729356548,81.25068729366548,81.25576312332207,81.25576312342207,81.26083763226129,81.26083763236129,81.26591028491663,81.26591028501663,81.27098043872832,81.27098043882832,81.28111806901916,81.28111806911916,81.28618418898319,81.28618418908319,81.30138029991598,81.30138030001598,81.30644263591672,81.30644263601673,81.3165647733768,81.3165647734768,81.3317384304143,81.3317384305143,81.33679098522697,81.33679098532697,81.34184064852053,81.34184064862053,81.34688874111235,81.34688874121235,81.3519345490471,81.3519345491471,81.35697867918681,81.35697867928681,81.36202170269576,81.36202170279576,81.37210632180297,81.37210632190298,81.38218783520449,81.38218783530449,81.39226592162039,81.3922659217204,81.39730344767325,81.39730344777325,81.40234040256185,81.40234040266185,81.4124131700105,81.4124131701105,81.41744709058892,81.41744709068892,81.42247951186114,81.42247951196114,81.42750747091252,81.42750747101252,81.43253396635545,81.43253396645545,81.44258438700213,81.44258438710213,81.44760777673935,81.44760777683935,81.45263016693912,81.45263016703912,81.457650700855,81.457650700955,81.46267030662897,81.46267030672897,81.4727079831729,81.4727079832729,81.47772583975629,81.47772583985629,81.48274069772725,81.48274069782725,81.4877547346496,81.4877547347496,81.49276673679924,81.49276673689924,81.4977770968516,81.49777709695161,81.5027863502732,81.5027863503732,81.50779481834392,81.50779481844393,81.51780929133947,81.51780929143948,81.52281290451393,81.52281290461393,81.53281902423205,81.53281902433206,81.53782128089138,81.53782128099138,81.54782436629934,81.54782436639934,81.55282287469313,81.55282287479314,81.5628150008867,81.5628150009867,81.5678075120557,81.5678075121557,81.58278183276366,81.58278183286366,81.59276032241031,81.59276032251032,81.60772525460563,81.60772525470563,81.61271112599,81.61271112609,81.6176958907436,81.6176958908436,81.62766417082898,81.62766417092898,81.63264597266794,81.63264597276795,81.6426069704089,81.64260697050891,81.65256339883572,81.65256339893573,81.65753834670348,81.65753834680348,81.67246215507149,81.67246215517149,81.68738314331591,81.68738314341591,81.69732743450203,81.69732743460203,81.70727029777748,81.70727029787749,81.7122402479579,81.7122402480579,81.72217647144872,81.72217647154872,81.73210741167007,81.73210741177007,81.737071007648,81.737071007748,81.74699712867084,81.74699712877084,81.75195829720062,81.75195829730062,81.75691250466589,81.75691250476589,81.76681859924157,81.76681859934158,81.776719838921,81.776719839021,81.78166812055699,81.78166812065699,81.7866128324163,81.7866128325163,81.7915532605436,81.7915532606436,81.80143276028306,81.80143276038307,81.80637054677563,81.80637054687563,81.81130686965975,81.81130686975975,81.81624237149524,81.81624237159524,81.8310472706022,81.8310472707022,81.83597923835877,81.83597923845878,81.85077321394911,81.85077321404911,81.85570168332455,81.85570168342456,81.86062876048709,81.86062876058709,81.86555523078759,81.8655552308876,81.87048084434169,81.87048084444169,81.87540531556725,81.87540531566725,81.88032910853524,81.88032910863524,81.8852509024283,81.8852509025283,81.89017194666825,81.89017194676825,81.89508959962036,81.89508959972036,81.9000055033819,81.9000055034819,81.9098358115988,81.9098358116988,81.91474935930772,81.91474935940772,81.91966140771046,81.91966140781047,81.9245727064601,81.9245727065601,81.93439387604869,81.93439387614869,81.94421254679361,81.94421254689361,81.95893816116062,81.95893816126062,81.9638451047827,81.96384510488271,81.96875090607625,81.96875090617625,81.97365610050777,81.97365610060777,81.97855901028221,81.97855901038221,81.98346113470579,81.9834611348058,81.99326388424674,81.99326388434675,81.99816233180034,81.99816233190035,82.0030597441187,82.0030597442187,82.0177504817676,82.01775048186761,82.02754202220979,82.02754202230979,82.03243350869887,82.03243350879887,82.0373243169304,82.0373243170304,82.04221380434454,82.04221380444454,82.05199042312023,82.05199042322023,82.0568758052912,82.0568758053912,82.06175872431626,82.06175872441626,82.06664017973289,82.06664017983289,82.07152049282098,82.07152049292098,82.08128044073959,82.08128044083959,82.08615832637955,82.08615832647955,82.09590478054233,82.09590478064233,82.10077149278129,82.1007714928813,82.10563574187434,82.10563574197434,82.11049717084381,82.11049717094382,82.12021713726368,82.12021713736368,82.12993246297385,82.12993246307386,82.13478796611405,82.13478796621405,82.1396398637798,82.13963986387981,82.14449115458353,82.14449115468354,82.14934176712968,82.14934176722969,82.15904095744929,82.1590409575493,82.16388375218452,82.16388375228452,82.16872411947162,82.16872411957162,82.17356230919495,82.17356230929495,82.17839924949644,82.17839924959644,82.18807148800214,82.18807148810214,82.19290582236665,82.19290582246666,82.20257088562124,82.20257088572124,82.20740107904481,82.20740107914482,82.21222988025549,82.2122298803555,82.22188551929966,82.22188551939966,82.23636428360957,82.23636428370958,82.24118483863613,82.24118483873613,82.25082498484953,82.25082498494953,82.25564118474854,82.25564118484854,82.26527197814704,82.26527197824704,82.27008621466888,82.27008621476888,82.27489773816045,82.27489773826045,82.28451814350882,82.28451814360882,82.28932377686887,82.28932377696887,82.29412808941156,82.29412808951156,82.29893111683462,82.29893111693463,82.30373214518276,82.30373214528277,82.3085321025979,82.3085321026979,82.3133310247778,82.3133310248778,82.31812773369616,82.31812773379616,82.32292247923735,82.32292247933735,82.32771572547233,82.32771572557233,82.33729782711698,82.33729782721699,82.3468742885145,82.3468742886145,82.35166060938273,82.35166060948274,82.36601796558794,82.36601796568794,82.37080257296337,82.37080257306337,82.38036843212416,82.38036843222416,82.38992722312713,82.38992722322713,82.39470376280728,82.39470376290728,82.4042549858837,82.4042549859837,82.4090273846229,82.4090273847229,82.4185712182616,82.4185712183616,82.42334226048567,82.42334226058567,82.42811169631024,82.42811169641024,82.44241854017548,82.44241854027548,82.44718540576083,82.44718540586084,82.45671803030078,82.45671803040078,82.461481397505,82.461481397605,82.46624355098514,82.46624355108514,82.47100416946132,82.47100416956133,82.47576421677323,82.47576421687323,82.48528245511316,82.48528245521317,82.49479751635187,82.49479751645187,82.49955038841266,82.49955038851266,82.50430240372704,82.50430240382704,82.50905234857096,82.50905234867096,82.5138012938774,82.51380129397741,82.51854963232182,82.51854963242182,82.52804541676647,82.52804541686648,82.53279186322925,82.53279186332925,82.53753734585233,82.53753734595233,82.54228190033346,82.54228190043347,82.54702466992626,82.54702467002626,82.55650935236545,82.55650935246545,82.56599150026321,82.56599150036321,82.57073087856817,82.57073087866817,82.57546947152225,82.57546947162226,82.58020677935674,82.58020677945674,82.58967957443961,82.58967957453962,82.5991459082267,82.5991459083267,82.603874827086,82.603874827186,82.61332952340112,82.61332952350112,82.62278068563734,82.62278068573734,82.6275048209959,82.6275048210959,82.63222784972368,82.63222784982368,82.63694937914526,82.63694937924527,82.64166955205171,82.64166955215171,82.64638626227477,82.64638626237478,82.65110033086312,82.65110033096312,82.6558134713095,82.6558134714095,82.66052332756136,82.66052332766137,82.66523229136905,82.66523229146905,82.66993904191519,82.66993904201519,82.68405708029209,82.6840570803921,82.69817158459009,82.69817158469009,82.70757694659156,82.70757694669156,82.71227673607319,82.7122767361732,82.71697481206202,82.71697481216202,82.72636832240494,82.72636832250494,82.73106054396001,82.73106054406001,82.73575037376472,82.73575037386472,82.74043866856546,82.74043866866546,82.74981422293169,82.74981422303169,82.75918792101406,82.75918792111406,82.76387311010909,82.76387311020909,82.77324241736615,82.77324241746615,82.77792600006167,82.77792600016167,82.78260894019739,82.7826089402974,82.7872852405485,82.7872852406485,82.8059874790383,82.8059874791383,82.81533490356433,82.81533490366434,82.82000702727673,82.82000702737673,82.82935006097746,82.82935006107746,82.83401893619309,82.83401893629309,82.84335593697124,82.84335593707124,82.84802341997396,82.84802342007396,82.85268868971514,82.85268868981514,82.8573507466573,82.8573507467573,82.86201091161784,82.86201091171785,82.86666882761908,82.86666882771908,82.87132485163868,82.87132485173868,82.87597702029946,82.87597702039946,82.88527921575502,82.88527921585502,82.88992760045252,82.88992760055253,82.89922254926144,82.89922254936144,82.90386872069742,82.90386872079742,82.90851414248029,82.90851414258029,82.91315885030781,82.91315885040781,82.91780295127332,82.91780295137332,82.92244619549241,82.92244619559241,82.9270885829651,82.9270885830651,82.93172900706062,82.93172900716063,82.93636764626781,82.93636764636781,82.94564260432735,82.94564260442735,82.95491352853924,82.95491352863924,82.95954692017472,82.95954692027472,82.96417792005981,82.96417792015981,82.96880788470968,82.96880788480968,82.97343727819528,82.97343727829528,82.97806560074787,82.97806560084787,82.98269217410989,82.98269217420989,82.98731617723271,82.98731617733272,82.99193703895206,82.99193703905206,83.00117455005426,83.00117455015426,83.01040470741653,83.01040470751653,83.01501689457855,83.01501689467855,83.02424019796959,83.02424019806959,83.03346046705046,83.03346046715046,83.03806419384176,83.03806419394176,83.04266652842017,83.04266652852017,83.05187001955066,83.05187001965066,83.05646660678862,83.05646660688862,83.06565685404765,83.06565685414765,83.07024994290445,83.07024994300446,83.08402574679151,83.08402574689151,83.09320321425005,83.09320321435005,83.09778991320661,83.09778991330661,83.1069621687912,83.1069621688912,83.11613256809193,83.11613256819193,83.1207162327383,83.1207162328383,83.13446362120304,83.13446362130304,83.13904093164695,83.13904093174695,83.14361592173601,83.14361592183602,83.15733835746178,83.15733835756178,83.16648312576596,83.16648312586597,83.17562535952872,83.17562535962873,83.18476448758577,83.18476448768577,83.18933237381925,83.18933237391926,83.19389833237332,83.19389833247332,83.20302917854849,83.20302917864849,83.21215670483134,83.21215670493135,83.21671866523555,83.21671866533555,83.22127769842287,83.22127769852287,83.23039465816676,83.23039465826676,83.23495079983499,83.23495079993499,83.2440618337496,83.2440618338496,83.24861619052949,83.24861619062949,83.25316744160367,83.25316744170367,83.26681898156467,83.26681898166467,83.2713671626309,83.2713671627309,83.28500984954574,83.28500984964575,83.2941011782931,83.2941011783931,83.29864486144074,83.29864486154074,83.30773054994096,83.30773055004096,83.31227219831588,83.31227219841588,83.32589485878357,83.32589485888357,83.33043411540812,83.33043411550813,83.3349720512153,83.3349720513153,83.33950627445475,83.33950627455475,83.34403749908178,83.34403749918178,83.34856790266016,83.34856790276017,83.35309609297703,83.35309609307703,83.35762317666311,83.35762317676311,83.36214797569212,83.36214797579213,83.3666699188998,83.3666699189998,83.37119096966332,83.37119096976332,83.37571066391169,83.37571066401169,83.38022910873822,83.38022910883822,83.38926510594712,83.38926510604712,83.39829814024138,83.39829814034138,83.40281190866047,83.40281190876047,83.40732485603093,83.40732485613093,83.42085969999242,83.42085970009242,83.42987816959784,83.42987816969784,83.43889506850152,83.43889506860152,83.44340116190077,83.44340116200077,83.4479066127402,83.4479066128402,83.45240620914589,83.4524062092459,83.45690462752528,83.45690462762528,83.46140208206496,83.46140208216497,83.46589728764535,83.46589728774535,83.47488594961553,83.47488594971553,83.47937844216564,83.47937844226564,83.48386989948051,83.48386989958051,83.48835975039587,83.48835975049587,83.4928463171167,83.4928463172167,83.49733174150899,83.49733174160899,83.50181605927051,83.50181605937051,83.50629666446429,83.50629666456429,83.51077662709827,83.51077662719827,83.51973244712305,83.51973244722305,83.52420723358084,83.52420723368084,83.53315262985772,83.53315262995773,83.53762309688575,83.53762309698575,83.55103199912614,83.55103199922614,83.55549971742612,83.55549971752612,83.56443308355561,83.56443308365562,83.57336123781116,83.57336123791116,83.57782167376672,83.57782167386672,83.58228111018481,83.58228111028481,83.59119916197237,83.59119916207237,83.60011546456934,83.60011546466934,83.61348691985242,83.61348691995242,83.62239901011293,83.62239901021293,83.62685216372407,83.62685216382407,83.63575711443123,83.63575711453123,83.64020641268357,83.64020641278357,83.65355241545893,83.65355241555893,83.65799914347207,83.65799914357207,83.66689002925912,83.66689002935912,83.67133304470451,83.67133304480451,83.6757745608437,83.67577456094371,83.68465655788684,83.68465655798684,83.68909546808905,83.68909546818905,83.69353245061185,83.69353245071186,83.7024057373999,83.7024057374999,83.71127077800384,83.71127077810384,83.71569787224526,83.71569787234526,83.72455113258616,83.72455113268616,83.72897629914816,83.72897629924816,83.73339946663523,83.73339946673524,83.74224390962776,83.74224390972776,83.75550709358427,83.75550709368427,83.76876570822664,83.76876570832664,83.7731822359291,83.7731822360291,83.77759601490351,83.77759601500351,83.78200904422484,83.78200904432484,83.79082999808682,83.79082999818682,83.7952368516945,83.7952368517945,83.81286244553907,83.81286244563907,83.8216733683286,83.8216733684286,83.82607658076407,83.82607658086407,83.83047854377769,83.83047854387769,83.83487890039181,83.83487890049182,83.83927647258012,83.83927647268013,83.8436734379064,83.8436734380064,83.84806815427338,83.84806815437338,83.85685630188773,83.85685630198773,83.8612488763887,83.8612488764887,83.8700324546889,83.8700324547889,83.8744223161596,83.8744223162596,83.87880985727546,83.87880985737546,83.8831947210588,83.8831947211588,83.88757894228236,83.88757894238236,83.8919613429198,83.8919613430198,83.89633924563863,83.89633924573863,83.90071586323786,83.90071586333787,83.90509048176216,83.90509048186216,83.90946427923782,83.90946427933783,83.9138364346162,83.9138364347162,83.9182078403415,83.9182078404415,83.92257856780921,83.92257856790921,83.93131730971437,83.93131730981437,83.93568482438309,83.93568482448309,83.9444179260411,83.9444179261411,83.94878308465721,83.94878308475721,83.95314724373584,83.95314724383584,83.96187356281817,83.96187356291817,83.96623486607547,83.96623486617547,83.97059392037346,83.97059392047346,83.97495143966748,83.97495143976748,83.98366351534088,83.98366351544088,83.98801746485823,83.98801746495823,83.99672368609788,83.99672368619788,84.00542658744521,84.00542658754522,84.00977625323056,84.00977625333056,84.0184743353794,84.0184743354794,84.0228226446496,84.0228226447496,84.02716692007215,84.02716692017215,84.03150876804656,84.03150876814657,84.04019153585345,84.04019153595345,84.04452917149139,84.04452917159139,84.04886616456953,84.04886616466953,84.053201729737,84.053201729837,84.06187153925457,84.06187153935457,84.06620517674263,84.06620517684263,84.07486963159518,84.07486963169518,84.0791998777954,84.0791998778954,84.08352866038719,84.08352866048719,84.08785555099733,84.08785555109733,84.09218012125265,84.09218012135265,84.09650294231739,84.09650294241739,84.10082494233349,84.10082494243349,84.10514612130096,84.10514612140096,84.11378462387708,84.11378462397708,84.11809816352246,84.11809816362246,84.12241077502591,84.12241077512591,84.12672199431645,84.12672199441646,84.13534304068463,84.13534304078463,84.13965115426947,84.13965115436947,84.14395812552577,84.14395812562577,84.16118376159167,84.16118376169167,84.16548798411993,84.16548798421994,84.17409550103451,84.17409550113452,84.18270041201212,84.18270041211213,84.20420356867676,84.20420356877676,84.20849576105762,84.20849576115762,84.21278631134122,84.21278631144122,84.22136648376647,84.22136648386648,84.22994497839669,84.22994497849669,84.24281095330255,84.24281095340255,84.25567168063671,84.25567168073671,84.25995555543793,84.25995555553793,84.26852226980512,84.26852226990512,84.27280328878498,84.27280328888499,84.28564523909384,84.28564523919384,84.29420459972107,84.29420459982107,84.29848304846173,84.29848304856174,84.30703737570387,84.30703737580387,84.31558802607601,84.31558802617602,84.31986154852487,84.31986154862487,84.32413421422733,84.32413421432733,84.328404702366,84.328404702466,84.34121431049817,84.34121431059818,84.34975143141672,84.34975143151672,84.35828676744694,84.35828676754694,84.36682014009999,84.36682014019999,84.37534976448752,84.37534976458753,84.38387524793411,84.38387524803412,84.39239894649236,84.39239894659237,84.39665979623402,84.39665979633402,84.40517974652677,84.40517974662677,84.41795590585149,84.41795590595149,84.42221343570084,84.42221343580084,84.42646985891942,84.42646985901942,84.43498163442358,84.43498163452358,84.44349176783047,84.44349176793047,84.45625364804849,84.45625364814849,84.46050578753506,84.46050578763506,84.46475735585736,84.46475735595736,84.46900756766453,84.46900756776454,84.48175631110439,84.48175631120439,84.49025223680012,84.49025223690012,84.49449859324847,84.49449859334847,84.50299002102558,84.50299002112558,84.50723416421239,84.5072341643124,84.51572016592895,84.51572016602896,84.52844427790976,84.52844427800976,84.53268224538293,84.53268224548293,84.53691821378116,84.53691821388117,84.54115346822407,84.54115346832407,84.54538797301386,84.54538797311386,84.55385505491404,84.55385505501404,84.57501686953299,84.57501686963299,84.57924687640418,84.57924687650419,84.58347602652896,84.58347602662896,84.58770453409394,84.58770453419395,84.59192982885992,84.59192982895992,84.59615333873755,84.59615333883755,84.60037584907772,84.60037584917772,84.60459696720498,84.60459696730499,84.6088163004439,84.60881630054391,84.61303388449227,84.61303388459227,84.62146791026044,84.62146791036044,84.62989975846484,84.62989975856485,84.63411466518069,84.63411466528069,84.64254040906698,84.64254040916698,84.64675113914411,84.64675113924412,84.65517085010782,84.65517085020782,84.65937726075519,84.6593772608552,84.66358274326062,84.66358274336062,84.66778758320623,84.66778758330624,84.67199113803225,84.67199113813226,84.68039421383666,84.68039421393667,84.69299216990976,84.69299217000976,84.69719047716407,84.69719047726407,84.70558477131786,84.70558477141786,84.71817169678103,84.71817169688103,84.72236561321003,84.72236561331003,84.73075205385513,84.73075205395513,84.73494411400026,84.73494411410026,84.74332734184634,84.74332734194634,84.75170932027059,84.7517093203706,84.7600870506606,84.7600870507606,84.76427268520771,84.76427268530772,84.76845742731065,84.76845742741065,84.7726403488275,84.7726403489275,84.78936964314448,84.78936964324448,84.79355003011989,84.79355003021989,84.79772970313995,84.79772970323995,84.80608783545601,84.80608783555601,84.81444414718597,84.81444414728597,84.81862135706012,84.81862135716013,84.82279653216158,84.82279653226158,84.82696910132606,84.82696910142606,84.83531102685602,84.83531102695602,84.83947884821752,84.83947884831753,84.8478132415187,84.8478132416187,84.8519783855477,84.8519783856477,84.85613881747148,84.85613881757148,84.86029814276449,84.8602981428645,84.86445618293791,84.86445618303792,84.87692869705864,84.87692869715865,84.88108452397053,84.88108452407053,84.89770401195699,84.89770401205699,84.90600699122342,84.90600699132342,84.9184569086578,84.9184569087578,84.92675481884136,84.92675481894136,84.93090122154281,84.93090122164281,84.93504641052019,84.93504641062019,84.9433347180045,84.9433347181045,84.94747744383596,84.94747744393597,84.9516194557121,84.95161945581211,84.95990094492295,84.95990094502295,84.968178578775,84.968178578875,84.9764529284325,84.9764529285325,84.98472342273118,84.98472342283118,84.98885651016563,84.98885651026563,84.99712143561271,84.99712143571271,85.00125280955436,85.00125280965436,85.01364507509544,85.01364507519544,85.0177693451815,85.01776934528151,85.021890795144,85.021890795244,85.03013258843822,85.03013258853822,85.038372918124,85.038372918224,85.04249222622049,85.04249222632049,85.04660996361524,85.04660996371524,85.0589618906799,85.0589618907799,85.06307787888407,85.06307787898407,85.06719301034185,85.06719301044185,85.0713071065644,85.0713071066644,85.07953401388988,85.07953401398989,85.0836463252241,85.0836463253241,85.0877579939985,85.0877579940985,85.09186844904883,85.09186844914883,85.09597679793093,85.09597679803093,85.10419110394474,85.10419110404474,85.11240216146179,85.11240216156179,85.1206121837436,85.1206121838436,85.12882038543931,85.12882038553931,85.13702723061988,85.13702723071988,85.14523097009474,85.14523097019475,85.14933132267709,85.14933132277709,85.15753049283781,85.15753049293781,85.16162870355414,85.16162870365415,85.16572587903525,85.16572587913525,85.16982105544142,85.16982105554142,85.18210472837603,85.18210472847603,85.18619740593853,85.18619740603853,85.19028919105686,85.19028919115686,85.19437840593598,85.19437840603598,85.19846069544835,85.19846069554835,85.2025423424009,85.2025423425009,85.2107040656043,85.2107040657043,85.2147835349931,85.2147835350931,85.21886132658686,85.21886132668686,85.22701566035254,85.22701566045254,85.23516745957677,85.23516745967677,85.23924139581172,85.23924139591172,85.24738751909105,85.24738751919105,85.25553178608651,85.25553178618651,85.26367437528693,85.26367437538693,85.2718148583191,85.2718148584191,85.27588269023593,85.27588269033593,85.27994977249966,85.27994977259966,85.28401564103933,85.28401564113933,85.29214627148788,85.29214627158788,85.29621024804591,85.29621024814591,85.30027322506646,85.30027322516646,85.30433520254955,85.30433520264955,85.308396358984,85.308396359084,85.31651731533778,85.31651731543778,85.32057615141738,85.32057615151739,85.32463323830642,85.32463323840642,85.32868875449373,85.32868875459373,85.3327429141659,85.3327429142659,85.3367943608078,85.3367943609078,85.34894745131164,85.34894745141165,85.35299607782997,85.35299607792997,85.3610904393475,85.36109043944751,85.36513585306682,85.36513585316682,85.3691783038715,85.3691783039715,85.37322018351189,85.37322018361189,85.37725849337562,85.37725849347562,85.38129619637732,85.38129619647732,85.38937074563431,85.38937074573431,85.39744383128287,85.39744383138287,85.40147957090738,85.40147957100739,85.40551431099445,85.40551431109445,85.40954762317082,85.40954762327083,85.4176123198442,85.4176123199442,85.4256746247672,85.4256746248672,85.4377656011569,85.4377656012569,85.4417941655303,85.4417941656303,85.44582108780644,85.44582108790644,85.44984686775403,85.44984686785403,85.45789699973857,85.45789699983857,85.461920816309,85.461920816409,85.46996780689005,85.46996780699006,85.47801347665374,85.47801347675374,85.48203454449613,85.48203454459613,85.48605286361048,85.48605286371048,85.49006921934766,85.49006921944766,85.50211650167087,85.50211650177087,85.5101440726668,85.5101440727668,85.51415593048537,85.51415593058537,85.52217871798058,85.52217871808058,85.5301985782589,85.5301985783589,85.53420700909186,85.53420700919186,85.54222219296274,85.54222219306274,85.55023523496763,85.55023523506763,85.55824545684892,85.55824545694892,85.56224803324814,85.56224803334814,85.56625000278532,85.56625000288533,85.5702513297627,85.5702513298627,85.57424947963884,85.57424947973884,85.5782470583507,85.5782470584507,85.59023697436272,85.59023697446273,85.59423148306665,85.59423148316665,85.6022180373286,85.6022180374286,85.60620972590895,85.60620972600896,85.61020009367193,85.61020009377194,85.61418981887512,85.61418981897512,85.62216759148644,85.62216759158645,85.62615528191692,85.62615528201692,85.63811424796518,85.63811424806518,85.65006478934049,85.65006478944049,85.65404666103498,85.65404666113498,85.66996900733456,85.66996900743456,85.67792150592703,85.67792150602703,85.68189605957934,85.68189605967935,85.69381900658094,85.69381900668094,85.70176543655306,85.70176543665306,85.7057371700818,85.7057371701818,85.70970740430434,85.70970740440434,85.72161553673274,85.72161553683274,85.72955029353513,85.72955029363513,85.73351649391003,85.73351649401003,85.73748048102338,85.73748048112338,85.74540713443272,85.74540713453273,85.74936944375104,85.74936944385104,85.75729184912613,85.75729184922614,85.76521029204912,85.76521029214912,85.77312516519544,85.77312516529544,85.78103703972934,85.78103703982934,85.79289965750513,85.79289965760513,85.79685079342245,85.79685079352245,85.80475181583526,85.80475181593526,85.80870073849105,85.80870073859106,85.81659365751081,85.81659365761081,85.82053736829265,85.82053736839265,85.82842325485237,85.82842325495237,85.83630392953813,85.83630392963813,85.84417957083878,85.84417957093878,85.84811614206727,85.84811614216727,85.85598739254262,85.85598739264262,85.86385628696534,85.86385628706535,85.8795874360262,85.8795874361262,85.88744719182066,85.88744719192066,85.89137483860745,85.89137483870745,85.89530141446124,85.89530141456125,85.89922716926641,85.89922716936641,85.90315228151178,85.90315228161178,85.90707575165987,85.90707575175988,85.91099822227049,85.9109982223705,85.91884205686097,85.91884205696097,85.92276206432568,85.92276206442568,85.92667957294672,85.92667957304673,85.93843038531706,85.93843038541706,85.94234514521007,85.94234514531007,85.95800282826694,85.95800282836694,85.96974303840055,85.96974303850055,85.97365476398339,85.97365476408339,85.97756359804711,85.97756359814711,85.98537830325992,85.98537830335992,85.99318983137148,85.99318983147148,85.99709438170319,85.9970943818032,86.00099771831084,86.00099771841084,86.00489962700782,86.00489962710782,86.00879917965219,86.00879917975219,86.01269816113228,86.01269816123228,86.02049473187958,86.02049473197958,86.02439203556465,86.02439203566465,86.03218460816207,86.03218460826207,86.03607887753695,86.03607887763695,86.03997154051234,86.03997154061234,86.04775579553011,86.04775579563011,86.05164685210599,86.05164685220599,86.05942800141806,86.05942800151806,86.07109752997349,86.07109753007349,86.07498612340346,86.07498612350346,86.07887296764288,86.07887296774288,86.08664333622937,86.08664333632937,86.09052557545687,86.09052557555687,86.09440652956476,86.09440652966477,86.09828691250839,86.09828691260839,86.10604371594353,86.10604371604353,86.1099188870132,86.1099188871132,86.11766433855851,86.11766433865851,86.12540411415891,86.12540411425891,86.13314167649777,86.13314167659777,86.13700935103643,86.13700935113643,86.14474341499414,86.14474341509414,86.15247572976129,86.1524757298613,86.15634008440765,86.15634008450765,86.16793043531642,86.16793043541642,86.17179193414144,86.17179193424144,86.17565275470888,86.17565275480888,86.17951071945498,86.17951071955498,86.18336797024575,86.18336797034576,86.18722464987225,86.18722464997225,86.19493718807662,86.19493718817662,86.20650376427274,86.20650376437274,86.21035787366004,86.21035787376005,86.21421119769647,86.21421119779647,86.21806395056862,86.21806395066862,86.2219157039033,86.22191570400331,86.22961749707989,86.22961749717989,86.24116633148594,86.24116633158594,86.24886080662033,86.24886080672033,86.25270599156593,86.25270599166593,86.26039482645318,86.26039482655318,86.2642364416221,86.2642364417221,86.27191817265377,86.27191817275377,86.27959683367749,86.27959683377749,86.28343423650995,86.28343423660995,86.28727071120046,86.28727071130047,86.29110547239817,86.29110547249817,86.29493944824502,86.29493944834502,86.30260650749454,86.30260650759455,86.30643859135975,86.30643859145975,86.31026878324333,86.31026878334333,86.31409676186536,86.31409676196536,86.31792217024818,86.31792217034818,86.32174682897791,86.32174682907791,86.32557059526347,86.32557059536347,86.33321702120381,86.33321702130381,86.3446835722575,86.3446835723575,86.3523260714435,86.3523260715435,86.35614630365015,86.35614630375015,86.35996592899475,86.35996592909476,86.36378323398453,86.36378323408454,86.3752286519603,86.3752286520603,86.38284830457559,86.38284830467559,86.39046474439186,86.39046474449187,86.39807814989796,86.39807814999796,86.4018831391582,86.4018831392582,86.4056847371306,86.4056847372306,86.41708771046171,86.41708771056172,86.42088627412394,86.42088627422395,86.42468237464026,86.42468237474026,86.42847586921961,86.42847586931961,86.43226786449276,86.43226786459276,86.44363988786009,86.44363988796009,86.45499995247556,86.45499995257556,86.46256587995188,86.46256588005188,86.4701290230024,86.4701290231024,86.47768913174274,86.47768913184274,86.49280245955443,86.49280245965443,86.4965787478102,86.4965787479102,86.50412964652672,86.50412964662672,86.50790240070359,86.50790240080359,86.51167404824969,86.51167404834969,86.51544430358288,86.51544430368288,86.51921373786745,86.51921373796745,86.5267511785259,86.5267511786259,86.53051864943328,86.53051864953328,86.534282550564,86.534282550664,86.5380453807617,86.5380453808617,86.54556972033976,86.54556972043976,86.55685185673042,86.55685185683042,86.56060933226311,86.56060933236311,86.5681210705295,86.5681210706295,86.5718745836101,86.5718745837101,86.58313237412382,86.58313237422382,86.5868817105657,86.58688171066571,86.59063029735448,86.59063029745448,86.59812597162583,86.59812597172584,86.6018718810821,86.6018718811821,86.60936220068844,86.60936220078844,86.613106075372,86.613106075472,86.62433612872097,86.62433612882097,86.62807832560951,86.62807832570951,86.63556175554686,86.63556175564686,86.64678177782336,86.64678177792337,86.6617302533482,86.6617302534482,86.66546377567941,86.66546377577941,86.66919633417092,86.66919633427092,86.67292725056515,86.67292725066515,86.67665720311969,86.67665720321969,86.68038515659929,86.68038515669929,86.68411228903024,86.68411228913024,86.69156537586586,86.69156537596587,86.6990162851377,86.6990162852377,86.70646490975248,86.70646490985249,86.7101866161229,86.7101866162229,86.71390621632507,86.71390621642507,86.72506294646112,86.72506294656112,86.7324976132491,86.7324976133491,86.7362136436746,86.7362136437746,86.74364338417077,86.74364338427077,86.75107001896122,86.75107001906122,86.75478226542344,86.75478226552345,86.75849379793033,86.75849379803033,86.76220450938858,86.76220450948858,86.76591350735404,86.76591350745404,86.76962136299096,86.76962136309096,86.77332782641497,86.77332782651497,86.78073907546795,86.78073907556795,86.78814807556164,86.78814807566164,86.79185118339556,86.79185118349557,86.79555368436748,86.79555368446748,86.79925554277958,86.79925554287958,86.80665808157747,86.80665808167747,86.81775905180186,86.81775905190186,86.82145755462389,86.82145755472389,86.82515495081515,86.82515495091515,86.82885156165553,86.82885156175553,86.83993975207943,86.83993975217943,86.85102430133111,86.85102430143111,86.85471784216355,86.85471784226355,86.85841016927193,86.85841016937194,86.8621016396339,86.86210163973391,86.86579193196958,86.86579193206958,86.86947947557722,86.86947947567722,86.87316551987864,86.87316551997864,86.87685017196718,86.87685017206718,86.88053393161152,86.88053393171153,86.88421672741619,86.88421672751619,86.89526368691948,86.89526368701948,86.90262631561416,86.90262631571416,86.90630639838854,86.90630639848854,86.91366449346683,86.91366449356683,86.9173417918154,86.9173417919154,86.92837100952862,86.92837100962862,86.9320452021715,86.9320452022715,86.9504129882846,86.9504129883846,86.95775537634552,86.95775537644552,86.96876678087312,86.96876678097313,86.97610631311271,86.97610631321271,86.97977486550843,86.97977486560843,86.98710972134057,86.98710972144058,86.99077359732885,86.99077359742886,87.00176394017407,87.00176394027407,87.00542556720305,87.00542556730305,87.01274739335032,87.01274739345033,87.01640744967756,87.01640744977756,87.02738651202849,87.02738651212849,87.03104464067633,87.03104464077633,87.03470180548445,87.03470180558445,87.04201488567887,87.04201488577887,87.04932582400728,87.04932582410729,87.05297852659457,87.05297852669457,87.05662573172577,87.05662573182578,87.06391899965965,87.06391899975965,87.06756484827571,87.06756484837571,87.07120955456324,87.07120955466324,87.0821425310973,87.0821425311973,87.08578602366077,87.08578602376078,87.0894289093622,87.0894289094622,87.09306761842491,87.09306761852491,87.09670468539036,87.09670468549037,87.1076142798872,87.1076142799872,87.1112492406844,87.1112492407844,87.11488327333967,87.11488327343967,87.11851348633391,87.11851348643391,87.12577262720276,87.12577262730277,87.1294012337975,87.1294012338975,87.13665630512094,87.13665630522094,87.14028162752113,87.14028162762114,87.14753055882869,87.1475305589287,87.15115391785172,87.15115391795172,87.15477631303503,87.15477631313503,87.16564157090556,87.16564157100557,87.16926178852509,87.16926178862509,87.17288061393174,87.17288061403174,87.18373555514768,87.18373555524768,87.18735070368434,87.18735070378435,87.19457971563806,87.19457971573806,87.19819265091319,87.19819265101319,87.20180487223297,87.20180487233297,87.20541620110859,87.20541620120859,87.20902638765567,87.20902638775567,87.21624311957763,87.21624311967763,87.2198479514597,87.2198479515597,87.2270562587087,87.2270562588087,87.23065916291138,87.23065916301138,87.2342608176922,87.2342608177922,87.2378614372378,87.2378614373378,87.24506164109376,87.24506164119376,87.25225984587476,87.25225984597476,87.26305417228276,87.26305417238277,87.2702474150742,87.2702474151742,87.2738416804173,87.27384168051731,87.2810285333085,87.2810285334085,87.28461994283028,87.28461994293028,87.2918009412877,87.2918009413877,87.29538678195787,87.29538678205788,87.29897176588163,87.29897176598163,87.3025523232823,87.3025523233823,87.30613120288795,87.30613120298796,87.30970940423602,87.30970940433602,87.31328482115829,87.31328482125829,87.31685895296096,87.31685895306096,87.32400461062932,87.32400461072932,87.3275749584687,87.3275749585687,87.33114473514382,87.33114473524383,87.34185224458308,87.34185224468308,87.34898740710801,87.34898740720801,87.35255275726006,87.35255275736006,87.35611692938579,87.3561169294858,87.35967888825,87.35967888835,87.3668016279521,87.3668016280521,87.3774842917457,87.3774842918457,87.3846045683019,87.3846045684019,87.38816370704252,87.38816370714252,87.39172134647696,87.39172134657696,87.39527791497838,87.39527791507838,87.40238912430182,87.40238912440182,87.40594180174666,87.40594180184667,87.40949219453444,87.40949219463444,87.41304190906465,87.41304190916465,87.42013976742332,87.42013976752332,87.42368430577736,87.42368430587736,87.42722805878051,87.42722805888052,87.4307692058467,87.4307692059467,87.43430971035308,87.43430971045308,87.43784564554532,87.43784564564533,87.44491651639234,87.44491651649234,87.45198613781751,87.45198613791752,87.45551935997948,87.45551936007948,87.46258087801161,87.46258087811161,87.46610756748225,87.46610756758226,87.4696334002065,87.4696334003065,87.4766830308823,87.4766830309823,87.48020479406114,87.48020479416114,87.48372512932931,87.48372512942932,87.49076369369742,87.49076369379742,87.49428153012191,87.49428153022191,87.49779815282234,87.49779815292234,87.50131402586968,87.50131402596968,87.50834391568046,87.50834391578046,87.51537055699445,87.51537055709446,87.51888243189191,87.51888243199191,87.52239352143849,87.52239352153849,87.5259035043543,87.5259035044543,87.53292193518196,87.53292193528196,87.5364294192541,87.5364294193541,87.5399359751843,87.53993597528431,87.54694708796978,87.54694708806979,87.55395534493391,87.55395534503391,87.56095896118836,87.56095896128836,87.56795679440461,87.56795679450461,87.57145306937798,87.57145306947798,87.57844133559273,87.57844133569273,87.58542467551567,87.58542467561567,87.58891506035754,87.58891506045754,87.59240430287086,87.59240430297086,87.5958918318914,87.5958918319914,87.59937875404988,87.59937875414988,87.60635013522096,87.60635013532097,87.60983416586035,87.60983416596035,87.61331651870469,87.6133165188047,87.62027986787825,87.62027986797825,87.62376000746106,87.62376000756106,87.62723954018183,87.62723954028183,87.63071850173834,87.63071850183834,87.64115178093341,87.64115178103341,87.65157924139251,87.65157924149251,87.65505266979517,87.65505266989517,87.6585250629626,87.6585250630626,87.66546724336047,87.66546724346047,87.66893645942666,87.66893645952666,87.67587360643941,87.67587360653941,87.6793407877329,87.6793407878329,87.68280714797774,87.68280714807774,87.68973876183665,87.68973876193665,87.69320355137975,87.69320355147975,87.6966666988256,87.6966666989256,87.70012777580096,87.70012777590097,87.70358749626118,87.70358749636118,87.71050433124466,87.71050433134467,87.71395787599124,87.71395787609124,87.72086396594693,87.72086396604693,87.72431551161857,87.72431551171857,87.72776466553985,87.72776466563985,87.73121196317724,87.73121196327725,87.7381040239106,87.7381040240106,87.74154807305122,87.74154807315122,87.74499115835214,87.74499115845214,87.75187500859914,87.75187500869914,87.7553139886569,87.7553139887569,87.7621911634215,87.7621911635215,87.76562832289315,87.76562832299315,87.77250114253023,87.77250114263023,87.77593591025149,87.77593591035149,87.78280294685035,87.78280294695035,87.78966630657925,87.78966630667925,87.79995608075198,87.79995608085198,87.80681090871462,87.80681090881463,87.81366338062465,87.81366338072465,87.81708829576229,87.81708829586229,87.82393430637654,87.82393430647654,87.83419809257508,87.83419809267508,87.83761615374151,87.83761615384151,87.8478690521212,87.8478690522212,87.85128557828365,87.85128557838365,87.85470146188631,87.85470146198631,87.85811331164132,87.85811331174132,87.86493347707244,87.86493347717244,87.86834018634904,87.86834018644905,87.8785583151039,87.8785583152039,87.88196323949217,87.88196323959217,87.8887723386156,87.8887723387156,87.89217412160039,87.89217412170039,87.89557533342091,87.89557533352091,87.90577739818073,87.90577739828073,87.90917568278438,87.90917568288438,87.91597082408101,87.91597082418102,87.91936589588565,87.91936589598565,87.92275996815282,87.92275996825282,87.92615079192322,87.92615079202322,87.93632097857734,87.93632097867734,87.93970848245543,87.93970848255543,87.94648195520763,87.94648195530763,87.95325392865364,87.95325392875364,87.95663893368805,87.95663893378806,87.96002336755821,87.96002336765821,87.97017413462721,87.97017413472722,87.97355578407155,87.97355578417155,87.98708084684496,87.98708084694496,87.99046064000542,87.99046064010543,87.9938382912999,87.9938382913999,87.99721526433679,87.99721526443679,88.00396785389545,88.00396785399545,88.007341971111,88.007341971211,88.01746232368274,88.01746232378274,88.02420673845282,88.02420673855282,88.03768949937262,88.03768949947262,88.0477933595761,88.0477933596761,88.05115987316735,88.05115987326735,88.05789193651013,88.05789193661013,88.06462285752437,88.06462285762437,88.07135188655697,88.07135188665697,88.07807902360794,88.07807902370794,88.08143953997435,88.08143954007436,88.08815857363227,88.08815857373227,88.09151252160962,88.09151252170962,88.10157347309747,88.10157347319748,88.1082786560219,88.1082786561219,88.11162928410693,88.11162928420693,88.11497884125896,88.11497884135896,88.11832432886558,88.11832432896558,88.12166899542358,88.12166899552358,88.12501259104856,88.12501259114856,88.13504127175527,88.13504127185527,88.14506563303222,88.14506563313222,88.14840401678326,88.14840401688326,88.15174033006382,88.15174033016382,88.15507607218012,88.15507607228012,88.15841110034108,88.15841110044109,88.16174512896458,88.16174512906458,88.1684126150473,88.1684126151473,88.1750783162417,88.1750783163417,88.17840656182698,88.17840656192698,88.18506126810921,88.18506126820921,88.1917129043835,88.1917129044835,88.19835932878384,88.19835932888384,88.20500022003033,88.20500022013033,88.20831907710296,88.20831907720296,88.21163732731354,88.21163732741354,88.21827036505135,88.21827036515135,88.23152922957809,88.23152922967809,88.24478277513758,88.24478277523758,88.24809335032832,88.24809335042832,88.25140296167936,88.25140296177936,88.25471196616836,88.25471196626836,88.26132876142229,88.2613287615223,88.27125156255283,88.27125156265284,88.28117079390668,88.28117079400668,88.28447597873465,88.28447597883465,88.28777905739437,88.28777905749438,88.29768697255616,88.29768697265617,88.30098619585708,88.30098619595708,88.30428416973616,88.30428416983617,88.30757989465593,88.30757989475593,88.31746564150458,88.31746564160458,88.32405394994065,88.32405395004065,88.33063997371964,88.33063997381964,88.33722224923314,88.33722224933314,88.34050987075986,88.34050987085986,88.3437969211223,88.3437969212223,88.34708218659642,88.34708218669643,88.35365096835407,88.35365096845408,88.35693352079792,88.35693352089793,88.36021535928643,88.36021535938643,88.3634965909129,88.3634965910129,88.37005734067303,88.37005734077303,88.3733357878737,88.3733357879737,88.38316905900523,88.38316905910523,88.38972231223434,88.38972231233434,88.39299815349803,88.39299815359803,88.39627317371308,88.39627317381309,88.39954712299514,88.39954712309515,88.40281960866876,88.40281960876877,88.40609080922277,88.40609080932278,88.40936143861252,88.40936143871252,88.41263056869606,88.41263056879606,88.4191677936279,88.41916779372791,88.43223985174123,88.43223985184123,88.43877297142991,88.43877297152991,88.45183671196358,88.45183671206358,88.46816130069891,88.46816130079891,88.47142493332638,88.47142493342638,88.4779508063684,88.4779508064684,88.48121236852539,88.48121236862539,88.48447328812257,88.48447328822257,88.48773356515996,88.48773356525996,88.49099309254424,88.49099309264425,88.49425154899554,88.49425154909554,88.5007663200321,88.5007663201321,88.51053205098893,88.51053205108893,88.51378240404715,88.51378240414715,88.52028164655518,88.52028164665518,88.5267790684771,88.5267790685771,88.53002603024748,88.53002603034749,88.53327213527147,88.53327213537148,88.54300880824617,88.54300880834617,88.54625237872872,88.54625237882873,88.5559815551724,88.5559815552724,88.55922341216214,88.55922341226214,88.56894705545197,88.56894705555197,88.57218594952707,88.57218594962707,88.5786608104604,88.57866081056041,88.58189538510574,88.58189538520574,88.59483086356347,88.59483086366347,88.59806226110756,88.59806226120756,88.6045239138672,88.6045239139672,88.61744322123658,88.61744322133659,88.62389677060317,88.62389677070317,88.6271214569671,88.6271214570671,88.63356975876198,88.63356975886198,88.64001495485114,88.64001495495114,88.65289913561806,88.65289913571806,88.65933883425114,88.65933883435115,88.66255670234163,88.66255670244163,88.6689907250298,88.6689907251298,88.67220670113865,88.67220670123865,88.67542167771003,88.67542167781004,88.67863576183724,88.67863576193724,88.68184870363592,88.68184870373592,88.68827365909134,88.68827365919134,88.69148335239323,88.69148335249324,88.6979003115489,88.69790031164891,88.7011071133317,88.7011071134317,88.70751875352012,88.70751875362012,88.71072305645924,88.71072305655925,88.713926038581,88.713926038681,88.71712823535188,88.71712823545188,88.72353195063607,88.72353195073607,88.72673239821638,88.72673239831639,88.72993184625923,88.72993184635924,88.73312833138743,88.73312833148744,88.73632381697817,88.73632381707817,88.7395174819828,88.7395174820828,88.74270989756559,88.74270989766559,88.74909173011878,88.74909173021878,88.76184832706733,88.76184832716733,88.76503595914937,88.76503595924937,88.76822173494755,88.76822173504755,88.78096080429262,88.78096080439262,88.78414265333646,88.78414265343646,88.79050510200229,88.79050510210229,88.79368502336672,88.79368502346672,88.79686423077581,88.79686423087581,88.80322150326548,88.80322150336548,88.8063984260175,88.8063984261175,88.80957299271691,88.80957299281691,88.81909508641567,88.81909508651567,88.82861589499481,88.82861589509481,88.83496067326611,88.83496067336611,88.83813099193128,88.83813099203128,88.84130041815229,88.84130041825229,88.84763662895956,88.84763662905956,88.85080259249719,88.85080259259719,88.85396759219512,88.85396759229512,88.85713191363548,88.85713191373549,88.8634589501167,88.8634589502167,88.86662120108659,88.86662120118659,88.86978220263464,88.86978220273464,88.87610249223795,88.87610249233795,88.87926088784903,88.87926088794903,88.88557710790691,88.88557710800691,88.88873368293189,88.88873368303189,88.891889472606,88.891889472706,88.90135537801989,88.90135537811989,88.90450963269002,88.90450963279002,88.91081635714195,88.91081635724196,88.91396843424832,88.91396843434832,88.91711983309712,88.91711983319712,88.92026962554641,88.92026962564641,88.9234175974096,88.9234175975096,88.92656431985095,88.92656431995096,88.93285658670736,88.93285658680736,88.93914689018659,88.9391468902866,88.94228954308252,88.94228954318253,88.94543144632537,88.94543144642537,88.95171343222495,88.95171343232495,88.95485230115762,88.95485230125762,88.95799049183272,88.95799049193272,88.96426573085438,88.96426573095438,88.96740274350317,88.96740274360317,88.97367323472186,88.97367323482186,88.97993933511523,88.97993933521524,88.98933584407055,88.98933584417055,88.99559866026938,88.99559866036938,89.00185958448658,89.00185958458658,89.00811786706903,89.00811786716903,89.01124563399624,89.01124563409624,89.01437254417705,89.01437254427705,89.02062493662798,89.02062493672798,89.02687565128389,89.02687565138389,89.0299998841322,89.0299998842322,89.03624674342929,89.03624674352929,89.04561355184266,89.04561355194267,89.04873464328749,89.04873464338749,89.05185516356805,89.05185516366805,89.05497304221386,89.05497304231386,89.07056015078588,89.07056015088588,89.07990893182705,89.07990893192705,89.086140048409,89.08614004850901,89.09236852335623,89.09236852345623,89.09859471364638,89.09859471374638,89.10170541704107,89.10170541714108,89.10792536022204,89.10792536032204,89.11725179446123,89.11725179456123,89.12035832121722,89.12035832131723,89.12657037519173,89.12657037529173,89.12967547403706,89.12967547413706,89.13588367265275,89.13588367275275,89.1389865939343,89.13898659403431,89.14208883695828,89.14208883705828,89.14519040172469,89.14519040182469,89.15449252578473,89.15449252588473,89.16069055053691,89.16069055063691,89.16378733180257,89.16378733190257,89.16688218538884,89.16688218548884,89.17307071453506,89.17307071463506,89.17925488855374,89.17925488865374,89.1823450300348,89.1823450301348,89.19779320289861,89.19779320299861,89.20088006018513,89.20088006028513,89.20396573944532,89.20396573954532,89.21013616982381,89.21013616992381,89.21630438694075,89.21630438704075,89.21938538979352,89.21938538989352,89.2224657143887,89.2224657144887,89.22554364723354,89.22554364733354,89.22862079472749,89.22862079482749,89.23784991685451,89.23784991695452,89.24399782194217,89.24399782204218,89.24706861523364,89.24706861533365,89.25320838123048,89.25320838133048,89.25934557698812,89.25934557708813,89.26547934576014,89.26547934586014,89.27773970805308,89.27773970815308,89.28080179108947,89.28080179118948,89.2869242793672,89.2869242794672,89.29304480426777,89.29304480436777,89.29610388869175,89.29610388879175,89.29916229485815,89.29916229495815,89.31139449161311,89.31139449171312,89.31445125568226,89.31445125578226,89.31750537811666,89.31750537821667,89.3205579655471,89.3205579656471,89.32360991041772,89.32360991051772,89.32971255073714,89.32971255083714,89.33581312058608,89.33581312068608,89.33886235242645,89.33886235252645,89.34191037054272,89.34191037064272,89.34495449760243,89.34495449770243,89.35104182357992,89.35104182367992,89.35408395156469,89.35408395166469,89.36016720799677,89.36016720809677,89.36320708702223,89.36320708712223,89.37232468932592,89.37232468942592,89.37840080620465,89.37840080630465,89.38447456703076,89.38447456713077,89.38750951976442,89.38750951986442,89.39357578405952,89.39357578415952,89.39660563201252,89.39660563211253,89.3996342305437,89.3996342306437,89.4026593663915,89.4026593664915,89.40568300293309,89.40568300303309,89.40870581842606,89.40870581852606,89.41172784856815,89.41172784866815,89.41474752265763,89.41474752275764,89.41776641139624,89.41776641149625,89.42682061446618,89.42682061456618,89.43586917728898,89.43586917738898,89.44189631683173,89.44189631693173,89.44490642391972,89.44490642401972,89.45694488889453,89.45694488899453,89.46295910584571,89.46295910594571,89.46596300152228,89.46596300162228,89.4689662189413,89.4689662190413,89.47497108307759,89.47497108317759,89.47797251560827,89.47797251570827,89.48097291290371,89.48097291300371,89.48397095414654,89.48397095424654,89.48696699631444,89.48696699641444,89.49595362351192,89.49595362361192,89.50493710930593,89.50493710940593,89.51391527613269,89.5139152762327,89.51989825753796,89.51989825763796,89.52587970393925,89.52587970403926,89.52886967748681,89.52886967758681,89.53484655457396,89.53484655467396,89.54380680112182,89.54380680122182,89.54977789517076,89.54977789527076,89.55574681165592,89.55574681175592,89.55872918157915,89.55872918167915,89.56171037347609,89.56171037357609,89.57065227137183,89.57065227147183,89.57959095646856,89.57959095656857,89.58256672230495,89.58256672240495,89.58851693316034,89.58851693326034,89.59149037864188,89.59149037874188,89.6004093228736,89.6004093229736,89.60337769927226,89.60337769937226,89.60634489764462,89.60634489774462,89.61524449368677,89.61524449378678,89.6241406627433,89.6241406628433,89.63006941924088,89.63006941934088,89.63599603387244,89.63599603397245,89.63895805606863,89.63895805616863,89.64784344439961,89.64784344449961,89.65376449017957,89.65376449027957,89.65968353688459,89.65968353698459,89.66560069160796,89.66560069170797,89.67151427655466,89.67151427665466,89.67742429172469,89.67742429182469,89.68037799634122,89.68037799644122,89.68332945199845,89.68332945209845,89.6892296859804,89.6892296860804,89.69217810732744,89.69217810742744,89.70397050759604,89.70397050769604,89.7128117734873,89.7128117735873,89.71870215488562,89.71870215498562,89.72459018023133,89.72459018033133,89.73341938027744,89.73341938037744,89.74224658124862,89.74224658134862,89.74812914483601,89.74812914493602,89.75106799918157,89.75106799928157,89.75400546131424,89.75400546141424,89.75987906476217,89.75987906486218,89.76575088332179,89.76575088342179,89.76868395462913,89.76868395472913,89.77454952607955,89.77454952617956,89.77748116947623,89.77748116957623,89.78334302835891,89.78334302845892,89.79213249679063,89.79213249689063,89.79506110587714,89.79506110597714,89.80384550522596,89.80384550532597,89.80969951059997,89.80969951069997,89.81262430002543,89.81262430012544,89.81554819700672,89.81554819710672,89.81846998781977,89.81846998791977,89.82139113607302,89.82139113617302,89.82431078502007,89.82431078512008,89.82722914884752,89.82722914894752,89.83014694151069,89.83014694161069,89.83598170578841,89.83598170588841,89.83889803484315,89.83889803494316,89.84472983620623,89.84472983630623,89.8476433451374,89.8476433452374,89.8505562829043,89.8505562830043,89.85929238317472,89.85929238327472,89.86220178686271,89.86220178696271,89.86511036950206,89.86511036960206,89.87092639245223,89.87092639255224,89.8767396309766,89.8767396310766,89.8825503706573,89.8825503707573,89.88835907556529,89.88835907566529,89.89706897367492,89.89706897377492,89.9028733948509,89.9028733949509,89.9057744095637,89.9057744096637,89.91447656125794,89.91447656135794,89.92027644881755,89.92027644891755,89.92317539305988,89.92317539315988,89.92897042572321,89.92897042582321,89.93476224558752,89.93476224568752,89.94055295882107,89.94055295892107,89.9434470838649,89.9434470839649,89.9492337632508,89.9492337633508,89.95212535375317,89.95212535385318,89.95790742812717,89.95790742822717,89.9636877533106,89.9636877534106,89.97235556375323,89.97235556385323,89.98679595296389,89.98679595306389,89.99545640966656,89.99545640976656,90.00411233275287,90.00411233285287,90.0098790213894,90.0098790214894,90.01564406792865,90.01564406802865,90.02140690120636,90.02140690130636,90.02428717551668,90.02428717561668,90.03004679599539,90.03004679609539,90.03292303645807,90.03292303655807,90.03579831308105,90.03579831318105,90.03867094806928,90.03867094816928,90.04154297619549,90.04154297629549,90.04728503337294,90.04728503347295,90.05015459835325,90.05015459845325,90.0530224855385,90.05302248563851,90.05588837364883,90.05588837374883,90.05875233407974,90.05875233417974,90.06161522357765,90.06161522367765,90.07306578203183,90.07306578213183,90.08165016679357,90.08165016689357,90.09023073189425,90.09023073199425,90.09308758846959,90.09308758856959,90.09594258876105,90.09594258886105,90.10450616172477,90.10450616182477,90.10735502200035,90.10735502210035,90.11304795905076,90.11304795915076,90.11589153605686,90.11589153615687,90.12726209581575,90.12726209591575,90.13010142478761,90.13010142488761,90.13861751972155,90.13861751982155,90.14429242939975,90.14429242949976,90.14712818859493,90.14712818869494,90.15563346710555,90.15563346720555,90.16130130862594,90.16130130872594,90.16413414060425,90.16413414070425,90.17546346944255,90.17546346954255,90.17829319571514,90.17829319581514,90.18395168442065,90.18395168452065,90.18677869766297,90.18677869776297,90.1924321172856,90.19243211738561,90.19525773831502,90.19525773841502,90.20090773095204,90.20090773105204,90.20373178127973,90.20373178137973,90.2093778471624,90.2093778472624,90.2150207716416,90.2150207717416,90.23194172726828,90.23194172736828,90.23757465637279,90.23757465647279,90.24320515802914,90.24320515812914,90.24601903449332,90.24601903459332,90.25445941446397,90.25445941456397,90.2600843829665,90.2600843830665,90.26851958676097,90.26851958686098,90.27132910809759,90.27132910819759,90.27694643727803,90.27694643737803,90.27975071104294,90.27975071114294,90.28255409236367,90.28255409246367,90.2937661540382,90.2937661541382,90.29656792895943,90.29656792905944,90.30217058635773,90.30217058645773,90.30777149456546,90.30777149466546,90.31056766493732,90.31056766503733,90.3161589704458,90.3161589705458,90.31895353441814,90.31895353451814,90.32174699175972,90.32174699185973,90.32733304969649,90.3273330497965,90.33012511482517,90.33012511492517,90.33291650169627,90.33291650179628,90.33849559856853,90.33849559866853,90.34128159507686,90.34128159517687,90.34406644925666,90.34406644935666,90.35241819167247,90.35241819177247,90.35519976165772,90.35519976175772,90.36076161650863,90.36076161660863,90.36910272098994,90.36910272108994,90.3718816136427,90.3718816137427,90.38021368658903,90.38021368668903,90.39131212227198,90.39131212237199,90.39685923394522,90.39685923404522,90.40517706348261,90.40517706358261,90.4107174639757,90.4107174640757,90.4134851475297,90.4134851476297,90.41901890823816,90.41901890833816,90.42731610446639,90.42731610456639,90.43560823161172,90.43560823171173,90.43836902549673,90.43836902559673,90.44664937237904,90.44664937247904,90.45216653362597,90.45216653372597,90.45492379343206,90.45492379353206,90.46319250284242,90.46319250294242,90.46870009708786,90.46870009718786,90.4797113588244,90.4797113589244,90.48246304977889,90.48246304987889,90.48796536075486,90.48796536085486,90.49621679244609,90.4962167925461,90.50446540401376,90.50446540411376,90.50996121799618,90.50996121809618,90.51270605497945,90.51270605507945,90.51819487219957,90.51819487229957,90.52642492092853,90.52642492102854,90.53190802650599,90.53190802660599,90.534646473589,90.53464647368901,90.53738263601495,90.53738263611496,90.54285317597851,90.54285317607851,90.54832150268052,90.54832150278052,90.55652127970328,90.55652127980328,90.56198582244203,90.56198582254203,90.56744872308349,90.56744872318349,90.57290841092596,90.57290841102596,90.5810923024425,90.5810923025425,90.59199971937556,90.59199971947557,90.59472513677373,90.59472513687373,90.59744923335451,90.59744923345451,90.60289631988532,90.60289631998532,90.60834087187467,90.60834087197468,90.6137823538561,90.6137823539561,90.6165012742607,90.6165012743607,90.6192192308256,90.61921923092561,90.62193658052847,90.62193658062847,90.6300859166068,90.6300859167068,90.63551601100063,90.63551601110063,90.64365802903679,90.64365802913679,90.64637038105231,90.64637038115231,90.64908109097055,90.64908109107056,90.65179108693347,90.65179108703347,90.65450036894106,90.65450036904106,90.65720865141117,90.65720865151117,90.66262400262734,90.66262400272734,90.67074542305207,90.67074542315207,90.67345181354054,90.67345181364054,90.67615706170048,90.67615706180048,90.6815666655762,90.6815666656762,90.68426902221702,90.68426902231703,90.68967091537512,90.68967091547512,90.69237034479909,90.69237034489909,90.70046552736525,90.70046552746525,90.70316120852371,90.70316120862371,90.70585614002906,90.70585614012906,90.71124489640899,90.71124489650899,90.71393825721259,90.71393825731259,90.71662951184797,90.71662951194797,90.72739324526987,90.72739324536987,90.73277154220413,90.73277154230414,90.73545822752536,90.73545822762536,90.73814237830516,90.73814237840516,90.74619272447629,90.74619272457629,90.75155645672172,90.75155645682172,90.75691779721679,90.75691779731679,90.75959637914497,90.75959637924497,90.76763012585455,90.76763012595455,90.7729836485387,90.7729836486387,90.7783352792412,90.7783352793412,90.78368437540227,90.78368437550228,90.7863564603369,90.7863564604369,90.79971470744627,90.79971470754627,90.80505534323662,90.80505534333662,90.80772442955885,90.80772442965885,90.81038991040663,90.81038991050663,90.82371520847728,90.82371520857728,90.82637794059703,90.82637794069703,90.831700727504,90.831700727604,90.83435988984706,90.83435988994707,90.83701826683925,90.83701826693925,90.84233409268171,90.84233409278171,90.8449904349012,90.84499043500121,90.84764592037429,90.84764592047429,90.85295542771203,90.85295542781203,90.85826315016143,90.85826315026144,90.86091510155562,90.86091510165562,90.86621786201545,90.86621786211545,90.87151690990753,90.87151691000753,90.87681138848546,90.87681138858547,90.88475025053103,90.88475025063103,90.887395347954,90.887395348054,90.89268486454236,90.89268486464236,90.89532789149484,90.89532789159485,90.90589835720755,90.90589835730755,90.90853988485384,90.90853988495384,90.9111806985448,90.9111806986448,90.91381933467197,90.91381933477197,90.91909517901567,90.91909517911567,90.9243683103291,90.9243683104291,90.93227343798509,90.93227343808509,90.93754082195805,90.93754082205805,90.94017089062122,90.94017089072122,90.94542992131677,90.94542992141677,90.95068716712399,90.95068716722399,90.96382647982988,90.96382647992988,90.96645319290296,90.96645319300296,90.96907747852791,90.96907747862791,90.98219676478665,90.98219676488665,90.98744240881967,90.98744240891968,90.99530759067467,90.99530759077467,90.9979279138475,90.9979279139475,91.00054745166948,91.00054745176948,91.0031630627371,91.0031630628371,91.01362029513368,91.01362029523368,91.02145206387898,91.02145206397898,91.03188698517133,91.03188698527133,91.03710057260982,91.03710057270982,91.03970554574295,91.03970554584295,91.04491038722858,91.04491038732858,91.04750943453243,91.04750943463243,91.05530497004449,91.05530497014449,91.05789951942974,91.05789951952974,91.0630874758717,91.0630874759717,91.068273540332,91.068273540432,91.07086551947805,91.07086551957805,91.07345646338885,91.07345646348885,91.07604622927335,91.07604622937335,91.08122444022499,91.08122444032499,91.09157754223594,91.09157754233594,91.09933837951871,91.09933837961871,91.1019235403913,91.1019235404913,91.10450755893535,91.10450755903535,91.10967295438873,91.10967295448873,91.11225322466727,91.11225322476727,91.11741265859358,91.11741265869358,91.1225695222807,91.1225695223807,91.1251460799915,91.1251460800915,91.13029766040913,91.13029766050913,91.13287150508967,91.13287150518967,91.14059168261612,91.14059168271612,91.14316324263957,91.14316324273958,91.15087645910151,91.15087645920151,91.1534456273746,91.1534456274746,91.15601322494594,91.15601322504594,91.16371466114484,91.16371466124484,91.17141384838445,91.17141384848445,91.17654401014202,91.17654401024203,91.18167181584701,91.18167181594701,91.18423427293995,91.18423427303995,91.19191853851304,91.19191853861304,91.19959598581269,91.1995959859127,91.20215301684507,91.20215301694508,91.2098223250539,91.20982232515391,91.21493278164425,91.21493278174425,91.22004088218198,91.22004088228198,91.22770058769154,91.22770058779155,91.23535586667805,91.23535586677805,91.23790657920571,91.23790657930572,91.24555678910934,91.24555678920935,91.25065339496618,91.25065339506618,91.25320057341494,91.25320057351495,91.25574578848654,91.25574578858654,91.26083514769672,91.26083514779673,91.26592265062304,91.26592265072304,91.26846525975766,91.26846525985766,91.2735494070939,91.2735494071939,91.27863184093734,91.27863184103734,91.28625185053028,91.28625185063028,91.2887903901195,91.2887903902195,91.29386639836491,91.29386639846491,91.30147898282237,91.30147898292238,91.30401563042996,91.30401563052996,91.3065516711755,91.3065516712755,91.30908710505899,91.309087105159,91.3166908364703,91.3166908365703,91.32175584980357,91.32175584990357,91.3268182215021,91.3268182216021,91.33187705912171,91.33187705922171,91.33440460379876,91.33440460389876,91.33692997091204,91.33692997101204,91.34197799210833,91.34197799220833,91.34450000363154,91.34450000373154,91.3470192664267,91.3470192665267,91.35205672108403,91.35205672118403,91.35709214096866,91.35709214106866,91.35960908340898,91.35960908350899,91.36212513340514,91.36212513350515,91.36715541281136,91.36715541291136,91.36966903535938,91.36966903545938,91.37469399579834,91.37469399589834,91.37720522659599,91.37720522669599,91.38222533213867,91.38222533223868,91.38473388560382,91.38473388570382,91.3922564759913,91.3922564760913,91.39476178095967,91.39476178105967,91.39726522964416,91.39726522974416,91.40226991375162,91.40226991385163,91.4047705780103,91.4047705781103,91.40727063540696,91.40727063550696,91.41476884421975,91.41476884431975,91.41726565311963,91.41726565321963,91.41976139108651,91.41976139118651,91.42225655788911,91.42225655798912,91.42974077317734,91.42974077327735,91.4372227038085,91.4372227039085,91.44719455367054,91.44719455377054,91.45217774772242,91.45217774782242,91.45466813102428,91.45466813112428,91.45715790746411,91.45715790756411,91.45964586331783,91.45964586341783,91.46213231986536,91.46213231996536,91.46710359086313,91.46710359096313,91.47207247011053,91.47207247021053,91.4795229153114,91.47952291541141,91.4844884746665,91.48448847476651,91.48697039759764,91.48697039769765,91.48945149948015,91.48945149958016,91.49441195405461,91.49441195415461,91.5043279369117,91.5043279370117,91.50680529052869,91.50680529062869,91.50928150181716,91.50928150191716,91.51423321043875,91.51423321053875,91.51670827939869,91.51670827949869,91.52165731068777,91.52165731078777,91.52660441429745,91.52660441439745,91.52907723429807,91.52907723439807,91.53896519440826,91.53896519450826,91.54637340915107,91.54637340925107,91.54884087448667,91.54884087458667,91.55130751877365,91.55130751887366,91.55377330631423,91.55377330641423,91.5611684556744,91.5611684557744,91.56609464039278,91.56609464049278,91.57347990147157,91.57347990157157,91.58086230672902,91.58086230682902,91.5857804237521,91.5857804238521,91.58823739394428,91.58823739404428,91.59560709079678,91.59560709089678,91.59806231179839,91.59806231189839,91.60297107600658,91.60297107610658,91.61278564150832,91.61278564160833,91.61769112152197,91.61769112162197,91.62014198739604,91.62014198749604,91.62259092559071,91.62259092569072,91.6299365621484,91.6299365622484,91.63727959276913,91.63727959286913,91.64461980326628,91.64461980336628,91.64706499319544,91.64706499329544,91.65439874239681,91.65439874249681,91.65684221883316,91.65684221893316,91.66172831495946,91.66172831505946,91.66905424634992,91.66905424644992,91.67149365324086,91.67149365334086,91.67393238187424,91.67393238197424,91.67880848262585,91.67880848272586,91.68368197744049,91.68368197754049,91.69098816796593,91.69098816806593,91.69342232728516,91.69342232738516,91.69585530857809,91.69585530867809,91.70071909360017,91.70071909370017,91.70558073675626,91.70558073685626,91.7104393099044,91.7104393100044,91.7201514942111,91.7201514943111,91.73228559350304,91.73228559360304,91.73470911488778,91.73470911498778,91.73713181522388,91.73713181532388,91.73955283776495,91.73955283786495,91.74197218251098,91.74197218261098,91.74680826606607,91.74680826616607,91.74922314859124,91.74922314869124,91.75405219968629,91.75405219978629,91.7564655472075,91.7564655473075,91.76370473302474,91.76370473312474,91.76852885782797,91.76852885792798,91.77093909964348,91.77093909974349,91.77334802064162,91.77334802074162,91.77816497019373,91.77816497029373,91.78057142804596,91.78057142814596,91.78297677926741,91.78297677936742,91.78538098816034,91.78538098826034,91.78778459019124,91.78778459029124,91.79739771319521,91.79739771329521,91.79979999440873,91.79979999450873,91.8046037714849,91.80460377158491,91.80940547809061,91.80940547819061,91.81420347212858,91.81420347222858,91.81660054146815,91.81660054156815,91.8213933950277,91.8213933951277,91.82378878657224,91.82378878667224,91.82857817744842,91.82857817754842,91.83097196259345,91.83097196269345,91.83336510517869,91.83336510527869,91.83815042650946,91.83815042660946,91.84054235537062,91.84054235547062,91.84293367736976,91.84293367746976,91.84771489345734,91.84771489355734,91.85249264686156,91.85249264696157,91.85965615341328,91.85965615351328,91.86681609018835,91.86681609028835,91.86919870193239,91.86919870203239,91.87872711413586,91.87872711423586,91.88110772680496,91.88110772690496,91.8858677384191,91.8858677385191,91.89300436455247,91.89300436465247,91.8953813003516,91.8953813004516,91.90013388683025,91.90013388693025,91.90250864506561,91.90250864516561,91.90725662653236,91.90725662663236,91.90962906441287,91.90962906451287,91.9167448073527,91.9167448074527,91.92148721996783,91.92148722006783,91.92622763350802,91.92622763360802,91.93096694041745,91.93096694051745,91.93807386600888,91.93807386610888,91.94281024570142,91.94281024580143,91.94517747170799,91.945177471808,91.95227750763044,91.95227750773044,91.96410610543452,91.96410610553453,91.97119907319916,91.97119907329916,91.98065080659964,91.98065080669964,91.98537283578996,91.98537283588996,91.98773270805658,91.98773270815659,91.99245048921266,91.99245048931266,91.99952285936587,91.99952285946587,92.0089486761877,92.0089486762877,92.01365892511498,92.01365892521498,92.02072072872924,92.02072072882925,92.02307217632291,92.02307217642291,92.02542305275232,92.02542305285232,92.03482463079051,92.03482463089051,92.04187269076456,92.04187269086457,92.04656987430923,92.04656987440923,92.05361229403614,92.05361229413614,92.05830533663985,92.05830533673985,92.06065089410201,92.06065089420201,92.06534075960448,92.06534075970448,92.07002923289406,92.07002923299406,92.07471627827296,92.07471627837296,92.0770588371227,92.07705883722271,92.08408522855234,92.08408522865234,92.08642650228249,92.0864265023825,92.0887671691506,92.0887671692506,92.09578763475096,92.09578763485096,92.09812630254413,92.09812630264413,92.10514037824424,92.10514037834425,92.10747661858927,92.10747661868928,92.10981221637451,92.10981221647451,92.11448184124323,92.11448184134323,92.11914935994372,92.11914936004372,92.12148228039645,92.12148228049645,92.12847943535512,92.12847943545512,92.1378045844747,92.1378045845747,92.1424643567598,92.1424643568598,92.14479267220062,92.14479267230062,92.1517767260789,92.1517767261789,92.1541021500006,92.1541021501006,92.15642632450047,92.15642632460047,92.16107345977615,92.16107345987615,92.17036576695031,92.17036576705031,92.17268776388642,92.17268776398642,92.18197418092907,92.18197418102908,92.18429364332374,92.18429364342374,92.18893071182919,92.18893071192919,92.19356574556193,92.19356574566193,92.19588181666875,92.19588181676875,92.20051335202035,92.20051335212035,92.20282770963436,92.20282770973436,92.20514142468858,92.20514142478858,92.20745389032095,92.20745389042095,92.209764571065,92.209764571165,92.21438329091833,92.21438329101834,92.2166904018857,92.2166904019857,92.22130401695838,92.22130401705839,92.23052889105116,92.23052889115117,92.23283357457039,92.2328335746704,92.23513700866778,92.23513700876778,92.23974212767199,92.23974212777199,92.24204341990337,92.24204342000337,92.24434385538835,92.24434385548835,92.246643576918,92.246643577018,92.24894237030571,92.24894237040571,92.25583660860283,92.25583660870284,92.26502464260034,92.26502464270034,92.2696161607554,92.2696161608554,92.27420596541768,92.27420596551768,92.27879391379608,92.27879391389608,92.2856748368262,92.2856748369262,92.290260429152,92.290260429252,92.29255169031093,92.29255169041093,92.29942415297033,92.29942415307033,92.30400285562715,92.30400285572715,92.30857720315642,92.30857720325642,92.3131470170693,92.3131470171693,92.31771490330279,92.31771490340279,92.32228004080822,92.32228004090823,92.3268432149365,92.3268432150365,92.33368504891205,92.33368504901205,92.33596370875931,92.33596370885931,92.34052024310296,92.34052024320296,92.34507509965158,92.34507509975158,92.35418124297215,92.35418124307215,92.35645661862488,92.35645661872488,92.36100551364645,92.36100551374645,92.3632777478957,92.3632777479957,92.37236418604901,92.37236418614901,92.3791746059898,92.3791746060898,92.38371282737913,92.38371282747913,92.38598045661647,92.38598045671647,92.39278163083567,92.39278163093567,92.3973148902354,92.39731489033541,92.39957969934916,92.39957969944916,92.40637077110038,92.40637077120039,92.41089357105446,92.41089357115446,92.41541283692963,92.41541283702963,92.4199301751254,92.4199301752254,92.42444551424623,92.42444551434623,92.42670200578034,92.42670200588034,92.43121370372896,92.43121370382896,92.43346841037473,92.43346841047473,92.44023103100584,92.44023103110584,92.44699150977095,92.44699150987095,92.44924261094229,92.4492426110423,92.45374395653855,92.45374395663855,92.45599284444833,92.45599284454833,92.45823969758543,92.45823969768543,92.46497929315699,92.46497929325699,92.46947085756516,92.46947085766516,92.47396038720062,92.47396038730062,92.47620120741513,92.47620120751513,92.48292252573012,92.48292252583012,92.48740156022215,92.48740156032216,92.48963898914883,92.48963898924883,92.49411238339371,92.49411238349371,92.49634749196555,92.49634749206555,92.50081649538512,92.50081649548513,92.50304964057979,92.50304964067979,92.50751478864056,92.50751478874056,92.50974636313347,92.50974636323348,92.5119772950666,92.5119772951666,92.51420740595107,92.51420740605107,92.52089527545861,92.52089527555862,92.52757746902124,92.52757746912124,92.52980183256527,92.52980183266527,92.53202423273214,92.53202423283214,92.53868957694885,92.53868957704886,92.54757110992102,92.54757111002102,92.54978958333355,92.54978958343355,92.55866240605063,92.55866240615063,92.56309542612132,92.56309542622132,92.57195646857538,92.57195646867538,92.57417008709162,92.57417008719162,92.58302170533526,92.58302170543526,92.58523218244802,92.58523218254803,92.58965245841598,92.58965245851599,92.59186168610692,92.59186168620693,92.59406902181621,92.59406902191621,92.59848222962637,92.59848222972637,92.60068699509645,92.60068699519645,92.60289076102907,92.60289076112907,92.60729725765907,92.60729725775907,92.61170239777394,92.61170239787394,92.61390209416115,92.61390209426115,92.61610107659303,92.61610107669303,92.62049743505727,92.62049743515728,92.62269438271645,92.62269438281645,92.62708692151965,92.62708692161965,92.63147746124791,92.63147746134791,92.63586396712853,92.63586396722853,92.6424357474986,92.6424357475986,92.64900031691978,92.64900031701978,92.65556306575485,92.65556306585485,92.65774951827059,92.6577495183706,92.66212117388025,92.66212117398025,92.66430516325009,92.6643051633501,92.67303947863218,92.67303947873218,92.67740435166614,92.67740435176614,92.68176629748324,92.68176629758324,92.6839433971841,92.6839433972841,92.68829681123495,92.68829681133495,92.69264829760638,92.69264829770638,92.69699849885822,92.69699849895822,92.6991721358757,92.6991721359757,92.71003707246633,92.71003707256634,92.71437988428045,92.71437988438045,92.7187212681839,92.7187212682839,92.72306143836327,92.72306143846328,92.72739968086326,92.72739968096326,92.73607323864633,92.73607323874633,92.73824034297249,92.73824034307249,92.74473969257382,92.74473969267382,92.7512357579806,92.7512357580806,92.75556421929248,92.75556421939248,92.75989053873836,92.75989053883836,92.76421482341154,92.76421482351154,92.76853800145395,92.76853800155395,92.77502114426916,92.77502114436916,92.77934146649024,92.77934146659024,92.78581964731588,92.78581964741588,92.78797636359191,92.78797636369191,92.7944445133451,92.7944445134451,92.79659858798641,92.79659858808641,92.80305891992867,92.80305892002868,92.80951553930319,92.8095155394032,92.81812095004955,92.81812095014955,92.82457135801265,92.82457135811265,92.8331676301307,92.8331676302307,92.84175847618822,92.84175847628822,92.84605252485296,92.84605252495297,92.85463687391692,92.85463687401692,92.8589261390809,92.85892613918091,92.86535805560085,92.86535805570085,92.86750006439699,92.867500064497,92.87178301105627,92.87178301115627,92.8739237347328,92.8739237348328,92.8803446206428,92.8803446207428,92.88676347178011,92.88676347188012,92.88890098265763,92.88890098275763,92.89317464789755,92.89317464799755,92.89531033818898,92.89531033828898,92.89958093342096,92.89958093352097,92.90171466033522,92.90171466043522,92.91024774740613,92.91024774750613,92.91238004640971,92.91238004650971,92.91451173855125,92.91451173865126,92.91877437318125,92.91877437328125,92.92090442322554,92.92090442332554,92.92303386640778,92.92303386650778,92.92516252423916,92.92516252433916,92.93367626312047,92.93367626322048,92.93793125842839,92.9379312585284,92.94431159167537,92.94431159177537,92.95068817665684,92.95068817675684,92.9528111585433,92.9528111586433,92.95493196286598,92.95493196296599,92.95705205323334,92.95705205333334,92.96129052047525,92.96129052057525,92.96764386621057,92.96764386631057,92.97187683599638,92.97187683609638,92.97822407741359,92.97822407751359,92.98245129985897,92.98245129995897,92.98667538090086,92.98667538100086,92.98878558298684,92.98878558308684,92.99089432146435,92.99089432156435,92.99300181052004,92.99300181062004,92.9951076217807,92.9951076218807,92.99931799488017,92.99931799498017,93.00352690437121,93.00352690447122,93.0077339218806,93.0077339219806,93.00983602057352,93.00983602067352,93.01193747670662,93.01193747680662,93.01613906815548,93.01613906825548,93.02033905320482,93.02033905330482,93.02453764604127,93.02453764614127,93.03083276871901,93.03083276881901,93.03502711352121,93.03502711362121,93.04341369695737,93.04341369705737,93.0517954968928,93.0517954969928,93.05807737569907,93.05807737579907,93.06226279605958,93.06226279615959,93.06853807077901,93.06853807087901,93.07062828211559,93.0706282822156,93.07689730972584,93.07689730982584,93.08107501936874,93.08107501946874,93.08524948051485,93.08524948061485,93.08733515823506,93.08733515833507,93.09359054929841,93.09359054939841,93.09775897752195,93.09775897762195,93.10192465701745,93.10192465711745,93.10608815894918,93.10608815904918,93.11649366528172,93.11649366538173,93.12065277638813,93.12065277648813,93.12480935295311,93.12480935305311,93.12688671309367,93.12688671319367,93.12896268102132,93.12896268112132,93.13311333175702,93.13311333185702,93.13933766576328,93.13933766586328,93.1434851750955,93.14348517519551,93.14970515397424,93.14970515407424,93.15177748072968,93.15177748082968,93.15799228343224,93.15799228353224,93.16213386693518,93.16213386703518,93.16834360055485,93.16834360065485,93.17041225044031,93.17041225054031,93.17454805090505,93.17454805100505,93.17868063857077,93.17868063867077,93.18281112006824,93.18281112016824,93.1848749507552,93.1848749508552,93.18900125561397,93.18900125571398,93.19312541860674,93.19312541870674,93.19518592940139,93.19518592950139,93.19930584435991,93.19930584445991,93.20754260426902,93.20754260436902,93.21165684328263,93.21165684338263,93.21576690565753,93.21576690575753,93.21782099085416,93.21782099095417,93.22192819740772,93.22192819750772,93.22808663333657,93.22808663343658,93.23219051999783,93.23219052009783,93.23424160658205,93.23424160668205,93.23629065839357,93.23629065849357,93.2444852949379,93.2444852950379,93.2485797930865,93.24857979318651,93.2526708642495,93.2526708643495,93.25880420464831,93.25880420474832,93.26084736632832,93.26084736642832,93.26493176200893,93.26493176210893,93.26901380163693,93.26901380173693,93.27513396955983,93.27513396965983,93.27717088413065,93.27717088423066,93.28328016423468,93.28328016433468,93.28531515112611,93.28531515122611,93.28734863871132,93.28734863881132,93.2934476735563,93.2934476736563,93.2995436740911,93.2995436741911,93.30157455573935,93.30157455583935,93.30360440215236,93.30360440225236,93.30563349891226,93.30563349901226,93.309690478708,93.30969047880801,93.3117174336019,93.3117174337019,93.313743745936,93.313743746036,93.31779522827566,93.31779522837566,93.31981939874376,93.31981939884376,93.32386670444471,93.32386670454471,93.32791261793275,93.32791261803276,93.33195728199897,93.33195728209897,93.3380225467566,93.3380225468566,93.34206385523274,93.34206385533274,93.34408284952465,93.34408284962466,93.34811941019781,93.34811941029781,93.35215279376973,93.35215279386973,93.35618317872924,93.35618317882924,93.35819642568072,93.35819642578072,93.36222188434841,93.36222188444842,93.36825791263507,93.36825791273507,93.37026855364957,93.37026855374957,93.37830908293485,93.37830908303485,93.38232779472463,93.38232779482463,93.38835418461427,93.38835418471427,93.39236946941844,93.39236946951844,93.39638218398342,93.39638218408342,93.39838700626193,93.39838700636193,93.40039104318957,93.40039104328957,93.40439751064537,93.40439751074537,93.40840144355971,93.40840144365971,93.41240219937282,93.41240219947282,93.41640002796906,93.41640002806906,93.42039492934842,93.42039492944842,93.42239011322991,93.42239011332991,93.42438419048064,93.42438419058064,93.43435311312581,93.43435311322581,93.43833830471262,93.43833830481262,93.44431400377347,93.44431400387347,93.44630329752344,93.44630329762344,93.45028077839262,93.45028077849263,93.45624516126142,93.45624516136142,93.45823177767889,93.45823177777889,93.46220411806966,93.46220411816967,93.4681603975454,93.4681603976454,93.47411296445341,93.47411296455341,93.48006342519317,93.48006342529317,93.4840280905641,93.4840280906641,93.48600885254781,93.48600885264781,93.4879885792963,93.4879885793963,93.49392526069809,93.4939252607981,93.49590113208778,93.49590113218778,93.49787639661542,93.49787639671543,93.50182471240917,93.50182471250918,93.50774361632313,93.50774361642313,93.5116878268737,93.5116878269737,93.51957221412721,93.51957221422721,93.52154112445237,93.52154112455237,93.52744657030826,93.52744657040826,93.52941298178975,93.52941298188975,93.53137875071143,93.53137875081143,93.53530925331957,93.53530925341957,93.53727341584175,93.53727341594175,93.54120102693079,93.5412010270308,93.54905285782102,93.54905285792103,93.55690201137874,93.55690201147874,93.56082416070947,93.56082416080947,93.56278346833537,93.56278346843537,93.57257800738996,93.57257800748997,93.57845250328208,93.57845250338208,93.58236622793986,93.58236622803986,93.58627809631375,93.58627809641375,93.59018860817251,93.59018860827251,93.59214273962223,93.59214273972223,93.59409540746353,93.59409540756353,93.597999993493,93.597999993593,93.60190315161182,93.60190315171182,93.60385360619158,93.60385360629158,93.60580313262939,93.60580313272939,93.60970072189659,93.60970072199659,93.61749218786323,93.61749218796324,93.62138477944309,93.62138477954309,93.6252762286944,93.6252762287944,93.62916635712836,93.62916635722836,93.63305309427446,93.63305309437446,93.636937011297,93.636937011397,93.63887782747972,93.63887782757972,93.64275667541938,93.64275667551938,93.64663223916449,93.64663223926449,93.6505055539503,93.6505055540503,93.65631311652974,93.65631311662975,93.66211846584767,93.66211846594767,93.66792028108667,93.66792028118667,93.67178674190126,93.67178674200126,93.67758470178146,93.67758470188146,93.68531205455903,93.68531205465904,93.68724226850503,93.68724226860503,93.69303126824578,93.69303126834578,93.69496005428111,93.69496005438111,93.69881619844111,93.69881619854111,93.70652641629064,93.70652641639064,93.70845277487783,93.70845277497783,93.71230374286165,93.71230374296165,93.71615321153926,93.71615321163927,93.71807585755872,93.71807585765872,93.72191990017579,93.7219199002758,93.72576019452735,93.72576019462736,93.73151410336338,93.73151410346338,93.73534765083703,93.73534765093703,93.73917916353797,93.73917916363797,93.74300860576844,93.74300860586844,93.74683672718153,93.74683672728153,93.75257607132873,93.75257607142873,93.75448675859633,93.75448675869633,93.76021749958173,93.76021749968173,93.76403498306034,93.76403498316034,93.76594013717408,93.76594013727409,93.7678445773325,93.7678445774325,93.77165231532078,93.77165231542078,93.77355486349755,93.77355486359755,93.784967118248,93.784967118348,93.79256824372234,93.79256824382234,93.7944672578202,93.7944672579202,93.80206088676353,93.80206088686353,93.80585448843618,93.80585448853618,93.80964655510486,93.80964655520486,93.81533038922475,93.81533038932476,93.81722247795587,93.81722247805587,93.81911353145175,93.81911353155175,93.82478415739794,93.82478415749794,93.82667185530374,93.82667185540375,93.82855891064975,93.82855891074975,93.83233180761768,93.83233180771768,93.83798783317728,93.83798783327728,93.83987139014214,93.83987139024214,93.843637718721,93.843637718821,93.84551934800645,93.84551934810645,93.8474003347321,93.8474003348321,93.84928064320019,93.84928064330019,93.85303979652792,93.85303979662793,93.85679727206063,93.85679727216063,93.8586742606364,93.85867426073641,93.86242659569069,93.8624265957907,93.86617678887897,93.86617678897898,93.86805027907361,93.86805027917362,93.87554241926605,93.87554241936606,93.87928607976302,93.87928607986302,93.88115625006535,93.88115625016535,93.88302513524808,93.88302513534808,93.88489291380003,93.88489291390003,93.88862697159774,93.88862697169775,93.89235981567138,93.89235981577139,93.8979562259605,93.8979562260605,93.89982064892239,93.89982064902239,93.90354838821538,93.90354838831539,93.90727487808654,93.90727487818654,93.91099886911402,93.91099886921403,93.91285940101933,93.91285940111933,93.9147187905961,93.9147187906961,93.92029521013585,93.92029521023585,93.92400984834845,93.92400984844845,93.92772191632183,93.92772191642183,93.93143230650018,93.93143230660019,93.93328644850524,93.93328644860524,93.93513969806614,93.93513969816614,93.93884505485937,93.93884505495937,93.94254880525311,93.94254880535311,93.94625066366521,93.94625066376521,93.94995102277112,93.94995102287112,93.9536496683842,93.9536496684842,93.9591967086619,93.95919670876191,93.96289264124472,93.96289264134472,93.97028257873096,93.97028257883096,93.97397594107457,93.97397594117457,93.97582155131337,93.97582155141338,93.98135641865262,93.98135641875263,93.98504232016299,93.98504232026299,93.9887257942252,93.9887257943252,93.99240784037676,93.99240784047676,94.00160865417475,94.00160865427475,94.00344741044233,94.00344741054234,94.00712295960035,94.00712295970035,94.01630703114573,94.01630703124573,94.01997897482929,94.01997897492929,94.02731982788625,94.02731982798625,94.03649194067977,94.03649194077977,94.03832502100244,94.03832502110244,94.04015645922784,94.04015645932785,94.04198704070684,94.04198704080684,94.04564649017205,94.04564649027205,94.04930162020747,94.04930162030747,94.0566049192138,94.0566049193138,94.06025480167752,94.06025480177752,94.06207785092774,94.06207785102774,94.06754474971908,94.06754474981908,94.0711864573942,94.0711864574942,94.07664757314734,94.07664757324734,94.08210601156796,94.08210601166796,94.08392327777997,94.08392327787998,94.08755645368883,94.08755645378884,94.0930018624246,94.0930018625246,94.09662893401536,94.09662893411536,94.09844059567799,94.09844059577799,94.1002511864076,94.1002511865076,94.10206042062208,94.10206042072208,94.10567660439399,94.105676604494,94.10748319697373,94.10748319707373,94.10928903990037,94.10928904000038,94.11289961912289,94.11289961922289,94.11470385565003,94.11470385575004,94.12011556569398,94.12011556579398,94.12372146850906,94.12372146860906,94.12912725272372,94.12912725282372,94.13632920577025,94.13632920587025,94.14172742205837,94.14172742215837,94.14712242554748,94.14712242564748,94.15071387076702,94.15071387086702,94.15609648713107,94.15609648723107,94.15788955025721,94.15788955035721,94.16326716893391,94.16326716903392,94.16685022517827,94.16685022527827,94.17222138255919,94.17222138265919,94.17401037613992,94.17401037623992,94.17579794180999,94.17579794190999,94.17937185942606,94.17937185952606,94.18115771160332,94.18115771170332,94.1865140901088,94.1865140902088,94.19186850523711,94.19186850533711,94.1936523940372,94.1936523941372,94.19721892221555,94.19721892231556,94.2007836298078,94.2007836299078,94.20434640972063,94.20434640982063,94.21681192707909,94.2168119271791,94.22037028046886,94.22037028056886,94.22392602792164,94.22392602802164,94.23103298921083,94.23103298931083,94.24168720188433,94.24168720198433,94.24701070274665,94.24701070284665,94.24878327535467,94.24878327545467,94.25587228066719,94.25587228076719,94.25941496273735,94.25941496283735,94.26295607410577,94.26295607420577,94.26649525779479,94.26649525789479,94.26826244004005,94.26826244014005,94.27002844425898,94.27002844435899,94.2717937345226,94.2717937346226,94.27355816803981,94.27355816813981,94.27884996928525,94.27884996938525,94.28061304628733,94.28061304638733,94.28590092077842,94.28590092087842,94.28766253417206,94.28766253427206,94.29118486851516,94.29118486861516,94.29470541796994,94.29470541806994,94.29646488949757,94.29646488959757,94.29822325439443,94.29822325449443,94.29998083394042,94.29998083404043,94.30173784232215,94.30173784242216,94.3052510023392,94.3052510024392,94.30700454803757,94.30700454813757,94.31050996163923,94.31050996173923,94.31226129407604,94.31226129417604,94.31401205534858,94.31401205544859,94.31576195987473,94.31576195997474,94.31926051950519,94.3192605196052,94.32450509260521,94.32450509270521,94.33149464393956,94.33149464403957,94.3384787335156,94.3384787336156,94.34545753982216,94.34545753992217,94.3472012329369,94.3472012330369,94.35068747683782,94.35068747693782,94.35242949215751,94.35242949225751,94.3541704365442,94.3541704366442,94.35765107589575,94.35765107599575,94.36113060861652,94.36113060871652,94.3628694111372,94.3628694112372,94.36460764249362,94.36460764259363,94.36808321276229,94.36808321286229,94.36982019469686,94.36982019479686,94.37155603430291,94.37155603440291,94.37502546455569,94.37502546465569,94.3784922888715,94.3784922889715,94.38368945533728,94.38368945543728,94.38715146045458,94.38715146055458,94.39061025277287,94.39061025287288,94.3992518610547,94.3992518611547,94.4044333205031,94.4044333206031,94.4078851160591,94.40788511615911,94.41133416288709,94.41133416298709,94.41822836548644,94.41822836558644,94.42339583141028,94.42339583151028,94.42511721342079,94.42511721352079,94.42855912069541,94.42855912079541,94.43199863621966,94.43199863631966,94.4371551429291,94.4371551430291,94.4405911600722,94.4405911601722,94.4440246069761,94.4440246070761,94.44574034873946,94.44574034883946,94.45088607472336,94.45088607482336,94.45603058698318,94.45603058708318,94.45945782247566,94.45945782257566,94.46630936624375,94.46630936634375,94.46973313905285,94.46973313915285,94.47144338336014,94.47144338346014,94.47315294940985,94.47315294950985,94.47486180150423,94.47486180160423,94.47998667999234,94.47998668009234,94.48340099289325,94.48340099299325,94.49022554914968,94.49022554924969,94.49363497145656,94.49363497155656,94.49533889725913,94.49533889735913,94.49704210910636,94.49704210920636,94.50044735477452,94.50044735487452,94.50214892452449,94.50214892462449,94.50555110018472,94.50555110028472,94.50725117062848,94.50725117072848,94.51064881267233,94.51064881277233,94.51234552752601,94.51234552762601,94.51743442266523,94.51743442276523,94.52082403271156,94.52082403281156,94.52251692790419,94.5225169280042,94.52928672378643,94.52928672388643,94.53267008672357,94.53267008682357,94.53774309635659,94.5377430964566,94.54112271102822,94.54112271112822,94.5445003623227,94.5445003624227,94.5495646973984,94.5495646974984,94.55800097212587,94.55800097222587,94.55968679916069,94.55968679926069,94.56474153153714,94.56474153163714,94.56979105203962,94.56979105213962,94.57483818079174,94.57483818089175,94.57820073193088,94.57820073203088,94.5815617837638,94.5815617838638,94.58827896113785,94.58827896123785,94.59163265923083,94.59163265933083,94.59330806251775,94.59330806261775,94.59498178800965,94.59498178810965,94.59832773968724,94.59832773978724,94.60334452533762,94.60334452543762,94.60668719282067,94.60668719292067,94.60835749132696,94.60835749142696,94.61169730298867,94.61169730308868,94.61503550825088,94.61503550835089,94.61837232130019,94.61837232140019,94.62170792062541,94.62170792072541,94.62837708450317,94.62837708460317,94.63504246441767,94.63504246451767,94.63837249489129,94.63837249499129,94.64502980711048,94.64502980721048,94.65168329966863,94.65168329976864,94.65999363267503,94.65999363277503,94.66995898554352,94.66995898564352,94.67161811064882,94.67161811074882,94.67493553981078,94.67493553991078,94.67659298712103,94.67659298722103,94.68322074158937,94.68322074168937,94.6865324948064,94.6865324949064,94.68984199906414,94.68984199916414,94.6931487902916,94.6931487903916,94.6964526543022,94.69645265440221,94.70635924864663,94.70635924874664,94.70965922160065,94.70965922170065,94.71460514718402,94.71460514728402,94.71789823046906,94.71789823056906,94.71954368332969,94.71954368342969,94.72283376800232,94.72283376810232,94.72776727076287,94.72776727086287,94.73269856026188,94.73269856036188,94.73762727952167,94.73762727962168,94.74255471366189,94.74255471376189,94.74583808715435,94.74583808725436,94.75076102337596,94.75076102347596,94.75568235319804,94.75568235329804,94.75896094318978,94.75896094328978,94.7622372842222,94.7622372843222,94.76551084082882,94.76551084092883,94.7687820770806,94.7687820771806,94.77532162236729,94.77532162246729,94.78022253306679,94.78022253316679,94.78348777209376,94.78348777219377,94.78838314963942,94.78838314973942,94.79001389502015,94.79001389512015,94.79490420348293,94.79490420358293,94.79653302118426,94.79653302128426,94.80141818916863,94.80141818926863,94.80467325433207,94.80467325443207,94.80792703437591,94.80792703447591,94.8144322027132,94.81443220281321,94.81605737924232,94.81605737934233,94.8193065185765,94.8193065186765,94.82255462267541,94.82255462277541,94.82742479759774,94.82742479769774,94.82904636865243,94.82904636875243,94.832287333198,94.832287333298,94.8371442285511,94.8371442286511,94.84038005261827,94.84038005271827,94.84361323505068,94.84361323515068,94.84684252642654,94.84684252652654,94.85168085894092,94.85168085904093,94.86135256198014,94.86135256208014,94.86618643227368,94.86618643237368,94.86940726327882,94.86940726337882,94.87423670704928,94.87423670714928,94.87906279522967,94.87906279532967,94.88227816447649,94.88227816457649,94.88388481386467,94.88388481396467,94.88709697031247,94.88709697041247,94.89030759175631,94.89030759185631,94.89672508637848,94.89672508647848,94.89993092432157,94.89993092442157,94.90473559384192,94.90473559394192,94.90793486339592,94.90793486349592,94.91113199108393,94.91113199118394,94.91272896637732,94.91272896647732,94.91432522771535,94.91432522781535,94.91592088219137,94.91592088229137,94.92070663189534,94.92070663199534,94.923895549097,94.923895549197,94.92867730065109,94.92867730075109,94.93186375470682,94.93186375480683,94.9350482453854,94.9350482454854,94.9366386522897,94.9366386523897,94.93981778830326,94.93981778840326,94.94299367582005,94.94299367592005,94.94616656472444,94.94616656482444,94.9509233278418,94.95092332794181,94.95409214720077,94.95409214730077,94.95567570013385,94.95567570023385,94.95884166367149,94.9588416637715,94.96358953804493,94.96358953814493,94.96517134178744,94.96517134188744,94.96675246727239,94.96675246737239,94.97149427302548,94.97149427312549,94.9746527400321,94.9746527401321,94.97938829867601,94.97938829877602,94.98254401695459,94.9825440170546,94.98412050172986,94.98412050182986,94.98727225755631,94.98727225765631,94.99042251407658,94.99042251417659,94.99514547140883,94.99514547150883,94.99986418070684,94.99986418080684,95.00615009336076,95.00615009346076,95.0092908185773,95.0092908186773,95.01085980682154,95.01085980692154,95.01399689086588,95.01399689096588,95.01556373724412,95.01556373734412,95.02026320544586,95.02026320554586,95.02182848112236,95.02182848122236,95.02339300714577,95.02339300724577,95.02495574828083,95.02495574838083,95.0265175255762,95.0265175256762,95.03120096548065,95.03120096558065,95.03432109308578,95.03432109318578,95.0374394001048,95.0374394002048,95.04055585083995,95.04055585093995,95.04211263044796,95.04211263054796,95.0452247974511,95.0452247975511,95.0514460971472,95.05144609724721,95.05300009232941,95.05300009242941,95.05766089984975,95.05766089994975,95.06231974399293,95.06231974409293,95.06697666045669,95.06697666055669,95.06852772842203,95.06852772852203,95.0700782252231,95.0700782253231,95.0731783977766,95.0731783978766,95.07937599415558,95.07937599425559,95.08092420629958,95.08092420639959,95.08401891709478,95.08401891719478,95.08865791327963,95.08865791337963,95.09020259134472,95.09020259144472,95.09483548321147,95.09483548331147,95.09792244759127,95.09792244769127,95.09946476960376,95.09946476970376,95.10409023633501,95.10409023643501,95.1087128829427,95.1087128830427,95.11025274180928,95.11025274190928,95.1133314243072,95.1133314244072,95.11486921270331,95.11486921280331,95.11794307600272,95.11794307610272,95.11947768729759,95.11947768739759,95.12254598174539,95.12254598184539,95.12714563899128,95.12714563909128,95.12867807272238,95.12867807282238,95.1302097211026,95.1302097212026,95.13327205402337,95.13327205412337,95.13480209600408,95.13480209610408,95.13786110903253,95.13786110913253,95.13938993728918,95.13938993738918,95.14091790879944,95.14091790889944,95.14702768867221,95.14702768877221,95.14855373250305,95.14855373260305,95.15313097155142,95.15313097165142,95.15618077455605,95.15618077465605,95.16075347998805,95.16075348008805,95.16380014158919,95.1638001416892,95.16836720677405,95.16836720687405,95.16988718198455,95.16988718208455,95.1729262042636,95.1729262043636,95.17596376293423,95.17596376303423,95.17899928683215,95.17899928693215,95.18658534784892,95.18658534794892,95.1896176946456,95.1896176947456,95.19113270786653,95.19113270796653,95.1956764624097,95.1956764625097,95.19718926236906,95.19718926246907,95.20172641282538,95.20172641292538,95.20474962099378,95.20474962109378,95.206260457576,95.206260457676,95.20777058020286,95.20777058030286,95.21078989731468,95.21078989741468,95.21380817919126,95.21380817929126,95.21682499745941,95.21682499755941,95.21984042351465,95.21984042361466,95.22285356491282,95.22285356501283,95.22586420746731,95.22586420756731,95.23037783309533,95.23037783319533,95.23338479877987,95.23338479887987,95.23639026515818,95.23639026525818,95.23939426792806,95.23939426802806,95.24239687848504,95.24239687858504,95.24839927947542,95.24839927957542,95.25289937564985,95.25289937574985,95.25439857476009,95.25439857486009,95.25739572355876,95.25739572365876,95.2588936375494,95.25889363764941,95.26039098037579,95.26039098047579,95.2663789594684,95.2663789595684,95.26937078929981,95.26937078939982,95.27385553543454,95.27385553553454,95.27833835388986,95.27833835398987,95.28132572150044,95.28132572160044,95.28729777938908,95.28729777948908,95.29028221978278,95.29028221988278,95.294757398916,95.294757399016,95.29624610288283,95.29624610298283,95.3007113580369,95.30071135813691,95.30517332899645,95.30517332909645,95.30963251553017,95.30963251563017,95.3126038191457,95.31260381924571,95.31557358775727,95.31557358785727,95.31854107171178,95.31854107181178,95.32002245763641,95.32002245773641,95.32890788166516,95.32890788176516,95.33038737560815,95.33038737570816,95.33482417964214,95.33482417974214,95.337779669147,95.337779669247,95.3392562715709,95.3392562716709,95.34220836978793,95.34220836988793,95.34958467072725,95.34958467082726,95.35547840771565,95.35547840781565,95.3569508691986,95.3569508692986,95.35842258102845,95.35842258112845,95.36283614581627,95.36283614591628,95.36577578551271,95.36577578561271,95.36724432024131,95.36724432034131,95.37018049725437,95.37018049735437,95.37311474658802,95.37311474668802,95.37604731812664,95.37604731822664,95.38044274844894,95.38044274854894,95.38629661103187,95.38629661113187,95.3892202581288,95.3892202582288,95.39360248027741,95.39360248037741,95.4023623552605,95.4023623553605,95.40382107310324,95.40382107320325,95.40673633122498,95.40673633132498,95.41256138571013,95.41256138581014,95.41401667656729,95.41401667666729,95.41838062145932,95.41838062155932,95.4198341631259,95.4198341632259,95.42274013982828,95.42274013992828,95.42419182521098,95.42419182531098,95.42709366097243,95.42709366107243,95.42854281181368,95.42854281191369,95.42999049904653,95.42999049914653,95.43288505246355,95.43288505256355,95.43722475857196,95.43722475867196,95.44300922469631,95.44300922479631,95.44734511114369,95.44734511124369,95.45456759040609,95.45456759050609,95.45745471868763,95.45745471878763,95.45889708695321,95.45889708705322,95.46178039557371,95.46178039567371,95.46610300245185,95.46610300255185,95.46754190803405,95.46754190813405,95.4704188267543,95.4704188268543,95.47185644721691,95.47185644731691,95.47473090279125,95.47473090289125,95.47760371626832,95.47760371636832,95.48047510183471,95.48047510193472,95.48334448832618,95.48334448842618,95.4847780035456,95.4847780036456,95.491944008941,95.491944009041,95.49624098482262,95.49624098492262,95.49910358873841,95.49910358883841,95.5019646576502,95.50196465775021,95.50482394167368,95.50482394177368,95.50768169069319,95.50768169079319,95.51625229611695,95.51625229621695,95.51910733210617,95.51910733220618,95.52338813689944,95.52338813699944,95.5276662643602,95.52766626446021,95.5305170523152,95.5305170524152,95.53479046767075,95.53479046777075,95.53763661491605,95.53763661501605,95.54190431862894,95.54190431872894,95.54474775284396,95.54474775294396,95.54616845256513,95.54616845266513,95.55042908812018,95.55042908822018,95.55326531138634,95.55326531148634,95.55751355981633,95.55751355991633,95.56034382155543,95.56034382165544,95.56175809567858,95.56175809577859,95.56317169154417,95.56317169164417,95.56458435926781,95.56458435936781,95.56599645582719,95.56599645592719,95.56740769564017,95.56740769574017,95.57022878305321,95.57022878315321,95.57304826406676,95.57304826416676,95.57868479864571,95.57868479874571,95.58431826321672,95.58431826331672,95.58572554057758,95.58572554067759,95.58713167560991,95.58713167570991,95.59134801023643,95.59134801033643,95.59275203910052,95.59275203920052,95.5969629476665,95.5969629477665,95.59836408501148,95.59836408511148,95.59976425851677,95.59976425861677,95.60256339180326,95.60256339190326,95.60396170902467,95.60396170912468,95.60675755811663,95.60675755821663,95.61094783337342,95.61094783347342,95.61373825640483,95.61373825650483,95.61652693024567,95.61652693034567,95.61792010698866,95.61792010708866,95.6193119629143,95.6193119630143,95.6234854602207,95.62348546032071,95.62626535241093,95.62626535251093,95.6290436738994,95.6290436739994,95.63043135318632,95.63043135328633,95.63320542664056,95.63320542674056,95.63597782229976,95.63597782239977,95.63874796899965,95.63874796909965,95.6401314359501,95.6401314360501,95.64151404615414,95.64151404625414,95.64704280917528,95.64704280927528,95.65118664163751,95.65118664173751,95.65256689578895,95.65256689588895,95.65670555207502,95.65670555217503,95.66221925063857,95.66221925073857,95.66359518536024,95.66359518546024,95.66772138312571,95.66772138322571,95.67047011116556,95.67047011126556,95.67321605477959,95.67321605487959,95.67458813414244,95.67458813424244,95.67595964234101,95.67595964244101,95.67870194478284,95.67870194488285,95.68144278361623,95.68144278371624,95.68555227482688,95.68555227492688,95.68829015074562,95.68829015084562,95.69376397490372,95.69376397500372,95.6951315206502,95.6951315207502,95.69923240869903,95.69923240879903,95.70333033383324,95.70333033393324,95.7074259386126,95.7074259387126,95.71288434133547,95.71288434143547,95.71697605505825,95.71697605515826,95.72106509144854,95.72106509154854,95.72515216446166,95.72515216456166,95.7333219910581,95.7333219911581,95.73876632886086,95.73876632896086,95.74284722616034,95.74284722626034,95.74556582528504,95.74556582538504,95.74692380403002,95.74692380413002,95.74828117591296,95.74828117601297,95.75099484874586,95.75099484884586,95.75642051661661,95.75642051671662,95.7591312979304,95.7591312980304,95.76048552840987,95.76048552850988,95.76454675623985,95.76454675633985,95.76725239707523,95.76725239717523,95.77266028745814,95.77266028755814,95.77536289398334,95.77536289408334,95.7780644652733,95.7780644653733,95.77941412643864,95.77941412653864,95.78346186051279,95.78346186061279,95.78615861260424,95.78615861270424,95.78750588201919,95.78750588211919,95.79154665502884,95.79154665512884,95.79423915908605,95.79423915918605,95.79962074021485,95.79962074031485,95.8036538739024,95.8036538740024,95.80499700237642,95.80499700247643,95.80768247397359,95.80768247407359,95.81305059704432,95.81305059714433,95.81573178490947,95.81573178500948,95.81975078228139,95.81975078238139,95.82644536296908,95.82644536306908,95.83179745774255,95.83179745784255,95.83313416061856,95.83313416071856,95.83714312691802,95.83714312701802,95.83847779502132,95.83847779512132,95.8398118205648,95.8398118206648,95.84247833664782,95.84247833674782,95.8451430678425,95.8451430679425,95.84780662101088,95.84780662111088,95.84913761224419,95.8491376123442,95.85046753254451,95.85046753264452,95.85578560734628,95.85578560744628,95.85844123561043,95.85844123571043,95.8637484582911,95.8637484583911,95.86507380927726,95.86507380937726,95.86772358310768,95.86772358320768,95.87037128646762,95.87037128656762,95.87434087813035,95.87434087823036,95.88095346104781,95.88095346114781,95.88227474248858,95.88227474258858,95.88491659141476,95.88491659151477,95.88755622707941,95.88755622717942,95.8888743671167,95.8888743672167,95.89282735931789,95.8928273594179,95.89546128333987,95.89546128343987,95.89809402933554,95.89809402943554,95.90204152408066,95.90204152418066,95.90467159274381,95.90467159284381,95.9072994838432,95.9072994839432,95.91255251731394,95.91255251741394,95.9151768029389,95.9151768030389,95.91779891100008,95.91779891110008,95.91910912613314,95.91910912623314,95.92041852021758,95.92041852031758,95.92434545304904,95.92434545314904,95.92696088562784,95.92696088572784,95.9308816070479,95.9308816071479,95.93610543974543,95.93610543984543,95.9439291050977,95.9439291051977,95.94783833183685,95.94783833193685,95.94914026508803,95.94914026518803,95.95044155577942,95.95044155587942,95.9517419897244,95.9517419898244,95.95304170971404,95.95304170981404,95.95953791791189,95.95953791801189,95.9673286699232,95.9673286700232,95.97251966067533,95.97251966077533,95.97511360319855,95.97511360329855,95.97900208953521,95.97900208963522,95.98159303344602,95.98159303354602,95.98288772005056,95.98288772015056,95.98418179979305,95.98418179989305,95.98547484430031,95.98547484440031,95.99064520174326,95.99064520184326,95.99581134684973,95.99581134694974,95.99710132134905,95.99710132144905,95.99839058189303,95.99839058199304,96.00483499263133,96.00483499273133,96.00612236119369,96.00612236129369,96.00869638436305,96.00869638446305,96.01255502736672,96.01255502746672,96.01769543438336,96.01769543448336,96.02154690213591,96.02154690223591,96.02539579964926,96.02539579974926,96.02924198413234,96.02924198423234,96.03564976896195,96.03564976906195,96.03949213378398,96.03949213388398,96.04333246383331,96.04333246393331,96.0484499528837,96.0484499529837,96.05484338721106,96.05484338731107,96.05739899033279,96.05739899043279,96.06250848308342,96.06250848318342,96.06506073061507,96.06506073071508,96.07016001380441,96.07016001390441,96.07653117272534,96.07653117282534,96.07907806559197,96.07907806569197,96.0828960488393,96.0828960489393,96.09052712473995,96.09052712483995,96.0930693768969,96.09306937699691,96.09433936064684,96.09433936074684,96.09560827346378,96.09560827356378,96.09814517095572,96.09814517105572,96.0994124416754,96.0994124417754,96.10067914123078,96.10067914133079,96.10447641977342,96.10447641987342,96.1057401207164,96.1057401208164,96.10953054528783,96.10953054538783,96.11205512705023,96.11205512715023,96.11583809078839,96.11583809088839,96.1183571036992,96.1183571037992,96.11961559276824,96.11961559286824,96.12087326078866,96.12087326088866,96.12338702612773,96.12338702622773,96.127155282386,96.127155282486,96.13092207503581,96.13092207513581,96.13217678014159,96.1321767802416,96.13719424404954,96.13719424414954,96.14095521796338,96.14095521806338,96.14220785259867,96.14220785269868,96.148467991465,96.148467991565,96.14971802016333,96.14971802026334,96.15221718511582,96.15221718521582,96.15596352294529,96.15596352304529,96.15721115989325,96.15721115999325,96.16095228584877,96.16095228594877,96.16344413275907,96.16344413285907,96.16718018963172,96.16718018973172,96.17340323851839,96.17340323861839,96.17589073030115,96.17589073040115,96.17962011169142,96.17962011179142,96.18334738691344,96.18334738701344,96.18583077345302,96.18583077355302,96.19079508338628,96.19079508348628,96.19327586398889,96.19327586408889,96.1969951072134,96.1969951073134,96.19947338897234,96.19947338907234,96.20071174450094,96.20071174460094,96.20813998569089,96.2081399857909,96.21061459057985,96.21061459067985,96.21432490936267,96.21432490946268,96.21926937133762,96.21926937143762,96.22421054911803,96.22421054921803,96.22544447812355,96.22544447822355,96.23161230256505,96.23161230266506,96.23777687850978,96.23777687860978,96.24147138318193,96.24147138328193,96.24639128509335,96.24639128519335,96.24884725574806,96.24884725584806,96.25375709088927,96.25375709098927,96.25621038421147,96.25621038431147,96.25866135717884,96.25866135727884,96.26111090223554,96.26111090233555,96.26723167702048,96.26723167712048,96.26967836625583,96.26967836635583,96.27457010262928,96.27457010272929,96.278236513159,96.278236513259,96.2806790971512,96.2806790972512,96.28434158092657,96.28434158102657,96.2880017443471,96.2880017444471,96.29043929495417,96.29043929505417,96.29531057650728,96.29531057660728,96.2977444145466,96.2977444146466,96.30139333317061,96.30139333327061,96.30260877073296,96.30260877083296,96.30625394109146,96.30625394119146,96.307467665161,96.307467665261,96.30989389957601,96.30989389967601,96.31110623143265,96.31110623153265,96.31352993130622,96.31352993140622,96.32079706847483,96.32079706857483,96.32442624623383,96.32442624633383,96.32805185421616,96.32805185431616,96.3316731427687,96.3316731428687,96.33649769594511,96.33649769604511,96.34011334425051,96.34011334435051,96.34252169408438,96.34252169418438,96.3461314165605,96.3461314166605,96.35094204760789,96.35094204770789,96.35334507847452,96.35334507857452,96.3557457889863,96.3557457890863,96.36174462020001,96.36174462030002,96.365340741827,96.365340741927,96.36893161588226,96.36893161598226,96.37132254520596,96.37132254530596,96.37490545865924,96.37490545875924,96.37609794255756,96.37609794265757,96.37848201791004,96.37848201801005,96.38086477244516,96.38086477254517,96.38562846092928,96.38562846102928,96.39039000754738,96.39039000764738,96.39276926370135,96.39276926380136,96.39514698485135,96.39514698495135,96.39871176383913,96.39871176393913,96.40108677195886,96.40108677205886,96.40346049495898,96.40346049505898,96.40583282574619,96.40583282584619,96.41057384614842,96.41057384624843,96.41294292843887,96.41294292853887,96.41886322456573,96.41886322466573,96.42359614157489,96.42359614167489,96.42596036896906,96.42596036906906,96.42832238310169,96.42832238320169,96.42950262266602,96.42950262276602,96.43186149539517,96.43186149549517,96.43539768047178,96.43539768057178,96.44010974998517,96.44010975008517,96.44246458886668,96.44246458896669,96.4471714822039,96.4471714823039,96.4518740918091,96.4518740919091,96.45304722751784,96.45304722761784,96.45656502824458,96.45656502834458,96.45773630766945,96.45773630776945,96.46124821826467,96.46124821836467,96.46241735582353,96.46241735592353,96.46475391744848,96.46475391754848,96.47175942568457,96.47175942578457,96.47525966311005,96.47525966321005,96.47875733029633,96.47875733039633,96.47992214842543,96.47992214852543,96.48341513920427,96.48341513930427,96.4869059881171,96.4869059882171,96.48806791472708,96.48806791482708,96.49039041143192,96.49039041153192,96.49155058885134,96.49155058895134,96.49387026543259,96.49387026553259,96.50082554691086,96.50082554701086,96.5054595097106,96.5054595098106,96.51009050959571,96.51009050969571,96.5124042603477,96.5124042604477,96.51702883463477,96.51702883473477,96.52164937507422,96.52164937517422,96.52511159437812,96.52511159447812,96.5308772477794,96.5308772478794,96.5320295002946,96.5320295003946,96.5377883711202,96.5377883712202,96.54239141965394,96.54239141975394,96.54469110548582,96.54469110558583,96.55043687523103,96.55043687533103,96.55617782577772,96.55617782587773,96.55847265671332,96.55847265681332,96.56076655950699,96.56076655960699,96.56305889159893,96.56305889169893,96.57337158373791,96.57337158383791,96.57680688692568,96.57680688702568,96.58138394748522,96.58138394758522,96.58367081781884,96.58367081791884,96.58709923133762,96.58709923143762,96.58938292457,96.58938292467,96.5928064831925,96.5928064832925,96.59622757866912,96.59622757876912,96.5996467464663,96.59964674656631,96.60534289631573,96.60534289641573,96.60875838724296,96.60875838734296,96.61330953122383,96.61330953132384,96.61558322908152,96.61558322918152,96.61899150905985,96.61899150915986,96.62239746868336,96.62239746878336,96.62466559768943,96.62466559778943,96.63033154722818,96.63033154732818,96.6348606299892,96.63486063008921,96.64165073790072,96.64165073800072,96.6439121914244,96.64391219152441,96.64956221975919,96.64956221985919,96.65069109023716,96.65069109033716,96.6574620641457,96.6574620642457,96.66197397628092,96.66197397638092,96.66535587560962,96.66535587570962,96.66760879736707,96.66760879746707,96.67323826378819,96.6732382638882,96.67998756915232,96.67998756925232,96.68673294776208,96.68673294786208,96.68785599950408,96.68785599960408,96.69346908065027,96.69346908075028,96.697957039584,96.697957039684,96.70019996596675,96.70019996606675,96.70580394418242,96.70580394428242,96.70916446054885,96.70916446064885,96.71252212109393,96.71252212119393,96.7169965148763,96.7169965149763,96.71923173054142,96.71923173064143,96.72593355787576,96.72593355797576,96.72928257956129,96.72928257966129,96.73486014142956,96.73486014152957,96.73708935986988,96.73708935996989,96.74265828287861,96.74265828297861,96.74710908043714,96.74710908053714,96.75155648670784,96.75155648680784,96.75488990846931,96.75488990856931,96.75822075999156,96.75822076009156,96.75932988960452,96.75932988970452,96.76265574343942,96.76265574353943,96.76487175370605,96.76487175380605,96.77040926268003,96.77040926278003,96.77151525088951,96.77151525098951,96.77925434823234,96.77925434833234,96.78588492282425,96.78588492292425,96.78919635476139,96.78919635486139,96.79581329280639,96.79581329290639,96.79801734432114,96.79801734442114,96.80132088705183,96.80132088715183,96.80682202000149,96.80682202010149,96.81011974399621,96.81011974409621,96.81231708433081,96.81231708443082,96.81780572306211,96.81780572316211,96.82438535694087,96.82438535704087,96.82767237160554,96.82767237170555,96.82986239389797,96.82986239399797,96.83314587448375,96.83314587458375,96.83533296955929,96.83533296965929,96.83642589238615,96.83642589248615,96.83861084559568,96.83861084569568,96.84407074762474,96.84407074772474,96.84625366606159,96.84625366616159,96.84734409004477,96.84734409014477,96.84952393847367,96.84952393857367,96.85497006116476,96.85497006126477,96.85714701807456,96.85714701817456,96.86149825456162,96.86149825466163,96.86475903136774,96.86475903146774,96.86584502882785,96.86584502892785,96.86910177178633,96.86910177188633,96.87235537334134,96.87235537344134,96.87560604767947,96.87560604777947,96.87668808268747,96.87668808278747,96.88101418794675,96.88101418804675,96.88641922250831,96.88641922260831,96.88965897332982,96.88965897342982,96.89289590402775,96.89289590412776,96.89936444645637,96.89936444655638,96.90259562981386,96.90259562991386,96.9058234218835,96.9058234219835,96.91119757787682,96.91119757797682,96.9154926617768,96.9154926618768,96.91978371182913,96.91978371192913,96.9229999735201,96.92299997362011,96.92514173243188,96.92514173253188,96.92835042619632,96.92835042629632,96.93048779428278,96.93048779438278,96.93262409143624,96.93262409153624,96.93796215698738,96.93796215708738,96.94223014628238,96.94223014638239,96.94649342347219,96.94649342357219,96.94968858778302,96.94968858788302,96.95288111045912,96.95288111055912,96.95819922095865,96.95819922105865,96.96138696013399,96.961386960234,96.96457220046567,96.96457220056567,96.9677547991626,96.96775479926261,96.97093482762034,96.97093482772034,96.97411282130537,96.97411282140537,96.97728853033331,96.97728853043331,96.9804607766779,96.9804607767779,96.98574600953425,96.98574600963425,96.98785831778845,96.98785831788845,96.98996959080742,96.98996959090742,96.99418995928158,96.99418995938159,96.9962981622926,96.9962981623926,96.99840400925102,96.99840400935102,97.00050703608588,97.00050703618588,97.00155769275689,97.00155769285689,97.00995973332606,97.00995973342606,97.01415761220717,97.01415761230717,97.01730412046192,97.01730412056192,97.02044830836184,97.02044830846184,97.02358931916051,97.02358931926051,97.02882082687789,97.02882082697789,97.03195705417582,97.03195705427582,97.03509210344745,97.03509210354746,97.03613626312492,97.03613626322492,97.0382236900357,97.0382236901357,97.03926624331366,97.03926624341366,97.04343452874613,97.04343452884613,97.04551760052937,97.04551760062937,97.0517632461024,97.0517632462024,97.05696366106494,97.05696366116494,97.06111816715945,97.06111816725945,97.06734578536027,97.06734578546028,97.07253306354465,97.07253306364466,97.0766792520595,97.07667925215951,97.08082190649544,97.08082190659545,97.08288998521665,97.08288998531665,97.08702407218857,97.08702407228857,97.09012303101801,97.09012303111801,97.09321981228368,97.09321981238368,97.09528328599296,97.09528328609296,97.10250162431443,97.10250162441443,97.10868294251176,97.10868294261176,97.11074159702254,97.11074159712254,97.11279875222712,97.11279875232712,97.11896632678426,97.11896632688426,97.12204754382361,97.12204754392361,97.12410041529618,97.12410041539619,97.1292291491431,97.1292291492431,97.13640365060698,97.13640365070698,97.13947419401408,97.13947419411409,97.14254245276412,97.14254245286412,97.14356415854631,97.14356415864631,97.14867118815104,97.14867118825104,97.15071252924494,97.15071252934494,97.15479335514887,97.15479335524887,97.15887143232476,97.15887143242476,97.16294618960835,97.16294618970835,97.16701830525719,97.16701830535719,97.17007028582559,97.17007028592559,97.17312055290118,97.17312055300118,97.17718460085473,97.17718460095473,97.18124543600929,97.1812454361093,97.18733308326667,97.18733308336667,97.1893609663025,97.1893609664025,97.19341416213497,97.19341416223497,97.19645114964133,97.19645114974134,97.19948688772585,97.19948688782586,97.20252041254884,97.20252041264884,97.20656279195798,97.20656279205798,97.20959260421321,97.20959260431322,97.21262009611362,97.21262009621363,97.2206900403587,97.2206900404587,97.22472231729985,97.22472231739985,97.23177248344214,97.23177248354214,97.23680408366573,97.23680408376573,97.23881522444896,97.23881522454896,97.24484589807061,97.24484589817061,97.25187625195237,97.25187625205237,97.25689582202855,97.25689582212856,97.26090889357118,97.26090889367119,97.26191128686156,97.26191128696156,97.26391514530037,97.26391514540038,97.26591757582851,97.26591757592851,97.26791897112142,97.26791897122142,97.2729197641723,97.2729197642723,97.27591780541513,97.27591780551514,97.27991284958556,97.27991284968556,97.28290749954056,97.28290749964056,97.2868971533482,97.28689715344821,97.28988716259352,97.28988716269352,97.29586382549407,97.29586382559407,97.30084091522782,97.30084091532783,97.305814113905,97.305814114005,97.3087953414997,97.3087953415997,97.31276753909941,97.31276753919941,97.31475226353524,97.31475226363524,97.32070418788344,97.32070418798344,97.32169512219068,97.32169512229068,97.325657431509,97.325657431609,97.32664686651005,97.32664686661005,97.33060296441694,97.33060296451694,97.33653425545596,97.33653425555596,97.34147382683686,97.34147382693686,97.34443431402907,97.34443431412907,97.34936463968837,97.34936463978838,97.35034961246858,97.35034961256858,97.3523186655848,97.3523186656848,97.35527008554426,97.35527008564426,97.36411831250005,97.36411831260006,97.36902907578319,97.36902907588319,97.37001015750681,97.37001015760681,97.37197149990541,97.37197150000542,97.37393130730005,97.37393130740006,97.37588957969074,97.37588957979074,97.37980366132616,97.37980366142617,97.38273673263352,97.38273673273352,97.38664610216374,97.38664610226374,97.38957613916091,97.38957613926091,97.39152841432677,97.39152841442677,97.39347936867526,97.39347936877526,97.3973790641107,97.3973790642107,97.39835304197874,97.39835304207874,97.4022480967045,97.4022480968045,97.40614065258657,97.40614065268657,97.40905751710781,97.40905751720781,97.41391544769616,97.41391544779616,97.41876895176142,97.41876895186142,97.42265008435814,97.42265008445814,97.42361949291204,97.42361949301204,97.42749594910131,97.42749594920132,97.4313697279581,97.4313697280581,97.437176148209,97.437176148309,97.43910978914063,97.43910978924063,97.44200887617403,97.44200887627403,97.44393980407537,97.44393980417537,97.44973051730891,97.44973051740891,97.45358883903268,97.45358883913268,97.45551662553055,97.45551662563055,97.46129627245641,97.46129627255641,97.46322045347982,97.46322045357982,97.467066316683,97.467066316783,97.4699494111169,97.4699494112169,97.47283032798701,97.47283032808701,97.4805072041224,97.4805072042224,97.48242481675675,97.48242481685675,97.48529880826011,97.48529880836011,97.48912760793078,97.48912760803078,97.4948669163802,97.4948669164802,97.49869025429257,97.49869025439257,97.50155514286541,97.50155514296542,97.50537230506413,97.50537230516413,97.50727877999523,97.50727878009523,97.5139476225931,97.5139476226931,97.52061221715671,97.52061221725671,97.52346657488837,97.52346657498838,97.53107330491208,97.53107330501209,97.53392419996038,97.53392420006038,97.5377224423427,97.5377224424427,97.54151818588136,97.54151818598136,97.54720805292384,97.54720805302384,97.5500490953885,97.5500490954885,97.55288763900948,97.55288763910949,97.55950661182719,97.55950661192719,97.56517673800465,97.56517673810465,97.56706500707473,97.56706500717473,97.57367141427854,97.57367141437854,97.5764999982226,97.5764999983226,97.57932697576717,97.57932697586718,97.58121017575436,97.58121017585437,97.58215095469933,97.58215095479933,97.58873372998154,97.58873373008154,97.59343266271678,97.59343266281678,97.60000547832207,97.60000547842208,97.60282006874158,97.60282006884158,97.6056316248509,97.6056316249509,97.609377712796,97.609377712896,97.61405711885283,97.61405711895283,97.61779753085301,97.61779753095301,97.62340254430393,97.62340254440393,97.62806809962724,97.62806809972724,97.63179730252867,97.63179730262867,97.63738753710415,97.63738753720415,97.64017978072165,97.64017978082165,97.64576009131797,97.64576009141797,97.6494771569787,97.6494771570787,97.6504054417053,97.6504054418053,97.6522610473188,97.6522610474188,97.65689718768232,97.65689718778232,97.659677008477,97.659677008577,97.66152858024284,97.66152858034285,97.66893122613405,97.66893122623405,97.67447905176262,97.67447905186262,97.67632633979646,97.67632633989646,97.6818633490017,97.6818633491017,97.68647400115977,97.68647400125977,97.68831664848392,97.68831664858392,97.69199926579975,97.69199926589975,97.69567984834286,97.69567984844286,97.69843864315293,97.69843864325293,97.70487188049023,97.70487188059023,97.70762617738168,97.70762617748169,97.71129658606128,97.71129658616128,97.7149636391508,97.7149636392508,97.71771201021298,97.71771201031298,97.72137406561515,97.72137406571515,97.72228867809709,97.72228867819709,97.72320254092591,97.72320254102591,97.73051190855259,97.73051190865259,97.73416421846444,97.73416421856444,97.74237466911946,97.74237466921946,97.74693302404921,97.74693302414921,97.75239792376563,97.75239792386563,97.75512866013102,97.75512866023102,97.75785725463042,97.75785725473042,97.76422013166946,97.76422013176946,97.76876181574215,97.76876181584215,97.77329971585156,97.77329971595157,97.77420651052257,97.77420651062258,97.77783251118032,97.77783251128032,97.78145594159888,97.78145594169888,97.78507669468492,97.78507669478492,97.78959949463899,97.789599494739,97.79231098990812,97.79231099000812,97.79502105726657,97.79502105736657,97.79863124381366,97.79863124391366,97.80494475084127,97.80494475094127,97.80764864248607,97.80764864258607,97.81485384402937,97.81485384412937,97.8220549403295,97.8220549404295,97.82475254916734,97.82475254926734,97.83014423276416,97.83014423286416,97.83463358391079,97.83463358401079,97.83732426738189,97.83732426748189,97.84090793048827,97.84090793058827,97.8444892018443,97.8444892019443,97.84717313843748,97.84717313853749,97.85164289151017,97.85164289161017,97.85521577389099,97.85521577399099,97.86235654096531,97.86235654106531,97.86592428286772,97.86592428296773,97.86948966871753,97.86948966881754,97.87305252002591,97.87305252012591,97.87750238944251,97.87750238954251,97.8801697979697,97.8801697980697,97.8828349218398,97.8828349219398,97.88638581439632,97.88638581449632,97.88904715430316,97.88904715440316,97.89348006728055,97.89348006738055,97.89525196163102,97.89525196173102,97.8996787702903,97.8996787703903,97.90675724476164,97.90675724486164,97.91294577390785,97.91294577400785,97.91647921025643,97.91647921035643,97.91912709210519,97.91912709220519,97.92353665874313,97.92353665884313,97.92970509004668,97.92970509014668,97.9341061963139,97.93410619641391,97.9376245682049,97.9376245683049,97.94289873475356,97.94289873485356,97.9481691530367,97.9481691531367,97.95255747950343,97.95255747960343,97.95781815229626,97.95781815239626,97.96482822984649,97.96482822994649,97.96745480012851,97.96745480022851,97.9709533240612,97.9709533241612,97.97532305199145,97.97532305209145,97.98143693710739,97.98143693720739,97.98667258576573,97.98667258586573,97.98928887509092,97.98928887519092,97.99364571612738,97.99364571622738,97.99887026278024,97.99887026288025,98.0014801622052,98.0014801623052,98.00321871484151,98.00321871494151,98.00669367824814,98.00669367834814,98.01190016183104,98.01190016193104,98.01536869963965,98.01536869973965,98.02403347577211,98.02403347587212,98.02749690880007,98.02749690890008,98.03268854211201,98.03268854221201,98.03701107759461,98.03701107769461,98.0430570655582,98.0430570656582,98.0482357405808,98.0482357406808,98.05427305398706,98.05427305408706,98.05944369701214,98.05944369711214,98.06202682311202,98.06202682321202,98.06632733307254,98.06632733317254,98.0714856603681,98.0714856604681,98.07406253935879,98.0740625394588,98.08178924957656,98.08178924967656,98.08607865753162,98.08607865763162,98.08779300708208,98.08779300718209,98.09550315353607,98.09550315363607,98.09892696204295,98.09892696214295,98.10234652251557,98.10234652261558,98.10490922949289,98.10490922959289,98.1083236494871,98.1083236495871,98.11002953866685,98.11002953876685,98.11343910376479,98.11343910386479,98.11684613432129,98.1168461344213,98.12280409159206,98.12280409169206,98.12365362704521,98.12365362714522,98.12874905487574,98.12874905497574,98.13299102049879,98.1329910205988,98.13807752388763,98.13807752398763,98.142312706935,98.142312707035,98.14569753348059,98.14569753358059,98.14907961129812,98.14907961139812,98.15330304978023,98.15330304988024,98.15752331116109,98.15752331126109,98.16173939590324,98.16173939600324,98.16342483026263,98.16342483036263,98.16595098272678,98.16595098282679,98.17099989636723,98.17099989646724,98.17604356243599,98.176043562536,98.18108215942185,98.18108215952185,98.1844387847317,98.1844387848317,98.18946892134684,98.18946892144685,98.19281926384976,98.19281926394976,98.19784136846738,98.19784136856738,98.20034963635038,98.20034963645038,98.20285640492719,98.20285640502719,98.20619632368219,98.2061963237822,98.21036864296231,98.21036864306231,98.21870756987987,98.21870756997987,98.22287321367759,98.22287321377759,98.22953413137124,98.22953413147124,98.23202940526714,98.23202940536714,98.23701756130859,98.23701756140859,98.24366341454467,98.24366341464467,98.24615326238003,98.24615326248004,98.24864103974492,98.24864103984493,98.25443971358045,98.25443971368045,98.25775089563322,98.25775089573322,98.2651945939562,98.2651945940562,98.27097970264036,98.27097970274036,98.27593487394533,98.27593487404533,98.27923488259711,98.27923488269711,98.28253332054716,98.28253332064716,98.28582818872053,98.28582818882053,98.29076679626172,98.29076679636172,98.29487885771157,98.29487885781157,98.29816583667848,98.29816583677848,98.30062923246868,98.30062923256868,98.30391085677059,98.30391085687059,98.30636996882878,98.30636996892878,98.31210381551989,98.31210381561989,98.31537690805554,98.31537690815554,98.32191770276407,98.32191770286407,98.32926937224434,98.32926937234434,98.331717239506,98.33171723960601,98.33905659325676,98.33905659335676,98.3439443671781,98.3439443672781,98.34882728620316,98.34882728630316,98.35532895615933,98.35532895625933,98.36020152283203,98.36020152293203,98.36507001995933,98.36507002005933,98.36993394777248,98.36993394787248,98.3747944842978,98.3747944843978,98.3780326287198,98.3780326288198,98.38126816720482,98.38126816730482,98.38692665591033,98.38692665601033,98.39015705391694,98.39015705401694,98.39419282923923,98.39419282933923,98.39822485629601,98.39822485639601,98.4030598689181,98.40305986901811,98.40708539898134,98.40708539908134,98.41512935525222,98.41512935535222,98.41914781715764,98.41914781725764,98.42155730932004,98.42155730942004,98.42797908767422,98.42797908777422,98.43118771004312,98.43118771014312,98.4343936550795,98.4343936551795,98.43839651706085,98.43839651716085,98.43919614703609,98.43919614713609,98.4447919147654,98.4447919148654,98.45357649260308,98.45357649270308,98.45916369286837,98.45916369296837,98.46474660940166,98.46474660950166,98.47271420814947,98.47271420824947,98.47988035633594,98.47988035643594,98.48544881527368,98.48544881537369,98.48862727302968,98.48862727312968,98.49418505833518,98.49418505843518,98.50053297800994,98.50053297810994,98.50449678663446,98.50449678673446,98.50766489203811,98.50766489213811,98.50924762392255,98.50924762402255,98.51162006180307,98.51162006190307,98.51794227908577,98.51794227918577,98.52031082590972,98.52031082600972,98.52504509943404,98.52504509953404,98.53056411696484,98.53056411706484,98.53765504995677,98.53765505005677,98.54159162118525,98.54159162128525,98.54552455124154,98.54552455134154,98.54945487536084,98.54945487546084,98.55338259354318,98.55338259364318,98.55730717032205,98.55730717042205,98.56044418297084,98.56044418307084,98.56671488837613,98.56671488847613,98.57141371401805,98.57141371411805,98.57532579657855,98.57532579667856,98.58158054508209,98.58158054518209,98.58626755476323,98.58626755486323,98.5886083644224,98.5886083645224,98.59172752818783,98.59172752828783,98.59640357865457,98.59640357875458,98.60107488131833,98.60107488141833,98.60574268560096,98.60574268570096,98.61273655636508,98.61273655646508,98.61506515738803,98.61506515748803,98.61816822146065,98.61816822156065,98.62204378520576,98.62204378530576,98.62901427393267,98.62901427403267,98.6313352713313,98.6313352714313,98.63442609106991,98.63442609116991,98.64214534045443,98.64214534055444,98.64754412790681,98.64754412800681,98.65448034677765,98.65448034687765,98.66141181784549,98.66141181794549,98.66525885907497,98.66525885917497,98.66833482854263,98.66833482864263,98.66987115333029,98.6698711534303,98.67140644288273,98.67140644298273,98.6775424249163,98.6775424250163,98.68290765646793,98.68290765656793,98.68443869798612,98.68443869808613,98.69208812253888,98.69208812263888,98.69743800405081,98.69743800415081,98.70431082368789,98.70431082378789,98.71041485622021,98.71041485632021,98.71498863258522,98.71498863268522,98.72184206833495,98.72184206843495,98.72412422656335,98.72412422666335,98.72944572835073,98.72944572845073,98.73400147734351,98.73400147744351,98.73931294805843,98.73931294815843,98.74386241424426,98.74386241434426,98.74689297615261,98.74689297625261,98.74916338981575,98.74916338991575,98.75370182539167,98.75370182549167,98.75748079097995,98.75748079107996,98.76276759453803,98.76276759463804,98.77106571890819,98.77106571900819,98.77709500031693,98.77709500041694,98.78086004377619,98.78086004387619,98.78687822317947,98.78687822327947,98.79063626987644,98.79063626997645,98.79514168501811,98.79514168511811,98.80038925673054,98.80038925683054,98.80638180513729,98.8063818052373,98.8116211306656,98.8116211307656,98.8198479308978,98.8198479309978,98.82358395207267,98.82358395217267,98.82955700949879,98.82955700959879,98.83477994975215,98.83477994985215,98.83999824929585,98.83999824939585,98.84670293245152,98.84670293255152,98.85414605961022,98.85414605971022,98.85860785208092,98.85860785218092,98.8675264036372,98.8675264037372,98.87198123504339,98.8719812351434,98.8764324966729,98.8764324967729,98.87939776736586,98.87939776746586,98.88384088990455,98.88384089000455,98.88532052663861,98.88532052673861,98.88901674480357,98.88901674490357,98.89344830126583,98.89344830136584,98.90304229026682,98.90304229036683,98.90672890573252,98.90672890583252,98.91114921739825,98.91114921749825,98.91630265409975,98.91630265419975,98.92365882149865,98.92365882159865,98.9280687094165,98.9280687095165,98.93320854526887,98.93320854536887,98.94274591761179,98.94274591771179,98.94787722169791,98.94787722179791,98.95446877863074,98.95446877873074,98.96178542860184,98.96178542870184,98.96617029238519,98.96617029248519,98.97274178717312,98.97274178727312,98.97857833633918,98.97857833643918,98.98222204308924,98.98222204318924,98.98658966485125,98.98658966495125,98.99022601786136,98.99022601796136,98.99313302979898,98.99313302989898,98.99894194889357,98.99894194899358,99.00184328488628,99.00184328498628,99.00546725077133,99.00546725087133,99.0141561585941,99.0141561586941,99.02066668159642,99.02066668169643,99.02644843469052,99.02644843479052,99.03078089415227,99.03078089425227,99.03510999802397,99.03510999812397,99.04015612723862,99.04015612733862,99.04447802016142,99.04447802026142,99.04735647388563,99.04735647398563,99.05095113190418,99.05095113200419,99.05669897211986,99.05669897221986,99.06531220067713,99.06531220077713,99.07033241331311,99.07033241341311,99.07678100069012,99.07678100079012,99.0803605942511,99.0803605943511,99.08608305335461,99.08608305345462,99.09108548850277,99.09108548860277,99.09536929190844,99.09536929200844,99.09822307847584,99.09822307857584,99.1039254040389,99.1039254041389,99.10891059716572,99.10891059726572,99.11531459803204,99.11531459813204,99.1174468256401,99.1174468257401,99.12241866780214,99.12241866790214,99.1288058907181,99.1288058908181,99.13377002216252,99.13377002226252,99.14014610737526,99.14014610747526,99.14510060042265,99.14510060052265,99.15075733993758,99.15075734003759,99.1564076538545,99.1564076539545,99.15852449572509,99.15852449582509,99.1641655639204,99.16416556402041,99.16768832663672,99.16768832673672,99.17402118185392,99.17402118195392,99.18034864670834,99.18034864680834,99.18667172073745,99.18667172083745,99.19088166546372,99.19088166556372,99.19508804041331,99.19508804051331,99.20279326057549,99.20279326067549,99.20629192729925,99.20629192739925,99.21258508659982,99.21258508669982,99.22306534405594,99.22306534415594,99.22934736565328,99.22934736575328,99.23213625368072,99.23213625378072,99.23561907059603,99.23561907069603,99.2418827078435,99.2418827079435,99.24605381339954,99.24605381349954,99.24952513563397,99.24952513573398,99.25507496033748,99.25507496043748,99.26131282379735,99.26131282389736,99.26754654631628,99.26754654641628,99.27100637386981,99.27100637396981,99.27446281013549,99.27446281023549,99.28067982609959,99.28067982619959,99.28550873440355,99.28550873450355,99.28964424928616,99.28964424938616,99.29239843908431,99.29239843918431,99.30202791166545,99.30202791176545,99.3089007670003,99.3089007671003,99.31370750699111,99.31370750709111,99.31919639560677,99.31919639570677,99.32399403266707,99.32399403276708,99.3274184123382,99.3274184124382,99.33357613431173,99.33357613441173,99.34041043605848,99.34041043615848,99.34450643351327,99.34450643361328,99.34928100981625,99.34928100991625,99.35609785535506,99.35609785545506,99.36154422793054,99.36154422803054,99.3676665734172,99.3676665735172,99.37038438719104,99.37038438729104,99.37785296685742,99.37785296695742,99.38395785613615,99.38395785623615,99.39141290634893,99.39141290644893,99.39818473700387,99.39818473710388,99.40495085601613,99.40495085611613,99.41441522642604,99.41441522652605,99.41846635178804,99.41846635188804,99.42453615016206,99.42453615026206,99.43396903514169,99.43396903524169,99.44002705325268,99.44002705335268,99.44406272148166,99.44406272158166,99.44943862666555,99.44943862676556,99.45548161601674,99.45548161611674,99.46084856106118,99.46084856116119,99.46621015144062,99.46621015154062,99.47290573166578,99.47290573176578,99.47691944576822,99.47691944586822,99.48092976858281,99.48092976868281,99.48493680720287,99.48493680730287,99.48693820249578,99.48693820259578,99.49160440037889,99.49160440047889,99.49559980152698,99.49559980162698,99.4989260837351,99.4989260838351,99.50357953751548,99.50357953761548,99.51154702916999,99.51154702926999,99.51751583856186,99.51751583866186,99.52546815866549,99.52546815876549,99.52944028486966,99.52944028496967,99.53340876990163,99.53340877000163,99.53803434372618,99.53803434382618,99.5433163994813,99.5433163995813,99.55123259344498,99.55123259354498,99.55518704925461,99.55518704935461,99.5591376854032,99.5591376855032,99.56505723187695,99.56505723197695,99.5722852442932,99.5722852443932,99.57556693999064,99.57556694009064,99.58146970851402,99.58146970861402,99.5867112116061,99.5867112117061,99.58998458972388,99.58998458982389,99.5952179894229,99.59521798952291,99.59783242246424,99.59783242256424,99.60370752521838,99.60370752531838,99.61153515302276,99.61153515312276,99.6187036929596,99.6187036930596,99.62195900800741,99.62195900810741,99.62586213042846,99.62586213052846,99.63236123014542,99.63236123024542,99.63820513305141,99.63820513315142,99.64144902481387,99.64144902491387,99.64598631806125,99.64598631816125,99.64922417690111,99.64922417700112,99.654399746218,99.654399746318,99.65763096527326,99.65763096537326,99.6608589715295,99.66085897162951,99.66924330600636,99.66924330610637,99.6750431578682,99.6750431579682,99.68019359595729,99.6801935960573,99.68598181034717,99.68598181044717,99.69304850455548,99.69304850465548,99.69882311809621,99.69882311819622,99.70587471214918,99.70587471224918,99.71035810176876,99.71035810186876,99.71739698741678,99.71739698751678,99.7276249330575,99.7276249331575,99.73145633866514,99.73145633876514,99.73783660051657,99.73783660061657,99.74357048290545,99.74357048300546,99.751845439425,99.751845439525,99.75884113077522,99.75884113087523,99.7651937625552,99.7651937626552,99.77281095202457,99.77281095212457,99.77598180615625,99.77598180625625,99.77978272587109,99.77978272597109,99.78547859013838,99.78547859023838,99.79180180695856,99.79180180705856,99.79622401060593,99.79622401070593,99.79937926481354,99.79937926491354,99.80442418030411,99.80442418040411,99.81009394950391,99.81009394960391,99.81449973217858,99.81449973227858,99.81952979739819,99.81952979749819,99.8245552576059,99.8245552577059,99.8289471538493,99.8289471539493,99.83270723531896,99.83270723541897,99.83834162803191,99.83834162813191,99.84647315092464,99.84647315102464,99.85272172371454,99.85272172381454,99.8589661555635,99.8589661556635,99.86395609649327,99.86395609659327,99.87018867668367,99.87018867678367,99.87516965747399,99.87516965757399,99.88014553348368,99.88014553358369,99.88325188175084,99.88325188185084,99.88511394866012,99.88511394876012,99.89069607984253,99.89069607994253,99.89503300152514,99.89503300162514,99.90060470895966,99.90060470905966,99.90431524192908,99.90431524202909,99.90863981218439,99.90863981228439,99.9166631351461,99.9166631352461,99.92467846180804,99.92467846190804,99.93206922034292,99.93206922044293,99.93699069295607,99.93699069305607,99.9394488054768,99.9394488055768,99.94743032635363,99.94743032645363,99.95663017631192,99.95663017641192,99.96275673413506,99.96275673423506,99.96826482814924,99.96826482824925,99.97193277368292,99.97193277378292,99.9768193338802,99.9768193339802,99.97925920484211,99.97925920494211,99.98352487378229,99.98352487388229,99.98778668736364,99.98778668746364,99.99204303918668,99.99204303928668,99.99325569232323,99.99325569242323,99.99385994842102,99.99385994852102,99.99446070613769,99.99446070623769,99.99503522599578,99.99503522609578,99.99556001886478,99.99556001896478,99.99602109122014,99.99602109132015,99.99643468554572,99.99643468564572,99.99682550469612,99.99682550479612,99.99713889539042,99.99713889549042,99.9974369717428,99.9974369718428,99.99771730630509,99.9977173064051,99.99826805145054,99.99826805155054,99.9985344638838,99.9985344639838,99.99878970291607,99.99878970301607,99.99903558913346,99.99903558923346,99.99926998066994,99.99926998076994,99.99948006202727,99.99948006212728,99.9996750075315,99.9996750076315,99.99985199705905,99.99985199715906,100.0,100.0000000001],[0.0,1e-10,0.18038246341647346,0.18038246351647347,0.3345680051019236,0.3345680052019236,0.48334293944270146,0.48334293954270147,0.6180663175345529,0.6180663176345529,0.7491892838300882,0.7491892839300882,0.8770070748550888,0.8770070749550888,1.0036088870391884,1.0036088871391884,1.1268086617823887,1.1268086618823887,1.2466002516076446,1.2466002517076447,1.3659683354341594,1.3659683355341594,1.4800591648795953,1.4800591649795953,1.5924228291236882,1.5924228292236882,1.702298065592353,1.702298065692353,1.8112726178601029,1.8112726179601029,1.9199488598635488,1.9199488599635488,2.0280278508279213,2.0280278509279213,2.1324623336702504,2.1324623337702504,2.2331305804636625,2.2331305805636625,2.332291316139381,2.332291316239381,2.4314159156840076,2.4314159157840076,2.529201626253735,2.529201626353735,2.6267151139296967,2.6267151140296967,2.723886825526093,2.723886825626093,2.820581674175549,2.820581674275549,2.9170045363726635,2.9170045364726636,3.0126986860985094,3.0126986861985094,3.108109894252664,3.108109894352664,3.2029110441239017,3.2029110442239017,3.297581717735547,3.297581717835547,3.3901311571044968,3.3901311572044968,3.482531086679157,3.482531086779157,3.573859069647531,3.573859069747531,3.6645181204182093,3.6645181205182094,3.755116129894382,3.755116129994382,3.8455352320257807,3.8455352321257807,3.9351200111828284,3.9351200112828284,4.024685441549945,4.024685441649945,4.114048793312657,4.114048793412657,4.202520563869308,4.202520563969308,4.29092514565441,4.29092514575441,4.377966484997857,4.377966485097857,4.464457034042206,4.464457034142206,4.5508576171244135,4.5508576172244135,4.636140068708688,4.636140068808688,4.720424422225025,4.720424422325025,4.8044194108442975,4.8044194109442975,4.888277893947645,4.888277894047645,4.972058193753189,4.972058193853189,5.05493631189546,5.05493631199546,5.136400470910456,5.136400471010456,5.217846305715027,5.217846305815027,5.299236616191672,5.299236616291672,5.380420986187308,5.380420986287308,5.461497223637995,5.461497223737995,5.542404444876847,5.542404444976847,5.623012055423558,5.623012055523558,5.702657034116054,5.702657034216054,5.782261581324907,5.782261581424907,5.861830938168367,5.861830938268367,5.94019464077981,5.94019464087981,6.017649265519757,6.017649265619757,6.0947511196538855,6.0947511197538855,6.171682066044779,6.171682066144779,6.2483267606840345,6.2483267607840345,6.323727103511411,6.323727103611411,6.398917683388076,6.398917683488076,6.474041429155328,6.474041429255328,6.548929166973714,6.548929167073714,6.623651671643061,6.623651671743061,6.697447286520243,6.697447286620243,6.771193918615584,6.771193918715584,6.844732876326133,6.844732876426133,6.918152943406906,6.918152943506906,6.991494630155354,6.991494630255354,7.064824849494698,7.064824849594698,7.138110381404754,7.138110381504754,7.210731158248952,7.210731158348952,7.283306498932682,7.283306499032682,7.355774928685493,7.355774928785493,7.427676293087798,7.427676293187798,7.499210148702138,7.499210148802138,7.569149325126837,7.569149325226837,7.638629962815019,7.638629962915019,7.707239550294195,7.707239550394195,7.775540345272567,7.775540345372567,7.843838933464308,7.843838933564308,7.9119080158463655,7.9119080159463655,7.978838356919627,7.978838357019627,8.045684485438752,8.045684485538752,8.112245859668594,8.112245859768594,8.178688461489374,8.178688461589374,8.245118571321543,8.245118571421543,8.311401496366766,8.311401496466766,8.377454876195403,8.377454876295403,8.442917980006996,8.442917980106996,8.507901107726223,8.507901107826223,8.572611894330972,8.572611894430972,8.63729915501395,8.63729915511395,8.701813813456818,8.701813813556818,8.766037688354071,8.766037688454071,8.830068193572734,8.830068193672734,8.893749750655276,8.893749750755276,8.957187773070265,8.957187773170265,9.020424859681068,9.020424859781068,9.083518150498682,9.083518150598682,9.146474502324425,9.146474502424425,9.209385536210412,9.209385536310412,9.272093663947008,9.272093664047008,9.334147912923287,9.334147913023287,9.395842179830291,9.395842179930291,9.45750161103404,9.45750161113404,9.51894426485814,9.51894426495814,9.580154023878801,9.580154023978801,9.641349911669208,9.641349911769208,9.702426790609126,9.702426790709126,9.763402669653743,9.763402669753743,9.82435289480377,9.82435289490377,9.885086224353437,9.885086224453437,9.94550552028405,9.94550552038405,10.005901132665276,10.005901132765276,10.066246146581594,10.066246146681594,10.126554118008025,10.126554118108025,10.18665362691158,10.18665362701158,10.246731107355723,10.246731107455723,10.306720513369125,10.306720513469125,10.366486127573943,10.366486127673943,10.426190227601404,10.426190227701404,10.485783554821342,10.485783554921342,10.54514903132157,10.54514903142157,10.604390967177334,10.604390967277334,10.66346195588296,10.66346195598296,10.722266947985672,10.722266948085672,10.780608869557891,10.780608869657891,10.83855120512218,10.83855120522218,10.89620594910009,10.89620594920009,10.953740502020386,10.953740502120386,11.010711772652973,11.010711772752973,11.067544173355389,11.067544173455389,11.124111956696535,11.124111956796535,11.180645101368947,11.180645101468947,11.236647316821944,11.236647316921944,11.292549281110823,11.292549281210823,11.34834185183382,11.34834185193382,11.404104552123485,11.404104552223485,11.459791078867456,11.459791078967456,11.515401668507158,11.515401668607158,11.57085912291739,11.57085912301739,11.626103425383155,11.626103425483155,11.681096035952208,11.681096036052208,11.73592920618713,11.73592920628713,11.790701295720643,11.790701295820643,11.845304723704455,11.845304723804455,11.8997331062201,11.8997331063201,11.953983014866918,11.953983014966918,12.00813673126074,12.00813673136074,12.062116584393516,12.062116584493516,12.116061759450655,12.116061759550655,12.169912870227623,12.169912870327623,12.223540662078856,12.223540662178856,12.276949233322386,12.276949233422386,12.33022500329898,12.33022500339898,12.383484577037972,12.383484577137972,12.436708132866586,12.436708132966587,12.489815004852057,12.489815004952057,12.542875100842316,12.542875100942316,12.595683268494442,12.595683268594442,12.648446748717276,12.648446748817276,12.700965354437816,12.700965354537816,12.753470128335003,12.753470128435003,12.805947080957871,12.805947081057871,12.85830522176477,12.85830522186477,12.910507468858912,12.910507468958912,12.96267882094021,12.96267882104021,13.014765487584523,13.014765487684523,13.066746228470521,13.066746228570521,13.118614896121157,13.118614896221157,13.170418424159257,13.170418424259257,13.222121267557293,13.222121267657293,13.273692176640283,13.273692176740283,13.325205275794907,13.325205275894907,13.376630339925693,13.376630340025693,13.428028213292626,13.428028213392626,13.479396925550498,13.479396925650498,13.530705857534798,13.530705857634798,13.581801874016055,13.581801874116055,13.63286423700118,13.63286423710118,13.683766765583131,13.683766765683131,13.734454723572068,13.734454723672068,13.785083925866939,13.785083925966939,13.835658352565062,13.835658352665062,13.886098992823646,13.886098992923646,13.93650542788944,13.93650542798944,13.986902838774187,13.986902838874187,14.037295875492575,14.037295875592575,14.087649662934444,14.087649663034444,14.137929917093198,14.137929917193198,14.188110944667338,14.188110944767338,14.238225886863244,14.238225886963244,14.288279354288699,14.288279354388699,14.338161874564015,14.338161874664015,14.387915021973056,14.387915022073056,14.437647993047179,14.437647993147179,14.487322760123893,14.487322760223893,14.536967459732754,14.536967459832754,14.58648842166263,14.58648842176263,14.635963750397515,14.635963750497515,14.685303874044314,14.685303874144314,14.73458173478258,14.73458173488258,14.783783106719918,14.783783106819918,14.832896995330076,14.832896995430076,14.881848094019253,14.881848094119253,14.930758958259306,14.930758958359306,14.979427824701066,14.979427824801066,15.02789023837206,15.02789023847206,15.076346110496967,15.076346110596967,15.124744527355647,15.124744527455647,15.173088680907332,15.173088681007332,15.221400481390722,15.221400481490722,15.269570732274458,15.269570732374458,15.317689320706872,15.317689320806872,15.365729922876003,15.365729922976003,15.413759451705072,15.413759451805072,15.461653539004628,15.461653539104628,15.509248291460374,15.509248291560374,15.55660128255925,15.55660128265925,15.603820566032393,15.603820566132393,15.65094562759775,15.65094562769775,15.698015440683509,15.698015440783509,15.744724798817144,15.744724798917144,15.79141177382923,15.79141177392923,15.837995148090343,15.837995148190343,15.884502388212669,15.884502388312669,15.930748991071043,15.930748991171043,15.976965447647753,15.976965447747753,16.023056314420987,16.023056314520986,16.06906036778441,16.069060367884408,16.115057249091276,16.115057249191274,16.16090923121164,16.16090923131164,16.206749115412435,16.206749115512434,16.252577335169605,16.252577335269603,16.298383171805227,16.298383171905225,16.34414640957748,16.344146409677478,16.38989159898764,16.389891599087637,16.43561728198025,16.435617282080248,16.481301075433763,16.48130107553376,16.526959806096247,16.526959806196245,16.57259213413296,16.57259213423296,16.618144584374996,16.618144584474994,16.663610300021023,16.66361030012102,16.708949243656452,16.70894924375645,16.754078384934264,16.754078385034262,16.7991810841794,16.799181084279397,16.8441261952149,16.8441261953149,16.889067247339273,16.88906724743927,16.934005619794167,16.934005619894165,16.978672399865758,16.978672399965756,17.023211383347242,17.02321138344724,17.067717028587825,17.067717028687824,17.11207505556552,17.112075055665517,17.156414837146595,17.156414837246594,17.200737870793418,17.200737870893416,17.245025595854134,17.245025595954132,17.28925692963503,17.28925692973503,17.33331778914864,17.33331778924864,17.377234143544786,17.377234143644785,17.42100812079629,17.42100812089629,17.464693393106586,17.464693393206584,17.50837042937392,17.508370429473917,17.552029220244638,17.552029220344636,17.59566547036619,17.595665470466187,17.63925920043818,17.639259200538177,17.682727458927403,17.6827274590274,17.72605586231385,17.726055862413848,17.769360109268064,17.769360109368062,17.812658248152136,17.812658248252134,17.855919778022265,17.855919778122264,17.899157979005146,17.899157979105144,17.942227281996907,17.942227282096905,17.985276408653746,17.985276408753744,18.02826867114792,18.02826867124792,18.07123583144416,18.071235831544158,18.11417887471507,18.114178874815067,18.157023361318736,18.157023361418734,18.199821465996237,18.199821466096235,18.242518531371534,18.242518531471532,18.2851953021912,18.2851953022912,18.32786647723048,18.327866477330478,18.370527879357535,18.370527879457534,18.41299362620556,18.412993626305557,18.45543832976677,18.45543832986677,18.497871290070556,18.497871290170554,18.540277847748566,18.540277847848564,18.582670573603227,18.582670573703226,18.62504864008955,18.625048640189547,18.667422568850938,18.667422568950936,18.709462011810025,18.709462011910023,18.75148340640702,18.751483406507017,18.79339611676241,18.793396116862407,18.835302482606235,18.835302482706233,18.877116124004626,18.877116124104624,18.91886553214928,18.918865532249278,18.960610605534477,18.960610605634475,19.002299366453666,19.002299366553665,19.043923421236265,19.043923421336263,19.085541801424668,19.085541801524666,19.12714579809306,19.12714579819306,19.168698211123942,19.16869821122394,19.210209483346905,19.210209483446903,19.251571285182482,19.25157128528248,19.292911019151745,19.292911019251743,19.33404398509501,19.33404398519501,19.374708954644767,19.374708954744765,19.415294401061978,19.415294401161976,19.455821958737012,19.45582195883701,19.496300415409497,19.496300415509495,19.536629047032456,19.536629047132454,19.576956575262102,19.5769565753621,19.61724193810432,19.61724193820432,19.65751004072253,19.65751004082253,19.69773231311123,19.69773231321123,19.737769767119538,19.737769767219536,19.777754849352245,19.777754849452243,19.817725784506372,19.81772578460637,19.85769112388011,19.857691123980107,19.89763589284989,19.89763589294989,19.93751453703453,19.93751453713453,19.977366345117456,19.977366345217455,20.01655863925277,20.01655863935277,20.055680000960646,20.055680001060644,20.133915512912935,20.133915513012933,20.17295234681144,20.172952346911437,20.211885422331356,20.211885422431354,20.25079047954243,20.25079047964243,20.28963965776344,20.28963965786344,20.328443202789465,20.328443202889463,20.36715804287428,20.36715804297428,20.405760376247787,20.405760376347786,20.444351636281237,20.444351636381235,20.482898524140627,20.482898524240625,20.521333614612985,20.521333614712983,20.559675350129446,20.559675350229444,20.59788617591037,20.59788617601037,20.63602165569059,20.63602165579059,20.674137944308494,20.674137944408493,20.712244578234888,20.712244578334886,20.750327962087837,20.750327962187836,20.78838218483173,20.788382184931727,20.82627629732412,20.82627629742412,20.864142352100764,20.864142352200762,20.90183087877428,20.90183087887428,20.93950640116943,20.93950640126943,20.977097947451874,20.977097947551872,21.014644451642887,21.014644451742885,21.05217160704397,21.05217160714397,21.08956875906831,21.08956875916831,21.126809032207287,21.126809032307285,21.163971122048462,21.16397112214846,21.201089115563907,21.201089115663905,21.2381093799571,21.238109380057097,21.27512132949352,21.275121329593517,21.3120433524747,21.312043352574698,21.348848297543697,21.348848297643695,21.3856437849557,21.3856437850557,21.422416219328785,21.422416219428783,21.4591727333126,21.459172733412597,21.4959148243695,21.495914824469498,21.532654393384536,21.532654393484535,21.569382692024988,21.569382692124986,21.606080883790682,21.60608088389068,21.67937004113592,21.679370041235916,21.71593511637944,21.71593511647944,21.752486517427233,21.75248651752723,21.789026608693536,21.789026608793534,21.825538996906232,21.82553899700623,21.86191807156224,21.861918071662238,21.898210726877476,21.898210726977474,21.934461374432907,21.934461374532905,21.97069996347567,21.97069996357567,22.006882988783605,22.006882988883604,22.043046547080895,22.043046547180893,22.07919899263122,22.079198992731218,22.11534489663546,22.115344896735458,22.151395396524787,22.151395396624785,22.187339734214373,22.18733973431437,22.22326231929288,22.223262319392877,22.25911590288224,22.259115902982238,22.294935478313334,22.294935478413333,22.36649550011202,22.366495500212018,22.402151576297857,22.402151576397856,22.43777470831184,22.437774708411837,22.473372501686462,22.47337250178646,22.508869531607207,22.508869531707205,22.544230922963923,22.54423092306392,22.57958167445652,22.57958167455652,22.614915520387243,22.61491552048724,22.650238253571,22.650238253671,22.685519609505416,22.685519609605414,22.720701147751658,22.720701147851656,22.755807261183385,22.755807261283383,22.790887011396244,22.790887011496242,22.825929876746834,22.825929876846832,22.860935187317782,22.86093518741778,22.89587512183477,22.89587512193477,22.930791806278318,22.930791806378316,22.96570336782433,22.965703367924327,23.000586714026976,23.000586714126975,23.035456858916742,23.03545685901674,23.070271794733813,23.07027179483381,23.10505717537278,23.10505717547278,23.13983865472824,23.13983865482824,23.174597947996673,23.17459794809667,23.209352394215898,23.209352394315896,23.24392615977966,23.24392615987966,23.278481955795137,23.278481955895135,23.31301245257816,23.31301245267816,23.347443565148954,23.347443565248952,23.38184618652806,23.381846186628056,23.41621858281169,23.41621858291169,23.45056753198736,23.45056753208736,23.484896659490254,23.48489665959025,23.51921861493659,23.51921861503659,23.55353808774797,23.55353808784797,23.58780869599822,23.587808696098218,23.62207327499214,23.622073275092138,23.656231573565606,23.656231573665604,23.690386404331505,23.690386404431504,23.724489572646085,23.724489572746084,23.758589943070472,23.75858994317047,23.792666432910952,23.79266643301095,23.82662564780473,23.826625647904727,23.860582537691158,23.860582537791156,23.928491076345768,23.928491076445766,23.962433740339804,23.962433740439803,23.996250893344172,23.99625089344417,24.030046884841017,24.030046884941015,24.063836058943448,24.063836059043446,24.097602495262194,24.097602495362192,24.131341110306618,24.131341110406616,24.165058682064235,24.165058682164233,24.198732709192782,24.19873270929278,24.232384195572166,24.232384195672164,24.266028352267377,24.266028352367375,24.299643505481146,24.299643505581145,24.33319856316611,24.33319856326611,24.366725247880098,24.366725247980096,24.400156449665364,24.400156449765362,24.43358280440142,24.43358280450142,24.466946029277054,24.466946029377052,24.500285334161877,24.500285334261875,24.533603083470137,24.533603083570135,24.5668684610028,24.5668684611028,24.600103495219283,24.60010349531928,24.666545111867457,24.666545111967455,24.69974334003548,24.699743340135477,24.732880566315895,24.732880566415893,24.765996197612846,24.765996197712845,24.799101346673297,24.799101346773295,24.832186398212638,24.832186398312636,24.865233028020448,24.865233028120446,24.898275047220476,24.898275047320475,24.93124707975876,24.931247079858757,24.964168080356185,24.964168080456183,24.996944457615434,24.996944457715433,25.029714805618354,25.029714805718353,25.06243841703297,25.062438417132967,25.09513960591913,25.095139606019128,25.1278389820877,25.127838982187697,25.1604652584491,25.1604652585491,25.19305220672017,25.19305220682017,25.22556018355536,25.225560183655357,25.258066347672955,25.258066347772953,25.29051600229002,25.290516002390017,25.322964395886153,25.32296439598615,25.35531888282973,25.35531888292973,25.38765480912146,25.38765480922146,25.419967209491425,25.419967209591423,25.452259867002414,25.452259867102413,25.48443288515245,25.48443288525245,25.5164811804509,25.516481180550898,25.54850346719262,25.54850346729262,25.580524650541022,25.58052465064102,25.612513993110884,25.612513993210882,25.644472598295522,25.64447259839552,25.676428247962345,25.676428248062344,25.708378538290212,25.70837853839021,25.740193111240234,25.740193111340233,25.771945066619583,25.77194506671958,25.80362159718442,25.80362159728442,25.835291546796267,25.835291546896265,25.866933438692367,25.866933438792366,25.898511451996868,25.898511452096866,25.930088164873528,25.930088164973526,25.961637371731104,25.961637371831102,25.993175189993384,25.993175190093382,26.02469153149291,26.024691531592907,26.056200503901362,26.05620050400136,26.08769016692679,26.08769016702679,26.119026379467517,26.119026379567515,26.15035963649043,26.150359636590427,26.181630275942673,26.18163027604267,26.21285559745516,26.21285559755516,26.244079342691485,26.244079342791483,26.275289847208015,26.275289847308013,26.306466264752473,26.30646626485247,26.337602526661616,26.337602526761614,26.36870154904635,26.368701549146348,26.399785912062747,26.399785912162745,26.430858965297656,26.430858965397654,26.492959517386293,26.49295951748629,26.52398461241887,26.52398461251887,26.554998437076865,26.554998437176863,26.58600713883732,26.58600713893732,26.617014579576846,26.617014579676844,26.64799813973246,26.64799813983246,26.67895817396631,26.67895817406631,26.709880692827426,26.709880692927424,26.7407513916096,26.740751391709598,26.771603490333025,26.771603490433023,26.802413935958775,26.802413936058773,26.833147656342177,26.833147656442176,26.863850442305832,26.86385044240583,26.894540105770407,26.894540105870405,26.925186855116383,26.92518685521638,26.95576692798056,26.955766928080557,26.98624631620467,26.98624631630467,27.01669800137518,27.016698001475177,27.047127894527694,27.047127894627693,27.077544389332807,27.077544389432806,27.107935506091657,27.107935506191655,27.138303648625396,27.138303648725394,27.16864069911177,27.168640699211767,27.198970813983017,27.198970814083015,27.229063187001618,27.229063187101616,27.259138378610015,27.259138378710013,27.289195482449415,27.289195482549413,27.319251955778345,27.319251955878343,27.349274184507586,27.349274184607584,27.379287743717914,27.379287743817912,27.40929180586435,27.409291805964347,27.439295158686505,27.439295158786503,27.46927226651051,27.46927226661051,27.499222183570662,27.49922218367066,27.529167174767796,27.529167174867794,27.559070591681067,27.559070591781065,27.58896518144781,27.58896518154781,27.61884893431592,27.61884893441592,27.64863389407536,27.648633894175358,27.708197547896486,27.708197547996484,27.737956853761332,27.73795685386133,27.76762154364935,27.76762154374935,27.797275672487064,27.797275672587062,27.82692172290943,27.826921723009427,27.85654247409934,27.856542474199337,27.886155383315323,27.88615538341532,27.94537008900033,27.94537008910033,27.974898943289798,27.974898943389796,28.00441061616906,28.00441061626906,28.033920515737638,28.033920515837636,28.063425095374157,28.063425095474155,28.092907882992687,28.092907883092685,28.122376405311922,28.12237640541192,28.15182616994479,28.15182617004479,28.181267619720884,28.181267619820883,28.210649446851022,28.21064944695102,28.240017363344,28.240017363443997,28.269326839398143,28.26932683949814,28.327767829927357,28.327767830027355,28.35680303392872,28.35680303402872,28.38575280376192,28.38575280386192,28.414699578670405,28.414699578770403,28.443589055940276,28.443589056040274,28.50131236793138,28.50131236803138,28.530141946706966,28.530141946806964,28.55894358598752,28.55894358608752,28.587734624810864,28.587734624910862,28.616428170953373,28.61642817105337,28.645108279341574,28.645108279441573,28.67377230971289,28.67377230981289,28.70242030147422,28.702420301574218,28.7309867997578,28.7309867998578,28.759534855610244,28.759534855710243,28.788076094068273,28.78807609416827,28.81661591387775,28.81661591397775,28.84513606964207,28.84513606974207,28.873636324919797,28.873636325019795,28.90210659154348,28.902106591643477,28.930567794579208,28.930567794679206,28.959009372976677,28.959009373076675,28.987422736030787,28.987422736130785,29.01582983338714,29.01582983348714,29.044181564043182,29.04418156414318,29.072518832365407,29.072518832465406,29.10078796615038,29.10078796625038,29.129044607946746,29.129044608046744,29.15729916117719,29.157299161277187,29.185546542351084,29.185546542451082,29.213783835357518,29.213783835457516,29.24201864572899,29.24201864582899,29.270249633630762,29.27024963373076,29.29845047525087,29.29845047535087,29.326596028984476,29.326596029084474,29.354739691186683,29.35473969128668,29.382787949273983,29.38278794937398,29.410798652581637,29.410798652681635,29.438769751950637,29.438769752050636,29.466738565719197,29.466738565819195,29.494700956162383,29.49470095626238,29.522656568618057,29.522656568718055,29.550580261481382,29.55058026158138,29.578499619585255,29.578499619685253,29.606416298019642,29.60641629811964,29.66223783281718,29.662237832917178,29.690132719233585,29.690132719333583,29.718022916228392,29.71802291632839,29.74591043355372,29.74591043365372,29.773787586863264,29.773787586963262,29.801646849438328,29.801646849538326,29.829487590768448,29.829487590868446,29.857303702783486,29.857303702883485,29.885117686825698,29.885117686925696,29.912909484780883,29.91290948488088,29.94066699872947,29.940666998829467,29.968390583333598,29.968390583433596,29.99604611221428,29.99604611231428,30.023667160053847,30.023667160153845,30.051286789244866,30.051286789344864,30.078873750112354,30.078873750212352,30.106424417221138,30.106424417321136,30.133960740216814,30.133960740316812,30.161431490124006,30.161431490224004,30.188877098426364,30.188877098526362,30.216321248673268,30.216321248773266,30.243705421612077,30.243705421712075,30.271071152119752,30.27107115221975,30.298434005923426,30.298434006023424,30.325761590547895,30.325761590647893,30.35306289076731,30.353062890867307,30.380363521069352,30.38036352116935,30.407649649630674,30.407649649730672,30.43493191631539,30.434931916415387,30.46220756264021,30.462207562740208,30.489481435654348,30.489481435754346,30.51672945773937,30.516729457839368,30.54397385438921,30.54397385448921,30.57120347345,30.57120347355,30.598428718344433,30.59842871844443,30.625643402188558,30.625643402288556,30.652829161365048,30.652829161465046,30.680009442981863,30.68000944308186,30.707175892775325,30.707175892875323,30.734319959447237,30.734319959547236,30.761460834159916,30.761460834259914,30.788600684293087,30.788600684393085,30.815734386949213,30.81573438704921,30.842847440387573,30.84284744048757,30.869954819231737,30.869954819331735,30.896981926212177,30.896981926312176,30.923990511947675,30.923990512047673,30.950953937371033,30.95095393747103,30.977900654267035,30.977900654367033,31.004845913107587,31.004845913207586,31.03177182315821,31.031771823258207,31.05866750811336,31.058667508213357,31.085558109577875,31.085558109677873,31.112434997439752,31.11243499753975,31.13924769145479,31.13924769155479,31.166019244661914,31.166019244761912,31.192771015603164,31.192771015703162,31.21946730162339,31.219467301723387,31.246156257959445,31.246156258059443,31.272842810474348,31.272842810574346,31.29947486324083,31.29947486334083,31.326087212555247,31.326087212655246,31.352672095257482,31.35267209535748,31.37922860498874,31.379228605088738,31.405781489284816,31.405781489384815,31.432287400551164,31.432287400651163,31.45878125330485,31.458781253404847,31.485246772494456,31.485246772594454,31.51166933815856,31.511669338258557,31.53802223241615,31.53802223251615,31.56435309821433,31.564353098314328,31.590667176671346,31.590667176771344,31.616978417831266,31.616978417931264,31.643259552116426,31.643259552216424,31.669531938068868,31.669531938168866,31.69579640323358,31.695796403333578,31.74831855557549,31.74831855567549,31.774565996957616,31.774565997057614,31.800808551883627,31.800808551983625,31.82700775921509,31.827007759315087,31.853178475354863,31.85317847545486,31.879325744386676,31.879325744486675,31.905462294740563,31.90546229484056,31.931598096363274,31.931598096463272,31.957733228068612,31.95773322816861,31.98384901098402,31.98384901108402,32.00994382942643,32.009943829526435,32.03602745630806,32.036027456408064,32.08819238506399,32.08819238516399,32.11422088168674,32.11422088178674,32.14024122108033,32.14024122118033,32.16624012311808,32.16624012321808,32.192213253040514,32.192213253140515,32.21816321170332,32.21816321180332,32.24409492496952,32.24409492506952,32.27001836278584,32.27001836288584,32.29591453102451,32.29591453112451,32.32169649805498,32.321696498154985,32.34743342299404,32.34743342309404,32.37313322662939,32.373133226729394,32.398829601864094,32.398829601964096,32.42452475548476,32.42452475558476,32.450190353927326,32.45019035402733,32.47584866209263,32.47584866219263,32.527144471577856,32.52714447167786,32.55277582542073,32.55277582552073,32.57832730146892,32.57832730156892,32.603868768163466,32.60386876826347,32.62937752712757,32.629377527227575,32.65486480932893,32.65486480942893,32.68025846072606,32.68025846082606,32.70564620108757,32.70564620118757,32.73102901558606,32.73102901568606,32.756407928801046,32.75640792890105,32.781777541986656,32.78177754208666,32.807144672537305,32.80714467263731,32.83245852495356,32.83245852505356,32.8577712345696,32.8577712346696,32.883058999615315,32.88305899971532,32.90834191761183,32.90834191771183,32.93362105254554,32.93362105264554,32.958886670911134,32.958886671011136,32.9841358565977,32.984135856697705,33.0093796041315,33.009379604231505,33.0346204355544,33.0346204356544,33.05985913900447,33.059859139104475,33.085081606810036,33.08508160691004,33.11027601689986,33.11027601699986,33.135446546405774,33.135446546505776,33.160608091137554,33.160608091237556,33.18574224807095,33.18574224817095,33.21086280962243,33.21086280972243,33.23597493809641,33.235974938196414,33.261009119714025,33.26100911981403,33.28603766614434,33.28603766624434,33.31105955280786,33.311059552907864,33.33606965680704,33.33606965690704,33.3610710124735,33.361071012573504,33.386066890580295,33.3860668906803,33.41103888810318,33.41103888820318,33.43599729024414,33.43599729034414,33.46095277627418,33.460952776374185,33.48590294237218,33.48590294247218,33.51081326809009,33.51081326819009,33.535717446330956,33.53571744643096,33.560604561382334,33.560604561482336,33.58548150945244,33.585481509552444,33.61035660539806,33.61035660549806,33.63517448251887,33.63517448261887,33.659984754107185,33.65998475420719,33.68478438583138,33.68478438593138,33.709576017954035,33.70957601805404,33.73433383895294,33.734333839052944,33.75908306924675,33.75908306934675,33.78377543537789,33.78377543547789,33.808463663784096,33.8084636638841,33.83312497727478,33.83312497737478,33.85776331654035,33.85776331664035,33.882400710040216,33.88240071014022,33.907034438698005,33.90703443879801,33.93166048300949,33.93166048310949,33.956238253863404,33.956238253963406,33.980812005213096,33.9808120053131,34.0053772840784,34.0053772841784,34.029917066676724,34.029917066776726,34.05445196281894,34.05445196291894,34.07898516446428,34.07898516456428,34.1034772253017,34.103477225401704,34.1279604589926,34.1279604590926,34.152430333742984,34.152430333842986,34.17682624173432,34.17682624183432,34.20121675097978,34.201216751079784,34.22560103393439,34.22560103403439,34.24995461891068,34.24995461901068,34.27430485430012,34.27430485440012,34.29865315875125,34.29865315885125,34.32298309958506,34.32298309968506,34.347294952649875,34.34729495274988,34.37155565555312,34.37155565565312,34.39581084148978,34.39581084158978,34.42006539691597,34.420065397015975,34.444315775210335,34.444315775310336,34.4685547649094,34.4685547650094,34.49277846472966,34.49277846482966,34.51698533780185,34.51698533790185,34.541174477767186,34.54117447786719,34.56534608166018,34.56534608176018,34.589512286807306,34.58951228690731,34.613669467773384,34.613669467873386,34.63781238344016,34.637812383540165,34.66192428587339,34.66192428597339,34.68596269443041,34.68596269453041,34.70999219702709,34.70999219712709,34.734016261471,34.734016261571,34.75802822799534,34.75802822809534,34.78203396822883,34.78203396832883,34.80601937449978,34.806019374599785,34.82999729345895,34.829997293558954,34.853964887809234,34.853964887909235,34.87786777602293,34.87786777612293,34.90175257646763,34.90175257656763,34.925631229435275,34.92563122953528,34.9494933709101,34.9494933710101,34.973310155038256,34.97331015513826,34.99711204335665,34.99711204345665,35.020912355398885,35.020912355498886,35.04470935726116,35.044709357361164,35.06848295142239,35.06848295152239,35.09224909767874,35.09224909777874,35.11600523458143,35.11600523468143,35.13972795442942,35.13972795452942,35.16342632081065,35.16342632091065,35.18710407738102,35.187104077481024,35.21077635639172,35.21077635649172,35.23437711187141,35.23437711197141,35.25797049826002,35.257970498360024,35.28155426936403,35.28155426946403,35.305137252329956,35.30513725242996,35.32871038356985,35.32871038366985,35.35228158387144,35.35228158397144,35.37576904450174,35.37576904460174,35.399240269487535,35.39924026958754,35.42268556473041,35.42268556483041,35.44611257516977,35.44611257526977,35.469483864246676,35.46948386434668,35.492842897776406,35.49284289787641,35.51619361644936,35.51619361654936,35.5395404732457,35.5395404733457,35.56287144905968,35.56287144915968,35.586172948509365,35.58617294860937,35.60943764191059,35.60943764201059,35.63269997089757,35.63269997099757,35.655928291726276,35.65592829182628,35.67914636675991,35.67914636685991,35.702263954188,35.702263954288,35.725374606000955,35.72537460610096,35.74847985906805,35.74847985916805,35.77158042271355,35.77158042281355,35.79466494774907,35.79466494784907,35.817742970645405,35.81774297074541,35.840815240133736,35.84081524023374,35.86387588458535,35.86387588468535,35.88693053918753,35.88693053928753,35.90998156835453,35.90998156845453,35.93302605597545,35.933026056075455,35.95605537193828,35.95605537203828,35.9790656543664,35.9790656544664,36.002048746030674,36.002048746130676,36.02502710886645,36.025027108966455,36.04800302847418,36.04800302857418,36.07097473154315,36.070974731643155,36.093917667572114,36.093917667672116,36.116845826012025,36.116845826112026,36.13977110774793,36.139771107847935,36.16266048979417,36.16266048989417,36.1855392319763,36.1855392320763,36.208415452116554,36.208415452216556,36.2312867463938,36.2312867464938,36.25414322367509,36.25414322377509,36.27698878524393,36.27698878534393,36.29979103862512,36.299791038725125,36.34538364450247,36.34538364460247,36.368167613080146,36.36816761318015,36.390943463835576,36.39094346393558,36.413704655222666,36.41370465532267,36.43644657663363,36.43644657673363,36.45916694246804,36.45916694256804,36.481874658686216,36.48187465878622,36.504568543081035,36.50456854318104,36.52724095071311,36.52724095081311,36.54989129047887,36.54989129057887,36.57252724672461,36.57252724682461,36.59515386353408,36.59515386363408,36.61774138869465,36.61774138879465,36.640291950179126,36.64029195027913,36.662825921356976,36.66282592145698,36.68533979501371,36.68533979511371,36.70784066439208,36.70784066449208,36.73033176085823,36.73033176095823,36.75281907426157,36.752819074361575,36.7752642222775,36.7752642223775,36.79770074018142,36.79770074028142,36.82012606652456,36.82012606662456,36.86497518234159,36.86497518244159,36.88739648918051,36.88739648928051,36.9097726751142,36.9097726752142,36.93214641781983,36.93214641791983,36.95451231855154,36.95451231865154,36.97686218067326,36.97686218077326,36.99919931436495,36.99919931446495,37.021533926014776,37.02153392611478,37.04383780027937,37.04383780037937,37.06614064996447,37.06614065006447,37.08841981610017,37.08841981620017,37.1106458223222,37.1106458224222,37.13285523823759,37.13285523833759,37.1550553541236,37.1550553542236,37.177245854725,37.177245854825,37.19942630656585,37.19942630666585,37.22157617864909,37.22157617874909,37.24370157904485,37.243701579144854,37.26581925568742,37.26581925578742,37.287929681459616,37.28792968155962,37.31002430506326,37.31002430516326,37.33211199305177,37.33211199315177,37.35419369119085,37.35419369129085,37.376249183738686,37.37624918383869,37.39830243009299,37.39830243019299,37.420340228940866,37.42034022904087,37.44237333836715,37.44237333846715,37.46440203422017,37.46440203432017,37.48641709528437,37.48641709538437,37.53044059705285,37.530440597152854,37.55244730385337,37.55244730395337,37.57445211912248,37.574452119222485,37.596449407672914,37.596449407772916,37.61844279493983,37.61844279503983,37.64042305970767,37.640423059807674,37.66237904982256,37.66237904992256,37.68432897127422,37.68432897137422,37.70627120837957,37.70627120847957,37.728206746311216,37.72820674641122,37.7501111527886,37.7501111528886,37.77201232789984,37.77201232799984,37.79390526696812,37.79390526706812,37.81578090640548,37.81578090650548,37.83765221108338,37.83765221118338,37.85952142719537,37.85952142729537,37.881376890297815,37.88137689039782,37.903207251202325,37.90320725130233,37.92500809633564,37.92500809643564,37.94676185021851,37.94676185031851,37.96848557604043,37.96848557614043,37.99016682121969,37.99016682131969,38.011820087497014,38.011820087597016,38.03345463549488,38.03345463559488,38.055084730512576,38.05508473061258,38.07670387041092,38.07670387051092,38.098287504688635,38.09828750478864,38.119865700813584,38.119865700913586,38.1414263608662,38.1414263609662,38.162985050573596,38.1629850506736,38.206093484621164,38.206093484721166,38.2276334560491,38.2276334561491,38.24916523084098,38.24916523094098,38.27069621749477,38.270696217594775,38.29221837700204,38.29221837710204,38.31373025130733,38.31373025140733,38.335239918825984,38.335239918925986,38.35674576387494,38.35674576397494,38.37825034790297,38.37825034800297,38.39974054841098,38.399740548510984,38.421218808627046,38.42121880872705,38.44268548321329,38.44268548331329,38.46415014804742,38.464150148147425,38.50707778321882,38.50707778331882,38.52853354209262,38.52853354219262,38.549974405156654,38.549974405256656,38.57140151521115,38.57140151531115,38.59280312899867,38.59280312909867,38.6141591095912,38.6141591096912,38.63549652953188,38.63549652963188,38.656830915140944,38.656830915240946,38.67816199057006,38.67816199067006,38.69943336453941,38.69943336463941,38.72070398977758,38.720703989877585,38.741946872555246,38.74194687265525,38.763182031579696,38.7631820316797,38.78437628623766,38.78437628633766,38.80552254328638,38.80552254338638,38.82661792602186,38.82661792612186,38.84769983159613,38.84769983169613,38.868778939280205,38.868778939380206,38.88982809771714,38.889828097817144,38.910863345516916,38.91086334561692,38.93189406152271,38.931894061622714,38.95292296481092,38.95292296491092,38.97394524773923,38.97394524783923,38.994945738649555,38.99494573874956,39.015939963862124,39.015939963962126,39.03692457379009,39.03692457389009,39.05789499723256,39.05789499733256,39.07884150068422,39.07884150078422,39.09976424177268,39.099764241872684,39.120670826030455,39.12067082613046,39.141558692008765,39.14155869210877,39.162444942304006,39.16244494240401,39.183323981135786,39.18332398123579,39.20416857833335,39.204168578433354,39.22500761915744,39.22500761925744,39.245838975635216,39.24583897573522,39.266662687173586,39.26666268727359,39.28747958131755,39.28747958141755,39.308294859778435,39.30829485987844,39.329104660679654,39.329104660779656,39.34990855054525,39.34990855064525,39.37067791996282,39.370677920062825,39.39144563429042,39.39144563439042,39.41221169352805,39.41221169362805,39.43296364509399,39.43296364519399,39.45370416865774,39.45370416875774,39.474421520961855,39.474421521061856,39.49513453850651,39.495134538606514,39.515836640338726,39.51583664043873,39.53648140512542,39.53648140522542,39.557122583883846,39.55712258398385,39.577760058393274,39.577760058493276,39.59838988796331,39.59838988806331,39.61899508821825,39.61899508831825,39.63959697829325,39.63959697839325,39.66018306619969,39.660183066299695,39.68075157862688,39.680751578726884,39.70131626858438,39.70131626868438,39.721875520389105,39.72187552048911,39.74241365009321,39.742413650193214,39.76293747509112,39.76293747519112,39.78344707419663,39.78344707429663,39.80394311732712,39.803943117427124,39.82443194899415,39.824431949094155,39.84491262343204,39.84491262353204,39.86539231269731,39.86539231279731,39.8858645934646,39.885864593564605,39.90632406698806,39.906324067088065,39.94723607841984,39.94723607851984,39.96768818285222,39.96768818295222,39.988124524522945,39.98812452462295,40.00855818652419,40.00855818662419,40.02897383957024,40.02897383967024,40.04938054724906,40.04938054734906,40.069783511271986,40.06978351137199,40.09018135239737,40.090181352497375,40.110572139686916,40.11057213978692,40.13095977442413,40.13095977452413,40.151343468470934,40.151343468570936,40.1717144340877,40.1717144341877,40.1920762573027,40.192076257402704,40.212412860099064,40.212412860199066,40.23274276372172,40.23274276382172,40.25307199742701,40.25307199752701,40.2733965811176,40.273396581217604,40.29372045548393,40.29372045558393,40.314039325173425,40.31403932527343,40.33434349608768,40.334343496187685,40.35462946101223,40.35462946111223,40.37490214581008,40.374902145910085,40.39516947126897,40.39516947136897,40.45596660059643,40.455966600696435,40.47620484376007,40.476204843860074,40.49644143183374,40.49644143193374,40.516630573994824,40.516630574094826,40.53679209191611,40.536792092016114,40.55692350296264,40.55692350306264,40.57704711144215,40.57704711154215,40.597167567369326,40.59716756746933,40.61727742283929,40.61727742293929,40.65747565701647,40.65747565711647,40.67756253826132,40.67756253836132,40.69763988303537,40.69763988313537,40.71770430234487,40.71770430244487,40.73774909701611,40.737749097116115,40.75778794124482,40.75778794134482,40.797819050741566,40.79781905084157,40.817805168532544,40.817805168632546,40.83777229219573,40.83777229229573,40.85771703273737,40.85771703283737,40.877651251635605,40.87765125173561,40.89757301795213,40.897573018052135,40.91749293214417,40.91749293224417,40.93741130946694,40.93741130956694,40.95732795288593,40.957327952985935,40.97723135558513,40.977231355685134,40.997106858196204,40.997106858296206,41.01697711968903,41.01697711978903,41.03682318534271,41.036823185442714,41.056668147603084,41.056668147703085,41.07650893273161,41.076508932831615,41.09634152122408,41.096341521324085,41.136005752443324,41.136005752543326,41.17566612178596,41.17566612188596,41.195491222966645,41.19549122306665,41.21527021806949,41.21527021816949,41.23503486905923,41.23503486915923,41.25478155050069,41.25478155060069,41.274521729802956,41.27452172990296,41.29425942647027,41.29425942657027,41.313966582786875,41.31396658288688,41.333671847572084,41.333671847672086,41.35337293522546,41.35337293532546,41.373066102091094,41.373066102191096,41.392753909617774,41.392753909717776,41.41243750060571,41.412437500705714,41.432114471233746,41.43211447133375,41.45174821248795,41.45174821258795,41.471342546838024,41.471342546938025,41.49092293114403,41.49092293124403,41.51049602517276,41.510496025272765,41.530062183586374,41.530062183686375,41.54962495300623,41.54962495310623,41.56917377238202,41.569173772482024,41.58872101548165,41.588721015581655,41.608252811074856,41.60825281117486,41.62776458796076,41.627764588060764,41.647264503368525,41.64726450346853,41.666756025105705,41.66675602520571,41.686244276069836,41.68624427616984,41.70571932572109,41.70571932582109,41.72518511474987,41.72518511484987,41.74464885461963,41.744648854719635,41.76410621057093,41.76410621067093,41.78355028639553,41.78355028649553,41.80295479768838,41.80295479778838,41.82235324031799,41.82235324041799,41.84175002785763,41.84175002795763,41.86114224419639,41.86114224429639,41.88052606686456,41.88052606696456,41.899903150952134,41.899903151052136,41.91927483629384,41.91927483639384,41.93864163517943,41.93864163527943,41.95800622727839,41.95800622737839,41.9773588790854,41.9773588791854,41.99670684147081,41.99670684157081,42.01604735595134,42.016047356051345,42.03538653059714,42.03538653069714,42.05472428659438,42.05472428669438,42.07405557985935,42.07405557995935,42.09337197731455,42.09337197741455,42.11268325187221,42.112683251972214,42.13198250732412,42.13198250742412,42.15124791224537,42.15124791234537,42.17050890359336,42.17050890369336,42.18975275293805,42.18975275303805,42.208992543371615,42.208992543471616,42.22823103337734,42.22823103347734,42.24745860767062,42.247458607770625,42.26666927640203,42.26666927650203,42.285876556139684,42.285876556239685,42.30506803370877,42.30506803380877,42.324254230752715,42.324254230852716,42.34342194595862,42.34342194605862,42.36258847895739,42.36258847905739,42.381749455582685,42.38174945568269,42.400903338965236,42.40090333906524,42.42005158716049,42.420051587260495,42.439179737834635,42.43917973793464,42.4582784515514,42.4582784516514,42.47736431861741,42.47736431871741,42.49643079748659,42.49643079758659,42.5154880945471,42.515488094647104,42.534536840309414,42.534536840409416,42.57262901190199,42.57262901200199,42.59165919701246,42.59165919711246,42.610684692701334,42.610684692801335,42.629689302731016,42.62968930283102,42.64868697714557,42.648686977245575,42.667681932483745,42.66768193258375,42.68664445593981,42.686644456039815,42.70560603363018,42.705606033730184,42.72452532771241,42.724525327812415,42.74343595227574,42.74343595237574,42.76234401539029,42.76234401549029,42.781224178416714,42.781224178516716,42.80008554434987,42.80008554444987,42.818932920832054,42.818932920932056,42.837770721436534,42.837770721536536,42.856599300825444,42.856599300925446,42.87542630393819,42.87542630403819,42.913076487693985,42.91307648779399,42.9318822110785,42.931882211178504,42.95068568826948,42.95068568836948,42.96946276283469,42.96946276293469,42.98823526619901,42.988235266299014,43.025778854279125,43.025778854379126,43.06331377284032,43.06331377294032,43.08205415957778,43.08205415967778,43.10077208437988,43.10077208447988,43.11948618671228,43.11948618681228,43.13818795468368,43.13818795478368,43.15688869807558,43.156888698175585,43.17558337280424,43.17558337290424,43.19425854111535,43.194258541215355,43.21291542462295,43.21291542472295,43.250226236120334,43.250226236220335,43.268855180132896,43.2688551802329,43.28748329660047,43.28748329670047,43.30611019145402,43.30611019155402,43.32471694937956,43.32471694947956,43.34331038777149,43.34331038787149,43.36189708758282,43.36189708768282,43.38047275346099,43.380472753560994,43.39903316886726,43.399033168967264,43.41755279812776,43.41755279822776,43.43606584643526,43.436065846535264,43.4545754663421,43.4545754664421,43.47307302773628,43.47307302783628,43.49156538741911,43.49156538751911,43.51005282123892,43.510052821338924,43.528528630022016,43.52852863012202,43.54700116803207,43.54700116813207,43.5654506924101,43.565450692510105,43.583899310429345,43.58389931052935,43.60233011652792,43.60233011662792,43.62074992810023,43.62074992820023,43.6391663900857,43.6391663901857,43.657500294607,43.657500294707,43.675824386809175,43.67582438690918,43.69414623281781,43.69414623291781,43.71246736950217,43.71246736960217,43.730784801937546,43.73078480203755,43.74909364366782,43.74909364376782,43.76740000276313,43.767400002863134,43.785698953360466,43.78569895346047,43.8039902196115,43.8039902197115,43.82227959433113,43.82227959443113,43.84056498895344,43.84056498905344,43.858843566181335,43.85884356628134,43.877096450107736,43.87709645020774,43.89532718735401,43.895327187454015,43.91354105844532,43.91354105854532,43.93175063418408,43.93175063428408,43.94994472300951,43.94994472310951,43.96812470416326,43.96812470426326,43.98629841961925,43.986298419719255,44.02263824499874,44.02263824509874,44.04080285745988,44.04080285755988,44.05896400211345,44.05896400221345,44.095283178275174,44.095283178375176,44.11343565340984,44.11343565350984,44.149715462074326,44.14971546217433,44.16783806677566,44.16783806687566,44.18595968630438,44.18595968640438,44.20407846853601,44.20407846863601,44.22218428589618,44.22218428599618,44.24028892104922,44.24028892114922,44.258366601879835,44.25836660197984,44.29451171774599,44.29451171784599,44.31257836464345,44.31257836474345,44.3487071660513,44.3487071661513,44.36675887773209,44.36675887783209,44.384798097424266,44.38479809752427,44.40282852937682,44.40282852947682,44.42085793674987,44.42085793684987,44.43888501911558,44.43888501921558,44.456897599740564,44.456897599840566,44.4748954815903,44.474895481690304,44.49288682189396,44.49288682199396,44.51087150243082,44.51087150253082,44.52884463674476,44.52884463684476,44.54681055959525,44.54681055969525,44.56476194129811,44.56476194139811,44.58270851535867,44.58270851545867,44.600650794066674,44.600650794166675,44.61858306342103,44.61858306352103,44.63651135267807,44.63651135277807,44.6544204507728,44.6544204508728,44.67232619928067,44.67232619938067,44.7081356471374,44.7081356472374,44.72603552401656,44.72603552411656,44.74389634865371,44.743896348753715,44.76175398133164,44.76175398143164,44.77960357500112,44.77960357510112,44.797441583040786,44.79744158314079,44.81527797539738,44.81527797549738,44.833112042746635,44.833112042846636,44.850938701597904,44.850938701697906,44.86875081930155,44.868750819401555,44.886552060699664,44.886552060799666,44.904350031324725,44.90435003142473,44.922135194705945,44.922135194805946,44.93991618095533,44.93991618105533,44.95769452694213,44.95769452704213,44.975466764858794,44.975466764958796,44.99322201839978,44.99322201849978,45.01095189389449,45.010951893994495,45.02868086303042,45.02868086313042,45.04638551810649,45.04638551820649,45.064083552822666,45.06408355292267,45.08175888916206,45.08175888926206,45.09942122122309,45.09942122132309,45.11707827275897,45.11707827285897,45.1347339844601,45.1347339845601,45.1523884351403,45.152388435240304,45.17003165485283,45.17003165495283,45.18765048169169,45.18765048179169,45.20526678648869,45.205266786588695,45.2228717815042,45.222871781604205,45.240458334088586,45.24045833418859,45.275629784167364,45.275629784267366,45.29321263250276,45.29321263260276,45.310783185884056,45.31078318598406,45.34591897271459,45.345918972814594,45.36345453276502,45.36345453286502,45.38098236906224,45.38098236916224,45.398482659933464,45.398482660033466,45.41598200503899,45.41598200513899,45.43347709419887,45.43347709429887,45.45096449901244,45.450964499112445,45.46844650508015,45.468446505180154,45.48592579207147,45.48592579217147,45.5033945968263,45.5033945969263,45.52084763881946,45.52084763891946,45.538299498605504,45.538299498705506,45.55574954567396,45.55574954577396,45.573184736339556,45.57318473643956,45.60804526594471,45.60804526604471,45.64289515568575,45.64289515578575,45.660312889092815,45.66031288919282,45.677727942830394,45.677727942930396,45.71254898671761,45.71254898681761,45.72994378530646,45.72994378540646,45.747336298294876,45.74733629839488,45.76472688034499,45.76472688044499,45.782105088627205,45.78210508872721,45.799478646894734,45.799478646994736,45.81684538776784,45.81684538786784,45.851572249154174,45.851572249254176,45.86889749455723,45.86889749465723,45.88622128190483,45.88622128200483,45.90354026161013,45.903540261710134,45.92085754681855,45.920857546918555,45.938164349790476,45.93816434989048,45.955439351390766,45.95543935149077,45.97271100340419,45.972711003504195,45.98996303077937,45.98996303087937,46.00720098988977,46.00720098998977,46.04166551951527,46.04166551961527,46.05888180482839,46.05888180492839,46.07609296724398,46.07609296734398,46.127708327314835,46.12770832741484,46.14489718542268,46.14489718552268,46.16207481256284,46.162074812662844,46.179238056183,46.179238056283005,46.19639901420272,46.19639901430272,46.21353412129332,46.21353412139332,46.23066859787346,46.23066859797346,46.247795232479675,46.247795232579676,46.26489128732828,46.264891287428284,46.281954003935994,46.281954004035995,46.29900005142325,46.29900005152325,46.316041567116535,46.31604156721654,46.33307969381606,46.33307969391606,46.350113682790656,46.35011368289066,46.36713856877039,46.367138568870395,46.38415943524591,46.384159435345914,46.40117703094838,46.401177031048384,46.41815845111286,46.41815845121286,46.43513660050429,46.435136600604295,46.469089667921025,46.46908966802103,46.486063679587524,46.486063679687526,46.50302374120996,46.50302374130996,46.519978325272724,46.519978325372726,46.53692885042436,46.53692885052436,46.55386589841478,46.55386589851478,46.587734201580716,46.58773420168072,46.604658560548,46.604658560648005,46.62158027925272,46.62158027935272,46.63849719031512,46.63849719041512,46.655402791596046,46.65540279169605,46.6891998470793,46.6891998471793,46.706087005929085,46.70608700602909,46.7229658499221,46.7229658500221,46.73983275362316,46.739832753723164,46.756698238675675,46.75669823877568,46.77355564531284,46.77355564541284,46.79039634342265,46.79039634352265,46.80723621398747,46.807236214087474,46.82406903071645,46.82406903081645,46.84089644869957,46.84089644879957,46.85771318741167,46.85771318751167,46.87451542438304,46.87451542448304,46.8913080066629,46.891308006762905,46.90809739698353,46.90809739708353,46.924877250833354,46.924877250933356,46.941654306792984,46.941654306892985,46.95841993475042,46.95841993485042,46.975184144059305,46.97518414415931,46.991946895312736,46.99194689541274,47.00870566646885,47.00870566656885,47.02546258550046,47.025462585600465,47.04221375112408,47.04221375122408,47.058963970982,47.058963971082,47.075713087446594,47.075713087546596,47.09245333735777,47.09245333745777,47.10918014951463,47.10918014961463,47.12590475488486,47.12590475498486,47.142622267012335,47.14262226711234,47.15933280411779,47.15933280421779,47.17604125265732,47.17604125275732,47.192745957540964,47.192745957640966,47.20944486960969,47.20944486970969,47.24281503010045,47.242815030200454,47.2594816679147,47.2594816680147,47.276138296375294,47.276138296475295,47.29278329979917,47.29278329989917,47.30942361380145,47.30942361390145,47.32606132694806,47.32606132704806,47.34269750322541,47.34269750332541,47.35932977821925,47.35932977831925,47.37595866421933,47.375958664319334,47.39257726501744,47.39257726511744,47.40919082173181,47.409190821831814,47.42578231057987,47.42578231067987,47.44237151382749,47.44237151392749,47.45895882554371,47.45895882564371,47.49213218795522,47.49213218805522,47.50871756873314,47.50871756883314,47.52529987577253,47.525299875872534,47.54187851796984,47.54187851806984,47.55845574151861,47.55845574161861,47.57502973370123,47.57502973380123,47.5915956080616,47.591595608161605,47.60814512855676,47.60814512865676,47.62468117189069,47.624681171990694,47.641191206667905,47.64119120676791,47.65769773423065,47.657697734330654,47.674197602026595,47.6741976021266,47.69069490837377,47.69069490847377,47.707190677851685,47.70719067795169,47.72368096976992,47.723680969869925,47.74016156758975,47.74016156768975,47.77309825213502,47.77309825223502,47.78954681214178,47.78954681224178,47.80597842717977,47.80597842727977,47.822400072271,47.822400072371,47.838817106754455,47.83881710685446,47.855232604368645,47.85523260446865,47.871645974010015,47.87164597411002,47.88803337450155,47.888033374601555,47.90441336649512,47.90441336659512,47.92079170339871,47.920791703498715,47.93714434700081,47.93714434710081,47.95348962151183,47.95348962161183,47.969833043898355,47.96983304399836,47.98615558062569,47.98615558072569,48.00247453132475,48.00247453142475,48.018784142587535,48.01878414268754,48.03509217757415,48.03509217767415,48.051390439648536,48.05139043974854,48.06768101737662,48.06768101747662,48.08396686627621,48.08396686637621,48.100251887630805,48.100251887730806,48.11650400422045,48.116504004320454,48.13274843646379,48.132748436563794,48.14898238647063,48.14898238657063,48.16520242584031,48.165202425940315,48.18141844570577,48.181418445805775,48.19763292870197,48.19763292880197,48.2138410671866,48.2138410672866,48.23004262471824,48.23004262481824,48.2462424483461,48.246242448446104,48.2624322232134,48.262432223313404,48.27862050061835,48.278620500718354,48.29479932036631,48.29479932046631,48.31096407184948,48.310964071949485,48.3271272864634,48.3271272865634,48.343282619696495,48.3432826197965,48.375587375127054,48.375587375227056,48.391724541777336,48.39172454187734,48.407856033833426,48.40785603393343,48.42397795001181,48.42397795011181,48.44009438863052,48.44009438873052,48.45620491621361,48.45620491631361,48.472307641229676,48.47230764132968,48.488405203941305,48.48840520404131,48.50448645219462,48.50448645229462,48.52056624239248,48.520566242492485,48.536623097772136,48.53662309787214,48.552669352694586,48.55266935279459,48.584755636248616,48.58475563634862,48.60079196063122,48.60079196073122,48.61682410788198,48.616824107981984,48.63284928011071,48.63284928021071,48.64886759553813,48.64886759563813,48.66488204908893,48.66488204918893,48.6808808975057,48.680880897605704,48.69686768740981,48.69686768750981,48.712852309934185,48.71285231003419,48.72882152435904,48.72882152445904,48.74478250274094,48.74478250284094,48.760739028142666,48.76073902824267,48.77669137641256,48.77669137651256,48.79264021746798,48.79264021756798,48.808584132660386,48.80858413276039,48.82452678683185,48.824526786931855,48.84046230835368,48.84046230845368,48.872320110677535,48.87232011077754,48.88822083590301,48.888220836003015,48.904117226369024,48.904117226469026,48.92000679944063,48.92000679954063,48.93589400809798,48.935894008197984,48.95177585741638,48.95177585751638,48.967651835106054,48.967651835206055,48.98351603013139,48.98351603023139,48.999378057777,48.999378057877,49.01521407686589,49.01521407696589,49.03104784976123,49.03104784986123,49.04687783959378,49.046877839693785,49.0626980171072,49.062698017207204,49.07850960391552,49.07850960401552,49.094316777150574,49.094316777250576,49.11011437450792,49.110114374607925,49.125890889171565,49.12589088927157,49.141662202123854,49.141662202223856,49.15742768285434,49.15742768295434,49.173190168660106,49.17319016876011,49.1889519845485,49.188951984648504,49.204709386863634,49.204709386963636,49.22045579465251,49.22045579475251,49.236200429130704,49.236200429230706,49.25194364496035,49.25194364506035,49.26768607265191,49.267686072751914,49.28341600835486,49.283416008454864,49.2991449194783,49.2991449195783,49.31486456997927,49.31486457007927,49.33057960987246,49.33057960997246,49.34629378281376,49.34629378291376,49.361998852760195,49.3619988528602,49.37769749938126,49.377697499481265,49.39337529975005,49.39337529985005,49.42472952124596,49.42472952134596,49.44039321394306,49.44039321404306,49.456036887932854,49.456036888032855,49.471670473755196,49.4716704738552,49.487299646004274,49.487299646104276,49.51854904513519,49.51854904523519,49.53415784401483,49.53415784411483,49.54976041660361,49.54976041670361,49.565360191302204,49.565360191402206,49.58095350326852,49.58095350336852,49.59654512073795,49.59654512083795,49.61212010849384,49.61212010859384,49.627689500469344,49.627689500569346,49.64325818312057,49.643258183220574,49.65882521068183,49.658825210781835,49.67439160773262,49.674391607832625,49.68995713783152,49.68995713793152,49.721085557766735,49.721085557866736,49.73663512806946,49.73663512816946,49.75217374325283,49.752173743352834,49.76771105800837,49.767711058108375,49.78324671767393,49.783246717773935,49.79877957944931,49.79877957954931,49.814302353057215,49.81430235315722,49.84533391082207,49.84533391092207,49.86083544410865,49.860835444208654,49.876327440924435,49.876327441024436,49.891809034317525,49.89180903441753,49.90728140649505,49.907281406595054,49.922748576961226,49.92274857706123,49.938206013922084,49.938206014022086,49.95365887968206,49.95365887978206,49.969110050945154,49.969110051045156,49.984551252261504,49.984551252361506,49.99997334122935,49.99997334132935,50.01539219883106,50.01539219893106,50.03080695811474,50.03080695821474,50.04621576695589,50.04621576705589,50.06161480288482,50.06161480298482,50.092392737814656,50.09239273791466,50.10777076986368,50.10777076996368,50.123140644683545,50.12314064478355,50.13849826395623,50.13849826405623,50.15385158787635,50.153851587976355,50.16919978889895,50.16919978899895,50.184539872099286,50.18453987219929,50.21521546729909,50.21521546739909,50.23054782674622,50.23054782684622,50.24587009802589,50.24587009812589,50.26119055658797,50.26119055668797,50.27650802022534,50.276508020325345,50.29180755372133,50.29180755382133,50.3071022401681,50.3071022402681,50.32239310414518,50.32239310424518,50.33767592911381,50.33767592921381,50.352953985847044,50.352953985947046,50.3682303874903,50.3682303875903,50.38350497641596,50.383504976515965,50.39877601872026,50.398776018820264,50.41404327796176,50.41404327806176,50.42929690241443,50.42929690251443,50.4445458374455,50.444545837545505,50.45977936437706,50.459779364477065,50.47500942350106,50.47500942360106,50.490233059299676,50.49023305939968,50.52066575034239,50.52066575044239,50.535859673335295,50.5358596734353,50.55105060140348,50.55105060150348,50.566235106146294,50.566235106246296,50.58141350281897,50.58141350291897,50.59658992914644,50.59658992924644,50.61174298717975,50.611742987279754,50.64202884809766,50.64202884819766,50.65715179925621,50.657151799356214,50.67227333176622,50.67227333186622,50.687393879103624,50.687393879203626,50.70250953998491,50.70250954008491,50.747852463717656,50.74785246381766,50.76296024321812,50.76296024331812,50.77806613118718,50.77806613128718,50.793171191611265,50.79317119171127,50.808264311743386,50.80826431184339,50.823355855599345,50.82335585569935,50.83844649309651,50.838446493196514,50.85353133777877,50.85353133787877,50.86861165066705,50.86861165076705,50.88368451565045,50.88368451575045,50.898753558164145,50.89875355826415,50.913806798509285,50.913806798609286,50.92885231510122,50.92885231520122,50.943896452451504,50.943896452551506,50.958939131746334,50.958939131846336,50.97397381143963,50.97397381153963,50.98900518095297,50.989005181052974,51.004024058477704,51.004024058577706,51.019040138112246,51.01904013821225,51.0340553902018,51.0340553903018,51.04906902660829,51.04906902670829,51.07909527484174,51.07909527494174,51.09409701036318,51.09409701046318,51.12409847165394,51.12409847175394,51.15409047528771,51.15409047538771,51.184070341595,51.184070341695005,51.1990488664499,51.1990488665499,51.214021440862275,51.21402144096228,51.24396178204472,51.24396178214472,51.25892324371013,51.25892324381013,51.273877572725894,51.273877572825896,51.28881397160027,51.28881397170027,51.30373839077579,51.30373839087579,51.31865571670857,51.31865571680857,51.333555112499965,51.33355511259997,51.348453798967086,51.34845379906709,51.36334472227409,51.363344722374094,51.3782337146428,51.378233714742805,51.39311971208679,51.393119712186795,51.40800220231631,51.40800220241631,51.4228796090552,51.4228796091552,51.43775634587672,51.43775634597672,51.452628590311164,51.452628590411166,51.4823658677166,51.482365867816604,51.497211276049335,51.497211276149336,51.526897048631064,51.526897048731065,51.541725827250254,51.541725827350255,51.58620593681696,51.58620593691696,51.601027149310546,51.60102714941055,51.61584666730726,51.61584666740726,51.63066326919307,51.63066326929307,51.645473920636356,51.64547392073636,51.6602813801204,51.660281380220404,51.67508387433453,51.675083874434534,51.689882270230626,51.68988227033063,51.70467956273341,51.70467956283341,51.71947307217339,51.719473072273395,51.734259291336116,51.73425929143612,51.74903313673094,51.74903313683094,51.763805169408165,51.76380516950817,51.778575822843756,51.77857582294376,51.79333646692569,51.79333646702569,51.80809285506198,51.80809285516198,51.82284254402457,51.82284254412457,51.837582341854215,51.83758234195422,51.85232064222151,51.85232064232151,51.86705460782934,51.86705460792934,51.88176051572143,51.88176051582143,51.91116720860808,51.91116720870808,51.92586125502203,51.92586125512203,51.94055084845581,51.94055084855581,51.955238589765095,51.9552385898651,51.96992045944566,51.96992045954566,51.98458266508107,51.98458266518107,51.99924167875724,51.99924167885724,52.013899667853906,52.01389966795391,52.02855537135013,52.02855537145013,52.04320161718936,52.04320161728936,52.05784561683505,52.057845616935055,52.072488394866724,52.072488394966726,52.08711517369531,52.08711517379531,52.11634130414721,52.116341304247214,52.1455339781375,52.1455339782375,52.16012024666863,52.160120246768635,52.17470564824788,52.17470564834788,52.18928860659907,52.18928860669907,52.21844632666539,52.218446326765395,52.23301829049033,52.23301829059033,52.24757133900128,52.247571339101285,52.26212107733228,52.26212107743228,52.2766671114143,52.2766671115143,52.29120317554957,52.29120317564957,52.30573722993273,52.30573723003273,52.320269865667335,52.32026986576734,52.334799703511756,52.33479970361176,52.34932745279025,52.34932745289025,52.36384554737723,52.36384554747723,52.37834397791905,52.37834397801905,52.40733914450582,52.40733914460582,52.42183477715744,52.421834777257445,52.43632351360085,52.43632351370085,52.450807442401945,52.45080744250195,52.465287391105726,52.46528739120573,52.47976359615361,52.47976359625361,52.49423523000062,52.49423523010062,52.508701228660335,52.50870122876034,52.52316182857419,52.52316182867419,52.53761943356332,52.53761943366332,52.55207585634533,52.552075856445335,52.56652743207814,52.56652743217814,52.58097636754837,52.58097636764837,52.595409973732885,52.595409973832886,52.60984160957219,52.609841609672195,52.624263945382125,52.62426394548213,52.63868100066691,52.63868100076691,52.65309553390983,52.65309553400983,52.66750821502825,52.66750821512825,52.68191723130459,52.68191723140459,52.69632045476602,52.69632045486602,52.71071359005998,52.71071359015998,52.72510262703592,52.725102627135925,52.739489575445944,52.739489575545946,52.75387577512479,52.75387577522479,52.76825811292703,52.76825811302703,52.782637337583836,52.78263733768384,52.79701293680547,52.797012936905475,52.811386959750934,52.811386959850935,52.82575325894319,52.82575325904319,52.84011782423166,52.840117824331664,52.85448073443016,52.85448073453016,52.88320147133653,52.88320147143653,52.89755023445644,52.89755023455644,52.911894426375476,52.91189442647548,52.92622541698162,52.926225417081625,52.9405528609664,52.9405528610664,52.954874472729365,52.95487447282937,52.96918997642219,52.969189976522195,52.98350063306581,52.98350063316581,52.99778638454601,52.99778638464601,53.01206839237032,53.01206839247032,53.026345198483284,53.026345198583286,53.04061810331274,53.04061810341274,53.05488698863797,53.05488698873797,53.069152957852296,53.0691529579523,53.083416286804045,53.08341628690405,53.11193849172737,53.111938491827374,53.126171595583656,53.12617159568366,53.140401665108314,53.140401665208316,53.154630079543,53.154630079643,53.16885534142535,53.16885534152535,53.18307461345828,53.18307461355828,53.19728895962819,53.19728895972819,53.22571615450565,53.22571615460565,53.23992502311588,53.23992502321588,53.25412963578047,53.254129635880474,53.28251856655402,53.28251856665402,53.29669496387525,53.29669496397525,53.310863598036356,53.31086359813636,53.32502805506563,53.32502805516563,53.339175212463985,53.33917521256399,53.35332142409665,53.35332142419665,53.36746432554936,53.367464325649365,53.38159635069653,53.381596350796535,53.3956997664313,53.3956997665313,53.409800660124205,53.40980066022421,53.4238976525336,53.4238976526336,53.43797533555997,53.43797533565997,53.45203753167301,53.45203753177301,53.466096890488956,53.46609689058896,53.49421367718254,53.494213677282545,53.522327784206645,53.52232778430665,53.53637067093666,53.536370671036664,53.55041028689364,53.55041028699364,53.56444584393949,53.56444584403949,53.57847828783991,53.57847828793991,53.592508879615835,53.59250887971584,53.60653375739066,53.60653375749066,53.62054303003145,53.62054303013145,53.63455048995465,53.63455049005465,53.648544393902824,53.648544394002826,53.67653137425419,53.67653137435419,53.690519051911515,53.69051905201152,53.704499833360615,53.704499833460616,53.718469383842034,53.718469383942036,53.732437476268004,53.732437476368005,53.74640190385189,53.74640190395189,53.77432768528113,53.77432768538113,53.78825696190653,53.78825696200653,53.80218154911033,53.80218154921033,53.816104126562024,53.816104126662026,53.830023709089,53.830023709189,53.84393442506255,53.84393442516255,53.85783410710294,53.85783410720294,53.87173244930859,53.87173244940859,53.885625826244315,53.88562582634432,53.89951415909631,53.899514159196315,53.913397684306005,53.91339768440601,53.92727998790167,53.92727998800167,53.94116166098687,53.94116166108687,53.9550426247478,53.9550426248478,53.96891988425973,53.968919884359735,53.98279336070887,53.98279336080887,53.99665493627296,53.99665493637296,54.0105141080159,54.0105141081159,54.024370481868644,54.024370481968646,54.03821022600784,54.03821022610784,54.05204595064282,54.05204595074282,54.065880690105196,54.0658806902052,54.07971014904242,54.07971014914242,54.093531293122865,54.09353129322287,54.1073285172125,54.1073285173125,54.12112262815671,54.121122628256714,54.134915241638566,54.13491524173857,54.14870080128458,54.148700801384585,54.1624788342119,54.162478834311905,54.176256157814954,54.176256157914956,54.19003225980398,54.19003225990398,54.20379933761195,54.20379933771195,54.23132864617869,54.23132864627869,54.24509044346151,54.24509044356151,54.25885058565436,54.25885058575436,54.28636882088103,54.28636882098103,54.30012419483848,54.30012419493848,54.31385777677794,54.31385777687794,54.3275827680123,54.3275827681123,54.34130393677695,54.341303936876955,54.35502163773405,54.35502163783405,54.36873772300807,54.36873772310807,54.396160711747456,54.39616071184746,54.40986666944711,54.409866669547114,54.42356395762786,54.423563957727865,54.43725643816631,54.43725643826631,54.450947933532156,54.45094793363216,54.46463611871805,54.46463611881805,54.47832237296564,54.478322373065645,54.49200630220589,54.49200630230589,54.505685660245256,54.50568566034526,54.51936411192583,54.51936411202583,54.533040238599064,54.533040238699066,54.54670655295316,54.54670655305316,54.56036206981553,54.560362069915534,54.57401439471867,54.57401439481867,54.587662306048536,54.58766230614854,54.60130422752897,54.60130422762897,54.6149403167876,54.614940316887605,54.64221127369083,54.642211273790835,54.65583932394102,54.65583932404102,54.6694664284255,54.6694664285255,54.683067130284215,54.68306713038422,54.69666483721822,54.696664837318224,54.71025686955802,54.710256869658025,54.723844764172895,54.7238447642729,54.7374310431047,54.7374310432047,54.75101026820066,54.75101026830066,54.76458732591689,54.764587326016894,54.77816367430885,54.77816367440885,54.79173379640996,54.79173379650996,54.81887029695628,54.81887029705628,54.832434980904615,54.83243498100462,54.845993714410426,54.84599371451043,54.859546142811574,54.859546142911576,54.87308328133392,54.87308328143392,54.88661852832487,54.88661852842487,54.913687682472016,54.91368768257202,54.92722009216587,54.92722009226587,54.940742019623215,54.94074201972322,54.95425275551979,54.95425275561979,54.981263666262635,54.981263666362636,54.99476356526057,54.99476356536057,55.008259996450946,55.00825999655095,55.02174708820504,55.02174708830504,55.035231657917265,55.03523165801727,55.048706651751786,55.04870665185179,55.06217857184779,55.06217857194779,55.0756496643988,55.075649664498805,55.102585110920224,55.102585111020225,55.11604741573162,55.11604741583162,55.12950857774279,55.12950857784279,55.15642258690836,55.156422587008365,55.16987094167569,55.16987094177569,55.18331504049738,55.18331504059738,55.19675712956695,55.19675712966695,55.2101981546501,55.2101981547501,55.223629761483174,55.223629761583176,55.23706034373674,55.23706034383674,55.25048430563041,55.25048430573041,55.26390487853032,55.26390487863032,55.27731800352535,55.27731800362535,55.29073014334778,55.29073014344778,55.304141100963086,55.30414110106309,55.31754997001247,55.31754997011247,55.33095414964026,55.330954149740265,55.34435107839769,55.344351078497695,55.35774438171995,55.35774438181995,55.37113654224198,55.37113654234198,55.38451944214154,55.38451944224154,55.39789576108811,55.39789576118811,55.411266602475,55.411266602575004,55.42463054765367,55.42463054775367,55.43799378350807,55.43799378360807,55.451355797748434,55.451355797848436,55.464712807311976,55.46471280741198,55.47804837951967,55.47804837961967,55.49138206019597,55.49138206029597,55.504714204003,55.504714204103,55.518044101616496,55.5180441017165,55.53135177373606,55.531351773836064,55.54465838186922,55.54465838196922,55.557963019657166,55.55796301975717,55.571266435831085,55.571266435931086,55.58456768462528,55.58456768472528,55.611164231771134,55.611164231871136,55.62444932373463,55.62444932383463,55.63772972627653,55.637729726376534,55.651003232610215,55.65100323271022,55.66427512326083,55.66427512336083,55.67753929015823,55.67753929025823,55.690793368888166,55.69079336898817,55.70403609842972,55.70403609852972,55.7172772516951,55.7172772517951,55.73049885913604,55.73049885923604,55.756939315534616,55.75693931563462,55.7701518593876,55.770151859487605,55.78335522143192,55.78335522153192,55.7965534605787,55.7965534606787,55.80972439074092,55.80972439084092,55.83606499004442,55.83606499014442,55.84922977272959,55.849229772829595,55.87555492452667,55.87555492462667,55.88870997370652,55.88870997380652,55.91501794409339,55.915017944193394,55.928170313603765,55.92817031370377,55.94131799369254,55.941317993792545,55.9544578712143,55.954457871314304,55.980734946588335,55.98073494668834,55.993862410935286,55.99386241103529,56.006984555350186,56.00698455545019,56.02010480823368,56.020104808333684,56.0332166674466,56.033216667546604,56.05943857315485,56.05943857325485,56.08565318858582,56.085653188685825,56.09875622065222,56.09875622075222,56.12495117203803,56.12495117213803,56.1380415544882,56.1380415545882,56.15111838096334,56.15111838106334,56.17726947246486,56.17726947256486,56.19034208240125,56.19034208250125,56.20340227916285,56.20340227926285,56.21645975684806,56.21645975694806,56.22951628876757,56.22951628886757,56.24257132322472,56.242571323324725,56.255623441570975,56.25562344167098,56.26866736328116,56.26866736338116,56.2817088417633,56.2817088418633,56.29474358166482,56.29474358176482,56.30777532664163,56.307775326741634,56.32080454957658,56.32080454967658,56.33382100467458,56.333821004774585,56.346833755523605,56.34683375562361,56.35984130466127,56.359841304761275,56.372847592778015,56.37284759287802,56.38585191054955,56.38585191064955,56.39885469145182,56.39885469155182,56.42484492397065,56.424844924070655,56.43783178448365,56.43783178458365,56.46380266821255,56.46380266831255,56.489756212903636,56.48975621300364,56.50272522208907,56.50272522218907,56.51569218211548,56.51569218221548,56.52865504382386,56.52865504392386,56.54161317670375,56.54161317680375,56.554567132451794,56.554567132551796,56.567520063620336,56.56752006372034,56.5804599116967,56.5804599117967,56.593392863564844,56.593392863664846,56.60631962854904,56.60631962864904,56.61924099478737,56.61924099488737,56.64507068357875,56.64507068367875,56.65797569595187,56.65797569605187,56.670873772709854,56.670873772809855,56.68376849988098,56.683768499980985,56.70955692964374,56.70955692974374,56.722448267821115,56.722448267921116,56.73533034537602,56.73533034547602,56.74819713305212,56.748197133152125,56.76104934017369,56.76104934027369,56.773899458729346,56.77389945882935,56.79959768608854,56.79959768618854,56.8124450855678,56.812445085667804,56.82528838672904,56.825288386829044,56.83811233910034,56.83811233920034,56.85092486346945,56.85092486356945,56.86372793018157,56.86372793028157,56.87652622865828,56.87652622875828,56.88931439956063,56.889314399660634,56.902100521303964,56.902100521403966,56.914884672702094,56.914884672802096,56.92766519866504,56.92766519876504,56.94044300555161,56.94044300565161,56.95321903912748,56.953219039227484,56.96599396931004,56.96599396941004,56.9787612151463,56.9787612152463,56.99152593894069,56.99152593904069,57.004278328374085,57.004278328474086,57.01701818641197,57.01701818651197,57.02975741393938,57.02975741403938,57.05522448039891,57.055224480498914,57.067947551095635,57.06794755119564,57.08066951839904,57.08066951849904,57.093389318322714,57.093389318422716,57.10610115805176,57.10610115815176,57.11881236727033,57.11881236737033,57.13151943876397,57.131519438863975,57.1442150034416,57.1442150035416,57.15690855836713,57.15690855846713,57.16959352258755,57.16959352268755,57.194960810765814,57.194960810865815,57.20763418935642,57.20763418945642,57.22030378488422,57.220303784984225,57.232972474053234,57.232972474153236,57.2456397445737,57.2456397446737,57.258291212925606,57.25829121302561,57.28359127292541,57.283591273025415,57.29623895821452,57.296238958314525,57.308884515530806,57.30888451563081,57.32152554105312,57.32152554115312,57.334145168626485,57.33414516872649,57.34675990974374,57.34675990984374,57.35937086779819,57.35937086789819,57.37197934321768,57.37197934331768,57.38457682411092,57.384576824210924,57.397172965169425,57.39717296526943,57.40975866339833,57.40975866349833,57.42234176077156,57.42234176087156,57.44749081351472,57.44749081361472,57.460044789185794,57.460044789285796,57.47258989830344,57.47258989840344,57.48512436755697,57.48512436765697,57.497656038920304,57.497656039020306,57.51018183865492,57.510181838754924,57.522702160829866,57.52270216092987,57.53522035503199,57.53522035513199,57.54773551490249,57.54773551500249,57.560246064165206,57.56024606426521,57.57275428842058,57.572754288520585,57.585254670702035,57.58525467080204,57.59775174280354,57.59775174290354,57.61024250980038,57.61024250990038,57.62273071534845,57.622730715448455,57.63521241875734,57.63521241885734,57.64769049673105,57.64769049683105,57.6601669984286,57.6601669985286,57.6726278161783,57.6726278162783,57.69754689022893,57.69754689032893,57.71000096939802,57.71000096949802,57.734905384080314,57.734905384180315,57.7473527640757,57.7473527641757,57.77224240116894,57.77224240126894,57.78468343665276,57.784683436752765,57.79711426574842,57.79711426584842,57.809542769836725,57.809542769936726,57.82196827900032,57.821968279100325,57.83437408471185,57.83437408481185,57.84675979290226,57.84675979300226,57.85914041760205,57.85914041770205,57.87151721983213,57.87151721993213,57.88388637712281,57.883886377222815,57.89625139668856,57.896251396788564,57.92097848030224,57.92097848040224,57.93333632781144,57.93333632791144,57.94568538758101,57.94568538768101,57.95803015199803,57.95803015209803,57.97037140920058,57.97037140930058,57.982709040967954,57.982709041067956,57.99504249560349,57.995042495703494,58.007357704842406,58.00735770494241,58.01965687547135,58.01965687557135,58.03195478507935,58.03195478517935,58.04424891162456,58.04424891172456,58.05653779705151,58.05653779715151,58.068824554505646,58.06882455460565,58.08111040560099,58.08111040570099,58.09339152786783,58.09339152796783,58.10567186199659,58.10567186209659,58.13023024465367,58.13023024475367,58.1424959982279,58.1424959983279,58.15476001789835,58.15476001799835,58.167022303665014,58.167022303765016,58.17928395892121,58.17928395902121,58.191544156121964,58.191544156221966,58.20380076729443,58.20380076739443,58.216050442851774,58.216050442951776,58.22829858153985,58.228298581639855,58.24054080919232,58.24054080929232,58.26501899879948,58.265018998899485,58.27724211410344,58.27724211420344,58.28945774209561,58.28945774219561,58.30167203025304,58.30167203035304,58.31388525442405,58.31388525452405,58.32609134594542,58.32609134604542,58.3382936938109,58.3382936939109,58.36268313906995,58.36268313916995,58.374864758903854,58.374864759003856,58.399219486680366,58.39921948678037,58.41138869333946,58.41138869343946,58.423548915224416,58.42354891532442,58.44786778271816,58.447867782818165,58.460025127899115,58.46002512799912,58.4721789658656,58.472178965965604,58.496480454914625,58.49648045501463,58.50862428352746,58.50862428362746,58.52076641764342,58.520766417743424,58.532902916572084,58.532902916672086,58.545037918038396,58.5450379181384,58.55717201314591,58.55717201324591,58.56930520189463,58.56930520199463,58.58143255842153,58.581432558521534,58.59355904799655,58.59355904809655,58.60568482824729,58.60568482834729,58.62993043830625,58.629930438406255,58.642048455396875,58.64204845549688,58.654163398748985,58.654163398848986,58.66626884503719,58.66626884513719,58.67837318793208,58.67837318803208,58.69047429946083,58.69047429956083,58.71467096614486,58.71467096624486,58.75095782864519,58.75095782874519,58.7630499553998,58.7630499554998,58.77514113638871,58.77514113648871,58.7872226232792,58.7872226233792,58.81138358730808,58.81138358740808,58.83554068946035,58.835540689560354,58.847615595397855,58.84761559549786,58.859681831816445,58.85968183191645,58.87174121143372,58.871741211533724,58.88379081813877,58.88379081823877,58.89583329219417,58.89583329229417,58.90786528401307,58.90786528411307,58.91989227115514,58.919892271255144,58.94394273822482,58.94394273832482,58.95596574526958,58.95596574536958,58.967982920092524,58.967982920192526,58.97999934618429,58.979999346284295,59.004022819524636,59.00402281962464,59.04004021761449,59.04004021771449,59.05203493050208,59.05203493060208,59.0640290128792,59.064029012979205,59.08801465559158,59.08801465569158,59.099981074322,59.099981074422004,59.11194418287247,59.11194418297247,59.13586247918568,59.135862479285684,59.14781341100277,59.147813411102774,59.15976205721943,59.15976205731943,59.171705265283315,59.171705265383316,59.183642444090864,59.183642444190866,59.19557359364208,59.19557359374208,59.20750115716502,59.20750115726502,59.21941449479557,59.219414494895574,59.23132507394283,59.23132507404283,59.24322083609413,59.24322083619413,59.25511249992741,59.25511250002741,59.266997898062925,59.26699789816293,59.27886709996084,59.27886710006084,59.290735159058535,59.29073515915854,59.32633673549596,59.32633673559596,59.33820172085513,59.338201720955134,59.35006359306888,59.35006359316888,59.37378316036454,59.37378316046454,59.38563856984601,59.38563856994601,59.397475300454914,59.397475300554916,59.40931025775314,59.40931025785314,59.421144072251145,59.42114407235115,59.43297607403156,59.43297607413156,59.44480228299706,59.44480228309706,59.456622147451,59.456622147551,59.46843464281387,59.46843464291387,59.49205809667034,59.49205809677034,59.503864444556164,59.503864444656166,59.515669137352006,59.51566913745201,59.527470638188625,59.52747063828863,59.539270050459315,59.53927005055932,59.55106701950195,59.55106701960195,59.562851141893844,59.562851141993846,59.58641584005625,59.58641584015625,59.598194800143695,59.5981948002437,59.60997009538906,59.60997009548906,59.62174184401305,59.62174184411305,59.63351142525731,59.63351142535731,59.64527249461028,59.645272494710284,59.66878162903786,59.66878162913786,59.68052902419509,59.68052902429509,59.69227448841402,59.69227448851402,59.704016642453006,59.70401664255301,59.71575517105681,59.715755171156815,59.72749054710829,59.72749054720829,59.73921835703417,59.73921835713417,59.75094553644958,59.75094553654958,59.77438220158045,59.77438220168045,59.78608317540461,59.786083175504615,59.797782021255955,59.79778202135596,59.80947621709261,59.80947621719261,59.821168836653094,59.821168836753095,59.83285656975747,59.83285656985747,59.85622376051635,59.85622376061635,59.867899592735675,59.86789959283568,59.87957317876146,59.87957317886146,59.902919208012825,59.90291920811283,59.91458893216201,59.91458893226201,59.926256488931465,59.926256489031466,59.937921405438345,59.93792140553835,59.96124717954098,59.961247179640985,59.97290425407394,59.97290425417394,59.99621343786993,59.996213437969935,60.00786097593209,60.00786097603209,60.01950449449002,60.019504494590024,60.031143599474696,60.0311435995747,60.04277781800325,60.042777818103254,60.054404667440735,60.05440466754074,60.06602789144304,60.066027891543044,60.0776468594997,60.0776468595997,60.08926488179066,60.089264881890664,60.100876283721725,60.10087628382173,60.11248532123854,60.11248532133854,60.12409250663086,60.12409250673086,60.13569259878044,60.13569259888044,60.14729186338503,60.14729186348503,60.15888931527203,60.15888931537203,60.17048259002719,60.17048259012719,60.182061757110674,60.182061757210676,60.193632648744284,60.193632648844286,60.205198890363214,60.205198890463215,60.216764067995726,60.21676406809573,60.22831403456324,60.22831403466324,60.23986321299268,60.23986321309268,60.251411760911644,60.251411761011646,60.262956880429954,60.262956880529956,60.27449813807166,60.27449813817166,60.28603801647172,60.28603801657172,60.29757304782257,60.29757304792257,60.30910382322777,60.309103823327774,60.32063341642585,60.32063341652585,60.33215776850568,60.33215776860568,60.34367711590868,60.343677116008685,60.35519441415268,60.35519441425268,60.37822692207473,60.37822692217473,60.38974055547664,60.389740555576644,60.40124985411909,60.40124985421909,60.4127550150366,60.412755015136604,60.435763169491906,60.43576316959191,60.458768486650115,60.45876848675012,60.470268564076996,60.470268564177,60.481765725392975,60.48176572549298,60.50474846239511,60.504748462495115,60.516233210536285,60.51623321063629,60.527715081973454,60.527715082073456,60.53919108178191,60.53919108188191,60.55066097352022,60.550660973620225,60.56212459956078,60.562124599660784,60.573585112455916,60.57358511255592,60.585037822784024,60.585037822884026,60.59648915387049,60.596489153970495,60.61938566856638,60.61938566866638,60.630827029706104,60.630827029806106,60.64225712047124,60.642257120571244,60.65368634428449,60.65368634438449,60.67654266393816,60.67654266403816,60.68796924748883,60.68796924758883,60.699388934831276,60.69938893493128,60.71080594250424,60.71080594260424,60.722220940425096,60.7222209405251,60.73363345571099,60.73363345581099,60.745044788789755,60.74504478888976,60.756440753175916,60.75644075327592,60.76783455018234,60.767834550282345,60.77922771667831,60.77922771677831,60.7906144992558,60.790614499355804,60.802000060219264,60.802000060319266,60.813383059733965,60.81338305983397,60.82476408890345,60.82476408900345,60.847517674758045,60.84751767485805,60.87024801053919,60.870248010639195,60.88159412519162,60.88159412529162,60.91562439073355,60.91562439083355,60.92696065365994,60.92696065375994,60.93829226657164,60.938292266671645,60.94961800785463,60.94961800795463,60.96094138472338,60.96094138482338,60.97225680139748,60.97225680149748,60.98357099645755,60.98357099655755,60.994880383875326,60.99488038397533,61.01749482395142,61.01749482405142,61.028797709230005,61.02879770933001,61.04009657500437,61.04009657510437,61.05139347043353,61.05139347053353,61.06268733153107,61.062687331631075,61.07398020745602,61.07398020755602,61.085271743546215,61.08527174364622,61.11914359333352,61.11914359343352,61.130414795461185,61.13041479556119,61.14167969248419,61.14167969258419,61.15294261916198,61.152942619261985,61.17546082757818,61.17546082767818,61.18671528177158,61.186715281871585,61.19795582532783,61.19795582542783,61.20919002437251,61.20919002447251,61.220421425527,61.220421425627,61.2316452999628,61.2316453000628,61.24286736168101,61.24286736178101,61.25407062630594,61.25407062640594,61.26527180236496,61.26527180246496,61.276471638589236,61.27647163868924,61.28766899217855,61.287668992278554,61.2988647694917,61.2988647695917,61.31005128618238,61.31005128628238,61.32122255240115,61.321222552501155,61.33238849868788,61.33238849878788,61.34354995258752,61.343549952687525,61.365869944275914,61.365869944375916,61.39933767626131,61.39933767636131,61.43277971494522,61.43277971504522,61.44391539672956,61.44391539682956,61.45505029037582,61.455050290475825,61.46618415944258,61.46618415954258,61.477313220867025,61.47731322096703,61.48843983906342,61.48843983916342,61.499564802169836,61.49956480226984,61.51068878010365,61.51068878020365,61.52181157583034,61.52181157593034,61.53292980035616,61.53292980045616,61.544047276150785,61.54404727625079,61.5551626239726,61.5551626240726,61.5662721395726,61.5662721396726,61.57737921194454,61.57737921204454,61.58848395930914,61.58848395940914,61.59958602700425,61.59958602710425,61.61068659723701,61.610686597337015,61.621781650503195,61.6217816506032,61.632871974940876,61.63287197504088,61.643957570550064,61.643957570650066,61.655034457233434,61.655034457333436,61.66610870365423,61.66610870375423,61.677182319564565,61.67718231966457,61.688254438012535,61.68825443811254,61.699319502624675,61.69931950272468,61.71038366087801,61.71038366097801,61.721443760220225,61.72144376032023,61.732500707010104,61.732500707110106,61.74355387073719,61.74355387083719,61.75460604929167,61.75460604939167,61.765656493942366,61.76565649404237,61.78775166924267,61.78775166934267,61.798795257092046,61.79879525719205,61.80983293390581,61.80983293400581,61.83190576549146,61.831905765591465,61.84293575795891,61.842935758058914,61.853964765253764,61.853964765353766,61.87602053364993,61.87602053374993,61.89807019397595,61.89807019407595,61.9090782367978,61.9090782368978,61.9200819842671,61.9200819843671,61.93107800798319,61.93107800808319,61.94207166728503,61.94207166738503,61.953063513869274,61.953063513969276,61.96405205027358,61.96405205037358,61.975039246843146,61.97503924694315,61.99701021158161,61.99701021168161,62.00799252169506,62.00799252179506,62.01896868433146,62.018968684431464,62.02994307365718,62.02994307375718,62.040916674844816,62.04091667494482,62.05188964552198,62.05188964562198,62.0738325131378,62.073832513237804,62.084799966848394,62.084799966948395,62.095762573509774,62.095762573609775,62.10672230346715,62.10672230356715,62.117678526210064,62.117678526310065,62.12862970486925,62.12862970496925,62.16147429547956,62.16147429557956,62.172415898261036,62.17241589836104,62.18335639764919,62.18335639774919,62.19429110422245,62.19429110432245,62.20522320994003,62.20522321004003,62.21615373938145,62.21615373948145,62.227083480684776,62.22708348078478,62.23801058172553,62.238010581825534,62.248934017924206,62.24893401802421,62.25985012443871,62.25985012453871,62.27076075339354,62.27076075349354,62.28167035776885,62.281670357868855,62.29257866171634,62.29257866181634,62.303485074132425,62.30348507423243,62.32529553455034,62.325295534650344,62.33619000667448,62.33619000677448,62.34707289316879,62.347072893268795,62.36883614411556,62.368836144215564,62.37971438059519,62.37971438069519,62.39058619374944,62.390586193849444,62.40144586957722,62.401445869677225,62.423161044100944,62.423161044200945,62.434016503389984,62.434016503489985,62.444869046568115,62.44486904666812,62.466569128247556,62.46656912834756,62.47741201673418,62.47741201683418,62.48825258021345,62.48825258031345,62.49908951825754,62.49908951835754,62.50992381603906,62.509923816139064,62.520755591778716,62.52075559187872,62.53158311157272,62.531583111672724,62.54240779406963,62.542407794169634,62.55322089093672,62.55322089103672,62.574833252847554,62.574833252947556,62.58563015347705,62.58563015357705,62.60722308778414,62.607223087884144,62.62881239665606,62.62881239675606,62.63960003666308,62.639600036763085,62.65038326309684,62.650383263196844,62.66116562257871,62.66116562267871,62.67194341085969,62.67194341095969,62.68271654912599,62.682716549225994,62.6934877958609,62.6934877959609,62.704248836207626,62.70424883630763,62.715007512140104,62.715007512240106,62.72576197153384,62.725761971633844,62.73651552456879,62.73651552466879,62.747265333947844,62.747265334047846,62.75801246365741,62.75801246375741,62.76874911113048,62.768749111230484,62.779482369609795,62.7794823697098,62.790209283577546,62.79020928367755,62.8009229568255,62.800922956925504,62.82234041218849,62.82234041228849,62.83304407608279,62.833044076182794,62.843741789534576,62.84374178963458,62.85443087287436,62.85443087297436,62.86511763120679,62.865117631306795,62.88648996566454,62.88648996576454,62.89716695108475,62.89716695118475,62.90784007462835,62.907840074728355,62.92917946491424,62.92917946501424,62.93984514055297,62.93984514065297,62.95050935813624,62.95050935823624,62.961166994766515,62.96116699486652,62.97182151825137,62.97182151835137,62.98247407139101,62.98247407149101,62.993123905454276,62.99312390555428,63.003754036065466,63.00375403616547,63.02500763752105,63.02500763762105,63.03562559139887,63.03562559149887,63.046239643993175,63.04623964409318,63.05684778555186,63.05684778565186,63.067454193206764,63.067454193306766,63.07805559618484,63.07805559628484,63.099254500857484,63.099254500957485,63.109846012702626,63.10984601280263,63.12042869740124,63.12042869750124,63.131003067243086,63.13100306734309,63.14157605784328,63.14157605794328,63.15214266452501,63.15214266462501,63.16270540933013,63.16270540943013,63.17326417403793,63.173264174137934,63.18379094033958,63.18379094043958,63.19431668206172,63.19431668216172,63.204839586486756,63.20483958658676,63.21536099344944,63.21536099354944,63.225871169444446,63.22587116954445,63.23637677423857,63.236376774338574,63.24687721672826,63.24687721682826,63.25737454607251,63.257374546172514,63.26787006269918,63.26787006279918,63.2783612839733,63.2783612840733,63.28884820989486,63.28884820999486,63.29932548112491,63.29932548122491,63.30979790530576,63.30979790540576,63.32026311802314,63.320263118123144,63.33072529640891,63.33072529650891,63.34118444046306,63.34118444056306,63.35164149595129,63.35164149605129,63.37255308488589,63.37255308498589,63.383006475532035,63.38300647563204,63.39345805346059,63.39345805356059,63.41435569235112,63.414355692451124,63.42479154692493,63.42479154702493,63.43521703748295,63.43521703758295,63.445637996246994,63.445637996346996,63.45605540838966,63.45605540848966,63.466469195097154,63.466469195197156,63.476876873734504,63.476876873834506,63.48728289728188,63.48728289738188,63.497687856842845,63.49768785694285,63.508087772320074,63.508087772420076,63.51848236786525,63.518482367965255,63.52887081593338,63.52887081603338,63.53925284067614,63.53925284077614,63.5496339590601,63.549633959160104,63.560004004104,63.560004004204,63.570370817781765,63.570370817881766,63.58073061703059,63.58073061713059,63.59108832771349,63.591088327813495,63.60144162482314,63.60144162492314,63.611793621504944,63.611793621604946,63.622142229192995,63.622142229293,63.632489812301536,63.63248981240154,63.642834479299175,63.64283447939918,63.65317626959281,63.653176269692814,63.66351463148579,63.66351463158579,63.67384499377722,63.673844993877225,63.684172873433695,63.6841728735337,63.7048272140981,63.704827214198104,63.71515209882986,63.71515209892986,63.72546744709082,63.72546744719082,63.73577794830258,63.73577794840258,63.746086360948404,63.746086361048405,63.75639087231073,63.75639087241073,63.76669116713431,63.76669116723431,63.77698858525389,63.77698858535389,63.787285097014674,63.787285097114676,63.80787638663246,63.80787638673246,63.82846172580772,63.82846172590772,63.83874870109771,63.83874870119771,63.84902413016477,63.84902413026477,63.859297904141876,63.85929790424188,63.86956198402056,63.86956198412056,63.87982417236783,63.879824172467835,63.89008474503205,63.89008474513205,63.90034062827467,63.90034062837467,63.92084884813854,63.92084884823854,63.93109933263529,63.93109933273529,63.94133559123965,63.94133559133965,63.951567515084555,63.95156751518456,63.97202336317283,63.97202336327283,63.982241691635814,63.982241691735815,63.99245844382263,63.99245844392263,64.00266979726358,64.00266979736358,64.01287973205596,64.01287973215597,64.02308856345505,64.02308856355505,64.03329640968154,64.03329640978154,64.04350338895613,64.04350338905613,64.06391269749061,64.06391269759061,64.07411427801934,64.07411427811934,64.08431057802291,64.08431057812291,64.09450013944586,64.09450013954586,64.10468556314389,64.10468556324389,64.1148679919172,64.1148679920172,64.12504943551791,64.12504943561791,64.13522433757254,64.13522433767254,64.14539805742002,64.14539805752003,64.15557043743279,64.15557043753279,64.17589718850311,64.17589718860312,64.18604820997383,64.18604821007384,64.19619793101671,64.19619793111671,64.20634694273532,64.20634694283532,64.21649331419134,64.21649331429134,64.2266370847917,64.2266370848917,64.23677912148827,64.23677912158827,64.24691528655613,64.24691528665613,64.257050624079,64.257050624179,64.26718446413952,64.26718446423952,64.27731436350962,64.27731436360962,64.28744111032739,64.2874411104274,64.29756395586165,64.29756395596165,64.3178061791226,64.3178061792226,64.32792441404908,64.32792441414908,64.33803527988448,64.33803527998448,64.3481443724092,64.3481443725092,64.35825129755419,64.35825129765419,64.36835711930586,64.36835711940586,64.38856655602258,64.38856655612258,64.39866784598027,64.39866784608027,64.40875743208744,64.40875743218744,64.41884638768414,64.41884638778414,64.43901988530429,64.43901988540429,64.44910202350658,64.44910202360659,64.45918148203938,64.45918148213939,64.46925499012967,64.46925499022967,64.49947405634506,64.49947405644507,64.50954114110998,64.50954114120998,64.51960432459137,64.51960432469137,64.52966569535518,64.52966569545518,64.53972190381455,64.53972190391455,64.54977543260443,64.54977543270444,64.55982703045602,64.55982703055602,64.56987626389335,64.56987626399335,64.57992305410264,64.57992305420264,64.58996661294577,64.58996661304577,64.6100484895138,64.6100484896138,64.63012331224598,64.63012331234599,64.64015701936309,64.64015701946309,64.65018635231384,64.65018635241384,64.6602119022018,64.6602119023018,64.68026095281867,64.68026095291867,64.69028354718881,64.69028354728881,64.70029309787368,64.70029309797368,64.71030122991002,64.71030123001002,64.72030857380825,64.72030857390826,64.73031410498889,64.7303141050889,64.74031770523125,64.74031770533125,64.75031992623195,64.75031992633195,64.77031408303137,64.77031408313137,64.78030353619513,64.78030353629514,64.79029153130344,64.79029153140344,64.80027806835629,64.8002780684563,64.8102507341789,64.8102507342789,64.82022079914583,64.82022079924583,64.83018987894015,64.83018987904015,64.84015222015387,64.84015222025387,64.8501109753393,64.8501109754393,64.86006578983434,64.86006578993434,64.87001859457725,64.87001859467725,64.87996954719567,64.87996954729567,64.88991427352325,64.88991427362325,64.90980041599843,64.90980041609843,64.91973493593783,64.91973493603783,64.92966752493892,64.92966752503892,64.93959723723601,64.93959723733602,64.94952470333958,64.94952470343958,64.95943408167413,64.95943408177413,64.96934093796683,64.96934093806684,64.97924688790073,64.97924688800073,64.98914948824778,64.98914948834778,64.99904787205608,64.99904787215608,65.00894428551918,65.00894428561918,65.01883754642995,65.01883754652995,65.02872745775386,65.02872745785386,65.03861606864994,65.03861606874995,65.04850314267676,65.04850314277677,65.06827559623352,65.06827559633352,65.07816085754276,65.07816085764276,65.08802578488945,65.08802578498945,65.09788586518694,65.09788586528694,65.10774027089022,65.10774027099022,65.11759377023472,65.11759377033472,65.12743659030819,65.1274365904082,65.14712029951684,65.14712029961684,65.15695886364468,65.15695886374468,65.17662968679566,65.17662968689567,65.18646021191505,65.18646021201505,65.19628872728232,65.19628872738232,65.20611357780751,65.20611357790752,65.2159366156151,65.21593661571511,65.23558095732653,65.23558095742654,65.255221988858,65.255221988958,65.27485943436119,65.27485943446119,65.28467037424922,65.28467037434922,65.29447926497824,65.29447926507824,65.30428657943108,65.30428657953108,65.31409290871133,65.31409290881133,65.32389746468088,65.32389746478088,65.33370048378119,65.33370048388119,65.35330537918156,65.35330537928157,65.36310445759145,65.36310445769145,65.37290125040089,65.37290125050089,65.38269138344354,65.38269138354354,65.39247552663676,65.39247552673676,65.40225379820126,65.40225379830126,65.41202994179294,65.41202994189294,65.43158021922417,65.43158021932418,65.44135384077398,65.44135384087399,65.4511263983374,65.4511263984374,65.46089824657652,65.46089824667652,65.47065338628829,65.47065338638829,65.48040529463393,65.48040529473393,65.49015629662077,65.49015629672077,65.50965723660804,65.50965723670804,65.51940540129777,65.51940540139778,65.5291524231873,65.5291524232873,65.53889684422116,65.53889684432116,65.54863444786058,65.54863444796058,65.55837122395504,65.55837122405504,65.568107369539,65.56810736963901,65.57783854985307,65.57783854995307,65.60702834713936,65.60702834723936,65.61674861174097,65.61674861184098,65.62646367463124,65.62646367473124,65.6361776735351,65.6361776736351,65.64589013556969,65.64589013566969,65.66531029140349,65.66531029150349,65.67501688180936,65.67501688190936,65.68472177771837,65.68472177781837,65.69442576726857,65.69442576736857,65.70412652545264,65.70412652555264,65.71382582558125,65.71382582568125,65.7332179236993,65.7332179237993,65.74290646574309,65.74290646584309,65.75259303744166,65.75259303754166,65.76224934463787,65.76224934473787,65.77190245987484,65.77190245997484,65.79120738992093,65.79120739002093,65.81051125598061,65.81051125608062,65.82016007586503,65.82016007596503,65.82980393047953,65.82980393057953,65.83944703636286,65.83944703646286,65.8490894723288,65.8490894724288,65.85873100193595,65.85873100203595,65.86837040357028,65.86837040367028,65.87800759841798,65.87800759851798,65.88764238944452,65.88764238954452,65.89726910205572,65.89726910215572,65.90689211041793,65.90689211051793,65.91651429123515,65.91651429133515,65.9261320190722,65.9261320191722,65.93574765834333,65.93574765844333,65.94535738657885,65.94535738667885,65.95496002157162,65.95496002167162,65.96456123791585,65.96456123801585,65.9741580800937,65.9741580801937,65.99334869071092,65.99334869081092,66.01252330212503,66.01252330222503,66.022101189582,66.022101189682,66.03167249608599,66.03167249618599,66.04124305385879,66.0412430539588,66.05081298112114,66.05081298122114,66.06037691853405,66.06037691863405,66.06993589067703,66.06993589077703,66.0794880060187,66.07948800611871,66.0890356683802,66.08903566848021,66.0985814786172,66.09858147871721,66.10812579139186,66.10812579149186,66.12720657496725,66.12720657506725,66.13674178474703,66.13674178484703,66.14627510299543,66.14627510309543,66.16533760176729,66.1653376018673,66.17486500898006,66.17486500908007,66.18438993355788,66.18438993365788,66.19391044456243,66.19391044466244,66.20342800004917,66.20342800014917,66.21293909280364,66.21293909290364,66.22244793936457,66.22244793946457,66.24146385917574,66.24146385927574,66.26047792686242,66.26047792696242,66.26998330561578,66.26998330571578,66.27948214282306,66.27948214292306,66.28897566009829,66.28897566019829,66.29846685236616,66.29846685246616,66.30795619250954,66.30795619260954,66.31744253772821,66.31744253782821,66.32692565158074,66.32692565168074,66.33639083529188,66.33639083539188,66.3458509355124,66.3458509356124,66.35530744970464,66.35530744980464,66.3647623482138,66.3647623483138,66.37421401535683,66.37421401545683,66.38366402740988,66.38366402750988,66.3931069856271,66.3931069857271,66.40254852519577,66.40254852529577,66.41198943425397,66.41198943435397,66.43086699642471,66.43086699652471,66.44030163978516,66.44030163988516,66.46859887069279,66.46859887079279,66.47802649962429,66.4780264997243,66.48744869040303,66.48744869050303,66.49686567947042,66.49686567957042,66.50627931895094,66.50627931905095,66.51569043638962,66.51569043648962,66.52509922882095,66.52509922892095,66.54390881408207,66.54390881418207,66.55330038569629,66.55330038579629,66.56268340601233,66.56268340611233,66.57206094876867,66.57206094886867,66.5814353389727,66.5814353390727,66.59080831052816,66.59080831062816,66.60018041513175,66.60018041523175,66.60954834260347,66.60954834270348,66.61891303870908,66.61891303880908,66.62827261191715,66.62827261201716,66.63763056944214,66.63763056954214,66.64698710831858,66.64698710841859,66.65634270142934,66.65634270152934,66.66569013731093,66.66569013741093,66.68438374805318,66.68438374815318,66.6937276767203,66.6937276768203,66.7030703443665,66.7030703444665,66.71240300265903,66.71240300275903,66.72172888296406,66.72172888306406,66.7403786732289,66.7403786733289,66.75902180372697,66.75902180382697,66.76833582255385,66.76833582265385,66.77764779222171,66.77764779232172,66.7962703129089,66.7962703130089,66.80557960290729,66.80557960300729,66.82418750363304,66.82418750373304,66.83348292240117,66.83348292250118,66.84277479454794,66.84277479464794,66.85206170142477,66.85206170152478,66.86134423413526,66.86134423423526,66.87062507234886,66.87062507244886,66.87989940842328,66.87989940852329,66.89844449454384,66.89844449464384,66.91698260564237,66.91698260574238,66.92624441032127,66.92624441042128,66.93550388999284,66.93550389009285,66.95402147009432,66.95402147019432,66.96327767899284,66.96327767909284,66.97253219339447,66.97253219349447,66.98178387049902,66.98178387059902,66.99103420776882,66.99103420786882,67.01877950557713,67.01877950567713,67.02801809958949,67.02801809968949,67.0372542897807,67.0372542898807,67.04648307147394,67.04648307157395,67.05571011926341,67.05571011936341,67.0649336992453,67.0649336993453,67.07415361438511,67.07415361448511,67.09258859761552,67.09258859771552,67.10180071018831,67.10180071028832,67.11100379858006,67.11100379868006,67.12020574417157,67.12020574427157,67.12940449780386,67.12940449790386,67.13860135990475,67.13860136000476,67.14779041943862,67.14779041953862,67.15697518361995,67.15697518371995,67.16615793804915,67.16615793814915,67.1753381704365,67.1753381705365,67.18451615663031,67.18451615673031,67.19368673432615,67.19368673442615,67.20285490820082,67.20285490830082,67.2120211905441,67.21202119064411,67.22118609364574,67.22118609374574,67.23034626791888,67.23034626801888,67.23949635402457,67.23949635412457,67.25779282198694,67.25779282208694,67.26693857333316,67.26693857343317,67.27608038398897,67.27608038408897,67.2852185692096,67.2852185693096,67.30349245701592,67.30349245711592,67.3126264256978,67.3126264257978,67.32175834522067,67.32175834532067,67.3400200168867,67.3400200169867,67.34914697113966,67.34914697123966,67.35827144275765,67.35827144285766,67.36739504742376,67.36739504752376,67.37651348978541,67.37651348988541,67.38563110460208,67.38563110470209,67.39474789187378,67.39474789197378,67.41297827445791,67.41297827455791,67.42209155451513,67.42209155461514,67.4494295425623,67.4494295426623,67.45854093108812,67.45854093118813,67.46765157088277,67.46765157098277,67.47675996448386,67.47675996458386,67.48586469324289,67.48586469334289,67.494967333436,67.494967333536,67.50406753040103,67.50406753050103,67.51316603286921,67.51316603296921,67.52226319550263,67.52226319560263,67.53135913652203,67.53135913662203,67.54045145210625,67.54045145220626,67.54953848716532,67.54953848726532,67.55861645863644,67.55861645873644,67.57677098293011,67.57677098303012,67.58584536837294,67.58584536847295,67.59491640422894,67.59491640432894,67.60398235659429,67.60398235669429,67.61304389538638,67.61304389548638,67.62210117823282,67.62210117833283,67.64021282781479,67.6402128279148,67.64926624878464,67.64926624888464,67.65831304939458,67.65831304949458,67.6673539389689,67.6673539390689,67.67639404040514,67.67639404050514,67.69447254878075,67.69447254888075,67.71254447620336,67.71254447630336,67.72156321909756,67.72156321919756,67.73057963698442,67.73057963708442,67.74860325154255,67.74860325164255,67.75760804439268,67.75760804449268,67.76661043342168,67.76661043352168,67.78461324113442,67.78461324123442,67.81161154166794,67.81161154176795,67.8206089260201,67.8206089261201,67.82960394595801,67.82960394605801,67.83859447350885,67.83859447360885,67.85657320360318,67.85657320370318,67.87453782602583,67.87453782612583,67.88351286666335,67.88351286676335,67.89248475474852,67.89248475484852,67.91042585124941,67.91042585134942,67.91939312872681,67.91939312882681,67.92835788416234,67.92835788426234,67.93732181205289,67.93732181215289,67.94628286323943,67.94628286333943,67.95524202289458,67.95524202299458,67.96419767533526,67.96419767543526,67.97314982056147,67.97314982066148,67.99105281058605,67.99105281068606,67.99999999054235,67.99999999064235,68.00894319040131,68.00894319050131,68.02682312738698,68.02682312748698,68.03575813060989,68.0357581307099,68.05362305356508,68.05362305366508,68.06255041184859,68.06255041194859,68.0714730413036,68.0714730414036,68.08039444914459,68.0803944492446,68.08930970950853,68.08930970960853,68.09821905883685,68.09821905893685,68.10712769884088,68.10712769894089,68.11602790576745,68.11602790586745,68.13382599461323,68.13382599471323,68.14271902948323,68.14271902958323,68.15160166093055,68.15160166103055,68.16048338601907,68.16048338611907,68.17824510229232,68.17824510239232,68.20488200210802,68.20488200220802,68.2226325268205,68.2226325269205,68.24037907143567,68.24037907153567,68.24924984140485,68.24924984150485,68.25811990204974,68.25811990214974,68.2669857461559,68.2669857462559,68.27584075336341,68.27584075346341,68.28469481480523,68.28469481490524,68.29354796988825,68.29354796998825,68.3023965143635,68.3023965144635,68.31123843847884,68.31123843857884,68.3200757913933,68.3200757914933,68.32890798200333,68.32890798210333,68.33773906922004,68.33773906932004,68.34656940770556,68.34656940780556,68.35539769703207,68.35539769713208,68.3642195630332,68.3642195631332,68.37303985275818,68.37303985285818,68.38185825095177,68.38185825105177,68.39949382572487,68.39949382582488,68.40830974128349,68.40830974138349,68.41712033691005,68.41712033701005,68.42592206598317,68.42592206608317,68.43472111538682,68.43472111548682,68.45231515528299,68.45231515538299,68.46110462880893,68.46110462890893,68.46989244724489,68.46989244734489,68.47867896525302,68.47867896535303,68.48746319766072,68.48746319776072,68.49624337115728,68.49624337125728,68.50502279592266,68.50502279602266,68.51379930457715,68.51379930467715,68.52256655260915,68.52256655270915,68.54009490119611,68.54009490129611,68.55761394975372,68.55761394985372,68.56636807528665,68.56636807538665,68.57512015166056,68.57512015176056,68.58386927251667,68.58386927261667,68.61011163040816,68.61011163050816,68.61885468260103,68.61885468270103,68.62758579450194,68.62758579460194,68.63631588182335,68.63631588192335,68.65377097297554,68.65377097307554,68.66248754372882,68.66248754382882,68.67119717886699,68.67119717896699,68.68861124743196,68.68861124753197,68.69731000626459,68.69731000636459,68.7060059278001,68.70600592790011,68.71468947556772,68.71468947566773,68.72337191994204,68.72337192004204,68.73204770454954,68.73204770464955,68.74072128237043,68.74072128247043,68.74939277162538,68.74939277172538,68.75806339392845,68.75806339402845,68.76673078486539,68.76673078496539,68.77539155544241,68.77539155554241,68.78405011923282,68.78405011933282,68.80136346375083,68.80136346385083,68.81001327920852,68.81001327930852,68.81865765651342,68.81865765661342,68.82729726558294,68.82729726568294,68.84457222777633,68.84457222787633,68.8532058075895,68.8532058076895,68.86183320051873,68.86183320061873,68.87045160867382,68.87045160877382,68.8790631994345,68.8790631995345,68.88766694822125,68.88766694832125,68.913477170002,68.913477170102,68.93067828365702,68.93067828375702,68.93927509682864,68.93927509692864,68.94786923033078,68.94786923043078,68.95646068416345,68.95646068426345,68.96504937951282,68.96504937961282,68.97363673502745,68.97363673512746,68.9822157362784,68.98221573637841,68.9907931612532,68.9907931613532,68.99936554214425,68.99936554224425,69.00793611031771,69.00793611041772,69.01650151618675,69.01650151628675,69.02505927711637,69.02505927721637,69.0336160922803,69.0336160923803,69.04217215871304,69.04217215881305,69.05072578191773,69.05072578201774,69.05927365171442,69.05927365181442,69.06781856599329,69.0678185660933,69.07636150992697,69.07636151002697,69.08490382335017,69.08490382345018,69.09344274777962,69.09344274787962,69.10197938660862,69.10197938670862,69.11904805365886,69.11904805375886,69.12757673229324,69.12757673239324,69.13610036684388,69.13610036694388,69.14461805095199,69.14461805105199,69.15313293716991,69.15313293726992,69.16164719287738,69.16164719297738,69.17016002993628,69.17016003003629,69.17867140893975,69.17867140903975,69.18718160573609,69.18718160583609,69.19568794065582,69.19568794075582,69.20419336921674,69.20419336931674,69.2126981672672,69.21269816736721,69.2211994186963,69.2211994187963,69.22969570485547,69.22969570495547,69.23819116346966,69.23819116356967,69.24667822841327,69.24667822851328,69.25515713612772,69.25515713622772,69.2721138087564,69.2721138088564,69.28058428339337,69.28058428349337,69.28905314234726,69.28905314244726,69.30598959923412,69.30598959933413,69.31445652724972,69.31445652734972,69.32292081500273,69.32292081510273,69.33138431461767,69.33138431471767,69.33984647439786,69.33984647449786,69.3482966938861,69.3482966939861,69.3567371010552,69.3567371011552,69.36517656245861,69.36517656255862,69.3820523327131,69.3820523328131,69.39047894746577,69.39047894756577,69.39890418297679,69.3989041830768,69.4073265023769,69.4073265024769,69.41574720609394,69.41574720619394,69.43258617029997,69.43258617039997,69.44100202696781,69.44100202706781,69.4494117361586,69.4494117362586,69.45781801694872,69.45781801704872,69.46622043586224,69.46622043596224,69.47462131790651,69.47462131800651,69.49141855020106,69.49141855030106,69.4998116296783,69.4998116297783,69.50820167482392,69.50820167492392,69.51659085301766,69.51659085311766,69.52497542060361,69.52497542070361,69.5417406150851,69.5417406151851,69.55850202650379,69.55850202660379,69.56688060424031,69.56688060434031,69.57525851205946,69.57525851215946,69.58363511945078,69.58363511955078,69.60038656092271,69.60038656102272,69.60876119796883,69.60876119806883,69.61713477102853,69.61713477112853,69.63386871583504,69.63386871593504,69.65058480931397,69.65058480941397,69.65892398014635,69.65892398024636,69.66726133826116,69.66726133836116,69.67559562263743,69.67559562273743,69.69225985663051,69.69225985673052,69.70058909692307,69.70058909702307,69.7089163668704,69.7089163669704,69.71724005078946,69.71724005088946,69.72556196139784,69.72556196149785,69.74220121141371,69.74220121151372,69.758834944463,69.758834944563,69.76714664868321,69.76714664878321,69.77545583086155,69.77545583096155,69.78376245159112,69.78376245169112,69.7920681659619,69.7920681660619,69.80867482646805,69.80867482656805,69.81697242301657,69.81697242311657,69.82526300513615,69.82526300523615,69.83354956775152,69.83354956785152,69.84183266255933,69.84183266265933,69.85011256540788,69.85011256550789,69.87494632351107,69.87494632361107,69.88321720217859,69.88321720227859,69.89148571643186,69.89148571653186,69.89975210271231,69.89975210281231,69.90801194744667,69.90801194754667,69.91626946717369,69.9162694672737,69.92452631698333,69.92452631708333,69.9327824180618,69.9327824181618,69.94103319920822,69.94103319930822,69.95752912631374,69.95752912641375,69.96577037098936,69.96577037108936,69.98225152050584,69.98225152060584,69.99048421388325,69.99048421398325,70.00694818198953,70.00694818208953,70.01517914146316,70.01517914156317,70.02340036743148,70.02340036753148,70.0316207658548,70.0316207659548,70.04804871605069,70.04804871615069,70.06446393781656,70.06446393791656,70.07267058323033,70.07267058333034,70.08907942107773,70.08907942117773,70.09727735756569,70.09727735766569,70.11366976273406,70.11366976283406,70.13005838483963,70.13005838493963,70.13825143487148,70.13825143497148,70.14644271159266,70.14644271169266,70.16282238833098,70.16282238843098,70.1791980455651,70.1791980456651,70.18738151971924,70.18738151981924,70.19556160487963,70.19556160497963,70.21191688874431,70.21191688884431,70.22009110227599,70.22009110237599,70.23643767721484,70.23643767731484,70.24460873819419,70.24460873829419,70.2527784593388,70.2527784594388,70.2609473135315,70.2609473136315,70.26911376390306,70.26911376400307,70.27727911088131,70.27727911098131,70.28544252692126,70.28544252702126,70.29360322388482,70.29360322398482,70.3017584432887,70.3017584433887,70.30990530842891,70.30990530852891,70.31804882398227,70.31804882408227,70.33433483050946,70.33433483060946,70.35061819677409,70.3506181968741,70.35875879621653,70.35875879631654,70.36689833167257,70.36689833177257,70.37503723661816,70.37503723671816,70.3831711368869,70.3831711369869,70.39943743996203,70.39943744006203,70.40756759657488,70.40756759667488,70.41569637394609,70.41569637404609,70.43195270707449,70.43195270717449,70.44007959291429,70.44007959301429,70.45633245823511,70.45633245833511,70.46445327541169,70.46445327551169,70.47257007308404,70.47257007318404,70.48879925485548,70.48879925495548,70.49689930459357,70.49689930469357,70.51308600572236,70.51308600582236,70.52925930850373,70.52925930860373,70.5373438319216,70.5373438320216,70.54542461168357,70.54542461178357,70.56158364916564,70.56158364926564,70.56966127637529,70.56966127647529,70.57773795781922,70.57773795791923,70.58581168374536,70.58581168384536,70.59388245415369,70.59388245425369,70.60194861395422,70.60194861405422,70.61001193645767,70.61001193655767,70.61807431319541,70.61807431329541,70.62612707464854,70.62612707474854,70.63417853567383,70.63417853577383,70.64222834160915,70.64222834170916,70.6502756255026,70.6502756256026,70.65832058438872,70.65832058448872,70.66636392759176,70.66636392769176,70.67440573392554,70.67440573402554,70.6904882037929,70.6904882038929,70.69852595121554,70.69852595131555,70.70656078252729,70.70656078262729,70.71459498332858,70.71459498342858,70.73065897135788,70.73065897145788,70.74671326528878,70.74671326538878,70.75473383130122,70.75473383140123,70.77077346586377,70.77077346596377,70.77879194331031,70.77879194341031,70.78680261818982,70.78680261828983,70.80282270692793,70.80282270702793,70.8188397219275,70.8188397220275,70.82684515568876,70.82684515578876,70.83484905258076,70.83484905268077,70.84284262486389,70.8428426249639,70.85882839018848,70.85882839028848,70.87481253983002,70.87481253993002,70.88279957056704,70.88279957066705,70.89876886380571,70.89876886390572,70.90674907714833,70.90674907724834,70.91472219724821,70.91472219734821,70.9226916919129,70.9226916920129,70.93065752173551,70.93065752183551,70.94658827502194,70.94658827512194,70.95455095229221,70.95455095239221,70.96251221091394,70.96251221101394,70.97046941062455,70.97046941072455,70.97842538872112,70.97842538882112,70.98638061808651,70.98638061818652,70.9943338376998,70.9943338377998,71.00228374713315,71.00228374723315,71.01023129215223,71.01023129225223,71.01817548758447,71.01817548768447,71.0261186584372,71.0261186585372,71.03405690342692,71.03405690352692,71.04198963145005,71.04198963155005,71.04992082260392,71.04992082270392,71.0578492158676,71.05784921596761,71.06577611166894,71.06577611176894,71.08162190367004,71.08162190377004,71.10537666133109,71.1053766614311,71.11329047404024,71.11329047414024,71.12120326216989,71.12120326226989,71.12911210960911,71.12911210970911,71.13701981424812,71.13701981434812,71.1449262972731,71.1449262973731,71.16073780526762,71.16073780536762,71.16863270266278,71.16863270276278,71.17652594496798,71.17652594506798,71.18441804447295,71.18441804457295,71.20020074602054,71.20020074612054,71.20808949593865,71.20808949603865,71.22386195169116,71.22386195179116,71.2317431354837,71.2317431355837,71.2396222307103,71.2396222308103,71.24749967084693,71.24749967094694,71.25537147579628,71.25537147589628,71.26324253201444,71.26324253211445,71.28685294218565,71.28685294228565,71.29472120051362,71.29472120061362,71.30258646391687,71.30258646401687,71.31044691967783,71.31044691977783,71.3183038288174,71.31830382891741,71.33401575556516,71.33401575566516,71.34186635960009,71.34186635970009,71.34971625431072,71.34971625441072,71.35756425748997,71.35756425758997,71.36541052676543,71.36541052686543,71.37325364348857,71.37325364358857,71.38109112502441,71.38109112512441,71.38892695147028,71.38892695157028,71.3967604923157,71.39676049241571,71.40459328442995,71.40459328452995,71.42024814998052,71.42024815008052,71.43589111464605,71.43589111474606,71.44371078426121,71.44371078436122,71.45152915344856,71.45152915354856,71.45934657687019,71.45934657697019,71.4671628574916,71.4671628575916,71.47497803471971,71.47497803481971,71.49060539425119,71.49060539435119,71.49841627612675,71.49841627622675,71.50622566053994,71.50622566063994,71.514033862746,71.514033862846,71.52183816366856,71.52183816376856,71.52964112475638,71.52964112485638,71.53743880531904,71.53743880541904,71.54523002314943,71.54523002324943,71.5608115130445,71.5608115131445,71.56859812026711,71.56859812036711,71.57638315121355,71.57638315131355,71.58416621181478,71.58416621191478,71.59194714444318,71.59194714454318,71.61528210035448,71.61528210045448,71.62305113209784,71.62305113219784,71.63858513667343,71.63858513677343,71.65410440306687,71.65410440316687,71.6618579873038,71.6618579874038,71.67736409179128,71.67736409189128,71.68510644506053,71.68510644516053,71.70832739679817,71.70832739689817,71.71606242038325,71.71606242048325,71.72379563125075,71.72379563135075,71.73152273404811,71.73152273414811,71.73924841819692,71.73924841829692,71.74697201377981,71.74697201387981,71.75469222036894,71.75469222046894,71.762408762116,71.762408762216,71.77783747144373,71.77783747154373,71.78554889029324,71.78554889039324,71.7932551074314,71.79325510753141,71.80866210355498,71.80866210365498,71.81636114863659,71.81636114873659,71.82405893269727,71.82405893279727,71.83175573158535,71.83175573168535,71.84714574333323,71.84714574343323,71.85483773457901,71.85483773467901,71.8702183674837,71.8702183675837,71.87790641803906,71.87790641813906,71.885591355449,71.885591355549,71.89327483480346,71.89327483490347,71.90095768364749,71.90095768374749,71.91631908598296,71.91631908608296,71.9239965360811,71.92399653618111,71.93167110947526,71.93167110957526,71.93934418540704,71.93934418550704,71.94701276895177,71.94701276905177,71.95468072198601,71.95468072208601,71.96234520721269,71.9623452073127,71.97000311148639,71.9700031115864,71.97765829668371,71.97765829678372,71.98531095983915,71.98531095993916,71.99296125858035,71.99296125868035,72.00060938994183,72.00060939004183,72.00825551155118,72.00825551165119,72.02354531154185,72.02354531164185,72.0311867043227,72.03118670442271,72.03882659964121,72.03882659974121,72.04646377588334,72.04646377598334,72.05409957288381,72.05409957298382,72.0693695512017,72.0693695513017,72.07700140751176,72.07700140761176,72.09226362266953,72.09226362276954,72.0998935874482,72.0998935875482,72.11514989157034,72.11514989167034,72.12277382709267,72.12277382719267,72.13039543760766,72.13039543770766,72.13801570828791,72.13801570838791,72.14563337811249,72.1456333782125,72.1608645800367,72.1608645801367,72.16847783628802,72.16847783638802,72.17608963448387,72.17608963458387,72.1913104723923,72.1913104724923,72.19891734472515,72.19891734482515,72.21413022243894,72.21413022253894,72.22173543968181,72.22173543978181,72.22933781962757,72.22933781972758,72.23693590422079,72.23693590432079,72.24452992990288,72.24452993000288,72.25212261575024,72.25212261585024,72.2597144740526,72.2597144741526,72.26729951496056,72.26729951506056,72.27488112746785,72.27488112756785,72.28246203065086,72.28246203075086,72.29004175162676,72.29004175172676,72.29762060565075,72.29762060575075,72.30519678000528,72.30519678010528,72.31276830434511,72.31276830444511,72.32790985556241,72.32790985566241,72.33547877904657,72.33547877914657,72.34304273726082,72.34304273736082,72.35816805283362,72.35816805293362,72.36572740044006,72.36572740054007,72.38839527627816,72.38839527637816,72.39594134375793,72.39594134385793,72.40348548029938,72.40348548039938,72.41102354817761,72.41102354827761,72.41855921223467,72.41855921233467,72.42609365467771,72.42609365477772,72.4336265602515,72.4336265603515,72.44115855946649,72.44115855956649,72.44868969172957,72.44868969182957,72.45621881424056,72.45621881434056,72.47127473425518,72.47127473435518,72.47880090124835,72.47880090134835,72.48632332458563,72.48632332468563,72.5013673437152,72.5013673438152,72.50888685094158,72.50888685104158,72.52392298869032,72.52392298879032,72.53894758021616,72.53894758031616,72.54644458610026,72.54644458620027,72.55393977926678,72.55393977936679,72.5614330809019,72.56143308100191,72.57641479771605,72.57641479781606,72.58390285823292,72.58390285833292,72.59138883018387,72.59138883028388,72.59887350170699,72.59887350180699,72.6138407167804,72.6138407168804,72.62131928023338,72.62131928033338,72.6287963856309,72.62879638573091,72.63627092957967,72.63627092967967,72.65869219701169,72.65869219711169,72.66616153924909,72.66616153934909,72.68109277581902,72.68109277591903,72.68854950784711,72.68854950794712,72.69600450597142,72.69600450607142,72.70345710027458,72.70345710037458,72.71090697550135,72.71090697560135,72.71835436809315,72.71835436819315,72.72580101195378,72.72580101205378,72.73324611894513,72.73324611904513,72.7406901619501,72.7406901620501,72.74813010663702,72.74813010673702,72.75556894793063,72.75556894803063,72.7630065676102,72.7630065677102,72.77787790568586,72.77787790578586,72.78531091475766,72.78531091485766,72.79274238696019,72.79274238706019,72.80017271636251,72.80017271646251,72.81503101075289,72.81503101085289,72.82245456216769,72.8224545622677,72.82987373941614,72.82987373951615,72.84470519770481,72.84470519780481,72.86693581453461,72.86693581463462,72.87433970190426,72.87433970200426,72.88914546689143,72.88914546699144,72.89654423136354,72.89654423146354,72.90393834582095,72.90393834592095,72.91132970179507,72.91132970189507,72.91872023022421,72.91872023032421,72.93349872563373,72.93349872573373,72.94088062395086,72.94088062405086,72.94825578368739,72.94825578378739,72.9556298794375,72.9556298795375,72.96300334467715,72.96300334477715,72.9703758641511,72.9703758642511,72.97774688616269,72.9777468862627,72.9998555386242,72.9998555387242,73.00722234409704,73.00722234419705,73.01458745507303,73.01458745517303,73.02195114740044,73.02195114750045,73.02931381514836,73.02931381524836,73.04403781080946,73.04403781090946,73.05139827177074,73.05139827187074,73.05875747171109,73.05875747181109,73.07347248259804,73.07347248269804,73.08081761427361,73.08081761437361,73.08815872644496,73.08815872654496,73.09549613436732,73.09549613446733,73.10282818295073,73.10282818305073,73.11015834000273,73.11015834010273,73.11748467458503,73.11748467468503,73.1248087235669,73.1248087236669,73.13945426008185,73.13945426018185,73.14677440778021,73.14677440788022,73.16141202350744,73.16141202360744,73.16872760000491,73.16872760010492,73.17603805360484,73.17603805370484,73.19065356205886,73.19065356215886,73.19795676478843,73.19795676488843,73.20525480521356,73.20525480531356,73.21254835325162,73.21254835335162,73.21984032501352,73.21984032511352,73.23442245581973,73.23442245591973,73.24171076273953,73.24171076283953,73.25628568208228,73.25628568218228,73.2708585128591,73.27085851295911,73.27814350959898,73.27814350969898,73.28542736353862,73.28542736363862,73.29999140657583,73.29999140667583,73.30726887659699,73.307268876697,73.32182196451485,73.32182196461486,73.3363731214944,73.3363731215944,73.34364688726659,73.34364688736659,73.35091935261094,73.35091935271095,73.36546270702348,73.36546270712348,73.37272662106963,73.37272662116963,73.3872522029684,73.3872522030684,73.39451296446222,73.39451296456222,73.40177250434202,73.40177250444202,73.40902956158685,73.40902956168685,73.41628551543836,73.41628551553836,73.42354040530347,73.42354040540347,73.43804683544683,73.43804683554683,73.44529443503467,73.44529443513467,73.45253927613922,73.45253927623922,73.45978309266427,73.45978309276427,73.47426946469342,73.47426946479342,73.48150859179687,73.48150859189687,73.49598247183741,73.49598247193741,73.50321675189166,73.50321675199166,73.51044874634546,73.51044874644546,73.51767971621976,73.51767971631976,73.53213909451958,73.53213909461958,73.53936427157896,73.53936427167896,73.54658858168646,73.54658858178647,73.55380985746234,73.55380985756234,73.5610280989066,73.5610280990066,73.56824444881946,73.56824444891946,73.57545713389024,73.57545713399024,73.58988128241776,73.58988128251777,73.60430137203416,73.60430137213416,73.61150751555886,73.61150751565886,73.61870873322053,73.61870873332053,73.62590857164055,73.62590857174055,73.63310777955012,73.63310777965012,73.64750450087236,73.64750450097236,73.65469700960823,73.65469700970823,73.6618877844403,73.6618877845403,73.66907619485814,73.66907619495814,73.67626283196527,73.67626283206528,73.6906298798887,73.6906298799887,73.69780906909097,73.69780906919097,73.7049861697273,73.70498616982731,73.71216157586677,73.71216157596677,73.71933540573008,73.71933540583008,73.72650836864149,73.72650836874149,73.73367404127562,73.73367404137562,73.74083005921825,73.74083005931826,73.75513980950308,73.75513980960308,73.76229212319672,73.76229212329672,73.76944037797924,73.76944037807924,73.77658618953369,73.77658618963369,73.78373109472936,73.78373109482936,73.7908729261865,73.7908729262865,73.79801322077438,73.79801322087438,73.80515189967919,73.80515189977919,73.81228876586641,73.81228876596641,73.8336967635724,73.8336967636724,73.84796501838717,73.84796501848717,73.85509368793832,73.85509368803832,73.86221041719753,73.86221041729753,73.86932320576632,73.86932320586632,73.88354795535892,73.88354795545892,73.89065924646536,73.89065924656536,73.8977666756952,73.8977666757952,73.9048718587315,73.9048718588315,73.9119704608148,73.9119704609148,73.91906792009789,73.91906792019789,73.92616439420836,73.92616439430836,73.93325866153221,73.93325866163221,73.9403513525799,73.9403513526799,73.94743797496434,73.94743797506435,73.95452317870024,73.95452317880024,73.96160649090474,73.96160649100474,73.96868775395023,73.96868775405024,73.98284894020647,73.98284894030647,73.99700531882038,73.99700531892039,74.01115346139134,74.01115346149135,74.01822408457272,74.01822408467272,74.03943398377162,74.03943398387162,74.04650232135255,74.04650232145255,74.05356793985709,74.05356793995709,74.0606322973407,74.0606322974407,74.07475553474825,74.07475553484825,74.08181335068578,74.08181335078578,74.08886848695383,74.08886848705383,74.10297568575139,74.10297568585139,74.11002439869407,74.11002439879407,74.11707232349865,74.11707232359865,74.13115950358892,74.13115950368892,74.14523691076695,74.14523691086696,74.1593115200548,74.1593115201548,74.17338368611459,74.17338368621459,74.18041671510942,74.18041671520942,74.18744521231025,74.18744521241025,74.20150090628412,74.20150090638413,74.20852766958119,74.20852766968119,74.2225771372642,74.2225771373642,74.2295994081742,74.2295994082742,74.23661955111136,74.23661955121136,74.24363524106836,74.24363524116836,74.2506503005149,74.25065030061491,74.2646786066904,74.2646786067904,74.27169130172268,74.27169130182268,74.27870316920999,74.27870316930999,74.29272469739796,74.29272469749796,74.29973234834652,74.29973234844653,74.31374520701559,74.31374520711559,74.32075009948086,74.32075009958086,74.32775424321495,74.32775424331496,74.34176044211722,74.34176044221722,74.34875225149031,74.34875225159031,74.36972369951229,74.3697236996123,74.3767079427598,74.3767079428598,74.38368954574473,74.38368954584473,74.39066657752878,74.39066657762878,74.40461957711045,74.40461957721045,74.41159448092168,74.41159448102168,74.41856851778101,74.41856851788101,74.43251529107185,74.43251529117185,74.43948424444055,74.43948424454055,74.44645229145046,74.44645229155046,74.45341931388086,74.45341931398086,74.46735056085146,74.46735056095146,74.47430954427341,74.47430954437341,74.48822211237145,74.48822211247145,74.49517483009565,74.49517483019565,74.50212565628844,74.50212565638844,74.50907143839751,74.50907143849751,74.51601083658811,74.51601083668811,74.52294767332994,74.52294767342994,74.53681815485436,74.53681815495436,74.54375018395389,74.54375018405389,74.55068122788082,74.55068122798082,74.55761081375228,74.55761081385228,74.57145623248567,74.57145623258567,74.58528048971155,74.58528048981155,74.59218587974388,74.59218587984388,74.59908977231386,74.59908977241386,74.60599260089742,74.60599260099742,74.61289168582509,74.61289168592509,74.626680161582,74.626680161682,74.6404559877227,74.6404559878227,74.64733962514394,74.64733962524394,74.66110477201362,74.66110477211362,74.6679849022204,74.6679849023204,74.67486420488218,74.67486420498219,74.68862194325388,74.68862194335388,74.70237759305965,74.70237759315965,74.70925287621723,74.70925287631724,74.71612441571891,74.71612441581891,74.72986532734254,74.72986532744254,74.75047182802531,74.75047182812531,74.75733690479471,74.75733690489471,74.76419973537058,74.76419973547058,74.77106126551861,74.77106126561861,74.77791952489359,74.77791952499359,74.78477510459909,74.7847751046991,74.79162883218011,74.79162883228011,74.80533498691429,74.80533498701429,74.81218607423271,74.81218607433271,74.81903393018501,74.81903393028502,74.82588044630256,74.82588044640256,74.8327239674954,74.8327239675954,74.83956543952922,74.83956543962923,74.84640628105258,74.84640628115258,74.86008654545074,74.86008654555074,74.86692415560796,74.86692415570796,74.87376022889592,74.87376022899592,74.88059496234914,74.88059496244914,74.89425930635802,74.89425930645803,74.90108659190636,74.90108659200637,74.90791245880615,74.90791245890615,74.92156111886719,74.92156111896719,74.92838072006923,74.92838072016923,74.93519846914675,74.93519846924676,74.94883282450161,74.94883282460161,74.95564726339919,74.95564726349919,74.96927290982822,74.96927290992822,74.9760834080354,74.9760834081354,74.982884685027,74.982884685127,74.98968363701125,74.98968363711126,74.99648109153316,74.99648109163316,75.00327531468892,75.00327531478892,75.01006524249215,75.01006524259215,75.01685020502543,75.01685020512544,75.02363225144782,75.02363225154782,75.03719413750596,75.03719413760597,75.04396641101613,75.04396641111613,75.0507370294363,75.05073702953631,75.0575067809046,75.0575067810046,75.06427527135197,75.06427527145198,75.07104285544054,75.07104285554054,75.07780921791509,75.07780921801509,75.08457487106534,75.08457487116534,75.0913394996361,75.09133949973611,75.0980996752267,75.09809967532671,75.10485886564469,75.10485886574469,75.11161600690366,75.11161600700366,75.11837082315529,75.11837082325529,75.12512496948955,75.12512496958955,75.13187489928507,75.13187489938507,75.1386240803494,75.1386240804494,75.14537227624113,75.14537227634113,75.1521169649184,75.15211696501841,75.16559991894756,75.16559991904757,75.17907530685113,75.17907530695113,75.18580848871238,75.18580848881238,75.19254096124938,75.19254096134938,75.19927272446208,75.19927272456208,75.21273487164586,75.21273487174587,75.2329245260967,75.2329245261967,75.23964892021833,75.23964892031833,75.2530967626959,75.2530967627959,75.27326584674279,75.27326584684279,75.2934303595888,75.2934303596888,75.30014943377837,75.30014943387837,75.30686255752542,75.30686255762542,75.31357177998898,75.31357178008898,75.32027584014807,75.32027584024807,75.33368100494847,75.33368100504848,75.34038029687217,75.34038029697217,75.34707501759499,75.34707501769499,75.35376824085546,75.35376824095546,75.36046075479166,75.36046075489166,75.36715149541716,75.36715149551716,75.373839635187,75.373839635287,75.38052162747978,75.38052162757978,75.3938822230716,75.3938822231716,75.40055712212164,75.40055712222164,75.40722914446768,75.40722914456768,75.42056901202793,75.42056901212793,75.43390312618017,75.43390312628017,75.44056628197278,75.44056628207278,75.45388904693662,75.45388904703663,75.46054853788715,75.46054853798715,75.46720605849248,75.46720605859248,75.47386235748377,75.47386235758377,75.48051696197818,75.48051696207818,75.48717042367237,75.48717042377237,75.50047533730864,75.50047533740864,75.50712600111265,75.50712600121265,75.51377359117812,75.51377359127812,75.52041980200195,75.52041980210196,75.52706372722535,75.52706372732536,75.54035051368572,75.54035051378573,75.55363481751114,75.55363481761114,75.56027263466439,75.56027263476439,75.5669092302036,75.56690923030361,75.57354444650119,75.57354444660119,75.58681204179925,75.58681204189925,75.6000728197029,75.6000728198029,75.6066967656259,75.6066967657259,75.613319844597,75.613319844697,75.6398052248663,75.6398052249663,75.65303089121835,75.65303089131835,75.65963848327613,75.65963848337613,75.6662449719406,75.6662449720406,75.67284728347322,75.67284728357322,75.68604985737946,75.68604985747946,75.69264877991833,75.69264878001833,75.70584075336735,75.70584075346736,75.71243352842917,75.71243352852917,75.71902393907676,75.71902393917676,75.73878902354244,75.73878902364244,75.7519564858971,75.7519564859971,75.75853822702577,75.75853822712577,75.76511453000165,75.76511453010166,75.77168929610829,75.7716892962083,75.77825795414479,75.7782579542448,75.78482542997416,75.78482543007416,75.79139117189975,75.79139117199975,75.80451938497788,75.80451938507788,75.81764212049634,75.81764212059635,75.82419718315091,75.82419718325092,75.83074854155649,75.83074854165649,75.83729789020995,75.83729789030996,75.84384649013225,75.84384649023225,75.8634903195539,75.8634903196539,75.87003434827531,75.87003434837531,75.88311972604866,75.88311972614866,75.90274164523161,75.90274164533162,75.9092767679927,75.9092767680927,75.91581102380188,75.91581102390188,75.92233692534738,75.92233692544738,75.92886168409268,75.92886168419268,75.93538459071347,75.93538459081347,75.94190261087815,75.94190261097815,75.94841980349784,75.94841980359784,75.96145221839203,75.96145221849203,75.9679673224458,75.9679673225458,75.98099563902196,75.98099563912196,75.98750877273052,75.98750877283052,76.00053389734744,76.00053389744744,76.00704139586871,76.00704139596871,76.01354739692763,76.01354739702764,76.0265540002996,76.0265540003996,76.03305322337101,76.03305322347101,76.03955071253864,76.03955071263864,76.04604646780248,76.04604646790249,76.05254119848682,76.05254119858682,76.05903458933642,76.05903458943642,76.06552451237845,76.06552451247845,76.07201278033051,76.07201278043051,76.07849836861308,76.07849836871308,76.09146682610185,76.09146682620185,76.097946424535,76.097946424635,76.10442424965747,76.10442424975747,76.11089829171713,76.11089829181714,76.11737087572133,76.11737087582134,76.12384251395986,76.12384251405986,76.13031151193579,76.13031151203579,76.13677897304247,76.13677897314247,76.14324119303089,76.14324119313089,76.14970254606743,76.14970254616743,76.16262296654006,76.16262296664006,76.16908132465187,76.16908132475187,76.1755388158118,76.1755388159118,76.18844906930316,76.18844906940316,76.20780384908299,76.207803849183,76.21425231606187,76.21425231616188,76.22069991608888,76.22069991618888,76.22714645212945,76.22714645222945,76.23358932332795,76.23358932342795,76.24646820892362,76.24646820902362,76.25934039534557,76.25934039544558,76.26576580928554,76.26576580938554,76.27218933169411,76.27218933179411,76.28503011081348,76.28503011091348,76.29144701286215,76.29144701296215,76.29786206278632,76.29786206288632,76.30427463007554,76.30427463017554,76.31068585753002,76.31068585763002,76.31709535108071,76.31709535118071,76.32991094918833,76.32991094928833,76.34912796243131,76.34912796253131,76.3555306779945,76.3555306780945,76.361930792702,76.361930792802,76.37472645091613,76.37472645101613,76.38112171857443,76.38112171867444,76.39391032295273,76.39391032305274,76.40669664173059,76.40669664183059,76.4130882445468,76.4130882446468,76.41947708887973,76.41947708897973,76.42586081031511,76.42586081041512,76.43862711038567,76.43862711048567,76.4450074428273,76.4450074429273,76.45776661024821,76.45776661034822,76.46414339606847,76.46414339616847,76.47051371915646,76.47051371925646,76.47688053502996,76.47688053512996,76.48961219643179,76.4896121965318,76.49597511102179,76.4959751111218,76.50869952155325,76.50869952165326,76.51505786494238,76.51505786504238,76.52777325129279,76.52777325139279,76.5341285997572,76.5341285998572,76.54048087448308,76.54048087458308,76.54683192759495,76.54683192769495,76.55953233932179,76.55953233942179,76.56587756021183,76.56587756031183,76.57856666215719,76.57856666225719,76.5849077059154,76.5849077060154,76.59124453313487,76.59124453323487,76.59758025696102,76.59758025706103,76.6039125129796,76.6039125130796,76.6102419317011,76.6102419318011,76.62289753777793,76.62289753787793,76.63554817858486,76.63554817868486,76.64819483929445,76.64819483939445,76.6545162978213,76.6545162979213,76.67347787551168,76.67347787561168,76.67979484165146,76.67979484175146,76.69874353328417,76.69874353338417,76.70505766212686,76.70505766222686,76.71137092401766,76.71137092411766,76.71768339777037,76.71768339787037,76.72399492575738,76.72399492585738,76.73661471095836,76.73661471105837,76.74292103723403,76.74292103733403,76.74922192535692,76.74922192545692,76.75552088254152,76.75552088264152,76.76181676598759,76.76181676608759,76.7681111519713,76.7681111520713,76.78069850529018,76.78069850539018,76.78699076330106,76.78699076340106,76.79328105096675,76.79328105106676,76.7995690924389,76.7995690925389,76.80585078939947,76.80585078949947,76.81213146178055,76.81213146188055,76.82469154552177,76.82469154562177,76.83724713687592,76.83724713697592,76.84352095245566,76.84352095255566,76.84979181251761,76.84979181261761,76.85605888951676,76.85605888961676,76.86231725759009,76.8623172576901,76.86857499515295,76.86857499525296,76.87483131406728,76.87483131416728,76.88108696306423,76.88108696316424,76.89359569960935,76.89359569970935,76.89984650155709,76.8998465016571,76.91234203678934,76.91234203688934,76.91858657303932,76.91858657313932,76.92482933597861,76.92482933607862,76.93731316736033,76.93731316746033,76.94355407817513,76.94355407827513,76.9497908906719,76.94979089077191,76.96226270294785,76.96226270304786,76.968497308658,76.968497308758,76.974728683002,76.974728683102,76.98095828403531,76.98095828413531,76.99341311193558,76.99341311203558,76.99963652608496,76.99963652618496,77.0058578122615,77.0058578123615,77.01829936003507,77.01829936013507,77.0307385828013,77.03073858290131,77.04317481064282,77.04317481074283,77.0493859298381,77.0493859299381,77.05559318715677,77.05559318725678,77.06179906523379,77.06179906533379,77.0679990716821,77.0679990717821,77.0803960108402,77.0803960109402,77.0865922342257,77.0865922343257,77.09278372878272,77.09278372888272,77.1113564391431,77.1113564392431,77.12373355662841,77.12373355672841,77.12991843082553,77.12991843092553,77.13610259569838,77.13610259579838,77.1422837262396,77.1422837263396,77.1484616648216,77.1484616649216,77.16081643859226,77.16081643869227,77.16698590468987,77.16698590478987,77.17315458264939,77.17315458274939,77.1793222360294,77.17932223612941,77.185486776264,77.185486776364,77.19164970081552,77.19164970091552,77.21013717404223,77.21013717414223,77.22245884601342,77.22245884611343,77.22861731758478,77.22861731768478,77.23477409465926,77.23477409475926,77.2409268522295,77.24092685232951,77.24707582673696,77.24707582683696,77.25322397369942,77.25322397379942,77.25936691895055,77.25936691905055,77.26550915487739,77.2655091549774,77.27164756833454,77.27164756843455,77.27778290805317,77.27778290815317,77.28391403123304,77.28391403133304,77.29617462250283,77.29617462260283,77.30230184439921,77.30230184449921,77.30842705654347,77.30842705664347,77.31454935257682,77.31454935267682,77.33291474321453,77.33291474331453,77.33903416254388,77.33903416264388,77.34514964118281,77.34514964128282,77.35126421346297,77.35126421356297,77.36348945673974,77.36348945683974,77.38181796251517,77.38181796261517,77.38792481104211,77.38792481114211,77.39402890108576,77.39402890118576,77.40013165129466,77.40013165139466,77.40623223412383,77.40623223422384,77.41233092542161,77.41233092552162,77.41842709467753,77.41842709477753,77.4245219635056,77.4245219636056,77.43061576834727,77.43061576844727,77.4367084303887,77.43670843048871,77.44889158709188,77.44889158719188,77.4549812936155,77.4549812937155,77.46106993615274,77.46106993625274,77.46715672656546,77.46715672666546,77.47324201951584,77.47324201961584,77.47932573619003,77.47932573629004,77.48540882235378,77.48540882245378,77.49148938647565,77.49148938657565,77.49756727092804,77.49756727102805,77.50972118770834,77.50972118780834,77.5340258293097,77.5340258294097,77.54009985188549,77.54009985198549,77.5583205009643,77.5583205010643,77.5643917256499,77.5643917257499,77.57045770921725,77.57045770931725,77.5765190821768,77.5765190822768,77.58257749961855,77.58257749971855,77.59469291585351,77.59469291595352,77.61286304528123,77.61286304538123,77.61891661567377,77.61891661577377,77.62496845216253,77.62496845226254,77.63101607211254,77.63101607221255,77.6370610912069,77.6370610913069,77.64914817387778,77.64914817397778,77.6551881488884,77.6551881489884,77.66122638999522,77.66122639009522,77.6672631730466,77.66726317314661,77.6732990891461,77.6732990892461,77.67933216794849,77.67933216804849,77.68535921749455,77.68535921759455,77.69741213437955,77.69741213447955,77.70343760764943,77.70343760774944,77.71548205205005,77.71548205215005,77.72149980156674,77.72149980166674,77.72751538370369,77.72751538380369,77.75157467791989,77.75157467801989,77.75758734394594,77.75758734404594,77.76961090268736,77.76961090278736,77.77562084963702,77.77562084973702,77.78162855039315,77.78162855049315,77.78763298037624,77.78763298047625,77.79363626755911,77.79363626765911,77.7996388454177,77.7996388455177,77.81164120324469,77.81164120334469,77.81763956456454,77.81763956466455,77.82363705893249,77.82363705903249,77.8296293121822,77.82962931228221,77.83561924042456,77.83561924052457,77.84759692952957,77.84759692962957,77.85358165606058,77.85358165616059,77.85956496394306,77.85956496404306,77.8655451192732,77.8655451193732,77.8775046417954,77.8775046418954,77.88945998718575,77.88945998728575,77.89543533487358,77.89543533497358,77.90140453508437,77.90140453518437,77.9073722378328,77.9073722379328,77.92527270581553,77.92527270591553,77.94316931192164,77.94316931202164,77.9491334286418,77.9491334287418,77.95509553560984,77.95509553570984,77.9610568150329,77.9610568151329,77.97297834929951,77.97297834939951,77.98489413015811,77.98489413025811,77.99680321184302,77.99680321194302,78.00275586115407,78.00275586125407,78.00870713122347,78.00870713132348,78.0146567462029,78.0146567463029,78.02060297218858,78.02060297228859,78.02654703079453,78.02654703089453,78.03248990719337,78.03248990729337,78.03843053739865,78.03843053749866,78.04436864556209,78.04436864566209,78.05624001483973,78.05624001493973,78.06217035984304,78.06217035994304,78.07402888246993,78.07402888256993,78.08588401610308,78.08588401620308,78.09180908058123,78.09180908068123,78.09773296285226,78.09773296295226,78.1036559387645,78.1036559388645,78.11550039312662,78.11550039322663,78.12142139869366,78.12142139879366,78.13326136066871,78.13326136076871,78.13917834673151,78.13917834683151,78.14508898828277,78.14508898838277,78.15099423108813,78.15099423118814,78.15689829168639,78.15689829178639,78.16280156414656,78.16280156424656,78.17460684804597,78.17460684814597,78.18050755905738,78.18050755915738,78.21000776452753,78.21000776462753,78.227702804319,78.227702804419,78.23359697378432,78.23359697388432,78.23948925171823,78.23948925181823,78.24537778599625,78.24537778609626,78.25126297068742,78.25126297078742,78.25714606681268,78.25714606691268,78.26890689972423,78.26890689982423,78.28066272795894,78.28066272805894,78.29241481253777,78.29241481263777,78.30416236532263,78.30416236542263,78.31003170843833,78.31003170853833,78.33350506139695,78.33350506149695,78.34523247725379,78.34523247735379,78.35109259915394,78.35109259925395,78.35695075070888,78.35695075080888,78.36280630140814,78.36280630150814,78.3745131074541,78.37451310755411,78.38036432339392,78.38036432349392,78.38621455416113,78.38621455426113,78.39791324238485,78.39791324248485,78.40376051763424,78.40376051773424,78.4096064530489,78.4096064531489,78.41545100922191,78.41545100932191,78.42713913639533,78.42713913649533,78.43298223451288,78.43298223461288,78.43881981566386,78.43881981576386,78.44465215569659,78.44465215579659,78.45048138258389,78.45048138268389,78.4563099001469,78.45630990024691,78.46213554100594,78.46213554110594,78.47378142397812,78.47378142407813,78.48541985904542,78.48541985914542,78.49123616046816,78.49123616056816,78.49705167375282,78.49705167385282,78.50867899607316,78.50867899617316,78.51448911061196,78.51448911071196,78.52610626595103,78.52610626605103,78.53191188810275,78.53191188820276,78.54352159553693,78.54352159563693,78.54932505030894,78.54932505040894,78.55512787457046,78.55512787467046,78.57253512574103,78.57253512584103,78.5783327482912,78.5783327483912,78.58412887337904,78.58412887347905,78.58992275227332,78.58992275237333,78.59571517311217,78.59571517321217,78.6015056630127,78.6015056631127,78.6072910300157,78.6072910301157,78.61885959664197,78.61885959674197,78.62464129880289,78.62464129890289,78.63620375735901,78.63620375745901,78.64198309510567,78.64198309520567,78.64775845275503,78.64775845285503,78.653533140487,78.653533140587,78.6593042421907,78.6593042422907,78.665074279908,78.665074280008,78.67084289897673,78.67084289907673,78.68237887609328,78.68237887619328,78.68814587947895,78.68814587957895,78.69967760064986,78.69967760074987,78.70544204258677,78.70544204268677,78.71696627644587,78.71696627654588,78.72272571370596,78.72272571380596,78.72848392935201,78.72848392945201,78.73424143567378,78.73424143577378,78.7457513648267,78.7457513649267,78.7515005562917,78.7515005563917,78.75724884139791,78.75724884149791,78.76874194380275,78.76874194390275,78.77448585474261,78.77448585484261,78.78022763770963,78.78022763780963,78.78596863253857,78.78596863263857,78.7917086421949,78.7917086422949,78.80318673056928,78.80318673066928,78.80892015927263,78.80892015937263,78.81464854389225,78.81464854399225,78.82037302722836,78.82037302732836,78.83182104813488,78.83182104823489,78.84898980872161,78.84898980882161,78.8547092873809,78.8547092874809,78.86042628340523,78.86042628350523,78.86614233366384,78.86614233376385,78.87185696527392,78.87185696537392,78.88328508569386,78.88328508579386,78.89470709804365,78.89470709814366,78.90041179911388,78.90041179921388,78.91752353791034,78.91752353801034,78.92322524405586,78.92322524415586,78.93462534616694,78.93462534626694,78.9460193402079,78.9460193403079,78.95171387429686,78.95171387439686,78.9744897644592,78.9744897645592,78.98018012141632,78.98018012151633,78.99155945608894,78.99155945618894,79.00862406422804,79.00862406432805,79.01999559633363,79.01999559643363,79.03704660909081,79.03704660919081,79.04840797421514,79.04840797431514,79.05976665966999,79.05976665976999,79.06544397294185,79.06544397304185,79.07111387771573,79.07111387781573,79.08244978597997,79.08244978607998,79.08811275513874,79.08811275523874,79.09943715658699,79.09943715668699,79.10509665793818,79.10509665803818,79.11641404495748,79.11641404505748,79.12206944799064,79.12206944809064,79.12772335356145,79.12772335366145,79.1333710328414,79.1333710329414,79.1390173328797,79.1390173329797,79.15030823845943,79.15030823855943,79.1672418580492,79.1672418581492,79.18417295559709,79.1841729556971,79.18981373866882,79.18981373876882,79.20109325565326,79.20109325575326,79.20673092557955,79.20673092567955,79.21236299972547,79.21236299982547,79.21799188191214,79.21799188201214,79.22361843909148,79.22361843919148,79.23487037124302,79.23487037134302,79.24049523392547,79.24049523402547,79.24611800804202,79.24611800814202,79.25174015164808,79.25174015174808,79.25735977321229,79.25735977331229,79.26297454772728,79.26297454782728,79.26858837647659,79.26858837657659,79.27420023488068,79.27420023498068,79.27981039878789,79.27981039888789,79.28541934108107,79.28541934118107,79.29663635871555,79.29663635881555,79.30223958700763,79.30223958710764,79.30784135724427,79.30784135734427,79.31344107832189,79.31344107842189,79.31903804091621,79.31903804101621,79.33022901058706,79.33022901068706,79.33582152020122,79.33582152030122,79.35259656640874,79.35259656650874,79.35818489889107,79.35818489899107,79.36935876596552,79.36935876606552,79.38610902523034,79.38610902533034,79.39168656022092,79.39168656032092,79.39726090325227,79.39726090335228,79.40283351237986,79.40283351247986,79.40840359946556,79.40840359956556,79.41397183442677,79.41397183452678,79.41953884777395,79.41953884787395,79.43067129819217,79.43067129829217,79.44180032020971,79.44180032030971,79.45292555916446,79.45292555926446,79.4584865038484,79.45848650394841,79.46404681802188,79.46404681812189,79.47516618534792,79.47516618544792,79.48072314993455,79.48072315003455,79.4862784594312,79.4862784595312,79.49182884306484,79.49182884316484,79.49737761101541,79.49737761111541,79.50847230961945,79.50847230971945,79.51401764916936,79.51401764926936,79.51956188532596,79.51956188542596,79.52510407232354,79.52510407242355,79.53064523474161,79.53064523484161,79.54172586508088,79.54172586518088,79.54726509656065,79.54726509666065,79.55280101786047,79.55280101796048,79.5583360328015,79.55833603290151,79.56386880154899,79.563868801649,79.56939975757889,79.5693997576789,79.57492949199477,79.57492949209477,79.58045835945875,79.58045835955875,79.59151337531033,79.59151337541033,79.59703688343535,79.59703688353535,79.60808267807136,79.60808267817136,79.61360468873401,79.61360468883402,79.62464713378318,79.62464713388319,79.63016500672092,79.63016500682092,79.64119535385049,79.6411953539505,79.64670660642834,79.64670660652834,79.66323681754047,79.66323681764047,79.67425211123268,79.67425211133268,79.69077195832901,79.69077195842901,79.69627588122268,79.69627588132268,79.70177858250231,79.70177858260232,79.71278260581994,79.71278260591994,79.71828203632654,79.71828203642654,79.72927802063572,79.72927802073572,79.74026896086117,79.74026896096117,79.74576082524217,79.74576082534217,79.76223527558496,79.76223527568496,79.7787066127823,79.7787066128823,79.78968415466035,79.78968415476035,79.80066012026224,79.80066012036224,79.80614646767665,79.80614646777666,79.81711510359438,79.81711510369438,79.82807790729026,79.82807790739027,79.83355724027575,79.83355724037575,79.8445147240396,79.8445147241396,79.84999137735561,79.84999137745561,79.8554603463253,79.8554603464253,79.86639572281592,79.86639572291593,79.87732573996759,79.87732574006759,79.8827881673912,79.8827881674912,79.88824665412439,79.88824665422439,79.89370041202909,79.89370041212909,79.90460643037613,79.90460643047614,79.91005727216994,79.91005727226994,79.91550649828065,79.91550649838065,79.92095481803258,79.92095481813259,79.93729800397769,79.93729800407769,79.9427424224461,79.9427424225461,79.95907354987854,79.95907354997854,79.96451410647036,79.96451410657036,79.96995312619292,79.96995312629292,79.9753914759981,79.9753914760981,79.98082888003758,79.98082888013758,79.98626502305613,79.98626502315614,79.99170041734351,79.99170041744351,79.99713360484425,79.99713360494425,80.0025659648,80.0025659649,80.00799458109987,80.00799458119987,80.01342126646142,80.01342126656142,80.02427298209457,80.02427298219457,80.02969706660046,80.02969706670046,80.03511949601636,80.03511949611637,80.0405410978873,80.0405410979873,80.05138272535298,80.05138272545298,80.06222159433538,80.06222159443539,80.0784772575464,80.0784772576464,80.08389405177502,80.08389405187502,80.08930958498271,80.08930958508272,80.09472444827303,80.09472444837303,80.10013678952149,80.10013678962149,80.10554826381805,80.10554826391805,80.11636955732119,80.1163695574212,80.12177697270664,80.12177697280664,80.12718324529186,80.12718324539186,80.14340040795753,80.14340040805753,80.15420932769278,80.15420932779278,80.15960905873192,80.15960905883192,80.16500804103987,80.16500804113987,80.17040556529237,80.17040556539237,80.18119801294169,80.1811980130417,80.18659100540022,80.18659100550022,80.19198127878235,80.19198127888235,80.19736993648142,80.19736993658142,80.20275733315957,80.20275733325957,80.21353137778466,80.21353137788466,80.21891609479331,80.21891609489332,80.22967524360864,80.22967524370864,80.2350476262563,80.2350476263563,80.24041728982758,80.24041728992758,80.24578384025342,80.24578384035343,80.2565137491459,80.2565137492459,80.26723853514082,80.26723853524082,80.27259854402058,80.27259854412058,80.27795457280303,80.27795457290303,80.2833099316681,80.2833099317681,80.28866454180199,80.288664541902,80.29937151587626,80.29937151597626,80.30471749589815,80.30471749599815,80.31006079625055,80.31006079635056,80.3154016927818,80.3154016928818,80.32074121007142,80.32074121017142,80.33141843193306,80.33141843203306,80.33675507251867,80.33675507261867,80.34742437359257,80.34742437369258,80.3527564429773,80.3527564430773,80.35808697549277,80.35808697559277,80.36874587314398,80.36874587324398,80.3847290376129,80.3847290377129,80.39005046713352,80.39005046723352,80.40069226218834,80.40069226228835,80.40600888406665,80.40600888416665,80.41664035451258,80.41664035461258,80.42195480901117,80.42195480911117,80.42726626858503,80.42726626868503,80.43788627162186,80.43788627172187,80.44319122905655,80.44319122915655,80.44849472843578,80.44849472853578,80.45379680916646,80.45379680926646,80.45909668311052,80.45909668321052,80.46439537484744,80.46439537494744,80.46969292378415,80.46969292388415,80.4749880294928,80.47498802959281,80.48028096782173,80.48028096792173,80.48557225106067,80.48557225116068,80.49614997048937,80.49614997058937,80.50672146362722,80.50672146372722,80.51200510192676,80.51200510202676,80.52785424351472,80.52785424361473,80.53313599028287,80.53313599038287,80.54369577957019,80.54369577967019,80.55424776629047,80.55424776639047,80.55952060709829,80.5595206071983,80.57006423955491,80.57006423965491,80.57533250916184,80.57533250926184,80.5858679843893,80.5858679844893,80.59113475653388,80.59113475663388,80.59639975536777,80.59639975546777,80.61219313618638,80.61219313628638,80.61745529772317,80.61745529782317,80.62797839918274,80.62797839928274,80.63323669884292,80.63323669894292,80.63849365866838,80.63849365876838,80.64374892399695,80.64374892409695,80.64900355881505,80.64900355891506,80.65951077929225,80.65951077939225,80.67001449255497,80.67001449265497,80.67526120658535,80.67526120668535,80.68050697485002,80.68050697495002,80.68575045751426,80.68575045761426,80.69099283678517,80.69099283688517,80.69623454613873,80.69623454623873,80.70671697967322,80.70671697977322,80.71195660046085,80.71195660056085,80.71719515726207,80.71719515736207,80.72243268948378,80.72243268958378,80.72766825136029,80.72766825146029,80.73813842934759,80.73813842944759,80.7486058094447,80.7486058095447,80.7538376276653,80.75383762776531,80.75906857893403,80.75906857903404,80.7642981115542,80.7642981116542,80.77475516704243,80.77475516714243,80.78520508988102,80.78520508998102,80.79042536187872,80.79042536197872,80.80086243806655,80.80086243816655,80.81129561297087,80.81129561307087,80.81651060444342,80.81651060454342,80.82172437430194,80.82172437440194,80.82693648907048,80.82693648917048,80.83214710637667,80.83214710647667,80.83735390121315,80.83735390131315,80.84255777993873,80.84255778003873,80.84776063408479,80.8477606341848,80.85295986279569,80.85295986289569,80.85815810633397,80.85815810643398,80.8633539066442,80.86335390674421,80.87893886434685,80.87893886444685,80.89451992076597,80.89451992086597,80.9049025366123,80.9049025367123,80.91009065257622,80.91009065267622,80.91527687700874,80.91527687710874,80.92564640976289,80.92564640986289,80.93082617146314,80.93082617156314,80.9360032929008,80.93600329300081,80.9411787198416,80.9411787199416,80.95152843092296,80.95152843102296,80.96187609284532,80.96187609294532,80.96704809138545,80.96704809148545,80.97739090625859,80.9773909063586,80.98256113148804,80.98256113158804,80.98773064739322,80.98773064749322,80.99289283361422,80.99289283371422,81.01353830772518,81.01353830782519,81.0238569661661,81.0238569662661,81.02901454177932,81.02901454187932,81.03932835317102,81.03932835327102,81.04448234275596,81.04448234285596,81.05478949438086,81.05478949448086,81.05994194709655,81.05994194719655,81.06509195658417,81.06509195668417,81.07023841945042,81.07023841955042,81.07538279375076,81.07538279385076,81.08052468541614,81.08052468551614,81.08566448851559,81.08566448861559,81.09080003566939,81.09080003576939,81.10106876556276,81.10106876566276,81.10620013558473,81.10620013568473,81.11646086587656,81.11646086597656,81.12158979267048,81.12158979277048,81.1267178919194,81.1267178920194,81.13184520303025,81.13184520313025,81.13697184422372,81.13697184432372,81.1420975396515,81.1420975397515,81.14722228931358,81.14722228941358,81.15234487159591,81.15234487169592,81.15746548353306,81.15746548363306,81.16770414595857,81.16770414605857,81.17793835540392,81.17793835550393,81.18305317452615,81.18305317462615,81.18816535338581,81.18816535348581,81.19327638944524,81.19327638954525,81.19838679499422,81.19838679509422,81.20349601833605,81.20349601843606,81.2086033107396,81.2086033108396,81.21370776584605,81.21370776594605,81.21880875314493,81.21880875324493,81.229006077728,81.229006077828,81.23919528448883,81.23919528458883,81.24428669591,81.24428669601001,81.25446833654523,81.25446833664523,81.26464662759362,81.26464662769362,81.26972869957851,81.26972869967851,81.27480923469415,81.27480923479415,81.28496900449758,81.28496900459758,81.29004319510165,81.29004319520165,81.30018834494365,81.30018834504365,81.30525867367112,81.30525867377112,81.32046583738381,81.32046583748381,81.33059687955414,81.33059687965414,81.33566015444575,81.33566015454575,81.34578544320807,81.34578544330807,81.35590868281137,81.35590868291138,81.36096860811614,81.36096860821614,81.37614440393314,81.37614440403314,81.38119731480899,81.38119731490899,81.38624766423605,81.38624766433605,81.40139591462706,81.40139591472706,81.41149086007319,81.41149086017319,81.42158300762912,81.42158300772913,81.43167172678442,81.43167172688442,81.43671423423756,81.43671423433756,81.44175461371788,81.44175461381788,81.45183419047139,81.45183419057139,81.46191010238283,81.46191010248283,81.46694606828989,81.46694606838989,81.4719788028308,81.4719788029308,81.48204305029861,81.48204305039862,81.48707259288031,81.48707259298031,81.49713029880203,81.49713029890204,81.5021578710385,81.50215787113851,81.50718201487432,81.50718201497432,81.52225200315372,81.52225200325373,81.52727275799579,81.52727275809579,81.54233297336296,81.54233297346296,81.5523689266736,81.5523689267736,81.55738471624574,81.55738471634574,81.56741444326552,81.56741444336552,81.57242798664413,81.57242798674413,81.58746609473809,81.58746609483809,81.5924769978541,81.59247699795411,81.59748644291469,81.59748644301469,81.60249178965724,81.60249178975724,81.60749382621984,81.60749382631984,81.61249495642363,81.61249495652363,81.61749364339938,81.61749364349939,81.6224911087611,81.6224911088611,81.62748605208095,81.62748605218096,81.63247784284847,81.63247784294848,81.6374686484434,81.6374686485434,81.64245795657595,81.64245795667595,81.64744588546688,81.64744588556688,81.65742075807611,81.65742075817612,81.6673923599123,81.6673923600123,81.67237512649878,81.67237512659878,81.67735698672645,81.67735698682645,81.69229815383623,81.69229815393624,81.70225367765553,81.70225367775554,81.71220746757105,81.71220746767105,81.71718176167315,81.71718176177315,81.72215534645096,81.72215534655096,81.72712246849649,81.72712246859649,81.73208829011419,81.7320882902142,81.73705304774548,81.73705304784548,81.74201532274179,81.7420153228418,81.75193794179614,81.75193794189614,81.75689722186776,81.75689722196776,81.76185535913915,81.76185535923915,81.76681172309985,81.76681172319985,81.77176446162538,81.77176446172538,81.77671593912997,81.77671593922997,81.78166619502053,81.78166619512054,81.78661235259307,81.78661235269307,81.79155780084133,81.79155780094133,81.80144416554387,81.80144416564387,81.80638389979103,81.80638389989103,81.81625875767757,81.81625875777758,81.82119372368933,81.82119372378934,81.83599586324132,81.83599586334132,81.84092779492147,81.84092779502147,81.8507893726813,81.8507893727813,81.86064519703315,81.86064519713315,81.86556908970485,81.86556908980485,81.87049187898323,81.87049187908323,81.8803365511812,81.8803365512812,81.89017929244086,81.89017929254086,81.90494009415042,81.90494009425042,81.91477818539539,81.9147781854954,81.91969403905865,81.91969403915866,81.9295242489228,81.9295242490228,81.93443584664041,81.93443584674041,81.94916855122732,81.94916855132732,81.95407731164782,81.95407731174782,81.96389199519174,81.96389199529175,81.96879665729422,81.96879665739422,81.97369966430674,81.97369966440674,81.98350453553152,81.98350453563152,81.98840466584002,81.98840466594002,81.99330266817572,81.99330266827572,82.00309792411589,82.0030979242159,82.01288407706122,82.01288407716122,82.01777116368446,82.01777116378446,82.02754431235142,82.02754431245143,82.03242927100183,82.03242927110183,82.0373120228656,82.03731202296561,82.04707543802725,82.04707543812725,82.06171671859657,82.06171671869657,82.07635295508214,82.07635295518214,82.08122837726175,82.08122837736175,82.08610076510975,82.08610076520975,82.09097232541276,82.09097232551277,82.10070981083148,82.10070981093148,82.10557455374007,82.10557455384007,82.12503151562233,82.12503151572233,82.13475792770099,82.13475792780099,82.13961865110535,82.13961865120535,82.14447799526808,82.14447799536808,82.14933556612012,82.14933556622012,82.15419006323363,82.15419006333363,82.15904389042977,82.15904389052977,82.16389523499096,82.16389523509096,82.17359650546479,82.17359650556479,82.17844548561172,82.17844548571172,82.18814171200181,82.18814171210181,82.19298769722403,82.19298769732403,82.19783112099749,82.1978311210975,82.20267158925313,82.20267158935313,82.2075113481845,82.2075113482845,82.21234909736376,82.21234909746376,82.21718188127309,82.2171818813731,82.22201324653389,82.22201324663389,82.22684240500804,82.22684240510804,82.2316706571234,82.2316706572234,82.23649709652118,82.23649709662118,82.24132270837396,82.24132270847396,82.24614757149557,82.24614757159557,82.25579430281407,82.25579430291407,82.2606156193143,82.2606156194143,82.27025612434194,82.27025612444194,82.2750748399865,82.2750748400865,82.27989245223775,82.27989245233775,82.2895254699536,82.2895254700536,82.29433992965252,82.29433992975252,82.29915190671647,82.29915190681648,82.30396218928355,82.30396218938355,82.31357948364466,82.31357948374466,82.31838582552132,82.31838582562132,82.32799665715015,82.32799665725015,82.3376038239369,82.3376038240369,82.34240543698506,82.34240543708506,82.35200728383975,82.35200728393976,82.35680739942556,82.35680739952556,82.36160306203121,82.36160306213121,82.36639604496736,82.36639604506736,82.37598098626017,82.37598098636018,82.38076931918165,82.38076931928165,82.38555694277885,82.38555694287885,82.39034299009988,82.39034299019988,82.3999136266865,82.3999136267865,82.40469754603471,82.40469754613471,82.4142622715857,82.4142622716857,82.41904244727802,82.41904244737802,82.42382100728726,82.42382100738726,82.4285974787306,82.4285974788306,82.43337138872515,82.43337138882515,82.43814336778141,82.43814336788141,82.44291444047887,82.44291444057887,82.44768460681753,82.44768460691753,82.45722068354922,82.45722068364923,82.4619824168104,82.4619824169104,82.46674312549207,82.46674312559207,82.47150229730448,82.47150229740448,82.48101910406005,82.48101910416005,82.4857748474718,82.4857748475718,82.49052932986262,82.49052932996263,82.50954477679095,82.50954477689095,82.51429622485014,82.51429622495014,82.52379809638904,82.52379809648905,82.53329709122394,82.53329709132394,82.55703450984717,82.55703450994717,82.56177267777969,82.56177267787969,82.5665090329946,82.5665090330946,82.57598071884493,82.57598071894493,82.58545055257076,82.58545055267076,82.59965335251776,82.59965335261776,82.61385035964985,82.61385035974985,82.6185793457737,82.6185793458737,82.62803617522117,82.62803617532117,82.63276200879268,82.63276200889268,82.64693828789319,82.6469382879932,82.65638699951842,82.65638699961842,82.66110999579283,82.66110999589283,82.67055315104456,82.67055315114456,82.67999224738517,82.67999224748517,82.68470980550681,82.68470980560681,82.68942641786276,82.68942641796276,82.69414062639756,82.69414062649756,82.70828120284293,82.70828120294293,82.71770536396687,82.71770536406687,82.7271275547456,82.72712755484561,82.73654757814461,82.73654757824461,82.74596346381868,82.74596346391868,82.75537477829188,82.75537477839188,82.76478412241987,82.76478412251987,82.76948769109055,82.76948769119055,82.7788928974936,82.77889289759361,82.7929965496698,82.7929965497698,82.79769645349839,82.79769645359839,82.80239513571296,82.80239513581296,82.81179131793496,82.81179131803496,82.82118568743937,82.82118568753937,82.83527357685391,82.83527357695391,82.83996753023997,82.83996753033998,82.84466085311558,82.84466085321559,82.84935267852883,82.84935267862883,82.86342606620265,82.86342606630265,82.87280475175922,82.87280475185922,82.87749232122681,82.87749232132681,82.88686604151346,82.88686604161346,82.89155116775301,82.89155116785301,82.90091889819023,82.90091889829023,82.91496509510019,82.91496509520019,82.91964340394532,82.91964340404532,82.92431950600381,82.92431950610381,82.92899481992423,82.92899482002423,82.93366930629965,82.93366930639965,82.94301615107769,82.94301615117769,82.96637676088358,82.96637676098358,82.97104628198908,82.97104628208908,82.97571485732888,82.97571485742888,82.98038272334442,82.98038272344442,82.98504704273857,82.98504704283857,82.98970939178753,82.98970939188753,82.99437063744315,82.99437063754316,82.99903034622953,82.99903034632953,83.0036880846707,83.0036880847707,83.00834389217357,83.00834389227357,83.01765424615837,83.01765424625837,83.02696219632202,83.02696219642202,83.03161504830707,83.03161504840708,83.04091625989011,83.04091625999011,83.04556450126739,83.0455645013674,83.05485905308363,83.05485905318363,83.0595025262255,83.0595025263255,83.06414497478787,83.06414497488787,83.06878671402595,83.06878671412595,83.07342703461549,83.0734270347155,83.0827032228144,83.0827032229144,83.09661015572516,83.09661015582516,83.10124468349979,83.10124468359979,83.11051117760029,83.11051117770029,83.12440593377767,83.12440593387767,83.12903561450308,83.12903561460308,83.13829343908468,83.13829343918468,83.14292107065108,83.14292107075109,83.1521753486113,83.1521753487113,83.16142824732987,83.16142824742987,83.17067645662685,83.17067645672685,83.17529699495051,83.17529699505052,83.17991654810157,83.17991654820158,83.18453409150052,83.18453409160053,83.20300162483375,83.20300162493375,83.2076163703425,83.20761637044251,83.21223032771317,83.21223032781317,83.22145690261979,83.2214569027198,83.23068146777429,83.23068146787429,83.23529270606858,83.23529270616858,83.23990169816932,83.23990169826932,83.24450781356607,83.24450781366608,83.25371649773821,83.25371649783821,83.2583173720167,83.25831737211671,83.26751774133207,83.26751774143207,83.27211566009275,83.27211566019275,83.2767083771421,83.2767083772421,83.28129987257742,83.28129987267742,83.28588994936419,83.28588994946419,83.29965840641381,83.29965840651381,83.30424603997253,83.30424604007253,83.32259235766863,83.32259235776863,83.33175804890834,83.33175804900834,83.34550156138765,83.34550156148765,83.35466165684697,83.35466165694697,83.359238886983,83.359238887083,83.36381477728428,83.36381477738428,83.3729642722864,83.3729642723864,83.37753744351129,83.37753744361129,83.3821098265981,83.3821098266981,83.39125179488153,83.39125179498153,83.40038950721932,83.40038950731932,83.40952359412192,83.40952359422192,83.41865342507887,83.41865342517887,83.42321595643965,83.42321595653965,83.43233963991956,83.43233964001956,83.43690027974894,83.43690027984894,83.45058015007807,83.45058015017807,83.45513294793352,83.45513294803352,83.45968263264355,83.45968263274355,83.46878078044958,83.46878078054958,83.47787731257255,83.47787731267255,83.48242463286833,83.48242463296833,83.48697021926033,83.48697021936033,83.50060555978777,83.50060555988777,83.50514921524147,83.50514921534148,83.50969192492947,83.50969192502947,83.51423349181725,83.51423349191725,83.52331520694426,83.52331520704426,83.52785480348683,83.52785480358683,83.53239369070512,83.53239369080512,83.53693123808868,83.53693123818869,83.54146646046489,83.54146646056489,83.55053426495473,83.55053426505474,83.55959848341631,83.55959848351631,83.56866155907765,83.56866155917766,83.5777226249869,83.5777226250869,83.58678219343378,83.58678219353378,83.59583833348,83.59583833358,83.60036472870969,83.6003647288097,83.60941582467218,83.60941582477219,83.61393985548763,83.61393985558763,83.61846274350285,83.61846274360285,83.62298342473143,83.62298342483143,83.63654341925817,83.63654341935818,83.64106134200347,83.64106134210347,83.64557827957616,83.64557827967616,83.65009237985176,83.65009237995176,83.65459883518795,83.65459883528796,83.65910458119987,83.65910458129987,83.66811433931993,83.66811433941993,83.67261768151069,83.67261768161069,83.67711917157696,83.67711917167696,83.68612077246786,83.68612077256786,83.69511957546857,83.69511957556857,83.6996168095892,83.6996168096892,83.70860934689215,83.70860934699215,83.71759983503608,83.71759983513608,83.72658847105552,83.72658847115552,83.73557478206762,83.73557478216762,83.74006527760763,83.74006527770763,83.74455494560266,83.74455494570266,83.74904327376295,83.74904327386295,83.7580187084695,83.75801870856951,83.76250494806388,83.76250494816388,83.76699008426493,83.76699008436493,83.77147411707267,83.77147411717267,83.77595724352162,83.77595724362162,83.78492199895716,83.78492199905716,83.78940256395734,83.78940256405734,83.7938811980192,83.79388119811921,83.7983580981773,83.7983580982773,83.80283350087304,83.80283350097304,83.80730590864405,83.80730590874406,83.82072175271547,83.82072175281547,83.82519104734106,83.82519104744107,83.83412644463301,83.83412644473302,83.83859219263724,83.83859219273724,83.84305466986842,83.84305466996842,83.84751651658912,83.84751651668913,83.85197442261943,83.85197442271944,83.85643165873236,83.85643165883236,83.86534518519251,83.86534518529251,83.8742570959696,83.8742570960696,83.8787121647028,83.8787121648028,83.8831661300427,83.8831661301427,83.88761851910641,83.88761851920641,83.89652116926102,83.89652116936102,83.90542117915307,83.90542117925307,83.91876845521129,83.9187684553113,83.92321560315676,83.92321560325676,83.92766093838463,83.92766093848464,83.93210501259158,83.93210501269158,83.94099158472932,83.94099158482932,83.94543349155653,83.94543349165653,83.9543165958867,83.9543165959867,83.96319824216141,83.96319824226141,83.96763711465701,83.96763711475701,83.972072952821,83.972072952921,83.97650662360525,83.97650662370525,83.9898056656128,83.98980566571281,83.99866733258713,83.99866733268713,84.00309603810146,84.00309603820146,84.01195242455063,84.01195242465063,84.02080557963365,84.02080557973365,84.02523050208518,84.02523050218518,84.03407849486376,84.03407849496377,84.04292412322809,84.04292412332809,84.051766638447,84.05176663854701,84.05618509816627,84.05618509826627,84.06060288796814,84.06060288806815,84.06501996844577,84.06501996854577,84.06943354170892,84.06943354180892,84.0738464844616,84.0738464845616,84.08708219957421,84.08708219967421,84.09149175333313,84.09149175343313,84.10030814177459,84.10030814187459,84.1047145823881,84.1047145824881,84.10911956494614,84.10911956504614,84.11352383817992,84.11352383827992,84.12233053252295,84.12233053262295,84.1267325595632,84.1267325596632,84.13993410888992,84.13993410898992,84.15312635818728,84.15312635828728,84.15752196190213,84.15752196200214,84.17509870216739,84.17509870226739,84.1838774964223,84.1838774965223,84.18826502172182,84.18826502182182,84.20142680948227,84.20142680958227,84.21019890456348,84.21019890466349,84.21458331671758,84.21458331681758,84.21896607378169,84.21896607388169,84.23211150767693,84.23211150777693,84.2408707167005,84.2408707168005,84.24524902078444,84.24524902088444,84.24962488164033,84.24962488174033,84.25837514529665,84.25837514539666,84.26274915402804,84.26274915412804,84.27149472826278,84.27149472836278,84.28023592833115,84.28023592843115,84.28897318770912,84.28897318780912,84.29770713690714,84.29770713700714,84.31080232699959,84.3108023270996,84.31516400137,84.31516400147,84.32388597086916,84.32388597096916,84.3282452020115,84.3282452021115,84.33695822614342,84.33695822624342,84.34131170387776,84.34131170397777,84.35001696484956,84.35001696494956,84.35871647241336,84.35871647251336,84.36741042360367,84.36741042370367,84.37175601995719,84.37175602005719,84.3804451240983,84.3804451241983,84.38913162738373,84.38913162748374,84.40649730427043,84.40649730437043,84.4151737193884,84.41517371948841,84.41950946401589,84.41950946411589,84.42384402643624,84.42384402653624,84.4281776824978,84.4281776825978,84.43251062923508,84.43251062933508,84.43684176325479,84.43684176335479,84.44117179388117,84.44117179398117,84.4498306335199,84.4498306336199,84.4541579450699,84.4541579451699,84.45848249813659,84.45848249823659,84.4714542658053,84.47145426590531,84.4757757845404,84.4757757846404,84.49306036201838,84.49306036211838,84.50602042583658,84.50602042593658,84.51033859498482,84.51033859508482,84.51465357217381,84.51465357227382,84.52328025577877,84.52328025587877,84.53190343216926,84.53190343226926,84.53621368052977,84.53621368062977,84.54052258905553,84.54052258915553,84.54482992130512,84.54482992140512,84.54913465269905,84.54913465279905,84.55343875358251,84.55343875368251,84.56204541848017,84.56204541858017,84.56634766723913,84.56634766733913,84.57494991856352,84.57494991866352,84.57924881773563,84.57924881783563,84.58354594359706,84.58354594369706,84.5921390131128,84.5921390132128,84.59643436566354,84.59643436576354,84.6050240067786,84.60502400687861,84.61790604492661,84.61790604502661,84.62219867840096,84.62219867850096,84.62648938093702,84.62648938103702,84.63506712116704,84.63506712126704,84.63935274021246,84.63935274031246,84.64363694060933,84.64363694070933,84.64792051049574,84.64792051059574,84.65648327610218,84.65648327620218,84.66076109258059,84.66076109268059,84.66931132679152,84.66931132689152,84.6778552953047,84.6778552954047,84.68639682058982,84.68639682068982,84.69066636161834,84.69066636171834,84.69920402502686,84.69920402512686,84.70773975749708,84.70773975759708,84.71200563368352,84.71200563378352,84.72480026731816,84.72480026741816,84.72906299095227,84.72906299105227,84.7333249658552,84.7333249659552,84.73758378820581,84.73758378830581,84.74184182241834,84.74184182251834,84.74609922612039,84.7460992262204,84.75461312716573,84.75461312726573,84.7673815158022,84.7673815159022,84.77163608220715,84.77163608230715,84.77588978166023,84.77588978176023,84.78014285060283,84.78014285070283,84.78439481615212,84.78439481625212,84.79289685571929,84.7928968558193,84.80564565912442,84.80564565922442,84.81413962027625,84.81413962037625,84.81838433495517,84.81838433505517,84.82687206981615,84.82687206991615,84.83111284380466,84.83111284390466,84.83959273669171,84.83959273679172,84.84806924058502,84.84806924068502,84.85230536455884,84.85230536465885,84.85654046395315,84.85654046405315,84.86077367181608,84.86077367191608,84.86500601272711,84.86500601282711,84.87346970937656,84.87346970947657,84.87769996172167,84.87769996182168,84.88192812550086,84.88192812560087,84.886153846052,84.886153846152,84.89037672930603,84.89037672940603,84.89459878501508,84.89459878511508,84.89881985555152,84.89881985565152,84.90726077501039,84.90726077511039,84.91991874550148,84.91991874560148,84.92835533020089,84.92835533030089,84.93257249945383,84.93257249955383,84.9367889987894,84.9367889988894,84.94100293667618,84.94100293677619,84.95363757828002,84.95363757838003,84.96204894256078,84.96204894266079,84.97045676022017,84.97045676032018,84.97886122829271,84.97886122839272,84.98306157079759,84.98306157089759,84.98725816964657,84.98725816974657,84.99984595644139,84.99984595654139,85.00403920570352,85.00403920580352,85.00822973588926,85.00822973598926,85.012417389371,85.012417389471,85.01660338776276,85.01660338786276,85.0291570087717,85.0291570088717,85.04169742846776,85.04169742856776,85.05004948535779,85.05004948545779,85.05839846850931,85.05839846860931,85.06674410207397,85.06674410217397,85.0834277636708,85.0834277637708,85.08759642302475,85.08759642312475,85.09593188960815,85.09593188970815,85.10009664767858,85.10009664777859,85.104260184135,85.104260184235,85.10842218372214,85.10842218382214,85.11258327695049,85.11258327705049,85.12090388713104,85.12090388723104,85.12506281297966,85.12506281307967,85.12921779813787,85.12921779823787,85.13337160108895,85.13337160118895,85.14167774893568,85.14167774903568,85.15413214336,85.15413214346,85.15828003527547,85.15828003537547,85.16657227248503,85.16657227258503,85.17071579023414,85.17071579033414,85.18314330914984,85.18314330924984,85.18728221629117,85.18728221639117,85.19142029588751,85.19142029598751,85.19969479999021,85.19969480009021,85.20382992406873,85.20382992416873,85.21209851713581,85.21209851723582,85.2162313950208,85.2162313951208,85.228628294772,85.228628294872,85.2327593205325,85.2327593206325,85.24102030806708,85.24102030816708,85.25340613447821,85.25340613457821,85.26990781499174,85.26990781509174,85.27402926487453,85.27402926497453,85.27814965077091,85.27814965087092,85.2822682239497,85.2822682240497,85.28638573314208,85.28638573324208,85.29050103554783,85.29050103564784,85.29461543159478,85.29461543169478,85.30284292326085,85.30284292336086,85.31106801110576,85.31106801120576,85.31929057690881,85.31929057700881,85.32339898310633,85.32339898320633,85.32750506429652,85.32750506439652,85.33982102226662,85.33982102236662,85.34802817997014,85.34802818007014,85.35213032046991,85.35213032056991,85.36033204002068,85.36033204012068,85.36853033117077,85.36853033127078,85.3726282945387,85.3726282946387,85.37672546976854,85.37672546986855,85.3808217386396,85.3808217387396,85.38491611597925,85.38491611607925,85.38900923229797,85.38900923239797,85.39310081174743,85.39310081184743,85.40128211852185,85.40128211862185,85.40946094266131,85.40946094276131,85.41354881786178,85.41354881796178,85.41763602314488,85.41763602324488,85.4217225191037,85.4217225192037,85.42989421059352,85.42989421069352,85.44214861497936,85.44214861507936,85.4462314066892,85.4462314067892,85.450312976785,85.450312976885,85.45439367992891,85.45439368002891,85.46663397664307,85.46663397674307,85.478870253853,85.478870253953,85.48294756800317,85.48294756810317,85.48702354231858,85.48702354241858,85.4910985708683,85.4910985709683,85.4951722989902,85.4951722990902,85.49924299278045,85.49924299288045,85.50331203148075,85.50331203158075,85.50737953331178,85.50737953341178,85.5114460499702,85.5114460500702,85.51551150264223,85.51551150274223,85.5277062843821,85.52770628448211,85.5358339189531,85.5358339190531,85.5398963767004,85.5398963768004,85.54801900659456,85.54801900669456,85.55207839060334,85.55207839070334,85.56425358711186,85.56425358721187,85.56830954271999,85.56830954281999,85.58858581354612,85.58858581364612,85.59669110440245,85.59669110450245,85.60884663686582,85.60884663696582,85.61694877516983,85.61694877526983,85.6209985044871,85.6209985045871,85.62909548048667,85.62909548058667,85.6331400474995,85.6331400475995,85.64527232988942,85.64527232998942,85.64931441426728,85.64931441436728,85.65739700674685,85.65739700684685,85.66143735722093,85.66143735732093,85.67355718702915,85.67355718712915,85.67759540953041,85.67759540963041,85.68163256804525,85.68163256814525,85.6897055058333,85.6897055059333,85.6977760792071,85.6977760793071,85.70180831185893,85.70180831195893,85.70583447584752,85.70583447594753,85.71388554280378,85.71388554290378,85.71791020933001,85.71791020943002,85.72193361483532,85.72193361493532,85.73400257033029,85.73400257043029,85.73802463600086,85.73802463610086,85.74204603175406,85.74204603185406,85.74606281689947,85.74606281699947,85.75007778932728,85.75007778942728,85.76212093330005,85.76212093340006,85.76613358072052,85.76613358082052,85.7701452035615,85.7701452036615,85.77415260986372,85.77415260996372,85.78216600381961,85.78216600391961,85.78617163681116,85.78617163691116,85.79418053837999,85.79418053847999,85.79818254593634,85.79818254603634,85.80618466951766,85.80618466961766,85.8101845096943,85.8101845097943,85.81418328588451,85.81418328598451,85.82617748648235,85.82617748658235,85.83017385885141,85.83017385895141,85.83416869435122,85.83416869445122,85.84615150635376,85.84615150645377,85.85014228294244,85.85014228304244,85.85812241747125,85.85812241757125,85.86211075083187,85.86211075093188,85.86609829605442,85.86609829615442,85.87008485610434,85.87008485620434,85.87407015513335,85.87407015523335,85.88203673368714,85.88203673378715,85.88601612168053,85.88601612178053,85.89397340020494,85.89397340030494,85.8979506602255,85.8979506603255,85.90192654100441,85.90192654110442,85.90590127898311,85.90590127908311,85.91384961214028,85.91384961224028,85.92179573851082,85.92179573861083,85.93371163759015,85.93371163769015,85.94165228640101,85.94165228650101,85.94562000995077,85.94562001005077,85.9535536049258,85.9535536050258,85.95751817592323,85.95751817602323,85.96544530816598,85.96544530826598,85.96940373168636,85.96940373178636,85.97336120944105,85.97336120954105,85.97731380073962,85.97731380083962,85.9812645399137,85.9812645400137,85.9852145303566,85.9852145304566,85.98916144706098,85.98916144716098,85.9931069451168,85.99310694521681,86.00099506452446,86.00099506462446,86.00493638544845,86.00493638554845,86.00887707586197,86.00887707596198,86.02069713735042,86.02069713745043,86.02857367112827,86.02857367122827,86.03250947508567,86.03250947518568,86.03644397861525,86.03644397871525,86.04037603891678,86.04037603901678,86.04823885909197,86.04823885919197,86.06003149337515,86.06003149347515,86.06789159447398,86.06789159457398,86.07182054163006,86.07182054173006,86.07574783369618,86.07574783379619,86.07967394355518,86.07967394365518,86.08752403530033,86.08752403540034,86.09144584980677,86.09144584990678,86.09536514227136,86.09536514237136,86.09928368600475,86.09928368610476,86.10711903956778,86.10711903966778,86.11103186930008,86.11103186940008,86.11494383208048,86.11494383218049,86.11885291815689,86.11885291825689,86.12276129490903,86.12276129500903,86.12666462757743,86.12666462767743,86.13447018952093,86.13447018962093,86.14227437222277,86.14227437232277,86.14617470996646,86.14617471006646,86.15396994730106,86.15396994740107,86.1578630735813,86.1578630736813,86.16175525409584,86.16175525419584,86.16953736893136,86.16953736903136,86.17342505705882,86.17342505715882,86.17731116891012,86.17731116901012,86.18508106760537,86.18508106770537,86.18896442097338,86.18896442107338,86.19284643450665,86.19284643460665,86.19672762049493,86.19672762059493,86.20448794331247,86.20448794341247,86.21224468010173,86.21224468020174,86.21612145251675,86.21612145261675,86.21999735797988,86.21999735807988,86.22387204182897,86.22387204192897,86.23161971503029,86.2316197151303,86.23549164039609,86.2354916404961,86.2393625411824,86.2393625412824,86.24710213596836,86.24710213606836,86.254838578202,86.254838578302,86.2625698975381,86.2625698976381,86.27029483295574,86.27029483305574,86.27415438455364,86.27415438465364,86.28186875892096,86.28186875902097,86.28957769513552,86.28957769523552,86.29343074459425,86.29343074469425,86.29728253303205,86.29728253313205,86.30113242993843,86.30113243003844,86.30498165692747,86.30498165702747,86.31267739182913,86.31267739192913,86.31652342685892,86.31652342695892,86.32036760976422,86.32036760986422,86.32805447811245,86.32805447821245,86.33189621778969,86.33189621788969,86.33573728754956,86.33573728764956,86.33957772679896,86.33957772689897,86.35109506444986,86.35109506454987,86.36260597877538,86.36260597887538,86.36644030995464,86.36644031005464,86.37027349833369,86.37027349843369,86.37793699838777,86.37793699848777,86.38176667955234,86.38176667965234,86.38942462323294,86.38942462333294,86.39325205820398,86.39325205830399,86.39707858681622,86.39707858691622,86.40473042242668,86.40473042252668,86.40855521713515,86.40855521723515,86.41237819912602,86.41237819922603,86.41619889551646,86.41619889561646,86.42001809444453,86.42001809454453,86.42765361559668,86.42765361569668,86.43146599713033,86.43146599723033,86.43908965680434,86.43908965690434,86.44289983155137,86.44289983165137,86.44670736603581,86.44670736613581,86.45051285136125,86.45051285146126,86.45812102412194,86.45812102422194,86.4619229234191,86.4619229235191,86.46572375872984,86.46572375882984,86.47332286790257,86.47332286800257,86.47711917141933,86.47711917151933,86.484710911501,86.484710911601,86.48850520526565,86.48850520536566,86.496092137705,86.49609213780501,86.49988379120708,86.49988379130708,86.50746433972796,86.50746433982796,86.51504082933772,86.51504082943772,86.52639975123398,86.52639975133398,86.53396682259365,86.53396682269366,86.54153129309765,86.54153129319765,86.54531207029419,86.54531207039419,86.55286940814852,86.55286940824853,86.56419964181859,86.56419964191859,86.56797285288954,86.56797285298954,86.57929106745384,86.57929106755384,86.58306258402791,86.58306258412792,86.58683339127771,86.58683339137771,86.59059974554734,86.59059974564734,86.59812855280309,86.59812855290309,86.60188923247851,86.60188923257851,86.61316906471819,86.6131690648182,86.61692777404842,86.61692777414842,86.62444436516388,86.62444436526388,86.62819960668655,86.62819960678655,86.63195421769875,86.63195421779875,86.64321631683156,86.64321631693156,86.64696769647762,86.64696769657762,86.65446887949356,86.65446887959357,86.65821671251825,86.65821671261826,86.66196344214961,86.66196344224961,86.66570658575272,86.66570658585272,86.67693349452013,86.67693349462013,86.68067297328113,86.68067297338114,86.68815023630629,86.68815023640629,86.69562584424145,86.69562584434145,86.69936256451916,86.69936256461916,86.70309865428642,86.70309865438642,86.71430412569799,86.714304125798,86.71803714172671,86.71803714182671,86.73296751134477,86.73296751144477,86.73669847821448,86.73669847831448,86.74042708066995,86.74042708076995,86.74415493439423,86.74415493449423,86.75160914438045,86.75160914448045,86.75533384555241,86.75533384565242,86.76650574228167,86.76650574238167,86.77395092808683,86.77395092818684,86.78883460052349,86.78883460062349,86.79998829126302,86.79998829136302,86.8037045987644,86.8037045988644,86.81113614978074,86.81113614988074,86.81856643977615,86.81856643987615,86.82599464120564,86.82599464130564,86.83342075406921,86.83342075416921,86.8371304412107,86.8371304413107,86.84454760870703,86.84454760880703,86.84825004497816,86.84825004507816,86.85935636861892,86.85935636871892,86.86675824623644,86.86675824633645,86.87045701766549,86.87045701776549,86.8741546068874,86.8741546069874,86.87784770372224,86.87784770382224,86.88153989419828,86.88153989429829,86.88523090246721,86.88523090256722,86.89630160226663,86.89630160236663,86.90736753383067,86.90736753393067,86.91105278869158,86.91105278879158,86.91473575795206,86.91473575805206,86.91841809670207,86.91841809680207,86.922099647314,86.922099647414,86.92578009453261,86.92578009463261,86.93314035845938,86.93314035855938,86.94049865204093,86.94049865214093,86.94417271534108,86.94417271544108,86.95151887159616,86.95151887169617,86.95886163885748,86.95886163895749,86.96619865271082,86.96619865281082,86.973529558494,86.973529558594,86.97719325777835,86.97719325787835,86.98085628714534,86.98085628724535,86.98817852340962,86.98817852350962,87.00281503574352,87.00281503584353,87.01744567644872,87.01744567654872,87.02110023333131,87.02110023343131,87.0247537262275,87.0247537263275,87.02840654920631,87.02840654930631,87.03571085532921,87.03571085542922,87.04666467425096,87.04666467435096,87.0576145524823,87.05761455258231,87.06126315892239,87.06126315902239,87.06490944035511,87.06490944045511,87.07584682659785,87.07584682669786,87.07948885208494,87.07948885218494,87.08312949833036,87.08312949843037,87.08676766194085,87.08676766204086,87.09768057649612,87.09768057659612,87.10495343590951,87.10495343600951,87.11222377328103,87.11222377338103,87.11948997292762,87.11948997302763,87.12311919117087,87.12311919127087,87.12674777890363,87.12674777900364,87.1303743962912,87.1303743963912,87.13762570012803,87.13762570022803,87.14124932259088,87.14124932269088,87.14487215691565,87.14487215701566,87.14849432132306,87.14849432142306,87.15573675860645,87.15573675870645,87.15935584927533,87.15935584937533,87.17021083568152,87.17021083578152,87.17744499751504,87.17744499761504,87.18106121147991,87.18106121157992,87.184676519086,87.184676519186,87.18829064448495,87.18829064458495,87.19190315420084,87.19190315430085,87.19551424526817,87.19551424536817,87.19912470582504,87.19912470592504,87.20273351129194,87.20273351139194,87.20994997942552,87.20994997952552,87.2243802754301,87.2243802755301,87.23159221176968,87.23159221186968,87.24601332596559,87.2460133260656,87.26403410322663,87.26403410332664,87.26763684003029,87.2676368401303,87.27484077676836,87.27484077686836,87.27844122797157,87.27844122807157,87.28204096985053,87.28204096995053,87.28564000240519,87.28564000250519,87.28923820741488,87.28923820751488,87.29283523021743,87.29283523031744,87.30002691140831,87.30002691150831,87.31080733995188,87.31080734005188,87.3143954173872,87.3143954174872,87.32156995657478,87.32156995667478,87.32874248601023,87.32874248611023,87.33232681978966,87.33232681988966,87.3359102078034,87.3359102079034,87.346658559127,87.346658559227,87.35023914925053,87.35023914935053,87.36097922512425,87.36097922522426,87.36455792371639,87.36455792381639,87.37529189151998,87.37529189161998,87.37886731933908,87.37886731943908,87.38601494361113,87.38601494371113,87.38958560319482,87.38958560329482,87.40386512838417,87.40386512848417,87.4074322807534,87.4074322808534,87.41456532447091,87.41456532457092,87.4288269983327,87.4288269984327,87.43595109668297,87.43595109678297,87.43951084055422,87.43951084065422,87.44662914608959,87.4466291461896,87.4537440232243,87.4537440233243,87.4679669206924,87.4679669207924,87.47507572916389,87.47507572926389,87.47862794631645,87.47862794641645,87.48573048909016,87.48573048919016,87.4892806176768,87.4892806177768,87.49282964287013,87.49282964297014,87.49637768289085,87.49637768299085,87.49992446189063,87.49992446199063,87.5070169953107,87.50701699541071,87.51056018828221,87.51056018838221,87.51764389455576,87.51764389465576,87.52118389556803,87.52118389566803,87.52826173021286,87.52826173031286,87.53179897274185,87.53179897284186,87.53533475721538,87.53533475731538,87.53886967473703,87.53886967483703,87.54593876104913,87.54593876114913,87.54947174763247,87.54947174773247,87.5530036308225,87.5530036309225,87.55653224323947,87.55653224333948,87.56005975226314,87.56005975236314,87.5635852515347,87.5635852516347,87.56710937156461,87.56710937166461,87.57415430144448,87.57415430154448,87.58823635863722,87.58823635873722,87.59175519814197,87.59175519824197,87.59527198848771,87.59527198858771,87.6093346968905,87.6093346969905,87.61284715247679,87.61284715257679,87.61987068440772,87.61987068450772,87.62338101202118,87.62338101212119,87.62689055149656,87.62689055159656,87.6339083694264,87.6339083695264,87.6374153868599,87.6374153869599,87.64091980343775,87.64091980353776,87.65143127986059,87.65143127996059,87.66194133763489,87.66194133773489,87.66894536314827,87.66894536324827,87.6724450903045,87.67244509040451,87.67594383228816,87.67594383238816,87.68293840014455,87.68293840024455,87.68643331965849,87.68643331975849,87.68992717518603,87.68992717528603,87.69342028198238,87.69342028208239,87.70040472226441,87.70040472236441,87.70389554346032,87.70389554356032,87.70738498541459,87.7073849855146,87.71436197779174,87.71436197789174,87.71784854304201,87.71784854314201,87.72482104303208,87.72482104313208,87.72830559853024,87.72830559863024,87.73178928707651,87.73178928717651,87.74223873703224,87.74223873713224,87.74572073108163,87.74572073118163,87.7526827488352,87.7526827489352,87.75616233906344,87.75616233916344,87.7596411805605,87.7596411806605,87.76311824874688,87.76311824884688,87.76659330718114,87.76659330728114,87.77006698637375,87.77006698647375,87.77701304433114,87.77701304443114,87.78395693490882,87.78395693500882,87.78742612171436,87.78742612181436,87.79089448097493,87.79089448107493,87.79782918974394,87.79782918984394,87.80129419941764,87.80129419951764,87.80475846036016,87.80475846046016,87.8116857212243,87.8116857213243,87.81514868173899,87.81514868183899,87.82207070148486,87.82207070158486,87.82898787418152,87.82898787428152,87.83936071711562,87.83936071721563,87.8462742643771,87.8462742644771,87.85318572307267,87.85318572317267,87.86009426565734,87.86009426575734,87.86354701978385,87.86354701988385,87.86699882814467,87.86699882824468,87.87390086859016,87.87390086869016,87.88080105691114,87.88080105701114,87.88424990975415,87.88424990985415,87.89114584212949,87.89114584222949,87.90148589851934,87.90148589861934,87.90493128355479,87.90493128365479,87.90837603807977,87.90837603817977,87.91181787649384,87.91181787659384,87.92902454652236,87.92902454662236,87.93934470242563,87.93934470252563,87.94622325635623,87.94622325645624,87.95309889417595,87.95309889427595,87.95997200995379,87.95997201005379,87.96340592758014,87.96340592768014,87.97027214714976,87.97027214724976,87.98056763433104,87.98056763443104,87.98399694134959,87.98399694144959,87.9908544519934,87.9908544520934,87.99428218273579,87.99428218283579,88.00113543743396,88.00113543753396,88.00456076435519,88.00456076445519,88.00798534254525,88.00798534264526,88.01140917200415,88.01140917210415,88.0216778230837,88.02167782318371,88.02851984681419,88.0285198469142,88.03193839574791,88.03193839584792,88.03535481670883,88.03535481680883,88.04218635820281,88.04218635830281,88.04901309205448,88.04901309215448,88.05242431130405,88.05242431140405,88.06947760966166,88.06947760976166,88.07288520347605,88.07288520357605,88.0762914968626,88.0762914969626,88.08310305905619,88.08310305915619,88.0899121780217,88.08991217812171,88.0933133091038,88.09331330920381,88.09671369145474,88.09671369155474,88.10011143354309,88.1001114336431,88.10350830867954,88.10350830877954,88.11369637264015,88.11369637274015,88.12048306907722,88.12048306917723,88.12387292978474,88.12387292988474,88.13065064144767,88.13065064154767,88.1374255158135,88.1374255159135,88.14419660711654,88.14419660721654,88.15773086893489,88.15773086903489,88.16111111435781,88.16111111445781,88.16786975307913,88.16786975317913,88.17462622442073,88.17462622452074,88.1780031596637,88.1780031597637,88.18137934617549,88.18137934627549,88.19488251594647,88.19488251604648,88.19825688974066,88.19825688984066,88.20162834742395,88.20162834752395,88.20499811061036,88.20499811071036,88.2083671644725,88.20836716457251,88.21510389295514,88.21510389305514,88.22183833583733,88.22183833593733,88.22520439477475,88.22520439487475,88.22856911387744,88.22856911397744,88.23192953762756,88.23192953772757,88.23864936054832,88.23864936064832,88.24200757751183,88.24200757761183,88.24872290804551,88.24872290814551,88.25207864237404,88.25207864247405,88.26214359916614,88.26214359926614,88.268851048319,88.268851048419,88.27555589661618,88.27555589671618,88.27890619279195,88.27890619289195,88.28560276563927,88.28560276573927,88.28894742662774,88.28894742672774,88.29229070837458,88.29229070847458,88.29563016765171,88.29563016775171,88.29896797183888,88.29896797193888,88.30230486966724,88.30230486976724,88.30564090054371,88.30564090064371,88.30897433056451,88.30897433066451,88.31230689363342,88.31230689373342,88.32230186376376,88.32230186386376,88.33229060760324,88.33229060770324,88.33894399048363,88.33894399058363,88.34226685945413,88.34226685955413,88.35555616795637,88.35555616805637,88.36219528553745,88.36219528563745,88.36551129770662,88.36551129780662,88.36882656114462,88.36882656124462,88.37545535411682,88.37545535421683,88.37876864720961,88.37876864730961,88.38208079750217,88.38208079760217,88.38539034693906,88.38539034703906,88.38869768958932,88.38869768968932,88.39861806245014,88.39861806255014,88.40853496750339,88.40853496760339,88.41844600092793,88.41844600102793,88.42505063746789,88.42505063756789,88.43165357951098,88.43165357961098,88.43495422298754,88.43495422308754,88.44155212094691,88.44155212104691,88.45144337210556,88.45144337220556,88.45803488614646,88.45803488624647,88.4646239963662,88.4646239964662,88.46791624617218,88.46791624627218,88.47120719555032,88.47120719565032,88.48107819156024,88.48107819166025,88.49094564094881,88.49094564104881,88.49423060047752,88.49423060057752,88.5007990614795,88.5007990615795,88.50408145955944,88.50408145965945,88.51392711693002,88.51392711703002,88.51720391922957,88.51720391932957,88.52047942110129,88.52047942120129,88.53030371992982,88.53030372002982,88.54012423569556,88.54012423579556,88.54666901314815,88.54666901324815,88.5532114261865,88.5532114262865,88.55648121405711,88.55648121415712,88.5662898289378,88.5662898290378,88.57282609449909,88.5728260945991,88.57936015327375,88.57936015337376,88.5858921234825,88.5858921235825,88.59242015300083,88.59242015310083,88.59894424182875,88.59894424192875,88.6022048478907,88.6022048479907,88.6054629713177,88.60546297141771,88.61197626265388,88.61197626275388,88.61523103649404,88.61523103659404,88.62824871320606,88.62824871330606,88.63800862115323,88.63800862125323,88.64451103618389,88.64451103628389,88.65101085035886,88.65101085045886,88.66075743877244,88.66075743887244,88.67050182039938,88.67050182049938,88.67699560531801,88.67699560541801,88.68023981810785,88.68023981820785,88.68348249402842,88.68348249412843,88.68996638781412,88.68996638791413,88.69644831125461,88.69644831135462,88.69968614012599,88.69968614022599,88.70616116735826,88.70616116745826,88.70939741995346,88.70939742005346,88.7158683488677,88.7158683489677,88.72557107739698,88.72557107749698,88.72880398040532,88.72880398050532,88.73850111315421,88.73850111325422,88.74496337254953,88.74496337264954,88.74819205901915,88.74819205911915,88.75141976031615,88.75141976041616,88.75464513660582,88.75464513670582,88.7578698035712,88.7578698036712,88.76109281544662,88.76109281554662,88.76431440867347,88.76431440877347,88.76753537138987,88.76753537148987,88.77397639046387,88.77397639056387,88.77719573749721,88.77719573759721,88.78363348579816,88.78363348589816,88.78684971968606,88.78684971978606,88.7900653230635,88.7900653231635,88.79970913827108,88.79970913837109,88.80292084036502,88.80292084046502,88.80613163610015,88.80613163620015,88.81255196654948,88.81255196664948,88.81896922326028,88.81896922336028,88.8253837214878,88.8253837215878,88.83179597352178,88.83179597362178,88.84141086406174,88.84141086416174,88.84781838726722,88.84781838736723,88.85102082873868,88.85102082883868,88.86062716798044,88.86062716808044,88.8670296865091,88.8670296866091,88.87022984238011,88.87022984248011,88.87662700156982,88.87662700166982,88.88302061413813,88.88302061423813,88.88941300509244,88.88941300519244,88.89260784103139,88.8926078411314,88.89899577900552,88.89899577910552,88.90218781705428,88.90218781715429,88.90857067153777,88.90857067163778,88.91495159508297,88.91495159518297,88.92452002488295,88.92452002498295,88.94046082692994,88.94046082702994,88.95002113890766,88.95002113900766,88.95957644620856,88.95957644630856,88.96594231631639,88.96594231641639,88.9723063737066,88.9723063738066,88.97866798786877,88.97866798796878,88.98184753392893,88.98184753402893,88.98820560146972,88.98820560156972,88.9913806945497,88.9913806946497,88.99455472364328,88.99455472374328,88.99772583662595,88.99772583672595,89.00089627969123,89.00089627979123,89.0072349590352,89.0072349591352,89.01040268302411,89.01040268312411,89.01356855488852,89.01356855498852,89.01673221996631,89.01673222006632,89.01989375707127,89.01989375717127,89.02305411196912,89.02305411206912,89.03569442816715,89.03569442826715,89.04517076403216,89.04517076413217,89.05464288335844,89.05464288345844,89.05779657848947,89.05779657858947,89.0609482244615,89.0609482245615,89.07040158610141,89.07040158620141,89.07354645408593,89.07354645418593,89.07983090952979,89.07983090962979,89.08296994529249,89.08296994539249,89.09552195061836,89.09552195071836,89.09865629695948,89.09865629705948,89.10805724741688,89.10805724751688,89.11432180237416,89.11432180247417,89.11745220802486,89.11745220812486,89.12684121819031,89.12684121829031,89.13309797058058,89.13309797068058,89.13622514486512,89.13622514496512,89.14873163521672,89.14873163531672,89.15185538110062,89.15185538120062,89.158101808882,89.158101808982,89.16122255984118,89.16122255994118,89.16746339184218,89.16746339194218,89.1705826059321,89.1705826060321,89.17681965487029,89.17681965497029,89.17993713505643,89.17993713515644,89.18616984923517,89.18616984933517,89.19239909560635,89.19239909570635,89.21107820460787,89.21107820470787,89.2172964170459,89.2172964171459,89.22351194981444,89.22351194991444,89.2266181990329,89.2266181991329,89.23593556744663,89.23593556754663,89.24214499214503,89.24214499224503,89.25145664655767,89.25145664665767,89.25455808813382,89.25455808823382,89.26075907975473,89.26075907985474,89.26385472851598,89.26385472861598,89.26694939210462,89.26694939220462,89.27932643077614,89.27932643087614,89.28241932105409,89.2824193211541,89.2886041164374,89.2886041165374,89.29478698088242,89.29478698098242,89.29787368427642,89.29787368437643,89.30404594826422,89.30404594836422,89.30713087834755,89.30713087844755,89.31021458681684,89.31021458691684,89.31638105798973,89.31638105808973,89.31946322958976,89.31946322968976,89.32254465245862,89.32254465255862,89.3287034392852,89.32870343938521,89.33177891171154,89.33177891181154,89.33485312311693,89.33485312321693,89.3440726441877,89.3440726442877,89.34714323015791,89.34714323025791,89.3532829834498,89.3532829835498,89.36249076126313,89.36249076136313,89.36555839171554,89.36555839181554,89.37475619958214,89.37475619968214,89.38700780607778,89.38700780617778,89.39313128431826,89.39313128441826,89.4023133688301,89.4023133689301,89.4084294385726,89.4084294386726,89.4114846952571,89.4114846953571,89.41759343531544,89.41759343541544,89.4267527426367,89.4267527427367,89.43590645417756,89.43590645427756,89.43895410532956,89.43895410542956,89.44809481259206,89.44809481269206,89.45418522843997,89.45418522853997,89.45722897830846,89.45722897840847,89.46635683892022,89.46635683902022,89.47243669371782,89.47243669381783,89.48459206855357,89.48459206865357,89.48762967094503,89.48762967104503,89.49370369352081,89.49370369362082,89.50281248119096,89.50281248129096,89.51191815571568,89.51191815581568,89.51798500623492,89.51798500633492,89.52101504250078,89.52101504260078,89.52707416926681,89.52707416936681,89.53615935220138,89.53615935230138,89.54221217386274,89.54221217396274,89.54523515629276,89.54523515639276,89.54825561668092,89.54825561678092,89.55429456711204,89.55429456721204,89.56033107431509,89.56033107441509,89.56938284019495,89.56938284029495,89.57541517026615,89.57541517036616,89.58144568761979,89.58144568771979,89.58747265835204,89.58747265845204,89.59650688815957,89.59650688825957,89.60854762831117,89.60854762841117,89.61155622722119,89.61155622732119,89.61456336807575,89.61456336817575,89.62057642817085,89.62057642827085,89.62658669037575,89.62658669047575,89.63259356358688,89.63259356368688,89.63559499044034,89.63559499054034,89.6385953533074,89.6385953534074,89.64159504625707,89.64159504635707,89.6505911301814,89.6505911302814,89.65658543259012,89.65658543269012,89.66557343809909,89.66557343819909,89.66856761408219,89.6685676141822,89.6715599773477,89.6715599774477,89.67455155247512,89.67455155257512,89.67754233946447,89.67754233956447,89.6805320230605,89.6805320231605,89.68651005041781,89.68651005051781,89.6954753181431,89.6954753182431,89.6984629131732,89.6984629132732,89.70144924718238,89.70144924728238,89.70742093002814,89.70742093012814,89.71040407207808,89.71040407217808,89.71636724303254,89.71636724313254,89.71934715371634,89.71934715381634,89.72828345736708,89.72828345746709,89.73125923032595,89.73125923042595,89.73423417573983,89.73423417583983,89.74018284495357,89.74018284505357,89.74315605646366,89.74315605656366,89.74612694296643,89.74612694306643,89.7580090703289,89.7580090704289,89.76394619331973,89.76394619341973,89.76691203573874,89.76691203583874,89.76987508026758,89.76987508036758,89.77876188884673,89.77876188894673,89.78468293382066,89.78468293392066,89.79060133853203,89.79060133863203,89.79355823558382,89.79355823568382,89.80242671995255,89.80242672005255,89.8083364945519,89.8083364946519,89.81424418058533,89.81424418068534,89.82014906872857,89.82014906882857,89.82309879372382,89.82309879382382,89.83784501487884,89.83784501497884,89.8437405635858,89.8437405636858,89.84668697840108,89.84668697850108,89.84962941311905,89.84962941321905,89.86433926170156,89.86433926180156,89.86727866208791,89.86727866218791,89.8731545073428,89.8731545074428,89.87608996703874,89.87608996713874,89.87902455978278,89.87902455988278,89.88489272069137,89.88489272079137,89.88782506724188,89.88782506734188,89.8907564680267,89.8907564681267,89.89661765391325,89.89661765401326,89.9024768694546,89.9024768695546,89.9054043689559,89.9054043690559,89.91125810693757,89.91125810703757,89.91710774660122,89.91710774670122,89.92295234218113,89.92295234228114,89.93171608299866,89.93171608309866,89.93463601637438,89.93463601647439,89.94047513439462,89.94047513449462,89.94339278216988,89.94339278226988,89.95506156055337,89.95506156065338,89.95797755323866,89.95797755333867,89.96089275778587,89.96089275788587,89.96380555851194,89.96380555861194,89.96962958368789,89.96962958378789,89.97545061393913,89.97545061403913,89.98417711523227,89.98417711533227,89.98999180097194,89.98999180107194,89.99289514404101,89.99289514414102,89.99870060856512,89.99870060866512,90.00450410274402,90.00450410284402,90.01900864135598,90.01900864145598,90.02190828017606,90.02190828027607,90.02480523932665,90.02480523942666,90.0392876706654,90.0392876707654,90.04507835760045,90.04507835770045,90.05376076256786,90.05376076266786,90.0566533475521,90.0566533476521,90.05954506558444,90.05954506568445,90.06243244885734,90.06243244895734,90.0739762285409,90.0739762286409,90.08262174864603,90.08262174874604,90.09414089901452,90.09414089911452,90.09989619854967,90.09989619864967,90.10277183856513,90.10277183866513,90.10851748340876,90.10851748350876,90.11138658187814,90.11138658197814,90.11999210397556,90.11999210407556,90.12285623717501,90.12285623727502,90.12858324255299,90.128583242653,90.13430815936505,90.13430815946505,90.13716945526741,90.13716945536741,90.14002960836955,90.14002960846955,90.14288846104384,90.14288846114384,90.14860470833699,90.14860470843699,90.1600335380812,90.1600335381812,90.16860075666781,90.16860075676782,90.17145452585147,90.17145452595148,90.1743070340142,90.17430703411421,90.18000913422877,90.18000913432877,90.18285750466657,90.18285750476657,90.18855302392814,90.18855302402814,90.19424570589261,90.19424570599261,90.19708997801237,90.19708997811237,90.20277682775505,90.20277682785505,90.20561810495009,90.20561810505009,90.2141404390729,90.21414043917291,90.21697919422608,90.21697919432609,90.22549384400259,90.22549384410259,90.2283299588932,90.2283299589932,90.23116433988004,90.23116433998004,90.23966598537817,90.23966598547817,90.24816514824134,90.24816514834134,90.25382835384153,90.25382835394153,90.25948895858603,90.25948895868603,90.26231766497867,90.26231766507867,90.27080035575591,90.27080035585591,90.27927551981448,90.27927551991448,90.28209823635768,90.28209823645768,90.2905644156421,90.2905644157421,90.29620586863119,90.29620586873119,90.30184472076462,90.30184472086462,90.31030029959182,90.31030029969182,90.3187509919629,90.3187509920629,90.32156673348408,90.32156673358408,90.33001183007478,90.33001183017478,90.33563799318507,90.33563799328508,90.33844983342274,90.33844983352274,90.34125950628068,90.34125950638068,90.34687766978944,90.34687766988944,90.35249378413917,90.35249378423917,90.3553005802931,90.3553005803931,90.36091299039386,90.36091299049386,90.3665235089632,90.3665235090632,90.374935267313,90.37493526741301,90.37773757107988,90.37773757117988,90.38334099640647,90.38334099650648,90.39174455812022,90.39174455822022,90.39454477332117,90.39454477342117,90.39734431860474,90.39734431870474,90.40014319397095,90.40014319407095,90.40853698277246,90.40853698287246,90.4141282707726,90.4141282708726,90.41971664266183,90.41971664276183,90.42530111326754,90.42530111336754,90.42809127970793,90.42809127980793,90.43087904232718,90.43087904242718,90.43645157264093,90.43645157274094,90.43923563101119,90.43923563111119,90.44201665504983,90.44201665514983,90.44757752091996,90.44757752101997,90.45313614059657,90.45313614069657,90.45591460318643,90.45591460328643,90.45869208060368,90.45869208070368,90.4642450256861,90.4642450257861,90.46701982343387,90.46701982353387,90.47256689688757,90.47256689698757,90.47533905437277,90.47533905447277,90.4808807684875,90.4808807685875,90.48364997045489,90.4836499705549,90.4919541873633,90.4919541874633,90.49471980330243,90.49471980340243,90.49748337008252,90.49748337018252,90.50300806041467,90.50300806051467,90.50576855345625,90.50576855355625,90.50852837658046,90.50852837668046,90.51680567857335,90.51680567867335,90.51956191566927,90.51956191576927,90.52231697055808,90.52231697065808,90.52507139493643,90.52507139503643,90.5333332494229,90.5333332495229,90.5415925818675,90.5415925819675,90.55260054587555,90.55260054597555,90.5581015132514,90.5581015133514,90.5608506571046,90.5608506572046,90.5635991310404,90.5635991311404,90.5663455952241,90.56634559532411,90.56909040431785,90.56909040441785,90.57457820978772,90.57457820988772,90.580063374995,90.58006337509501,90.58828795055017,90.58828795065017,90.59376945091539,90.59376945101539,90.5965092553323,90.5965092554323,90.5992481533904,90.59924815349041,90.60472401856832,90.60472401866832,90.61567031077139,90.6156703108714,90.61840507110458,90.61840507120458,90.62113857041682,90.62113857051682,90.62660478090322,90.62660478100322,90.62933701919454,90.62933701929454,90.63480027416313,90.63480027426313,90.64026140115891,90.64026140125891,90.64299115681526,90.64299115691526,90.65390651459857,90.65390651469858,90.66208447178614,90.66208447188615,90.66480831640688,90.66480831650688,90.66753125466882,90.66753125476882,90.67025324716505,90.67025324726505,90.67841678142571,90.67841678152571,90.68385481597566,90.68385481607567,90.69200743452387,90.69200743462387,90.70015690051976,90.70015690061976,90.70558602910937,90.70558602920937,90.70829828810028,90.70829828820028,90.71643372523829,90.71643372533829,90.7191440532909,90.7191440533909,90.72456285727164,90.72456285737164,90.73539719446006,90.73539719456006,90.74081237300561,90.74081237310561,90.74351789341593,90.74351789351593,90.74622128585341,90.74622128595341,90.75433016273803,90.75433016283803,90.76243616291865,90.76243616301865,90.77053904995384,90.77053905005384,90.77323830466639,90.7732383047664,90.78133405905193,90.78133405915193,90.78403142223308,90.78403142233309,90.78942520282969,90.78942520292969,90.79751232710323,90.79751232720324,90.80020519789731,90.80020519799731,90.80289731996021,90.80289732006021,90.80828006662367,90.80828006672367,90.81365993658311,90.81365993668311,90.82172526883866,90.82172526893866,90.82441234681784,90.82441234691784,90.82709812436917,90.82709812446917,90.83246727565069,90.83246727575069,90.83783406251797,90.83783406261797,90.84319746039148,90.84319746049148,90.85391877857884,90.85391877867885,90.86731365817727,90.86731365827727,90.869988992899,90.869988992999,90.87266342126195,90.87266342136195,90.8753359975004,90.8753359976004,90.87800672161436,90.87800672171436,90.88334529313828,90.88334529323828,90.88601109138921,90.88601109148921,90.89134189975302,90.89134189985302,90.89400600350707,90.89400600360707,90.90199736900357,90.90199736910357,90.90732273921459,90.9073227393146,90.90998341456799,90.90998341466799,90.91264263186595,90.91264263196595,90.91796008128924,90.91796008138924,90.9206165795108,90.92061657961081,90.92327185611835,90.92327185621835,90.92592587170495,90.92592587180495,90.92857921737418,90.92857921747418,90.93919118140256,90.93919118150257,90.94184306901634,90.94184306911635,90.94714597729201,90.94714597739201,90.95244659996725,90.95244660006725,90.95774312432445,90.95774312442445,90.96038925853023,90.96038925863023,90.96568010829323,90.96568010839323,90.96832439037452,90.96832439047452,90.97361141766781,90.97361141776781,90.97625392643842,90.97625392653842,90.97889572588474,90.97889572598474,90.98417826079097,90.98417826089097,90.98681872040255,90.98681872050255,90.98945851009677,90.98945851019677,90.99473651320902,90.99473651330902,91.00001069385158,91.00001069395158,91.0079185167113,91.0079185168113,91.0158223988806,91.0158223989806,91.0184525732902,91.0184525733902,91.02897102473509,91.02897102483509,91.03159899235806,91.03159899245806,91.03685358776926,91.03685358786926,91.04473173723017,91.04473173733017,91.047355645942,91.047355646042,91.05260204471715,91.05260204481715,91.05522354960785,91.05522354970785,91.06046486489235,91.06046486499235,91.06308380833427,91.06308380843427,91.07093890475626,91.07093890485626,91.07617407256372,91.07617407266372,91.08140703358455,91.08140703368456,91.08663877299135,91.08663877309135,91.09448413590802,91.09448413600802,91.09971264394869,91.0997126440487,91.1023258339826,91.1023258340826,91.11016359136677,91.11016359146677,91.1232212266796,91.1232212267796,91.13105118149676,91.13105118159676,91.14148498691155,91.14148498701155,91.14669765337675,91.14669765347675,91.14930272558841,91.14930272568841,91.15451070263202,91.15451070273203,91.16231791966548,91.16231791976548,91.17272311566788,91.17272311576788,91.17792277785472,91.17792277795472,91.18571833044456,91.18571833054456,91.18831410262685,91.18831410272685,91.19090924429868,91.19090924439868,91.20128768301315,91.20128768311315,91.2090680639449,91.2090680640449,91.21425330320483,91.21425330330483,91.22202745784573,91.22202745794573,91.22720812590477,91.22720812600477,91.22979739594787,91.22979739604787,91.23497455679245,91.23497455689245,91.24015018076777,91.24015018086777,91.24532422846691,91.24532422856691,91.24791018833008,91.24791018843008,91.25566664927102,91.25566664937102,91.25825119048562,91.25825119058563,91.26083506178287,91.26083506188287,91.26858498117772,91.26858498127773,91.27116664568834,91.27116664578834,91.27890951124735,91.27890951134735,91.28148849608849,91.28148849618849,91.28406677160534,91.28406677170534,91.28922158873526,91.28922158883526,91.29437408085785,91.29437408095785,91.2969494008569,91.2969494009569,91.30467358754335,91.30467358764335,91.31496765607608,91.31496765617608,91.32011159690046,91.32011159700046,91.32268183340888,91.32268183350888,91.33039155776152,91.33039155786152,91.3329586023107,91.3329586024107,91.33552426761823,91.33552426771823,91.34065425839854,91.34065425849855,91.35091207257948,91.35091207267948,91.35347533406585,91.35347533416585,91.3637266461076,91.3637266462076,91.36628710970378,91.36628710980378,91.37140598773713,91.37140598783714,91.37652261957696,91.37652261967696,91.37907933951726,91.37907933961726,91.38419210948048,91.38419210958048,91.38674693788937,91.38674693798937,91.38930105697399,91.389301057074,91.39185379681697,91.39185379691698,91.39440456631473,91.39440456641474,91.39950318919938,91.39950318929938,91.40205001800673,91.40205001810673,91.40714300570407,91.40714300580407,91.41732638024308,91.41732638034308,91.41987052938096,91.41987052948096,91.42241329927718,91.42241329937718,91.42749690813135,91.42749690823135,91.43003731361333,91.43003731371333,91.43257677332961,91.43257677342962,91.43511544490781,91.43511544500781,91.4376530919065,91.4376530920065,91.44526366848834,91.44526366858834,91.45540637510229,91.45540637520229,91.46047496992595,91.46047497002596,91.46554167321824,91.46554167331824,91.47060632735152,91.47060632745152,91.4782022051581,91.4782022052581,91.4832642584357,91.4832642585357,91.48579359057761,91.48579359067762,91.49338012894792,91.49338012904792,91.49843457669303,91.49843457679303,91.50348421679581,91.50348421689581,91.50852885222179,91.50852885232179,91.51357135967493,91.51357135977493,91.51861083279645,91.51861083289646,91.52364813853826,91.52364813863826,91.53120086578481,91.53120086588481,91.53371628728958,91.53371628738958,91.5387462633472,91.53874626344721,91.54377438728034,91.54377438738034,91.5538266944562,91.55382669455621,91.55633849052577,91.55633849062578,91.56136003350592,91.56136003360592,91.56386836176794,91.56386836186795,91.57389891633272,91.57389891643273,91.58141696550364,91.58141696560364,91.58642672581945,91.58642672591945,91.58892997059083,91.58892997069083,91.59643781337358,91.59643781347359,91.60144209612972,91.60144209622972,91.60394222775567,91.60394222785567,91.61143891838455,91.61143891848455,91.61643165491778,91.61643165501778,91.62142049016748,91.62142049026748,91.62640719744437,91.62640719754437,91.63139169793463,91.63139169803463,91.63388264775192,91.63388264785192,91.63886312873795,91.63886312883795,91.64135210821004,91.64135210831004,91.64881739153633,91.64881739163633,91.65628031044837,91.65628031054837,91.65876530982312,91.65876530992313,91.66373436280696,91.66373436290696,91.66621691895368,91.66621691905368,91.66869722890685,91.66869722900685,91.67613709477997,91.67613709487998,91.68109535027207,91.68109535037208,91.68605135957064,91.68605135967064,91.68852500975702,91.68852500985702,91.69594469929523,91.69594469939523,91.70088912296397,91.70088912306397,91.70335902949445,91.70335902959445,91.70829722687236,91.70829722697236,91.71076457195407,91.71076457205407,91.71569792228276,91.71569792238276,91.71816309998475,91.71816310008475,91.7230921943678,91.7230921944678,91.725555638166,91.725555638266,91.72801837263994,91.72801837273994,91.73048020075507,91.73048020085507,91.7378629660241,91.7378629661241,91.74523946559536,91.74523946569536,91.74769494919893,91.74769494929893,91.75014826542278,91.75014826552278,91.7575061649353,91.7575061650353,91.76731052387032,91.76731052397032,91.76975950533472,91.76975950543472,91.77955424898515,91.77955424908515,91.78444787715446,91.78444787725446,91.79422961652654,91.79422961662654,91.79667323865196,91.79667323875196,91.80644457460133,91.80644457470133,91.8088847289192,91.8088847290192,91.81376428882375,91.81376428892375,91.81620306389996,91.81620306399996,91.81863975041026,91.81863975051026,91.82351150774778,91.82351150784778,91.82594535696099,91.82594535706099,91.82837810278087,91.82837810288088,91.83324245162042,91.83324245172042,91.8381053029976,91.8381053030976,91.84053355643043,91.84053355653043,91.84296102172516,91.84296102182516,91.84781417900393,91.84781417910393,91.85023939810513,91.85023939820513,91.85508833884516,91.85508833894517,91.85993507279856,91.85993507289857,91.8647773537718,91.8647773538718,91.87203196778857,91.87203196788857,91.87927862161071,91.87927862171071,91.88652326568074,91.88652326578074,91.88893689915211,91.88893689925212,91.89376278685323,91.89376278695323,91.89617370124823,91.89617370134823,91.90581554611062,91.90581554621062,91.91063394649994,91.91063394659994,91.91544911552313,91.91544911562313,91.91785242438563,91.91785242448563,91.92265817515873,91.92265817525873,91.927461797959,91.927461798059,91.93226400211073,91.93226400221073,91.93466348850353,91.93466348860353,91.94665733443922,91.94665733453922,91.95145138136179,91.9514513814618,91.9562438520082,91.9562438521082,91.96103498281987,91.96103498291987,91.96582398565872,91.96582398575872,91.97539875997026,91.97539876007026,91.9777910348996,91.9777910349996,91.98496569230788,91.98496569240788,91.99213672428098,91.99213672438098,91.9969149296281,91.9969149297281,92.00169077056096,92.00169077066096,92.00646436530029,92.00646436540029,92.01123673842558,92.01123673852558,92.0183935050994,92.0183935051994,92.02316272567236,92.02316272577237,92.0303140147865,92.0303140148865,92.03269482230672,92.03269482240673,92.0398350380808,92.0398350381808,92.04221292949013,92.04221292959014,92.0493445151522,92.0493445152522,92.05647200249624,92.05647200259624,92.06597154902778,92.06597154912778,92.0730921795705,92.0730921796705,92.0825816379346,92.0825816380346,92.09206510644925,92.09206510654926,92.09680532354078,92.09680532364078,92.1062816199989,92.1062816200989,92.11101655656526,92.11101655666526,92.11811677433164,92.11811677443164,92.12048134620737,92.12048134630737,92.12520930775172,92.12520930785172,92.1275724609789,92.1275724610789,92.13466050201188,92.13466050211188,92.14174629685135,92.14174629695135,92.14410590345716,92.14410590355716,92.14882361920642,92.14882361930643,92.15118121606012,92.15118121616013,92.15589554281563,92.15589554291563,92.1582509722896,92.1582509723896,92.16767068043337,92.16767068053338,92.17002453363118,92.17002453373118,92.17237771691161,92.17237771701161,92.1770832559275,92.1770832560275,92.17943462649035,92.17943462659035,92.18178532713583,92.18178532723583,92.1841351608294,92.1841351609294,92.19353351043114,92.19353351053114,92.19823061636954,92.19823061646954,92.20527389115944,92.20527389125944,92.2123130282244,92.2123130283244,92.21465659622022,92.21465659632022,92.2169977603949,92.2169977604949,92.2193381364315,92.2193381365315,92.22401699697328,92.22401699707328,92.23103048014366,92.23103048024366,92.2357032720222,92.23570327212221,92.24271001661197,92.24271001671197,92.24737646397895,92.24737646407895,92.25203944353837,92.25203944363837,92.25436890386251,92.25436890396251,92.25669674850359,92.25669674860359,92.25902321390302,92.25902321400302,92.26134782717796,92.26134782727796,92.26599567448619,92.2659956745862,92.27064190611135,92.27064190621135,92.27528604917059,92.27528604927059,92.27760656412751,92.27760656422751,92.27992636976013,92.27992636986014,92.28456452296994,92.28456452306995,92.28920090286908,92.28920090296909,92.29383574589895,92.29383574599895,92.30078495640868,92.30078495650868,92.30541511001695,92.30541511011695,92.31467309222614,92.31467309232615,92.3239257939102,92.32392579401021,92.33086038445849,92.33086038455849,92.33548068634073,92.33548068644073,92.34240798661176,92.34240798671176,92.3447153790696,92.3447153791696,92.3516357831324,92.3516357832324,92.35624757312334,92.35624757322334,92.36085577708602,92.36085577718602,92.36315816486702,92.36315816496702,92.37006351549245,92.37006351559245,92.37466505968833,92.37466505978833,92.37926356955258,92.37926356965258,92.38385967559567,92.38385967569567,92.39534635467515,92.39534635477516,92.39993761366904,92.39993761376904,92.40452607477275,92.40452607487275,92.40681928074508,92.40681928084508,92.40911094984817,92.40911094994817,92.41369286940578,92.41369286950578,92.4205639360246,92.42056393612461,92.42514238777466,92.42514238787466,92.43200864675117,92.43200864685117,92.43429629635004,92.43429629645004,92.44115684132547,92.44115684142547,92.44572875152942,92.44572875162942,92.45258370072446,92.45258370082446,92.45486729141219,92.45486729151219,92.45943281769769,92.45943281779769,92.4639947973618,92.46399479746181,92.46855445201858,92.46855445211858,92.47083272277425,92.47083272287425,92.47538776682325,92.47538776692325,92.47994044645799,92.47994044655799,92.48221505237159,92.48221505247159,92.48676304258473,92.48676304268473,92.49585563401727,92.49585563411728,92.50039735853268,92.50039735863268,92.50493447244028,92.50493447254028,92.50720198511114,92.50720198521114,92.51173594646642,92.51173594656642,92.51853426794021,92.51853426804021,92.5230645644534,92.52306456455341,92.5253287669443,92.5253287670443,92.52759072324166,92.52759072334166,92.53663681452733,92.53663681462733,92.54115674702474,92.54115674712475,92.54567289645935,92.54567289655935,92.55244351487954,92.55244351497954,92.55469896903772,92.55469896913772,92.55920774938126,92.55920774948126,92.56371392886912,92.56371392896912,92.57047000614168,92.57047000624168,92.57271856409163,92.57271856419163,92.57946262225843,92.57946262235843,92.58170905223557,92.58170905233557,92.58395382712273,92.58395382722273,92.59068657550804,92.59068657560805,92.5974159743065,92.5974159744065,92.59965787248966,92.59965787258966,92.6018986278726,92.6018986279726,92.60413855571055,92.60413855581055,92.60861707155172,92.60861707165172,92.61085463497541,92.61085463507541,92.61309148907485,92.61309148917485,92.61756393625276,92.61756393635277,92.61979842593794,92.61979842603795,92.62426626250809,92.62426626260809,92.62873256220897,92.62873256230897,92.63319748266821,92.63319748276821,92.63989295212221,92.63989295222221,92.64435416833246,92.64435416843246,92.64658294401653,92.64658294411653,92.65103891910853,92.65103891920853,92.65549138698606,92.65549138708606,92.65994054468364,92.65994054478364,92.66216297585615,92.66216297595615,92.66660669540096,92.66660669550096,92.67326989060048,92.67326989070048,92.67548944506899,92.67548944516899,92.68436541674951,92.68436541684952,92.68880168838943,92.68880168848943,92.69545424372484,92.69545424382484,92.69988673230196,92.69988673240196,92.704316383582,92.704316383682,92.70652951472515,92.70652951482515,92.70874177891639,92.70874177901639,92.7131645339882,92.71316453408821,92.71758449116982,92.71758449126982,92.72200094113697,92.72200094123697,92.72641415973797,92.72641415983797,92.73082414697285,92.73082414707285,92.73302663825187,92.73302663835187,92.73522790791687,92.73522790801687,92.74623264055876,92.74623264065876,92.75063190911571,92.75063190921571,92.75722850664724,92.75722850674724,92.75942449578707,92.75942449588707,92.76381525245273,92.76381525255273,92.77039935799564,92.77039935809564,92.77259239161768,92.77259239171768,92.77697747368913,92.77697747378913,92.7835526338648,92.7835526339648,92.79012369572246,92.79012369582246,92.79669243257275,92.79669243267276,92.80106904215982,92.80106904225983,92.80325561304957,92.80325561314957,92.8054410411391,92.8054410412391,92.81199456692441,92.81199456702441,92.8141757390683,92.8141757391683,92.81635624129481,92.81635624139481,92.82071480251979,92.82071480261979,92.82724870366684,92.82724870376684,92.83160273309784,92.83160273319784,92.84030633897966,92.84030633907966,92.84247982677724,92.84247982687724,92.84899887152143,92.84899887162143,92.85116960083572,92.85116960093572,92.85333962082574,92.85333962092574,92.85767851800554,92.85767851810554,92.85984676468487,92.85984676478488,92.86418246990544,92.86418247000545,92.87285013669072,92.87285013679072,92.88151484795816,92.88151484805816,92.8858445239224,92.8858445240224,92.88800741126276,92.88800741136276,92.89881964117797,92.89881964127797,92.90530452013627,92.90530452023627,92.90962489607114,92.90962489617114,92.913943222847,92.913943222947,92.91826005216048,92.91826005226048,92.92041722549975,92.92041722559975,92.92257278315594,92.92257278325594,92.92688307092335,92.92688307102335,92.9311917824146,92.9311917825146,92.93334489684274,92.93334489694274,92.93549698669138,92.93549698679138,92.93979955070557,92.93979955080557,92.94840058041594,92.94840058051594,92.95269762746355,92.95269762756355,92.95699341349024,92.95699341359024,92.96128774146148,92.96128774156148,92.96557832577682,92.96557832587682,92.96986579694673,92.96986579704674,92.97200827151076,92.97200827161076,92.97629014690028,92.97629014700028,92.98056839685462,92.98056839695462,92.984844164174,92.98484416427401,92.99125515518706,92.99125515528706,92.99766370297206,92.99766370307206,93.00406834947354,93.00406834957354,93.00833655066734,93.00833655076734,93.01473694122316,93.01473694132316,93.0232671961337,93.0232671962337,93.02539796684721,93.02539796694721,93.0317884662701,93.0317884663701,93.03391766070743,93.03391766080743,93.03817447330593,93.03817447340593,93.04668581290248,93.04668581300248,93.04881232767033,93.04881232777034,93.05306342626773,93.05306342636773,93.05731286977516,93.05731286987516,93.05943528622497,93.05943528632497,93.06367873988297,93.06367873998298,93.06791805581602,93.06791805591602,93.07426981825216,93.07426981835216,93.07850168628035,93.07850168638035,93.08273130811499,93.08273130821499,93.08695864434918,93.08695864444918,93.09118452252794,93.09118452262794,93.09752020694718,93.09752020704718,93.09962942208412,93.09962942218412,93.10595560943948,93.10595560953948,93.11016974436079,93.11016974446079,93.11227285142758,93.11227285152758,93.1143751703563,93.1143751704563,93.11857854719278,93.11857854729278,93.12067877755557,93.12067877765557,93.13327681014547,93.13327681024548,93.14166772224299,93.14166772234299,93.14376405132226,93.14376405142227,93.15214668796992,93.15214668806992,93.15633445967237,93.15633445977237,93.16052053687794,93.16052053697794,93.16679494356126,93.16679494366126,93.16888362770113,93.16888362780114,93.1709711690408,93.1709711691408,93.17723099516958,93.17723099526958,93.17931483226026,93.17931483236026,93.18139796002666,93.18139796012666,93.18556287572471,93.18556287582471,93.1918065844297,93.1918065845297,93.19388585031949,93.19388585041949,93.19804351514719,93.19804351524719,93.20012065306416,93.20012065316416,93.20219708165685,93.20219708175685,93.20427276151837,93.20427276161837,93.20842250555833,93.20842250565833,93.2125703974738,93.2125703975738,93.21464241249322,93.21464241259322,93.2187846298145,93.2187846299145,93.22292448272152,93.22292448282153,93.22499263586434,93.22499263596434,93.23326323868353,93.23326323878354,93.2373958801271,93.2373958802271,93.23946036842784,93.23946036852784,93.24152343808004,93.24152343818004,93.2435852861182,93.2435852862182,93.24770732710455,93.24770732720455,93.25182802825617,93.25182802835617,93.25800592743126,93.25800592753126,93.26006407122043,93.26006407132043,93.26417913718475,93.26417913728476,93.26829282390743,93.26829282400743,93.27240375214681,93.27240375224682,93.27445760058345,93.27445760068345,93.27651018799914,93.27651018809914,93.28266601930791,93.28266601940791,93.28676662293842,93.28676662303842,93.29086438927182,93.29086438937182,93.29496030348074,93.29496030358074,93.29700709808152,93.29700709818152,93.2990529075097,93.2990529076097,93.30314326534513,93.30314326544513,93.30723184986986,93.30723184996987,93.3113183458287,93.3113183459287,93.31540318669755,93.31540318679755,93.319486136035,93.319486136135,93.32560953546167,93.32560953556167,93.32968948987441,93.32968948997441,93.33784727072705,93.33784727082706,93.3419243878427,93.3419243879427,93.3439617641934,93.3439617642934,93.35007172586576,93.35007172596576,93.35414060693843,93.35414060703843,93.35820680834162,93.35820680844162,93.36227143346865,93.36227143356865,93.37242824775427,93.37242824785427,93.37445805797938,93.37445805807938,93.37851551104986,93.37851551114986,93.38865384349744,93.38865384359744,93.3927073164706,93.39270731657061,93.40081091283007,93.40081091293007,93.41093604396477,93.41093604406477,93.41295958849211,93.41295958859212,93.41498132030186,93.41498132040186,93.41700210634592,93.41700210644592,93.42104178690263,93.42104178700264,93.42507669922394,93.42507669932394,93.43313883952025,93.43313883962026,93.43716795902667,93.43716795912667,93.43918043021394,93.43918043031394,93.44521536114082,93.44521536124083,93.44923545646618,93.44923545656619,93.45526400347461,93.45526400357461,93.46128959496521,93.46128959506521,93.46329568223402,93.46329568233402,93.46730635930929,93.46730635940929,93.47331756727989,93.4733175673799,93.47732150577455,93.47732150587456,93.47932140615941,93.47932140625942,93.48132012433716,93.48132012443716,93.48331734505254,93.48331734515254,93.48730926444144,93.48730926454144,93.48930356904592,93.48930356914592,93.4912970461054,93.4912970462054,93.49528277861035,93.49528277871035,93.49727448235916,93.49727448245916,93.50324849021224,93.50324849031225,93.50722906041275,93.50722906051276,93.51319652671977,93.51319652681977,93.52114679081524,93.52114679091524,93.52710590285857,93.52710590295857,93.53306146828052,93.53306146838052,93.53702607868486,93.53702607878486,93.54296796991109,93.54296797001109,93.5449473392989,93.5449473393989,93.55088371355853,93.55088371365854,93.5548390633404,93.5548390634404,93.5607683049504,93.5607683050504,93.56274318195113,93.56274318205114,93.5647164826757,93.56471648277571,93.56866174429011,93.56866174439011,93.57063315348329,93.57063315358329,93.57654608063497,93.57654608073497,93.58245684040692,93.58245684050692,93.58442608222038,93.58442608232038,93.58836318660562,93.58836318670562,93.59229828123875,93.59229828133876,93.59623124789907,93.59623124799907,93.60999198119548,93.60999198129548,93.61392006139968,93.61392006149968,93.61784526489987,93.61784526499987,93.62569066722345,93.62569066732345,93.63745189420403,93.63745189430404,93.643328527597,93.64332852769701,93.64528527742185,93.64528527752185,93.65311085807264,93.65311085817264,93.65702163864592,93.65702163874592,93.66093068531542,93.66093068541542,93.6648376040121,93.66483760411211,93.66678840339442,93.66678840349442,93.6687379023489,93.6687379024489,93.67068661316529,93.67068661326529,93.67263437821597,93.67263437831598,93.67847601827808,93.67847601837808,93.68042228586641,93.68042228596642,93.68625959116906,93.68625959126906,93.68820424307432,93.68820424317433,93.69209256171226,93.69209256181226,93.69597891000498,93.69597891010498,93.697921197496,93.697921197596,93.69986226337298,93.69986226347298,93.70180246229808,93.70180246239808,93.70374203071272,93.70374203081272,93.70762022177628,93.70762022187628,93.70955596772122,93.70955596782122,93.71342560748658,93.71342560758659,93.71535891020345,93.71535891030345,93.71729158240986,93.71729158250986,93.71922330885057,93.71922330895057,93.72308538249034,93.72308538259034,93.72887488721828,93.72887488731828,93.73659068023414,93.73659068033415,93.7443004439937,93.7443004440937,93.75200437553141,93.75200437563142,93.75392924517081,93.75392924527081,93.75777772342865,93.75777772352865,93.75970074094354,93.75970074104355,93.76162257625131,93.76162257635131,93.76546486762521,93.76546486772521,93.76930593738508,93.76930593748509,93.77122540827861,93.77122540837861,93.77314424866167,93.77314424876167,93.77698094425517,93.77698094435517,93.77889840539659,93.77889840549659,93.78081460551707,93.78081460561707,93.78464452312306,93.78464452322306,93.78847156402507,93.78847156412508,93.7942087363843,93.79420873648431,93.79803045735424,93.79803045745425,93.80184863170281,93.80184863180281,93.81138813683516,93.81138813693516,93.81710797015658,93.81710797025659,93.82091842075194,93.82091842085194,93.82472583701568,93.82472583711568,93.8323363741906,93.8323363742906,93.8380407600056,93.8380407601056,93.83994100032992,93.83994100042992,93.84374053521282,93.84374053531282,93.84753742983315,93.84753742993315,93.85322971772858,93.85322971782858,93.85702275047231,93.85702275057231,93.86081294591895,93.86081294601895,93.8627069599524,93.8627069600524,93.86838734696278,93.86838734706278,93.87406639413842,93.87406639423843,93.87784973278373,93.87784973288373,93.88541317870822,93.88541317880822,93.88919269488383,93.88919269498383,93.89108064025405,93.89108064035405,93.89296783689308,93.89296783699308,93.89485424539403,93.89485424549403,93.9005116187724,93.9005116188724,93.90428069211842,93.90428069221842,93.91181434642337,93.91181434652337,93.91557802102352,93.91557802112352,93.9174589913717,93.9174589914717,93.9193391735818,93.9193391736818,93.92309823757417,93.92309823767417,93.92497660706668,93.92497660716668,93.92873228206528,93.92873228216529,93.93060899646783,93.93060899656783,93.9343596667896,93.9343596668896,93.93623267694316,93.93623267704317,93.94185032816216,93.94185032826216,93.94559213193051,93.94559213203051,93.94746092554531,93.94746092564532,93.95493412965934,93.95493412975934,93.95866903721947,93.95866903731947,93.96426915236611,93.96426915246612,93.96799992220132,93.96799992230132,93.97172852465678,93.97172852475678,93.97731906392573,93.97731906402574,93.98663190054549,93.98663190064549,93.98849289159328,93.98849289169328,93.99407283040502,93.99407283050502,93.99964701580876,93.99964701590876,94.00521856094993,94.00521856104993,94.00893049428494,94.00893049438494,94.01264077252999,94.01264077263,94.02005589086731,94.02005589096731,94.0237580512901,94.0237580513901,94.02560753552187,94.02560753562187,94.02745516762916,94.02745516772916,94.03114877675375,94.03114877685375,94.0366868260264,94.0366868261264,94.04037680971582,94.04037680981583,94.0422206587603,94.0422206588603,94.04590748989739,94.04590748999739,94.04959254772379,94.04959254782379,94.05327606868093,94.05327606878093,94.05695824980333,94.05695824990333,94.06432036585458,94.06432036595459,94.071678304774,94.071678304874,94.07535433841934,94.07535433851935,94.08270337137844,94.08270337147844,94.09004818779877,94.09004818789877,94.09922199686075,94.09922199696075,94.11022278881224,94.11022278891224,94.11205430349573,94.11205430359573,94.11571642650392,94.11571642660392,94.11754608906293,94.11754608916293,94.12486249310538,94.12486249320538,94.12851835041582,94.12851835051582,94.1321717250913,94.1321717251913,94.13582210484205,94.13582210494205,94.13946925322666,94.13946925332667,94.15040518141396,94.15040518151396,94.15404803444602,94.15404803454602,94.15950786101396,94.15950786111397,94.16314310851354,94.16314310861354,94.16495953035275,94.16495953045275,94.16859146767237,94.16859146777237,94.17403758063766,94.17403758073766,94.1794812503749,94.17948125047491,94.18492208281505,94.18492208291505,94.19036149660664,94.19036149670664,94.19398602542829,94.19398602552829,94.19942047394997,94.19942047404997,94.20485314916095,94.20485314926096,94.20847239745744,94.20847239755744,94.21208916311897,94.21208916321898,94.21570285504198,94.21570285514198,94.21931398551622,94.21931398561622,94.22653301509857,94.22653301519857,94.2319431495604,94.2319431496604,94.23554765967474,94.23554765977474,94.24095168606642,94.24095168616643,94.24275187226112,94.24275187236113,94.24815030287242,94.24815030297242,94.2499483610943,94.2499483611943,94.2553411171114,94.2553411172114,94.25893439625807,94.25893439635807,94.26252625675619,94.26252625685619,94.26970733748985,94.26970733758985,94.27150137620751,94.27150137630751,94.2750881138081,94.2750881139081,94.27867370860845,94.27867370870845,94.28404991372581,94.28404991382581,94.28583997234615,94.28583997244615,94.2894176857657,94.2894176858657,94.29477923151471,94.29477923161471,94.29835127034005,94.29835127044005,94.30192039305449,94.30192039315449,94.30548522041637,94.30548522051637,94.31082627457525,94.31082627467525,94.32150290533332,94.32150290543332,94.32683903362917,94.32683903372917,94.33039452155478,94.33039452165478,94.33572576339452,94.33572576349452,94.34105330098527,94.34105330108527,94.34460275965453,94.34460275975454,94.34637634618896,94.34637634628896,94.34992225823686,94.34992225833686,94.35346647578787,94.35346647588787,94.36055077316497,94.36055077326498,94.36408971019084,94.36408971029084,94.36939360363911,94.36939360373911,94.37292528979462,94.37292528989462,94.37645461153588,94.37645461163588,94.37821751879927,94.37821751889928,94.37997963792458,94.37997963802458,94.38174108713253,94.38174108723253,94.3870240949216,94.3870240950216,94.39054435307492,94.39054435317492,94.39582294729072,94.39582294739073,94.39934048636765,94.39934048646765,94.40285585806484,94.40285585816484,94.40461151445787,94.40461151455787,94.40812097511945,94.40812097521945,94.41162684975274,94.41162684985274,94.41512941420609,94.4151294143061,94.42038042358901,94.42038042368901,94.42387849565529,94.42387849575529,94.42562658592273,94.42562658602273,94.42912150543667,94.42912150553667,94.43436270250047,94.43436270260047,94.43610886182961,94.43610886192961,94.43785427242756,94.43785427252756,94.44308877031766,94.44308877041766,94.44657541438174,94.44657541448174,94.45180301606361,94.45180301616361,94.45528662579606,94.45528662589606,94.45702691349648,94.45702691359648,94.46050614906258,94.46050614916258,94.46398372953871,94.46398372963871,94.46919742058341,94.46919742068341,94.47440642220653,94.47440642230653,94.48134546573499,94.48134546583499,94.48481252456772,94.48481252466772,94.48654453681827,94.48654453691827,94.49000757614677,94.49000757624677,94.49173722398308,94.49173722408308,94.49692498528486,94.49692498538487,94.49865289921738,94.49865289931738,94.50037998560492,94.50037998570492,94.50210510164725,94.50210510174725,94.50382915370317,94.50382915380317,94.50899922130499,94.50899922140499,94.51244354235403,94.51244354245404,94.51588585365096,94.51588585375096,94.51932611578887,94.51932611588887,94.52104465087821,94.52104465097821,94.52448018418762,94.52448018428763,94.53134790121959,94.53134790131959,94.53306336257042,94.53306336267042,94.53820844619501,94.53820844629502,94.5433513624399,94.54335136253991,94.54849215071197,94.54849215081197,94.55020438069664,94.55020438079664,94.55191598017085,94.55191598027085,94.55533827276047,94.55533827286047,94.5621798236081,94.5621798237081,94.56388890104044,94.56388890114044,94.56730516437374,94.56730516447374,94.57242617037993,94.57242617047993,94.57413134652876,94.57413134662876,94.57924561395433,94.57924561405433,94.58265332598943,94.58265332608943,94.58435590128259,94.5843559013826,94.5894619720721,94.5894619721721,94.5945649297162,94.5945649298162,94.59626478593297,94.59626478603298,94.5996633555663,94.59966335566631,94.60136092618265,94.60136092628265,94.60475417588393,94.60475417598393,94.6064482392858,94.6064482393858,94.60983534151002,94.60983534161002,94.61491292110784,94.61491292120785,94.61660458068856,94.61660458078856,94.6182953733174,94.6182953734174,94.62167589458863,94.62167589468864,94.62336491390677,94.62336491400677,94.62674177033593,94.62674177043593,94.62842944981934,94.62842944991934,94.63011618353703,94.63011618363703,94.63686079340049,94.63686079350049,94.63854539914537,94.63854539924537,94.64359823120739,94.6435982313074,94.64696492065528,94.64696492075528,94.65201274804049,94.65201274814049,94.65537596968082,94.65537596978082,94.66041757077517,94.66041757087517,94.66209547734634,94.66209547744634,94.66545026590917,94.66545026600917,94.66880343878896,94.66880343888896,94.67215436547518,94.67215436557518,94.68052864785916,94.68052864795916,94.68387606733093,94.68387606743093,94.68554849634242,94.68554849644242,94.69056436472837,94.69056436482838,94.69223435051181,94.69223435061181,94.6972429286205,94.6972429287205,94.7005802599248,94.70058026002481,94.70224807832852,94.70224807842852,94.70391510859416,94.70391510869416,94.70724814454591,94.70724814464592,94.71058003769745,94.71058003779746,94.71391031516592,94.71391031526592,94.71723905576513,94.71723905586514,94.72056527432248,94.72056527442248,94.72388873439652,94.72388873449653,94.7288713433554,94.7288713434554,94.73219074451832,94.73219074461832,94.73550849059127,94.73550849069127,94.73882462098115,94.73882462108115,94.74213921450178,94.74213921460178,94.74876528839759,94.7487652884976,94.7537329621398,94.7537329622398,94.75538793389276,94.75538793399276,94.75869649815705,94.75869649825705,94.76035005126147,94.76035005136147,94.76200297385542,94.76200297395542,94.76861312736197,94.76861312746198,94.77191581999755,94.77191582009755,94.77686654877095,94.77686654887096,94.78181514957154,94.78181514967154,94.7851129163441,94.7851129164441,94.79170549437141,94.79170549447142,94.79500002977782,94.79500002987783,94.79994019750093,94.79994019760093,94.80158358362408,94.80158358372408,94.80651279622784,94.80651279632784,94.81143838339642,94.81143838349642,94.81636089682647,94.81636089692647,94.81964093049217,94.81964093059217,94.82291926966099,94.82291926976099,94.82619508678793,94.82619508688794,94.82783039449575,94.82783039459575,94.83763904878333,94.83763904888333,94.83927226792522,94.83927226802523,94.84417007322638,94.84417007332638,94.84743264963355,94.84743264973355,94.84906267681619,94.84906267691619,94.85232150956746,94.85232150966746,94.86046423698274,94.86046423708274,94.86697035626237,94.86697035636237,94.86859581224414,94.86859581234414,94.87022044068092,94.87022044078093,94.8750925920875,94.8750925921875,94.87833767182921,94.87833767192922,94.87995879305153,94.87995879315153,94.88320005032357,94.88320005042357,94.88643917962277,94.88643917972277,94.88967645679747,94.88967645689748,94.89452858949674,94.89452858959675,94.90099069126445,94.90099069136446,94.90421811671314,94.90421811681314,94.90905566885787,94.90905566895788,94.91872572906361,94.91872572916361,94.9203360133873,94.9203360134873,94.9235541782135,94.9235541783135,94.92998447860957,94.92998447870957,94.93159097987045,94.93159097997045,94.93640835568027,94.93640835578027,94.93801292600286,94.93801292610286,94.94122084503398,94.94122084513398,94.94282336619754,94.94282336629755,94.9460267140278,94.9460267141278,94.94762643730117,94.94762643740117,94.94922454489146,94.94922454499147,94.95241985371327,94.95241985381327,94.95721047223519,94.95721047233519,94.96359596698126,94.96359596708126,94.96838236896443,94.96838236906443,94.97635529202978,94.97635529212978,94.97954240421552,94.97954240431552,94.98113464017712,94.98113464027712,94.98431753582412,94.98431753592412,94.98908927843895,94.98908927853896,94.99067769193083,94.99067769203083,94.993853533742,94.993853533842,94.99544052858533,94.99544052868534,94.9986136513201,94.9986136514201,95.0017849613373,95.0017849614373,95.00495469507831,95.00495469517831,95.00812222203271,95.00812222213271,95.00970468508207,95.00970468518207,95.01761526642508,95.01761526652508,95.02235871488274,95.02235871498274,95.02551875452535,95.02551875462535,95.02867709967107,95.02867709977107,95.0318334744716,95.0318334745716,95.03498815477523,95.03498815487524,95.04444927957526,95.04444927967526,95.04760096495419,95.04760096505419,95.05232656208427,95.05232656218428,95.05704920369655,95.05704920379655,95.06019619965389,95.06019619975389,95.06491363955482,95.06491363965482,95.06805551261462,95.06805551271462,95.07276664741089,95.07276664751089,95.07590552554598,95.07590552564598,95.07747384151675,95.07747384161675,95.082177173746,95.082177173846,95.08530809168646,95.08530809178646,95.08999774971997,95.08999774981997,95.09312208670742,95.09312208680743,95.09468330943545,95.09468330953545,95.09624378343231,95.09624378353232,95.09780323284966,95.09780323294966,95.09936205175653,95.09936205185653,95.10091992489771,95.10091992499771,95.10403413431081,95.10403413441081,95.1071465704132,95.10714657051321,95.11336876294855,95.11336876304856,95.11958756649013,95.11958756659013,95.12114106546495,95.12114106556496,95.12269330341884,95.12269330351884,95.12734773168006,95.12734773178006,95.1288976446266,95.1288976447266,95.1335460830384,95.1335460831384,95.1350928040257,95.1350928041257,95.13663846102659,95.1366384611266,95.13972843519365,95.13972843529365,95.14127204303553,95.14127204313553,95.1443583917674,95.1443583918674,95.14898405298861,95.14898405308861,95.15206441187105,95.15206441197105,95.15514283981518,95.15514283991519,95.15668077306287,95.15668077316288,95.15821724825511,95.15821724835511,95.16282438823137,95.16282438833137,95.1658931220771,95.1658931221771,95.16896012201903,95.16896012211903,95.17049198660348,95.17049198670348,95.17355429712383,95.17355429722383,95.1766147555197,95.1766147556197,95.17967273128058,95.17967273138058,95.18119994585034,95.18119994595034,95.18272621465441,95.18272621475441,95.18882943774615,95.18882943784615,95.19340383058508,95.19340383068509,95.19492749853347,95.19492749863348,95.19949617737129,95.19949617747129,95.2055827707495,95.2055827708495,95.20710167046249,95.2071016705625,95.21165659629077,95.21165659639077,95.21469092790919,95.21469092800919,95.21772218578907,95.21772218588907,95.21923682955641,95.21923682965641,95.22075084281329,95.22075084291329,95.22377808118895,95.22377808128896,95.22680370388156,95.22680370398156,95.2313401872787,95.2313401873787,95.23436253919826,95.23436253929826,95.24040511506455,95.24040511516455,95.24191475415506,95.24191475425506,95.24644174048831,95.24644174058831,95.25096545604852,95.25096545614852,95.25548661015995,95.25548661025995,95.26151216224366,95.26151216234366,95.26602902100254,95.26602902110254,95.27054292424361,95.27054292434362,95.27505466010497,95.27505466020497,95.28407336359226,95.28407336369226,95.29008338935573,95.29008338945573,95.29458830782266,95.29458830792267,95.29758938001399,95.29758938011399,95.2990884580542,95.2990884581542,95.30058686617703,95.30058686627703,95.30358250021558,95.30358250031558,95.3095719161682,95.3095719162682,95.31256435824751,95.31256435834752,95.31405929856278,95.31405929866278,95.31854250382554,95.31854250392554,95.32152927131067,95.32152927141067,95.32749906262501,95.32749906272501,95.33048248052329,95.33048248062329,95.33346475562134,95.33346475572134,95.33495465185288,95.33495465195288,95.33942296130587,95.33942296140587,95.34239991647186,95.34239991657186,95.34388717244083,95.34388717254083,95.34834779754752,95.34834779764752,95.35132006329192,95.35132006339192,95.35726081171792,95.35726081181792,95.36171300374713,95.36171300384713,95.3631956885152,95.3631956886152,95.3661601910995,95.3661601911995,95.37208608312264,95.37208608322264,95.37504585687843,95.37504585697843,95.3794824437736,95.3794824438736,95.38687261754059,95.38687261764059,95.39278081586377,95.39278081596377,95.39425640738912,95.39425640748912,95.39868192094423,95.39868192104423,95.40015526627603,95.40015526637603,95.40162790228356,95.40162790238357,95.40457147980176,95.40457147990176,95.40751308697473,95.40751308707473,95.41045339371989,95.41045339381989,95.41192268014056,95.41192268024056,95.41339078435412,95.41339078445412,95.41926142789767,95.41926142799767,95.4221929863101,95.4221929864101,95.42805165015479,95.42805165025479,95.42951471028462,95.42951471038462,95.43243980596478,95.43243980606478,95.43536261604449,95.43536261614449,95.43974466378432,95.43974466388433,95.44704431989253,95.44704431999253,95.44850288763529,95.44850288773529,95.45141923498271,95.45141923508271,95.45433313910209,95.45433313920209,95.45578823903728,95.45578823913728,95.4601519625667,95.4601519626667,95.46305956158142,95.46305956168142,95.4659658601683,95.4659658602683,95.47032351503447,95.47032351513447,95.47322685810353,95.47322685820353,95.47612779735145,95.47612779745145,95.48192664151567,95.48192664161567,95.48482360066626,95.48482360076626,95.48771815599571,95.48771815609571,95.48916450759819,95.48916450769819,95.49060995284187,95.49060995294187,95.49494490933127,95.49494490943127,95.49783209556963,95.49783209566964,95.50216019525772,95.50216019535772,95.50792680457434,95.50792680467434,95.51656337931223,95.51656337941223,95.52087878997717,95.52087879007718,95.52231599917789,95.5223159992779,95.52375249905434,95.52375249915434,95.52518805316508,95.52518805326508,95.52662281913774,95.52662281923774,95.53379400873847,95.53379400883847,95.54239425031075,95.54239425041075,95.54812460527558,95.54812460537558,95.55098806855766,95.55098806865766,95.5552805838113,95.5552805839113,95.55814073691343,95.55814073701343,95.55956994651261,95.55956994661261,95.56099848619442,95.56099848629442,95.562425883076,95.562425883176,95.56813346085023,95.56813346095024,95.57383638860979,95.5738363887098,95.57526039649761,95.57526039659761,95.57668361624737,95.57668361634737,95.58379762643018,95.58379762653018,95.58521875761402,95.58521875771402,95.5880602318436,95.5880602319436,95.59231980292537,95.59231980302538,95.59799431830704,95.59799431840705,95.6022459686011,95.6022459687011,95.60649478159806,95.60649478169806,95.6107405996703,95.6107405997703,95.61781417836949,95.61781417846949,95.62205577990298,95.62205578000298,95.62629513524294,95.62629513534294,95.63194435139215,95.63194435149215,95.63900208851587,95.63900208861587,95.64182322878291,95.64182322888291,95.6474636177856,95.6474636178856,95.65028105380365,95.65028105390365,95.65591017243176,95.65591017253176,95.6629433196473,95.6629433197473,95.66575484462972,95.66575484472972,95.6699695312477,95.6699695313477,95.67839350573779,95.67839350583779,95.68119990782267,95.68119990792268,95.6826018478442,95.6826018479442,95.68400260565859,95.6840026057586,95.68680309670786,95.68680309680786,95.68820204180466,95.68820204190466,95.689600356391,95.689600356491,95.69379218700458,95.69379218710458,95.69518719141097,95.69518719151097,95.69937145589894,95.69937145599894,95.7021583515663,95.7021583516663,95.70633438001131,95.70633438011131,95.70911512820162,95.70911512830162,95.7105043792,95.7105043793,95.71189272383958,95.71189272393958,95.71466767921498,95.71466767931499,95.7188274720155,95.7188274721155,95.72298564913295,95.72298564923295,95.7243707229995,95.7243707230995,95.72990952100332,95.72990952110332,95.7340612747954,95.7340612748954,95.7354440630615,95.7354440631615,95.74235465480517,95.74235465490517,95.74373456636727,95.74373456646727,95.74649340431887,95.74649340441887,95.75062900128026,95.75062900138026,95.75200627257978,95.75200627267978,95.75613611613315,95.75613611623315,95.7588868756694,95.7588868757694,95.7630111234424,95.7630111235424,95.76988077141267,95.76988077151267,95.77262672330663,95.77262672340663,95.77674360198854,95.77674360208854,95.78085815566311,95.78085815576311,95.78359957576308,95.78359957586308,95.78907969688665,95.78907969698665,95.79181824028262,95.79181824038263,95.79592392740376,95.79592392750376,95.79865971231645,95.79865971241645,95.8000267378209,95.8000267379209,95.80822680228168,95.80822680238168,95.81095852828325,95.81095852838325,95.81505436367834,95.81505436377834,95.82051257456321,95.82051257466321,95.82596716001291,95.82596716011291,95.82732929906125,95.82732929916125,95.83413798455082,95.83413798465082,95.84094308401212,95.84094308411213,95.84502146214871,95.84502146224871,95.85045256108353,95.85045256118353,95.85316371668112,95.85316371678113,95.85858370286898,95.85858370296899,95.86129190294878,95.86129190304878,95.86399754157979,95.8639975416798,95.86670160393464,95.86670160403465,95.87345835112458,95.87345835122458,95.87615926092711,95.87615926102711,95.88155926781457,95.88155926791457,95.88560663271758,95.88560663281758,95.88830301072613,95.88830301082614,95.8923460408697,95.8923460409697,95.8963865095645,95.8963865096645,95.89907733119956,95.89907733129957,95.90445475793095,95.90445475803095,95.90714148124799,95.90714148134799,95.91116953676797,95.91116953686797,95.91251126303997,95.91251126313998,95.91653518083503,95.91653518093503,95.91787501557563,95.91787501567563,95.92055334522209,95.92055334532209,95.92189164309342,95.92189164319342,95.92456717484968,95.92456717494969,95.9325893959521,95.93258939605211,95.93659565945411,95.93659565955411,95.9405979822657,95.9405979823657,95.9445955368419,95.9445955369419,95.94992137993577,95.94992138003578,95.95391270822111,95.95391270832111,95.9565712950086,95.9565712951086,95.96055608174784,95.96055608184784,95.96586655614911,95.96586655624911,95.96851927130788,95.96851927140789,95.97116942501788,95.97116942511788,95.97779155822329,95.97779155832329,95.98176133093206,95.98176133103206,95.98572531082593,95.98572531092593,95.9883646670442,95.9883646671442,95.99231985919843,95.99231985929843,95.99363624683107,95.99363624693108,95.99626803692375,95.99626803702375,95.99889836896097,95.99889836906097,96.0041570232833,96.0041570233833,96.00941331319137,96.00941331329138,96.01203978335198,96.01203978345198,96.01466455901573,96.01466455911573,96.01859973246266,96.01859973256266,96.02122151320168,96.02122151330168,96.02384187529216,96.02384187539216,96.02646070051337,96.02646070061337,96.03169433145158,96.03169433155158,96.03430957064451,96.03430957074451,96.04084500866081,96.04084500876081,96.04606969423178,96.04606969433178,96.04867957408575,96.04867957418575,96.05128701071166,96.05128701081166,96.05258988177619,96.05258988187619,96.05519385059453,96.05519385069454,96.05909745911127,96.05909745921127,96.06429913105022,96.06429913115022,96.0668986468884,96.0668986469884,96.07209460482625,96.07209460492625,96.0772858339356,96.0772858340356,96.0785808630262,96.0785808631262,96.0824641769873,96.0824641770873,96.08375715691888,96.08375715701888,96.0876339687408,96.0876339688408,96.08892458425814,96.08892458435814,96.09150392376141,96.09150392386141,96.09923733166342,96.09923733176342,96.1031012574277,96.1031012575277,96.10696234589486,96.10696234599486,96.10824819317679,96.1082481932768,96.11210411933952,96.11210411943952,96.115957681088,96.11595768118801,96.1172403364107,96.1172403365107,96.11980414959375,96.11980414969375,96.12108487397813,96.12108487407814,96.12364557401575,96.12364557411576,96.13132353640366,96.13132353650366,96.13643898603635,96.13643898613635,96.141551164896,96.141551164996,96.14410532338754,96.14410532348754,96.14921040900444,96.14921040910444,96.1543110416412,96.1543110417412,96.15813299905255,96.15813299915256,96.16449772636015,96.16449772646016,96.16576970241182,96.16576970251182,96.17212694240763,96.17212694250763,96.17720826566135,96.17720826576135,96.17974689783264,96.17974689793265,96.18608967549464,96.18608967559464,96.19242713322457,96.19242713332457,96.19496040605691,96.19496040615691,96.19749265430973,96.19749265440973,96.20002316865877,96.20002316875878,96.2114073897875,96.2114073898875,96.21519963439314,96.21519963449315,96.22025226942066,96.22025226952066,96.22277675451336,96.22277675461336,96.22656139358651,96.22656139368651,96.22908237146476,96.22908237156476,96.23286165119895,96.23286165129895,96.23663821185674,96.23663821195674,96.24041264454172,96.24041264464172,96.24670064660697,96.24670064670697,96.25047102038081,96.25047102048082,96.25549504599593,96.25549504609593,96.25800498994101,96.25800499004102,96.26176740352022,96.26176740362023,96.26552725565067,96.26552725575067,96.26803105211872,96.26803105221872,96.27428571594305,96.27428571604305,96.27928538809141,96.27928538819141,96.28678101473388,96.28678101483388,96.28927744211084,96.28927744221085,96.29551453045595,96.29551453055595,96.29676069498541,96.29676069508541,96.30423519952726,96.30423519962726,96.30921591695473,96.30921591705473,96.31294920883178,96.31294920893178,96.31543621795866,96.31543621805866,96.32165060792698,96.32165060802699,96.32910119247802,96.32910119257802,96.3365474422696,96.3365474423696,96.33778718347371,96.33778718357371,96.34398348567302,96.34398348577302,96.3489377610678,96.3489377611678,96.35141373626153,96.35141373636154,96.35759998970029,96.35759998980029,96.36130967684178,96.36130967694179,96.36501621143093,96.36501621153093,96.36995551220214,96.36995551230214,96.37242297550458,96.37242297560458,96.37982114887312,96.37982114897312,96.38351814699146,96.38351814709146,96.38967523932118,96.38967523942118,96.3921360822637,96.3921360823637,96.3982836381226,96.39828363822261,96.40319689093019,96.40319689103019,96.40810640008188,96.40810640018188,96.41178617738312,96.41178617748312,96.41546311738728,96.41546311748728,96.41668748989875,96.41668748999875,96.42035891293632,96.42035891303632,96.42280517532433,96.42280517542433,96.4289180531076,96.4289180532076,96.43013895781151,96.43013895791151,96.4386821775935,96.43868217769351,96.4460016947814,96.4460016948814,96.44965719742969,96.44965719752969,96.4569616611802,96.4569616612802,96.45939472225531,96.45939472235531,96.4630415159778,96.46304151607781,96.46911423812575,96.46911423822576,96.47275460852286,96.47275460862286,96.47518026110001,96.47518026120001,96.4812391908315,96.4812391909315,96.4885024743672,96.4885024744672,96.49213102269306,96.49213102279306,96.49454859685486,96.49454859695486,96.49817324389721,96.49817324399721,96.50058758669287,96.50058758679288,96.50179406846988,96.50179406856988,96.50420604685128,96.50420604695128,96.51023325402495,96.51023325412496,96.51264298621282,96.51264298631283,96.51384670950654,96.51384670960654,96.51625305270065,96.51625305280065,96.52226504880933,96.52226504890933,96.52466820004422,96.52466820014422,96.52947154699616,96.52947154709616,96.5330711312475,96.5330711313475,96.5342699680851,96.5342699681851,96.53786509935625,96.53786509945625,96.54145676281985,96.54145676291985,96.54504519491731,96.54504519501731,96.54623965758856,96.54623965768856,96.55101526207999,96.55101526217999,96.55698190084202,96.55698190094202,96.56055827442681,96.56055827452681,96.56413153486618,96.56413153496618,96.5712721841162,96.5712721842162,96.57483910004399,96.57483910014399,96.5784022723159,96.5784022724159,96.58433482410584,96.58433482420584,96.58907618399759,96.58907618409759,96.59381309090917,96.59381309100917,96.59736353475104,96.59736353485104,96.59972783077843,96.59972783087844,96.60326992035664,96.60326992045664,96.60562936933482,96.60562936943482,96.60798763610589,96.60798763620589,96.61388034751576,96.61388034761576,96.61859179756726,96.61859179766726,96.62329804590742,96.62329804600742,96.62682520026895,96.62682520036896,96.63034943851957,96.63034943861958,96.63622012147002,96.63622012157002,96.6397390791955,96.6397390792955,96.64325527843768,96.64325527853768,96.64676856156895,96.64676856166895,96.65027900740311,96.65027900750312,96.65378720704376,96.65378720714376,96.65729288464253,96.65729288474253,96.6607947397716,96.6607947398716,96.66662912896335,96.66662912906335,96.66896091429484,96.66896091439484,96.6712915568261,96.6712915569261,96.6759504380675,96.6759504381675,96.67827769160502,96.67827769170502,96.68060234428685,96.68060234438686,96.68292388382328,96.68292388392328,96.68408370782578,96.68408370792578,96.69335875322447,96.69335875332447,96.69799280811625,96.69799280821626,96.70146625086745,96.70146625096746,96.70493713216987,96.70493713226988,96.70840450625784,96.70840450635784,96.71417958805885,96.71417958815886,96.71764168162164,96.71764168172164,96.72110247475662,96.72110247485662,96.72225512670256,96.72225512680257,96.72455944542187,96.72455944552188,96.72571032405715,96.72571032415715,96.7303117106254,96.7303117107254,96.7326112217024,96.7326112218024,96.739505814243,96.739505814343,96.74524657263052,96.74524657273052,96.74983274813378,96.74983274823379,96.75670744018778,96.75670744028778,96.76243369683458,96.76243369693458,96.76701069052918,96.76701069062918,96.77158378294027,96.77158378304027,96.77386674311754,96.77386674321754,96.77843037787163,96.77843037797163,96.78185133062651,96.78185133072651,96.78526987956025,96.78526987966025,96.78754775624688,96.78754775634688,96.79551610811134,96.79551610821134,96.8023396894107,96.8023396895107,96.80461224616526,96.80461224626526,96.80688314782986,96.80688314792987,96.81369155747112,96.81369155757112,96.81709292499465,96.81709292509466,96.81935909783076,96.81935909793076,96.82502072715477,96.82502072725477,96.83294068734097,96.83294068744097,96.83633027220016,96.83633027230016,96.83971733501748,96.83971733511748,96.84084520002074,96.84084520012074,96.84648286994704,96.84648287004704,96.84873631435312,96.84873631445312,96.85324115400624,96.85324115410624,96.85774295932774,96.85774295942774,96.86224109980716,96.86224109990717,96.86673632417568,96.86673632427568,96.87010541744472,96.87010541754472,96.87347261918237,96.87347261928237,96.87795893759055,96.87795893769055,96.88244170937735,96.88244170947735,96.88916188696024,96.88916188706024,96.89140047496346,96.89140047506346,96.89587481367278,96.89587481377278,96.89922735604209,96.89922735614209,96.90257851916975,96.90257851926975,96.90592723906934,96.90592723916934,96.91038963748672,96.91038963758672,96.91373425906829,96.9137342591683,96.9170763192011,96.9170763193011,96.92598476217063,96.92598476227063,96.93043600843413,96.93043600853413,96.93821871437322,96.93821871447322,96.94377311751107,96.94377311761107,96.94599322367625,96.94599322377626,96.95265050784015,96.95265050794015,96.96041134294744,96.96041134304744,96.96595246595861,96.96595246605861,96.97038251130769,96.97038251140769,96.9714890571758,96.9714890572758,96.97370112433254,96.97370112443254,96.97591161521309,96.9759116153131,96.97812096329343,96.97812096339344,96.98364135827303,96.98364135837303,96.98695090770993,96.98695090780993,96.9913610525724,96.9913610526724,96.99466685835341,96.99466685845341,96.99907105277337,96.99907105287338,97.00237173565684,97.00237173575684,97.00896939717477,97.00896939727477,97.01446362597002,97.01446362607003,97.01995355941271,97.01995355951271,97.02324454819777,97.02324454829777,97.0276294726416,97.02762947274161,97.0298204176977,97.02982041779771,97.03639077023108,97.03639077033108,97.03748466648298,97.03748466658298,97.04185867521437,97.04185867531437,97.04295091637628,97.04295091647629,97.04731806830637,97.04731806840637,97.05386564364915,97.05386564374915,97.05931845578814,97.05931845588815,97.06258654916189,97.0625865492619,97.06802915491272,97.06802915501272,97.06911647021161,97.06911647031161,97.07129011563681,97.07129011573682,97.0745481996569,97.0745481997569,97.08431579195039,97.08431579205039,97.08973680271775,97.08973680281775,97.0908198226641,97.0908198227641,97.09298495619801,97.09298495629801,97.09514839523503,97.09514839533503,97.09731013977517,97.09731013987518,97.10163090977908,97.10163090987908,97.10486873865045,97.10486873875045,97.109184306943,97.10918430704301,97.11241878622752,97.11241878632752,97.11457391040778,97.11457391050779,97.11672757653258,97.11672757663258,97.12103246555412,97.12103246565412,97.12210764352655,97.12210764362655,97.12640740965055,97.12640740975056,97.13070441729127,97.13070441739127,97.13392435542815,97.13392435552815,97.1392870439774,97.1392870440774,97.14464484607053,97.14464484617054,97.14892924350193,97.14892924360193,97.14999937739061,97.14999937749062,97.15427861251757,97.15427861261757,97.1585548921267,97.1585548922267,97.16496462211883,97.16496462221883,97.16709917589512,97.16709917599512,97.17029948939374,97.17029948949374,97.17243104824533,97.17243104834533,97.17882343919962,97.17882343929962,97.18308265561927,97.18308265571928,97.18521074666329,97.18521074676329,97.1915909214773,97.1915909215773,97.19371503242401,97.19371503252401,97.19796049583411,97.19796049593411,97.20114315503969,97.20114315513969,97.20432341042411,97.20432341052411,97.2127979439722,97.21279794407221,97.21491480404855,97.21491480414855,97.21808741449357,97.21808741459357,97.2223140414035,97.2223140415035,97.22864968641584,97.22864968651584,97.23287028406943,97.23287028416944,97.2360328457539,97.2360328458539,97.24024662601309,97.24024662611309,97.24235119113533,97.24235119123533,97.24971295252445,97.24971295262445,97.25707002449198,97.25707002459198,97.26022096113972,97.26022096123972,97.2686180601212,97.2686180602212,97.27176517429925,97.27176517439925,97.27595806889923,97.27595806899923,97.28014820501593,97.28014820511594,97.28642927146605,97.28642927156605,97.28956550933856,97.28956550943856,97.29269898872776,97.29269898882777,97.30000569867181,97.30000569877181,97.30626497310394,97.30626497320394,97.30834944070509,97.30834944080509,97.31564227941887,97.31564227951887,97.31876476428184,97.31876476438184,97.32188547583411,97.32188547593411,97.32396434765487,97.32396434775487,97.32500287720644,97.32500287730645,97.3322696285497,97.3322696286497,97.33745679874792,97.33745679884792,97.34471255556491,97.34471255566491,97.34781959292145,97.34781959302146,97.35092328069115,97.35092328079115,97.35505860180419,97.35505860190419,97.36022421642586,97.36022421652586,97.36435327184115,97.36435327194116,97.37054066808014,97.37054066818014,97.375690992823,97.375690992923,97.37980767447038,97.37980767457039,97.38597875625106,97.38597875635107,97.38906112488561,97.38906112498562,97.39522125154694,97.39522125164694,97.39932453484693,97.39932453494693,97.40034927198205,97.40034927208205,97.40239768226591,97.40239768236592,97.40751553571977,97.40751553581977,97.41058419075168,97.41058419085168,97.41262814805536,97.41262814815536,97.4207999577659,97.4207999578659,97.42692422414444,97.42692422424444,97.42896345261964,97.42896345271964,97.43507577870625,97.43507577880625,97.44016549563055,97.44016549573055,97.4421996012082,97.4421996013082,97.44626485684569,97.44626485694569,97.45032786628965,97.45032786638966,97.45337331065504,97.45337331075504,97.46047498647687,97.46047498657687,97.46351546557231,97.46351546567232,97.4675672440486,97.4675672441486,97.4716153182759,97.4716153183759,97.47464925582527,97.47464925592527,97.47869181308597,97.47869181318597,97.47970145737683,97.47970145747684,97.48071027412269,97.48071027422269,97.48877911359271,97.48877911369271,97.49281091276859,97.49281091286859,97.5018744613128,97.5018744614128,97.50690644712253,97.50690644722253,97.51293917126279,97.51293917136279,97.51595364180152,97.51595364190152,97.518965747926,97.518965748026,97.52598975273978,97.52598975283978,97.53100333552528,97.53100333562529,97.53601274117895,97.53601274127895,97.5370137553578,97.5370137554578,97.54101651164534,97.54101651174534,97.54501643063578,97.54501643073579,97.5490133941084,97.5490133942084,97.55400613064162,97.55400613074163,97.55699936085902,97.55699936095903,97.55999101480026,97.55999101490026,97.56397631382927,97.56397631392927,97.57094581889463,97.57094581899463,97.57393065544144,97.57393065554145,97.5818845055652,97.5818845056652,97.58983382389498,97.58983382399498,97.59281172482666,97.59281172492666,97.59876362540653,97.59876362550654,97.60371943767059,97.6037194377706,97.60668969366287,97.60668969376287,97.6106457133621,97.6106457134621,97.61459909279877,97.61459909289877,97.61756190088617,97.61756190098617,97.62249607875985,97.62249607885985,97.62644019757403,97.62644019767403,97.63432291823582,97.63432291833583,97.63826136245581,97.63826136255581,97.64219720582012,97.64219720592013,97.64613025129425,97.64613025139425,97.65104247952232,97.65104247962232,97.65398704221312,97.65398704231312,97.65692908286204,97.65692908296204,97.66084892702328,97.66084892712328,97.66378679054037,97.66378679064037,97.66868030048897,97.66868030058897,97.67063630158263,97.67063630168263,97.67552307295064,97.67552307305064,97.68333702856471,97.68333702866471,97.69016857005869,97.69016857015869,97.6940691442438,97.6940691443438,97.69699215135803,97.69699215145803,97.70185988919134,97.70185988929134,97.7086692445983,97.7086692446983,97.71352764299532,97.71352764309532,97.71741158746688,97.71741158756689,97.72323376051835,97.72323376061836,97.72905179584487,97.72905179594487,97.73389608657021,97.73389608667021,97.73970336381191,97.73970336391191,97.74744181579767,97.74744181589767,97.75034129699013,97.75034129709013,97.754203331223,97.754203331323,97.75902709095129,97.7590270910513,97.76577623260872,97.76577623270872,97.77155588561062,97.77155588571063,97.7744440176147,97.7744440177147,97.7792535514506,97.7792535515506,97.7850209489053,97.7850209490053,97.78790202707353,97.78790202717353,97.78982122211873,97.78982122221873,97.79365724779487,97.79365724789487,97.79940470535608,97.79940470545608,97.80323363778949,97.80323363788949,97.81279871800261,97.81279871810261,97.81662201524873,97.81662201534873,97.82235307953782,97.82235307963782,97.82712474333884,97.82712474343884,97.83379893306461,97.83379893316462,97.83951569264751,97.83951569274751,97.84618030649557,97.84618030659557,97.85188819952505,97.85188819962505,97.85473972251518,97.85473972261518,97.85948707225634,97.85948707235634,97.8651813699039,97.8651813700039,97.86802599668579,97.86802599678579,97.87655554227206,97.87655554237206,97.88129063646605,97.88129063656605,97.88318311363024,97.88318311373024,97.89169437441298,97.89169437451298,97.89547392999549,97.89547393009549,97.89924879615641,97.89924879625642,97.90207777839738,97.90207777849739,97.9058469699641,97.9058469700641,97.90773010769202,97.90773010779202,97.91149393991978,97.91149394001978,97.91525497425735,97.91525497435735,97.92183198655752,97.92183198665752,97.92276979206214,97.92276979216214,97.9283946547446,97.9283946548446,97.93307737716299,97.93307737726299,97.9386923881194,97.93869238821941,97.94336762322601,97.94336762332601,97.94710414646921,97.94710414656922,97.9508376353808,97.9508376354808,97.95549990561594,97.95549990571594,97.96015866863661,97.96015866873661,97.9648128210495,97.96481282114951,97.96667337862134,97.96667337872134,97.96946200819247,97.96946200829247,97.97503552367884,97.97503552377884,97.9806032463503,97.9806032464503,97.98616537324138,97.98616537334138,97.9898707650303,97.9898707651303,97.9954235524851,97.9954235525851,97.9991220086589,97.9991220087589,98.00466592956026,98.00466592966026,98.00743481627242,98.00743481637242,98.01020204789461,98.01020204799461,98.01388899725241,98.01388899735241,98.01849483680083,98.01849483690083,98.02770021079301,98.02770021089302,98.03229868125035,98.03229868135035,98.03965169430676,98.03965169440676,98.04240623690582,98.04240623700582,98.04791268184134,98.04791268194134,98.0552490651842,98.0552490652842,98.05799761793382,98.05799761803382,98.06074388508301,98.06074388518302,98.06714506377693,98.06714506387694,98.0708002905769,98.0708002906769,98.07901741822718,98.07901741832718,98.08540362229752,98.08540362239752,98.09087365525363,98.09087365535363,98.0945165476926,98.09451654779261,98.0981577062278,98.0981577063278,98.10179492407258,98.10179492417258,98.10724667222516,98.10724667232516,98.11178599291941,98.11178599301941,98.11541450183837,98.11541450193837,98.11813385407203,98.11813385417203,98.12175645195538,98.12175645205538,98.12447107536055,98.12447107546055,98.13080069111655,98.13080069121655,98.13441387074981,98.13441387084981,98.1416342795738,98.1416342796738,98.14974981622521,98.14974981632521,98.15245202645558,98.15245202655558,98.16055396772506,98.16055396782507,98.16594960044617,98.16594960054617,98.1713398738283,98.1713398739283,98.17851709268537,98.17851709278537,98.1838959380653,98.1838959381653,98.18927029105816,98.18927029115817,98.1946395999673,98.1946396000673,98.20000516522055,98.20000516532055,98.20357976549467,98.20357976559467,98.20715148906477,98.20715148916477,98.21339791684615,98.21339791694615,98.21696396582206,98.21696396592206,98.22141907396217,98.22141907406217,98.22587004437733,98.22587004447733,98.23120743369412,98.23120743379413,98.23565123205275,98.23565123215275,98.24453098679605,98.24453098689605,98.24896698258765,98.24896698268765,98.25162683039606,98.25162683049606,98.25871585660167,98.25871585670167,98.26225786736606,98.26225786746606,98.26579692261264,98.26579692271264,98.27021569758713,98.27021569768714,98.27109841223975,98.27109841233975,98.27727560209057,98.27727560219057,98.28697293187398,98.28697293197398,98.2931406640678,98.29314066416781,98.29930366743312,98.29930366753312,98.3080991308085,98.3080991309085,98.31600986977912,98.31600986987912,98.32215691334828,98.32215691344828,98.32566562527867,98.32566562537868,98.33180088618349,98.33180088628349,98.33880837950443,98.33880837960443,98.34318404332579,98.3431840434258,98.34668132725399,98.34668132735399,98.34842851116264,98.34842851126264,98.35104745460455,98.35104745470456,98.35802657495452,98.35802657505452,98.3606412230439,98.3606412231439,98.36586740607721,98.36586740617722,98.37195987108414,98.37195987118415,98.37978757970775,98.37978757980775,98.38413317606127,98.38413317616127,98.38847475291057,98.38847475301057,98.39281345305585,98.39281345315585,98.39714927649715,98.39714927659715,98.40148163213087,98.40148163223087,98.40494459264556,98.40494459274557,98.41186684883286,98.41186684893286,98.41705390081037,98.41705390091037,98.42137246402764,98.42137246412764,98.4282771053288,98.4282771054288,98.43345111362105,98.43345111372105,98.4360351425459,98.4360351426459,98.43947839960853,98.43947839970853,98.4446403099812,98.4446403100812,98.44979697923563,98.44979697933563,98.45494978661344,98.45494978671344,98.46267034786472,98.46267034796472,98.46524089962837,98.46524089972837,98.46866638417723,98.46866638427723,98.47294463413157,98.47294463423157,98.48063938386063,98.48063938396064,98.4832015419537,98.4832015420537,98.48661350993444,98.48661351003445,98.49513481947774,98.49513481957774,98.50109456203154,98.50109456213154,98.50875148113263,98.50875148123264,98.51640315911547,98.51640315921547,98.52064992295341,98.52064992305341,98.52404549766203,98.52404549776203,98.5257414525953,98.5257414526953,98.52743626472835,98.52743626482835,98.53420979925944,98.53420979935945,98.54013249932335,98.54013249942335,98.54182262203481,98.54182262213482,98.55026685167363,98.55026685177363,98.55617260676875,98.55617260686876,98.5637595392081,98.56375953930811,98.57049780455999,98.57049780466,98.57554681415232,98.57554681425232,98.58311234864273,98.58311234874273,98.58563163202409,98.58563163212409,98.59150605863043,98.59150605873043,98.59653516773618,98.59653516783618,98.60239852100246,98.60239852110246,98.60742069449307,98.60742069459307,98.61076614361963,98.61076614371963,98.61327246212953,98.61327246222953,98.61828245888677,98.61828245898677,98.62245407375852,98.62245407385852,98.62829019685405,98.62829019695405,98.63745052875481,98.63745052885481,98.64410627604944,98.64410627614944,98.64826252222858,98.64826252232858,98.65490601397603,98.65490601407603,98.65905453640197,98.65905453650197,98.66402808177287,98.66402808187287,98.66982089668075,98.66982089678075,98.67643609427103,98.67643609437103,98.68221980618405,98.68221980628405,98.69130140309035,98.69130140319035,98.69542561145643,98.69542561155643,98.70201929287705,98.70201929297706,98.70778491702107,98.70778491712107,98.71354541826757,98.71354541836757,98.72094674418844,98.72094674428844,98.72916324132825,98.72916324142825,98.73408863146231,98.73408863156232,98.74393385535693,98.74393385545693,98.74885156114468,98.74885156124468,98.75376532624202,98.75376532634202,98.75703870014091,98.75703870024091,98.76194348046411,98.76194348056411,98.76357685723362,98.76357685733362,98.76765712690158,98.76765712700158,98.77254913938783,98.77254913948784,98.7831399813156,98.7831399814156,98.78720965052636,98.78720965062637,98.79208924983782,98.79208924993782,98.79777814873948,98.79777814883948,98.80589865066082,98.80589865076082,98.81076674315626,98.81076674325627,98.81644062802746,98.81644062812747,98.82696897060528,98.82696897070528,98.83263343722639,98.8326334373264,98.83990988266805,98.83990988276805,98.84798676114652,98.84798676124652,98.85282722940217,98.85282722950217,98.8600815281637,98.8600815282637,98.86652451758292,98.86652451768292,98.87054681969491,98.87054681979491,98.87536825441585,98.87536825451585,98.87938243870559,98.87938243880559,98.88259150053693,98.88259150063693,98.88900398901234,98.88900398911234,98.89220678514593,98.89220678524593,98.89620729523993,98.89620729533993,98.90579901452026,98.90579901462026,98.91298600628954,98.91298600638954,98.9193685061109,98.9193685062109,98.92415112503127,98.92415112513127,98.92893003970266,98.92893003980267,98.9345004814505,98.9345004815505,98.93927143592725,98.93927143602725,98.9424489722353,98.9424489723353,98.946417129261,98.946417129361,98.95276219252342,98.95276219262342,98.96227036916699,98.96227036926699,98.96781220150243,98.96781220160243,98.97493082229303,98.97493082239303,98.9788823496052,98.9788823497052,98.98519939455879,98.98519939465879,98.99072160225597,98.99072160235598,98.99545050956601,98.99545050966601,98.99860081570328,98.99860081580329,99.00489563516295,99.00489563526295,99.01039880932544,99.01039880942544,99.01746821089277,99.01746821099277,99.01982198527678,99.01982198537678,99.02531042125712,99.02531042135712,99.03236130157951,99.03236130167952,99.03784122566856,99.03784122576856,99.04487981103686,99.04487981113687,99.0503490952618,99.0503490953618,99.05659359210487,99.05659359220488,99.06283099570521,99.06283099580521,99.06516778571353,99.06516778581353,99.07139498292568,99.07139498302568,99.07528377444646,99.07528377454646,99.08227463805386,99.08227463815386,99.08925955121873,99.08925955131873,99.0962396173344,99.0962396174344,99.10088699175978,99.10088699185978,99.10553042549475,99.10553042559475,99.11403624812472,99.11403624822472,99.11789843998521,99.11789844008521,99.12484548311521,99.12484548321521,99.13641468025195,99.13641468035195,99.14334942842785,99.14334942852786,99.14642809281342,99.14642809291342,99.15027278800848,99.15027278810848,99.15718724162876,99.15718724172876,99.16179174134244,99.16179174144244,99.16562374751436,99.16562374761436,99.17175022067956,99.17175022077956,99.17863622251504,99.17863622261504,99.18551765314966,99.18551765324966,99.18933697029844,99.18933697039844,99.19315254379134,99.19315254389134,99.20001553199482,99.20001553209482,99.205346182731,99.20534618283101,99.20991139376126,99.20991139386126,99.21295175463601,99.21295175473601,99.22358176702649,99.22358176712649,99.23116873887274,99.23116873897274,99.23647491792146,99.23647491802146,99.24253412350129,99.24253412360129,99.24783025378945,99.24783025388945,99.25161043988243,99.25161043998243,99.25840797321814,99.25840797331814,99.26595238560793,99.26595238570793,99.27047397319531,99.27047397329531,99.2757446466234,99.2757446467234,99.28326978903706,99.28326978913707,99.28928206099407,99.28928206109407,99.2960405420878,99.2960405421878,99.29904074732723,99.29904074742723,99.30728534158968,99.30728534168968,99.31402455270728,99.31402455280728,99.32225421175308,99.32225421185308,99.32972966206061,99.32972966216062,99.3371988072635,99.3371988073635,99.34764656272236,99.34764656282236,99.35211861583124,99.35211861593125,99.35881908996207,99.35881909006207,99.36923208853148,99.36923208863148,99.37591955838394,99.37591955848394,99.38037454830334,99.38037454840334,99.38630903103159,99.38630903113159,99.39297991057741,99.39297991067741,99.39890450217271,99.39890450227271,99.4048231827324,99.4048231828324,99.41221445989272,99.41221445999273,99.41664521456607,99.41664521466608,99.42107222558353,99.42107222568353,99.4254956111658,99.4254956112658,99.42770495924614,99.42770495934614,99.43285599331328,99.43285599341328,99.4372665322448,99.43726653234481,99.44093842816523,99.44093842826523,99.44607539396758,99.44607539406758,99.45487073912223,99.45487073922223,99.46145973112127,99.46145973122127,99.47023832834167,99.47023832844167,99.4746231739717,99.4746231740717,99.4790040000975,99.4790040001975,99.48411018910772,99.48411018920773,99.489941071085,99.489941071185,99.49867978851842,99.49867978861842,99.5030451277309,99.5030451278309,99.50740625040464,99.50740625050464,99.51394086087596,99.51394086097596,99.52191989201147,99.52191989211147,99.52554256870862,99.52554256880862,99.532058657935,99.532058658035,99.53784477366916,99.53784477376917,99.54145826855766,99.54145826865766,99.5472354389246,99.5472354390246,99.55012152176965,99.55012152186966,99.55660707064533,99.55660707074533,99.56524801954959,99.56524801964959,99.57316139878279,99.57316139888279,99.57675495377778,99.57675495387778,99.58106362586213,99.58106362596213,99.58823800742208,99.58823800752208,99.59468911466354,99.59468911476354,99.5982700594492,99.59827005954921,99.60327879518552,99.60327879528552,99.60685308020439,99.60685308030439,99.61256641138662,99.61256641148663,99.61613336672133,99.61613336682133,99.61969677543466,99.61969677553466,99.62895227500891,99.62895227510892,99.63535475413066,99.63535475423066,99.64104034285239,99.6410403429524,99.6474299753234,99.6474299754234,99.6552309266591,99.6552309267591,99.66160554509965,99.66160554519965,99.6693898273149,99.6693898274149,99.67433905862596,99.67433905872596,99.68210931198334,99.68210931208334,99.69339998112164,99.69339998122165,99.69762948473559,99.69762948483559,99.70467268071167,99.70467268081167,99.71100233587458,99.71100233597458,99.72013709269456,99.72013709279456,99.72785966369794,99.72785966379794,99.73487235873023,99.73487235883023,99.74328100393461,99.74328100403461,99.74678132219442,99.74678132229442,99.75097717231222,99.75097717241222,99.75726485912223,99.75726485922223,99.76424508286551,99.76424508296552,99.76912677074289,99.76912677084289,99.77260986818558,99.77260986828558,99.7781789700987,99.7781789701987,99.78443785046179,99.78443785056179,99.78930141116325,99.78930141126325,99.79485411980423,99.79485411990423,99.80040174495458,99.80040174505459,99.80524997637033,99.80524997647034,99.8094007449898,99.8094007450898,99.81562057311089,99.8156205732109,99.82459699299005,99.82459699309005,99.8314948168968,99.8314948169968,99.83838806960264,99.83838806970265,99.84389648488339,99.84389648498339,99.85077665449707,99.85077665459707,99.85627517864486,99.85627517874487,99.86176806760537,99.86176806770537,99.86519717758941,99.86519717768941,99.86725272052291,99.86725272062291,99.87341485693635,99.87341485703635,99.87820240171973,99.87820240181973,99.88435303131716,99.88435303141716,99.88844910315369,99.88844910325369,99.89322301314826,99.89322301324826,99.90207999070098,99.90207999080098,99.91092814110718,99.91092814120718,99.91908682831863,99.91908682841863,99.92451966115723,99.92451966125724,99.92723318116907,99.92723318126907,99.93604401323705,99.93604401333705,99.94619976353627,99.94619976363627,99.95296289464467,99.95296289474467,99.95904330113893,99.95904330123894,99.96309236053882,99.96309236063883,99.96848665342519,99.96848665352519,99.97118003650903,99.97118003660903,99.97588892511176,99.97588892521176,99.98059355776884,99.98059355786884,99.98529216116961,99.98529216126961,99.98663081370307,99.98663081380307,99.98729785436937,99.98729785446938,99.98796103315907,99.98796103325907,99.98859524787422,99.98859524797422,99.98917456877192,99.98917456887192,99.98968354834572,99.98968354844573,99.99014011673704,99.99014011683704,99.9905715435235,99.9905715436235,99.99091749673491,99.99091749683491,99.99124654438444,99.99124654448444,99.99155600680261,99.99155600690261,99.9921639765196,99.99216397661961,99.99245807024518,99.99245807034518,99.99273982960975,99.99273982970975,99.99301126436542,99.99301126446542,99.99327001009797,99.99327001019797,99.99350191972881,99.99350191982882,99.9937171208323,99.9937171209323,99.993912500263,99.993912500363,99.99407588128754,99.99407588138754,99.99407615713586,99.99407615723587]],"filenames":["B6Eve","B6Eve\_broken"]}

{"coord\_y":[[9177662,9177662,9177662,8545921,8545921,8105515,8105515,7678524,7678524,7576305,7576305,7378928,7378928,7044041,7044041,7031885,7031885,6717911,6717911,6568745,6568745,6065545,6065545,5726933,5726933,5492634,5492634,5484239,5484239,5331103,5331103,5239597,5239597,5153703,5153703,5032164,5032164,4995302,4995302,4930397,4930397,4894719,4894719,4737590,4737590,4687971,4687971,4610929,4610929,4599233,4599233,4577433,4577433,4487000,4487000,4460343,4460343,4339348,4339348,4322873,4322873,4322607,4322607,4282844,4282844,4268630,4268630,4254679,4254679,4243097,4243097,4212005,4212005,4197143,4197143,4190444,4190444,4132050,4132050,4121404,4121404,4120018,4120018,4093307,4093307,4059383,4059383,4052957,4052957,4014129,4014129,3991828,3991828,3955295,3955295,3931750,3931750,3917426,3917426,3902112,3902112,3892790,3892790,3838091,3838091,3830475,3830475,3817316,3817316,3815258,3815258,3806206,3806206,3803369,3803369,3799092,3799092,3779752,3779752,3775352,3775352,3769984,3769984,3765926,3765926,3749294,3749294,3728928,3728928,3708854,3708854,3708510,3708510,3697295,3697295,3694626,3694626,3689986,3689986,3686968,3686968,3663345,3663345,3638833,3638833,3606267,3606267,3598289,3598289,3559489,3559489,3556693,3556693,3521952,3521952,3517958,3517958,3514244,3514244,3504872,3504872,3502077,3502077,3475268,3475268,3473192,3473192,3433515,3433515,3432172,3432172,3430570,3430570,3418776,3418776,3416165,3416165,3412672,3412672,3407297,3407297,3406991,3406991,3404775,3404775,3351511,3351511,3349211,3349211,3337798,3337798,3322174,3322174,3321597,3321597,3317426,3317426,3312148,3312148,3311016,3311016,3307065,3307065,3276704,3276704,3274586,3274586,3271197,3271197,3267823,3267823,3243538,3243538,3232772,3232772,3212681,3212681,3210577,3210577,3201269,3201269,3192631,3192631,3190374,3190374,3132265,3132265,3132065,3132065,3130918,3130918,3128809,3128809,3126809,3126809,3126350,3126350,3124726,3124726,3078505,3078505,3076147,3076147,3073465,3073465,3025583,3025583,3014745,3014745,3012150,3012150,3009817,3009817,3005418,3005418,3000898,3000898,2997064,2997064,2994220,2994220,2993266,2993266,2989348,2989348,2980124,2980124,2980015,2980015,2977842,2977842,2941234,2941234,2924104,2924104,2918614,2918614,2910562,2910562,2895199,2895199,2887686,2887686,2882559,2882559,2873313,2873313,2866269,2866269,2851370,2851370,2842036,2842036,2841561,2841561,2839786,2839786,2834966,2834966,2827103,2827103,2802283,2802283,2795603,2795603,2795403,2795403,2794252,2794252,2791386,2791386,2780916,2780916,2772159,2772159,2771104,2771104,2760613,2760613,2756147,2756147,2756120,2756120,2745829,2745829,2742641,2742641,2728229,2728229,2723223,2723223,2716440,2716440,2711999,2711999,2706266,2706266,2689651,2689651,2675702,2675702,2668282,2668282,2667768,2667768,2658370,2658370,2651871,2651871,2650157,2650157,2643903,2643903,2642853,2642853,2624267,2624267,2624248,2624248,2617590,2617590,2603784,2603784,2593181,2593181,2583820,2583820,2583408,2583408,2576963,2576963,2573282,2573282,2560632,2560632,2544737,2544737,2544111,2544111,2530857,2530857,2523272,2523272,2521447,2521447,2511634,2511634,2508805,2508805,2500778,2500778,2500275,2500275,2494664,2494664,2489798,2489798,2474762,2474762,2471936,2471936,2461260,2461260,2455247,2455247,2452784,2452784,2449995,2449995,2443779,2443779,2443407,2443407,2440898,2440898,2420981,2420981,2416063,2416063,2414770,2414770,2412943,2412943,2402324,2402324,2397047,2397047,2385965,2385965,2383102,2383102,2379882,2379882,2372340,2372340,2362326,2362326,2358386,2358386,2350464,2350464,2348559,2348559,2346215,2346215,2338545,2338545,2331880,2331880,2328218,2328218,2327577,2327577,2317381,2317381,2316702,2316702,2309639,2309639,2306952,2306952,2301170,2301170,2296669,2296669,2296294,2296294,2295112,2295112,2294499,2294499,2293484,2293484,2288634,2288634,2286899,2286899,2286284,2286284,2286015,2286015,2285735,2285735,2281879,2281879,2272901,2272901,2271386,2271386,2270123,2270123,2268893,2268893,2265226,2265226,2264978,2264978,2256195,2256195,2252146,2252146,2249058,2249058,2239984,2239984,2231669,2231669,2227605,2227605,2220894,2220894,2218949,2218949,2210495,2210495,2202575,2202575,2202519,2202519,2189003,2189003,2187457,2187457,2174888,2174888,2172107,2172107,2164150,2164150,2154790,2154790,2152358,2152358,2146480,2146480,2144492,2144492,2144338,2144338,2140171,2140171,2134174,2134174,2127527,2127527,2123602,2123602,2123094,2123094,2122261,2122261,2118515,2118515,2116945,2116945,2113871,2113871,2113099,2113099,2105097,2105097,2104306,2104306,2097739,2097739,2081089,2081089,2077920,2077920,2071952,2071952,2071923,2071923,2071400,2071400,2069509,2069509,2067256,2067256,2066989,2066989,2061832,2061832,2060156,2060156,2060098,2060098,2059482,2059482,2057946,2057946,2054864,2054864,2054812,2054812,2052044,2052044,2047492,2047492,2045570,2045570,2043046,2043046,2040752,2040752,2035964,2035964,2026746,2026746,2024728,2024728,2019778,2019778,2016279,2016279,2016141,2016141,2015861,2015861,2002346,2002346,2001001,2001001,1996023,1996023,1994860,1994860,1991308,1991308,1988578,1988578,1988510,1988510,1986541,1986541,1975709,1975709,1966462,1966462,1965509,1965509,1962210,1962210,1956557,1956557,1951019,1951019,1943830,1943830,1941283,1941283,1940234,1940234,1939680,1939680,1929652,1929652,1929047,1929047,1922563,1922563,1922033,1922033,1918314,1918314,1916270,1916270,1911814,1911814,1911731,1911731,1910899,1910899,1906360,1906360,1905676,1905676,1904056,1904056,1898073,1898073,1886477,1886477,1883902,1883902,1874736,1874736,1872784,1872784,1863127,1863127,1861138,1861138,1855142,1855142,1851959,1851959,1844968,1844968,1843082,1843082,1842844,1842844,1842549,1842549,1842200,1842200,1841691,1841691,1823858,1823858,1817481,1817481,1816196,1816196,1814273,1814273,1810922,1810922,1806830,1806830,1804357,1804357,1802899,1802899,1798623,1798623,1791305,1791305,1785297,1785297,1781834,1781834,1774795,1774795,1774084,1774084,1768111,1768111,1765749,1765749,1762213,1762213,1761786,1761786,1755144,1755144,1754975,1754975,1752047,1752047,1750167,1750167,1747078,1747078,1746514,1746514,1745345,1745345,1744263,1744263,1740988,1740988,1740466,1740466,1740277,1740277,1738364,1738364,1735414,1735414,1731481,1731481,1726284,1726284,1723102,1723102,1722951,1722951,1721867,1721867,1721040,1721040,1718233,1718233,1718060,1718060,1713434,1713434,1713349,1713349,1711128,1711128,1708918,1708918,1705511,1705511,1705237,1705237,1701917,1701917,1699969,1699969,1699075,1699075,1688141,1688141,1687570,1687570,1686065,1686065,1685643,1685643,1685095,1685095,1683730,1683730,1682013,1682013,1679784,1679784,1677607,1677607,1672525,1672525,1672200,1672200,1670635,1670635,1665921,1665921,1664440,1664440,1662798,1662798,1662703,1662703,1662019,1662019,1659368,1659368,1659110,1659110,1658184,1658184,1656392,1656392,1654374,1654374,1649774,1649774,1641521,1641521,1639876,1639876,1638175,1638175,1637215,1637215,1635474,1635474,1628599,1628599,1624655,1624655,1620797,1620797,1608723,1608723,1605987,1605987,1597597,1597597,1597152,1597152,1597058,1597058,1596447,1596447,1591866,1591866,1580807,1580807,1580488,1580488,1578664,1578664,1573842,1573842,1570495,1570495,1561813,1561813,1553720,1553720,1552537,1552537,1551827,1551827,1548852,1548852,1548666,1548666,1547340,1547340,1545223,1545223,1544196,1544196,1544180,1544180,1542725,1542725,1541621,1541621,1537847,1537847,1533216,1533216,1533059,1533059,1527337,1527337,1527068,1527068,1518981,1518981,1517873,1517873,1515099,1515099,1515067,1515067,1512256,1512256,1509615,1509615,1506673,1506673,1504694,1504694,1503217,1503217,1502011,1502011,1500402,1500402,1499001,1499001,1495695,1495695,1495575,1495575,1491376,1491376,1487334,1487334,1485427,1485427,1485388,1485388,1481575,1481575,1480948,1480948,1480325,1480325,1480278,1480278,1479297,1479297,1476014,1476014,1472333,1472333,1471469,1471469,1471204,1471204,1470551,1470551,1470360,1470360,1467273,1467273,1466774,1466774,1466642,1466642,1465412,1465412,1462511,1462511,1453002,1453002,1449242,1449242,1444715,1444715,1438204,1438204,1433571,1433571,1431707,1431707,1430578,1430578,1421471,1421471,1421449,1421449,1419391,1419391,1417510,1417510,1416945,1416945,1416180,1416180,1415807,1415807,1414324,1414324,1414261,1414261,1413116,1413116,1411414,1411414,1410598,1410598,1409769,1409769,1406926,1406926,1406704,1406704,1404122,1404122,1399307,1399307,1391461,1391461,1390099,1390099,1387096,1387096,1385631,1385631,1380258,1380258,1377236,1377236,1375582,1375582,1374901,1374901,1369807,1369807,1368232,1368232,1363105,1363105,1359163,1359163,1357890,1357890,1357445,1357445,1357052,1357052,1356598,1356598,1353923,1353923,1353048,1353048,1352934,1352934,1352121,1352121,1351940,1351940,1351529,1351529,1350615,1350615,1350352,1350352,1350163,1350163,1348701,1348701,1347240,1347240,1346467,1346467,1345607,1345607,1344384,1344384,1342535,1342535,1341402,1341402,1338940,1338940,1337112,1337112,1336607,1336607,1335995,1335995,1334556,1334556,1332753,1332753,1332375,1332375,1331669,1331669,1330274,1330274,1321765,1321765,1320175,1320175,1318842,1318842,1316375,1316375,1316233,1316233,1316073,1316073,1314580,1314580,1312790,1312790,1312554,1312554,1312025,1312025,1307281,1307281,1301385,1301385,1300822,1300822,1298261,1298261,1296626,1296626,1295715,1295715,1294502,1294502,1294242,1294242,1293986,1293986,1293304,1293304,1291461,1291461,1291016,1291016,1290461,1290461,1290161,1290161,1288327,1288327,1287838,1287838,1286271,1286271,1285220,1285220,1284834,1284834,1284780,1284780,1282305,1282305,1280671,1280671,1279363,1279363,1279127,1279127,1278898,1278898,1278787,1278787,1277791,1277791,1277765,1277765,1276382,1276382,1275837,1275837,1273293,1273293,1272028,1272028,1271730,1271730,1271464,1271464,1271346,1271346,1270729,1270729,1270170,1270170,1269444,1269444,1262549,1262549,1262037,1262037,1260998,1260998,1260926,1260926,1260056,1260056,1257979,1257979,1257774,1257774,1257206,1257206,1255178,1255178,1252132,1252132,1252068,1252068,1251675,1251675,1249245,1249245,1246712,1246712,1242196,1242196,1241286,1241286,1241175,1241175,1240076,1240076,1238333,1238333,1237891,1237891,1236961,1236961,1236184,1236184,1228171,1228171,1226500,1226500,1222956,1222956,1222381,1222381,1222089,1222089,1220958,1220958,1220046,1220046,1218810,1218810,1218138,1218138,1215373,1215373,1213861,1213861,1208803,1208803,1206953,1206953,1206906,1206906,1202220,1202220,1202183,1202183,1201365,1201365,1199827,1199827,1198140,1198140,1197376,1197376,1196171,1196171,1195858,1195858,1194811,1194811,1190010,1190010,1189851,1189851,1189599,1189599,1188079,1188079,1187703,1187703,1187554,1187554,1187257,1187257,1185330,1185330,1185309,1185309,1184512,1184512,1183781,1183781,1182785,1182785,1181150,1181150,1181048,1181048,1180986,1180986,1180180,1180180,1179944,1179944,1176652,1176652,1175600,1175600,1172738,1172738,1172265,1172265,1168633,1168633,1167229,1167229,1166723,1166723,1163245,1163245,1162688,1162688,1162613,1162613,1161834,1161834,1160842,1160842,1158648,1158648,1156236,1156236,1155986,1155986,1155060,1155060,1153750,1153750,1150817,1150817,1149709,1149709,1147293,1147293,1146797,1146797,1145001,1145001,1143663,1143663,1140915,1140915,1140368,1140368,1139470,1139470,1139021,1139021,1138081,1138081,1135645,1135645,1134074,1134074,1133507,1133507,1127378,1127378,1126712,1126712,1124836,1124836,1124753,1124753,1123857,1123857,1123754,1123754,1119028,1119028,1115116,1115116,1114433,1114433,1112280,1112280,1112176,1112176,1110820,1110820,1108895,1108895,1107978,1107978,1107636,1107636,1107325,1107325,1106254,1106254,1103190,1103190,1102824,1102824,1100026,1100026,1099047,1099047,1098353,1098353,1095919,1095919,1093330,1093330,1092934,1092934,1089734,1089734,1088594,1088594,1087160,1087160,1085851,1085851,1085504,1085504,1082835,1082835,1080417,1080417,1078239,1078239,1073572,1073572,1073356,1073356,1072548,1072548,1067302,1067302,1063401,1063401,1062006,1062006,1061725,1061725,1060522,1060522,1058970,1058970,1057958,1057958,1057682,1057682,1056119,1056119,1053863,1053863,1053813,1053813,1051294,1051294,1050984,1050984,1050937,1050937,1050743,1050743,1049608,1049608,1049048,1049048,1048849,1048849,1046683,1046683,1046410,1046410,1044720,1044720,1041112,1041112,1040497,1040497,1038051,1038051,1035523,1035523,1034114,1034114,1032347,1032347,1031286,1031286,1030862,1030862,1028854,1028854,1028438,1028438,1023362,1023362,1021228,1021228,1019935,1019935,1016571,1016571,1016001,1016001,1014672,1014672,1014171,1014171,1014132,1014132,1009918,1009918,1007924,1007924,1007906,1007906,1003876,1003876,1001487,1001487,999106,999106,995035,995035,994554,994554,993874,993874,993740,993740,990609,990609,990084,990084,987278,987278,985330,985330,983247,983247,981713,981713,980783,980783,980208,980208,978741,978741,978202,978202,973650,973650,972864,972864,972090,972090,969591,969591,962607,962607,961611,961611,961010,961010,960462,960462,959767,959767,959745,959745,959618,959618,958807,958807,958109,958109,953933,953933,950767,950767,949563,949563,949453,949453,945019,945019,940948,940948,940858,940858,940841,940841,939436,939436,938129,938129,938001,938001,937590,937590,936808,936808,933972,933972,933147,933147,932468,932468,932377,932377,932313,932313,932027,932027,929064,929064,927538,927538,926120,926120,923829,923829,921357,921357,921193,921193,920932,920932,918190,918190,917248,917248,916816,916816,916327,916327,914827,914827,913186,913186,912523,912523,909830,909830,908957,908957,907795,907795,906887,906887,906336,906336,904818,904818,903339,903339,902851,902851,902303,902303,902152,902152,901692,901692,901554,901554,899121,899121,897340,897340,895473,895473,894309,894309,894158,894158,889257,889257,887240,887240,887080,887080,886585,886585,885925,885925,885209,885209,883473,883473,883306,883306,882624,882624,878057,878057,877353,877353,876897,876897,873403,873403,871145,871145,869660,869660,867503,867503,866910,866910,865411,865411,864687,864687,864593,864593,864503,864503,863930,863930,861960,861960,861699,861699,860545,860545,860309,860309,858861,858861,857616,857616,855626,855626,854521,854521,853763,853763,851374,851374,850743,850743,847453,847453,847126,847126,847070,847070,845480,845480,845043,845043,843840,843840,843274,843274,841747,841747,840899,840899,840735,840735,839697,839697,839051,839051,838483,838483,837762,837762,836717,836717,836153,836153,835799,835799,831589,831589,831572,831572,826935,826935,826726,826726,825963,825963,825565,825565,824922,824922,824885,824885,823419,823419,823110,823110,822694,822694,822157,822157,819863,819863,819467,819467,816431,816431,816304,816304,814048,814048,813907,813907,813623,813623,812734,812734,812578,812578,810915,810915,810779,810779,809341,809341,808166,808166,807155,807155,805955,805955,804859,804859,803320,803320,801299,801299,799090,799090,797600,797600,794621,794621,793532,793532,792006,792006,790201,790201,790122,790122,789177,789177,788506,788506,788336,788336,787821,787821,787015,787015,786885,786885,785525,785525,784140,784140,783859,783859,779907,779907,779122,779122,778789,778789,777494,777494,774835,774835,773453,773453,772750,772750,772395,772395,771857,771857,771518,771518,771352,771352,770630,770630,769956,769956,769665,769665,769481,769481,767553,767553,762721,762721,761852,761852,760707,760707,760017,760017,758343,758343,755180,755180,755025,755025,754688,754688,752306,752306,750124,750124,748735,748735,748716,748716,748389,748389,746754,746754,744485,744485,744100,744100,736471,736471,735496,735496,733100,733100,730878,730878,726979,726979,724444,724444,723735,723735,723286,723286,720866,720866,719094,719094,716995,716995,715277,715277,709802,709802,709581,709581,709034,709034,708416,708416,707418,707418,707274,707274,705869,705869,705252,705252,703521,703521,703088,703088,701794,701794,700054,700054,698769,698769,696077,696077,693287,693287,692805,692805,692788,692788,692632,692632,691453,691453,690607,690607,688708,688708,687301,687301,685847,685847,684032,684032,683807,683807,683770,683770,682512,682512,680406,680406,679362,679362,677205,677205,675822,675822,675322,675322,672315,672315,671734,671734,671596,671596,666337,666337,666083,666083,665798,665798,665721,665721,664270,664270,663651,663651,663520,663520,663248,663248,663229,663229,661857,661857,660769,660769,659208,659208,657764,657764,656889,656889,654803,654803,654402,654402,652801,652801,652078,652078,649304,649304,647220,647220,645032,645032,644246,644246,643620,643620,642404,642404,640592,640592,640464,640464,638736,638736,635131,635131,631542,631542,630293,630293,628261,628261,628196,628196,626161,626161,625812,625812,625453,625453,625205,625205,625112,625112,624917,624917,623692,623692,623375,623375,623176,623176,622697,622697,621517,621517,619883,619883,619413,619413,618003,618003,617869,617869,615646,615646,615506,615506,611748,611748,610005,610005,608470,608470,608308,608308,608208,608208,605825,605825,605532,605532,604853,604853,603879,603879,602467,602467,601994,601994,601009,601009,596024,596024,595612,595612,595577,595577,594954,594954,594595,594595,594058,594058,593886,593886,593076,593076,590371,590371,589091,589091,588915,588915,588750,588750,588226,588226,586960,586960,586206,586206,585634,585634,585193,585193,584956,584956,583043,583043,582206,582206,580706,580706,580388,580388,579844,579844,578281,578281,577594,577594,576465,576465,575044,575044,574781,574781,574416,574416,572249,572249,571318,571318,569574,569574,569058,569058,566816,566816,566780,566780,565760,565760,562343,562343,562081,562081,561460,561460,559686,559686,559629,559629,559125,559125,558861,558861,558400,558400,557007,557007,555852,555852,555678,555678,555360,555360,555012,555012,554691,554691,554610,554610,554333,554333,553973,553973,553224,553224,552029,552029,551267,551267,549479,549479,548613,548613,547079,547079,546837,546837,546641,546641,545738,545738,544658,544658,541303,541303,540677,540677,537764,537764,536047,536047,535984,535984,533681,533681,533551,533551,533467,533467,532914,532914,532755,532755,532511,532511,531996,531996,530531,530531,528671,528671,528255,528255,528075,528075,527007,527007,526965,526965,526607,526607,525619,525619,523796,523796,522152,522152,519173,519173,518244,518244,516707,516707,515562,515562,514874,514874,514299,514299,511817,511817,510313,510313,509488,509488,508504,508504,508110,508110,507173,507173,505410,505410,505074,505074,504478,504478,504233,504233,503730,503730,503591,503591,503273,503273,502411,502411,500869,500869,500094,500094,499940,499940,499739,499739,498503,498503,497231,497231,496692,496692,495143,495143,494622,494622,493755,493755,492848,492848,491581,491581,490838,490838,490790,490790,490756,490756,488337,488337,487924,487924,487176,487176,483260,483260,482796,482796,482266,482266,482048,482048,481174,481174,479816,479816,478033,478033,477749,477749,475561,475561,474894,474894,473442,473442,472681,472681,469507,469507,468180,468180,467688,467688,464828,464828,464164,464164,464116,464116,462625,462625,462091,462091,461170,461170,460862,460862,460638,460638,459458,459458,457092,457092,456719,456719,455494,455494,455417,455417,453241,453241,452790,452790,450993,450993,448882,448882,448204,448204,446281,446281,446243,446243,445741,445741,443960,443960,443765,443765,443293,443293,443187,443187,443078,443078,442817,442817,442436,442436,441699,441699,440546,440546,439650,439650,439613,439613,436189,436189,435018,435018,431637,431637,430831,430831,430737,430737,430141,430141,426746,426746,426097,426097,425275,425275,425228,425228,425058,425058,423903,423903,423752,423752,420972,420972,420915,420915,419965,419965,418601,418601,416973,416973,416603,416603,416034,416034,415663,415663,413589,413589,413253,413253,412418,412418,411502,411502,411463,411463,411302,411302,411135,411135,411091,411091,410201,410201,409501,409501,409357,409357,408975,408975,408827,408827,404376,404376,404056,404056,402873,402873,402574,402574,401127,401127,400069,400069,398767,398767,397832,397832,395935,395935,395093,395093,394237,394237,393910,393910,393371,393371,392865,392865,392499,392499,392093,392093,390237,390237,389274,389274,388025,388025,386482,386482,383331,383331,382944,382944,382493,382493,381995,381995,381727,381727,381052,381052,379942,379942,379832,379832,378073,378073,377751,377751,376721,376721,376107,376107,375990,375990,375839,375839,374859,374859,374824,374824,374462,374462,371991,371991,371610,371610,370487,370487,370145,370145,368820,368820,367447,367447,367236,367236,366727,366727,364970,364970,364563,364563,363274,363274,361106,361106,360670,360670,359002,359002,358894,358894,358618,358618,357828,357828,356759,356759,356740,356740,356273,356273,356170,356170,355113,355113,354495,354495,354458,354458,354354,354354,353473,353473,352434,352434,352153,352153,349351,349351,349060,349060,348063,348063,347806,347806,346800,346800,345135,345135,344350,344350,344221,344221,343910,343910,343522,343522,343460,343460,343402,343402,342852,342852,341843,341843,341678,341678,341110,341110,340375,340375,340350,340350,339872,339872,338320,338320,335515,335515,334205,334205,333774,333774,333381,333381,332991,332991,332730,332730,332696,332696,332229,332229,331734,331734,331326,331326,330309,330309,329558,329558,329125,329125,328190,328190,326141,326141,326050,326050,323608,323608,323266,323266,322836,322836,322781,322781,321830,321830,320987,320987,320951,320951,320897,320897,320115,320115,319643,319643,319308,319308,318348,318348,316969,316969,316904,316904,316209,316209,316038,316038,315313,315313,313596,313596,312075,312075,311769,311769,311622,311622,311259,311259,311118,311118,310851,310851,310812,310812,310658,310658,310507,310507,309987,309987,309639,309639,309124,309124,308809,308809,308581,308581,308541,308541,308274,308274,308122,308122,307918,307918,307747,307747,307163,307163,307081,307081,306954,306954,306491,306491,306022,306022,305593,305593,304365,304365,303663,303663,303579,303579,302677,302677,301766,301766,300298,300298,299730,299730,299559,299559,299478,299478,298723,298723,298452,298452,297608,297608,297585,297585,296715,296715,296591,296591,296079,296079,295701,295701,295046,295046,294547,294547,293481,293481,292266,292266,292063,292063,291472,291472,291367,291367,291063,291063,290825,290825,290592,290592,289990,289990,289577,289577,288593,288593,287922,287922,287143,287143,286884,286884,286825,286825,286673,286673,286537,286537,285811,285811,285621,285621,285596,285596,284820,284820,284147,284147,283872,283872,283157,283157,282700,282700,282536,282536,282310,282310,281918,281918,280767,280767,280245,280245,280095,280095,279862,279862,279831,279831,278727,278727,278534,278534,278380,278380,278313,278313,278190,278190,277629,277629,277289,277289,277196,277196,276871,276871,276812,276812,276714,276714,276161,276161,275578,275578,275487,275487,275093,275093,275001,275001,274934,274934,274690,274690,274387,274387,273953,273953,273500,273500,273382,273382,273334,273334,272908,272908,272722,272722,270942,270942,270832,270832,270618,270618,270322,270322,269753,269753,269590,269590,269195,269195,269024,269024,268549,268549,268043,268043,265568,265568,265414,265414,265268,265268,265222,265222,265096,265096,264108,264108,263911,263911,263071,263071,262583,262583,262509,262509,262436,262436,262083,262083,262007,262007,261765,261765,261383,261383,261310,261310,261287,261287,261247,261247,260710,260710,260416,260416,260119,260119,259644,259644,259062,259062,258878,258878,257544,257544,257477,257477,256979,256979,256880,256880,256807,256807,256763,256763,256531,256531,256333,256333,256036,256036,256019,256019,255847,255847,255617,255617,255189,255189,255121,255121,255072,255072,254750,254750,254507,254507,254336,254336,254005,254005,253819,253819,253458,253458,252565,252565,252289,252289,251184,251184,250936,250936,250897,250897,250212,250212,249770,249770,249729,249729,249521,249521,249411,249411,249272,249272,248965,248965,248873,248873,248760,248760,248739,248739,248606,248606,247973,247973,247549,247549,247083,247083,246941,246941,246664,246664,246532,246532,246360,246360,246274,246274,246169,246169,244942,244942,244725,244725,244706,244706,244689,244689,244556,244556,243532,243532,243456,243456,243125,243125,243051,243051,242397,242397,242199,242199,242001,242001,241920,241920,241707,241707,241299,241299,241045,241045,240910,240910,240646,240646,239971,239971,239848,239848,239806,239806,238883,238883,238760,238760,238106,238106,237887,237887,237851,237851,237829,237829,237723,237723,237176,237176,236828,236828,236618,236618,236293,236293,236277,236277,235973,235973,235667,235667,235348,235348,235176,235176,235030,235030,234943,234943,234773,234773,234739,234739,233962,233962,233769,233769,233540,233540,233184,233184,232107,232107,231885,231885,231786,231786,231734,231734,231545,231545,231392,231392,231254,231254,231229,231229,231137,231137,230991,230991,230852,230852,229840,229840,229776,229776,229742,229742,229376,229376,229228,229228,228583,228583,228508,228508,228444,228444,228387,228387,227753,227753,227673,227673,227605,227605,227046,227046,226940,226940,226423,226423,226392,226392,225672,225672,225408,225408,225385,225385,225348,225348,225255,225255,225188,225188,224375,224375,224259,224259,223776,223776,223715,223715,223198,223198,222911,222911,222480,222480,222095,222095,222005,222005,221526,221526,220670,220670,220356,220356,220050,220050,219946,219946,219722,219722,219362,219362,219136,219136,218703,218703,218314,218314,218187,218187,217949,217949,217915,217915,217535,217535,217140,217140,216838,216838,216426,216426,216298,216298,216147,216147,216076,216076,215836,215836,215730,215730,215163,215163,215138,215138,214891,214891,214792,214792,214435,214435,213836,213836,213320,213320,213183,213183,212769,212769,212665,212665,212600,212600,212556,212556,211365,211365,211177,211177,210920,210920,210768,210768,210656,210656,210282,210282,209884,209884,209770,209770,209545,209545,209100,209100,208907,208907,208589,208589,208284,208284,208143,208143,207953,207953,207909,207909,207864,207864,207580,207580,207431,207431,207295,207295,206737,206737,206578,206578,206408,206408,206273,206273,206252,206252,206207,206207,206076,206076,205876,205876,205224,205224,205060,205060,204956,204956,204881,204881,204617,204617,204071,204071,203878,203878,203641,203641,203258,203258,202838,202838,202797,202797,202508,202508,202236,202236,202143,202143,201893,201893,201823,201823,201594,201594,201303,201303,200967,200967,200884,200884,200240,200240,200173,200173,199900,199900,199727,199727,199686,199686,199667,199667,199335,199335,199219,199219,199201,199201,198844,198844,198786,198786,198767,198767,198452,198452,198377,198377,198348,198348,198320,198320,197595,197595,197555,197555,197451,197451,197045,197045,196730,196730,196459,196459,196085,196085,195817,195817,195687,195687,195555,195555,195422,195422,195373,195373,195315,195315,195186,195186,195015,195015,194714,194714,194600,194600,193977,193977,193872,193872,193734,193734,193668,193668,193637,193637,193467,193467,193374,193374,193308,193308,193159,193159,193034,193034,192920,192920,192708,192708,192375,192375,192345,192345,192323,192323,192255,192255,192124,192124,192071,192071,191828,191828,191196,191196,191157,191157,191061,191061,190986,190986,190891,190891,190871,190871,190817,190817,190750,190750,190246,190246,190200,190200,190140,190140,189966,189966,189933,189933,189886,189886,189667,189667,189630,189630,189224,189224,189180,189180,188987,188987,188770,188770,188739,188739,188583,188583,187924,187924,187871,187871,187844,187844,187753,187753,187565,187565,187534,187534,187489,187489,187154,187154,187127,187127,187111,187111,187087,187087,187049,187049,186824,186824,186749,186749,186290,186290,185915,185915,185795,185795,185758,185758,185702,185702,185510,185510,185197,185197,185002,185002,184904,184904,184837,184837,184747,184747,184675,184675,184639,184639,184623,184623,184581,184581,184543,184543,184329,184329,184220,184220,184157,184157,184017,184017,183808,183808,183696,183696,183597,183597,183579,183579,183348,183348,183172,183172,183020,183020,182740,182740,182705,182705,182689,182689,182475,182475,182415,182415,182370,182370,182128,182128,182045,182045,182023,182023,181592,181592,181396,181396,181334,181334,181311,181311,181233,181233,181107,181107,181034,181034,180596,180596,180496,180496,180480,180480,180458,180458,180360,180360,180303,180303,180026,180026,179796,179796,179748,179748,179696,179696,179570,179570,179466,179466,179426,179426,179395,179395,179368,179368,179267,179267,179140,179140,179018,179018,178970,178970,178837,178837,178643,178643,178601,178601,178561,178561,178472,178472,178357,178357,178199,178199,178007,178007,177935,177935,177881,177881,177828,177828,177781,177781,177758,177758,177737,177737,177411,177411,177351,177351,177167,177167,177051,177051,176997,176997,176955,176955,176846,176846,176784,176784,176456,176456,176386,176386,176258,176258,176031,176031,175957,175957,175847,175847,175496,175496,175312,175312,175195,175195,175073,175073,174824,174824,174755,174755,174681,174681,174637,174637,174592,174592,174571,174571,174550,174550,174529,174529,174469,174469,174374,174374,174301,174301,174120,174120,173922,173922,173855,173855,173773,173773,173663,173663,173595,173595,173513,173513,173323,173323,173251,173251,173204,173204,173148,173148,173056,173056,173002,173002,172925,172925,172853,172853,172825,172825,172591,172591,172532,172532,172494,172494,172412,172412,172303,172303,172177,172177,172103,172103,172047,172047,171813,171813,171759,171759,171736,171736,171705,171705,171687,171687,171661,171661,171548,171548,171523,171523,171395,171395,171288,171288,171269,171269,171244,171244,171142,171142,171005,171005,170956,170956,170863,170863,170845,170845,170827,170827,170803,170803,170741,170741,170639,170639,170617,170617,170586,170586,170505,170505,170354,170354,170255,170255,170121,170121,170052,170052,170003,170003,169831,169831,169720,169720,169550,169550,169384,169384,169301,169301,169214,169214,169156,169156,169086,169086,169011,169011,168993,168993,168912,168912,168869,168869,168840,168840,168792,168792,168691,168691,168656,168656,168598,168598,168577,168577,168412,168412,168377,168377,168305,168305,168204,168204,168060,168060,167924,167924,167705,167705,167676,167676,167648,167648,167542,167542,167495,167495,167456,167456,167321,167321,167300,167300,167240,167240,167094,167094,167047,167047,166882,166882,166843,166843,166688,166688,166658,166658,166614,166614,166538,166538,166472,166472,166343,166343,166198,166198,166179,166179,166145,166145,166063,166063,165989,165989,165955,165955,165837,165837,165815,165815,165603,165603,165574,165574,165527,165527,165382,165382,165347,165347,165329,165329,165295,165295,165259,165259,165136,165136,165098,165098,164965,164965,164926,164926,164838,164838,164812,164812,164778,164778,164690,164690,164648,164648,164580,164580,164525,164525,164428,164428,164383,164383,164287,164287,164250,164250,164226,164226,164159,164159,164120,164120,163987,163987,163965,163965,163922,163922,163889,163889,163867,163867,163752,163752,163660,163660,163638,163638,163616,163616,163589,163589,163496,163496,163336,163336,163314,163314,163053,163053,163005,163005,162859,162859,162837,162837,162790,162790,162727,162727,162693,162693,162649,162649,162568,162568,162532,162532,162477,162477,162411,162411,162308,162308,162288,162288,162253,162253,162227,162227,162195,162195,162125,162125,161995,161995,161974,161974,161909,161909,161886,161886,161819,161819,161655,161655,161566,161566,161522,161522,161467,161467,161446,161446,161351,161351,161327,161327,161275,161275,161197,161197,161166,161166,161130,161130,161018,161018,160999,160999,160912,160912,160805,160805,160777,160777,160617,160617,160536,160536,160500,160500,160436,160436,160415,160415,160355,160355,160301,160301,160256,160256,160229,160229,160207,160207,160187,160187,160163,160163,160089,160089,160031,160031,159893,159893,159844,159844,159766,159766,159728,159728,159703,159703,159674,159674,159624,159624,159567,159567,159406,159406,159380,159380,159355,159355,159262,159262,159206,159206,159131,159131,159035,159035,158814,158814,158798,158798,158762,158762,158745,158745,158681,158681,158622,158622,158482,158482,158463,158463,158418,158418,158387,158387,158371,158371,158267,158267,158235,158235,158211,158211,158129,158129,158084,158084,157979,157979,157927,157927,157873,157873,157848,157848,157814,157814,157775,157775,157615,157615,157517,157517,157482,157482,157333,157333,157265,157265,157237,157237,157117,157117,157078,157078,157012,157012,156949,156949,156931,156931,156854,156854,156773,156773,156746,156746,156558,156558,156538,156538,156512,156512,156433,156433,156392,156392,156371,156371,156320,156320,156279,156279,156236,156236,156134,156134,156038,156038,156017,156017,155885,155885,155867,155867,155770,155770,155692,155692,155585,155585,155551,155551,155486,155486,155435,155435,155361,155361,155336,155336,155288,155288,155188,155188,155165,155165,155069,155069,154989,154989,154969,154969,154899,154899,154865,154865,154810,154810,154762,154762,154698,154698,154665,154665,154638,154638,154609,154609,154556,154556,154534,154534,154507,154507,154460,154460,154422,154422,154382,154382,154366,154366,154302,154302,154234,154234,154202,154202,154170,154170,154136,154136,154107,154107,154065,154065,153932,153932,153815,153815,153740,153740,153708,153708,153687,153687,153617,153617,153573,153573,153466,153466,153400,153400,153325,153325,153272,153272,153228,153228,153191,153191,153169,153169,153097,153097,152944,152944,152926,152926,152904,152904,152748,152748,152708,152708,152653,152653,152610,152610,152579,152579,152537,152537,152510,152510,152453,152453,152370,152370,152341,152341,152323,152323,152258,152258,152216,152216,152194,152194,152061,152061,152002,152002,151953,151953,151870,151870,151834,151834,151754,151754,151735,151735,151660,151660,151632,151632,151476,151476,151438,151438,151395,151395,151368,151368,151347,151347,151296,151296,151275,151275,151254,151254,151129,151129,151091,151091,151056,151056,151021,151021,150979,150979,150893,150893,150838,150838,150808,150808,150751,150751,150687,150687,150568,150568,150490,150490,150446,150446,150396,150396,150356,150356,150326,150326,150303,150303,150274,150274,150253,150253,150209,150209,150151,150151,149990,149990,149853,149853,149823,149823,149803,149803,149782,149782,149738,149738,149691,149691,149665,149665,149572,149572,149524,149524,149429,149429,149344,149344,149291,149291,149170,149170,149153,149153,149084,149084,149035,149035,148942,148942,148868,148868,148730,148730,148708,148708,148658,148658,148592,148592,148498,148498,148482,148482,148457,148457,148421,148421,148382,148382,148348,148348,148313,148313,148293,148293,148276,148276,148136,148136,148003,148003,147924,147924,147906,147906,147833,147833,147707,147707,147645,147645,147596,147596,147576,147576,147486,147486,147439,147439,147361,147361,147325,147325,147292,147292,147270,147270,147254,147254,147237,147237,147122,147122,147084,147084,147027,147027,146990,146990,146941,146941,146811,146811,146771,146771,146718,146718,146700,146700,146658,146658,146557,146557,146540,146540,146449,146449,146422,146422,146386,146386,146364,146364,146345,146345,146300,146300,146280,146280,146169,146169,146153,146153,146122,146122,146104,146104,145983,145983,145893,145893,145870,145870,145792,145792,145759,145759,145705,145705,145685,145685,145660,145660,145622,145622,145493,145493,145365,145365,145266,145266,145249,145249,145198,145198,145139,145139,145076,145076,145052,145052,145016,145016,144997,144997,144863,144863,144764,144764,144722,144722,144688,144688,144619,144619,144529,144529,144506,144506,144479,144479,144400,144400,144373,144373,144312,144312,144261,144261,144178,144178,144142,144142,144101,144101,144069,144069,143881,143881,143797,143797,143705,143705,143680,143680,143617,143617,143583,143583,143513,143513,143475,143475,143317,143317,143282,143282,143253,143253,143226,143226,143210,143210,143142,143142,143091,143091,143063,143063,142921,142921,142840,142840,142781,142781,142761,142761,142738,142738,142685,142685,142669,142669,142605,142605,142482,142482,142458,142458,142408,142408,142365,142365,142334,142334,142313,142313,142189,142189,142152,142152,142100,142100,142030,142030,141960,141960,141917,141917,141868,141868,141811,141811,141744,141744,141596,141596,141537,141537,141456,141456,141412,141412,141348,141348,141301,141301,141270,141270,141243,141243,141184,141184,141141,141141,141116,141116,141100,141100,141073,141073,141015,141015,140973,140973,140848,140848,140807,140807,140748,140748,140720,140720,140692,140692,140640,140640,140614,140614,140583,140583,140565,140565,140481,140481,140458,140458,140401,140401,140355,140355,140324,140324,140302,140302,140245,140245,140166,140166,140145,140145,140128,140128,140096,140096,140023,140023,139889,139889,139855,139855,139781,139781,139751,139751,139720,139720,139669,139669,139638,139638,139608,139608,139555,139555,139493,139493,139431,139431,139363,139363,139347,139347,139303,139303,139280,139280,139258,139258,139223,139223,139125,139125,139073,139073,139045,139045,139016,139016,138977,138977,138782,138782,138718,138718,138655,138655,138616,138616,138516,138516,138396,138396,138371,138371,138322,138322,138281,138281,138258,138258,138220,138220,138159,138159,138120,138120,138061,138061,138022,138022,138005,138005,137981,137981,137949,137949,137930,137930,137874,137874,137853,137853,137758,137758,137709,137709,137675,137675,137643,137643,137601,137601,137580,137580,137542,137542,137514,137514,137494,137494,137458,137458,137426,137426,137409,137409,137345,137345,137323,137323,137296,137296,137220,137220,137191,137191,137158,137158,137137,137137,137025,137025,137006,137006,136969,136969,136909,136909,136854,136854,136785,136785,136744,136744,136712,136712,136694,136694,136644,136644,136392,136392,136331,136331,136262,136262,136183,136183,136114,136114,136043,136043,136017,136017,135916,135916,135899,135899,135880,135880,135830,135830,135661,135661,135593,135593,135532,135532,135497,135497,135457,135457,135424,135424,135329,135329,135308,135308,135269,135269,135218,135218,135170,135170,135038,135038,135022,135022,134916,134916,134884,134884,134861,134861,134785,134785,134716,134716,134620,134620,134583,134583,134547,134547,134491,134491,134461,134461,134432,134432,134370,134370,134315,134315,134273,134273,134165,134165,134102,134102,134079,134079,134047,134047,134031,134031,133946,133946,133827,133827,133805,133805,133768,133768,133689,133689,133667,133667,133651,133651,133606,133606,133577,133577,133534,133534,133507,133507,133436,133436,133402,133402,133359,133359,133343,133343,133298,133298,133259,133259,133142,133142,133118,133118,133060,133060,133032,133032,133015,133015,132991,132991,132962,132962,132935,132935,132909,132909,132859,132859,132839,132839,132785,132785,132764,132764,132742,132742,132706,132706,132660,132660,132535,132535,132471,132471,132397,132397,132372,132372,132337,132337,132306,132306,132264,132264,132226,132226,132129,132129,132055,132055,132029,132029,131937,131937,131912,131912,131850,131850,131811,131811,131785,131785,131701,131701,131655,131655,131607,131607,131544,131544,131443,131443,131376,131376,131333,131333,131310,131310,131281,131281,131246,131246,131218,131218,131201,131201,131183,131183,130997,130997,130935,130935,130916,130916,130880,130880,130849,130849,130789,130789,130773,130773,130750,130750,130688,130688,130598,130598,130545,130545,130482,130482,130429,130429,130321,130321,130276,130276,130215,130215,130172,130172,130153,130153,130132,130132,130112,130112,130095,130095,130071,130071,130047,130047,129992,129992,129942,129942,129891,129891,129816,129816,129795,129795,129728,129728,129699,129699,129683,129683,129653,129653,129604,129604,129532,129532,129444,129444,129333,129333,129239,129239,129201,129201,129174,129174,129119,129119,128964,128964,128925,128925,128899,128899,128764,128764,128687,128687,128666,128666,128586,128586,128511,128511,128487,128487,128459,128459,128442,128442,128402,128402,128308,128308,128224,128224,128159,128159,128101,128101,128077,128077,128025,128025,127998,127998,127960,127960,127906,127906,127886,127886,127816,127816,127794,127794,127712,127712,127683,127683,127631,127631,127598,127598,127581,127581,127494,127494,127453,127453,127408,127408,127375,127375,127314,127314,127282,127282,127246,127246,127225,127225,127177,127177,127158,127158,127121,127121,127017,127017,126933,126933,126910,126910,126848,126848,126817,126817,126753,126753,126673,126673,126648,126648,126610,126610,126575,126575,126552,126552,126500,126500,126444,126444,126421,126421,126329,126329,126308,126308,126291,126291,126229,126229,126211,126211,126047,126047,126014,126014,125987,125987,125924,125924,125887,125887,125848,125848,125819,125819,125774,125774,125682,125682,125650,125650,125619,125619,125515,125515,125497,125497,125384,125384,125352,125352,125249,125249,125231,125231,125180,125180,125154,125154,125104,125104,125001,125001,124950,124950,124922,124922,124904,124904,124875,124875,124843,124843,124815,124815,124746,124746,124709,124709,124638,124638,124611,124611,124566,124566,124495,124495,124462,124462,124420,124420,124395,124395,124347,124347,124293,124293,124277,124277,124061,124061,124016,124016,123991,123991,123962,123962,123906,123906,123862,123862,123822,123822,123788,123788,123720,123720,123643,123643,123622,123622,123487,123487,123449,123449,123433,123433,123398,123398,123347,123347,123312,123312,123267,123267,123189,123189,123172,123172,123109,123109,123074,123074,123049,123049,123014,123014,122973,122973,122908,122908,122833,122833,122815,122815,122764,122764,122638,122638,122602,122602,122546,122546,122523,122523,122477,122477,122456,122456,122437,122437,122375,122375,122347,122347,122298,122298,122281,122281,122253,122253,122208,122208,122173,122173,122110,122110,122067,122067,121988,121988,121967,121967,121925,121925,121881,121881,121847,121847,121825,121825,121809,121809,121696,121696,121628,121628,121611,121611,121510,121510,121492,121492,121452,121452,121426,121426,121398,121398,121330,121330,121303,121303,121262,121262,121209,121209,121144,121144,121095,121095,121072,121072,121049,121049,120949,120949,120835,120835,120809,120809,120770,120770,120734,120734,120683,120683,120651,120651,120615,120615,120574,120574,120556,120556,120506,120506,120378,120378,120237,120237,120191,120191,120171,120171,120152,120152,120124,120124,120033,120033,120004,120004,119986,119986,119924,119924,119908,119908,119869,119869,119852,119852,119782,119782,119757,119757,119714,119714,119690,119690,119629,119629,119592,119592,119560,119560,119544,119544,119510,119510,119435,119435,119402,119402,119382,119382,119359,119359,119317,119317,119283,119283,119266,119266,119235,119235,119211,119211,119184,119184,119147,119147,119115,119115,119099,119099,119061,119061,119014,119014,118993,118993,118953,118953,118927,118927,118891,118891,118834,118834,118785,118785,118718,118718,118662,118662,118642,118642,118621,118621,118579,118579,118520,118520,118495,118495,118471,118471,118453,118453,118363,118363,118313,118313,118285,118285,118246,118246,118196,118196,118147,118147,118117,118117,118091,118091,118072,118072,117972,117972,117955,117955,117909,117909,117834,117834,117808,117808,117790,117790,117754,117754,117656,117656,117623,117623,117607,117607,117555,117555,117507,117507,117484,117484,117450,117450,117432,117432,117414,117414,117395,117395,117301,117301,117252,117252,117227,117227,117176,117176,117126,117126,117105,117105,117085,117085,117060,117060,117040,117040,117016,117016,116959,116959,116886,116886,116807,116807,116753,116753,116726,116726,116678,116678,116546,116546,116515,116515,116479,116479,116461,116461,116417,116417,116336,116336,116268,116268,116237,116237,116215,116215,116153,116153,116119,116119,116067,116067,116050,116050,116030,116030,115970,115970,115925,115925,115864,115864,115832,115832,115780,115780,115757,115757,115732,115732,115691,115691,115533,115533,115454,115454,115435,115435,115413,115413,115394,115394,115350,115350,115332,115332,115301,115301,115277,115277,115248,115248,115222,115222,115198,115198,115180,115180,115146,115146,115087,115087,115023,115023,115005,115005,114985,114985,114961,114961,114937,114937,114880,114880,114863,114863,114822,114822,114803,114803,114774,114774,114718,114718,114683,114683,114648,114648,114623,114623,114551,114551,114357,114357,114339,114339,114303,114303,114278,114278,114231,114231,114207,114207,114156,114156,114123,114123,114087,114087,114065,114065,114039,114039,114005,114005,113952,113952,113931,113931,113897,113897,113880,113880,113844,113844,113816,113816,113788,113788,113765,113765,113734,113734,113700,113700,113651,113651,113607,113607,113569,113569,113493,113493,113465,113465,113414,113414,113345,113345,113324,113324,113241,113241,113225,113225,113125,113125,113108,113108,113092,113092,113070,113070,113053,113053,113025,113025,112985,112985,112938,112938,112921,112921,112877,112877,112852,112852,112806,112806,112774,112774,112746,112746,112719,112719,112701,112701,112683,112683,112642,112642,112565,112565,112510,112510,112475,112475,112402,112402,112384,112384,112358,112358,112305,112305,112288,112288,112254,112254,112224,112224,112180,112180,112124,112124,112107,112107,112089,112089,112000,112000,111984,111984,111926,111926,111898,111898,111842,111842,111819,111819,111782,111782,111764,111764,111724,111724,111707,111707,111613,111613,111563,111563,111544,111544,111424,111424,111356,111356,111339,111339,111322,111322,111284,111284,111260,111260,111218,111218,111158,111158,111127,111127,111105,111105,111043,111043,111017,111017,110996,110996,110943,110943,110879,110879,110854,110854,110783,110783,110749,110749,110683,110683,110648,110648,110621,110621,110494,110494,110475,110475,110447,110447,110328,110328,110299,110299,110275,110275,110235,110235,110200,110200,110104,110104,110082,110082,110025,110025,109995,109995,109972,109972,109954,109954,109908,109908,109880,109880,109861,109861,109811,109811,109741,109741,109702,109702,109664,109664,109639,109639,109622,109622,109579,109579,109498,109498,109423,109423,109407,109407,109378,109378,109344,109344,109304,109304,109238,109238,109222,109222,109194,109194,109175,109175,109124,109124,109090,109090,109045,109045,109028,109028,109000,109000,108975,108975,108954,108954,108931,108931,108882,108882,108790,108790,108753,108753,108717,108717,108701,108701,108596,108596,108555,108555,108425,108425,108396,108396,108364,108364,108345,108345,108324,108324,108282,108282,108252,108252,108189,108189,108172,108172,108153,108153,108073,108073,108053,108053,108037,108037,108017,108017,107999,107999,107965,107965,107943,107943,107927,107927,107899,107899,107858,107858,107825,107825,107734,107734,107715,107715,107678,107678,107615,107615,107584,107584,107530,107530,107497,107497,107471,107471,107423,107423,107401,107401,107380,107380,107348,107348,107295,107295,107233,107233,107161,107161,107140,107140,107115,107115,107086,107086,107058,107058,107037,107037,107016,107016,106999,106999,106934,106934,106760,106760,106697,106697,106674,106674,106614,106614,106589,106589,106494,106494,106454,106454,106409,106409,106340,106340,106267,106267,106225,106225,106124,106124,105997,105997,105955,105955,105925,105925,105859,105859,105816,105816,105785,105785,105740,105740,105686,105686,105655,105655,105616,105616,105593,105593,105566,105566,105538,105538,105438,105438,105408,105408,105382,105382,105300,105300,105258,105258,105174,105174,105147,105147,105075,105075,105030,105030,105009,105009,104972,104972,104934,104934,104896,104896,104877,104877,104859,104859,104830,104830,104806,104806,104699,104699,104667,104667,104587,104587,104560,104560,104514,104514,104487,104487,104431,104431,104408,104408,104377,104377,104351,104351,104319,104319,104256,104256,104197,104197,104146,104146,104130,104130,104097,104097,104035,104035,104010,104010,103991,103991,103971,103971,103948,103948,103900,103900,103868,103868,103829,103829,103794,103794,103764,103764,103735,103735,103718,103718,103700,103700,103681,103681,103628,103628,103606,103606,103575,103575,103553,103553,103521,103521,103489,103489,103467,103467,103433,103433,103409,103409,103376,103376,103299,103299,103257,103257,103218,103218,103193,103193,103166,103166,103143,103143,103114,103114,103090,103090,103034,103034,103012,103012,102953,102953,102925,102925,102863,102863,102834,102834,102812,102812,102794,102794,102767,102767,102718,102718,102636,102636,102619,102619,102573,102573,102547,102547,102529,102529,102511,102511,102475,102475,102448,102448,102424,102424,102388,102388,102323,102323,102169,102169,102148,102148,102131,102131,102099,102099,102082,102082,102065,102065,102048,102048,101931,101931,101885,101885,101852,101852,101826,101826,101800,101800,101693,101693,101667,101667,101648,101648,101593,101593,101556,101556,101523,101523,101501,101501,101474,101474,101438,101438,101413,101413,101374,101374,101337,101337,101271,101271,101244,101244,101209,101209,101189,101189,101164,101164,101132,101132,101030,101030,100982,100982,100955,100955,100928,100928,100893,100893,100864,100864,100839,100839,100815,100815,100778,100778,100726,100726,100686,100686,100644,100644,100606,100606,100562,100562,100450,100450,100426,100426,100302,100302,100255,100255,100236,100236,100158,100158,100122,100122,100050,100050,100016,100016,100000,100000,99963,99963,99923,99923,99876,99876,99843,99843,99781,99781,99761,99761,99745,99745,99726,99726,99702,99702,99660,99660,99630,99630,99587,99587,99521,99521,99457,99457,99438,99438,99397,99397,99293,99293,99271,99271,99198,99198,99180,99180,99052,99052,99036,99036,99010,99010,98976,98976,98844,98844,98793,98793,98769,98769,98716,98716,98655,98655,98615,98615,98565,98565,98545,98545,98511,98511,98490,98490,98446,98446,98402,98402,98378,98378,98356,98356,98325,98325,98285,98285,98255,98255,98229,98229,98185,98185,98147,98147,98045,98045,98000,98000,97941,97941,97828,97828,97805,97805,97776,97776,97744,97744,97696,97696,97679,97679,97617,97617,97598,97598,97551,97551,97514,97514,97489,97489,97472,97472,97456,97456,97381,97381,97354,97354,97301,97301,97272,97272,97210,97210,97183,97183,97149,97149,97126,97126,97103,97103,97077,97077,97059,97059,97013,97013,96955,96955,96917,96917,96849,96849,96744,96744,96725,96725,96688,96688,96621,96621,96569,96569,96506,96506,96478,96478,96451,96451,96401,96401,96336,96336,96315,96315,96285,96285,96252,96252,96218,96218,96160,96160,96110,96110,96042,96042,95989,95989,95973,95973,95942,95942,95849,95849,95829,95829,95750,95750,95728,95728,95707,95707,95689,95689,95576,95576,95478,95478,95432,95432,95404,95404,95382,95382,95363,95363,95294,95294,95278,95278,95240,95240,95196,95196,95170,95170,95106,95106,95078,95078,94987,94987,94944,94944,94894,94894,94862,94862,94843,94843,94824,94824,94808,94808,94761,94761,94730,94730,94706,94706,94678,94678,94618,94618,94599,94599,94573,94573,94519,94519,94485,94485,94450,94450,94362,94362,94325,94325,94306,94306,94288,94288,94267,94267,94241,94241,94218,94218,94138,94138,94090,94090,93954,93954,93936,93936,93898,93898,93861,93861,93831,93831,93717,93717,93694,93694,93664,93664,93625,93625,93580,93580,93518,93518,93460,93460,93444,93444,93424,93424,93396,93396,93380,93380,93361,93361,93234,93234,93191,93191,93148,93148,93048,93048,92992,92992,92971,92971,92954,92954,92866,92866,92838,92838,92776,92776,92739,92739,92712,92712,92695,92695,92674,92674,92635,92635,92613,92613,92588,92588,92529,92529,92502,92502,92421,92421,92386,92386,92323,92323,92296,92296,92276,92276,92220,92220,92182,92182,92096,92096,92080,92080,92030,92030,91985,91985,91954,91954,91934,91934,91917,91917,91879,91879,91839,91839,91799,91799,91783,91783,91766,91766,91743,91743,91713,91713,91672,91672,91636,91636,91620,91620,91578,91578,91560,91560,91521,91521,91499,91499,91478,91478,91442,91442,91424,91424,91397,91397,91366,91366,91348,91348,91330,91330,91309,91309,91279,91279,91222,91222,91136,91136,91052,91052,91025,91025,90999,90999,90957,90957,90933,90933,90910,90910,90862,90862,90831,90831,90814,90814,90774,90774,90731,90731,90663,90663,90610,90610,90557,90557,90521,90521,90502,90502,90412,90412,90376,90376,90333,90333,90311,90311,90246,90246,90218,90218,90180,90180,90142,90142,90109,90109,90089,90089,90061,90061,90036,90036,90004,90004,89980,89980,89913,89913,89851,89851,89832,89832,89792,89792,89762,89762,89725,89725,89703,89703,89685,89685,89654,89654,89626,89626,89543,89543,89515,89515,89464,89464,89429,89429,89348,89348,89331,89331,89295,89295,89243,89243,89162,89162,89133,89133,89105,89105,89079,89079,89059,89059,89036,89036,88995,88995,88929,88929,88909,88909,88893,88893,88856,88856,88810,88810,88785,88785,88714,88714,88688,88688,88661,88661,88642,88642,88610,88610,88584,88584,88549,88549,88514,88514,88476,88476,88460,88460,88425,88425,88403,88403,88387,88387,88360,88360,88320,88320,88299,88299,88280,88280,88235,88235,88184,88184,88149,88149,88129,88129,88087,88087,88035,88035,88014,88014,87969,87969,87929,87929,87910,87910,87893,87893,87877,87877,87792,87792,87752,87752,87730,87730,87713,87713,87681,87681,87635,87635,87618,87618,87594,87594,87561,87561,87545,87545,87519,87519,87480,87480,87452,87452,87431,87431,87415,87415,87341,87341,87307,87307,87288,87288,87265,87265,87218,87218,87202,87202,87140,87140,87109,87109,87056,87056,87023,87023,87004,87004,86983,86983,86938,86938,86922,86922,86903,86903,86884,86884,86831,86831,86806,86806,86750,86750,86696,86696,86665,86665,86581,86581,86564,86564,86529,86529,86472,86472,86439,86439,86415,86415,86381,86381,86308,86308,86289,86289,86222,86222,86200,86200,86147,86147,86077,86077,86022,86022,85980,85980,85952,85952,85882,85882,85832,85832,85778,85778,85745,85745,85715,85715,85694,85694,85675,85675,85655,85655,85629,85629,85555,85555,85512,85512,85494,85494,85461,85461,85440,85440,85418,85418,85384,85384,85275,85275,85258,85258,85219,85219,85203,85203,85156,85156,85121,85121,85102,85102,85057,85057,85018,85018,84927,84927,84875,84875,84840,84840,84743,84743,84701,84701,84678,84678,84656,84656,84590,84590,84568,84568,84515,84515,84465,84465,84384,84384,84322,84322,84288,84288,84192,84192,84148,84148,84129,84129,84099,84099,84079,84079,84050,84050,83984,83984,83928,83928,83910,83910,83856,83856,83815,83815,83795,83795,83775,83775,83758,83758,83679,83679,83659,83659,83620,83620,83590,83590,83545,83545,83512,83512,83479,83479,83448,83448,83360,83360,83333,83333,83295,83295,83268,83268,83153,83153,83120,83120,83081,83081,83063,83063,83029,83029,83008,83008,82975,82975,82959,82959,82922,82922,82891,82891,82869,82869,82799,82799,82771,82771,82742,82742,82679,82679,82615,82615,82594,82594,82576,82576,82528,82528,82486,82486,82455,82455,82435,82435,82413,82413,82385,82385,82326,82326,82287,82287,82261,82261,82238,82238,82164,82164,82148,82148,82124,82124,82084,82084,82068,82068,82039,82039,82015,82015,81978,81978,81932,81932,81907,81907,81848,81848,81830,81830,81788,81788,81752,81752,81736,81736,81714,81714,81695,81695,81673,81673,81616,81616,81600,81600,81547,81547,81501,81501,81478,81478,81457,81457,81404,81404,81381,81381,81343,81343,81321,81321,81291,81291,81266,81266,81246,81246,81228,81228,81208,81208,81139,81139,81121,81121,81100,81100,81073,81073,81043,81043,81002,81002,80985,80985,80952,80952,80917,80917,80881,80881,80853,80853,80784,80784,80768,80768,80736,80736,80709,80709,80685,80685,80667,80667,80572,80572,80545,80545,80471,80471,80454,80454,80403,80403,80385,80385,80338,80338,80282,80282,80228,80228,80198,80198,80173,80173,80139,80139,80106,80106,80029,80029,79977,79977,79950,79950,79805,79805,79674,79674,79657,79657,79570,79570,79538,79538,79509,79509,79474,79474,79438,79438,79401,79401,79381,79381,79356,79356,79321,79321,79269,79269,79253,79253,79193,79193,79176,79176,79154,79154,79133,79133,79110,79110,79053,79053,79027,79027,78912,78912,78887,78887,78847,78847,78825,78825,78804,78804,78782,78782,78744,78744,78703,78703,78659,78659,78556,78556,78531,78531,78505,78505,78486,78486,78470,78470,78431,78431,78329,78329,78307,78307,78284,78284,78253,78253,78230,78230,78214,78214,78195,78195,78096,78096,78044,78044,78012,78012,77953,77953,77920,77920,77898,77898,77878,77878,77845,77845,77735,77735,77710,77710,77691,77691,77636,77636,77578,77578,77531,77531,77490,77490,77460,77460,77384,77384,77338,77338,77285,77285,77267,77267,77239,77239,77172,77172,77127,77127,77107,77107,77083,77083,77065,77065,77038,77038,77013,77013,76950,76950,76891,76891,76867,76867,76824,76824,76776,76776,76712,76712,76648,76648,76607,76607,76583,76583,76548,76548,76531,76531,76505,76505,76437,76437,76398,76398,76379,76379,76347,76347,76310,76310,76282,76282,76242,76242,76193,76193,76165,76165,76138,76138,76121,76121,76073,76073,76047,76047,76004,76004,75981,75981,75935,75935,75915,75915,75895,75895,75867,75867,75846,75846,75830,75830,75814,75814,75782,75782,75761,75761,75701,75701,75637,75637,75619,75619,75552,75552,75514,75514,75493,75493,75475,75475,75449,75449,75390,75390,75374,75374,75294,75294,75262,75262,75191,75191,75150,75150,75113,75113,75079,75079,75035,75035,74994,74994,74975,74975,74940,74940,74908,74908,74853,74853,74816,74816,74793,74793,74769,74769,74668,74668,74634,74634,74591,74591,74521,74521,74491,74491,74469,74469,74449,74449,74412,74412,74388,74388,74361,74361,74338,74338,74289,74289,74264,74264,74184,74184,74144,74144,74121,74121,74097,74097,74079,74079,74039,74039,74021,74021,73997,73997,73977,73977,73916,73916,73889,73889,73831,73831,73802,73802,73760,73760,73676,73676,73658,73658,73622,73622,73606,73606,73582,73582,73514,73514,73489,73489,73466,73466,73419,73419,73403,73403,73381,73381,73271,73271,73161,73161,73091,73091,73049,73049,73011,73011,72973,72973,72845,72845,72807,72807,72778,72778,72681,72681,72649,72649,72631,72631,72609,72609,72580,72580,72547,72547,72522,72522,72483,72483,72457,72457,72418,72418,72386,72386,72322,72322,72281,72281,72251,72251,72223,72223,72177,72177,72135,72135,72101,72101,72076,72076,72037,72037,72017,72017,71970,71970,71926,71926,71910,71910,71877,71877,71843,71843,71811,71811,71782,71782,71709,71709,71662,71662,71630,71630,71604,71604,71559,71559,71542,71542,71496,71496,71471,71471,71453,71453,71415,71415,71370,71370,71354,71354,71299,71299,71283,71283,71245,71245,71226,71226,71197,71197,71178,71178,71128,71128,71112,71112,71092,71092,71076,71076,71059,71059,71042,71042,71025,71025,70963,70963,70938,70938,70881,70881,70841,70841,70804,70804,70743,70743,70674,70674,70649,70649,70572,70572,70548,70548,70523,70523,70507,70507,70482,70482,70444,70444,70414,70414,70365,70365,70347,70347,70294,70294,70272,70272,70213,70213,70181,70181,70129,70129,70073,70073,70051,70051,70034,70034,69993,69993,69943,69943,69913,69913,69897,69897,69877,69877,69849,69849,69828,69828,69781,69781,69763,69763,69746,69746,69695,69695,69653,69653,69612,69612,69583,69583,69563,69563,69542,69542,69526,69526,69503,69503,69457,69457,69427,69427,69398,69398,69366,69366,69350,69350,69334,69334,69306,69306,69289,69289,69271,69271,69197,69197,69151,69151,69121,69121,69098,69098,69074,69074,69032,69032,68971,68971,68952,68952,68923,68923,68858,68858,68827,68827,68801,68801,68778,68778,68745,68745,68729,68729,68690,68690,68656,68656,68602,68602,68579,68579,68562,68562,68514,68514,68497,68497,68465,68465,68449,68449,68427,68427,68375,68375,68335,68335,68316,68316,68287,68287,68244,68244,68205,68205,68188,68188,68155,68155,68105,68105,68085,68085,68026,68026,68001,68001,67934,67934,67890,67890,67867,67867,67820,67820,67773,67773,67711,67711,67648,67648,67632,67632,67605,67605,67587,67587,67558,67558,67518,67518,67481,67481,67459,67459,67412,67412,67381,67381,67349,67349,67332,67332,67315,67315,67295,67295,67275,67275,67238,67238,67180,67180,67149,67149,67123,67123,67102,67102,67075,67075,67057,67057,67039,67039,67022,67022,67005,67005,66988,66988,66948,66948,66903,66903,66879,66879,66829,66829,66744,66744,66706,66706,66688,66688,66660,66660,66605,66605,66585,66585,66550,66550,66524,66524,66488,66488,66459,66459,66440,66440,66413,66413,66394,66394,66377,66377,66350,66350,66331,66331,66313,66313,66281,66281,66252,66252,66211,66211,66156,66156,66129,66129,66107,66107,66055,66055,66023,66023,66005,66005,65956,65956,65928,65928,65871,65871,65855,65855,65823,65823,65807,65807,65776,65776,65744,65744,65724,65724,65706,65706,65685,65685,65664,65664,65646,65646,65622,65622,65603,65603,65586,65586,65569,65569,65542,65542,65513,65513,65483,65483,65445,65445,65427,65427,65396,65396,65369,65369,65352,65352,65319,65319,65295,65295,65241,65241,65223,65223,65204,65204,65142,65142,65107,65107,65082,65082,65065,65065,65046,65046,65027,65027,64975,64975,64937,64937,64909,64909,64880,64880,64863,64863,64832,64832,64814,64814,64779,64779,64729,64729,64669,64669,64629,64629,64613,64613,64597,64597,64561,64561,64526,64526,64485,64485,64466,64466,64442,64442,64422,64422,64396,64396,64363,64363,64327,64327,64305,64305,64277,64277,64259,64259,64242,64242,64224,64224,64185,64185,64156,64156,64120,64120,64029,64029,63991,63991,63972,63972,63922,63922,63906,63906,63859,63859,63832,63832,63814,63814,63784,63784,63756,63756,63740,63740,63703,63703,63652,63652,63611,63611,63583,63583,63560,63560,63523,63523,63505,63505,63485,63485,63444,63444,63375,63375,63325,63325,63283,63283,63266,63266,63231,63231,63211,63211,63186,63186,63161,63161,63140,63140,63114,63114,63060,63060,63037,63037,62998,62998,62941,62941,62925,62925,62898,62898,62875,62875,62772,62772,62755,62755,62717,62717,62677,62677,62643,62643,62612,62612,62581,62581,62557,62557,62532,62532,62513,62513,62495,62495,62472,62472,62424,62424,62352,62352,62311,62311,62256,62256,62217,62217,62179,62179,62146,62146,62119,62119,62076,62076,62027,62027,62010,62010,61956,61956,61922,61922,61905,61905,61887,61887,61834,61834,61798,61798,61762,61762,61734,61734,61713,61713,61688,61688,61620,61620,61600,61600,61559,61559,61543,61543,61512,61512,61489,61489,61395,61395,61317,61317,61289,61289,61273,61273,61249,61249,61217,61217,61180,61180,61151,61151,61134,61134,61058,61058,60987,60987,60968,60968,60941,60941,60922,60922,60890,60890,60847,60847,60819,60819,60802,60802,60780,60780,60754,60754,60730,60730,60707,60707,60670,60670,60651,60651,60611,60611,60584,60584,60562,60562,60545,60545,60526,60526,60508,60508,60472,60472,60416,60416,60378,60378,60342,60342,60307,60307,60278,60278,60239,60239,60222,60222,60175,60175,60154,60154,60136,60136,60095,60095,60074,60074,60042,60042,60004,60004,59987,59987,59968,59968,59948,59948,59932,59932,59878,59878,59855,59855,59827,59827,59805,59805,59772,59772,59749,59749,59732,59732,59715,59715,59695,59695,59669,59669,59652,59652,59630,59630,59612,59612,59588,59588,59571,59571,59500,59500,59471,59471,59410,59410,59390,59390,59345,59345,59302,59302,59282,59282,59244,59244,59180,59180,59155,59155,59113,59113,59072,59072,59037,59037,58990,58990,58966,58966,58944,58944,58908,58908,58886,58886,58868,58868,58845,58845,58824,58824,58801,58801,58761,58761,58742,58742,58713,58713,58675,58675,58640,58640,58613,58613,58577,58577,58553,58553,58534,58534,58499,58499,58443,58443,58426,58426,58394,58394,58378,58378,58329,58329,58306,58306,58289,58289,58229,58229,58213,58213,58193,58193,58154,58154,58132,58132,58107,58107,58089,58089,58070,58070,58052,58052,58019,58019,58003,58003,57969,57969,57949,57949,57908,57908,57873,57873,57835,57835,57814,57814,57784,57784,57758,57758,57721,57721,57699,57699,57664,57664,57591,57591,57558,57558,57541,57541,57517,57517,57496,57496,57479,57479,57457,57457,57400,57400,57370,57370,57331,57331,57283,57283,57260,57260,57235,57235,57195,57195,57170,57170,57125,57125,57060,57060,57041,57041,57006,57006,56964,56964,56939,56939,56914,56914,56891,56891,56862,56862,56841,56841,56817,56817,56781,56781,56763,56763,56734,56734,56703,56703,56685,56685,56666,56666,56650,56650,56618,56618,56592,56592,56558,56558,56525,56525,56472,56472,56437,56437,56397,56397,56375,56375,56356,56356,56324,56324,56304,56304,56283,56283,56260,56260,56235,56235,56185,56185,56161,56161,56139,56139,56096,56096,56079,56079,56006,56006,55989,55989,55924,55924,55891,55891,55860,55860,55837,55837,55810,55810,55785,55785,55726,55726,55708,55708,55680,55680,55651,55651,55632,55632,55596,55596,55576,55576,55549,55549,55518,55518,55487,55487,55458,55458,55410,55410,55350,55350,55333,55333,55284,55284,55258,55258,55233,55233,55184,55184,55155,55155,55137,55137,55085,55085,55067,55067,55050,55050,55022,55022,55004,55004,54979,54979,54953,54953,54925,54925,54886,54886,54868,54868,54830,54830,54810,54810,54783,54783,54766,54766,54741,54741,54700,54700,54684,54684,54657,54657,54638,54638,54612,54612,54585,54585,54538,54538,54519,54519,54503,54503,54478,54478,54415,54415,54389,54389,54368,54368,54305,54305,54263,54263,54247,54247,54218,54218,54191,54191,54169,54169,54147,54147,54130,54130,54108,54108,54071,54071,54048,54048,54031,54031,54006,54006,53990,53990,53963,53963,53928,53928,53908,53908,53859,53859,53837,53837,53749,53749,53716,53716,53679,53679,53655,53655,53630,53630,53611,53611,53560,53560,53524,53524,53502,53502,53440,53440,53369,53369,53349,53349,53329,53329,53296,53296,53246,53246,53214,53214,53197,53197,53149,53149,53129,53129,53101,53101,53034,53034,53003,53003,52974,52974,52926,52926,52880,52880,52862,52862,52833,52833,52794,52794,52764,52764,52745,52745,52710,52710,52692,52692,52673,52673,52641,52641,52618,52618,52580,52580,52549,52549,52519,52519,52482,52482,52453,52453,52432,52432,52389,52389,52353,52353,52322,52322,52295,52295,52276,52276,52245,52245,52228,52228,52212,52212,52185,52185,52148,52148,52119,52119,52087,52087,52059,52059,52007,52007,51983,51983,51957,51957,51940,51940,51915,51915,51892,51892,51868,51868,51839,51839,51821,51821,51804,51804,51788,51788,51761,51761,51744,51744,51726,51726,51701,51701,51656,51656,51611,51611,51587,51587,51560,51560,51525,51525,51509,51509,51469,51469,51441,51441,51422,51422,51398,51398,51378,51378,51350,51350,51304,51304,51280,51280,51243,51243,51160,51160,51135,51135,51119,51119,51069,51069,51027,51027,51003,51003,50985,50985,50950,50950,50907,50907,50870,50870,50816,50816,50790,50790,50750,50750,50720,50720,50682,50682,50632,50632,50608,50608,50587,50587,50559,50559,50542,50542,50523,50523,50501,50501,50453,50453,50418,50418,50397,50397,50346,50346,50274,50274,50254,50254,50229,50229,50207,50207,50167,50167,50141,50141,50115,50115,50075,50075,50049,50049,50027,50027,50008,50008,49988,49988,49972,49972,49945,49945,49914,49914,49898,49898,49858,49858,49829,49829,49790,49790,49754,49754,49726,49726,49687,49687,49655,49655,49636,49636,49608,49608,49590,49590,49554,49554,49504,49504,49471,49471,49451,49451,49427,49427,49411,49411,49389,49389,49371,49371,49348,49348,49324,49324,49305,49305,49288,49288,49257,49257,49235,49235,49219,49219,49199,49199,49122,49122,49083,49083,49060,49060,49044,49044,49020,49020,48988,48988,48970,48970,48924,48924,48889,48889,48865,48865,48846,48846,48823,48823,48799,48799,48769,48769,48749,48749,48727,48727,48709,48709,48693,48693,48675,48675,48658,48658,48626,48626,48570,48570,48551,48551,48511,48511,48474,48474,48425,48425,48396,48396,48376,48376,48321,48321,48298,48298,48265,48265,48243,48243,48221,48221,48204,48204,48162,48162,48145,48145,48112,48112,48083,48083,48063,48063,48044,48044,48028,48028,47994,47994,47972,47972,47945,47945,47909,47909,47890,47890,47870,47870,47844,47844,47811,47811,47765,47765,47748,47748,47732,47732,47712,47712,47691,47691,47666,47666,47646,47646,47624,47624,47557,47557,47530,47530,47500,47500,47467,47467,47423,47423,47403,47403,47386,47386,47360,47360,47326,47326,47302,47302,47279,47279,47243,47243,47225,47225,47165,47165,47137,47137,47119,47119,47087,47087,47068,47068,46991,46991,46962,46962,46933,46933,46886,46886,46856,46856,46831,46831,46815,46815,46790,46790,46774,46774,46753,46753,46734,46734,46717,46717,46692,46692,46666,46666,46632,46632,46615,46615,46571,46571,46542,46542,46497,46497,46477,46477,46461,46461,46430,46430,46401,46401,46372,46372,46345,46345,46303,46303,46261,46261,46232,46232,46211,46211,46139,46139,46117,46117,46094,46094,46077,46077,46061,46061,46035,46035,46019,46019,46002,46002,45985,45985,45965,45965,45938,45938,45911,45911,45873,45873,45835,45835,45814,45814,45779,45779,45749,45749,45730,45730,45703,45703,45682,45682,45653,45653,45628,45628,45606,45606,45590,45590,45565,45565,45547,45547,45526,45526,45507,45507,45488,45488,45467,45467,45425,45425,45368,45368,45338,45338,45308,45308,45275,45275,45192,45192,45168,45168,45150,45150,45127,45127,45111,45111,45076,45076,45051,45051,45031,45031,45007,45007,44984,44984,44959,44959,44917,44917,44892,44892,44825,44825,44809,44809,44760,44760,44736,44736,44716,44716,44699,44699,44681,44681,44658,44658,44642,44642,44621,44621,44589,44589,44552,44552,44519,44519,44459,44459,44433,44433,44400,44400,44382,44382,44360,44360,44343,44343,44327,44327,44311,44311,44292,44292,44258,44258,44232,44232,44216,44216,44192,44192,44162,44162,44141,44141,44117,44117,44089,44089,44032,44032,44012,44012,43975,43975,43952,43952,43936,43936,43892,43892,43874,43874,43848,43848,43827,43827,43777,43777,43750,43750,43713,43713,43691,43691,43665,43665,43645,43645,43610,43610,43582,43582,43555,43555,43532,43532,43516,43516,43499,43499,43472,43472,43450,43450,43434,43434,43412,43412,43391,43391,43370,43370,43335,43335,43289,43289,43271,43271,43255,43255,43230,43230,43205,43205,43180,43180,43155,43155,43136,43136,43115,43115,43078,43078,43040,43040,42989,42989,42965,42965,42944,42944,42928,42928,42906,42906,42880,42880,42861,42861,42843,42843,42827,42827,42803,42803,42773,42773,42749,42749,42733,42733,42710,42710,42691,42691,42669,42669,42619,42619,42579,42579,42560,42560,42534,42534,42513,42513,42495,42495,42465,42465,42440,42440,42412,42412,42378,42378,42361,42361,42342,42342,42323,42323,42303,42303,42286,42286,42270,42270,42247,42247,42229,42229,42183,42183,42163,42163,42131,42131,42109,42109,42087,42087,42066,42066,42048,42048,42031,42031,42014,42014,41997,41997,41977,41977,41961,41961,41945,41945,41923,41923,41891,41891,41870,41870,41854,41854,41834,41834,41815,41815,41797,41797,41781,41781,41703,41703,41683,41683,41651,41651,41630,41630,41611,41611,41592,41592,41540,41540,41498,41498,41465,41465,41445,41445,41408,41408,41389,41389,41364,41364,41342,41342,41289,41289,41265,41265,41248,41248,41227,41227,41195,41195,41154,41154,41138,41138,41113,41113,41086,41086,41068,41068,41011,41011,40973,40973,40938,40938,40910,40910,40886,40886,40863,40863,40804,40804,40788,40788,40767,40767,40734,40734,40718,40718,40688,40688,40666,40666,40628,40628,40595,40595,40554,40554,40535,40535,40519,40519,40498,40498,40478,40478,40455,40455,40434,40434,40405,40405,40379,40379,40348,40348,40308,40308,40291,40291,40272,40272,40250,40250,40229,40229,40212,40212,40175,40175,40157,40157,40135,40135,40115,40115,40085,40085,40065,40065,40039,40039,40021,40021,40005,40005,39987,39987,39962,39962,39945,39945,39922,39922,39880,39880,39862,39862,39835,39835,39819,39819,39798,39798,39773,39773,39682,39682,39658,39658,39639,39639,39618,39618,39599,39599,39573,39573,39557,39557,39533,39533,39503,39503,39482,39482,39461,39461,39444,39444,39422,39422,39390,39390,39347,39347,39331,39331,39303,39303,39250,39250,39223,39223,39191,39191,39171,39171,39151,39151,39097,39097,39078,39078,39055,39055,39027,39027,38990,38990,38954,38954,38931,38931,38911,38911,38873,38873,38845,38845,38829,38829,38781,38781,38755,38755,38731,38731,38709,38709,38686,38686,38665,38665,38621,38621,38605,38605,38544,38544,38517,38517,38484,38484,38453,38453,38436,38436,38420,38420,38400,38400,38383,38383,38361,38361,38344,38344,38326,38326,38309,38309,38276,38276,38253,38253,38236,38236,38218,38218,38192,38192,38171,38171,38152,38152,38134,38134,38116,38116,38098,38098,38065,38065,38041,38041,38024,38024,38007,38007,37985,37985,37958,37958,37936,37936,37909,37909,37886,37886,37867,37867,37851,37851,37830,37830,37808,37808,37788,37788,37763,37763,37741,37741,37724,37724,37704,37704,37671,37671,37648,37648,37625,37625,37608,37608,37569,37569,37543,37543,37516,37516,37494,37494,37467,37467,37445,37445,37426,37426,37388,37388,37370,37370,37336,37336,37320,37320,37302,37302,37285,37285,37255,37255,37221,37221,37182,37182,37147,37147,37127,37127,37107,37107,37077,37077,37057,37057,37039,37039,37013,37013,36993,36993,36962,36962,36925,36925,36907,36907,36889,36889,36870,36870,36854,36854,36829,36829,36801,36801,36780,36780,36737,36737,36719,36719,36703,36703,36680,36680,36659,36659,36621,36621,36601,36601,36541,36541,36513,36513,36490,36490,36471,36471,36453,36453,36429,36429,36409,36409,36383,36383,36367,36367,36346,36346,36323,36323,36305,36305,36286,36286,36268,36268,36251,36251,36222,36222,36194,36194,36158,36158,36136,36136,36116,36116,36094,36094,36063,36063,36045,36045,36015,36015,35970,35970,35954,35954,35928,35928,35907,35907,35891,35891,35870,35870,35853,35853,35831,35831,35810,35810,35794,35794,35776,35776,35723,35723,35704,35704,35685,35685,35667,35667,35637,35637,35621,35621,35605,35605,35576,35576,35546,35546,35525,35525,35500,35500,35484,35484,35419,35419,35400,35400,35384,35384,35342,35342,35292,35292,35274,35274,35254,35254,35231,35231,35198,35198,35182,35182,35166,35166,35148,35148,35131,35131,35115,35115,35090,35090,35041,35041,35017,35017,34994,34994,34968,34968,34950,34950,34915,34915,34895,34895,34877,34877,34856,34856,34837,34837,34817,34817,34793,34793,34776,34776,34758,34758,34742,34742,34725,34725,34706,34706,34690,34690,34671,34671,34654,34654,34637,34637,34615,34615,34586,34586,34566,34566,34548,34548,34518,34518,34474,34474,34411,34411,34394,34394,34376,34376,34349,34349,34322,34322,34303,34303,34285,34285,34269,34269,34250,34250,34231,34231,34205,34205,34189,34189,34167,34167,34129,34129,34105,34105,34085,34085,34064,34064,34048,34048,34031,34031,34000,34000,33980,33980,33961,33961,33942,33942,33896,33896,33878,33878,33833,33833,33800,33800,33775,33775,33758,33758,33716,33716,33699,33699,33681,33681,33642,33642,33620,33620,33597,33597,33557,33557,33500,33500,33482,33482,33422,33422,33405,33405,33388,33388,33371,33371,33355,33355,33338,33338,33316,33316,33300,33300,33281,33281,33258,33258,33242,33242,33216,33216,33197,33197,33179,33179,33156,33156,33137,33137,33094,33094,33078,33078,33062,33062,33028,33028,33011,33011,32995,32995,32972,32972,32948,32948,32913,32913,32863,32863,32830,32830,32811,32811,32767,32767,32751,32751,32715,32715,32699,32699,32671,32671,32654,32654,32630,32630,32607,32607,32590,32590,32549,32549,32517,32517,32500,32500,32484,32484,32464,32464,32441,32441,32425,32425,32402,32402,32366,32366,32350,32350,32313,32313,32295,32295,32278,32278,32247,32247,32231,32231,32200,32200,32175,32175,32158,32158,32142,32142,32125,32125,32105,32105,32089,32089,32066,32066,32047,32047,32029,32029,31999,31999,31980,31980,31957,31957,31939,31939,31923,31923,31905,31905,31885,31885,31863,31863,31842,31842,31820,31820,31788,31788,31755,31755,31728,31728,31712,31712,31692,31692,31662,31662,31645,31645,31623,31623,31607,31607,31591,31591,31571,31571,31551,31551,31520,31520,31503,31503,31483,31483,31460,31460,31444,31444,31425,31425,31407,31407,31389,31389,31370,31370,31344,31344,31323,31323,31300,31300,31284,31284,31260,31260,31244,31244,31216,31216,31177,31177,31159,31159,31140,31140,31118,31118,31094,31094,31070,31070,31050,31050,31034,31034,31015,31015,30982,30982,30965,30965,30938,30938,30905,30905,30884,30884,30864,30864,30831,30831,30806,30806,30787,30787,30769,30769,30730,30730,30707,30707,30689,30689,30672,30672,30649,30649,30632,30632,30616,30616,30600,30600,30584,30584,30568,30568,30546,30546,30522,30522,30502,30502,30483,30483,30460,30460,30439,30439,30422,30422,30391,30391,30368,30368,30348,30348,30311,30311,30292,30292,30263,30263,30239,30239,30211,30211,30184,30184,30159,30159,30118,30118,30097,30097,30059,30059,30039,30039,30023,30023,29977,29977,29952,29952,29933,29933,29916,29916,29898,29898,29872,29872,29848,29848,29827,29827,29802,29802,29779,29779,29757,29757,29730,29730,29705,29705,29688,29688,29664,29664,29635,29635,29618,29618,29601,29601,29580,29580,29564,29564,29547,29547,29522,29522,29475,29475,29449,29449,29432,29432,29411,29411,29394,29394,29370,29370,29341,29341,29317,29317,29298,29298,29282,29282,29266,29266,29250,29250,29226,29226,29205,29205,29188,29188,29168,29168,29151,29151,29107,29107,29086,29086,29069,29069,29048,29048,29027,29027,28983,28983,28964,28964,28948,28948,28931,28931,28914,28914,28897,28897,28869,28869,28847,28847,28829,28829,28806,28806,28784,28784,28765,28765,28743,28743,28723,28723,28686,28686,28666,28666,28640,28640,28621,28621,28605,28605,28589,28589,28572,28572,28549,28549,28531,28531,28510,28510,28492,28492,28471,28471,28451,28451,28431,28431,28415,28415,28399,28399,28371,28371,28355,28355,28339,28339,28322,28322,28303,28303,28286,28286,28266,28266,28249,28249,28226,28226,28201,28201,28181,28181,28165,28165,28146,28146,28130,28130,28113,28113,28096,28096,28080,28080,28060,28060,28044,28044,28027,28027,28004,28004,27987,27987,27971,27971,27951,27951,27933,27933,27914,27914,27894,27894,27877,27877,27851,27851,27828,27828,27812,27812,27795,27795,27779,27779,27759,27759,27739,27739,27717,27717,27701,27701,27684,27684,27662,27662,27634,27634,27616,27616,27592,27592,27570,27570,27550,27550,27525,27525,27505,27505,27483,27483,27460,27460,27441,27441,27424,27424,27403,27403,27387,27387,27371,27371,27355,27355,27338,27338,27322,27322,27305,27305,27284,27284,27267,27267,27251,27251,27229,27229,27206,27206,27189,27189,27172,27172,27156,27156,27140,27140,27118,27118,27096,27096,27077,27077,27061,27061,27045,27045,27028,27028,27011,27011,26993,26993,26964,26964,26947,26947,26931,26931,26915,26915,26891,26891,26875,26875,26844,26844,26825,26825,26807,26807,26790,26790,26761,26761,26744,26744,26717,26717,26698,26698,26679,26679,26663,26663,26646,26646,26629,26629,26610,26610,26594,26594,26576,26576,26541,26541,26518,26518,26499,26499,26480,26480,26460,26460,26444,26444,26427,26427,26410,26410,26391,26391,26375,26375,26354,26354,26334,26334,26313,26313,26291,26291,26267,26267,26247,26247,26228,26228,26201,26201,26183,26183,26150,26150,26129,26129,26113,26113,26088,26088,26064,26064,26040,26040,26022,26022,26004,26004,25980,25980,25964,25964,25946,25946,25930,25930,25911,25911,25893,25893,25861,25861,25840,25840,25821,25821,25805,25805,25786,25786,25770,25770,25753,25753,25727,25727,25711,25711,25695,25695,25674,25674,25658,25658,25641,25641,25621,25621,25600,25600,25584,25584,25565,25565,25547,25547,25523,25523,25507,25507,25491,25491,25474,25474,25456,25456,25438,25438,25421,25421,25402,25402,25386,25386,25370,25370,25349,25349,25333,25333,25316,25316,25296,25296,25274,25274,25258,25258,25240,25240,25224,25224,25200,25200,25181,25181,25162,25162,25140,25140,25108,25108,25090,25090,25073,25073,25056,25056,25031,25031,25014,25014,24996,24996,24980,24980,24961,24961,24940,24940,24921,24921,24896,24896,24878,24878,24861,24861,24845,24845,24828,24828,24811,24811,24794,24794,24778,24778,24755,24755,24739,24739,24716,24716,24696,24696,24669,24669,24651,24651,24634,24634,24617,24617,24595,24595,24571,24571,24555,24555,24539,24539,24513,24513,24492,24492,24475,24475,24455,24455,24437,24437,24419,24419,24403,24403,24385,24385,24366,24366,24347,24347,24317,24317,24301,24301,24284,24284,24262,24262,24246,24246,24227,24227,24204,24204,24188,24188,24172,24172,24150,24150,24129,24129,24108,24108,24091,24091,24074,24074,24058,24058,24041,24041,24025,24025,24005,24005,23989,23989,23962,23962,23945,23945,23922,23922,23906,23906,23889,23889,23872,23872,23851,23851,23823,23823,23798,23798,23777,23777,23761,23761,23739,23739,23716,23716,23697,23697,23671,23671,23651,23651,23634,23634,23616,23616,23598,23598,23581,23581,23557,23557,23535,23535,23516,23516,23496,23496,23478,23478,23458,23458,23433,23433,23417,23417,23400,23400,23384,23384,23368,23368,23350,23350,23334,23334,23315,23315,23297,23297,23281,23281,23261,23261,23244,23244,23215,23215,23198,23198,23179,23179,23163,23163,23142,23142,23124,23124,23108,23108,23090,23090,23066,23066,23047,23047,23028,23028,23012,23012,22995,22995,22978,22978,22955,22955,22938,22938,22910,22910,22892,22892,22872,22872,22852,22852,22831,22831,22809,22809,22775,22775,22756,22756,22740,22740,22719,22719,22702,22702,22685,22685,22669,22669,22652,22652,22636,22636,22620,22620,22600,22600,22582,22582,22564,22564,22543,22543,22526,22526,22510,22510,22494,22494,22477,22477,22461,22461,22433,22433,22416,22416,22400,22400,22383,22383,22365,22365,22348,22348,22330,22330,22313,22313,22292,22292,22269,22269,22253,22253,22236,22236,22219,22219,22201,22201,22184,22184,22164,22164,22147,22147,22130,22130,22112,22112,22096,22096,22079,22079,22063,22063,22047,22047,22030,22030,22014,22014,21998,21998,21982,21982,21964,21964,21947,21947,21928,21928,21912,21912,21890,21890,21873,21873,21855,21855,21839,21839,21819,21819,21802,21802,21780,21780,21756,21756,21740,21740,21721,21721,21704,21704,21685,21685,21658,21658,21633,21633,21615,21615,21598,21598,21582,21582,21564,21564,21547,21547,21530,21530,21514,21514,21496,21496,21478,21478,21453,21453,21436,21436,21420,21420,21400,21400,21382,21382,21364,21364,21340,21340,21323,21323,21305,21305,21281,21281,21265,21265,21249,21249,21232,21232,21215,21215,21199,21199,21180,21180,21164,21164,21140,21140,21120,21120,21104,21104,21082,21082,21065,21065,21046,21046,21009,21009,20993,20993,20975,20975,20959,20959,20942,20942,20926,20926,20908,20908,20891,20891,20875,20875,20859,20859,20843,20843,20826,20826,20810,20810,20793,20793,20776,20776,20754,20754,20734,20734,20718,20718,20702,20702,20684,20684,20664,20664,20648,20648,20632,20632,20614,20614,20596,20596,20580,20580,20561,20561,20545,20545,20528,20528,20506,20506,20485,20485,20465,20465,20445,20445,20428,20428,20402,20402,20385,20385,20368,20368,20348,20348,20331,20331,20312,20312,20294,20294,20273,20273,20257,20257,20240,20240,20221,20221,20203,20203,20187,20187,20171,20171,20151,20151,20134,20134,20116,20116,20099,20099,20080,20080,20064,20064,20045,20045,20029,20029,20011,20011,19993,19993,19976,19976,19959,19959,19943,19943,19923,19923,19906,19906,19890,19890,19874,19874,19854,19854,19833,19833,19817,19817,19794,19794,19777,19777,19761,19761,19745,19745,19723,19723,19705,19705,19689,19689,19666,19666,19649,19649,19632,19632,19612,19612,19596,19596,19580,19580,19562,19562,19541,19541,19525,19525,19504,19504,19486,19486,19461,19461,19443,19443,19426,19426,19410,19410,19394,19394,19375,19375,19358,19358,19334,19334,19317,19317,19300,19300,19278,19278,19262,19262,19245,19245,19228,19228,19212,19212,19191,19191,19175,19175,19155,19155,19136,19136,19119,19119,19103,19103,19082,19082,19064,19064,19046,19046,19027,19027,19011,19011,18995,18995,18978,18978,18962,18962,18946,18946,18924,18924,18902,18902,18885,18885,18866,18866,18850,18850,18834,18834,18818,18818,18802,18802,18783,18783,18766,18766,18750,18750,18734,18734,18716,18716,18700,18700,18682,18682,18666,18666,18649,18649,18632,18632,18608,18608,18589,18589,18573,18573,18557,18557,18538,18538,18522,18522,18504,18504,18487,18487,18471,18471,18454,18454,18436,18436,18415,18415,18398,18398,18380,18380,18364,18364,18348,18348,18332,18332,18314,18314,18297,18297,18281,18281,18264,18264,18248,18248,18231,18231,18214,18214,18198,18198,18182,18182,18166,18166,18150,18150,18134,18134,18112,18112,18095,18095,18077,18077,18061,18061,18043,18043,18026,18026,18004,18004,17985,17985,17968,17968,17951,17951,17934,17934,17918,17918,17900,17900,17884,17884,17857,17857,17840,17840,17824,17824,17806,17806,17789,17789,17772,17772,17756,17756,17739,17739,17723,17723,17705,17705,17689,17689,17673,17673,17657,17657,17640,17640,17624,17624,17605,17605,17588,17588,17568,17568,17547,17547,17531,17531,17515,17515,17499,17499,17483,17483,17467,17467,17450,17450,17434,17434,17413,17413,17397,17397,17381,17381,17365,17365,17349,17349,17332,17332,17315,17315,17298,17298,17280,17280,17261,17261,17243,17243,17227,17227,17207,17207,17190,17190,17169,17169,17153,17153,17136,17136,17119,17119,17098,17098,17082,17082,17064,17064,17042,17042,17026,17026,16947,16947,16927,16927,16829,16829,16094,16094,14701,14701,12916,12916,11586,11586,10948,10948,8779,8779,8350,8350,7853,7853,7581,7581,7463,7463,7150,7150,6888,6888,6566,6566,5885,5885,5461,5461,4958,4958,4146,4146,0.0],[4577433,4577433,4577433,3912653,3912653,3775352,3775352,3418776,3418776,3327411,3327411,3243538,3243538,3212681,3212681,3126350,3126350,3039863,3039863,3029116,3029116,2895199,2895199,2851370,2851370,2788223,2788223,2765367,2765367,2757797,2757797,2742641,2742641,2650157,2650157,2554584,2554584,2516329,2516329,2515412,2515412,2481436,2481436,2474528,2474528,2465855,2465855,2453754,2453754,2446852,2446852,2428360,2428360,2421180,2421180,2405699,2405699,2402388,2402388,2348559,2348559,2344765,2344765,2317563,2317563,2300588,2300588,2299039,2299039,2294499,2294499,2273327,2273327,2272836,2272836,2267708,2267708,2245083,2245083,2243378,2243378,2208784,2208784,2194807,2194807,2192524,2192524,2164150,2164150,2138822,2138822,2131479,2131479,2128015,2128015,2126031,2126031,2103137,2103137,2067256,2067256,2066791,2066791,2065382,2065382,2060156,2060156,2057412,2057412,2053123,2053123,2045520,2045520,2021092,2021092,2020066,2020066,2019173,2019173,1988578,1988578,1965509,1965509,1956557,1956557,1952220,1952220,1944956,1944956,1913379,1913379,1908056,1908056,1906360,1906360,1900371,1900371,1896178,1896178,1872657,1872657,1871414,1871414,1866144,1866144,1863127,1863127,1861138,1861138,1860847,1860847,1859713,1859713,1842844,1842844,1841691,1841691,1838978,1838978,1824588,1824588,1815262,1815262,1774795,1774795,1763159,1763159,1741055,1741055,1733219,1733219,1733163,1733163,1727339,1727339,1698442,1698442,1696305,1696305,1689079,1689079,1686065,1686065,1685748,1685748,1682013,1682013,1676188,1676188,1661209,1661209,1649029,1649029,1642118,1642118,1641521,1641521,1637141,1637141,1629762,1629762,1624855,1624855,1616000,1616000,1609820,1609820,1604721,1604721,1601072,1601072,1597597,1597597,1596447,1596447,1591298,1591298,1574705,1574705,1565570,1565570,1564686,1564686,1559185,1559185,1553275,1553275,1552923,1552923,1549903,1549903,1547340,1547340,1546689,1546689,1541185,1541185,1533216,1533216,1532615,1532615,1531331,1531331,1530391,1530391,1525101,1525101,1524542,1524542,1522307,1522307,1516628,1516628,1515067,1515067,1512256,1512256,1506474,1506474,1503339,1503339,1499001,1499001,1492251,1492251,1480500,1480500,1470360,1470360,1463062,1463062,1460012,1460012,1445718,1445718,1442194,1442194,1435479,1435479,1434600,1434600,1421127,1421127,1418583,1418583,1415807,1415807,1415049,1415049,1413116,1413116,1411189,1411189,1407303,1407303,1401894,1401894,1395507,1395507,1391461,1391461,1389911,1389911,1385631,1385631,1381189,1381189,1376660,1376660,1374219,1374219,1369807,1369807,1368927,1368927,1366540,1366540,1360873,1360873,1355310,1355310,1351940,1351940,1351529,1351529,1350615,1350615,1347654,1347654,1346467,1346467,1340074,1340074,1338940,1338940,1332726,1332726,1332375,1332375,1331669,1331669,1328654,1328654,1324698,1324698,1323914,1323914,1321765,1321765,1319077,1319077,1316233,1316233,1314580,1314580,1312025,1312025,1308677,1308677,1307210,1307210,1304976,1304976,1304286,1304286,1303546,1303546,1302029,1302029,1296626,1296626,1295772,1295772,1291716,1291716,1286271,1286271,1284780,1284780,1283390,1283390,1279995,1279995,1279127,1279127,1278898,1278898,1278787,1278787,1277791,1277791,1275925,1275925,1273407,1273407,1271730,1271730,1270170,1270170,1265832,1265832,1262549,1262549,1262037,1262037,1260560,1260560,1259797,1259797,1256657,1256657,1255499,1255499,1252068,1252068,1250488,1250488,1248547,1248547,1246327,1246327,1242196,1242196,1241175,1241175,1235034,1235034,1229795,1229795,1229629,1229629,1228171,1228171,1226794,1226794,1225973,1225973,1222381,1222381,1221070,1221070,1219091,1219091,1218810,1218810,1215373,1215373,1207777,1207777,1201642,1201642,1198249,1198249,1195858,1195858,1194456,1194456,1185309,1185309,1184741,1184741,1182112,1182112,1180180,1180180,1173566,1173566,1172801,1172801,1169614,1169614,1167411,1167411,1167229,1167229,1163552,1163552,1163245,1163245,1162949,1162949,1162381,1162381,1161300,1161300,1160842,1160842,1160347,1160347,1159284,1159284,1158648,1158648,1157978,1157978,1155951,1155951,1153750,1153750,1150533,1150533,1145209,1145209,1144538,1144538,1140539,1140539,1140436,1140436,1140368,1140368,1133476,1133476,1130233,1130233,1129387,1129387,1125641,1125641,1125178,1125178,1124753,1124753,1123857,1123857,1122426,1122426,1118100,1118100,1114433,1114433,1110820,1110820,1108569,1108569,1108360,1108360,1107897,1107897,1107325,1107325,1106246,1106246,1103062,1103062,1099513,1099513,1098900,1098900,1098745,1098745,1097816,1097816,1097224,1097224,1092938,1092938,1092426,1092426,1090983,1090983,1090346,1090346,1089734,1089734,1087233,1087233,1086056,1086056,1083492,1083492,1082977,1082977,1082835,1082835,1082587,1082587,1077622,1077622,1077088,1077088,1076790,1076790,1076120,1076120,1075769,1075769,1075397,1075397,1075292,1075292,1066804,1066804,1066346,1066346,1063588,1063588,1063427,1063427,1061074,1061074,1059444,1059444,1059334,1059334,1057905,1057905,1056263,1056263,1056119,1056119,1055754,1055754,1054445,1054445,1053401,1053401,1049608,1049608,1049048,1049048,1043801,1043801,1031925,1031925,1029907,1029907,1028438,1028438,1027192,1027192,1023390,1023390,1023362,1023362,1022292,1022292,1021854,1021854,1020691,1020691,1016001,1016001,1014672,1014672,1014313,1014313,1014171,1014171,1013649,1013649,1011971,1011971,1011290,1011290,994554,994554,992754,992754,992571,992571,990609,990609,987976,987976,987265,987265,985847,985847,984689,984689,982438,982438,979583,979583,979302,979302,978176,978176,975339,975339,972970,972970,969648,969648,967736,967736,967249,967249,967004,967004,966414,966414,965674,965674,961611,961611,960899,960899,956394,956394,956064,956064,953933,953933,952790,952790,952299,952299,949000,949000,945019,945019,943035,943035,941916,941916,939436,939436,939225,939225,936943,936943,933972,933972,933732,933732,933147,933147,932743,932743,932377,932377,932313,932313,932027,932027,931263,931263,928556,928556,927885,927885,927538,927538,927251,927251,926548,926548,923165,923165,920972,920972,919906,919906,919600,919600,918190,918190,917696,917696,917414,917414,917248,917248,914827,914827,912133,912133,911581,911581,909830,909830,908967,908967,906969,906969,904818,904818,903982,903982,903339,903339,900782,900782,897340,897340,897070,897070,896641,896641,896359,896359,895309,895309,892776,892776,890862,890862,890193,890193,889257,889257,888304,888304,886645,886645,886055,886055,885925,885925,885209,885209,884874,884874,883473,883473,882723,882723,882624,882624,882061,882061,881938,881938,877353,877353,876897,876897,876255,876255,873733,873733,873010,873010,872243,872243,871648,871648,871145,871145,870963,870963,870900,870900,869660,869660,869507,869507,866810,866810,866722,866722,865411,865411,865340,865340,864734,864734,861758,861758,861699,861699,861568,861568,861338,861338,858153,858153,857616,857616,857443,857443,856866,856866,856160,856160,855626,855626,854521,854521,853949,853949,853763,853763,853027,853027,851502,851502,850782,850782,848359,848359,848236,848236,846634,846634,846027,846027,845480,845480,844151,844151,843381,843381,842663,842663,842447,842447,840899,840899,840351,840351,840085,840085,839575,839575,838600,838600,838483,838483,836707,836707,835412,835412,831742,831742,831589,831589,830403,830403,829834,829834,829788,829788,827933,827933,826935,826935,824931,824931,824885,824885,823451,823451,823419,823419,821036,821036,820565,820565,819968,819968,819467,819467,816431,816431,813266,813266,812606,812606,812578,812578,811770,811770,810990,810990,810915,810915,810779,810779,807335,807335,805746,805746,803832,803832,803665,803665,802953,802953,801332,801332,801299,801299,800601,800601,800312,800312,799767,799767,799580,799580,799090,799090,795196,795196,795121,795121,793532,793532,792382,792382,792342,792342,792006,792006,791141,791141,790122,790122,789177,789177,788805,788805,788518,788518,787368,787368,787301,787301,787015,787015,786885,786885,786853,786853,786247,786247,785650,785650,784698,784698,783383,783383,782911,782911,781854,781854,779907,779907,779122,779122,778789,778789,777700,777700,776008,776008,773453,773453,772750,772750,772197,772197,771857,771857,771213,771213,770630,770630,769841,769841,769665,769665,763632,763632,763196,763196,762737,762737,762721,762721,761852,761852,761632,761632,761391,761391,761373,761373,760707,760707,760017,760017,759892,759892,758837,758837,758613,758613,758338,758338,755831,755831,755674,755674,755180,755180,752779,752779,752511,752511,752306,752306,751664,751664,751465,751465,751198,751198,749332,749332,748896,748896,748851,748851,748716,748716,748163,748163,747801,747801,747325,747325,747114,747114,745601,745601,745248,745248,743765,743765,739256,739256,736805,736805,734637,734637,734561,734561,733107,733107,731702,731702,731587,731587,730878,730878,730609,730609,728135,728135,727794,727794,727386,727386,726979,726979,724911,724911,724443,724443,724270,724270,724234,724234,723735,723735,723230,723230,722469,722469,722239,722239,721741,721741,721025,721025,720866,720866,719461,719461,719094,719094,717365,717365,717048,717048,716995,716995,716813,716813,716557,716557,716494,716494,716397,716397,715632,715632,714229,714229,714181,714181,711760,711760,710807,710807,709802,709802,709744,709744,709581,709581,709409,709409,708599,708599,708489,708489,708421,708421,708126,708126,707868,707868,707749,707749,707681,707681,707418,707418,706964,706964,706494,706494,705869,705869,705815,705815,705252,705252,704382,704382,703521,703521,701794,701794,700919,700919,700883,700883,700054,700054,699133,699133,698769,698769,697105,697105,696467,696467,696430,696430,694908,694908,694440,694440,694367,694367,693472,693472,692805,692805,692788,692788,692420,692420,692322,692322,692154,692154,692109,692109,691453,691453,691361,691361,690986,690986,690875,690875,690607,690607,689873,689873,689734,689734,689383,689383,688815,688815,688734,688734,688708,688708,688552,688552,688028,688028,687884,687884,685847,685847,685377,685377,684231,684231,683807,683807,683770,683770,683279,683279,682512,682512,682383,682383,682035,682035,680406,680406,679362,679362,678860,678860,677452,677452,677266,677266,677205,677205,675822,675822,675322,675322,674625,674625,673905,673905,673813,673813,672621,672621,672315,672315,671596,671596,670506,670506,668738,668738,668179,668179,667753,667753,667681,667681,666917,666917,666695,666695,666494,666494,666337,666337,666062,666062,665938,665938,664838,664838,664115,664115,663520,663520,663248,663248,663229,663229,663212,663212,662721,662721,662189,662189,661905,661905,661857,661857,660506,660506,660299,660299,659755,659755,659101,659101,658513,658513,658050,658050,657840,657840,657148,657148,654250,654250,653107,653107,652165,652165,652078,652078,652047,652047,651297,651297,651112,651112,650592,650592,650428,650428,648401,648401,648147,648147,647317,647317,646772,646772,644396,644396,644246,644246,644121,644121,644022,644022,643786,643786,643723,643723,642371,642371,642342,642342,641709,641709,641586,641586,641490,641490,641147,641147,640730,640730,640592,640592,640518,640518,640464,640464,640052,640052,639340,639340,638734,638734,638506,638506,637811,637811,637466,637466,637252,637252,635274,635274,635131,635131,634962,634962,634663,634663,634441,634441,634302,634302,633696,633696,633351,633351,633277,633277,633142,633142,632131,632131,631975,631975,631542,631542,631284,631284,631237,631237,629785,629785,629592,629592,629322,629322,629119,629119,628261,628261,628043,628043,626600,626600,626495,626495,625812,625812,625229,625229,625205,625205,625112,625112,624917,624917,623692,623692,623590,623590,623375,623375,622728,622728,622604,622604,622561,622561,621517,621517,621293,621293,620954,620954,619077,619077,618940,618940,618782,618782,618003,618003,617918,617918,617869,617869,617403,617403,616944,616944,615646,615646,615506,615506,615490,615490,615384,615384,615095,615095,614707,614707,614280,614280,613830,613830,613385,613385,613248,613248,613019,613019,612657,612657,611870,611870,610005,610005,609779,609779,609641,609641,609334,609334,609176,609176,608660,608660,608470,608470,608208,608208,606566,606566,606107,606107,605951,605951,605532,605532,604381,604381,604003,604003,603963,603963,603879,603879,603285,603285,603096,603096,602842,602842,601994,601994,601376,601376,600853,600853,600714,600714,598899,598899,598712,598712,598468,598468,598448,598448,598198,598198,598149,598149,596024,596024,595612,595612,594954,594954,594490,594490,593076,593076,592765,592765,592554,592554,592456,592456,592053,592053,591305,591305,590371,590371,590311,590311,589448,589448,589188,589188,586638,586638,586462,586462,586325,586325,586206,586206,585799,585799,585634,585634,585488,585488,585193,585193,585041,585041,584949,584949,584783,584783,584398,584398,583915,583915,583225,583225,583105,583105,583043,583043,582936,582936,582206,582206,581831,581831,581758,581758,580847,580847,580577,580577,580513,580513,580388,580388,580012,580012,579735,579735,578636,578636,578335,578335,578172,578172,577966,577966,577594,577594,577105,577105,576558,576558,576237,576237,575886,575886,575341,575341,574781,574781,574416,574416,574179,574179,573187,573187,572249,572249,571828,571828,571318,571318,570988,570988,570740,570740,570644,570644,569574,569574,569355,569355,569071,569071,569045,569045,568969,568969,567824,567824,567762,567762,567563,567563,567156,567156,566833,566833,566769,566769,565989,565989,565963,565963,565362,565362,564013,564013,563592,563592,563356,563356,563112,563112,562857,562857,562081,562081,561460,561460,561264,561264,561080,561080,560679,560679,560503,560503,560351,560351,559686,559686,559629,559629,559237,559237,559118,559118,559006,559006,558660,558660,558507,558507,558448,558448,558400,558400,558209,558209,558110,558110,557777,557777,557161,557161,557007,557007,556812,556812,556642,556642,555852,555852,555770,555770,555561,555561,555122,555122,555012,555012,554959,554959,554610,554610,553973,553973,553224,553224,552029,552029,551267,551267,550189,550189,549479,549479,549004,549004,548891,548891,548613,548613,547712,547712,547574,547574,547129,547129,547079,547079,546861,546861,546604,546604,546396,546396,546376,546376,546152,546152,545891,545891,545835,545835,545738,545738,545706,545706,545341,545341,545038,545038,544744,544744,544693,544693,544658,544658,544467,544467,544089,544089,543740,543740,543093,543093,541935,541935,541464,541464,541387,541387,541303,541303,539788,539788,539769,539769,539065,539065,538869,538869,537831,537831,536613,536613,535322,535322,534980,534980,534909,534909,534149,534149,533796,533796,533681,533681,533635,533635,533467,533467,532914,532914,532755,532755,532511,532511,532151,532151,531544,531544,530941,530941,530531,530531,530056,530056,530015,530015,529832,529832,528958,528958,528817,528817,528622,528622,528428,528428,528255,528255,528214,528214,528075,528075,527925,527925,527049,527049,527007,527007,526965,526965,526607,526607,526317,526317,525729,525729,525619,525619,525342,525342,523887,523887,523796,523796,523702,523702,523508,523508,522883,522883,522799,522799,522398,522398,521952,521952,521855,521855,521717,521717,521181,521181,520818,520818,520457,520457,520113,520113,519930,519930,519723,519723,519698,519698,519510,519510,519185,519185,519021,519021,518998,518998,518598,518598,518530,518530,518073,518073,517846,517846,517751,517751,517621,517621,517442,517442,517362,517362,517262,517262,516939,516939,516707,516707,516067,516067,515897,515897,515880,515880,515762,515762,515744,515744,515617,515617,515244,515244,514782,514782,514445,514445,514309,514309,514199,514199,513571,513571,513529,513529,512325,512325,511624,511624,510860,510860,510662,510662,510582,510582,510313,510313,509780,509780,509730,509730,509488,509488,509160,509160,508662,508662,508511,508511,507336,507336,507173,507173,506691,506691,506123,506123,505856,505856,505540,505540,505493,505493,505454,505454,505410,505410,505074,505074,504366,504366,504233,504233,503619,503619,503591,503591,503485,503485,503277,503277,503257,503257,503189,503189,503087,503087,501917,501917,501553,501553,501097,501097,500932,500932,500869,500869,500094,500094,500046,500046,499940,499940,499739,499739,499603,499603,499496,499496,499328,499328,498231,498231,497231,497231,496877,496877,496692,496692,496516,496516,496430,496430,496076,496076,496036,496036,495644,495644,495136,495136,494835,494835,494622,494622,494539,494539,494204,494204,493969,493969,493917,493917,493755,493755,493418,493418,492414,492414,492260,492260,492218,492218,492102,492102,491889,491889,491718,491718,491581,491581,491457,491457,491401,491401,491098,491098,490979,490979,490790,490790,490756,490756,490720,490720,490556,490556,490178,490178,490048,490048,489743,489743,488884,488884,488772,488772,488337,488337,488234,488234,488201,488201,487924,487924,487495,487495,487409,487409,487008,487008,486874,486874,486405,486405,486375,486375,486234,486234,486054,486054,485911,485911,485401,485401,484654,484654,484328,484328,483836,483836,483603,483603,483386,483386,483260,483260,482915,482915,482796,482796,482266,482266,482090,482090,482021,482021,481198,481198,481174,481174,480101,480101,479881,479881,479816,479816,479108,479108,478631,478631,478276,478276,478033,478033,477799,477799,477759,477759,477672,477672,477219,477219,477162,477162,476492,476492,476376,476376,476342,476342,476164,476164,475561,475561,474991,474991,474894,474894,474581,474581,474555,474555,474401,474401,473906,473906,473442,473442,473376,473376,472733,472733,472712,472712,472681,472681,472170,472170,471832,471832,471661,471661,471381,471381,470994,470994,469959,469959,469792,469792,469705,469705,469399,469399,469267,469267,469142,469142,468847,468847,468764,468764,468180,468180,468157,468157,467705,467705,467426,467426,467341,467341,465246,465246,464997,464997,464940,464940,464922,464922,464828,464828,464610,464610,464547,464547,464359,464359,464164,464164,464116,464116,464015,464015,463842,463842,463190,463190,462628,462628,462200,462200,462091,462091,461698,461698,461340,461340,461181,461181,460999,460999,460950,460950,460862,460862,460784,460784,460642,460642,460008,460008,459884,459884,459859,459859,459787,459787,459458,459458,459428,459428,458744,458744,458498,458498,458464,458464,458358,458358,458085,458085,457768,457768,457545,457545,457519,457519,457460,457460,457092,457092,456719,456719,456553,456553,456384,456384,456091,456091,455908,455908,455539,455539,455417,455417,455308,455308,455054,455054,454953,454953,454466,454466,454381,454381,454341,454341,454232,454232,453241,453241,453160,453160,452956,452956,452662,452662,452621,452621,452562,452562,452374,452374,452005,452005,451729,451729,451646,451646,451321,451321,451215,451215,451148,451148,450993,450993,450562,450562,449918,449918,449895,449895,449278,449278,449110,449110,448534,448534,448204,448204,448070,448070,448036,448036,448004,448004,447719,447719,447100,447100,447036,447036,446749,446749,446281,446281,446243,446243,446187,446187,445875,445875,445741,445741,444987,444987,444791,444791,444092,444092,444068,444068,443960,443960,443765,443765,443628,443628,443559,443559,443293,443293,442893,442893,442863,442863,442817,442817,442440,442440,442194,442194,442167,442167,441997,441997,441929,441929,441699,441699,441415,441415,441357,441357,441308,441308,440994,440994,440876,440876,440703,440703,440546,440546,439650,439650,439613,439613,439491,439491,439448,439448,439182,439182,438375,438375,438290,438290,437792,437792,437435,437435,437156,437156,436885,436885,436755,436755,436316,436316,436189,436189,435904,435904,435539,435539,435481,435481,434825,434825,434809,434809,434610,434610,433834,433834,432988,432988,432565,432565,432450,432450,432364,432364,432259,432259,432028,432028,431926,431926,431843,431843,430925,430925,430842,430842,430771,430771,430737,430737,430383,430383,430244,430244,430141,430141,429799,429799,429665,429665,429477,429477,429410,429410,429288,429288,429001,429001,428648,428648,428533,428533,428322,428322,428019,428019,427983,427983,427778,427778,427354,427354,427333,427333,427154,427154,427017,427017,426746,426746,426378,426378,426133,426133,426052,426052,425810,425810,425739,425739,425449,425449,425413,425413,425376,425376,425275,425275,425228,425228,425082,425082,425058,425058,425030,425030,424805,424805,424464,424464,424408,424408,424228,424228,424051,424051,423998,423998,423903,423903,423756,423756,423058,423058,422937,422937,422683,422683,422388,422388,422269,422269,422203,422203,422164,422164,422065,422065,421979,421979,421718,421718,421590,421590,421030,421030,420972,420972,420924,420924,420901,420901,420875,420875,420797,420797,420704,420704,420668,420668,420586,420586,420380,420380,419965,419965,419623,419623,418963,418963,418874,418874,418705,418705,418640,418640,418601,418601,418462,418462,418216,418216,417605,417605,417403,417403,416973,416973,416720,416720,416603,416603,416564,416564,416510,416510,415851,415851,415663,415663,415621,415621,414969,414969,414782,414782,414735,414735,414205,414205,414114,414114,413877,413877,413837,413837,413589,413589,413394,413394,413274,413274,413253,413253,412418,412418,412223,412223,411957,411957,411604,411604,411502,411502,411463,411463,411302,411302,411135,411135,411091,411091,410836,410836,410798,410798,410558,410558,410201,410201,410162,410162,409962,409962,409823,409823,409501,409501,409357,409357,409114,409114,408975,408975,408825,408825,408627,408627,408496,408496,408082,408082,408045,408045,407463,407463,407194,407194,407047,407047,406942,406942,406836,406836,406659,406659,406485,406485,406387,406387,405991,405991,405685,405685,405630,405630,405239,405239,405030,405030,404917,404917,404811,404811,404722,404722,404597,404597,404565,404565,404384,404384,404056,404056,403501,403501,403391,403391,403218,403218,403158,403158,403022,403022,402873,402873,402574,402574,402519,402519,401859,401859,401802,401802,401706,401706,401457,401457,401239,401239,401127,401127,400884,400884,400349,400349,400217,400217,400069,400069,399993,399993,399976,399976,399864,399864,399585,399585,399540,399540,399504,399504,399484,399484,399167,399167,399141,399141,398906,398906,398789,398789,398767,398767,398536,398536,398373,398373,397844,397844,397821,397821,397486,397486,396978,396978,396722,396722,396610,396610,396387,396387,396093,396093,395935,395935,395864,395864,395700,395700,395657,395657,395235,395235,395093,395093,395075,395075,395033,395033,395017,395017,394995,394995,394934,394934,394590,394590,394312,394312,394279,394279,394237,394237,394166,394166,393910,393910,393569,393569,393371,393371,393129,393129,392865,392865,392631,392631,392499,392499,392252,392252,392136,392136,392093,392093,391840,391840,391355,391355,391273,391273,391169,391169,391018,391018,390770,390770,390270,390270,390237,390237,390030,390030,389719,389719,389610,389610,389480,389480,389274,389274,389162,389162,389078,389078,388822,388822,388776,388776,388700,388700,388245,388245,388122,388122,388025,388025,387821,387821,387700,387700,387658,387658,387612,387612,387522,387522,387426,387426,387080,387080,386961,386961,386570,386570,386482,386482,386319,386319,385960,385960,385565,385565,385489,385489,385326,385326,385171,385171,385121,385121,384528,384528,384027,384027,383764,383764,383728,383728,383703,383703,383579,383579,383483,383483,383379,383379,383331,383331,383310,383310,383007,383007,382967,382967,382944,382944,382797,382797,382682,382682,382493,382493,382396,382396,381995,381995,381799,381799,381764,381764,381727,381727,381524,381524,381440,381440,381123,381123,381052,381052,381031,381031,380990,380990,380965,380965,380688,380688,380646,380646,380447,380447,380338,380338,380099,380099,379948,379948,379832,379832,379666,379666,379485,379485,379030,379030,378726,378726,378546,378546,378091,378091,378073,378073,377876,377876,377827,377827,377751,377751,377662,377662,377533,377533,377516,377516,377402,377402,377231,377231,376721,376721,376607,376607,376299,376299,376159,376159,376107,376107,376064,376064,375990,375990,375839,375839,375758,375758,375632,375632,375528,375528,375500,375500,375404,375404,375219,375219,374905,374905,374859,374859,374824,374824,374570,374570,374462,374462,374292,374292,374041,374041,374003,374003,373893,373893,373181,373181,373055,373055,372880,372880,372767,372767,372720,372720,372571,372571,372072,372072,371991,371991,371965,371965,371907,371907,371667,371667,371610,371610,371579,371579,371173,371173,370487,370487,370318,370318,370145,370145,370123,370123,370061,370061,369858,369858,369782,369782,369302,369302,369218,369218,369124,369124,368871,368871,368820,368820,368784,368784,368713,368713,368660,368660,368415,368415,367916,367916,367875,367875,367845,367845,367670,367670,367548,367548,367447,367447,367352,367352,367236,367236,367093,367093,366956,366956,366880,366880,366850,366850,366727,366727,366660,366660,366271,366271,366221,366221,365985,365985,365851,365851,365787,365787,365740,365740,365647,365647,365500,365500,365244,365244,365140,365140,365087,365087,365068,365068,364970,364970,364891,364891,364799,364799,364759,364759,364563,364563,364519,364519,364477,364477,364360,364360,364118,364118,364002,364002,363667,363667,363577,363577,363429,363429,363274,363274,363151,363151,362519,362519,362424,362424,362292,362292,362193,362193,362091,362091,362017,362017,361950,361950,361844,361844,361183,361183,361106,361106,361064,361064,360984,360984,360832,360832,360707,360707,360670,360670,360568,360568,360460,360460,359952,359952,359744,359744,359547,359547,359441,359441,359002,359002,358978,358978,358894,358894,358618,358618,357892,357892,357828,357828,357729,357729,357239,357239,356846,356846,356774,356774,356740,356740,356703,356703,356356,356356,356273,356273,356170,356170,356091,356091,356044,356044,355899,355899,355503,355503,355457,355457,355113,355113,355092,355092,354955,354955,354780,354780,354495,354495,354458,354458,354365,354365,354298,354298,353473,353473,353354,353354,353303,353303,353227,353227,353002,353002,352722,352722,352688,352688,352562,352562,352434,352434,352312,352312,352281,352281,352265,352265,352247,352247,352153,352153,352057,352057,351755,351755,351694,351694,351623,351623,351201,351201,351099,351099,351074,351074,350940,350940,350729,350729,350122,350122,350043,350043,350005,350005,349826,349826,349635,349635,349617,349617,349586,349586,349357,349357,349240,349240,349223,349223,349181,349181,349137,349137,349060,349060,348507,348507,348289,348289,348192,348192,348104,348104,348063,348063,347835,347835,347806,347806,347586,347586,347464,347464,347439,347439,347355,347355,347306,347306,347247,347247,347131,347131,347108,347108,347049,347049,346800,346800,346526,346526,346445,346445,346333,346333,346181,346181,346033,346033,346017,346017,345829,345829,345805,345805,345135,345135,345059,345059,344915,344915,344810,344810,344769,344769,344590,344590,344535,344535,344517,344517,344359,344359,344273,344273,344221,344221,344070,344070,343910,343910,343522,343522,343474,343474,343454,343454,343402,343402,343136,343136,342852,342852,342595,342595,342577,342577,342489,342489,342252,342252,342188,342188,341945,341945,341867,341867,341846,341846,341678,341678,341623,341623,341594,341594,341384,341384,341269,341269,341161,341161,341110,341110,341083,341083,340844,340844,340818,340818,340650,340650,340564,340564,340375,340375,340350,340350,340320,340320,340267,340267,340148,340148,339964,339964,339872,339872,339843,339843,339608,339608,339441,339441,339302,339302,339127,339127,339109,339109,339078,339078,338951,338951,338407,338407,338359,338359,338320,338320,338263,338263,337699,337699,337672,337672,337622,337622,337591,337591,337536,337536,337391,337391,337126,337126,337007,337007,336832,336832,336791,336791,336595,336595,336339,336339,336051,336051,336011,336011,335515,335515,335457,335457,335285,335285,335052,335052,334922,334922,334229,334229,334205,334205,334073,334073,333973,333973,333826,333826,333774,333774,333758,333758,333639,333639,333441,333441,333381,333381,333126,333126,332991,332991,332943,332943,332730,332730,332696,332696,332547,332547,332506,332506,332229,332229,332185,332185,331841,331841,331778,331778,331734,331734,331419,331419,331350,331350,331326,331326,331288,331288,331214,331214,331006,331006,330944,330944,330773,330773,330697,330697,330633,330633,330309,330309,330215,330215,330083,330083,330051,330051,330001,330001,329962,329962,329585,329585,329558,329558,329488,329488,329125,329125,329105,329105,329053,329053,328949,328949,328829,328829,328723,328723,328697,328697,328365,328365,328190,328190,328033,328033,327896,327896,327573,327573,327481,327481,327305,327305,327220,327220,327202,327202,327134,327134,326899,326899,326511,326511,326141,326141,326088,326088,326050,326050,326019,326019,325915,325915,325424,325424,325134,325134,324894,324894,324773,324773,324516,324516,324464,324464,324414,324414,324322,324322,324253,324253,324208,324208,324180,324180,323985,323985,323921,323921,323608,323608,323290,323290,323274,323274,322993,322993,322864,322864,322836,322836,322781,322781,322579,322579,322563,322563,322458,322458,322166,322166,322115,322115,321897,321897,321830,321830,321603,321603,321507,321507,321484,321484,321448,321448,321047,321047,320987,320987,320951,320951,320897,320897,320782,320782,320239,320239,320115,320115,320019,320019,319956,319956,319677,319677,319643,319643,319378,319378,319312,319312,318881,318881,318573,318573,318348,318348,318078,318078,318007,318007,317858,317858,317719,317719,317665,317665,317588,317588,317471,317471,317412,317412,317213,317213,317129,317129,316969,316969,316904,316904,316739,316739,316647,316647,316607,316607,316209,316209,316149,316149,316038,316038,315946,315946,315868,315868,315752,315752,315707,315707,315448,315448,315389,315389,315313,315313,314813,314813,314303,314303,314174,314174,314077,314077,313883,313883,313778,313778,313718,313718,313596,313596,313373,313373,313264,313264,313175,313175,313083,313083,312977,312977,312514,312514,312107,312107,312075,312075,311979,311979,311846,311846,311792,311792,311769,311769,311649,311649,311629,311629,311578,311578,311259,311259,311215,311215,311171,311171,311155,311155,311118,311118,311027,311027,310851,310851,310812,310812,310662,310662,310507,310507,310177,310177,309987,309987,309953,309953,309926,309926,309745,309745,309650,309650,309274,309274,309124,309124,308921,308921,308809,308809,308581,308581,308541,308541,308508,308508,308419,308419,308274,308274,308165,308165,308122,308122,307979,307979,307941,307941,307918,307918,307895,307895,307747,307747,307725,307725,307707,307707,307565,307565,307510,307510,307432,307432,307191,307191,307163,307163,307081,307081,306954,306954,306929,306929,306853,306853,306829,306829,306583,306583,306540,306540,306491,306491,306416,306416,306196,306196,306022,306022,305774,305774,305593,305593,305327,305327,305200,305200,305118,305118,305099,305099,304951,304951,304932,304932,304700,304700,304602,304602,304381,304381,304365,304365,304307,304307,303663,303663,303579,303579,303385,303385,303270,303270,303212,303212,303074,303074,302921,302921,302768,302768,302677,302677,302316,302316,302246,302246,301870,301870,301766,301766,301607,301607,301196,301196,301167,301167,301108,301108,301089,301089,301010,301010,300905,300905,300846,300846,300372,300372,300327,300327,300298,300298,300252,300252,300105,300105,299944,299944,299757,299757,299730,299730,299601,299601,299559,299559,299478,299478,299425,299425,299363,299363,299037,299037,298956,298956,298906,298906,298813,298813,298723,298723,298668,298668,298452,298452,298125,298125,298105,298105,298056,298056,297972,297972,297880,297880,297800,297800,297608,297608,297592,297592,297150,297150,296927,296927,296873,296873,296755,296755,296715,296715,296591,296591,296389,296389,296289,296289,296232,296232,296212,296212,296134,296134,296079,296079,296012,296012,295914,295914,295813,295813,295701,295701,295571,295571,295469,295469,295357,295357,295233,295233,295046,295046,294954,294954,294846,294846,294822,294822,294654,294654,294594,294594,294547,294547,294367,294367,294346,294346,294300,294300,294194,294194,293836,293836,293626,293626,293508,293508,293481,293481,293095,293095,293075,293075,293059,293059,292972,292972,292874,292874,292839,292839,292716,292716,292608,292608,292578,292578,292445,292445,292318,292318,292266,292266,292220,292220,292173,292173,292063,292063,291958,291958,291916,291916,291877,291877,291829,291829,291755,291755,291472,291472,291440,291440,291367,291367,291218,291218,291063,291063,290904,290904,290825,290825,290627,290627,290592,290592,290447,290447,290339,290339,290053,290053,290031,290031,289990,289990,289964,289964,289789,289789,289721,289721,289670,289670,289607,289607,289577,289577,289187,289187,289132,289132,289116,289116,288954,288954,288923,288923,288858,288858,288808,288808,288593,288593,288228,288228,287922,287922,287721,287721,287672,287672,287554,287554,287405,287405,287345,287345,287143,287143,287112,287112,286990,286990,286884,286884,286825,286825,286723,286723,286673,286673,286596,286596,286571,286571,286537,286537,286485,286485,286021,286021,285861,285861,285811,285811,285621,285621,285596,285596,285243,285243,285082,285082,285011,285011,284820,284820,284774,284774,284297,284297,284244,284244,284210,284210,284147,284147,284107,284107,283872,283872,283485,283485,283350,283350,283236,283236,283164,283164,282978,282978,282700,282700,282582,282582,282562,282562,282536,282536,282414,282414,282352,282352,282310,282310,282285,282285,282255,282255,282139,282139,282120,282120,282066,282066,281918,281918,281856,281856,281797,281797,281729,281729,281691,281691,281551,281551,281431,281431,281311,281311,281090,281090,281023,281023,281007,281007,280969,280969,280790,280790,280767,280767,280664,280664,280584,280584,280488,280488,280463,280463,280419,280419,280283,280283,280245,280245,280095,280095,280032,280032,279900,279900,279875,279875,279831,279831,279719,279719,279343,279343,279234,279234,279038,279038,278978,278978,278932,278932,278848,278848,278814,278814,278727,278727,278690,278690,278534,278534,278489,278489,278469,278469,278453,278453,278380,278380,278313,278313,278190,278190,278117,278117,278028,278028,277900,277900,277689,277689,277657,277657,277629,277629,277482,277482,277416,277416,277376,277376,277356,277356,277289,277289,277196,277196,277010,277010,276871,276871,276845,276845,276812,276812,276764,276764,276714,276714,276461,276461,276167,276167,276109,276109,276049,276049,275886,275886,275578,275578,275487,275487,275471,275471,275397,275397,275271,275271,275152,275152,275093,275093,275001,275001,274934,274934,274870,274870,274762,274762,274690,274690,274396,274396,274054,274054,273985,273985,273966,273966,273903,273903,273750,273750,273638,273638,273616,273616,273500,273500,273382,273382,273334,273334,273075,273075,273015,273015,272908,272908,272885,272885,272790,272790,272722,272722,272456,272456,272370,272370,272209,272209,271873,271873,271635,271635,271619,271619,271468,271468,271249,271249,271190,271190,271174,271174,270942,270942,270844,270844,270682,270682,270655,270655,270618,270618,270451,270451,270372,270372,270322,270322,270253,270253,269753,269753,269590,269590,269444,269444,269345,269345,269195,269195,269151,269151,269024,269024,268938,268938,268773,268773,268549,268549,268338,268338,268303,268303,268141,268141,268043,268043,267942,267942,267130,267130,267104,267104,267032,267032,266994,266994,266709,266709,266593,266593,266462,266462,266383,266383,266337,266337,266228,266228,266119,266119,265874,265874,265751,265751,265568,265568,265491,265491,265414,265414,265361,265361,265306,265306,265268,265268,265222,265222,265096,265096,264823,264823,264560,264560,264445,264445,264355,264355,264263,264263,264108,264108,264066,264066,264039,264039,263911,263911,263776,263776,263620,263620,263457,263457,263434,263434,263153,263153,263071,263071,262893,262893,262840,262840,262728,262728,262695,262695,262609,262609,262583,262583,262509,262509,262436,262436,262349,262349,262146,262146,262083,262083,262051,262051,262007,262007,261765,261765,261642,261642,261589,261589,261490,261490,261383,261383,261310,261310,261287,261287,261247,261247,261140,261140,261045,261045,260752,260752,260710,260710,260464,260464,260416,260416,260375,260375,260256,260256,260181,260181,260119,260119,259758,259758,259648,259648,259449,259449,259303,259303,259263,259263,259126,259126,259090,259090,259062,259062,259037,259037,259015,259015,258899,258899,258878,258878,258744,258744,258573,258573,258468,258468,258392,258392,258367,258367,258201,258201,258171,258171,258137,258137,257685,257685,257595,257595,257562,257562,257544,257544,257477,257477,257411,257411,257367,257367,257218,257218,257197,257197,257159,257159,257059,257059,256979,256979,256880,256880,256807,256807,256763,256763,256576,256576,256531,256531,256476,256476,256448,256448,256407,256407,256333,256333,256036,256036,256020,256020,255909,255909,255847,255847,255779,255779,255628,255628,255612,255612,255465,255465,255366,255366,255320,255320,255189,255189,255121,255121,255072,255072,255012,255012,254950,254950,254868,254868,254750,254750,254689,254689,254618,254618,254507,254507,254411,254411,254365,254365,254336,254336,254005,254005,253969,253969,253949,253949,253903,253903,253854,253854,253819,253819,253559,253559,253495,253495,253458,253458,253421,253421,253069,253069,253003,253003,252978,252978,252807,252807,252716,252716,252616,252616,252565,252565,252518,252518,252360,252360,252289,252289,252101,252101,252052,252052,251979,251979,251922,251922,251463,251463,251399,251399,251376,251376,251291,251291,251184,251184,251134,251134,251054,251054,250969,250969,250936,250936,250897,250897,250869,250869,250851,250851,250335,250335,250212,250212,250068,250068,250045,250045,249774,249774,249729,249729,249666,249666,249521,249521,249462,249462,249411,249411,249318,249318,249272,249272,249237,249237,249187,249187,249140,249140,248965,248965,248913,248913,248873,248873,248848,248848,248803,248803,248764,248764,248739,248739,248664,248664,248606,248606,248437,248437,248285,248285,248136,248136,248082,248082,248035,248035,248018,248018,247991,247991,247973,247973,247549,247549,247467,247467,247444,247444,247420,247420,247372,247372,247343,247343,247277,247277,247104,247104,247083,247083,247067,247067,246941,246941,246853,246853,246664,246664,246532,246532,246505,246505,246466,246466,246360,246360,246317,246317,246274,246274,246251,246251,246169,246169,246132,246132,245972,245972,245859,245859,245809,245809,245041,245041,244960,244960,244942,244942,244922,244922,244851,244851,244725,244725,244706,244706,244689,244689,244666,244666,244612,244612,244556,244556,244495,244495,244290,244290,244196,244196,244175,244175,244062,244062,244009,244009,243859,243859,243679,243679,243643,243643,243532,243532,243456,243456,243125,243125,243051,243051,242884,242884,242865,242865,242849,242849,242697,242697,242571,242571,242397,242397,242284,242284,242237,242237,242199,242199,242001,242001,241968,241968,241920,241920,241815,241815,241770,241770,241707,241707,241595,241595,241520,241520,241356,241356,241299,241299,241267,241267,241251,241251,241211,241211,241045,241045,240910,240910,240851,240851,240804,240804,240728,240728,240646,240646,240191,240191,240062,240062,239971,239971,239930,239930,239848,239848,239806,239806,239627,239627,239591,239591,239575,239575,239471,239471,239416,239416,239268,239268,239238,239238,239100,239100,238968,238968,238883,238883,238819,238819,238760,238760,238564,238564,238323,238323,238106,238106,237967,237967,237887,237887,237851,237851,237829,237829,237723,237723,237641,237641,237511,237511,237470,237470,237434,237434,237410,237410,237203,237203,237176,237176,237114,237114,237082,237082,236828,236828,236656,236656,236618,236618,236484,236484,236355,236355,236303,236303,236277,236277,236235,236235,235973,235973,235883,235883,235793,235793,235667,235667,235556,235556,235513,235513,235348,235348,235267,235267,235176,235176,235030,235030,234971,234971,234943,234943,234888,234888,234845,234845,234773,234773,234739,234739,234606,234606,234441,234441,234380,234380,234192,234192,234148,234148,234060,234060,233967,233967,233849,233849,233769,233769,233540,233540,233511,233511,233430,233430,233382,233382,233184,233184,233075,233075,233024,233024,232960,232960,232903,232903,232715,232715,232654,232654,232606,232606,232571,232571,232451,232451,232195,232195,232107,232107,232085,232085,231985,231985,231893,231893,231838,231838,231786,231786,231734,231734,231679,231679,231608,231608,231545,231545,231523,231523,231392,231392,231371,231371,231350,231350,231277,231277,231261,231261,231229,231229,231213,231213,231194,231194,231137,231137,231044,231044,230991,230991,230929,230929,230886,230886,230852,230852,230821,230821,230729,230729,230595,230595,230365,230365,230332,230332,230274,230274,230189,230189,230060,230060,229948,229948,229840,229840,229776,229776,229742,229742,229574,229574,229424,229424,229404,229404,229376,229376,229228,229228,228862,228862,228803,228803,228583,228583,228508,228508,228447,228447,228400,228400,228340,228340,228320,228320,228260,228260,228146,228146,228090,228090,227793,227793,227753,227753,227673,227673,227605,227605,227556,227556,227492,227492,227471,227471,227398,227398,227350,227350,227261,227261,227172,227172,227152,227152,227046,227046,226945,226945,226786,226786,226737,226737,226614,226614,226543,226543,226423,226423,226392,226392,226236,226236,226086,226086,226068,226068,225854,225854,225801,225801,225672,225672,225408,225408,225385,225385,225350,225350,225255,225255,225188,225188,225158,225158,225107,225107,225089,225089,224982,224982,224707,224707,224683,224683,224660,224660,224543,224543,224375,224375,224259,224259,224128,224128,224100,224100,224081,224081,224029,224029,223866,223866,223826,223826,223778,223778,223749,223749,223715,223715,223580,223580,223355,223355,223287,223287,223198,223198,223044,223044,223002,223002,222969,222969,222911,222911,222808,222808,222789,222789,222715,222715,222480,222480,222337,222337,222236,222236,222147,222147,222095,222095,222020,222020,221921,221921,221866,221866,221563,221563,221537,221537,221419,221419,221194,221194,221018,221018,220892,220892,220742,220742,220670,220670,220356,220356,220328,220328,220159,220159,220103,220103,220050,220050,220028,220028,219946,219946,219778,219778,219722,219722,219640,219640,219500,219500,219362,219362,219241,219241,219136,219136,219088,219088,218931,218931,218703,218703,218530,218530,218331,218331,218314,218314,218187,218187,218155,218155,218087,218087,218019,218019,217949,217949,217915,217915,217703,217703,217663,217663,217535,217535,217489,217489,217358,217358,217164,217164,217140,217140,217121,217121,217059,217059,216913,216913,216838,216838,216788,216788,216772,216772,216686,216686,216628,216628,216512,216512,216426,216426,216298,216298,216147,216147,216076,216076,216060,216060,216024,216024,215987,215987,215957,215957,215859,215859,215836,215836,215820,215820,215730,215730,215604,215604,215583,215583,215370,215370,215163,215163,215138,215138,214949,214949,214908,214908,214891,214891,214859,214859,214792,214792,214772,214772,214738,214738,214435,214435,214186,214186,214162,214162,214087,214087,213836,213836,213801,213801,213727,213727,213686,213686,213638,213638,213563,213563,213407,213407,213320,213320,213222,213222,213183,213183,213081,213081,212985,212985,212908,212908,212886,212886,212769,212769,212681,212681,212665,212665,212617,212617,212600,212600,212567,212567,212533,212533,212517,212517,212490,212490,212159,212159,212046,212046,211617,211617,211571,211571,211493,211493,211383,211383,211365,211365,211315,211315,211224,211224,211179,211179,211065,211065,211049,211049,210920,210920,210856,210856,210791,210791,210768,210768,210656,210656,210562,210562,210384,210384,210282,210282,210194,210194,210113,210113,209968,209968,209884,209884,209824,209824,209770,209770,209604,209604,209545,209545,209528,209528,209509,209509,209374,209374,209233,209233,209132,209132,209100,209100,208915,208915,208887,208887,208871,208871,208624,208624,208603,208603,208291,208291,208273,208273,208254,208254,208143,208143,208033,208033,207959,207959,207929,207929,207909,207909,207864,207864,207795,207795,207763,207763,207666,207666,207580,207580,207460,207460,207431,207431,207392,207392,207351,207351,207317,207317,207295,207295,207234,207234,207206,207206,207157,207157,207088,207088,206949,206949,206737,206737,206652,206652,206628,206628,206596,206596,206578,206578,206551,206551,206535,206535,206408,206408,206379,206379,206313,206313,206278,206278,206252,206252,206230,206230,206207,206207,206076,206076,205974,205974,205876,205876,205549,205549,205224,205224,205206,205206,205155,205155,205060,205060,205008,205008,204980,204980,204956,204956,204881,204881,204806,204806,204689,204689,204617,204617,204593,204593,204349,204349,204316,204316,204274,204274,204210,204210,204151,204151,204110,204110,204071,204071,204042,204042,203968,203968,203894,203894,203878,203878,203768,203768,203641,203641,203532,203532,203497,203497,203479,203479,203281,203281,203260,203260,203194,203194,203148,203148,203109,203109,202847,202847,202821,202821,202797,202797,202681,202681,202568,202568,202508,202508,202328,202328,202236,202236,202143,202143,202123,202123,202063,202063,202027,202027,201924,201924,201893,201893,201874,201874,201823,201823,201739,201739,201679,201679,201594,201594,201568,201568,201443,201443,201303,201303,201264,201264,201193,201193,201155,201155,200967,200967,200884,200884,200823,200823,200797,200797,200697,200697,200668,200668,200637,200637,200604,200604,200343,200343,200301,200301,200272,200272,200240,200240,200187,200187,200073,200073,199995,199995,199942,199942,199900,199900,199757,199757,199738,199738,199686,199686,199667,199667,199591,199591,199469,199469,199379,199379,199335,199335,199219,199219,199201,199201,199153,199153,199109,199109,199029,199029,198886,198886,198844,198844,198786,198786,198767,198767,198510,198510,198452,198452,198434,198434,198401,198401,198377,198377,198348,198348,198320,198320,198257,198257,198211,198211,198173,198173,198143,198143,198044,198044,198010,198010,197876,197876,197712,197712,197691,197691,197595,197595,197555,197555,197505,197505,197451,197451,197276,197276,197149,197149,197056,197056,196776,196776,196757,196757,196730,196730,196472,196472,196332,196332,196286,196286,196240,196240,196085,196085,196049,196049,195996,195996,195910,195910,195817,195817,195715,195715,195687,195687,195555,195555,195422,195422,195373,195373,195341,195341,195316,195316,195226,195226,195194,195194,195117,195117,195094,195094,195015,195015,194978,194978,194962,194962,194867,194867,194825,194825,194752,194752,194714,194714,194600,194600,194584,194584,194496,194496,194329,194329,194260,194260,194196,194196,194136,194136,194081,194081,194030,194030,193977,193977,193910,193910,193872,193872,193803,193803,193768,193768,193734,193734,193668,193668,193637,193637,193620,193620,193541,193541,193467,193467,193408,193408,193374,193374,193308,193308,193212,193212,193196,193196,193159,193159,193104,193104,193034,193034,193014,193014,192992,192992,192920,192920,192811,192811,192708,192708,192674,192674,192653,192653,192480,192480,192393,192393,192375,192375,192345,192345,192323,192323,192255,192255,192137,192137,192112,192112,192071,192071,191945,191945,191889,191889,191828,191828,191593,191593,191491,191491,191442,191442,191288,191288,191227,191227,191196,191196,191157,191157,191134,191134,191112,191112,191061,191061,191014,191014,190986,190986,190891,190891,190871,190871,190817,190817,190750,190750,190530,190530,190246,190246,190200,190200,190152,190152,190040,190040,190019,190019,189966,189966,189933,189933,189886,189886,189778,189778,189741,189741,189676,189676,189630,189630,189544,189544,189356,189356,189224,189224,189180,189180,189119,189119,189050,189050,188987,188987,188968,188968,188929,188929,188902,188902,188798,188798,188770,188770,188739,188739,188646,188646,188622,188622,188583,188583,188554,188554,188500,188500,188382,188382,188271,188271,188108,188108,187924,187924,187883,187883,187844,187844,187753,187753,187635,187635,187565,187565,187544,187544,187489,187489,187325,187325,187154,187154,187127,187127,187111,187111,187087,187087,187049,187049,186960,186960,186942,186942,186899,186899,186863,186863,186837,186837,186816,186816,186781,186781,186749,186749,186677,186677,186392,186392,186290,186290,186196,186196,186060,186060,186012,186012,185915,185915,185857,185857,185795,185795,185758,185758,185702,185702,185642,185642,185512,185512,185377,185377,185328,185328,185197,185197,185083,185083,185043,185043,185002,185002,184950,184950,184919,184919,184900,184900,184866,184866,184837,184837,184747,184747,184675,184675,184639,184639,184623,184623,184581,184581,184548,184548,184513,184513,184331,184331,184276,184276,184251,184251,184220,184220,184157,184157,184129,184129,184102,184102,184017,184017,183917,183917,183847,183847,183821,183821,183802,183802,183702,183702,183597,183597,183579,183579,183521,183521,183495,183495,183443,183443,183348,183348,183326,183326,183249,183249,183172,183172,183124,183124,183031,183031,183011,183011,182924,182924,182865,182865,182740,182740,182705,182705,182689,182689,182657,182657,182519,182519,182475,182475,182415,182415,182370,182370,182227,182227,182181,182181,182128,182128,182085,182085,182045,182045,182023,182023,181838,181838,181593,181593,181536,181536,181499,181499,181396,181396,181334,181334,181311,181311,181233,181233,181194,181194,181153,181153,181107,181107,181051,181051,181034,181034,180899,180899,180596,180596,180496,180496,180480,180480,180458,180458,180360,180360,180303,180303,180136,180136,180107,180107,180082,180082,180026,180026,179986,179986,179832,179832,179796,179796,179748,179748,179696,179696,179671,179671,179570,179570,179466,179466,179426,179426,179395,179395,179368,179368,179299,179299,179267,179267,179140,179140,179101,179101,179033,179033,178970,178970,178870,178870,178850,178850,178643,178643,178601,178601,178561,178561,178541,178541,178472,178472,178357,178357,178339,178339,178313,178313,178222,178222,178199,178199,178145,178145,178032,178032,178016,178016,177979,177979,177956,177956,177935,177935,177881,177881,177828,177828,177781,177781,177758,177758,177739,177739,177688,177688,177426,177426,177351,177351,177234,177234,177167,177167,177051,177051,177029,177029,176997,176997,176975,176975,176955,176955,176846,176846,176823,176823,176797,176797,176739,176739,176593,176593,176456,176456,176434,176434,176386,176386,176258,176258,176096,176096,176031,176031,175957,175957,175909,175909,175884,175884,175847,175847,175507,175507,175312,175312,175233,175233,175195,175195,175168,175168,175073,175073,174831,174831,174755,174755,174681,174681,174637,174637,174592,174592,174571,174571,174550,174550,174529,174529,174469,174469,174374,174374,174321,174321,174301,174301,174210,174210,174153,174153,174120,174120,174037,174037,173969,173969,173922,173922,173896,173896,173855,173855,173773,173773,173739,173739,173663,173663,173611,173611,173595,173595,173564,173564,173513,173513,173474,173474,173440,173440,173323,173323,173251,173251,173215,173215,173148,173148,173056,173056,173009,173009,172987,172987,172925,172925,172853,172853,172825,172825,172591,172591,172532,172532,172494,172494,172412,172412,172303,172303,172177,172177,172103,172103,172047,172047,171855,171855,171813,171813,171791,171791,171759,171759,171736,171736,171705,171705,171687,171687,171661,171661,171548,171548,171523,171523,171471,171471,171412,171412,171395,171395,171288,171288,171269,171269,171244,171244,171155,171155,171005,171005,170956,170956,170863,170863,170845,170845,170827,170827,170803,170803,170741,170741,170640,170640,170617,170617,170597,170597,170530,170530,170505,170505,170354,170354,170255,170255,170124,170124,170052,170052,170003,170003,169887,169887,169849,169849,169831,169831,169786,169786,169720,169720,169564,169564,169492,169492,169384,169384,169311,169311,169215,169215,169156,169156,169086,169086,169011,169011,168993,168993,168943,168943,168912,168912,168869,168869,168840,168840,168792,168792,168769,168769,168691,168691,168656,168656,168598,168598,168577,168577,168543,168543,168443,168443,168412,168412,168377,168377,168313,168313,168204,168204,168091,168091,168069,168069,167924,167924,167705,167705,167676,167676,167648,167648,167542,167542,167495,167495,167456,167456,167321,167321,167300,167300,167240,167240,167098,167098,167047,167047,167020,167020,166882,166882,166843,166843,166688,166688,166658,166658,166614,166614,166538,166538,166472,166472,166343,166343,166249,166249,166198,166198,166179,166179,166145,166145,166063,166063,165998,165998,165955,165955,165837,165837,165815,165815,165603,165603,165574,165574,165527,165527,165403,165403,165382,165382,165347,165347,165329,165329,165295,165295,165279,165279,165259,165259,165136,165136,165098,165098,164965,164965,164926,164926,164882,164882,164838,164838,164812,164812,164778,164778,164690,164690,164648,164648,164580,164580,164525,164525,164428,164428,164383,164383,164287,164287,164250,164250,164226,164226,164159,164159,164120,164120,163987,163987,163965,163965,163922,163922,163889,163889,163867,163867,163752,163752,163660,163660,163638,163638,163616,163616,163589,163589,163496,163496,163336,163336,163314,163314,163053,163053,163005,163005,162859,162859,162837,162837,162790,162790,162727,162727,162693,162693,162649,162649,162568,162568,162532,162532,162477,162477,162411,162411,162308,162308,162288,162288,162253,162253,162227,162227,162195,162195,162125,162125,161995,161995,161974,161974,161909,161909,161886,161886,161819,161819,161655,161655,161566,161566,161522,161522,161467,161467,161446,161446,161351,161351,161327,161327,161275,161275,161197,161197,161166,161166,161130,161130,161018,161018,160999,160999,160912,160912,160805,160805,160777,160777,160689,160689,160617,160617,160536,160536,160500,160500,160436,160436,160415,160415,160355,160355,160301,160301,160256,160256,160229,160229,160207,160207,160187,160187,160163,160163,160089,160089,160031,160031,159893,159893,159844,159844,159766,159766,159728,159728,159703,159703,159674,159674,159624,159624,159567,159567,159406,159406,159380,159380,159355,159355,159262,159262,159206,159206,159131,159131,159035,159035,158814,158814,158798,158798,158762,158762,158745,158745,158681,158681,158622,158622,158482,158482,158463,158463,158418,158418,158387,158387,158371,158371,158267,158267,158235,158235,158211,158211,158129,158129,158084,158084,157979,157979,157927,157927,157873,157873,157848,157848,157814,157814,157775,157775,157615,157615,157517,157517,157482,157482,157333,157333,157265,157265,157237,157237,157117,157117,157078,157078,157012,157012,156949,156949,156931,156931,156854,156854,156773,156773,156746,156746,156558,156558,156538,156538,156512,156512,156433,156433,156392,156392,156371,156371,156320,156320,156279,156279,156236,156236,156134,156134,156038,156038,156017,156017,155885,155885,155867,155867,155770,155770,155692,155692,155585,155585,155551,155551,155486,155486,155435,155435,155361,155361,155336,155336,155288,155288,155188,155188,155165,155165,155069,155069,154989,154989,154969,154969,154899,154899,154865,154865,154810,154810,154762,154762,154698,154698,154665,154665,154638,154638,154609,154609,154556,154556,154534,154534,154507,154507,154460,154460,154422,154422,154382,154382,154366,154366,154302,154302,154234,154234,154202,154202,154170,154170,154136,154136,154107,154107,154065,154065,153932,153932,153815,153815,153740,153740,153708,153708,153687,153687,153617,153617,153573,153573,153466,153466,153400,153400,153325,153325,153272,153272,153228,153228,153191,153191,153169,153169,153097,153097,152944,152944,152926,152926,152904,152904,152748,152748,152708,152708,152653,152653,152610,152610,152579,152579,152537,152537,152510,152510,152453,152453,152370,152370,152341,152341,152323,152323,152258,152258,152216,152216,152194,152194,152061,152061,152002,152002,151953,151953,151870,151870,151834,151834,151754,151754,151735,151735,151660,151660,151632,151632,151476,151476,151438,151438,151395,151395,151368,151368,151347,151347,151296,151296,151275,151275,151254,151254,151129,151129,151091,151091,151056,151056,151021,151021,150979,150979,150893,150893,150838,150838,150808,150808,150751,150751,150687,150687,150568,150568,150490,150490,150446,150446,150396,150396,150356,150356,150326,150326,150303,150303,150274,150274,150253,150253,150209,150209,150151,150151,149990,149990,149853,149853,149823,149823,149803,149803,149782,149782,149738,149738,149691,149691,149665,149665,149572,149572,149524,149524,149429,149429,149344,149344,149291,149291,149170,149170,149153,149153,149084,149084,149035,149035,148942,148942,148868,148868,148730,148730,148708,148708,148658,148658,148592,148592,148498,148498,148482,148482,148457,148457,148421,148421,148382,148382,148348,148348,148313,148313,148293,148293,148276,148276,148136,148136,148003,148003,147924,147924,147906,147906,147833,147833,147707,147707,147645,147645,147596,147596,147576,147576,147486,147486,147439,147439,147361,147361,147325,147325,147292,147292,147270,147270,147254,147254,147237,147237,147122,147122,147084,147084,147027,147027,146990,146990,146941,146941,146811,146811,146771,146771,146718,146718,146700,146700,146658,146658,146557,146557,146540,146540,146449,146449,146422,146422,146386,146386,146364,146364,146345,146345,146300,146300,146280,146280,146169,146169,146153,146153,146122,146122,146104,146104,145983,145983,145893,145893,145870,145870,145792,145792,145759,145759,145705,145705,145685,145685,145660,145660,145622,145622,145493,145493,145365,145365,145266,145266,145249,145249,145198,145198,145139,145139,145076,145076,145052,145052,145016,145016,144997,144997,144863,144863,144764,144764,144722,144722,144688,144688,144619,144619,144529,144529,144506,144506,144479,144479,144400,144400,144373,144373,144312,144312,144261,144261,144178,144178,144142,144142,144101,144101,144069,144069,143881,143881,143797,143797,143705,143705,143680,143680,143617,143617,143583,143583,143513,143513,143475,143475,143317,143317,143282,143282,143253,143253,143226,143226,143210,143210,143142,143142,143091,143091,143063,143063,142921,142921,142840,142840,142781,142781,142761,142761,142738,142738,142685,142685,142669,142669,142605,142605,142482,142482,142458,142458,142408,142408,142365,142365,142334,142334,142313,142313,142189,142189,142152,142152,142100,142100,142030,142030,141960,141960,141917,141917,141868,141868,141811,141811,141744,141744,141596,141596,141537,141537,141456,141456,141412,141412,141348,141348,141301,141301,141270,141270,141243,141243,141184,141184,141141,141141,141116,141116,141100,141100,141073,141073,141015,141015,140973,140973,140848,140848,140807,140807,140748,140748,140720,140720,140692,140692,140640,140640,140614,140614,140583,140583,140565,140565,140481,140481,140458,140458,140401,140401,140355,140355,140324,140324,140302,140302,140245,140245,140166,140166,140145,140145,140128,140128,140096,140096,140023,140023,139889,139889,139855,139855,139781,139781,139751,139751,139720,139720,139669,139669,139638,139638,139608,139608,139555,139555,139493,139493,139431,139431,139363,139363,139347,139347,139303,139303,139280,139280,139258,139258,139223,139223,139125,139125,139073,139073,139045,139045,139016,139016,138977,138977,138782,138782,138718,138718,138655,138655,138616,138616,138516,138516,138396,138396,138371,138371,138322,138322,138281,138281,138258,138258,138220,138220,138159,138159,138120,138120,138061,138061,138022,138022,138005,138005,137981,137981,137949,137949,137930,137930,137874,137874,137853,137853,137758,137758,137709,137709,137675,137675,137643,137643,137601,137601,137580,137580,137542,137542,137514,137514,137494,137494,137458,137458,137426,137426,137409,137409,137345,137345,137323,137323,137296,137296,137220,137220,137191,137191,137158,137158,137137,137137,137025,137025,137006,137006,136969,136969,136909,136909,136854,136854,136785,136785,136744,136744,136712,136712,136694,136694,136644,136644,136392,136392,136331,136331,136262,136262,136183,136183,136114,136114,136043,136043,136017,136017,135916,135916,135899,135899,135880,135880,135830,135830,135661,135661,135593,135593,135532,135532,135497,135497,135457,135457,135424,135424,135329,135329,135308,135308,135269,135269,135218,135218,135170,135170,135038,135038,135022,135022,134916,134916,134884,134884,134861,134861,134785,134785,134716,134716,134620,134620,134583,134583,134547,134547,134491,134491,134461,134461,134432,134432,134370,134370,134315,134315,134273,134273,134165,134165,134102,134102,134079,134079,134047,134047,134031,134031,133946,133946,133827,133827,133805,133805,133768,133768,133689,133689,133667,133667,133651,133651,133606,133606,133577,133577,133534,133534,133507,133507,133436,133436,133402,133402,133359,133359,133343,133343,133298,133298,133259,133259,133142,133142,133118,133118,133060,133060,133032,133032,133015,133015,132991,132991,132962,132962,132935,132935,132909,132909,132859,132859,132839,132839,132785,132785,132764,132764,132742,132742,132706,132706,132660,132660,132535,132535,132471,132471,132397,132397,132372,132372,132337,132337,132306,132306,132264,132264,132226,132226,132129,132129,132055,132055,132029,132029,131937,131937,131912,131912,131850,131850,131811,131811,131785,131785,131701,131701,131655,131655,131607,131607,131544,131544,131443,131443,131376,131376,131333,131333,131310,131310,131281,131281,131246,131246,131218,131218,131201,131201,131183,131183,130997,130997,130935,130935,130916,130916,130880,130880,130849,130849,130789,130789,130773,130773,130750,130750,130688,130688,130598,130598,130545,130545,130482,130482,130429,130429,130321,130321,130276,130276,130215,130215,130172,130172,130153,130153,130132,130132,130112,130112,130095,130095,130071,130071,130047,130047,129992,129992,129942,129942,129891,129891,129816,129816,129795,129795,129728,129728,129699,129699,129683,129683,129653,129653,129604,129604,129532,129532,129444,129444,129333,129333,129239,129239,129201,129201,129174,129174,129119,129119,128964,128964,128925,128925,128899,128899,128764,128764,128687,128687,128666,128666,128586,128586,128511,128511,128487,128487,128459,128459,128442,128442,128402,128402,128308,128308,128224,128224,128159,128159,128101,128101,128077,128077,128025,128025,127998,127998,127960,127960,127906,127906,127886,127886,127816,127816,127794,127794,127712,127712,127683,127683,127631,127631,127598,127598,127581,127581,127494,127494,127453,127453,127408,127408,127375,127375,127314,127314,127282,127282,127246,127246,127225,127225,127177,127177,127158,127158,127121,127121,127017,127017,126933,126933,126910,126910,126848,126848,126817,126817,126753,126753,126673,126673,126648,126648,126610,126610,126575,126575,126552,126552,126500,126500,126444,126444,126421,126421,126329,126329,126308,126308,126291,126291,126229,126229,126211,126211,126047,126047,126014,126014,125987,125987,125924,125924,125887,125887,125848,125848,125819,125819,125774,125774,125682,125682,125650,125650,125619,125619,125515,125515,125497,125497,125384,125384,125352,125352,125249,125249,125231,125231,125180,125180,125154,125154,125104,125104,125001,125001,124950,124950,124922,124922,124904,124904,124875,124875,124843,124843,124815,124815,124746,124746,124709,124709,124638,124638,124611,124611,124566,124566,124495,124495,124462,124462,124420,124420,124395,124395,124347,124347,124293,124293,124277,124277,124061,124061,124016,124016,123991,123991,123962,123962,123906,123906,123862,123862,123822,123822,123788,123788,123720,123720,123643,123643,123622,123622,123487,123487,123449,123449,123433,123433,123398,123398,123347,123347,123312,123312,123267,123267,123189,123189,123172,123172,123109,123109,123074,123074,123049,123049,123014,123014,122973,122973,122908,122908,122833,122833,122815,122815,122764,122764,122638,122638,122602,122602,122546,122546,122523,122523,122477,122477,122456,122456,122437,122437,122375,122375,122347,122347,122298,122298,122281,122281,122253,122253,122208,122208,122173,122173,122110,122110,122067,122067,121988,121988,121967,121967,121925,121925,121881,121881,121847,121847,121825,121825,121809,121809,121696,121696,121628,121628,121611,121611,121510,121510,121492,121492,121452,121452,121426,121426,121398,121398,121330,121330,121303,121303,121262,121262,121209,121209,121144,121144,121095,121095,121072,121072,121049,121049,120949,120949,120835,120835,120809,120809,120770,120770,120734,120734,120683,120683,120651,120651,120615,120615,120574,120574,120556,120556,120506,120506,120378,120378,120237,120237,120191,120191,120171,120171,120152,120152,120124,120124,120033,120033,120004,120004,119986,119986,119924,119924,119908,119908,119869,119869,119852,119852,119782,119782,119757,119757,119714,119714,119690,119690,119629,119629,119592,119592,119560,119560,119544,119544,119510,119510,119435,119435,119402,119402,119382,119382,119359,119359,119317,119317,119283,119283,119266,119266,119235,119235,119211,119211,119184,119184,119147,119147,119115,119115,119099,119099,119061,119061,119014,119014,118993,118993,118953,118953,118927,118927,118891,118891,118834,118834,118785,118785,118718,118718,118662,118662,118642,118642,118621,118621,118579,118579,118520,118520,118495,118495,118471,118471,118453,118453,118363,118363,118313,118313,118285,118285,118246,118246,118196,118196,118147,118147,118117,118117,118091,118091,118072,118072,117972,117972,117955,117955,117909,117909,117834,117834,117808,117808,117790,117790,117754,117754,117656,117656,117623,117623,117607,117607,117555,117555,117507,117507,117484,117484,117450,117450,117432,117432,117414,117414,117395,117395,117301,117301,117252,117252,117227,117227,117176,117176,117126,117126,117105,117105,117085,117085,117060,117060,117040,117040,117016,117016,116959,116959,116886,116886,116807,116807,116753,116753,116726,116726,116678,116678,116546,116546,116515,116515,116479,116479,116461,116461,116417,116417,116336,116336,116268,116268,116237,116237,116215,116215,116153,116153,116119,116119,116067,116067,116050,116050,116030,116030,115970,115970,115925,115925,115864,115864,115832,115832,115780,115780,115757,115757,115732,115732,115691,115691,115533,115533,115454,115454,115435,115435,115413,115413,115394,115394,115350,115350,115332,115332,115301,115301,115277,115277,115248,115248,115222,115222,115198,115198,115180,115180,115146,115146,115087,115087,115023,115023,115005,115005,114985,114985,114961,114961,114937,114937,114880,114880,114863,114863,114822,114822,114803,114803,114774,114774,114718,114718,114683,114683,114648,114648,114623,114623,114551,114551,114357,114357,114339,114339,114303,114303,114278,114278,114231,114231,114207,114207,114156,114156,114123,114123,114087,114087,114065,114065,114039,114039,114005,114005,113952,113952,113931,113931,113897,113897,113880,113880,113844,113844,113816,113816,113788,113788,113765,113765,113734,113734,113700,113700,113651,113651,113607,113607,113569,113569,113493,113493,113465,113465,113414,113414,113345,113345,113324,113324,113241,113241,113225,113225,113125,113125,113108,113108,113092,113092,113070,113070,113053,113053,113025,113025,112985,112985,112938,112938,112921,112921,112877,112877,112852,112852,112806,112806,112774,112774,112746,112746,112719,112719,112701,112701,112683,112683,112642,112642,112565,112565,112510,112510,112475,112475,112402,112402,112384,112384,112358,112358,112305,112305,112288,112288,112254,112254,112224,112224,112180,112180,112124,112124,112107,112107,112089,112089,112000,112000,111984,111984,111926,111926,111898,111898,111842,111842,111819,111819,111782,111782,111764,111764,111724,111724,111707,111707,111613,111613,111563,111563,111544,111544,111424,111424,111356,111356,111339,111339,111322,111322,111284,111284,111260,111260,111218,111218,111158,111158,111127,111127,111105,111105,111043,111043,111017,111017,110996,110996,110943,110943,110879,110879,110854,110854,110783,110783,110749,110749,110683,110683,110648,110648,110621,110621,110494,110494,110475,110475,110447,110447,110328,110328,110299,110299,110275,110275,110235,110235,110200,110200,110104,110104,110082,110082,110025,110025,109995,109995,109972,109972,109954,109954,109908,109908,109880,109880,109861,109861,109811,109811,109741,109741,109702,109702,109664,109664,109639,109639,109622,109622,109579,109579,109498,109498,109423,109423,109407,109407,109378,109378,109344,109344,109304,109304,109238,109238,109222,109222,109194,109194,109175,109175,109124,109124,109090,109090,109045,109045,109028,109028,109000,109000,108975,108975,108954,108954,108931,108931,108882,108882,108790,108790,108753,108753,108717,108717,108701,108701,108596,108596,108555,108555,108425,108425,108396,108396,108364,108364,108345,108345,108324,108324,108282,108282,108252,108252,108189,108189,108172,108172,108153,108153,108073,108073,108053,108053,108037,108037,108017,108017,107999,107999,107965,107965,107943,107943,107927,107927,107899,107899,107858,107858,107825,107825,107734,107734,107715,107715,107678,107678,107615,107615,107584,107584,107530,107530,107497,107497,107471,107471,107423,107423,107401,107401,107380,107380,107348,107348,107295,107295,107233,107233,107161,107161,107140,107140,107115,107115,107086,107086,107058,107058,107037,107037,107016,107016,106999,106999,106934,106934,106760,106760,106697,106697,106674,106674,106614,106614,106589,106589,106494,106494,106454,106454,106409,106409,106340,106340,106267,106267,106225,106225,106124,106124,105997,105997,105955,105955,105925,105925,105859,105859,105816,105816,105785,105785,105740,105740,105686,105686,105655,105655,105616,105616,105593,105593,105566,105566,105538,105538,105438,105438,105408,105408,105382,105382,105300,105300,105258,105258,105174,105174,105147,105147,105075,105075,105030,105030,105009,105009,104972,104972,104934,104934,104896,104896,104877,104877,104859,104859,104830,104830,104806,104806,104699,104699,104667,104667,104587,104587,104560,104560,104514,104514,104487,104487,104431,104431,104408,104408,104377,104377,104351,104351,104319,104319,104256,104256,104197,104197,104146,104146,104130,104130,104097,104097,104035,104035,104010,104010,103991,103991,103971,103971,103948,103948,103900,103900,103868,103868,103829,103829,103794,103794,103764,103764,103735,103735,103718,103718,103700,103700,103681,103681,103628,103628,103606,103606,103575,103575,103553,103553,103521,103521,103489,103489,103467,103467,103433,103433,103409,103409,103376,103376,103299,103299,103257,103257,103218,103218,103193,103193,103166,103166,103143,103143,103114,103114,103090,103090,103034,103034,103012,103012,102953,102953,102925,102925,102863,102863,102834,102834,102812,102812,102794,102794,102767,102767,102718,102718,102636,102636,102619,102619,102573,102573,102547,102547,102529,102529,102511,102511,102475,102475,102448,102448,102424,102424,102388,102388,102323,102323,102169,102169,102148,102148,102131,102131,102099,102099,102082,102082,102065,102065,102048,102048,101931,101931,101885,101885,101852,101852,101826,101826,101800,101800,101693,101693,101667,101667,101648,101648,101593,101593,101556,101556,101523,101523,101501,101501,101474,101474,101438,101438,101413,101413,101374,101374,101337,101337,101271,101271,101244,101244,101209,101209,101189,101189,101164,101164,101132,101132,101030,101030,100982,100982,100955,100955,100928,100928,100893,100893,100864,100864,100839,100839,100815,100815,100778,100778,100726,100726,100686,100686,100644,100644,100606,100606,100562,100562,100450,100450,100426,100426,100302,100302,100255,100255,100236,100236,100158,100158,100122,100122,100050,100050,100016,100016,100000,100000,99963,99963,99923,99923,99876,99876,99843,99843,99781,99781,99761,99761,99745,99745,99726,99726,99702,99702,99660,99660,99630,99630,99587,99587,99521,99521,99457,99457,99438,99438,99397,99397,99293,99293,99271,99271,99198,99198,99180,99180,99052,99052,99036,99036,99010,99010,98976,98976,98844,98844,98793,98793,98769,98769,98716,98716,98655,98655,98615,98615,98565,98565,98545,98545,98511,98511,98490,98490,98446,98446,98402,98402,98378,98378,98356,98356,98325,98325,98285,98285,98255,98255,98229,98229,98185,98185,98147,98147,98045,98045,98000,98000,97941,97941,97828,97828,97805,97805,97776,97776,97744,97744,97696,97696,97679,97679,97617,97617,97598,97598,97551,97551,97514,97514,97489,97489,97472,97472,97456,97456,97381,97381,97354,97354,97301,97301,97272,97272,97210,97210,97183,97183,97149,97149,97126,97126,97103,97103,97077,97077,97059,97059,97013,97013,96955,96955,96917,96917,96849,96849,96744,96744,96725,96725,96688,96688,96621,96621,96569,96569,96506,96506,96478,96478,96451,96451,96401,96401,96336,96336,96315,96315,96285,96285,96252,96252,96218,96218,96160,96160,96110,96110,96042,96042,95989,95989,95973,95973,95942,95942,95849,95849,95829,95829,95750,95750,95728,95728,95707,95707,95689,95689,95576,95576,95478,95478,95432,95432,95404,95404,95382,95382,95363,95363,95294,95294,95278,95278,95240,95240,95196,95196,95170,95170,95106,95106,95078,95078,94987,94987,94944,94944,94894,94894,94862,94862,94843,94843,94824,94824,94808,94808,94761,94761,94730,94730,94706,94706,94678,94678,94618,94618,94599,94599,94573,94573,94519,94519,94485,94485,94450,94450,94362,94362,94325,94325,94306,94306,94288,94288,94267,94267,94241,94241,94218,94218,94138,94138,94090,94090,93954,93954,93936,93936,93898,93898,93861,93861,93831,93831,93717,93717,93694,93694,93664,93664,93625,93625,93580,93580,93518,93518,93460,93460,93444,93444,93424,93424,93396,93396,93380,93380,93361,93361,93234,93234,93191,93191,93148,93148,93048,93048,92992,92992,92971,92971,92954,92954,92866,92866,92838,92838,92776,92776,92739,92739,92712,92712,92695,92695,92674,92674,92635,92635,92613,92613,92588,92588,92529,92529,92502,92502,92421,92421,92386,92386,92323,92323,92296,92296,92276,92276,92220,92220,92182,92182,92096,92096,92080,92080,92030,92030,91985,91985,91954,91954,91934,91934,91917,91917,91879,91879,91839,91839,91799,91799,91783,91783,91766,91766,91743,91743,91713,91713,91672,91672,91636,91636,91620,91620,91578,91578,91560,91560,91521,91521,91499,91499,91478,91478,91442,91442,91424,91424,91397,91397,91366,91366,91348,91348,91330,91330,91309,91309,91279,91279,91222,91222,91136,91136,91052,91052,91025,91025,90999,90999,90957,90957,90933,90933,90910,90910,90862,90862,90831,90831,90814,90814,90774,90774,90731,90731,90663,90663,90610,90610,90557,90557,90521,90521,90502,90502,90412,90412,90376,90376,90333,90333,90311,90311,90246,90246,90218,90218,90180,90180,90142,90142,90109,90109,90089,90089,90061,90061,90036,90036,90004,90004,89980,89980,89913,89913,89851,89851,89832,89832,89792,89792,89762,89762,89725,89725,89703,89703,89685,89685,89654,89654,89626,89626,89543,89543,89515,89515,89464,89464,89429,89429,89348,89348,89331,89331,89295,89295,89243,89243,89162,89162,89133,89133,89105,89105,89079,89079,89059,89059,89036,89036,88995,88995,88929,88929,88909,88909,88893,88893,88856,88856,88810,88810,88785,88785,88714,88714,88688,88688,88661,88661,88642,88642,88610,88610,88584,88584,88549,88549,88514,88514,88476,88476,88460,88460,88425,88425,88403,88403,88387,88387,88360,88360,88320,88320,88299,88299,88280,88280,88235,88235,88184,88184,88149,88149,88129,88129,88087,88087,88035,88035,88014,88014,87969,87969,87929,87929,87910,87910,87893,87893,87877,87877,87792,87792,87752,87752,87730,87730,87713,87713,87681,87681,87635,87635,87618,87618,87594,87594,87561,87561,87545,87545,87519,87519,87480,87480,87452,87452,87431,87431,87415,87415,87341,87341,87307,87307,87288,87288,87265,87265,87218,87218,87202,87202,87140,87140,87109,87109,87056,87056,87023,87023,87004,87004,86983,86983,86938,86938,86922,86922,86903,86903,86884,86884,86831,86831,86806,86806,86750,86750,86696,86696,86665,86665,86581,86581,86564,86564,86529,86529,86472,86472,86439,86439,86415,86415,86381,86381,86308,86308,86289,86289,86222,86222,86200,86200,86147,86147,86077,86077,86022,86022,85980,85980,85952,85952,85882,85882,85832,85832,85778,85778,85745,85745,85715,85715,85694,85694,85675,85675,85655,85655,85629,85629,85555,85555,85512,85512,85494,85494,85461,85461,85440,85440,85418,85418,85384,85384,85275,85275,85258,85258,85219,85219,85203,85203,85156,85156,85121,85121,85102,85102,85057,85057,85018,85018,84927,84927,84875,84875,84840,84840,84743,84743,84701,84701,84678,84678,84656,84656,84590,84590,84568,84568,84515,84515,84465,84465,84384,84384,84322,84322,84288,84288,84192,84192,84148,84148,84129,84129,84099,84099,84079,84079,84050,84050,83984,83984,83928,83928,83910,83910,83856,83856,83815,83815,83795,83795,83775,83775,83758,83758,83679,83679,83659,83659,83620,83620,83590,83590,83545,83545,83512,83512,83479,83479,83448,83448,83360,83360,83333,83333,83295,83295,83268,83268,83153,83153,83120,83120,83081,83081,83063,83063,83029,83029,83008,83008,82975,82975,82959,82959,82922,82922,82891,82891,82869,82869,82799,82799,82771,82771,82742,82742,82679,82679,82615,82615,82594,82594,82576,82576,82528,82528,82486,82486,82455,82455,82435,82435,82413,82413,82385,82385,82326,82326,82287,82287,82261,82261,82238,82238,82164,82164,82148,82148,82124,82124,82084,82084,82068,82068,82039,82039,82015,82015,81978,81978,81932,81932,81907,81907,81848,81848,81830,81830,81788,81788,81752,81752,81736,81736,81714,81714,81695,81695,81673,81673,81616,81616,81600,81600,81547,81547,81501,81501,81478,81478,81457,81457,81404,81404,81381,81381,81343,81343,81321,81321,81291,81291,81266,81266,81246,81246,81228,81228,81208,81208,81139,81139,81121,81121,81100,81100,81073,81073,81043,81043,81002,81002,80985,80985,80952,80952,80917,80917,80881,80881,80853,80853,80784,80784,80768,80768,80736,80736,80709,80709,80685,80685,80667,80667,80572,80572,80545,80545,80471,80471,80454,80454,80403,80403,80385,80385,80338,80338,80282,80282,80228,80228,80198,80198,80173,80173,80139,80139,80106,80106,80029,80029,79977,79977,79950,79950,79805,79805,79674,79674,79657,79657,79570,79570,79538,79538,79509,79509,79474,79474,79438,79438,79401,79401,79381,79381,79356,79356,79321,79321,79269,79269,79253,79253,79193,79193,79176,79176,79154,79154,79133,79133,79110,79110,79053,79053,79027,79027,78912,78912,78887,78887,78847,78847,78825,78825,78804,78804,78782,78782,78744,78744,78703,78703,78659,78659,78556,78556,78531,78531,78505,78505,78486,78486,78470,78470,78431,78431,78329,78329,78307,78307,78284,78284,78253,78253,78230,78230,78214,78214,78195,78195,78096,78096,78044,78044,78012,78012,77953,77953,77920,77920,77898,77898,77878,77878,77845,77845,77735,77735,77710,77710,77691,77691,77636,77636,77578,77578,77531,77531,77490,77490,77460,77460,77384,77384,77338,77338,77285,77285,77267,77267,77239,77239,77172,77172,77127,77127,77107,77107,77083,77083,77065,77065,77038,77038,77013,77013,76950,76950,76891,76891,76867,76867,76824,76824,76776,76776,76712,76712,76648,76648,76607,76607,76583,76583,76548,76548,76531,76531,76505,76505,76437,76437,76398,76398,76379,76379,76347,76347,76310,76310,76282,76282,76242,76242,76193,76193,76165,76165,76138,76138,76121,76121,76073,76073,76047,76047,76004,76004,75981,75981,75935,75935,75915,75915,75895,75895,75867,75867,75846,75846,75830,75830,75814,75814,75782,75782,75761,75761,75701,75701,75637,75637,75619,75619,75552,75552,75514,75514,75493,75493,75475,75475,75449,75449,75390,75390,75374,75374,75294,75294,75262,75262,75191,75191,75150,75150,75113,75113,75079,75079,75035,75035,74994,74994,74975,74975,74940,74940,74908,74908,74853,74853,74816,74816,74793,74793,74769,74769,74668,74668,74634,74634,74591,74591,74521,74521,74491,74491,74469,74469,74449,74449,74412,74412,74388,74388,74361,74361,74338,74338,74289,74289,74264,74264,74184,74184,74144,74144,74121,74121,74097,74097,74079,74079,74039,74039,74021,74021,73997,73997,73977,73977,73916,73916,73889,73889,73831,73831,73802,73802,73760,73760,73676,73676,73658,73658,73622,73622,73606,73606,73582,73582,73514,73514,73489,73489,73466,73466,73419,73419,73403,73403,73381,73381,73271,73271,73161,73161,73091,73091,73049,73049,73011,73011,72973,72973,72845,72845,72807,72807,72778,72778,72681,72681,72649,72649,72631,72631,72609,72609,72580,72580,72547,72547,72522,72522,72483,72483,72457,72457,72418,72418,72386,72386,72322,72322,72281,72281,72251,72251,72223,72223,72177,72177,72135,72135,72101,72101,72076,72076,72037,72037,72017,72017,71970,71970,71926,71926,71910,71910,71877,71877,71843,71843,71811,71811,71782,71782,71709,71709,71662,71662,71630,71630,71604,71604,71559,71559,71542,71542,71496,71496,71471,71471,71453,71453,71415,71415,71370,71370,71354,71354,71299,71299,71283,71283,71245,71245,71226,71226,71197,71197,71178,71178,71128,71128,71112,71112,71092,71092,71076,71076,71059,71059,71042,71042,71025,71025,70963,70963,70938,70938,70881,70881,70841,70841,70804,70804,70743,70743,70674,70674,70649,70649,70572,70572,70548,70548,70523,70523,70507,70507,70482,70482,70444,70444,70414,70414,70365,70365,70347,70347,70294,70294,70272,70272,70213,70213,70181,70181,70129,70129,70073,70073,70051,70051,70034,70034,69993,69993,69943,69943,69913,69913,69897,69897,69877,69877,69849,69849,69828,69828,69781,69781,69763,69763,69746,69746,69695,69695,69653,69653,69612,69612,69583,69583,69563,69563,69542,69542,69526,69526,69503,69503,69457,69457,69427,69427,69398,69398,69366,69366,69350,69350,69334,69334,69306,69306,69289,69289,69271,69271,69197,69197,69151,69151,69121,69121,69098,69098,69074,69074,69032,69032,68971,68971,68952,68952,68923,68923,68858,68858,68827,68827,68801,68801,68778,68778,68745,68745,68729,68729,68690,68690,68656,68656,68602,68602,68579,68579,68562,68562,68514,68514,68497,68497,68465,68465,68449,68449,68427,68427,68375,68375,68335,68335,68316,68316,68287,68287,68244,68244,68205,68205,68188,68188,68155,68155,68105,68105,68085,68085,68026,68026,68001,68001,67934,67934,67890,67890,67867,67867,67820,67820,67773,67773,67711,67711,67648,67648,67632,67632,67605,67605,67587,67587,67558,67558,67518,67518,67481,67481,67459,67459,67412,67412,67381,67381,67349,67349,67332,67332,67315,67315,67295,67295,67275,67275,67238,67238,67180,67180,67149,67149,67123,67123,67102,67102,67075,67075,67057,67057,67039,67039,67022,67022,67005,67005,66988,66988,66948,66948,66903,66903,66879,66879,66829,66829,66744,66744,66706,66706,66688,66688,66660,66660,66605,66605,66585,66585,66550,66550,66524,66524,66488,66488,66459,66459,66440,66440,66413,66413,66394,66394,66377,66377,66350,66350,66331,66331,66313,66313,66281,66281,66252,66252,66211,66211,66156,66156,66129,66129,66107,66107,66055,66055,66023,66023,66005,66005,65956,65956,65928,65928,65871,65871,65855,65855,65823,65823,65807,65807,65776,65776,65744,65744,65724,65724,65706,65706,65685,65685,65664,65664,65646,65646,65622,65622,65603,65603,65586,65586,65569,65569,65542,65542,65513,65513,65483,65483,65445,65445,65427,65427,65396,65396,65369,65369,65352,65352,65319,65319,65295,65295,65241,65241,65223,65223,65204,65204,65142,65142,65107,65107,65082,65082,65065,65065,65046,65046,65027,65027,64975,64975,64937,64937,64909,64909,64880,64880,64863,64863,64832,64832,64814,64814,64779,64779,64729,64729,64669,64669,64629,64629,64613,64613,64597,64597,64561,64561,64526,64526,64485,64485,64466,64466,64442,64442,64422,64422,64396,64396,64363,64363,64327,64327,64305,64305,64277,64277,64259,64259,64242,64242,64224,64224,64185,64185,64156,64156,64120,64120,64029,64029,63991,63991,63972,63972,63922,63922,63906,63906,63859,63859,63832,63832,63814,63814,63784,63784,63756,63756,63740,63740,63703,63703,63652,63652,63611,63611,63583,63583,63560,63560,63523,63523,63505,63505,63485,63485,63444,63444,63375,63375,63325,63325,63283,63283,63266,63266,63231,63231,63211,63211,63186,63186,63161,63161,63140,63140,63114,63114,63060,63060,63037,63037,62998,62998,62941,62941,62925,62925,62898,62898,62875,62875,62772,62772,62755,62755,62717,62717,62677,62677,62643,62643,62612,62612,62581,62581,62557,62557,62532,62532,62513,62513,62495,62495,62472,62472,62424,62424,62352,62352,62311,62311,62256,62256,62217,62217,62179,62179,62146,62146,62119,62119,62076,62076,62027,62027,62010,62010,61956,61956,61922,61922,61905,61905,61887,61887,61834,61834,61798,61798,61762,61762,61734,61734,61713,61713,61688,61688,61620,61620,61600,61600,61559,61559,61543,61543,61512,61512,61489,61489,61395,61395,61317,61317,61289,61289,61273,61273,61249,61249,61217,61217,61180,61180,61151,61151,61134,61134,61058,61058,60987,60987,60968,60968,60941,60941,60922,60922,60890,60890,60847,60847,60819,60819,60802,60802,60780,60780,60754,60754,60730,60730,60707,60707,60670,60670,60651,60651,60611,60611,60584,60584,60562,60562,60545,60545,60526,60526,60508,60508,60472,60472,60416,60416,60378,60378,60342,60342,60307,60307,60278,60278,60239,60239,60222,60222,60175,60175,60154,60154,60136,60136,60095,60095,60074,60074,60042,60042,60004,60004,59987,59987,59968,59968,59948,59948,59932,59932,59878,59878,59855,59855,59827,59827,59805,59805,59772,59772,59749,59749,59732,59732,59715,59715,59695,59695,59669,59669,59652,59652,59630,59630,59612,59612,59588,59588,59571,59571,59500,59500,59471,59471,59410,59410,59390,59390,59345,59345,59302,59302,59282,59282,59244,59244,59180,59180,59155,59155,59113,59113,59072,59072,59037,59037,58990,58990,58966,58966,58944,58944,58908,58908,58886,58886,58868,58868,58845,58845,58824,58824,58801,58801,58761,58761,58742,58742,58713,58713,58675,58675,58640,58640,58613,58613,58577,58577,58553,58553,58534,58534,58499,58499,58443,58443,58426,58426,58394,58394,58378,58378,58329,58329,58306,58306,58289,58289,58229,58229,58213,58213,58193,58193,58154,58154,58132,58132,58107,58107,58089,58089,58070,58070,58052,58052,58019,58019,58003,58003,57969,57969,57949,57949,57908,57908,57873,57873,57835,57835,57814,57814,57784,57784,57758,57758,57721,57721,57699,57699,57664,57664,57591,57591,57558,57558,57541,57541,57517,57517,57496,57496,57479,57479,57457,57457,57400,57400,57370,57370,57331,57331,57283,57283,57260,57260,57235,57235,57195,57195,57170,57170,57125,57125,57060,57060,57041,57041,57006,57006,56964,56964,56939,56939,56914,56914,56891,56891,56862,56862,56841,56841,56817,56817,56781,56781,56763,56763,56734,56734,56703,56703,56685,56685,56666,56666,56650,56650,56618,56618,56592,56592,56558,56558,56525,56525,56472,56472,56437,56437,56397,56397,56375,56375,56356,56356,56324,56324,56304,56304,56283,56283,56260,56260,56235,56235,56185,56185,56161,56161,56139,56139,56096,56096,56079,56079,56006,56006,55989,55989,55924,55924,55891,55891,55860,55860,55837,55837,55810,55810,55785,55785,55726,55726,55708,55708,55680,55680,55651,55651,55632,55632,55596,55596,55576,55576,55549,55549,55518,55518,55487,55487,55458,55458,55410,55410,55350,55350,55333,55333,55284,55284,55258,55258,55233,55233,55184,55184,55155,55155,55137,55137,55085,55085,55067,55067,55050,55050,55022,55022,55004,55004,54979,54979,54953,54953,54925,54925,54886,54886,54868,54868,54830,54830,54810,54810,54783,54783,54766,54766,54741,54741,54700,54700,54684,54684,54657,54657,54638,54638,54612,54612,54585,54585,54538,54538,54519,54519,54503,54503,54478,54478,54415,54415,54389,54389,54368,54368,54305,54305,54263,54263,54247,54247,54218,54218,54191,54191,54169,54169,54147,54147,54130,54130,54108,54108,54071,54071,54048,54048,54031,54031,54006,54006,53990,53990,53963,53963,53928,53928,53908,53908,53859,53859,53837,53837,53749,53749,53716,53716,53679,53679,53655,53655,53630,53630,53611,53611,53560,53560,53524,53524,53502,53502,53440,53440,53369,53369,53349,53349,53329,53329,53296,53296,53246,53246,53214,53214,53197,53197,53149,53149,53129,53129,53101,53101,53034,53034,53003,53003,52974,52974,52926,52926,52880,52880,52862,52862,52833,52833,52794,52794,52764,52764,52745,52745,52710,52710,52692,52692,52673,52673,52641,52641,52618,52618,52580,52580,52549,52549,52519,52519,52482,52482,52453,52453,52432,52432,52389,52389,52353,52353,52322,52322,52295,52295,52276,52276,52245,52245,52228,52228,52212,52212,52185,52185,52148,52148,52119,52119,52087,52087,52059,52059,52007,52007,51983,51983,51957,51957,51940,51940,51915,51915,51892,51892,51868,51868,51839,51839,51821,51821,51804,51804,51788,51788,51761,51761,51744,51744,51726,51726,51701,51701,51656,51656,51611,51611,51587,51587,51560,51560,51525,51525,51509,51509,51469,51469,51441,51441,51422,51422,51398,51398,51378,51378,51350,51350,51304,51304,51280,51280,51243,51243,51160,51160,51135,51135,51119,51119,51069,51069,51027,51027,51003,51003,50985,50985,50950,50950,50907,50907,50870,50870,50816,50816,50790,50790,50750,50750,50720,50720,50682,50682,50632,50632,50608,50608,50587,50587,50559,50559,50542,50542,50523,50523,50501,50501,50453,50453,50418,50418,50397,50397,50346,50346,50274,50274,50254,50254,50229,50229,50207,50207,50167,50167,50141,50141,50115,50115,50075,50075,50049,50049,50027,50027,50008,50008,49988,49988,49972,49972,49945,49945,49914,49914,49898,49898,49858,49858,49829,49829,49790,49790,49754,49754,49726,49726,49687,49687,49655,49655,49636,49636,49608,49608,49590,49590,49554,49554,49504,49504,49471,49471,49451,49451,49427,49427,49411,49411,49389,49389,49371,49371,49348,49348,49324,49324,49305,49305,49288,49288,49257,49257,49235,49235,49219,49219,49199,49199,49122,49122,49083,49083,49060,49060,49044,49044,49020,49020,48988,48988,48970,48970,48924,48924,48889,48889,48865,48865,48846,48846,48823,48823,48799,48799,48769,48769,48749,48749,48727,48727,48709,48709,48693,48693,48675,48675,48658,48658,48626,48626,48570,48570,48551,48551,48511,48511,48474,48474,48425,48425,48396,48396,48376,48376,48321,48321,48298,48298,48265,48265,48243,48243,48221,48221,48204,48204,48162,48162,48145,48145,48112,48112,48083,48083,48063,48063,48044,48044,48028,48028,47994,47994,47972,47972,47945,47945,47909,47909,47890,47890,47870,47870,47844,47844,47811,47811,47765,47765,47748,47748,47732,47732,47712,47712,47691,47691,47666,47666,47646,47646,47624,47624,47557,47557,47530,47530,47500,47500,47467,47467,47423,47423,47403,47403,47386,47386,47360,47360,47326,47326,47302,47302,47279,47279,47243,47243,47225,47225,47165,47165,47137,47137,47119,47119,47087,47087,47068,47068,46991,46991,46962,46962,46933,46933,46886,46886,46856,46856,46831,46831,46815,46815,46790,46790,46774,46774,46753,46753,46734,46734,46717,46717,46692,46692,46666,46666,46632,46632,46615,46615,46571,46571,46542,46542,46497,46497,46477,46477,46461,46461,46430,46430,46401,46401,46372,46372,46345,46345,46303,46303,46261,46261,46232,46232,46211,46211,46139,46139,46117,46117,46094,46094,46077,46077,46061,46061,46035,46035,46019,46019,46002,46002,45985,45985,45965,45965,45938,45938,45911,45911,45873,45873,45835,45835,45814,45814,45779,45779,45749,45749,45730,45730,45703,45703,45682,45682,45653,45653,45628,45628,45606,45606,45590,45590,45565,45565,45547,45547,45526,45526,45507,45507,45488,45488,45467,45467,45425,45425,45368,45368,45338,45338,45308,45308,45275,45275,45192,45192,45168,45168,45150,45150,45127,45127,45111,45111,45076,45076,45051,45051,45031,45031,45007,45007,44984,44984,44959,44959,44917,44917,44892,44892,44825,44825,44809,44809,44760,44760,44736,44736,44716,44716,44699,44699,44681,44681,44658,44658,44642,44642,44621,44621,44589,44589,44552,44552,44519,44519,44459,44459,44433,44433,44400,44400,44382,44382,44360,44360,44343,44343,44327,44327,44311,44311,44292,44292,44258,44258,44232,44232,44216,44216,44192,44192,44162,44162,44141,44141,44117,44117,44089,44089,44032,44032,44012,44012,43975,43975,43952,43952,43936,43936,43892,43892,43874,43874,43848,43848,43827,43827,43777,43777,43750,43750,43713,43713,43691,43691,43665,43665,43645,43645,43610,43610,43582,43582,43555,43555,43532,43532,43516,43516,43499,43499,43472,43472,43450,43450,43434,43434,43412,43412,43391,43391,43370,43370,43335,43335,43289,43289,43271,43271,43255,43255,43230,43230,43205,43205,43180,43180,43155,43155,43136,43136,43115,43115,43078,43078,43040,43040,42989,42989,42965,42965,42944,42944,42928,42928,42906,42906,42880,42880,42861,42861,42843,42843,42827,42827,42803,42803,42773,42773,42749,42749,42733,42733,42710,42710,42691,42691,42669,42669,42619,42619,42579,42579,42560,42560,42534,42534,42513,42513,42495,42495,42465,42465,42440,42440,42412,42412,42378,42378,42361,42361,42342,42342,42323,42323,42303,42303,42286,42286,42270,42270,42247,42247,42229,42229,42183,42183,42163,42163,42131,42131,42109,42109,42087,42087,42066,42066,42048,42048,42031,42031,42014,42014,41997,41997,41977,41977,41961,41961,41945,41945,41923,41923,41891,41891,41870,41870,41854,41854,41834,41834,41815,41815,41797,41797,41781,41781,41703,41703,41683,41683,41651,41651,41630,41630,41611,41611,41592,41592,41540,41540,41498,41498,41465,41465,41445,41445,41408,41408,41389,41389,41364,41364,41342,41342,41289,41289,41265,41265,41248,41248,41227,41227,41195,41195,41154,41154,41138,41138,41113,41113,41086,41086,41068,41068,41011,41011,40973,40973,40938,40938,40910,40910,40886,40886,40863,40863,40804,40804,40788,40788,40767,40767,40734,40734,40718,40718,40688,40688,40666,40666,40628,40628,40595,40595,40554,40554,40535,40535,40519,40519,40498,40498,40478,40478,40455,40455,40434,40434,40405,40405,40379,40379,40348,40348,40308,40308,40291,40291,40272,40272,40250,40250,40229,40229,40212,40212,40175,40175,40157,40157,40135,40135,40115,40115,40085,40085,40065,40065,40039,40039,40021,40021,40005,40005,39987,39987,39962,39962,39945,39945,39922,39922,39880,39880,39862,39862,39835,39835,39819,39819,39798,39798,39773,39773,39682,39682,39658,39658,39639,39639,39618,39618,39599,39599,39573,39573,39557,39557,39533,39533,39503,39503,39482,39482,39461,39461,39444,39444,39422,39422,39390,39390,39347,39347,39331,39331,39303,39303,39250,39250,39223,39223,39191,39191,39171,39171,39151,39151,39097,39097,39078,39078,39055,39055,39027,39027,38990,38990,38954,38954,38931,38931,38911,38911,38873,38873,38845,38845,38829,38829,38781,38781,38755,38755,38731,38731,38709,38709,38686,38686,38665,38665,38621,38621,38605,38605,38544,38544,38517,38517,38484,38484,38453,38453,38436,38436,38420,38420,38400,38400,38383,38383,38361,38361,38344,38344,38326,38326,38309,38309,38276,38276,38253,38253,38236,38236,38218,38218,38192,38192,38171,38171,38152,38152,38134,38134,38116,38116,38098,38098,38065,38065,38041,38041,38024,38024,38007,38007,37985,37985,37958,37958,37936,37936,37909,37909,37886,37886,37867,37867,37851,37851,37830,37830,37808,37808,37788,37788,37763,37763,37741,37741,37724,37724,37704,37704,37671,37671,37648,37648,37625,37625,37608,37608,37569,37569,37543,37543,37516,37516,37494,37494,37467,37467,37445,37445,37426,37426,37388,37388,37370,37370,37336,37336,37320,37320,37302,37302,37285,37285,37255,37255,37221,37221,37182,37182,37147,37147,37127,37127,37107,37107,37077,37077,37057,37057,37039,37039,37013,37013,36993,36993,36962,36962,36925,36925,36907,36907,36889,36889,36870,36870,36854,36854,36829,36829,36801,36801,36780,36780,36737,36737,36719,36719,36703,36703,36680,36680,36659,36659,36621,36621,36601,36601,36541,36541,36513,36513,36490,36490,36471,36471,36453,36453,36429,36429,36409,36409,36383,36383,36367,36367,36346,36346,36323,36323,36305,36305,36286,36286,36268,36268,36251,36251,36222,36222,36194,36194,36158,36158,36136,36136,36116,36116,36094,36094,36063,36063,36045,36045,36015,36015,35970,35970,35954,35954,35928,35928,35907,35907,35891,35891,35870,35870,35853,35853,35831,35831,35810,35810,35794,35794,35776,35776,35723,35723,35704,35704,35685,35685,35667,35667,35637,35637,35621,35621,35605,35605,35576,35576,35546,35546,35525,35525,35500,35500,35484,35484,35419,35419,35400,35400,35384,35384,35342,35342,35292,35292,35274,35274,35254,35254,35231,35231,35198,35198,35182,35182,35166,35166,35148,35148,35131,35131,35115,35115,35090,35090,35041,35041,35017,35017,34994,34994,34968,34968,34950,34950,34915,34915,34895,34895,34877,34877,34856,34856,34837,34837,34817,34817,34793,34793,34776,34776,34758,34758,34742,34742,34725,34725,34706,34706,34690,34690,34671,34671,34654,34654,34637,34637,34615,34615,34586,34586,34566,34566,34548,34548,34518,34518,34474,34474,34411,34411,34394,34394,34376,34376,34349,34349,34322,34322,34303,34303,34285,34285,34269,34269,34250,34250,34231,34231,34205,34205,34189,34189,34167,34167,34129,34129,34105,34105,34085,34085,34064,34064,34048,34048,34031,34031,34000,34000,33980,33980,33961,33961,33942,33942,33896,33896,33878,33878,33833,33833,33800,33800,33775,33775,33758,33758,33716,33716,33699,33699,33681,33681,33642,33642,33620,33620,33597,33597,33557,33557,33500,33500,33482,33482,33422,33422,33405,33405,33388,33388,33371,33371,33355,33355,33338,33338,33316,33316,33300,33300,33281,33281,33258,33258,33242,33242,33216,33216,33197,33197,33179,33179,33156,33156,33137,33137,33094,33094,33078,33078,33062,33062,33028,33028,33011,33011,32995,32995,32972,32972,32948,32948,32913,32913,32863,32863,32830,32830,32811,32811,32767,32767,32751,32751,32715,32715,32699,32699,32671,32671,32654,32654,32630,32630,32607,32607,32590,32590,32549,32549,32517,32517,32500,32500,32484,32484,32464,32464,32441,32441,32425,32425,32402,32402,32366,32366,32350,32350,32313,32313,32295,32295,32278,32278,32247,32247,32231,32231,32200,32200,32175,32175,32158,32158,32142,32142,32125,32125,32105,32105,32089,32089,32066,32066,32047,32047,32029,32029,31999,31999,31980,31980,31957,31957,31939,31939,31923,31923,31905,31905,31885,31885,31863,31863,31842,31842,31820,31820,31788,31788,31755,31755,31728,31728,31712,31712,31692,31692,31662,31662,31645,31645,31623,31623,31607,31607,31591,31591,31571,31571,31551,31551,31520,31520,31503,31503,31483,31483,31460,31460,31444,31444,31425,31425,31407,31407,31389,31389,31370,31370,31344,31344,31323,31323,31300,31300,31284,31284,31260,31260,31244,31244,31216,31216,31177,31177,31159,31159,31140,31140,31118,31118,31094,31094,31070,31070,31050,31050,31034,31034,31015,31015,30982,30982,30965,30965,30938,30938,30905,30905,30884,30884,30864,30864,30831,30831,30806,30806,30787,30787,30769,30769,30730,30730,30707,30707,30689,30689,30672,30672,30649,30649,30632,30632,30616,30616,30600,30600,30584,30584,30568,30568,30546,30546,30522,30522,30502,30502,30483,30483,30460,30460,30439,30439,30422,30422,30391,30391,30368,30368,30348,30348,30311,30311,30292,30292,30263,30263,30239,30239,30211,30211,30184,30184,30159,30159,30118,30118,30097,30097,30059,30059,30039,30039,30023,30023,29977,29977,29952,29952,29933,29933,29916,29916,29898,29898,29872,29872,29848,29848,29827,29827,29802,29802,29779,29779,29757,29757,29730,29730,29705,29705,29688,29688,29664,29664,29635,29635,29618,29618,29601,29601,29580,29580,29564,29564,29547,29547,29522,29522,29475,29475,29449,29449,29432,29432,29411,29411,29394,29394,29370,29370,29341,29341,29317,29317,29298,29298,29282,29282,29266,29266,29250,29250,29226,29226,29205,29205,29188,29188,29168,29168,29151,29151,29107,29107,29086,29086,29069,29069,29048,29048,29027,29027,28983,28983,28964,28964,28948,28948,28931,28931,28914,28914,28897,28897,28869,28869,28847,28847,28829,28829,28806,28806,28784,28784,28765,28765,28743,28743,28723,28723,28686,28686,28666,28666,28640,28640,28621,28621,28605,28605,28589,28589,28572,28572,28549,28549,28531,28531,28510,28510,28492,28492,28471,28471,28451,28451,28431,28431,28415,28415,28399,28399,28371,28371,28355,28355,28339,28339,28322,28322,28303,28303,28286,28286,28266,28266,28249,28249,28226,28226,28201,28201,28181,28181,28165,28165,28146,28146,28130,28130,28113,28113,28096,28096,28080,28080,28060,28060,28044,28044,28027,28027,28004,28004,27987,27987,27971,27971,27951,27951,27933,27933,27914,27914,27894,27894,27877,27877,27851,27851,27828,27828,27812,27812,27795,27795,27779,27779,27759,27759,27739,27739,27717,27717,27701,27701,27684,27684,27662,27662,27634,27634,27616,27616,27592,27592,27570,27570,27550,27550,27525,27525,27505,27505,27483,27483,27460,27460,27441,27441,27424,27424,27403,27403,27387,27387,27371,27371,27355,27355,27338,27338,27322,27322,27305,27305,27284,27284,27267,27267,27251,27251,27229,27229,27206,27206,27189,27189,27172,27172,27156,27156,27140,27140,27118,27118,27096,27096,27077,27077,27061,27061,27045,27045,27028,27028,27011,27011,26993,26993,26964,26964,26947,26947,26931,26931,26915,26915,26891,26891,26875,26875,26844,26844,26825,26825,26807,26807,26790,26790,26761,26761,26744,26744,26717,26717,26698,26698,26679,26679,26663,26663,26646,26646,26629,26629,26610,26610,26594,26594,26576,26576,26541,26541,26518,26518,26499,26499,26480,26480,26460,26460,26444,26444,26427,26427,26410,26410,26391,26391,26375,26375,26354,26354,26334,26334,26313,26313,26291,26291,26267,26267,26247,26247,26228,26228,26201,26201,26183,26183,26150,26150,26129,26129,26113,26113,26088,26088,26064,26064,26040,26040,26022,26022,26004,26004,25980,25980,25964,25964,25946,25946,25930,25930,25911,25911,25893,25893,25861,25861,25840,25840,25821,25821,25805,25805,25786,25786,25770,25770,25753,25753,25727,25727,25711,25711,25695,25695,25674,25674,25658,25658,25641,25641,25621,25621,25600,25600,25584,25584,25565,25565,25547,25547,25523,25523,25507,25507,25491,25491,25474,25474,25456,25456,25438,25438,25421,25421,25402,25402,25386,25386,25370,25370,25349,25349,25333,25333,25316,25316,25296,25296,25274,25274,25258,25258,25240,25240,25224,25224,25200,25200,25181,25181,25162,25162,25140,25140,25108,25108,25090,25090,25073,25073,25056,25056,25031,25031,25014,25014,24996,24996,24980,24980,24961,24961,24940,24940,24921,24921,24896,24896,24878,24878,24861,24861,24845,24845,24828,24828,24811,24811,24794,24794,24778,24778,24755,24755,24739,24739,24716,24716,24696,24696,24669,24669,24651,24651,24634,24634,24617,24617,24595,24595,24571,24571,24555,24555,24539,24539,24513,24513,24492,24492,24475,24475,24455,24455,24437,24437,24419,24419,24403,24403,24385,24385,24366,24366,24347,24347,24317,24317,24301,24301,24284,24284,24262,24262,24246,24246,24227,24227,24204,24204,24188,24188,24172,24172,24150,24150,24129,24129,24108,24108,24091,24091,24074,24074,24058,24058,24041,24041,24025,24025,24005,24005,23989,23989,23962,23962,23945,23945,23922,23922,23906,23906,23889,23889,23872,23872,23851,23851,23823,23823,23798,23798,23777,23777,23761,23761,23739,23739,23716,23716,23697,23697,23671,23671,23651,23651,23634,23634,23616,23616,23598,23598,23581,23581,23557,23557,23535,23535,23516,23516,23496,23496,23478,23478,23458,23458,23433,23433,23417,23417,23400,23400,23384,23384,23368,23368,23350,23350,23334,23334,23315,23315,23297,23297,23281,23281,23261,23261,23244,23244,23215,23215,23198,23198,23179,23179,23163,23163,23142,23142,23124,23124,23108,23108,23090,23090,23066,23066,23047,23047,23028,23028,23012,23012,22995,22995,22978,22978,22955,22955,22938,22938,22910,22910,22892,22892,22872,22872,22852,22852,22831,22831,22809,22809,22775,22775,22756,22756,22740,22740,22719,22719,22702,22702,22685,22685,22669,22669,22652,22652,22636,22636,22620,22620,22600,22600,22582,22582,22564,22564,22543,22543,22526,22526,22510,22510,22494,22494,22477,22477,22461,22461,22433,22433,22416,22416,22400,22400,22383,22383,22365,22365,22348,22348,22330,22330,22313,22313,22292,22292,22269,22269,22253,22253,22236,22236,22219,22219,22201,22201,22184,22184,22164,22164,22147,22147,22130,22130,22112,22112,22096,22096,22079,22079,22063,22063,22047,22047,22030,22030,22014,22014,21998,21998,21982,21982,21964,21964,21947,21947,21928,21928,21912,21912,21890,21890,21873,21873,21855,21855,21839,21839,21819,21819,21802,21802,21780,21780,21756,21756,21740,21740,21721,21721,21704,21704,21685,21685,21658,21658,21633,21633,21615,21615,21598,21598,21582,21582,21564,21564,21547,21547,21530,21530,21514,21514,21496,21496,21478,21478,21453,21453,21436,21436,21420,21420,21400,21400,21382,21382,21364,21364,21340,21340,21323,21323,21305,21305,21281,21281,21265,21265,21249,21249,21232,21232,21215,21215,21199,21199,21180,21180,21164,21164,21140,21140,21120,21120,21104,21104,21082,21082,21065,21065,21046,21046,21009,21009,20993,20993,20975,20975,20959,20959,20942,20942,20926,20926,20908,20908,20891,20891,20875,20875,20859,20859,20843,20843,20826,20826,20810,20810,20793,20793,20776,20776,20754,20754,20734,20734,20718,20718,20702,20702,20684,20684,20664,20664,20648,20648,20632,20632,20614,20614,20596,20596,20580,20580,20561,20561,20545,20545,20528,20528,20506,20506,20485,20485,20465,20465,20445,20445,20428,20428,20402,20402,20385,20385,20368,20368,20348,20348,20331,20331,20312,20312,20294,20294,20273,20273,20257,20257,20240,20240,20221,20221,20203,20203,20187,20187,20171,20171,20151,20151,20134,20134,20116,20116,20099,20099,20080,20080,20064,20064,20045,20045,20029,20029,20011,20011,19993,19993,19976,19976,19959,19959,19943,19943,19923,19923,19906,19906,19890,19890,19874,19874,19854,19854,19833,19833,19817,19817,19794,19794,19777,19777,19761,19761,19745,19745,19723,19723,19705,19705,19689,19689,19666,19666,19649,19649,19632,19632,19612,19612,19596,19596,19580,19580,19562,19562,19541,19541,19525,19525,19504,19504,19486,19486,19461,19461,19443,19443,19426,19426,19410,19410,19394,19394,19375,19375,19358,19358,19334,19334,19317,19317,19300,19300,19278,19278,19262,19262,19245,19245,19228,19228,19212,19212,19191,19191,19175,19175,19155,19155,19136,19136,19119,19119,19103,19103,19082,19082,19064,19064,19046,19046,19027,19027,19011,19011,18995,18995,18978,18978,18962,18962,18946,18946,18924,18924,18902,18902,18885,18885,18866,18866,18850,18850,18834,18834,18818,18818,18802,18802,18783,18783,18766,18766,18750,18750,18734,18734,18716,18716,18700,18700,18682,18682,18666,18666,18649,18649,18632,18632,18608,18608,18589,18589,18573,18573,18557,18557,18538,18538,18522,18522,18504,18504,18487,18487,18471,18471,18454,18454,18436,18436,18415,18415,18398,18398,18380,18380,18364,18364,18348,18348,18332,18332,18314,18314,18297,18297,18281,18281,18264,18264,18248,18248,18231,18231,18214,18214,18198,18198,18182,18182,18166,18166,18150,18150,18134,18134,18112,18112,18095,18095,18077,18077,18061,18061,18043,18043,18026,18026,18004,18004,17985,17985,17968,17968,17951,17951,17934,17934,17918,17918,17900,17900,17884,17884,17857,17857,17840,17840,17824,17824,17806,17806,17789,17789,17772,17772,17756,17756,17739,17739,17723,17723,17705,17705,17689,17689,17673,17673,17657,17657,17640,17640,17624,17624,17605,17605,17588,17588,17568,17568,17547,17547,17531,17531,17515,17515,17499,17499,17483,17483,17467,17467,17450,17450,17434,17434,17413,17413,17397,17397,17381,17381,17365,17365,17349,17349,17332,17332,17315,17315,17298,17298,17280,17280,17261,17261,17243,17243,17227,17227,17207,17207,17190,17190,17169,17169,17153,17153,17136,17136,17119,17119,17098,17098,17082,17082,17064,17064,17042,17042,17026,17026,16947,16947,16927,16927,16829,16829,16094,16094,14701,14701,12916,12916,11586,11586,10948,10948,8779,8779,8350,8350,7853,7853,7581,7581,7463,7463,7150,7150,6888,6888,6566,6566,5885,5885,5461,5461,4958,4958,4146,4146,7,7,0.0]],"coord\_x":[[0.0,1e-10,0.3477536979352335,0.3477536980352335,0.6715698975307698,0.6715698976307698,0.9786985338784042,0.9786985339784042,1.2696479189924388,1.2696479190924388,1.556724091598839,1.556724091698839,1.8363213897775947,1.8363213898775947,2.103229378561383,2.103229378661383,2.3696767604986673,2.3696767605986673,2.6242272541483334,2.6242272542483334,2.873125651642551,2.873125651742551,3.1029571385443964,3.1029571386443964,3.319958170896353,3.319958170996353,3.5280813056233447,3.5280813057233447,3.7358863427461575,3.7358863428461575,3.937888855185994,3.937888855285994,4.136424084794619,4.136424084894619,4.331704677641676,4.331704677741676,4.522379997741344,4.522379997841344,4.711658568122697,4.711658568222697,4.8984778025899045,4.8984778026899045,5.083945150657838,5.083945150757838,5.263458674014983,5.263458674114983,5.441092068125938,5.441092068225938,5.6158062394112624,5.6158062395112625,5.79007723385579,5.79007723395579,5.9635221975950055,5.9635221976950055,6.133540535885889,6.133540535985889,6.302548805345969,6.302548805445969,6.466972414934975,6.466972415034975,6.630771765080402,6.630771765180402,6.794561036135571,6.794561036235571,6.956843634762512,6.956843634862512,7.1185876462130615,7.1185876463130615,7.2798030359035275,7.2798030360035275,7.440579568363308,7.440579568463308,7.600177984002465,7.600177984102465,7.759213258891972,7.759213258991972,7.91799469985052,7.91799469995052,8.074563515258331,8.074563515358332,8.23072893970794,8.23072893980794,8.386841846792526,8.386841846892526,8.541942638915202,8.541942639015202,8.695758006008196,8.695758006108196,8.849329883499706,8.849329883599706,9.001430516944156,9.001430517044156,9.152686136133598,9.152686136233598,9.302557471847937,9.302557471947937,9.451536656509223,9.451536656609223,9.599973085949273,9.599973086049273,9.747829247764498,9.747829247864498,9.89533218672491,9.895332186824911,10.040762508550005,10.040762508650005,10.185904250106674,10.185904250206674,10.33054737982613,10.33054737992613,10.475112529215702,10.475112529315702,10.619334686405983,10.619334686505983,10.763449345930624,10.763449346030624,10.907401944293493,10.907401944393493,11.050621724590272,11.050621724690272,11.193674783093318,11.193674783193318,11.336524441008008,11.336524441108008,11.479220335959297,11.479220336059297,11.621286022530278,11.621286022630278,11.762580014544172,11.762580014644172,11.903113376265479,11.903113376365479,12.04363370337382,12.04363370347382,12.183729079364726,12.183729079464726,12.323723323431206,12.323723323531206,12.46354175178793,12.46354175188793,12.603245824150681,12.603245824250681,12.74205478993767,12.74205479003767,12.8799349637683,12.8799349638683,13.016581168977405,13.016581169077405,13.152925077261418,13.152925077361418,13.28779880245524,13.28779880255524,13.42256658352741,13.42256658362741,13.556017982364105,13.556017982464105,13.689318043281672,13.689318043381672,13.822477375848802,13.822477375948802,13.955281590995279,13.955281591095279,14.08797989991142,14.08797990001142,14.21966238051661,14.21966238061661,14.351266198748212,14.351266198848212,14.481366603204824,14.481366603304824,14.611416119623032,14.611416119723032,14.741404934151795,14.741404934251795,14.87094685849072,14.87094685859072,15.000389848601591,15.000389848701591,15.129700484343028,15.129700484443028,15.258807454256894,15.258807454356894,15.387902829427832,15.387902829527832,15.516914237440835,15.516914237540835,15.643907402358067,15.643907402458067,15.770813417246757,15.770813417346757,15.897286978537293,15.897286978637293,16.02316852589481,16.02316852599481,16.14902820996256,16.149028210062557,16.274729849348113,16.27472984944811,16.400231498363816,16.400231498463814,16.525690254408953,16.52569025450895,16.65099930186158,16.65099930196158,16.775157931046138,16.775157931146136,16.8992363064218,16.899236306521797,17.02318626812497,17.02318626822497,17.1470083845254,17.147008384625398,17.269910310298375,17.269910310398373,17.39240429815514,17.392404298255137,17.514137011566927,17.514137011666925,17.635790001648257,17.635790001748255,17.75709029935321,17.757090299453207,17.87806329186401,17.878063291964008,17.99895076367289,17.99895076377289,18.117636408956308,18.117636409056306,18.236314475976375,18.236314476076373,18.35494908165612,18.354949081756118,18.473503774548824,18.473503774648822,18.591982684808013,18.59198268490801,18.71044420295281,18.71044420305281,18.828844185599195,18.828844185699193,18.945492793693727,18.945492793793726,19.06205205406335,19.06205205416335,19.178509689921427,19.178509690021425,19.29315301375052,19.293153013850517,19.407385671488598,19.407385671588596,19.521520001259688,19.521520001359686,19.635565930588786,19.635565930688784,19.749445176015467,19.749445176115465,19.8631531526904,19.8631531527904,19.976715854056888,19.976715854156886,20.090170792518517,20.090170792618515,20.203589582663962,20.20358958276396,20.31685991463035,20.316859914730347,20.429780737090965,20.429780737190963,20.542697429398054,20.54269742949805,20.65553178387383,20.655531783973828,20.766979013025743,20.76697901312574,20.877777163921607,20.877777164021605,20.98836729148847,20.988367291588467,21.0986523181728,21.098652318272798,21.208355220557785,21.208355220657783,21.317773445479972,21.31777344557997,21.42699740162114,21.426997401721138,21.535871014647572,21.53587101474757,21.64447772123876,21.64447772133876,21.752519885101584,21.752519885201583,21.86020837141379,21.86020837151379,21.967878859350538,21.967878859450536,22.07548209020004,22.07548209030004,22.18290268490277,22.18290268500277,22.29002534018184,22.290025340281836,22.396207532978984,22.396207533078982,22.50213661178019,22.50213661188019,22.608058112318044,22.608058112418043,22.713935999950312,22.71393600005031,22.81970529106875,22.81970529116875,22.92507786010074,22.925077860200737,23.03011861487188,23.03011861497188,23.135119394303846,23.135119394403844,23.239722655931708,23.239722656031706,23.344156694938928,23.344156695038926,23.448589710880597,23.448589710980595,23.552632787281517,23.552632787381516,23.65655506616461,23.65655506626461,23.759931255390597,23.759931255490596,23.863117760684897,23.863117760784895,23.966047249177628,23.966047249277626,24.06880846233264,24.06880846243264,24.171352444568683,24.17135244466868,24.273266862576797,24.273266862676795,24.374652734607462,24.37465273470746,24.47575745306779,24.475757453167788,24.5768426953913,24.576842695491298,24.677571835119924,24.677571835219922,24.778054719180943,24.77805471928094,24.878472657525037,24.878472657625036,24.978653623574132,24.97865362367413,25.078794803740628,25.078794803840626,25.178231735893874,25.178231735993872,25.277667948112096,25.277667948212095,25.376851879943352,25.37685188004335,25.475512684255452,25.47551268435545,25.57377172693597,25.573771727035968,25.671676069000323,25.67167606910032,25.769564799842172,25.76956479994217,25.86720932114752,25.86720932124752,25.96471436451588,25.96471436461588,26.061740082727262,26.06174008282726,26.15816351845878,26.15816351855878,26.254563234226012,26.25456323432601,26.350460738480937,26.350460738580935,26.446070837098258,26.446070837198256,26.541611784062496,26.541611784162495,26.636780903535392,26.63678090363539,26.731842828473184,26.731842828573182,26.82660059981136,26.82660059991136,26.921339311817206,26.921339311917205,27.01586541564473,27.015865415744727,27.11020714032491,27.110207140424908,27.203979131166324,27.203979131266323,27.29764404114658,27.297644041246578,27.390904423429134,27.390904423529133,27.48393696522403,27.483936965324027,27.576876180705746,27.576876180805744,27.669709717305025,27.669709717405023,27.762307721479342,27.76230772157934,27.854891630083827,27.854891630183825,27.947380469374565,27.947380469474563,28.03911462730944,28.03911462740944,28.13066243574851,28.130662435848507,28.222161250715008,28.222161250815006,28.31359083824579,28.31359083834579,28.404618057883923,28.40461805798392,28.49544532504353,28.49544532514353,28.58585268063083,28.585852680730827,28.67615155337825,28.67615155347825,28.766328416085713,28.76632841618571,28.85621950248219,28.85621950258219,28.94573114523266,28.94573114533266,29.035093496195103,29.0350934962951,29.124155672146195,29.124155672246193,29.21314566513886,29.21314566523886,29.30204684088505,29.30204684098505,29.39065739023171,29.390657390331707,29.479015393952178,29.479015394052176,29.567234639670684,29.567234639770682,29.655429597055143,29.65542959715514,29.743238214573946,29.743238214673944,29.83102110388867,29.831021103988668,29.918536366833138,29.918536366933136,30.005949815809476,30.005949815909474,30.093144177192322,30.09314417729232,30.180167989758445,30.180167989858443,30.26717759308078,30.26717759318078,30.35414240886671,30.35414240896671,30.44108399727547,30.44108399737547,30.52798712599772,30.527987126097717,30.614706481833693,30.61470648193369,30.701360096235092,30.70136009633509,30.787990407476688,30.787990407576686,30.874610525954076,30.874610526054074,30.96122003486277,30.961220034962768,31.047683434854047,31.047683434954045,31.133806646603478,31.133806646703476,31.21987245300802,31.219872453108017,31.305890402679495,31.305890402779493,31.39186174603136,31.391861746131358,31.477694141924676,31.477694142024674,31.563517140771435,31.563517140871433,31.649007340183115,31.649007340283113,31.73434411765324,31.73434411775324,31.81956388673722,31.81956388683722,31.904439830012944,31.904439830112942,31.989000706989827,31.989000707089826,32.07340759365541,32.07340759375541,32.15756019169423,32.15756019179423,32.24163909112196,32.24163909122196,32.325397657357826,32.32539765745783,32.408856124364966,32.40885612446497,32.49231246945837,32.49231246955837,32.575256675514474,32.575256675614476,32.65814230159488,32.65814230169488,32.740551671714954,32.740551671814956,32.82285566608313,32.82285566618313,32.904858159243865,32.90485815934387,32.98650598967975,32.98650598977975,33.06806166843328,33.06806166853328,33.14939462202691,33.14939462212691,33.230652247682826,33.23065224778283,33.31190403807596,33.31190403817596,33.39299793535216,33.392997935452165,33.473864598401775,33.47386459850178,33.5544793978689,33.554479397968905,33.63494547391775,33.634945474017755,33.715392301177694,33.715392301277696,33.79580756497077,33.795807565070774,33.876080887891284,33.876080887991286,33.95629472144448,33.95629472154448,34.036392077089964,34.036392077189966,34.11646018063891,34.11646018073891,34.19622507787116,34.196225077971164,34.275960003071866,34.27596000317187,34.35544609599542,34.35544609609542,34.43430129849496,34.43430129859496,34.51303642341169,34.51303642351169,34.59154541295002,34.59154541305002,34.67005330364016,34.67005330374016,34.74854137717164,34.74854137727164,34.82695779822313,34.82695779832313,34.905288850137964,34.905288850237966,34.98360978507122,34.983609785171225,35.06173531448396,35.06173531458396,35.13979733804982,35.13979733814982,35.217857163919305,35.21785716401931,35.29589364873766,35.29589364883766,35.373871932493486,35.37387193259349,35.45173343521106,35.45173343531106,35.529592967580164,35.529592967680166,35.60734761678448,35.60734761688448,35.684929784714924,35.684929784814926,35.762439125534556,35.76243912563456,35.839852828670686,35.83985282877069,35.91717960912618,35.91717960922618,35.99432496595704,35.99432496605704,36.071121040630025,36.07112104073003,36.147840650625795,36.1478406507258,36.22437269860362,36.22437269870362,36.30077216486411,36.30077216496411,36.37716640212288,36.37716640222288,36.453550029812966,36.45355002991297,36.52942155635707,36.52942155645707,36.60524211908014,36.60524211918014,36.68087405882838,36.68087405892838,36.75646193097524,36.75646193107524,36.831915213164976,36.83191521326498,36.90726505205996,36.907265052159964,36.98261231434541,36.98261231444541,37.057884968628166,37.05788496872817,37.1327471841678,37.1327471842678,37.20725901870138,37.20725901880138,37.28173474281009,37.28173474291009,37.356085463464815,37.35608546356482,37.43022198450591,37.430221984605915,37.504148663434805,37.50414866353481,37.57780294168753,37.57780294178753,37.65136071075647,37.65136071085647,37.72487873183413,37.72487873193413,37.79837576112231,37.79837576122231,37.87149281628605,37.87149281638605,37.94458694720314,37.944586947303144,38.017435390822385,38.01743539092239,38.09026375204375,38.09026375214375,38.162951195458085,38.16295119555809,38.23556118902097,38.23556118912097,38.308002338876385,38.30800233897639,38.38044034375251,38.38044034385251,38.45284682305309,38.452846823153095,38.52508131366692,38.52508131376692,38.59728988662007,38.59728988672007,38.66943707564008,38.669437075740085,38.74135756091193,38.74135756101193,38.81283865847466,38.81283865857466,38.884222185896746,38.88422218599675,38.955258401509425,38.95525840160943,39.02622065327179,39.026220653371794,39.09681698858824,39.09681698868824,39.16733795807565,39.16733795817565,39.237631731227786,39.23763173132779,39.30780489631868,39.30780489641868,39.377713163214125,39.37771316331413,39.44754996708616,39.44754996718616,39.517377752824814,39.517377752924816,39.58719436062502,39.58719436072502,39.65699774435568,39.65699774445568,39.72678184140611,39.72678184150611,39.795890222604804,39.795890222704806,39.864756970876535,39.864756970976536,39.93357502880623,39.93357502890623,40.002320221733804,40.002320221833806,40.07093844085892,40.07093844095892,40.139401608715865,40.13940160881587,40.20777107134647,40.20777107144647,40.276085288437244,40.276085288537246,40.34423748225756,40.34423748235756,40.41211238742185,40.41211238752185,40.479759641555056,40.47975964165506,40.547275678058334,40.547275678158336,40.614524997582954,40.614524997682956,40.68174737638136,40.68174737648136,40.74874343034477,40.74874343044477,40.815649985018,40.815649985118,40.88242255599518,40.88242255609518,40.9491789473801,40.9491789474801,41.01568366463912,41.01568366473912,41.0821819782656,41.082181978365604,41.14856934611662,41.148569346216625,41.21488547829214,41.21488547839214,41.281084564190195,41.2810845642902,41.347262279385596,41.3472622794856,41.41339569963171,41.41339569973171,41.47948812147309,41.47948812157309,41.545456449252086,41.54545644935209,41.611404997763735,41.61140499786374,41.67734638481652,41.677346384916525,41.74321528578035,41.74321528588035,41.80897240735975,41.80897240745975,41.87458050239033,41.87458050249033,41.93999167624773,41.93999167634773,42.0052822799352,42.0052822800352,42.070567162033846,42.07056716213385,42.135810969945126,42.13581097004513,42.26626668127897,42.26626668137897,42.33137279214515,42.331372792245155,42.396472347813535,42.39647234791354,42.4613966182506,42.461396618350605,42.52631766792574,42.52631766802574,42.59115456098636,42.59115456108636,42.655907714236946,42.65590771433695,42.72053177177134,42.72053177187134,42.78514544708494,42.785145447184945,42.84963332322691,42.84963332332691,42.91404738708383,42.914047387183835,42.97842757610358,42.97842757620358,43.04239346146589,43.042393461565894,43.10633771088634,43.106337710986345,43.17022493387507,43.17022493397507,43.23409616672812,43.234096166828124,43.297946635139596,43.2979466352396,43.361745381903695,43.3617453820037,43.42547906927692,43.42547906937692,43.48912829690509,43.489128297005095,43.552695035136686,43.55269503523669,43.61606920969652,43.61606920979652,43.6794310695784,43.6794310696784,43.74273362954956,43.742733629649564,43.80585756985352,43.805857569953524,43.86892539311737,43.868925393217374,43.9319309988391,43.9319309989391,43.994933004885745,43.99493300498575,44.05790909327172,44.05790909337172,44.12078473177697,44.120784731876974,44.1836505943225,44.1836505944225,44.246481369508714,44.246481369608716,44.309244243455296,44.3092442435553,44.37193065272466,44.37193065282466,44.43444276193694,44.434442762036944,44.496642154112024,44.496642154212026,44.558779215071034,44.558779215171036,44.62085182290025,44.62085182300025,44.68288805506537,44.68288805516537,44.74485831844802,44.74485831854802,44.80656807902796,44.80656807912796,44.86812839625461,44.868128396354614,44.92954252878121,44.929542528881214,44.99049916154928,44.99049916164928,45.0513521236747,45.0513521237747,45.11188717765253,45.11188717775253,45.172405369994394,45.172405370094395,45.23292000055248,45.232920000652484,45.29341147951604,45.29341147961604,45.35372937835753,45.35372937845753,45.41362823712699,45.413628237226995,45.473515008566416,45.47351500866642,45.53333266624407,45.53333266634407,45.59296761199232,45.59296761209232,45.65247573550339,45.65247573560339,45.71165488660236,45.71165488670236,45.77052738327482,45.77052738337482,45.88822702071357,45.888227020813574,45.9470277891234,45.947027789223405,46.00571583086589,46.00571583096589,46.064396824823454,46.064396824923456,46.123027574895,46.123027574995,46.181578109048964,46.181578109148965,46.24008972882062,46.24008972892062,46.29860074233121,46.29860074243121,46.35705662397592,46.35705662407592,46.415470673606926,46.41547067370693,46.473741721408494,46.473741721508496,46.531837294522155,46.53183729462216,46.589926918699085,46.58992691879909,46.64779972876153,46.64779972886153,46.70566234605976,46.70566234615976,46.76321853627938,46.763218536379384,46.82073274292003,46.82073274302003,46.878141839048,46.878141839148,46.935549722653825,46.93554972275383,46.99285109376825,46.99285109386825,47.05005239391512,47.05005239401512,47.10714221780808,47.107142217908084,47.164157054785186,47.16415705488519,47.22111592628744,47.22111592638744,47.27802910086168,47.27802910096168,47.33488130830726,47.33488130840726,47.39168043001806,47.391680430118065,47.44835428303567,47.44835428313567,47.50502358909526,47.50502358919526,47.561533789515785,47.56153378961579,47.61789083323398,47.61789083333398,47.674175618211116,47.67417561831112,47.7304589254269,47.730458925526904,47.786597753051886,47.78659775315189,47.84271282282127,47.84271282292127,47.89880428630031,47.89880428640031,47.95489396888746,47.95489396898746,48.010946480092876,48.01094648019288,48.06687459410537,48.06687459420537,48.12266323018089,48.12266323028089,48.17841912815872,48.178419128258724,48.23416498493762,48.23416498503762,48.28988609868667,48.289886098786674,48.34559997519422,48.34559997529422,48.40119688120694,48.401196881306944,48.4567748794526,48.456774879552604,48.51234787604445,48.51234787614445,48.567874266316686,48.56787426641669,48.62329073387901,48.62329073397901,48.678346892910284,48.678346893010286,48.73326058059055,48.733260580690555,48.78800273427986,48.78800273437986,48.842498177605755,48.84249817770576,48.896818070461116,48.89681807056112,48.95106733390204,48.95106733400204,49.00527381804634,49.00527381814634,49.05913522596894,49.05913522606894,49.11299580028256,49.11299580038256,49.1667783942663,49.166778394366304,49.22048971468322,49.22048971478322,49.27417962650617,49.27417962660617,49.3278405514718,49.3278405515718,49.38148734297628,49.38148734307628,49.43507794165801,49.43507794175801,49.488666153186784,49.488666153286786,49.542210979157865,49.54221097925787,49.59569131410783,49.59569131420783,49.64914072974332,49.649140729843324,49.70255873347722,49.70255873357722,49.75586901219758,49.75586901229758,49.80917087904562,49.80917087914562,49.86237491051378,49.86237491061378,49.915396495291766,49.91539649539177,49.968120784798465,49.96812078489847,50.020793466331746,50.02079346643175,50.0733523602408,50.0733523603408,50.125855743370806,50.12585574347081,50.178155536455876,50.17815553655588,50.2303408219817,50.2303408220817,50.28246343526961,50.282463435369614,50.33456024457081,50.33456024467081,50.386464035504446,50.38646403560445,50.43830814761419,50.43830814771419,50.48995799094291,50.489957991042914,50.54145846670098,50.54145846680098,50.592910706812816,50.59291070691282,50.644346085288696,50.6443460853887,50.69576657247709,50.69576657257709,50.74716985700768,50.74716985710768,50.798471782265935,50.79847178236594,50.84974055262203,50.84974055272203,50.90100500336801,50.901005003468015,50.952238648473475,50.95223864857348,51.003465435250604,51.003465435350606,51.05467664869655,51.05467664879655,51.105853229478974,51.105853229578976,51.15701984484509,51.15701984494509,51.20817929875234,51.20817929885234,51.259283355554494,51.259283355654496,51.310332053142865,51.31033205324287,51.36135146074338,51.36135146084338,51.41233828181148,51.41233828191148,51.46327876179919,51.46327876189919,51.514149180742216,51.51414918084222,51.564976668823356,51.56497666892336,51.61571086848264,51.61571086858264,51.66637580281488,51.666375802914885,51.71702160203217,51.71702160213217,51.7676442117636,51.767644211863605,51.81821229589022,51.81821229599022,51.86871206197272,51.868712062072724,51.91919750513749,51.91919750523749,51.96965619703263,51.96965619713263,52.02006203054089,52.020062030640894,52.070145446834864,52.070145446934866,52.120168615935185,52.12016861603519,52.170141275910275,52.17014127601028,52.22002045800693,52.22002045810693,52.26989425953659,52.269894259636594,52.31976199845558,52.319761998555585,52.369573165638656,52.36957316573866,52.41931650736473,52.41931650746473,52.469050906740044,52.469050906840046,52.5187652616088,52.5187652617088,52.56829986007085,52.56829986017085,52.617611051329305,52.61761105142931,52.66690090977642,52.666900909876425,52.71609372856133,52.716093728661335,52.765224595043335,52.76522459514334,52.81432094253577,52.814320942635774,52.86337132786099,52.86337132796099,52.91241186144384,52.912411861543845,52.96144269484961,52.96144269494961,53.01044768637734,53.01044768647734,53.059382844208294,53.059382844308296,53.10830114040329,53.10830114050329,53.15719840691749,53.15719840701749,53.20608430603665,53.20608430613665,53.25490071248089,53.25490071258089,53.30369859007123,53.30369859017123,53.352437091968206,53.35243709206821,53.40113577009128,53.40113577019128,53.44981982216608,53.44981982226608,53.498501828109774,53.498501828209776,53.54709005304449,53.547090053144494,53.595616363567636,53.59561636366764,53.644093112248456,53.64409311234846,53.69256091857852,53.692560918678524,53.74102004779704,53.741020047897045,53.789474971079414,53.789474971179416,53.837892154610294,53.837892154710296,53.886308352966935,53.88630835306694,53.9346721476325,53.9346721477325,53.983015291530435,53.98301529163044,54.03126203991854,54.03126204001854,54.07946085579094,54.07946085589094,54.12764838005095,54.12764838015095,54.175825825220706,54.17582582532071,54.22399879921508,54.22399879931508,54.27214839426701,54.272148394367015,54.36842609999429,54.36842610009429,54.416527004704186,54.41652700480419,54.464366648785045,54.46436664888505,54.51218689251173,54.51218689261173,54.559967767160295,54.5599677672603,54.607745913634055,54.60774591373406,54.65549109466224,54.65549109476224,54.70315757542552,54.70315757552552,54.75081628846886,54.75081628856886,54.79845347924429,54.79845347934429,54.893650865639486,54.89365086573949,54.94109579587368,54.94109579597368,54.988538301063606,54.98853830116361,55.03596591496604,55.03596591506604,55.08330145296876,55.08330145306876,55.130541012266136,55.13054101236614,55.17760945437703,55.17760945447703,55.22464341538967,55.22464341548967,55.27167317046616,55.27167317056616,55.31866128298552,55.318661283085525,55.36558335093979,55.36558335103979,55.41248867093204,55.412488671032044,55.45935875199971,55.45935875209971,55.50619939151426,55.50619939161426,55.55273640790763,55.55273640800763,55.59921010791069,55.599210108010695,55.692022842177074,55.692022842277076,55.73834046784642,55.73834046794642,55.784647029251275,55.78464702935128,55.83091073557687,55.83091073567687,55.877139885021585,55.877139885121586,55.92332220079879,55.923322200898795,55.96947905361113,55.96947905371113,56.015531136932644,56.015531137032646,56.061525928583215,56.06152592868322,56.10732906595362,56.10732906605362,56.153062104388034,56.153062104488036,56.198793361930555,56.19879336203056,56.244347060762756,56.24434706086276,56.28989935761623,56.28989935771623,56.3354206593726,56.335420659472604,56.380883684283795,56.3808836843838,56.426282786543624,56.426282786643625,56.47165293983745,56.47165293993745,56.51697743409458,56.51697743419458,56.56229006836956,56.562290068469565,56.6075630304359,56.6075630305359,56.65265407629049,56.65265407639049,56.69773909742571,56.69773909752571,56.74281456994911,56.74281457004911,56.78783244767104,56.787832447771045,56.83283607825787,56.83283607835787,56.8778340630385,56.8778340631385,56.92282079409806,56.92282079419806,56.96773450859022,56.96773450869022,57.012647427364726,57.01264742746473,57.05753014675978,57.05753014685978,57.10238516760229,57.102385167702295,57.14720244869331,57.14720244879331,57.19195777748142,57.19195777758142,57.23670924135523,57.23670924145523,57.281458355967395,57.2814583560674,57.326176930178256,57.32617693027826,57.370886562038365,57.37088656213837,57.41547145568371,57.41547145578371,57.460016487663815,57.460016487763816,57.50445307469537,57.50445307479537,57.54887173913409,57.548871739234094,57.593152782310355,57.59315278241036,57.63738062607789,57.637380626177894,57.68158929683915,57.68158929693915,57.72566618160072,57.725666181700724,57.76972196089886,57.769721960998865,57.81377489834824,57.813774898448244,57.857798318461874,57.857798318561876,57.901784150389275,57.90178415048928,57.945686848767714,57.945686848867716,57.989498153290135,57.98949815339014,58.033299984983366,58.03329998508337,58.07706672931728,58.07706672941728,58.12078383602624,58.12078383612624,58.164389807503156,58.16438980760316,58.207953795401096,58.2079537955011,58.25142623787775,58.25142623797775,58.2948798862613,58.2948798863613,58.33826548183995,58.33826548193995,58.38160037883678,58.38160037893678,58.42483115049516,58.42483115059516,58.46804119560327,58.46804119570327,58.511217214308935,58.51121721440894,58.55437621981337,58.55437621991337,58.59749960748006,58.59749960758006,58.64053069189913,58.64053069199913,58.683502249059565,58.68350224915957,58.726452321843404,58.726452321943405,58.76917015874684,58.76917015884684,58.81186276003331,58.81186276013331,58.854484277209544,58.854484277309545,58.89710264940649,58.89710264950649,58.93968707098362,58.93968707108362,58.98226758975512,58.98226758985512,59.02466903416363,59.02466903426363,59.06692224774098,59.066922247840985,59.109149581548984,59.109149581648985,59.15129533535202,59.15129533545202,59.1934371484581,59.193437148558104,59.23552758093866,59.23552758103866,59.27754507263447,59.27754507273447,59.3195278179928,59.3195278180928,59.36149760452081,59.36149760462081,59.4034556068493,59.4034556069493,59.445373027577546,59.44537302767755,59.487174349311246,59.48717434941125,59.52896180282302,59.52896180292302,59.5706432364305,59.5706432365305,59.612287574438874,59.612287574538875,59.65390561587342,59.65390561597342,59.69543142984298,59.69543142994298,59.73685914319345,59.73685914329345,59.77827185158249,59.77827185168249,59.8195633077579,59.819563307857905,59.860811567832215,59.86081156793222,59.90200549175829,59.90200549185829,59.94314981595073,59.943149816050735,59.98428099185627,59.98428099195627,60.02531103583737,60.02531103593737,60.06624945861455,60.06624945871455,60.10710535410384,60.10710535420384,60.14778441081781,60.147784410917815,60.18845528300737,60.18845528310737,60.22909553901299,60.22909553911299,60.2695370171709,60.2695370172709,60.30983068130214,60.30983068140214,60.3500714870465,60.3500714871465,60.39030164533085,60.39030164543085,60.430486220361146,60.43048622046115,60.47061198806783,60.47061198816783,60.510699409761955,60.51069940986196,60.550776373452656,60.55077637355266,60.59079411301527,60.59079411311527,60.630726369767274,60.630726369867276,60.67065673195344,60.670656732053445,60.7104916459127,60.7104916460127,60.750314813563755,60.750314813663756,60.79013620032293,60.79013620042293,60.82995023616665,60.829950236266654,60.869721265365854,60.869721265465856,60.90947107542767,60.90947107552767,60.94921334511745,60.94921334521745,60.98887354221514,60.98887354231514,61.02852339498335,61.028523395083354,61.06810921142624,61.06810921152624,61.10755831599827,61.10755831609827,61.1469841174105,61.1469841175105,61.18631723666193,61.18631723676193,61.225554566664606,61.22555456676461,61.264738507801965,61.26473850790197,61.30385549498261,61.303855495082615,61.34293227947619,61.34293227957619,61.38199299805145,61.38199299815145,61.46003823576398,61.46003823586398,61.49900710578743,61.49900710588743,61.53778363948701,61.53778363958701,61.576479313116636,61.57647931321664,61.61512599327369,61.615125993373695,61.65364520704118,61.65364520714118,61.692142822758115,61.692142822858116,61.73059008091507,61.73059008101507,61.76901835552234,61.76901835562234,61.80744515236825,61.807445152468254,61.84571227520535,61.84571227530535,61.88390384275683,61.88390384285683,61.92209472826461,61.92209472836461,61.96013291176586,61.96013291186586,61.99808057291138,61.99808057301138,62.035938014831686,62.03593801493169,62.07364120120148,62.073641201301484,62.11132616184792,62.11132616194792,62.14898535639896,62.14898535649896,62.18663947351355,62.18663947361355,62.22417495291538,62.22417495301538,62.261690539375905,62.26169053947591,62.299099802801614,62.299099802901615,62.33643525394228,62.33643525404228,62.373691777470135,62.37369177757014,62.41089017571809,62.41089017581809,62.44805333504145,62.44805333514145,62.48519470685768,62.485194706957685,62.52228049211223,62.52228049221223,62.55934585394705,62.55934585404705,62.596238734507985,62.59623873460799,62.63310183249395,62.63310183259395,62.669935602600745,62.66993560270075,62.70667468230696,62.70667468240696,62.743149129056945,62.74314912915695,62.77958583605544,62.77958583615544,62.81599977037256,62.81599977047256,62.852392940248095,62.852392940348096,62.888759775658485,62.88875977575849,62.92512577745991,62.92512577755991,62.961486967064104,62.961486967164106,62.99781742681041,62.99781742691041,63.034121438417614,63.034121438517616,63.07026721588605,63.07026721598605,63.10629302944562,63.10629302954562,63.14227322185982,63.14227322195982,63.17824924622917,63.17824924632917,63.21405726050003,63.21405726060003,63.249711019220356,63.24971101932036,63.28536136772218,63.28536136782218,63.32101107207162,63.32101107217162,63.356607539121015,63.35660753922102,63.39215448221941,63.39215448231941,63.427696575229255,63.42769657532926,63.46322309490792,63.46322309500792,63.49871998357687,63.49871998367687,63.5341094124715,63.5341094125715,63.56946758102981,63.569467581129814,63.604800021384044,63.604800021484046,63.640129013628446,63.64012901372845,63.675455580828576,63.67545558092858,63.71077131111211,63.71077131121211,63.7459747694241,63.7459747695241,63.78112040558671,63.781120405686714,63.81621231186217,63.81621231196217,63.851217409130925,63.85121740923093,63.88612883906466,63.88612883916466,63.92103405482245,63.92103405492245,63.955929380946564,63.955929381046566,63.99072080908013,63.99072080918013,64.02547654359331,64.02547654369332,64.06021590905765,64.06021590915765,64.0949367456681,64.0949367457681,64.1296007453034,64.12960074540341,64.16420256528792,64.16420256538792,64.19877926332941,64.19877926342942,64.23325392005489,64.2332539201549,64.26769549766084,64.26769549776084,64.3020930455567,64.3020930456567,64.33645618813696,64.33645618823697,64.37079845260168,64.37079845270168,64.40508319804756,64.40508319814757,64.43931190223597,64.43931190233597,64.47352211546179,64.47352211556179,64.50771156424602,64.50771156434602,64.54189529144143,64.54189529154144,64.57606158863113,64.57606158873114,64.61022265681912,64.61022265691912,64.64429153543344,64.64429153553344,64.67829292961261,64.67829292971261,64.7122235807034,64.7122235808034,64.74611012630147,64.74611012640148,64.77999095031072,64.77999095041072,64.81368606897654,64.81368606907654,64.84730476085646,64.84730476095646,64.8809173901257,64.8809173902257,64.91451126319315,64.91451126329315,64.94808012799153,64.94808012809153,64.98162186260711,64.98162186270712,65.01509781789682,65.01509781799682,65.04856744533662,65.04856744543662,65.08201123089837,65.08201123099838,65.11528196681651,65.11528196691651,65.14852602724766,65.14852602734766,65.18175280923835,65.18175280933835,65.2148471989683,65.2148471990683,65.24785603010501,65.24785603020501,65.28080859263633,65.28080859273634,65.3136794235974,65.31367942369741,65.34652778500765,65.34652778510765,65.37931934733408,65.37931934743408,65.41208347634716,65.41208347644717,65.44484404357648,65.44484404367648,65.47760120058729,65.47760120068729,65.51033664587358,65.51033664597358,65.54299744526588,65.54299744536588,65.5756483550245,65.5756483551245,65.64090637217956,65.64090637227956,65.67350461300788,65.67350461310788,65.70604798720954,65.70604798730955,65.73854418672184,65.73854418682184,65.77096498251379,65.77096498261379,65.80334390840072,65.80334390850072,65.83569411266954,65.83569411276954,65.86795379458265,65.86795379468265,65.90018956707486,65.90018956717486,65.93230067713496,65.93230067723496,65.96439939673446,65.96439939683447,65.99649599442024,65.99649599452025,66.02853234491236,66.02853234501237,66.06055213689908,66.06055213699908,66.09252634563171,66.09252634573171,66.12447910787908,66.12447910797908,66.15637401008576,66.15637401018576,66.18823678045581,66.18823678055581,66.22009333664992,66.22009333674993,66.25191056165724,66.25191056175724,66.28370330887394,66.28370330897394,66.31547453382271,66.31547453392271,66.3472184391321,66.34721843923211,66.37892274801548,66.37892274811549,66.41060568619622,66.41060568629622,66.44227521085082,66.44227521095083,66.47378521306186,66.47378521316186,66.50529457112053,66.50529457122053,66.5366282271434,66.5366282272434,66.56795396388105,66.56795396398105,66.59925078954402,66.59925078964402,66.63053253446292,66.63053253456292,66.66178991526515,66.66178991536515,66.69304589408866,66.69304589418866,66.72424632424179,66.72424632434179,66.75543504597806,66.75543504607806,66.78660800492655,66.78660800502655,66.81776061623795,66.81776061633795,66.8488263048687,66.8488263049687,66.87987698853802,66.87987698863802,66.91081263416964,66.91081263426965,66.94174346760406,66.94174346770406,66.97258881822786,66.97258881832786,67.003428826176,67.00342882627601,67.0342580729902,67.0342580730902,67.06505363442378,67.06505363452378,67.09584328481195,67.09584328491195,67.12656992194036,67.12656992204036,67.15729140584968,67.15729140594968,67.18795840204551,67.18795840214551,67.21858087594416,67.21858087604416,67.24916504172154,67.24916504182154,67.27970373791884,67.27970373801884,67.31020090523296,67.31020090533296,67.34063975781059,67.3406397579106,67.37100203203705,67.37100203213706,67.4012806043448,67.4012806044448,67.43150271859058,67.43150271869058,67.46161195460374,67.46161195470374,67.49167992697295,67.49167992707295,67.52169007719279,67.52169007729279,67.55163183358587,67.55163183368587,67.58157059656494,67.58157059666495,67.61147355224966,67.61147355234966,67.64135108286085,67.64135108296085,67.67122217194819,67.67122217204819,67.7010737470074,67.7010737471074,67.7308947816653,67.7308947817653,67.76071089045202,67.76071089055202,67.79047546704795,67.79047546714796,67.82018756417018,67.82018756427019,67.8498890138324,67.8498890139324,67.87944071701078,67.87944071711078,67.9089626755055,67.90896267560551,67.93847201619177,67.93847201629177,67.96793228762282,67.96793228772282,67.99729180604261,67.99729180614261,68.02659895866265,68.02659895876265,68.055879473687,68.055879473787,68.08514653729391,68.08514653739391,68.1143932153724,68.1143932154724,68.1436270482945,68.1436270483945,68.17285459125804,68.17285459135805,68.20205477669087,68.20205477679087,68.2312294233762,68.23122942347621,68.26039304368835,68.26039304378835,68.28954969199823,68.28954969209823,68.3186332858494,68.3186332859494,68.34753378885799,68.34753378895799,68.37640136431233,68.37640136441233,68.40522555420897,68.40522555430897,68.43402359909705,68.43402359919705,68.46275821392088,68.46275821402088,68.4913729785098,68.4913729786098,68.51998186994463,68.51998187004463,68.54857799200572,68.54857799210572,68.57708385695028,68.57708385705028,68.60550704304168,68.60550704314169,68.6338775980941,68.6338775981941,68.6622474332115,68.6622474333115,68.69060487786834,68.69060487796834,68.71890037022226,68.71890037032226,68.74710988717847,68.74710988727847,68.77530481597772,68.77530481607772,68.80321067192143,68.80321067202144,68.8310795838313,68.8310795839313,68.85885770814622,68.85885770824622,68.88655163795531,68.88655163805531,68.91409782952036,68.91409782962036,68.94154796659743,68.94154796669743,68.9689712387309,68.9689712388309,68.99637749766318,68.99637749776318,69.02369205960888,69.02369205970888,69.05093947814129,69.05093947824129,69.07810736279984,69.07810736289984,69.10521015017619,69.10521015027619,69.13210548259329,69.13210548269329,69.1589924410294,69.1589924411294,69.18585867291523,69.18585867301523,69.21270148796731,69.21270148806731,69.23950648748526,69.23950648758526,69.2663060306536,69.2663060307536,69.2930523365219,69.2930523366219,69.31977526344775,69.31977526354775,69.3464326005043,69.3464326006043,69.3730735306207,69.3730735307207,69.39966542937321,69.39966542947322,69.42619139723456,69.42619139733456,69.45266867475388,69.45266867485388,69.47904394884849,69.47904394894849,69.50531350616934,69.50531350626935,69.53156479987551,69.53156479997551,69.55781544942931,69.55781544952931,69.58406018793768,69.58406018803768,69.61026025258361,69.61026025268362,69.63642826117555,69.63642826127555,69.66252431415697,69.66252431425697,69.68856705405572,69.68856705415573,69.7145546999799,69.7145547000799,69.74047357316415,69.74047357326415,69.76638392080216,69.76638392090216,69.79229286646142,69.79229286656143,69.81815414484421,69.81815414494422,69.84393562411393,69.84393562421393,69.86967754484893,69.86967754494893,69.89533773401368,69.89533773411368,69.92094551948738,69.92094551958738,69.94653435930267,69.94653435940268,69.97200925992848,69.97200926002849,69.99746214569927,69.99746214579928,70.02290980246833,70.02290980256834,70.04815818880257,70.04815818890258,70.07339695074236,70.07339695084237,70.09862491365686,70.09862491375686,70.12384995893999,70.12384995903999,70.14902002392249,70.14902002402249,70.17416663417991,70.17416663427991,70.19930828067484,70.19930828077484,70.22443962073163,70.22443962083163,70.24957024085339,70.24957024095339,70.27464887408854,70.27464887418854,70.29968628157108,70.29968628167109,70.32466454070816,70.32466454080816,70.34958808478383,70.34958808488383,70.37447847395735,70.37447847405736,70.3992898218441,70.39928982194411,70.42408597531285,70.42408597541285,70.44882146478345,70.44882146488345,70.47352955883206,70.47352955893206,70.49813254236796,70.49813254246796,70.52265656039975,70.52265656049975,70.54709767223046,70.54709767233047,70.57150900148622,70.57150900158622,70.59589661077767,70.59589661087767,70.62023814422795,70.62023814432796,70.64451101861228,70.64451101871228,70.66877904290806,70.66877904300806,70.69298159100846,70.69298159110846,70.74125001322976,70.74125001332976,70.76517997119741,70.76517997129741,70.78906260291045,70.78906260301045,70.81286823946783,70.81286823956783,70.83667141308962,70.83667141318962,70.86039747788182,70.86039747798182,70.88411031860446,70.88411031870446,70.90780955634438,70.90780955644438,70.93149939703775,70.93149939713776,70.95518571383866,70.95518571393866,70.9788646418328,70.9788646419328,71.00249715296393,71.00249715306393,71.02611765254763,71.02611765264763,71.0497306117593,71.0497306118593,71.07332542103023,71.07332542113024,71.0968755185474,71.09687551864741,71.12036370165299,71.12036370175299,71.1438340758397,71.14383407593971,71.16725102326978,71.16725102336979,71.19066289326342,71.19066289336342,71.2139905308599,71.2139905309599,71.23731286367205,71.23731286377205,71.26049280091581,71.26049280101581,71.28360669359446,71.28360669369447,71.3066624231019,71.3066624232019,71.32971201421603,71.32971201431603,71.35275781619846,71.35275781629846,71.37571332317307,71.37571332327308,71.39865772799187,71.39865772809188,71.44452039262092,71.44452039272092,71.46740216309311,71.46740216319311,71.49023043102605,71.49023043112605,71.51304077636617,71.51304077646617,71.53581379875926,71.53581379885927,71.55839793293833,71.55839793303834,71.58096645589491,71.58096645599491,71.60353365265539,71.60353365275539,71.62607724312552,71.62607724322552,71.64860723061294,71.64860723071294,71.67111687046327,71.67111687056327,71.69361999300712,71.69361999310712,71.71609242358439,71.71609242368439,71.73846235814982,71.73846235824982,71.76078379182981,71.76078379192981,71.78309855663805,71.78309855673805,71.80540706937903,71.80540706947903,71.82769572707002,71.82769572717002,71.849936414354,71.849936414454,71.87214853158514,71.87214853168514,71.8943389749831,71.8943389750831,71.91651270831038,71.91651270841038,71.93867746139557,71.93867746149557,71.9829342162378,71.9829342163378,72.00499476820191,72.00499476830191,72.04905895937237,72.04905895947238,72.07105062492262,72.07105062502262,72.09302167759655,72.09302167769656,72.11493350614239,72.1149335062424,72.13681930335362,72.13681930345362,72.15866232126824,72.15866232136824,72.18045149562172,72.18045149572173,72.20223070455891,72.20223070465892,72.22399608316549,72.22399608326549,72.24567935128866,72.24567935138866,72.2673273425959,72.2673273426959,72.28890925144674,72.28890925154674,72.31047160837814,72.31047160847814,72.33194901297735,72.33194901307735,72.35342505348916,72.35342505358916,72.39633806856521,72.39633806866522,72.43908326299935,72.43908326309935,72.46038125221372,72.46038125231372,72.48165571092038,72.48165571102038,72.50286295043111,72.50286295053111,72.52406803013679,72.52406803023679,72.54525401261883,72.54525401271883,72.56642999179324,72.56642999189324,72.58758850307062,72.58758850317062,72.60869423174377,72.60869423184377,72.62975619594606,72.62975619604606,72.65081156705922,72.65081156715922,72.67185488873368,72.67185488883368,72.69288502422988,72.69288502432988,72.71390299661341,72.71390299671341,72.7349178998003,72.7349178999003,72.75592230709242,72.75592230719242,72.77691307351053,72.77691307361053,72.79787545933237,72.79787545943238,72.81879256503069,72.81879256513069,72.83968079754565,72.83968079764566,72.86050128038625,72.86050128048625,72.88128894934653,72.88128894944653,72.9020184930269,72.9020184931269,72.92273886700862,72.92273886710862,72.94345181429225,72.94345181439225,72.96413054571686,72.96413054581686,72.98476835451936,72.98476835461936,73.00527903795414,73.00527903805414,73.02576600142463,73.02576600152463,73.04614258748941,73.04614258758942,73.06645411416332,73.06645411426332,73.08676325368427,73.08676325378427,73.10698512950272,73.10698512960272,73.12720207945,73.12720207955,73.14741584652667,73.14741584662667,73.16760865970518,73.16760865980518,73.18779544816431,73.18779544826431,73.20797299114217,73.20797299124217,73.2281310200919,73.2281310201919,73.24823353826255,73.24823353836256,73.26826557858406,73.26826557868407,73.28828185611779,73.2882818562178,73.30829131321451,73.30829131331451,73.32826030238492,73.32826030248492,73.34822770012003,73.34822770022004,73.36818153276376,73.36818153286376,73.38809792878651,73.38809792888651,73.40794524893882,73.40794524903882,73.42773027576638,73.42773027586638,73.44740242436131,73.44740242446132,73.467039371923,73.467039372023,73.4866180805308,73.4866180806308,73.50615340358094,73.50615340368094,73.52566265740514,73.52566265750514,73.5451501237222,73.5451501238222,73.56454354379107,73.56454354389108,73.58387997531953,73.58387997541953,73.62252112334434,73.62252112344434,73.6417890094808,73.6417890095808,73.66104196643845,73.66104196653845,73.6802594192323,73.6802594193323,73.69941006963471,73.69941006973471,73.71854798855468,73.71854798865468,73.73766332424987,73.73766332434987,73.75676937657245,73.75676937667245,73.7758563695627,73.7758563696627,73.79493809565993,73.79493809575993,73.81400777231842,73.81400777241842,73.83304478666187,73.83304478676187,73.85202337259489,73.85202337269489,73.87097259275741,73.87097259285741,73.88991597765715,73.88991597775716,73.90885174640222,73.90885174650222,73.92774068147979,73.92774068157979,73.94658141880244,73.94658141890244,73.96540173270535,73.96540173280535,73.9841633529586,73.9841633530586,74.00290523183583,74.00290523193583,74.02161425894143,74.02161425904143,74.04028891862272,74.04028891872272,74.0589155700057,74.0589155701057,74.0775140681403,74.0775140682403,74.09611074749172,74.09611074759172,74.11470613853837,74.11470613863837,74.13320987048976,74.13320987058977,74.15169795332736,74.15169795342736,74.17015769346,74.17015769356,74.18846905119624,74.18846905129624,74.20676282736149,74.20676282746149,74.22503652112886,74.22503652122886,74.24330195458917,74.24330195468917,74.26153427103864,74.26153427113864,74.27971513107995,74.27971513117996,74.2978284309035,74.2978284310035,74.31593096959307,74.31593096969307,74.33395060208159,74.33395060218159,74.35194496106182,74.35194496116182,74.36988430185012,74.36988430195012,74.40573380711282,74.40573380721283,74.42352404556968,74.42352404566968,74.4412640022492,74.4412640023492,74.45898531640088,74.45898531650089,74.47659826138664,74.47659826148664,74.49418604653806,74.49418604663806,74.51177201290629,74.5117720130063,74.52930148332122,74.52930148342122,74.546810719773,74.54681071987301,74.56428505832207,74.56428505842207,74.58174772634555,74.58174772644556,74.5992019067141,74.5992019068141,74.61661137532886,74.61661137542886,74.63393119308817,74.63393119318818,74.65123687738635,74.65123687748635,74.66849614482149,74.66849614492149,74.68575249462523,74.68575249472524,74.70292639292371,74.70292639302372,74.72008320223834,74.72008320233834,74.73717192085675,74.73717192095675,74.7541806509055,74.7541806510055,74.77116369064147,74.77116369074147,74.78807386537532,74.78807386547533,74.80498260023914,74.80498260033914,74.82187231366194,74.82187231376194,74.8386945426496,74.83869454274961,74.85550938283049,74.85550938293049,74.87230633830987,74.87230633840987,74.88909927730968,74.88909927740968,74.90588808615595,74.90588808625596,74.92266700536855,74.92266700546855,74.93943148798948,74.93943148808948,74.95616804470994,74.95616804480994,74.97286091274219,74.9728609128422,74.98951983015463,74.98951983025464,75.00617734558834,75.00617734568834,75.02270512115348,75.02270512125348,75.03918852598669,75.03918852608669,75.05554382027795,75.05554382037795,75.0718685741679,75.0718685742679,75.08818976627407,75.08818976637407,75.10448837515546,75.10448837525546,75.12065834301644,75.12065834311645,75.13680371941287,75.13680371951287,75.15291794914691,75.15291794924691,75.16903039798906,75.16903039808906,75.18513640530738,75.18513640540738,75.20119864815483,75.20119864825483,75.21725516941345,75.21725516951345,75.23320635281148,75.23320635291148,75.24915537640447,75.24915537650448,75.26506840324653,75.26506840334653,75.28092974633253,75.28092974643253,75.29672940235486,75.29672940245486,75.31251503858998,75.31251503868998,75.32827911466586,75.32827911476586,75.34402913306324,75.34402913316325,75.35970056486966,75.35970056496966,75.37535926519364,75.37535926529364,75.39098632626813,75.39098632636814,75.40657867889648,75.40657867899648,75.42216955376347,75.42216955386347,75.43775432812846,75.43775432822846,75.45333277464356,75.45333277474356,75.4689095539407,75.4689095540407,75.48445260996594,75.48445261006594,75.49996914206946,75.49996914216946,75.51548021782335,75.51548021792335,75.53097681909425,75.53097681919425,75.54646781245027,75.54646781255028,75.57728106912879,75.57728106922879,75.59259128301255,75.59259128311255,75.6078566714686,75.6078566715686,75.62311073042092,75.62311073052092,75.63830996063791,75.63830996073791,75.65346910184176,75.65346910194177,75.6685789085512,75.6685789086512,75.68365328687946,75.68365328697946,75.69865578537984,75.69865578547984,75.71362637939151,75.71362637949152,75.72856453843605,75.72856453853605,75.74349030701998,75.74349030711998,75.7583956521842,75.75839565228421,75.77328182434213,75.77328182444214,75.78815412827814,75.78815412837814,75.80301104833953,75.80301104843953,75.81779764211703,75.81779764221703,75.83254774655649,75.83254774665649,75.8620004776155,75.8620004777155,75.87664478949857,75.87664478959857,75.89116970584254,75.89116970594254,75.90567995824692,75.90567995834692,75.92017312166745,75.92017312176745,75.93464741521223,75.93464741531223,75.94911155388411,75.94911155398411,75.96355011591719,75.96355011601719,75.97794661858867,75.97794661868868,75.9923389532153,75.9923389533153,76.00666463701576,76.00666463711576,76.02097811981221,76.02097811991221,76.03525257455242,76.03525257465242,76.04950376402414,76.04950376412414,76.06375052021178,76.06375052031179,76.07799155481061,76.07799155491061,76.092195455919,76.092195456019,76.10639803083131,76.10639803093132,76.12058688908695,76.12058688918695,76.13468211789889,76.13468211799889,76.14876291011915,76.14876291021915,76.16280115039068,76.16280115049068,76.17682643183188,76.17682643193189,76.19080150727838,76.19080150737838,76.20472455794697,76.20472455804698,76.21863961354774,76.21863961364774,76.23253538246826,76.23253538256826,76.24636457634524,76.24636457644525,76.26017834845631,76.26017834855631,76.27394327866007,76.27394327876007,76.2876260604891,76.2876260605891,76.30129232170403,76.30129232180403,76.3148953802026,76.3148953803026,76.32849434643897,76.32849434653897,76.3420828546719,76.3420828547719,76.35564142876461,76.35564142886462,76.36915949703969,76.36915949713969,76.38267684537976,76.38267684547976,76.3961764984749,76.3961764985749,76.40967224876442,76.40967224886442,76.42312794793213,76.42312794803213,76.43656023026608,76.43656023036608,76.44999111062131,76.44999111072131,76.46341805027959,76.46341805037959,76.47681160768781,76.47681160778781,76.49016579601792,76.49016579611792,76.50350933688803,76.50350933698803,76.51674670628857,76.51674670638857,76.52997304931594,76.52997304941594,76.54316161470051,76.54316161480051,76.55634044201668,76.55634044211668,76.56948115066817,76.56948115076817,76.58255877027727,76.58255877037728,76.59560664520271,76.59560664530271,76.6086496321483,76.6086496322483,76.62168083489436,76.62168083499436,76.63469733580953,76.63469733590954,76.64771148746306,76.64771148756306,76.66072344142022,76.66072344152022,76.67371455515315,76.67371455525316,76.6866674365475,76.6866674366475,76.69961406587456,76.69961406597456,76.71253917293372,76.71253917303372,76.72543642987505,76.72543642997505,76.73833273953346,76.73833273963346,76.75121093714246,76.75121093724246,76.76403032742786,76.76403032752786,76.77674343256975,76.77674343266975,76.78940690008669,76.78940690018669,76.80205403644611,76.80205403654611,76.81468628151804,76.81468628161804,76.82730374897643,76.82730374907644,76.83991132680114,76.83991132690115,76.85251761632111,76.85251761642111,76.87771196963764,76.87771196973765,76.89028180771088,76.89028180781088,76.90283618612688,76.90283618622688,76.91535202907372,76.91535202917372,76.92783941564169,76.92783941574169,76.9403103952695,76.94031039536951,76.95274594651615,76.95274594661615,76.96510385845475,76.96510385855476,76.97745832228352,76.97745832238353,76.98972025551679,76.98972025561679,77.00196922991972,77.00196923001972,77.01420191105645,77.01420191115645,77.02643250817074,77.02643250827074,77.0386270706428,77.0386270707428,77.05078969073485,77.05078969083485,77.06295094673948,77.06295094683948,77.07511015661301,77.07511015671301,77.08723973547684,77.08723973557684,77.09935142963916,77.09935142973916,77.11145043021035,77.11145043031036,77.12351305511747,77.12351305521747,77.13552342789878,77.13552342799878,77.1475313377445,77.1475313378445,77.15951291312507,77.15951291322507,77.17148800909047,77.17148800919047,77.19541061614268,77.19541061624268,77.20729318151257,77.20729318161257,77.21911811418967,77.21911811428967,77.23093145212383,77.23093145222383,77.24273922003444,77.24273922013444,77.25453323339705,77.25453323349706,77.26632190408401,77.26632190418401,77.2781004577894,77.27810045788941,77.28987753373343,77.28987753383343,77.30164877441469,77.30164877451469,77.3134142935071,77.3134142936071,77.3251601091148,77.3251601092148,77.33689273854428,77.33689273864428,77.34860585394563,77.34860585404563,77.36030703358219,77.36030703368219,77.37199957399854,77.37199957409854,77.38369059876221,77.38369059886222,77.39537150654431,77.39537150664431,77.40704665484627,77.40704665494627,77.41871407331959,77.4187140734196,77.43037501237777,77.43037501247777,77.44201382290694,77.44201382300695,77.45364952634816,77.45364952644816,77.46528041759214,77.46528041769214,77.47689376515646,77.47689376525646,77.48848934169322,77.48848934179323,77.5000686628551,77.5000686629551,77.51160145348,77.51160145358,77.52310764440053,77.52310764450053,77.53461065245045,77.53461065255046,77.55758222534337,77.55758222544337,77.56901653643601,77.56901653653601,77.58039522307564,77.58039522317564,77.59175238744737,77.59175238754737,77.60310307240393,77.60310307250393,77.61445068816383,77.61445068826383,77.62576969597959,77.62576969607959,77.63707843524848,77.63707843534849,77.64835519424605,77.64835519434605,77.65963108174333,77.65963108184333,77.67087400379503,77.67087400389504,77.68211222732346,77.68211222742346,77.6933310504977,77.6933310505977,77.7045355507542,77.7045355508542,77.71571523219824,77.71571523229824,77.72687600587521,77.72687600597521,77.73799638740851,77.73799638750852,77.74907073099196,77.74907073109196,77.7601373826381,77.76013738273811,77.77118164051605,77.77118164061605,77.78222191980574,77.78222191990574,77.79325068013512,77.79325068023512,77.80427042233111,77.80427042243112,77.81528133585032,77.81528133595032,77.82626943879681,77.82626943889682,77.8372418926295,77.83724189272951,77.8481770614065,77.8481770615065,77.85908680510995,77.85908680520996,77.86996703147764,77.86996703157764,77.88083744399431,77.88083744409431,77.89170562092328,77.89170562102328,77.9025680383721,77.9025680384721,77.91342530260184,77.91342530270184,77.93511182937824,77.93511182947825,77.95676398873034,77.95676398883035,77.96758559723101,77.96758559733101,77.97837780206989,77.97837780216989,77.98914450605258,77.98914450615258,77.99990078992316,77.99990079002316,78.01062998150226,78.01062998160226,78.0213418567496,78.0213418568496,78.032047517821,78.032047517921,78.04274461545481,78.04274461555481,78.05342685969245,78.05342685979245,78.06406549102451,78.06406549112451,78.0746843430892,78.0746843431892,78.0852975114564,78.0852975115564,78.10651498162265,78.10651498172265,78.11711814668223,78.11711814678223,78.12767947972809,78.12767947982809,78.13823349974983,78.13823349984983,78.14878168450878,78.14878168460878,78.1593273305495,78.1593273306495,78.16986831595827,78.16986831605827,78.18038804433834,78.18038804443835,78.1908948896707,78.1908948897707,78.20139821111061,78.20139821121062,78.21188921787258,78.21188921797258,78.22237798904686,78.22237798914686,78.2328630468721,78.2328630469721,78.24332715079916,78.24332715089916,78.25376916408855,78.25376916418855,78.27464917418777,78.27464917428777,78.28507281018855,78.28507281028855,78.29549296018817,78.29549296028817,78.30591057146958,78.30591057156958,78.3163189372697,78.3163189373697,78.32671582200084,78.32671582210084,78.3370962619005,78.3370962620005,78.34745953703367,78.34745953713367,78.35781834099147,78.35781834109147,78.36817532616607,78.36817532626607,78.37851616963972,78.37851616973973,78.38884996532846,78.38884996542846,78.39911631447337,78.39911631457338,78.40937849557343,78.40937849567344,78.41963256793171,78.41963256803172,78.42987542446023,78.42987542456024,78.44009672082952,78.44009672092952,78.45031184091417,78.45031184101417,78.4605119939287,78.4605119940287,78.47070566752808,78.47070566762808,78.48088134275199,78.48088134285199,78.49103784496961,78.49103784506961,78.50110056617827,78.50110056627827,78.51115745212414,78.51115745222414,78.52120880593777,78.52120880603778,78.53125841675083,78.53125841685083,78.55135233359259,78.55135233369259,78.56135973347878,78.56135973357878,78.57135966877557,78.57135966887557,78.58132777536628,78.58132777546628,78.59127739099442,78.59127739109442,78.60122420266511,78.60122420276511,78.61116824826968,78.61116824836968,78.62109891823944,78.62109891833944,78.63102670846912,78.63102670856912,78.64094532900015,78.64094532910015,78.65084947504816,78.65084947514816,78.66075085503007,78.66075085513008,78.67065136351168,78.67065136361168,78.69045071325698,78.69045071335698,78.70032935844883,78.70032935854883,78.71019686359354,78.71019686369354,78.72005311501718,78.72005311511718,78.72989136806537,78.72989136816537,78.7397075683672,78.7397075684672,78.74951679666675,78.74951679676676,78.75927547794974,78.75927547804974,78.76903162051451,78.76903162061451,78.77876889320353,78.77876889330354,78.7885024146522,78.7885024147522,78.80796612311366,78.80796612321366,78.81769521127826,78.81769521137826,78.82741550865738,78.82741550875738,78.83712830355577,78.83712830365577,78.8468298447331,78.8468298448331,78.85653074175804,78.85653074185804,78.86622512147649,78.86622512157649,78.8759107861921,78.8759107862921,78.88558023342412,78.88558023352412,78.89524710404662,78.89524710414662,78.90491211799458,78.90491211809459,78.91456493093855,78.91456493103856,78.92420853629255,78.92420853639256,78.93384566223139,78.9338456623314,78.94347024614437,78.94347024624437,78.95308778227245,78.95308778237245,78.96269163963517,78.96269163973517,78.97226166005203,78.97226166015203,78.98182122246546,78.98182122256546,78.99133891497387,78.99133891507387,79.00084721043572,79.00084721053572,79.01035402813623,79.01035402823624,79.01983489028476,79.01983489038476,79.02929900447128,79.02929900457129,79.03876156511382,79.03876156521382,79.04821624436246,79.04821624446247,79.05766675556626,79.05766675566626,79.06711199987704,79.06711199997704,79.07654561155357,79.07654561165357,79.08597573722895,79.08597573732895,79.09540158118556,79.09540158128556,79.1048266294245,79.1048266295245,79.1142466381183,79.1142466382183,79.12364266160861,79.12364266170862,79.13302261918062,79.13302261928062,79.142384919399,79.14238491949901,79.15174183905042,79.15174183915042,79.1610882628071,79.1610882629071,79.17042968490995,79.17042968500995,79.17976458970632,79.17976458980633,79.18909623584946,79.18909623594946,79.19842390340433,79.19842390350433,79.20770507831355,79.20770507841355,79.22625905415097,79.22625905425097,79.23553128670945,79.23553128680945,79.24480287511551,79.24480287521551,79.25406942397646,79.25406942407646,79.26329717212904,79.26329717222904,79.28174971291152,79.28174971301152,79.29096203929818,79.29096203939818,79.30017156172741,79.30017156182741,79.30935630323548,79.30935630333548,79.31853354226283,79.31853354236283,79.32770327880945,79.32770327890945,79.33686994615942,79.33686994625943,79.34602854265893,79.34602854275893,79.3551716795012,79.3551716796012,79.364305191949,79.364305192049,79.37343358906905,79.37343358916905,79.38255198288148,79.38255198298148,79.39164480005509,79.39164480015509,79.40073295659674,79.40073295669674,79.4098195217031,79.4098195218031,79.41887111312407,79.41887111322407,79.4279180439131,79.4279180440131,79.43694019378096,79.43694019388096,79.44595404545045,79.44595404555045,79.45496653303253,79.45496653313253,79.46397818700565,79.46397818710565,79.47298582449919,79.47298582459919,79.48197273544247,79.48197273554247,79.49094646020751,79.49094646030751,79.49991222779603,79.49991222789603,79.50886568070662,79.50886568080662,79.51781852735613,79.51781852745613,79.52675985504534,79.52675985514534,79.53568958799163,79.53568958809163,79.54460723360788,79.54460723370788,79.5624356286207,79.5624356287207,79.57134122479823,79.57134122489823,79.58024352443118,79.58024352453118,79.5891393825403,79.5891393826403,79.59803395234462,79.59803395244462,79.60689908059584,79.60689908069584,79.61575689582293,79.61575689592293,79.62460603393848,79.62460603403848,79.63344168274526,79.63344168284526,79.64223652260388,79.64223652270388,79.6510229505902,79.6510229506902,79.65980562733615,79.65980562743616,79.66858633373363,79.66858633383363,79.67735987867225,79.67735987877225,79.6861276262394,79.6861276263394,79.69489014480483,79.69489014490483,79.70365171608735,79.70365171618735,79.71240980136874,79.71240980146874,79.72116235451786,79.72116235461786,79.72990964077397,79.72990964087397,79.7386185810175,79.7386185811175,79.75603365754715,79.75603365764715,79.76473888444166,79.76473888454166,79.77343024311422,79.77343024321422,79.78211599387191,79.78211599397191,79.79077730473028,79.79077730483029,79.7994357737399,79.7994357738399,79.80809181770526,79.80809181780526,79.81674570186556,79.81674570196556,79.82537556293101,79.82537556303102,79.83400239269115,79.83400239279115,79.85125347560187,79.85125347570187,79.8598565475064,79.8598565476064,79.86845560293133,79.86845560303134,79.87703506854552,79.87703506864553,79.88561335952889,79.88561335962889,79.89416436876418,79.89416436886418,79.90270537469185,79.90270537479185,79.91124550911924,79.91124550921924,79.91978424156791,79.91978424166791,79.9368579552249,79.9368579553249,79.94539062506287,79.94539062516287,79.95389248926034,79.95389248936034,79.96238995806506,79.96238995816506,79.97086912536378,79.97086912546378,79.97934598129218,79.97934598139219,79.98780324740983,79.98780324750983,79.99624963871956,79.99624963881956,80.00467969887177,80.00467969897177,80.01309517086702,80.01309517096702,80.02150723264377,80.02150723274377,80.0299011444798,80.0299011445798,80.03826262134866,80.03826262144867,80.04661220034409,80.04661220044409,80.05495018459656,80.05495018469657,80.06328422815211,80.06328422825212,80.0716097840527,80.0716097841527,80.07992169907926,80.07992169917927,80.08822505066823,80.08822505076823,80.09651199531704,80.09651199541705,80.10478420024364,80.10478420034364,80.1213237600083,80.1213237601083,80.12958213460426,80.12958213470426,80.13783922089547,80.13783922099547,80.14608190848632,80.14608190858632,80.15430962900705,80.15430962910705,80.1625259063501,80.1625259064501,80.17072657247066,80.17072657257066,80.17892238850267,80.17892238860267,80.18711248294586,80.18711248304587,80.19529988710555,80.19529988720555,80.20347819734923,80.20347819744923,80.21165249111331,80.21165249121331,80.21980530050081,80.21980530060081,80.22795716260538,80.22795716270538,80.23609966555472,80.23609966565472,80.24423841726369,80.24423841736369,80.25236364177259,80.25236364187259,80.26046616938274,80.26046616948274,80.26854914507345,80.26854914517345,80.27662692965376,80.27662692975376,80.29276631922214,80.29276631932214,80.30082447610037,80.30082447620038,80.30888017004301,80.30888017014301,80.31693419676772,80.31693419686772,80.32494309493417,80.32494309503417,80.33294486953307,80.33294486963307,80.34093690606356,80.34093690616356,80.3489231831139,80.3489231832139,80.35690521633677,80.35690521643677,80.36487307820717,80.36487307830717,80.3728258593335,80.3728258594335,80.38077432084972,80.38077432094973,80.38871425681967,80.38871425691967,80.39663733115366,80.39663733125366,80.40455309246352,80.40455309256352,80.41245680433465,80.41245680443465,80.42034895935417,80.42034895945417,80.42823577169803,80.42823577179803,80.43611538469169,80.43611538479169,80.44399333046742,80.44399333056742,80.45186957113391,80.45186957123391,80.45973505066642,80.45973505076643,80.47546021236,80.47546021246,80.48331489286726,80.48331489296726,80.49114843001975,80.49114843011975,80.49897594245287,80.49897594255287,80.50679701336216,80.50679701346216,80.51461296894368,80.51461296904368,80.52242812880755,80.52242812890755,80.53024158356216,80.53024158366216,80.53805007455428,80.53805007465428,80.54585098728303,80.54585098738303,80.55362719487329,80.55362719497329,80.56917294118202,80.56917294128202,80.57693899389936,80.57693899399936,80.58470220476795,80.58470220486795,80.59245541232892,80.59245541242892,80.60018793123093,80.60018793133094,80.60791313710882,80.60791313720883,80.6233544928399,80.6233544929399,80.63105620610139,80.6310562062014,80.63874200500985,80.63874200510985,80.6464262503743,80.6464262504743,80.65409954514824,80.65409954524824,80.66176253348401,80.66176253358401,80.66942199792732,80.66942199802732,80.67707198954145,80.67707198964145,80.68471932876339,80.6847193288634,80.6923579908738,80.6923579909738,80.69998562661104,80.69998562671104,80.70760053086583,80.70760053096583,80.71521229014135,80.71521229024135,80.72279964740886,80.72279964750886,80.73038446595817,80.73038446605817,80.73795894017799,80.73795894027799,80.74552685920001,80.74552685930001,80.75309322467805,80.75309322477806,80.76065887022108,80.76065887032108,80.76821193584692,80.76821193594692,80.77576060608004,80.77576060618004,80.78330859426944,80.78330859436944,80.79084305525878,80.79084305535878,80.79837531855173,80.79837531865174,80.80590686190966,80.80590686200966,80.81342646950283,80.81342646960283,80.82094323524723,80.82094323534723,80.82845890214344,80.82845890224344,80.83597350808279,80.83597350818279,80.84346064281749,80.84346064291749,80.85094626189952,80.85094626199952,80.8584279402846,80.8584279403846,80.86589423479508,80.86589423489508,80.87334859354078,80.87334859364078,80.88079268373964,80.88079268383964,80.88822260258604,80.88822260268604,80.89564236655954,80.89564236665954,80.90305720466186,80.90305720476186,80.91046704111038,80.91046704121038,80.92528148500568,80.92528148510569,80.93268442523456,80.93268442533456,80.94008516776704,80.94008516786704,80.94748102231966,80.94748102241967,80.95487039745713,80.95487039755713,80.96224836730825,80.96224836740825,80.96962201754926,80.96962201764926,80.97697206149994,80.97697206159994,80.98431812686233,80.98431812696234,80.99165896322303,80.99165896332303,80.99899729875682,80.99899729885682,81.0063344596598,81.0063344597598,81.01366517903892,81.01366517913893,81.02099237452558,81.02099237462558,81.02831706918535,81.02831706928535,81.03563611803891,81.03563611813891,81.04295043047787,81.04295043057787,81.05026042330672,81.05026042340673,81.05756238317643,81.05756238327643,81.06485172523765,81.06485172533765,81.07213993055937,81.07213993065938,81.07942730227212,81.07942730237212,81.08671209737534,81.08671209747534,81.09399192871605,81.09399192881605,81.10126975181699,81.10126975191699,81.10853836732794,81.10853836742794,81.1157830355267,81.1157830356267,81.12302622596413,81.12302622606413,81.13026577883512,81.13026577893513,81.14474158803257,81.14474158813258,81.15197469937972,81.15197469947972,81.15920705290054,81.15920705300054,81.16643736029026,81.16643736039026,81.18089520900357,81.18089520910357,81.18810388045142,81.18810388055142,81.1953108088987,81.1953108089987,81.20251546386696,81.20251546396696,81.20971352574611,81.20971352584611,81.21691033721181,81.21691033731182,81.22410536778563,81.22410536788563,81.23129210016107,81.23129210026107,81.2384774305578,81.2384774306578,81.24564737707992,81.24564737717992,81.2528156563841,81.2528156564841,81.25997662266415,81.25997662276416,81.26712936652847,81.26712936662847,81.27428093576196,81.27428093586197,81.28857793583563,81.28857793593563,81.29569862364596,81.29569862374596,81.30281730321651,81.30281730331652,81.3099349597215,81.3099349598215,81.31704916811668,81.31704916821668,81.32415625294429,81.3241562530443,81.33126216314109,81.33126216324109,81.33836636822865,81.33836636832865,81.34545787972507,81.34545787982508,81.35254836815595,81.35254836825595,81.35963825032577,81.35963825042577,81.36672722310398,81.36672722320398,81.37381475601215,81.37381475611215,81.38798095526037,81.38798095536038,81.40213582500489,81.40213582510489,81.41627023343835,81.41627023353836,81.42331479759332,81.42331479769332,81.43035481479028,81.43035481489028,81.43739343000853,81.43739343010853,81.44442992331302,81.44442992341303,81.4514591414847,81.4514591415847,81.45847649967425,81.45847649977425,81.46548646905701,81.46548646915701,81.47249272509075,81.47249272519075,81.47949644240624,81.47949644250625,81.48649674950325,81.48649674960325,81.49349432842543,81.49349432852543,81.50049054326023,81.50049054336023,81.5144820635382,81.5144820636382,81.5214760806766,81.52147608077661,81.52846865794498,81.52846865804499,81.53545312647158,81.53545312657158,81.54243346484465,81.54243346494465,81.54941141606476,81.54941141616476,81.55638406250053,81.55638406260053,81.56334878965109,81.56334878975109,81.57030927297417,81.57030927307417,81.5772660050569,81.5772660051569,81.58422205509592,81.58422205519592,81.59116935224077,81.59116935234077,81.59810998051387,81.59810998061387,81.60504484930684,81.60504484940684,81.6119691085311,81.6119691086311,81.61889204155928,81.61889204165928,81.62581436832639,81.62581436842639,81.6327285863517,81.6327285864517,81.63964053089803,81.63964053099804,81.6465507703351,81.6465507704351,81.66036151114082,81.66036151124082,81.66725943589994,81.66725943599994,81.67415652705009,81.67415652715009,81.68103728704271,81.68103728714271,81.68791062033725,81.68791062043725,81.69478160437016,81.69478160447017,81.70165171690277,81.70165171700278,81.70851887391268,81.70851887401268,81.71538125661668,81.71538125671668,81.72224087325456,81.72224087335456,81.7290838934957,81.7290838935957,81.73592312460516,81.73592312470517,81.74276174945355,81.74276174955355,81.74959954069298,81.74959954079299,81.75643361858336,81.75643361868336,81.76326553666868,81.76326553676869,81.77008695885927,81.77008695895927,81.776899666047,81.776899666147,81.78371055445153,81.78371055455153,81.79051947250758,81.79051947260758,81.80413219329193,81.80413219339194,81.81093239634514,81.81093239644514,81.81773108374566,81.81773108384566,81.82452859651538,81.82452859661538,81.83132508621954,81.83132508631954,81.8381177489007,81.8381177490007,81.85169780737,81.85169780747,81.8584810351133,81.8584810352133,81.86526244407338,81.86526244417338,81.87203881348834,81.87203881358835,81.87880783198786,81.87880783208786,81.89234382285576,81.89234382295577,81.90587712344019,81.9058771235402,81.91940546026211,81.91940546036211,81.92616364184504,81.92616364194504,81.93967344981307,81.93967344991307,81.94641836943512,81.94641836953512,81.95316056088237,81.95316056098237,81.96664282186312,81.96664282196312,81.97338095893947,81.97338095903947,81.98011731512393,81.98011731522394,81.98685279980812,81.98685279990812,81.99358748877464,81.99358748887464,82.00030982517191,82.00030982527191,82.00702988809016,82.00702988819016,82.01374297900614,82.01374297910614,82.02045167452937,82.02045167462937,82.0271583239215,82.0271583240215,82.03386338187832,82.03386338197832,82.04056430968161,82.04056430978162,82.04726288822327,82.04726288832327,82.05394903841304,82.05394903851304,82.06063253621062,82.06063253631062,82.06731118391966,82.06731118401966,82.07398123029981,82.07398123039981,82.08731825386343,82.08731825396343,82.09398132824128,82.09398132834129,82.10063110276695,82.10063110286696,82.10727390529034,82.10727390539034,82.11391227452967,82.11391227462967,82.12054602102836,82.12054602112836,82.13380392752259,82.1338039276226,82.14042562458255,82.14042562468255,82.14704451768506,82.14704451778506,82.15366174356964,82.15366174366964,82.16027726434496,82.16027726444496,82.16689198940263,82.16689198950263,82.18012060590901,82.18012060600901,82.1933472141756,82.1933472142756,82.19995807431897,82.19995807441897,82.20656533478724,82.20656533488724,82.21977701387502,82.21977701397502,82.22637464994884,82.22637465004884,82.23296478354193,82.23296478364193,82.2395523784168,82.2395523785168,82.24613686620371,82.24613686630371,82.25271718594577,82.25271718604577,82.25929492907828,82.25929492917828,82.26586956512283,82.26586956522283,82.27243700181718,82.27243700191718,82.27900171033674,82.27900171043675,82.2855646379644,82.2855646380644,82.29212544367833,82.29212544377833,82.29868276339111,82.29868276349112,82.3052380369728,82.3052380370728,82.31834509813501,82.31834509823501,82.3248947259105,82.3248947260105,82.33144329272912,82.33144329282912,82.3379829929796,82.3379829930796,82.34452045764242,82.34452045774242,82.35105648243518,82.35105648253518,82.35758940013997,82.35758940023997,82.36411818769123,82.36411818779123,82.3706422009366,82.3706422010366,82.37716341022451,82.37716341032451,82.39020370688661,82.39020370698661,82.39671392769266,82.39671392779266,82.40322210236761,82.40322210246761,82.40972940554228,82.40972940564228,82.41623553408613,82.41623553418613,82.42274098058627,82.42274098068627,82.42924544191217,82.42924544201217,82.43574562151929,82.43574562161929,82.44224485384348,82.44224485394348,82.44873923607913,82.44873923617914,82.45522956394389,82.45522956404389,82.46171917187363,82.46171917197363,82.46820783252045,82.46820783262045,82.47469262825297,82.47469262835297,82.48117223287508,82.48117223297508,82.48764998082268,82.48764998092268,82.4941242048778,82.49412420497781,82.50059774688924,82.50059774698924,82.50707060685697,82.50707060695697,82.52001500059636,82.52001500069636,82.53942816204126,82.53942816214126,82.54589389844143,82.54589389854144,82.55235880123266,82.55235880133266,82.57175176660573,82.57175176670573,82.58467607794722,82.58467607804722,82.59113101532213,82.59113101542214,82.59758220145667,82.59758220155668,82.60402831015477,82.60402831025478,82.61047180435202,82.61047180445202,82.61691344187474,82.61691344197475,82.623348562091,82.623348562191,82.62977947637107,82.62977947647107,82.6362039491273,82.63620394922731,82.64262213192495,82.64262213202495,82.64903716974332,82.64903716984333,82.65544891101712,82.65544891111712,82.66185845459455,82.66185845469455,82.66826534577982,82.66826534587982,82.68107571793182,82.68107571803182,82.68747908522462,82.68747908532463,82.69387938332076,82.69387938342076,82.70027805209028,82.70027805219028,82.70667562201162,82.70667562211162,82.71307137314975,82.71307137324975,82.71946329726488,82.71946329736488,82.72585389518395,82.72585389528395,82.73224229540662,82.73224229550662,82.74501807278642,82.74501807288642,82.75778649925078,82.75778649935079,82.76416652549246,82.76416652559246,82.77692354667047,82.77692354677048,82.78329701771435,82.78329701781435,82.78966503240862,82.78966503250862,82.80239587068675,82.80239587078675,82.81511287863427,82.81511287873427,82.8214663430629,82.8214663431629,82.82781874653466,82.82781874663466,82.83416713352685,82.83416713362685,82.84686193716274,82.84686193726274,82.85320706550168,82.85320706560168,82.86589167637338,82.86589167647338,82.87223089366691,82.87223089376691,82.87856783748143,82.87856783758143,82.89757260631431,82.89757260641431,82.9039022371047,82.9039022372047,82.91022561582783,82.91022561592783,82.9165475167896,82.91654751688961,82.92286354459728,82.92286354469728,82.92917843566545,82.92917843576545,82.93549165951568,82.93549165961568,82.94811496223686,82.94811496233686,82.95442280552011,82.95442280562011,82.96072576082351,82.96072576092351,82.96702322188597,82.96702322198597,82.97331996301341,82.97331996311341,82.98591170226773,82.98591170236773,82.99220404800242,82.99220404810242,83.00478582184043,83.00478582194043,83.01736354130755,83.01736354140755,83.02364732360466,83.02364732370467,83.0299302722928,83.0299302723928,83.0362051880218,83.0362051881218,83.0424790049026,83.0424790050026,83.04875104089152,83.04875104099152,83.05501758263951,83.05501758273951,83.06754877156965,83.06754877166965,83.07381330507785,83.07381330517785,83.08634055331105,83.08634055341105,83.09260243442708,83.09260243452708,83.09885965491114,83.09885965501114,83.10511543552516,83.10511543562517,83.11762180564283,83.11762180574283,83.12387106895038,83.12387106905038,83.13011699782204,83.13011699792204,83.13636194151948,83.13636194161948,83.14260559691215,83.14260559701215,83.14884591786894,83.14884591796894,83.15508464739042,83.15508464749043,83.16132080030238,83.16132080040238,83.1675548691919,83.1675548692919,83.17378526262371,83.17378526272371,83.18001395094626,83.18001395104626,83.18623900170239,83.18623900180239,83.19246265047981,83.19246265057981,83.19868538986563,83.19868538996563,83.20490559053323,83.20490559063323,83.21112431343947,83.21112431353947,83.21733799680058,83.21733799690058,83.22355084655271,83.22355084665271,83.23597434836063,83.23597434846063,83.24218431837271,83.24218431847271,83.24839345477581,83.24839345487581,83.260807180624,83.260807180724,83.27941757663207,83.27941757673207,83.28561803592363,83.28561803602364,83.29181766160623,83.29181766170623,83.29801626422326,83.29801626432327,83.30421134294785,83.30421134304785,83.3165949073079,83.3165949074079,83.3289720301441,83.3289720302441,83.33515032301537,83.33515032311537,83.34132679710342,83.34132679720342,83.35367372056017,83.35367372066017,83.359843828907,83.359843829007,83.36601215636196,83.36601215646196,83.37217809666396,83.37217809676396,83.37834274866118,83.37834274876118,83.38450573344048,83.38450573354048,83.39682844434581,83.39682844444582,83.41530637231051,83.41530637241051,83.42146283978332,83.42146283988332,83.42761680642921,83.42761680652922,83.43992034432827,83.43992034442827,83.44606965034222,83.44606965044223,83.45836640569557,83.45836640579557,83.47066096335257,83.47066096345257,83.47680674547405,83.47680674557405,83.48294987520336,83.48294987530336,83.48908807906149,83.48908807916149,83.50136338792956,83.50136338802956,83.50749833313445,83.50749833323445,83.5197667836742,83.5197667837742,83.52589831866058,83.52589831876058,83.532023639471,83.53202363957101,83.53814558795425,83.53814558805425,83.5503875903549,83.5503875904549,83.55650578759777,83.55650578769777,83.56874081799614,83.56874081809615,83.57485461984628,83.57485461994628,83.5870809731331,83.5870809732331,83.59319189524317,83.59319189534317,83.59929986183053,83.59929986193053,83.60540665378707,83.60540665388707,83.61761860837352,83.61761860847352,83.62371979241517,83.62371979251517,83.63592087219372,83.63592087229372,83.64201803975581,83.64201803985581,83.648111152947,83.648111153047,83.65420320518132,83.65420320528132,83.66028919480496,83.66028919490496,83.67245806696425,83.67245806706426,83.68462216481765,83.68462216491766,83.69678243564805,83.69678243574805,83.7028607712257,83.7028607713257,83.72109308767517,83.72109308777517,83.72716710364271,83.72716710374272,83.74538702963162,83.74538702973162,83.75145831742435,83.75145831752435,83.75752877160812,83.75752877170812,83.76359846796555,83.76359846806555,83.76966725493138,83.76966725503138,83.78180168388376,83.78180168398376,83.78786546919576,83.78786546929577,83.79392402550607,83.79392402560607,83.79998072514186,83.79998072524187,83.80603446925494,83.80603446935494,83.81208677349798,83.81208677359798,83.82419001789665,83.82419001799666,83.8302402760086,83.8302402761086,83.8362886395547,83.8362886396547,83.84233484329575,83.84233484339575,83.84837494653479,83.84837494663479,83.8544140645996,83.8544140646996,83.86649108820707,83.86649108830707,83.87856379220445,83.87856379230445,83.88459631718014,83.88459631728014,83.89062600030708,83.89062600040708,83.89665204586761,83.89665204596761,83.90266971744714,83.90266971754714,83.90868678276559,83.90868678286559,83.91470248399665,83.91470248409665,83.92071754107532,83.92071754117532,83.93274519229706,83.93274519239706,83.93875558874377,83.93875558884378,83.9507705463744,83.9507705464744,83.95677491810174,83.95677491820175,83.96277758471983,83.96277758481983,83.97478128862939,83.97478128872939,83.9807821743556,83.9807821744556,83.98677911938485,83.98677911948485,83.9987712664428,83.9987712665428,84.00476608955832,84.00476608965832,84.01075780558587,84.01075780568587,84.01674781650415,84.01674781660415,84.02872363240458,84.02872363250458,84.03470769438614,84.03470769448614,84.04068971023659,84.0406897103366,84.05265275676328,84.05265275686328,84.06461356770227,84.06461356780227,84.07657149890117,84.07657149900118,84.08254373879191,84.08254373889191,84.0885122653336,84.08851226543361,84.0944794656792,84.0944794657792,84.10044102021861,84.10044102031861,84.11236117377472,84.11236117387472,84.11831909074772,84.11831909084772,84.12427246076271,84.12427246086271,84.14213086569842,84.14213086579842,84.1540319599222,84.1540319600222,84.15997896419596,84.15997896429596,84.16592528642603,84.16592528652603,84.17186869102471,84.17186869112471,84.17780902642674,84.17780902652675,84.18968863627391,84.18968863637392,84.19562082504284,84.19562082514284,84.20155225598542,84.20155225608542,84.20748270175376,84.20748270185376,84.21341015410809,84.21341015420809,84.21933605291844,84.21933605301844,84.237112498936,84.237112499036,84.2489602800771,84.2489602801771,84.25488189716864,84.25488189726865,84.26080188493356,84.26080188503356,84.26671800778418,84.26671800788418,84.27263049306838,84.27263049316838,84.27854218263494,84.27854218273494,84.28444887054768,84.28444887064768,84.29035487641671,84.29035487651672,84.29625720682803,84.29625720692803,84.30215658171664,84.30215658181665,84.30805190223435,84.30805190233436,84.31984095183448,84.31984095193448,84.32573252111185,84.32573252121185,84.33162215793205,84.33162215803205,84.3375089907948,84.3375089908948,84.35516804951305,84.35516804961306,84.3610521163097,84.3610521164097,84.36693239397465,84.36693239407465,84.37281180013933,84.37281180023933,84.38456686122832,84.38456686132832,84.40219045367408,84.40219045377408,84.40806243314067,84.40806243324067,84.41393176021509,84.4139317603151,84.41979979898474,84.41979979908474,84.42566575373198,84.42566575383198,84.431529889696,84.431529889796,84.43739160061575,84.43739160071576,84.44325206112205,84.44325206122205,84.44911149856279,84.44911149866279,84.45496983715535,84.45496983725535,84.46668443031805,84.46668443041806,84.47253992706186,84.47253992716186,84.4783944007401,84.4783944008401,84.48424709352646,84.48424709362646,84.49009834644279,84.4900983465428,84.49594808370645,84.49594808380645,84.50179721470903,84.50179721480903,84.50764392066735,84.50764392076735,84.51348805001612,84.51348805011612,84.52517452782179,84.52517452792179,84.54854441423646,84.54854441433646,84.55438483023619,84.55438483033619,84.57190471414799,84.57190471424799,84.57774243986424,84.57774243996424,84.58357512603534,84.58357512613534,84.5894033789224,84.5894033790224,84.5952287899607,84.5952287900607,84.6068782479499,84.6068782480499,84.6243495551936,84.6243495552936,84.63017030559995,84.63017030569995,84.63598938878835,84.63598938888835,84.64180441760584,84.64180441770584,84.64761694559644,84.64761694569644,84.65923915972888,84.65923915982889,84.66504683763094,84.66504683773094,84.67085284831506,84.67085284841507,84.67665745702045,84.67665745712046,84.68246123211689,84.6824612322169,84.68826227903851,84.68826227913851,84.69405752858867,84.69405752868867,84.70564689094948,84.70564689104948,84.71144062484697,84.71144062494697,84.72302184057469,84.72302184067469,84.72880814777409,84.72880814787409,84.73459237095106,84.73459237105106,84.75772634602761,84.75772634612761,84.76350776524714,84.76350776534714,84.77506889857698,84.77506889867698,84.78084770329566,84.78084770339566,84.78662434820929,84.78662434830929,84.79239784814364,84.79239784824364,84.79817024922978,84.79817024932979,84.80394196827224,84.80394196837224,84.81548271607366,84.81548271617366,84.82125038074521,84.82125038084521,84.8270172118078,84.8270172119078,84.83277900332527,84.83277900342527,84.83853855925504,84.83853855935504,84.85005558709216,84.85005558719216,84.85581014136812,84.85581014146813,84.86156333155668,84.86156333165668,84.8673134904399,84.8673134905399,84.87881305037999,84.87881305047999,84.89030859384052,84.89030859394052,84.89605412998309,84.89605413008309,84.90179375508025,84.90179375518025,84.90753194030736,84.90753194040737,84.92474395727051,84.92474395737051,84.94195226088462,84.94195226098462,84.94768699800191,84.94768699810191,84.95341980266205,84.95341980276206,84.95915181160454,84.95915181170454,84.97061484431529,84.9706148444153,84.98207234489378,84.98207234499378,84.99352340394843,84.99352340404843,84.99924711469255,84.99924711479255,85.00496949924059,85.00496949934059,85.01069029235332,85.01069029245332,85.01640782681281,85.01640782691281,85.02212327724988,85.02212327734988,85.02783759094744,85.02783759104744,85.03354974483995,85.03354974493995,85.03925947368819,85.03925947378819,85.0506742707527,85.0506742708527,85.05637653501154,85.05637653511154,85.06777897950678,85.06777897960679,85.0791781653488,85.0791781654488,85.08487535217118,85.08487535227118,85.09057140225407,85.09057140235407,85.09626658083667,85.09626658093667,85.10765549813183,85.10765549823184,85.1133487821486,85.11334878224861,85.12473337983366,85.12473337993366,85.13042279893611,85.13042279903611,85.13610611753657,85.13610611763657,85.14178424502663,85.14178424512663,85.14746123577719,85.14746123587719,85.15313746870142,85.15313746880142,85.16448872202773,85.16448872212773,85.17016249201636,85.17016249211636,85.1985281211976,85.1985281212976,85.21554261072649,85.21554261082649,85.22121009075654,85.22121009085654,85.22687575200338,85.22687575210338,85.23253781357515,85.23253781367515,85.23819665438498,85.23819665448498,85.24385348695502,85.24385348705502,85.25516199887602,85.25516199897602,85.2664656985998,85.2664656986998,85.27776579864849,85.27776579874849,85.28906154119575,85.28906154129575,85.29470514969624,85.29470514979624,85.31727571878392,85.31727571888392,85.32855209886831,85.32855209896832,85.33418684080068,85.33418684090068,85.33981968816722,85.33981968826723,85.34545003470686,85.34545003480686,85.35670659763258,85.35670659773258,85.36233277612737,85.36233277622738,85.36795800733924,85.36795800743924,85.37920676465372,85.37920676475372,85.38482915401683,85.38482915411683,85.39045025507518,85.39045025517518,85.39607002993743,85.39607003003744,85.40730863237901,85.40730863247902,85.41292700526256,85.41292700536256,85.41854007336174,85.41854007346174,85.4241481019158,85.4241481020158,85.42975313705584,85.42975313715584,85.43535749015217,85.43535749025217,85.44095907718238,85.44095907728239,85.45215706013241,85.45215706023241,85.46334788162356,85.46334788172356,85.4689404884117,85.4689404885117,85.47453233737352,85.47453233747352,85.48571247351335,85.48571247361335,85.49129913136476,85.49129913146476,85.50246949154486,85.50246949164486,85.50805182978615,85.50805182988616,85.5192150285074,85.5192150286074,85.52479528272627,85.52479528282628,85.53037493068409,85.53037493078409,85.5471126999267,85.5471127000267,85.55268734623068,85.55268734633069,85.55826055266465,85.55826055276465,85.56383159929355,85.56383159939355,85.56940124394374,85.56940124404375,85.5749690319194,85.5749690320194,85.58053189402389,85.5805318941239,85.59165553421043,85.59165553431043,85.59721487242246,85.59721487252246,85.60833263945491,85.60833263955492,85.61388970418793,85.61388970428793,85.61944294189796,85.61944294199796,85.6249955354556,85.6249955355556,85.63054468090343,85.63054468100343,85.63609280328569,85.6360928033857,85.64163956158055,85.64163956168055,85.65273186564814,85.65273186574814,85.65827707039901,85.65827707049901,85.6693652822044,85.6693652823044,85.67490802401969,85.67490802411969,85.68598903647488,85.68598903657488,85.69152696609294,85.69152696619294,85.69706372108017,85.69706372118017,85.70259979402371,85.70259979412371,85.71366705193093,85.71366705203093,85.71919512980662,85.71919512990662,85.72472233618204,85.72472233628204,85.735773414497,85.735773414597,85.74129641493626,85.74129641503626,85.74681736924441,85.74681736934441,85.75233756572622,85.75233756582622,85.75785681492512,85.75785681502512,85.76889345664839,85.76889345674839,85.7744063779974,85.7744063780974,85.77991444925784,85.77991444935785,85.78541876927794,85.78541876937794,85.79642649992653,85.79642650002653,85.81293495092011,85.81293495102011,85.81843445874298,85.81843445884299,85.82393157941289,85.82393157951289,85.8294277906912,85.8294277907912,85.8349226378821,85.8349226379821,85.84591123341572,85.84591123351572,85.85689395579526,85.85689395589526,85.86237925437433,85.86237925447433,85.87883287663257,85.87883287673257,85.88431529547158,85.88431529557158,85.895276950279,85.895276950379,85.9062327319323,85.90623273203231,85.91170825455167,85.91170825465167,85.93360818522405,85.93360818532405,85.93907969136383,85.93907969146383,85.9500213774473,85.9500213775473,85.9664296816907,85.9664296817907,85.97736386529347,85.97736386539347,85.99375909703258,85.99375909713258,86.00468350467563,86.00468350477563,86.01560533570911,86.01560533580911,86.02106429982305,86.02106429992305,86.02651614036944,86.02651614046944,86.03741607022185,86.03741607032185,86.04286124189649,86.04286124199649,86.0537501074844,86.0537501075844,86.05919194472317,86.05919194482317,86.0700740656567,86.0700740657567,86.07551196219852,86.07551196229852,86.0809484188703,86.08094841897031,86.08637888871404,86.08637888881404,86.09180803236168,86.09180803246169,86.10266469033036,86.10266469043036,86.11894704383685,86.11894704393686,86.13522697229908,86.13522697239908,86.14065081116237,86.14065081126238,86.15149651854051,86.15149651864051,86.15691736398979,86.1569173640898,86.16233282887208,86.16233282897208,86.16774522455773,86.16774522465774,86.17315538465569,86.17315538475569,86.18397456811208,86.18397456821208,86.18938309888341,86.18938309898341,86.19478962141496,86.19478962151496,86.20019553768543,86.20019553778543,86.20559902891164,86.20559902901164,86.21099785950588,86.21099785960588,86.21639578070851,86.21639578080851,86.22179180734531,86.22179180744531,86.2271862046555,86.2271862047555,86.23257942733485,86.23257942743486,86.24336503908461,86.24336503918461,86.24875276752304,86.24875276762305,86.25413909398276,86.25413909408276,86.259523450094,86.259523450194,86.26490515381306,86.26490515391306,86.27566571940244,86.27566571950244,86.28104314140272,86.28104314150272,86.29717302025058,86.29717302035058,86.30254642577128,86.30254642587128,86.31329054652917,86.31329054662918,86.3293965917388,86.3293965918388,86.3347596150387,86.33475961513871,86.34011956914196,86.34011956924196,86.34547785602727,86.34547785612727,86.35083371786831,86.35083371796831,86.35618779881746,86.35618779891746,86.3615407051358,86.3615407052358,86.37224500211978,86.37224500221978,86.38294600255922,86.38294600265922,86.39364336543224,86.39364336553224,86.3989904364878,86.3989904365878,86.40433690128228,86.40433690138228,86.41502861834911,86.41502861844911,86.42037186238167,86.42037186248167,86.42571351497892,86.42571351507893,86.43105043116158,86.43105043126158,86.43638579380026,86.43638579390026,86.44705379090279,86.44705379100279,86.45238585699691,86.45238585709691,86.45771686213415,86.45771686223415,86.46304589692292,86.46304589702292,86.46837394653747,86.46837394663747,86.47902841643992,86.47902841653992,86.48435460937993,86.48435460947994,86.48967761944935,86.48967761954935,86.49499975801848,86.49499975811848,86.50031973678254,86.50031973688255,86.50563797254604,86.50563797264604,86.51095503367871,86.51095503377871,86.51627126120243,86.51627126130244,86.52690110174899,86.52690110184899,86.53221217605362,86.53221217615362,86.54283315003207,86.54283315013207,86.54814278446666,86.54814278456666,86.55876053768318,86.55876053778319,86.56406619352951,86.56406619362951,86.5746723141118,86.5746723142118,86.57997160421691,86.57997160431691,86.59586606431375,86.59586606441376,86.60645771041303,86.60645771051303,86.62234220509357,86.62234220519358,86.62763444741378,86.62763444751378,86.63292551510317,86.63292551520317,86.64350632428584,86.64350632438584,86.64879424699593,86.64879424709594,86.65936732635,86.65936732645,86.66993555561551,86.66993555571551,86.67521620319279,86.67521620329279,86.69105704707643,86.69105704717643,86.70689489754604,86.70689489764604,86.71745024376386,86.71745024386387,86.72800407432901,86.72800407442901,86.73327941712193,86.73327941722194,86.74382619990217,86.74382620000218,86.75436737476753,86.75436737486753,86.75963597290607,86.75963597300607,86.77017203244365,86.77017203254366,86.77543805397266,86.77543805407267,86.7806966866949,86.7806966867949,86.79121148920379,86.79121148930379,86.8017211384936,86.8017211385936,86.80697348125726,86.80697348135726,86.81222203488923,86.81222203498923,86.81746604156321,86.81746604166321,86.82795261504111,86.82795261514111,86.83319381775765,86.83319381785765,86.83843346693018,86.83843346703019,86.84367224460244,86.84367224470245,86.85938687250996,86.85938687260996,86.86462189894186,86.86462189904186,86.88032493210645,86.88032493220645,86.8855562451893,86.88555624528931,86.89078608051081,86.89078608061081,86.89601527167993,86.89601527177993,86.90124355345745,86.90124355355745,86.90647062271283,86.90647062281283,86.91169697203318,86.91169697213319,86.91692119943981,86.91692119953981,86.92214463112879,86.92214463122879,86.92736446314267,86.92736446324267,86.93258243848202,86.93258243858202,86.94301679772543,86.94301679782544,86.94823227223789,86.94823227233789,86.95344615531504,86.95344615541504,86.95865924267453,86.95865924277453,86.96908390174085,86.96908390184085,86.97950590841499,86.979505908515,86.995136379708,86.995136379808,87.00034484432685,87.00034484442685,87.00555209642357,87.00555209652357,87.0107587043679,87.0107587044679,87.01596288726796,87.01596288736796,87.02116623655904,87.02116623665904,87.03157134370592,87.03157134380592,87.03677079019138,87.03677079029138,87.04196913782866,87.04196913792866,87.05756258930519,87.0575625894052,87.06795579857861,87.06795579867861,87.07314785625731,87.07314785635731,87.07833919400097,87.07833919410098,87.08352912976594,87.08352912986594,87.09390650046895,87.09390650056895,87.09909207873248,87.09909207883248,87.10427504249515,87.10427504259515,87.10945645271384,87.10945645281384,87.11463665041039,87.11463665051039,87.12499632586847,87.12499632596847,87.13017394695548,87.13017394705548,87.14051929949584,87.14051929959584,87.1456850606007,87.1456850607007,87.1508482072047,87.1508482073047,87.1560083603947,87.1560083604947,87.16632559757801,87.16632559767801,87.17663790889014,87.17663790899014,87.18179177212154,87.18179177222154,87.18694180832996,87.18694180842996,87.19209120038599,87.19209120048599,87.197239872507,87.197239872607,87.20753505694395,87.20753505704396,87.2126754308666,87.2126754309666,87.2178132281797,87.2178132282797,87.22294871412247,87.22294871422247,87.22808287386917,87.22808287396917,87.23834945036197,87.23834945046197,87.24348084404254,87.24348084414254,87.25373980438069,87.25373980448069,87.2588668026685,87.25886680276851,87.26399232319497,87.26399232329497,87.2742412802255,87.2742412803255,87.28960973306312,87.28960973316312,87.29472650069542,87.29472650079542,87.30495901289446,87.30495901299446,87.31007115778611,87.31007115788611,87.32029374246017,87.32029374256017,87.3254038033294,87.3254038034294,87.33051098445856,87.33051098455856,87.34072254275944,87.34072254285944,87.34582347182133,87.34582347192134,87.35092299890451,87.35092299900451,87.35602116190027,87.35602116200027,87.36111720298231,87.36111720308232,87.36621210732484,87.36621210742484,87.37130591281918,87.37130591291918,87.37639736905189,87.37639736915189,87.38148674126217,87.38148674136217,87.38657452203715,87.38657452213715,87.39674542295515,87.39674542305515,87.40691033704509,87.40691033714509,87.41199076690462,87.41199076700462,87.42723035137395,87.42723035147395,87.43230896245028,87.43230896255028,87.44246262281915,87.44246262291915,87.45260878070731,87.45260878080731,87.45767882834605,87.45767882844605,87.46781695327505,87.46781695337505,87.47288260552104,87.47288260562104,87.48301288694748,87.48301288704748,87.48807709932343,87.48807709942344,87.49313960659013,87.49313960669014,87.50832557484625,87.50832557494626,87.51338535393815,87.51338535403815,87.52350373749111,87.52350373759111,87.52855980323397,87.52855980333398,87.53361458067205,87.53361458077205,87.53866772878351,87.53866772888351,87.54372027063391,87.54372027073391,87.55382338398623,87.55382338408623,87.56392312501136,87.56392312511136,87.56896805070708,87.56896805080709,87.57401206701121,87.57401206711121,87.57905388561896,87.57905388571896,87.58409464326984,87.58409464336984,87.58913475676835,87.58913475686835,87.59921403648242,87.59921403658242,87.60425214174113,87.60425214184113,87.60928922393428,87.60928922403428,87.61432532095321,87.61432532105322,87.61935952340629,87.61935952350629,87.62942701892086,87.62942701902087,87.63949182415193,87.63949182425193,87.64452242692992,87.64452242702993,87.64955219609895,87.64955219619895,87.65458060118057,87.65458060128057,87.66463547888665,87.66463547898665,87.67468349826441,87.67468349836442,87.6797029988866,87.6797029989866,87.68973866569507,87.68973866579508,87.69977058126321,87.69977058136321,87.70478500444895,87.70478500454895,87.70979825300387,87.70979825310387,87.71480991012348,87.71480991022348,87.71982012737305,87.71982012747306,87.72482666916491,87.72482666926491,87.72983040699931,87.72983040709931,87.73483315965949,87.73483315975949,87.73983242631853,87.73983242641853,87.74483074569464,87.74483074579464,87.74982671580912,87.74982671590912,87.76481227689091,87.76481227699091,87.77979408673234,87.77979408683234,87.78977738774107,87.78977738784107,87.79476596904877,87.79476596914877,87.79975273157328,87.79975273167328,87.80972345266484,87.80972345276484,87.8147040010134,87.8147040011134,87.81968201064373,87.81968201074373,87.82465839094745,87.82465839104745,87.83461005270668,87.83461005280668,87.84455974411745,87.84455974421745,87.8495328278766,87.8495328279766,87.85947785865541,87.85947785875541,87.86444923730531,87.86444923740531,87.86941993391152,87.86941993401152,87.87438358273279,87.87438358283279,87.89423503303863,87.89423503313863,87.90415683644027,87.90415683654027,87.90911605197748,87.90911605207748,87.91903319474716,87.91903319484716,87.92398896217455,87.92398896227455,87.93389970131169,87.93389970141169,87.93885399097773,87.93885399107774,87.94380593138214,87.94380593148215,87.94875446156803,87.94875446166803,87.95370098351414,87.95370098361414,87.95864511830729,87.95864511840729,87.96358724486066,87.96358724496066,87.9685252791518,87.9685252792518,87.97839907425511,87.97839907435511,87.98333309206669,87.98333309216669,87.99319919523269,87.99319919533269,87.99813086378263,87.99813086388264,88.00306173661491,88.00306173671491,88.00799185162087,88.00799185172087,88.01292132247443,88.01292132257443,88.01784988393639,88.0178498840364,88.02277753600676,88.02277753610676,88.0277031040547,88.0277031041547,88.0326267775368,88.0326267776368,88.04247166156541,88.04247166166542,88.05231226387524,88.05231226397524,88.05723036733377,88.05723036743377,88.06214593207409,88.06214593217409,88.06706039796623,88.06706039806623,88.07197425759729,88.07197425769729,88.07688698048885,88.07688698058885,88.08179784670588,88.08179784680588,88.08670598474812,88.08670598484812,88.09161078835447,88.09161078845447,88.10141592439182,88.10141592449182,88.1112132548179,88.1112132549179,88.11610885083428,88.11610885093428,88.12589890612755,88.12589890622755,88.1356857406589,88.1356857407589,88.14057235643321,88.14057235653321,88.14545749444616,88.14545749454616,88.15522652005863,88.15522652015864,88.1601055575696,88.1601055576696,88.16986052550355,88.16986052560355,88.17473584966547,88.17473584976547,88.18935814669351,88.18935814679351,88.19909954953606,88.19909954963606,88.20396809115228,88.20396809125228,88.21370396186258,88.21370396196258,88.22343786222442,88.22343786232442,88.22830318307871,88.22830318317871,88.24289531861861,88.24289531871861,88.24775389481852,88.24775389491852,88.25261000808284,88.25261000818284,88.26717565759232,88.26717565769232,88.27688235198872,88.27688235208872,88.28658635610164,88.28658635620164,88.29628706367,88.29628706377,88.30113563656228,88.30113563666228,88.30598216332346,88.30598216342347,88.31567408010633,88.31567408020634,88.32536247299673,88.32536247309673,88.33020475593044,88.33020475603044,88.33504393177617,88.33504393187617,88.34472110883682,88.34472110893682,88.3495572154859,88.3495572155859,88.35922810258796,88.35922810268796,88.3640623146712,88.3640623147712,88.36889323020988,88.36889323030988,88.38338362756429,88.38338362766429,88.38821128444972,88.38821128454973,88.40269228475756,88.40269228485757,88.41234225585278,88.41234225595278,88.41716513843231,88.41716513853231,88.42680912269948,88.42680912279948,88.43162984547395,88.43162984557395,88.4460895887531,88.4460895888531,88.45090777280934,88.45090777290935,88.45572455488687,88.45572455498687,88.46053739626745,88.46053739636746,88.46534705477744,88.46534705487744,88.47015584178713,88.47015584188713,88.47496227953519,88.47496227963519,88.47976754265243,88.47976754275243,88.48457038072539,88.48457038082539,88.48937018749301,88.48937018759301,88.49416904697772,88.49416904707772,88.49896646659238,88.49896646669238,88.50376256001095,88.50376256011096,88.5133537995652,88.5133537996652,88.52294189414015,88.52294189424015,88.52773302379623,88.52773302389623,88.53252328195204,88.53252328205204,88.54688981259196,88.54688981269196,88.55646244750967,88.55646244760968,88.56603341520945,88.56603341530945,88.57081639823242,88.57081639833243,88.57559869921171,88.57559869931171,88.58037478601504,88.58037478611504,88.58514962240491,88.58514962250491,88.58992343572925,88.58992343582925,88.59469486190062,88.59469486200062,88.60423585756884,88.60423585766884,88.60900440400013,88.60900440410013,88.61377185158325,88.61377185168325,88.6185375940571,88.6185375941571,88.62329985052982,88.62329985062982,88.6280608944804,88.6280608945804,88.63282076380015,88.63282076390016,88.63757669242298,88.63757669252298,88.64233193900209,88.64233193910209,88.6518380746589,88.6518380747589,88.65658782699708,88.65658782709708,88.66608289838939,88.66608289848939,88.67082806587825,88.67082806597826,88.68506091595265,88.68506091605265,88.68980316581012,88.68980316591012,88.69928546782869,88.69928546792869,88.70876223771499,88.708762237815,88.71349675774384,88.71349675784384,88.71823021681583,88.71823021691583,88.7276962634595,88.72769626355951,88.73716045342866,88.73716045352866,88.75135355551178,88.75135355561179,88.76081327430555,88.76081327440555,88.76554006450579,88.76554006460579,88.77499220503621,88.77499220513621,88.77971490297423,88.77971490307424,88.79388098854851,88.79388098864851,88.79860095831172,88.79860095841173,88.80803816966333,88.80803816976334,88.8127541987296,88.81275419882961,88.81746863636057,88.81746863646057,88.82689641277433,88.82689641287433,88.83160808433917,88.83160808443917,88.83631770977291,88.83631770987292,88.84573624070538,88.84573624080538,88.85514601874367,88.85514601884367,88.85984514828266,88.85984514838266,88.86924242218642,88.86924242228642,88.87393950559431,88.87393950569431,88.87863446708846,88.87863446718846,88.88802238183699,88.88802238193699,88.90210055955639,88.90210055965639,88.91617388718724,88.91617388728724,88.92086180089647,88.92086180099648,88.92554679697432,88.92554679707432,88.93023099733452,88.93023099743452,88.93959397959661,88.93959397969661,88.94427162475901,88.94427162485901,88.96298027295144,88.96298027305144,88.97233260775351,88.97233260785352,88.97700638800161,88.97700638810161,88.98167884205361,88.98167884215361,88.98634959099635,88.98634959109636,88.9910173844164,88.9910173845164,88.99568453368404,88.99568453378404,89.00034929579874,89.00034929589874,89.00967745594073,89.00967745604073,89.01433994457643,89.01433994467644,89.02366325462988,89.02366325472988,89.0283228635255,89.0283228636255,89.03298000948553,89.03298000958553,89.0376343135968,89.0376343136968,89.04228793566438,89.04228793576438,89.04693962527479,89.04693962537479,89.0515865405793,89.0515865406793,89.0562320917964,89.0562320918964,89.06087552109976,89.06087552119976,89.06551807890284,89.06551807900284,89.07015889370535,89.07015889380536,89.07479891279021,89.07479891289022,89.07943821194004,89.07943821204005,89.08871393049965,89.08871393059965,89.09334981943097,89.09334981953097,89.10261955116253,89.10261955126253,89.10725293926696,89.10725293936696,89.1118852664145,89.1118852665145,89.12114779879587,89.12114779889588,89.1257770946381,89.1257770947381,89.13040400332734,89.13040400342734,89.13502928268998,89.13502928278999,89.14427669643597,89.14427669653597,89.14889818666693,89.14889818676693,89.15813938623697,89.15813938633697,89.16737706191455,89.16737706201455,89.1719940051875,89.1719940052875,89.18122656553732,89.18122656563732,89.18584206894022,89.18584206904022,89.19045329062435,89.19045329072435,89.19506193569892,89.19506193579892,89.20427824067384,89.20427824077385,89.20888241457305,89.20888241467306,89.21348590642856,89.21348590652856,89.21808788263138,89.21808788273138,89.22729043305833,89.22729043315833,89.23189036313005,89.23189036323005,89.24108722985949,89.24108722995949,89.24568356025613,89.24568356035613,89.25027833710877,89.25027833720877,89.25487110572162,89.25487110582162,89.2594614113989,89.2594614114989,89.26404986040164,89.26404986050164,89.2686374379041,89.2686374380041,89.27322414390629,89.27322414400629,89.28239346364843,89.28239346374843,89.28697206090882,89.28697206100883,89.29154967299498,89.29154967309498,89.29612580731978,89.29612580741978,89.30527659820804,89.30527659830804,89.30984943598828,89.30984943608829,89.31442106124639,89.3144210613464,89.33270517512587,89.33270517522587,89.33727388275258,89.33727388285259,89.3464103128318,89.3464103129318,89.35554397684487,89.35554397694487,89.37836845564613,89.37836845574613,89.3829243938991,89.3829243939991,89.3874785891515,89.3874785892515,89.39658599448207,89.39658599458207,89.40569161892074,89.40569161902074,89.41934817995858,89.41934818005858,89.43299917097285,89.43299917107285,89.43754628054901,89.43754628064902,89.44663940085316,89.44663940095316,89.45118347912398,89.45118347922399,89.46481453930565,89.46481453940565,89.47389985399855,89.47389985409855,89.47844120409457,89.47844120419457,89.4875211761118,89.4875211762118,89.4965972453234,89.4965972454234,89.5011333664177,89.5011333665177,89.5056685781204,89.50566857822041,89.5102014784528,89.5102014785528,89.52379820910147,89.52379820920147,89.53285991750403,89.53285991760403,89.54191973134074,89.54191973144074,89.55097746115504,89.55097746125504,89.56003121238108,89.56003121248108,89.56908056821436,89.56908056831436,89.57812802948182,89.57812802958182,89.58265069915868,89.58265069925868,89.59169418183788,89.59169418193788,89.60525540832275,89.60525540842275,89.60977455410715,89.60977455420715,89.61429252526072,89.61429252536072,89.62332733082839,89.62332733092839,89.63236039339547,89.63236039349547,89.64590646335364,89.64590646345364,89.6504198875492,89.6504198876492,89.65493270548372,89.65493270558372,89.65944408354818,89.65944408364818,89.67297620950178,89.67297620960179,89.6819941913248,89.6819941914248,89.68650147712705,89.68650147722705,89.69551468464415,89.69551468474415,89.70001962118478,89.70001962128478,89.70902706922173,89.70902706932173,89.72253305016677,89.72253305026678,89.72703143150959,89.72703143160959,89.73152769093866,89.73152769103866,89.73602319254141,89.73602319264141,89.7405178984265,89.7405178985265,89.74950526406558,89.74950526416558,89.77196742609601,89.77196742619601,89.77645735767518,89.77645735777519,89.78094637986277,89.78094637996277,89.78543472000663,89.78543472010664,89.789919649932,89.789919650032,89.79440268529153,89.79440268539153,89.7988846596942,89.7988846597942,89.8033651563355,89.8033651564355,89.80784375841097,89.80784375851097,89.81232050381192,89.81232050391192,89.82127278209168,89.82127278219168,89.83022274900112,89.83022274910113,89.8346966525533,89.8346966526533,89.84364014004758,89.84364014014758,89.84810961031572,89.84810961041572,89.85704669417746,89.85704669427746,89.86151157959625,89.86151157969626,89.86597547984081,89.86597547994081,89.87043869804168,89.87043869814168,89.87490055215515,89.87490055225516,89.88381997866328,89.88381997876328,89.8971920516949,89.8971920517949,89.9016483357848,89.9016483358848,89.91055844102901,89.91055844112901,89.92391880564375,89.92391880574375,89.92837042910169,89.92837042920169,89.93727219825621,89.93727219835621,89.94172185136567,89.94172185146567,89.9506202103017,89.9506202104017,89.95951724304162,89.95951724314162,89.96840976671484,89.96840976681484,89.9728525993873,89.9728525994873,89.97729448477683,89.97729448487684,89.9817344377092,89.98173443780921,89.99949171072049,89.9994917108205,90.00392897336937,90.00392897346937,90.00836547819192,90.00836547829192,90.01723719953226,90.01723719963226,90.02610698841544,90.02610698851544,90.03054087873713,90.03054087883713,90.03497260925377,90.03497260935377,90.0394015737043,90.0394015738043,90.04825609238681,90.04825609248681,90.0526800172922,90.0526800173922,90.0615265409069,90.0615265410069,90.06594762396352,90.06594762406353,90.07036370536635,90.07036370546635,90.07477861213835,90.07477861223835,90.07919215482293,90.07919215492294,90.09243107776747,90.09243107786747,90.09684227119043,90.09684227129043,90.11448299051136,90.11448299061136,90.12329616976731,90.12329616986732,90.13651110750834,90.13651110760834,90.1453189061973,90.1453189062973,90.14972009631263,90.14972009641264,90.15411999812319,90.1541199982232,90.16291760404795,90.16291760414795,90.16731489135766,90.16731489145766,90.17171142084104,90.17171142094104,90.18050178952429,90.18050178962429,90.18928806594533,90.18928806604534,90.19807085636523,90.19807085646524,90.20684955452293,90.20684955462293,90.21123661117711,90.21123661127712,90.22000939828939,90.2200093983894,90.22439463616037,90.22439463626037,90.23754837942484,90.23754837952484,90.24192607692378,90.24192607702378,90.2463007810087,90.2463007811087,90.25504901454772,90.25504901464772,90.26379569454275,90.26379569464275,90.26816812514866,90.26816812524866,90.27253888853664,90.27253888863665,90.28564981461317,90.28564981471317,90.29001872132663,90.29001872142663,90.29438671864848,90.29438671874848,90.29875361712214,90.29875361722215,90.30748604998207,90.30748605008208,90.31185105388991,90.31185105398991,90.31621537575404,90.31621537585404,90.3205784093134,90.3205784094134,90.32493920728507,90.32493920738507,90.33365826451019,90.3336582646102,90.34237387362548,90.34237387372548,90.35108838389259,90.35108838399259,90.35980096170253,90.35980096180253,90.36851209964244,90.36851209974245,90.37721994103781,90.37721994113781,90.38157225135453,90.38157225145453,90.39027524266133,90.39027524276133,90.39462527949905,90.39462527959905,90.39897421748857,90.39897421758857,90.40332103356435,90.40332103366435,90.41635951144325,90.41635951154325,90.42070367512686,90.42070367522686,90.42504689152756,90.42504689162756,90.42938737975344,90.42938737985344,90.43372051706388,90.43372051716388,90.43805297233061,90.43805297243061,90.44671621564615,90.44671621574615,90.45104635954256,90.45104635964256,90.45537472254709,90.4553747226471,90.46403012236004,90.46403012246004,90.47268283188953,90.47268283198953,90.47700710263184,90.47700710273185,90.48565378744195,90.48565378754195,90.49429850190359,90.49429850200359,90.50294143547335,90.50294143557335,90.5115821334554,90.51158213355541,90.51589992478256,90.51589992488256,90.52021692039204,90.52021692049205,90.52453262769679,90.52453262779679,90.53316286767541,90.53316286777542,90.53747656674035,90.53747656684035,90.54178920484843,90.54178920494843,90.54610078199963,90.54610078209963,90.55041148765055,90.55041148775055,90.55903145908235,90.55903145918235,90.56333970179767,90.56333970189768,90.56764608783848,90.56764608793848,90.57195080666135,90.57195080676135,90.57625408561418,90.57625408571418,90.58055448482693,90.58055448492694,90.59345435626912,90.59345435636912,90.59775176206786,90.59775176216786,90.60634350446867,90.60634350456867,90.6106375000489,90.6106375001489,90.61492835064983,90.61492835074984,90.6192185949897,90.6192185950897,90.62350505019789,90.6235050502979,90.6277908612537,90.6277908613537,90.63636157397372,90.63636157407372,90.64493073314974,90.64493073324974,90.64921446018312,90.64921446028312,90.65349712625964,90.65349712635964,90.65777827668349,90.65777827678349,90.66633853140007,90.66633853150007,90.67489624739844,90.67489624749844,90.68773018794947,90.68773018804947,90.69200629882818,90.69200629892818,90.69628066670633,90.69628066680633,90.70055382206233,90.70055382216233,90.70909861712168,90.70909861722168,90.71336968845527,90.71336968855528,90.72191114907875,90.72191114917875,90.7304512077235,90.7304512078235,90.7347193614257,90.7347193615257,90.73898459749651,90.73898459759651,90.74324774954489,90.7432477496449,90.75603531112422,90.75603531122422,90.76455615887137,90.76455615897137,90.76881453661385,90.76881453671385,90.77733030692455,90.77733030702456,90.7858429701473,90.7858429702473,90.79009771032337,90.79009771042337,90.7986054097836,90.7986054098836,90.80711083576485,90.80711083586485,90.81561326833206,90.81561326843206,90.81986179433218,90.81986179443219,90.82410967617992,90.82410967627992,90.82835687598394,90.82835687608394,90.83260070346078,90.83260070356079,90.83684392467656,90.83684392477656,90.84957059490985,90.84957059500985,90.85381055747237,90.85381055757237,90.86228786809657,90.86228786819657,90.86652483724508,90.86652483734508,90.87076040441487,90.87076040451487,90.87499528954096,90.87499528964096,90.88346327890125,90.88346327900125,90.88769600422226,90.88769600432227,90.90038982268393,90.90038982278394,90.91307469879484,90.91307469889485,90.91730124783123,90.91730124793123,90.93420198762722,90.93420198772722,90.94264314993524,90.94264315003524,90.94686193125169,90.94686193135169,90.95951751737475,90.95951751747475,90.96795223815892,90.96795223825892,90.97216802606135,90.97216802616136,90.97638222252849,90.97638222262849,90.98902208375509,90.98902208385509,90.99744441407869,90.99744441417869,91.00165432882702,91.00165432892702,91.00586189431372,91.00586189441373,91.01427562330841,91.01427562340841,91.01848140790322,91.01848140800323,91.02689062783122,91.02689062793122,91.03529564182304,91.03529564192304,91.0436968666832,91.04369686678321,91.05209490867274,91.05209490877274,91.06468645847048,91.06468645857048,91.06888038308315,91.06888038318316,91.0772669061124,91.0772669062124,91.08145848146344,91.08145848156344,91.08983640316379,91.08983640326379,91.09402244638257,91.09402244648257,91.10239290349352,91.10239290359353,91.11075782847222,91.11075782857222,91.11911741077526,91.11911741087526,91.12329587573069,91.12329587583069,91.13165079740178,91.13165079750178,91.14000321824594,91.14000321834594,91.15670101214937,91.15670101224937,91.16504373281644,91.16504373291644,91.1692127249427,91.1692127250427,91.17338058032944,91.17338058042944,91.17754756421589,91.17754756431589,91.18171386605864,91.18171386615865,91.18587842490082,91.18587842500082,91.19004192278614,91.19004192288614,91.19836774392594,91.19836774402594,91.2025286273104,91.2025286274104,91.20668685830269,91.20668685840269,91.21915973249634,91.21915973259634,91.22331504585723,91.22331504595724,91.23993485943078,91.23993485953078,91.25239647990334,91.25239648000334,91.25654857250231,91.25654857260231,91.26069759590463,91.26069759600463,91.26899249772995,91.26899249782996,91.2772840272281,91.27728402732811,91.2814285036724,91.2814285037724,91.28557169181194,91.28557169191194,91.28971336429879,91.28971336439879,91.29385253595875,91.29385253605875,91.29799110135764,91.29799110145764,91.30626675439406,91.30626675449406,91.31040353890106,91.31040353900106,91.31867494811,91.31867494821,91.32280851185507,91.32280851195507,91.3269403704909,91.3269403705909,91.33520295102304,91.33520295112304,91.33933310454961,91.33933310464961,91.34759238853718,91.34759238863718,91.35997898420258,91.35997898430259,91.36410652322829,91.36410652332829,91.36823220557947,91.36823220567948,91.3764800463894,91.3764800464894,91.38060084076072,91.38060084086072,91.38472027104466,91.38472027114466,91.38883909506751,91.38883909516751,91.39707253717707,91.39707253727707,91.40118582906767,91.40118582916767,91.40940722173849,91.40940722183849,91.41762258968996,91.41762258978996,91.42583560837977,91.42583560847977,91.42994094309387,91.42994094319387,91.43815024843465,91.43815024853465,91.44635769710091,91.44635769720091,91.45045950792253,91.45045950802253,91.46276206064736,91.46276206074737,91.46686084016365,91.46686084026365,91.47095889974493,91.47095889984493,91.47505392802086,91.47505392812086,91.47914819847045,91.47914819857046,91.48324186265899,91.48324186275899,91.49142831953576,91.49142831963576,91.50370563664362,91.50370563674362,91.50779657265734,91.50779657275734,91.5118866750621,91.5118866751621,91.51597617120578,91.51597617130578,91.52006460639261,91.52006460649261,91.52823965798304,91.52823965808304,91.54049814310646,91.54049814320646,91.54866542697697,91.54866542707697,91.5527468901615,91.5527468902615,91.56090818720394,91.56090818730394,91.5649858612568,91.5649858613568,91.57313961792721,91.57313961802721,91.58129011594438,91.58129011604439,91.58536331882186,91.58536331892186,91.5894355365251,91.5894355366251,91.59350593544515,91.59350593554515,91.59757550075622,91.59757550085622,91.60571368409545,91.60571368419545,91.60978124116673,91.60978124126673,91.61384678999822,91.61384679009822,91.61790998956808,91.61790998966808,91.62197046096313,91.62197046106313,91.62603013664052,91.62603013674052,91.630088865035,91.63008886513501,91.63820514719315,91.63820514729315,91.65037629283145,91.65037629293145,91.65848840694476,91.65848840704476,91.66254338409887,91.66254338419887,91.6665977171006,91.6665977172006,91.67064958716675,91.67064958726675,91.68279830114554,91.68279830124554,91.6908861648161,91.69088616491611,91.69897061826818,91.69897061836818,91.70705185095831,91.70705185105831,91.71109064852018,91.71109064862019,91.71512584640695,91.71512584650695,91.72722950761012,91.72722950771012,91.73126148473496,91.73126148483496,91.73529084735895,91.73529084745896,91.73931744391682,91.73931744401682,91.74334244903939,91.74334244913939,91.75541325847092,91.75541325857093,91.76747137431134,91.76747137441134,91.77550221155019,91.77550221165019,91.78353009326634,91.78353009336634,91.79155475422056,91.79155475432056,91.80759676310487,91.80759676320487,91.81160509604805,91.81160509614806,91.81961998104255,91.81961998114255,91.82362456274538,91.82362456284538,91.8276279698174,91.8276279699174,91.83162989912806,91.83162989922806,91.83563095693843,91.83563095703843,91.8436315569065,91.8436315570065,91.84763053069446,91.84763053079446,91.85162571535074,91.85162571545074,91.85561976326751,91.85561976336751,91.86360645712234,91.86360645722235,91.87558185621828,91.87558185631828,91.87957022043754,91.87957022053754,91.88754353865755,91.88754353875755,91.89152769694066,91.89152769704066,91.90347725415856,91.90347725425856,91.9074569791576,91.9074569792576,91.91143590843899,91.91143590853899,91.91939217556646,91.91939217566646,91.92336826299909,91.92336826309909,91.93131884642905,91.93131884652905,91.93529277405663,91.93529277415664,91.94721288972141,91.94721288982142,91.9511850364571,91.9511850365571,91.95912830686294,91.95912830696294,91.97103777506771,91.97103777516772,91.98690480185122,91.98690480195123,91.99086774099695,91.99086774109695,91.9948296570771,91.9948296571771,91.99878983015671,91.99878983025671,92.00274898017075,92.00274898027075,92.00670600827105,92.00670600837105,92.01066216487106,92.01066216497107,92.01857322765764,92.01857322775764,92.0264819790739,92.02648197917391,92.03438830544589,92.03438830554589,92.03833870256575,92.03833870266575,92.04228686409793,92.04228686419793,92.05412915099811,92.05412915109811,92.06202066186525,92.06202066196525,92.06596503426574,92.06596503436575,92.07385131613117,92.07385131623117,92.08173430145203,92.08173430155203,92.08567465737296,92.08567465747296,92.08961425546755,92.08961425556755,92.09355298206187,92.09355298216187,92.09748988987296,92.09748988997296,92.10142558516193,92.10142558526194,92.10535980268955,92.10535980278955,92.11322645685289,92.11322645695289,92.12109072386328,92.12109072396328,92.1250213796071,92.1250213797071,92.12895139119857,92.12895139129857,92.13288072074631,92.13288072084632,92.14073812942837,92.14073812952837,92.15252123009176,92.15252123019177,92.15644699785574,92.15644699795574,92.1603715909889,92.1603715910889,92.16429535051309,92.16429535061309,92.17606488608509,92.17606488618509,92.18783055674277,92.18783055684277,92.19175105761371,92.19175105771372,92.1956702701799,92.1956702702799,92.19958857335448,92.19958857345448,92.2035056261156,92.2035056262156,92.20741976124533,92.20741976134534,92.21133230493976,92.21133230503976,92.21524337087283,92.21524337097283,92.219153489523,92.219153489623,92.2230625851076,92.2230625852076,92.23478835620874,92.23478835630874,92.24260340239864,92.24260340249864,92.24650961824318,92.24650961834318,92.25431985223587,92.25431985233587,92.25822311255769,92.25822311265769,92.26993005167441,92.26993005177441,92.27383001545167,92.27383001555167,92.29332646201081,92.29332646211081,92.30112002382413,92.30112002392413,92.31280805517378,92.31280805527378,92.32059858568175,92.32059858578175,92.32449256263097,92.32449256273097,92.33227812937645,92.33227812947645,92.33616714256317,92.33616714266317,92.34783281803593,92.34783281813593,92.3517194440697,92.3517194441697,92.35949118048455,92.35949118058456,92.36337613930039,92.36337613940039,92.37502984111705,92.37502984121706,92.37891275380179,92.37891275390179,92.38279464342095,92.38279464352095,92.39055709646321,92.39055709656321,92.39831727602646,92.39831727612646,92.40219442923105,92.40219442933105,92.40606574717283,92.40606574727283,92.4138071705343,92.4138071706343,92.41767704860605,92.41767704870605,92.42154571415567,92.42154571425567,92.43315049828242,92.43315049838242,92.43701787552726,92.43701787562726,92.44088460861973,92.44088460871973,92.44474690842814,92.44474690852815,92.44860746523598,92.44860746533598,92.46018743055023,92.46018743065024,92.46404575177039,92.46404575187039,92.4679030878163,92.4679030879163,92.47175636949132,92.47175636959132,92.47946156875396,92.47946156885396,92.48331314531973,92.48331314541973,92.49101402497224,92.49101402507225,92.49486211553688,92.49486211563688,92.50255647788292,92.50255647798292,92.50640248442512,92.50640248452513,92.51024746790176,92.51024746800177,92.52178037220061,92.52178037230061,92.52562304430694,92.52562304440694,92.52946423865191,92.52946423875191,92.54098619236022,92.54098619246022,92.54482348389956,92.54482348399956,92.55249670289086,92.55249670299087,92.55633164516857,92.55633164526857,92.56016582961995,92.56016582971995,92.5639990667884,92.5639990668884,92.56783109143473,92.56783109153473,92.57549127581305,92.57549127591305,92.57931761676186,92.57931761686186,92.58696885878943,92.58696885888943,92.59079315360712,92.59079315370712,92.59461612222874,92.59461612232874,92.59843799200218,92.59843799210218,92.60608063270084,92.60608063280084,92.61372115148578,92.61372115158578,92.62517876573823,92.62517876583823,92.63281401763012,92.63281401773013,92.63662914274917,92.63662914284917,92.64425761209537,92.64425761219537,92.64806970590908,92.64806970600908,92.65569196107933,92.65569196117933,92.65949814384763,92.65949814394763,92.66330341722431,92.66330341732431,92.66710399207771,92.66710399217772,92.67090278603925,92.67090278613925,92.67470086006574,92.67470086016574,92.67849597856954,92.67849597866955,92.68228973298594,92.68228973308594,92.6898744757526,92.6898744758526,92.69366421368942,92.69366421378942,92.69745334536518,92.69745334546518,92.70881880793526,92.70881880803526,92.7163924106548,92.7163924107548,92.72017684380728,92.72017684390728,92.7239600265463,92.7239600266463,92.72774086002367,92.72774086012367,92.73530127656498,92.73530127666498,92.7466403667786,92.74664036687861,92.75419816881906,92.75419816891906,92.75797600888241,92.75797600898241,92.76175225751047,92.76175225761047,92.76552736939902,92.76552736949903,92.77307554704501,92.77307554714501,92.77684652878003,92.77684652888003,92.78061508547079,92.78061508557079,92.78438292222651,92.78438292232651,92.79191692852004,92.79191692862004,92.79567927103484,92.79567927113484,92.79944077994068,92.79944078004068,92.80319952278039,92.80319952288039,92.80695758357639,92.8069575836764,92.81071079428386,92.81071079438387,92.81821615474192,92.81821615484192,92.82572018900389,92.82572018910389,92.82947051997128,92.82947052007128,92.83696595290435,92.83696595300435,92.84070934976077,92.84070934986077,92.84445183722559,92.8444518373256,92.85193465235018,92.85193465245018,92.85567282020489,92.85567282030489,92.85940947240692,92.85940947250693,92.86688054122332,92.86688054132333,92.87061454103319,92.8706145411332,92.87434725253829,92.87434725263829,92.87807916832573,92.87807916842573,92.88554102955216,92.88554102965216,92.89299944266875,92.89299944276875,92.89672711462872,92.89672711472872,92.90045395297972,92.90045395307972,92.90417961669989,92.9041796167999,92.91162931481364,92.91162931491364,92.91535232614164,92.91535232624165,92.91907435229541,92.91907435239541,92.92651628268922,92.92651628278922,92.93395518177769,92.93395518187769,92.94138915499498,92.94138915509498,92.94881698981895,92.94881698991895,92.9525281032735,92.9525281033735,92.95994578322457,92.95994578332457,92.96735823417394,92.96735823427395,92.97106309556122,92.97106309566122,92.97476674442636,92.97476674452636,92.97846857450831,92.97846857460831,92.98216976043787,92.98216976053787,92.98956951779611,92.98956951789611,92.99326763452902,92.99326763462902,92.99696397037003,92.99696397047003,93.00435520218201,93.00435520228201,93.00804918876138,93.00804918886138,93.01174253118837,93.01174253128838,93.01543526735429,93.0154352674543,93.02650964882905,93.02650964892905,93.03757785401918,93.03757785411918,93.041264717031,93.041264717131,93.04495048119463,93.04495048129463,93.05231924345578,93.05231924355579,93.05600163529223,93.05600163539224,93.0633650548777,93.06336505497771,93.0670452869091,93.06704528700911,93.0707246474402,93.0707246475402,93.0780821938716,93.0780821939716,93.08175988718476,93.08175988728476,93.08543583749736,93.08543583759736,93.08910959011358,93.08910959021358,93.09278190285976,93.09278190295976,93.10012376228602,93.10012376238602,93.1037895198344,93.1037895199344,93.1111199739743,93.1111199740743,93.11478360960895,93.11478360970895,93.11844470652538,93.11844470662538,93.12210383309333,93.12210383319334,93.12941939594575,93.12941939604575,93.13307507440388,93.13307507450388,93.13672972979646,93.13672972989646,93.14403657764602,93.14403657774602,93.14768687553718,93.14768687563718,93.15498663771051,93.15498663781051,93.1586350031445,93.1586350032445,93.1659301425772,93.1659301426772,93.16957596929296,93.16957596939297,93.17686497033233,93.17686497043233,93.18415006856608,93.18415006866609,93.19507212694748,93.19507212704748,93.20234816915652,93.20234816925652,93.20962171053866,93.20962171063866,93.21325707925101,93.21325707935101,93.22052376230481,93.22052376240481,93.23141823580761,93.23141823590761,93.23504632938713,93.23504632948713,93.24592924603832,93.24592924613832,93.24955571029123,93.24955571039123,93.25318149250045,93.25318149260045,93.25680299299086,93.25680299309086,93.26404224273132,93.26404224283132,93.26765828687212,93.26765828697212,93.27850429738079,93.2785042974808,93.28211844695575,93.28211844705575,93.28934595038803,93.28934595048803,93.29295676552712,93.29295676562712,93.29656697440514,93.29656697450514,93.30739593382127,93.30739593392127,93.31100303561131,93.31100303571131,93.31821572353873,93.31821572363873,93.32181941511027,93.32181941521027,93.32542204572493,93.32542204582494,93.32902122822979,93.3290212283298,93.33981635070005,93.33981635080005,93.34341200931243,93.34341200941243,93.35060169721058,93.35060169731058,93.35778979367343,93.35778979377343,93.36138279989365,93.36138279999365,93.3649751998528,93.3649751999528,93.37574970944674,93.37574970954674,93.37933915388317,93.37933915398317,93.3936953023023,93.3936953024023,93.39728277639027,93.39728277649027,93.40086797699922,93.40086797709922,93.40445245767317,93.40445245777317,93.41161997915101,93.41161997925101,93.41520142851962,93.41520142861962,93.42594365471169,93.42594365481169,93.43310249907799,93.43310249917799,93.44741374628677,93.44741374638677,93.45813846669049,93.45813846679049,93.46171184520863,93.46171184530863,93.46885757917934,93.46885757927934,93.47600210062791,93.47600210072791,93.4831446138367,93.4831446139367,93.4902851188057,93.4902851189057,93.49385213158261,93.49385213168262,93.50098403522271,93.50098403532272,93.50454407599734,93.50454407609735,93.51522325103834,93.51522325113834,93.52234045284753,93.52234045294753,93.5258969697297,93.5258969698297,93.52945234987237,93.52945234997237,93.53300341040493,93.53300341050493,93.53655359943721,93.53655359953721,93.54010265172998,93.54010265182998,93.55074757302062,93.55074757312062,93.56138790946193,93.56138790956193,93.56493142962245,93.56493142972245,93.5684727520866,93.56847275218661,93.5720134682897,93.5720134683897,93.57555342666645,93.57555342676645,93.57909232408633,93.57909232418633,93.58616951266502,93.58616951276503,93.59324480667789,93.59324480677789,93.59677756570446,93.59677756580446,93.60384118919175,93.60384118929176,93.61090155402582,93.61090155412582,93.61795638672763,93.61795638682763,93.62500534627534,93.62500534637535,93.6285281398856,93.6285281399856,93.63205028934348,93.63205028944348,93.6390909128015,93.63909091290151,93.65316450567157,93.65316450577157,93.66723245273545,93.66723245283545,93.67074645556022,93.67074645566022,93.67425943531944,93.67425943541944,93.67777177092627,93.67777177102627,93.68479515383518,93.68479515393518,93.69532768948031,93.69532768958031,93.70585643599377,93.70585643609377,93.7093647172297,93.7093647173297,93.71287076287796,93.71287076297796,93.723387497844,93.723387497944,93.72688945123005,93.72688945133005,93.73039007842,93.73039007852,93.733888318457,93.733888318557,93.74438152291535,93.74438152301535,93.75137466855348,93.75137466865348,93.75836538914733,93.75836538924733,93.76535213115292,93.76535213125293,93.76884176986103,93.76884176996103,93.77233080230806,93.77233080240806,93.77581794018926,93.77581794028926,93.78279035927712,93.78279035937712,93.78627461741823,93.78627461751823,93.78975811773302,93.78975811783302,93.79324097389542,93.79324097399542,93.800204867437,93.800204867537,93.8036847680767,93.80368476817671,93.8141222722994,93.8141222723994,93.82107820866447,93.82107820876448,93.82455534323805,93.82455534333805,93.82803160631133,93.82803160641133,93.83150673264511,93.83150673274511,93.8349803054349,93.8349803055349,93.83845251413729,93.83845251423729,93.84192411657861,93.84192411667861,93.84539412758463,93.84539412768463,93.85233305074848,93.85233305084849,93.86620835835797,93.86620835845797,93.87314292402039,93.87314292412039,93.88700940295305,93.88700940305306,93.90433710210625,93.90433710220626,93.9078012778495,93.9078012779495,93.91472815157462,93.91472815167462,93.91819012962148,93.91819012972148,93.92165142562465,93.92165142572465,93.9251120395841,93.92511203968411,93.92857185782592,93.92857185792592,93.93203053932822,93.93203053942823,93.93894562885383,93.93894562895383,93.94931144270524,93.94931144280524,93.95276152287863,93.95276152297863,93.95966012968145,93.95966012978145,93.96655680402712,93.96655680412712,93.97000328452543,93.97000328462543,93.97344885563214,93.97344885573214,93.9837838259517,93.9837838260517,93.98722670677492,93.98722670687492,93.99755371991796,93.99755372001796,94.00099478195797,94.00099478205797,94.01131592194692,94.01131592204692,94.01475383900764,94.01475383910764,94.0216265660411,94.02162656614111,94.02505989825251,94.02505989835251,94.03879023368407,94.03879023378407,94.04222019356828,94.04222019366829,94.04907890081455,94.04907890091455,94.06279207147962,94.06279207157962,94.069642177397,94.069642177497,94.07306501371365,94.07306501381365,94.07990954960745,94.07990954970745,94.0867507889567,94.08675078905671,94.10042667456608,94.10042667466608,94.10726207865255,94.10726207875256,94.11067767772771,94.11067767782771,94.11750705709481,94.11750705719481,94.12092064793018,94.12092064803018,94.12433317780868,94.12433317790868,94.12774476040425,94.12774476050426,94.1311551304777,94.1311551305777,94.13797488545035,94.13797488555035,94.14138180741396,94.14138180751397,94.14819307473165,94.14819307483165,94.15159692749862,94.15159692759862,94.1584025490101,94.15840254911011,94.1618037493849,94.1618037494849,94.16520354778096,94.16520354788096,94.16860251256806,94.16860251266806,94.17539972220723,94.17539972230723,94.17879683031981,94.17879683041981,94.18219287747552,94.18219287757552,94.18558577965194,94.18558577975195,94.18897762087148,94.18897762097149,94.19236752963388,94.19236752973389,94.19575611220019,94.19575611230019,94.20253009446219,94.20253009456219,94.21607055650549,94.21607055660549,94.21945406163535,94.21945406173535,94.22283559641673,94.22283559651673,94.23635745382349,94.23635745392349,94.23973482056005,94.23973482066005,94.24648822783705,94.24648822793705,94.24986354844249,94.24986354854249,94.25323811122159,94.2532381113216,94.25998602425767,94.25998602435767,94.2633581619925,94.2633581620925,94.26672779890043,94.26672779900044,94.27683500451495,94.27683500461495,94.28694084604209,94.28694084614209,94.29367549711729,94.2936754972173,94.29704062495853,94.29704062505853,94.30040480551683,94.30040480561684,94.30713036267602,94.30713036277602,94.31049086777662,94.31049086787662,94.31385034981164,94.31385034991165,94.31720911191167,94.31720911201167,94.32392493100245,94.32392493110245,94.3272814954061,94.3272814955061,94.33063673361366,94.33063673371366,94.33734539124558,94.33734539134558,94.340697863387,94.34069786348701,94.34740220140881,94.34740220150881,94.35075274109309,94.35075274119309,94.3541024471684,94.3541024472684,94.36415001185036,94.36415001195036,94.36749808859905,94.36749808869905,94.3741923475306,94.3741923476306,94.37753811290897,94.37753811300897,94.38088315835232,94.38088315845232,94.38422649868642,94.38422649878642,94.38756790656336,94.38756790666336,94.39090798824422,94.39090798834422,94.39758690119248,94.39758690129248,94.40426373011832,94.40426373021832,94.40759949218906,94.40759949228907,94.41093445854216,94.41093445864216,94.4176024587912,94.4176024588912,94.42093420438236,94.42093420448236,94.42426523003851,94.42426523013852,94.43092606882868,94.43092606892868,94.43425584407137,94.43425584417137,94.44091164331641,94.44091164341641,94.44756278192949,94.4475627820295,94.45753668589167,94.45753668599167,94.4641843385036,94.4641843386036,94.47082998287574,94.47082998297574,94.47747282329046,94.47747282339046,94.48079278468211,94.48079278478211,94.48411183668217,94.48411183678218,94.49074842502961,94.49074842512961,94.49738323248516,94.49738323258516,94.50069944263646,94.50069944273646,94.50733015782981,94.50733015792981,94.51727253621644,94.51727253631644,94.52058541193186,94.52058541203186,94.52389768138622,94.52389768148622,94.52720714688314,94.52720714698314,94.54375204932344,94.54375204942345,94.55367529259512,94.55367529269512,94.56028929771777,94.56028929781777,94.56690049888299,94.56690049898299,94.57350927500393,94.57350927510393,94.57681112434618,94.57681112444618,94.58341326948668,94.58341326958669,94.59331279279407,94.59331279289407,94.59661020885225,94.59661020895226,94.60320398001176,94.60320398011176,94.60649988041727,94.60649988051728,94.61308955931457,94.61308955941458,94.61638314834977,94.61638314844977,94.61967601744995,94.61967601754995,94.62296816661511,94.62296816671511,94.63284188593578,94.63284188603578,94.6394207658078,94.6394207659078,94.6427078375365,94.64270783763651,94.64599286313411,94.64599286323411,94.65256166391588,94.65256166401588,94.65912584195699,94.659125842057,94.6624058659008,94.6624058660008,94.67880329533628,94.67880329543628,94.68207983327893,94.68207983337894,94.68535512080814,94.68535512090814,94.6919047106923,94.6919047107923,94.69845195131482,94.69845195141482,94.70172227508154,94.70172227518154,94.70499187891322,94.70499187901322,94.70825894402668,94.70825894412668,94.71152517553118,94.71152517563118,94.72132140710909,94.72132140720909,94.72784708757237,94.72784708767237,94.73110657442248,94.73110657452249,94.73762361566556,94.73762361576556,94.74413792873382,94.74413792883382,94.75064860423568,94.75064860433568,94.76366233908473,94.76366233918473,94.76691258045355,94.76691258055355,94.77341128229932,94.77341128239932,94.77990790012265,94.77990790022265,94.78315495862087,94.78315495872087,94.78640129718407,94.78640129728407,94.79938513578419,94.7993851358842,94.80262973134683,94.80262973144683,94.805871522952,94.805871523052,94.80911168523058,94.80911168533058,94.81235116546544,94.81235116556545,94.81882879973908,94.81882879983908,94.82530423631636,94.82530423641636,94.82854083681114,94.82854083691115,94.83177614900117,94.83177614910117,94.83500733103767,94.83500733113767,94.84146870993642,94.84146871003642,94.84469777005918,94.84469777015919,94.85115482934782,94.85115482944782,94.85438150231762,94.85438150241762,94.86405936142198,94.86405936152198,94.87050884244726,94.87050884254727,94.87695582264566,94.87695582274566,94.88017726661374,94.88017726671374,94.88661628963561,94.88661628973561,94.8898323151454,94.8898323152454,94.89304701445911,94.89304701455912,94.8962580383151,94.8962580384151,94.89946747073577,94.89946747083577,94.90267603165616,94.90267603175616,94.90588375896758,94.90588375906758,94.9090889854521,94.9090889855521,94.91229337832765,94.91229337842765,94.92190394245343,94.92190394255343,94.93150851975118,94.93150851985118,94.93790601388987,94.93790601398987,94.9411010855015,94.9411010856015,94.95387928792557,94.95387928802558,94.96026306540762,94.96026306550762,94.96345154393012,94.96345154403012,94.96663930251762,94.96663930261762,94.97301315247466,94.97301315257467,94.97619901649632,94.97619901659633,94.97938378166978,94.97938378176978,94.98256604601634,94.98256604611635,94.98574618844917,94.98574618854917,94.99528502431234,94.99528502441234,95.00482052573963,95.00482052583963,95.01435038136071,95.01435038146072,95.02070100394059,95.02070100404059,95.02704999719384,95.02704999729384,95.03022369810282,95.03022369820282,95.03656784126753,95.03656784136753,95.0460786754476,95.0460786755476,95.052416680219,95.052416680319,95.05875237362007,95.05875237372007,95.06191800367857,95.06191800377857,95.06508238332363,95.06508238342363,95.07457374136689,95.07457374146689,95.08406168919166,95.08406168929166,95.08722030935657,95.08722030945657,95.09353614770765,95.09353614780765,95.09669230493697,95.09669230503697,95.10615929886356,95.10615929896356,95.1093100755259,95.1093100756259,95.11245960177477,95.11245960187478,95.12190605860769,95.1219060587077,95.13134887787419,95.13134887797419,95.13764194354391,95.13764194364391,95.14393273573462,95.14393273583462,95.14707676774258,95.14707676784258,95.15650814383142,95.15650814393142,95.16279302497672,95.16279302507672,95.16907578420827,95.16907578430828,95.17535653520005,95.17535653530005,95.18163349706015,95.18163349716015,95.18790666978856,95.18790666988856,95.19104187311972,95.19104187321972,95.19417468929791,95.19417468939791,95.20043747980554,95.20043747990555,95.20356707522181,95.20356707532181,95.21608409279948,95.21608409289948,95.2254686352208,95.22546863532081,95.23172096772501,95.23172096782501,95.2379707994023,95.2379707995023,95.24734253455857,95.24734253465857,95.2567121478011,95.2567121479011,95.26295618210693,95.26295618220693,95.26607562265032,95.26607562275032,95.26919358543233,95.26919358553234,95.27542810901767,95.27542810911767,95.28166073803716,95.28166073813716,95.28477404018722,95.28477404028722,95.29100003822627,95.29100003832627,95.29411182472367,95.29411182482367,95.30033388206579,95.30033388216579,95.3096634441865,95.3096634442865,95.31277200992197,95.31277201002197,95.3220961914757,95.3220961915757,95.32830991272813,95.32830991282813,95.3314144240927,95.3314144241927,95.33451798817435,95.33451798827436,95.33761931666832,95.33761931676833,95.34071996311859,95.34071996321859,95.34381901813356,95.34381901823356,95.34691670906112,95.34691670916112,95.3500137937276,95.3500137938276,95.35620709156031,95.35620709166031,95.3593026226828,95.3593026227828,95.36549277553621,95.36549277563621,95.3685853132447,95.3685853133447,95.3716772446921,95.3716772447921,95.38095015929426,95.38095015939426,95.38403833950132,95.38403833960132,95.38712564820808,95.38712564830809,95.39329905309948,95.39329905319948,95.39946950246816,95.39946950256817,95.40563729944469,95.4056372995447,95.41180293661616,95.41180293671616,95.42104803899181,95.42104803909181,95.42720912920527,95.42720912930527,95.43028840495288,95.43028840505288,95.4395252849128,95.4395252850128,95.44568156292902,95.44568156302903,95.44875864098027,95.44875864108027,95.45490976577742,95.45490976587742,95.46105748035606,95.46105748045606,95.46720402030387,95.46720402040387,95.47027598302735,95.47027598312735,95.47641824125637,95.47641824135637,95.47948751369637,95.47948751379637,95.48562488394553,95.48562488404554,95.49176039752017,95.49176039762017,95.50096082603338,95.50096082613338,95.51628854570548,95.51628854580548,95.52548116860743,95.52548116870743,95.53466897931216,95.53466897941216,95.5407900184038,95.5407900185038,95.54690931449487,95.54690931459487,95.55302626132429,95.5530262614243,95.55608352221688,95.55608352231688,95.56219705882779,95.5621970589278,95.56525003800158,95.56525003810158,95.56830199410982,95.56830199420982,95.57135114626061,95.57135114636061,95.57439965425901,95.57439965435901,95.58049454834209,95.58049454844209,95.58354044183965,95.58354044193965,95.58658455444531,95.58658455454531,95.58962654513724,95.58962654523724,95.59266648969805,95.59266648979805,95.59570529751937,95.59570529761938,95.60785946784777,95.60785946794778,95.61697134435371,95.61697134445372,95.62607916648876,95.62607916658877,95.62911157067755,95.62911157077755,95.63214200451786,95.63214200461786,95.64123179038613,95.64123179048613,95.64425570691996,95.64425570701997,95.65029846255118,95.65029846265118,95.65331677117014,95.65331677127014,95.66538602705768,95.66538602715768,95.66839982660994,95.66839982670994,95.67743921702693,95.67743921712693,95.68346283754317,95.68346283764318,95.68647284796377,95.68647284806377,95.69550075731178,95.69550075741178,95.70151687534731,95.70151687544731,95.70452377867991,95.70452377877992,95.7165492700966,95.71654927019661,95.71955287688466,95.71955287698466,95.72555906739521,95.72555906749521,95.72855979444319,95.72855979454319,95.73456060438676,95.73456060448676,95.73755985367337,95.73755985377338,95.74355702605054,95.74355702615054,95.74655460811923,95.74655460821923,95.75254761245154,95.75254761255154,95.75853728234797,95.75853728244797,95.77649799383893,95.77649799393893,95.78247705416668,95.78247705426668,95.78845353788489,95.78845353798489,95.7914403209283,95.7914403210283,95.80039934386244,95.80039934396244,95.80636995442656,95.80636995452656,95.81532348311977,95.81532348321977,95.81830564342253,95.81830564352254,95.82426814524486,95.82426814534486,95.82724473552406,95.82724473562406,95.83022037852034,95.83022037862034,95.84212139696149,95.84212139706149,95.84509533484851,95.84509533494851,95.85104226333965,95.85104226343965,95.85698733515628,95.85698733525628,95.85995532410656,95.85995532420657,95.86589020315898,95.86589020325899,95.86885648700002,95.86885648710002,95.87182159621025,95.87182159631026,95.8777509052391,95.8777509053391,95.88071453668796,95.88071453678796,95.88367744820182,95.88367744830182,95.8895993684239,95.8895993685239,95.89255655834891,95.89255655844892,95.8955125357518,95.89551253585181,95.90437747454644,95.90437747464644,95.90732996594818,95.90732996604818,95.91323358466427,95.91323358476427,95.9220872318465,95.9220872319465,95.92503688139949,95.9250368814995,95.93388094207859,95.93388094217859,95.9456613524585,95.9456613525585,95.95154932206076,95.95154932216076,95.96037826410448,95.96037826420448,95.96625911013919,95.96625911023919,95.96919686181872,95.96919686191872,95.97507066006851,95.97507066016851,95.98387770093115,95.98387770103115,95.99267936122679,95.9926793613268,95.99560979988217,95.99560979998218,96.0043989560433,96.0043989561433,96.0102551348308,96.0102551349308,96.01318182224584,96.01318182234584,96.02195862583768,96.02195862593769,96.0278046497523,96.0278046498523,96.03949252953669,96.03949252963669,96.04241330590631,96.04241330600631,96.04825372190604,96.04825372200604,96.05701218610058,96.05701218620058,96.0657676568811,96.06576765698111,96.07160117666119,96.07160117676119,96.074514677898,96.074514677998,96.08034077097999,96.08034077107999,96.08907653827579,96.08907653837579,96.09489656874712,96.09489656884712,96.09780328743821,96.09780328753821,96.10070758108505,96.10070758118505,96.10651427381286,96.10651427391286,96.11231861727904,96.11231861737905,96.12102225273824,96.12102225283824,96.12682257972484,96.12682257982485,96.13262116371088,96.13262116381088,96.1384163374784,96.1384163375784,96.14710311130165,96.14710311140165,96.15868076524558,96.15868076534558,96.16157365360606,96.16157365370606,96.16446513998783,96.16446514008783,96.17024693812054,96.17024693822054,96.17602604596975,96.17602604606975,96.18180189516573,96.18180189526574,96.18468788730657,96.18468788740657,96.18757285638185,96.18757285648185,96.19045718130475,96.19045718140475,96.19910727633336,96.19910727643337,96.2048710381993,96.2048710382993,96.21351336550798,96.21351336560798,96.21639238564653,96.21639238574653,96.21926966278451,96.21926966288451,96.22214618209615,96.22214618219616,96.22502194358147,96.22502194368147,96.2278966441099,96.2278966442099,96.23364475686202,96.23364475696202,96.24226522088094,96.24226522098094,96.24513791316959,96.24513791326959,96.24800939293611,96.24800939303611,96.25375140518622,96.25375140528622,96.25661981575608,96.25661981585608,96.26235364348177,96.26235364358178,96.26521894696366,96.26521894706366,96.27381156086476,96.27381156096476,96.27667288575839,96.27667288585839,96.27953341493435,96.27953341503435,96.28525329865548,96.28525329875548,96.28811216061352,96.28811216071352,96.29096878698387,96.29096878708387,96.30239392837785,96.30239392847785,96.30810270994317,96.30810271004317,96.31095448622497,96.31095448632497,96.31380357222329,96.3138035723233,96.32234859463054,96.32234859473054,96.32804191653862,96.32804191663863,96.33373269972847,96.33373269982847,96.33657587468137,96.33657587478137,96.34510327762634,96.34510327772634,96.35078576261782,96.35078576271782,96.35646623936952,96.35646623946953,96.36214402583774,96.36214402593774,96.36498030457098,96.36498030467098,96.3791593868669,96.3791593869669,96.38482819309304,96.38482819319304,96.38766128895568,96.38766128905569,96.39049055779533,96.39049055789533,96.40463466640588,96.40463466650588,96.40746101761414,96.40746101771414,96.41311087818191,96.41311087828191,96.41593344025848,96.41593344035849,96.4187551687261,96.4187551688261,96.42439764048709,96.42439764058709,96.42721720914965,96.42721720924965,96.4300358684206,96.4300358685206,96.43567163341854,96.43567163351854,96.44130550385063,96.44130550395063,96.44412041188122,96.44412041198122,96.44974901542028,96.44974901552028,96.45537367826239,96.45537367836239,96.46099349101597,96.46099349111597,96.46942017884098,96.46942017894098,96.47222781173876,96.47222781183876,96.4778423575993,96.4778423576993,96.48064779280071,96.48064779290071,96.49186779060578,96.49186779070578,96.49467163437188,96.49467163447189,96.49747472031164,96.49747472041165,96.5002754948811,96.5002754949811,96.50587552836733,96.50587552846733,96.51147268211346,96.51147268221347,96.51986356264415,96.51986356274415,96.52545461588831,96.52545461598831,96.52824629654172,96.52824629664173,96.53382848321776,96.53382848331776,96.53940877532796,96.53940877542796,96.5533554701782,96.5533554702782,96.55614358904785,96.55614358914785,96.55892913130796,96.55892913140796,96.5728545691295,96.5728545692295,96.57842254656175,96.57842254666176,96.58677102670899,96.58677102680899,96.58955236303294,96.58955236313294,96.59233286574792,96.59233286584792,96.59510920041805,96.59510920051805,96.60620900696637,96.60620900706637,96.61452202084111,96.61452202094111,96.62559814531645,96.62559814541645,96.63113209634624,96.63113209644624,96.63389713940398,96.63389713950399,96.63942180706118,96.63942180716118,96.64218056016034,96.64218056026034,96.65045511434857,96.65045511444858,96.65320909314183,96.65320909324183,96.65871583820619,96.6587158383062,96.66422057503077,96.66422057513077,96.66697182564921,96.66697182574921,96.66972197741947,96.66972197751947,96.67247087877628,96.67247087887628,96.67796727951118,96.67796727961118,96.6889565570885,96.6889565571885,96.69719428091685,96.69719428101685,96.69993829429379,96.69993829439379,96.7026810951486,96.70268109524861,96.70816389290077,96.70816389300077,96.71090271516732,96.71090271526732,96.7163791850696,96.7163791851696,96.72185292679707,96.72185292689707,96.72458780836668,96.72458780846668,96.73005594217926,96.73005594227926,96.7327879440088,96.73278794410881,96.74098250962736,96.74098250972736,96.74371208641261,96.74371208651262,96.75189926322442,96.75189926332442,96.75462630129145,96.75462630139145,96.75735167214054,96.75735167224055,96.76552634481781,96.76552634491782,96.77369863034212,96.77369863044213,96.77914402936466,96.77914402946466,96.78458692756028,96.78458692766029,96.78730684205978,96.78730684215978,96.79546328901367,96.79546328911367,96.80361249872608,96.80361249882608,96.80632665374542,96.80632665384542,96.81446722423759,96.81446722433759,96.81989170725328,96.81989170735328,96.82531368944205,96.82531368954206,96.83344406717003,96.83344406727004,96.84156974637473,96.84156974647473,96.844277194631,96.844277194731,96.85239749326871,96.85239749336871,96.8578072744535,96.85780727455351,96.86051097146941,96.86051097156941,96.86321258446291,96.86321258456292,96.8686146737104,96.8686146738104,96.8740147926094,96.8740147927094,96.87671363953677,96.87671363963678,96.88211019665201,96.88211019675201,96.88750493498404,96.88750493508404,96.89559317756778,96.89559317766778,96.89828770488504,96.89828770498504,96.90367562278006,96.90367562288006,96.91175598396991,96.91175598406991,96.91444850304738,96.91444850314738,96.91714037797246,96.91714037807246,96.91983160874517,96.91983160884517,96.92790257288846,96.92790257298846,96.93327882025791,96.93327882035791,96.93865226366992,96.93865226376992,96.94402195584158,96.94402195594158,96.94670481263329,96.94670481273329,96.94938535805466,96.94938535815466,96.95474356915733,96.95474356925733,96.95742055279493,96.95742055289493,96.96009461880115,96.96009461890115,96.96544161407407,96.96544161417407,96.97078644954192,96.97078644964192,96.97345805261254,96.97345805271254,96.97612870840025,96.97612870850026,96.9814680875185,96.9814680876185,96.98413616669666,96.98413616679666,96.98946990000871,96.98946990010872,96.99213544046864,96.99213544056865,96.99746402056162,96.99746402066162,97.00012671917281,97.00012671927281,97.00811155635311,97.00811155645312,97.01077080685447,97.01077080695447,97.01342808700736,97.01342808710736,97.0187402980515,97.0187402981515,97.02139462268167,97.02139462278167,97.02404830315946,97.02404830325946,97.03200726057042,97.03200726067043,97.0346574929384,97.0346574930384,97.03730658856686,97.03730658866687,97.03995507793427,97.03995507803427,97.04789918194905,97.04789918204905,97.05584086091957,97.05584086101958,97.06642545923393,97.06642545933393,97.07171485970538,97.07171485980538,97.07435827163633,97.07435827173633,97.0770010394149,97.0770010395149,97.07964187473631,97.07964187483631,97.08228111862242,97.08228111872242,97.08755786339407,97.08755786349407,97.0928320694475,97.0928320695475,97.10074032827664,97.10074032837665,97.1060110104376,97.1060110105376,97.10864544212649,97.10864544222649,97.11127900231509,97.11127900241509,97.11654426601775,97.11654426611776,97.12706956442139,97.12706956452139,97.12969914602172,97.12969914612172,97.13232751509992,97.13232751519992,97.13758349543,97.13758349553,97.14021065198605,97.14021065208605,97.14546379046736,97.14546379056736,97.15071488281757,97.15071488291757,97.15333965222067,97.15333965232067,97.16383520594064,97.16383520604064,97.17169863934207,97.17169863944207,97.17431772504766,97.17431772514766,97.17693593925297,97.17693593935297,97.17955324406668,97.17955324416668,97.18740280924615,97.18740280934615,97.19263169728474,97.19263169738474,97.20047076656948,97.20047076666948,97.20830680454888,97.20830680464888,97.21352712914988,97.21352712924988,97.21613507480835,97.21613507490835,97.22395762347898,97.22395762357898,97.22656371246293,97.22656371256294,97.23177410953893,97.23177410963893,97.24219175871166,97.24219175881166,97.24739866978652,97.24739866988652,97.25000013602983,97.25000013612983,97.25259955614203,97.25259955624203,97.26039656606517,97.26039656616517,97.26819080992219,97.26819081002219,97.27598206036518,97.27598206046518,97.27857750188912,97.27857750198912,97.28636189400378,97.28636189410378,97.28895551674451,97.28895551684451,97.29414185283437,97.29414185293437,97.30191794675066,97.30191794685066,97.30450724988128,97.30450724998128,97.30709583307689,97.30709583317689,97.31227155959806,97.31227155969806,97.31744452005312,97.31744452015312,97.32519966007123,97.32519966017124,97.3277833931783,97.3277833932783,97.3303658758719,97.3303658759719,97.33552852988879,97.33552852998879,97.34068891042668,97.34068891052668,97.34584603231133,97.34584603241133,97.35615500918759,97.35615500928759,97.36903472244927,97.36903472254927,97.37160716394393,97.37160716404394,97.37417873393831,97.37417873403831,97.3767485230408,97.3767485231408,97.3793165312514,97.3793165313514,97.3844497816065,97.38444978170651,97.38701305340251,97.38701305350251,97.3921388391682,97.3921388392682,97.39470048163759,97.39470048173759,97.40238449965416,97.40238449975416,97.40750505641813,97.40750505651813,97.41006340234297,97.41006340244297,97.41262034628907,97.41262034638908,97.41773328689838,97.41773328699838,97.42028761634364,97.42028761644364,97.42284077115808,97.42284077125808,97.42539271345038,97.42539271355038,97.4279440115903,97.4279440116903,97.43814784006256,97.43814784016256,97.44069773622375,97.44069773632376,97.44579669493717,97.44579669503717,97.45089345595422,97.45089345605422,97.45598627627433,97.45598627637433,97.45853064030328,97.45853064040328,97.46361800427377,97.46361800437377,97.46616058741084,97.46616058751084,97.47124427592361,97.47124427602361,97.47378515395141,97.47378515405141,97.47632534993552,97.47632535003552,97.48140471883818,97.48140471893818,97.48394362651752,97.48394362661752,97.48648189004446,97.48648189014446,97.49155690144569,97.49155690154569,97.4966282373892,97.4966282374892,97.50423192581425,97.50423192591425,97.51183182510762,97.51183182520762,97.51436084315327,97.51436084325327,97.52447475553087,97.52447475563088,97.5270016516628,97.5270016517628,97.53205415562188,97.53205415572188,97.53962931188539,97.5396293119854,97.5421523052117,97.5421523053117,97.54719692777688,97.54719692787688,97.54971760973287,97.54971760983287,97.55475734431819,97.55475734441819,97.55727556333858,97.55727556343858,97.5648285531818,97.5648285532818,97.56986237672172,97.56986237682172,97.5748940783479,97.5748940784479,97.57992460534325,97.57992460544325,97.58746823603124,97.58746823613124,97.59249565593862,97.59249565603862,97.59500834282676,97.59500834292676,97.60254466049061,97.60254466059061,97.61510009986347,97.61510009996347,97.6226289150466,97.6226289151466,97.63266143687581,97.63266143697581,97.63767362447385,97.63767362457385,97.64017850575074,97.64017850585074,97.6451861842821,97.6451861843821,97.65269313617546,97.65269313627546,97.66269814890869,97.66269814900869,97.66769783237221,97.66769783247221,97.6751935684358,97.6751935685358,97.67768950736195,97.67768950746195,97.68018484002701,97.68018484012701,97.69016412455616,97.69016412465616,97.69764527246281,97.69764527256281,97.70263108770439,97.70263108780439,97.71010624878299,97.710106248883,97.71508766863184,97.71508766873184,97.7175773554907,97.7175773555907,97.72255540301235,97.72255540311235,97.72753197277265,97.72753197287265,97.73250702688027,97.73250702698027,97.73499353086854,97.73499353096854,97.74245167874592,97.74245167884592,97.74493681864678,97.74493681874678,97.74742131439524,97.74742131449524,97.75487317231403,97.75487317241404,97.75735554614877,97.75735554624877,97.76480062152187,97.76480062162187,97.76728041874706,97.76728041884706,97.76975953392856,97.76975953402857,97.77471609707361,97.77471609717361,97.77967042463096,97.77967042473097,97.7821466979637,97.7821466980637,97.78957381285265,97.78957381295265,97.79947197207264,97.79947197217264,97.80441807721425,97.80441807731425,97.80688946256713,97.80688946266713,97.81430267134283,97.81430267144283,97.81677098749905,97.81677098759906,97.81923797745918,97.81923797755918,97.82417066907468,97.82417066917468,97.83403396828324,97.83403396838324,97.83649864687304,97.83649864697304,97.84635569401433,97.84635569411434,97.84881768232066,97.84881768242066,97.85373968858482,97.85373968868483,97.85865953504393,97.85865953514393,97.86111792367515,97.86111792377515,97.86603405678522,97.86603405688523,97.86849062663325,97.86849062673325,97.87094651443756,97.87094651453756,97.8734010760458,97.8734010761458,97.8758537430882,97.8758537431882,97.88075627321554,97.88075627331554,97.88320515112626,97.88320515122626,97.88810226279531,97.88810226289532,97.89789398530652,97.89789398540653,97.9003402866077,97.9003402867077,97.90278526171281,97.90278526181281,97.90767335524846,97.90767335534846,97.91011605687456,97.91011605697456,97.91255784910905,97.91255784920905,97.9149988835172,97.9149988836172,97.91743893275111,97.91743893285111,97.92475680697385,97.92475680707385,97.93450942564617,97.93450942574617,97.93938308259014,97.93938308269014,97.94425492075092,97.94425492085092,97.94912478856322,97.94912478866323,97.95642852932482,97.95642852942483,97.96129589631022,97.96129589641022,97.9637279504763,97.9637279505763,97.97102271099583,97.97102271109583,97.97588276495709,97.9758827650571,97.98073819617771,97.98073819627771,97.98558881520111,97.98558881530111,97.9904373880934,97.9904373881934,97.99528304335429,97.9952830434543,98.00012661459277,98.00012661469277,98.00738886436251,98.00738886446251,98.00980754289378,98.00980754299378,98.01464406634734,98.01464406644735,98.01947880890901,98.01947880900902,98.02914450490069,98.02914450500069,98.03155969743081,98.03155969753081,98.03638811214257,98.03638811224258,98.03879997023682,98.03879997033683,98.04844475022165,98.04844475032165,98.05567365563265,98.05567365573265,98.0604907408407,98.06049074094071,98.06289771095508,98.06289771105509,98.07011680251505,98.07011680261505,98.07492862083008,98.07492862093008,98.07733259753044,98.07733259763044,98.08454096584775,98.08454096594775,98.08934168200697,98.08934168210698,98.09413864692584,98.09413864702584,98.09893356571361,98.09893356581361,98.10372636258762,98.10372636268762,98.10612151061119,98.10612151071119,98.1109104425709,98.1109104426709,98.11330369602862,98.11330369612863,98.12048186496648,98.12048186506648,98.12765776042532,98.12765776052532,98.13004718686005,98.13004718696006,98.1348251303379,98.1348251304379,98.137212207511,98.137212207611,98.13959712487903,98.13959712497903,98.14675085391758,98.14675085401758,98.15151841517464,98.15151841527464,98.15628381662664,98.15628381672664,98.15866233036215,98.15866233046215,98.16579665904652,98.16579665914652,98.17055092045139,98.17055092055139,98.1729258345118,98.1729258346118,98.17767410908864,98.17767410918864,98.18004656021346,98.18004656031346,98.18479017415832,98.18479017425832,98.18716054126072,98.18716054136073,98.19190006294339,98.19190006304339,98.19426876282785,98.19426876292785,98.19663678066861,98.19663678076861,98.19900392700909,98.19900392710909,98.20610275152966,98.20610275162966,98.21319555133086,98.21319555143086,98.21555659716932,98.21555659726933,98.21791555898538,98.21791555908538,98.22499047408508,98.22499047418508,98.23441775791171,98.23441775801172,98.23677255168292,98.23677255178292,98.24619059002826,98.24619059012826,98.25089600952585,98.25089600962585,98.26030154373666,98.26030154383666,98.26265118428879,98.26265118438879,98.27204671519198,98.27204671529198,98.27439302130823,98.27439302140823,98.27908491360573,98.27908491370573,98.2814298935259,98.2814298936259,98.28377286520629,98.28377286530629,98.28845725502306,98.28845725512306,98.29079749852863,98.29079749862863,98.29313668107734,98.29313668117734,98.29781394732657,98.29781394742658,98.30248977370577,98.30248977380577,98.30482463664437,98.30482463674437,98.30715874175662,98.30715874185663,98.31182524687189,98.31182524697189,98.3141571921791,98.3141571922791,98.31881964292347,98.31881964302347,98.32347997175411,98.32347997185411,98.32813601886596,98.32813601896596,98.33511162082443,98.33511162092444,98.34207956873692,98.34207956883692,98.34904558419225,98.34904558429226,98.35136638945232,98.35136638955233,98.3560066737764,98.35600667387641,98.35832486453563,98.35832486463563,98.36759588457195,98.36759588467196,98.37222896954584,98.37222896964585,98.37685894743176,98.37685894753176,98.37916982516684,98.37916982526684,98.38379074702806,98.38379074712806,98.38840962275816,98.38840962285816,98.39302713440085,98.39302713450085,98.39533433667822,98.39533433677822,98.40686689995522,98.40686690005522,98.41147656809535,98.41147656819535,98.41608472058282,98.41608472068282,98.4206915847655,98.42069158486551,98.4252964028171,98.4252964029171,98.4345029318323,98.4345029319323,98.43680319999869,98.4368032000987,98.44370192047546,98.44370192057546,98.4505971549511,98.4505971550511,98.4551915907819,98.4551915908819,98.45978375313368,98.45978375323368,98.46437375568041,98.46437375578041,98.46896258359634,98.46896258369634,98.47584410141529,98.47584410151529,98.48042989802587,98.48042989812588,98.4873061489518,98.4873061490518,98.48959539074502,98.48959539084503,98.49646099421095,98.49646099431095,98.49874743204673,98.49874743214673,98.50560473731429,98.50560473741429,98.5124581018849,98.5124581019849,98.52159229637641,98.52159229647641,98.5284390678579,98.5284390679579,98.53756356217232,98.53756356227233,98.5466822970066,98.5466822971066,98.55124020560804,98.55124020570804,98.56035204422267,98.56035204432267,98.56490487538767,98.56490487548767,98.57173201916709,98.57173201926709,98.5740056497378,98.5740056498378,98.57855177413973,98.57855177423973,98.58082404062304,98.58082404072304,98.58763947598558,98.58763947608558,98.59445275154306,98.59445275164306,98.59672160780786,98.59672160790787,98.60125788046744,98.60125788056745,98.6035248042751,98.6035248043751,98.60805781828142,98.60805781838143,98.61032265806665,98.61032265816665,98.61938008475042,98.61938008485042,98.62164340888297,98.62164340898298,98.62390608886315,98.62390608896315,98.62843065310585,98.62843065320585,98.63069159008545,98.63069159018545,98.63295188291266,98.63295188301267,98.63521134213092,98.63521134223092,98.644248231721,98.644248231821,98.6487646872219,98.6487646873219,98.65553707804861,98.65553707814861,98.66230549028705,98.66230549038706,98.66455892478594,98.66455892488594,98.6668100479145,98.6668100480145,98.66906041321673,98.66906041331673,98.67355932503797,98.67355932513797,98.68030307002921,98.68030307012921,98.68479614658767,98.68479614668767,98.69153341216374,98.69153341226374,98.69602038822022,98.69602038832022,98.70050402984081,98.70050402994082,98.7027438992483,98.7027438993483,98.7049822151118,98.7049822152118,98.70721920477921,98.70721920487921,98.70945441355474,98.70945441365474,98.7139235049097,98.7139235050097,98.71839104272067,98.71839104282067,98.72285657229186,98.72285657239186,98.72508784037045,98.72508784047045,98.72731842640532,98.72731842650532,98.73177819649636,98.73177819659637,98.73623626147815,98.73623626157816,98.74069284869859,98.74069284879859,98.74737479295219,98.74737479305219,98.75182687110592,98.75182687120592,98.76072879182571,98.76072879192571,98.76962563510905,98.76962563520905,98.77629352168414,98.77629352178414,98.78073612700868,98.78073612710868,98.78739700369017,98.78739700379018,98.78961565396027,98.78961565406027,98.79626989966133,98.79626989976133,98.80070432046145,98.80070432056145,98.80513529315176,98.80513529325177,98.80734913122463,98.80734913132463,98.81398890244267,98.81398890254268,98.81841347150045,98.81841347160045,98.82283512292683,98.82283512302683,98.82725446298289,98.8272544630829,98.83829936501323,98.83829936511323,98.84271404443732,98.84271404453732,98.84712603357794,98.84712603367794,98.849331042974,98.849331043074,98.85153457460872,98.85153457470872,98.85594027379074,98.85594027389074,98.86254707956321,98.86254707966322,98.86694944430937,98.86694944440937,98.87355162734119,98.87355162744119,98.8757512940616,98.8757512941616,98.8823479828525,98.8823479829525,98.88674405764006,98.88674405774006,98.89333536586398,98.89333536596398,98.89553112977876,98.89553112987876,98.89992106617302,98.89992106627302,98.90430759234877,98.90430759244877,98.90869188293682,98.90869188303682,98.91088253152384,98.91088253162384,98.91526238882784,98.91526238892784,98.91963997265283,98.91963997275283,98.92182709734739,98.92182709744739,98.9262001721057,98.9262001722057,98.93494306296905,98.93494306306906,98.93931011300799,98.939310113108,98.94367272976287,98.94367272986287,98.9458530340204,98.9458530341204,98.95021261946994,98.95021261956994,98.95674947787167,98.95674947797167,98.96110553942874,98.96110553952875,98.96328266081568,98.96328266091568,98.96545762239757,98.96545762249757,98.97415580150715,98.97415580160715,98.97850189764792,98.97850189774792,98.98284435622227,98.98284435632227,98.98935457702834,98.98935457712834,98.99152328654294,98.99152328664294,98.99585865944107,98.99585865954107,99.0001915315123,99.0001915316123,99.00668777042246,99.00668777052246,99.00884984895664,99.00884984905665,99.01533453101521,99.01533453111522,99.01749456341828,99.01749456351828,99.01965300438606,99.01965300448606,99.02612681163671,99.02612681173672,99.03259739812545,99.03259739822545,99.03475307302709,99.03475307312709,99.03690764908056,99.03690764918056,99.03906142941636,99.03906142951637,99.04336770178323,99.04336770188323,99.04551920864003,99.04551920874003,99.04767003345313,99.04767003355313,99.05197047055721,99.05197047065721,99.05411902189131,99.05411902199131,99.05841502571133,99.05841502581133,99.06270955176998,99.06270955186999,99.06700275163256,99.06700275173256,99.07344071369756,99.07344071379757,99.07773035177637,99.07773035187637,99.07987340886953,99.07987340896953,99.0841580074032,99.0841580075032,99.08843923360968,99.08843923370968,99.09271727694555,99.09271727704555,99.09485423353672,99.09485423363672,99.09912704787088,99.09912704797088,99.10553397694746,99.10553397704746,99.10766816747251,99.10766816757251,99.11620276976765,99.11620276986766,99.12046842264294,99.12046842274295,99.12686512106399,99.12686512116399,99.13112713637288,99.13112713647288,99.13538642350696,99.13538642360696,99.13751443774737,99.13751443784737,99.13964161837882,99.13964161847882,99.14389427453246,99.14389427463246,99.14814424040262,99.14814424050262,99.15239083394557,99.15239083404558,99.15663432040057,99.15663432050057,99.16087469976759,99.16087469986759,99.16299248335248,99.16299248345248,99.16510909230655,99.16510909240655,99.17569058353293,99.17569058363293,99.17992065646179,99.17992065656179,99.18626354921304,99.18626354931304,99.18837508073067,99.18837508083067,99.19259696913511,99.19259696923511,99.19892785033895,99.19892785043895,99.20103654000782,99.20103654010782,99.20525297206265,99.20525297216265,99.21157525193759,99.21157525203759,99.21789359111558,99.21789359121559,99.22420969470589,99.2242096948059,99.2284179801276,99.2284179802276,99.23052045562054,99.23052045572054,99.23262183226527,99.23262183236527,99.2389233098073,99.2389233099073,99.24102059418983,99.24102059428984,99.24311723441998,99.24311723451999,99.24730816561862,99.24730816571862,99.25359077328491,99.25359077338491,99.25777734698214,99.25777734708214,99.26614621265777,99.26614621275777,99.26823610823354,99.26823610833354,99.27450443087342,99.27450443097342,99.276591674057,99.276591674157,99.27867823519689,99.2786782352969,99.28285025862847,99.28285025872847,99.2849351146591,99.28493511475911,99.28910406889403,99.28910406899404,99.29743837768879,99.2974383777888,99.3057698446348,99.3057698447348,99.30993300149825,99.30993300159825,99.31201270430981,99.31201270440981,99.32240909645382,99.32240909655383,99.32864456732207,99.32864456742207,99.33279878183478,99.33279878193478,99.33695102599903,99.33695102609903,99.34110183029323,99.34110183039323,99.34317603886385,99.34317603896385,99.34524869389048,99.34524869399048,99.3493932082261,99.3493932083261,99.35353620690904,99.35353620700904,99.35560651267404,99.35560651277405,99.3576758332648,99.35767583336481,99.36181292090234,99.36181292100234,99.37008315548046,99.37008315558046,99.37421493833365,99.37421493843365,99.3783455086647,99.3783455087647,99.38247467701703,99.38247467711703,99.38660024569427,99.38660024579427,99.39072282095749,99.3907228210575,99.39278289606696,99.39278289616696,99.3969000907632,99.3969000908632,99.40101379945828,99.40101379955829,99.40512512100042,99.40512512110043,99.41128954564975,99.41128954574975,99.41745162103744,99.41745162113745,99.42360994518477,99.42360994528477,99.42771399159409,99.42771399169409,99.43386822347921,99.43386822357921,99.44207040525244,99.44207040535244,99.44411922664084,99.44411922674084,99.45026394780544,99.45026394790544,99.45231125354117,99.45231125364117,99.45640434935994,99.45640434945994,99.46458834330113,99.46458834340113,99.46663307242731,99.46663307252732,99.47072067400516,99.47072067410517,99.47480668414771,99.47480668424771,99.47684747257695,99.47684747267695,99.48092772323935,99.48092772333935,99.48500399531349,99.48500399541349,99.49111146931372,99.49111146941372,99.495180579929,99.495180580029,99.49924753073921,99.49924753083921,99.50331228385306,99.50331228395306,99.50737563498818,99.50737563508818,99.51346764933119,99.51346764943119,99.51549574416931,99.51549574426932,99.52157862670498,99.52157862680498,99.5256306862277,99.5256306863277,99.52765290791174,99.52765290801175,99.52967437176943,99.52967437186943,99.53371608696268,99.53371608706269,99.5357355425806,99.5357355426806,99.54784905552611,99.54784905562612,99.55591725360328,99.55591725370329,99.55793295798084,99.55793295808084,99.56599319888149,99.56599319898149,99.57001990911331,99.57001990921331,99.5740449900185,99.57404499011851,99.58007808336396,99.58007808346396,99.58208643682606,99.58208643692606,99.58409369143996,99.58409369153996,99.59011276499821,99.59011276509821,99.59211645782834,99.59211645792834,99.59411946861478,99.59411946871478,99.59812420188288,99.59812420198288,99.60412777789257,99.60412777799257,99.60612707532997,99.60612707542997,99.61012483659579,99.61012483669579,99.61212208790207,99.61212208800207,99.61411865716465,99.61411865726465,99.61611450649222,99.61611450659223,99.62010465160337,99.62010465170337,99.62409301582262,99.62409301592263,99.62608534125773,99.62608534135774,99.63006824912739,99.63006824922739,99.63404888351803,99.63404888361804,99.6360374956041,99.6360374957041,99.64399001149121,99.64399001159121,99.64796371177088,99.64796371187089,99.64994879996449,99.6499488000645,99.6519325240707,99.6519325241707,99.65391507354609,99.65391507364609,99.65787858106155,99.65787858116155,99.66184080027224,99.66184080037225,99.66778109778295,99.66778109788295,99.66976008547456,99.66976008557457,99.67371688622697,99.67371688632697,99.67767236078328,99.67767236088328,99.68162518294743,99.68162518304743,99.6836000404855,99.6836000405855,99.68557368550145,99.68557368560145,99.69149276387478,99.69149276397478,99.69543565851393,99.69543565861393,99.69937582497828,99.69937582507828,99.70331421055073,99.70331421065073,99.70528228554312,99.70528228564312,99.70724941325258,99.70724941335259,99.71118245614937,99.71118245624938,99.71511379393692,99.71511379403692,99.71904312348467,99.71904312358467,99.72297086159712,99.72297086169712,99.72689678092637,99.72689678102637,99.73278467474599,99.73278467484599,99.73670771433517,99.73670771443517,99.7445517473824,99.7445517474824,99.74847205879676,99.74847205889677,99.75043107776445,99.75043107786445,99.75630605064505,99.75630605074505,99.76021844277422,99.76021844287422,99.76412825829384,99.76412825839384,99.76803655816079,99.76803655826079,99.7778027419245,99.7778027420245,99.77975448575937,99.77975448585937,99.78365588940667,99.78365588950668,99.79340430214282,99.79340430224282,99.79730187876713,99.79730187886713,99.80509381125383,99.80509381135383,99.81482953039887,99.81482953049887,99.81677524951436,99.81677524961437,99.81871922562928,99.81871922572928,99.82066229235261,99.82066229245261,99.82454660701606,99.82454660711606,99.82842633683018,99.82842633693018,99.83617840765163,99.83617840775163,99.84005256744219,99.84005256754219,99.84198763909768,99.84198763919768,99.84779046691119,99.8477904670112,99.85165594959021,99.85165594969021,99.8574526390104,99.8574526391104,99.86324648658184,99.86324648668185,99.86517541984401,99.86517541994401,99.86903184649832,99.86903184659832,99.87481186373914,99.87481186383914,99.87866181097829,99.87866181107829,99.88058479530373,99.88058479540373,99.88250664288968,99.88250664298968,99.88442705060558,99.88442705070558,99.88826544099311,99.88826544109311,99.89018304475158,99.89018304485158,99.89209985279238,99.89209985289239,99.8959322942432,99.8959322943432,99.89784739717474,99.89784739727475,99.90359164501255,99.90359164511256,99.90741912270086,99.90741912280086,99.91315708058008,99.91315708068008,99.92080157795327,99.92080157805327,99.92653150287335,99.92653150297335,99.9322580175749,99.9322580176749,99.93607014927991,99.93607014937992,99.94178351569457,99.94178351579457,99.94368675864398,99.94368675874398,99.94939482027428,99.94939482037428,99.95319804751986,99.95319804761986,99.95889925082183,99.95889925092183,99.96079817416113,99.96079817426113,99.96269558184775,99.96269558194776,99.96648910891625,99.96648910901625,99.96838469781969,99.96838469791969,99.97407021411651,99.97407021421651,99.97975364639092,99.97975364649092,99.98164715127193,99.98164715137193,99.98543283483785,99.98543283493785,99.98921658594662,99.98921658604662,99.99299829092428,99.99299829102428,100.00622978717074,100.00622978727074,100.01000679362512,100.01000679372513,100.01378103401339,100.01378103411339,100.02132470259268,100.02132470269268,100.03263359342687,100.03263359352687,100.03828421182095,100.03828421192095,100.04016570515455,100.04016570525455,100.04769031440152,100.04769031450152,100.05145068656785,100.05145068666785,100.05520939151624,100.05520939161624,100.05896605033352,100.05896605043353,100.06084182207829,100.06084182217829,100.0627163434096,100.0627163435096,100.06459010691458,100.06459010701458,100.06646296102795,100.06646296112795,100.07207993193276,100.07207993203276,100.0739513461761,100.0739513462761,100.07956414903607,100.07956414913608,100.08143400973542,100.08143400983542,100.0851727838512,100.08517278395121,100.08890966340114,100.08890966350114,100.09077725062149,100.09077725072149,100.092643663211,100.09264366331101,100.09450924219156,100.09450924229156,100.09637421491105,100.09637421501105,100.10010325095843,100.10010325105843,100.10196454822018,100.10196454832018,100.10568536185183,100.10568536195183,100.10754430985195,100.10754430995195,100.109402651591,100.109402651691,100.11126008393845,100.11126008403845,100.11497362243728,100.11497362253728,100.12054046313001,100.12054046323001,100.1279595071685,100.1279595072685,100.13537275383551,100.13537275393551,100.14278039258765,100.14278039268765,100.14463123184599,100.144631231946,100.14833169784053,100.14833169794053,100.15018075620698,100.15018075630698,100.15202867783393,100.15202867793393,100.15572319489173,100.15572319499174,100.15941653731872,100.15941653741872,100.16126218546667,100.16126218556667,100.16310722735354,100.16310722745354,100.16679636384436,100.16679636394436,100.16864007953515,100.16864007963515,100.1704825827038,100.1704825828038,100.17416520188814,100.17416520198815,100.17784505500636,100.17784505510636,100.18336157603045,100.18336157613045,100.18703631382091,100.18703631392091,100.19070764139286,100.19070764149286,100.19988025767957,100.19988025777957,100.20538010652429,100.20538010662429,100.20904400739815,100.20904400749815,100.21270499064063,100.21270499074063,100.22002282697206,100.22002282707206,100.2255078224206,100.2255078225206,100.22733497960596,100.22733497970596,100.23098838458509,100.23098838468509,100.23463925084599,100.23463925094599,100.24011261366029,100.24011261376029,100.24375976657215,100.24375976667216,100.2474041913092,100.2474041914092,100.24922536166652,100.24922536176652,100.25468728130316,100.25468728140316,100.26014791263503,100.26014791273504,100.26378574428297,100.26378574438297,100.27105830049086,100.27105830059087,100.27469245668108,100.27469245678108,100.27650779177561,100.27650779187562,100.27832240693513,100.27832240703513,100.28013626426831,100.28013626436831,100.28557605537597,100.28557605547597,100.28920017036724,100.28920017046724,100.29644408073968,100.29644408083968,100.30006300462055,100.30006300472056,100.30187163295201,100.30187163305202,100.30367950345715,100.30367950355715,100.30729399405396,100.30729399415397,100.30910012155853,100.30910012165853,100.3127113535021,100.3127113536021,100.31451588957135,100.31451588967136,100.3181223093177,100.3181223094177,100.31992328360319,100.31992328370319,100.32532488026355,100.32532488036355,100.32892277446362,100.32892277456362,100.33071969437822,100.33071969447822,100.33790547947075,100.33790547957075,100.34149674269038,100.34149674279038,100.34688147771479,100.34688147781479,100.35046876234618,100.35046876244618,100.35405396295513,100.35405396305514,100.35942949038957,100.35942949048957,100.36838415582228,100.36838415592229,100.37017357325615,100.37017357335615,100.37553890792638,100.37553890802639,100.38089871046436,100.38089871056437,100.38625597428413,100.38625597438413,100.3898251468661,100.3898251469661,100.39339272801277,100.39339272811277,100.4005226613044,100.4005226614044,100.40408243683981,100.40408243693982,100.40586079000919,100.40586079010919,100.40763736228668,100.40763736238668,100.41118891540636,100.41118891550636,100.41651397160688,100.41651397170688,100.42006203872542,100.42006203882542,100.4218349734365,100.4218349735365,100.4253800092497,100.4253800093497,100.42892333995364,100.42892334005364,100.43246519289623,100.43246519299623,100.43600575753405,100.43600575763405,100.44308472700463,100.44308472710463,100.45015967999565,100.45015968009565,100.45369433358805,100.45369433368805,100.46076072314148,100.46076072324148,100.467823058324,100.467823058424,100.4766440431912,100.4766440432912,100.48722174528591,100.48722174538591,100.48898282001484,100.48898282011484,100.49250409797243,100.49250409807243,100.49426339180948,100.49426339190948,100.50129840735264,100.50129840745264,100.50481366059086,100.50481366069086,100.50832652667613,100.50832652677613,100.51183651302132,100.51183651312132,100.51534339227855,100.51534339237855,100.52585872526586,100.52585872536586,100.52936147436957,100.52936147446957,100.53461131630631,100.53461131640631,100.53810675238587,100.53810675248587,100.5398533147405,100.5398533148405,100.54334556794946,100.54334556804946,100.54858222370797,100.54858222380797,100.55381653020486,100.55381653030486,100.55904810852692,100.55904810862692,100.5642783227616,100.5642783228616,100.567763452403,100.567763452503,100.57298889233176,100.57298889243177,100.57821262715126,100.57821262725126,100.58169267935622,100.58169267945622,100.58517034440823,100.58517034450823,100.58864505393753,100.58864505403753,100.59211730053123,100.59211730063123,100.59905868663067,100.59905868673067,100.60426074761699,100.60426074771699,100.60772662846948,100.60772662856948,100.6129228163017,100.6129228164017,100.61465376743382,100.61465376753382,100.61984457469906,100.61984457479906,100.62157347970007,100.62157347980008,100.6267588306157,100.6267588307157,100.63021391244291,100.63021391254291,100.63366763018273,100.63366763028273,100.64057252694413,100.64057252704413,100.64229756703084,100.64229756713084,100.64574635889947,100.64574635899947,100.64919405191992,100.64919405201992,100.6543634884825,100.6543634885825,100.65608470154622,100.65608470164622,100.65952481630332,100.65952481640332,100.66468015729608,100.66468015739608,100.66811481570356,100.66811481580356,100.6715466701536,100.6715466702536,100.67497439445012,100.67497439455012,100.68011003195816,100.68011003205817,100.69037604008122,100.69037604018122,100.69550694117467,100.69550694127467,100.69892568522911,100.69892568532912,100.70405188779928,100.70405188789928,100.70917452858568,100.70917452868568,100.71258747526866,100.71258747536866,100.71429284976196,100.71429284986196,100.71770238622643,100.71770238632644,100.72111029336429,100.72111029346429,100.72792212905173,100.72792212915174,100.73132495875315,100.73132495885315,100.73642486474948,100.73642486484948,100.73982072244861,100.73982072254861,100.74321430666872,100.74321430676872,100.74490941261519,100.74490941271519,100.74660376073531,100.74660376083531,100.74829746470306,100.74829746480306,100.75337728830152,100.75337728840152,100.75676215751878,100.75676215761878,100.76183773728977,100.76183773738977,100.76521999200618,100.76521999210618,100.76860016270017,100.76860016280017,100.77028829664434,100.77028829674434,100.77366278364082,100.77366278374082,100.77703382252746,100.77703382262746,100.7804016785435,100.7804016786435,100.78545073439275,100.78545073449276,100.78881427079868,100.78881427089868,100.79049512961005,100.79049512971005,100.79385563471064,100.79385563481064,100.79889525562201,100.79889525572202,100.80057425775885,100.80057425785886,100.80225253996068,100.80225254006068,100.80728571934822,100.80728571944822,100.81063826727228,100.81063826737228,100.81566481567938,100.81566481577939,100.81901444597206,100.81901444607206,100.8206878023027,100.8206878024027,100.82403322665922,100.82403322675923,100.82737705958044,100.82737705968044,100.83239023235272,100.83239023245272,100.83739889605832,100.83739889615832,100.84407106435219,100.8440710644522,100.84740478029182,100.84740478039183,100.84907017944596,100.84907017954596,100.85240003047129,100.85240003057129,100.85406315614641,100.85406315624641,100.85905139643226,100.85905139653227,100.86071285488944,100.86071285498944,100.86237351762898,100.86237351772898,100.86403228580267,100.86403228590267,100.86569003091081,100.86569003101081,100.87066125799545,100.87066125809545,100.87397311064532,100.87397311074533,100.87728303083803,100.87728303093803,100.88059098068229,100.88059098078229,100.88224342100608,100.88224342110608,100.88554682389231,100.88554682399231,100.89215040890285,100.89215040900285,100.89379989370394,100.89379989380394,100.89874709769376,100.89874709779376,100.90369221766115,100.90369221776115,100.90863529149742,100.90863529159742,100.91028166921053,100.91028166931054,100.91192744066258,100.91192744076258,100.91521811206638,100.91521811216639,100.9217965372426,100.9217965373426,100.92343988365037,100.92343988375038,100.92672475768272,100.92672475778272,100.93164881007799,100.93164881017799,100.9332884052454,100.93328840534541,100.93820597822551,100.93820597832551,100.94148262984211,100.94148262994212,100.94311972418262,100.94311972428262,100.94802941576884,100.94802941586885,100.95293611394104,100.95293611404104,100.95457059378069,100.95457059388069,100.9578384546118,100.9578384547118,100.95947073675508,100.95947073685508,100.96273348225843,100.96273348235843,100.96436239207452,100.96436239217452,100.96761922653246,100.96761922663246,100.97250152269666,100.97250152279666,100.97412812114243,100.97412812124243,100.97575388597923,100.97575388607923,100.97900439258727,100.97900439268727,100.98062845231482,100.98062845241482,100.9838754350304,100.9838754351304,100.98549820645317,100.98549820655317,100.98712006848434,100.98712006858435,100.99360528102133,100.99360528112133,100.9952250969214,100.9952250970214,101.00008359733869,101.00008359743869,101.00332080409454,101.00332080419454,101.0081744923146,101.0081744924146,101.01140836463458,101.01140836473458,101.01625606602659,101.01625606612659,101.0178694404028,101.0178694405028,101.021095203981,101.021095204081,101.02431941401521,101.02431941411521,101.02754146424436,101.02754146434437,101.03559367218587,101.03559367228587,101.03881235008784,101.03881235018784,101.04042045757103,101.04042045767103,101.04524341593319,101.04524341603319,101.04684917415473,101.04684917425473,101.05166512262329,101.05166512272329,101.05487410034817,101.05487410044817,101.0564777745473,101.0564777746473,101.0580806909201,101.0580806910201,101.06128553849145,101.06128553859145,101.0644892872146,101.0644892873146,101.06769148239378,101.06769148249379,101.0708921998116,101.0708921999116,101.07409049218515,101.07409049228515,101.07728613216653,101.07728613226654,101.08207711025734,101.08207711035735,101.0852688474331,101.0852688475331,101.08845899317355,101.08845899327355,101.09164758537,101.09164758547,101.0948346998051,101.0948346999051,101.1012059352613,101.1012059353613,101.10598255254307,101.10598255264307,101.10757387417293,101.10757387427293,101.11075519123658,101.11075519133658,101.11234514877904,101.11234514887904,101.11393450006042,101.11393450016043,101.12029042742465,101.12029042752465,101.12346609868209,101.12346609878209,101.12822642269765,101.12822642279765,101.13298470058211,101.13298470068212,101.13615563542496,101.13615563552496,101.1424946632619,101.1424946633619,101.14566249101678,101.14566249111678,101.15041266015945,101.15041266025945,101.15199284174219,101.15199284184219,101.1567324770988,101.1567324771988,101.16146862645427,101.16146862655427,101.16620182028703,101.16620182038703,101.16935570403734,101.16935570413735,101.17250795846104,101.17250795856104,101.17565778784045,101.17565778794045,101.17723020170325,101.17723020180325,101.1866616156834,101.18666161578341,101.18823202130643,101.18823202140644,101.19294145728358,101.19294145738358,101.19607855518056,101.19607855528056,101.19764589160692,101.19764589170693,101.20077938982882,101.20077938992883,101.20860894839305,101.20860894849305,101.21486484268104,101.21486484278104,101.21642778371465,101.21642778381465,101.21798992903061,101.21798992913061,101.22267469776055,101.22267469786055,101.2257949719129,101.2257949720129,101.22735374490168,101.22735374500168,101.2304703435963,101.2304703436963,101.23358489615981,101.23358489625981,101.23669766783144,101.23669766793144,101.24136318777246,101.24136318787247,101.24757675745963,101.24757675755963,101.25068005630207,101.25068005640208,101.2553315564559,101.2553315565559,101.26462970667502,101.26462970677503,101.26617805955169,101.2661780596517,101.26927245393469,101.26927245403469,101.27545544532923,101.27545544542923,101.27700016063947,101.27700016073948,101.28163226043914,101.28163226053914,101.28317511907487,101.28317511917487,101.2862596617155,101.2862596618155,101.28780055000277,101.28780055010277,101.29088069725067,101.29088069735067,101.29241889525444,101.29241889535444,101.29395553971422,101.29395553981422,101.29702795713351,101.29702795723351,101.30163432872908,101.30163432882908,101.30777423769646,101.30777423779647,101.31237655492114,101.31237655502115,101.32004283980147,101.32004283990148,101.32310737582686,101.32310737592687,101.32463837448046,101.32463837458046,101.32769885613496,101.32769885623496,101.33228707778981,101.33228707788982,101.33381440098567,101.33381440108568,101.33686810009448,101.33686810019448,101.33839405920293,101.33839405930293,101.34144514381087,101.34144514391087,101.34449448541825,101.34449448551825,101.34754231137296,101.34754231147296,101.35058801541393,101.35058801551394,101.35210961702096,101.35210961712096,101.35971595783818,101.35971595793818,101.3642769735276,101.3642769736276,101.36731547821837,101.36731547831837,101.37035235358253,101.37035235368253,101.37338733438087,101.37338733448087,101.37642068585257,101.37642068595257,101.38551793631024,101.38551793641024,101.38854840804187,101.38854840814187,101.3930922589648,101.3930922590648,101.39763326803896,101.39763326813896,101.40065923070391,101.40065923080391,101.40519523812426,101.40519523822427,101.40821627491802,101.40821627501802,101.4127462197277,101.4127462198277,101.41576437678138,101.41576437688138,101.41727237540569,101.41727237550569,101.42179481773465,101.42179481783465,101.42480532074235,101.42480532084235,101.42931461478439,101.42931461488439,101.43231878994219,101.43231879004219,101.4338199681295,101.4338199682295,101.43532042638178,101.43532042648178,101.43681989945982,101.43681989955982,101.4383187662768,101.4383187663768,101.43981672370217,101.43981672380217,101.44281116079156,101.44281116089157,101.4458038927717,101.44580389287171,101.45178678012245,101.45178678022245,101.45776640881995,101.45776640891995,101.45926016030916,101.45926016040916,101.46075269927624,101.46075269937624,101.4652281184811,101.4652281185811,101.46671842186049,101.4667184219605,101.4711880815852,101.4711880816852,101.47267531576793,101.47267531586793,101.47416152688511,101.47416152698511,101.4771326608147,101.4771326609147,101.47861690158341,101.47861690168341,101.48158454951185,101.48158454961185,101.48603230805548,101.48603230815549,101.48899419650378,101.48899419660378,101.49195422827756,101.49195422837757,101.49343301269666,101.49343301279666,101.49491039513703,101.49491039523703,101.49934034476179,101.49934034486179,101.50229105527164,101.50229105537164,101.50524009856356,101.50524009866356,101.50671304771987,101.50671304781987,101.5096575819451,101.50965758204511,101.51260033527844,101.51260033537844,101.51554070145882,101.51554070155882,101.51700917943975,101.51700917953976,101.51847674802909,101.51847674812909,101.52434524149454,101.52434524159455,101.52874370343507,101.52874370353507,101.5302087711975,101.5302087712975,101.5346017388971,101.5346017389971,101.54045424222687,101.54045424232687,101.54191472513997,101.54191472523998,101.54629446877001,101.54629446887002,101.54921210016035,101.54921210026035,101.55212677602798,101.55212677612798,101.55358316667886,101.55358316677886,101.55503895106868,101.55503895116868,101.55794976202199,101.55794976212199,101.56085901943132,101.56085901953132,101.56522102992513,101.56522103002513,101.56812714235515,101.56812714245515,101.57393732108412,101.57393732118412,101.57538889953778,101.57538889963779,101.57974177822425,101.57974177832425,101.58409151193142,101.58409151203142,101.58843878270301,101.58843878280301,101.59423259238314,101.59423259248314,101.5985757330012,101.5985757331012,101.6029160317705,101.6029160318705,101.60725424651739,101.60725424661739,101.61592609116182,101.61592609126183,101.62170497166314,101.62170497176314,101.62603663121222,101.62603663131222,101.6289222823312,101.6289222824312,101.63036370591198,101.63036370601198,101.63180448534037,101.63180448544037,101.63468490745764,101.63468490755764,101.64044397080029,101.6404439709003,101.6433213237209,101.6433213238209,101.64475876871342,101.64475876881342,101.64906955014696,101.64906955024696,101.65194144671797,101.65194144681797,101.65768164018488,101.65768164028488,101.66055031599396,101.66055031609396,101.66341789295484,101.66341789305484,101.66485048785881,101.66485048795882,101.66914694637462,101.66914694647463,101.67200940800775,101.67200940810775,101.6734394641935,101.67343946429351,101.67772853390255,101.67772853400255,101.68058648646898,101.68058648656898,101.68629875403545,101.68629875413545,101.6905797150027,101.6905797151027,101.6920053757957,101.6920053758957,101.69485586377274,101.69485586387275,101.70055384631279,101.70055384641279,101.7033997873318,101.7033997874318,101.70766574333763,101.70766574343763,101.71477169142574,101.71477169152574,101.72045266076456,101.72045266086457,101.72187150112055,101.72187150122055,101.72612680966637,101.72612680976637,101.7275434902173,101.7275434903173,101.72895948872453,101.72895948882453,101.73178985641236,101.73178985651236,101.73461832953436,101.73461832963436,101.7374455522429,101.7374455523429,101.7388583299882,101.7388583300882,101.740269970994,101.74026997109401,101.74591482990795,101.74591483000795,101.74873364074418,101.74873364084418,101.75436698069782,101.75436698079783,101.75577377161508,101.75577377171508,101.75858636827536,101.75858636837536,101.76139676723926,101.76139676733926,101.76561028166269,101.76561028176269,101.77262919339621,101.77262919349621,101.77403166470336,101.77403166480336,101.77683584949132,101.77683584959132,101.77963768501763,101.77963768511763,101.7810368218889,101.7810368219889,101.78523271685005,101.78523271695005,101.78802848976568,101.78802848986568,101.79082301226786,101.79082301236787,101.79501307196622,101.79501307206623,101.79780475261965,101.79780475271966,101.80059412190275,101.80059412200275,101.80616994283757,101.80616994293757,101.80895548509768,101.80895548519769,101.81173871598747,101.81173871608748,101.81312944098642,101.81312944108642,101.81451929448508,101.81451929458508,101.81868752878499,101.818687528885,101.82146367399855,101.82146367409855,101.82562531520934,101.82562531530934,101.83117014104705,101.83117014114706,101.8394745535929,101.8394745536929,101.84362399379968,101.84362399389968,101.84500592801315,101.84500592811315,101.8463871801829,101.8463871802829,101.84776752296106,101.84776752306107,101.84914710791288,101.84914710801289,101.85604249395378,101.85604249405378,101.86431197070557,101.86431197080557,101.86982193653186,101.86982193663187,101.87257527117274,101.87257527127274,101.87670269652449,101.87670269662449,101.87945284829475,101.87945284839475,101.88082709057092,101.88082709067092,101.8822006886947,101.8822006887947,101.88357318797028,101.88357318807029,101.88906125261549,101.88906125271549,101.89454484608531,101.89454484618531,101.89591408670766,101.89591408680766,101.89728256950367,101.89728256960368,101.90412297524395,101.90412297534395,101.90548944980019,101.90548944990019,101.9082216410863,101.9082216411863,101.91231738929724,101.91231738939724,101.91777366312769,101.91777366322769,101.92186179518397,101.92186179528397,101.92594719906545,101.92594719916545,101.93002972320686,101.93002972330686,101.93683125245614,101.93683125255615,101.94090972222665,101.94090972232665,101.9449860321921,101.9449860322921,101.9504179797972,101.9504179798972,101.95720427673714,101.95720427683715,101.95991691610382,101.95991691620382,101.96534037605394,101.96534037615395,101.96804945363684,101.96804945373684,101.97346207667039,101.97346207677039,101.98022472942868,101.98022472952869,101.98292812331405,101.98292812341406,101.98698071331522,101.98698071341522,101.99508070220716,101.99508070230716,101.99777917022136,101.99777917032137,101.99912719170632,101.99912719180632,102.00047407645178,102.00047407655178,102.00316686076847,102.00316686086848,102.00451200251337,102.00451200261337,102.00585653799719,102.00585653809719,102.00988715103463,102.00988715113463,102.01122850364784,102.01122850374784,102.01525184155247,102.01525184165247,102.01793155336489,102.01793155346489,102.0219469719843,102.02194697208431,102.0246207727513,102.0246207728513,102.02595659323227,102.02595659333227,102.02729154221295,102.02729154231295,102.02995977295639,102.02995977305639,102.0339595803533,102.0339595804533,102.03795783420624,102.03795783430624,102.03928963820763,102.03928963830764,102.04461541434317,102.04461541444317,102.04860749191147,102.04860749201147,102.04993709821649,102.04993709831649,102.05658190897967,102.05658190907967,102.05790874921856,102.05790874931856,102.06056148241343,102.06056148251344,102.06453802454187,102.06453802464188,102.06586232606254,102.06586232616255,102.06983333605874,102.06983333615874,102.07247830153366,102.07247830163367,102.07644393096288,102.07644393106288,102.08304937264795,102.08304937274795,102.08568971538224,102.08568971548225,102.08964825913522,102.08964825923522,102.09360456730049,102.09360456740049,102.09624055253336,102.09624055263336,102.10150990849824,102.10150990859825,102.10414312766498,102.10414312776498,102.108090910284,102.108090910384,102.11072147705858,102.11072147715858,102.1120359268369,102.1120359269369,102.11992061726701,102.11992061736701,102.12254728123595,102.12254728133595,102.12648559102578,102.12648559112579,102.13173387941855,102.13173387951855,102.13697868181016,102.13697868191016,102.13828843306521,102.13828843316521,102.14483525688325,102.14483525698326,102.15137863259147,102.15137863269148,102.15530015652797,102.15530015662797,102.1605223756948,102.1605223757948,102.1631292603964,102.1631292604964,102.16834079421191,102.16834079431192,102.17094483706475,102.17094483716475,102.173546416982,102.17354641708201,102.17614648124659,102.17614648134659,102.18264336430914,102.18264336440915,102.18524039726839,102.18524039736839,102.19043272018631,102.19043272028631,102.19432442365651,102.19432442375651,102.19691709911433,102.19691709921433,102.2008046345397,102.2008046346397,102.20468970702947,102.20468970712948,102.20727703981163,102.20727703991163,102.21244765100504,102.21244765110504,102.21503104309025,102.21503104319025,102.21890417981525,102.21890417991526,102.2201943033682,102.2201943034682,102.22406346150495,102.22406346160496,102.22535176627471,102.22535176637471,102.22792708750944,102.22792708760944,102.22921391451784,102.22921391461784,102.23178654546909,102.2317865455691,102.23950023238669,102.23950023248669,102.24335241521352,102.24335241531352,102.24720080890869,102.24720080900869,102.25104461775452,102.25104461785452,102.25616562921428,102.25616562931428,102.26000345123207,102.26000345133207,102.26255978891712,102.26255978901712,102.26639132097631,102.26639132107631,102.27149755482256,102.27149755492256,102.2740482467014,102.2740482468014,102.27659647564467,102.27659647574467,102.28296392196917,102.28296392206917,102.2867810174367,102.2867810175367,102.29059254288065,102.29059254298065,102.2931303896031,102.29313038970311,102.29693346528343,102.29693346538343,102.29819922471971,102.29819922481971,102.30072979630935,102.30072979640936,102.30325896592029,102.3032589660203,102.308315372685,102.308315372785,102.3133695059707,102.3133695060707,102.31589496223258,102.31589496233258,102.31841878916785,102.31841878926785,102.32220261605924,102.32220261615925,102.32472356325444,102.32472356335444,102.32724314636224,102.32724314646224,102.32976125170867,102.32976125180868,102.33479359748723,102.33479359758724,102.33730825472384,102.33730825482384,102.34359234015149,102.34359234025149,102.34861608460115,102.34861608470115,102.35112558861869,102.35112558871869,102.35363274337458,102.35363274347458,102.35488550608922,102.35488550618922,102.35738932640923,102.35738932650924,102.36114280235591,102.36114280245592,102.36614441827658,102.36614441837658,102.3686439568778,102.3686439569778,102.37364007855756,102.37364007865756,102.37863165327929,102.37863165337929,102.37987687562189,102.37987687572189,102.38361083754043,102.38361083764043,102.38485408953457,102.38485408963457,102.38858179938585,102.38858179948585,102.38982277790097,102.38982277800098,102.39230291614803,102.39230291624803,102.39973889760509,102.39973889770509,102.4034542169958,102.4034542170958,102.4071668082117,102.4071668083117,102.4084032018775,102.4084032019775,102.4121108293309,102.4121108294309,102.4158161833053,102.4158161834053,102.41704950777444,102.41704950787444,102.41951471684268,102.41951471694269,102.42074618463731,102.42074618473731,102.42320840029151,102.42320840039152,102.43059106866592,102.43059106876592,102.43550977838552,102.43550977848552,102.44042534312584,102.44042534322584,102.44288126882148,102.44288126892148,102.44779001312477,102.44779001322478,102.45269447570928,102.45269447580928,102.45636944084764,102.45636944094764,102.4624893810911,102.4624893811911,102.4637124370134,102.4637124371134,102.46982517790667,102.46982517800667,102.47471107374596,102.47471107384597,102.47715207026279,102.47715207036279,102.48325090504281,102.48325090514281,102.48934462449508,102.48934462459508,102.49178046779282,102.49178046789282,102.49421532591634,102.49421532601635,102.49664851682194,102.49664851692194,102.50759490142869,102.50759490152869,102.51124129651421,102.51124129661422,102.51609960747491,102.51609960757492,102.51852700100902,102.51852700110902,102.52216608307042,102.52216608317042,102.52459010427734,102.52459010437734,102.52822403311966,102.52822403321966,102.5318553474611,102.5318553475611,102.53548461567146,102.53548461577147,102.54153078152119,102.54153078162119,102.5451561469259,102.5451561470259,102.54998694879063,102.54998694889063,102.55240036042888,102.55240036052888,102.55601807178762,102.55601807188762,102.55963332021076,102.55963332031077,102.56204082080359,102.56204082090359,102.56805493059933,102.56805493069933,102.5728623156303,102.5728623157303,102.58006966088206,102.58006966098206,102.58247007579865,102.58247007589866,102.58846728606713,102.58846728616713,102.58966552317695,102.58966552327695,102.59685255868293,102.59685255878293,102.60164171799055,102.60164171809055,102.6052314276662,102.6052314277662,102.60762278655808,102.60762278665808,102.61359817142811,102.61359817152811,102.62076220690481,102.62076220700482,102.62792207433668,102.62792207443668,102.62911413516187,102.62911413526187,102.6350721279175,102.6350721280175,102.63983586215157,102.63983586225157,102.64221661147475,102.64221661157475,102.64816494194461,102.64816494204462,102.65173195472153,102.65173195482153,102.65529593619311,102.65529593629311,102.66004527172682,102.66004527182682,102.66241783652558,102.66241783662558,102.669531476551,102.669531476651,102.67308628832392,102.67308628842392,102.67900657921938,102.67900657931938,102.68137277827694,102.68137277837694,102.68728389947374,102.68728389957374,102.69200818884705,102.69200818894706,102.6967288785453,102.6967288786453,102.7002671318128,102.7002671319128,102.70380265690548,102.70380265700548,102.70497994011714,102.70497994021714,102.70851016042549,102.70851016052549,102.71086233969584,102.71086233979584,102.7167401165339,102.7167401166339,102.71791406530967,102.71791406540967,102.72612871332612,102.72612871342612,102.73316672228329,102.73316672238329,102.73668163449966,102.73668163459966,102.74370516897383,102.74370516907383,102.74604465465308,102.74604465475308,102.74955119288845,102.74955119298845,102.75539035847473,102.75539035857473,102.75889072042547,102.75889072052547,102.76122308253716,102.76122308263716,102.76704898616258,102.76704898626258,102.77403292421073,102.77403292431073,102.77752191876644,102.77752191886644,102.7798465131582,102.7798465132582,102.78333175647356,102.78333175657356,102.78565324377733,102.78565324387733,102.78681332433119,102.7868133244312,102.78913253815597,102.78913253825597,102.79492793927139,102.79492793937139,102.7972449932911,102.7972449933911,102.79840242145278,102.79840242155278,102.80071621681927,102.80071621691927,102.80649699188642,102.80649699198642,102.80880771805624,102.80880771815625,102.81342632854712,102.81342632864713,102.81688747298503,102.81688747308503,102.81804020262342,102.81804020272342,102.82149706534253,102.82149706544253,102.82495059362576,102.82495059372576,102.82840101482101,102.82840101492101,102.82954953852324,102.82954953862324,102.83414147352714,102.83414147362714,102.8398786356887,102.83987863578871,102.84331746214103,102.84331746224103,102.84675329517934,102.84675329527934,102.85361931544975,102.85361931554975,102.85704904798605,102.85704904808605,102.86047518084726,102.86047518094726,102.86617956701983,102.86617956711983,102.87073857446946,102.87073857456946,102.8752933002003,102.8752933003003,102.8787071941662,102.8787071942662,102.8809805594977,102.8809805595977,102.88438642050444,102.88438642060444,102.88665512520399,102.88665512530399,102.88892269316402,102.88892269326402,102.89458877121535,102.89458877131536,102.89911901915556,102.89911901925556,102.90364426544195,102.90364426554196,102.90703576563965,102.90703576573965,102.91042446187991,102.91042446197991,102.91606935868518,102.91606935878518,102.91945297748899,102.91945297758899,102.92283394390063,102.92283394400063,102.92621210635482,102.92621210645483,102.92958754063422,102.92958754073422,102.93296081510854,102.93296081520855,102.93633166453861,102.93633166463862,102.93969883851095,102.93969883861095,102.94530883741348,102.94530883751348,102.94755094240865,102.94755094250866,102.94979194855564,102.94979194865564,102.9542716494793,102.9542716495793,102.95650939697305,102.95650939707305,102.95874464363989,102.95874464373989,102.9609768968927,102.9609768969927,102.96209211412751,102.96209211422752,102.97101044178746,102.97101044188746,102.97546627118156,102.97546627128156,102.97880612551451,102.97880612561451,102.98214351691188,102.98214351701188,102.98547753598206,102.98547753608206,102.99103050845287,102.99103050855287,102.99435945008659,102.9943594501866,102.99768714130687,102.99768714140687,102.99879546232202,102.99879546242202,103.00101115706943,103.00101115716943,103.00211777297532,103.00211777307533,103.00654219046783,103.00654219056783,103.00875326247458,103.00875326257459,103.01538268936315,103.01538268946315,103.02090265849706,103.02090265859707,103.02531244994131,103.02531245004131,103.03192274171492,103.03192274181492,103.03742876684427,103.03742876694427,103.0418297296117,103.04182972971171,103.04622694113877,103.04622694123877,103.04842209879249,103.04842209889249,103.05281021640354,103.05281021650354,103.05609959950257,103.05609959960258,103.05938667123128,103.05938667133128,103.06157694090514,103.06157694100514,103.06923883039272,103.06923883049272,103.0757999771285,103.0757999772285,103.07798513147459,103.0779851315746,103.08016869438536,103.08016869448537,103.0867152529642,103.0867152530642,103.08998580407881,103.08998580417881,103.09216482003157,103.09216482013157,103.09760870340145,103.09760870350145,103.10522406235205,103.10522406245205,103.10848328396294,103.10848328406294,103.11174008052957,103.11174008062957,103.11282456790649,103.11282456800649,103.11824541335577,103.11824541345577,103.12041219041322,103.12041219051322,103.12474377417968,103.12474377427968,103.12907244031473,103.12907244041473,103.13339758255734,103.13339758265734,103.1377199208425,103.1377199209425,103.14095943896868,103.14095943906868,103.14419713831165,103.14419713841166,103.14851091315923,103.14851091325923,103.15282127778829,103.1528212778883,103.1592829977089,103.1592829978089,103.16143548973994,103.16143548983995,103.16573774562723,103.16573774572723,103.16896134940036,103.16896134950036,103.17218362697743,103.17218362707743,103.17540355529285,103.17540355539285,103.17969433011115,103.17969433021115,103.18291031772962,103.18291031782962,103.1861238424125,103.1861238425125,103.19468966715266,103.19468966725266,103.19896971872832,103.19896971882832,103.20645310222265,103.20645310232266,103.21179388331963,103.21179388341963,103.2139286043231,103.2139286044231,103.22032984970217,103.22032984980217,103.2277922035157,103.2277922036157,103.23312021523893,103.23312021533893,103.23737988128617,103.23737988138618,103.23844386946072,103.23844386956073,103.24057086063559,103.24057086073559,103.24269633615778,103.24269633625778,103.2448207128318,103.2448207129318,103.2501287937224,103.2501287938224,103.25331105806896,103.25331105816896,103.25755158900124,103.25755158910124,103.2607302536727,103.2607302537727,103.26496506301616,103.26496506311616,103.26813880181645,103.26813880191645,103.27448271763326,103.27448271773326,103.27976563868954,103.27976563878954,103.28504442959229,103.28504442969229,103.28820884712866,103.28820884722866,103.29242512761822,103.29242512771822,103.2945318090473,103.2945318091473,103.3008494661816,103.3008494662816,103.30190129124347,103.30190129134347,103.30610707583828,103.30610707593829,103.30715730946486,103.30715730956486,103.31135650097055,103.31135650107055,103.31765225692376,103.31765225702377,103.32289535420612,103.32289535430613,103.32603775688746,103.32603775698746,103.33127104031878,103.33127104041878,103.33231653753076,103.33231653763076,103.33440658467178,103.33440658477178,103.33753936295867,103.33753936305867,103.34693129418676,103.34693129428676,103.35214381317651,103.35214381327651,103.35318518023495,103.35318518033495,103.35526704285157,103.35526704295157,103.35734727614154,103.35734727624154,103.35942588010491,103.35942588020491,103.36358047353079,103.36358047363079,103.36669377568084,103.36669377578085,103.3708433674529,103.37084336755291,103.37395344884105,103.37395344894105,103.3760256870632,103.3760256871632,103.37809652330664,103.37809652340664,103.38223584653186,103.38223584663186,103.38326967321827,103.38326967331827,103.38740407057232,103.38740407067232,103.39153581553418,103.39153581563419,103.39463191502644,103.39463191512644,103.39978835486738,103.39978835496738,103.40494009618504,103.40494009628505,103.40905971592555,103.40905971602555,103.41008869252342,103.41008869262342,103.41420334850143,103.41420334860143,103.41831516263069,103.41831516273069,103.42447837475788,103.42447837485788,103.42653083371269,103.42653083381269,103.4296080633292,103.4296080634292,103.43165764254393,103.43165764264393,103.43780418249175,103.43780418259175,103.44189958968084,103.44189958978085,103.4439458344597,103.4439458345597,103.45008062809931,103.45008062819932,103.45212304585519,103.45212304595519,103.45620522897474,103.45620522907474,103.45926548328134,103.45926548338134,103.46232342621762,103.46232342631762,103.47047202966895,103.47047202976896,103.47250747542253,103.47250747552253,103.47555806744336,103.47555806754336,103.4796221385135,103.47962213861351,103.48571411496519,103.48571411506519,103.48977238866387,103.48977238876387,103.49281331839892,103.49281331849892,103.4968650368998,103.49686503699981,103.49888866056256,103.49888866066256,103.50596728901128,103.50596728911128,103.51304140839332,103.51304140849332,103.51607116018994,103.51607116028994,103.52414530720384,103.52414530730384,103.52717138354274,103.52717138364274,103.53120301964573,103.53120301974573,103.53523200335655,103.53523200345656,103.54127150033453,103.54127150043453,103.54428711866998,103.54428711876999,103.54730008461328,103.54730008471329,103.5543257788925,103.5543257789925,103.56034432197231,103.56034432207231,103.56234862106352,103.56234862116352,103.56936097759925,103.56936097769925,103.57236337186517,103.57236337196517,103.57536406102182,103.57536406112182,103.57736297954604,103.57736297964604,103.57836156730787,103.57836156740787,103.5853488397919,103.5853488398919,103.590336511708,103.590336511808,103.59731321251466,103.59731321261467,103.60030075338442,103.60030075348442,103.60328507349223,103.60328507359223,103.60726135038145,103.60726135048145,103.61222829574729,103.61222829584729,103.61619854791714,103.61619854801714,103.62214797723519,103.62214797733519,103.62710022077012,103.62710022087012,103.6310585750665,103.6310585751665,103.63699231737941,103.63699231747941,103.63995613828487,103.63995613838487,103.64587934681172,103.64587934691173,103.64982481806041,103.64982481816041,103.65081014386138,103.65081014396138,103.65277977239774,103.65277977249774,103.65770079348768,103.65770079358768,103.66065142821489,103.66065142831489,103.66261677503248,103.66261677513248,103.67047429738848,103.67047429748848,103.67636302481708,103.67636302491708,103.67832382467664,103.67832382477664,103.68420107103626,103.68420107113626,103.68909503772603,103.68909503782604,103.69105091171441,103.69105091181441,103.69495981784243,103.69495981794243,103.6988665641654,103.6988665642654,103.70179488090706,103.70179488100706,103.70862342666518,103.70862342676519,103.71154696910094,103.71154696920094,103.71544291639862,103.71544291649862,103.71933530191254,103.71933530201254,103.7222525543897,103.7222525544897,103.72613963511927,103.72613963521927,103.7271104485459,103.7271104486459,103.7280804662549,103.7280804663549,103.73583897860021,103.73583897870022,103.7397157150003,103.7397157151003,103.7484306799632,103.74843068006321,103.75326913587392,103.75326913597392,103.7590698417737,103.7590698418737,103.76196837594038,103.76196837604039,103.76486463662806,103.76486463672806,103.77161849860086,103.77161849870086,103.77643925926665,103.77643925936665,103.78125600345287,103.78125600355287,103.78221851868113,103.78221851878114,103.78606732918078,103.78606732928078,103.78991341150561,103.78991341160561,103.7937566519817,103.7937566520817,103.79855736814092,103.79855736824092,103.80143547888787,103.80143547898787,103.80431207398215,103.80431207408215,103.80814409862846,103.80814409872846,103.8148455569102,103.8148455570102,103.81771559680668,103.81771559690668,103.8253635422897,103.8253635423897,103.83300713027128,103.83300713037129,103.83587050129601,103.83587050139602,103.84159349210512,103.84159349220512,103.84635870410054,103.84635870420054,103.84921472420982,103.84921472430982,103.85301859560778,103.85301859570778,103.85681992828754,103.85681992838754,103.85966878693795,103.85966878703795,103.86441319660047,103.86441319670047,103.86820562482077,103.86820562492078,103.87578517647704,103.87578517657704,103.87957214834773,103.87957214844774,103.88335661939152,103.88335661949152,103.88713840015183,103.88713840025183,103.89186170435092,103.89186170445092,103.89469301932166,103.89469301942167,103.89752190924814,103.89752190934814,103.90129099641733,103.90129099651733,103.90411586986424,103.90411586996424,103.90882117568786,103.90882117578786,103.91070194908644,103.91070194918645,103.9154007754949,103.9154007755949,103.92291420680343,103.92291420690343,103.92948300758519,103.92948300768519,103.93323356590048,103.93323356600048,103.93604415432097,103.93604415442097,103.94072467922344,103.94072467932344,103.94727214719387,103.94727214729387,103.95194369185427,103.95194369195427,103.95567826003389,103.9556782601339,103.96127651262822,103.96127651272822,103.9668707866343,103.9668707867343,103.97152876620329,103.9715287663033,103.9771126958799,103.9771126959799,103.98455352742552,103.98455352752552,103.9873414947299,103.9873414948299,103.99105499533741,103.99105499543741,103.99569323353037,103.99569323363038,104.0021828035688,104.0021828036688,104.00774017143235,104.00774017153235,104.0105172260375,104.01051722613751,104.01514178546513,104.01514178556513,104.02068736912916,104.02068736922917,104.02345764118863,104.02345764128863,104.02530302409735,104.02530302419736,104.0289915164358,104.0289915165358,104.03451792709356,104.03451792719356,104.03819959899498,104.03819959909498,104.04739680674626,104.04739680684627,104.0510730601894,104.0510730602894,104.05658370805938,104.05658370815938,104.0611718539316,104.0611718540316,104.06758935468555,104.06758935478555,104.07308624800756,104.07308624810756,104.07949454117154,104.07949454127154,104.08498290894728,104.08498290904728,104.08772476251916,104.08772476261916,104.09228952944895,104.09228952954895,104.0977648247204,104.0977648248204,104.10050004731185,104.10050004741186,104.10870154704138,104.10870154714138,104.11325452977165,104.11325452987165,104.11507422236761,104.11507422246761,104.12325814052616,104.12325814062616,104.12689233460769,104.12689233470769,104.13052201962252,104.13052201972252,104.13324219936122,104.13324219946122,104.13686642802645,104.13686642812645,104.13867714038034,104.13867714048034,104.14229621582648,104.14229621592648,104.14591260098913,104.14591260108914,104.15223666175596,104.15223666185597,104.15313839931216,104.15313839941216,104.15854693008349,104.15854693018349,104.16304955525378,104.16304955535378,104.16844861319592,104.16844861329592,104.17294403901603,104.17294403911603,104.17653685577966,104.17653685587966,104.1801267549119,104.1801267550119,104.18460971448879,104.18460971458879,104.18908930173849,104.1890893018385,104.19356445570413,104.19356445580414,104.19535345633354,104.19535345643354,104.19803483536387,104.19803483546387,104.20339399374947,104.20339399384947,104.2087475821115,104.2087475822115,104.21409578990655,104.21409579000655,104.21765867252995,104.21765867262995,104.22299790008293,104.22299790018293,104.22655411383457,104.22655411393457,104.23188481584128,104.23188481594129,104.23454721132192,104.23454721142193,104.23720801536727,104.23720801546727,104.24075316485442,104.24075316495443,104.24518186406573,104.24518186416573,104.25403319987764,104.25403319997764,104.25845481341271,104.25845481351271,104.26552502998912,104.26552503008912,104.26817363303047,104.26817363313047,104.27346830039494,104.27346830049494,104.28052252683568,104.28052252693568,104.28316537039689,104.28316537049689,104.28580601626172,104.28580601636172,104.29196100597318,104.29196100607318,104.29547565295033,104.29547565305033,104.30337675032061,104.30337675042061,104.30951734133168,104.30951734143169,104.31477699711948,104.31477699721948,104.3182797841145,104.3182797842145,104.32178090389156,104.32178090399157,104.32527823453697,104.32527823463697,104.33052030875379,104.33052030885379,104.3348850474224,104.3348850475224,104.3383740040868,104.3383740041868,104.34098877018228,104.34098877028228,104.34447204314917,104.34447204324917,104.34708226228663,104.34708226238664,104.35316844136686,104.35316844146686,104.35664265830904,104.35664265840904,104.36358537060457,104.36358537070457,104.37138878416023,104.37138878426023,104.37398706753294,104.37398706763294,104.38177740858433,104.38177740868433,104.38696552556607,104.38696552566607,104.39214848932875,104.39214848942875,104.39904967274111,104.39904967284112,104.40422164802192,104.40422164812192,104.40938930369263,104.40938930379264,104.41455210927478,104.41455210937478,104.41971131518186,104.41971131528186,104.42314843652493,104.42314843662493,104.42658279180188,104.42658279190188,104.43258898231242,104.43258898241243,104.43601788123976,104.43601788133977,104.44030164616446,104.44030164626446,104.4445814325009,104.4445814326009,104.44971354611648,104.44971354621649,104.45398643623328,104.45398643633328,104.46252467609483,104.46252467619483,104.46679006373091,104.46679006383091,104.46934761393808,104.46934761403809,104.47616399658354,104.47616399668354,104.47956978180765,104.47956978190766,104.48297272518302,104.48297272528302,104.48722155431366,104.48722155441367,104.48807031980904,104.48807031990904,104.49400993527604,104.49400993537604,104.50333430628636,104.50333430638636,104.50926482783734,104.50926482793734,104.51519080243031,104.51519080253031,104.52364799276532,104.52364799286532,104.5312544851478,104.5312544852478,104.53716511375748,104.53716511385748,104.54053888081894,104.54053888091894,104.54643817992492,104.54643818002492,104.553176165436,104.553176165536,104.55738354146612,104.55738354156613,104.56074632004571,104.56074632014571,104.56242630735679,104.56242630745679,104.56494452637718,104.56494452647718,104.5716552301402,104.5716552302402,104.57416931900705,104.57416931910706,104.57919450332675,104.57919450342675,104.58505265246272,104.58505265256272,104.59257930784081,104.59257930794081,104.59675777279624,104.59675777289624,104.60093237283736,104.60093237293736,104.60510420681236,104.60510420691236,104.60927327472123,104.60927327482123,104.61343900819423,104.61343900829424,104.61676878343694,104.61676878353694,104.62342481002987,104.62342481012988,104.62841236827204,104.62841236837204,104.63256483978417,104.63256483988417,104.63920392895852,104.63920392905852,104.64417894517483,104.64417894527483,104.64666359248857,104.64666359258857,104.64997442207289,104.64997442217289,104.65493780565495,104.65493780575495,104.65989614969189,104.65989614979189,104.66485078037978,104.66485078047978,104.67227440926759,104.67227440936759,104.674746097751,104.674746097851,104.67803983835147,104.67803983845147,104.68215354704655,104.68215354714656,104.68955235712188,104.68955235722188,104.69201597475482,104.69201597485483,104.69529671863364,104.69529671873364,104.70349029907797,104.70349029917797,104.7092208302591,104.7092208303591,104.71658326467035,104.71658326477035,104.72394065953647,104.72394065963647,104.72802409306948,104.72802409316948,104.73128907416051,104.73128907426052,104.73291980275981,104.73291980285981,104.73454943251092,104.73454943261092,104.74106245727441,104.74106245737441,104.7467573706178,104.7467573707178,104.74838249130221,104.74838249140221,104.75650195633096,104.75650195643097,104.76218057640814,104.76218057650814,104.76947571584083,104.76947571594083,104.77595482787582,104.77595482797582,104.78080965283537,104.78080965293537,104.78808421728306,104.78808421738306,104.79050660916337,104.79050660926337,104.79615510564372,104.79615510574372,104.80099079548832,104.80099079558832,104.80662864450866,104.80662864460867,104.81145766548151,104.81145766558151,104.81467444881764,104.81467444891764,104.81708437445472,104.81708437455472,104.8219016870107,104.8219016871107,104.82591286180265,104.82591286190265,104.83152452792311,104.83152452802311,104.84033255395997,104.84033255405997,104.84673232157768,104.84673232167768,104.8507287187561,104.8507287188561,104.8571167021743,104.8571167022743,104.86110567265462,104.86110567275462,104.86588793574258,104.86588793584258,104.87145795930593,104.87145795940593,104.8778187367587,104.8778187368587,104.88338000742787,104.88338000752788,104.89211232661387,104.89211232671387,104.89607791815175,104.89607791825175,104.90241800694557,104.90241800704557,104.90796188550036,104.90796188560036,104.91350083818396,104.91350083828397,104.92061750951473,104.92061750961473,104.92851800062392,104.92851800072393,104.93325396052282,104.93325396062282,104.94272053764493,104.94272053774493,104.94744910873705,104.94744910883705,104.95217389069748,104.95217389079748,104.95532137081527,104.95532137091527,104.96003751355549,104.9600375136555,104.96160807074376,104.96160807084377,104.96553141346347,104.96553141356347,104.97023527941707,104.97023527951707,104.98041879814355,104.98041879824355,104.98433194809904,104.98433194819904,104.98902387828785,104.98902387838785,104.99449398244892,104.99449398254892,105.0023021703105,105.0023021704105,105.00698303623481,105.00698303633482,105.0124387038042,105.0124387039042,105.0225621269023,105.0225621270023,105.02800873844697,105.02800873854697,105.03500533219493,105.03500533229493,105.04277157436886,105.04277157446886,105.04742587848013,105.04742587858013,105.05440117730807,105.05440117740807,105.0605963697066,105.0605963698066,105.06446397429936,105.06446397439936,105.06909997690464,105.06909997700464,105.07295977588613,105.07295977598613,105.07604541737497,105.07604541747497,105.08221128189433,105.08221128199433,105.0852908986638,105.0852908987638,105.08913754935838,105.08913754945839,105.0983603716398,105.0983603717398,105.1052709520987,105.10527095219871,105.11140798132602,105.11140798142603,105.1160066609843,105.1160066610843,105.12060177885878,105.12060177895879,105.12595798172168,105.12595798182168,105.1305454455502,105.1305454456502,105.13360077398562,105.13360077408562,105.13741631590915,105.13741631600915,105.14351734838554,105.14351734848555,105.15265984107542,105.15265984117542,105.15798853484235,105.15798853494235,105.16483337386668,105.16483337396669,105.16863292565455,105.16863292575455,105.17470701740471,105.17470701750472,105.1800168412959,105.1800168413959,105.18456387508944,105.18456387518944,105.18759302062497,105.18759302072498,105.1936457416725,105.1936457417725,105.19893726405769,105.19893726415769,105.2057347768274,105.2057347769274,105.20799802517732,105.20799802527732,105.21327537621004,105.21327537631004,105.22005508006087,105.22005508016088,105.22532424656917,105.22532424666917,105.23209212832917,105.23209212842917,105.23735106418195,105.23735106428195,105.24335539801798,105.24335539811798,105.24935291141699,105.249352911517,105.25159982860939,105.25159982870939,105.25758752815736,105.25758752825736,105.26132675696894,105.26132675706894,105.26804875234436,105.26804875244436,105.27476502613094,105.27476502623094,105.28147663928556,105.28147663938556,105.28594527594471,105.28594527604471,105.2904101234722,105.2904101235722,105.29858881262905,105.29858881272905,105.30230246480181,105.30230246490181,105.30898232503299,105.30898232513299,105.32010657148061,105.32010657158061,105.32677460962095,105.32677460972096,105.32973486874263,105.32973486884264,105.33343169717077,105.33343169727077,105.34008022128299,105.34008022138299,105.34450763218952,105.34450763228952,105.34819225961365,105.34819225971366,105.35408310895599,105.35408310905599,105.3607042755375,105.3607042756375,105.36732104672629,105.36732104682629,105.37099347314643,105.37099347324643,105.37466229989147,105.37466229999147,105.381261337944,105.381261338044,105.38638697214442,105.38638697224442,105.39077660540815,105.39077660550815,105.39370003416994,105.39370003426994,105.40392121686527,105.40392121696527,105.41121639418927,105.41121639428927,105.41631849788199,105.41631849798199,105.42214466674662,105.42214466684662,105.42723710815356,105.42723710825356,105.43087190849616,105.43087190859616,105.43740800907156,105.43740800917156,105.44466226377347,105.44466226387347,105.44900995134954,105.44900995144954,105.45407791496585,105.45407791506585,105.46131364081386,105.46131364091386,105.46709468112023,105.46709468122023,105.47359323140073,105.47359323150073,105.47647804891075,105.47647804901075,105.4844055565288,105.4844055566288,105.4908855779554,105.4908855780554,105.4987987247644,105.4987987248644,105.50598666966198,105.50598666976198,105.51316855194888,105.51316855204888,105.52321448730422,105.52321448740422,105.52751454549512,105.52751454559512,105.53395731975735,105.53395731985735,105.54396983497129,105.5439698350713,105.55040010509899,105.55040010519899,105.55468375634975,105.55468375644975,105.56038999919684,105.56038999929685,105.56680431708018,105.56680431718019,105.57250104920678,105.57250104930678,105.57819209763585,105.57819209773585,105.58529910668084,105.58529910678084,105.58955945477177,105.58955945487178,105.59381620318763,105.59381620328763,105.59806946560235,105.59806946570235,105.60019384227635,105.60019384237636,105.60514676785499,105.60514676795499,105.60938767770044,105.60938767780044,105.61291835270458,105.61291835280458,105.61785775108315,105.61785775118315,105.6263148277442,105.6263148278442,105.63265040747132,105.63265040757132,105.64109138032276,105.64109138042276,105.64530758502968,105.64530758512969,105.6495199248223,105.6495199249223,105.65442973008247,105.65442973018247,105.6600363566578,105.6600363567578,105.66843898349669,105.66843898359669,105.67263643200181,105.67263643210181,105.67682982613604,105.67682982623604,105.68311311584604,105.68311311594604,105.69078527388046,105.69078527398047,105.69426862262998,105.69426862272998,105.70053410342109,105.7005341035211,105.7060976854606,105.70609768556061,105.7095722055333,105.70957220563331,105.7151271862439,105.7151271863439,105.7179022705006,105.7179022706006,105.72413838552123,105.72413838562123,105.73244700400323,105.73244700410324,105.74005603510393,105.74005603520393,105.74351138217037,105.74351138227037,105.747654342962,105.747654343062,105.75455279819955,105.75455279829956,105.76075579620934,105.76075579630934,105.76419901805441,105.76419901815441,105.76901511808823,105.76901511818824,105.77245193630077,105.77245193640077,105.77794553307822,105.77794553317823,105.78137530350585,105.78137530360586,105.78480166371496,105.78480166381496,105.79370119728179,105.7937011973818,105.7998574374067,105.7998574375067,105.80532435869715,105.80532435879715,105.8114682462528,105.8114682463528,105.8189691734268,105.8189691735268,105.82509862439076,105.82509862449076,105.83258352353776,105.83258352363777,105.8373424076833,105.8373424077833,105.84481381752153,105.84481381762153,105.8556702481423,105.8556702482423,105.85973708527857,105.85973708537857,105.86650940032264,105.86650940042264,105.87259561729418,105.87259561739418,105.88137905186646,105.88137905196646,105.88880461321143,105.88880461331144,105.89554760037633,105.89554760047633,105.90363284954604,105.90363284964604,105.90699854575703,105.90699854585704,105.91103302370878,105.91103302380878,105.91707888642797,105.91707888652797,105.92379065114787,105.92379065124787,105.92848458957647,105.92848458967647,105.93183372728203,105.93183372738203,105.93718864184015,105.93718864194015,105.94320680600678,105.94320680610679,105.94788331442969,105.94788331452969,105.95322246620003,105.95322246630003,105.95855672999052,105.95855673009052,105.96321849869119,105.96321849879119,105.96720962897658,105.96720962907658,105.97319024284832,105.97319024294832,105.98182143010987,105.98182143020988,105.98845396408642,105.98845396418642,105.99508210267022,105.99508210277023,106.00037866460053,106.00037866470053,106.00699422326718,106.00699422336719,106.01228127447699,106.01228127457699,106.0175629072285,106.0175629073285,106.0208601338301,106.0208601339301,106.0228366206948,106.0228366207948,106.02876176167881,106.02876176177881,106.03336517775168,106.03336517785168,106.03927925447118,106.03927925457118,106.04321779160891,106.04321779170891,106.04780809728618,106.04780809738618,106.05632443596664,106.05632443606665,106.06483228699216,106.06483228709216,106.07267719153968,106.07267719163968,106.07790107792445,106.07790107802445,106.08051023610506,106.08051023620506,106.0889822040536,106.0889822041536,106.09874736475176,106.09874736485176,106.10525038620763,106.10525038630763,106.11109694060067,106.11109694070068,106.11499027339751,106.11499027349751,106.12017710207449,106.1201771021745,106.12276689779223,106.12276689789223,106.12729468279684,106.12729468289685,106.13181837553927,106.13181837563927,106.1363362709102,106.13633627101021,106.13762343894047,106.13762343904047,106.13826482525921,106.13826482535922,106.13890249822893,106.13890249832893,106.13951232108082,106.13951232118082,106.14006936132847,106.14006936142847,106.14055876557572,106.14055876567572,106.14099777437167,106.14099777447167,106.14141260850752,106.14141260860752,106.14174525637733,106.14174525647734,106.14206164887227,106.14206164897227,106.14235920938276,106.14235920948276,106.1429437966177,106.1429437967177,106.14322657951466,106.14322657961466,106.14349750242947,106.14349750252947,106.1437584978193,106.14375849791931,106.14400729220513,106.14400729230513,106.14423028260424,106.14423028270424,106.14443720708506,106.14443720718506,106.14462507223354,106.14462507233354,106.14478216963282,106.14478216973282],[0.0,1e-10,0.1734449637392148,0.17344496383921482,0.32170053792438147,0.3217005380243815,0.4647535964274272,0.4647535965274272,0.5942955207663521,0.5942955208663521,0.7203755049497284,0.7203755050497284,0.8432774307227031,0.8432774308227031,0.9650101441344918,0.9650101442344918,1.0834716622792893,1.0834716623792893,1.198656074111681,1.198656074211681,1.31343326796288,1.31343326806288,1.4231361703478653,1.4231361704478653,1.5311783342106866,1.5311783343106866,1.6368277750942222,1.6368277751942222,1.7416111720419483,1.7416111721419483,1.8461077317218202,1.8461077318218202,1.9500300106049155,1.9500300107049156,2.0504479489490106,2.0504479490490106,2.1472445004766416,2.1472445005766416,2.2425915196817146,2.2425915197817146,2.337903792549321,2.337903792649321,2.4319286700387748,2.431928670138775,2.5256917943120674,2.5256917944120674,2.6191262871951224,2.6191262872951224,2.712102257254095,2.712102257354095,2.8048167014448078,2.8048167015448078,2.8968304594060403,2.8968304595060403,2.9885721577129543,2.9885721578129543,3.0797272605451456,3.0797272606451456,3.1707569052275524,3.1707569053275524,3.2597468982202202,3.2597468983202202,3.3485931315571102,3.3485931316571103,3.436408645295563,3.436408645395563,3.523580953932057,3.523580954032057,3.6106945689188934,3.6106945690188934,3.697636157327651,3.697636157427651,3.783775510778019,3.783775510878019,3.869896259591859,3.869896259691859,3.9558227017333665,3.9558227018333665,4.040891852833236,4.040891852933236,4.125896399238034,4.125896399338034,4.209590133430923,4.209590133530923,4.292754260689492,4.292754260789492,4.375831882071903,4.375831882171903,4.457834375232639,4.457834375332639,4.538877157122539,4.538877157222539,4.61964170307349,4.61964170317349,4.700274993503193,4.700274993603193,4.780833107560449,4.780833107660449,4.860523737811858,4.860523737911858,4.938854789726692,4.938854789826692,5.017168222179234,5.017168222279234,5.095428265766464,5.095428265866464,5.17349028933232,5.17349028943232,5.251448339124993,5.251448339224993,5.3292438730600935,5.3292438731600935,5.406751319313886,5.406751319413886,5.483333156481928,5.483333156581928,5.559876117158975,5.559876117258975,5.636385240890158,5.636385240990158,5.7117350797851465,5.7117350798851465,5.786210803893855,5.786210803993855,5.860347324934951,5.860347325034951,5.93431951133527,5.93431951143527,6.0080164552106625,6.0080164553106625,6.080516904976803,6.080516905076803,6.152815659263844,6.152815659363844,6.225050149877664,6.225050149977664,6.297057709395423,6.297057709495423,6.368906390622018,6.368906390722018,6.43986383018716,6.43986383028716,6.510774170845573,6.510774170945573,6.581484824264673,6.581484824364673,6.652081159581116,6.652081159681116,6.7226021290685285,6.7226021291685285,6.7931120721827645,6.7931120722827645,6.863579046543798,6.863579046643798,6.933406832282448,6.933406832382448,7.003190929332877,7.003190929432877,7.072872227240944,7.072872227340944,7.14200826910087,7.14200826920087,7.2107909365407155,7.2107909366407155,7.278040256065336,7.278040256165336,7.344848672228165,7.344848672328165,7.410819538725387,7.410819538825387,7.476493488864498,7.476493488964498,7.54216531708987,7.54216531718987,7.607616466286446,7.607616466386446,7.67197267010268,7.67197267020268,7.736247900175004,7.736247900275004,7.800249327592439,7.800249327692439,7.864136550581167,7.864136550681167,7.928011762022482,7.928011762122482,7.991745449395708,7.991745449495708,8.05525841984882,8.05525841994882,8.118203816268224,8.118203816368224,8.180687696449521,8.180687696549521,8.242909709740706,8.242909709840706,8.305109101915788,8.305109102015788,8.367142530123472,8.367142530223472,8.428896358304803,8.428896358404803,8.490464253794803,8.490464253894803,8.551696621674916,8.551696621774916,8.612694821217469,8.612694821317469,8.673499812935875,8.673499813035875,8.734166539239432,8.734166539339432,8.794701593217257,8.794701593317257,8.85519307218081,8.85519307228081,8.91548944875438,8.91548944885438,8.975157094708994,8.975157094808994,9.034478603485026,9.034478603585026,9.093766616337046,9.093766616437046,9.152846189055582,9.152846189155582,9.211701824092081,9.211701824192081,9.270544121385083,9.270544121485083,9.329271986901475,9.329271987001475,9.387902736973018,9.387902737073018,9.446508819797353,9.446508819897353,9.504906348814254,9.504906348914254,9.563001921927915,9.563001922027915,9.621074722360206,9.621074722460206,9.679098870341779,9.679098870441779,9.7370874004856,9.7370874005856,9.794875485563773,9.794875485663773,9.852642389395879,9.852642389495879,9.910324606135033,9.910324606235033,9.96779163808632,9.96779163818632,10.02519952169215,10.02519952179215,10.082500892806573,10.082500892906573,10.139583176327505,10.139583176427506,10.196546670570402,10.196546670670402,10.253345792281205,10.253345792381205,10.309889147603895,10.309889147703895,10.365987242063367,10.365987242163367,10.421701118570919,10.421701118670919,10.477138464248775,10.477138464348775,10.53246024141052,10.53246024151052,10.587240400090533,10.587240400190533,10.641887029770293,10.641887029870293,10.696279219258027,10.696279219358027,10.75063810227833,10.75063810237833,10.80448647558796,10.80448647568796,10.85823845338776,10.85823845348776,10.91188524489224,10.91188524499224,10.965503314778617,10.965503314878617,11.019048140749703,11.019048140849703,11.072519950153398,11.072519950253398,11.125844513900171,11.125844514000171,11.178964123514604,11.178964123614604,11.231841721288907,11.231841721388907,11.28456601079561,11.28456601089561,11.337231568761338,11.337231568861338,11.389734951891343,11.389734951991343,11.442070021792311,11.442070021892311,11.494233481919686,11.494233482019686,11.546304449342855,11.546304449442855,11.59820824027649,11.59820824037649,11.65007868685138,11.65007868695138,11.701858686853168,11.701858686953168,11.75342395676289,11.75342395686289,11.804778437277493,11.804778437377493,11.856005224054622,11.856005224154622,11.907216437500564,11.907216437600564,11.958393018282989,11.958393018382989,12.009457402876496,12.009457402976496,12.060476810477011,12.060476810577011,12.111253978889495,12.111253978989495,12.161988178548777,12.161988178648777,12.212486921565727,12.212486921665727,12.262972364730496,12.262972364830496,12.313431056625634,12.313431056725634,12.363775506200748,12.363775506300748,12.41397005772677,12.41397005782677,12.464134902460453,12.464134902560453,12.514218318754425,12.514218318854425,12.564199883188952,12.564199883288952,12.61407368471862,12.61407368481862,12.66388485190169,12.66388485200169,12.713599206770443,12.713599206870443,12.763186701510692,12.763186701610692,12.81271860968926,12.81271860978926,12.862165868666189,12.862165868766189,12.911586982634557,12.911586982734557,12.960980057028523,12.960980057128523,13.010315650294968,13.010315650394968,13.059446516776974,13.059446516876974,13.108545024074468,13.108545024174468,13.157489844191193,13.157489844291193,13.206228346088174,13.206228346188174,13.254910352031871,13.254910352131871,13.303539689045273,13.303539689145273,13.352040385038284,13.352040385138285,13.400508191368349,13.400508191468349,13.448967320586878,13.448967320686878,13.497422243869245,13.497422243969245,13.545839427400123,13.545839427500123,13.59418590573393,13.59418590583393,13.642436973732144,13.642436973832144,13.690624497992154,13.690624498092154,13.738752911798022,13.738752911898022,13.786716953071796,13.786716953171796,13.834556597152657,13.834556597252657,13.882376840879337,13.882376840979337,13.930141119131166,13.930141119231166,13.977876486308308,13.977876486408308,14.025492874750833,14.025492874850833,14.073065385048553,14.073065385148553,14.120507890238477,14.120507890338477,14.167890527147923,14.167890527247923,14.215199617011546,14.215199617111546,14.262424588151966,14.262424588251966,14.309493030262859,14.309493030362859,14.356522785339344,14.356522785439344,14.403319849839622,14.403319849939622,14.449918401731406,14.449918401831406,14.49651066366461,14.49651066376461,14.54304768005798,14.54304768015798,14.589532520108175,14.589532520208175,14.635986251387314,14.635986251487314,14.682303877056658,14.682303877156658,14.728571827209732,14.728571827309732,14.774764790446946,14.774764790546946,14.82094710622415,14.82094710632415,14.866999189545657,14.866999189645657,14.912763450425075,14.912763450525075,14.958295248076187,14.958295248176187,15.00369848048954,15.00369848058954,15.049011114764525,15.049011114864525,15.094270625413417,15.094270625513417,15.139183544187928,15.139183544287928,15.184074940694519,15.184074940794519,15.228866720929357,15.228866721029357,15.273585295140219,15.273585295240219,15.318053256182045,15.318053256282045,15.362492230366554,15.362492230466554,15.406810444924554,15.406810445024554,15.45104518491174,15.45104518501174,15.495273028679275,15.495273028779275,15.539361546075094,15.539361546175094,15.583438430836667,15.583438430936667,15.62750409976848,15.62750409986848,15.671548246432375,15.671548246532375,15.715551432582854,15.715551432682854,15.759537264510259,15.759537264610259,15.80350434023587,15.80350434033587,15.847431137491766,15.847431137591766,15.891333835870206,15.891333835970206,15.935211147066417,15.935211147166417,15.979011652563562,15.979011652663562,16.02272875927252,16.02272875937252,16.066323969615475,16.066323969715473,16.10971744658801,16.10971744668801,16.153085498487002,16.153085498587,16.19630202301028,16.196302023110277,16.23951464472793,16.239514644827928,16.28272468983604,16.28272468993604,16.325673587989062,16.32567358808906,16.368499604601837,16.368499604701835,16.411293565160634,16.41129356526063,16.45394558484686,16.453945584946858,16.496580060853425,16.496580060953423,16.53919843305037,16.539198433150368,16.581782854627498,16.581782854727496,16.624313053730347,16.624313053830345,16.666679334996903,16.6666793350969,16.708906668804907,16.708906668904906,16.75099710128547,16.75099710138547,16.79300224041201,16.79300224051201,16.83499946025335,16.834999460353348,16.876979136415027,16.876979136515025,16.918937138743523,16.91893713884352,16.960854256341236,16.960854256441234,17.002650727986392,17.00265072808639,17.044312723348376,17.044312723448375,17.08595149133319,17.08595149143319,17.127584386163903,17.1275843862639,17.169182079961352,17.16918208006135,17.210757342099278,17.210757342199276,17.252170202053584,17.252170202153582,17.293563661653707,17.293563661753705,17.33490244408375,17.334902444183747,17.37621708974502,17.376217089845017,17.41750854592043,17.41750854602043,17.458705235912635,17.458705236012634,17.499857327825012,17.49985732792501,17.540912266401225,17.540912266501223,17.58194769094931,17.581947691049308,17.622977734930412,17.62297773503041,17.66399838186496,17.66399838196496,17.704830898411807,17.704830898511805,17.745643180995504,17.745643181095502,17.786444171966806,17.786444172066805,17.827219775755882,17.82721977585588,17.867982079692776,17.867982079792775,17.90873028805984,17.908730288159838,17.94947451783864,17.94947451793864,17.989897126120805,17.989897126220804,18.030302380179894,18.030302380279892,18.070603129987365,18.070603130087363,18.110897779292838,18.110897779392836,18.15110327032998,18.151103270429978,18.19124699852081,18.19124699862081,18.231386558666795,18.231386558766793,18.271471972121134,18.271471972221132,18.311495168033357,18.311495168133355,18.351512907595968,18.351512907695966,18.39151681682796,18.391516816927957,18.43147112632632,18.431471126426317,18.47138587728998,18.471385877389977,18.51115690648918,18.51115690658918,18.550906716551,18.550906716651,18.59045771087379,18.590457710973787,18.62955870791877,18.629558708018767,18.66858324028653,18.66858324038653,18.707552110309976,18.707552110409974,18.746473767752743,18.74647376785274,18.7852513624092,18.785251362509197,18.82402789610878,18.82402789620878,18.862763886099437,18.862763886199435,18.90148327969335,18.901483279793347,18.940158605685873,18.94015860578587,18.978656221402808,18.978656221502806,19.01710347955977,19.017103479659767,19.055537134734017,19.055537134834015,19.093965409341283,19.09396540944128,19.132373904681202,19.1323739047812,19.1707188183916,19.170718818491597,19.20903792811529,19.209037928215288,19.246722888761724,19.246722888861722,19.284339645037996,19.284339645137994,19.35956622347957,19.359566223579566,19.397101702881397,19.397101702981395,19.4345374144462,19.4345374145462,19.47194618528479,19.471946185384787,19.509301226236218,19.509301226336216,19.54661238904284,19.546612389142837,19.58383825849544,19.58383825859544,19.6209559482387,19.620955948338697,19.658062990521945,19.658062990621943,19.695127367182526,19.695127367282524,19.732084246177465,19.732084246277463,19.76895136064301,19.768951360743007,19.80569260015428,19.80569260025428,19.84236139146791,19.84236139156791,19.87901172971028,19.87901172981028,19.915652784580047,19.915652784680045,19.952271483572925,19.952271483672924,19.988862142991405,19.988862143091403,20.0252988499899,20.025298850089897,20.061708578370858,20.061708578470856,20.09794760636983,20.097947606469827,20.134174130234264,20.134174130334262,20.170319907702694,20.170319907802693,20.20642237539607,20.206422375496068,20.242506238452915,20.242506238552913,20.27846509805578,20.278465098155777,20.31427311232663,20.31427311242663,20.350005950225036,20.350005950325034,20.38569638773999,20.38569638783999,20.421292854789385,20.421292854889384,20.456881326770944,20.456881326870942,20.492383330767662,20.49238333086766,20.527772759662295,20.527772759762293,20.563153094640903,20.5631530947409,20.59851126319921,20.59851126329921,20.633854123665547,20.633854123765545,20.669183115909952,20.66918311600995,20.704509683110082,20.70450968321008,20.73982541339362,20.73982541349362,20.77511219471116,20.775112194811157,20.845582655073333,20.84558265517333,20.88074143952286,20.88074143962286,20.915887075685475,20.915887075785474,20.95102183704018,20.95102183714018,20.986129960799204,20.986129960899202,21.02110989823364,21.021109898333638,21.056006740010424,21.056006740110423,21.090863189643542,21.09086318974354,21.125708044533738,21.125708044633736,21.160499472667304,21.160499472767302,21.19527218249039,21.195272182590386,21.230034206962152,21.23003420706215,21.264789941475332,21.26478994157533,21.29945394111064,21.29945394121064,21.334015861538607,21.334015861638605,21.36855686595972,21.36855686605972,21.403031522685193,21.40303152278519,21.437473479204304,21.437473479304302,21.506281306478478,21.506281306578476,21.54056605192436,21.540566052024356,21.574819120229435,21.574819120329433,21.609047824417836,21.609047824517834,21.643179640509285,21.643179640609283,21.677181034688456,21.677181034788454,21.711172198212104,21.711172198312102,21.74514710636086,21.74514710646086,21.779111329158294,21.779111329258292,21.81303576607313,21.81303576617313,21.84686422428262,21.846864224382617,21.880620158511835,21.880620158611833,21.914350743450143,21.91435074355014,21.94804586211596,21.94804586221596,21.981704870356914,21.981704870456912,22.015301016903365,22.015301017003363,22.048874807572926,22.048874807672924,22.082443672371312,22.08244367247131,22.115985406986898,22.115985407086896,22.14951444801137,22.14951444811137,22.182990403301066,22.182990403401064,22.21643794010319,22.21643794020319,22.24988172566496,22.249881725764958,22.283304178415392,22.28330417851539,22.316721970533866,22.316721970633864,22.349966030965007,22.349966031065005,22.383192812955702,22.3831928130557,22.416395268721043,22.41639526882104,22.44950216258552,22.44950216268552,22.482581661027982,22.48258166112798,22.51563209683049,22.51563209693049,22.548659987299526,22.548659987399525,22.581668818436235,22.581668818536233,22.614670753353295,22.614670753453293,22.647670301117397,22.647670301217396,22.680622863648722,22.68062286374872,22.71356962880858,22.71356962890858,22.74641420108715,22.746414201187147,22.779255438929837,22.779255439029836,22.81204700125626,22.81204700135626,22.84483587329919,22.844835873399187,22.877601783204167,22.877601783304165,22.910254928550472,22.91025492865047,22.94290583830909,22.94290583840909,23.008202618281196,23.008202618381194,23.040839849274466,23.040839849374464,23.07335639642386,23.07335639652386,23.105852595936156,23.105852596036154,23.138342240250655,23.138342240350653,23.170810021275383,23.17081002137538,23.20325105103048,23.20325105113048,23.23567184682243,23.23567184692243,23.268050772709362,23.26805077280936,23.300408024763108,23.300408024863106,23.33275822903194,23.332758229131937,23.365080545291637,23.365080545391635,23.397345077293277,23.397345077393275,23.429582327546854,23.429582327646852,23.461727767139926,23.461727767239925,23.493868546101037,23.493868546201035,23.525948623172702,23.5259486232727,23.558005700215098,23.558005700315096,23.590042050707225,23.590042050807224,23.622028043639382,23.62202804373938,23.653984860257633,23.65398486035763,23.717871135963442,23.71787113606344,23.749792562091844,23.749792562191843,23.781655332461906,23.781655332561904,23.813497338390384,23.813497338490382,23.845329265228607,23.845329265328605,23.87714186749528,23.877141867595277,23.908917525728118,23.908917525828116,23.94068875067689,23.94068875077689,23.972392680647108,23.972392680747106,24.00404754136212,24.004047541462118,24.035563340944634,24.035563341044632,24.06707334315568,24.06707334325568,24.098538406265057,24.098538406365055,24.129981909215196,24.129981909315195,24.161423669164765,24.161423669264764,24.19279514072175,24.192795140821747,24.22412879674461,24.224128796844607,24.255386518568688,24.255386518668686,24.286642497392194,24.286642497492192,24.31784414006747,24.31784414016747,24.349044570220613,24.34904457032061,24.380154705365918,24.380154705465916,24.411246993701035,24.411246993801033,24.442316660920046,24.442316661020044,24.47336734458936,24.473367344689358,24.504302990221,24.504302990320998,24.5351187098351,24.535118709935098,24.56590942118014,24.56590942128014,24.596699071568313,24.59669907166831,24.627458105772547,24.627458105872545,24.658187584749708,24.658187584849706,24.688914221878115,24.688914221978113,24.71963570578744,24.719635705887438,24.750226692001853,24.75022669210185,24.780757468913936,24.780757469013935,24.81121572184575,24.811215721945747,24.84166764692766,24.84166764702766,24.872092593392043,24.87209259349204,24.90245611803196,24.90245611813196,24.932818392258426,24.932818392358424,24.963154218345792,24.96315421844579,24.99347909384262,24.993479093942618,25.023783318571812,25.02378331867181,25.05408045762477,25.05408045772477,25.084359029932518,25.084359030032516,25.114490053452812,25.11449005355281,25.144618235124348,25.144618235224346,25.174686207493558,25.174686207593556,25.204710604848493,25.20471060494849,25.234733486550763,25.23473348665076,25.2647436367706,25.264743636870598,25.294721011001442,25.29472101110144,25.324659773980507,25.324659774080505,25.35456272966524,25.354562729765238,25.384451589780134,25.384451589880133,25.414329575087123,25.41432957518712,25.474041743338926,25.474041743438924,25.50387361491342,25.50387361501342,25.53369464957132,25.533694649671318,25.563510758358042,25.56351075845804,25.593325654622628,25.593325654722626,25.62311758874926,25.623117588849258,25.652886901759786,25.652886901859784,25.682620142236757,25.682620142336756,25.712303555632197,25.712303555732195,25.741969084326122,25.74196908442612,25.77159456189824,25.77159456199824,25.80114626507663,25.801146265176627,25.83066822357136,25.83066822367136,25.860177564257615,25.860177564357613,25.889645641299918,25.889645641399916,25.91904960623427,25.91904960633427,25.948356758854306,25.948356758954304,25.97763727387866,25.977637273978658,26.00689683500485,26.006896835104847,26.03614351308334,26.036143513183337,26.065365789153837,26.065365789253836,26.094565974586665,26.094565974686663,26.123736263770574,26.123736263870573,26.152899884082732,26.15289988418273,26.18183490608089,26.18183490618089,26.210753407464946,26.210753407564944,26.239654516734607,26.239654516834605,26.268555019743204,26.268555019843202,26.297422595197535,26.297422595297533,26.32628183456218,26.32628183466218,26.355131942119485,26.355131942219483,26.38398136763309,26.38398136773309,26.412805557529737,26.412805557629735,26.441603602417818,26.441603602517816,26.470396910891303,26.4703969109913,26.49915024402561,26.49915024412561,26.527895089504966,26.527895089604964,26.556629514872213,26.55662951497221,26.585268946708347,26.585268946808345,26.64254178566125,26.64254178576125,26.671156550250178,26.671156550350176,26.69968033778757,26.699680337887568,26.728193970452068,26.728193970552066,26.756699835396635,26.756699835496633,26.78518137411584,26.78518137421584,26.81365537246301,26.81365537256301,26.87059268380603,26.87059268390603,26.89898585997456,26.898985860074557,26.92736251552898,26.927362515628978,26.955737465974146,26.955737466074144,26.98410730109155,26.98410730119155,27.01245618231079,27.012456182410787,27.040791346873362,27.04079134697336,27.06910847516916,27.069108475269157,27.09741760839712,27.097417608497118,27.125669412062823,27.12566941216282,27.153907840093716,27.153907840193714,27.182090075301854,27.182090075401852,27.238283428531656,27.238283428631654,27.26620194017516,27.26620194027516,27.294038303443937,27.294038303543935,27.32187178697264,27.321871787072638,27.34965017652678,27.349650176626778,27.405153453095792,27.40515345319579,27.432874247848456,27.432874247948455,27.460568177657542,27.46056817775754,27.48825191470242,27.48825191480242,27.51584190862964,27.515841908729637,27.543418981617844,27.543418981717842,27.570980594948814,27.570980595048812,27.59852678651386,27.59852678661386,27.625994618835854,27.625994618935852,27.653444718021603,27.6534447181216,27.680888262009557,27.680888262109555,27.708330441910103,27.7083304420101,27.73575371404359,27.735753714143588,27.763157851062115,27.763157851162113,27.790533152788583,27.79053315288858,27.817899739512203,27.8178997396122,27.845247456360074,27.845247456460072,27.872568043025147,27.872568043125145,27.899882604970855,27.899882605070854,27.92714392961652,27.927143929716518,27.954391348148935,27.954391348248933,27.98157325259468,27.981573252694677,28.008743145493007,28.008743145593005,28.035911030151546,28.035911030251544,28.06307201859044,28.063072018690438,28.090223306852238,28.090223306952236,28.117372207961083,28.11737220806108,28.144517433612204,28.144517433712203,28.171633672406003,28.171633672506,28.198696749682394,28.198696749782393,28.22575800817558,28.225758008275577,28.252727531790896,28.252727531890894,28.27966094498134,28.27966094508134,28.306556277398442,28.30655627749844,28.333449412119176,28.333449412219174,28.360336370555277,28.360336370655276,28.387216811684894,28.387216811784892,28.41406656084794,28.414066560947937,28.44091214196614,28.44091214206614,28.467755146474804,28.467755146574802,28.5214297880971,28.5214297881971,28.548251838707596,28.548251838807595,28.575069380251396,28.575069380351394,28.601884345185656,28.601884345285654,28.628689344703613,28.62868934480361,28.655477141563757,28.655477141663756,28.68224712950503,28.682247129605027,28.708993435373326,28.708993435473324,28.73573769511052,28.735737695210517,28.762460622036375,28.762460622136373,28.789150583516655,28.789150583616653,28.815807920573203,28.8158079206732,28.842399819325713,28.84239981942571,28.86895856317606,28.86895856327606,28.895515942939006,28.895515943039005,28.922041910800356,28.922041910900354,28.948532980758976,28.948532980858975,28.975010258278296,28.975010258378294,29.001424484646527,29.001424484746526,29.027814536354672,29.02781453645467,29.054203186084095,29.054203186184093,29.080534165229412,29.08053416532941,29.106847411238487,29.106847411338485,29.13315789118144,29.13315789128144,29.159434458395893,29.15943445849589,29.18568575210207,29.18568575220207,29.211936401655862,29.21193640175586,29.23817310720509,29.238173107305087,29.26440609940527,29.26440609950527,29.29063272586424,29.290632725964237,29.316857647213954,29.316857647313952,29.343057711859874,29.343057711959872,29.369254290504653,29.36925429060465,29.39543665990565,29.395436660005647,29.421614823370483,29.42161482347048,29.447782831962428,29.447782832062426,29.47392302832787,29.47392302842787,29.500057957800287,29.500057957900285,29.52617958742052,29.526179587520517,29.55227969477283,29.55227969487283,29.57837673292849,29.578376733028488,29.604472785909913,29.60447278600991,29.63056292784592,29.630562927945917,29.656633214731947,29.656633214831945,29.68269804526836,29.682698045368358,29.70868569119254,29.708685691292537,29.73465552819784,29.73465552829784,29.760581941754136,29.760581941854134,29.786492289392132,29.78649228949213,29.812401235051404,29.812401235151402,29.83829157607415,29.838291576174147,29.86415285445694,29.864152854556938,29.890009244859872,29.89000924495987,29.91585244908457,29.91585244918457,29.94163392835427,29.94163392845427,29.967375849089276,29.967375849189274,29.99309874838327,29.99309874848327,30.018768296703268,30.018768296803266,30.044430797238352,30.04443079733835,30.070090986403113,30.07009098650311,30.095698771876798,30.095698771976796,30.1212876116921,30.1212876117921,30.14685004125963,30.146850041359627,30.172385189079087,30.172385189179085,30.197916850897407,30.197916850997405,30.22340334626615,30.223403346366148,30.248878246891966,30.248878246991964,30.274325903661033,30.27432590376103,30.299732258894835,30.299732258994833,30.32507162228061,30.32507162238061,30.350389804420317,30.350389804520315,30.375691844859084,30.375691844959082,30.400991157123045,30.400991157223043,30.426261520421004,30.426261520521003,30.45152347184664,30.45152347194664,30.476777807117614,30.476777807217612,30.527279960353074,30.527279960453072,30.552517926575202,30.5525179266752,30.577751194274057,30.577751194374056,30.602942781524476,30.602942781624474,30.62810697335288,30.62810697345288,30.653248619847815,30.653248619947814,30.67837995990459,30.678379960004587,30.703510580026347,30.703510580126345,30.72864055599572,30.72864055609572,30.753751927328565,30.753751927428564,30.778843140480895,30.778843140580893,30.803923592499263,30.80392359259926,30.854082260948317,30.854082261048315,30.879109703014542,30.87910970311454,30.9041293015782,30.9041293016782,30.92912828726554,30.92912828736554,30.954102492031726,30.954102492131724,30.979054416703654,30.979054416803653,31.003988797695925,31.003988797795923,31.028915221511674,31.028915221611673,31.05381542453623,31.053815424636227,31.07860581852482,31.07860581862482,31.10335290273836,31.10335290283836,31.128064293331512,31.12806429343151,31.152772387380107,31.152772387480105,31.17747930679788,31.17747930689788,31.20215780772809,31.202157807828087,31.226829298764695,31.226829298864693,31.27615227422266,31.27615227432266,31.300797847598606,31.300797847698604,31.32536661527548,31.32536661537548,31.3499257585579,31.3499257586579,31.374453452047415,31.374453452147414,31.398960494769295,31.398960494869293,31.423377507722556,31.423377507822554,31.447788836978308,31.447788837078306,31.47219542981946,31.47219542991946,31.496598271420257,31.496598271520256,31.5209921706703,31.5209921707703,31.545383682767383,31.54538368286738,31.569723965804215,31.569723965904213,31.59406314999286,31.594063150092857,31.618378348977995,31.618378349077993,31.64268888733117,31.642688887431166,31.666995788117934,31.666995788217932,31.69128969218305,31.69128969228305,31.715567795569083,31.71556779566908,31.739840669953402,31.7398406700534,31.76411074038028,31.764110740480277,31.78837876467605,31.78837876477605,31.81263117774932,31.81263117784932,31.836856612205057,31.836856612305056,31.86105908452284,31.861059084622838,31.885252917620402,31.8852529177204,31.909420416252818,31.909420416352816,31.933574842380953,31.93357484248095,31.9577211597673,31.957721159867297,31.981792528129102,31.9817925282291,32.00585847803261,32.00585847813261,32.029918024303576,32.02991802440358,32.05396624107084,32.05396624117084,32.078006045965786,32.07800604606579,32.1020405839677,32.1020405840677,32.12605215983165,32.12605215993165,32.15005066319133,32.15005066329133,32.17404636259357,32.17404636269357,32.19803694666804,32.19803694676804,32.221989222621275,32.22198922272128,32.2459355875291,32.2459355876291,32.26986554549676,32.26986554559676,32.2937857275047,32.2937857276047,32.31770412862075,32.31770412872075,32.341567511544866,32.34156751164487,32.36542358144485,32.36542358154485,32.38926942068931,32.38926942078931,32.41310756799647,32.41310756809647,32.43691320455385,32.436913204653855,32.46071058080418,32.460710580904184,32.48445327988443,32.48445327998443,32.50819200037642,32.50819200047642,32.531904841099056,32.53190484119906,32.55559559118403,32.55559559128403,32.5792854318774,32.579285431977404,32.602971748678314,32.602971748778316,32.626650676672455,32.62665067677246,32.65028318780357,32.650283187903575,32.67391183402038,32.67391183412038,32.69753233360408,32.69753233370408,32.721128317505844,32.721128317605846,32.74471960288433,32.74471960298433,32.76830925893619,32.768309259036194,32.79185935645336,32.79185935655336,32.81540096631558,32.81540096641558,32.83892973102141,32.83892973112141,32.86238737372569,32.86238737382569,32.88583982531957,32.88583982541957,32.90928629008541,32.90928629018541,32.9327032375155,32.9327032376155,32.95611696418365,32.95611696428365,32.97952883417729,32.97952883427729,33.002923046817315,33.00292304691732,33.026299867342956,33.02629986744296,33.04962750493944,33.04962750503944,33.07294983775158,33.07294983785158,33.09627156430265,33.09627156440265,33.11958927437414,33.119589274474144,33.142896033855095,33.1428960339551,33.16618809150515,33.16618809160515,33.189463969562944,33.189463969662945,33.2127227965282,33.2127227966282,33.235964761857495,33.2359647619575,33.2592015360764,33.2592015361764,33.28242963318375,33.282429633283755,33.305644013634456,33.30564401373446,33.328828573618864,33.328828573718866,33.35194246629752,33.351942466397524,33.375047795538585,33.37504779563859,33.398147895777946,33.39814789587795,33.42123636338306,33.42123636348306,33.44431884416012,33.444318844260124,33.46738177301774,33.467381773117744,33.490437502525175,33.49043750262518,33.513483304507616,33.51348330460762,33.53646688894794,33.536466889047944,33.55943308127388,33.55943308137388,33.5823933625544,33.5823933626544,33.6053377673732,33.6053377674732,33.62823855928641,33.62823855938641,33.65112502828189,33.65112502838189,33.6740099816247,33.6740099817247,33.696891752096896,33.6968917521969,33.719751015126946,33.71975101522695,33.74260311669812,33.74260311679812,33.765445593874844,33.765445593974846,33.788255939214956,33.78825593931496,33.81104286772131,33.811042867821314,33.833809979069,33.833809979169004,33.856571823523666,33.85657182362367,33.879264895238414,33.879264895338416,33.901950881276925,33.901950881376926,33.924627621834155,33.92462762193416,33.94730360456504,33.947303604665045,33.96997011446674,33.96997011456674,33.99263476769392,33.99263476779392,34.01521890187299,34.01521890197299,34.03778742482956,34.03778742492956,34.060331015299695,34.0603310153997,34.08285702419886,34.08285702429886,34.10532945477613,34.105329454876134,34.127790101153884,34.127790101253886,34.15024275246381,34.15024275256381,34.172691690424685,34.17269169052469,34.19512535818491,34.19512535828491,34.217530683240206,34.21753068334021,34.239900617805645,34.23990061790565,34.26226827889208,34.26226827899208,34.28460323977215,34.284603239872155,34.306928348909864,34.306928349009866,34.32915683518985,34.32915683528985,34.351378652598086,34.35137865269809,34.37359527889592,34.37359527899592,34.39580739612707,34.39580739622707,34.418004091592294,34.418004091692296,34.44019453499025,34.44019453509025,34.462379446255966,34.46237944635597,34.48455317958323,34.48455317968323,34.506721153430355,34.50672115353036,34.52888564127633,34.528885641376334,34.55104383916373,34.551043839263734,34.57318744889418,34.57318744899418,34.595312757118634,34.595312757218636,34.617411920334526,34.61741192043453,34.63950653659241,34.63950653669241,34.661598803588646,34.66159880368865,34.683687016213995,34.683687016313996,34.705747568178104,34.705747568278106,34.72779391089844,34.72779391099844,34.74983748755265,34.74983748765265,34.77184654521729,34.77184654531729,34.7938453722264,34.7938453723264,34.81584177419125,34.81584177429125,34.837833439741495,34.837833439841496,34.85981085815665,34.85981085825665,34.881777780677055,34.88177778077706,34.903703060640346,34.90370306074035,34.947542177389266,34.94754217748927,34.96944987578158,34.96944987588158,34.99134976856264,34.991349768662644,35.01323556577387,35.01323556587387,35.0351028341312,35.0351028342312,35.056949375938274,35.056949376038276,35.07878375463266,35.07878375473266,35.10060483347487,35.10060483357487,35.122405261549446,35.12240526164945,35.144184470486636,35.14418447058664,35.16594984909321,35.16594984919321,35.18770624745771,35.18770624755771,35.20942505763599,35.20942505773599,35.23110832575915,35.231108325859154,35.25277564163795,35.252775641737955,35.274423632945215,35.27442363304522,35.296059120117945,35.29605912021795,35.317685210244115,35.317685210344116,35.33930766280388,35.339307662903884,35.36088957165472,35.36088957175472,35.38246318230718,35.38246318240718,35.40402603182569,35.40402603192569,35.44715025310134,35.44715025320134,35.46870923770554,35.46870923780554,35.49022483675204,35.49022483685204,35.511738086536916,35.51173808663692,35.53324379594975,35.53324379604975,35.554734083596664,35.554734083696665,35.57621213234827,35.57621213244827,35.59768775605559,35.597687756155594,35.61913382453585,35.61913382463585,35.64057890784188,35.64057890794188,35.66200121846653,35.66200121856653,35.68337241370487,35.683372413804875,35.704727656698864,35.704727656798866,35.726073957342095,35.7260739574421,35.747411012504045,35.74741101260405,35.76873840538022,35.76873840548022,35.79003639459459,35.79003639469459,35.81131085330125,35.81131085340125,35.83257788530983,35.83257788540983,35.85383794531613,35.85383794541613,35.8750828109044,35.875082811004404,35.896321007620934,35.896321007720935,35.91755344485731,35.91755344495731,35.93876068436805,35.93876068446805,35.95996576407373,35.95996576417373,35.981155990383236,35.98115599048324,36.00234170762606,36.00234170772606,36.0235231810414,36.0235231811414,36.04469154406114,36.04469154416114,36.08702190435941,36.08702190445941,36.10818223442,36.10818223452,36.129340745697384,36.129340745797386,36.15049201973327,36.15049201983327,36.171639542528794,36.171639542628796,36.192774447515845,36.19277444761585,36.21388601145176,36.213886011551764,36.234991740124904,36.234991740224906,36.25609007999128,36.25609008009128,36.27718197833381,36.27718197843381,36.2982439425361,36.2982439426361,36.31930279965041,36.31930279975041,36.34035373747952,36.34035373757952,36.361388041020575,36.36138804112058,36.382418176516786,36.38241817661679,36.40344630377321,36.40344630387321,36.42446120696008,36.424461207060084,36.445451973378184,36.445451973478185,36.46641435920004,36.46641435930004,36.48733146489836,36.48733146499836,36.50821969741331,36.50821969751331,36.529067083088805,36.52906708318881,36.549887565929396,36.5498875660294,36.570690050394525,36.57069005049453,36.59148825314087,36.59148825324087,36.61227592210115,36.61227592220115,36.63302945098503,36.63302945108503,36.6537777508672,36.6537777509672,36.67450918911341,36.67450918921341,36.69523873279378,36.695238732893785,36.736689218825624,36.736689218925626,36.757400764130544,36.757400764230546,36.77810442804157,36.778104428141575,36.798807334126266,36.79880733422627,36.819501752556,36.819501752656,36.84018628135207,36.84018628145207,36.8608686882344,36.8608686883344,36.881547419659,36.881547419759,36.90222493856147,36.90222493866147,36.92288862713332,36.92288862723332,36.94354083463619,36.94354083473619,36.96418190209193,36.96418190219193,36.984821037090526,36.98482103719053,37.026097677761086,37.02609767786109,37.046728249322086,37.04672824942209,37.06734449796536,37.06734449806536,37.087947522539075,37.08794752263908,37.108526031430856,37.10852603153086,37.12906066217783,37.12906066227783,37.14957744611461,37.149577446214614,37.17009131242,37.17009131252,37.190601995854784,37.190601995954786,37.21105527394468,37.21105527404468,37.23150783209955,37.231507832199554,37.25193371476743,37.251933714867434,37.27235217073723,37.27235217083723,37.292731295520234,37.292731295620236,37.31306426867942,37.31306426877942,37.33334832414868,37.33334832424868,37.3536194207876,37.353619420887604,37.37388782714304,37.37388782724304,37.39412743609774,37.39412743619774,37.41435366941762,37.414353669517624,37.43457554523608,37.43457554533608,37.45479567805397,37.45479567815397,37.47500944513064,37.47500944523064,37.49520225830914,37.495202258409144,37.515389046768284,37.515389046868286,37.53556658974613,37.535566589846134,37.55573049184995,37.55573049194995,37.575871393924494,37.575871394024496,37.59598944753504,37.59598944763504,37.61609196570571,37.61609196580571,37.63617648550092,37.63617648560092,37.65625945175214,37.656259451852144,37.6763354838924,37.6763354839924,37.69637839902181,37.69637839912181,37.71641597147556,37.716415971575564,37.73644615512254,37.736446155222545,37.75646898785407,37.75646898795407,37.776485265387805,37.77648526548781,37.79649998937755,37.79649998947755,37.816509446474264,37.816509446574265,37.83651321987347,37.83651321997347,37.85648380047918,37.856483800579184,37.876452789649605,37.87645278974961,37.89642018738472,37.89642018748472,37.91637402002843,37.91637402012843,37.93631686419029,37.93631686429029,37.956237428257886,37.95623742835789,37.976153824280644,37.976153824380646,37.99605972440866,37.99605972450866,38.0159104926708,38.0159104927708,38.03575781282311,38.03575781292311,38.05560157119164,38.05560157129164,38.07543797864472,38.07543797874472,38.095250704024835,38.09525070412484,38.115060246534334,38.115060246634336,38.13485459462582,38.13485459472582,38.15463204319003,38.15463204329003,38.174405816296506,38.17440581639651,38.19417436040128,38.19417436050128,38.21392259476027,38.21392259486027,38.233657074571276,38.23365707467128,38.25337787561693,38.25337787571693,38.27308564204962,38.273085642149624,38.29278647437135,38.29278647447135,38.3124794631905,38.3124794632905,38.33217150472674,38.33217150482674,38.35185642269543,38.35185642279543,38.37152902598616,38.371529026086165,38.410867563695895,38.4108675637959,38.43053308131041,38.43053308141041,38.450183442398206,38.45018344249821,38.46983122687647,38.469831226976474,38.48946169502298,38.48946169512298,38.50908356184058,38.509083561940585,38.528701828983095,38.5287018290831,38.548315170254426,38.54831517035443,38.56792172898005,38.567921729080055,38.58752525640035,38.58752525650035,38.60712499468897,38.60712499478897,38.62671249408227,38.62671249418227,38.64629120269008,38.64629120279008,38.665845660855176,38.66584566095518,38.685393677496414,38.685393677596416,38.70494104998527,38.704941050085274,38.72448395129875,38.72448395139875,38.74402617056853,38.74402617066853,38.76356357764108,38.76356357774108,38.78308685125248,38.78308685135248,38.80259261907553,38.802592619175535,38.82208561752485,38.82208561762485,38.84157346275508,38.84157346285508,38.900032337813805,38.90003233791381,38.91949221925227,38.91949221935227,38.938950509255434,38.938950509355436,38.95836317811322,38.95836317821322,38.977749285157955,38.97774928525796,38.99710644323669,38.99710644333669,39.0164560988347,39.016456098934704,39.03580272312738,39.035802723227384,39.05513915465585,39.05513915475585,39.093791366945155,39.09379136704516,39.11310570783595,39.11310570793595,39.13241087902809,39.132410879128095,39.151703621868336,39.15170362196834,39.17097749483284,39.17097749493284,39.19024564620851,39.19024564630851,39.228737161423446,39.22873716152345,39.2479546142173,39.2479546143173,39.26715380339647,39.26715380349647,39.28633147030772,39.286331470407724,39.305499020237406,39.30549902033741,39.32465459651099,39.32465459661099,39.343808391892686,39.34380839199269,39.362960709513025,39.36296070961303,39.382111359915434,39.382111360015436,39.40124927883541,39.40124927893541,39.42036037070312,39.420360370803124,39.4394664230257,39.4394664231257,39.458549210079795,39.4585492101798,39.47763093617702,39.47763093627702,39.49670864579467,39.49670864589467,39.51577847401843,39.515778474118434,39.55391722107435,39.55391722117435,39.59205225478123,39.59205225488123,39.611114883654814,39.611114883754816,39.63013317968778,39.630133179787784,39.64913768328145,39.649137683381454,39.668124908434685,39.668124908534686,39.68710588152065,39.68710588162065,39.706084467453664,39.706084467553666,39.72503368761618,39.725033687716184,39.743981088995504,39.743981089095506,39.76292447389525,39.76292447399525,39.78186024264032,39.78186024274032,39.800790858166316,39.80079085826632,39.81971741932142,39.81971741942142,39.838637614735305,39.83863761483531,39.85751624337471,39.85751624347471,39.87635698069735,39.876356980797354,39.89518430449387,39.89518430459387,39.91400461839678,39.91400461849678,39.93281826342795,39.93281826352795,39.95162864980587,39.95162864990587,39.970425622657665,39.970425622757666,39.98922107985678,39.989221079956785,40.008001683659735,40.00800168375974,40.02676303867378,40.02676303877378,40.04551298840147,40.04551298850147,40.0642548672787,40.0642548673787,40.08299360117663,40.082993601276634,40.101719641483456,40.10171964158346,40.12043677733084,40.12043677743084,40.13915194282975,40.13915194292975,40.15786096993535,40.15786097003535,40.176557227667196,40.1765572277672,40.195215442517025,40.19521544261703,40.21386782210407,40.21386782220407,40.23251861025581,40.23251861035581,40.251165003014805,40.25116500311481,40.26980332492334,40.26980332502334,40.2884351674167,40.2884351675167,40.30706181879967,40.30706181889967,40.32568377165935,40.32568377175935,40.3443036026053,40.3443036027053,40.36291195248228,40.36291195258228,40.38151579329256,40.38151579339256,40.40011247264397,40.40011247274397,40.41870786369061,40.41870786379061,40.43730189064985,40.43730189074985,40.45588970343314,40.45588970353314,40.47446319329869,40.47446319339869,40.49303175729307,40.49303175739307,40.511588764435835,40.51158876453584,40.53011322293751,40.53011322303751,40.548633437611706,40.54863343771171,40.56713716956311,40.56713716966311,40.585636998708885,40.58563699880889,40.60413557744121,40.604135577541214,40.6226236602788,40.6226236603788,40.641095487741495,40.6410954878415,40.65956405655095,40.65956405665095,40.67801743094238,40.678017431042385,40.69646572789737,40.69646572799737,40.7148962538248,40.7148962539248,40.733325643012726,40.73332564311273,40.75174968952499,40.751749689624994,40.77016691560024,40.770166915700244,40.78857872321719,40.788578723317194,40.806971206262595,40.8069712063626,40.82533538449439,40.82533538459439,40.84368721015691,40.84368721025691,40.86202039329159,40.86202039339159,40.88034474774946,40.88034474784946,40.898660879791606,40.89866087989161,40.935288028548115,40.93528802864812,40.953586313780065,40.953586313880066,40.971880089945316,40.97188009004532,40.990153783712685,40.99015378381269,41.00842080860831,41.00842080870831,41.02668521900307,41.02668521910307,41.04491844484415,41.04491844494415,41.06315076129362,41.06315076139362,41.08134242036021,41.08134242046021,41.09952574333711,41.09952574343711,41.11770660337843,41.11770660347843,41.135860636367475,41.13586063646748,41.153996595198436,41.15399659529844,41.172119102611944,41.172119102711946,41.19023240243548,41.19023240253548,41.20833683569089,41.208336835790895,41.22643975329364,41.22643975339364,41.2626419130414,41.2626419131414,41.2807243693331,41.2807243694331,41.29880466581974,41.29880466591974,41.31685957512415,41.31685957522415,41.334910089035816,41.33491008913582,41.371009752771755,41.37100975287176,41.407101080418,41.407101080518004,41.42512071290651,41.425120713006514,41.44311874734447,41.44311874744447,41.46111310632471,41.46111310642471,41.4790956053228,41.4790956054228,41.49707711914665,41.497077119246654,41.515052797707725,41.515052797807726,41.533009720067,41.533009720167,41.5509490608553,41.550949060955304,41.58682490058315,41.586824900683155,41.60473737642787,41.60473737652787,41.62264905655494,41.62264905665494,41.64055956205119,41.64055956215119,41.658450705084576,41.65845070518458,41.6763290408529,41.6763290409529,41.69420089720606,41.69420089730606,41.71206214399052,41.712062144090524,41.729908726835404,41.729908726935406,41.74771609216744,41.74771609226744,41.765517129649574,41.765517129749576,41.783314870587155,41.78331487068716,41.80110101678181,41.80110101688181,41.81888216132265,41.81888216142265,41.83665856944889,41.836658569548895,41.85442379963669,41.85442379973669,41.87218588484521,41.87218588494521,41.88992584152473,41.88992584162473,41.907664926703966,41.90766492680397,41.92538688500803,41.92538688510803,41.943098271634724,41.943098271734726,41.960806437499485,41.96080643759949,41.97843522105565,41.97843522115565,41.99605456967393,41.99605456977393,42.01367175848716,42.013671758587165,42.03128826525669,42.03128826535669,42.04890121024244,42.04890121034244,42.06650589492114,42.066505895021145,42.08410819244689,42.08410819254689,42.10170336640508,42.101703366505085,42.11929115155651,42.11929115165651,42.13687711792473,42.136877118024735,42.15445925726996,42.154459257369965,42.17203484141739,42.172034841517394,42.189585720426294,42.189585720526296,42.207115304515185,42.207115304615186,42.2246286711205,42.2246286712205,42.242137907572285,42.24213790767229,42.25963225273659,42.25963225283659,42.27711303280949,42.27711303290949,42.29458778816303,42.29458778826303,42.32952998584597,42.329529985945975,42.34699598830534,42.34699598840534,42.364458656328836,42.36445865642884,42.399380998961796,42.3993809990618,42.4168353308956,42.4168353309956,42.45171982010312,42.451719820203124,42.46914543041883,42.46914543051883,42.48657009345161,42.48657009355161,42.50399202830959,42.50399202840959,42.521401496924355,42.52140149702436,42.53880982879962,42.53880982889962,42.55619224301422,42.556192243114225,42.590947219701064,42.590947219801066,42.608319024346976,42.60831902444698,42.64305831402868,42.64305831412868,42.660415757865536,42.66041575796554,42.67776119015498,42.677761190254984,42.695098172680794,42.695098172780796,42.71243417003237,42.71243417013237,42.729767931796246,42.72976793189625,42.74708774955557,42.74708774965557,42.76439343385373,42.764393433953735,42.781692828193314,42.781692828293316,42.79898581890037,42.79898581900037,42.816267707451615,42.81626770755162,42.83354266189189,42.83354266199189,42.85080363443628,42.850803634536284,42.868059984240034,42.868059984340036,42.88531220389025,42.88531220399025,42.90255479914602,42.90255479924602,42.91979356737879,42.91979356747879,42.937013882540306,42.93701388264031,42.95423097693989,42.95423097703989,42.98866319539059,42.988663195490595,43.00587464398399,43.00587464408399,43.023048542282474,43.023048542382476,43.040219371384296,43.0402193714843,43.05738247065751,43.05738247075751,43.074534429883585,43.074534429983586,43.09168483556568,43.09168483566568,43.10883300566009,43.10883300576009,43.12597405218694,43.12597405228694,43.14310111681792,43.14310111691792,43.16021772344546,43.160217723545465,43.17733118509372,43.17733118519372,43.194432332064025,43.19443233216403,43.211529462554765,43.21152946265477,43.22862405432727,43.22862405442727,43.24571277294569,43.24571277304569,43.26278516040658,43.26278516050658,43.27983314585948,43.27983314595948,43.29688025981209,43.29688025991209,43.31390399482226,43.31390399492226,43.330921364091225,43.33092136419123,43.34791690796173,43.347916908061734,43.36489994769771,43.36489994779771,43.381877909997236,43.38187791009724,43.398854583992,43.398854584092,43.415830045464624,43.415830045564626,43.432794707911974,43.432794708011976,43.44973591563425,43.44973591573425,43.466674698312254,43.466674698412255,43.483602606182345,43.483602606282346,43.5005127809162,43.500512781016205,43.5343315389486,43.5343315390486,43.55123815189868,43.55123815199868,43.56813294275793,43.56813294285793,43.60191740914866,43.601917409248664,43.61877855251863,43.61877855261863,43.63563226919051,43.63563226929051,43.652459499831984,43.652459499931986,43.66928582108185,43.66928582118185,43.68610805006951,43.68610805016951,43.7029228902504,43.7029228903504,43.719732539320894,43.719732539420896,43.73653957389053,43.736539573990534,43.75333652936991,43.75333652946991,43.77011832832259,43.77011832842259,43.78689899053576,43.78689899063576,43.803677909748366,43.80367790984837,43.820442543934554,43.820442544034556,43.85396233947774,43.85396233957774,43.8874719043654,43.8874719044654,43.90421975269826,43.90421975279826,43.920965024421584,43.920965024521585,43.954446852865374,43.954446852965376,43.97117264845188,43.971172648551885,43.987896246342025,43.98789624644203,44.00461798755764,44.00461798765764,44.02132783089979,44.021327830999795,44.038033203066576,44.03803320316658,44.05473202003555,44.05473202013555,44.088123288232296,44.0881232883323,44.104782205644725,44.104782205744726,44.12143972107844,44.12143972117844,44.13809261377151,44.138092613871514,44.15474387713796,44.15474387723796,44.17138506141415,44.17138506151415,44.187995667397715,44.18799566749772,44.204603052619355,44.20460305271936,44.221191567965256,44.22119156806526,44.23776655611107,44.23776655621107,44.27090558181216,44.27090558191216,44.287459729733754,44.287459729833756,44.304008951784176,44.30400895188418,44.35363918792973,44.35363918802973,44.37016696349487,44.37016696359487,44.38668394003472,44.38668394013472,44.40318708624397,44.40318708634397,44.41968803475684,44.41968803485684,44.43616412656591,44.436164126665915,44.45263961211393,44.45263961221393,44.4691075572899,44.469107557389904,44.48554609880407,44.485546098904074,44.50195258426427,44.50195258436427,44.51834304169748,44.51834304179748,44.534729141629256,44.53472914172926,44.55111198290779,44.55111198300779,44.56749084559807,44.567490845698075,44.58386095539418,44.58386095549418,44.600227200275974,44.600227200375976,44.61659030017848,44.61659030027848,44.632918615852205,44.63291861595221,44.64924378654664,44.64924378664664,44.68189102084753,44.68189102094753,44.698212212953706,44.69821221305371,44.71451999153375,44.71451999163375,44.73082250322076,44.73082250332076,44.74712111210214,44.747121112202144,44.7634067621532,44.763406762253204,44.79597249223175,44.79597249233175,44.8122459412788,44.812245941378805,44.82851685160764,44.82851685170764,44.84478313919583,44.844783139295835,44.861038551976115,44.86103855207612,44.89353577455397,44.89353577465397,44.909773454198,44.909773454298005,44.926003138774206,44.92600313887421,44.942221342281435,44.94222134238144,44.95843818170126,44.95843818180126,44.974647253401145,44.974647253501146,44.99084025918273,44.990840259282734,45.00703246924666,45.00703246934666,45.02321789676489,45.02321789686489,45.039398133172725,45.03939813327273,45.05556810103372,45.05556810113372,45.07172412489015,45.071724124990155,45.08787086537397,45.087870865473974,45.10401453666114,45.10401453676114,45.12014903824964,45.120149038349645,45.136280849554666,45.13628084965467,45.15240167237783,45.15240167247783,45.16852113111358,45.16852113121358,45.184639187870616,45.18463918797062,45.20075341760466,45.20075341770466,45.21686586644682,45.21686586654682,45.232972783156725,45.23297278325673,45.24907879047504,45.24907879057504,45.26518373683648,45.26518373693648,45.28128015765164,45.281280157751645,45.2973636575278,45.2973636576278,45.31344503549021,45.31344503559021,45.32951959301561,45.32951959311561,45.345587443777944,45.345587443877946,45.36165328630049,45.36165328640049,45.37771552914794,45.37771552924794,45.39377220197183,45.39377220207183,45.425858947915245,45.42585894801525,45.441884587750714,45.441884587850716,45.45790060319172,45.45790060329172,45.47390544069429,45.47390544079429,45.489905769130154,45.489905769230155,45.505903596739124,45.505903596839126,45.521899946586736,45.52189994668674,45.53789254519398,45.537892545293985,45.553881885147995,45.553881885247996,45.56986133546833,45.56986133556833,45.585835935700125,45.58583593580013,45.60178931679454,45.60178931689454,45.617740500192575,45.61774050029258,45.6336898648074,45.6336898649074,45.66558738151493,45.665587381614934,45.68153488945524,45.681534889555245,45.69747944187285,45.69747944197285,45.71342047039799,45.713420470497994,45.72936013483574,45.72936013493574,45.745296692185505,45.74529669228551,45.76122544392402,45.76122544402402,45.77713847076608,45.777138470866085,45.793038538777814,45.793038538877816,45.80891359852048,45.808913598620485,45.82478528593596,45.82478528603596,45.84065056971891,45.84065056981891,45.85651339056627,45.85651339066627,45.87237473365227,45.872374733752274,45.88823080984525,45.88823080994525,45.9040775647743,45.9040775648743,45.93574750623338,45.935747506333385,45.951563455521914,45.951563455621915,45.96736311154424,45.96736311164424,45.98315318106342,45.98315318116342,45.998938817298544,45.998938817398546,46.014722975772315,46.01472297587232,46.03050508811498,46.03050508821498,46.0462622300799,46.0462622301799,46.06201224847727,46.06201224857727,46.077760675439336,46.07776067553934,46.09348439726288,46.09348439736288,46.10920103341019,46.10920103351019,46.124915888665605,46.12491588876561,46.14061066152314,46.14061066162314,46.156301986270854,46.156301986370856,46.171984330776496,46.1719843308765,46.18766515962947,46.18766515972947,46.203336591435885,46.20333659153589,46.21900063443554,46.21900063453554,46.234660130477174,46.234660130577176,46.250318830801156,46.25031883090116,46.26594589187565,46.265945891975655,46.28156556414338,46.28156556424338,46.297175157320844,46.297175157420845,46.312771374863495,46.312771374963496,46.32836372749184,46.32836372759184,46.34395460235883,46.34395460245883,46.35953937672382,46.359539376823825,46.37511782323892,46.37511782333892,46.39069460253607,46.390694602636074,46.40626171954745,46.40626171964745,46.4218273966888,46.4218273967888,46.437383979914124,46.437383980014125,46.45292703593936,46.45292703603936,46.46846861420325,46.468468614303255,46.48400261420379,46.48400261430379,46.51506493050735,46.51506493060735,46.53058146261086,46.53058146271086,46.54609253836476,46.54609253846476,46.561594406528684,46.561594406628686,46.57709100779958,46.57709100789958,46.59258192537296,46.59258192547296,46.60806534046563,46.60806534056563,46.6235437917958,46.6235437918958,46.63900655612083,46.639006556220835,46.65446791846714,46.65446791856714,46.6699072280671,46.669907228167105,46.68533634490286,46.68533634500286,46.71618859174631,46.71618859184631,46.73160815997024,46.731608160070245,46.74702371171459,46.74702371181459,46.76243255669588,46.76243255679588,46.77783480858805,46.77783480868805,46.79323334713118,46.79323334723118,46.80861688071287,46.80861688081287,46.82398881955164,46.82398881965164,46.83935867436798,46.83935867446798,46.85471371367947,46.854713713779475,46.87006083370576,46.87006083380576,46.885403672013254,46.885403672113256,46.90074249384117,46.90074249394117,46.9160779433419,46.9160779434419,46.93140865642803,46.931408656528035,46.946738156992026,46.94673815709203,46.96206079922769,46.962060799327695,46.992693352216584,46.992693352316586,47.007982536419554,47.007982536519556,47.02326755257768,47.02326755267768,47.038546013538,47.038546013638005,47.05382220101932,47.053822201119324,47.06909323528156,47.06909323538156,47.0843586237376,47.0843586238376,47.09961268268993,47.09961268278993,47.11486465761985,47.11486465771985,47.130091624280695,47.1300916243807,47.14531643113649,47.145316431236495,47.160537600425876,47.16053760052588,47.17574933477739,47.175749334877395,47.190952808821855,47.19095280892186,47.206152039038834,47.206152039138836,47.22134206166585,47.22134206176585,47.236511812438394,47.236511812538396,47.251676561557126,47.25167656165713,47.266835702760986,47.26683570286099,47.281991964224765,47.28199196432477,47.29714758153616,47.29714758163616,47.31229895502008,47.312298955120085,47.32743975682662,47.32743975692662,47.34257885352391,47.34257885362391,47.3577165861338,47.3577165862338,47.37285356091735,47.37285356101735,47.38797852415349,47.38797852425349,47.4031025022154,47.4031025023154,47.41821757581786,47.418217575917865,47.43332821613627,47.43332821623627,47.4484380228457,47.4484380229457,47.463539076660965,47.463539076760966,47.4786339541916,47.4786339542916,47.493708787215674,47.493708787315676,47.52385712706772,47.52385712716772,47.538918395000394,47.538918395100396,47.55396041414415,47.55396041424415,47.56899273311082,47.56899273321082,47.58402080825001,47.58402080835001,47.614068357199486,47.61406835729949,47.62907684252791,47.62907684262791,47.64407934102829,47.64407934112829,47.65907914924518,47.65907914934518,47.674072743286125,47.67407274338613,47.689064708000444,47.689064708100446,47.70404068257909,47.70404068267909,47.71901127659076,47.719011276690765,47.73398118855873,47.73398118865873,47.74894950909139,47.74894950919139,47.76391722336299,47.76391722346299,47.778884104025614,47.778884104125616,47.80881532663265,47.80881532673265,47.82376686131199,47.82376686141199,47.83870786220527,47.83870786230527,47.853647612685094,47.853647612785096,47.86858577172962,47.86858577182962,47.88352124049066,47.88352124059066,47.8984470090746,47.8984470091746,47.92828509482504,47.928285094925045,47.94319043998926,47.94319044008926,47.95808661545482,47.95808661555482,47.97297278761275,47.97297278771275,47.98785009320256,47.98785009330256,48.00272239713856,48.002722397238564,48.01758534191932,48.01758534201932,48.03244389130734,48.03244389140734,48.047300811368736,48.04730081146874,48.062148144926994,48.062148145026995,48.076977101196626,48.07697710129663,48.09180295037828,48.09180295047828,48.10662485886299,48.106624858962995,48.12144104575887,48.12144104585887,48.1362478356082,48.136247835708204,48.16584205284399,48.165842052943994,48.180628646621486,48.18062864672149,48.195407396896414,48.195407396996416,48.21017436297183,48.21017436307183,48.22493719889371,48.224937198993715,48.239695108944424,48.239695109044426,48.25444521338388,48.25444521348388,48.28394102687005,48.283941026970055,48.29868370461143,48.29868370471143,48.313416682175706,48.31341668227571,48.32814791673942,48.32814791683942,48.34287627156306,48.34287627166306,48.357587385837576,48.35758738593758,48.37229383948013,48.37229383958013,48.386996617664956,48.38699661776496,48.401691666021165,48.40169166612117,48.416382129528046,48.41638212962805,48.43107100159962,48.43107100169962,48.445758130670626,48.44575813077063,48.46044184952313,48.46044184962313,48.475121930809216,48.47512193090922,48.489788901699704,48.489788901799706,48.50445136352351,48.50445136362351,48.51909900984245,48.51909900994245,48.533743321725524,48.533743321825526,48.54838145732396,48.548381457423964,48.57764370873364,48.57764370883364,48.592253274279244,48.592253274379246,48.60685996008477,48.60685996018477,48.62146046960567,48.621460469705674,48.636055105972474,48.636055106072476,48.650647847773435,48.65064784787344,48.66521812002356,48.66521812012356,48.694339188387,48.694339188487,48.708880511671126,48.70888051177113,48.72342047086784,48.723420470967845,48.73795948278165,48.73795948288165,48.75249379617217,48.75249379627217,48.79609283353812,48.79609283363812,48.81061956866529,48.81061956876529,48.825144485009254,48.825144485109256,48.83966860563557,48.839668605735575,48.85418124519291,48.854181245292914,48.86869236909757,48.868692369197575,48.88320262150196,48.88320262160196,48.897707303882775,48.89770730398278,48.91220762876217,48.91220762886217,48.92670079218269,48.926700792282695,48.94119028014549,48.94119028024549,48.95566457369027,48.95566457379027,48.97013144053697,48.97013144063697,48.98459698118758,48.98459698128758,48.99906111985947,48.99906111995947,49.013517566594054,49.013517566694055,49.02797083045803,49.02797083055803,49.0424120827746,49.0424120828746,49.056850644807675,49.056850644907676,49.071288411123106,49.07128841122311,49.08572462389454,49.085724623994544,49.11459606426318,49.114596064363184,49.12902083385696,49.129020833956965,49.15786844058736,49.15786844068736,49.18670695340174,49.18670695350174,49.215533795690554,49.215533795790556,49.22993624729877,49.22993624739877,49.244332977318145,49.244332977418146,49.27312181461625,49.273121814716255,49.287507859284304,49.287507859384306,49.30188704562402,49.30188704572402,49.31624899141461,49.31624899151461,49.330599418244915,49.33059941834492,49.3449430246382,49.3449430247382,49.35926939048235,49.359269390582355,49.37359507428281,49.37359507438281,49.38791329349387,49.38791329359387,49.4022296560304,49.4022296561304,49.41654313882686,49.41654313892686,49.430853249296135,49.43085324939614,49.44515847178554,49.44515847188554,49.459463050122565,49.45946305022257,49.47376330884948,49.47376330894948,49.50235689219233,49.50235689229233,49.516631346932535,49.516631347032536,49.54517540632439,49.54517540642439,49.55943387092892,49.55943387102892,49.602203277914455,49.60220327801446,49.61645446738617,49.61645446748617,49.63070402753126,49.63070402763126,49.64495078371891,49.64495078381891,49.65919181831773,49.65919181841773,49.6734297837199,49.6734297838199,49.68766297481615,49.68766297491615,49.70189222521546,49.70189222531546,49.716120414657894,49.716120414757896,49.73034496653393,49.73034496663393,49.74456250851636,49.74456250861636,49.75876815262533,49.75876815272533,49.772972053733724,49.772972053833726,49.787174628646035,49.78717462874604,49.801367579163895,49.801367579263896,49.81555643741954,49.81555643751954,49.82973885415134,49.82973885425134,49.84391176016263,49.84391176026263,49.858083226303876,49.85808322640388,49.872250524400286,49.87225052450029,49.88639084387917,49.88639084397917,49.91466655696575,49.91466655706575,49.92879547115828,49.92879547125828,49.94292010363203,49.94292010373203,49.95704295521388,49.95704295531388,49.971160160989534,49.971160161089536,49.98525845899813,49.985258459098134,49.999353687810064,49.999353687910066,50.01344793144777,50.01344793154777,50.0275399773891,50.0275399774891,50.04162292941441,50.04162292951441,50.05570372163466,50.05570372173466,50.069783339224095,50.0697833393241,50.08384757293892,50.08384757303892,50.11194966801212,50.111949668112125,50.140019593357394,50.140019593457396,50.15404487479859,50.154044874898595,50.16806932263083,50.16806932273083,50.18209142120143,50.18209142130143,50.21012773694873,50.21012773704873,50.22413926384195,50.22413926394195,50.23813260290313,50.23813260300313,50.252122759093695,50.252122759193696,50.26610935350049,50.26610935360049,50.280086361404145,50.28008636150415,50.29406143685065,50.29406143695065,50.30803514820974,50.30803514830974,50.322006169285345,50.32200616938535,50.33597518212117,50.33597518222117,50.34993491158438,50.34993491168438,50.36387573328053,50.36387573338053,50.39175574734621,50.39175574744621,50.405693878758875,50.40569387885888,50.4196253791911,50.419625379291105,50.433552256882685,50.433552256982686,50.44747530755128,50.44747530765128,50.46139475854478,50.46139475864478,50.47530981414553,50.475309814245534,50.489219451287994,50.489219451387996,50.50312389732005,50.50312389742005,50.51702546361204,50.517025463712045,50.53092589316453,50.53092589326453,50.544821662085056,50.54482166218506,50.55871489228736,50.558714892387364,50.572593382767444,50.572593382867446,50.58646997868169,50.58646997878169,50.600337632245186,50.60033763234519,50.61420020837223,50.614200208472234,50.628060359455006,50.62806035955501,50.641918729645894,50.641918729745896,50.65577357594432,50.655773576044325,50.669622852219184,50.669622852319186,50.68346242831696,50.68346242841696,50.697298063717795,50.6972980638178,50.71113169087884,50.71113169097884,50.72496459810486,50.72496459820486,50.73879379198185,50.73879379208185,50.75261999244481,50.75261999254481,50.76644270690662,50.766442707006625,50.78026390571577,50.78026390581577,50.794077677826834,50.794077677926836,50.80788978271996,50.80788978281996,50.82170029617778,50.82170029627778,50.849316435113565,50.84931643521357,50.863113345588665,50.86311334568867,50.87690586067103,50.87690586077103,50.89068568216228,50.89068568226228,50.90446209343502,50.904462093535024,50.91823289679288,50.91823289689288,50.93199782699664,50.931997827096644,50.94575809656844,50.94575809666844,50.95949441882805,50.95949441892805,50.97322714141257,50.973227141512574,50.98695486234328,50.986954862443284,51.00067883203363,51.00067883213363,51.01439893680966,51.014398936909664,51.02811623762826,51.02811623772826,51.04183099972864,51.04183099982864,51.0692562422106,51.0692562423106,51.08294194167102,51.08294194177102,51.09662472350005,51.09662472360005,51.11030591389378,51.110305913993784,51.12398407298217,51.12398407308217,51.13765647259041,51.13765647269041,51.15132413578406,51.15132413588406,51.17865802230131,51.178658022401315,51.19232041860193,51.19232041870193,51.20597872264033,51.20597872274033,51.233275816689016,51.23327581678902,51.24690699054462,51.246906990644625,51.260530699810836,51.26053069991084,51.27415039259746,51.274150392697464,51.28775345109604,51.28775345119604,51.30135560020301,51.30135560030301,51.314954566439376,51.31495456653938,51.32854307467232,51.32854307477232,51.34210407380929,51.34210407390929,51.355662647901994,51.355662648001996,51.369217470754336,51.36921747085434,51.382753726861466,51.38275372696147,51.39627509168111,51.39627509178111,51.40979372832595,51.40979372842595,51.43682914494111,51.43682914504111,51.463861984946725,51.46386198504673,51.47736478302116,51.47736478312116,51.4908644361163,51.4908644362163,51.50436018640582,51.50436018650582,51.51785294328131,51.517852943381314,51.53134391926492,51.53134391936492,51.54482940100759,51.54482940110759,51.55829987778883,51.558299877888835,51.5717686115695,51.571768611669505,51.5852243107372,51.585224310837205,51.612134913354964,51.612134913454966,51.62558462569462,51.62558462579462,51.63902770705385,51.63902770715385,51.65245998938779,51.652459989487795,51.66589086974302,51.665890869843025,51.679318226205794,51.679318226305796,51.706169983608625,51.70616998370863,51.719563541016846,51.71956354111685,51.732952589358376,51.73295258945838,51.74633970524275,51.74633970534275,51.75972394138705,51.75972394148705,51.77309965198508,51.77309965208508,51.786464753014414,51.786464753114416,51.799828565738984,51.799828565838986,51.81318760415764,51.81318760425764,51.82654179248775,51.826541792587754,51.83989135807722,51.83989135817722,51.853239749035865,51.85323974913587,51.866587533733444,51.866587533833446,51.879934636387325,51.87993463648733,51.89327817725743,51.89327817735743,51.90661808056112,51.90661808066112,51.91994654068715,51.91994654078715,51.933272689442866,51.93327268954287,51.94659614791509,51.94659614801509,51.95990361625164,51.95990361635164,51.973207219673874,51.973207219773876,51.9865098758132,51.9865098759132,51.99980745451607,51.999807454616075,52.01309703815111,52.01309703825111,52.02636362175688,52.02636362185688,52.03962721194862,52.03962721204862,52.05288936227033,52.05288936237033,52.06614473004634,52.06614473014634,52.079392860580846,52.07939286068085,52.09264030907165,52.09264030917165,52.10588658293164,52.10588658303164,52.119124179680085,52.11912417978009,52.14559471254502,52.14559471264502,52.158827231857025,52.15882723195703,52.17205815973372,52.172058159833725,52.19851804513865,52.19851804523865,52.21174438816602,52.21174438826602,52.22494977729522,52.224949777395224,52.23814690611738,52.23814690621738,52.2513403594818,52.2513403595818,52.26453047841036,52.26453047851036,52.27771904379492,52.27771904389492,52.30408734588724,52.30408734598724,52.3172661732034,52.3172661733034,52.33043666442987,52.33043666452987,52.343602532915696,52.3436025330157,52.356767454118604,52.356767454218605,52.369929192450904,52.369929192550906,52.38308907410868,52.383089074208684,52.39624672017877,52.396246720278775,52.40939997085612,52.40939997095612,52.42255235003318,52.42255235013318,52.43570249362256,52.43570249372256,52.448843202274055,52.44884320237406,52.46197352870476,52.46197352880476,52.475100785938814,52.475100786038816,52.488223799345384,52.488223799445386,52.501341053271815,52.50134105337182,52.51445269928336,52.51445269938336,52.54067481667563,52.540674816775635,52.55377873285856,52.55377873295856,52.56688173964989,52.56688173974989,52.579959359258986,52.57995935935899,52.59303409912801,52.593034099228014,52.60610338264742,52.60610338274742,52.61916868757857,52.619168687678574,52.63223243896574,52.63223243906574,52.645289407807205,52.64528940790721,52.658344292626246,52.65834429272625,52.67139849540159,52.67139849550159,52.68444671134888,52.684446711448885,52.71053954356838,52.71053954366838,52.723582530513966,52.72358253061397,52.736619795870716,52.73661979597072,52.74965099861679,52.74965099871679,52.76266749953196,52.76266749963196,52.77568218166392,52.77568218176392,52.80171025762308,52.80171025772308,52.81472221158023,52.814722211680234,52.827724086447134,52.827724086547136,52.84071520018008,52.84071520028008,52.86668727277307,52.86668727287307,52.8796679663939,52.879667966493905,52.89264532557886,52.89264532567886,52.90561370452174,52.90561370462174,52.918579658420356,52.91857965852036,52.931536404729,52.931536404829004,52.94449019551494,52.94449019561494,52.95744319058322,52.95744319068322,52.98334270130462,52.983342701404624,52.99628724660926,52.996287246709265,53.009230693065724,53.009230693165726,53.03510959091081,53.03510959101081,53.04804072268932,53.048040722789324,53.06096776220563,53.06096776230563,53.073892869264775,53.07389286936478,53.08681695325837,53.08681695335837,53.099731981227265,53.09973198132727,53.11264602402192,53.11264602412192,53.12555370107536,53.12555370117536,53.138458119475565,53.13845811957557,53.151355376416895,53.1513553765169,53.16425168607531,53.16425168617531,53.17714685899422,53.17714685909422,53.19004002367334,53.19004002377334,53.20292867928577,53.20292867938577,53.215810362895915,53.21581036299592,53.22868856050492,53.22868856060492,53.24156565926574,53.24156565936574,53.254433853567114,53.254433853667116,53.2672957200186,53.267295720118604,53.28015231957705,53.28015231967705,53.29300228815507,53.293002288255074,53.305851574689385,53.305851574789386,53.318699686592886,53.31869968669289,53.331542986299155,53.33154298639916,53.34436567312911,53.34436567322911,53.35718654117586,53.35718654127586,53.37000593146126,53.37000593156126,53.382823161941594,53.382823162041596,53.395619021719284,53.395619021819286,53.40841385843142,53.408413858531425,53.42120680057772,53.42120680067772,53.4339985680932,53.4339985681932,53.44678825158626,53.446788251686264,53.47236189698355,53.47236189708355,53.48513604503673,53.485136045136734,53.49790568402323,53.49790568412323,53.51066869202929,53.51066869212929,53.523430146491364,53.523430146591366,53.536184174255354,53.536184174355355,53.54892850184225,53.54892850194225,53.561661916729925,53.56166191682993,53.574393815964925,53.57439381606493,53.587106921106816,53.58710692120682,53.61253047899843,53.61253047909843,53.625234869137465,53.62523486923747,53.637930430599695,53.6379304306997,53.65062106619075,53.65062106629075,53.66328544309929,53.66328544319929,53.68861298439424,53.68861298449424,53.701271450257366,53.70127145035737,53.72658413815614,53.72658413825614,53.73923324486403,53.73923324496403,53.764529412148704,53.764529412248706,53.77717594224705,53.77717594234705,53.7898179632787,53.7898179633787,53.80245248182964,53.80245248192964,53.82771894232197,53.827718942421974,53.84034152510813,53.84034152520813,53.85295899256653,53.85295899266653,53.865574641241714,53.865574641341716,53.87818221906644,53.87818221916644,53.90339563171531,53.903395631815314,53.928602034470586,53.92860203457059,53.94120112464035,53.94120112474035,53.96638861962856,53.96638861972856,53.978975546685646,53.97897554678565,53.99154943912978,53.99154943922978,54.016694761082434,54.016694761182436,54.02926459915567,54.02926459925567,54.04182250146412,54.04182250156412,54.05437778927171,54.05437778937171,54.06693216768771,54.06693216778771,54.07948510623366,54.079485106333664,54.092035240822185,54.092035240922186,54.104577494016816,54.10457749411682,54.117117397949805,54.11711739804981,54.12965082246763,54.12965082256763,54.142181367245385,54.14218136734539,54.15470948697887,54.15470948707887,54.16722532992572,54.16722533002572,54.1797376110888,54.1797376111888,54.19224489059806,54.19224489069806,54.20475095758519,54.20475095768519,54.21725513000648,54.21725513010648,54.229757824666414,54.229757824766416,54.25474847426407,54.25474847436407,54.26723586083204,54.26723586093204,54.29220790579316,54.292207905893164,54.31716327857492,54.31716327867492,54.329633500376396,54.3296335004764,54.3421017518294,54.3421017519294,54.35456606258546,54.354566062685464,54.36702582638351,54.36702582648351,54.37948157370199,54.37948157380199,54.391936335846225,54.39193633594623,54.4043785180733,54.4043785181733,54.41681406931994,54.41681406941994,54.429243671629855,54.42924367172986,54.441668082829366,54.44166808292937,54.46650436320255,54.466504363302555,54.478913049505614,54.478913049605616,54.49131506693693,54.49131506703693,54.503713863606315,54.50371386370632,54.52851047177086,54.52851047187086,54.540906009787,54.540906009887,54.55329264334371,54.55329264344371,54.56566457506952,54.56566457516952,54.57802248700813,54.57802248710813,54.590378390706945,54.59037839080695,54.61508826564742,54.615088265747424,54.62744155484539,54.62744155494539,54.63979090334641,54.63979090344641,54.6521216472109,54.6521216473109,54.66444140259353,54.664441402693534,54.67675206406014,54.67675206416014,54.689058140677425,54.689058140777426,54.701354479226296,54.7013544793263,54.713648847426704,54.713648847526706,54.72594132106127,54.72594132116127,54.73823030869469,54.73823030879469,54.75051668182726,54.75051668192726,54.76280134985058,54.76280134995058,54.77508495691702,54.77508495701702,54.7873611751767,54.7873611752767,54.7996349683921,54.7996349684921,54.81189690162536,54.81189690172536,54.82414678541989,54.82414678551989,54.836396062953355,54.83639606305336,54.86088366742974,54.86088366752974,54.87311740952333,54.87311740962333,54.885350090660054,54.885350090760056,54.89758068777435,54.89758068787435,54.90980363084267,54.90980363094267,54.922025967649915,54.92202596774992,54.93424432586891,54.93424432596891,54.9464516198234,54.9464516199234,54.95865698132074,54.958656981420745,54.97085408251103,54.970854082611034,54.99524574617338,54.99524574627338,55.00743170731654,55.00743170741654,55.019614030893294,55.019614030993296,55.03179548296976,55.031795483069764,55.04397557095883,55.04397557105883,55.05614046452987,55.05614046462987,55.08046748560584,55.08046748570584,55.09262874161047,55.09262874171047,55.104787951484,55.104787951584,55.116942803856105,55.11694280395611,55.12907708124321,55.12907708134321,55.14120666010703,55.14120666020703,55.15333260140445,55.15333260150445,55.16545615554891,55.16545615564891,55.177569138016,55.177569138116,55.18968083217831,55.18968083227831,55.20178248514169,55.201782485241694,55.213881637278156,55.21388163737816,55.23806345882831,55.23806345892831,55.250134609281695,55.250134609381696,55.26219723418881,55.26219723428881,55.2742496284404,55.2742496285404,55.286299332408504,55.286299332508506,55.29834339057041,55.29834339067041,55.31038218183928,55.31038218193928,55.32241892697705,55.32241892707705,55.33445275448343,55.33445275458343,55.346482148705746,55.34648214880575,55.35850930734038,55.35850930744038,55.370528925602976,55.37052892570298,55.38254536099496,55.38254536109496,55.39455573377627,55.394555733876274,55.406563643621986,55.40656364372199,55.41856530140044,55.41856530150044,55.43056347317775,55.43056347327775,55.44256012930239,55.442560129402395,55.45454170468296,55.45454170478296,55.47850239250852,55.47850239260852,55.49047748847392,55.49047748857392,55.51442408072964,55.51442408082964,55.52639273517119,55.526392735271195,55.55032511818313,55.55032511828313,55.562287672122686,55.56228767222269,55.57424041221121,55.57424041231121,55.586190916712034,55.586190916812036,55.59813854147279,55.598138541572794,55.610067220575175,55.61006722067518,55.621976575106004,55.621976575206006,55.63388104165697,55.63388104175697,55.64578183275022,55.64578183285022,55.65767527292801,55.65767527302801,55.66956473451754,55.669564734617545,55.693340815847854,55.693340815947856,55.705223381217735,55.70522338131774,55.71709749682398,55.717097496923984,55.7289674822767,55.7289674823767,55.740834095402235,55.74083409550224,55.75269722252662,55.75269722262662,55.764556333171434,55.764556333271436,55.77639790013658,55.77639790023658,55.78822404533582,55.78822404543582,55.80004897801291,55.80004897811291,55.811870273123596,55.8118702732236,55.82368652868916,55.82368652878916,55.835500738123606,55.83550073822361,55.84731407605778,55.84731407615778,55.85912286703393,55.859122867133934,55.87093090018375,55.87093090028375,55.89454476878703,55.89454476888703,55.906338782149646,55.90633878224965,55.918131128294334,55.918131128394336,55.92992180722108,55.92992180732108,55.94171187988675,55.941711879986755,55.953500550573715,55.953500550673716,55.965285773150846,55.96528577325085,55.97706432685624,55.97706432695624,55.98884140280026,55.98884140290026,56.000612795046784,56.000612795146786,56.024149554820454,56.024149554920456,56.035902569778344,56.035902569878346,56.04764838538605,56.04764838548605,56.059392912688985,56.05939291278899,56.071136416926365,56.071136417026366,56.08287306283542,56.08287306293542,56.094606109069375,56.094606109169376,56.11805753759771,56.11805753769771,56.12977065299905,56.12977065309905,56.15318869927732,56.153188699377324,56.16488987891389,56.16488987901389,56.17658241933023,56.17658241943023,56.19996598451025,56.19996598461025,56.21165575886047,56.211655758960475,56.223342160883504,56.223342160983506,56.246709015992835,56.24670901609284,56.25838579362141,56.25838579372141,56.27006094192336,56.27006094202336,56.281730671767015,56.28173067186702,56.293398961740635,56.29339896184064,56.30506638021397,56.305066380313974,56.31673292718702,56.31673292728702,56.328393866245186,56.32839386634519,56.34005397169439,56.34005397179439,56.351713395099885,56.35171339519989,56.37502652032205,56.375026520422054,56.386678479138155,56.38667847923816,56.39832748243154,56.39832748253154,56.409967353917594,56.409967354017596,56.42160616444678,56.42160616454678,56.433241867887986,56.43324186798799,56.45650793209474,56.45650793219474,56.49139920384796,56.491399203947964,56.50302626806895,56.50302626816895,56.51465242289834,56.51465242299834,56.5262692564638,56.5262692565638,56.54950099113757,56.54950099123757,56.572729012462304,56.572729012562306,56.58433951817786,56.584339518277865,56.595941687803744,56.595941687903746,56.60753726434051,56.60753726444051,56.619123443830716,56.61912344393072,56.630702764992584,56.630702765092586,56.6422720070642,56.6422720071642,56.65383643693859,56.65383643703859,56.67696192436017,56.67696192446017,56.68852252721157,56.68852252731157,56.700077522148085,56.70007752224809,56.71163179714958,56.71163179724958,56.73473132901918,56.734731329119185,56.76936349994842,56.76936350004842,56.780896896834385,56.78089689693439,56.792429687459276,56.79242968755928,56.8154928436648,56.8154928437648,56.82699903458533,56.82699903468533,56.838502042635255,56.83850204273526,56.86150044258043,56.86150044268043,56.87299174221348,56.87299174231348,56.884480844150154,56.884480844250156,56.895964717085114,56.895964717185116,56.90744279264861,56.90744279274861,56.91891507084064,56.91891507094064,56.930383900922855,56.93038390102286,56.941839052239715,56.941839052339716,56.9532915511644,56.9532915512644,56.96472980295398,56.964729803053984,56.976164114046625,56.97616411414663,56.987592400419906,56.98759240051991,56.99900511346199,56.99900511356199,57.0104167276559,57.0104167277559,57.04464906941071,57.04464906951071,57.0560577280819,57.056057728181905,57.067463393339075,57.06746339343908,57.09027070737385,57.09027070747385,57.10167015845508,57.10167015855508,57.11305164905216,57.11305164915216,57.124431434539986,57.12443143463999,57.13581012117963,57.13581012127963,57.1471870648187,57.147187064918704,57.158558438434206,57.15855843853421,57.16992371154772,57.16992371164772,57.181281898985,57.181281899085,57.203996796098195,57.2039967961982,57.215349072490056,57.21534907259006,57.22669975744662,57.22669975754662,57.23804737320652,57.23804737330652,57.24939298072663,57.249392980826634,57.2607362389851,57.260736239085105,57.27206714467432,57.27206714477432,57.29472554583423,57.294725545934234,57.306051487760946,57.30605148786095,57.31737390579521,57.31737390589521,57.32869291361096,57.32869291371096,57.34000983740428,57.34000983750428,57.351318576673194,57.351318576773195,57.37392355107648,57.37392355117648,57.38521914205847,57.38521914215847,57.39651287636594,57.39651287646594,57.407803427802804,57.407803427902806,57.41909049323853,57.41909049333853,57.43037452736891,57.43037452746891,57.44165128636648,57.44165128646648,57.45292743910297,57.45292743920297,57.47546273137474,57.47546273147474,57.4867136863856,57.4867136864856,57.49796259526534,57.497962595365344,57.50920703296971,57.509207033069714,57.52044995502141,57.520449955121414,57.531688178549835,57.53168817864984,57.55415666843017,57.55415666853017,57.56538344878093,57.56538344888093,57.57660806932663,57.576608069426634,57.59905621156986,57.59905621166986,57.610277118766525,57.61027711886653,57.621495941940765,57.62149594204077,57.632712226396784,57.632712226496785,57.6551408925032,57.6551408926032,57.666349636587185,57.66634963668719,57.688762350449245,57.68876235054925,57.699961924834575,57.69996192493458,57.71115763430559,57.711157634405595,57.722349099949135,57.722349100049136,57.733535867069406,57.73353586716941,57.74471554851343,57.744715548613435,57.755891743956326,57.75589174405633,57.76706384713701,57.76706384723701,57.778235040926084,57.778235041026086,57.78939986897395,57.789399869073954,57.80056242354281,57.80056242364281,57.81172319721978,57.81172319731978,57.82287715045973,57.822877150559734,57.83403030798203,57.834030308082035,57.84518172250376,57.84518172260376,57.856329120545915,57.85632912064592,57.86746295349668,57.86746295359668,57.87858882927091,57.87858882937091,57.88971023386977,57.88971023396977,57.90083061540307,57.900830615503075,57.911936370888114,57.911936370988116,57.92304136854682,57.92304136864682,57.93414575994445,57.93414576004445,57.94524685479753,57.94524685489753,57.95634423630156,57.95634423640156,57.96744029160951,57.967440291709515,57.9785316862855,57.9785316863855,57.98961898869928,57.98961898879928,58.00070515437355,58.00070515447355,58.0117862805027,58.0117862806027,58.02286259443461,58.022862594534615,58.03393693801806,58.03393693811806,58.056083616945166,58.05608361704517,58.06715443663615,58.06715443673615,58.0782210882823,58.0782210883823,58.089283761340184,58.089283761440186,58.11140702343353,58.11140702353353,58.133527557352075,58.13352755745208,58.1445853424301,58.1445853425301,58.15564032355068,58.15564032365068,58.17773914574472,58.17773914584472,58.18878219110052,58.188782191200524,58.1998224703902,58.199822470490204,58.21085710387369,58.210857103973694,58.221885864203074,58.221885864303076,58.2329085998131,58.2329085999131,58.2439283420091,58.2439283421091,58.25494058172438,58.25494058182438,58.26595149524357,58.26595149534357,58.28796741123655,58.28796741133655,58.298968738252604,58.298968738352606,58.309959228352064,58.309959228452065,58.32094888484256,58.32094888494256,58.34292615169244,58.34292615179244,58.35391326946471,58.353913269564714,58.36489375625655,58.36489375635655,58.37587166643885,58.37587166653885,58.38684764416399,58.386847644263995,58.39782123473618,58.39782123483618,58.40879368856886,58.408793688668865,58.41975136478801,58.419751364888015,58.43070695698474,58.430706957084745,58.441661942920405,58.44166194302041,58.45261079046275,58.45261079056275,58.46355846337428,58.46355846347428,58.47450367335021,58.474503673450215,58.48544698876031,58.48544698886031,58.50732547294741,58.50732547304741,58.52918160125761,58.529181601357614,58.54009134496106,58.540091345061064,58.572812808351486,58.57281280845149,58.583713079225745,58.58371307932575,58.59460887892463,58.59460887902463,58.60549903281732,58.60549903291732,58.616386913231004,58.616386913331006,58.6272671395987,58.6272671396987,58.63814619133558,58.63814619143558,58.64902062033181,58.64902062043181,58.670765310279435,58.67076531037944,58.6816334872084,58.6816334873084,58.69249779922306,58.692497799323064,58.70336021667188,58.703360216771884,58.71421971648932,58.71421971658932,58.72507826902383,58.725078269123834,58.735935533253574,58.735935533353576,58.76850467355062,58.768504673650625,58.77934238586092,58.77934238596092,58.79017403556054,58.79017403566054,58.80100379069432,58.80100379079432,58.82265595004642,58.82265595014642,58.83347755854709,58.83347755864709,58.84428579141295,58.84428579151295,58.85508792377681,58.85508792387681,58.865887365857176,58.86588736595718,58.87667957069605,58.87667957079605,58.887470032534345,58.88747003263435,58.89824242021455,58.89824242031455,58.909012799654974,58.909012799754976,58.91978189079062,58.91978189089062,58.93054859477331,58.930548594873315,58.94131378310333,58.941313783203334,58.95207006697392,58.95207006707392,58.962811686904914,58.962811687004915,58.97354819150815,58.97354819160815,58.98428037650128,58.98428037660128,59.00574194253009,59.005741942630095,59.0379225073738,59.0379225074738,59.07007836707898,59.07007836717898,59.08078577115094,59.080785771250945,59.09149241739658,59.09149241749658,59.10219807846797,59.10219807856797,59.11289911679872,59.112899116898724,59.12359780586784,59.12359780596784,59.13429490350165,59.13429490360165,59.144991053852536,59.14499105395254,59.155686067463925,59.15568606756393,59.166376685682565,59.166376685782566,59.17706658396619,59.17706658406619,59.18775443611871,59.18775443621871,59.19843668035635,59.19843668045635,59.20911657533235,59.209116575432354,59.21979423472066,59.21979423482066,59.230469317499434,59.230469317599436,59.24114296040817,59.24114296050817,59.25181129853256,59.25181129863256,59.26247508969894,59.26247508979894,59.27313433390731,59.27313433400731,59.283785204134674,59.283785204234675,59.29443353564381,59.29443353574381,59.30508126089189,59.30508126099189,59.315727546269926,59.31572754636993,59.32636704910226,59.32636704920226,59.337005680434316,59.33700568053432,59.34764040896074,59.347640409060745,59.35827210618182,59.358272106281824,59.3689001658365,59.3689001659365,59.37952727820826,59.37952727830826,59.39015272336208,59.39015272346208,59.41139811942879,59.41139811952879,59.42201697149349,59.42201697159349,59.432630139860684,59.432630139960686,59.45385405155079,59.453854051650794,59.46445983111122,59.46445983121122,59.47506466338872,59.47506466348872,59.496272168138674,59.496272168238676,59.51747379973453,59.517473799834534,59.52805847383152,59.52805847393152,59.53863901777498,59.538639017874985,59.54921213502036,59.549212135120364,59.559782978786735,59.55978297888674,59.570352079552535,59.57035207965254,59.58091799744773,59.58091799754773,59.591482627038154,59.591482627138156,59.612608589674444,59.612608589774446,59.623168520741594,59.623168520841595,59.63372254076332,59.633722540863324,59.6442748556758,59.644274855775805,59.65482641276194,59.654826412861944,59.66537736358702,59.66537736368702,59.686476309714465,59.686476309814466,59.69702195575519,59.69702195585519,59.70756294116396,59.70756294126396,59.718101160506606,59.71810116060661,59.728636007522056,59.72863600762206,59.73916600444897,59.73916600454897,59.770747393900784,59.770747394000786,59.78126818323772,59.78126818333772,59.79178791161779,59.79178791171779,59.80230206997429,59.80230207007429,59.81281372750389,59.81281372760389,59.823323869380815,59.82332386948082,59.83383325343141,59.83383325353141,59.84434009876378,59.84434009886378,59.854843420203686,59.85484342030369,59.86533969385868,59.86533969395868,59.87583070062065,59.87583070072065,59.886320722208374,59.886320722308376,59.896809493382655,59.89680949348266,59.907296445773724,59.907296445873726,59.928268077076865,59.92826807717687,59.93874354839895,59.938743548498955,59.94920787967392,59.94920787977392,59.97013411717958,59.970134117279585,59.98059397727918,59.98059397737918,59.991047661094136,59.99104766119414,60.00148967438353,60.00148967448353,60.02236968448275,60.02236968458275,60.032807643401256,60.03280764350126,60.043242798362314,60.043242798462316,60.06410829608722,60.06410829618722,60.07453416767567,60.074534167775674,60.084957803676446,60.08495780377645,60.09537795367607,60.09537795377607,60.10579556495748,60.105795565057484,60.116210751194615,60.116210751294616,60.12662184516954,60.12662184526954,60.13703021096965,60.13703021106965,60.147427436722644,60.147427436822646,60.16820858837644,60.16820858847644,60.17859024079824,60.178590240898245,60.199352712032876,60.19935271213288,60.220111697266375,60.22011169736638,60.23048444522874,60.23048444532874,60.24085294936363,60.24085294946363,60.251220619889544,60.251220619989546,60.26158389502272,60.26158389512272,60.27194269898052,60.27194269908052,60.28229968415511,60.282299684255115,60.292646855478665,60.29264685557867,60.30299175332321,60.302991753423214,60.31333259679687,60.31333259689687,60.323672568770235,60.323672568870236,60.33400894106851,60.33400894116851,60.34434273675725,60.34434273685725,60.35466645335573,60.354666453455735,60.36498691130097,60.36498691140097,60.37530126874421,60.37530126884421,60.38560289470502,60.385602894805025,60.406196635906134,60.406196636006136,60.41648863747249,60.41648863757249,60.42677491745002,60.42677491755002,60.43705289922917,60.437052899329174,60.44732864542063,60.447328645520635,60.46787900106406,60.467879001164064,60.47814535020897,60.47814535030897,60.48840798600484,60.48840798610484,60.508926664507456,60.50892666460746,60.51918213884446,60.51918213894446,60.529436211202736,60.52943621130274,60.53968395571112,60.53968395581112,60.549928706805474,60.549928706905476,60.560171563333995,60.560171563434,60.57041180536166,60.57041180546166,60.580633101730946,60.58063310183095,60.60106929083698,60.601069290936984,60.61127887878939,60.611278878889394,60.62148471550144,60.62148471560144,60.63168486851597,60.63168486861597,60.64188335431257,60.64188335441257,60.652077027911936,60.65207702801194,60.672460623870315,60.672460623970316,60.68264478674918,60.68264478684918,60.692820461973085,60.69282046207309,60.70298814212916,60.70298814222916,60.713154496089146,60.71315449618915,60.72331471165582,60.72331471175582,60.73347121387344,60.733471213973445,60.74362388906808,60.743623889168084,60.75374579651351,60.753745796613515,60.763866718784705,60.76386671888471,60.7739849128811,60.7739849129811,60.78410166710744,60.784101667207445,60.79420762230852,60.79420762240852,60.80430918211685,60.804309182216855,60.81440577816269,60.81440577826269,60.8244993807945,60.8244993808945,60.834591240425745,60.83459124052575,60.844678969903455,60.84467897000346,60.854762569227645,60.85476256932765,60.86483688517923,60.86483688527923,60.87490654049885,60.87490654059885,60.884969261707504,60.884969261807505,60.89502906528477,60.89502906538477,60.90508595123064,60.90508595133064,60.91514082893673,60.91514082903673,60.93524815930463,60.93524815940463,60.94529951311825,60.94529951321825,60.95534912393131,60.955349124031315,60.97544304077308,60.97544304087308,60.985477532950746,60.98547753305075,60.99550205971211,60.99550205981211,61.00552222897205,61.00552222907205,61.015538988013475,61.01553898811348,61.02555226105376,61.02555226115376,61.03555966093995,61.035559661039954,61.04556546939084,61.04556546949084,61.055570254776164,61.055570254876166,61.065570190072954,61.065570190172956,61.07556501004198,61.07556501014198,61.08555391896559,61.08555391906559,61.09553665160457,61.09553665170457,61.10551851274327,61.10551851284327,61.11548972642195,61.11548972652195,61.12545783301266,61.12545783311266,61.13541919494899,61.13541919504899,61.14537854864553,61.14537854874553,61.15533365851459,61.155333658614595,61.16528751797021,61.16528751807021,61.175238118772576,61.17523811887258,61.18518773440071,61.18518773450071,61.1951345460714,61.1951345461714,61.20507859167597,61.20507859177597,61.215019340735985,61.21501934083599,61.224952397858694,61.224952397958695,61.23488306782845,61.23488306792845,61.254743043680556,61.25474304378056,61.26467083391024,61.26467083401024,61.27458945444126,61.274589454541264,61.28450341434033,61.28450341444033,61.294415365999605,61.29441536609961,61.30432356641852,61.30432356651852,61.31422771246655,61.31422771256655,61.324129092448445,61.324129092548446,61.33402960093006,61.33402960103006,61.35382895067536,61.35382895077536,61.37362257883182,61.37362257893182,61.38351391761478,61.38351391771478,61.39339415424193,61.393394154341934,61.40327279943378,61.40327279953378,61.4131421233617,61.413142123461704,61.42300962850642,61.42300962860642,61.43287558010716,61.43287558020716,61.4427370226412,61.4427370227412,61.46245649749076,61.462456497590765,61.472312748914405,61.47231274901441,61.4821553215727,61.4821553216727,61.49199372618615,61.491993726286154,61.51166284347575,61.51166284357575,61.52148817558492,61.52148817568492,61.53131199204142,61.53131199214142,61.54113061738752,61.541130617487525,61.55094787864622,61.55094787874622,61.56076407894805,61.56076407904805,61.57057933196696,61.57057933206696,61.5803937513769,61.580393751476905,61.600018119021414,61.600018119121415,61.60982734732096,61.60982734742096,61.61963149818406,61.619631498284065,61.629429169632,61.629429169732,61.63922286249167,61.63922286259167,61.64901367561128,61.64901367571128,61.65880354144796,61.65880354154796,61.66858711732606,61.668587117426064,61.67836955646466,61.67836955656466,61.68815070729849,61.68815070739849,61.707695692634395,61.7076956927344,61.71745630637454,61.71745630647454,61.72721566970124,61.72721566980124,61.73697435098423,61.73697435108423,61.746730493548995,61.746730493649,61.75648413528686,61.75648413538686,61.76623610980678,61.766236109906785,61.775982438520515,61.77598243862052,61.78572797151659,61.785727971616595,61.79547206464264,61.79547206474264,61.805212368637,61.805212368737,61.814949641326024,61.814949641426026,61.82468316277469,61.82468316287469,61.84414687123615,61.84414687133615,61.85387595940075,61.853875959500755,61.86359796188912,61.86359796198912,61.87331825926824,61.87331825936824,61.88303647262494,61.88303647272494,61.89275362502476,61.89275362512476,61.912185807910674,61.912185808010676,61.92189860280907,61.921898602909074,61.931600143986394,61.931600144086396,61.941301078902654,61.941301079002656,61.960698704907685,61.96069870500769,61.97039308462614,61.97039308472614,61.98008488773505,61.980084887835055,61.98977096925514,61.98977096935514,62.01882781183667,62.018827811936674,62.02850771707213,62.02850771717213,62.038183871067226,62.03818387116723,62.047858282061746,62.04785828216175,62.057527729293774,62.057527729393776,62.06719459991626,62.067194600016265,62.07685961386423,62.07685961396423,62.0865223543332,62.0865223544332,62.09618274554052,62.09618274564052,62.105840029659866,62.10584002975987,62.125149558353435,62.12514955845344,62.14445230450131,62.14445230460131,62.15410011579147,62.154100115891474,62.163743721145465,62.16374372124547,62.17338368893306,62.17338368903306,62.19266165415977,62.19266165425977,62.20229878009861,62.20229878019861,62.2119233640116,62.2119233641116,62.22154658383718,62.22154658393718,62.231169045836424,62.231169045936426,62.24078976483511,62.24078976493511,62.25040862715926,62.25040862725926,62.260026163287336,62.26002616338734,62.2792513459098,62.2792513460098,62.28885660525124,62.28885660535124,62.29846046261397,62.29846046271397,62.308062917997965,62.30806291809797,62.31765203563847,62.31765203573847,62.32723865245207,62.32723865255207,62.336824321982746,62.33682432208275,62.346403512098256,62.34640351219826,62.35597925410394,62.355979254203945,62.365551206977955,62.36555120707796,62.375121227394814,62.375121227494816,62.38468946691978,62.38468946701978,62.394251719616705,62.394251719716706,62.41337304213994,62.41337304223994,62.42292548098582,62.42292548108582,62.43247606315718,62.432476063257184,62.442023879262415,62.44202387936242,62.4515695355626,62.4515695356626,62.46109779974839,62.46109779984839,62.470623638889904,62.470623638989906,62.48014860653114,62.48014860663114,62.489670353410446,62.48967035351045,62.499188045918864,62.499188046018865,62.50870384386144,62.50870384396144,62.518216610498676,62.51821661059868,62.52772615637399,62.52772615647399,62.53723445183584,62.537234451935845,62.546741269536355,62.54674126963636,62.56575327561074,62.56575327571074,62.575258350310676,62.57525835041068,62.58474387309116,62.584743873191165,62.59422473523969,62.594224735339694,62.6037001410386,62.6037001411386,62.613174675337234,62.613174675437236,62.62263894108902,62.622638941189024,62.64156561591807,62.64156561601807,62.65102578940765,62.65102578950765,62.66994007377612,62.66994007387612,62.67939251743708,62.67939251753708,62.68884302864088,62.688843028740884,62.69829001595223,62.69829001605223,62.70773526026301,62.70773526036301,62.726624081666614,62.726624081766616,62.74550972019962,62.74550972029962,62.7643919106228,62.7643919107228,62.773825522299326,62.77382552239933,62.78325716362739,62.78325716372739,62.792687289302776,62.79268728940278,62.80211646769524,62.802116467795244,62.81154394097846,62.81154394107846,62.82096993650032,62.82096993660032,62.839820828695856,62.83982082879586,62.84924303508605,62.84924303518605,62.858663043779856,62.85866304387986,62.86807664884114,62.86807664894114,62.877484494422276,62.87748449452228,62.886886694197216,62.88688669429722,62.89628684784105,62.89628684794105,62.91508522267156,62.91508522277156,62.924482951271116,62.92448295137112,62.933879656805125,62.93387965690513,62.943275680295436,62.94327568039544,62.95265563786744,62.95265563796744,62.96203248835146,62.96203248845146,62.9714084673352,62.971408467435204,62.99015940223713,62.990159402337135,62.999532653046074,62.999532653146076,63.00890480500682,63.00890480510682,63.01827445614067,63.01827445624067,63.027637552076705,63.02763755217671,63.036999852295104,63.036999852395105,63.04636154625243,63.04636154635243,63.05571846590384,63.05571846600384,63.08378562518298,63.083785625282985,63.09313204893966,63.09313204903966,63.102473471042515,63.10247347114252,63.11181387007982,63.11181387017982,63.121152791355776,63.12115279145578,63.13982604905835,63.13982604915835,63.14915932452811,63.14915932462811,63.158490970671245,63.15849097077125,63.16782174531409,63.167821745414095,63.17714941286897,63.17714941296897,63.18647567844513,63.18647567854513,63.205121957530174,63.205121957630176,63.214437878776856,63.21443787887686,63.223751905457696,63.2237519055577,63.23303683160727,63.233036831707274,63.24231868856019,63.24231868866019,63.26088115205257,63.26088115215257,63.27944259247939,63.27944259257939,63.28872031927878,63.28872031937878,63.29799327177226,63.297993271872265,63.30726550433072,63.307265504430724,63.316537092736795,63.316537092836796,63.32580780964258,63.325807809742585,63.33507648041727,63.33507648051727,63.344343029278214,63.344343029378216,63.35360726676884,63.35360726686884,63.36286373653953,63.36286373663953,63.37211664452644,63.372116644626445,63.3813687567957,63.381368756895704,63.39061658734617,63.390616587446175,63.39986240965685,63.39986240975685,63.40910254827002,63.40910254837002,63.41833586644617,63.41833586654617,63.42756782053491,63.42756782063491,63.4367955686875,63.4367955687875,63.45524810946996,63.455248109569965,63.47368526637783,63.47368526647783,63.48289478880705,63.48289478890705,63.49209798338638,63.49209798348638,63.50130045803069,63.50130045813069,63.510502326413935,63.51050232651394,63.519698435317025,63.51969843541703,63.52888976991421,63.52888977001421,63.53807451142227,63.538074511522275,63.54725497121155,63.54725497131155,63.556433650108936,63.55643365020894,63.56561088913628,63.56561088923628,63.583957826818946,63.58395782691895,63.59312631295212,63.593126313052124,63.602292980302096,63.6022929804021,63.62062233641378,63.62062233651378,63.629783320066245,63.62978332016625,63.63894191656575,63.63894191666575,63.648096269237776,63.64809626933778,63.65724778006105,63.65724778016105,63.66639307670837,63.66639307680837,63.675536213550636,63.67553621365064,63.69382078212592,63.69382078222592,63.71210356980931,63.71210356990931,63.7212433722157,63.721243372315705,63.730376884663514,63.730376884763515,63.73950528178356,63.73950528188356,63.748631443315915,63.74863144341592,63.757755823956394,63.757755824056396,63.766877324856786,63.76687732495679,63.775995718669215,63.77599571876922,63.78509687193252,63.78509687203252,63.79419313721595,63.79419313731595,63.803285954389565,63.80328595448957,63.81237721801919,63.812377218119195,63.821465374560844,63.821465374660846,63.83055193966719,63.83055193976719,63.839631722227836,63.83963172232784,63.84871014070109,63.84871014080109,63.85778795291326,63.857787953013265,63.875939485075406,63.87593948517541,63.88501127256822,63.88501127266822,63.912220193522806,63.91222019362281,63.921285236361236,63.92128523646124,63.93034505019796,63.93034505029796,63.93939986238086,63.93939986248086,63.94845145380184,63.94845145390184,63.95750062017855,63.95750062027855,63.96654755096757,63.96654755106757,63.98463372060831,63.98463372070831,63.993664092891905,63.99366409299191,64.00268624275976,64.00268624285977,64.0117031257346,64.0117031258346,64.02071697740408,64.02071697750408,64.02972946498618,64.02972946508618,64.0387411189593,64.0387411190593,64.04774875645283,64.04774875655283,64.0567532868584,64.0567532869584,64.06575289139279,64.06575289149279,64.07475094238319,64.07475094248319,64.0837476292862,64.0837476293862,64.0927434067976,64.0927434068976,64.10173134080642,64.10173134090643,64.11970599630193,64.11970599640193,64.12869055798357,64.12869055808358,64.13767390714308,64.13767390724308,64.14664763190812,64.14664763200813,64.15561483936668,64.15561483946668,64.17354735971797,64.17354735981797,64.19147347643673,64.19147347653673,64.20042927860894,64.20042927870894,64.20938311043268,64.20938311053268,64.22728940999278,64.22728941009278,64.23624066520698,64.23624066530698,64.25413290708856,64.25413290718856,64.26307082455926,64.26307082465927,64.27200533181147,64.27200533191147,64.28093506475776,64.28093506485776,64.28986059176788,64.28986059186788,64.29878448945139,64.2987844895514,64.30770213506763,64.30770213516763,64.3255339781903,64.3255339782903,64.34335911454988,64.34335911464989,64.35226471072741,64.35226471082741,64.36116807131722,64.36116807141723,64.37897346630079,64.3789734664008,64.38787368191133,64.38787368201133,64.39677226819525,64.39677226829525,64.40566812630435,64.40566812640435,64.41456269610869,64.41456269620869,64.44124091128077,64.44124091138077,64.45012418947272,64.45012418957272,64.45900515629434,64.45900515639434,64.4678789995484,64.4678789996484,64.47675117558454,64.47675117568454,64.4856200171848,64.4856200172848,64.4944853348926,64.4944853349926,64.51221130967625,64.51221130977625,64.52106912490333,64.52106912500334,64.52991826301887,64.52991826311887,64.53876630228623,64.53876630238624,64.54761127235693,64.54761127245693,64.55645442364444,64.55645442374444,64.56529007245122,64.56529007255122,64.57412159110447,64.57412159120447,64.58295117730057,64.58295117740057,64.59177833845239,64.59177833855239,64.60060333979916,64.60060333989917,64.60942121757839,64.60942121767839,64.61823678398729,64.61823678408729,64.62705053161298,64.62705053171298,64.63586295304259,64.63586295314259,64.64467082751418,64.64467082761418,64.65346900180869,64.65346900190869,64.67106178861393,64.67106178871393,64.67985579486358,64.67985579496359,64.68864601198158,64.68864601208158,64.69743274309842,64.69743274319842,64.71500381817916,64.71500381827916,64.72378649492511,64.72378649502511,64.7325672013226,64.7325672014226,64.75012653009513,64.75012653019513,64.75890246218671,64.75890246228671,64.76767600712532,64.76767600722532,64.77644871845497,64.77644871855497,64.78521646602212,64.78521646612212,64.79398341787162,64.79398341797162,64.80274957400346,64.80274957410346,64.8202788170705,64.8202788171705,64.82904160087516,64.82904160097516,64.85532817139723,64.85532817149723,64.86408913641868,64.86408913651869,64.87284938150512,64.87284938160512,64.88160746678649,64.88160746688649,64.89036202817542,64.89036202827542,64.89911458132454,64.89911458142454,64.90786478521203,64.90786478531203,64.91661335977291,64.91661335987291,64.925360646029,64.92536064612901,64.93410675765429,64.9341067577543,64.94284938327843,64.94284938337843,64.95158693146612,64.95158693156613,64.96031576465096,64.96031576475096,64.97777206693324,64.97777206703324,64.98649745200825,64.98649745210825,64.99521961632135,64.99521961642135,65.00393689265458,65.00393689275458,65.01264992516033,65.01264992526033,65.02135886540387,65.02135886550387,65.03877394193351,65.03877394203352,65.04747916882802,65.04747916892802,65.05617802998131,65.05617803008131,65.06487120743708,65.06487120753708,65.07356362706652,65.07356362716652,65.09094683699877,65.09094683709877,65.10832371908111,65.10832371918112,65.11699560161688,65.11699560171688,65.12566524856494,65.12566524866494,65.14299567589295,65.14299567599295,65.15165414490257,65.15165414500257,65.16031030254187,65.16031030264188,65.17762072325463,65.17762072335464,65.20358067062627,65.20358067072627,65.21223201606833,65.21223201616833,65.22088108803139,65.22088108813139,65.22952584038435,65.22952584048436,65.24681310950258,65.24681310960258,65.2640868135294,65.2640868136294,65.27271667459488,65.27271667469488,65.281343504355,65.281343504455,65.29859458726571,65.29859458736571,65.30721698374178,65.30721698384178,65.31583695517358,65.31583695527358,65.32445613088773,65.32445613098773,65.33307254053575,65.33307254063575,65.34168713140056,65.34168713150056,65.35029834993819,65.35029835003819,65.35890619614862,65.35890619624863,65.37612063815604,65.37612063825604,65.38472371006056,65.38472371016056,65.3933229549421,65.3933229550421,65.4105152305292,65.4105152306292,65.41910659401685,65.41910659411685,65.43628443301228,65.43628443311228,65.44486844558446,65.44486844568446,65.45344791119865,65.45344791129865,65.46202620218202,65.46202620228202,65.47059858211996,65.47059858221996,65.4791652783604,65.4791652784604,65.48773129255713,65.48773129265713,65.49628919801208,65.49628919811208,65.51340277333428,65.51340277343428,65.52195378256958,65.52195378266958,65.53049478849725,65.53049478859725,65.53903492292464,65.53903492302464,65.55611352456148,65.55611352466148,65.58172597066712,65.58172597076712,65.59879381117001,65.59879381127001,65.6158578246499,65.6158578247499,65.62438742529123,65.62438742539123,65.63291634388885,65.63291634398885,65.6414412081156,65.6414412082156,65.64995565223022,65.64995565233022,65.65846918695324,65.65846918705324,65.66698185017597,65.66698185027597,65.67549008011466,65.67549008021466,65.68399194431213,65.68399194441213,65.69248941311685,65.69248941321685,65.70098191815906,65.70098191825906,65.70947336224442,65.70947336234443,65.71796408639476,65.71796408649476,65.72645284019661,65.72645284029662,65.73493541771386,65.73493541781386,65.74341647957841,65.74341647967842,65.75189572265978,65.75189572275978,65.76885303419168,65.76885303429168,65.77732989012007,65.77732989022007,65.78580163072071,65.78580163082071,65.79426484577509,65.79426484587509,65.80272548421992,65.80272548431992,65.81964285830395,65.81964285840395,65.82809428915881,65.82809428925881,65.83654412857837,65.83654412867837,65.84499271758447,65.84499271768448,65.85343910889421,65.85343910899421,65.86188159739831,65.86188159749831,65.8703233659674,65.8703233660674,65.87876233057904,65.87876233067904,65.88719239073124,65.88719239083125,65.90404659999025,65.90404660009025,65.92089186689849,65.92089186699849,65.92930930924221,65.92930930934222,65.93772478123748,65.93772478133748,65.94613741138397,65.94613741148397,65.97137048962624,65.97137048972624,65.97977728450996,65.97977728460997,65.98817259832471,65.98817259842471,65.99656692696522,65.99656692706522,66.01335069626636,66.01335069636636,66.02173202818521,66.02173202828521,66.03010669123232,66.03010669133232,66.04685101567273,66.04685101577273,66.0552152207164,66.05521522081641,66.06357669758529,66.06357669768529,66.0719262765807,66.0719262766807,66.08027479461924,66.08027479471924,66.08861690902526,66.08861690912526,66.09695690151754,66.09695690161755,66.10529488577002,66.10529488587002,66.11363203641355,66.11363203651355,66.12196607996908,66.12196608006909,66.13029375778342,66.13029375788342,66.138619313684,66.138619313784,66.15526678791878,66.15526678801878,66.16358393194704,66.16358393204705,66.17189584697361,66.17189584707361,66.18020317715084,66.18020317725085,66.19681374524309,66.19681374534309,66.20511527804885,66.20511527814885,66.2134108619179,66.2134108620179,66.22169780656671,66.22169780666671,66.22997819601773,66.22997819611773,66.2382510450967,66.2382510451967,66.26306860715941,66.26306860725941,66.27960816692405,66.27960816702405,66.2878743471313,66.2878743472313,66.29613795072898,66.29613795082898,66.30439897771713,66.30439897781713,66.31265735231311,66.31265735241311,66.32091443860432,66.32091443870432,66.32916349193637,66.32916349203637,66.33741102961577,66.33741102971577,66.3456537172066,66.3456537173066,66.35389466179687,66.35389466189687,66.36213064262465,66.36213064272465,66.37035927253697,66.37035927263697,66.3785869930577,66.3785869931577,66.3868139936434,66.3868139937434,66.39503864496747,66.39503864506747,66.40325776415929,66.4032577642593,66.41147404150234,66.41147404160235,66.41968842427957,66.41968842437957,66.42790220079573,66.42790220089573,66.43611271865865,66.43611271875865,66.4443210388252,66.4443210389252,66.46073324587421,66.46073324597421,66.46893391199477,66.46893391209477,66.47712972802678,66.47712972812678,66.48531982246996,66.48531982256996,66.49350722662966,66.49350722672966,66.50169402452828,66.50169402462828,66.5098794583395,66.5098794584395,66.51806349017201,66.51806349027201,66.526246385265,66.526246385365,66.53442556700897,66.53442556710897,66.54260387725263,66.54260387735263,66.55078158123523,66.55078158133523,66.55895587499933,66.55895587509933,66.5671253944575,66.56712539455751,66.57529411819804,66.57529411829805,66.5834547710881,66.5834547711881,66.5916075804756,66.5916075805756,66.6079121004024,66.6079121005024,66.6160568010481,66.6160568011481,66.62419994814982,66.62419994824982,66.64048502983113,66.64048502993113,66.64862632025833,66.64862632035833,66.6567650719673,66.6567650720673,66.66490306584994,66.66490306594994,66.67303977142781,66.67303977152781,66.68116499593671,66.68116499603671,66.68928078550773,66.68928078560774,66.69739566568715,66.69739566578716,66.71362239474065,66.71362239484066,66.7217249223508,66.7217249224508,66.72982612376487,66.72982612386487,66.7379245212215,66.7379245213215,66.74602136513414,66.74602136523414,66.76221270369778,66.76221270379779,66.77030488697847,66.77030488707847,66.77839115921374,66.77839115931374,66.78647413490445,66.78647413500445,66.79455339724612,66.79455339734612,66.80263118182643,66.80263118192643,66.81878239348562,66.81878239358562,66.82685267558522,66.82685267568522,66.83492004005342,66.83492004015342,66.84298657091266,66.84298657101266,66.85104866848783,66.85104866858784,66.8671690745065,66.86716907460651,66.88328584295878,66.88328584305879,66.89134218105382,66.89134218115382,66.89939787499645,66.89939787509645,66.90745231852564,66.90745231862564,66.92355950047477,66.92355950057477,66.93161204943813,66.93161204953813,66.93966357533593,66.93966357543593,66.95575393354041,66.95575393364041,66.97182712697841,66.97182712707841,66.97984557375668,66.97984557385668,66.98786227753438,66.98786227763438,66.99587602578937,66.99587602588937,67.01189935425451,67.01189935435451,67.01990825242096,67.01990825252096,67.02791525602157,67.02791525612157,67.03591881151236,67.03591881161236,67.04392066189389,67.0439206619939,67.05991996726421,67.05991996736421,67.07591396785018,67.07591396795019,67.08390600438067,67.08390600448067,67.09189561586689,67.09189561596689,67.09988276441752,67.09988276451752,67.10786904146786,67.10786904156787,67.12383701071923,67.12383701081923,67.13181548215832,67.13181548225832,67.13978720894302,67.13978720904302,67.14775507081342,67.14775507091342,67.15571959824794,67.15571959834794,67.16368105648581,67.16368105658582,67.18755970961058,67.18755970971058,67.19551249073692,67.19551249083692,67.20346299838424,67.20346299848424,67.21141145990046,67.21141146000046,67.2193536314581,67.2193536315581,67.22729356742805,67.22729356752805,67.23523285924561,67.23523285934561,67.24317143112816,67.24317143122816,67.25110488768294,67.25110488778294,67.26696638233422,67.26696638243422,67.27489066919034,67.27489066929034,67.29073795459783,67.29073795469783,67.29865401903822,67.29865401913823,67.3144847838316,67.3144847839316,67.32239918105407,67.32239918115407,67.33030421912127,67.33030421922128,67.33820846147084,67.33820846157084,67.3540045936007,67.35400459370071,67.36978848683526,67.36978848693526,67.37767950511527,67.37767950521527,67.3934572599565,67.39345726005651,67.40133990425552,67.40133990435552,67.41710185841767,67.41710185851767,67.43286017501342,67.43286017511342,67.44073812078915,67.44073812088915,67.44861436145563,67.44861436155563,67.46436407672246,67.46436407682246,67.48010992707499,67.480109927175,67.48797866526075,67.48797866536076,67.49584414479328,67.49584414489328,67.51157040533504,67.51157040543504,67.51943023906136,67.51943023916137,67.53514812562213,67.53514812572213,67.5430049280431,67.5430049281431,67.55086044215932,67.55086044225932,67.55871512266657,67.55871512276657,67.5665674918035,67.5665674919035,67.57441879998355,67.57441880008355,67.58226825148908,67.58226825158908,67.59011508849375,67.59011508859375,67.5979566586054,67.5979566587054,67.60579019575789,67.60579019585789,67.61362051214846,67.61362051224846,67.62928015975537,67.62928015985537,67.64493726864406,67.64493726874406,67.65276478107718,67.65276478117718,67.66059127044475,67.66059127054476,67.66841715355127,67.66841715365128,67.67623822446055,67.67623822456055,67.69187892640907,67.69187892650908,67.69969639764327,67.69969639774327,67.70751254268137,67.70751254278137,67.72314365812676,67.72314365822676,67.73095798438165,67.73095798448165,67.74658576539116,67.74658576549116,67.75439425638328,67.75439425648328,67.76219888246109,67.7621988825611,67.77780389078923,67.77780389088923,67.78559241305742,67.78559241315742,67.80115657454608,67.80115657464609,67.81670785298707,67.81670785308707,67.82448144607645,67.82448144617645,67.83225143949075,67.83225143959075,67.84778900127506,67.84778900137506,67.855555963384,67.85555596348401,67.86332201610136,67.86332201620137,67.87108522696995,67.87108522706995,67.87884559598979,67.87884559608979,67.88660153172556,67.88660153182556,67.89435473928653,67.89435473938653,67.90210703745589,67.9021070375559,67.90985009014398,67.90985009024398,67.9175918924186,67.9175918925186,67.92533210325792,67.92533210335792,67.93306988905297,67.93306988915297,67.94080543926033,67.94080543936033,67.94853943592369,67.9485394360237,67.95627195482572,67.95627195492573,67.97173589378157,67.97173589388157,67.97946450987796,67.97946450997796,67.98719032201691,67.98719032211692,67.9949155278948,67.9949155279948,68.0103616958231,68.0103616959231,68.02579854248746,68.02579854258747,68.03351063796975,68.03351063806976,68.04893338906429,68.04893338916429,68.05664347630679,68.05664347640679,68.06434606106856,68.06434606116856,68.07975001806999,68.07975001816999,68.09515101954871,68.09515101964871,68.10284856476535,68.10284856486535,68.11054463222065,68.11054463232065,68.11823077215097,68.11823077225097,68.1336017258155,68.1336017259155,68.14897112593604,68.14897112603605,68.15665097590777,68.15665097600777,68.17200609100189,68.17200609110189,68.17967938577581,68.17967938587582,68.18734586011273,68.18734586021273,68.1950088484485,68.1950088485485,68.2026683128918,68.2026683129918,68.21798637027814,68.21798637037814,68.22564280341611,68.22564280351611,68.23329787246668,68.23329787256668,68.24094903871162,68.24094903881162,68.24859903032574,68.24859903042574,68.25624830200483,68.25624830210484,68.26389564122678,68.26389564132678,68.27153979757813,68.27153979767813,68.27918168045046,68.27918168055047,68.28682034256087,68.28682034266087,68.29445801949704,68.29445801959704,68.30209096001863,68.30209096011863,68.30971859575587,68.30971859585587,68.31734475373175,68.31734475383175,68.32496822142413,68.32496822152413,68.3325902492465,68.3325902493465,68.3478266129539,68.3478266130539,68.37066776393453,68.37066776403454,68.37827721183972,68.37827721193972,68.38588567457067,68.38588567467067,68.39349034816996,68.39349034826996,68.40109392292105,68.40109392302105,68.40869632304133,68.40869632314133,68.42389972130316,68.42389972140316,68.4314909813763,68.4314909814763,68.43908065001415,68.43908065011415,68.4466692198038,68.4466692199038,68.46184491951307,68.46184491961307,68.4694302685408,68.4694302686408,68.48459611650773,68.48459611660773,68.49217419040265,68.49217419050265,68.49975025605777,68.49975025615777,68.5073247302776,68.5073247303776,68.51489378603912,68.51489378613913,68.52246212186563,68.52246212196563,68.54516447695299,68.54516447705299,68.552730122496,68.552730122596,68.56029288829895,68.56029288839895,68.56785103136124,68.56785103146125,68.57540576420504,68.57540576430505,68.59051341110943,68.59051341120943,68.59806208134253,68.59806208144254,68.60561006953195,68.60561006963195,68.61315623893815,68.61315623903815,68.62070074112641,68.62070074122641,68.62824221200934,68.62824221210934,68.63577826443398,68.63577826453398,68.6433127254233,68.6433127255233,68.65084498871626,68.65084498881626,68.6583765320742,68.6583765321742,68.67342931235191,68.67342931245192,68.68847064945197,68.68847064955197,68.69598957500143,68.69598957510144,68.70350725013743,68.70350725023744,68.71102401588183,68.71102401598183,68.71853968277804,68.71853968287805,68.72605428871739,68.7260542888174,68.74108062085601,68.74108062095601,68.74859109664183,68.74859109674183,68.75610013255762,68.75610013265762,68.7636080317339,68.7636080318339,68.77111217966981,68.77111217976982,68.77861503930097,68.77861503940098,68.78611282149568,68.78611282159568,68.79360438951444,68.79360438961444,68.80858661616035,68.80858661626036,68.81607375089504,68.81607375099505,68.82355936997708,68.82355937007708,68.83104309449327,68.83104309459327,68.83852477287834,68.83852477297835,68.86096226766156,68.86096226776156,68.86843250286898,68.86843250296899,68.88336907047821,68.88336907057821,68.89829146673496,68.89829146683496,68.90574684854622,68.90574684864622,68.92065658910317,68.92065658920318,68.92810117188915,68.92810117198916,68.950429047093,68.950429047193,68.95786658209406,68.95786658219406,68.96530237409455,68.96530237419455,68.97273229294095,68.97273229304095,68.98016084769993,68.98016084779994,68.98758739421913,68.98758739431914,68.9950106820851,68.9950106821851,69.0024304460586,69.0024304461586,69.01726576806944,69.01726576816944,69.02468060617177,69.02468060627177,69.03209044262027,69.03209044272027,69.04690488651559,69.0469048866156,69.05430782674445,69.05430782684445,69.06170955445118,69.06170955455119,69.06911033487499,69.069110334975,69.08390844761277,69.08390844771277,69.09130460529593,69.09130460539593,69.10609369990034,69.10609370000034,69.11348606845182,69.11348606855182,69.12087544358928,69.12087544368929,69.12826341674803,69.12826341684803,69.1356507836457,69.1356507837457,69.15042138728752,69.15042138738752,69.15780356307481,69.15780356317481,69.16518297279596,69.16518297289596,69.17256094264708,69.17256094274708,69.17993459288809,69.17993459298809,69.18730763686803,69.18730763696803,69.19467734641209,69.1946773465121,69.20204072810627,69.20204072820627,69.20940149529957,69.20940149539958,69.21675983744862,69.21675983754862,69.22411590611866,69.22411590621866,69.23146989076626,69.23146989086626,69.23882194295672,69.23882194305672,69.253523698076,69.253523698176,69.26087120330844,69.26087120340844,69.26821726867085,69.26821726877085,69.27556071953241,69.27556071963241,69.28290284419786,69.28290284429787,69.29758553998481,69.29758554008481,69.3049238755186,69.3049238756186,69.31959910671615,69.31959910681616,69.32693562346674,69.32693562356674,69.34160517096676,69.34160517106676,69.34893589034589,69.34893589044589,69.35626437413733,69.35626437423733,69.36359156962399,69.363591569724,69.37091626428375,69.37091626438375,69.385561675015,69.385561675115,69.39288212584728,69.39288212594728,69.40020117470084,69.40020117480084,69.41483662001579,69.41483662011579,69.42215093245476,69.42215093255476,69.43677872372372,69.43677872382372,69.44409144472739,69.4440914448274,69.45140143755624,69.45140143765624,69.45870730023157,69.45870730033157,69.46600926010127,69.46600926020128,69.47330993166621,69.47330993176621,69.48060980751349,69.48060980761349,69.48790312816297,69.48790312826297,69.4951931522679,69.4951931523679,69.50248249432912,69.50248249442912,69.50977069965084,69.50977069975085,69.51705807136359,69.5170580714636,69.52434286646681,69.52434286656681,69.53162319039464,69.53162319049464,69.54618239838027,69.54618239848027,69.5534602214812,69.5534602215812,69.56073327027622,69.56073327037622,69.57527686703935,69.57527686713935,69.5825454825503,69.5825454826503,69.60434155312345,69.60434155322345,69.61159739926066,69.61159739936066,69.61885138872336,69.61885138882336,69.62609954292326,69.62609954302326,69.63334538575285,69.63334538585285,69.64059005395161,69.64059005405161,69.64783324438902,69.64783324448902,69.65507556332615,69.65507556342615,69.66231704865432,69.66231704875432,69.66955660152531,69.66955660162532,69.68403347167964,69.68403347177964,69.69127018270189,69.69127018280189,69.69850329404903,69.69850329414903,69.71296872102569,69.71296872112569,69.72019902841541,69.72019902851541,69.73465687712871,69.73465687722872,69.74910362368621,69.74910362378621,69.75631229513407,69.75631229523407,69.76351922358134,69.76351922368134,69.77072433324541,69.77072433334541,69.78512985405027,69.78512985415027,69.79232992416921,69.79232992426921,69.79952798604837,69.79952798614838,69.80672479751406,69.80672479761407,69.82111637431437,69.82111637441437,69.82830731262597,69.82830731272597,69.83549684895885,69.83549684905886,69.84268392235614,69.84268392245615,69.86424286906903,69.86424286916903,69.87142494081252,69.87142494091252,69.88578192284061,69.88578192294061,69.89295186936273,69.89295186946273,69.90012014866691,69.90012014876692,69.90728611660077,69.90728611670077,69.91444947003379,69.91444947013379,69.92161043631383,69.92161043641383,69.92877068265886,69.92877068275887,69.93592945124254,69.93592945134255,69.94308719676066,69.94308719686066,69.95024100158184,69.95024100168185,69.95739374544615,69.95739374554616,69.96454531467965,69.96454531477966,69.97884470190627,69.97884470200627,69.9859918378557,69.98599183795571,69.99313749604379,69.99313749614379,70.00028205538368,70.00028205548368,70.01456890058446,70.01456890068447,70.02170694261787,70.02170694271787,70.02884077871512,70.02884077881512,70.04310181992919,70.0431018200292,70.0644774484516,70.0644774485516,70.07159658271794,70.07159658281795,70.08583291879349,70.08583291889349,70.09294712718865,70.09294712728865,70.10005686440844,70.10005686450845,70.10716394923607,70.10716394933607,70.11427023834604,70.11427023844604,70.12848035363038,70.12848035373038,70.13557834454198,70.13557834464198,70.14266985603841,70.14266985613841,70.1497603444693,70.1497603445693,70.15685022663911,70.15685022673911,70.16393919941731,70.16393919951732,70.17102673232549,70.17102673242549,70.19228508722253,70.19228508732253,70.19936856575981,70.19936856585981,70.20645041497046,70.20645041507046,70.21353090009373,70.21353090019373,70.22061040004274,70.22061040014275,70.23476811163601,70.23476811173602,70.24184548967129,70.2418454897713,70.24892165518445,70.24892165528445,70.26307072755749,70.26307072765749,70.27013336587056,70.27013336597057,70.27719213926932,70.27719213936932,70.28424735088429,70.28424735098429,70.29129740928019,70.29129740938019,70.29834564889289,70.29834564899289,70.30539021304786,70.30539021314786,70.31243257950645,70.31243257960645,70.32651484948806,70.32651484958807,70.3335534647063,70.3335534648063,70.34762811853324,70.34762811863324,70.35466233835874,70.35466233845874,70.36169163231305,70.36169163241306,70.37574502911129,70.37574502921129,70.38276735106332,70.38276735116332,70.38978470925285,70.38978470935285,70.39679774783228,70.39679774793228,70.40380927075904,70.40380927085904,70.41783057361198,70.41783057371198,70.42483857264628,70.42483857274628,70.43885294138826,70.43885294148826,70.45286530189045,70.45286530199046,70.45987011805414,70.45987011815414,70.46687383536964,70.46687383546964,70.48087774610819,70.4808777462082,70.48787532503039,70.48787532513039,70.50186870198289,70.50186870208289,70.51586022226086,70.51586022236086,70.52285423939927,70.52285423949927,70.52984700612423,70.52984700622423,70.54383102392148,70.54383102402149,70.55081556823072,70.55081556833072,70.5647824970441,70.56478249714411,70.571764010048,70.571764010148,70.57874434842105,70.57874434852106,70.58572229964118,70.58572229974118,70.59269918990441,70.59269919000441,70.5996750571021,70.5996750572021,70.61362357073556,70.61362357083556,70.62059242803964,70.62059242813964,70.62755863295156,70.62755863305156,70.63452385268924,70.63452385278924,70.64845307964247,70.64845307974247,70.65541379031345,70.65541379041345,70.66933100571926,70.66933100581926,70.67628705575828,70.67628705585828,70.68324090810093,70.68324090820093,70.69019377526935,70.69019377536935,70.7040970466706,70.7040970467706,70.71104434381544,70.71104434391545,70.71799080735133,70.71799080745133,70.72493435325583,70.72493435335583,70.73187498152893,70.73187498162893,70.73881379101883,70.73881379111883,70.74574907661628,70.74574907671628,70.75961847318034,70.75961847328034,70.77348396693877,70.77348396703877,70.78041296257763,70.78041296267763,70.78733722180189,70.7873372219019,70.79426015483007,70.79426015493007,70.80118248159718,70.80118248169718,70.81502550580478,70.81502550590479,70.82194139104804,70.82194139114804,70.82885560907337,70.82885560917337,70.83576755361969,70.83576755371969,70.84267779305675,70.84267779315675,70.85649228510283,70.85649228520283,70.86339536308103,70.86339536318103,70.87029643281944,70.87029643291945,70.87719587323123,70.87719587333123,70.88409379799035,70.88409379809035,70.8909908891405,70.8909908892405,70.89788097039704,70.89788097049704,70.90476176828099,70.904761768381,70.9185211663525,70.9185211664525,70.92539840245267,70.92539840255267,70.93227173574722,70.93227173584722,70.93914271978012,70.93914271988012,70.94601283231275,70.94601283241275,70.95287998932265,70.95287998942266,70.95974566857122,70.95974566867122,70.96660979427578,70.96660979437578,70.97347217697978,70.97347217707978,70.99405682426487,70.99405682436488,71.00777632277985,71.00777632287985,71.01463082408996,71.01463082418996,71.0214738443311,71.0214738444311,71.02831307544055,71.02831307554055,71.04199074194183,71.04199074204183,71.04882853318125,71.04882853328125,71.05566261107163,71.05566261117163,71.06249452915696,71.06249452925697,71.06932011939239,71.06932011949239,71.07614461077962,71.07614461087962,71.08296815488396,71.08296815498396,71.08978957707454,71.08978957717454,71.09660948361245,71.09660948371246,71.10342355488758,71.10342355498759,71.11023626207532,71.11023626217532,71.11704715047985,71.11704715057985,71.1238560685359,71.1238560686359,71.13747261634323,71.13747261644323,71.15108454140993,71.15108454150993,71.16468854719143,71.16468854729143,71.17148723459196,71.17148723469197,71.19188140222771,71.19188140232771,71.19867789193187,71.19867789203187,71.20547176713517,71.20547176723517,71.21226442981634,71.21226442991635,71.22584448828565,71.22584448838565,71.23263086100823,71.23263086110823,71.23941465712127,71.23941465722127,71.25297929382465,71.25297929392465,71.25975691365306,71.25975691375307,71.26653377565515,71.26653377575515,71.28007916356961,71.28007916366961,71.29361515443753,71.29361515453753,71.30714845502195,71.30714845512195,71.32067940634474,71.32067940644474,71.32744194542909,71.32744194552909,71.334200127012,71.334200127112,71.34771523976438,71.34771523986439,71.35447175412936,71.35447175422937,71.3679808800537,71.3679808801537,71.37473307480856,71.37473307490856,71.38148322343233,71.38148322353233,71.38822909033729,71.38822909043729,71.3949743509812,71.3949743510812,71.40846312926843,71.40846312936843,71.41520611643332,71.41520611653333,71.42194830788057,71.42194830798057,71.43543056886132,71.43543056896132,71.44216870593768,71.44216870603768,71.45564263082873,71.45564263092874,71.46237811551292,71.46237811561292,71.46911288026207,71.46911288036208,71.48258040152061,71.48258040162061,71.48930330628762,71.48930330638763,71.50946819356568,71.50946819366568,71.51618382319988,71.51618382329988,71.52289691411586,71.52289691421586,71.52960560963909,71.5296056097391,71.54302197762,71.54302197772,71.54972862701213,71.54972862711213,71.55643444279528,71.55643444289528,71.56984482394815,71.56984482404815,71.57654575175144,71.57654575185144,71.58324580805446,71.58324580815446,71.58994487918322,71.58994487928322,71.60334033115728,71.60334033125729,71.61003167245744,71.61003167255744,71.62340916394736,71.62340916404736,71.63009448052816,71.63009448062816,71.63677797832574,71.63677797842574,71.64345662603478,71.64345662613478,71.65012913535051,71.65012913545051,71.65679918173065,71.65679918183065,71.67013620529427,71.67013620539427,71.67680162893376,71.67680162903376,71.68346610529034,71.68346610539034,71.69012917966819,71.69012917976819,71.70344210435435,71.70344210445435,71.71673468140342,71.71673468150343,71.72337449051278,71.72337449061278,71.73001285975212,71.73001285985212,71.7366502059259,71.7366502060259,71.74328395242458,71.74328395252458,71.75654212415803,71.75654212425803,71.7697881327788,71.7697881328788,71.77640702588131,71.77640702598131,71.78964276595524,71.78964276605524,71.79625828673056,71.79625828683056,71.80287301178824,71.80287301188824,71.81610162829462,71.81610162839462,71.8293282365612,71.8293282366612,71.83593909670456,71.83593909680457,71.84254635717284,71.84254635727284,71.85575879408695,71.85575879418695,71.87557276988052,71.87557276998052,71.88217381617284,71.88217381627284,71.8887727026601,71.88877270276011,71.89537033873393,71.89537033883393,71.90196482982846,71.90196482992846,71.90855674431344,71.90855674441345,71.91514687790654,71.91514687800654,71.92832589467928,71.92832589477928,71.93491348955415,71.93491348965415,71.94149797734104,71.94149797744105,71.94808117682318,71.94808117692318,71.95466149656524,71.95466149666524,71.96123984595883,71.96123984605883,71.96781758909134,71.96781758919134,71.98097171126898,71.98097171136898,71.98754634731351,71.98754634741351,71.9941195055967,71.9941195056967,72.00069137557513,72.00069137567513,72.0138301896608,72.0138301897608,72.02039489818034,72.02039489828034,72.0269582426125,72.0269582427125,72.04008197595408,72.04008197605408,72.04663929566686,72.04663929576687,72.05319483448777,72.05319483458777,72.06630481328138,72.06630481338138,72.07285716923167,72.07285716933167,72.08595877404427,72.08595877414427,72.09250734086288,72.09250734096288,72.09904704111338,72.09904704121338,72.10558450577618,72.10558450587618,72.11212053056894,72.11212053066895,72.11865344827375,72.11865344837375,72.12518223582501,72.12518223592501,72.13170624907036,72.13170624917036,72.13822745835829,72.13822745845829,72.15126775502037,72.15126775512037,72.15777956726173,72.15777956736173,72.16428978806779,72.16428978816779,72.17079917526488,72.17079917536488,72.17730734993982,72.17730735003983,72.1838146531145,72.1838146532145,72.19032078165834,72.19032078175834,72.19682622815849,72.19682622825849,72.20333068948439,72.20333068958439,72.2098308690915,72.2098308691915,72.2163301014157,72.21633010151571,72.22282736339143,72.22282736349143,72.22932238977945,72.22932238987946,72.23581677201511,72.23581677211511,72.24230709987987,72.24230709997987,72.2487967078096,72.2487967079096,72.25528536845643,72.25528536855643,72.26177065677606,72.26177065687607,72.27473505713068,72.27473505723069,72.2876921823525,72.2876921824525,72.29416640640763,72.29416640650763,72.30063994841906,72.30063994851906,72.3071128083868,72.3071128084868,72.32005720212618,72.32005720222618,72.33947036357108,72.33947036367108,72.34593613786258,72.34593613796258,72.35886677705398,72.35886677715398,72.37826015923153,72.37826015933153,72.39764914601633,72.39764914611634,72.40410980498008,72.40410980508008,72.41056474235498,72.41056474245498,72.41701592848953,72.41701592858954,72.42346215086158,72.42346215096158,72.43635175375692,72.43635175385693,72.44279339127965,72.44279339137965,72.44923063340964,72.44923063350964,72.45566643566958,72.45566643576959,72.46210155588582,72.46210155598583,72.4685349709928,72.46853497109281,72.4749658852729,72.4749658853729,72.48139088850756,72.48139088860756,72.49423763632366,72.49423763642366,72.5006558191213,72.50065581922131,72.50707123585283,72.50707123595284,72.51989805283631,72.51989805293631,72.53271933768755,72.53271933778755,72.5391262288728,72.5391262289728,72.55193660102482,72.55193660112482,72.55833996831761,72.55833996841761,72.56474144104457,72.56474144114458,72.57114173914071,72.57114173924072,72.57754040791023,72.57754040801024,72.58393797783158,72.58393797793158,72.5967311852171,72.5967311853171,72.60312606485493,72.60312606495494,72.60951798897008,72.60951798907008,72.61590858688913,72.61590858698914,72.62229698711181,72.62229698721181,72.63507276449162,72.63507276459163,72.64784615471847,72.64784615481847,72.65422868178705,72.65422868188705,72.66061003422482,72.66061003432482,72.66699006046649,72.66699006056649,72.67974738477504,72.67974738487504,72.6924981538858,72.6924981539858,72.69886734321088,72.69886734331088,72.70523569892698,72.70523569902699,72.7307024529197,72.7307024530197,72.74341946086722,72.74341946096722,72.74977292529584,72.74977292539585,72.75612532876761,72.75612532886761,72.7624737157598,72.7624737158598,72.77516851939569,72.7751685194957,72.78151364773464,72.78151364783464,72.79419825860633,72.79419825870633,72.80053747589986,72.80053747599986,72.80687441971438,72.80687441981438,72.82587934011254,72.82587934021254,72.8385403825852,72.8385403826852,72.84486899031003,72.84486899041003,72.85119236903316,72.85119236913316,72.85751426999494,72.85751427009494,72.86383029780261,72.86383029790261,72.87014518887078,72.87014518897078,72.87645841272101,72.87645841282101,72.88908171544219,72.88908171554219,72.90169975127033,72.90169975137033,72.90800270657373,72.90800270667373,72.91430210009334,72.91430210019334,72.92059956115581,72.92059956125581,72.92689630228325,72.92689630238326,72.94578463109974,72.94578463119974,72.95207697683445,72.95207697693445,72.9646590916943,72.9646590917943,72.98352635293868,72.98352635303868,72.98981013523579,72.98981013533579,72.99609308392394,72.99609308402394,73.00236799965293,73.00236799975293,73.00864181653374,73.00864181663374,73.01491385252265,73.01491385262266,73.0211811899883,73.0211811900883,73.02744773173627,73.02744773183628,73.03997892066641,73.03997892076642,73.04624345417461,73.04624345427462,73.05877070240781,73.05877070250781,73.06503334135017,73.06503334145017,73.07755752038672,73.07755752048672,73.08381474087078,73.08381474097078,73.0900705214848,73.0900705215848,73.10257689160247,73.10257689170247,73.10882615491002,73.10882615501002,73.11507375099963,73.11507375109963,73.1213196798713,73.12131967997131,73.12756462356873,73.12756462366873,73.1338082789614,73.1338082790614,73.14004859991819,73.14004860001819,73.14628732943969,73.14628732953969,73.15252348235163,73.15252348245163,73.16499317367467,73.16499317377468,73.17122356710648,73.17122356720648,73.17745225542902,73.17745225552902,73.18367730618516,73.18367730628516,73.18990095496258,73.18990095506258,73.1961236943484,73.1961236944484,73.20234389501599,73.20234389511599,73.20856261792223,73.20856261802223,73.21477630128334,73.21477630138334,73.22098915103548,73.22098915113548,73.2334126528434,73.2334126529434,73.23962262285546,73.23962262295547,73.24583175925856,73.24583175935857,73.25824548510676,73.25824548520676,73.27685588111484,73.27685588121484,73.2830563404064,73.2830563405064,73.28925596608899,73.28925596618899,73.29545456870603,73.29545456880604,73.30164964743062,73.30164964753062,73.31403321179067,73.31403321189067,73.32641033462687,73.32641033472687,73.33258862749813,73.33258862759813,73.33876510158619,73.33876510168619,73.35111202504294,73.35111202514294,73.35728213338977,73.35728213348978,73.36345046084473,73.36345046094473,73.36961640114673,73.36961640124673,73.37578105314395,73.37578105324395,73.38194403792325,73.38194403802325,73.39426674882858,73.39426674892859,73.41274467679328,73.41274467689328,73.41890114426609,73.41890114436609,73.42505511091198,73.42505511101199,73.43735864881104,73.43735864891104,73.44350795482498,73.44350795492498,73.45580471017834,73.45580471027834,73.46809926783533,73.46809926793533,73.47424504995682,73.47424505005682,73.48038817968612,73.48038817978612,73.48652638354426,73.48652638364426,73.49880169241233,73.49880169251233,73.50493663761722,73.50493663771722,73.51720508815696,73.51720508825696,73.52333662314334,73.52333662324334,73.52946194395378,73.52946194405378,73.53558389243702,73.53558389253702,73.54782589483766,73.54782589493766,73.55394409208054,73.55394409218054,73.5661791224789,73.5661791225789,73.57229292432905,73.57229292442905,73.58451927761587,73.58451927771587,73.59063019972594,73.59063019982594,73.59673816631329,73.59673816641329,73.60284495826983,73.60284495836983,73.61505691285629,73.61505691295629,73.62115809689794,73.62115809699795,73.63335917667649,73.6333591767765,73.63945634423858,73.63945634433858,73.64554945742977,73.64554945752977,73.65164150966409,73.65164150976409,73.65773022746252,73.65773022756252,73.66381621708616,73.66381621718617,73.67598508924547,73.67598508934547,73.68814918709886,73.68814918719886,73.70030945792927,73.70030945802927,73.70638779350692,73.70638779360692,73.72462010995639,73.72462011005639,73.73069412592393,73.73069412602393,73.74891405191282,73.74891405201282,73.75498533970556,73.75498533980556,73.76105579388933,73.76105579398933,73.76712549024676,73.76712549034676,73.7731942772126,73.7731942773126,73.78532870616496,73.78532870626496,73.79139249147698,73.79139249157699,73.79745104778729,73.7974510478873,73.80350774742307,73.80350774752307,73.80956149153614,73.80956149163615,73.81561379577919,73.81561379587919,73.82771704017787,73.82771704027788,73.83376729828981,73.83376729838982,73.8398156618359,73.8398156619359,73.84586186557695,73.84586186567695,73.851901968816,73.851901968916,73.8579410868808,73.8579410869808,73.87001811048829,73.87001811058829,73.88209081448566,73.88209081458567,73.88812333946136,73.88812333956136,73.8941530225883,73.8941530226883,73.90017906814882,73.90017906824882,73.90619673972834,73.90619673982835,73.9122138050468,73.9122138051468,73.91822950627785,73.91822950637786,73.92424456335652,73.92424456345653,73.93627221457827,73.93627221467827,73.94228261102498,73.94228261112498,73.95429756865562,73.95429756875562,73.96030194038296,73.96030194048296,73.96630460700105,73.96630460710105,73.9783083109106,73.97830831101061,73.9843091966368,73.9843091967368,73.99030614166607,73.99030614176607,74.002298288724,74.00229828882401,74.00829311183954,74.00829311193954,74.01428482786707,74.01428482796707,74.02027483878537,74.02027483888537,74.0322506546858,74.0322506547858,74.03823471666735,74.03823471676735,74.04421673251781,74.04421673261781,74.05617977904448,74.05617977914449,74.06814058998347,74.06814059008347,74.08009852118239,74.0800985212824,74.08607076107312,74.08607076117312,74.09203928761481,74.09203928771481,74.09800648796042,74.09800648806042,74.10396804249983,74.10396804259983,74.11588819605593,74.11588819615594,74.12184611302892,74.12184611312892,74.12779948304392,74.12779948314392,74.14565788797962,74.14565788807963,74.1575589822034,74.1575589823034,74.16350598647718,74.16350598657718,74.16945230870725,74.16945230880725,74.17539571330593,74.17539571340593,74.18133604870795,74.18133604880795,74.19321565855512,74.19321565865512,74.19914784732404,74.19914784742404,74.20507927826662,74.20507927836663,74.21100972403498,74.21100972413498,74.21693717638931,74.21693717648931,74.22286307519964,74.22286307529964,74.2406395212172,74.2406395213172,74.2524873023583,74.2524873024583,74.25840891944985,74.25840891954985,74.26432890721478,74.26432890731478,74.27024503006538,74.27024503016538,74.2761575153496,74.2761575154496,74.28206920491616,74.28206920501616,74.2879758928289,74.2879758929289,74.29388189869793,74.29388189879793,74.29978422910925,74.29978422920925,74.30568360399785,74.30568360409785,74.31157892451557,74.31157892461557,74.3233679741157,74.3233679742157,74.32925954339305,74.32925954349305,74.33514918021325,74.33514918031325,74.34103601307602,74.34103601317602,74.35869507179426,74.35869507189426,74.3645791385909,74.3645791386909,74.37045941625585,74.37045941635586,74.37633882242054,74.37633882252054,74.38809388350954,74.38809388360954,74.4057174759553,74.4057174760553,74.41158945542189,74.41158945552189,74.41745878249631,74.41745878259631,74.42332682126596,74.42332682136596,74.42919277601318,74.42919277611318,74.43505691197721,74.43505691207721,74.44091862289696,74.44091862299696,74.44677908340326,74.44677908350326,74.452638520844,74.452638520944,74.45849685943656,74.45849685953657,74.47021145259926,74.47021145269926,74.47606694934306,74.47606694944307,74.48192142302132,74.48192142312132,74.48777411580768,74.48777411590768,74.493625368724,74.493625368824,74.49947510598766,74.49947510608766,74.50532423699025,74.50532423709025,74.51117094294857,74.51117094304857,74.51701507229734,74.51701507239734,74.528701550103,74.528701550203,74.55207143651766,74.55207143661767,74.5579118525174,74.55791185261741,74.5754317364292,74.5754317365292,74.58126946214544,74.58126946224544,74.58710214831656,74.58710214841656,74.59293040120362,74.59293040130362,74.5987558122419,74.5987558123419,74.6104052702311,74.6104052703311,74.62787657747482,74.62787657757482,74.63369732788115,74.63369732798115,74.63951641106955,74.63951641116955,74.64533143988706,74.64533143998706,74.65114396787766,74.65114396797766,74.6627661820101,74.6627661821101,74.66857385991216,74.66857386001216,74.67437987059627,74.67437987069627,74.68018447930167,74.68018447940167,74.6859882543981,74.6859882544981,74.69178930131972,74.69178930141972,74.69758455086988,74.69758455096988,74.7091739132307,74.7091739133307,74.71496764712818,74.71496764722818,74.72654886285589,74.7265488629559,74.7323351700553,74.7323351701553,74.73811939323228,74.73811939333228,74.76125336830881,74.76125336840882,74.76703478752836,74.76703478762836,74.7785959208582,74.7785959209582,74.78437472557688,74.78437472567688,74.7901513704905,74.79015137059051,74.79592487042484,74.79592487052484,74.801697271511,74.801697271611,74.80746899055345,74.80746899065345,74.81900973835486,74.81900973845487,74.82477740302643,74.82477740312643,74.83054423408902,74.83054423418902,74.83630602560648,74.83630602570648,74.84206558153625,74.84206558163625,74.85358260937338,74.85358260947338,74.85933716364933,74.85933716374933,74.86509035383789,74.86509035393789,74.8708405127211,74.8708405128211,74.88234007266121,74.88234007276121,74.89383561612173,74.89383561622174,74.8995811522643,74.8995811523643,74.90532077736147,74.90532077746147,74.91105896258858,74.91105896268859,74.92827097955173,74.92827097965173,74.94547928316582,74.94547928326583,74.95121402028312,74.95121402038312,74.95694682494326,74.95694682504326,74.96267883388576,74.96267883398576,74.9741418665965,74.9741418666965,74.985599367175,74.985599367275,74.99705042622965,74.99705042632965,75.00277413697377,75.00277413707377,75.0084965215218,75.00849652162181,75.01421731463454,75.01421731473454,75.01993484909401,75.01993484919402,75.02565029953108,75.02565029963108,75.03136461322865,75.03136461332865,75.03707676712116,75.03707676722117,75.0427864959694,75.0427864960694,75.05420129303391,75.05420129313391,75.05990355729274,75.05990355739274,75.07130600178799,75.07130600188799,75.08270518763,75.08270518773,75.08840237445239,75.08840237455239,75.09409842453528,75.09409842463528,75.09979360311789,75.09979360321789,75.11118252041305,75.11118252051305,75.11687580442981,75.11687580452981,75.12826040211488,75.12826040221488,75.13394982121733,75.13394982131733,75.13963313981778,75.13963313991778,75.14531126730785,75.14531126740785,75.1509882580584,75.1509882581584,75.15666449098262,75.15666449108262,75.16801574430893,75.16801574440893,75.17368951429756,75.17368951439757,75.20205514347882,75.20205514357882,75.21906963300769,75.2190696331077,75.22473711303775,75.22473711313775,75.2304027742846,75.2304027743846,75.23606483585635,75.23606483595636,75.24172367666618,75.24172367676618,75.24738050923622,75.24738050933622,75.25868902115724,75.25868902125724,75.26999272088102,75.26999272098102,75.2812928209297,75.28129282102971,75.29258856347697,75.29258856357697,75.29823217197746,75.29823217207746,75.32080274106512,75.32080274116512,75.33207912114953,75.33207912124954,75.3377138630819,75.3377138631819,75.34334671044843,75.34334671054843,75.34897705698806,75.34897705708806,75.3602336199138,75.3602336200138,75.36585979840858,75.36585979850858,75.37148502962046,75.37148502972046,75.38273378693493,75.38273378703494,75.38835617629805,75.38835617639805,75.39397727735638,75.39397727745639,75.39959705221864,75.39959705231864,75.41083565466023,75.41083565476023,75.41645402754376,75.41645402764377,75.42206709564296,75.42206709574296,75.42767512419701,75.42767512429701,75.43328015933704,75.43328015943705,75.43888451243339,75.43888451253339,75.44448609946359,75.44448609956359,75.45568408241363,75.45568408251363,75.46687490390478,75.46687490400478,75.47246751069292,75.47246751079292,75.47805935965472,75.47805935975472,75.48923949579455,75.48923949589455,75.49482615364596,75.49482615374596,75.50599651382608,75.50599651392608,75.51157885206737,75.51157885216737,75.52274205078861,75.52274205088861,75.52832230500749,75.5283223051075,75.5339019529653,75.5339019530653,75.5506397222079,75.5506397223079,75.5562143685119,75.5562143686119,75.56178757494587,75.56178757504587,75.56735862157477,75.56735862167477,75.57292826622495,75.57292826632495,75.57849605420061,75.57849605430062,75.5840589163051,75.5840589164051,75.59518255649165,75.59518255659165,75.60074189470367,75.60074189480368,75.61185966173612,75.61185966183612,75.61741672646914,75.61741672656915,75.62296996417918,75.62296996427918,75.62852255773682,75.62852255783682,75.63407170318463,75.63407170328463,75.6396198255669,75.6396198256669,75.64516658386177,75.64516658396177,75.65625888792935,75.65625888802936,75.66180409268023,75.66180409278023,75.6728923044856,75.6728923045856,75.67843504630089,75.67843504640089,75.68951605875608,75.68951605885609,75.69505398837414,75.69505398847414,75.70059074336139,75.70059074346139,75.70612681630492,75.70612681640492,75.71719407421213,75.71719407431213,75.72272215208784,75.72272215218784,75.72824935846326,75.72824935856326,75.73930043677821,75.73930043687821,75.74482343721746,75.74482343731746,75.75034439152562,75.75034439162562,75.75586458800743,75.75586458810743,75.76138383720632,75.76138383730633,75.7724204789296,75.7724204790296,75.7779334002786,75.7779334003786,75.78344147153905,75.78344147163905,75.78894579155914,75.78894579165915,75.79995352220773,75.79995352230773,75.81646197320133,75.81646197330133,75.82196148102419,75.82196148112419,75.8274586016941,75.8274586017941,75.8329548129724,75.8329548130724,75.83844966016332,75.83844966026332,75.84943825569694,75.84943825579694,75.86042097807646,75.86042097817646,75.86590627665555,75.86590627675555,75.88235989891379,75.88235989901379,75.8878423177528,75.8878423178528,75.8988039725602,75.8988039726602,75.90975975421352,75.90975975431353,75.91523527683287,75.91523527693288,75.93713520750525,75.93713520760525,75.94260671364503,75.94260671374504,75.95354839972852,75.95354839982852,75.96995670397192,75.96995670407192,75.98089088757467,75.98089088767468,75.9972861193138,75.9972861194138,76.00821052695683,76.00821052705683,76.01913235799033,76.01913235809033,76.02459132210426,76.02459132220426,76.03004316265064,76.03004316275064,76.04094309250306,76.04094309260306,76.0463882641777,76.0463882642777,76.05727712976561,76.05727712986561,76.06271896700437,76.06271896710437,76.07360108793792,76.07360108803792,76.07903898447972,76.07903898457973,76.08447544115151,76.08447544125151,76.08990591099524,76.08990591109524,76.09533505464289,76.09533505474289,76.10619171261156,76.10619171271156,76.12247406611806,76.12247406621806,76.13875399458028,76.13875399468029,76.14417783344359,76.1441778335436,76.15502354082172,76.15502354092172,76.160444386271,76.160444386371,76.1658598511533,76.1658598512533,76.17127224683894,76.17127224693894,76.17668240693689,76.17668240703689,76.1875015903933,76.1875015904933,76.19291012116463,76.19291012126463,76.19831664369617,76.19831664379618,76.20372255996664,76.20372256006664,76.20912605119284,76.20912605129284,76.21452488178709,76.21452488188709,76.21992280298973,76.21992280308973,76.22531882962653,76.22531882972653,76.23071322693671,76.23071322703672,76.23610644961607,76.23610644971608,76.24689206136583,76.24689206146583,76.25227978980426,76.25227978990426,76.25766611626396,76.25766611636396,76.26305047237521,76.26305047247521,76.26843217609428,76.26843217619428,76.27919274168366,76.27919274178366,76.28457016368392,76.28457016378393,76.30070004253179,76.30070004263179,76.30607344805249,76.30607344815249,76.3168175688104,76.3168175689104,76.33292361402002,76.33292361412002,76.33828663731992,76.33828663741993,76.34364659142317,76.34364659152317,76.34900487830848,76.34900487840848,76.35436074014952,76.35436074024952,76.35971482109868,76.35971482119868,76.365067727417,76.365067727517,76.375772024401,76.375772024501,76.38647302484043,76.38647302494043,76.39717038771346,76.39717038781346,76.40251745876901,76.40251745886901,76.4078639235635,76.4078639236635,76.41855564063032,76.41855564073032,76.42389888466288,76.42389888476288,76.42924053726013,76.42924053736013,76.43457745344278,76.43457745354279,76.43991281608146,76.43991281618146,76.450580813184,76.450580813284,76.45591287927812,76.45591287937812,76.46124388441537,76.46124388451537,76.46657291920414,76.46657291930414,76.47190096881867,76.47190096891867,76.48255543872114,76.48255543882114,76.48788163166115,76.48788163176116,76.49320464173056,76.49320464183056,76.49852678029968,76.49852678039969,76.50384675906375,76.50384675916375,76.50916499482724,76.50916499492725,76.51448205595993,76.51448205605993,76.51979828348364,76.51979828358364,76.53042812403021,76.53042812413021,76.53573919833484,76.53573919843484,76.54636017231329,76.54636017241329,76.55166980674787,76.55166980684787,76.56228755996439,76.56228756006439,76.56759321581073,76.56759321591073,76.578199336393,76.578199336493,76.58349862649813,76.58349862659813,76.59939308659497,76.59939308669497,76.60998473269424,76.60998473279425,76.62586922737479,76.6258692274748,76.63116146969499,76.63116146979499,76.63645253738437,76.63645253748437,76.64703334656706,76.64703334666706,76.65232126927714,76.65232126937714,76.66289434863121,76.66289434873121,76.67346257789673,76.67346257799673,76.678743225474,76.678743225574,76.69458406935763,76.69458406945763,76.71042191982725,76.71042191992726,76.72097726604507,76.72097726614507,76.73153109661021,76.73153109671021,76.73680643940315,76.73680643950316,76.74735322218338,76.74735322228338,76.75789439704873,76.75789439714873,76.76316299518727,76.76316299528727,76.77369905472487,76.77369905482487,76.77896507625387,76.77896507635387,76.78422370897611,76.78422370907612,76.794738511485,76.794738511585,76.80524816077481,76.80524816087481,76.81050050353846,76.81050050363847,76.81574905717045,76.81574905727045,76.82099306384441,76.82099306394441,76.83147963732232,76.83147963742232,76.83672084003885,76.83672084013885,76.8419604892114,76.8419604893114,76.84719926688365,76.84719926698365,76.86291389479118,76.86291389489118,76.86814892122307,76.86814892132307,76.88385195438767,76.88385195448767,76.88908326747053,76.88908326757053,76.89431310279203,76.89431310289203,76.89954229396115,76.89954229406115,76.90477057573865,76.90477057583865,76.90999764499404,76.90999764509404,76.9152239943144,76.9152239944144,76.92044822172102,76.92044822182102,76.92567165340999,76.92567165351,76.93089148542387,76.93089148552387,76.93610946076323,76.93610946086324,76.94654382000664,76.94654382010664,76.95175929451909,76.9517592946191,76.95697317759624,76.95697317769624,76.96218626495573,76.96218626505573,76.97261092402206,76.97261092412207,76.98303293069621,76.98303293079621,76.9986634019892,76.9986634020892,77.00387186660807,77.00387186670807,77.00907911870478,77.00907911880478,77.0142857266491,77.01428572674911,77.01948990954916,77.01948990964917,77.02469325884026,77.02469325894026,77.03509836598714,77.03509836608714,77.0402978124726,77.0402978125726,77.04549616010988,77.04549616020988,77.0610896115864,77.0610896116864,77.07148282085981,77.07148282095982,77.07667487853851,77.07667487863851,77.0818662162822,77.0818662163822,77.08705615204715,77.08705615214716,77.09743352275015,77.09743352285015,77.10261910101369,77.10261910111369,77.10780206477637,77.10780206487637,77.11298347499505,77.11298347509505,77.1181636726916,77.1181636727916,77.12852334814968,77.12852334824969,77.1337009692367,77.1337009693367,77.14404632177704,77.14404632187704,77.1492120828819,77.1492120829819,77.15437522948592,77.15437522958592,77.1595353826759,77.1595353827759,77.16985261985921,77.16985261995922,77.18016493117136,77.18016493127136,77.18531879440276,77.18531879450276,77.19046883061117,77.19046883071117,77.19561822266719,77.1956182227672,77.2007668947882,77.2007668948882,77.21106207922517,77.21106207932517,77.2162024531478,77.2162024532478,77.22134025046091,77.22134025056091,77.22647573640369,77.22647573650369,77.23160989615037,77.23160989625038,77.24187647264318,77.24187647274319,77.24700786632376,77.24700786642376,77.25726682666189,77.25726682676189,77.26239382494971,77.26239382504971,77.26751934547619,77.26751934557619,77.2777683025067,77.2777683026067,77.29313675534432,77.29313675544432,77.29825352297662,77.29825352307662,77.30848603517568,77.30848603527568,77.31359818006733,77.31359818016733,77.32382076474138,77.32382076484139,77.32893082561061,77.32893082571061,77.33403800673977,77.33403800683978,77.34424956504066,77.34424956514066,77.34935049410254,77.34935049420254,77.35445002118571,77.35445002128571,77.35954818418149,77.3595481842815,77.36464422526352,77.36464422536352,77.36973912960605,77.36973912970605,77.3748329351004,77.3748329352004,77.3799243913331,77.37992439143311,77.38501376354338,77.38501376364339,77.39010154431836,77.39010154441836,77.40027244523635,77.40027244533636,77.41043735932631,77.41043735942631,77.41551778918584,77.41551778928584,77.43075737365517,77.43075737375517,77.4358359847315,77.4358359848315,77.44598964510037,77.44598964520037,77.45613580298851,77.45613580308851,77.46120585062725,77.46120585072725,77.47134397555625,77.47134397565625,77.47640962780225,77.47640962790226,77.48653990922868,77.48653990932868,77.49160412160464,77.49160412170464,77.49666662887134,77.49666662897134,77.51185259712746,77.51185259722746,77.51691237621935,77.51691237631935,77.52703075977233,77.52703075987233,77.53208682551518,77.53208682561518,77.53714160295327,77.53714160305327,77.54219475106473,77.54219475116473,77.54724729291512,77.54724729301512,77.55735040626745,77.55735040636745,77.56745014729256,77.56745014739256,77.57249507298829,77.57249507308829,77.57753908929242,77.57753908939242,77.58258090790017,77.58258090800017,77.58762166555105,77.58762166565106,77.59266177904955,77.59266177914955,77.60274105876363,77.60274105886363,77.60777916402233,77.60777916412233,77.6128162462155,77.6128162463155,77.61785234323442,77.61785234333442,77.6228865456875,77.62288654578751,77.63295404120207,77.63295404130207,77.64301884643315,77.64301884653315,77.64804944921114,77.64804944931115,77.65307921838016,77.65307921848016,77.65810762346179,77.65810762356179,77.66816250116787,77.66816250126787,77.67821052054562,77.67821052064562,77.6832300211678,77.6832300212678,77.6932656879763,77.6932656880763,77.70329760354443,77.70329760364443,77.70831202673016,77.70831202683016,77.71332527528507,77.71332527538507,77.7183369324047,77.7183369325047,77.72334714965427,77.72334714975428,77.72835369144612,77.72835369154612,77.73335742928053,77.73335742938053,77.7383601819407,77.73836018204071,77.74335944859973,77.74335944869974,77.74835776797585,77.74835776807585,77.75335373809033,77.75335373819033,77.76833929917211,77.76833929927211,77.78332110901354,77.78332110911354,77.79330441002227,77.79330441012227,77.79829299132999,77.79829299142999,77.8032797538545,77.8032797539545,77.81325047494606,77.81325047504606,77.81823102329462,77.81823102339462,77.82320903292495,77.82320903302495,77.82818541322865,77.82818541332865,77.8381370749879,77.8381370750879,77.84808676639867,77.84808676649867,77.85305985015782,77.85305985025782,77.86300488093663,77.86300488103663,77.86797625958653,77.86797625968653,77.87294695619272,77.87294695629272,77.87791060501401,77.87791060511401,77.89776205531984,77.89776205541985,77.90768385872147,77.90768385882147,77.9126430742587,77.9126430743587,77.92256021702836,77.92256021712836,77.92751598445577,77.92751598455577,77.9374267235929,77.93742672369291,77.94238101325895,77.94238101335895,77.94733295366335,77.94733295376335,77.95228148384925,77.95228148394925,77.95722800579536,77.95722800589536,77.9621721405885,77.96217214068851,77.96711426714187,77.96711426724187,77.97205230143302,77.97205230153303,77.98192609653633,77.98192609663633,77.9868601143479,77.98686011444791,77.9967262175139,77.9967262176139,78.00165788606384,78.00165788616384,78.00658875889613,78.00658875899613,78.01151887390208,78.01151887400208,78.01644834475564,78.01644834485565,78.02137690621761,78.02137690631761,78.02630455828798,78.02630455838798,78.03123012633591,78.03123012643591,78.03615379981801,78.03615379991801,78.04599868384663,78.04599868394664,78.05583928615646,78.05583928625646,78.06075738961499,78.060757389715,78.0656729543553,78.06567295445531,78.07058742024743,78.07058742034744,78.0755012798785,78.0755012799785,78.08041400277006,78.08041400287006,78.0853248689871,78.0853248690871,78.09023300702933,78.09023300712933,78.09513781063569,78.0951378107357,78.10494294667302,78.10494294677302,78.1147402770991,78.1147402771991,78.1196358731155,78.1196358732155,78.12942592840876,78.12942592850877,78.1392127629401,78.1392127630401,78.14409937871442,78.14409937881442,78.14898451672738,78.14898451682738,78.15875354233985,78.15875354243985,78.16363257985081,78.16363257995081,78.17338754778476,78.17338754788476,78.17826287194667,78.17826287204667,78.19288516897471,78.19288516907471,78.20262657181726,78.20262657191726,78.20749511343348,78.20749511353348,78.21723098414378,78.21723098424378,78.22696488450562,78.22696488460562,78.23183020535991,78.23183020545991,78.24642234089981,78.24642234099981,78.25128091709972,78.25128091719972,78.25613703036404,78.25613703046405,78.27070267987354,78.27070267997354,78.28040937426994,78.28040937436994,78.29011337838284,78.29011337848284,78.2998140859512,78.2998140860512,78.30466265884348,78.30466265894349,78.30950918560467,78.30950918570467,78.31920110238754,78.31920110248754,78.32888949527795,78.32888949537795,78.33373177821166,78.33373177831166,78.33857095405739,78.33857095415739,78.34824813111803,78.34824813121803,78.3530842377671,78.35308423786711,78.36275512486918,78.36275512496918,78.3675893369524,78.36758933705241,78.3724202524911,78.3724202525911,78.38691064984549,78.38691064994549,78.39173830673093,78.39173830683093,78.40621930703878,78.40621930713878,78.415869278134,78.415869278234,78.42069216071353,78.42069216081353,78.43033614498069,78.43033614508069,78.43515686775517,78.43515686785517,78.4496166110343,78.4496166111343,78.45443479509056,78.45443479519056,78.45925157716809,78.45925157726809,78.46406441854867,78.46406441864868,78.46887407705866,78.46887407715866,78.47368286406835,78.47368286416835,78.4784893018164,78.4784893019164,78.48329456493363,78.48329456503363,78.4880974030066,78.48809740310661,78.49289720977423,78.49289720987423,78.49769606925892,78.49769606935892,78.5024934888736,78.5024934889736,78.50728958229217,78.50728958239218,78.51688082184641,78.51688082194642,78.52646891642136,78.52646891652137,78.53126004607745,78.53126004617745,78.53605030423324,78.53605030433324,78.55041683487318,78.55041683497318,78.55998946979088,78.55998946989088,78.56956043749065,78.56956043759065,78.57434342051364,78.57434342061364,78.57912572149291,78.57912572159292,78.58390180829625,78.58390180839625,78.58867664468613,78.58867664478613,78.59345045801045,78.59345045811045,78.59822188418183,78.59822188428183,78.60776287985004,78.60776287995004,78.61253142628135,78.61253142638135,78.61729887386446,78.61729887396446,78.62206461633832,78.62206461643832,78.62682687281104,78.62682687291104,78.63158791676162,78.63158791686162,78.63634778608137,78.63634778618137,78.64110371470419,78.64110371480419,78.64585896128331,78.64585896138331,78.6553650969401,78.6553650970401,78.66011484927829,78.66011484937829,78.66960992067061,78.66960992077061,78.67435508815946,78.67435508825946,78.68858793823387,78.68858793833387,78.69333018809132,78.69333018819133,78.70281249010989,78.7028124902099,78.71228925999621,78.71228926009621,78.71702378002506,78.71702378012506,78.72175723909704,78.72175723919705,78.73122328574071,78.73122328584071,78.74068747570986,78.74068747580986,78.75488057779299,78.75488057789299,78.76434029658677,78.76434029668677,78.76906708678699,78.76906708688699,78.77851922731742,78.77851922741742,78.78324192525544,78.78324192535544,78.79740801082971,78.79740801092971,78.80212798059293,78.80212798069293,78.81156519194455,78.81156519204455,78.81628122101083,78.81628122111083,78.82099565864179,78.8209956587418,78.83042343505554,78.83042343515554,78.83513510662038,78.83513510672039,78.83984473205412,78.83984473215412,78.84926326298658,78.84926326308658,78.85867304102487,78.85867304112487,78.86337217056386,78.86337217066387,78.87276944446764,78.87276944456764,78.87746652787553,78.87746652797553,78.88216148936968,78.88216148946968,78.8915494041182,78.8915494042182,78.90562758183759,78.9056275819376,78.91970090946845,78.91970090956845,78.92438882317768,78.92438882327768,78.92907381925554,78.92907381935554,78.93375801961572,78.93375801971573,78.94312100187781,78.94312100197781,78.94779864704022,78.94779864714022,78.96650729523265,78.96650729533265,78.97585963003473,78.97585963013474,78.98053341028282,78.98053341038282,78.98520586433482,78.98520586443482,78.98987661327756,78.98987661337756,78.9945444066976,78.9945444067976,78.99921155596526,78.99921155606526,79.00387631807996,79.00387631817996,79.01320447822195,79.01320447832195,79.01786696685764,79.01786696695764,79.02719027691109,79.02719027701109,79.03184988580671,79.03184988590671,79.03650703176675,79.03650703186675,79.04116133587802,79.04116133597802,79.04581495794558,79.04581495804558,79.05046664755601,79.05046664765601,79.05511356286051,79.05511356296051,79.05975911407761,79.05975911417761,79.06440254338098,79.06440254348098,79.06904510118406,79.06904510128406,79.07368591598656,79.07368591608656,79.07832593507142,79.07832593517142,79.08296523422126,79.08296523432126,79.09224095278086,79.09224095288086,79.09687684171219,79.09687684181219,79.10614657344374,79.10614657354374,79.11077996154816,79.11077996164816,79.11541228869572,79.11541228879572,79.1246748210771,79.1246748211771,79.1293041169193,79.1293041170193,79.13393102560856,79.13393102570856,79.1385563049712,79.1385563050712,79.14780371871719,79.14780371881719,79.15242520894815,79.15242520904815,79.16166640851817,79.16166640861817,79.17090408419575,79.17090408429576,79.1755210274687,79.1755210275687,79.18475358781852,79.18475358791852,79.18936909122144,79.18936909132144,79.19398031290555,79.19398031300555,79.19858895798014,79.19858895808014,79.20780526295506,79.20780526305506,79.21240943685426,79.21240943695426,79.21701292870976,79.21701292880977,79.2216149049126,79.2216149050126,79.23081745533953,79.23081745543954,79.23541738541127,79.23541738551127,79.24461425214069,79.2446142522407,79.24921058253733,79.24921058263733,79.25380535938997,79.25380535948997,79.25839812800284,79.25839812810284,79.2629884336801,79.26298843378011,79.26757688268286,79.26757688278286,79.27216446018532,79.27216446028532,79.2767511661875,79.2767511662875,79.28592048592964,79.28592048602964,79.29049908319004,79.29049908329004,79.2950766952762,79.2950766953762,79.299652829601,79.299652829701,79.30880362048924,79.30880362058925,79.31337645826949,79.31337645836949,79.3179480835276,79.3179480836276,79.33623219740709,79.33623219750709,79.3408009050338,79.3408009051338,79.349937335113,79.349937335213,79.35907099912608,79.35907099922608,79.38189547792734,79.38189547802735,79.38645141618031,79.38645141628031,79.3910056114327,79.3910056115327,79.40011301676327,79.40011301686327,79.40921864120195,79.40921864130195,79.4228752022398,79.4228752023398,79.43652619325407,79.43652619335407,79.44107330283023,79.44107330293023,79.45016642313438,79.45016642323438,79.4547105014052,79.4547105015052,79.46834156158685,79.46834156168686,79.47742687627975,79.47742687637975,79.48196822637577,79.48196822647577,79.491048198393,79.491048198493,79.5001242676046,79.5001242677046,79.50466038869892,79.50466038879892,79.50919560040163,79.50919560050163,79.513728500734,79.513728500834,79.52732523138268,79.52732523148268,79.53638693978523,79.53638693988523,79.54544675362196,79.54544675372196,79.55450448343625,79.55450448353625,79.56355823466228,79.56355823476228,79.57260759049558,79.57260759059558,79.58165505176304,79.58165505186304,79.5861777214399,79.5861777215399,79.59522120411908,79.59522120421909,79.60878243060395,79.60878243070395,79.61330157638835,79.61330157648835,79.61781954754194,79.61781954764194,79.6268543531096,79.6268543532096,79.63588741567668,79.63588741577668,79.64943348563486,79.64943348573486,79.65394690983042,79.65394690993043,79.65845972776492,79.65845972786492,79.66297110582939,79.66297110592939,79.67650323178299,79.67650323188299,79.68552121360601,79.68552121370601,79.69002849940827,79.69002849950827,79.69904170692537,79.69904170702537,79.70354664346598,79.70354664356599,79.71255409150294,79.71255409160294,79.72606007244798,79.72606007254798,79.7305584537908,79.73055845389081,79.73505471321988,79.73505471331988,79.73955021482263,79.73955021492263,79.74404492070771,79.74404492080771,79.7530322863468,79.7530322864468,79.77549444837722,79.77549444847722,79.7799843799564,79.7799843800564,79.78447340214397,79.78447340224398,79.78896174228785,79.78896174238785,79.79344667221322,79.79344667231322,79.79792970757275,79.79792970767275,79.80241168197541,79.80241168207542,79.80689217861672,79.80689217871672,79.81137078069219,79.81137078079219,79.81584752609314,79.81584752619314,79.82479980437289,79.82479980447289,79.83374977128233,79.83374977138233,79.83822367483452,79.83822367493453,79.84716716232879,79.84716716242879,79.85163663259692,79.85163663269692,79.86057371645866,79.86057371655866,79.86503860187746,79.86503860197746,79.86950250212203,79.86950250222203,79.8739657203229,79.8739657204229,79.87842757443636,79.87842757453636,79.88734700094449,79.88734700104449,79.9007190739761,79.90071907407611,79.905175358066,79.905175358166,79.91408546331022,79.91408546341022,79.92744582792496,79.92744582802496,79.9318974513829,79.9318974514829,79.94079922053741,79.94079922063742,79.94524887364689,79.94524887374689,79.9541472325829,79.9541472326829,79.96304426532282,79.96304426542282,79.97193678899606,79.97193678909606,79.97637962166851,79.97637962176852,79.98082150705804,79.98082150715804,79.98526145999041,79.98526146009041,80.0030187330017,80.0030187331017,80.00745599565059,80.00745599575059,80.01189250047314,80.01189250057314,80.02076422181347,80.02076422191347,80.02963401069665,80.02963401079666,80.03406790101835,80.03406790111835,80.03849963153499,80.03849963163499,80.0429285959855,80.0429285960855,80.05178311466803,80.05178311476803,80.05620703957341,80.05620703967341,80.0650535631881,80.0650535632881,80.06947464624473,80.06947464634473,80.07389072764755,80.07389072774755,80.07830563441955,80.07830563451955,80.08271917710415,80.08271917720415,80.09595810004869,80.09595810014869,80.10036929347164,80.10036929357165,80.11801001279258,80.11801001289258,80.12682319204852,80.12682319214852,80.14003812978954,80.14003812988955,80.14884592847851,80.14884592857851,80.15324711859384,80.15324711869384,80.15764702040441,80.15764702050441,80.16644462632917,80.16644462642917,80.17084191363887,80.17084191373887,80.17523844312224,80.17523844322224,80.1840288118055,80.1840288119055,80.19281508822654,80.19281508832654,80.20159787864645,80.20159787874645,80.21037657680414,80.21037657690414,80.21476363345832,80.21476363355832,80.2235364205706,80.2235364206706,80.22792165844159,80.22792165854159,80.24107540170604,80.24107540180604,80.245453099205,80.245453099305,80.2498278032899,80.2498278033899,80.25857603682893,80.25857603692893,80.26732271682397,80.26732271692397,80.27169514742988,80.27169514752988,80.27606591081785,80.27606591091785,80.28917683689438,80.28917683699439,80.29354574360784,80.29354574370784,80.29791374092969,80.29791374102969,80.30228063940336,80.30228063950337,80.3110130722633,80.3110130723633,80.31537807617113,80.31537807627113,80.31974239803525,80.31974239813525,80.3241054315946,80.3241054316946,80.32846622956627,80.32846622966628,80.3371852867914,80.3371852868914,80.3459008959067,80.3459008960067,80.35461540617379,80.3546154062738,80.36332798398375,80.36332798408375,80.37203912192366,80.37203912202366,80.38074696331901,80.38074696341901,80.38509927363573,80.38509927373573,80.39380226494255,80.39380226504255,80.39815230178026,80.39815230188026,80.40250123976978,80.40250123986978,80.40684805584557,80.40684805594557,80.41988653372445,80.41988653382445,80.42423069740806,80.42423069750807,80.42857391380876,80.42857391390876,80.43291440203465,80.43291440213466,80.4372475393451,80.4372475394451,80.44157999461183,80.44157999471183,80.45024323792735,80.45024323802735,80.45457338182378,80.45457338192378,80.4589017448283,80.4589017449283,80.46755714464126,80.46755714474126,80.47620985417073,80.47620985427073,80.48053412491305,80.48053412501305,80.48918080972317,80.48918080982317,80.4978255241848,80.4978255242848,80.50646845775455,80.50646845785455,80.51510915573661,80.51510915583661,80.51942694706376,80.51942694716377,80.52374394267326,80.52374394277327,80.52805964997799,80.52805965007799,80.53668988995663,80.53668989005664,80.54100358902157,80.54100358912157,80.54531622712965,80.54531622722965,80.54962780428085,80.54962780438085,80.55393850993175,80.55393851003176,80.56255848136355,80.56255848146355,80.56686672407888,80.56686672417888,80.57117311011969,80.57117311021969,80.57547782894255,80.57547782904255,80.57978110789539,80.57978110799539,80.58408150710815,80.58408150720815,80.59698137855034,80.59698137865034,80.60127878434908,80.60127878444908,80.60987052674989,80.60987052684989,80.61416452233011,80.61416452243012,80.61845537293105,80.61845537303105,80.62274561727091,80.62274561737091,80.62703207247911,80.62703207257911,80.6313178835349,80.6313178836349,80.63988859625492,80.63988859635492,80.64845775543095,80.64845775553096,80.65274148246434,80.65274148256434,80.65702414854086,80.65702414864086,80.66130529896469,80.6613052990647,80.66986555368128,80.66986555378128,80.67842326967964,80.67842326977964,80.69125721023067,80.69125721033068,80.6955333211094,80.6955333212094,80.69980768898753,80.69980768908754,80.70408084434355,80.70408084444355,80.7126256394029,80.7126256395029,80.71689671073649,80.7168967108365,80.72543817135995,80.72543817145996,80.73397823000471,80.73397823010471,80.73824638370691,80.73824638380691,80.74251161977772,80.74251161987772,80.74677477182611,80.74677477192611,80.75956233340543,80.75956233350543,80.76808318115259,80.76808318125259,80.77234155889506,80.77234155899507,80.78085732920577,80.78085732930577,80.78936999242852,80.78936999252852,80.79362473260457,80.79362473270457,80.80213243206482,80.80213243216483,80.81063785804606,80.81063785814607,80.81914029061328,80.81914029071328,80.82338881661339,80.82338881671339,80.82763669846112,80.82763669856112,80.83188389826516,80.83188389836516,80.83612772574199,80.83612772584199,80.84037094695776,80.84037094705776,80.85309761719105,80.85309761729106,80.85733757975359,80.85733757985359,80.86581489037779,80.8658148904778,80.8700518595263,80.8700518596263,80.87428742669609,80.87428742679609,80.87852231182217,80.87852231192217,80.88699030118245,80.88699030128245,80.89122302650348,80.89122302660348,80.90391684496514,80.90391684506514,80.91660172107605,80.91660172117605,80.92082827011245,80.92082827021245,80.93772900990842,80.93772901000843,80.94617017221644,80.94617017231644,80.95038895353291,80.95038895363291,80.96304453965595,80.96304453975596,80.97147926044013,80.97147926054014,80.97569504834257,80.97569504844257,80.9799092448097,80.9799092449097,80.99254910603631,80.99254910613631,81.00097143635989,81.00097143645989,81.00518135110823,81.00518135120824,81.00938891659493,81.00938891669493,81.01780264558963,81.01780264568963,81.02200843018444,81.02200843028444,81.03041765011243,81.03041765021243,81.03882266410426,81.03882266420426,81.04722388896441,81.04722388906441,81.05562193095396,81.05562193105396,81.0682134807517,81.0682134808517,81.07240740536436,81.07240740546436,81.08079392839362,81.08079392849362,81.08498550374465,81.08498550384465,81.09336342544499,81.093363425545,81.09754946866379,81.09754946876379,81.10591992577473,81.10591992587473,81.11428485075344,81.11428485085344,81.12264443305646,81.12264443315647,81.12682289801191,81.12682289811191,81.13517781968298,81.13517781978298,81.14353024052716,81.14353024062716,81.16022803443057,81.16022803453058,81.16857075509766,81.16857075519766,81.1727397472239,81.1727397473239,81.17690760261064,81.17690760271064,81.1810745864971,81.1810745865971,81.18524088833985,81.18524088843985,81.18940544718204,81.18940544728204,81.19356894506736,81.19356894516736,81.20189476620716,81.20189476630716,81.20605564959162,81.20605564969162,81.2102138805839,81.2102138806839,81.22268675477754,81.22268675487754,81.22684206813844,81.22684206823844,81.24346188171198,81.24346188181198,81.25592350218456,81.25592350228456,81.26007559478353,81.26007559488353,81.26422461818584,81.26422461828584,81.27251952001117,81.27251952011117,81.28081104950931,81.28081104960931,81.2849555259536,81.28495552605361,81.28909871409314,81.28909871419314,81.29324038658001,81.29324038668001,81.29737955823997,81.29737955833997,81.30151812363884,81.30151812373884,81.30979377667526,81.30979377677527,81.31393056118227,81.31393056128228,81.32220197039122,81.32220197049122,81.32633553413629,81.32633553423629,81.3304673927721,81.33046739287211,81.33872997330425,81.33872997340426,81.34286012683081,81.34286012693082,81.3511194108184,81.3511194109184,81.36350600648379,81.36350600658379,81.3676335455095,81.36763354560951,81.3717592278607,81.3717592279607,81.38000706867061,81.38000706877061,81.38412786304194,81.38412786314194,81.38824729332586,81.38824729342586,81.39236611734871,81.39236611744872,81.40059955945827,81.40059955955827,81.40471285134888,81.40471285144888,81.41293424401971,81.41293424411971,81.42114961197116,81.42114961207116,81.42936263066099,81.42936263076099,81.43346796537507,81.43346796547507,81.44167727071586,81.44167727081586,81.44988471938211,81.44988471948211,81.45398653020375,81.45398653030375,81.46628908292857,81.46628908302857,81.47038786244487,81.47038786254487,81.47448592202613,81.47448592212614,81.47858095030207,81.47858095040208,81.48267522075167,81.48267522085168,81.48676888494019,81.48676888504019,81.49495534181698,81.49495534191698,81.50723265892483,81.50723265902484,81.51132359493856,81.51132359503856,81.5154136973433,81.51541369744331,81.519503193487,81.519503193587,81.52359162867381,81.52359162877381,81.53176668026424,81.53176668036424,81.54402516538768,81.54402516548768,81.55219244925817,81.55219244935817,81.5562739124427,81.55627391254271,81.56443520948515,81.56443520958516,81.568512883538,81.56851288363801,81.57666664020843,81.57666664030843,81.58481713822559,81.58481713832559,81.58889034110308,81.58889034120308,81.59296255880632,81.59296255890632,81.59703295772636,81.59703295782636,81.60110252303743,81.60110252313743,81.60924070637665,81.60924070647665,81.61330826344793,81.61330826354794,81.61737381227944,81.61737381237944,81.6214370118493,81.6214370119493,81.62549748324435,81.62549748334435,81.62955715892174,81.62955715902174,81.63361588731622,81.63361588741623,81.64173216947435,81.64173216957435,81.65390331511267,81.65390331521267,81.66201542922596,81.66201542932596,81.66607040638009,81.66607040648009,81.67012473938182,81.67012473948182,81.67417660944797,81.67417660954797,81.68632532342676,81.68632532352676,81.69441318709733,81.69441318719733,81.70249764054938,81.70249764064938,81.71057887323953,81.71057887333953,81.71461767080139,81.71461767090139,81.71865286868817,81.71865286878817,81.73075652989132,81.73075652999133,81.73478850701618,81.73478850711618,81.73881786964017,81.73881786974017,81.74284446619804,81.74284446629804,81.7468694713206,81.7468694714206,81.75894028075213,81.75894028085213,81.77099839659256,81.77099839669256,81.77902923383141,81.77902923393141,81.78705711554755,81.78705711564756,81.79508177650177,81.79508177660178,81.81112378538607,81.81112378548607,81.81513211832926,81.81513211842926,81.82314700332375,81.82314700342376,81.8271515850266,81.8271515851266,81.8311549920986,81.8311549921986,81.83515692140926,81.83515692150927,81.83915797921964,81.83915797931964,81.84715857918772,81.84715857928772,81.85115755297568,81.85115755307568,81.85515273763195,81.85515273773196,81.85914678554872,81.85914678564872,81.86713347940355,81.86713347950355,81.8791088784995,81.8791088785995,81.88309724271875,81.88309724281875,81.89107056093876,81.89107056103876,81.89505471922186,81.89505471932186,81.90700427643978,81.90700427653978,81.91098400143882,81.91098400153882,81.91496293072021,81.91496293082021,81.92291919784768,81.92291919794768,81.92689528528031,81.92689528538031,81.93484586871027,81.93484586881027,81.93881979633784,81.93881979643784,81.95073991200263,81.95073991210263,81.95471205873832,81.95471205883833,81.96265532914414,81.96265532924414,81.97456479734892,81.97456479744892,81.99043182413244,81.99043182423245,81.99439476327817,81.99439476337817,81.99835667935832,81.99835667945833,82.00231685243791,82.00231685253792,82.00627600245195,82.00627600255196,82.01023303055226,82.01023303065226,82.01418918715227,82.01418918725227,82.02210024993886,82.02210025003886,82.03000900135511,82.03000900145511,82.0379153277271,82.03791532782711,82.04186572484697,82.04186572494697,82.04581388637915,82.04581388647915,82.05765617327933,82.05765617337933,82.06554768414645,82.06554768424645,82.06949205654696,82.06949205664696,82.07737833841237,82.07737833851237,82.08526132373323,82.08526132383324,82.08920167965417,82.08920167975417,82.09314127774877,82.09314127784877,82.09708000434307,82.09708000444307,82.10101691215418,82.10101691225418,82.10495260744314,82.10495260754314,82.10888682497075,82.10888682507075,82.1167534791341,82.1167534792341,82.12461774614448,82.12461774624448,82.12854840188832,82.12854840198833,82.13247841347977,82.13247841357978,82.13640774302753,82.13640774312753,82.14426515170958,82.14426515180958,82.15604825237297,82.15604825247297,82.15997402013696,82.15997402023696,82.1638986132701,82.16389861337011,82.1678223727943,82.1678223728943,82.17959190836629,82.17959190846629,82.19135757902399,82.19135757912399,82.19527807989493,82.19527807999494,82.19919729246111,82.19919729256111,82.20311559563568,82.20311559573568,82.2070326483968,82.2070326484968,82.21094678352654,82.21094678362654,82.21485932722096,82.21485932732097,82.21877039315405,82.21877039325405,82.2226805118042,82.2226805119042,82.2265896073888,82.2265896074888,82.23831537848994,82.23831537858995,82.24613042467986,82.24613042477986,82.25003664052439,82.25003664062439,82.25784687451709,82.25784687461709,82.2617501348389,82.26175013493891,82.27345707395561,82.27345707405561,82.27735703773288,82.27735703783289,82.29685348429201,82.29685348439202,82.30464704610533,82.30464704620533,82.31633507745498,82.31633507755498,82.32412560796296,82.32412560806296,82.32801958491217,82.32801958501217,82.33580515165765,82.33580515175765,82.33969416484437,82.33969416494438,82.35135984031714,82.35135984041715,82.3552464663509,82.35524646645091,82.36301820276577,82.36301820286577,82.3669031615816,82.3669031616816,82.37855686339827,82.37855686349828,82.38243977608299,82.38243977618299,82.38632166570217,82.38632166580217,82.39408411874442,82.39408411884442,82.40184429830768,82.40184429840768,82.40572145151225,82.40572145161225,82.40959276945405,82.40959276955405,82.4173341928155,82.4173341929155,82.42120407088726,82.42120407098726,82.42507273643689,82.42507273653689,82.43667752056362,82.43667752066362,82.44054489780848,82.44054489790848,82.44441163090094,82.44441163100095,82.44827393070935,82.44827393080935,82.45213448751718,82.45213448761719,82.46371445283145,82.46371445293146,82.4675727740516,82.4675727741516,82.47143011009751,82.47143011019752,82.47528339177254,82.47528339187254,82.48298859103517,82.48298859113517,82.48684016760093,82.48684016770093,82.49454104725346,82.49454104735347,82.49838913781808,82.49838913791808,82.50608350016412,82.50608350026413,82.50992950670633,82.50992950680633,82.51377449018298,82.51377449028298,82.52530739448183,82.52530739458183,82.52915006658816,82.52915006668816,82.53299126093313,82.53299126103313,82.54451321464143,82.54451321474143,82.54835050618078,82.54835050628078,82.55602372517207,82.55602372527207,82.55985866744979,82.55985866754979,82.56369285190115,82.56369285200115,82.56752608906962,82.56752608916962,82.57135811371593,82.57135811381593,82.57901829809425,82.57901829819426,82.58284463904306,82.58284463914306,82.59049588107064,82.59049588117064,82.59432017588834,82.59432017598834,82.59814314450995,82.59814314460995,82.60196501428338,82.60196501438338,82.60960765498206,82.60960765508206,82.61724817376698,82.61724817386698,82.62870578801943,82.62870578811943,82.63634103991133,82.63634104001133,82.64015616503038,82.64015616513038,82.64778463437658,82.64778463447658,82.65159672819028,82.65159672829029,82.65921898336055,82.65921898346055,82.66302516612883,82.66302516622883,82.66683043950552,82.66683043960552,82.67063101435893,82.67063101445893,82.67442980832045,82.67442980842046,82.67822788234696,82.67822788244696,82.68202300085076,82.68202300095076,82.68581675526715,82.68581675536716,82.69340149803381,82.69340149813381,82.69719123597064,82.69719123607064,82.70098036764638,82.70098036774638,82.71234583021648,82.71234583031648,82.71991943293601,82.71991943303601,82.72370386608848,82.72370386618849,82.7274870488275,82.7274870489275,82.73126788230488,82.73126788240488,82.73882829884619,82.73882829894619,82.75016738905983,82.75016738915983,82.75772519110028,82.75772519120028,82.76150303116363,82.76150303126363,82.76527927979168,82.76527927989169,82.76905439168023,82.76905439178023,82.77660256932622,82.77660256942622,82.78037355106125,82.78037355116125,82.78414210775199,82.78414210785199,82.78790994450773,82.78790994460773,82.79544395080126,82.79544395090126,82.79920629331606,82.79920629341606,82.80296780222189,82.80296780232189,82.8067265450616,82.8067265451616,82.81048460585761,82.81048460595761,82.81423781656507,82.81423781666507,82.82174317702312,82.82174317712312,82.8292472112851,82.8292472113851,82.83299754225249,82.83299754235249,82.84049297518555,82.84049297528556,82.84423637204198,82.84423637214198,82.8479788595068,82.8479788596068,82.85546167463139,82.85546167473139,82.8591998424861,82.85919984258611,82.86293649468814,82.86293649478814,82.87040756350453,82.87040756360453,82.8741415633144,82.8741415634144,82.87787427481949,82.87787427491949,82.88160619060695,82.88160619070695,82.88906805183336,82.88906805193336,82.89652646494996,82.89652646504996,82.90025413690992,82.90025413700992,82.90398097526092,82.90398097536092,82.90770663898111,82.90770663908111,82.91515633709484,82.91515633719484,82.91887934842286,82.91887934852286,82.92260137457663,82.92260137467663,82.93004330497044,82.93004330507044,82.9374822040589,82.9374822041589,82.94491617727618,82.94491617737619,82.95234401210016,82.95234401220016,82.9560551255547,82.9560551256547,82.96347280550579,82.96347280560579,82.97088525645515,82.97088525655515,82.97459011784244,82.97459011794244,82.97829376670758,82.97829376680758,82.98199559678952,82.98199559688952,82.98569678271907,82.98569678281908,82.99309654007733,82.99309654017733,82.99679465681022,82.99679465691023,83.00049099265124,83.00049099275124,83.00788222446322,83.00788222456322,83.0115762110426,83.0115762111426,83.01526955346958,83.01526955356958,83.0189622896355,83.0189622897355,83.03003667111027,83.03003667121027,83.0411048763004,83.0411048764004,83.04479173931222,83.04479173941222,83.04847750347585,83.04847750357585,83.055846265737,83.055846265837,83.05952865757344,83.05952865767344,83.06689207715893,83.06689207725893,83.07057230919031,83.07057230929031,83.07425166972142,83.07425166982142,83.0816092161528,83.0816092162528,83.08528690946596,83.08528690956597,83.08896285977856,83.08896285987856,83.09263661239478,83.09263661249479,83.09630892514097,83.09630892524098,83.10365078456722,83.10365078466722,83.10731654211561,83.10731654221561,83.11464699625552,83.11464699635552,83.11831063189017,83.11831063199017,83.12197172880659,83.12197172890659,83.12563085537454,83.12563085547454,83.13294641822696,83.13294641832697,83.1366020966851,83.1366020967851,83.14025675207768,83.14025675217768,83.14756359992724,83.14756360002724,83.15121389781838,83.15121389791838,83.15851365999171,83.15851366009171,83.16216202542572,83.16216202552572,83.1694571648584,83.1694571649584,83.17310299157417,83.17310299167417,83.18039199261355,83.18039199271355,83.18767709084729,83.18767709094729,83.19859914922868,83.19859914932869,83.20587519143773,83.20587519153773,83.21314873281986,83.21314873291986,83.21678410153221,83.21678410163221,83.22405078458601,83.22405078468601,83.23494525808881,83.23494525818882,83.23857335166835,83.23857335176835,83.24945626831953,83.24945626841954,83.25308273257244,83.25308273267244,83.25670851478165,83.25670851488165,83.26033001527206,83.26033001537206,83.26756926501254,83.26756926511254,83.27118530915334,83.27118530925334,83.282031319662,83.282031319762,83.28564546923697,83.28564546933697,83.29287297266924,83.29287297276925,83.29648378780833,83.29648378790833,83.30009399668634,83.30009399678634,83.31092295610247,83.31092295620248,83.31453005789253,83.31453005799253,83.32174274581995,83.32174274591995,83.32534643739149,83.32534643749149,83.32894906800615,83.32894906810616,83.332548250511,83.332548250611,83.34334337298125,83.34334337308125,83.34693903159364,83.34693903169364,83.3541287194918,83.3541287195918,83.36131681595464,83.36131681605464,83.36490982217485,83.36490982227485,83.368502222134,83.368502222234,83.37927673172794,83.37927673182794,83.38286617616438,83.38286617626439,83.39722232458351,83.39722232468351,83.40080979867147,83.40080979877148,83.40439499928044,83.40439499938044,83.40797947995438,83.40797948005438,83.41514700143223,83.41514700153223,83.41872845080083,83.41872845090083,83.42947067699289,83.4294706770929,83.43662952135921,83.43662952145921,83.45094076856797,83.45094076866798,83.4616654889717,83.4616654890717,83.46523886748983,83.46523886758983,83.47238460146055,83.47238460156055,83.47952912290913,83.47952912300913,83.48667163611792,83.48667163621792,83.4938121410869,83.49381214118691,83.49737915386383,83.49737915396383,83.50451105750392,83.50451105760392,83.50807109827856,83.50807109837857,83.51875027331954,83.51875027341954,83.52586747512873,83.52586747522874,83.5294239920109,83.5294239921109,83.53297937215358,83.53297937225358,83.53653043268615,83.53653043278615,83.54008062171842,83.54008062181842,83.5436296740112,83.5436296741112,83.55427459530183,83.55427459540184,83.56491493174313,83.56491493184313,83.56845845190367,83.56845845200367,83.57199977436782,83.57199977446783,83.5755404905709,83.5755404906709,83.57908044894766,83.57908044904767,83.58261934636754,83.58261934646754,83.58969653494624,83.58969653504624,83.5967718289591,83.59677182905911,83.60030458798566,83.60030458808566,83.60736821147297,83.60736821157298,83.61442857630702,83.61442857640702,83.62148340900883,83.62148340910883,83.62853236855655,83.62853236865655,83.63205516216681,83.63205516226681,83.63557731162469,83.63557731172469,83.64261793508271,83.64261793518271,83.65669152795279,83.65669152805279,83.67075947501665,83.67075947511665,83.67427347784142,83.67427347794143,83.67778645760065,83.67778645770065,83.68129879320749,83.68129879330749,83.68832217611639,83.68832217621639,83.69885471176151,83.69885471186151,83.70938345827497,83.70938345837497,83.71289173951092,83.71289173961092,83.71639778515917,83.71639778525918,83.72691452012522,83.72691452022522,83.73041647351126,83.73041647361126,83.73391710070122,83.73391710080122,83.73741534073822,83.73741534083823,83.74790854519655,83.74790854529655,83.75490169083469,83.75490169093469,83.76189241142855,83.76189241152855,83.76887915343414,83.76887915353414,83.77236879214225,83.77236879224225,83.77585782458927,83.77585782468927,83.77934496247046,83.77934496257046,83.78631738155832,83.78631738165832,83.78980163969945,83.78980163979945,83.79328514001423,83.79328514011424,83.79676799617663,83.79676799627663,83.80373188971822,83.80373188981822,83.80721179035791,83.80721179045791,83.81764929458062,83.81764929468062,83.82460523094569,83.8246052310457,83.82808236551925,83.82808236561925,83.83155862859255,83.83155862869255,83.83503375492631,83.83503375502632,83.83850732771612,83.83850732781612,83.8419795364185,83.84197953651851,83.84545113885983,83.84545113895983,83.84892114986584,83.84892114996585,83.85586007302969,83.85586007312969,83.86973538063917,83.86973538073917,83.87666994630159,83.87666994640159,83.89053642523426,83.89053642533426,83.90786412438747,83.90786412448747,83.9113283001307,83.9113283002307,83.91825517385583,83.91825517395583,83.9217171519027,83.9217171520027,83.92517844790585,83.92517844800585,83.92863906186531,83.92863906196531,83.93209888010713,83.93209888020714,83.93555756160944,83.93555756170944,83.94247265113505,83.94247265123505,83.95283846498644,83.95283846508644,83.95628854515985,83.95628854525985,83.96318715196267,83.96318715206267,83.97008382630833,83.97008382640833,83.97353030680664,83.97353030690664,83.97697587791335,83.97697587801335,83.98731084823291,83.98731084833291,83.99075372905612,83.99075372915613,84.00108074219916,84.00108074229917,84.00452180423919,84.00452180433919,84.01484294422812,84.01484294432812,84.01828086128884,84.01828086138885,84.02515358832233,84.02515358842233,84.02858692053373,84.02858692063373,84.04231725596529,84.04231725606529,84.04574721584949,84.04574721594949,84.05260592309577,84.05260592319577,84.06631909376084,84.06631909386084,84.0731691996782,84.0731691997782,84.07659203599486,84.07659203609487,84.08343657188867,84.08343657198867,84.09027781123791,84.09027781133791,84.1039536968473,84.1039536969473,84.11078910093377,84.11078910103377,84.11420470000893,84.11420470010893,84.12103407937603,84.12103407947603,84.1244476702114,84.1244476703114,84.12786020008988,84.12786020018989,84.13127178268546,84.13127178278546,84.13468215275891,84.13468215285891,84.14150190773155,84.14150190783155,84.14490882969517,84.14490882979517,84.15172009701287,84.15172009711287,84.15512394977982,84.15512394987982,84.16192957129132,84.16192957139133,84.1653307716661,84.16533077176611,84.16873057006218,84.16873057016218,84.17212953484928,84.17212953494928,84.17892674448845,84.17892674458845,84.18232385260103,84.18232385270103,84.18571989975673,84.18571989985674,84.18911280193315,84.18911280203315,84.1925046431527,84.1925046432527,84.19589455191509,84.19589455201509,84.19928313448139,84.19928313458139,84.2060571167434,84.2060571168434,84.21959757878669,84.21959757888669,84.22298108391655,84.22298108401655,84.22636261869795,84.22636261879795,84.2398844761047,84.23988447620471,84.24326184284125,84.24326184294125,84.25001525011825,84.25001525021825,84.2533905707237,84.2533905708237,84.2567651335028,84.2567651336028,84.26351304653888,84.26351304663888,84.2668851842737,84.2668851843737,84.27025482118164,84.27025482128164,84.28036202679617,84.28036202689617,84.29046786832329,84.29046786842329,84.2972025193985,84.2972025194985,84.30056764723973,84.30056764733973,84.30393182779805,84.30393182789805,84.31065738495724,84.31065738505724,84.31401789005783,84.31401789015783,84.31737737209286,84.31737737219287,84.32073613419288,84.32073613429289,84.32745195328366,84.32745195338366,84.3308085176873,84.3308085177873,84.33416375589486,84.33416375599487,84.34087241352678,84.34087241362678,84.34422488566823,84.34422488576823,84.35092922369002,84.35092922379002,84.35427976337431,84.35427976347431,84.35762946944962,84.35762946954962,84.36767703413157,84.36767703423158,84.37102511088027,84.37102511098027,84.37771936981181,84.37771936991182,84.38106513519018,84.38106513529019,84.38441018063352,84.38441018073352,84.38775352096762,84.38775352106762,84.39109492884457,84.39109492894457,84.39443501052543,84.39443501062543,84.40111392347369,84.40111392357369,84.40779075239952,84.40779075249952,84.41112651447027,84.41112651457027,84.41446148082336,84.41446148092336,84.4211294810724,84.4211294811724,84.42446122666357,84.42446122676357,84.42779225231972,84.42779225241972,84.43445309110989,84.43445309120989,84.43778286635259,84.4377828664526,84.44443866559763,84.44443866569763,84.4510898042107,84.4510898043107,84.46106370817287,84.46106370827287,84.4677113607848,84.4677113608848,84.47435700515696,84.47435700525696,84.48099984557166,84.48099984567166,84.48431980696333,84.48431980706333,84.48763885896338,84.48763885906338,84.49427544731083,84.49427544741083,84.50091025476638,84.50091025486638,84.50422646491768,84.50422646501768,84.51085718011102,84.51085718021102,84.52079955849766,84.52079955859766,84.52411243421308,84.52411243431308,84.52742470366744,84.52742470376744,84.53073416916435,84.53073416926435,84.54727907160466,84.54727907170466,84.55720231487633,84.55720231497634,84.56381631999898,84.56381632009898,84.5704275211642,84.57042752126421,84.57703629728515,84.57703629738515,84.58033814662738,84.58033814672739,84.5869402917679,84.5869402918679,84.59683981507528,84.59683981517529,84.60013723113346,84.60013723123346,84.60673100229297,84.60673100239298,84.6100269026985,84.6100269027985,84.61661658159578,84.61661658169578,84.61991017063097,84.61991017073098,84.62320303973115,84.62320303983115,84.62649518889631,84.62649518899632,84.63636890821698,84.63636890831698,84.642947788089,84.642947788189,84.64623485981771,84.64623485991771,84.64951988541533,84.64951988551533,84.65608868619708,84.65608868629708,84.66265286423821,84.66265286433821,84.665932888182,84.665932888282,84.68233031761748,84.68233031771749,84.68560685556014,84.68560685566014,84.68888214308934,84.68888214318935,84.69543173297352,84.69543173307352,84.70197897359604,84.70197897369604,84.70524929736274,84.70524929746274,84.70851890119444,84.70851890129444,84.7117859663079,84.7117859664079,84.7150521978124,84.7150521979124,84.72484842939029,84.7248484294903,84.73137410985359,84.73137410995359,84.7346335967037,84.7346335968037,84.74115063794677,84.74115063804678,84.74766495101504,84.74766495111504,84.7541756265169,84.7541756266169,84.76718936136594,84.76718936146594,84.77043960273477,84.77043960283477,84.77693830458053,84.77693830468053,84.78343492240387,84.78343492250387,84.78668198090209,84.78668198100209,84.78992831946528,84.78992831956528,84.80291215806541,84.80291215816541,84.80615675362803,84.80615675372803,84.80939854523322,84.80939854533322,84.81263870751178,84.81263870761178,84.81587818774665,84.81587818784665,84.8223558220203,84.8223558221203,84.82883125859757,84.82883125869758,84.83206785909236,84.83206785919236,84.83530317128238,84.83530317138238,84.83853435331888,84.83853435341888,84.84499573221763,84.84499573231763,84.84822479234039,84.84822479244039,84.85468185162904,84.85468185172904,84.85790852459884,84.85790852469884,84.86758638370318,84.86758638380319,84.87403586472848,84.87403586482849,84.88048284492686,84.88048284502686,84.88370428889496,84.88370428899496,84.89014331191682,84.89014331201682,84.89335933742662,84.89335933752662,84.89657403674032,84.89657403684032,84.8997850605963,84.8997850606963,84.90299449301698,84.90299449311698,84.90620305393738,84.90620305403738,84.9094107812488,84.9094107813488,84.9126160077333,84.91261600783331,84.91582040060885,84.91582040070885,84.92543096473464,84.92543096483465,84.93503554203238,84.93503554213238,84.94143303617109,84.94143303627109,84.9446281077827,84.9446281078827,84.95740631020679,84.9574063103068,84.96379008768882,84.96379008778882,84.96697856621134,84.96697856631134,84.97016632479883,84.97016632489883,84.97654017475587,84.97654017485587,84.97972603877753,84.97972603887753,84.982910803951,84.982910804051,84.98609306829756,84.98609306839757,84.98927321073037,84.98927321083038,84.99881204659354,84.99881204669354,85.00834754802084,85.00834754812084,85.01787740364193,85.01787740374193,85.0242280262218,85.02422802632181,85.03057701947506,85.03057701957506,85.03375072038403,85.03375072048404,85.04009486354873,85.04009486364873,85.04960569772882,85.04960569782882,85.05594370250022,85.05594370260022,85.06227939590129,85.06227939600129,85.06544502595979,85.06544502605979,85.06860940560485,85.06860940570485,85.07810076364811,85.07810076374811,85.08758871147288,85.08758871157288,85.09074733163777,85.09074733173777,85.09706316998886,85.09706317008886,85.10021932721818,85.10021932731819,85.10968632114478,85.10968632124478,85.1128370978071,85.1128370979071,85.11598662405599,85.115986624156,85.1254330808889,85.1254330809889,85.13487590015541,85.13487590025541,85.14116896582512,85.14116896592512,85.14745975801584,85.14745975811584,85.1506037900238,85.1506037901238,85.16003516611264,85.16003516621264,85.16632004725793,85.16632004735793,85.1726028064895,85.1726028065895,85.17888355748126,85.17888355758126,85.18516051934135,85.18516051944135,85.19143369206978,85.19143369216978,85.19456889540093,85.19456889550094,85.19770171157911,85.19770171167912,85.20396450208675,85.20396450218675,85.20709409750302,85.20709409760302,85.2196111150807,85.2196111151807,85.22899565750203,85.22899565760203,85.23524799000623,85.23524799010623,85.24149782168352,85.24149782178353,85.25086955683979,85.25086955693979,85.26023917008231,85.26023917018232,85.26648320438815,85.26648320448815,85.26960264493152,85.26960264503153,85.27272060771355,85.27272060781355,85.27895513129887,85.27895513139887,85.28518776031837,85.28518776041837,85.28830106246843,85.28830106256844,85.29452706050749,85.29452706060749,85.29763884700488,85.29763884710488,85.30386090434699,85.30386090444699,85.3131904664677,85.3131904665677,85.31629903220318,85.31629903230318,85.32562321375691,85.32562321385691,85.33183693500933,85.33183693510934,85.33494144637392,85.33494144647392,85.33804501045557,85.33804501055558,85.34114633894954,85.34114633904954,85.3442469853998,85.34424698549981,85.34734604041476,85.34734604051476,85.35044373134232,85.35044373144233,85.35354081600882,85.35354081610882,85.35973411384151,85.35973411394151,85.36282964496401,85.36282964506401,85.36901979781742,85.36901979791742,85.3721123355259,85.3721123356259,85.37520426697331,85.37520426707331,85.38447718157548,85.38447718167548,85.38756536178252,85.38756536188252,85.39065267048929,85.39065267058929,85.39682607538069,85.39682607548069,85.40299652474938,85.40299652484939,85.4091643217259,85.4091643218259,85.41532995889736,85.41532995899736,85.42457506127303,85.42457506137303,85.43073615148647,85.43073615158647,85.4338154272341,85.4338154273341,85.44305230719402,85.44305230729402,85.44920858521024,85.44920858531025,85.45228566326148,85.45228566336148,85.45843678805863,85.45843678815864,85.46458450263727,85.46458450273727,85.47073104258509,85.47073104268509,85.47380300530857,85.47380300540857,85.47994526353759,85.47994526363759,85.48301453597757,85.48301453607758,85.48915190622674,85.48915190632674,85.49528741980139,85.49528741990139,85.50448784831458,85.50448784841458,85.5198155679867,85.5198155680867,85.52900819088865,85.52900819098865,85.53819600159336,85.53819600169336,85.54431704068502,85.54431704078502,85.55043633677607,85.55043633687607,85.55655328360551,85.55655328370551,85.55961054449809,85.55961054459809,85.56572408110901,85.56572408120901,85.5687770602828,85.5687770603828,85.57182901639102,85.57182901649102,85.57487816854182,85.57487816864182,85.57792667654023,85.57792667664023,85.5840215706233,85.58402157072331,85.58706746412085,85.58706746422085,85.59011157672651,85.59011157682652,85.59315356741844,85.59315356751844,85.59619351197927,85.59619351207927,85.59923231980058,85.59923231990058,85.61138649012898,85.61138649022898,85.62049836663492,85.62049836673492,85.62960618876997,85.62960618886997,85.63263859295876,85.63263859305876,85.63566902679908,85.63566902689908,85.64475881266735,85.64475881276735,85.64778272920118,85.64778272930118,85.6538254848324,85.6538254849324,85.65684379345134,85.65684379355135,85.6689130493389,85.6689130494389,85.67192684889115,85.67192684899115,85.68096623930813,85.68096623940814,85.6869898598244,85.6869898599244,85.68999987024497,85.68999987034498,85.69902777959298,85.69902777969298,85.70504389762851,85.70504389772852,85.70805080096113,85.70805080106113,85.72007629237783,85.72007629247783,85.72307989916587,85.72307989926587,85.72908608967641,85.72908608977642,85.73208681672439,85.73208681682439,85.73808762666796,85.73808762676796,85.74108687595458,85.74108687605458,85.74708404833174,85.74708404843174,85.75008163040043,85.75008163050043,85.75607463473274,85.75607463483274,85.76206430462919,85.7620643047292,85.78002501612013,85.78002501622014,85.7860040764479,85.7860040765479,85.7919805601661,85.7919805602661,85.7949673432095,85.79496734330951,85.80392636614366,85.80392636624366,85.80989697670776,85.80989697680776,85.81885050540097,85.81885050550098,85.82183266570374,85.82183266580374,85.82779516752606,85.82779516762606,85.83077175780527,85.83077175790527,85.83374740080156,85.83374740090156,85.84564841924269,85.8456484193427,85.84862235712973,85.84862235722973,85.85456928562087,85.85456928572087,85.86051435743748,85.86051435753748,85.86348234638778,85.86348234648779,85.86941722544019,85.86941722554019,85.87238350928124,85.87238350938124,85.87534861849146,85.87534861859146,85.8812779275203,85.8812779276203,85.88424155896918,85.88424155906918,85.88720447048303,85.88720447058303,85.8931263907051,85.89312639080511,85.89608358063013,85.89608358073014,85.89903955803301,85.89903955813301,85.90790449682764,85.90790449692764,85.9108569882294,85.9108569883294,85.91676060694547,85.91676060704548,85.92561425412772,85.92561425422772,85.9285639036807,85.9285639037807,85.9374079643598,85.93740796445981,85.9491883747397,85.94918837483971,85.95507634434198,85.95507634444198,85.96390528638568,85.96390528648568,85.9697861324204,85.9697861325204,85.97272388409992,85.97272388419992,85.97859768234973,85.97859768244973,85.98740472321235,85.98740472331235,85.996206383508,85.996206383608,85.9991368221634,85.9991368222634,86.00792597832451,86.00792597842451,86.01378215711202,86.01378215721202,86.01670884452705,86.01670884462705,86.0254856481189,86.0254856482189,86.03133167203352,86.03133167213352,86.04301955181789,86.0430195519179,86.04594032818751,86.04594032828751,86.05178074418725,86.05178074428726,86.0605392083818,86.0605392084818,86.06929467916231,86.06929467926231,86.0751281989424,86.0751281990424,86.0780417001792,86.0780417002792,86.0838677932612,86.08386779336121,86.09260356055701,86.09260356065701,86.09842359102832,86.09842359112832,86.10133030971943,86.10133030981943,86.10423460336625,86.10423460346625,86.11004129609407,86.11004129619407,86.11584563956025,86.11584563966025,86.12454927501945,86.12454927511945,86.13034960200605,86.13034960210605,86.13614818599208,86.13614818609209,86.14194335975961,86.14194335985961,86.15063013358285,86.15063013368285,86.16220778752678,86.16220778762678,86.16510067588727,86.16510067598728,86.16799216226903,86.16799216236903,86.17377396040175,86.17377396050175,86.17955306825097,86.17955306835097,86.18532891744694,86.18532891754694,86.18821490958778,86.18821490968779,86.19109987866307,86.19109987876307,86.19398420358596,86.19398420368596,86.20263429861458,86.20263429871459,86.2083980604805,86.2083980605805,86.2170403877892,86.2170403878892,86.21991940792775,86.21991940802775,86.22279668506573,86.22279668516573,86.22567320437737,86.22567320447737,86.22854896586269,86.22854896596269,86.23142366639112,86.23142366649112,86.23717177914324,86.23717177924324,86.24579224316214,86.24579224326214,86.2486649354508,86.2486649355508,86.25153641521732,86.25153641531732,86.25727842746743,86.25727842756743,86.26014683803729,86.26014683813729,86.26588066576298,86.26588066586298,86.26874596924488,86.26874596934488,86.27733858314596,86.27733858324596,86.28019990803959,86.2801999081396,86.28306043721557,86.28306043731557,86.2887803209367,86.2887803210367,86.29163918289473,86.29163918299473,86.29449580926507,86.29449580936507,86.30592095065907,86.30592095075907,86.31162973222439,86.31162973232439,86.31448150850619,86.31448150860619,86.3173305945045,86.3173305946045,86.32587561691174,86.32587561701175,86.33156893881983,86.33156893891983,86.33725972200969,86.33725972210969,86.34010289696259,86.3401028970626,86.34863029990754,86.34863030000754,86.35431278489904,86.35431278499904,86.35999326165073,86.35999326175073,86.36567104811894,86.36567104821894,86.3685073268522,86.3685073269522,86.3826864091481,86.38268640924811,86.38835521537425,86.38835521547425,86.39118831123689,86.39118831133689,86.39401758007654,86.39401758017654,86.4081616886871,86.4081616887871,86.41098803989536,86.41098803999536,86.41663790046312,86.41663790056312,86.4194604625397,86.4194604626397,86.42228219100731,86.42228219110731,86.4279246627683,86.4279246628683,86.43074423143086,86.43074423153087,86.43356289070182,86.43356289080182,86.43919865569974,86.43919865579974,86.44483252613183,86.44483252623183,86.44764743416243,86.44764743426244,86.45327603770149,86.45327603780149,86.45890070054361,86.45890070064361,86.46452051329717,86.46452051339718,86.4729472011222,86.4729472012222,86.47575483401998,86.47575483411998,86.48136937988052,86.48136937998052,86.48417481508193,86.48417481518193,86.49539481288699,86.49539481298699,86.49819865665309,86.49819865675309,86.50100174259286,86.50100174269286,86.50380251716231,86.50380251726232,86.50940255064853,86.50940255074853,86.51499970439468,86.51499970449468,86.52339058492537,86.52339058502537,86.52898163816951,86.52898163826951,86.53177331882294,86.53177331892294,86.53735550549897,86.53735550559897,86.54293579760916,86.54293579770916,86.55688249245941,86.55688249255941,86.55967061132907,86.55967061142907,86.56245615358917,86.56245615368917,86.57638159141072,86.57638159151072,86.58194956884297,86.58194956894297,86.59029804899019,86.5902980490902,86.59307938531414,86.59307938541414,86.59585988802912,86.59585988812913,86.59863622269927,86.59863622279927,86.60973602924757,86.60973602934757,86.61804904312231,86.61804904322231,86.62912516759765,86.62912516769765,86.63465911862745,86.63465911872746,86.63742416168519,86.63742416178519,86.64294882934239,86.64294882944239,86.64570758244155,86.64570758254155,86.65398213662979,86.6539821367298,86.65673611542304,86.65673611552305,86.66224286048741,86.66224286058741,86.66774759731199,86.66774759741199,86.67049884793043,86.67049884803043,86.67324899970069,86.67324899980069,86.6759979010575,86.6759979011575,86.68149430179238,86.68149430189239,86.6924835793697,86.69248357946971,86.70072130319805,86.70072130329805,86.703465316575,86.70346531667501,86.70620811742981,86.70620811752981,86.71169091518199,86.71169091528199,86.71442973744854,86.71442973754854,86.71990620735082,86.71990620745082,86.72537994907829,86.72537994917829,86.72811483064788,86.72811483074788,86.73358296446048,86.73358296456048,86.73631496629001,86.73631496639001,86.74450953190858,86.74450953200858,86.74723910869383,86.74723910879383,86.75542628550562,86.75542628560562,86.75815332357266,86.75815332367266,86.76087869442176,86.76087869452176,86.76905336709902,86.76905336719902,86.77722565262333,86.77722565272333,86.78267105164586,86.78267105174587,86.7881139498415,86.7881139499415,86.79083386434098,86.79083386444098,86.79899031129489,86.79899031139489,86.80713952100729,86.80713952110729,86.80985367602662,86.80985367612662,86.81799424651881,86.81799424661881,86.8234187295345,86.8234187296345,86.82884071172327,86.82884071182328,86.83697108945124,86.83697108955124,86.84509676865594,86.84509676875594,86.84780421691221,86.84780421701221,86.85592451554993,86.85592451564993,86.86133429673471,86.86133429683471,86.86403799375063,86.86403799385063,86.86673960674412,86.86673960684412,86.8721416959916,86.8721416960916,86.87754181489062,86.87754181499062,86.88024066181798,86.88024066191798,86.88563721893323,86.88563721903323,86.89103195726526,86.89103195736526,86.899120199849,86.899120199949,86.90181472716625,86.90181472726626,86.90720264506126,86.90720264516126,86.91528300625112,86.91528300635112,86.91797552532859,86.91797552542859,86.92066740025368,86.92066740035368,86.92335863102637,86.92335863112638,86.93142959516967,86.93142959526968,86.93680584253913,86.93680584263913,86.94217928595114,86.94217928605114,86.9475489781228,86.9475489782228,86.95023183491449,86.95023183501449,86.95291238033587,86.95291238043588,86.95827059143855,86.95827059153855,86.96094757507615,86.96094757517615,86.96362164108236,86.96362164118236,86.96896863635527,86.96896863645527,86.97431347182314,86.97431347192314,86.97698507489376,86.97698507499376,86.97965573068146,86.97965573078146,86.98499510979971,86.98499510989971,86.98766318897788,86.98766318907788,86.99299692228992,86.99299692238992,86.99566246274986,86.99566246284986,87.00099104284284,87.00099104294284,87.00365374145402,87.00365374155402,87.01163857863432,87.01163857873432,87.01429782913569,87.01429782923569,87.01695510928857,87.01695510938858,87.0222673203327,87.0222673204327,87.02492164496289,87.02492164506289,87.02757532544068,87.02757532554068,87.03553428285164,87.03553428295164,87.03818451521961,87.03818451531961,87.04083361084807,87.04083361094807,87.04348210021547,87.04348210031547,87.05142620423027,87.05142620433027,87.05936788320079,87.0593678833008,87.06995248151514,87.06995248161515,87.07524188198659,87.07524188208659,87.07788529391755,87.07788529401755,87.08052806169611,87.08052806179612,87.08316889701753,87.08316889711753,87.08580814090364,87.08580814100364,87.09108488567529,87.09108488577529,87.0963590917287,87.0963590918287,87.10426735055785,87.10426735065785,87.10953803271882,87.10953803281882,87.11217246440769,87.1121724645077,87.11480602459629,87.11480602469629,87.12007128829896,87.12007128839896,87.1305965867026,87.1305965868026,87.13322616830293,87.13322616840293,87.13585453738114,87.13585453748114,87.1411105177112,87.1411105178112,87.14373767426726,87.14373767436726,87.14899081274856,87.14899081284857,87.15424190509877,87.15424190519877,87.15686667450188,87.15686667460189,87.16736222822186,87.16736222832186,87.17522566162327,87.17522566172327,87.17784474732888,87.17784474742888,87.18046296153418,87.18046296163418,87.18308026634789,87.18308026644789,87.19092983152737,87.19092983162737,87.19615871956596,87.19615871966596,87.2039977888507,87.2039977889507,87.2118338268301,87.2118338269301,87.21705415143109,87.21705415153109,87.21966209708955,87.21966209718956,87.2274846457602,87.2274846458602,87.23009073474414,87.23009073484414,87.23530113182015,87.23530113192015,87.24571878099287,87.24571878109288,87.25092569206774,87.25092569216774,87.25352715831104,87.25352715841105,87.25612657842323,87.25612657852324,87.26392358834637,87.26392358844637,87.2717178322034,87.2717178323034,87.27950908264638,87.27950908274639,87.28210452417032,87.28210452427032,87.28988891628498,87.28988891638498,87.29248253902571,87.29248253912571,87.29766887511559,87.29766887521559,87.30544496903187,87.30544496913187,87.3080342721625,87.3080342722625,87.3106228553581,87.3106228554581,87.31579858187928,87.31579858197928,87.32097154233432,87.32097154243432,87.32872668235245,87.32872668245246,87.33131041545951,87.33131041555951,87.33389289815311,87.33389289825311,87.33905555217001,87.33905555227001,87.3442159327079,87.3442159328079,87.34937305459255,87.34937305469255,87.35968203146881,87.35968203156881,87.37256174473048,87.37256174483048,87.37513418622514,87.37513418632514,87.37770575621953,87.37770575631953,87.38027554532202,87.38027554542202,87.38284355353262,87.38284355363263,87.38797680388771,87.38797680398771,87.39054007568372,87.39054007578372,87.39566586144942,87.39566586154942,87.3982275039188,87.3982275040188,87.40591152193537,87.40591152203537,87.41103207869935,87.41103207879935,87.41359042462418,87.41359042472418,87.4161473685703,87.4161473686703,87.4212603091796,87.4212603092796,87.42381463862485,87.42381463872485,87.42636779343928,87.42636779353928,87.42891973573158,87.42891973583158,87.4314710338715,87.4314710339715,87.44167486234376,87.44167486244376,87.44422475850496,87.44422475860496,87.44932371721838,87.44932371731838,87.45442047823543,87.45442047833544,87.45951329855554,87.45951329865554,87.46205766258448,87.46205766268449,87.46714502655499,87.46714502665499,87.46968760969204,87.46968760979205,87.47477129820481,87.47477129830482,87.47731217623263,87.47731217633263,87.47985237221673,87.47985237231673,87.48493174111938,87.48493174121938,87.48747064879872,87.48747064889872,87.49000891232568,87.49000891242568,87.49508392372691,87.49508392382691,87.50015525967041,87.50015525977041,87.50775894809546,87.50775894819546,87.51535884738882,87.51535884748883,87.51788786543449,87.51788786553449,87.52800177781208,87.52800177791208,87.530528673944,87.530528674044,87.53558117790308,87.53558117800308,87.54315633416661,87.54315633426661,87.5456793274929,87.54567932759291,87.5507239500581,87.5507239501581,87.55324463201407,87.55324463211407,87.55828436659941,87.55828436669941,87.5608025856198,87.5608025857198,87.56835557546302,87.56835557556302,87.57338939900293,87.57338939910294,87.57842110062911,87.57842110072912,87.58345162762447,87.58345162772447,87.59099525831245,87.59099525841245,87.59602267821984,87.59602267831984,87.59853536510798,87.59853536520798,87.60607168277183,87.60607168287183,87.61862712214469,87.61862712224469,87.62615593732782,87.62615593742782,87.63618845915701,87.63618845925701,87.64120064675507,87.64120064685507,87.64370552803196,87.64370552813196,87.6487132065633,87.6487132066633,87.65622015845666,87.65622015855666,87.6662251711899,87.6662251712899,87.67122485465342,87.67122485475342,87.67872059071702,87.67872059081702,87.68121652964315,87.68121652974315,87.68371186230821,87.68371186240822,87.69369114683737,87.69369114693737,87.70117229474403,87.70117229484403,87.70615810998561,87.70615811008561,87.71363327106421,87.71363327116421,87.71861469091304,87.71861469101304,87.72110437777192,87.72110437787192,87.72608242529357,87.72608242539357,87.73105899505386,87.73105899515386,87.73603404916149,87.73603404926149,87.73852055314975,87.73852055324976,87.74597870102713,87.74597870112713,87.74846384092798,87.74846384102798,87.75094833667646,87.75094833677646,87.75840019459525,87.75840019469526,87.76088256842999,87.76088256853,87.76832764380308,87.76832764390308,87.77080744102828,87.77080744112828,87.77328655620977,87.77328655630977,87.77824311935481,87.77824311945481,87.78319744691218,87.78319744701218,87.78567372024492,87.78567372034492,87.79310083513387,87.79310083523387,87.80299899435384,87.80299899445384,87.80794509949547,87.80794509959547,87.81041648484835,87.81041648494835,87.81782969362405,87.81782969372405,87.82029800978026,87.82029800988026,87.82276499974039,87.82276499984039,87.82769769135588,87.82769769145588,87.83756099056444,87.83756099066444,87.84002566915424,87.84002566925425,87.84988271629554,87.84988271639554,87.85234470460186,87.85234470470186,87.85726671086603,87.85726671096603,87.86218655732515,87.86218655742515,87.86464494595637,87.86464494605637,87.86956107906643,87.86956107916643,87.87201764891445,87.87201764901445,87.87447353671878,87.87447353681878,87.87692809832701,87.87692809842702,87.8793807653694,87.8793807654694,87.88428329549676,87.88428329559676,87.88673217340748,87.88673217350748,87.89162928507653,87.89162928517653,87.90142100758774,87.90142100768774,87.90386730888892,87.90386730898892,87.90631228399401,87.90631228409401,87.91120037752968,87.91120037762968,87.91364307915576,87.91364307925576,87.91608487139025,87.91608487149026,87.9185259057984,87.9185259058984,87.92096595503232,87.92096595513232,87.92828382925507,87.92828382935507,87.93803644792737,87.93803644802738,87.94291010487136,87.94291010497136,87.94778194303214,87.94778194313214,87.95265181084444,87.95265181094445,87.95995555160603,87.95995555170603,87.96482291859144,87.96482291869144,87.96725497275752,87.96725497285752,87.97454973327703,87.97454973337703,87.9794097872383,87.9794097873383,87.98426521845893,87.98426521855893,87.98911583748232,87.98911583758232,87.9939644103746,87.9939644104746,87.99881006563551,87.99881006573551,88.00365363687398,88.00365363697398,88.01091588664373,88.01091588674373,88.01333456517499,88.01333456527499,88.01817108862855,88.01817108872855,88.02300583119022,88.02300583129022,88.03267152718189,88.03267152728189,88.03508671971201,88.03508671981201,88.0399151344238,88.0399151345238,88.04232699251804,88.04232699261804,88.05197177250285,88.05197177260285,88.05920067791385,88.05920067801385,88.06401776312191,88.06401776322191,88.0664247332363,88.0664247333363,88.07364382479625,88.07364382489625,88.0784556431113,88.0784556432113,88.08085961981165,88.08085961991165,88.08806798812897,88.08806798822897,88.0928687042882,88.0928687043882,88.09766566920706,88.09766566930706,88.10246058799481,88.10246058809481,88.10725338486884,88.10725338496884,88.1096485328924,88.1096485329924,88.1144374648521,88.1144374649521,88.11683071830983,88.11683071840983,88.12400888724768,88.12400888734769,88.13118478270654,88.13118478280654,88.13357420914126,88.13357420924126,88.13835215261912,88.13835215271912,88.1407392297922,88.1407392298922,88.14312414716024,88.14312414726024,88.15027787619879,88.15027787629879,88.15504543745585,88.15504543755586,88.15981083890786,88.15981083900786,88.16218935264335,88.16218935274335,88.16932368132773,88.16932368142773,88.17407794273261,88.17407794283261,88.17645285679302,88.17645285689302,88.18120113136985,88.18120113146985,88.18357358249467,88.18357358259468,88.18831719643954,88.18831719653954,88.19068756354194,88.19068756364194,88.1954270852246,88.1954270853246,88.19779578510907,88.19779578520907,88.20016380294982,88.20016380304982,88.20253094929029,88.2025309493903,88.20962977381086,88.20962977391086,88.21672257361206,88.21672257371206,88.21908361945054,88.21908361955055,88.2214425812666,88.2214425813666,88.2285174963663,88.2285174964663,88.23794478019292,88.23794478029292,88.24029957396414,88.24029957406414,88.24971761230947,88.24971761240947,88.25442303180705,88.25442303190705,88.26382856601786,88.26382856611787,88.26617820657,88.26617820667,88.27557373747318,88.27557373757318,88.27792004358945,88.27792004368945,88.28261193588693,88.28261193598694,88.28495691580711,88.28495691590712,88.28729988748749,88.2872998875875,88.29198427730428,88.29198427740428,88.29432452080985,88.29432452090985,88.29666370335856,88.29666370345856,88.30134096960778,88.30134096970778,88.30601679598698,88.30601679608698,88.30835165892557,88.30835165902558,88.31068576403784,88.31068576413784,88.31535226915311,88.31535226925311,88.31768421446031,88.31768421456032,88.32234666520469,88.32234666530469,88.32700699403533,88.32700699413533,88.33166304114717,88.33166304124717,88.33863864310564,88.33863864320564,88.34560659101813,88.34560659111813,88.35257260647346,88.35257260657346,88.35489341173354,88.35489341183354,88.35953369605761,88.35953369615761,88.36185188681684,88.36185188691684,88.37112290685316,88.37112290695316,88.37575599182705,88.37575599192705,88.38038596971296,88.38038596981296,88.38269684744806,88.38269684754806,88.38731776930926,88.38731776940926,88.39193664503937,88.39193664513937,88.39655415668207,88.39655415678207,88.39886135895944,88.39886135905944,88.41039392223642,88.41039392233643,88.41500359037656,88.41500359047656,88.41961174286402,88.41961174296402,88.42421860704673,88.42421860714673,88.4288234250983,88.4288234251983,88.4380299541135,88.4380299542135,88.44033022227991,88.44033022237991,88.44722894275668,88.44722894285668,88.4541241772323,88.4541241773323,88.4587186130631,88.4587186131631,88.4633107754149,88.4633107755149,88.46790077796163,88.46790077806163,88.47248960587754,88.47248960597754,88.47937112369651,88.47937112379651,88.48395692030708,88.48395692040708,88.49083317123302,88.49083317133302,88.49312241302624,88.49312241312624,88.49998801649217,88.49998801659217,88.50227445432795,88.50227445442795,88.50913175959549,88.50913175969549,88.51598512416611,88.51598512426611,88.52511931865762,88.52511931875762,88.53196609013912,88.53196609023912,88.54109058445354,88.54109058455354,88.55020931928782,88.55020931938782,88.55476722788926,88.55476722798926,88.56387906650389,88.56387906660389,88.56843189766887,88.56843189776887,88.57525904144829,88.57525904154829,88.57753267201902,88.57753267211902,88.58207879642094,88.58207879652095,88.58435106290426,88.58435106300426,88.59116649826679,88.59116649836679,88.59797977382428,88.59797977392428,88.60024863008908,88.60024863018909,88.60478490274865,88.60478490284865,88.6070518265563,88.6070518266563,88.61158484056264,88.61158484066264,88.61384968034787,88.61384968044787,88.62290710703162,88.62290710713162,88.6251704311642,88.6251704312642,88.62743311114436,88.62743311124436,88.63195767538706,88.63195767548706,88.63421861236667,88.63421861246667,88.63647890519388,88.63647890529388,88.63873836441213,88.63873836451214,88.64777525400221,88.64777525410221,88.65229170950312,88.65229170960312,88.65906410032981,88.65906410042982,88.66583251256826,88.66583251266826,88.66808594706714,88.66808594716714,88.6703370701957,88.6703370702957,88.67258743549793,88.67258743559793,88.67708634731919,88.67708634741919,88.68383009231042,88.68383009241042,88.68832316886889,88.68832316896889,88.69506043444495,88.69506043454496,88.69954741050142,88.69954741060143,88.70403105212202,88.70403105222202,88.70627092152951,88.70627092162951,88.708509237393,88.708509237493,88.71074622706043,88.71074622716043,88.71298143583594,88.71298143593594,88.7174505271909,88.71745052729091,88.72191806500189,88.72191806510189,88.72638359457306,88.72638359467307,88.72861486265165,88.72861486275166,88.73084544868654,88.73084544878654,88.73530521877757,88.73530521887757,88.73976328375936,88.73976328385936,88.74421987097979,88.74421987107979,88.75090181523339,88.75090181533339,88.75535389338714,88.75535389348714,88.76425581410692,88.76425581420692,88.77315265739027,88.77315265749027,88.77982054396534,88.77982054406534,88.7842631492899,88.7842631493899,88.79092402597138,88.79092402607138,88.79314267624149,88.79314267634149,88.79979692194253,88.79979692204253,88.80423134274267,88.80423134284267,88.80866231543297,88.80866231553297,88.81087615350584,88.81087615360585,88.81751592472389,88.8175159248239,88.82194049378167,88.82194049388167,88.82636214520805,88.82636214530805,88.83078148526411,88.83078148536411,88.84182638729443,88.84182638739443,88.84624106671853,88.84624106681854,88.85065305585914,88.85065305595914,88.8528580652552,88.8528580653552,88.85506159688993,88.85506159698993,88.85946729607195,88.85946729617196,88.86607410184442,88.86607410194442,88.87047646659057,88.87047646669058,88.8770786496224,88.8770786497224,88.8792783163428,88.87927831644281,88.8858750051337,88.8858750052337,88.89027107992128,88.89027108002128,88.8968623881452,88.8968623882452,88.89905815205998,88.89905815215998,88.90344808845423,88.90344808855423,88.90783461462998,88.90783461472998,88.91221890521804,88.91221890531804,88.91440955380506,88.91440955390506,88.91878941110906,88.91878941120906,88.92316699493405,88.92316699503405,88.92535411962861,88.92535411972861,88.92972719438691,88.92972719448692,88.93847008525026,88.93847008535026,88.9428371352892,88.9428371353892,88.94719975204407,88.94719975214407,88.94938005630162,88.94938005640162,88.95373964175116,88.95373964185116,88.96027650015289,88.96027650025289,88.96463256170996,88.96463256180996,88.9668096830969,88.9668096831969,88.96898464467878,88.96898464477879,88.97768282378836,88.97768282388836,88.98202891992914,88.98202892002914,88.98637137850349,88.9863713786035,88.99288159930954,88.99288159940954,88.99505030882416,88.99505030892416,88.99938568172229,88.99938568182229,89.0037185537935,89.0037185538935,89.01021479270368,89.01021479280368,89.01237687123786,89.01237687133786,89.01886155329642,89.01886155339642,89.0210215856995,89.0210215857995,89.02318002666728,89.02318002676728,89.02965383391792,89.02965383401792,89.03612442040665,89.03612442050665,89.0382800953083,89.0382800954083,89.04043467136177,89.04043467146177,89.04258845169758,89.04258845179758,89.04689472406443,89.04689472416443,89.04904623092125,89.04904623102125,89.05119705573435,89.05119705583435,89.05549749283843,89.05549749293843,89.05764604417253,89.05764604427253,89.06194204799255,89.06194204809255,89.0662365740512,89.0662365741512,89.07052977391378,89.07052977401378,89.07696773597877,89.07696773607877,89.08125737405757,89.08125737415757,89.08340043115075,89.08340043125075,89.08768502968441,89.08768502978441,89.0919662558909,89.0919662559909,89.09624429922677,89.09624429932677,89.09838125581794,89.09838125591794,89.1026540701521,89.1026540702521,89.10906099922867,89.10906099932868,89.11119518975372,89.11119518985372,89.11972979204886,89.11972979214886,89.12399544492415,89.12399544502415,89.1303921433452,89.1303921434452,89.13465415865409,89.13465415875409,89.13891344578816,89.13891344588816,89.14104146002857,89.14104146012858,89.14316864066004,89.14316864076004,89.14742129681368,89.14742129691368,89.15167126268383,89.15167126278384,89.1559178562268,89.1559178563268,89.16016134268177,89.16016134278178,89.1644017220488,89.1644017221488,89.16651950563369,89.16651950573369,89.16863611458776,89.16863611468776,89.17921760581415,89.17921760591415,89.183447678743,89.183447678843,89.18979057149426,89.18979057159426,89.19190210301188,89.19190210311189,89.19612399141631,89.19612399151632,89.20245487262017,89.20245487272017,89.20456356228904,89.20456356238904,89.20877999434386,89.20877999444386,89.2151022742188,89.2151022743188,89.22142061339679,89.22142061349679,89.2277367169871,89.2277367170871,89.23194500240882,89.23194500250882,89.23404747790174,89.23404747800174,89.23614885454649,89.23614885464649,89.24245033208852,89.24245033218853,89.24454761647104,89.24454761657104,89.24664425670119,89.24664425680119,89.25083518789984,89.25083518799984,89.25711779556613,89.25711779566613,89.26130436926334,89.26130436936334,89.26967323493899,89.269673235039,89.27176313051476,89.27176313061476,89.27803145315463,89.27803145325463,89.2801186963382,89.28011869643821,89.2822052574781,89.2822052575781,89.28637728090968,89.28637728100968,89.28846213694031,89.28846213704031,89.29263109117524,89.29263109127524,89.30096539997,89.30096540007,89.309296866916,89.309296867016,89.31346002377947,89.31346002387947,89.31553972659103,89.31553972669103,89.32593611873504,89.32593611883505,89.33217158960329,89.3321715897033,89.336325804116,89.336325804216,89.34047804828023,89.34047804838023,89.34462885257443,89.34462885267443,89.34670306114505,89.34670306124505,89.34877571617169,89.34877571627169,89.35292023050731,89.35292023060731,89.35706322919026,89.35706322929026,89.35913353495525,89.35913353505525,89.36120285554601,89.36120285564601,89.36533994318354,89.36533994328354,89.37361017776168,89.37361017786168,89.37774196061486,89.37774196071486,89.38187253094591,89.38187253104591,89.38600169929825,89.38600169939825,89.39012726797549,89.39012726807549,89.3942498432387,89.3942498433387,89.39630991834817,89.39630991844817,89.4004271130444,89.4004271131444,89.4045408217395,89.4045408218395,89.40865214328164,89.40865214338164,89.41481656793097,89.41481656803097,89.42097864331865,89.42097864341865,89.42713696746598,89.42713696756599,89.43124101387531,89.43124101397531,89.43739524576043,89.43739524586043,89.44559742753364,89.44559742763364,89.44764624892204,89.44764624902204,89.45379097008666,89.45379097018666,89.45583827582239,89.45583827592239,89.45993137164116,89.45993137174116,89.46811536558235,89.46811536568235,89.47016009470853,89.47016009480853,89.47424769628637,89.47424769638637,89.47833370642893,89.47833370652893,89.48037449485817,89.48037449495817,89.48445474552057,89.48445474562057,89.4885310175947,89.4885310176947,89.49463849159494,89.49463849169494,89.4987076022102,89.4987076023102,89.50277455302043,89.50277455312043,89.50683930613427,89.50683930623427,89.5109026572694,89.5109026573694,89.5169946716124,89.5169946717124,89.51902276645052,89.51902276655052,89.52510564898618,89.52510564908619,89.52915770850892,89.52915770860892,89.53117993019295,89.53117993029295,89.53320139405065,89.53320139415065,89.5372431092439,89.5372431093439,89.5392625648618,89.5392625649618,89.55137607780733,89.55137607790734,89.5594442758845,89.5594442759845,89.56145998026204,89.56145998036205,89.5695202211627,89.5695202212627,89.57354693139452,89.57354693149452,89.57757201229973,89.57757201239973,89.58360510564516,89.58360510574516,89.58561345910726,89.58561345920727,89.58762071372118,89.58762071382118,89.59363978727941,89.59363978737942,89.59564348010956,89.59564348020956,89.597646490896,89.597646490996,89.60165122416409,89.60165122426409,89.60765480017378,89.60765480027378,89.60965409761117,89.60965409771117,89.61365185887699,89.613651858977,89.61564911018328,89.61564911028329,89.61764567944587,89.61764567954587,89.61964152877343,89.61964152887343,89.62363167388457,89.62363167398458,89.62762003810384,89.62762003820384,89.62961236353895,89.62961236363896,89.6335952714086,89.6335952715086,89.63757590579924,89.63757590589924,89.6395645178853,89.6395645179853,89.64751703377242,89.64751703387242,89.65149073405209,89.65149073415209,89.65347582224571,89.65347582234571,89.6554595463519,89.65545954645191,89.65744209582729,89.6574420959273,89.66140560334276,89.66140560344276,89.66536782255346,89.66536782265347,89.67130812006417,89.67130812016417,89.67328710775578,89.67328710785578,89.67724390850817,89.67724390860818,89.6811993830645,89.6811993831645,89.68515220522863,89.68515220532863,89.68712706276672,89.68712706286672,89.68910070778267,89.68910070788267,89.695019786156,89.695019786256,89.69896268079513,89.69896268089514,89.70290284725948,89.70290284735948,89.70684123283195,89.70684123293195,89.70880930782432,89.70880930792433,89.71077643553379,89.71077643563379,89.71470947843059,89.7147094785306,89.71864081621813,89.71864081631813,89.72257014576589,89.72257014586589,89.72649788387832,89.72649788397833,89.73042380320757,89.73042380330757,89.7363116970272,89.7363116971272,89.74023473661637,89.74023473671637,89.7480787696636,89.74807876976361,89.75199908107797,89.75199908117797,89.75395810004565,89.75395810014565,89.75983307292627,89.75983307302627,89.76374546505544,89.76374546515544,89.76765528057506,89.76765528067506,89.771563580442,89.77156358054201,89.78132976420572,89.78132976430572,89.78328150804057,89.78328150814058,89.78718291168788,89.78718291178788,89.79693132442404,89.79693132452404,89.80082890104833,89.80082890114834,89.80862083353503,89.80862083363503,89.81835655268007,89.81835655278007,89.82030227179557,89.82030227189557,89.8222462479105,89.8222462480105,89.82418931463383,89.82418931473383,89.82807362929726,89.82807362939727,89.83195335911138,89.83195335921138,89.83970542993285,89.83970543003285,89.8435795897234,89.8435795898234,89.84551466137889,89.84551466147889,89.8513174891924,89.8513174892924,89.85518297187141,89.85518297197142,89.86097966129161,89.86097966139161,89.86677350886305,89.86677350896305,89.86870244212523,89.86870244222523,89.87255886877954,89.87255886887954,89.87833888602036,89.87833888612036,89.8821888332595,89.8821888333595,89.88411181758495,89.88411181768495,89.88603366517088,89.88603366527089,89.88795407288679,89.88795407298679,89.89179246327433,89.89179246337433,89.89371006703279,89.89371006713279,89.8956268750736,89.8956268751736,89.8994593165244,89.8994593166244,89.90137441945596,89.90137441955596,89.90711866729376,89.90711866739376,89.91094614498206,89.91094614508206,89.9166841028613,89.9166841029613,89.92432860023447,89.92432860033448,89.93005852515455,89.93005852525455,89.93578503985611,89.93578503995612,89.93959717156113,89.93959717166113,89.94531053797577,89.94531053807577,89.9472137809252,89.9472137810252,89.95292184255548,89.95292184265548,89.95672506980107,89.95672506990107,89.96242627310303,89.96242627320304,89.96432519644235,89.96432519654235,89.96622260412897,89.96622260422897,89.97001613119747,89.97001613129747,89.97191172010089,89.9719117202009,89.97759723639773,89.97759723649773,89.98328066867212,89.98328066877212,89.98517417355313,89.98517417365314,89.98895985711906,89.98895985721906,89.99274360822783,89.99274360832783,89.9965253132055,89.9965253133055,90.00975680945194,90.00975680955194,90.01353381590633,90.01353381600633,90.0173080562946,90.0173080563946,90.0248517248739,90.0248517249739,90.03616061570807,90.03616061580807,90.04181123410217,90.04181123420217,90.04369272743575,90.04369272753576,90.05121733668273,90.05121733678273,90.05497770884907,90.05497770894907,90.05873641379745,90.05873641389745,90.06249307261474,90.06249307271474,90.0643688443595,90.06436884445951,90.06624336569081,90.06624336579081,90.06811712919578,90.06811712929579,90.06998998330916,90.06998998340916,90.07560695421398,90.07560695431398,90.0774783684573,90.07747836855731,90.08309117131728,90.08309117141728,90.08496103201664,90.08496103211664,90.08869980613241,90.08869980623241,90.09243668568236,90.09243668578236,90.09430427290269,90.09430427300269,90.09617068549223,90.09617068559223,90.09803626447278,90.09803626457278,90.09990123719227,90.09990123729227,90.10363027323963,90.10363027333963,90.1054915705014,90.1054915706014,90.10921238413303,90.10921238423303,90.11107133213315,90.11107133223315,90.11292967387222,90.11292967397222,90.11478710621967,90.11478710631967,90.11850064471848,90.11850064481848,90.12406748541123,90.12406748551123,90.1314865294497,90.1314865295497,90.13889977611673,90.13889977621673,90.14630741486887,90.14630741496887,90.14815825412721,90.14815825422721,90.15185872012174,90.15185872022174,90.1537077784882,90.1537077785882,90.15555570011514,90.15555570021515,90.15925021717294,90.15925021727294,90.16294355959994,90.16294355969994,90.16478920774787,90.16478920784787,90.16663424963474,90.16663424973474,90.17032338612557,90.17032338622558,90.17216710181636,90.17216710191636,90.174009604985,90.174009605085,90.17769222416935,90.17769222426935,90.18137207728758,90.18137207738758,90.18688859831167,90.18688859841167,90.19056333610212,90.19056333620212,90.19423466367407,90.19423466377407,90.20340727996077,90.20340728006077,90.20890712880549,90.2089071289055,90.21257102967937,90.21257102977937,90.21623201292184,90.21623201302184,90.22354984925326,90.22354984935326,90.22903484470181,90.22903484480182,90.23086200188717,90.23086200198718,90.2345154068663,90.2345154069663,90.23816627312719,90.2381662732272,90.2436396359415,90.24363963604151,90.24728678885336,90.24728678895336,90.25093121359042,90.25093121369042,90.25275238394772,90.25275238404772,90.25821430358437,90.25821430368437,90.26367493491624,90.26367493501624,90.26731276656419,90.26731276666419,90.27458532277208,90.27458532287208,90.2782194789623,90.2782194790623,90.28003481405683,90.28003481415683,90.28184942921635,90.28184942931635,90.28366328654953,90.28366328664953,90.28910307765719,90.28910307775719,90.29272719264846,90.29272719274846,90.29997110302088,90.29997110312088,90.30359002690176,90.30359002700176,90.30539865523323,90.30539865533324,90.30720652573837,90.30720652583837,90.31082101633517,90.31082101643517,90.31262714383973,90.31262714393974,90.31623837578331,90.31623837588332,90.31804291185257,90.31804291195257,90.32164933159892,90.32164933169892,90.32345030588439,90.3234503059844,90.32885190254477,90.32885190264477,90.33244979674484,90.33244979684484,90.33424671665944,90.33424671675944,90.34143250175195,90.34143250185195,90.3450237649716,90.3450237650716,90.350408499996,90.35040850009601,90.35399578462739,90.35399578472739,90.35758098523635,90.35758098533636,90.36295651267078,90.36295651277078,90.3719111781035,90.3719111782035,90.37370059553737,90.37370059563737,90.3790659302076,90.3790659303076,90.38442573274558,90.38442573284559,90.38978299656534,90.38978299666535,90.3933521691473,90.39335216924731,90.39691975029397,90.39691975039398,90.4040496835856,90.4040496836856,90.40760945912102,90.40760945922102,90.4093878122904,90.4093878123904,90.4111643845679,90.4111643846679,90.41471593768757,90.41471593778758,90.42004099388808,90.42004099398808,90.42358906100662,90.42358906110663,90.42536199571771,90.42536199581771,90.42890703153091,90.42890703163091,90.43245036223485,90.43245036233485,90.43599221517744,90.43599221527744,90.43953277981527,90.43953277991527,90.44661174928585,90.44661174938585,90.45368670227685,90.45368670237686,90.45722135586927,90.45722135596927,90.46428774542268,90.46428774552268,90.47135008060522,90.47135008070522,90.4801710654724,90.4801710655724,90.49074876756711,90.49074876766711,90.49250984229606,90.49250984239606,90.49603112025365,90.49603112035365,90.49779041409069,90.49779041419069,90.50482542963384,90.50482542973384,90.50834068287206,90.50834068297206,90.51185354895733,90.51185354905734,90.51536353530254,90.51536353540254,90.51887041455976,90.51887041465976,90.52938574754708,90.52938574764708,90.53288849665077,90.53288849675077,90.53813833858753,90.53813833868753,90.54163377466709,90.54163377476709,90.54338033702172,90.54338033712172,90.54687259023066,90.54687259033066,90.55210924598919,90.55210924608919,90.55734355248606,90.55734355258606,90.56257513080814,90.56257513090814,90.56780534504281,90.56780534514282,90.57129047468422,90.57129047478422,90.57651591461297,90.57651591471297,90.58173964943248,90.58173964953248,90.58521970163744,90.58521970173744,90.58869736668944,90.58869736678945,90.59217207621873,90.59217207631873,90.59564432281245,90.59564432291245,90.60258570891187,90.60258570901188,90.6077877698982,90.6077877699982,90.6112536507507,90.6112536508507,90.61644983858291,90.61644983868291,90.61818078971503,90.61818078981503,90.62337159698028,90.62337159708028,90.62510050198128,90.62510050208128,90.63028585289692,90.63028585299692,90.63374093472413,90.63374093482413,90.63719465246395,90.63719465256395,90.64409954922534,90.64409954932535,90.64582458931204,90.64582458941204,90.64927338118068,90.64927338128068,90.65272107420112,90.65272107430113,90.65789051076372,90.65789051086372,90.65961172382742,90.65961172392743,90.66305183858452,90.66305183868452,90.66820717957728,90.66820717967728,90.67164183798477,90.67164183808477,90.67507369243481,90.67507369253481,90.67850141673132,90.67850141683132,90.68363705423937,90.68363705433937,90.69390306236242,90.69390306246243,90.69903396345588,90.69903396355588,90.70245270751032,90.70245270761032,90.7075789100805,90.7075789101805,90.7127015508669,90.7127015509669,90.71611449754987,90.71611449764987,90.71781987204318,90.71781987214318,90.72122940850765,90.72122940860766,90.7246373156455,90.7246373157455,90.73144915133295,90.73144915143295,90.73485198103435,90.73485198113436,90.7399518870307,90.7399518871307,90.74334774472982,90.74334774482982,90.74674132894994,90.74674132904994,90.7484364348964,90.7484364349964,90.75013078301653,90.75013078311653,90.75182448698428,90.75182448708428,90.75690431058273,90.75690431068273,90.7602891798,90.7602891799,90.76536475957099,90.76536475967099,90.76874701428738,90.76874701438739,90.77212718498137,90.77212718508137,90.77381531892556,90.77381531902556,90.77718980592202,90.77718980602202,90.78056084480868,90.78056084490868,90.78392870082472,90.78392870092472,90.78897775667396,90.78897775677396,90.7923412930799,90.7923412931799,90.79402215189126,90.79402215199126,90.79738265699184,90.79738265709184,90.80242227790323,90.80242227800323,90.80410128004007,90.80410128014007,90.8057795622419,90.8057795623419,90.81081274162942,90.81081274172942,90.81416528955349,90.81416528965349,90.81919183796059,90.81919183806059,90.82254146825328,90.82254146835328,90.82421482458392,90.82421482468392,90.82756024894043,90.82756024904043,90.83090408186165,90.83090408196165,90.83591725463393,90.83591725473393,90.84092591833952,90.84092591843952,90.8475980866334,90.8475980867334,90.85093180257304,90.85093180267305,90.85259720172716,90.85259720182717,90.8559270527525,90.85592705285251,90.85759017842761,90.85759017852762,90.86257841871348,90.86257841881348,90.86423987717066,90.86423987727066,90.86590053991019,90.86590054001019,90.86755930808388,90.86755930818389,90.86921705319202,90.86921705329202,90.87418828027666,90.87418828037666,90.87750013292653,90.87750013302653,90.88081005311925,90.88081005321925,90.88411800296349,90.88411800306349,90.88577044328729,90.88577044338729,90.88907384617352,90.88907384627352,90.89567743118407,90.89567743128407,90.89732691598516,90.89732691608516,90.90227411997496,90.90227412007496,90.90721923994235,90.90721924004235,90.91216231377864,90.91216231387864,90.91380869149175,90.91380869159175,90.91545446294379,90.91545446304379,90.9187451343476,90.9187451344476,90.92532355952382,90.92532355962382,90.9269669059316,90.9269669060316,90.93025177996392,90.93025178006393,90.93517583235919,90.9351758324592,90.93681542752661,90.93681542762661,90.94173300050672,90.94173300060672,90.94500965212332,90.94500965222332,90.94664674646383,90.94664674656383,90.95155643805005,90.95155643815005,90.95646313622224,90.95646313632224,90.95809761606189,90.9580976161619,90.96136547689301,90.96136547699301,90.9629977590363,90.9629977591363,90.96626050453965,90.96626050463965,90.96788941435574,90.96788941445574,90.97114624881367,90.97114624891367,90.97602854497788,90.97602854507788,90.97765514342365,90.97765514352365,90.97928090826044,90.97928090836044,90.98253141486848,90.98253141496848,90.98415547459602,90.98415547469602,90.9874024573116,90.9874024574116,90.98902522873438,90.98902522883438,90.99064709076555,90.99064709086555,90.99713230330255,90.99713230340255,90.99875211920262,90.99875211930262,91.00361061961989,91.0036106197199,91.00684782637576,91.00684782647576,91.0117015145958,91.0117015146958,91.0149353869158,91.0149353870158,91.01978308830779,91.01978308840779,91.02139646268401,91.02139646278401,91.02462222626221,91.02462222636221,91.02784643629643,91.02784643639643,91.03106848652558,91.03106848662559,91.03912069446709,91.03912069456709,91.04233937236904,91.04233937246904,91.04394747985224,91.04394747995224,91.04877043821439,91.04877043831439,91.05037619643595,91.05037619653595,91.05519214490451,91.05519214500451,91.05840112262938,91.05840112272938,91.06000479682851,91.06000479692851,91.0616077132013,91.0616077133013,91.06481256077265,91.06481256087265,91.06801630949582,91.06801630959582,91.07121850467499,91.07121850477499,91.07441922209281,91.07441922219282,91.07761751446637,91.07761751456637,91.08081315444774,91.08081315454774,91.08560413253856,91.08560413263857,91.0887958697143,91.0887958698143,91.09198601545475,91.09198601555475,91.09517460765122,91.09517460775122,91.09836172208632,91.09836172218633,91.10473295754251,91.10473295764251,91.10950957482427,91.10950957492427,91.11110089645415,91.11110089655415,91.11428221351778,91.11428221361778,91.11587217106025,91.11587217116025,91.11746152234164,91.11746152244164,91.12381744970585,91.12381744980586,91.1269931209633,91.12699312106331,91.13175344497887,91.13175344507887,91.13651172286332,91.13651172296332,91.13968265770617,91.13968265780618,91.14602168554312,91.14602168564312,91.149189513298,91.149189513398,91.15393968244067,91.15393968254067,91.1555198640234,91.1555198641234,91.16025949938002,91.16025949948002,91.16499564873548,91.16499564883549,91.16972884256825,91.16972884266825,91.17288272631856,91.17288272641856,91.17603498074224,91.17603498084225,91.17918481012167,91.17918481022167,91.18075722398447,91.18075722408447,91.19018863796462,91.19018863806463,91.19175904358764,91.19175904368764,91.1964684795648,91.1964684796648,91.19960557746178,91.19960557756178,91.20117291388814,91.20117291398815,91.20430641211003,91.20430641221003,91.21213597067427,91.21213597077427,91.21839186496224,91.21839186506224,91.21995480599585,91.21995480609586,91.22151695131183,91.22151695141183,91.22620172004177,91.22620172014177,91.22932199419411,91.22932199429411,91.23088076718288,91.23088076728288,91.2339973658775,91.2339973659775,91.23711191844102,91.23711191854102,91.24022469011265,91.24022469021266,91.24489021005368,91.24489021015368,91.25110377974084,91.25110377984085,91.25420707858328,91.25420707868328,91.25885857873712,91.25885857883712,91.26815672895624,91.26815672905624,91.2697050818329,91.2697050819329,91.2727994762159,91.2727994763159,91.27898246761043,91.27898246771043,91.2805271829207,91.2805271830207,91.28515928272034,91.28515928282035,91.28670214135607,91.28670214145608,91.28978668399672,91.28978668409673,91.29132757228398,91.29132757238398,91.29440771953188,91.29440771963188,91.29594591753565,91.29594591763565,91.29748256199544,91.29748256209544,91.30055497941471,91.30055497951471,91.3051613510103,91.3051613511103,91.31130125997768,91.31130126007768,91.31590357720236,91.31590357730236,91.32356986208268,91.32356986218268,91.32663439810808,91.32663439820809,91.32816539676166,91.32816539686166,91.33122587841618,91.33122587851618,91.33581410007102,91.33581410017102,91.33734142326688,91.33734142336688,91.34039512237568,91.34039512247568,91.34192108148414,91.34192108158415,91.34497216609209,91.3449721661921,91.34802150769947,91.34802150779947,91.35106933365417,91.35106933375417,91.35411503769514,91.35411503779514,91.35563663930216,91.35563663940216,91.36324298011938,91.36324298021938,91.3678039958088,91.3678039959088,91.37084250049959,91.3708425005996,91.37387937586375,91.37387937596375,91.37691435666207,91.37691435676207,91.37994770813378,91.37994770823379,91.38904495859146,91.38904495869146,91.39207543032309,91.39207543042309,91.396619281246,91.396619281346,91.40116029032018,91.40116029042018,91.40418625298511,91.40418625308511,91.40872226040547,91.40872226050547,91.41174329719922,91.41174329729922,91.4162732420089,91.4162732421089,91.41929139906259,91.41929139916259,91.4207993976869,91.42079939778691,91.42532184001585,91.42532184011586,91.42833234302356,91.42833234312356,91.43284163706561,91.43284163716561,91.4358458122234,91.43584581232341,91.4373469904107,91.4373469905107,91.43884744866298,91.43884744876298,91.44034692174102,91.44034692184103,91.441845788558,91.441845788658,91.44334374598338,91.44334374608339,91.44633818307277,91.44633818317277,91.44933091505291,91.44933091515291,91.45531380240365,91.45531380250365,91.46129343110115,91.46129343120116,91.46278718259038,91.46278718269038,91.46427972155745,91.46427972165745,91.46875514076231,91.46875514086231,91.4702454441417,91.4702454442417,91.47471510386642,91.47471510396642,91.47620233804915,91.47620233814915,91.47768854916632,91.47768854926632,91.48065968309591,91.48065968319591,91.48214392386463,91.48214392396463,91.48511157179307,91.48511157189307,91.48955933033669,91.48955933043669,91.492521218785,91.492521218885,91.49548125055877,91.49548125065877,91.49696003497787,91.49696003507788,91.49843741741824,91.49843741751825,91.502867367043,91.502867367143,91.50581807755285,91.50581807765285,91.50876712084477,91.50876712094477,91.51024007000109,91.51024007010109,91.51318460422631,91.51318460432631,91.51612735755965,91.51612735765966,91.51906772374004,91.51906772384004,91.52053620172097,91.52053620182097,91.5220037703103,91.52200377041031,91.52787226377575,91.52787226387575,91.53227072571627,91.53227072581628,91.53373579347871,91.53373579357871,91.5381287611783,91.5381287612783,91.54398126450808,91.54398126460808,91.54544174742118,91.54544174752118,91.54982149105123,91.54982149115123,91.55273912244157,91.55273912254157,91.55565379830918,91.55565379840918,91.55711018896007,91.55711018906007,91.5585659733499,91.5585659734499,91.56147678430321,91.56147678440321,91.56438604171252,91.56438604181253,91.56874805220633,91.56874805230633,91.57165416463637,91.57165416473637,91.57746434336534,91.57746434346534,91.57891592181899,91.57891592191899,91.58326880050546,91.58326880060547,91.58761853421264,91.58761853431264,91.59196580498423,91.59196580508423,91.59775961466434,91.59775961476434,91.6021027552824,91.6021027553824,91.60644305405171,91.60644305415171,91.61078126879859,91.61078126889859,91.61945311344303,91.61945311354303,91.62523199394435,91.62523199404436,91.62956365349343,91.62956365359344,91.63244930461242,91.63244930471242,91.6338907281932,91.6338907282932,91.63533150762157,91.63533150772157,91.63821192973884,91.63821192983885,91.6439709930815,91.6439709931815,91.64684834600212,91.64684834610212,91.64828579099463,91.64828579109464,91.65259657242818,91.65259657252818,91.65546846899917,91.65546846909918,91.6612086624661,91.6612086625661,91.66407733827516,91.66407733837517,91.66694491523606,91.66694491533606,91.66837751014002,91.66837751024002,91.67267396865584,91.67267396875584,91.67553643028897,91.67553643038897,91.67696648647471,91.67696648657471,91.68125555618376,91.68125555628376,91.6841135087502,91.6841135088502,91.68982577631665,91.68982577641665,91.69410673728392,91.69410673738392,91.69553239807692,91.69553239817692,91.69838288605395,91.69838288615395,91.70408086859399,91.70408086869399,91.70692680961301,91.70692680971301,91.71119276561883,91.71119276571883,91.71829871370696,91.71829871380696,91.72397968304577,91.72397968314577,91.72539852340176,91.72539852350177,91.72965383194757,91.72965383204757,91.7310705124985,91.7310705125985,91.73248651100573,91.73248651110573,91.73531687869357,91.73531687879357,91.73814535181556,91.73814535191556,91.74097257452411,91.74097257462411,91.74238535226941,91.74238535236941,91.74379699327521,91.74379699337521,91.74944185218916,91.74944185228917,91.75226066302538,91.75226066312538,91.75789400297904,91.75789400307904,91.75930079389629,91.75930079399629,91.76211339055656,91.76211339065657,91.76492378952047,91.76492378962047,91.76913730394391,91.76913730404391,91.77615621567743,91.77615621577743,91.77755868698458,91.77755868708458,91.78036287177252,91.78036287187253,91.78316470729885,91.78316470739885,91.78456384417012,91.78456384427012,91.78875973913125,91.78875973923125,91.7915555120469,91.7915555121469,91.79435003454907,91.79435003464907,91.79854009424744,91.79854009434744,91.80133177490086,91.80133177500086,91.80412114418397,91.80412114428397,91.80969696511877,91.80969696521878,91.81248250737889,91.81248250747889,91.81526573826868,91.81526573836868,91.81665646326763,91.81665646336764,91.8180463167663,91.8180463168663,91.8222145510662,91.8222145511662,91.82499069627976,91.82499069637976,91.82915233749056,91.82915233759056,91.83469716332826,91.83469716342826,91.8430015758741,91.8430015759741,91.8471510160809,91.8471510161809,91.84853295029436,91.84853295039436,91.84991420246412,91.84991420256412,91.85129454524228,91.85129454534228,91.8526741301941,91.8526741302941,91.859569516235,91.859569516335,91.86783899298679,91.86783899308679,91.87334895881308,91.87334895891308,91.87610229345394,91.87610229355394,91.88022971880571,91.88022971890571,91.88297987057597,91.88297987067597,91.88435411285212,91.88435411295212,91.8857277109759,91.8857277110759,91.88710021025149,91.88710021035149,91.8925882748967,91.8925882749967,91.89807186836653,91.89807186846653,91.89944110898887,91.89944110908887,91.90080959178489,91.9008095918849,91.90764999752517,91.90764999762517,91.90901647208139,91.90901647218139,91.9117486633675,91.9117486634675,91.91584441157846,91.91584441167846,91.92130068540891,91.92130068550891,91.92538881746519,91.92538881756519,91.92947422134667,91.92947422144667,91.93355674548806,91.93355674558806,91.94035827473735,91.94035827483735,91.94443674450787,91.94443674460787,91.94851305447332,91.94851305457333,91.95394500207841,91.95394500217841,91.96073129901836,91.96073129911836,91.96344393838503,91.96344393848503,91.96886739833516,91.96886739843517,91.97157647591806,91.97157647601806,91.97698909895159,91.9769890990516,91.98375175170989,91.98375175180989,91.98645514559527,91.98645514569527,91.99050773559644,91.99050773569644,91.99860772448837,91.99860772458837,92.00130619250257,92.00130619260257,92.00265421398754,92.00265421408754,92.004001098733,92.004001098833,92.00669388304968,92.00669388314968,92.00803902479457,92.00803902489457,92.0093835602784,92.0093835603784,92.01341417331584,92.01341417341584,92.01475552592906,92.01475552602906,92.01877886383369,92.01877886393369,92.0214585756461,92.0214585757461,92.02547399426552,92.02547399436553,92.02814779503251,92.02814779513251,92.02948361551348,92.02948361561349,92.03081856449417,92.03081856459417,92.0334867952376,92.0334867953376,92.03748660263452,92.03748660273452,92.04148485648744,92.04148485658745,92.04281666048884,92.04281666058884,92.04814243662437,92.04814243672438,92.05213451419267,92.05213451429267,92.0534641204977,92.0534641205977,92.06010893126087,92.06010893136087,92.06143577149977,92.06143577159978,92.06408850469465,92.06408850479465,92.06806504682308,92.06806504692308,92.06938934834376,92.06938934844376,92.07336035833994,92.07336035843994,92.07600532381488,92.07600532391488,92.07997095324409,92.07997095334409,92.08657639492915,92.08657639502916,92.08921673766345,92.08921673776345,92.09317528141642,92.09317528151642,92.0971315895817,92.09713158968171,92.09976757481458,92.09976757491458,92.10503693077946,92.10503693087946,92.1076701499462,92.1076701500462,92.11161793256521,92.11161793266521,92.11424849933978,92.11424849943978,92.1155629491181,92.1155629492181,92.12344763954822,92.12344763964822,92.12607430351717,92.12607430361717,92.13001261330699,92.13001261340699,92.13526090169975,92.13526090179975,92.14050570409138,92.14050570419138,92.14181545534642,92.14181545544642,92.14836227916446,92.14836227926446,92.15490565487268,92.15490565497268,92.15882717880918,92.15882717890918,92.16404939797602,92.16404939807602,92.16665628267761,92.16665628277761,92.17186781649312,92.17186781659312,92.17447185934597,92.17447185944597,92.17707343926322,92.17707343936323,92.1796735035278,92.1796735036278,92.18617038659036,92.18617038669036,92.1887674195496,92.18876741964961,92.19395974246751,92.19395974256751,92.19785144593773,92.19785144603773,92.20044412139553,92.20044412149554,92.20433165682091,92.20433165692091,92.20821672931069,92.2082167294107,92.21080406209283,92.21080406219284,92.21597467328625,92.21597467338626,92.21855806537145,92.21855806547146,92.22243120209646,92.22243120219646,92.22372132564942,92.22372132574942,92.22759048378616,92.22759048388616,92.22887878855592,92.22887878865592,92.23145410979066,92.23145410989066,92.23274093679906,92.23274093689906,92.23531356775031,92.23531356785031,92.2430272546679,92.2430272547679,92.24687943749473,92.24687943759473,92.25072783118989,92.2507278312899,92.25457164003572,92.25457164013572,92.2596926514955,92.2596926515955,92.26353047351328,92.26353047361329,92.26608681119832,92.26608681129832,92.26991834325753,92.26991834335753,92.27502457710376,92.27502457720377,92.27757526898262,92.27757526908262,92.28012349792587,92.28012349802587,92.28649094425039,92.28649094435039,92.2903080397179,92.2903080398179,92.29411956516185,92.29411956526185,92.29665741188433,92.29665741198433,92.30046048756464,92.30046048766464,92.30172624700093,92.30172624710093,92.30425681859057,92.30425681869058,92.3067859882015,92.3067859883015,92.3118423949662,92.31184239506621,92.3168965282519,92.3168965283519,92.3194219845138,92.3194219846138,92.32194581144905,92.32194581154906,92.32572963834046,92.32572963844046,92.32825058553566,92.32825058563566,92.33077016864344,92.33077016874344,92.33328827398988,92.33328827408988,92.33832061976844,92.33832061986844,92.34083527700506,92.34083527710506,92.34711936243271,92.34711936253271,92.35214310688237,92.35214310698237,92.35465261089989,92.35465261099989,92.35715976565578,92.35715976575578,92.35841252837042,92.35841252847042,92.36091634869044,92.36091634879044,92.36466982463712,92.36466982473712,92.3696714405578,92.3696714406578,92.37217097915902,92.37217097925902,92.37716710083876,92.37716710093876,92.38215867556049,92.38215867566049,92.3834038979031,92.3834038980031,92.38713785982165,92.38713785992165,92.38838111181578,92.38838111191578,92.39210882166707,92.39210882176707,92.3933498001822,92.3933498002822,92.39582993842923,92.39582993852923,92.40326591988631,92.40326591998631,92.40698123927702,92.40698123937702,92.41069383049292,92.41069383059292,92.41193022415871,92.41193022425871,92.41563785161212,92.41563785171212,92.41934320558651,92.41934320568652,92.42057653005565,92.42057653015566,92.42304173912389,92.42304173922389,92.42427320691851,92.42427320701852,92.42673542257273,92.42673542267273,92.43411809094712,92.43411809104713,92.43903680066674,92.43903680076674,92.44395236540706,92.44395236550706,92.44640829110268,92.44640829120269,92.45131703540599,92.451317035506,92.4562214979905,92.4562214980905,92.45989646312886,92.45989646322886,92.46601640337231,92.46601640347231,92.4672394592946,92.4672394593946,92.47335220018788,92.47335220028788,92.47823809602717,92.47823809612717,92.48067909254401,92.48067909264401,92.48677792732403,92.48677792742403,92.49287164677628,92.49287164687628,92.49530749007404,92.49530749017404,92.49774234819756,92.49774234829756,92.50017553910314,92.50017553920314,92.5111219237099,92.5111219238099,92.51476831879542,92.51476831889542,92.51962662975612,92.51962662985612,92.52205402329024,92.52205402339024,92.52569310535162,92.52569310545162,92.52811712655856,92.52811712665856,92.53175105540086,92.53175105550086,92.53538236974232,92.53538236984232,92.53901163795267,92.53901163805267,92.5450578038024,92.5450578039024,92.54868316920712,92.54868316930713,92.55351397107185,92.55351397117185,92.55592738271008,92.55592738281008,92.55954509406882,92.55954509416883,92.56316034249197,92.56316034259197,92.56556784308479,92.5655678431848,92.57158195288055,92.57158195298055,92.57638933791152,92.57638933801152,92.58359668316328,92.58359668326328,92.58599709807987,92.58599709817987,92.59199430834835,92.59199430844835,92.59319254545817,92.59319254555817,92.60037958096414,92.60037958106415,92.60516874027175,92.60516874037175,92.60875844994742,92.60875845004742,92.61114980883929,92.61114980893929,92.61712519370931,92.61712519380931,92.62428922918602,92.62428922928602,92.63144909661789,92.63144909671789,92.63264115744308,92.63264115754308,92.6385991501987,92.6385991502987,92.64336288443278,92.64336288453278,92.64574363375597,92.64574363385597,92.65169196422582,92.65169196432582,92.65525897700275,92.65525897710275,92.65882295847432,92.65882295857432,92.66357229400802,92.66357229410802,92.6659448588068,92.6659448589068,92.6730584988322,92.6730584989322,92.67661331060513,92.67661331070514,92.6825336015006,92.6825336016006,92.68489980055814,92.68489980065814,92.69081092175495,92.69081092185495,92.69553521112827,92.69553521122828,92.7002559008265,92.7002559009265,92.703794154094,92.703794154194,92.7073296791867,92.7073296792867,92.70850696239835,92.70850696249835,92.7120371827067,92.7120371828067,92.71438936197706,92.71438936207706,92.7202671388151,92.7202671389151,92.72144108759089,92.72144108769089,92.72965573560732,92.72965573570733,92.7366937445645,92.7366937446645,92.74020865678088,92.74020865688088,92.74723219125505,92.74723219135505,92.74957167693428,92.74957167703428,92.75307821516967,92.75307821526967,92.75891738075593,92.75891738085593,92.76241774270667,92.76241774280668,92.76475010481838,92.76475010491838,92.77057600844378,92.77057600854378,92.77755994649195,92.77755994659195,92.78104894104766,92.78104894114766,92.78337353543941,92.78337353553941,92.78685877875476,92.78685877885476,92.78918026605855,92.78918026615855,92.7903403466124,92.7903403467124,92.79265956043717,92.79265956053717,92.7984549615526,92.7984549616526,92.80077201557232,92.80077201567232,92.801929443734,92.801929443834,92.80424323910047,92.80424323920047,92.81002401416762,92.81002401426763,92.81233474033745,92.81233474043745,92.81695335082834,92.81695335092834,92.82041449526623,92.82041449536624,92.82156722490464,92.82156722500464,92.82502408762373,92.82502408772373,92.82847761590696,92.82847761600696,92.83192803710222,92.83192803720222,92.83307656080446,92.83307656090446,92.83766849580834,92.83766849590835,92.84340565796991,92.84340565806991,92.84684448442225,92.84684448452225,92.85028031746054,92.85028031756055,92.85714633773095,92.85714633783095,92.86057607026726,92.86057607036726,92.86400220312846,92.86400220322847,92.86970658930105,92.86970658940105,92.87426559675067,92.87426559685068,92.8788203224815,92.8788203225815,92.88223421644742,92.88223421654742,92.8845075817789,92.8845075818789,92.88791344278566,92.88791344288566,92.89018214748519,92.8901821475852,92.89244971544522,92.89244971554523,92.89811579349656,92.89811579359656,92.90264604143677,92.90264604153677,92.90717128772317,92.90717128782318,92.91056278792087,92.91056278802087,92.91395148416112,92.91395148426112,92.91959638096638,92.91959638106638,92.92297999977019,92.9229799998702,92.92636096618183,92.92636096628183,92.92973912863603,92.92973912873603,92.93311456291542,92.93311456301542,92.93648783738976,92.93648783748976,92.93985868681982,92.93985868691982,92.94322586079217,92.94322586089217,92.9488358596947,92.9488358597947,92.95107796468987,92.95107796478987,92.95331897083686,92.95331897093686,92.95779867176051,92.95779867186052,92.96003641925425,92.96003641935425,92.9622716659211,92.9622716660211,92.96450391917392,92.96450391927392,92.96561913640872,92.96561913650872,92.97453746406867,92.97453746416868,92.97899329346276,92.97899329356277,92.98233314779573,92.98233314789573,92.9856705391931,92.9856705392931,92.98900455826326,92.98900455836326,92.99455753073407,92.99455753083407,92.99788647236781,92.99788647246781,93.00121416358809,93.00121416368809,93.00232248460324,93.00232248470324,93.00453817935063,93.00453817945063,93.00564479525653,93.00564479535653,93.01006921274904,93.01006921284905,93.01228028475579,93.01228028485579,93.01890971164437,93.01890971174437,93.02442968077828,93.02442968087828,93.02883947222251,93.02883947232252,93.03544976399613,93.03544976409613,93.04095578912548,93.04095578922548,93.04535675189291,93.04535675199291,93.04975396341999,93.04975396351999,93.0519491210737,93.0519491211737,93.05633723868475,93.05633723878475,93.05962662178379,93.0596266218838,93.0629136935125,93.0629136936125,93.06510396318635,93.06510396328635,93.07276585267392,93.07276585277393,93.07932699940972,93.07932699950972,93.0815121537558,93.0815121538558,93.08369571666658,93.08369571676658,93.0902422752454,93.0902422753454,93.09351282636001,93.09351282646001,93.09569184231279,93.09569184241279,93.10113572568265,93.10113572578265,93.10875108463325,93.10875108473326,93.11201030624416,93.11201030634416,93.11526710281078,93.11526710291078,93.11635159018769,93.11635159028769,93.12177243563697,93.12177243573697,93.12393921269444,93.12393921279444,93.12827079646088,93.12827079656088,93.13259946259595,93.13259946269595,93.13692460483854,93.13692460493854,93.1412469431237,93.1412469432237,93.14448646124988,93.14448646134989,93.14772416059286,93.14772416069286,93.15203793544043,93.15203793554043,93.15634830006951,93.15634830016951,93.1628100199901,93.1628100200901,93.16496251202115,93.16496251212115,93.16926476790843,93.16926476800843,93.17248837168158,93.17248837178158,93.17571064925863,93.17571064935863,93.17893057757405,93.17893057767405,93.18322135239235,93.18322135249235,93.18643734001083,93.18643734011083,93.18965086469372,93.18965086479372,93.19821668943386,93.19821668953387,93.20249674100953,93.20249674110953,93.20998012450386,93.20998012460386,93.21532090560083,93.21532090570084,93.21745562660432,93.21745562670432,93.22385687198337,93.22385687208337,93.23131922579692,93.23131922589693,93.23664723752013,93.23664723762013,93.24090690356738,93.24090690366738,93.24197089174193,93.24197089184193,93.2440978829168,93.2440978830168,93.246223358439,93.246223358539,93.248347735113,93.248347735213,93.25365581600362,93.25365581610362,93.25683808035016,93.25683808045017,93.26107861128246,93.26107861138246,93.26425727595392,93.26425727605393,93.26849208529737,93.26849208539737,93.27166582409767,93.27166582419767,93.27800973991447,93.27800974001447,93.28329266097076,93.28329266107076,93.28857145187351,93.28857145197351,93.29173586940988,93.29173586950988,93.29595214989943,93.29595214999944,93.29805883132852,93.29805883142852,93.30437648846281,93.30437648856281,93.30542831352469,93.30542831362469,93.30963409811949,93.30963409821949,93.31068433174606,93.31068433184606,93.31488352325177,93.31488352335177,93.32117927920497,93.32117927930497,93.32642237648734,93.32642237658735,93.32956477916868,93.32956477926868,93.3347980626,93.3347980627,93.33584355981198,93.33584355991198,93.337933606953,93.337933607053,93.34106638523987,93.34106638533987,93.35045831646796,93.35045831656797,93.35567083545772,93.35567083555772,93.35671220251616,93.35671220261617,93.35879406513277,93.35879406523277,93.36087429842276,93.36087429852276,93.36295290238613,93.36295290248613,93.367107495812,93.367107495912,93.37022079796206,93.37022079806206,93.37437038973412,93.37437038983413,93.37748047112227,93.37748047122227,93.37955270934441,93.37955270944441,93.38162354558784,93.38162354568784,93.38576286881306,93.38576286891306,93.38679669549948,93.38679669559949,93.39093109285352,93.39093109295352,93.39506283781539,93.39506283791539,93.39815893730764,93.39815893740764,93.40331537714859,93.40331537724859,93.40846711846626,93.40846711856626,93.41258673820677,93.41258673830677,93.41361571480464,93.41361571490464,93.41773037078265,93.41773037088265,93.4218421849119,93.4218421850119,93.42800539703909,93.42800539713909,93.4300578559939,93.4300578560939,93.4331350856104,93.4331350857104,93.43518466482513,93.43518466492513,93.44133120477295,93.44133120487295,93.44542661196205,93.44542661206205,93.4474728567409,93.4474728568409,93.45360765038053,93.45360765048054,93.45565006813639,93.4556500682364,93.45973225125594,93.45973225135594,93.46279250556255,93.46279250566255,93.46585044849883,93.46585044859883,93.47399905195016,93.47399905205016,93.47603449770374,93.47603449780374,93.47908508972456,93.47908508982457,93.48314916079471,93.48314916089471,93.4892411372464,93.4892411373464,93.49329941094507,93.49329941104507,93.49634034068013,93.49634034078014,93.50039205918101,93.50039205928101,93.50241568284376,93.50241568294376,93.5094943112925,93.5094943113925,93.51656843067454,93.51656843077454,93.51959818247116,93.51959818257116,93.52767232948506,93.52767232958506,93.53069840582394,93.53069840592394,93.53473004192693,93.53473004202694,93.53875902563776,93.53875902573776,93.54479852261574,93.54479852271574,93.5478141409512,93.5478141410512,93.55082710689449,93.55082710699449,93.55785280117372,93.55785280127373,93.56387134425353,93.56387134435353,93.56587564334473,93.56587564344473,93.57288799988046,93.57288799998047,93.57589039414637,93.57589039424637,93.57889108330303,93.57889108340304,93.58089000182726,93.58089000192726,93.58188858958908,93.58188858968909,93.58887586207311,93.58887586217311,93.59386353398922,93.59386353408922,93.60084023479588,93.60084023489588,93.60382777566562,93.60382777576562,93.60681209577345,93.60681209587345,93.61078837266265,93.61078837276266,93.6157553180285,93.6157553181285,93.61972557019836,93.61972557029836,93.62567499951639,93.6256749996164,93.63062724305134,93.63062724315134,93.63458559734772,93.63458559744772,93.64051933966063,93.64051933976063,93.64348316056608,93.64348316066608,93.64940636909293,93.64940636919293,93.65335184034161,93.65335184044162,93.65433716614258,93.65433716624258,93.65630679467895,93.65630679477896,93.66122781576888,93.66122781586888,93.66417845049611,93.66417845059611,93.66614379731368,93.66614379741368,93.67400131966969,93.67400131976969,93.67989004709828,93.67989004719828,93.68185084695786,93.68185084705786,93.68772809331747,93.68772809341748,93.69262206000724,93.69262206010724,93.69457793399563,93.69457793409563,93.69848684012365,93.69848684022365,93.70239358644662,93.70239358654662,93.70532190318826,93.70532190328827,93.7121504489464,93.7121504490464,93.71507399138214,93.71507399148214,93.71896993867983,93.71896993877984,93.72286232419374,93.72286232429374,93.7257795766709,93.7257795767709,93.72966665740047,93.72966665750047,93.73063747082712,93.73063747092712,93.73160748853611,93.73160748863612,93.73936600088142,93.73936600098142,93.74324273728152,93.74324273738152,93.75195770224443,93.75195770234443,93.75679615815514,93.75679615825514,93.76259686405491,93.76259686415492,93.76549539822159,93.76549539832159,93.76839165890927,93.76839165900927,93.77514552088208,93.77514552098208,93.77996628154786,93.77996628164786,93.78478302573407,93.78478302583407,93.78574554096235,93.78574554106235,93.78959435146199,93.78959435156199,93.79344043378683,93.79344043388683,93.7972836742629,93.7972836743629,93.80208439042212,93.80208439052213,93.80496250116909,93.80496250126909,93.80783909626336,93.80783909636337,93.81167112090968,93.81167112100968,93.8183725791914,93.8183725792914,93.82124261908788,93.82124261918788,93.8288905645709,93.8288905646709,93.83653415255249,93.83653415265249,93.83939752357722,93.83939752367722,93.84512051438632,93.84512051448633,93.84988572638174,93.84988572648174,93.85274174649103,93.85274174659104,93.856545617889,93.856545617989,93.86034695056874,93.86034695066874,93.86319580921915,93.86319580931915,93.86794021888167,93.86794021898167,93.87173264710198,93.87173264720198,93.87931219875826,93.87931219885826,93.88309917062895,93.88309917072895,93.88688364167274,93.88688364177274,93.89066542243303,93.89066542253303,93.89538872663212,93.89538872673212,93.89822004160288,93.89822004170288,93.90104893152936,93.90104893162936,93.90481801869855,93.90481801879855,93.90764289214545,93.90764289224545,93.91234819796908,93.91234819806908,93.91422897136765,93.91422897146765,93.91892779777612,93.91892779787612,93.92644122908465,93.92644122918465,93.93301002986641,93.93301002996641,93.9367605881817,93.9367605882817,93.93957117660219,93.93957117670219,93.94425170150465,93.94425170160466,93.95079916947508,93.95079916957508,93.95547071413547,93.95547071423547,93.9592052823151,93.9592052824151,93.96480353490944,93.96480353500944,93.97039780891551,93.97039780901551,93.97505578848451,93.97505578858451,93.98063971816111,93.98063971826112,93.98808054970674,93.98808054980674,93.99086851701111,93.99086851711111,93.99458201761861,93.99458201771861,93.99922025581158,93.99922025591158,94.00570982585,94.00570982595,94.01126719371356,94.01126719381357,94.01404424831873,94.01404424841873,94.01866880774634,94.01866880784634,94.02421439141038,94.02421439151038,94.02698466346983,94.02698466356983,94.02883004637856,94.02883004647856,94.032518538717,94.032518538817,94.03804494937476,94.03804494947477,94.0417266212762,94.0417266213762,94.05092382902747,94.05092382912747,94.0546000824706,94.0546000825706,94.0601107303406,94.0601107304406,94.06469887621282,94.06469887631282,94.07111637696676,94.07111637706676,94.07661327028877,94.07661327038878,94.08302156345275,94.08302156355275,94.08850993122849,94.08850993132849,94.09125178480038,94.09125178490038,94.09581655173015,94.09581655183015,94.1012918470016,94.1012918471016,94.10402706959307,94.10402706969307,94.1122285693226,94.1122285694226,94.11678155205286,94.11678155215286,94.11860124464881,94.11860124474882,94.12678516280737,94.12678516290737,94.1304193568889,94.1304193569889,94.13404904190374,94.13404904200374,94.13676922164244,94.13676922174244,94.14039345030766,94.14039345040766,94.14220416266154,94.14220416276154,94.1458232381077,94.1458232382077,94.14943962327034,94.14943962337034,94.15576368403717,94.15576368413717,94.15666542159336,94.15666542169336,94.1620739523647,94.1620739524647,94.16657657753498,94.16657657763498,94.17197563547712,94.17197563557713,94.17647106129724,94.17647106139724,94.18006387806086,94.18006387816087,94.1836537771931,94.1836537772931,94.18813673677,94.18813673687,94.19261632401971,94.19261632411971,94.19709147798535,94.19709147808535,94.19888047861474,94.19888047871474,94.20156185764509,94.20156185774509,94.20692101603068,94.20692101613068,94.21227460439272,94.21227460449272,94.21762281218777,94.21762281228777,94.22118569481115,94.22118569491116,94.22652492236413,94.22652492246414,94.23008113611579,94.23008113621579,94.23541183812249,94.23541183822249,94.23807423360314,94.23807423370314,94.24073503764849,94.2407350377485,94.24428018713563,94.24428018723563,94.24870888634693,94.24870888644693,94.25756022215886,94.25756022225886,94.26198183569392,94.26198183579392,94.26905205227033,94.26905205237033,94.27170065531168,94.27170065541168,94.27699532267616,94.27699532277616,94.2840495491169,94.2840495492169,94.2866923926781,94.28669239277811,94.28933303854294,94.28933303864294,94.29548802825438,94.29548802835438,94.29900267523155,94.29900267533155,94.30690377260181,94.30690377270182,94.3130443636129,94.3130443637129,94.3183040194007,94.3183040195007,94.32180680639571,94.32180680649572,94.32530792617278,94.32530792627279,94.32880525681819,94.32880525691819,94.334047331035,94.334047331135,94.33841206970361,94.33841206980361,94.341901026368,94.34190102646801,94.3445157924635,94.3445157925635,94.34799906543037,94.34799906553037,94.35060928456785,94.35060928466785,94.35669546364807,94.35669546374807,94.36016968059025,94.36016968069025,94.36711239288577,94.36711239298577,94.37491580644145,94.37491580654145,94.37751408981414,94.37751408991414,94.38530443086553,94.38530443096553,94.39049254784729,94.39049254794729,94.39567551160997,94.39567551170997,94.40257669502232,94.40257669512232,94.40774867030314,94.40774867040314,94.41291632597384,94.41291632607384,94.418079131556,94.418079131656,94.42323833746306,94.42323833756306,94.42667545880614,94.42667545890615,94.43010981408308,94.43010981418308,94.43611600459363,94.43611600469363,94.43954490352097,94.43954490362097,94.44382866844568,94.44382866854568,94.44810845478212,94.44810845488212,94.4532405683977,94.4532405684977,94.4575134585145,94.4575134586145,94.46605169837605,94.46605169847605,94.47031708601212,94.47031708611212,94.4728746362193,94.4728746363193,94.47969101886476,94.47969101896476,94.48309680408887,94.48309680418888,94.48649974746422,94.48649974756422,94.49074857659488,94.49074857669488,94.49159734209024,94.49159734219025,94.49753695755724,94.49753695765725,94.50686132856757,94.50686132866757,94.51279185011855,94.51279185021855,94.51871782471152,94.51871782481152,94.52717501504652,94.52717501514653,94.53478150742902,94.53478150752902,94.5406921360387,94.5406921361387,94.54406590310015,94.54406590320015,94.54996520220612,94.54996520230613,94.5567031877172,94.5567031878172,94.56091056374733,94.56091056384733,94.56427334232693,94.56427334242693,94.565953329638,94.56595332973801,94.56847154865838,94.56847154875838,94.57518225242141,94.57518225252142,94.57769634128826,94.57769634138826,94.58272152560797,94.58272152570797,94.58857967474394,94.58857967484394,94.59610633012201,94.59610633022201,94.60028479507744,94.60028479517744,94.60445939511857,94.60445939521857,94.60863122909358,94.60863122919358,94.61280029700245,94.61280029710245,94.61696603047545,94.61696603057545,94.62029580571814,94.62029580581815,94.6269518323111,94.6269518324111,94.63193939055324,94.63193939065324,94.63609186206538,94.63609186216539,94.64273095123973,94.64273095133973,94.64770596745603,94.64770596755604,94.65019061476977,94.65019061486977,94.65350144435409,94.6535014444541,94.65846482793616,94.65846482803616,94.6634231719731,94.6634231720731,94.66837780266098,94.66837780276099,94.6758014315488,94.67580143164881,94.6782731200322,94.6782731201322,94.68156686063267,94.68156686073267,94.68568056932777,94.68568056942777,94.6930793794031,94.6930793795031,94.69554299703604,94.69554299713604,94.69882374091486,94.69882374101486,94.70701732135917,94.70701732145918,94.71274785254032,94.71274785264032,94.72011028695157,94.72011028705157,94.72746768181769,94.72746768191769,94.73155111535068,94.73155111545069,94.73481609644173,94.73481609654173,94.73644682504103,94.73644682514103,94.73807645479214,94.73807645489214,94.74458947955563,94.74458947965563,94.750284392899,94.750284392999,94.75190951358343,94.75190951368343,94.76002897861217,94.76002897871217,94.76570759868935,94.76570759878935,94.77300273812203,94.77300273822203,94.77948185015703,94.77948185025703,94.78433667511659,94.7843366752166,94.79161123956428,94.79161123966428,94.79403363144458,94.79403363154458,94.79968212792494,94.79968212802494,94.80451781776954,94.80451781786954,94.81015566678988,94.81015566688988,94.81498468776272,94.81498468786272,94.81820147109885,94.81820147119885,94.82061139673594,94.82061139683594,94.8254287092919,94.8254287093919,94.82943988408385,94.82943988418386,94.83505155020433,94.83505155030433,94.84385957624119,94.84385957634119,94.8502593438589,94.8502593439589,94.8542557410373,94.85425574113731,94.8606437244555,94.8606437245555,94.86463269493584,94.86463269503584,94.8694149580238,94.8694149581238,94.87498498158715,94.87498498168715,94.88134575903992,94.88134575913992,94.88690702970909,94.8869070298091,94.89563934889507,94.89563934899508,94.89960494043297,94.89960494053297,94.90594502922677,94.90594502932677,94.91148890778156,94.91148890788156,94.91702786046518,94.91702786056518,94.92414453179593,94.92414453189593,94.93204502290514,94.93204502300514,94.93678098280402,94.93678098290403,94.94624755992614,94.94624756002614,94.95097613101825,94.95097613111825,94.9557009129787,94.9557009130787,94.95884839309647,94.95884839319648,94.9635645358367,94.9635645359367,94.96513509302498,94.96513509312499,94.96905843574469,94.96905843584469,94.97376230169827,94.97376230179827,94.98394582042475,94.98394582052475,94.98785897038024,94.98785897048025,94.99255090056906,94.99255090066906,94.99802100473012,94.99802100483012,95.0058291925917,95.0058291926917,95.01051005851603,95.01051005861603,95.01596572608541,95.01596572618541,95.0260891491835,95.02608914928351,95.03153576072818,95.03153576082818,95.03853235447613,95.03853235457613,95.04629859665008,95.04629859675008,95.05095290076135,95.05095290086135,95.05792819958928,95.05792819968929,95.06412339198782,95.06412339208782,95.06799099658056,95.06799099668056,95.07262699918584,95.07262699928584,95.07648679816735,95.07648679826735,95.07957243965618,95.07957243975618,95.08573830417555,95.08573830427555,95.088817920945,95.088817921045,95.0926645716396,95.0926645717396,95.101887393921,95.101887394021,95.10879797437993,95.10879797447993,95.11493500360723,95.11493500370723,95.1195336832655,95.1195336833655,95.12412880114,95.12412880124,95.12948500400289,95.12948500410289,95.1340724678314,95.1340724679314,95.13712779626682,95.13712779636683,95.14094333819035,95.14094333829036,95.14704437066675,95.14704437076675,95.15618686335664,95.15618686345664,95.16151555712355,95.16151555722355,95.1683603961479,95.1683603962479,95.17215994793575,95.17215994803576,95.17823403968593,95.17823403978593,95.18354386357711,95.18354386367712,95.18809089737064,95.18809089747064,95.19112004290619,95.1911200430062,95.19717276395372,95.19717276405372,95.2024642863389,95.2024642864389,95.2092617991086,95.2092617992086,95.21152504745854,95.21152504755854,95.21680239849125,95.21680239859126,95.22358210234208,95.22358210244208,95.22885126885038,95.22885126895038,95.23561915061039,95.23561915071039,95.24087808646316,95.24087808656316,95.24688242029919,95.24688242039919,95.2528799336982,95.2528799337982,95.2551268508906,95.2551268509906,95.26111455043858,95.26111455053858,95.26485377925015,95.26485377935015,95.27157577462556,95.27157577472556,95.27829204841214,95.27829204851214,95.28500366156676,95.28500366166676,95.28947229822593,95.28947229832593,95.29393714575342,95.29393714585342,95.30211583491025,95.30211583501026,95.30582948708302,95.30582948718302,95.3125093473142,95.3125093474142,95.32363359376181,95.32363359386181,95.33030163190217,95.33030163200218,95.33326189102385,95.33326189112385,95.33695871945197,95.33695871955197,95.3436072435642,95.3436072436642,95.34803465447072,95.34803465457072,95.35171928189486,95.35171928199486,95.3576101312372,95.3576101313372,95.36423129781872,95.36423129791872,95.3708480690075,95.3708480691075,95.37452049542763,95.37452049552763,95.37818932217267,95.37818932227268,95.38478836022522,95.38478836032522,95.38991399442563,95.38991399452563,95.39430362768935,95.39430362778936,95.39722705645114,95.39722705655115,95.40744823914648,95.40744823924648,95.41474341647049,95.41474341657049,95.4198455201632,95.4198455202632,95.42567168902784,95.42567168912784,95.43076413043477,95.43076413053477,95.43439893077736,95.43439893087736,95.44093503135277,95.44093503145277,95.44818928605467,95.44818928615467,95.45253697363074,95.45253697373074,95.45760493724705,95.45760493734706,95.46484066309507,95.46484066319508,95.47062170340145,95.47062170350145,95.47712025368195,95.47712025378195,95.48000507119195,95.48000507129196,95.48793257881,95.48793257891,95.49441260023661,95.49441260033662,95.50232574704562,95.50232574714562,95.5095136919432,95.5095136920432,95.51669557423008,95.51669557433009,95.52674150958542,95.52674150968542,95.53104156777633,95.53104156787633,95.53748434203855,95.53748434213855,95.54749685725251,95.54749685735251,95.5539271273802,95.55392712748021,95.55821077863095,95.55821077873095,95.56391702147805,95.56391702157805,95.57033133936139,95.57033133946139,95.57602807148798,95.57602807158798,95.58171911991705,95.58171912001706,95.58882612896204,95.58882612906204,95.59308647705299,95.593086477153,95.59734322546885,95.59734322556885,95.60159648788355,95.60159648798356,95.60372086455756,95.60372086465756,95.60867379013621,95.60867379023621,95.61291469998166,95.61291470008166,95.6164453749858,95.6164453750858,95.62138477336435,95.62138477346436,95.62984185002541,95.62984185012542,95.63617742975254,95.63617742985254,95.64461840260397,95.64461840270397,95.64883460731089,95.64883460741089,95.6530469471035,95.6530469472035,95.65795675236367,95.65795675246368,95.66356337893902,95.66356337903902,95.67196600577789,95.67196600587789,95.67616345428301,95.67616345438302,95.68035684841725,95.68035684851725,95.68664013812725,95.68664013822725,95.69431229616167,95.69431229626167,95.6977956449112,95.6977956450112,95.70406112570231,95.70406112580231,95.70962470774181,95.70962470784181,95.71309922781452,95.71309922791453,95.71865420852512,95.71865420862513,95.7214292927818,95.72142929288181,95.72766540780243,95.72766540790244,95.73597402628444,95.73597402638444,95.74358305738515,95.74358305748515,95.74703840445157,95.74703840455157,95.75118136524321,95.75118136534321,95.75807982048076,95.75807982058076,95.76428281849054,95.76428281859054,95.76772604033562,95.76772604043562,95.77254214036945,95.77254214046945,95.77597895858199,95.77597895868199,95.78147255535944,95.78147255545944,95.78490232578706,95.78490232588706,95.78832868599618,95.78832868609618,95.79722821956301,95.79722821966301,95.80338445968792,95.80338445978792,95.80885138097837,95.80885138107837,95.814995268534,95.81499526863401,95.82249619570801,95.82249619580801,95.82862564667197,95.82862564677197,95.83611054581898,95.83611054591898,95.8408694299645,95.8408694300645,95.84834083980274,95.84834083990275,95.85919727042352,95.85919727052352,95.86326410755979,95.86326410765979,95.87003642260385,95.87003642270385,95.87612263957539,95.87612263967539,95.88490607414768,95.88490607424768,95.89233163549264,95.89233163559264,95.89907462265754,95.89907462275754,95.90715987182726,95.90715987192726,95.91052556803824,95.91052556813824,95.91456004599,95.91456004609,95.92060590870919,95.9206059088092,95.92731767342907,95.92731767352907,95.93201161185767,95.93201161195768,95.93536074956324,95.93536074966325,95.94071566412136,95.94071566422136,95.946733828288,95.946733828388,95.9514103367109,95.9514103368109,95.95674948848125,95.95674948858125,95.96208375227172,95.96208375237173,95.96674552097241,95.96674552107241,95.9707366512578,95.9707366513578,95.97671726512952,95.97671726522952,95.98534845239108,95.98534845249108,95.99198098636764,95.99198098646764,95.99860912495143,95.99860912505143,96.00390568688175,96.00390568698175,96.01052124554839,96.01052124564839,96.0158082967582,96.0158082968582,96.02108992950971,96.02108992960972,96.02438715611132,96.02438715621132,96.02636364297602,96.02636364307602,96.03228878396001,96.03228878406001,96.03689220003288,96.03689220013288,96.0428062767524,96.0428062768524,96.04674481389011,96.04674481399012,96.0513351195674,96.0513351196674,96.05985145824786,96.05985145834786,96.06835930927336,96.06835930937336,96.07620421382089,96.07620421392089,96.08142810020566,96.08142810030566,96.08403725838626,96.08403725848626,96.09250922633481,96.09250922643481,96.10227438703296,96.10227438713297,96.10877740848883,96.10877740858884,96.11462396288188,96.11462396298188,96.11851729567871,96.11851729577872,96.1237041243557,96.1237041244557,96.12629392007344,96.12629392017344,96.13082170507806,96.13082170517806,96.13534539782047,96.13534539792047,96.13986329319142,96.13986329329143,96.14115046122167,96.14115046132167,96.14179184754043,96.14179184764043,96.14242952051013,96.14242952061014,96.14303934336203,96.14303934346204,96.14359638360969,96.14359638370969,96.14408578785692,96.14408578795693,96.14452479665287,96.14452479675288,96.14493963078874,96.14493963088874,96.14527227865855,96.14527227875855,96.14558867115348,96.14558867125348,96.14588623166398,96.14588623176398,96.1464708188989,96.1464708189989,96.14675360179587,96.14675360189587,96.14702452471069,96.14702452481069,96.14728552010051,96.14728552020051,96.14753431448634,96.14753431458634,96.14775730488546,96.14775730498546,96.14796422936627,96.14796422946627,96.14815209451476,96.14815209461476,96.14830919191404,96.14830919201404,96.14830945715325,96.14830945725325]],"filenames":["B6Eve","B6Eve\_broken"]}
